# R/LinkedCharts: A novel approach for simple but powerful interactive data analysis

**DOI:** 10.1101/2022.05.31.494177

**Authors:** Svetlana Ovchinnikova, Simon Anders

## Abstract

In any research project involving data-rich assays, exploratory data analysis is a crucial step. Typically, this involves jumping back and forth between visualizations that provide overview of the whole data and others that dive into details. In data quality assessment, for example, it might be very helpful to have one chart showing a summary statistic for all samples, and clicking on one of the data points would display details on this sample in a second plot. Setting up such interactively linked charts is usually too cumbersome and time-consuming to use them in *ad hoc* analysis. We present R/LinkedCharts, a framework that renders this task radically simple: Producing linked charts is as quickly done as is producing conventional static plots in R, requiring a data scientist to write only very few lines of simple R code to obtain complex and general visualization. We expect that the convenience of our new tool will enable data scientists and bioinformaticians to perform much deeper and more thorough EDA with much less effort. Furthermore, R/LinkedCharts apps, typically first written as quick-and-dirty hacks, can also later be polished to provide interactive data access in publication quality, thus contributing to open science.

## Introduction

Effective data visualization has been crucial for scientific success since the first quantitative experiments. The continuously growing amount and complexity of available data over the last decades make this task more and more challenging. For a while, the only possible approach was to develop ever more elaborate types of plots employing color, shape, transparency, and other visual aspects to combine multiple layers of information (Bertin 2011; Tufte 1983; Wilkinson 1999). Yet, there are certain limits to how much information one can learn from a static image (Hegarty 2011). Excessive details and multiple overlapping layers make it harder to grasp the crux of a plot. Thus, a researcher has to decide what information to keep and what to dismiss to convey the message better (O’Donoghue et al. 2018). This is, arguably, an essential step since data often contain much noise and information irrelevant to the point one tries to make. Yet, it may be useful to provide the reader with a way to estimate the relevance of the omitted pieces of data on his or her own to avoid biases (Bresciani and Eppler 2009) and thus boost confidence in reported data patterns.

With the advance of modern computer technology, interactive visualization emerged to offer new ways of presenting information, starting already in the 1970s (Newman 1979; Becker and Cleveland 1987). In an interactive figure, there is no need to fix all the parameters or to exclude any data that do not contribute to the main idea. Instead, the user can experiment with the data and explore details quickly and intuitively, navigating to the plot’s most exciting or suspicious parts. Numerous tools (Caldarola and Rinaldi 2017) now provide means of interactive inspection for many specific types of data. Examples from biology include metabolic maps (Noronha et al. 2017), genome assemblies (Wick et al. 2015), scRNA-Seq data (Hillje et al. 2020), QTL data (Broman 2015) and many more. While such solutions are each tailored for one very specific type of data, there are also a number of general low-level frameworks to create interactive apps, such as D3 (Bostock et al. 2011) and Vega-Lite (Satyanarayan et al. 2015), and more highlevel but still general-purpose packages, such as Vega (Satyanarayan et al. 2016), Shiny (RStudio, Inc 2013), BPG (P’ng et al. 2019), and plotly (Sievert et al. 2019; Sievert 2020). However, the gap between these general-purpose frameworks and special-purpose apps is still wide, as we discuss below.

The advantage of interactivity lies beyond just simplifying navigation through big or complex data. When it merely takes a click or two to add changes to a plot, this urges a researcher not to put aside ideas or concerns and thus go through the data more thoroughly. At the same time, readers can check conclusions and claims of a paper on the fly, without going through all the scripts and analysis, thus making the findings more credible. Therefore, we believe that further integration of interactive tools in a researchers’ routine can significantly improve the quality of research (Shander 2016; Yuk and Diamond 2014).

In practice, however, interactivity is still seriously underused during data analysis. Even though many authors now accompany their papers with an interactive resource to present their data and results (for example Travaglini et al. (2020); Roider et al. (2020); Kalucka et al. (2020)), these chiefly serve to present and communicate research that has already been completed. Commonly, it is only after most of the work on a project has been finished and the paper is being written up that researches spend a couple of days implementing a nice-looking interactive app to share their data and results with the scientific community (Batch and Elmqvist 2017) to be added as supplement to their publication.

Visualization frameworks must be simple, to encourage practitioners to use interactivity as soon as there is merely the smallest hint that it might be useful. If they are too complicated, one will prefer to do most of the analysis by more conventional static means, and wait for a special occasion when it is worthwhile to invest time and effort into an interactive app. Furthermore, a good framework should be similar in design to common static visualization tools to facilitate seamless switching from static to interactive data visualization.

However, too simple frameworks lack flexibility and work only for particular data structures and precisely matching data flow patterns. While much of routine work may involve only standardized steps and data types, research that goes beyond routine should not be forced into a predefined mold.

Here, we present “LinkedCharts”, a framework to perform interactive data exploration.

With LinkedCharts, we aimed to find a balance between the simplicity of usage and possibility for flexible customization. With only basic coding skills, users can create with just a few lines of code fully functional apps for *ad hoc* analysis. These apps can then be improved by customizing many plot settings such as colors, labels, axes, etc., and, by investing a bit more effort, turned into a “publication quality” app for deployment on a public server. Since the library is JavaScript-based, LinkedCharts apps can be combined with various existing web solutions.

LinkedCharts is not fixed on any specific task. It is a toolbox, and its building blocks can be combined in any manner, the same way as one combines plots for a complex paper figure. All blocks share the same interface and very similar interactivity capabilities, which means that understanding one of them is enough to grasp the entire concept of LinkedCharts.

We hope that LinkedCharts can become a valuable asset for the scientific community that can be used both for everyday routine and for presenting one’s research to a greater audience. LinkedCharts available as an R package (“rlc”, can be downloaded from CRAN) and as a JavaScript library. This paper’s focus is on the R implementation of LinkedCharts, which we also refer to as R/LinkedCharts.

## Results

### Linking charts

As its name suggests, the central concept of LinkedCharts is linking and focusing (Buja et al. 1991): one can connect two or more plots thus that interacting with one of them affects what is displayed in the others. We illustrate the concept of linking charts with a simple example based on data from Conway et al. (2015).

In that study, three samples were taken from each of 17 patients with oral cancer: of normal, cancerous, and dysplasic tissue. mRNA from all these samples was sequenced to obtain gene expression values. The goal was to find genes that differentially expressed between the tissue types – a standard task in bioinformatics, readily addressed using available software tools (Ritchie et al. 2015; Love et al. 2014). Here, we have used the function `voom` from the “limma” package (Law et al. 2014) to compare normal and cancerous tissues. It is common to visualize such a comparison with an MA plot (Dudoit et al. 2002), where each dot represents a gene, showing the gene’s average expression on the X-axis and log fold change between the two groups on the Y-axis (Fig 1(A)). Red dots correspond to genes that are considered significantly different between the two conditions (adjusted p-value < 0.1).

**Figure 1:**
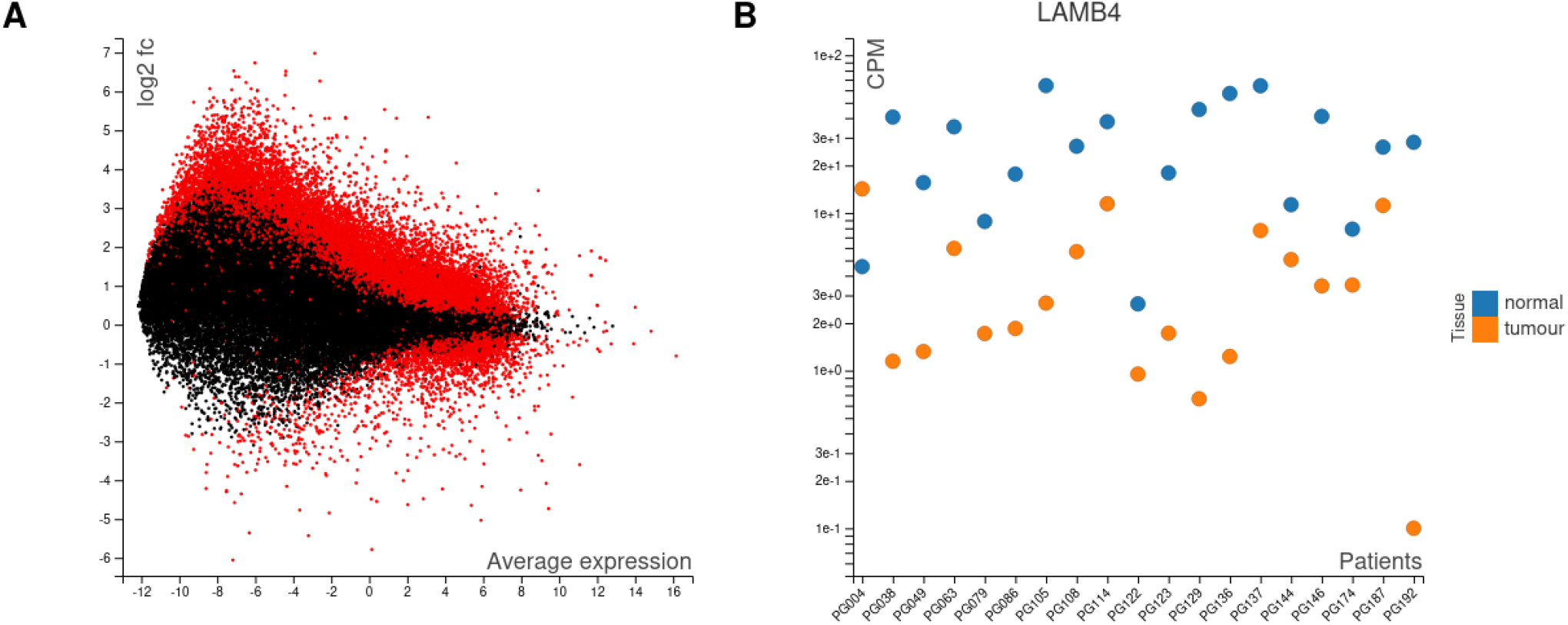
An example for two linked charts, based on a study by Conway et al. (2015) comparing cancerous and normal tissues from 19 patients. The MA plot (A) shows all genes with their average expression on the X-axis and log_2_-fold change between tumour and notmal on the Y-axis. Red indicates genes where the difference was reported as significant by the “voom” method (Law et al. 2014). The plot to the right (B) shows, for one selected gene (here, LAMB4), the individual expression values (as counts per million, CPM) for each sample. This figure is a screenshot of a LinkedCharts app, the live version of which is provided in the Supplement(as Interactive Supplementary Figure 1): When the user clicks on any point in the MA plot (A), the expression plot (B) changes, showing the selected gene. Thus, one can rapidly gain an impression of the details hidden in a summarizing plot like the MA plot.

About these genes, one may now wonder: How does the difference in expression look like for every single patient? Is it consistent across all the patients or only detected in some of them? Are there any artifacts or outliers that cause the p-value to be too small?

To investigate such questions, we can add another plot that shows expression values (as “counts per million”, CPM) from each individual sample (Fig 1(B)). While this second plot can show expression for only one selected gene at a time, the *linking* between the two charts overcomes this limitation: A mouse click on a point in the MA plot causes the plot to the right to switch to displaying the expression values for the thus selected gene. Fig 1 depicts a LinkedCharts app, which is provided in this paper’s online supplement (https://anders-biostat.github.io/lc-paper/) – and we encourage the reader to pause for a moment and try it out there.

In the Supplement, we also provide full code to generate the app and links to necessary data files to immediately get this app in one’s R session and experiment with it. For this and all further examples, we provide two versions of the code: minimal with only essential parameters needed to make the app functional, and more extended with custom colors, labels, etc. In the paper, we only focus on the minimal code.

To illustrate typical usage of the “rlc” package and to explain the linking mechanism, we briefly discuss the code that generates the app shown in Fig 1. This is a minimal, but complete code for the app, suitable to illustrate the design principles of R/LinekdCharts. For now, we concentrate only on the highlighted lines.

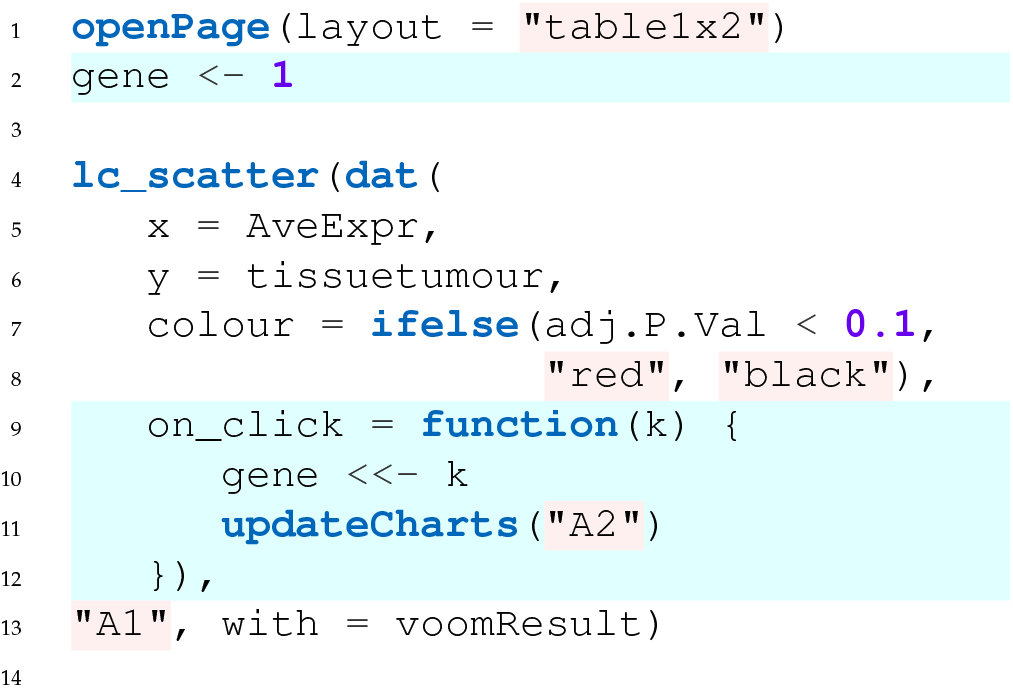
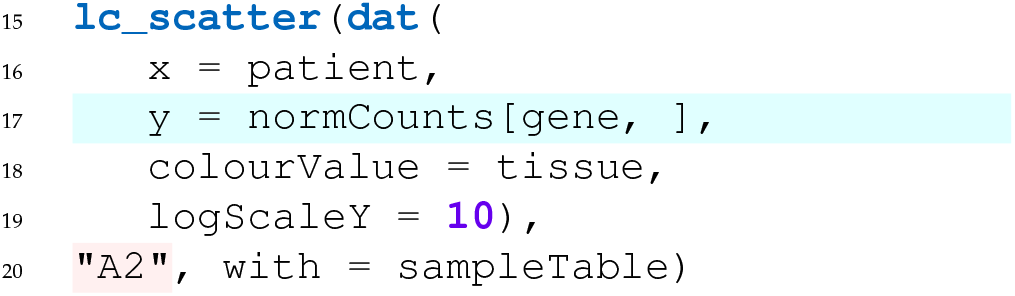

In Line 2, we introduce a global variable, `gene`, which stores the index of the gene to be shown in the righthand plot. This index tells the chart which line of the `normCounts` matrix (where the normalized counts are stored) to use as *y* values of the expression plot (Line 17). Almost every chart type in R/LinkedCharts has the `on_click` argument, which allows the user to define a function that is called whenever someone clicks on an element of the plot (point, line, cell of a heatmap, etc.) and is passed the index of the clicked element (`k`). Here, our callback function simply changes the value of `gene` to the clicked point index (Line 10). Then, we tell R/LinkedCharts to update the second plot (Line 11; “A2” is its ID set in Line 20). Updating means that the package will reevaluate all arguments inside the **dat**() function and redraw the chart accordingly. In our case, a new value of `gene` will yield new *y* values for the expression plot.

This simple logic is not limited to just two plots, but provides a basis to create many simple and complex apps. For example, the tutorial at https://anders-biostat.github.io/linked-charts/rlc/tutorials/citeseq1.html gives detailed instructions to generate an app for single-cell data exploration. The app consists of four charts, three of which are scatter plots and one is an information table to show genes that define a selected cell cluster.

Besides mouse clicks, LinkedCharts can react to other events, such as moving the mouse cursor over or out of an element, selecting or deselecting elements with the *Shift* key pressed, clicking on any position of a plot or on a heatmap label. The complete list can be found on the man page of any function of the “rlc” package. Understanding how to define these callback functions (above is shown a very typical example) is all one needs to generate apps.

### Basic syntax, chart types, and HTML5 integration

We aimed to make R/LinkedCharts simple and familiar to any user with at least some basic knowledge of R. Every chart has a set of properties to define each of its specific aspects. In the previous example, we set the properties `x`, `y` and `color`, which received vectors of coordinates and colors to specify the scatter plots’ data points. This principle will be familiar to most users from other plotting libraries. For example, Figure 2 shows a comparison of the syntax in R/LinkedCharts (“rlc” package) and ggplot (from the widely used “ggplot2” package by Wickham (2016)) for a simple scatter plot. Lines are arranged to match the same aspects of the plots; above each code block, its output is shown. One can see that the input data structure is identical, and there is hardly any difference between the two.

**Figure 2:**
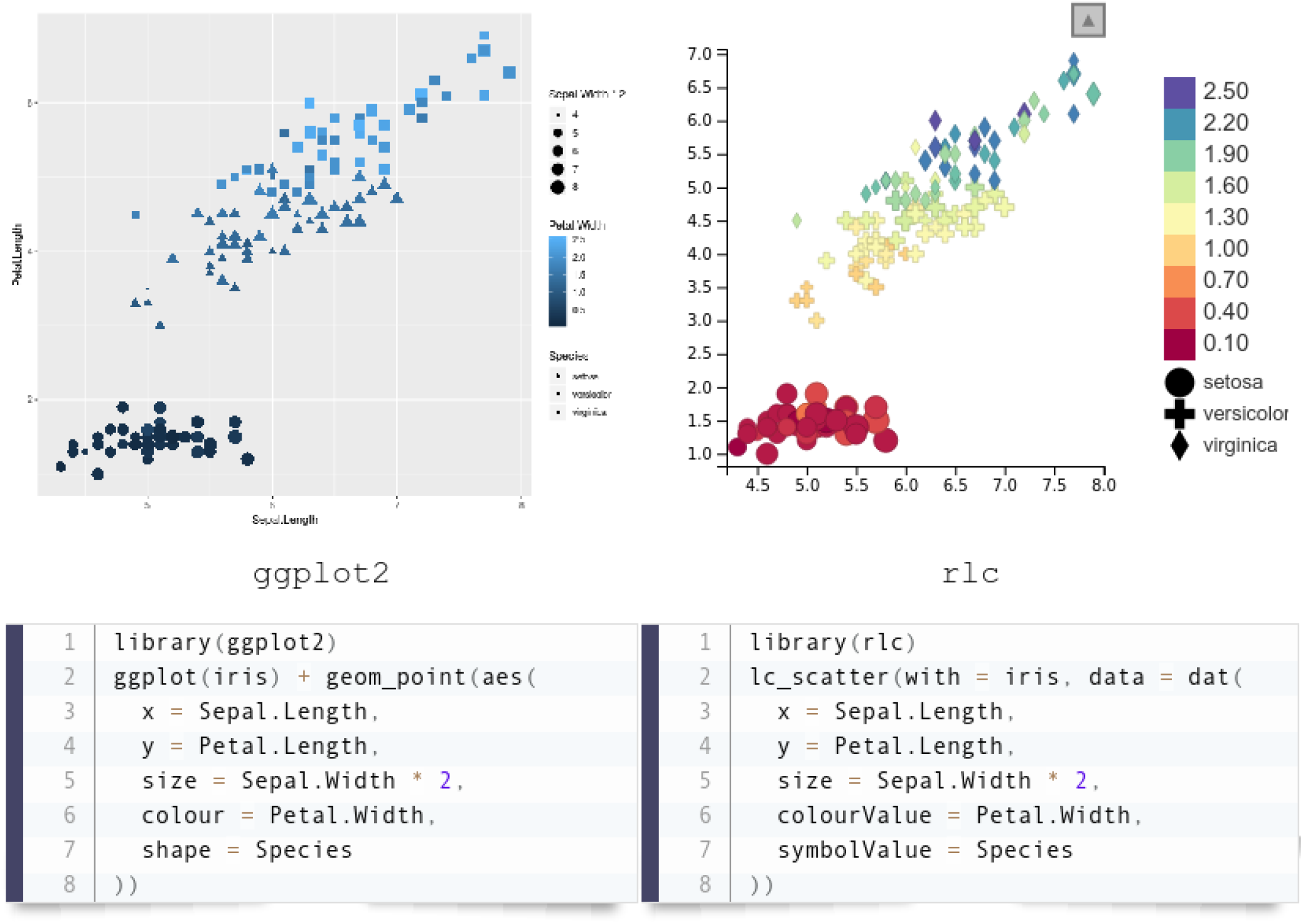
Typical syntax of an R/LinkedCharts plot with comparison to the “ggplot2” (Wickham 2016) package, one of the most widely used plotting libraries. Lines of the code are arranged to put the same aspects of the charts next to each other. The “iris” dataset, one of the built-in example datasets of R, was used here. Both pieces of code are complete and fully functional, and their output is shown above the code.

R/LinkedCharts is not limited to scatter plots. There are 15 main functions in the “rlc” package, each generating a specific type of plot (such as scatter plot, heatmap, bar plot, etc.) or a navigation element (such as sliders or text fields). Figure 3 shows them all. Each plot is defined by its properties: some of them are required (such as `x` and `y` for a scatter plot or `value` for a heatmap), others are optional (`palette`, `title`, `ticks` etc.). A full list of all the properties with live examples is available at https://anders-biostat.github.io/linked-charts/rlc/tutorials/props.html and also on the R man page of each plotting function.

**Figure 3:**
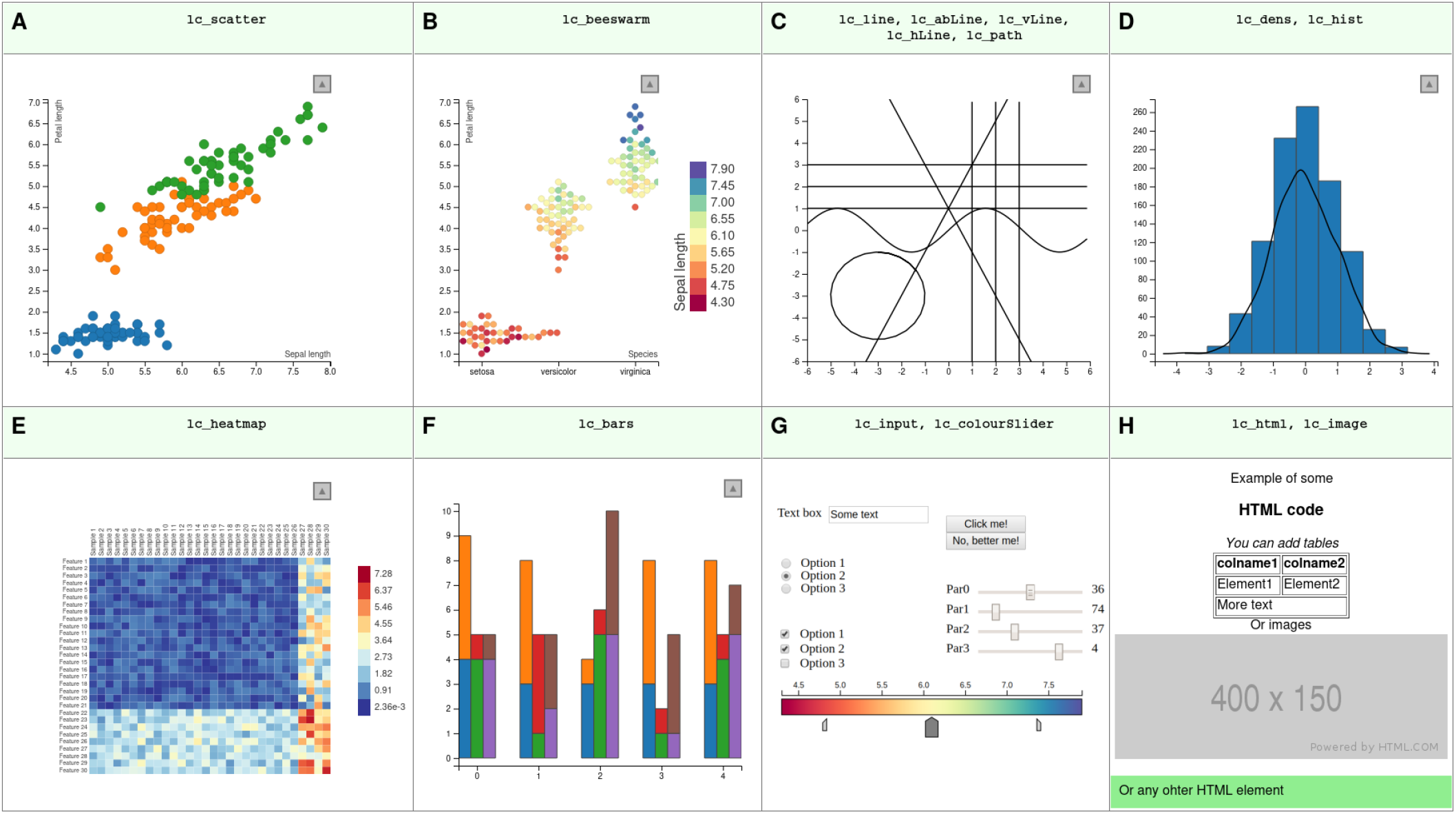
Gallery of all available plotting functions in the “rlc” package. A scatter plot (A); a bee swarm plot (based on the d3beeswarm plugin of Lebeau (2017)) (B); a collection of various lines (C); a histogram and a density plot (D); a heatmap (E); a bar chart (F); a collection of interactive elements to gather input from the user (G); functions to add custom HTML code and static plots to the page (H). All these examples with code to create them can be found in the Supplement.

LinkedCharts apps are displayed as HTML pages, using a standard Web browser. This means that the layout as well as decorations (such as headlines) can easily be specified by producing a standard HTML5 page, in which the elements where the charts are to be placed are marked by their `id` attribute. As knowledge of HTML5 is wide-spread, this allows practitioners to improve the appearance of LinkedCharts app without having to learn anything new. Furthermore, it facilitates integrating LinkedCharts with other web-based apps. For example, one can easily link a LinkedChars app with a web-browser-based genome browser, such as *IGV*.*js* Robinson et al. (2020), so that the user’s interaction with the LinkedCharts app controls what genomic region is displayed in IGV’s genome track.

Once one has developed a rough prototype of a LinkedCharts app, the app’s appearance can be easily improved by using HTML5 to specify layout, decorations, and add further static elements. To facilitate this, the web server integrated in R/LinkedCharts provides basic functionality to also serve, e.g., images and CSS style sheets.

### Use cases

#### Back-tracking in analysis pipelines

Most analysis of big data comprises multiple steps of data summarization, each reducing the total amount of data and thus losing information.

For example, in the oral-cancer example, we first have for each gene expression values from 28 samples, but the differential expression data analysis summarizes this to just 3 values: the gene’s average expression over all samples, the fold change between tumor and healthy and the associated p-value. The LinkedCharts app shown in Figure 1 allows to “undo” this summarization by inspecting the original values for each gene.

As an example of an analysis pipeline with multiple data-reduction steps, we use the drug-screening study of He et al. (2018). A collection of drugs was tested against various pancreatic cancer cell lines at several concentrations per drug. Figure 4 illustrates a possible analysis pipeline: Panel A shows the viability readout from the microtiter plates. For each combination of one cell line and one drug, the values for the different tested concentrations can be shown as a scatter plot, with each point depicting the viability value from one well (panel B). Here, we can fit dose-response curves, which can then be further summarized to a single number, such as the area under the curve, or, in the case of this study, a refined variant of that, called the drug sensitivity score (DSS) (Yadav et al. 2014). If two drugs show effect on the same subset of cell lines, they likely have similar mode of action. Hence, to assess similarity for each pair of drugs, we compare their activity over all cell lines, as shown by the scatter plots in panel D, where each point represents a cell line, with its *x* and *y* coordinates denoting the drug sensitivity scores of the that cell line for the two compared drugs. Again, we summarize each such plot into a single number, the correlation coefficient, and finally, we depict all the correlation coefficients in a correlation-matrix heatmap (panel C).

**Figure 4:**
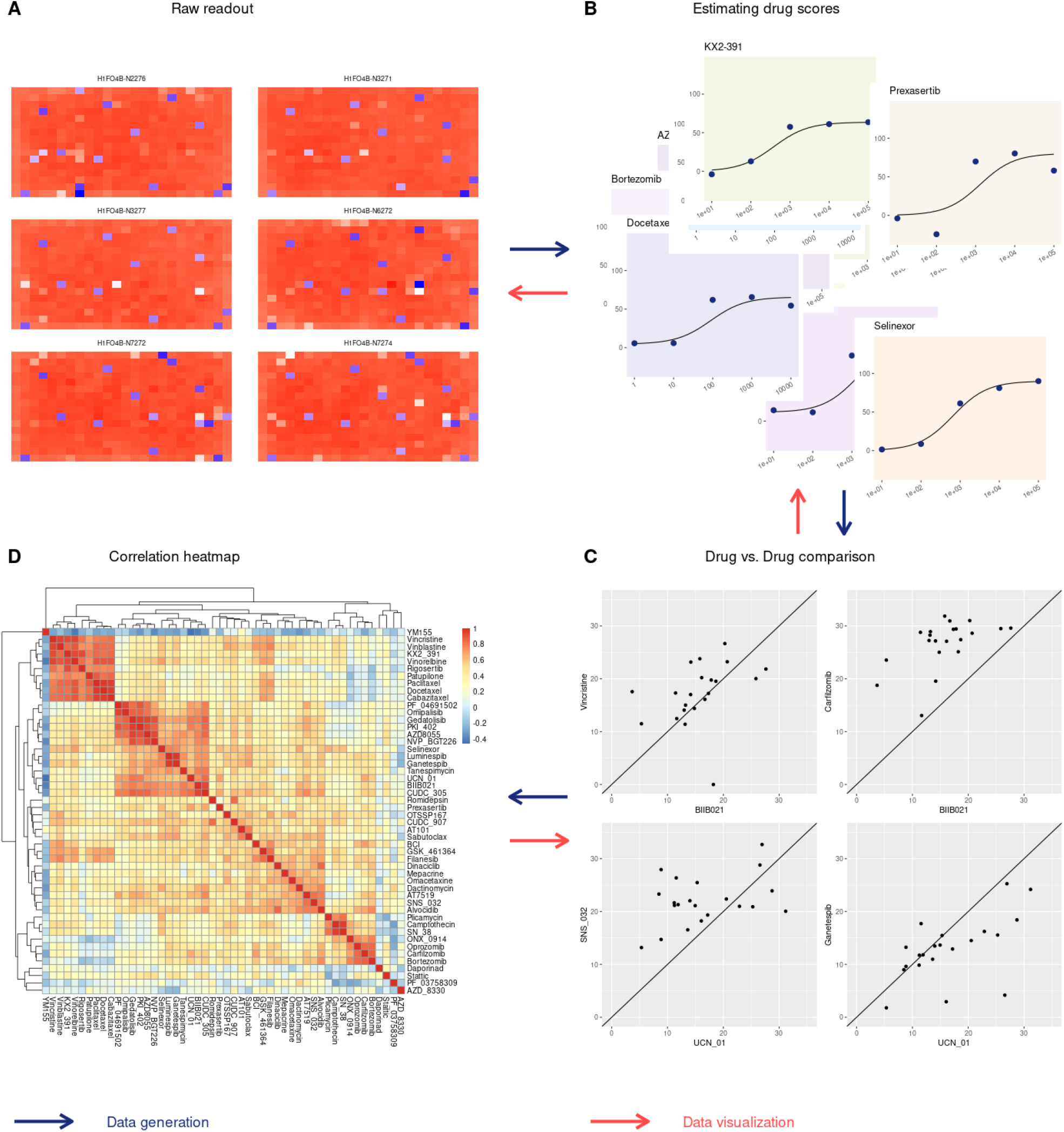
LinkedCharts can be used to “walk backwards” through an analysis pipeline. This is illustrated here using a drug screening experiment (He et al. 2018) as an example. For an interactive version, see Interactive Supplementary Figure 4. The *blue arrows* show the direction of a typical analysis pipeline used in drug screening experiments. We start with reading intensity values from plates with different cell lines cultured in the presence of studied drugs (A). These values are then normalized and turned into a fraction of the cells that remained viable. A sigmoid curve is fitted to the obtained viability values at different drug concentrations, and the area under the fitted curve yields a single score for each drug (B). Different drugs’ scores are compared to each other across all the tested cell lines (C). A drug-drug correlation heatmap is then produced to identify clusters of similar drugs (D). The *red arrows* illustrate the direction of interactive data exploration: We start by showing the summary heatmap plot (D). Suppose the researcher is interested in a particular drug combination or a cluster of drugs. In that case, he or she can examine the corresponding drug scores simply by clicking on the heatmap cell (D) to see the underlying correlation plots (C). Similarly, one can click in a point in (C) to examine the individual viability values at the tested concentrations and check the sigmoid fit (B). And finally, if needed, it is possible to take one more step back and to look at the raw read-outs to inspect them for the presence of any artifacts (A).

Often, such an analysis pipeline is fully automated and no one ever looks at the intermediate plots. Inspecting them is, however, crucial to note problems with data quality or mistakes in the design or programming of the analysis pipeline.

LinkedCharts allows to “walk” such an analysis pipeline backward: In the Supplement, we show an app that depicts the plots of Figure 4 in an interactive fashion, as follows. As each cell of the final heatmap (panel C of Figure 4) summarizes on scatter plot comparing two drugs (panel D), we can click on any cell in the heatmap and then see the corresponding scatter plot. Each point in that scatter plot represents a pair of drug sensitivity scores, which are, themselves, summaries of a dose response curves. Again, clicking on a point in the correlation scatter plot will display these two dose response curves. Finally, each value in a drug response curve stems from a well in a microtiter plate, and hovering over a point there hence highlights the well in a heatmap depicting the plate.

Thus, LinkedCharts allows to explore the “parentage” of any result value. If we find a specific drug-drug correlation value suspicious or surprising, or if we just wish to double-check it before drawing further conclusions from it, we can check its provenance in arbitrary detail. Similarly, we can perform random spot checks.

Each layer in the backwards journey can inform about another type of problem: From the correlation scatter plots, we may find that the correlation coefficient was unduly influenced by a single out-lying cell line, from the dose response plot, we may find that specific dose response curves fail to have the expected sigmoid shape, and from inspecting plate plots, we may trace back a surprising final result to, say, a normalization issue or a plate-edge effect.

Once such an analysis pipeline has been developed, all the intermediate results are typically available in suitable data structures, which can be readily explored with LinkedCharts. The Supplementprovides code for the example just described.

### Quality assurance thresholds

Typically, analysis pipelines include steps to exclude bad-quality data. Often, this is done by calculating quality metrics and setting thresholds. In the drug screen example, the goodness of fit of the dose response curves might be quantified by the residual sum of squares, and if this value exceeds a threshold, the drug sensitivity score might be discarded as unreliable. In the oral-cancer sample, the log fold change of some genes might be unduly influenced by a single outlying sample, and one might use a threshold on an outlier-detection score such as Cook’s distance to flag such genes.

Typically, the thresholds on such quality metrics are chosen a priori, often simply taking over values from previous work or from tutorials, even though the characteristics of the assay might have changed. Doing otherwise seems to cause a chicken-and-egg problem: One cannot run the analysis without first somehow deciding on thresholds, and therefore, one cannot use analysis results to guide the choice of thresholds.

The approach of “walking the pipeline backwards” with LinkedCharts opens another approach: Typically, outliers tend to cause false positive results. Therefore, one can run the analysis first without excluding any outliers, then inspect the provenance of the statistically significant items found and will be so guided to specifically those places in the raw data where outliers can actually cause false positives. This provides the analyst with a better “feel” for the data and the analysis procedure and helps build an intuition that will allow to more critically judge whether traditionally used standard values for quality-assurance thresholds are appropriate for the specific data set under analysis.

#### Exploratory analysis

Analyzing complex data sets from many different angles and asking many different questions on it is a crucial to all computational biology, not only to ensure that one does not overlook potential problems but also in order to not miss the chance of serendipitous discoveries. The importance of such exploratory data analysis (EDA) has been argued since long, and it therefore forms a large part of computational biologists’ everyday work. An important element is to pick examples and study them in detail, similarly to the quality-assurance applications discussed in the previous section, but now with the aim of getting a “feel” for the data and looking for insights.

The standard approach in inspecting examples is to pick, e.g., a gene from a result list, produce a plot showing the provenance of this result, then pick another gene, change the code for plotting to now show underlying data for this gene, etc. At that point, it is trivial to alter this code into a linked charts app, using the similarity between code for static and dynamic plots (Figure 5).

**Figure 5:**
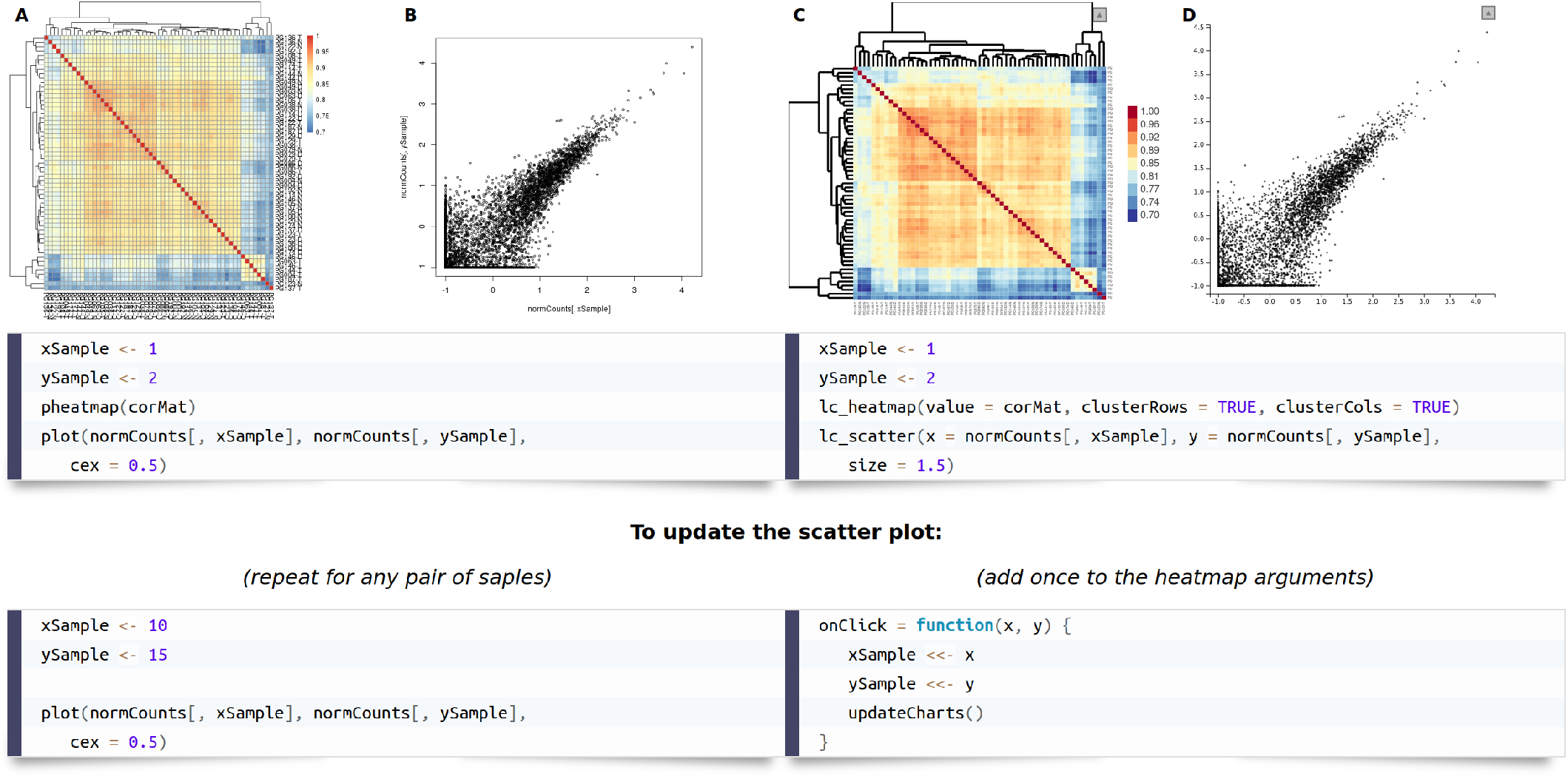
An example of an R/LinkedCharts app (C, D) for a simple exploratory analysis and the code to generate it in comparison with static plots (A, B) produced for the same purpose. The heatmaps (A, C) show Spearman correlation of gene expression for all samples from Conway et al. (2015). Here, we can see, inter alia, two outlier samples in the heatmap’s bottom-right corner and some more or less pronounced clusters of samples with similar gene expression levels. The scatter plots (B, D) show expression values for two samples plotted against each other. Browsing through several such plots can help the researcher get a feeling of the data and explore unexpected patterns like the outliers just mentioned. The code is split into two pieces, where the upper one is responsible for generating the plots and the lower part shows the code to update them to show a specific sample pair. For static plots, one has to execute the same lines of code for any pair of samples, while for R/LinkedCharts the provided code should be added to the list of arguments for the heatmap. After that, switching between pairs of samples can be done simply by clicking on the corresponding cell of the heatmap. The static heatmap (A) was generated with the “pheatmap” package (Kolde 2019); scatter plot (B) was made with a base R function. The live version of the app can be found in the Supplement. For simplicity, gene expression for all the samples is subset to 8000 randomly selected genes.

Figure 5 illustrates this with another example based on the oral cancer dataset. A bioinformatician had produced a correlation heatmap depicting correlations between all sample pairs (Figure 5A), using, e.g., the pheatmap package (Kolde 2019), and now wishes to inspect a specific correlation value (panel B) and writes to this end the short code shown in the figure. To inspect other sample pairs, she would simply change the sample indices in the code. This is routine practice for most bioinformaticians, but cumbersome. As the code example below the plots in Figure 5 shows, however, it is now virtually none effort to transform the code into a LinkedCharts app, by merely making a few simple substitutions.

### Public apps and concurrent use

Technically, an R/LinkedCharts app is provided by a web server running inside the R session and can hence be used from any web browser. Importantly, there is no need for that web browser to be running on the same computer as the R session. This allows a bioinformatician to easily share a LinkedCharts application with colleagues. They only have to direct their operating system’s firewall to open the TCP/IP port the app is listening at for incoming connections and tell their colleagues their computer’s IP address or DNS name and the port number, which they simply enter into their browser’s address line.

As now multiple users might use the app simultaneously, we have to make sure that each user gets their own copy of any global variable, such as the variable `gene` in the initial code example. To do so, a trivial change is required: one only has to list all such session variables at the beginning. In the initial code example, one would simply amend the first line to

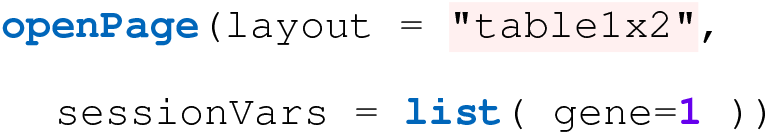

### LinkedCharts for Open Science

Analyses in computational biology are often complex and involved, making them difficult to explain and even more so to verify. It is not uncommon that neither the peer reviewers nor the readers of a publication are effectively able to double-check a result unless they would be willing to redo the whole analysis themselves. The importance of making all raw data and code available to do that has been often stressed (Gentleman 2005), but even verifying a complex analysis is a demanding task. Published interactive apps for data exploration are hence the next step towards open science. Traditionally, publications illustrate the characteristics of typical data by showing “typical examples” – but whether an example can be considered typical can be quite controversial. A LinkedCharts app in a paper’s online supplement allows readers to chose their own examples rather than relying on the authors potentially “cherry-picked” ones. A second, less obvious, advantage is that interactivity can help clarify the details of a complex analysis.

To illustrate this, we describe a LinkedCharts app that we have provided as online supplement to the paper of Wang et al. (2020), a big-data study aiming at elucidating to which extent evolution of expression regulation acts on transcription and to which extent on translation. Using RNA-Seq and ribosomal footprinting data from three organs, taken from animals of six species, changes in transcript abundance and in translation of transcripts into proteins were quantified and compared. A core idea of the analysis was that the evolutionary changes to transcription and translation may either compensate for each other (thus compensating deleterious changes in one layer by an opposite change in the other), or reinforce each other (in case of adaptive changes). To this extent, a score denoted as Δ was calculated, which is negative if the between-species difference is lower in the ribosome footprinting data than in the transcriptional data (thus indicating that transcriptional difference are at least partially compensated on the translational layer) and positive if the variance at the ribosomal layer is higher (indicating reinforcement).

The definition of this Δ-score is technical, and it is hard for the reader to form an intuition on its meaning. By “playing around” a bit with the app, available at (static picture: Figure 6), this is quickly remedied: The reader can click on any gene in the upper scatter plot, inspecting examples of genes with positive, negative, or near-zero Δ-score to see the data from the individual samples. After a few clicks, the relationship between the transcriptional and the translational data on the one hand and the Δ score on the other hand will be clearer than after reading several paragraphs of text. The use of HTML design elements to position explanatory labels renders the app nearly self-explanatory. Here, it is not a simple picture, but an interactive one, that is worth the proverbial thousand word.

**Figure 6:**
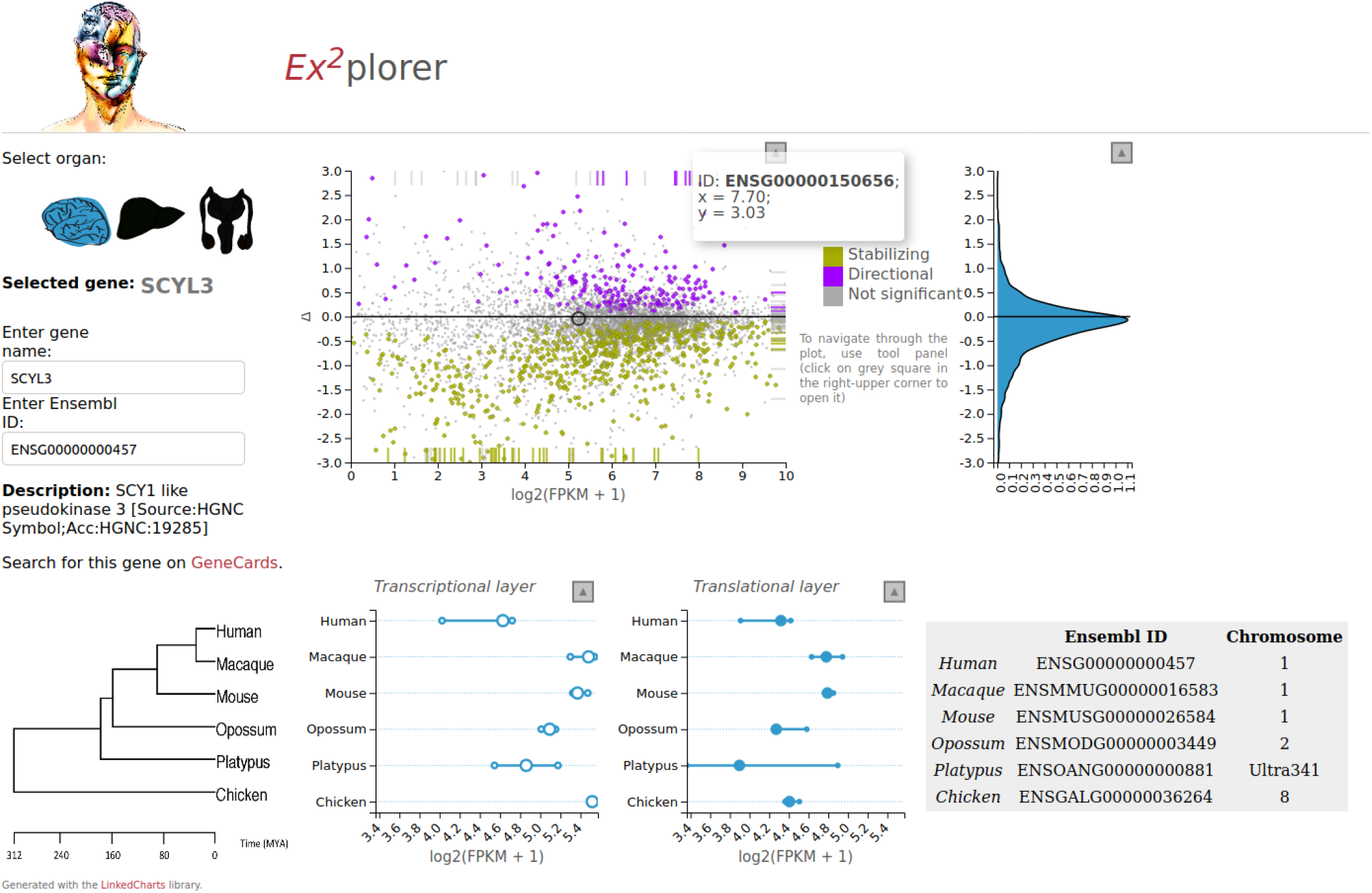
A LinkedCharts app as a paper supplement Wang et al. (2020). The main chart (upper row, centre) shows for every gene in the study its average expression and so-called Δ-score, which indicates whether evolutionary changes in the translatome compensate for changes in the transcriptome or introduces additional variance. The two plots below show expression values for the selected gene in all the tested samples. The user selects a gene by clicking on the corresponding point of the main plot or by entering the gene name (upper-left corner). The density plot to the right shows the distribution of Δ-scores, and its Y-axis is linked to the Y-axis of the main plot. In the upper-right and bottom-left corners, some additional information on the selected gene is displayed. Icons in the upper left corner allow switching between the three studied tissues. Detailed information on the data, study goal and the source code for the app can be found in the related publication. The app is written in JavaScript and, thus, can be downloaded and opened in any modern browser without installation requirements. Though largely customized, the app is based on the same principles as other examples throughout this paper. For the live version, see.

The app, when considered as a supplement to the publication, hence offers multiple benefits: It helps the reader in understanding the publication’s core quantitative analysis, it explains the meaning of calculated scores, and it allows the reader to verify that the examples discussed as typical in the paper are, in fact, typical, and not cherry-picked. Finally, it makes the data easily accessible; for example, readers can input their “favorite” gene to inspect its evolutionary analysis. All these aspects greatly contribute to making the publication’s results truly open science – the data is not only available somehow, but it is easily accessible.

### Javascript stand-alone apps

Apps based on R/LinkedCharts (“rlc” package) require a connection to a running R session. Typically, however, the server-side R session is only needed to provide data to the app and trigger updating of displayed charts, while all the visualization and interactivity handling happens on the client side, by the part of LinkedCharts written in JavaScript and running in the browser. In fact, this JavaScript component is the larger part of LinkedCharts, the code in the R package “rlc” is merely a thin layer to handle communication between server and client.

The JavaScript part of LinkedCharts can be used independent of R. The JavaScript library *linked-charts*.*js* offers a user-friendly and well documented interface providing access to all the functionality described so far and all the chart types shown in Figure 3. Bioinformaticians familiar with JavaScript will find its use not much more difficult then use of the R package discussed so far. The LinkedCharts documentation offers multiple tutorials describing the JavaScript API. In the Supplement, there is also a JavaScript version of the code for every app.

The advantage of working with the JavaScript library is that one can generate stand-alone apps, containing all code and data in a single HTML file. Such an HTML page can then be sent, as a single file, to a collaborator or used as a supplement file for a paper. Unlike a link to an app deployed somewhere on a server, this kind of interactive supplement will be available to any user with an installed web browser at any moment, without a need for the research team to maintain a running app on the server. The Supplementto this paper is an example of such an app contained within an HTML page. Another example is the app for Wang et al. (2020) (Fig. 6), discussed in the preceding section, which has also been implemented using the JavaScript rather than the R interface of LinkedCharts.

As Javascript-based LinkedCharts do not require that a running R session is maintained on a server, they are especially suitable for inclusion on web pages served by existing web servers, as the HTML/JavaScript code for the app can simply be inserted into existing pages. Thus, a JS app may be used in a blog or even as a main figure of a scientific publication: Some journals have recently begun allowing for interactive plots to be included into online articles as main figures (Ingraham 2017). So far, only the Plotly platform is typically supported, but as Plotly figures are integrated by simply inserting HTML snippets, extension of such services to other HTML/JS-based interactive apps should be straight-forward.

If one wishes to pack all parts of the app into a single file, one obviously has to include the data. This can be done inside an HTML file using the JSON format. In our “Data import Tutorial”, we explain this and also alternative ways of loading the data on demand, discussing also specific issues arising from web browser’s security models.

Unfortunately, we can not provide users with an automated converter of R/LinkendCharts apps into their JavaScript counterparts. Unrestricted customization is an essential part of the “rlc” package, and therefore providing such a converter is equivalent to developing a tool to translate any R code (including various packages and their chains of dependencies) into a JavaScript equivalent which is a major computer science project in its own.

### Apps with complex user interfaces

In all examples discussed so far, user input is constrained to selecting data points in one chart in order to affect the display in a linked chart. However, the LinkedCharts library also provides for more general means of data input by the user, by leveraging the HTML5 tag `<input>` and thus offering buttons, checkboxes, radio buttons, scrolls and text fields via the “rlc” function `lc_input` (Figure 3G). As for any other LinkedCharts element, `lc_input` can be provided a callback function that is run every time the user changes the state of an input element (e.g., clicks a button or enters new text). This allows to easily add functionality to enter, say, a gene name rather than clicking on its point (as in Figure 6), but also to build up complex apps.

Figure 7 shown a screenshot of an example of a more complex LinkedCharts app, which was developed as part of an effort to establish LAMP-based testing for SARS-CoV-2 at Heidelberg University campus (Dao Thi et al. 2020). In this project (see also (Herbst et al. 2021)), a colorimetric assay based on loop-mediated DNA amplification (LAMP, (Notomi et al. 2000)) was carried out on microtiter plates. Each patient sample was placed in a well of a 96-well plate, and was then split into four aliquots placed in adjacent wells of a 384-well plate. In the assay, successful amplification and hence detection of the virus template was indicated by a color change mediated by phenol red. The plots in the top-left corner of Fig. 7 show the well color (indicated as differential absorbance on the Y-axis) at several incubation times, representing each well or sample as a line. Moving the mouse over one of the lines or one of the wells in the plate highlighted all other lines representing aliquots of the same sample, as well as the sample position in the plate diagrams. This allowed the lab technicians to inspect each assay run for quality issues and double-check all automatic calls of assay results, and verifying correct performance of internal and external controls. Such continuous quality control is vital for reliable medical diagnostics and has to be offered in an easy-to-use manner and quick-to-grasp to avoid mistakes from repetition and fatigue.

**Figure 7:**
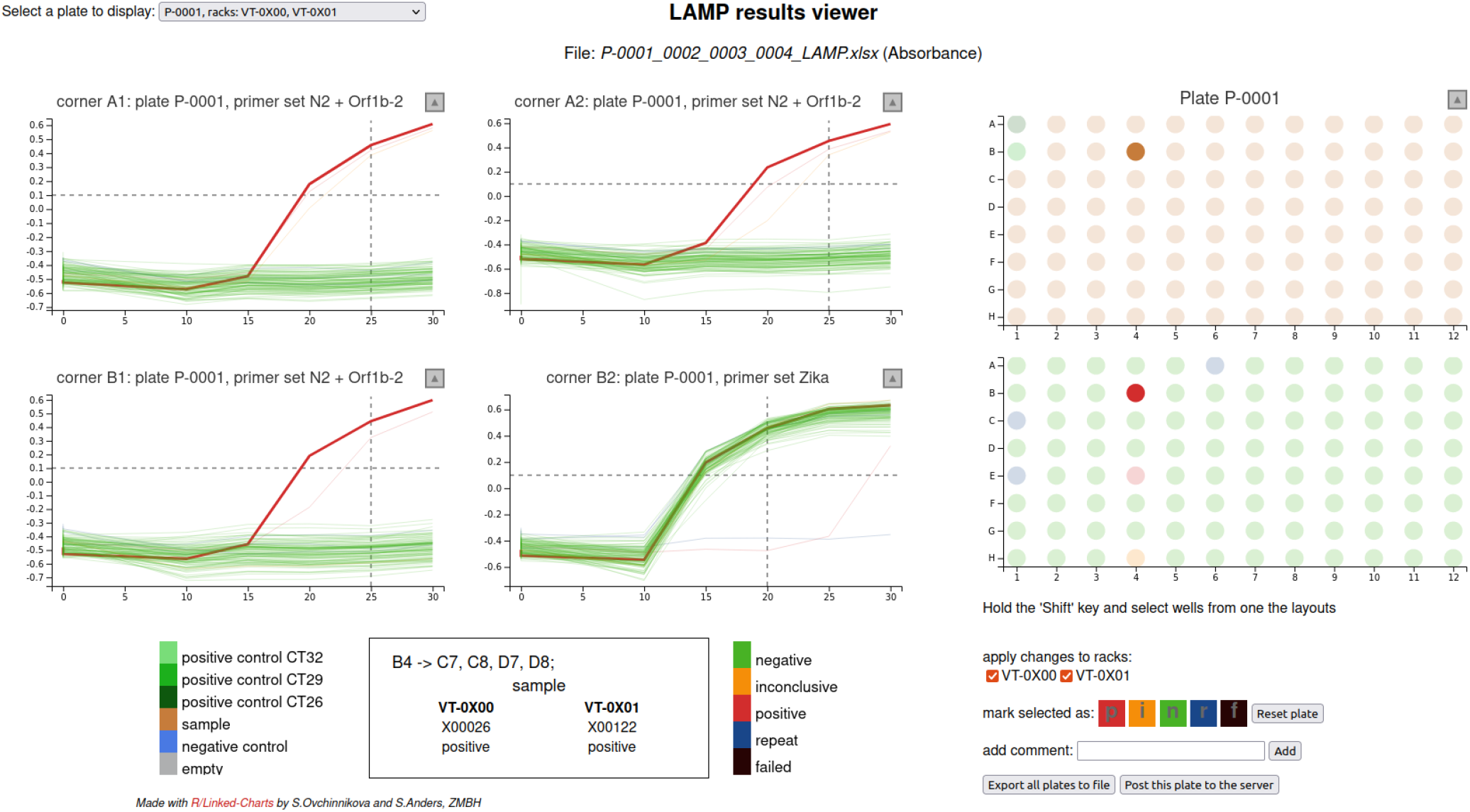
An example of an app that was used as a GUI to perform manual inspection and classification of LAMP testing for SARS-CoV-19 viral RNA (Dao Thi et al. 2020). The app was used during our SARS-CoV-2 surveillance study (Deckert et al. 2021) and for voluntary testing for Covid-19 infection on campus (University of Heidelberg) in 2020/21. To the right, the app shows a 96-well plate layout colored either by content type (sample, empty, positive or negative control) or by the assigned result. To the left, it shows the results of three tests and one control for each sample. Accumulation of the LAMP product is indicated by the change of color from red to yellow and is measured as a difference in absorbance on two wave lengths. This difference is plotted as a function of time. Besides exploration (highlighting the corresponding lines for each sample), the app allows to manually reassign status, store results and send them to the server, where they can be queried by the test subjects. The app is provided as an R script; the code and some example data are available on GitHub at https://github.com/anders-biostat/lampplateanalysis.

Here, LinkedCharts turned out to be well suited to quickly develop the app, to continuously refine it while the assay was finalized, and to turn it into a production tool, well integrated into the testing campaign’s databases and result reporting services.

## Discussion

Interactivity has already proved itself to be useful for data visualization. Static plots can accommodate only a limited number of accents. Any change, no matter how small, requires one to make a new plot. And yet, it is crucial to look at the data from different angles and on different scales to find peculiar patterns and make sure that there are no hidden artifacts that can influence conclusions. Sometimes, even the visualize transition from one state to another is what helps to understand the data. Thus interactive visualization is helpful during the research and for presenting data to the scientific community.

The field of interactive visualization is an actively developing one with some already well known and established tools such as “shiny” (RStudio, Inc 2013) or “plotly”(Sievert 2020). Yet, possible benefits of interactivity are far from being explored. We have described LinkedCharts as a library for spontaneous and highly customizable visualization. A simple interface makes R/LinkedCharts suitable for generating minor on-the-fly apps for data exploration. It can be used by researchers with only basic experience in R or by bioinformaticians who are not willing to invest much time into an acquaintance with an unfamiliar visualization library. The linking mechanism is ensured by userdefined callback functions, allowing R/LinkedCharts to do without requirements for fixed data structures. There are no restrictions on the linking scheme. Every chart can be linked to any other or multiple ones. Any kind of backwards or partial linking is also possible.

We believe easy exploratory analysis to be the main niche for R/Linked charts. As we have shown, it provides users with a possibility to generate visualizations with the same effort as one generally puts into routine data digging and exploration. However, interactive apps are much more engaging than the static plots commonly used in the early stages of any project. When checking an idea or concern takes just a click, the researcher is more likely to go through the data thoroughly and, with this, maybe, save time on the further steps of the analysis.

Once the need to present final or intermediate results to colleagues arises, the same essential apps that were previously used as “quick and dirty” solutions can be prettified and shared by deploying the app on a server. The required changes for a R/LinkedCharts app to work on the server are minimal, and therefore there is no need to start from scratch. One can utilize the same scripts as personal drafts for exploration and as a basis for result presentation.

LinkedCharts is a JavaScript-based library, and unlike in existing alternatives, we gave users complete freedom to employ the functionality of various web solutions. For example, R/LinkedCharts can be embedded into any predefined HTML layout. Therefore, someone with experience in web design may provide the researcher with a custom HTML page with allocated containers for the chart and any predefined functionality made available by hundreds of JavaScript packages explicitly developed to ensure user interaction with a web page. Moreover, the “jrc” package on top of which R/LinkedCharts is built allows running any JavaScript code from the R session and any R code from the web page, providing endless possibilities for custom functionality and design. Such an app can also load locally stored scripts, images, stylesheets, etc.

The JavaScript basis of R/LinkedCharts offers an interface of its own that is also very simple and similar to the “rlc” syntax. Therefore a user who is familiar with JavaScript or willing to learn its essentials gets a way to convert R app into a stand-alone one fully contained within an HTML page. Such an app doesn’t require any side resources, can be shared by email between collaborators and opened in any browser. It does not need to have a constantly running R session and can be published on any hosting, including the most simple ones that do not allow to run other software in the background. An example of such stand-alone apps is the Supplementto this paper. There, one can also find examples of conversion from R apps to JavaScript ones. More information can be found in our online tutorials.

The structure of the *linked-charts*.*js* library and even principles of JavaScript as a language to manipulate DOM elements can allow an experienced user to customize not just the ecosystem in which LinkedCharts will be embedded but the charts themselves without a need to dive into the source code. One can go as far as defining custom types of charts (see https://anders-biostat.github.io/linked-charts/js/tutorials/layers.html for more details on that).

Overall, R/LinkedCharts serves two primary purposes: to facilitate data exploration and presentation. It offers an easy way to utilize interactivity for everyday research tasks. And it also provides the user with a possibility to fully employ the power of JavaScript for presenting the data. The latter aspect addresses more experienced users and thus has no limits for possible customization of a LinkedCharts app.

## Implementation

The JavaScript foundation of R/LinkedCharts is built on top of the D3 library (Bostock et al. 2011).

*linked-charts*.*js* is by itself a fully functional tool for interactive data visualization that can be used by those familiar with JavaScript to create standalone apps. The library is open-source and available on GitHub at https://github.com/anders-biostat/linked-charts. The minified version can be downloaded from https://github.com/anders-biostat/linked-charts/raw/master/lib/linked-charts.min.js, and its stylesheet is at https://github.com/anders-biostat/linked-charts/raw/master/lib/linked-charts.css.

The “jrc” package (Ovchinnikova and Anders 2020) package is used as a bridge between R and JavaScript. It allows one to run JavaScript code from an R session and vice versa. It also manages client connections to the app and is responsible for all the functionality necessary to make an R/LinkedCharts app public. “jrc” in turn is based mainly on “httpuv” (Cheng and Chang 2020) package to run a local server and ensure a WebSocket connection (Fette and Melnikov 2011).

R/LinkedCharts (“rlc” package) is an R (R Core Team 2019) interface to the JavaScript version of LinkedCharts. In addition to providing access to *linked-charts*.*js* functionality, it also ensures proper storing of charts and serving them to each connected client by extending “App” class of the “jrc” package. “rlc” is open source and is available on CRAN or GitHub https://github.com/anders-biostat/rlc.

## Supporting information

Supplemental Examples

## Acknowledgement

The authors acknowledge funding by the Deutsche Forschungsgemeinschaft via CRC 1366.

## Supplement

The interactive supplement to this paper can be found online at https://anders-biostat.github.io/lc-paper/ or offline in the zip file accompanying the paper. To see the supplement offline, unpack the zip file and use a web browser to open the file *index*.*html* contained within.

## Notes

https://anders-biostat.github.io/linked-charts/

### Competing Interest Statement

The authors have declared no competing interest.

