## Supplemental Examples for "R/LinkedCharts: A novel approach for simple but powerful interactive data analysis": JS_code_full.html

```
lc.scatter()
   .x(i => +iris[i].sepal_length)
   .y(i => +iris[i].petal_length)
   .size(i => +iris[i].sepal_width * 2)
   .colourValue(i => +iris[i].petal_width)
   .symbolValue(i => iris[i].species)
   .place();
```
