## Supplemental Examples for "R/LinkedCharts: A novel approach for simple but powerful interactive data analysis": JS_code_full.html

```
// a function to build a sigmoid curve
function get_curve (drug, cellLine, x) {
  var max = curves[drug][cellLine].max,
    min = curves[drug][cellLine].min,
    IC50 = curves[drug][cellLine].IC50,
    slope = curves[drug][cellLine].Slope,
    minConc = curves[drug][cellLine].minConc;

  return min + (max - min)/ 
    (1 + Math.pow(10, -(x - Math.log10(IC50/minConc)) * slope));
}

var selDrugs = [drugs[0], drugs[1]],
   selCellLine = cellLines[0];

lc.heatmap()
   .nrows(drugs.length)
   .ncols(drugs.length)
   .rowLabel(i => drugs[i])
   .colLabel(i => drugs[i])
   .set_paddings({left: 40, top: 40, right: 50})
   .showLegend(false)
   .clusterRows(true)
   .clusterCols(true)
   .value((row, col) => corMat[row][col])
   .on_click((row, col) => {
      selDrugs = [drugs[row], drugs[col]];
      dssplot.update();
      viability.update();
      plateHms[0].update();
      plateHms[1].update();
   })
  .place("#A1");

  var dssplot = lc.scatter()
   .width(300)
   .height(250)
   .title("DSS value")
   .x(i => scoreMat[i][drugs.indexOf(selDrugs[0])])
   .y(i => scoreMat[i][drugs.indexOf(selDrugs[1])])
   .domainX([0, 45])
   .domainY([0, 45])
   .set_paddings({bottom: 20})
   .colour(i => i == cellLines.indexOf(selCellLine) ? "black" : "grey")
   .axisTitleX(() => selDrugs[0])
   .axisTitleY(() => selDrugs[1])
   .label(i => cellLines[i])
   .on_click(d => {
      selCellLine = cellLines[d];
      dssplot.get_layer("layer0").updateElementStyle();
      viability.update();
      plateHms[0].update();
      plateHms[1].update();
   });       

//add another layer to the 'dssplot'
lc.xLine("line", dssplot)
   .lineFun(x => x)
   .place("#A2");

//here, data are transformed from an array in a nested format
//which easier and faster by the linked-charts.js to access
curves = lc.separateBy(curves, ["Drug", "CellLine"]);

var viability = lc.xLine()
   .width(300)
   .height(150)
   .title(() => selCellLine)
   .axisTitleX("Concentration order")
   .axisTitleY("Cell viability, %")
   .set_paddings({bottom: 20})
   .nelements(2)
   .addColourScaleToLegend(false)
   .lineFun((x, i) => get_curve(selDrugs[i], selCellLine, x))
   .colourValue(i => selDrugs[i]);

lc.scatter("points", viability)
   .nelements(10)
   .x(i => i % 5)
   .y(i => curves[selDrugs[+(i < 5)]][selCellLine]["D" + (i % 5 + 1)])
   .domainY([-10, 105])
   .on_mouseover(i => {
      let well = curves[selDrugs[+(i < 5)]][selCellLine]["W" + (i % 5 + 1)];
      plateHms[+(i < 5)].mark([well[0], well.slice(1)]);
   })
   .on_mouseout(() => {
      plateHms.forEach(el => el.mark("__clear__"));
   })    
   .colourValue(i => selDrugs[+(i < 5)])
   .legend.container("#A2")
   .legend.width(300)
   .place("#A2");

plates = lc.separateBy(plates, ["Barcode", "DWell"]);


//since heatmaps for readouts are so similar to each other
//it is easier to make them both within a "for" loop
//however, since we need the index variable inside the callback
//functions, we are using a "placeHeatmap" function that produces
//and returns a heatmap
var plateHms = [];
for(let i = 0; i < 2; i++)
   plateHms[i] = placeHeatmap(i);

function placeHeatmap(i) {
   return lc.heatmap()
      .width(300)
      .height(225)
      .showPanel(false)
      .title(() => "Plate " + curves[selDrugs[i]][selCellLine].Barcode)
      .titleSize(15)
      .paddings({top: 40, left: 15, bottom: 5, right: 5})
      .rowIds("ABCDEFGHIJKLMNOP".split(""))
      .colIds(d3.range(24).map(el => el + 1))
      .value((row, col) => 
         plates[curves[selDrugs[i]][selCellLine].Barcode][row+col].rawIntensity)
      .informText((row, col) => {
         let well = plates[curves[selDrugs[i]][selCellLine].Barcode][row + col];

         return "<b>" + well.ProductName + "</b><br>" + 
            "Concentration:" + well.Concentration + "<br>" +
            "Value: " + well.rawIntensity;
      })
      .showLegend(false)
      .place("#A3");   
}
```
