## Supplemental Examples for "R/LinkedCharts: A novel approach for simple but powerful interactive data analysis": JS_code_min.html

```
// a function to build a sigmoid curve
function get_curve ( drug, cellLine, x ){
  var max = curves[drug][cellLine].max,
    min = curves[drug][cellLine].min,
    IC50 = curves[drug][cellLine].IC50,
    slope = curves[drug][cellLine].Slope,
    minConc = curves[drug][cellLine].minConc;

  return min + (max - min)/ 
    (1 + Math.pow(10, -(x - Math.log10(IC50/minConc)) * slope));
}

selDrugs = [drugs[0], drugs[1]];
selCellLine = cellLines[0];

lc.heatmap()
  .nrows(drugs.length)
  .ncols(drugs.length)
  .rowLabel(i => drugs[i])
  .colLabel(i => drugs[i])
  .value((row, col) => corMat[row][col])
  .on_click((row, col) => {
    selDrugs = [drugs[row], drugs[col]];
    dssplot.update();
    viability.update();
    plateHms[0].update();
    plateHms[1].update();
  })
  .place("#A1");

var dssplot = lc.scatter()
   //sizes and paddings are, in fact, optional parameters
   //however, we leave them here to roughly keep the layout of the app
  .width(300)
  .height(250)
  .x(i => scoreMat[i][drugs.indexOf(selDrugs[0])])
  .y(i => scoreMat[i][drugs.indexOf(selDrugs[1])])
  .label(i => cellLines[i])
  .on_click(d => {
    selCellLine = cellLines[d];
    dssplot.get_layer("layer0").updateElementStyle();
    viability.update();
    plateHms[0].update();
    plateHms[1].update();
  })
  .place("#A2");    

//here, data are transformed from an array in a nested format
//which easier and faster by the linked-charts.js to access
curves = lc.separateBy(curves, ["Drug", "CellLine"]);

var viability = lc.scatter()
  .nelements(10)
  .x(i => i % 5)
  .y(i => curves[selDrugs[+(i < 5)]][selCellLine]["D" + (i % 5 + 1)])
  .colourValue(i => selDrugs[+(i < 5)]);

lc.xLine("line", viability)
  .width(300)
  .height(150)
  .nelements(2)
  .lineFun((x, i) => get_curve(selDrugs[i], selCellLine, x))
  .colourValue(i => selDrugs[i])
  .place("#A2");

plates = lc.separateBy(plates, ["Barcode", "DWell"]);

//since heatmaps for readouts are so similar to each other
//it is easier to make them both within a "for" loop
//however, since we need the index variable inside the callback
//functions, we are using a "placeHeatmap" function that produces
//and returns a heatmap
var plateHms = [];
for(let i = 0; i < 2; i++)
  plateHms[i] = placeHeatmap(i);

function placeHeatmap(i) {
  return lc.heatmap()
    .width(300)
    .height(225)
    .paddings({top: 40, left: 15, bottom: 5, right: 5})
    .rowIds("ABCDEFGHIJKLMNOP".split(""))
    .colIds(d3.range(24).map(el => el + 1))
    .value((row, col) => 
      plates[curves[selDrugs[i]][selCellLine].Barcode][row + col].rawIntensity)
    .informText((row, col) => {
      let well = plates[curves[selDrugs[i]][selCellLine].Barcode][row + col];
        return "<b>" + well.ProductName + "</b><br>" + 
          "Concentration:" + well.Concentration + "<br>" +
          "Value: " + well.rawIntensity;
    })
    .place("#A3");
}
```
