## Supplemental Examples for "R/LinkedCharts: A novel approach for simple but powerful interactive data analysis": R_code_min.html

```
library(tidyverse)

openPage(useViewer = FALSE, layout = "table1x3")

selDrugs <- colnames(scoreMatrix)[1:2]
selCellLine <- rownames(scoreMatrix)[1]

#this function calculates the sigmoid curve for a give drug and
#cell line based on the "curves" table
getCurve <- function(drug, cellLine, x) {
  row <- paste(cellLine, drug, sep = "_")
  curves[row, "min"] + (curves[row, "max"] - curves[row, "min"])/
    (1 + 10^(-(x - log10(curves[row, "IC50"]/curves[row, "minConc"])) * curves[row, "Slope"]))
}

lc_heatmap(value = cor(scoreMatrix),
           on_click = function(d) {
             selDrugs <<-  colnames(scoreMatrix)[d]
             updateCharts()
           },
           place = "A1")

lc_scatter(dat(
  x = scoreMatrix[, selDrugs[1]],
  y = scoreMatrix[, selDrugs[2]]), 
  width = 300,
  height = 250,
  paddings = list(bottom = 20),
  label = rownames(scoreMatrix),
  on_click = function(d) {
    selCellLine <<- rownames(scoreMatrix)[d]
    updateCharts()
  },
  place = "A2"
)

lc_scatter(dat(
  x = c(0:4, 0:4),
  y = c(unlist(curves[paste(selCellLine, selDrugs[1], sep = "_"), paste0("D", 1:5)]),
        unlist(curves[paste(selCellLine, selDrugs[2], sep = "_"), paste0("D", 1:5)])),
  colourValue = rep(selDrugs, each = 5)),
  width = 300,
  height = 150,
# Note that to put several charts can be put into the same
# container, but they have to get different IDs.
# Chart with the same ID will either replace the existing one,
# or will be added as a new layer.  
  place = "A2", chartId = "viability"
)

x <- seq(0, 4, length.out = 50)
lc_line(dat(
  x = x,
  y = cbind(getCurve(selDrugs[1], selCellLine, x),
            getCurve(selDrugs[2], selCellLine, x)),
  colourValue = selDrugs),
# That is how a new layer can be added to a chart  
  chartId = "viability", addLayer = TRUE
)

# This part is quite different from the JavaScript version.
# In JavaScript, LinkedCharts query for each value separately
# and thus one can use a function that returns the value based 
# the ID of the element. In R, a full matrix (for heatmaps) or
# vector has to be provided. The following two functions make such matrices,
# based on the long tibble "plates". 

getInfo <- function(drug) {
  plates %>% 
    filter(Barcode == barcodes[paste(selCellLine, drug, sep = "_")]) %>%
    transmute(DRow, DCol, text = str_c(
      "<b>", ProductName, "</b><br>",
      "Concentration: ", str_replace_na(Concentration), "<br>",
      "Value: ", rawIntensity
    )) %>%
    pivot_wider(names_from = DCol, values_from = text) %>%
    select(-DRow) %>%
    as.matrix()
}

getPlate <- function(drug) {
  plates %>% 
    filter(Barcode == barcodes[paste(selCellLine, drug, sep = "_")]) %>%
    select(DRow, DCol, rawIntensity) %>%
    pivot_wider(names_from = DCol, values_from = rawIntensity) %>%
    select(-DRow) %>%
    as.matrix()
}

placeHeatmap <- function(i) {
  lc_heatmap(dat(
    value = getPlate(selDrugs[ind]),
    informText = getInfo(selDrugs[ind])),
    width = 300,
    height = 225,
    paddings = list(top = 40, left = 15, bottom = 5, right = 5),
    rowLabel = LETTERS[1:16],
    place = "A3", chartId = paste0("plate", i),
# Due do the usage of non-standard evaluation, "rlc" cannot access
# the "i" argument from inside the "dat" function. However, one can
# specify a list of variables for each give chart with the "with" argument    
    with = list(ind = i)
  )
}

for(i in 1:2)
  placeHeatmap(i)
```
