## Supplemental Examples for "R/LinkedCharts: A novel approach for simple but powerful interactive data analysis": rawData.js

```
curves = [{"CellLine":"Pa16C","Drug":"PF_04691502","minConc":1,"D1":1.03,"D2":-4.31,"D3":3.72,"D4":61.31,"D5":92.33,"IC50":672.29,"Slope":1.73,"max":92.33,"min":0,"Barcode":"H1FO4B-N1177","W5":"B2","W4":"C2","W3":"D2","W2":"E2","W1":"F2"},{"CellLine":"Pa16C","Drug":"SNS_032","minConc":1,"D1":9.57,"D2":14.12,"D3":59.43,"D4":72.99,"D5":75.3,"IC50":48.5,"Slope":1.57,"max":75.3,"min":9.25,"Barcode":"H1FO4B-N1177","W5":"G9","W4":"H9","W3":"I9","W2":"J9","W1":"K9"},{"CellLine":"Pa16C","Drug":"Alvocidib","minConc":1,"D1":10.64,"D2":8.94,"D3":23.46,"D4":74.14,"D5":79.59,"IC50":225.23,"Slope":1.67,"max":79.59,"min":8.94,"Barcode":"H1FO4B-N1177","W5":"P9","W4":"O9","W3":"N9","W2":"M9","W1":"L9"},{"CellLine":"Pa16C","Drug":"KX2_391","minConc":1,"D1":14.49,"D2":14.07,"D3":57.47,"D4":50.01,"D5":52.12,"IC50":30.02,"Slope":2.5,"max":57.47,"min":13.04,"Barcode":"H1FO4B-N1177","W5":"F17","W4":"G17","W3":"H17","W2":"I17","W1":"J17"},{"CellLine":"Pa16C","Drug":"AZD8055","minConc":1,"D1":4.62,"D2":3.87,"D3":32.38,"D4":45.46,"D5":52.79,"IC50":80.04,"Slope":1.06,"max":52.79,"min":1.92,"Barcode":"H1FO4B-N1177","W5":"G23","W4":"H23","W3":"I23","W2":"J23","W1":"K23"},{"CellLine":"Pa16C","Drug":"PF_03758309","minConc":1,"D1":3.55,"D2":11.83,"D3":25.15,"D4":53.58,"D5":88.14,"IC50":457.59,"Slope":0.76,"max":88.14,"min":3.55,"Barcode":"H1FO4B-N2177","W5":"P5","W4":"O5","W3":"N5","W2":"M5","W1":"L5"},{"CellLine":"Pa16C","Drug":"AZD_8330","minConc":1,"D1":8.22,"D2":24.69,"D3":45.07,"D4":49.39,"D5":50.08,"IC50":12.88,"Slope":1.01,"max":50.08,"min":5.05,"Barcode":"H1FO4B-N2177","W5":"F6","W4":"H6","W3":"I6","W2":"J6","W1":"K6"},{"CellLine":"Pa16C","Drug":"Rigosertib","minConc":1,"D1":4.05,"D2":2.43,"D3":58.26,"D4":49.98,"D5":48.27,"IC50":30.88,"Slope":2.5,"max":58.26,"min":1.79,"Barcode":"H1FO4B-N2177","W5":"F8","W4":"G8","W3":"H8","W2":"J8","W1":"K8"},{"CellLine":"Pa16C","Drug":"Omipalisib","minConc":0.1,"D1":4.96,"D2":-2.87,"D3":1.46,"D4":54.88,"D5":57.1,"IC50":32.86,"Slope":2.5,"max":57.1,"min":0,"Barcode":"H1FO4B-N2177","W5":"P17","W4":"O17","W3":"N17","W2":"M17","W1":"K17"},{"CellLine":"Pa16C","Drug":"UCN_01","minConc":1,"D1":2.18,"D2":-4.34,"D3":21.57,"D4":44.16,"D5":89.54,"IC50":732.98,"Slope":0.86,"max":89.54,"min":0,"Barcode":"H1FO4B-N2177","W5":"P21","W4":"O21","W3":"N21","W2":"M21","W1":"L21"},{"CellLine":"Pa16C","Drug":"Dinaciclib","minConc":0.1,"D1":5.22,"D2":-5.82,"D3":10.19,"D4":77.53,"D5":77.94,"IC50":21.16,"Slope":2.5,"max":77.94,"min":0,"Barcode":"H1FO4B-N3177","W5":"A3","W4":"B3","W3":"C3","W2":"D3","W1":"E3"},{"CellLine":"Pa16C","Drug":"NVP_BGT226","minConc":0.1,"D1":6.55,"D2":10.53,"D3":27.42,"D4":41.05,"D5":95.55,"IC50":104.47,"Slope":0.87,"max":95.55,"min":6.55,"Barcode":"H1FO4B-N3177","W5":"G12","W4":"H12","W3":"I12","W2":"J12","W1":"K12"},{"CellLine":"Pa16C","Drug":"Gedatolisib","minConc":0.1,"D1":0.41,"D2":-1.81,"D3":9.36,"D4":42.77,"D5":62.78,"IC50":49.84,"Slope":1.16,"max":62.78,"min":0,"Barcode":"H1FO4B-N3177","W5":"P12","W4":"O12","W3":"N12","W2":"M12","W1":"L12"},{"CellLine":"Pa16C","Drug":"PKI_402","minConc":0.25,"D1":13.25,"D2":10.72,"D3":15.08,"D4":45.53,"D5":59.95,"IC50":133.18,"Slope":1.42,"max":59.95,"min":10.72,"Barcode":"H1FO4B-N3177","W5":"A16","W4":"C16","W3":"D16","W2":"E16","W1":"F16"},{"CellLine":"Pa16C","Drug":"AT7519","minConc":1,"D1":4.25,"D2":3.91,"D3":5.93,"D4":76.08,"D5":74.02,"IC50":288.45,"Slope":2.5,"max":76.08,"min":3.17,"Barcode":"H1FO4B-N3177","W5":"F22","W4":"G22","W3":"H22","W2":"I22","W1":"J22"},{"CellLine":"Pa16C","Drug":"GSK_461364","minConc":1,"D1":6.96,"D2":24.69,"D3":22.32,"D4":24.25,"D5":50.77,"IC50":329.01,"Slope":0.39,"max":50.77,"min":6.96,"Barcode":"H1FO4B-N4177","W5":"A6","W4":"B6","W3":"C6","W2":"D6","W1":"E6"},{"CellLine":"Pa16C","Drug":"Daporinad","minConc":0.1,"D1":7.46,"D2":2.61,"D3":5.45,"D4":33.45,"D5":33.12,"IC50":24.04,"Slope":2.5,"max":33.45,"min":2.61,"Barcode":"H1FO4B-N4177","W5":"A14","W4":"B14","W3":"C14","W2":"D14","W1":"E14"},{"CellLine":"Pa16C","Drug":"Mepacrine","minConc":5,"D1":0.5,"D2":1.24,"D3":1.82,"D4":61.86,"D5":101.1,"IC50":4065.9,"Slope":2.31,"max":100,"min":0.5,"Barcode":"H1FO4B-N4177","W5":"P20","W4":"O20","W3":"N20","W2":"M20","W1":"L20"},{"CellLine":"Pa16C","Drug":"Camptothecin","minConc":0.5,"D1":11.99,"D2":9.16,"D3":28.61,"D4":63.89,"D5":88.18,"IC50":186.97,"Slope":0.93,"max":88.18,"min":9.16,"Barcode":"H1FO4B-N5177","W5":"O3","W4":"N3","W3":"M3","W2":"L3","W1":"K3"},{"CellLine":"Pa16C","Drug":"Omacetaxine","minConc":1,"D1":7.99,"D2":5.58,"D3":28.54,"D4":83.6,"D5":91.07,"IC50":198.92,"Slope":1.47,"max":91.07,"min":5.58,"Barcode":"H1FO4B-N5177","W5":"A5","W4":"B5","W3":"C5","W2":"E5","W1":"F5"},{"CellLine":"Pa16C","Drug":"Vinorelbine","minConc":1,"D1":2.06,"D2":50.67,"D3":59.15,"D4":54.17,"D5":86.56,"IC50":25.15,"Slope":0.41,"max":86.56,"min":0,"Barcode":"H1FO4B-N5177","W5":"P14","W4":"O14","W3":"N14","W2":"M14","W1":"L14"},{"CellLine":"Pa16C","Drug":"Docetaxel","minConc":0.1,"D1":7.23,"D2":13.16,"D3":51.93,"D4":55.75,"D5":50.21,"IC50":2.77,"Slope":1.92,"max":55.75,"min":7.14,"Barcode":"H1FO4B-N5177","W5":"P17","W4":"O17","W3":"N17","W2":"M17","W1":"K17"},{"CellLine":"Pa16C","Drug":"Carfilzomib","minConc":0.1,"D1":3.33,"D2":-0.46,"D3":92.91,"D4":99.27,"D5":101.44,"IC50":3.88,"Slope":2.5,"max":100,"min":0,"Barcode":"H1FO4B-N5177","W5":"A18","W4":"B18","W3":"C18","W2":"D18","W1":"E18"},{"CellLine":"Pa16C","Drug":"Patupilone","minConc":0.1,"D1":13.19,"D2":8.91,"D3":44.62,"D4":51.69,"D5":49.7,"IC50":5.26,"Slope":2.5,"max":51.69,"min":8.91,"Barcode":"H1FO4B-N5177","W5":"F18","W4":"G18","W3":"H18","W2":"I18","W1":"J18"},{"CellLine":"Pa16C","Drug":"Plicamycin","minConc":1,"D1":11.85,"D2":8.82,"D3":26.14,"D4":88.05,"D5":91.02,"IC50":193.44,"Slope":2,"max":91.02,"min":8.82,"Barcode":"H1FO4B-N5177","W5":"P18","W4":"N18","W3":"M18","W2":"L18","W1":"K18"},{"CellLine":"Pa16C","Drug":"Vincristine","minConc":0.1,"D1":6.21,"D2":-1.85,"D3":55.57,"D4":52.32,"D5":47.65,"IC50":3.35,"Slope":2.5,"max":55.57,"min":0,"Barcode":"H1FO4B-N5177","W5":"A19","W4":"B19","W3":"C19","W2":"D19","W1":"E19"},{"CellLine":"Pa16C","Drug":"Dactinomycin","minConc":0.1,"D1":9.36,"D2":22.66,"D3":35.86,"D4":90.29,"D5":81.62,"IC50":15.08,"Slope":1.52,"max":90.29,"min":9.36,"Barcode":"H1FO4B-N5177","W5":"P19","W4":"O19","W3":"N19","W2":"M19","W1":"L19"},{"CellLine":"Pa16C","Drug":"Vinblastine","minConc":0.1,"D1":4.31,"D2":0.3,"D3":46.19,"D4":54.79,"D5":47.83,"IC50":5.15,"Slope":2.5,"max":54.79,"min":0.3,"Barcode":"H1FO4B-N5177","W5":"A21","W4":"B21","W3":"C21","W2":"D21","W1":"E21"},{"CellLine":"Pa16C","Drug":"Paclitaxel","minConc":0.1,"D1":9.47,"D2":3.54,"D3":49.85,"D4":59.62,"D5":57.74,"IC50":5.39,"Slope":2.5,"max":59.62,"min":3.54,"Barcode":"H1FO4B-N5177","W5":"F21","W4":"G21","W3":"H21","W2":"I21","W1":"J21"},{"CellLine":"Pa16C","Drug":"Oprozomib","minConc":0.25,"D1":-0.33,"D2":-5.47,"D3":42.85,"D4":97.06,"D5":102.1,"IC50":28.1,"Slope":2.5,"max":100,"min":0,"Barcode":"H1FO4B-N5177","W5":"A22","W4":"B22","W3":"C22","W2":"D22","W1":"E22"},{"CellLine":"Pa16C","Drug":"Bortezomib","minConc":0.1,"D1":-0.24,"D2":3.55,"D3":99.86,"D4":99.23,"D5":99.98,"IC50":2.91,"Slope":2.5,"max":99.98,"min":0,"Barcode":"H1FO4B-N5177","W5":"F22","W4":"G22","W3":"H22","W2":"I22","W1":"J22"},{"CellLine":"Pa16C","Drug":"Filanesib","minConc":0.1,"D1":3.72,"D2":-0.57,"D3":51.84,"D4":52.95,"D5":53.97,"IC50":3.55,"Slope":2.5,"max":53.97,"min":0,"Barcode":"H1FO4B-N5177","W5":"G23","W4":"H23","W3":"I23","W2":"J23","W1":"K23"},{"CellLine":"Pa16C","Drug":"CUDC_907","minConc":1,"D1":10.19,"D2":6.47,"D3":64.46,"D4":97.96,"D5":99.89,"IC50":81.2,"Slope":2.36,"max":99.89,"min":6.47,"Barcode":"H1FO4B-N6177","W5":"F17","W4":"G17","W3":"H17","W2":"I17","W1":"J17"},{"CellLine":"Pa16C","Drug":"YM155","minConc":1,"D1":15.1,"D2":27.4,"D3":98.58,"D4":100.45,"D5":103.02,"IC50":20.29,"Slope":2.5,"max":100,"min":15.04,"Barcode":"H1FO4B-N6177","W5":"F18","W4":"G18","W3":"H18","W2":"I18","W1":"J18"},{"CellLine":"Pa16C","Drug":"AT101","minConc":10,"D1":5.12,"D2":2.47,"D3":10.8,"D4":40.91,"D5":103.33,"IC50":11847.55,"Slope":2.5,"max":100,"min":2.47,"Barcode":"H1FO4B-N6177","W5":"A21","W4":"B21","W3":"C21","W2":"D21","W1":"E21"},{"CellLine":"Pa16C","Drug":"Selinexor","minConc":1,"D1":7.39,"D2":14.29,"D3":32.29,"D4":62.55,"D5":58.26,"IC50":106.36,"Slope":1.16,"max":62.55,"min":7.39,"Barcode":"H1FO4B-N6177","W5":"P21","W4":"O21","W3":"N21","W2":"M21","W1":"L21"},{"CellLine":"Pa16C","Drug":"BIIB021","minConc":1,"D1":10.3,"D2":6,"D3":46.18,"D4":55.67,"D5":65.82,"IC50":66.58,"Slope":1.52,"max":65.82,"min":6,"Barcode":"H1FO4B-N7177","W5":"P13","W4":"N13","W3":"M13","W2":"L13","W1":"K13"},{"CellLine":"Pa16C","Drug":"Tanespimycin","minConc":1,"D1":10.44,"D2":5.58,"D3":38.34,"D4":58.68,"D5":58.3,"IC50":82.65,"Slope":2.5,"max":58.68,"min":5.58,"Barcode":"H1FO4B-N7177","W5":"A14","W4":"B14","W3":"C14","W2":"D14","W1":"E14"},{"CellLine":"Pa16C","Drug":"CUDC_305","minConc":1,"D1":16.06,"D2":8.51,"D3":29.88,"D4":51.35,"D5":72.72,"IC50":310.71,"Slope":0.77,"max":72.72,"min":8.51,"Barcode":"H1FO4B-N7177","W5":"A15","W4":"B15","W3":"C15","W2":"D15","W1":"F15"},{"CellLine":"Pa16C","Drug":"Luminespib","minConc":0.1,"D1":5.81,"D2":10.08,"D3":27.11,"D4":60.92,"D5":58.25,"IC50":12.91,"Slope":1.7,"max":60.92,"min":5.81,"Barcode":"H1FO4B-N7177","W5":"G15","W4":"H15","W3":"I15","W2":"J15","W1":"K15"},{"CellLine":"Pa16C","Drug":"Ganetespib","minConc":0.1,"D1":12.89,"D2":13.85,"D3":21.37,"D4":53.95,"D5":51.85,"IC50":17.09,"Slope":2.5,"max":53.95,"min":12.89,"Barcode":"H1FO4B-N7177","W5":"A16","W4":"C16","W3":"D16","W2":"E16","W1":"F16"},{"CellLine":"Pa16C","Drug":"Stattic","minConc":5,"D1":10.05,"D2":6.58,"D3":5.16,"D4":92.4,"D5":103.4,"IC50":1991.11,"Slope":2.5,"max":100,"min":5.16,"Barcode":"H1FO4B-N7177","W5":"F21","W4":"G21","W3":"H21","W2":"I21","W1":"J21"},{"CellLine":"Pa16C","Drug":"ONX_0914","minConc":1,"D1":2.93,"D2":6.26,"D3":39.04,"D4":101.26,"D5":102.06,"IC50":123.16,"Slope":2.5,"max":100,"min":2.93,"Barcode":"H1FO4B-N7177","W5":"G23","W4":"H23","W3":"I23","W2":"J23","W1":"K23"},{"CellLine":"Pa16C","Drug":"SN_38","minConc":1,"D1":4.8,"D2":33.06,"D3":47.83,"D4":74.77,"D5":83.23,"IC50":36.36,"Slope":0.58,"max":83.23,"min":0,"Barcode":"H1FO4B-N8177","W5":"G2","W4":"H2","W3":"I2","W2":"J2","W1":"K2"},{"CellLine":"Pa16C","Drug":"Cabazitaxel","minConc":0.1,"D1":6.06,"D2":5.93,"D3":47.76,"D4":51.85,"D5":45.48,"IC50":4.03,"Slope":2.5,"max":51.85,"min":5.33,"Barcode":"H1FO4B-N8177","W5":"A3","W4":"B3","W3":"C3","W2":"D3","W1":"E3"},{"CellLine":"Pa16C","Drug":"OTSSP167","minConc":0.1,"D1":10.71,"D2":6.67,"D3":11.13,"D4":79.42,"D5":86.07,"IC50":34.74,"Slope":2.26,"max":86.07,"min":6.67,"Barcode":"H1FO4B-N8177","W5":"A5","W4":"B5","W3":"C5","W2":"E5","W1":"F5"},{"CellLine":"Pa16C","Drug":"Prexasertib","minConc":1,"D1":10.71,"D2":8.45,"D3":2.7,"D4":6.83,"D5":28.41,"IC50":1927.44,"Slope":2.5,"max":28.41,"min":2.7,"Barcode":"H1FO4B-N8177","W5":"P6","W4":"O6","W3":"N6","W2":"M6","W1":"L6"},{"CellLine":"Pa16C","Drug":"Romidepsin","minConc":0.1,"D1":13.69,"D2":30.47,"D3":93,"D4":99.28,"D5":100.04,"IC50":2.31,"Slope":1.66,"max":100,"min":13.22,"Barcode":"H1FO4B-N8177","W5":"F18","W4":"G18","W3":"H18","W2":"I18","W1":"J18"},{"CellLine":"Pa16C","Drug":"Sabutoclax","minConc":2.5,"D1":12.1,"D2":14.6,"D3":28.09,"D4":42.85,"D5":87.66,"IC50":2606.88,"Slope":0.93,"max":87.66,"min":12.1,"Barcode":"H1FO4B-N8177","W5":"F19","W4":"G19","W3":"I19","W2":"J19","W1":"K19"},{"CellLine":"Pa16C","Drug":"BCI","minConc":5,"D1":6.43,"D2":-0.54,"D3":-3.94,"D4":94.84,"D5":99.99,"IC50":1912.18,"Slope":2.5,"max":99.99,"min":0,"Barcode":"H1FO4B-N8177","W5":"A22","W4":"B22","W3":"C22","W2":"D22","W1":"E22"},{"CellLine":"Pa14C","Drug":"PF_04691502","minConc":1,"D1":7.02,"D2":12.14,"D3":10.72,"D4":56.38,"D5":77.69,"IC50":592.18,"Slope":1.61,"max":77.69,"min":7.02,"Barcode":"H1FO4B-N1029","W5":"B2","W4":"C2","W3":"D2","W2":"E2","W1":"F2"},{"CellLine":"Pa14C","Drug":"SNS_032","minConc":1,"D1":1.82,"D2":10.43,"D3":63.25,"D4":81.49,"D5":79.15,"IC50":42.65,"Slope":1.44,"max":81.49,"min":1.52,"Barcode":"H1FO4B-N1029","W5":"G9","W4":"H9","W3":"I9","W2":"J9","W1":"K9"},{"CellLine":"Pa14C","Drug":"Alvocidib","minConc":1,"D1":-3.63,"D2":-0.68,"D3":21.28,"D4":80.18,"D5":85.3,"IC50":192.42,"Slope":1.69,"max":85.3,"min":0,"Barcode":"H1FO4B-N1029","W5":"P9","W4":"O9","W3":"N9","W2":"M9","W1":"L9"},{"CellLine":"Pa14C","Drug":"KX2_391","minConc":1,"D1":4.51,"D2":14.32,"D3":26.59,"D4":34.93,"D5":49.91,"IC50":82.37,"Slope":0.49,"max":49.91,"min":0,"Barcode":"H1FO4B-N1029","W5":"F17","W4":"G17","W3":"H17","W2":"I17","W1":"J17"},{"CellLine":"Pa14C","Drug":"AZD8055","minConc":1,"D1":-1.78,"D2":9.85,"D3":17.26,"D4":37.11,"D5":39.73,"IC50":94.2,"Slope":0.79,"max":39.73,"min":0,"Barcode":"H1FO4B-N1029","W5":"G23","W4":"H23","W3":"I23","W2":"J23","W1":"K23"},{"CellLine":"Pa14C","Drug":"PF_03758309","minConc":1,"D1":5.47,"D2":20.47,"D3":40.26,"D4":68.26,"D5":93.77,"IC50":180.67,"Slope":0.64,"max":93.77,"min":4.77,"Barcode":"H1FO4B-N2029","W5":"P5","W4":"O5","W3":"N5","W2":"M5","W1":"L5"},{"CellLine":"Pa14C","Drug":"AZD_8330","minConc":1,"D1":11.74,"D2":22.91,"D3":33.52,"D4":36.79,"D5":31.07,"IC50":8.16,"Slope":0.79,"max":36.79,"min":6.92,"Barcode":"H1FO4B-N2029","W5":"F6","W4":"H6","W3":"I6","W2":"J6","W1":"K6"},{"CellLine":"Pa14C","Drug":"Rigosertib","minConc":1,"D1":11.72,"D2":14.59,"D3":39.55,"D4":46.4,"D5":47.28,"IC50":44.59,"Slope":1.58,"max":47.28,"min":11.58,"Barcode":"H1FO4B-N2029","W5":"F8","W4":"G8","W3":"H8","W2":"J8","W1":"K8"},{"CellLine":"Pa14C","Drug":"Omipalisib","minConc":0.1,"D1":6.67,"D2":-0.51,"D3":8.36,"D4":30.1,"D5":41.97,"IC50":39.46,"Slope":1.06,"max":41.97,"min":0,"Barcode":"H1FO4B-N2029","W5":"P17","W4":"O17","W3":"N17","W2":"M17","W1":"K17"},{"CellLine":"Pa14C","Drug":"UCN_01","minConc":1,"D1":9.88,"D2":12.11,"D3":19.93,"D4":37.86,"D5":87.9,"IC50":1389.18,"Slope":1.57,"max":87.9,"min":9.88,"Barcode":"H1FO4B-N2029","W5":"P21","W4":"O21","W3":"N21","W2":"M21","W1":"L21"},{"CellLine":"Pa14C","Drug":"Dinaciclib","minConc":0.1,"D1":3.3,"D2":-2.59,"D3":68.91,"D4":82.77,"D5":83.6,"IC50":5.45,"Slope":2.5,"max":83.6,"min":0,"Barcode":"H1FO4B-N3029","W5":"A3","W4":"B3","W3":"C3","W2":"D3","W1":"E3"},{"CellLine":"Pa14C","Drug":"NVP_BGT226","minConc":0.1,"D1":-1.17,"D2":5.06,"D3":21.58,"D4":39.51,"D5":55.53,"IC50":21.6,"Slope":0.73,"max":55.53,"min":0,"Barcode":"H1FO4B-N3029","W5":"G12","W4":"H12","W3":"I12","W2":"J12","W1":"K12"},{"CellLine":"Pa14C","Drug":"Gedatolisib","minConc":0.1,"D1":-10.85,"D2":-10.83,"D3":5.75,"D4":36.72,"D5":51.32,"IC50":51.19,"Slope":1.42,"max":51.32,"min":0,"Barcode":"H1FO4B-N3029","W5":"P12","W4":"O12","W3":"N12","W2":"M12","W1":"L12"},{"CellLine":"Pa14C","Drug":"PKI_402","minConc":0.25,"D1":9.62,"D2":14.78,"D3":4.91,"D4":34.01,"D5":57.54,"IC50":229.62,"Slope":2.5,"max":57.54,"min":4.91,"Barcode":"H1FO4B-N3029","W5":"A16","W4":"C16","W3":"D16","W2":"E16","W1":"F16"},{"CellLine":"Pa14C","Drug":"AT7519","minConc":1,"D1":3.73,"D2":5.49,"D3":2.62,"D4":78.05,"D5":80.46,"IC50":340.11,"Slope":2.5,"max":80.46,"min":2.62,"Barcode":"H1FO4B-N3029","W5":"F22","W4":"G22","W3":"H22","W2":"I22","W1":"J22"},{"CellLine":"Pa14C","Drug":"GSK_461364","minConc":1,"D1":8.84,"D2":18.07,"D3":25.72,"D4":16.29,"D5":63.27,"IC50":2059.39,"Slope":2.5,"max":63.27,"min":8.84,"Barcode":"H1FO4B-N4029","W5":"A6","W4":"B6","W3":"C6","W2":"D6","W1":"E6"},{"CellLine":"Pa14C","Drug":"Daporinad","minConc":0.1,"D1":-2.31,"D2":-4.96,"D3":6.25,"D4":27.77,"D5":45.47,"IC50":64.19,"Slope":1.17,"max":45.47,"min":0,"Barcode":"H1FO4B-N4029","W5":"A14","W4":"B14","W3":"C14","W2":"D14","W1":"E14"},{"CellLine":"Pa14C","Drug":"Mepacrine","minConc":5,"D1":-0.2,"D2":-1.64,"D3":0.69,"D4":49.33,"D5":96.95,"IC50":4927.19,"Slope":2.44,"max":96.95,"min":0,"Barcode":"H1FO4B-N4029","W5":"P20","W4":"O20","W3":"N20","W2":"M20","W1":"L20"},{"CellLine":"Pa14C","Drug":"Camptothecin","minConc":0.5,"D1":8.21,"D2":6.45,"D3":16.41,"D4":48.88,"D5":86.75,"IC50":414.25,"Slope":1.13,"max":86.75,"min":6.45,"Barcode":"H1FO4B-N5029","W5":"O3","W4":"N3","W3":"M3","W2":"L3","W1":"K3"},{"CellLine":"Pa14C","Drug":"Omacetaxine","minConc":1,"D1":9.5,"D2":8.79,"D3":48.77,"D4":76.94,"D5":90.05,"IC50":109.31,"Slope":0.94,"max":90.05,"min":5.31,"Barcode":"H1FO4B-N5029","W5":"A5","W4":"B5","W3":"C5","W2":"E5","W1":"F5"},{"CellLine":"Pa14C","Drug":"Vinorelbine","minConc":1,"D1":-4.43,"D2":16.72,"D3":32.58,"D4":45.84,"D5":74.02,"IC50":191.78,"Slope":0.56,"max":74.02,"min":0,"Barcode":"H1FO4B-N5029","W5":"P14","W4":"O14","W3":"N14","W2":"M14","W1":"L14"},{"CellLine":"Pa14C","Drug":"Docetaxel","minConc":0.1,"D1":-0.59,"D2":5.05,"D3":18.9,"D4":32.39,"D5":27.42,"IC50":6.56,"Slope":1,"max":32.39,"min":0,"Barcode":"H1FO4B-N5029","W5":"P17","W4":"O17","W3":"N17","W2":"M17","W1":"K17"},{"CellLine":"Pa14C","Drug":"Carfilzomib","minConc":0.1,"D1":3.08,"D2":-2.95,"D3":44.33,"D4":97.08,"D5":100.93,"IC50":11.11,"Slope":2.21,"max":100,"min":0,"Barcode":"H1FO4B-N5029","W5":"A18","W4":"B18","W3":"C18","W2":"D18","W1":"E18"},{"CellLine":"Pa14C","Drug":"Patupilone","minConc":0.1,"D1":4.64,"D2":12.68,"D3":23.26,"D4":35.38,"D5":33.09,"IC50":3.85,"Slope":0.75,"max":35.38,"min":3.08,"Barcode":"H1FO4B-N5029","W5":"F18","W4":"G18","W3":"H18","W2":"I18","W1":"J18"},{"CellLine":"Pa14C","Drug":"Plicamycin","minConc":1,"D1":3.69,"D2":2.52,"D3":1.09,"D4":69.26,"D5":90.65,"IC50":629.77,"Slope":2.5,"max":90.65,"min":1.09,"Barcode":"H1FO4B-N5029","W5":"P18","W4":"N18","W3":"M18","W2":"L18","W1":"K18"},{"CellLine":"Pa14C","Drug":"Vincristine","minConc":0.1,"D1":3.94,"D2":-2.51,"D3":24.4,"D4":46.17,"D5":44.77,"IC50":9.57,"Slope":2.5,"max":46.17,"min":0,"Barcode":"H1FO4B-N5029","W5":"A19","W4":"B19","W3":"C19","W2":"D19","W1":"E19"},{"CellLine":"Pa14C","Drug":"Dactinomycin","minConc":0.1,"D1":-3.14,"D2":8.74,"D3":25.51,"D4":90.27,"D5":89.54,"IC50":15.18,"Slope":2.21,"max":90.27,"min":0,"Barcode":"H1FO4B-N5029","W5":"P19","W4":"O19","W3":"N19","W2":"M19","W1":"L19"},{"CellLine":"Pa14C","Drug":"Vinblastine","minConc":0.1,"D1":5.47,"D2":-4.7,"D3":20.73,"D4":42.06,"D5":45.3,"IC50":10.74,"Slope":2.5,"max":45.3,"min":0,"Barcode":"H1FO4B-N5029","W5":"A21","W4":"B21","W3":"C21","W2":"D21","W1":"E21"},{"CellLine":"Pa14C","Drug":"Paclitaxel","minConc":0.1,"D1":5.43,"D2":6.53,"D3":16.75,"D4":30.94,"D5":34.96,"IC50":15.65,"Slope":1.03,"max":34.96,"min":5.11,"Barcode":"H1FO4B-N5029","W5":"F21","W4":"G21","W3":"H21","W2":"I21","W1":"J21"},{"CellLine":"Pa14C","Drug":"Oprozomib","minConc":0.25,"D1":2.69,"D2":-0.82,"D3":20.29,"D4":89.13,"D5":99.91,"IC50":61.77,"Slope":1.52,"max":99.91,"min":0,"Barcode":"H1FO4B-N5029","W5":"A22","W4":"B22","W3":"C22","W2":"D22","W1":"E22"},{"CellLine":"Pa14C","Drug":"Bortezomib","minConc":0.1,"D1":-0.18,"D2":5.8,"D3":81.13,"D4":96.9,"D5":94.73,"IC50":4.24,"Slope":1.91,"max":96.9,"min":0,"Barcode":"H1FO4B-N5029","W5":"F22","W4":"G22","W3":"H22","W2":"I22","W1":"J22"},{"CellLine":"Pa14C","Drug":"Filanesib","minConc":0.1,"D1":-2.03,"D2":3.4,"D3":21.9,"D4":21.49,"D5":24.73,"IC50":3.02,"Slope":1.68,"max":24.73,"min":0,"Barcode":"H1FO4B-N5029","W5":"G23","W4":"H23","W3":"I23","W2":"J23","W1":"K23"},{"CellLine":"Pa14C","Drug":"CUDC_907","minConc":1,"D1":-0.01,"D2":5.42,"D3":56.13,"D4":97.7,"D5":99.46,"IC50":82.72,"Slope":1.42,"max":99.46,"min":0,"Barcode":"H1FO4B-N6029","W5":"F17","W4":"G17","W3":"H17","W2":"I17","W1":"J17"},{"CellLine":"Pa14C","Drug":"YM155","minConc":1,"D1":0.88,"D2":13.49,"D3":89.01,"D4":103.28,"D5":101.55,"IC50":30,"Slope":1.74,"max":100,"min":0.68,"Barcode":"H1FO4B-N6029","W5":"F18","W4":"G18","W3":"H18","W2":"I18","W1":"J18"},{"CellLine":"Pa14C","Drug":"AT101","minConc":10,"D1":4.11,"D2":-3.86,"D3":5.67,"D4":48.19,"D5":103.95,"IC50":10287.02,"Slope":2.27,"max":100,"min":0,"Barcode":"H1FO4B-N6029","W5":"A21","W4":"B21","W3":"C21","W2":"D21","W1":"E21"},{"CellLine":"Pa14C","Drug":"Selinexor","minConc":1,"D1":4.06,"D2":-6.04,"D3":11.24,"D4":40.41,"D5":56.8,"IC50":416.32,"Slope":1.12,"max":56.8,"min":0,"Barcode":"H1FO4B-N6029","W5":"P21","W4":"O21","W3":"N21","W2":"M21","W1":"L21"},{"CellLine":"Pa14C","Drug":"BIIB021","minConc":1,"D1":-6.11,"D2":-7.46,"D3":-9.15,"D4":29.51,"D5":38.97,"IC50":650.5,"Slope":2.5,"max":38.97,"min":0,"Barcode":"H1FO4B-N7029","W5":"P13","W4":"N13","W3":"M13","W2":"L13","W1":"K13"},{"CellLine":"Pa14C","Drug":"Tanespimycin","minConc":1,"D1":2.44,"D2":-1.52,"D3":20.34,"D4":30.18,"D5":41.61,"IC50":168.41,"Slope":0.8,"max":41.61,"min":0,"Barcode":"H1FO4B-N7029","W5":"A14","W4":"B14","W3":"C14","W2":"D14","W1":"E14"},{"CellLine":"Pa14C","Drug":"CUDC_305","minConc":1,"D1":2.59,"D2":6.94,"D3":-0.24,"D4":25.11,"D5":46.62,"IC50":940.15,"Slope":2.5,"max":46.62,"min":0,"Barcode":"H1FO4B-N7029","W5":"A15","W4":"B15","W3":"C15","W2":"D15","W1":"F15"},{"CellLine":"Pa14C","Drug":"Luminespib","minConc":0.1,"D1":-0.74,"D2":1.67,"D3":4.38,"D4":39.82,"D5":38.8,"IC50":22.58,"Slope":2.5,"max":39.82,"min":0,"Barcode":"H1FO4B-N7029","W5":"G15","W4":"H15","W3":"I15","W2":"J15","W1":"K15"},{"CellLine":"Pa14C","Drug":"Ganetespib","minConc":0.1,"D1":6.02,"D2":1.3,"D3":14.34,"D4":39.43,"D5":33.95,"IC50":12.99,"Slope":2.5,"max":39.43,"min":1.3,"Barcode":"H1FO4B-N7029","W5":"A16","W4":"C16","W3":"D16","W2":"E16","W1":"F16"},{"CellLine":"Pa14C","Drug":"Stattic","minConc":5,"D1":8.84,"D2":5.74,"D3":11.33,"D4":43.06,"D5":99.67,"IC50":6235.69,"Slope":1.81,"max":99.67,"min":5.74,"Barcode":"H1FO4B-N7029","W5":"F21","W4":"G21","W3":"H21","W2":"I21","W1":"J21"},{"CellLine":"Pa14C","Drug":"ONX_0914","minConc":1,"D1":5.5,"D2":4.01,"D3":18.28,"D4":97.88,"D5":97.46,"IC50":197.52,"Slope":2.5,"max":97.88,"min":4.01,"Barcode":"H1FO4B-N7029","W5":"G23","W4":"H23","W3":"I23","W2":"J23","W1":"K23"},{"CellLine":"Pa14C","Drug":"SN_38","minConc":1,"D1":6.82,"D2":17.07,"D3":42.53,"D4":84.33,"D5":87.22,"IC50":110.15,"Slope":1.09,"max":87.22,"min":6.82,"Barcode":"H1FO4B-N8029","W5":"G2","W4":"H2","W3":"I2","W2":"J2","W1":"K2"},{"CellLine":"Pa14C","Drug":"Cabazitaxel","minConc":0.1,"D1":5.7,"D2":13.54,"D3":25.86,"D4":35.12,"D5":36.2,"IC50":3.31,"Slope":0.78,"max":36.2,"min":3.92,"Barcode":"H1FO4B-N8029","W5":"A3","W4":"B3","W3":"C3","W2":"D3","W1":"E3"},{"CellLine":"Pa14C","Drug":"OTSSP167","minConc":0.1,"D1":8.72,"D2":4.85,"D3":-1.93,"D4":58.82,"D5":74.76,"IC50":59.65,"Slope":2.5,"max":74.76,"min":0,"Barcode":"H1FO4B-N8029","W5":"A5","W4":"B5","W3":"C5","W2":"E5","W1":"F5"},{"CellLine":"Pa14C","Drug":"Prexasertib","minConc":1,"D1":7.69,"D2":8.24,"D3":23.76,"D4":20.65,"D5":18.55,"IC50":27.66,"Slope":2.5,"max":23.76,"min":7.39,"Barcode":"H1FO4B-N8029","W5":"P6","W4":"O6","W3":"N6","W2":"M6","W1":"L6"},{"CellLine":"Pa14C","Drug":"Romidepsin","minConc":0.1,"D1":5.23,"D2":18.5,"D3":98,"D4":100.52,"D5":99.28,"IC50":2.09,"Slope":2.45,"max":100,"min":5.18,"Barcode":"H1FO4B-N8029","W5":"F18","W4":"G18","W3":"H18","W2":"I18","W1":"J18"},{"CellLine":"Pa14C","Drug":"Sabutoclax","minConc":2.5,"D1":3.88,"D2":6.03,"D3":9.94,"D4":41.07,"D5":81.38,"IC50":2556.84,"Slope":1.39,"max":81.38,"min":3.88,"Barcode":"H1FO4B-N8029","W5":"F19","W4":"G19","W3":"I19","W2":"J19","W1":"K19"},{"CellLine":"Pa14C","Drug":"BCI","minConc":5,"D1":5.26,"D2":0.64,"D3":1.29,"D4":64.79,"D5":105.01,"IC50":3930.26,"Slope":2.5,"max":100,"min":0.64,"Barcode":"H1FO4B-N8029","W5":"A22","W4":"B22","W3":"C22","W2":"D22","W1":"E22"},{"CellLine":"Pa02C","Drug":"PF_04691502","minConc":1,"D1":-2.36,"D2":11.86,"D3":49.8,"D4":87.7,"D5":98.48,"IC50":97.14,"Slope":0.92,"max":98.48,"min":0,"Barcode":"H1FO4B-N1039","W5":"B2","W4":"C2","W3":"D2","W2":"E2","W1":"F2"},{"CellLine":"Pa02C","Drug":"SNS_032","minConc":1,"D1":3.04,"D2":8.38,"D3":85.7,"D4":100.32,"D5":96.42,"IC50":41.46,"Slope":1.99,"max":100,"min":2.99,"Barcode":"H1FO4B-N1039","W5":"G9","W4":"H9","W3":"I9","W2":"J9","W1":"K9"},{"CellLine":"Pa02C","Drug":"Alvocidib","minConc":1,"D1":1.77,"D2":-4.35,"D3":16.28,"D4":97.68,"D5":102.33,"IC50":199.52,"Slope":2.38,"max":100,"min":0,"Barcode":"H1FO4B-N1039","W5":"P9","W4":"O9","W3":"N9","W2":"M9","W1":"L9"},{"CellLine":"Pa02C","Drug":"KX2_391","minConc":1,"D1":-12.69,"D2":3.58,"D3":55.69,"D4":58.48,"D5":56.56,"IC50":30.05,"Slope":2.5,"max":58.48,"min":0,"Barcode":"H1FO4B-N1039","W5":"F17","W4":"G17","W3":"H17","W2":"I17","W1":"J17"},{"CellLine":"Pa02C","Drug":"AZD8055","minConc":1,"D1":19.28,"D2":31.58,"D3":58.91,"D4":78.94,"D5":79.69,"IC50":43.45,"Slope":0.92,"max":79.69,"min":17.97,"Barcode":"H1FO4B-N1039","W5":"G23","W4":"H23","W3":"I23","W2":"J23","W1":"K23"},{"CellLine":"Pa02C","Drug":"PF_03758309","minConc":1,"D1":5.5,"D2":31.63,"D3":61.31,"D4":82.79,"D5":101.11,"IC50":45.71,"Slope":0.6,"max":100,"min":0,"Barcode":"H1FO4B-N2039","W5":"P5","W4":"O5","W3":"N5","W2":"M5","W1":"L5"},{"CellLine":"Pa02C","Drug":"AZD_8330","minConc":1,"D1":36.85,"D2":70.36,"D3":82.22,"D4":85,"D5":82.73,"IC50":1.4,"Slope":0.8,"max":85,"min":0,"Barcode":"H1FO4B-N2039","W5":"F6","W4":"H6","W3":"I6","W2":"J6","W1":"K6"},{"CellLine":"Pa02C","Drug":"Rigosertib","minConc":1,"D1":9.69,"D2":10.54,"D3":48.26,"D4":64.77,"D5":60.51,"IC50":67.8,"Slope":2.18,"max":64.77,"min":9.69,"Barcode":"H1FO4B-N2039","W5":"F8","W4":"G8","W3":"H8","W2":"J8","W1":"K8"},{"CellLine":"Pa02C","Drug":"Omipalisib","minConc":0.1,"D1":-13.75,"D2":0.86,"D3":39.34,"D4":82.84,"D5":89.03,"IC50":12.05,"Slope":1.4,"max":89.03,"min":0,"Barcode":"H1FO4B-N2039","W5":"P17","W4":"O17","W3":"N17","W2":"M17","W1":"K17"},{"CellLine":"Pa02C","Drug":"UCN_01","minConc":1,"D1":-0.69,"D2":9.25,"D3":69.24,"D4":87.58,"D5":102.04,"IC50":55.37,"Slope":1.18,"max":100,"min":0,"Barcode":"H1FO4B-N2039","W5":"P21","W4":"O21","W3":"N21","W2":"M21","W1":"L21"},{"CellLine":"Pa02C","Drug":"Dinaciclib","minConc":0.1,"D1":-11,"D2":-1.63,"D3":32.88,"D4":102.45,"D5":100.43,"IC50":13.28,"Slope":2.5,"max":100,"min":0,"Barcode":"H1FO4B-N3039","W5":"A3","W4":"B3","W3":"C3","W2":"D3","W1":"E3"},{"CellLine":"Pa02C","Drug":"NVP_BGT226","minConc":0.1,"D1":-2.53,"D2":-3.32,"D3":38.34,"D4":62.83,"D5":76.1,"IC50":12.04,"Slope":1.08,"max":76.1,"min":0,"Barcode":"H1FO4B-N3039","W5":"G12","W4":"H12","W3":"I12","W2":"J12","W1":"K12"},{"CellLine":"Pa02C","Drug":"Gedatolisib","minConc":0.1,"D1":-20.01,"D2":-4.57,"D3":24.06,"D4":78.81,"D5":92.04,"IC50":23.45,"Slope":1.31,"max":92.04,"min":0,"Barcode":"H1FO4B-N3039","W5":"P12","W4":"O12","W3":"N12","W2":"M12","W1":"L12"},{"CellLine":"Pa02C","Drug":"PKI_402","minConc":0.25,"D1":6.48,"D2":10.93,"D3":44.71,"D4":86.45,"D5":90.27,"IC50":28.25,"Slope":1.31,"max":90.27,"min":6.48,"Barcode":"H1FO4B-N3039","W5":"A16","W4":"C16","W3":"D16","W2":"E16","W1":"F16"},{"CellLine":"Pa02C","Drug":"AT7519","minConc":1,"D1":1.57,"D2":15.86,"D3":29.13,"D4":98.62,"D5":98.34,"IC50":157.16,"Slope":1.98,"max":98.62,"min":1.57,"Barcode":"H1FO4B-N3039","W5":"F22","W4":"G22","W3":"H22","W2":"I22","W1":"J22"},{"CellLine":"Pa02C","Drug":"GSK_461364","minConc":1,"D1":5.18,"D2":38.41,"D3":53.72,"D4":55.48,"D5":89.3,"IC50":95.28,"Slope":0.58,"max":89.3,"min":5.18,"Barcode":"H1FO4B-N4039","W5":"A6","W4":"B6","W3":"C6","W2":"D6","W1":"E6"},{"CellLine":"Pa02C","Drug":"Daporinad","minConc":0.1,"D1":3.34,"D2":-5.62,"D3":38.91,"D4":69.58,"D5":69.33,"IC50":9.11,"Slope":2.5,"max":69.58,"min":0,"Barcode":"H1FO4B-N4039","W5":"A14","W4":"B14","W3":"C14","W2":"D14","W1":"E14"},{"CellLine":"Pa02C","Drug":"Mepacrine","minConc":5,"D1":-4.29,"D2":-2.74,"D3":42.02,"D4":101.02,"D5":106.36,"IC50":568.73,"Slope":2.5,"max":100,"min":0,"Barcode":"H1FO4B-N4039","W5":"P20","W4":"O20","W3":"N20","W2":"M20","W1":"L20"},{"CellLine":"Pa02C","Drug":"Camptothecin","minConc":0.5,"D1":6.6,"D2":0.62,"D3":11.73,"D4":46.2,"D5":91.09,"IC50":449.31,"Slope":1.15,"max":91.09,"min":0.62,"Barcode":"H1FO4B-N5039","W5":"O3","W4":"N3","W3":"M3","W2":"L3","W1":"K3"},{"CellLine":"Pa02C","Drug":"Omacetaxine","minConc":1,"D1":0.27,"D2":4.03,"D3":62.18,"D4":96.01,"D5":100.59,"IC50":72.46,"Slope":1.5,"max":100,"min":0,"Barcode":"H1FO4B-N5039","W5":"A5","W4":"B5","W3":"C5","W2":"E5","W1":"F5"},{"CellLine":"Pa02C","Drug":"Vinorelbine","minConc":1,"D1":-13.33,"D2":50.47,"D3":63,"D4":62.14,"D5":84.28,"IC50":15.29,"Slope":0.77,"max":84.28,"min":0,"Barcode":"H1FO4B-N5039","W5":"P14","W4":"O14","W3":"N14","W2":"M14","W1":"L14"},{"CellLine":"Pa02C","Drug":"Docetaxel","minConc":0.1,"D1":-3,"D2":33.06,"D3":57.45,"D4":72.43,"D5":67.96,"IC50":1.56,"Slope":0.95,"max":72.43,"min":0,"Barcode":"H1FO4B-N5039","W5":"P17","W4":"O17","W3":"N17","W2":"M17","W1":"K17"},{"CellLine":"Pa02C","Drug":"Carfilzomib","minConc":0.1,"D1":0.09,"D2":-18.3,"D3":100.99,"D4":101.41,"D5":103.25,"IC50":3.83,"Slope":2.5,"max":100,"min":0,"Barcode":"H1FO4B-N5039","W5":"A18","W4":"B18","W3":"C18","W2":"D18","W1":"E18"},{"CellLine":"Pa02C","Drug":"Patupilone","minConc":0.1,"D1":-8.98,"D2":22.62,"D3":59.66,"D4":64.97,"D5":62.32,"IC50":1.52,"Slope":1.66,"max":64.97,"min":0,"Barcode":"H1FO4B-N5039","W5":"F18","W4":"G18","W3":"H18","W2":"I18","W1":"J18"},{"CellLine":"Pa02C","Drug":"Plicamycin","minConc":1,"D1":-18.04,"D2":-17.86,"D3":29.38,"D4":84.35,"D5":100.34,"IC50":223.79,"Slope":1.31,"max":100,"min":0,"Barcode":"H1FO4B-N5039","W5":"P18","W4":"N18","W3":"M18","W2":"L18","W1":"K18"},{"CellLine":"Pa02C","Drug":"Vincristine","minConc":0.1,"D1":3.94,"D2":2.55,"D3":59.76,"D4":61.86,"D5":68.18,"IC50":4.71,"Slope":2.5,"max":68.18,"min":2.55,"Barcode":"H1FO4B-N5039","W5":"A19","W4":"B19","W3":"C19","W2":"D19","W1":"E19"},{"CellLine":"Pa02C","Drug":"Dactinomycin","minConc":0.1,"D1":1.94,"D2":52.87,"D3":59.55,"D4":101.86,"D5":101.6,"IC50":1.99,"Slope":0.64,"max":100,"min":0,"Barcode":"H1FO4B-N5039","W5":"P19","W4":"O19","W3":"N19","W2":"M19","W1":"L19"},{"CellLine":"Pa02C","Drug":"Vinblastine","minConc":0.1,"D1":0.79,"D2":11.63,"D3":43.89,"D4":65.3,"D5":64.39,"IC50":4.75,"Slope":1.05,"max":65.3,"min":0.15,"Barcode":"H1FO4B-N5039","W5":"A21","W4":"B21","W3":"C21","W2":"D21","W1":"E21"},{"CellLine":"Pa02C","Drug":"Paclitaxel","minConc":0.1,"D1":11.67,"D2":19.49,"D3":49.81,"D4":71.41,"D5":71.19,"IC50":5.83,"Slope":1.15,"max":71.41,"min":11.67,"Barcode":"H1FO4B-N5039","W5":"F21","W4":"G21","W3":"H21","W2":"I21","W1":"J21"},{"CellLine":"Pa02C","Drug":"Oprozomib","minConc":0.25,"D1":15.6,"D2":6.59,"D3":-32.14,"D4":102.59,"D5":106.19,"IC50":100.75,"Slope":2.5,"max":100,"min":0,"Barcode":"H1FO4B-N5039","W5":"A22","W4":"B22","W3":"C22","W2":"D22","W1":"E22"},{"CellLine":"Pa02C","Drug":"Bortezomib","minConc":0.1,"D1":13.06,"D2":-6.8,"D3":103.65,"D4":104.58,"D5":100.08,"IC50":3.3,"Slope":2.5,"max":100,"min":0,"Barcode":"H1FO4B-N5039","W5":"F22","W4":"G22","W3":"H22","W2":"I22","W1":"J22"},{"CellLine":"Pa02C","Drug":"Filanesib","minConc":0.1,"D1":8.39,"D2":9.69,"D3":59.02,"D4":58.62,"D5":57.57,"IC50":2.81,"Slope":2.5,"max":59.02,"min":7.31,"Barcode":"H1FO4B-N5039","W5":"G23","W4":"H23","W3":"I23","W2":"J23","W1":"K23"},{"CellLine":"Pa02C","Drug":"CUDC_907","minConc":1,"D1":-9.62,"D2":7.98,"D3":83.46,"D4":102.99,"D5":101.58,"IC50":40.36,"Slope":1.8,"max":100,"min":0,"Barcode":"H1FO4B-N6039","W5":"F17","W4":"G17","W3":"H17","W2":"I17","W1":"J17"},{"CellLine":"Pa02C","Drug":"YM155","minConc":1,"D1":-1.75,"D2":6.86,"D3":78.52,"D4":102.84,"D5":99.93,"IC50":46.84,"Slope":1.73,"max":100,"min":0,"Barcode":"H1FO4B-N6039","W5":"F18","W4":"G18","W3":"H18","W2":"I18","W1":"J18"},{"CellLine":"Pa02C","Drug":"AT101","minConc":10,"D1":9.65,"D2":8.07,"D3":23.68,"D4":51.98,"D5":104,"IC50":8961.58,"Slope":1.11,"max":100,"min":8.07,"Barcode":"H1FO4B-N6039","W5":"A21","W4":"B21","W3":"C21","W2":"D21","W1":"E21"},{"CellLine":"Pa02C","Drug":"Selinexor","minConc":1,"D1":17.12,"D2":-1.11,"D3":28.4,"D4":69.09,"D5":89.82,"IC50":248.66,"Slope":0.93,"max":89.82,"min":0,"Barcode":"H1FO4B-N6039","W5":"P21","W4":"O21","W3":"N21","W2":"M21","W1":"L21"},{"CellLine":"Pa02C","Drug":"BIIB021","minConc":1,"D1":0.5,"D2":-2.24,"D3":21.64,"D4":82.31,"D5":89.15,"IC50":205.11,"Slope":1.6,"max":89.15,"min":0,"Barcode":"H1FO4B-N7039","W5":"P13","W4":"N13","W3":"M13","W2":"L13","W1":"K13"},{"CellLine":"Pa02C","Drug":"Tanespimycin","minConc":1,"D1":0.06,"D2":4.21,"D3":23.57,"D4":78.81,"D5":91.14,"IC50":229.31,"Slope":1.24,"max":91.14,"min":0.06,"Barcode":"H1FO4B-N7039","W5":"A14","W4":"B14","W3":"C14","W2":"D14","W1":"E14"},{"CellLine":"Pa02C","Drug":"CUDC_305","minConc":1,"D1":-3.42,"D2":2.77,"D3":19.12,"D4":79.96,"D5":96.33,"IC50":292.44,"Slope":1.29,"max":96.33,"min":0,"Barcode":"H1FO4B-N7039","W5":"A15","W4":"B15","W3":"C15","W2":"D15","W1":"F15"},{"CellLine":"Pa02C","Drug":"Luminespib","minConc":0.1,"D1":4.91,"D2":0.45,"D3":2.62,"D4":-3.86,"D5":46.94,"IC50":357.14,"Slope":2.5,"max":46.94,"min":0,"Barcode":"H1FO4B-N7039","W5":"G15","W4":"H15","W3":"I15","W2":"J15","W1":"K15"},{"CellLine":"Pa02C","Drug":"Ganetespib","minConc":0.1,"D1":4.41,"D2":7.78,"D3":4.28,"D4":9.42,"D5":60.59,"IC50":241.19,"Slope":2.5,"max":60.59,"min":4.28,"Barcode":"H1FO4B-N7039","W5":"A16","W4":"C16","W3":"D16","W2":"E16","W1":"F16"},{"CellLine":"Pa02C","Drug":"Stattic","minConc":5,"D1":7.45,"D2":-2.17,"D3":1.68,"D4":60.25,"D5":103.43,"IC50":4230.06,"Slope":2.5,"max":100,"min":0,"Barcode":"H1FO4B-N7039","W5":"F21","W4":"G21","W3":"H21","W2":"I21","W1":"J21"},{"CellLine":"Pa02C","Drug":"ONX_0914","minConc":1,"D1":8.37,"D2":2.17,"D3":-26.72,"D4":104.65,"D5":102.83,"IC50":382.23,"Slope":2.5,"max":100,"min":0,"Barcode":"H1FO4B-N7039","W5":"G23","W4":"H23","W3":"I23","W2":"J23","W1":"K23"},{"CellLine":"Pa02C","Drug":"SN_38","minConc":1,"D1":5.79,"D2":11.91,"D3":44.69,"D4":73.43,"D5":97.79,"IC50":159.86,"Slope":0.71,"max":97.79,"min":2.24,"Barcode":"H1FO4B-N8039","W5":"G2","W4":"H2","W3":"I2","W2":"J2","W1":"K2"},{"CellLine":"Pa02C","Drug":"Cabazitaxel","minConc":0.1,"D1":1.34,"D2":25.61,"D3":56.33,"D4":68.72,"D5":54.68,"IC50":1.87,"Slope":0.95,"max":68.72,"min":0,"Barcode":"H1FO4B-N8039","W5":"A3","W4":"B3","W3":"C3","W2":"D3","W1":"E3"},{"CellLine":"Pa02C","Drug":"OTSSP167","minConc":0.1,"D1":0.28,"D2":4.55,"D3":7.28,"D4":35.08,"D5":101.5,"IC50":128.09,"Slope":2.5,"max":100,"min":0.28,"Barcode":"H1FO4B-N8039","W5":"A5","W4":"B5","W3":"C5","W2":"E5","W1":"F5"},{"CellLine":"Pa02C","Drug":"Prexasertib","minConc":1,"D1":1.14,"D2":6.37,"D3":76.19,"D4":82.61,"D5":44.77,"IC50":33.16,"Slope":2.23,"max":82.61,"min":1.11,"Barcode":"H1FO4B-N8039","W5":"P6","W4":"O6","W3":"N6","W2":"M6","W1":"L6"},{"CellLine":"Pa02C","Drug":"Romidepsin","minConc":0.1,"D1":-0.43,"D2":15.66,"D3":103.5,"D4":103.35,"D5":99.2,"IC50":1.93,"Slope":2.5,"max":100,"min":0,"Barcode":"H1FO4B-N8039","W5":"F18","W4":"G18","W3":"H18","W2":"I18","W1":"J18"},{"CellLine":"Pa02C","Drug":"Sabutoclax","minConc":2.5,"D1":-6.63,"D2":-3.32,"D3":-0.98,"D4":51.93,"D5":101.42,"IC50":2424.01,"Slope":2.5,"max":100,"min":0,"Barcode":"H1FO4B-N8039","W5":"F19","W4":"G19","W3":"I19","W2":"J19","W1":"K19"},{"CellLine":"Pa02C","Drug":"BCI","minConc":5,"D1":7.68,"D2":11.03,"D3":18.43,"D4":97.76,"D5":106.34,"IC50":1131.17,"Slope":2.48,"max":100,"min":7.68,"Barcode":"H1FO4B-N8039","W5":"A22","W4":"B22","W3":"C22","W2":"D22","W1":"E22"},{"CellLine":"Pa18C","Drug":"PF_04691502","minConc":1,"D1":2.52,"D2":8.7,"D3":48.75,"D4":73.15,"D5":77.55,"IC50":66.3,"Slope":1.15,"max":77.55,"min":1.5,"Barcode":"H1FO4B-N1059","W5":"B2","W4":"C2","W3":"D2","W2":"E2","W1":"F2"},{"CellLine":"Pa18C","Drug":"SNS_032","minConc":1,"D1":-0.27,"D2":11.73,"D3":41.62,"D4":34.83,"D5":31.7,"IC50":14.53,"Slope":2.5,"max":41.62,"min":0,"Barcode":"H1FO4B-N1059","W5":"G9","W4":"H9","W3":"I9","W2":"J9","W1":"K9"},{"CellLine":"Pa18C","Drug":"Alvocidib","minConc":1,"D1":-3.24,"D2":2.56,"D3":29.14,"D4":31.72,"D5":95.5,"IC50":1061.82,"Slope":0.77,"max":95.5,"min":0,"Barcode":"H1FO4B-N1059","W5":"P9","W4":"O9","W3":"N9","W2":"M9","W1":"L9"},{"CellLine":"Pa18C","Drug":"KX2_391","minConc":1,"D1":-2.89,"D2":-1.89,"D3":46.84,"D4":44.2,"D5":53.45,"IC50":47.38,"Slope":2.5,"max":53.45,"min":0,"Barcode":"H1FO4B-N1059","W5":"F17","W4":"G17","W3":"H17","W2":"I17","W1":"J17"},{"CellLine":"Pa18C","Drug":"AZD8055","minConc":1,"D1":7.78,"D2":23.64,"D3":52.32,"D4":57.88,"D5":65.29,"IC50":20.96,"Slope":0.74,"max":65.29,"min":1.33,"Barcode":"H1FO4B-N1059","W5":"G23","W4":"H23","W3":"I23","W2":"J23","W1":"K23"},{"CellLine":"Pa18C","Drug":"PF_03758309","minConc":1,"D1":8.87,"D2":21.47,"D3":42.95,"D4":59.73,"D5":78.52,"IC50":90.07,"Slope":0.56,"max":78.52,"min":3.89,"Barcode":"H1FO4B-N2059","W5":"P5","W4":"O5","W3":"N5","W2":"M5","W1":"L5"},{"CellLine":"Pa18C","Drug":"AZD_8330","minConc":1,"D1":22.26,"D2":25.03,"D3":38.97,"D4":35.69,"D5":38.45,"IC50":18.92,"Slope":2.5,"max":38.97,"min":22.23,"Barcode":"H1FO4B-N2059","W5":"F6","W4":"H6","W3":"I6","W2":"J6","W1":"K6"},{"CellLine":"Pa18C","Drug":"Rigosertib","minConc":1,"D1":13.07,"D2":5.6,"D3":64.81,"D4":51.38,"D5":55.23,"IC50":31.65,"Slope":2.5,"max":64.81,"min":5.6,"Barcode":"H1FO4B-N2059","W5":"F8","W4":"G8","W3":"H8","W2":"J8","W1":"K8"},{"CellLine":"Pa18C","Drug":"Omipalisib","minConc":0.1,"D1":1.81,"D2":7.26,"D3":54.79,"D4":70.95,"D5":61.19,"IC50":4.69,"Slope":1.57,"max":70.95,"min":1.65,"Barcode":"H1FO4B-N2059","W5":"P17","W4":"O17","W3":"N17","W2":"M17","W1":"K17"},{"CellLine":"Pa18C","Drug":"UCN_01","minConc":1,"D1":4.63,"D2":9.43,"D3":22.98,"D4":35.23,"D5":64.32,"IC50":508.54,"Slope":0.68,"max":64.32,"min":4.63,"Barcode":"H1FO4B-N2059","W5":"P21","W4":"O21","W3":"N21","W2":"M21","W1":"L21"},{"CellLine":"Pa18C","Drug":"Dinaciclib","minConc":0.1,"D1":2.19,"D2":1.28,"D3":47.99,"D4":37.62,"D5":33.93,"IC50":3.05,"Slope":2.5,"max":47.99,"min":0.46,"Barcode":"H1FO4B-N3059","W5":"A3","W4":"B3","W3":"C3","W2":"D3","W1":"E3"},{"CellLine":"Pa18C","Drug":"NVP_BGT226","minConc":0.1,"D1":3.48,"D2":7.54,"D3":37.41,"D4":61.4,"D5":67.44,"IC50":8.87,"Slope":1.03,"max":67.44,"min":2.2,"Barcode":"H1FO4B-N3059","W5":"G12","W4":"H12","W3":"I12","W2":"J12","W1":"K12"},{"CellLine":"Pa18C","Drug":"Gedatolisib","minConc":0.1,"D1":-3.95,"D2":5.3,"D3":4.21,"D4":63.06,"D5":69.14,"IC50":34.51,"Slope":2.19,"max":69.14,"min":0,"Barcode":"H1FO4B-N3059","W5":"P12","W4":"O12","W3":"N12","W2":"M12","W1":"L12"},{"CellLine":"Pa18C","Drug":"PKI_402","minConc":0.25,"D1":2.58,"D2":3.98,"D3":14.16,"D4":69.66,"D5":69.9,"IC50":46.71,"Slope":2.5,"max":69.9,"min":2.58,"Barcode":"H1FO4B-N3059","W5":"A16","W4":"C16","W3":"D16","W2":"E16","W1":"F16"},{"CellLine":"Pa18C","Drug":"AT7519","minConc":1,"D1":5.96,"D2":8.98,"D3":23.61,"D4":34.69,"D5":39.68,"IC50":89.37,"Slope":0.83,"max":39.68,"min":4.76,"Barcode":"H1FO4B-N3059","W5":"F22","W4":"G22","W3":"H22","W2":"I22","W1":"J22"},{"CellLine":"Pa18C","Drug":"GSK_461364","minConc":1,"D1":8.56,"D2":4.11,"D3":8.41,"D4":1.94,"D5":61.15,"IC50":3159.79,"Slope":2.5,"max":61.15,"min":1.94,"Barcode":"H1FO4B-N4059","W5":"A6","W4":"B6","W3":"C6","W2":"D6","W1":"E6"},{"CellLine":"Pa18C","Drug":"Daporinad","minConc":0.1,"D1":-3.17,"D2":1.4,"D3":10.3,"D4":70.24,"D5":76.41,"IC50":27.12,"Slope":1.86,"max":76.41,"min":0,"Barcode":"H1FO4B-N4059","W5":"A14","W4":"B14","W3":"C14","W2":"D14","W1":"E14"},{"CellLine":"Pa18C","Drug":"Mepacrine","minConc":5,"D1":7.89,"D2":5.96,"D3":11.67,"D4":15.41,"D5":98.87,"IC50":11610.92,"Slope":2.5,"max":98.87,"min":5.96,"Barcode":"H1FO4B-N4059","W5":"P20","W4":"O20","W3":"N20","W2":"M20","W1":"L20"},{"CellLine":"Pa18C","Drug":"Camptothecin","minConc":0.5,"D1":5.44,"D2":7.84,"D3":17.71,"D4":21.17,"D5":71.38,"IC50":840.13,"Slope":2.19,"max":71.38,"min":5.44,"Barcode":"H1FO4B-N5059","W5":"O3","W4":"N3","W3":"M3","W2":"L3","W1":"K3"},{"CellLine":"Pa18C","Drug":"Omacetaxine","minConc":1,"D1":3.97,"D2":8.85,"D3":31.79,"D4":62.43,"D5":70.6,"IC50":137.75,"Slope":0.99,"max":70.6,"min":3.82,"Barcode":"H1FO4B-N5059","W5":"A5","W4":"B5","W3":"C5","W2":"E5","W1":"F5"},{"CellLine":"Pa18C","Drug":"Vinorelbine","minConc":1,"D1":0.08,"D2":15.55,"D3":61.41,"D4":60.95,"D5":76.11,"IC50":33.09,"Slope":1.04,"max":76.11,"min":0,"Barcode":"H1FO4B-N5059","W5":"P14","W4":"O14","W3":"N14","W2":"M14","W1":"L14"},{"CellLine":"Pa18C","Drug":"Docetaxel","minConc":0.1,"D1":3.11,"D2":10.22,"D3":49.04,"D4":65.88,"D5":65.61,"IC50":4.65,"Slope":1.34,"max":65.88,"min":2.87,"Barcode":"H1FO4B-N5059","W5":"P17","W4":"O17","W3":"N17","W2":"M17","W1":"K17"},{"CellLine":"Pa18C","Drug":"Carfilzomib","minConc":0.1,"D1":2.13,"D2":0.93,"D3":8.42,"D4":64.59,"D5":82.94,"IC50":44.58,"Slope":1.55,"max":82.94,"min":0.93,"Barcode":"H1FO4B-N5059","W5":"A18","W4":"B18","W3":"C18","W2":"D18","W1":"E18"},{"CellLine":"Pa18C","Drug":"Patupilone","minConc":0.1,"D1":8.64,"D2":14.24,"D3":60.9,"D4":64.78,"D5":60.31,"IC50":2.86,"Slope":2.08,"max":64.78,"min":8.59,"Barcode":"H1FO4B-N5059","W5":"F18","W4":"G18","W3":"H18","W2":"I18","W1":"J18"},{"CellLine":"Pa18C","Drug":"Plicamycin","minConc":1,"D1":-7.38,"D2":7.33,"D3":20.64,"D4":50.88,"D5":70.67,"IC50":287.6,"Slope":0.84,"max":70.67,"min":0,"Barcode":"H1FO4B-N5059","W5":"P18","W4":"N18","W3":"M18","W2":"L18","W1":"K18"},{"CellLine":"Pa18C","Drug":"Vincristine","minConc":0.1,"D1":0.89,"D2":2.79,"D3":8.65,"D4":60.56,"D5":57.75,"IC50":21.13,"Slope":2.5,"max":60.56,"min":0.89,"Barcode":"H1FO4B-N5059","W5":"A19","W4":"B19","W3":"C19","W2":"D19","W1":"E19"},{"CellLine":"Pa18C","Drug":"Dactinomycin","minConc":0.1,"D1":2.29,"D2":17.13,"D3":27.14,"D4":57.76,"D5":76.33,"IC50":19.89,"Slope":0.68,"max":76.33,"min":2.29,"Barcode":"H1FO4B-N5059","W5":"P19","W4":"O19","W3":"N19","W2":"M19","W1":"L19"},{"CellLine":"Pa18C","Drug":"Vinblastine","minConc":0.1,"D1":6,"D2":6.48,"D3":11.43,"D4":65.29,"D5":61.32,"IC50":24.1,"Slope":2.5,"max":65.29,"min":6,"Barcode":"H1FO4B-N5059","W5":"A21","W4":"B21","W3":"C21","W2":"D21","W1":"E21"},{"CellLine":"Pa18C","Drug":"Paclitaxel","minConc":0.1,"D1":-0.72,"D2":7.86,"D3":38.53,"D4":62.69,"D5":62.93,"IC50":6.4,"Slope":1.15,"max":62.93,"min":0,"Barcode":"H1FO4B-N5059","W5":"F21","W4":"G21","W3":"H21","W2":"I21","W1":"J21"},{"CellLine":"Pa18C","Drug":"Oprozomib","minConc":0.25,"D1":1.05,"D2":-5.07,"D3":1.2,"D4":45.6,"D5":58.56,"IC50":142.89,"Slope":2.25,"max":58.56,"min":0,"Barcode":"H1FO4B-N5059","W5":"A22","W4":"B22","W3":"C22","W2":"D22","W1":"E22"},{"CellLine":"Pa18C","Drug":"Bortezomib","minConc":0.1,"D1":1.33,"D2":2.39,"D3":34.24,"D4":59.95,"D5":70.21,"IC50":11.77,"Slope":0.98,"max":70.21,"min":0,"Barcode":"H1FO4B-N5059","W5":"F22","W4":"G22","W3":"H22","W2":"I22","W1":"J22"},{"CellLine":"Pa18C","Drug":"Filanesib","minConc":0.1,"D1":3.41,"D2":3.44,"D3":10.08,"D4":11.03,"D5":11.91,"IC50":5.67,"Slope":2.29,"max":11.91,"min":3.35,"Barcode":"H1FO4B-N5059","W5":"G23","W4":"H23","W3":"I23","W2":"J23","W1":"K23"},{"CellLine":"Pa18C","Drug":"CUDC_907","minConc":1,"D1":-3.69,"D2":2.94,"D3":60.24,"D4":97.33,"D5":95.69,"IC50":76.2,"Slope":1.8,"max":97.33,"min":0,"Barcode":"H1FO4B-N6059","W5":"F17","W4":"G17","W3":"H17","W2":"I17","W1":"J17"},{"CellLine":"Pa18C","Drug":"YM155","minConc":1,"D1":2.97,"D2":26.75,"D3":103.8,"D4":103.41,"D5":102.85,"IC50":15.54,"Slope":2.5,"max":100,"min":2.74,"Barcode":"H1FO4B-N6059","W5":"F18","W4":"G18","W3":"H18","W2":"I18","W1":"J18"},{"CellLine":"Pa18C","Drug":"AT101","minConc":10,"D1":3.93,"D2":7.97,"D3":10.38,"D4":50.96,"D5":103.17,"IC50":10080.73,"Slope":1.76,"max":100,"min":3.93,"Barcode":"H1FO4B-N6059","W5":"A21","W4":"B21","W3":"C21","W2":"D21","W1":"E21"},{"CellLine":"Pa18C","Drug":"Selinexor","minConc":1,"D1":10.57,"D2":2.09,"D3":35.55,"D4":63.12,"D5":72.04,"IC50":119.22,"Slope":1.07,"max":72.04,"min":2.09,"Barcode":"H1FO4B-N6059","W5":"P21","W4":"O21","W3":"N21","W2":"M21","W1":"L21"},{"CellLine":"Pa18C","Drug":"BIIB021","minConc":1,"D1":-21.78,"D2":1.81,"D3":22.09,"D4":48.34,"D5":53.09,"IC50":133.15,"Slope":1.3,"max":53.09,"min":0,"Barcode":"H1FO4B-N7059","W5":"P13","W4":"N13","W3":"M13","W2":"L13","W1":"K13"},{"CellLine":"Pa18C","Drug":"Tanespimycin","minConc":1,"D1":-0.49,"D2":-1.04,"D3":23.66,"D4":49.06,"D5":52.14,"IC50":114.44,"Slope":1.58,"max":52.14,"min":0,"Barcode":"H1FO4B-N7059","W5":"A14","W4":"B14","W3":"C14","W2":"D14","W1":"E14"},{"CellLine":"Pa18C","Drug":"CUDC_305","minConc":1,"D1":2.19,"D2":1.79,"D3":7.69,"D4":47.91,"D5":46.49,"IC50":212.72,"Slope":2.5,"max":47.91,"min":1.79,"Barcode":"H1FO4B-N7059","W5":"A15","W4":"B15","W3":"C15","W2":"D15","W1":"F15"},{"CellLine":"Pa18C","Drug":"Luminespib","minConc":0.1,"D1":6.21,"D2":7.63,"D3":6.3,"D4":41.98,"D5":43.25,"IC50":34.16,"Slope":2.5,"max":43.25,"min":6.21,"Barcode":"H1FO4B-N7059","W5":"G15","W4":"H15","W3":"I15","W2":"J15","W1":"K15"},{"CellLine":"Pa18C","Drug":"Ganetespib","minConc":0.1,"D1":4.55,"D2":3.29,"D3":8.72,"D4":47.37,"D5":41.62,"IC50":21.6,"Slope":2.5,"max":47.37,"min":3.29,"Barcode":"H1FO4B-N7059","W5":"A16","W4":"C16","W3":"D16","W2":"E16","W1":"F16"},{"CellLine":"Pa18C","Drug":"Stattic","minConc":5,"D1":3.26,"D2":-2.52,"D3":6.01,"D4":25.23,"D5":102.88,"IC50":7685.83,"Slope":2.5,"max":100,"min":0,"Barcode":"H1FO4B-N7059","W5":"F21","W4":"G21","W3":"H21","W2":"I21","W1":"J21"},{"CellLine":"Pa18C","Drug":"ONX_0914","minConc":1,"D1":2.8,"D2":0.91,"D3":-0.39,"D4":65.5,"D5":81.53,"IC50":571.35,"Slope":2.5,"max":81.53,"min":0,"Barcode":"H1FO4B-N7059","W5":"G23","W4":"H23","W3":"I23","W2":"J23","W1":"K23"},{"CellLine":"Pa18C","Drug":"SN_38","minConc":1,"D1":5.99,"D2":8.39,"D3":14.93,"D4":64.68,"D5":72.58,"IC50":302.5,"Slope":1.67,"max":72.58,"min":5.99,"Barcode":"H1FO4B-N8059","W5":"G2","W4":"H2","W3":"I2","W2":"J2","W1":"K2"},{"CellLine":"Pa18C","Drug":"Cabazitaxel","minConc":0.1,"D1":4.46,"D2":9.62,"D3":55.6,"D4":67.59,"D5":66.49,"IC50":4.2,"Slope":1.67,"max":67.59,"min":4.36,"Barcode":"H1FO4B-N8059","W5":"A3","W4":"B3","W3":"C3","W2":"D3","W1":"E3"},{"CellLine":"Pa18C","Drug":"OTSSP167","minConc":0.1,"D1":6.67,"D2":11.25,"D3":10.33,"D4":26.13,"D5":35.94,"IC50":51.97,"Slope":1.08,"max":35.94,"min":6.67,"Barcode":"H1FO4B-N8059","W5":"A5","W4":"B5","W3":"C5","W2":"E5","W1":"F5"},{"CellLine":"Pa18C","Drug":"Prexasertib","minConc":1,"D1":1.89,"D2":8.06,"D3":13.45,"D4":13.27,"D5":10.54,"IC50":9.45,"Slope":2.5,"max":13.45,"min":1.84,"Barcode":"H1FO4B-N8059","W5":"P6","W4":"O6","W3":"N6","W2":"M6","W1":"L6"},{"CellLine":"Pa18C","Drug":"Romidepsin","minConc":0.1,"D1":-1.21,"D2":16.79,"D3":81.63,"D4":95.73,"D5":95.39,"IC50":2.95,"Slope":1.45,"max":95.73,"min":0,"Barcode":"H1FO4B-N8059","W5":"F18","W4":"G18","W3":"H18","W2":"I18","W1":"J18"},{"CellLine":"Pa18C","Drug":"Sabutoclax","minConc":2.5,"D1":-4.06,"D2":-5.65,"D3":3.65,"D4":61.36,"D5":70.69,"IC50":1016.23,"Slope":2.1,"max":70.69,"min":0,"Barcode":"H1FO4B-N8059","W5":"F19","W4":"G19","W3":"I19","W2":"J19","W1":"K19"},{"CellLine":"Pa18C","Drug":"BCI","minConc":5,"D1":3.85,"D2":2,"D3":5.44,"D4":57.08,"D5":101.86,"IC50":4351.36,"Slope":1.87,"max":100,"min":2,"Barcode":"H1FO4B-N8059","W5":"A22","W4":"B22","W3":"C22","W2":"D22","W1":"E22"},{"CellLine":"Pa03C","Drug":"PF_04691502","minConc":1,"D1":1.06,"D2":4.53,"D3":31.49,"D4":73.94,"D5":94.12,"IC50":220.52,"Slope":0.91,"max":94.12,"min":0,"Barcode":"H1FO4B-N1069","W5":"B2","W4":"C2","W3":"D2","W2":"E2","W1":"F2"},{"CellLine":"Pa03C","Drug":"SNS_032","minConc":1,"D1":6.15,"D2":19.94,"D3":58.68,"D4":76.49,"D5":71.77,"IC50":34.54,"Slope":1.07,"max":76.49,"min":4.67,"Barcode":"H1FO4B-N1069","W5":"G9","W4":"H9","W3":"I9","W2":"J9","W1":"K9"},{"CellLine":"Pa03C","Drug":"Alvocidib","minConc":1,"D1":3.51,"D2":7.7,"D3":35.9,"D4":74.47,"D5":74.11,"IC50":109.78,"Slope":1.6,"max":74.47,"min":3.51,"Barcode":"H1FO4B-N1069","W5":"P9","W4":"O9","W3":"N9","W2":"M9","W1":"L9"},{"CellLine":"Pa03C","Drug":"KX2_391","minConc":1,"D1":10.15,"D2":7.49,"D3":34.57,"D4":48.61,"D5":54.17,"IC50":84.27,"Slope":1.29,"max":54.17,"min":7.49,"Barcode":"H1FO4B-N1069","W5":"F17","W4":"G17","W3":"H17","W2":"I17","W1":"J17"},{"CellLine":"Pa03C","Drug":"AZD8055","minConc":1,"D1":12.37,"D2":27.3,"D3":49.68,"D4":68.84,"D5":70.15,"IC50":34.91,"Slope":0.78,"max":70.15,"min":9.4,"Barcode":"H1FO4B-N1069","W5":"G23","W4":"H23","W3":"I23","W2":"J23","W1":"K23"},{"CellLine":"Pa03C","Drug":"PF_03758309","minConc":1,"D1":16.07,"D2":34.23,"D3":44.79,"D4":59.48,"D5":88.69,"IC50":203.59,"Slope":0.51,"max":88.69,"min":15.33,"Barcode":"H1FO4B-N2069","W5":"P5","W4":"O5","W3":"N5","W2":"M5","W1":"L5"},{"CellLine":"Pa03C","Drug":"AZD_8330","minConc":1,"D1":4.13,"D2":24.34,"D3":28.52,"D4":38.85,"D5":33.77,"IC50":8.84,"Slope":0.66,"max":38.85,"min":0,"Barcode":"H1FO4B-N2069","W5":"F6","W4":"H6","W3":"I6","W2":"J6","W1":"K6"},{"CellLine":"Pa03C","Drug":"Rigosertib","minConc":1,"D1":12.46,"D2":13.97,"D3":42.48,"D4":49.83,"D5":51,"IC50":51.24,"Slope":1.88,"max":51,"min":12.35,"Barcode":"H1FO4B-N2069","W5":"F8","W4":"G8","W3":"H8","W2":"J8","W1":"K8"},{"CellLine":"Pa03C","Drug":"Omipalisib","minConc":0.1,"D1":4.03,"D2":0.75,"D3":25.11,"D4":70.17,"D5":72.99,"IC50":14.87,"Slope":1.71,"max":72.99,"min":0.75,"Barcode":"H1FO4B-N2069","W5":"P17","W4":"O17","W3":"N17","W2":"M17","W1":"K17"},{"CellLine":"Pa03C","Drug":"UCN_01","minConc":1,"D1":10.41,"D2":30.7,"D3":57.67,"D4":47.25,"D5":92.74,"IC50":102.49,"Slope":0.57,"max":92.74,"min":10.41,"Barcode":"H1FO4B-N2069","W5":"P21","W4":"O21","W3":"N21","W2":"M21","W1":"L21"},{"CellLine":"Pa03C","Drug":"Dinaciclib","minConc":0.1,"D1":4.71,"D2":8.04,"D3":48.91,"D4":73.6,"D5":78.86,"IC50":7.34,"Slope":1.23,"max":78.86,"min":3.33,"Barcode":"H1FO4B-N3069","W5":"A3","W4":"B3","W3":"C3","W2":"D3","W1":"E3"},{"CellLine":"Pa03C","Drug":"NVP_BGT226","minConc":0.1,"D1":-16.18,"D2":14.08,"D3":42.22,"D4":66.86,"D5":73.04,"IC50":6.95,"Slope":0.97,"max":73.04,"min":0,"Barcode":"H1FO4B-N3069","W5":"G12","W4":"H12","W3":"I12","W2":"J12","W1":"K12"},{"CellLine":"Pa03C","Drug":"Gedatolisib","minConc":0.1,"D1":-1.23,"D2":6.25,"D3":28.53,"D4":69.2,"D5":84.19,"IC50":19.74,"Slope":0.95,"max":84.19,"min":0,"Barcode":"H1FO4B-N3069","W5":"P12","W4":"O12","W3":"N12","W2":"M12","W1":"L12"},{"CellLine":"Pa03C","Drug":"PKI_402","minConc":0.25,"D1":2.22,"D2":15.37,"D3":33.39,"D4":67.04,"D5":81.55,"IC50":37.44,"Slope":0.76,"max":81.55,"min":2.22,"Barcode":"H1FO4B-N3069","W5":"A16","W4":"C16","W3":"D16","W2":"E16","W1":"F16"},{"CellLine":"Pa03C","Drug":"AT7519","minConc":1,"D1":2.3,"D2":-2.88,"D3":11.72,"D4":77.68,"D5":71.29,"IC50":198.29,"Slope":2.5,"max":77.68,"min":0,"Barcode":"H1FO4B-N3069","W5":"F22","W4":"G22","W3":"H22","W2":"I22","W1":"J22"},{"CellLine":"Pa03C","Drug":"GSK_461364","minConc":1,"D1":9.51,"D2":7.38,"D3":17.97,"D4":11.55,"D5":60.62,"IC50":2519.33,"Slope":2.5,"max":60.62,"min":7.38,"Barcode":"H1FO4B-N4069","W5":"A6","W4":"B6","W3":"C6","W2":"D6","W1":"E6"},{"CellLine":"Pa03C","Drug":"Daporinad","minConc":0.1,"D1":9.85,"D2":7.9,"D3":59.21,"D4":82.46,"D5":80.05,"IC50":7.29,"Slope":2.5,"max":82.46,"min":7.9,"Barcode":"H1FO4B-N4069","W5":"A14","W4":"B14","W3":"C14","W2":"D14","W1":"E14"},{"CellLine":"Pa03C","Drug":"Mepacrine","minConc":5,"D1":7.69,"D2":17.66,"D3":15.98,"D4":66.31,"D5":102.19,"IC50":3151.05,"Slope":1.29,"max":100,"min":7.69,"Barcode":"H1FO4B-N4069","W5":"P20","W4":"O20","W3":"N20","W2":"M20","W1":"L20"},{"CellLine":"Pa03C","Drug":"Camptothecin","minConc":0.5,"D1":7.34,"D2":9.82,"D3":26.08,"D4":59.44,"D5":80.7,"IC50":169.47,"Slope":0.92,"max":80.7,"min":7.19,"Barcode":"H1FO4B-N5069","W5":"O3","W4":"N3","W3":"M3","W2":"L3","W1":"K3"},{"CellLine":"Pa03C","Drug":"Omacetaxine","minConc":1,"D1":-1.55,"D2":19.05,"D3":61.65,"D4":92.5,"D5":96.05,"IC50":50.55,"Slope":0.94,"max":96.05,"min":0,"Barcode":"H1FO4B-N5069","W5":"A5","W4":"B5","W3":"C5","W2":"E5","W1":"F5"},{"CellLine":"Pa03C","Drug":"Vinorelbine","minConc":1,"D1":-2.62,"D2":17.81,"D3":45.82,"D4":52.88,"D5":70.89,"IC50":61.07,"Slope":0.64,"max":70.89,"min":0,"Barcode":"H1FO4B-N5069","W5":"P14","W4":"O14","W3":"N14","W2":"M14","W1":"L14"},{"CellLine":"Pa03C","Drug":"Docetaxel","minConc":0.1,"D1":-0.45,"D2":-4.57,"D3":23.69,"D4":27.69,"D5":20.1,"IC50":5.23,"Slope":2.5,"max":27.69,"min":0,"Barcode":"H1FO4B-N5069","W5":"P17","W4":"O17","W3":"N17","W2":"M17","W1":"K17"},{"CellLine":"Pa03C","Drug":"Carfilzomib","minConc":0.1,"D1":-2.97,"D2":-1.57,"D3":80.3,"D4":102.1,"D5":104.19,"IC50":5.73,"Slope":2.5,"max":100,"min":0,"Barcode":"H1FO4B-N5069","W5":"A18","W4":"B18","W3":"C18","W2":"D18","W1":"E18"},{"CellLine":"Pa03C","Drug":"Patupilone","minConc":0.1,"D1":0.6,"D2":-1.93,"D3":27.08,"D4":27.1,"D5":20.91,"IC50":3.53,"Slope":2.5,"max":27.1,"min":0,"Barcode":"H1FO4B-N5069","W5":"F18","W4":"G18","W3":"H18","W2":"I18","W1":"J18"},{"CellLine":"Pa03C","Drug":"Plicamycin","minConc":1,"D1":0.96,"D2":-11.58,"D3":27.87,"D4":78.01,"D5":81.6,"IC50":130.51,"Slope":2.5,"max":81.6,"min":0,"Barcode":"H1FO4B-N5069","W5":"P18","W4":"N18","W3":"M18","W2":"L18","W1":"K18"},{"CellLine":"Pa03C","Drug":"Vincristine","minConc":0.1,"D1":12.62,"D2":-0.63,"D3":34.06,"D4":54.14,"D5":46.09,"IC50":8.1,"Slope":2.5,"max":54.14,"min":0,"Barcode":"H1FO4B-N5069","W5":"A19","W4":"B19","W3":"C19","W2":"D19","W1":"E19"},{"CellLine":"Pa03C","Drug":"Dactinomycin","minConc":0.1,"D1":0.8,"D2":39.12,"D3":47.39,"D4":81.22,"D5":81.67,"IC50":2.58,"Slope":0.63,"max":81.67,"min":0,"Barcode":"H1FO4B-N5069","W5":"P19","W4":"O19","W3":"N19","W2":"M19","W1":"L19"},{"CellLine":"Pa03C","Drug":"Vinblastine","minConc":0.1,"D1":0.55,"D2":1.96,"D3":27.07,"D4":45.66,"D5":46.23,"IC50":8.22,"Slope":1.66,"max":46.23,"min":0.55,"Barcode":"H1FO4B-N5069","W5":"A21","W4":"B21","W3":"C21","W2":"D21","W1":"E21"},{"CellLine":"Pa03C","Drug":"Paclitaxel","minConc":0.1,"D1":3.54,"D2":0.43,"D3":22.84,"D4":27.59,"D5":38.5,"IC50":11.46,"Slope":0.72,"max":38.5,"min":0,"Barcode":"H1FO4B-N5069","W5":"F21","W4":"G21","W3":"H21","W2":"I21","W1":"J21"},{"CellLine":"Pa03C","Drug":"Oprozomib","minConc":0.25,"D1":14.87,"D2":-3.83,"D3":13.13,"D4":99.46,"D5":105.31,"IC50":52.8,"Slope":2.5,"max":100,"min":0,"Barcode":"H1FO4B-N5069","W5":"A22","W4":"B22","W3":"C22","W2":"D22","W1":"E22"},{"CellLine":"Pa03C","Drug":"Bortezomib","minConc":0.1,"D1":4.33,"D2":7.63,"D3":97.46,"D4":101.98,"D5":100.58,"IC50":2.97,"Slope":2.5,"max":100,"min":3.06,"Barcode":"H1FO4B-N5069","W5":"F22","W4":"G22","W3":"H22","W2":"I22","W1":"J22"},{"CellLine":"Pa03C","Drug":"Filanesib","minConc":0.1,"D1":0.12,"D2":-2.83,"D3":30.54,"D4":32.68,"D5":33.94,"IC50":4.54,"Slope":2.5,"max":33.94,"min":0,"Barcode":"H1FO4B-N5069","W5":"G23","W4":"H23","W3":"I23","W2":"J23","W1":"K23"},{"CellLine":"Pa03C","Drug":"CUDC_907","minConc":1,"D1":4.08,"D2":11.12,"D3":98.89,"D4":100.42,"D5":100.81,"IC50":25.53,"Slope":2.5,"max":100,"min":3.42,"Barcode":"H1FO4B-N6069","W5":"F17","W4":"G17","W3":"H17","W2":"I17","W1":"J17"},{"CellLine":"Pa03C","Drug":"YM155","minConc":1,"D1":6.94,"D2":54.18,"D3":100.6,"D4":103.7,"D5":101.58,"IC50":9.85,"Slope":2.5,"max":100,"min":6.63,"Barcode":"H1FO4B-N6069","W5":"F18","W4":"G18","W3":"H18","W2":"I18","W1":"J18"},{"CellLine":"Pa03C","Drug":"AT101","minConc":10,"D1":9.7,"D2":-3.03,"D3":18.76,"D4":54.19,"D5":103.69,"IC50":6881.73,"Slope":1.01,"max":100,"min":0,"Barcode":"H1FO4B-N6069","W5":"A21","W4":"B21","W3":"C21","W2":"D21","W1":"E21"},{"CellLine":"Pa03C","Drug":"Selinexor","minConc":1,"D1":3.5,"D2":7.96,"D3":38.68,"D4":67.2,"D5":76.31,"IC50":106.58,"Slope":0.94,"max":76.31,"min":1.87,"Barcode":"H1FO4B-N6069","W5":"P21","W4":"O21","W3":"N21","W2":"M21","W1":"L21"},{"CellLine":"Pa03C","Drug":"BIIB021","minConc":1,"D1":-2.05,"D2":-4,"D3":-4.78,"D4":72.52,"D5":75.1,"IC50":376.31,"Slope":2.5,"max":75.1,"min":0,"Barcode":"H1FO4B-N7069","W5":"P13","W4":"N13","W3":"M13","W2":"L13","W1":"K13"},{"CellLine":"Pa03C","Drug":"Tanespimycin","minConc":1,"D1":5.06,"D2":-11.76,"D3":67.62,"D4":80.75,"D5":76.43,"IC50":54.24,"Slope":2.5,"max":80.75,"min":0,"Barcode":"H1FO4B-N7069","W5":"A14","W4":"B14","W3":"C14","W2":"D14","W1":"E14"},{"CellLine":"Pa03C","Drug":"CUDC_305","minConc":1,"D1":5.22,"D2":8.46,"D3":6.08,"D4":74.34,"D5":76.79,"IC50":334.74,"Slope":2.5,"max":76.79,"min":5.22,"Barcode":"H1FO4B-N7069","W5":"A15","W4":"B15","W3":"C15","W2":"D15","W1":"F15"},{"CellLine":"Pa03C","Drug":"Luminespib","minConc":0.1,"D1":-1.67,"D2":2.43,"D3":39.38,"D4":75.48,"D5":74.93,"IC50":9.48,"Slope":1.76,"max":75.48,"min":0,"Barcode":"H1FO4B-N7069","W5":"G15","W4":"H15","W3":"I15","W2":"J15","W1":"K15"},{"CellLine":"Pa03C","Drug":"Ganetespib","minConc":0.1,"D1":7.97,"D2":-9.18,"D3":-0.47,"D4":80.03,"D5":77.74,"IC50":32.03,"Slope":2.5,"max":80.03,"min":0,"Barcode":"H1FO4B-N7069","W5":"A16","W4":"C16","W3":"D16","W2":"E16","W1":"F16"},{"CellLine":"Pa03C","Drug":"Stattic","minConc":5,"D1":11.3,"D2":4.8,"D3":3.39,"D4":97.77,"D5":98.99,"IC50":1625.2,"Slope":2.5,"max":98.99,"min":3.39,"Barcode":"H1FO4B-N7069","W5":"F21","W4":"G21","W3":"H21","W2":"I21","W1":"J21"},{"CellLine":"Pa03C","Drug":"ONX_0914","minConc":1,"D1":6.5,"D2":5.59,"D3":11.27,"D4":102.27,"D5":100.98,"IC50":256.49,"Slope":2.5,"max":100,"min":5.15,"Barcode":"H1FO4B-N7069","W5":"G23","W4":"H23","W3":"I23","W2":"J23","W1":"K23"},{"CellLine":"Pa03C","Drug":"SN_38","minConc":1,"D1":6.05,"D2":26.18,"D3":49.43,"D4":72.89,"D5":80.23,"IC50":38.23,"Slope":0.64,"max":80.23,"min":0,"Barcode":"H1FO4B-N8069","W5":"G2","W4":"H2","W3":"I2","W2":"J2","W1":"K2"},{"CellLine":"Pa03C","Drug":"Cabazitaxel","minConc":0.1,"D1":10.13,"D2":16.86,"D3":35.57,"D4":38.11,"D5":27.83,"IC50":2.11,"Slope":1.48,"max":38.11,"min":9.82,"Barcode":"H1FO4B-N8069","W5":"A3","W4":"B3","W3":"C3","W2":"D3","W1":"E3"},{"CellLine":"Pa03C","Drug":"OTSSP167","minConc":0.1,"D1":18.16,"D2":17.76,"D3":17.12,"D4":79.11,"D5":80.95,"IC50":33.42,"Slope":2.5,"max":80.95,"min":16.76,"Barcode":"H1FO4B-N8069","W5":"A5","W4":"B5","W3":"C5","W2":"E5","W1":"F5"},{"CellLine":"Pa03C","Drug":"Prexasertib","minConc":1,"D1":11.81,"D2":31.75,"D3":60.61,"D4":62.21,"D5":18.03,"IC50":6.84,"Slope":0.2,"max":62.21,"min":0,"Barcode":"H1FO4B-N8069","W5":"P6","W4":"O6","W3":"N6","W2":"M6","W1":"L6"},{"CellLine":"Pa03C","Drug":"Romidepsin","minConc":0.1,"D1":8.89,"D2":37.57,"D3":102.46,"D4":101.74,"D5":100.51,"IC50":1.36,"Slope":2.5,"max":100,"min":8.71,"Barcode":"H1FO4B-N8069","W5":"F18","W4":"G18","W3":"H18","W2":"I18","W1":"J18"},{"CellLine":"Pa03C","Drug":"Sabutoclax","minConc":2.5,"D1":-1.82,"D2":12.92,"D3":18.76,"D4":53.9,"D5":85.14,"IC50":1073.68,"Slope":0.79,"max":85.14,"min":0,"Barcode":"H1FO4B-N8069","W5":"F19","W4":"G19","W3":"I19","W2":"J19","W1":"K19"},{"CellLine":"Pa03C","Drug":"BCI","minConc":5,"D1":3.03,"D2":11.82,"D3":5.26,"D4":97.29,"D5":105.2,"IC50":1600.27,"Slope":2.5,"max":100,"min":3.03,"Barcode":"H1FO4B-N8069","W5":"A22","W4":"B22","W3":"C22","W2":"D22","W1":"E22"},{"CellLine":"Pa01C","Drug":"PF_04691502","minConc":1,"D1":-3.48,"D2":4.94,"D3":42.09,"D4":73.67,"D5":84.37,"IC50":108.35,"Slope":1.02,"max":84.37,"min":0,"Barcode":"H1FO4B-N1080","W5":"B2","W4":"C2","W3":"D2","W2":"E2","W1":"F2"},{"CellLine":"Pa01C","Drug":"SNS_032","minConc":1,"D1":-4.8,"D2":4.87,"D3":61.34,"D4":65.72,"D5":71.02,"IC50":38.65,"Slope":1.93,"max":71.02,"min":0,"Barcode":"H1FO4B-N1080","W5":"G9","W4":"H9","W3":"I9","W2":"J9","W1":"K9"},{"CellLine":"Pa01C","Drug":"Alvocidib","minConc":1,"D1":5.17,"D2":2.85,"D3":35.18,"D4":65.28,"D5":61.32,"IC50":97.2,"Slope":2.5,"max":65.28,"min":2.85,"Barcode":"H1FO4B-N1080","W5":"P9","W4":"O9","W3":"N9","W2":"M9","W1":"L9"},{"CellLine":"Pa01C","Drug":"KX2_391","minConc":1,"D1":-9.51,"D2":-14.25,"D3":49.62,"D4":58.93,"D5":61.35,"IC50":58.67,"Slope":2.5,"max":61.35,"min":0,"Barcode":"H1FO4B-N1080","W5":"F17","W4":"G17","W3":"H17","W2":"I17","W1":"J17"},{"CellLine":"Pa01C","Drug":"AZD8055","minConc":1,"D1":5.24,"D2":23.47,"D3":41.73,"D4":60.86,"D5":69.68,"IC50":41.1,"Slope":0.59,"max":69.68,"min":0,"Barcode":"H1FO4B-N1080","W5":"G23","W4":"H23","W3":"I23","W2":"J23","W1":"K23"},{"CellLine":"Pa01C","Drug":"PF_03758309","minConc":1,"D1":15.16,"D2":16.38,"D3":31.24,"D4":73.88,"D5":87.63,"IC50":288.89,"Slope":1.19,"max":87.63,"min":15.09,"Barcode":"H1FO4B-N2080","W5":"P5","W4":"O5","W3":"N5","W2":"M5","W1":"L5"},{"CellLine":"Pa01C","Drug":"AZD_8330","minConc":1,"D1":17.01,"D2":16.45,"D3":-0.72,"D4":11.95,"D5":5.67,"IC50":10000,"Slope":-0.28,"max":17.01,"min":0,"Barcode":"H1FO4B-N2080","W5":"F6","W4":"H6","W3":"I6","W2":"J6","W1":"K6"},{"CellLine":"Pa01C","Drug":"Rigosertib","minConc":1,"D1":-9.71,"D2":9.96,"D3":55.07,"D4":61.66,"D5":63.54,"IC50":30.18,"Slope":1.59,"max":63.54,"min":0,"Barcode":"H1FO4B-N2080","W5":"F8","W4":"G8","W3":"H8","W2":"J8","W1":"K8"},{"CellLine":"Pa01C","Drug":"Omipalisib","minConc":0.1,"D1":8.61,"D2":16.5,"D3":37.23,"D4":68.06,"D5":71.3,"IC50":10.69,"Slope":1.04,"max":71.3,"min":8.61,"Barcode":"H1FO4B-N2080","W5":"P17","W4":"O17","W3":"N17","W2":"M17","W1":"K17"},{"CellLine":"Pa01C","Drug":"UCN_01","minConc":1,"D1":-4.86,"D2":9.21,"D3":31.16,"D4":56.46,"D5":87.43,"IC50":285.56,"Slope":0.7,"max":87.43,"min":0,"Barcode":"H1FO4B-N2080","W5":"P21","W4":"O21","W3":"N21","W2":"M21","W1":"L21"},{"CellLine":"Pa01C","Drug":"Dinaciclib","minConc":0.1,"D1":-5.2,"D2":-2.94,"D3":39.58,"D4":65.65,"D5":72.47,"IC50":9.2,"Slope":1.81,"max":72.47,"min":0,"Barcode":"H1FO4B-N3080","W5":"A3","W4":"B3","W3":"C3","W2":"D3","W1":"E3"},{"CellLine":"Pa01C","Drug":"NVP_BGT226","minConc":0.1,"D1":4.63,"D2":-9.21,"D3":32.8,"D4":57.42,"D5":89.81,"IC50":33.66,"Slope":0.81,"max":89.81,"min":0,"Barcode":"H1FO4B-N3080","W5":"G12","W4":"H12","W3":"I12","W2":"J12","W1":"K12"},{"CellLine":"Pa01C","Drug":"Gedatolisib","minConc":0.1,"D1":3.46,"D2":-1.81,"D3":18.69,"D4":64.43,"D5":79.29,"IC50":28.03,"Slope":1.19,"max":79.29,"min":0,"Barcode":"H1FO4B-N3080","W5":"P12","W4":"O12","W3":"N12","W2":"M12","W1":"L12"},{"CellLine":"Pa01C","Drug":"PKI_402","minConc":0.25,"D1":-12.14,"D2":-2.29,"D3":25.98,"D4":68.78,"D5":78.32,"IC50":45.65,"Slope":1.27,"max":78.32,"min":0,"Barcode":"H1FO4B-N3080","W5":"A16","W4":"C16","W3":"D16","W2":"E16","W1":"F16"},{"CellLine":"Pa01C","Drug":"AT7519","minConc":1,"D1":-8.34,"D2":4.74,"D3":37.06,"D4":69.28,"D5":70.39,"IC50":91.25,"Slope":1.4,"max":70.39,"min":0,"Barcode":"H1FO4B-N3080","W5":"F22","W4":"G22","W3":"H22","W2":"I22","W1":"J22"},{"CellLine":"Pa01C","Drug":"GSK_461364","minConc":1,"D1":7.56,"D2":16.06,"D3":19.17,"D4":22.2,"D5":72.41,"IC50":1692.23,"Slope":2.31,"max":72.41,"min":7.56,"Barcode":"H1FO4B-N4080","W5":"A6","W4":"B6","W3":"C6","W2":"D6","W1":"E6"},{"CellLine":"Pa01C","Drug":"Daporinad","minConc":0.1,"D1":-5.93,"D2":18.9,"D3":88.85,"D4":96.75,"D5":94.23,"IC50":2.33,"Slope":1.7,"max":96.75,"min":0,"Barcode":"H1FO4B-N4080","W5":"A14","W4":"B14","W3":"C14","W2":"D14","W1":"E14"},{"CellLine":"Pa01C","Drug":"Mepacrine","minConc":5,"D1":0.89,"D2":3.83,"D3":16.7,"D4":80.28,"D5":99.47,"IC50":1717.08,"Slope":1.33,"max":99.47,"min":0.89,"Barcode":"H1FO4B-N4080","W5":"P20","W4":"O20","W3":"N20","W2":"M20","W1":"L20"},{"CellLine":"Pa01C","Drug":"Camptothecin","minConc":0.5,"D1":2.39,"D2":5.83,"D3":10.34,"D4":40.35,"D5":64.01,"IC50":307.21,"Slope":1.1,"max":64.01,"min":2.39,"Barcode":"H1FO4B-N5080","W5":"O3","W4":"N3","W3":"M3","W2":"L3","W1":"K3"},{"CellLine":"Pa01C","Drug":"Omacetaxine","minConc":1,"D1":3.2,"D2":4.71,"D3":52.91,"D4":80.19,"D5":79.75,"IC50":74.04,"Slope":2,"max":80.19,"min":3.2,"Barcode":"H1FO4B-N5080","W5":"A5","W4":"B5","W3":"C5","W2":"E5","W1":"F5"},{"CellLine":"Pa01C","Drug":"Vinorelbine","minConc":1,"D1":-1.29,"D2":21.56,"D3":61.1,"D4":65.41,"D5":92.19,"IC50":64.28,"Slope":0.61,"max":92.19,"min":0,"Barcode":"H1FO4B-N5080","W5":"P14","W4":"O14","W3":"N14","W2":"M14","W1":"L14"},{"CellLine":"Pa01C","Drug":"Docetaxel","minConc":0.1,"D1":-9.17,"D2":4.76,"D3":35.84,"D4":43.97,"D5":42.66,"IC50":3.96,"Slope":1.63,"max":43.97,"min":0,"Barcode":"H1FO4B-N5080","W5":"P17","W4":"O17","W3":"N17","W2":"M17","W1":"K17"},{"CellLine":"Pa01C","Drug":"Carfilzomib","minConc":0.1,"D1":1.82,"D2":8.41,"D3":75.2,"D4":95.85,"D5":97.75,"IC50":4.82,"Slope":1.62,"max":97.75,"min":1.53,"Barcode":"H1FO4B-N5080","W5":"A18","W4":"B18","W3":"C18","W2":"D18","W1":"E18"},{"CellLine":"Pa01C","Drug":"Patupilone","minConc":0.1,"D1":-7.05,"D2":15.44,"D3":39.03,"D4":41.79,"D5":37.72,"IC50":1.27,"Slope":2.3,"max":41.79,"min":0,"Barcode":"H1FO4B-N5080","W5":"F18","W4":"G18","W3":"H18","W2":"I18","W1":"J18"},{"CellLine":"Pa01C","Drug":"Plicamycin","minConc":1,"D1":-10.68,"D2":-7.27,"D3":9.25,"D4":52.22,"D5":81.54,"IC50":614.8,"Slope":1.27,"max":81.54,"min":0,"Barcode":"H1FO4B-N5080","W5":"P18","W4":"N18","W3":"M18","W2":"L18","W1":"K18"},{"CellLine":"Pa01C","Drug":"Vincristine","minConc":0.1,"D1":-3.49,"D2":-1.88,"D3":42.63,"D4":63.57,"D5":64.79,"IC50":7.71,"Slope":2.5,"max":64.79,"min":0,"Barcode":"H1FO4B-N5080","W5":"A19","W4":"B19","W3":"C19","W2":"D19","W1":"E19"},{"CellLine":"Pa01C","Drug":"Dactinomycin","minConc":0.1,"D1":-2.19,"D2":23.28,"D3":39.83,"D4":73.74,"D5":76.31,"IC50":5.72,"Slope":0.74,"max":76.31,"min":0,"Barcode":"H1FO4B-N5080","W5":"P19","W4":"O19","W3":"N19","W2":"M19","W1":"L19"},{"CellLine":"Pa01C","Drug":"Vinblastine","minConc":0.1,"D1":7.29,"D2":2.94,"D3":35.5,"D4":64.1,"D5":65.13,"IC50":9.57,"Slope":2.09,"max":65.13,"min":2.94,"Barcode":"H1FO4B-N5080","W5":"A21","W4":"B21","W3":"C21","W2":"D21","W1":"E21"},{"CellLine":"Pa01C","Drug":"Paclitaxel","minConc":0.1,"D1":4.81,"D2":8.2,"D3":32.05,"D4":45.05,"D5":41.86,"IC50":5.79,"Slope":1.38,"max":45.05,"min":4.77,"Barcode":"H1FO4B-N5080","W5":"F21","W4":"G21","W3":"H21","W2":"I21","W1":"J21"},{"CellLine":"Pa01C","Drug":"Oprozomib","minConc":0.25,"D1":2.77,"D2":1.09,"D3":18.15,"D4":81.4,"D5":98.21,"IC50":78.52,"Slope":1.36,"max":98.21,"min":1.09,"Barcode":"H1FO4B-N5080","W5":"A22","W4":"B22","W3":"C22","W2":"D22","W1":"E22"},{"CellLine":"Pa01C","Drug":"Bortezomib","minConc":0.1,"D1":-2.69,"D2":7.32,"D3":80.01,"D4":94.45,"D5":93.74,"IC50":3.91,"Slope":1.83,"max":94.45,"min":0,"Barcode":"H1FO4B-N5080","W5":"F22","W4":"G22","W3":"H22","W2":"I22","W1":"J22"},{"CellLine":"Pa01C","Drug":"Filanesib","minConc":0.1,"D1":12.54,"D2":1.17,"D3":30.35,"D4":28.14,"D5":30.85,"IC50":3.28,"Slope":2.5,"max":30.85,"min":1.17,"Barcode":"H1FO4B-N5080","W5":"G23","W4":"H23","W3":"I23","W2":"J23","W1":"K23"},{"CellLine":"Pa01C","Drug":"CUDC_907","minConc":1,"D1":-15.34,"D2":-18.68,"D3":45.3,"D4":93.77,"D5":93.77,"IC50":103.13,"Slope":2.5,"max":93.77,"min":0,"Barcode":"H1FO4B-N6080","W5":"F17","W4":"G17","W3":"H17","W2":"I17","W1":"J17"},{"CellLine":"Pa01C","Drug":"YM155","minConc":1,"D1":38.46,"D2":96.53,"D3":101.94,"D4":102.9,"D5":100.38,"IC50":3.16,"Slope":2.5,"max":100,"min":35,"Barcode":"H1FO4B-N6080","W5":"F18","W4":"G18","W3":"H18","W2":"I18","W1":"J18"},{"CellLine":"Pa01C","Drug":"AT101","minConc":10,"D1":2.1,"D2":-9.77,"D3":18.27,"D4":90.74,"D5":104.76,"IC50":2473.07,"Slope":1.7,"max":100,"min":0,"Barcode":"H1FO4B-N6080","W5":"A21","W4":"B21","W3":"C21","W2":"D21","W1":"E21"},{"CellLine":"Pa01C","Drug":"Selinexor","minConc":1,"D1":4.94,"D2":5.17,"D3":41.02,"D4":70.23,"D5":68.7,"IC50":91.89,"Slope":2.5,"max":70.23,"min":4.93,"Barcode":"H1FO4B-N6080","W5":"P21","W4":"O21","W3":"N21","W2":"M21","W1":"L21"},{"CellLine":"Pa01C","Drug":"BIIB021","minConc":1,"D1":-19.86,"D2":-6.2,"D3":30.42,"D4":52.35,"D5":63.8,"IC50":129.44,"Slope":1.22,"max":63.8,"min":0,"Barcode":"H1FO4B-N7080","W5":"P13","W4":"N13","W3":"M13","W2":"L13","W1":"K13"},{"CellLine":"Pa01C","Drug":"Tanespimycin","minConc":1,"D1":-1.29,"D2":0.42,"D3":42.04,"D4":57.89,"D5":59.14,"IC50":68.55,"Slope":2.38,"max":59.14,"min":0,"Barcode":"H1FO4B-N7080","W5":"A14","W4":"B14","W3":"C14","W2":"D14","W1":"E14"},{"CellLine":"Pa01C","Drug":"CUDC_305","minConc":1,"D1":-9.74,"D2":11.02,"D3":29.16,"D4":55.65,"D5":61.6,"IC50":101.3,"Slope":0.91,"max":61.6,"min":0,"Barcode":"H1FO4B-N7080","W5":"A15","W4":"B15","W3":"C15","W2":"D15","W1":"F15"},{"CellLine":"Pa01C","Drug":"Luminespib","minConc":0.1,"D1":-7.75,"D2":2.04,"D3":35.86,"D4":55.05,"D5":54.64,"IC50":7.03,"Slope":1.79,"max":55.05,"min":0,"Barcode":"H1FO4B-N7080","W5":"G15","W4":"H15","W3":"I15","W2":"J15","W1":"K15"},{"CellLine":"Pa01C","Drug":"Ganetespib","minConc":0.1,"D1":4.7,"D2":-14.89,"D3":32.74,"D4":52.86,"D5":52.25,"IC50":8.31,"Slope":2.5,"max":52.86,"min":0,"Barcode":"H1FO4B-N7080","W5":"A16","W4":"C16","W3":"D16","W2":"E16","W1":"F16"},{"CellLine":"Pa01C","Drug":"Stattic","minConc":5,"D1":-0.73,"D2":-14.3,"D3":-4.54,"D4":94.6,"D5":103.04,"IC50":1938.7,"Slope":2.5,"max":100,"min":0,"Barcode":"H1FO4B-N7080","W5":"F21","W4":"G21","W3":"H21","W2":"I21","W1":"J21"},{"CellLine":"Pa01C","Drug":"ONX_0914","minConc":1,"D1":-6.81,"D2":5.58,"D3":14.84,"D4":98.72,"D5":96.99,"IC50":198.4,"Slope":2.5,"max":98.72,"min":0,"Barcode":"H1FO4B-N7080","W5":"G23","W4":"H23","W3":"I23","W2":"J23","W1":"K23"},{"CellLine":"Pa01C","Drug":"SN_38","minConc":1,"D1":17.71,"D2":16.65,"D3":33.46,"D4":69.42,"D5":67.22,"IC50":135.53,"Slope":2.5,"max":69.42,"min":16.65,"Barcode":"H1FO4B-N8080","W5":"G2","W4":"H2","W3":"I2","W2":"J2","W1":"K2"},{"CellLine":"Pa01C","Drug":"Cabazitaxel","minConc":0.1,"D1":10.89,"D2":15.57,"D3":34.07,"D4":41.17,"D5":40.26,"IC50":3.78,"Slope":1.24,"max":41.17,"min":10.59,"Barcode":"H1FO4B-N8080","W5":"A3","W4":"B3","W3":"C3","W2":"D3","W1":"E3"},{"CellLine":"Pa01C","Drug":"OTSSP167","minConc":0.1,"D1":4.06,"D2":-1.04,"D3":16.84,"D4":74.29,"D5":81.51,"IC50":23.21,"Slope":1.61,"max":81.51,"min":0,"Barcode":"H1FO4B-N8080","W5":"A5","W4":"B5","W3":"C5","W2":"E5","W1":"F5"},{"CellLine":"Pa01C","Drug":"Prexasertib","minConc":1,"D1":11.71,"D2":34.04,"D3":73.65,"D4":65.38,"D5":39.13,"IC50":12.55,"Slope":2.5,"max":73.65,"min":11.6,"Barcode":"H1FO4B-N8080","W5":"P6","W4":"O6","W3":"N6","W2":"M6","W1":"L6"},{"CellLine":"Pa01C","Drug":"Romidepsin","minConc":0.1,"D1":1.93,"D2":0.26,"D3":75.16,"D4":88.83,"D5":87.11,"IC50":5.1,"Slope":2.5,"max":88.83,"min":0.26,"Barcode":"H1FO4B-N8080","W5":"F18","W4":"G18","W3":"H18","W2":"I18","W1":"J18"},{"CellLine":"Pa01C","Drug":"Sabutoclax","minConc":2.5,"D1":4.04,"D2":2.21,"D3":48.07,"D4":59.44,"D5":78.09,"IC50":229.6,"Slope":0.83,"max":78.09,"min":0,"Barcode":"H1FO4B-N8080","W5":"F19","W4":"G19","W3":"I19","W2":"J19","W1":"K19"},{"CellLine":"Pa01C","Drug":"BCI","minConc":5,"D1":0.88,"D2":1.74,"D3":7.93,"D4":90.52,"D5":105.42,"IC50":1705.89,"Slope":2.1,"max":100,"min":0.88,"Barcode":"H1FO4B-N8080","W5":"A22","W4":"B22","W3":"C22","W2":"D22","W1":"E22"},{"CellLine":"MDA-PATC53","Drug":"PF_04691502","minConc":1,"D1":-0.62,"D2":8.98,"D3":39.6,"D4":74.25,"D5":89.18,"IC50":133.96,"Slope":0.85,"max":89.18,"min":0,"Barcode":"H1FO4B-N1089","W5":"B2","W4":"C2","W3":"D2","W2":"E2","W1":"F2"},{"CellLine":"MDA-PATC53","Drug":"SNS_032","minConc":1,"D1":4.87,"D2":11.64,"D3":49.9,"D4":82.34,"D5":83.34,"IC50":76.91,"Slope":1.26,"max":83.34,"min":4.87,"Barcode":"H1FO4B-N1089","W5":"G9","W4":"H9","W3":"I9","W2":"J9","W1":"K9"},{"CellLine":"MDA-PATC53","Drug":"Alvocidib","minConc":1,"D1":4.77,"D2":5.36,"D3":32.54,"D4":81.53,"D5":86.8,"IC50":156.76,"Slope":1.46,"max":86.8,"min":4.34,"Barcode":"H1FO4B-N1089","W5":"P9","W4":"O9","W3":"N9","W2":"M9","W1":"L9"},{"CellLine":"MDA-PATC53","Drug":"KX2_391","minConc":1,"D1":-0.95,"D2":-1.96,"D3":53.44,"D4":69.76,"D5":75.06,"IC50":69.86,"Slope":2.5,"max":75.06,"min":0,"Barcode":"H1FO4B-N1089","W5":"F17","W4":"G17","W3":"H17","W2":"I17","W1":"J17"},{"CellLine":"MDA-PATC53","Drug":"AZD8055","minConc":1,"D1":9.33,"D2":23.12,"D3":8.07,"D4":65.45,"D5":72.42,"IC50":439.94,"Slope":2.5,"max":72.42,"min":8.07,"Barcode":"H1FO4B-N1089","W5":"G23","W4":"H23","W3":"I23","W2":"J23","W1":"K23"},{"CellLine":"MDA-PATC53","Drug":"PF_03758309","minConc":1,"D1":14.43,"D2":24.08,"D3":43.43,"D4":63,"D5":96.22,"IC50":322.44,"Slope":0.64,"max":96.22,"min":14.43,"Barcode":"H1FO4B-N2089","W5":"P5","W4":"O5","W3":"N5","W2":"M5","W1":"L5"},{"CellLine":"MDA-PATC53","Drug":"AZD_8330","minConc":1,"D1":22.76,"D2":37.8,"D3":50.21,"D4":56.79,"D5":54.5,"IC50":7.29,"Slope":0.68,"max":56.79,"min":14.02,"Barcode":"H1FO4B-N2089","W5":"F6","W4":"H6","W3":"I6","W2":"J6","W1":"K6"},{"CellLine":"MDA-PATC53","Drug":"Rigosertib","minConc":1,"D1":9.78,"D2":9.52,"D3":61.71,"D4":69.66,"D5":69.38,"IC50":47.3,"Slope":2.5,"max":69.66,"min":9.05,"Barcode":"H1FO4B-N2089","W5":"F8","W4":"G8","W3":"H8","W2":"J8","W1":"K8"},{"CellLine":"MDA-PATC53","Drug":"Omipalisib","minConc":0.1,"D1":5.21,"D2":0.07,"D3":39.93,"D4":71.75,"D5":71.97,"IC50":9.16,"Slope":2.5,"max":71.97,"min":0.07,"Barcode":"H1FO4B-N2089","W5":"P17","W4":"O17","W3":"N17","W2":"M17","W1":"K17"},{"CellLine":"MDA-PATC53","Drug":"UCN_01","minConc":1,"D1":10.33,"D2":15.19,"D3":68.04,"D4":80.91,"D5":98.78,"IC50":59.44,"Slope":0.83,"max":98.78,"min":3.84,"Barcode":"H1FO4B-N2089","W5":"P21","W4":"O21","W3":"N21","W2":"M21","W1":"L21"},{"CellLine":"MDA-PATC53","Drug":"Dinaciclib","minConc":0.1,"D1":-4.1,"D2":-3.07,"D3":41.7,"D4":86.26,"D5":90,"IC50":10.8,"Slope":2.05,"max":90,"min":0,"Barcode":"H1FO4B-N3089","W5":"A3","W4":"B3","W3":"C3","W2":"D3","W1":"E3"},{"CellLine":"MDA-PATC53","Drug":"NVP_BGT226","minConc":0.1,"D1":3.76,"D2":15.66,"D3":3.86,"D4":-0.86,"D5":88.97,"IC50":322.14,"Slope":2.5,"max":88.97,"min":0,"Barcode":"H1FO4B-N3089","W5":"G12","W4":"H12","W3":"I12","W2":"J12","W1":"K12"},{"CellLine":"MDA-PATC53","Drug":"Gedatolisib","minConc":0.1,"D1":3.78,"D2":5.38,"D3":9.29,"D4":57.51,"D5":74.17,"IC50":47.55,"Slope":1.58,"max":74.17,"min":3.78,"Barcode":"H1FO4B-N3089","W5":"P12","W4":"O12","W3":"N12","W2":"M12","W1":"L12"},{"CellLine":"MDA-PATC53","Drug":"PKI_402","minConc":0.25,"D1":-6.1,"D2":-0.59,"D3":15.87,"D4":65.69,"D5":80.36,"IC50":76.23,"Slope":1.28,"max":80.36,"min":0,"Barcode":"H1FO4B-N3089","W5":"A16","W4":"C16","W3":"D16","W2":"E16","W1":"F16"},{"CellLine":"MDA-PATC53","Drug":"AT7519","minConc":1,"D1":-8.87,"D2":8.31,"D3":11.83,"D4":91.96,"D5":86.79,"IC50":212.04,"Slope":2.5,"max":91.96,"min":0,"Barcode":"H1FO4B-N3089","W5":"F22","W4":"G22","W3":"H22","W2":"I22","W1":"J22"},{"CellLine":"MDA-PATC53","Drug":"GSK_461364","minConc":1,"D1":4.89,"D2":28.2,"D3":38.51,"D4":48.85,"D5":82.91,"IC50":112.19,"Slope":0.48,"max":82.91,"min":4.89,"Barcode":"H1FO4B-N4089","W5":"A6","W4":"B6","W3":"C6","W2":"D6","W1":"E6"},{"CellLine":"MDA-PATC53","Drug":"Daporinad","minConc":0.1,"D1":-3.94,"D2":-13.9,"D3":49.37,"D4":96.61,"D5":96.09,"IC50":9.85,"Slope":2.5,"max":96.61,"min":0,"Barcode":"H1FO4B-N4089","W5":"A14","W4":"B14","W3":"C14","W2":"D14","W1":"E14"},{"CellLine":"MDA-PATC53","Drug":"Mepacrine","minConc":5,"D1":-2.92,"D2":-1.23,"D3":11.28,"D4":96.49,"D5":103.12,"IC50":1207.14,"Slope":2.34,"max":100,"min":0,"Barcode":"H1FO4B-N4089","W5":"P20","W4":"O20","W3":"N20","W2":"M20","W1":"L20"},{"CellLine":"MDA-PATC53","Drug":"Camptothecin","minConc":0.5,"D1":0.12,"D2":0.31,"D3":28.01,"D4":72.13,"D5":87.97,"IC50":108.94,"Slope":1.05,"max":87.97,"min":0,"Barcode":"H1FO4B-N5089","W5":"O3","W4":"N3","W3":"M3","W2":"L3","W1":"K3"},{"CellLine":"MDA-PATC53","Drug":"Omacetaxine","minConc":1,"D1":2.92,"D2":-7.39,"D3":49.27,"D4":91.3,"D5":96.67,"IC50":98.72,"Slope":2.5,"max":96.67,"min":0,"Barcode":"H1FO4B-N5089","W5":"A5","W4":"B5","W3":"C5","W2":"E5","W1":"F5"},{"CellLine":"MDA-PATC53","Drug":"Vinorelbine","minConc":1,"D1":3.03,"D2":41.28,"D3":63.92,"D4":73.57,"D5":93.51,"IC50":27.64,"Slope":0.55,"max":93.51,"min":0,"Barcode":"H1FO4B-N5089","W5":"P14","W4":"O14","W3":"N14","W2":"M14","W1":"L14"},{"CellLine":"MDA-PATC53","Drug":"Docetaxel","minConc":0.1,"D1":-4.09,"D2":12.04,"D3":32.9,"D4":51.77,"D5":51.59,"IC50":4.78,"Slope":0.96,"max":51.77,"min":0,"Barcode":"H1FO4B-N5089","W5":"P17","W4":"O17","W3":"N17","W2":"M17","W1":"K17"},{"CellLine":"MDA-PATC53","Drug":"Carfilzomib","minConc":0.1,"D1":0.19,"D2":10.28,"D3":96.9,"D4":100.39,"D5":103.4,"IC50":2.44,"Slope":2.44,"max":100,"min":0.15,"Barcode":"H1FO4B-N5089","W5":"A18","W4":"B18","W3":"C18","W2":"D18","W1":"E18"},{"CellLine":"MDA-PATC53","Drug":"Patupilone","minConc":0.1,"D1":-3.53,"D2":19.55,"D3":43.81,"D4":47.96,"D5":47.24,"IC50":1.33,"Slope":1.51,"max":47.96,"min":0,"Barcode":"H1FO4B-N5089","W5":"F18","W4":"G18","W3":"H18","W2":"I18","W1":"J18"},{"CellLine":"MDA-PATC53","Drug":"Plicamycin","minConc":1,"D1":-6.88,"D2":-4.69,"D3":33.26,"D4":90.36,"D5":93.99,"IC50":137.5,"Slope":1.93,"max":93.99,"min":0,"Barcode":"H1FO4B-N5089","W5":"P18","W4":"N18","W3":"M18","W2":"L18","W1":"K18"},{"CellLine":"MDA-PATC53","Drug":"Vincristine","minConc":0.1,"D1":-5.75,"D2":16.57,"D3":64.53,"D4":77.05,"D5":76.71,"IC50":2.79,"Slope":1.34,"max":77.05,"min":0,"Barcode":"H1FO4B-N5089","W5":"A19","W4":"B19","W3":"C19","W2":"D19","W1":"E19"},{"CellLine":"MDA-PATC53","Drug":"Dactinomycin","minConc":0.1,"D1":-1.55,"D2":30.99,"D3":64.25,"D4":90.66,"D5":92.9,"IC50":3.03,"Slope":0.82,"max":92.9,"min":0,"Barcode":"H1FO4B-N5089","W5":"P19","W4":"O19","W3":"N19","W2":"M19","W1":"L19"},{"CellLine":"MDA-PATC53","Drug":"Vinblastine","minConc":0.1,"D1":4.73,"D2":5.56,"D3":54.57,"D4":75.52,"D5":79.28,"IC50":6.73,"Slope":1.78,"max":79.28,"min":3.93,"Barcode":"H1FO4B-N5089","W5":"A21","W4":"B21","W3":"C21","W2":"D21","W1":"E21"},{"CellLine":"MDA-PATC53","Drug":"Paclitaxel","minConc":0.1,"D1":3.15,"D2":5.1,"D3":38.07,"D4":56.06,"D5":50.81,"IC50":6.8,"Slope":1.73,"max":56.06,"min":3.15,"Barcode":"H1FO4B-N5089","W5":"F21","W4":"G21","W3":"H21","W2":"I21","W1":"J21"},{"CellLine":"MDA-PATC53","Drug":"Oprozomib","minConc":0.25,"D1":2.27,"D2":4.13,"D3":36.13,"D4":98.85,"D5":104.84,"IC50":33.84,"Slope":2.09,"max":100,"min":2.27,"Barcode":"H1FO4B-N5089","W5":"A22","W4":"B22","W3":"C22","W2":"D22","W1":"E22"},{"CellLine":"MDA-PATC53","Drug":"Bortezomib","minConc":0.1,"D1":1.8,"D2":-7.36,"D3":98.41,"D4":100.79,"D5":98.73,"IC50":3.65,"Slope":2.5,"max":100,"min":0,"Barcode":"H1FO4B-N5089","W5":"F22","W4":"G22","W3":"H22","W2":"I22","W1":"J22"},{"CellLine":"MDA-PATC53","Drug":"Filanesib","minConc":0.1,"D1":0.14,"D2":9.25,"D3":39.88,"D4":50.39,"D5":51.51,"IC50":3.58,"Slope":1.2,"max":51.51,"min":0,"Barcode":"H1FO4B-N5089","W5":"G23","W4":"H23","W3":"I23","W2":"J23","W1":"K23"},{"CellLine":"MDA-PATC53","Drug":"CUDC_907","minConc":1,"D1":3.53,"D2":33.68,"D3":88.95,"D4":102.24,"D5":97.71,"IC50":17.75,"Slope":1.23,"max":100,"min":0.74,"Barcode":"H1FO4B-N6089","W5":"F17","W4":"G17","W3":"H17","W2":"I17","W1":"J17"},{"CellLine":"MDA-PATC53","Drug":"YM155","minConc":1,"D1":0.66,"D2":20.35,"D3":82.38,"D4":102.65,"D5":102.11,"IC50":29.27,"Slope":1.29,"max":100,"min":0,"Barcode":"H1FO4B-N6089","W5":"F18","W4":"G18","W3":"H18","W2":"I18","W1":"J18"},{"CellLine":"MDA-PATC53","Drug":"AT101","minConc":10,"D1":0.76,"D2":2.05,"D3":42.17,"D4":89.4,"D5":105.84,"IC50":1346.14,"Slope":1.19,"max":100,"min":0,"Barcode":"H1FO4B-N6089","W5":"A21","W4":"B21","W3":"C21","W2":"D21","W1":"E21"},{"CellLine":"MDA-PATC53","Drug":"Selinexor","minConc":1,"D1":1.1,"D2":15.45,"D3":38.37,"D4":85.02,"D5":96.92,"IC50":141.04,"Slope":0.88,"max":96.92,"min":1.1,"Barcode":"H1FO4B-N6089","W5":"P21","W4":"O21","W3":"N21","W2":"M21","W1":"L21"},{"CellLine":"MDA-PATC53","Drug":"BIIB021","minConc":1,"D1":11.05,"D2":7.42,"D3":27.77,"D4":71.72,"D5":79.99,"IC50":206.33,"Slope":1.32,"max":79.99,"min":7.42,"Barcode":"H1FO4B-N7089","W5":"P13","W4":"N13","W3":"M13","W2":"L13","W1":"K13"},{"CellLine":"MDA-PATC53","Drug":"Tanespimycin","minConc":1,"D1":-8.91,"D2":-2.25,"D3":26.16,"D4":78.47,"D5":80.84,"IC50":145.38,"Slope":1.99,"max":80.84,"min":0,"Barcode":"H1FO4B-N7089","W5":"A14","W4":"B14","W3":"C14","W2":"D14","W1":"E14"},{"CellLine":"MDA-PATC53","Drug":"CUDC_305","minConc":1,"D1":0.87,"D2":-6.36,"D3":9.28,"D4":78.92,"D5":82.44,"IC50":249.89,"Slope":2.27,"max":82.44,"min":0,"Barcode":"H1FO4B-N7089","W5":"A15","W4":"B15","W3":"C15","W2":"D15","W1":"F15"},{"CellLine":"MDA-PATC53","Drug":"Luminespib","minConc":0.1,"D1":-1.33,"D2":-10.22,"D3":33.44,"D4":77.47,"D5":75.74,"IC50":11.19,"Slope":2.5,"max":77.47,"min":0,"Barcode":"H1FO4B-N7089","W5":"G15","W4":"H15","W3":"I15","W2":"J15","W1":"K15"},{"CellLine":"MDA-PATC53","Drug":"Ganetespib","minConc":0.1,"D1":-10.49,"D2":-0.48,"D3":6.44,"D4":78.56,"D5":81.64,"IC50":27.01,"Slope":2.47,"max":81.64,"min":0,"Barcode":"H1FO4B-N7089","W5":"A16","W4":"C16","W3":"D16","W2":"E16","W1":"F16"},{"CellLine":"MDA-PATC53","Drug":"Stattic","minConc":5,"D1":5.21,"D2":1.67,"D3":-9.33,"D4":13.81,"D5":102.76,"IC50":10187.93,"Slope":2.5,"max":100,"min":0,"Barcode":"H1FO4B-N7089","W5":"F21","W4":"G21","W3":"H21","W2":"I21","W1":"J21"},{"CellLine":"MDA-PATC53","Drug":"ONX_0914","minConc":1,"D1":11.17,"D2":1.47,"D3":-0.11,"D4":102.67,"D5":101.56,"IC50":301.85,"Slope":2.5,"max":100,"min":0,"Barcode":"H1FO4B-N7089","W5":"G23","W4":"H23","W3":"I23","W2":"J23","W1":"K23"},{"CellLine":"MDA-PATC53","Drug":"SN_38","minConc":1,"D1":2.58,"D2":32.95,"D3":68.96,"D4":84.94,"D5":87.21,"IC50":19.57,"Slope":0.88,"max":87.21,"min":0,"Barcode":"H1FO4B-N8089","W5":"G2","W4":"H2","W3":"I2","W2":"J2","W1":"K2"},{"CellLine":"MDA-PATC53","Drug":"Cabazitaxel","minConc":0.1,"D1":1.83,"D2":22.16,"D3":45.79,"D4":46.85,"D5":47.78,"IC50":1.12,"Slope":1.41,"max":47.78,"min":0.31,"Barcode":"H1FO4B-N8089","W5":"A3","W4":"B3","W3":"C3","W2":"D3","W1":"E3"},{"CellLine":"MDA-PATC53","Drug":"OTSSP167","minConc":0.1,"D1":2.92,"D2":-1.45,"D3":18.41,"D4":84.46,"D5":97.66,"IC50":27.59,"Slope":1.45,"max":97.66,"min":0,"Barcode":"H1FO4B-N8089","W5":"A5","W4":"B5","W3":"C5","W2":"E5","W1":"F5"},{"CellLine":"MDA-PATC53","Drug":"Prexasertib","minConc":1,"D1":11.56,"D2":80.88,"D3":89.95,"D4":83.02,"D5":39.87,"IC50":2.93,"Slope":1.78,"max":89.95,"min":0,"Barcode":"H1FO4B-N8089","W5":"P6","W4":"O6","W3":"N6","W2":"M6","W1":"L6"},{"CellLine":"MDA-PATC53","Drug":"Romidepsin","minConc":0.1,"D1":-5.92,"D2":19.85,"D3":99.83,"D4":102.42,"D5":99.68,"IC50":1.75,"Slope":2.5,"max":100,"min":0,"Barcode":"H1FO4B-N8089","W5":"F18","W4":"G18","W3":"H18","W2":"I18","W1":"J18"},{"CellLine":"MDA-PATC53","Drug":"Sabutoclax","minConc":2.5,"D1":-3.39,"D2":-2.66,"D3":26.73,"D4":79.66,"D5":95.85,"IC50":596.08,"Slope":1.17,"max":95.85,"min":0,"Barcode":"H1FO4B-N8089","W5":"F19","W4":"G19","W3":"I19","W2":"J19","W1":"K19"},{"CellLine":"MDA-PATC53","Drug":"BCI","minConc":5,"D1":-2.04,"D2":-3.23,"D3":2.5,"D4":104.41,"D5":104.02,"IC50":1390.86,"Slope":2.5,"max":100,"min":0,"Barcode":"H1FO4B-N8089","W5":"A22","W4":"B22","W3":"C22","W2":"D22","W1":"E22"},{"CellLine":"Pa04C","Drug":"PF_04691502","minConc":1,"D1":5.4,"D2":9.99,"D3":9.08,"D4":56.6,"D5":91.99,"IC50":787.18,"Slope":1.58,"max":91.99,"min":5.4,"Barcode":"H1FO4B-N1100","W5":"B2","W4":"C2","W3":"D2","W2":"E2","W1":"F2"},{"CellLine":"Pa04C","Drug":"SNS_032","minConc":1,"D1":3.32,"D2":13.26,"D3":46.84,"D4":90.85,"D5":86.49,"IC50":93.15,"Slope":1.17,"max":90.85,"min":3.32,"Barcode":"H1FO4B-N1100","W5":"G9","W4":"H9","W3":"I9","W2":"J9","W1":"K9"},{"CellLine":"Pa04C","Drug":"Alvocidib","minConc":1,"D1":4.52,"D2":-1.4,"D3":25.21,"D4":78.19,"D5":91.44,"IC50":225.8,"Slope":1.23,"max":91.44,"min":0,"Barcode":"H1FO4B-N1100","W5":"P9","W4":"O9","W3":"N9","W2":"M9","W1":"L9"},{"CellLine":"Pa04C","Drug":"KX2_391","minConc":1,"D1":3.77,"D2":3.12,"D3":41.72,"D4":61.12,"D5":62.05,"IC50":77.42,"Slope":2.5,"max":62.05,"min":3.12,"Barcode":"H1FO4B-N1100","W5":"F17","W4":"G17","W3":"H17","W2":"I17","W1":"J17"},{"CellLine":"Pa04C","Drug":"AZD8055","minConc":1,"D1":0.53,"D2":1.91,"D3":10.05,"D4":28.72,"D5":44.23,"IC50":456.71,"Slope":0.91,"max":44.23,"min":0.53,"Barcode":"H1FO4B-N1100","W5":"G23","W4":"H23","W3":"I23","W2":"J23","W1":"K23"},{"CellLine":"Pa04C","Drug":"PF_03758309","minConc":1,"D1":12.36,"D2":14.44,"D3":35.37,"D4":65.06,"D5":91.86,"IC50":338.13,"Slope":0.8,"max":91.86,"min":11.3,"Barcode":"H1FO4B-N2100","W5":"P5","W4":"O5","W3":"N5","W2":"M5","W1":"L5"},{"CellLine":"Pa04C","Drug":"AZD_8330","minConc":1,"D1":21.83,"D2":32.95,"D3":57.48,"D4":72.09,"D5":72.91,"IC50":36.15,"Slope":0.93,"max":72.91,"min":20.24,"Barcode":"H1FO4B-N2100","W5":"F6","W4":"H6","W3":"I6","W2":"J6","W1":"K6"},{"CellLine":"Pa04C","Drug":"Rigosertib","minConc":1,"D1":12.97,"D2":9.83,"D3":43.19,"D4":61.59,"D5":63.75,"IC50":79.12,"Slope":2.04,"max":63.75,"min":9.83,"Barcode":"H1FO4B-N2100","W5":"F8","W4":"G8","W3":"H8","W2":"J8","W1":"K8"},{"CellLine":"Pa04C","Drug":"Omipalisib","minConc":0.1,"D1":-15.32,"D2":-2.49,"D3":4.8,"D4":25.68,"D5":43.95,"IC50":72.78,"Slope":1.24,"max":43.95,"min":0,"Barcode":"H1FO4B-N2100","W5":"P17","W4":"O17","W3":"N17","W2":"M17","W1":"K17"},{"CellLine":"Pa04C","Drug":"UCN_01","minConc":1,"D1":-2.32,"D2":12.4,"D3":17.33,"D4":44.24,"D5":87.88,"IC50":701.03,"Slope":0.81,"max":87.88,"min":0,"Barcode":"H1FO4B-N2100","W5":"P21","W4":"O21","W3":"N21","W2":"M21","W1":"L21"},{"CellLine":"Pa04C","Drug":"Dinaciclib","minConc":0.1,"D1":2.4,"D2":6.17,"D3":45.23,"D4":89.9,"D5":90.4,"IC50":10.24,"Slope":1.62,"max":90.4,"min":2.4,"Barcode":"H1FO4B-N3100","W5":"A3","W4":"B3","W3":"C3","W2":"D3","W1":"E3"},{"CellLine":"Pa04C","Drug":"NVP_BGT226","minConc":0.1,"D1":8.63,"D2":3.66,"D3":13.11,"D4":37.48,"D5":96.99,"IC50":136.32,"Slope":1.7,"max":96.99,"min":3.66,"Barcode":"H1FO4B-N3100","W5":"G12","W4":"H12","W3":"I12","W2":"J12","W1":"K12"},{"CellLine":"Pa04C","Drug":"Gedatolisib","minConc":0.1,"D1":8.44,"D2":-0.28,"D3":5.64,"D4":17.42,"D5":55.78,"IC50":147.1,"Slope":1.99,"max":55.78,"min":0,"Barcode":"H1FO4B-N3100","W5":"P12","W4":"O12","W3":"N12","W2":"M12","W1":"L12"},{"CellLine":"Pa04C","Drug":"PKI_402","minConc":0.25,"D1":8.48,"D2":2.59,"D3":-9.47,"D4":26.92,"D5":60.32,"IC50":273.26,"Slope":2.5,"max":60.32,"min":0,"Barcode":"H1FO4B-N3100","W5":"A16","W4":"C16","W3":"D16","W2":"E16","W1":"F16"},{"CellLine":"Pa04C","Drug":"AT7519","minConc":1,"D1":0.61,"D2":8.25,"D3":21.76,"D4":71.3,"D5":89.4,"IC50":280.17,"Slope":1.05,"max":89.4,"min":0.61,"Barcode":"H1FO4B-N3100","W5":"F22","W4":"G22","W3":"H22","W2":"I22","W1":"J22"},{"CellLine":"Pa04C","Drug":"GSK_461364","minConc":1,"D1":12.06,"D2":30.05,"D3":36.39,"D4":21.48,"D5":62.03,"IC50":476.92,"Slope":0.35,"max":62.03,"min":12.06,"Barcode":"H1FO4B-N4100","W5":"A6","W4":"B6","W3":"C6","W2":"D6","W1":"E6"},{"CellLine":"Pa04C","Drug":"Daporinad","minConc":0.1,"D1":8.59,"D2":2.52,"D3":47.94,"D4":59.9,"D5":54.19,"IC50":5.87,"Slope":2.5,"max":59.9,"min":2.52,"Barcode":"H1FO4B-N4100","W5":"A14","W4":"B14","W3":"C14","W2":"D14","W1":"E14"},{"CellLine":"Pa04C","Drug":"Mepacrine","minConc":5,"D1":-6.16,"D2":8.64,"D3":16.43,"D4":92.05,"D5":102.05,"IC50":1252.32,"Slope":1.74,"max":100,"min":0,"Barcode":"H1FO4B-N4100","W5":"P20","W4":"O20","W3":"N20","W2":"M20","W1":"L20"},{"CellLine":"Pa04C","Drug":"Camptothecin","minConc":0.5,"D1":4.87,"D2":9.12,"D3":16.96,"D4":52.09,"D5":91.64,"IC50":376.32,"Slope":1.04,"max":91.64,"min":4.87,"Barcode":"H1FO4B-N5100","W5":"O3","W4":"N3","W3":"M3","W2":"L3","W1":"K3"},{"CellLine":"Pa04C","Drug":"Omacetaxine","minConc":1,"D1":3.51,"D2":22.16,"D3":75.61,"D4":86.42,"D5":91.51,"IC50":27.83,"Slope":1.16,"max":91.51,"min":1.45,"Barcode":"H1FO4B-N5100","W5":"A5","W4":"B5","W3":"C5","W2":"E5","W1":"F5"},{"CellLine":"Pa04C","Drug":"Vinorelbine","minConc":1,"D1":14.11,"D2":35.98,"D3":57.02,"D4":60.07,"D5":89.08,"IC50":36.05,"Slope":0.4,"max":89.08,"min":0,"Barcode":"H1FO4B-N5100","W5":"P14","W4":"O14","W3":"N14","W2":"M14","W1":"L14"},{"CellLine":"Pa04C","Drug":"Docetaxel","minConc":0.1,"D1":7.12,"D2":14.78,"D3":33.81,"D4":30.35,"D5":26.12,"IC50":1.44,"Slope":2.5,"max":33.81,"min":7.08,"Barcode":"H1FO4B-N5100","W5":"P17","W4":"O17","W3":"N17","W2":"M17","W1":"K17"},{"CellLine":"Pa04C","Drug":"Carfilzomib","minConc":0.1,"D1":4.68,"D2":6.07,"D3":89.71,"D4":100.11,"D5":102.42,"IC50":4.31,"Slope":2.5,"max":100,"min":4.17,"Barcode":"H1FO4B-N5100","W5":"A18","W4":"B18","W3":"C18","W2":"D18","W1":"E18"},{"CellLine":"Pa04C","Drug":"Patupilone","minConc":0.1,"D1":-2.81,"D2":16.93,"D3":39.58,"D4":38.02,"D5":31.4,"IC50":1.12,"Slope":2.5,"max":39.58,"min":0,"Barcode":"H1FO4B-N5100","W5":"F18","W4":"G18","W3":"H18","W2":"I18","W1":"J18"},{"CellLine":"Pa04C","Drug":"Plicamycin","minConc":1,"D1":-6.22,"D2":-0.16,"D3":-5.23,"D4":60.56,"D5":95.33,"IC50":802.8,"Slope":2.5,"max":95.33,"min":0,"Barcode":"H1FO4B-N5100","W5":"P18","W4":"N18","W3":"M18","W2":"L18","W1":"K18"},{"CellLine":"Pa04C","Drug":"Vincristine","minConc":0.1,"D1":12.17,"D2":-2.13,"D3":51.26,"D4":58.76,"D5":60.64,"IC50":5.16,"Slope":2.5,"max":60.64,"min":0,"Barcode":"H1FO4B-N5100","W5":"A19","W4":"B19","W3":"C19","W2":"D19","W1":"E19"},{"CellLine":"Pa04C","Drug":"Dactinomycin","minConc":0.1,"D1":-1.11,"D2":4.38,"D3":19.07,"D4":93.13,"D5":92.15,"IC50":17.16,"Slope":2.5,"max":93.13,"min":0,"Barcode":"H1FO4B-N5100","W5":"P19","W4":"O19","W3":"N19","W2":"M19","W1":"L19"},{"CellLine":"Pa04C","Drug":"Vinblastine","minConc":0.1,"D1":17.9,"D2":6.17,"D3":43.16,"D4":62.08,"D5":60.67,"IC50":7.65,"Slope":2.5,"max":62.08,"min":6.17,"Barcode":"H1FO4B-N5100","W5":"A21","W4":"B21","W3":"C21","W2":"D21","W1":"E21"},{"CellLine":"Pa04C","Drug":"Paclitaxel","minConc":0.1,"D1":10.13,"D2":8.2,"D3":37.94,"D4":39.45,"D5":49.59,"IC50":6.01,"Slope":1.63,"max":49.59,"min":8.2,"Barcode":"H1FO4B-N5100","W5":"F21","W4":"G21","W3":"H21","W2":"I21","W1":"J21"},{"CellLine":"Pa04C","Drug":"Oprozomib","minConc":0.25,"D1":6.3,"D2":11.4,"D3":26.23,"D4":99.46,"D5":104.05,"IC50":42.13,"Slope":2.5,"max":100,"min":6.3,"Barcode":"H1FO4B-N5100","W5":"A22","W4":"B22","W3":"C22","W2":"D22","W1":"E22"},{"CellLine":"Pa04C","Drug":"Bortezomib","minConc":0.1,"D1":6.26,"D2":10.89,"D3":95.06,"D4":99.91,"D5":98.18,"IC50":3.17,"Slope":2.5,"max":99.91,"min":6.09,"Barcode":"H1FO4B-N5100","W5":"F22","W4":"G22","W3":"H22","W2":"I22","W1":"J22"},{"CellLine":"Pa04C","Drug":"Filanesib","minConc":0.1,"D1":1.04,"D2":14.76,"D3":39.34,"D4":47.6,"D5":39.62,"IC50":2.2,"Slope":1.04,"max":47.6,"min":0,"Barcode":"H1FO4B-N5100","W5":"G23","W4":"H23","W3":"I23","W2":"J23","W1":"K23"},{"CellLine":"Pa04C","Drug":"CUDC_907","minConc":1,"D1":10.5,"D2":17.41,"D3":44.45,"D4":82.68,"D5":88.67,"IC50":122.07,"Slope":1.09,"max":88.67,"min":10.5,"Barcode":"H1FO4B-N6100","W5":"F17","W4":"G17","W3":"H17","W2":"I17","W1":"J17"},{"CellLine":"Pa04C","Drug":"YM155","minConc":1,"D1":2.03,"D2":4.6,"D3":23.43,"D4":64.59,"D5":101.15,"IC50":463.3,"Slope":0.92,"max":100,"min":2.03,"Barcode":"H1FO4B-N6100","W5":"F18","W4":"G18","W3":"H18","W2":"I18","W1":"J18"},{"CellLine":"Pa04C","Drug":"AT101","minConc":10,"D1":7.29,"D2":3.88,"D3":12.52,"D4":28.7,"D5":103.84,"IC50":15167.02,"Slope":2.5,"max":100,"min":3.88,"Barcode":"H1FO4B-N6100","W5":"A21","W4":"B21","W3":"C21","W2":"D21","W1":"E21"},{"CellLine":"Pa04C","Drug":"Selinexor","minConc":1,"D1":0.28,"D2":4.14,"D3":37.5,"D4":47.25,"D5":61.29,"IC50":87.79,"Slope":0.82,"max":61.29,"min":0,"Barcode":"H1FO4B-N6100","W5":"P21","W4":"O21","W3":"N21","W2":"M21","W1":"L21"},{"CellLine":"Pa04C","Drug":"BIIB021","minConc":1,"D1":12.18,"D2":12.69,"D3":20.89,"D4":52.28,"D5":49.35,"IC50":166.76,"Slope":2.5,"max":52.28,"min":12.18,"Barcode":"H1FO4B-N7100","W5":"P13","W4":"N13","W3":"M13","W2":"L13","W1":"K13"},{"CellLine":"Pa04C","Drug":"Tanespimycin","minConc":1,"D1":10.98,"D2":-10.11,"D3":32.67,"D4":48.49,"D5":38.37,"IC50":75.63,"Slope":2.5,"max":48.49,"min":0,"Barcode":"H1FO4B-N7100","W5":"A14","W4":"B14","W3":"C14","W2":"D14","W1":"E14"},{"CellLine":"Pa04C","Drug":"CUDC_305","minConc":1,"D1":5.99,"D2":10.45,"D3":12.4,"D4":48.53,"D5":51.8,"IC50":259.95,"Slope":1.87,"max":51.8,"min":5.99,"Barcode":"H1FO4B-N7100","W5":"A15","W4":"B15","W3":"C15","W2":"D15","W1":"F15"},{"CellLine":"Pa04C","Drug":"Luminespib","minConc":0.1,"D1":-0.14,"D2":5.75,"D3":12.42,"D4":48.44,"D5":42.63,"IC50":15.98,"Slope":2.24,"max":48.44,"min":0,"Barcode":"H1FO4B-N7100","W5":"G15","W4":"H15","W3":"I15","W2":"J15","W1":"K15"},{"CellLine":"Pa04C","Drug":"Ganetespib","minConc":0.1,"D1":7.99,"D2":10.48,"D3":9.09,"D4":53.06,"D5":40.7,"IC50":29.96,"Slope":2.5,"max":53.06,"min":7.99,"Barcode":"H1FO4B-N7100","W5":"A16","W4":"C16","W3":"D16","W2":"E16","W1":"F16"},{"CellLine":"Pa04C","Drug":"Stattic","minConc":5,"D1":8.75,"D2":-1.23,"D3":10.81,"D4":66.99,"D5":99.57,"IC50":2785.29,"Slope":1.28,"max":99.57,"min":0,"Barcode":"H1FO4B-N7100","W5":"F21","W4":"G21","W3":"H21","W2":"I21","W1":"J21"},{"CellLine":"Pa04C","Drug":"ONX_0914","minConc":1,"D1":9.75,"D2":6.58,"D3":10.43,"D4":100.55,"D5":99.17,"IC50":282.5,"Slope":2.5,"max":100,"min":6.58,"Barcode":"H1FO4B-N7100","W5":"G23","W4":"H23","W3":"I23","W2":"J23","W1":"K23"},{"CellLine":"Pa04C","Drug":"SN_38","minConc":1,"D1":6.77,"D2":22.75,"D3":53.48,"D4":90.22,"D5":88,"IC50":64.72,"Slope":0.93,"max":90.22,"min":6.77,"Barcode":"H1FO4B-N8100","W5":"G2","W4":"H2","W3":"I2","W2":"J2","W1":"K2"},{"CellLine":"Pa04C","Drug":"Cabazitaxel","minConc":0.1,"D1":11.59,"D2":20.87,"D3":37.76,"D4":32.01,"D5":23.02,"IC50":0.41,"Slope":0.36,"max":37.76,"min":0,"Barcode":"H1FO4B-N8100","W5":"A3","W4":"B3","W3":"C3","W2":"D3","W1":"E3"},{"CellLine":"Pa04C","Drug":"OTSSP167","minConc":0.1,"D1":19.47,"D2":14.83,"D3":15.52,"D4":38.99,"D5":82.37,"IC50":126.32,"Slope":2.5,"max":82.37,"min":14.83,"Barcode":"H1FO4B-N8100","W5":"A5","W4":"B5","W3":"C5","W2":"E5","W1":"F5"},{"CellLine":"Pa04C","Drug":"Prexasertib","minConc":1,"D1":9.72,"D2":14.32,"D3":29.98,"D4":35.06,"D5":19.22,"IC50":31.47,"Slope":1.16,"max":35.06,"min":9.14,"Barcode":"H1FO4B-N8100","W5":"P6","W4":"O6","W3":"N6","W2":"M6","W1":"L6"},{"CellLine":"Pa04C","Drug":"Romidepsin","minConc":0.1,"D1":-2.96,"D2":10.28,"D3":71.32,"D4":81.11,"D5":81.24,"IC50":3.13,"Slope":1.71,"max":81.24,"min":0,"Barcode":"H1FO4B-N8100","W5":"F18","W4":"G18","W3":"H18","W2":"I18","W1":"J18"},{"CellLine":"Pa04C","Drug":"Sabutoclax","minConc":2.5,"D1":-5.23,"D2":-3.76,"D3":3.8,"D4":32.97,"D5":71.83,"IC50":2727.87,"Slope":1.67,"max":71.83,"min":0,"Barcode":"H1FO4B-N8100","W5":"F19","W4":"G19","W3":"I19","W2":"J19","W1":"K19"},{"CellLine":"Pa04C","Drug":"BCI","minConc":5,"D1":16.78,"D2":6.35,"D3":17.22,"D4":95.65,"D5":105.77,"IC50":1257.49,"Slope":2.2,"max":100,"min":6.35,"Barcode":"H1FO4B-N8100","W5":"A22","W4":"B22","W3":"C22","W2":"D22","W1":"E22"},{"CellLine":"PANC-1","Drug":"PF_04691502","minConc":1,"D1":-6.47,"D2":-1.22,"D3":17.49,"D4":71.48,"D5":73.63,"IC50":176.26,"Slope":2.06,"max":73.63,"min":0,"Barcode":"H1FO4B-N1151","W5":"B2","W4":"C2","W3":"D2","W2":"E2","W1":"F2"},{"CellLine":"PANC-1","Drug":"SNS_032","minConc":1,"D1":7.97,"D2":15.59,"D3":47.74,"D4":94.96,"D5":92.51,"IC50":108.54,"Slope":1.38,"max":94.96,"min":7.97,"Barcode":"H1FO4B-N1151","W5":"G9","W4":"H9","W3":"I9","W2":"J9","W1":"K9"},{"CellLine":"PANC-1","Drug":"Alvocidib","minConc":1,"D1":5.13,"D2":11.38,"D3":36.82,"D4":94.96,"D5":97.36,"IC50":146.72,"Slope":1.62,"max":97.36,"min":5.13,"Barcode":"H1FO4B-N1151","W5":"P9","W4":"O9","W3":"N9","W2":"M9","W1":"L9"},{"CellLine":"PANC-1","Drug":"KX2_391","minConc":1,"D1":-3.8,"D2":-4.71,"D3":36.83,"D4":26.85,"D5":24.04,"IC50":37.39,"Slope":2.5,"max":36.83,"min":0,"Barcode":"H1FO4B-N1151","W5":"F17","W4":"G17","W3":"H17","W2":"I17","W1":"J17"},{"CellLine":"PANC-1","Drug":"AZD8055","minConc":1,"D1":12.57,"D2":6.6,"D3":28.28,"D4":46.45,"D5":55.19,"IC50":147.07,"Slope":0.92,"max":55.19,"min":6.6,"Barcode":"H1FO4B-N1151","W5":"G23","W4":"H23","W3":"I23","W2":"J23","W1":"K23"},{"CellLine":"PANC-1","Drug":"PF_03758309","minConc":1,"D1":5.97,"D2":18.66,"D3":53.45,"D4":78.77,"D5":99.19,"IC50":89.54,"Slope":0.65,"max":99.19,"min":0.59,"Barcode":"H1FO4B-N2151","W5":"P5","W4":"O5","W3":"N5","W2":"M5","W1":"L5"},{"CellLine":"PANC-1","Drug":"AZD_8330","minConc":1,"D1":2.55,"D2":11.01,"D3":11.43,"D4":21.82,"D5":29.55,"IC50":111.57,"Slope":0.68,"max":29.55,"min":2.55,"Barcode":"H1FO4B-N2151","W5":"F6","W4":"H6","W3":"I6","W2":"J6","W1":"K6"},{"CellLine":"PANC-1","Drug":"Rigosertib","minConc":1,"D1":-10.87,"D2":-11.01,"D3":34.06,"D4":31.46,"D5":29.56,"IC50":41.86,"Slope":2.5,"max":34.06,"min":0,"Barcode":"H1FO4B-N2151","W5":"F8","W4":"G8","W3":"H8","W2":"J8","W1":"K8"},{"CellLine":"PANC-1","Drug":"Omipalisib","minConc":0.1,"D1":9.77,"D2":17.01,"D3":21.84,"D4":57.64,"D5":65.07,"IC50":25.04,"Slope":1.26,"max":65.07,"min":9.77,"Barcode":"H1FO4B-N2151","W5":"P17","W4":"O17","W3":"N17","W2":"M17","W1":"K17"},{"CellLine":"PANC-1","Drug":"UCN_01","minConc":1,"D1":14.63,"D2":7.81,"D3":30.67,"D4":59.77,"D5":97.96,"IC50":517.57,"Slope":0.82,"max":97.96,"min":7.81,"Barcode":"H1FO4B-N2151","W5":"P21","W4":"O21","W3":"N21","W2":"M21","W1":"L21"},{"CellLine":"PANC-1","Drug":"Dinaciclib","minConc":0.1,"D1":3.6,"D2":-11.01,"D3":60.8,"D4":98.77,"D5":97.95,"IC50":8.32,"Slope":2.5,"max":98.77,"min":0,"Barcode":"H1FO4B-N3151","W5":"A3","W4":"B3","W3":"C3","W2":"D3","W1":"E3"},{"CellLine":"PANC-1","Drug":"NVP_BGT226","minConc":0.1,"D1":3.5,"D2":6.07,"D3":7.22,"D4":33.41,"D5":47.05,"IC50":55.97,"Slope":1.37,"max":47.05,"min":3.5,"Barcode":"H1FO4B-N3151","W5":"G12","W4":"H12","W3":"I12","W2":"J12","W1":"K12"},{"CellLine":"PANC-1","Drug":"Gedatolisib","minConc":0.1,"D1":-4.9,"D2":14.22,"D3":14.04,"D4":40.71,"D5":64.5,"IC50":40.81,"Slope":0.74,"max":64.5,"min":0,"Barcode":"H1FO4B-N3151","W5":"P12","W4":"O12","W3":"N12","W2":"M12","W1":"L12"},{"CellLine":"PANC-1","Drug":"PKI_402","minConc":0.25,"D1":-4.74,"D2":-4.56,"D3":3.99,"D4":36.02,"D5":64.42,"IC50":209.79,"Slope":1.48,"max":64.42,"min":0,"Barcode":"H1FO4B-N3151","W5":"A16","W4":"C16","W3":"D16","W2":"E16","W1":"F16"},{"CellLine":"PANC-1","Drug":"AT7519","minConc":1,"D1":10.04,"D2":-8,"D3":9.87,"D4":93.15,"D5":94.1,"IC50":232.28,"Slope":2.5,"max":94.1,"min":0,"Barcode":"H1FO4B-N3151","W5":"F22","W4":"G22","W3":"H22","W2":"I22","W1":"J22"},{"CellLine":"PANC-1","Drug":"GSK_461364","minConc":1,"D1":-5.97,"D2":23.71,"D3":39.99,"D4":42.64,"D5":73.65,"IC50":123.79,"Slope":0.49,"max":73.65,"min":0,"Barcode":"H1FO4B-N4151","W5":"A6","W4":"B6","W3":"C6","W2":"D6","W1":"E6"},{"CellLine":"PANC-1","Drug":"Daporinad","minConc":0.1,"D1":-15.79,"D2":-7.61,"D3":38.29,"D4":74.24,"D5":72.2,"IC50":9.77,"Slope":2.5,"max":74.24,"min":0,"Barcode":"H1FO4B-N4151","W5":"A14","W4":"B14","W3":"C14","W2":"D14","W1":"E14"},{"CellLine":"PANC-1","Drug":"Mepacrine","minConc":5,"D1":-0.54,"D2":3.73,"D3":22.44,"D4":89.2,"D5":102.86,"IC50":1165.47,"Slope":1.45,"max":100,"min":0,"Barcode":"H1FO4B-N4151","W5":"P20","W4":"O20","W3":"N20","W2":"M20","W1":"L20"},{"CellLine":"PANC-1","Drug":"Camptothecin","minConc":0.5,"D1":-15.03,"D2":-1.83,"D3":3.7,"D4":54.32,"D5":74.01,"IC50":278.61,"Slope":1.74,"max":74.01,"min":0,"Barcode":"H1FO4B-N5151","W5":"O3","W4":"N3","W3":"M3","W2":"L3","W1":"K3"},{"CellLine":"PANC-1","Drug":"Omacetaxine","minConc":1,"D1":-11.06,"D2":0.98,"D3":66.61,"D4":93.66,"D5":96.56,"IC50":69.48,"Slope":2.19,"max":96.56,"min":0,"Barcode":"H1FO4B-N5151","W5":"A5","W4":"B5","W3":"C5","W2":"E5","W1":"F5"},{"CellLine":"PANC-1","Drug":"Vinorelbine","minConc":1,"D1":-0.76,"D2":41.72,"D3":40.38,"D4":33.71,"D5":56.37,"IC50":14.94,"Slope":0.42,"max":56.37,"min":0,"Barcode":"H1FO4B-N5151","W5":"P14","W4":"O14","W3":"N14","W2":"M14","W1":"L14"},{"CellLine":"PANC-1","Drug":"Docetaxel","minConc":0.1,"D1":0.52,"D2":17.27,"D3":38.12,"D4":38.45,"D5":39.42,"IC50":1.17,"Slope":1.61,"max":39.42,"min":0,"Barcode":"H1FO4B-N5151","W5":"P17","W4":"O17","W3":"N17","W2":"M17","W1":"K17"},{"CellLine":"PANC-1","Drug":"Carfilzomib","minConc":0.1,"D1":-7.21,"D2":-6.33,"D3":87.88,"D4":98.45,"D5":99.99,"IC50":4.76,"Slope":2.5,"max":99.99,"min":0,"Barcode":"H1FO4B-N5151","W5":"A18","W4":"B18","W3":"C18","W2":"D18","W1":"E18"},{"CellLine":"PANC-1","Drug":"Patupilone","minConc":0.1,"D1":-9.99,"D2":-3.55,"D3":35.62,"D4":45.76,"D5":36.21,"IC50":6.13,"Slope":2.5,"max":45.76,"min":0,"Barcode":"H1FO4B-N5151","W5":"F18","W4":"G18","W3":"H18","W2":"I18","W1":"J18"},{"CellLine":"PANC-1","Drug":"Plicamycin","minConc":1,"D1":-0.71,"D2":-3.3,"D3":3.16,"D4":75.07,"D5":100.19,"IC50":573.58,"Slope":1.99,"max":100,"min":0,"Barcode":"H1FO4B-N5151","W5":"P18","W4":"N18","W3":"M18","W2":"L18","W1":"K18"},{"CellLine":"PANC-1","Drug":"Vincristine","minConc":0.1,"D1":-2.72,"D2":7.87,"D3":33.68,"D4":27.68,"D5":29.47,"IC50":1.61,"Slope":2.5,"max":33.68,"min":0,"Barcode":"H1FO4B-N5151","W5":"A19","W4":"B19","W3":"C19","W2":"D19","W1":"E19"},{"CellLine":"PANC-1","Drug":"Dactinomycin","minConc":0.1,"D1":3.02,"D2":23.15,"D3":48.07,"D4":98.46,"D5":99.32,"IC50":8.52,"Slope":0.9,"max":99.32,"min":3.02,"Barcode":"H1FO4B-N5151","W5":"P19","W4":"O19","W3":"N19","W2":"M19","W1":"L19"},{"CellLine":"PANC-1","Drug":"Vinblastine","minConc":0.1,"D1":-1.43,"D2":7.19,"D3":51.15,"D4":33.18,"D5":30.68,"IC50":2.05,"Slope":2.5,"max":51.15,"min":0,"Barcode":"H1FO4B-N5151","W5":"A21","W4":"B21","W3":"C21","W2":"D21","W1":"E21"},{"CellLine":"PANC-1","Drug":"Paclitaxel","minConc":0.1,"D1":-5.5,"D2":7.83,"D3":35.35,"D4":48.72,"D5":44.26,"IC50":4.38,"Slope":1.23,"max":48.72,"min":0,"Barcode":"H1FO4B-N5151","W5":"F21","W4":"G21","W3":"H21","W2":"I21","W1":"J21"},{"CellLine":"PANC-1","Drug":"Oprozomib","minConc":0.25,"D1":-6.5,"D2":1.67,"D3":25.04,"D4":94.69,"D5":101.95,"IC50":47.13,"Slope":1.73,"max":100,"min":0,"Barcode":"H1FO4B-N5151","W5":"A22","W4":"B22","W3":"C22","W2":"D22","W1":"E22"},{"CellLine":"PANC-1","Drug":"Bortezomib","minConc":0.1,"D1":3.36,"D2":-11.19,"D3":98.39,"D4":100.21,"D5":98.76,"IC50":3.79,"Slope":2.5,"max":100,"min":0,"Barcode":"H1FO4B-N5151","W5":"F22","W4":"G22","W3":"H22","W2":"I22","W1":"J22"},{"CellLine":"PANC-1","Drug":"Filanesib","minConc":0.1,"D1":13.24,"D2":2.1,"D3":52.39,"D4":48.92,"D5":47.61,"IC50":3.16,"Slope":2.5,"max":52.39,"min":2.1,"Barcode":"H1FO4B-N5151","W5":"G23","W4":"H23","W3":"I23","W2":"J23","W1":"K23"},{"CellLine":"PANC-1","Drug":"CUDC_907","minConc":1,"D1":-3.4,"D2":-0.06,"D3":58.95,"D4":86.51,"D5":96.41,"IC50":77.53,"Slope":1.59,"max":96.41,"min":0,"Barcode":"H1FO4B-N6151","W5":"F17","W4":"G17","W3":"H17","W2":"I17","W1":"J17"},{"CellLine":"PANC-1","Drug":"YM155","minConc":1,"D1":31.04,"D2":94,"D3":100.27,"D4":99.55,"D5":99.57,"IC50":3.85,"Slope":2.5,"max":100,"min":28.66,"Barcode":"H1FO4B-N6151","W5":"F18","W4":"G18","W3":"H18","W2":"I18","W1":"J18"},{"CellLine":"PANC-1","Drug":"AT101","minConc":10,"D1":-0.37,"D2":12.33,"D3":28.15,"D4":84.52,"D5":101.62,"IC50":2170.35,"Slope":1.05,"max":100,"min":0,"Barcode":"H1FO4B-N6151","W5":"A21","W4":"B21","W3":"C21","W2":"D21","W1":"E21"},{"CellLine":"PANC-1","Drug":"Selinexor","minConc":1,"D1":16.42,"D2":9.59,"D3":39.4,"D4":62.27,"D5":77.45,"IC50":167.04,"Slope":0.84,"max":77.45,"min":9.59,"Barcode":"H1FO4B-N6151","W5":"P21","W4":"O21","W3":"N21","W2":"M21","W1":"L21"},{"CellLine":"PANC-1","Drug":"BIIB021","minConc":1,"D1":11.4,"D2":3.78,"D3":20.12,"D4":60.39,"D5":67.74,"IC50":220.7,"Slope":1.36,"max":67.74,"min":3.78,"Barcode":"H1FO4B-N7151","W5":"P13","W4":"N13","W3":"M13","W2":"L13","W1":"K13"},{"CellLine":"PANC-1","Drug":"Tanespimycin","minConc":1,"D1":-0.26,"D2":-7.86,"D3":9.79,"D4":56.38,"D5":55.2,"IC50":186.17,"Slope":2.5,"max":56.38,"min":0,"Barcode":"H1FO4B-N7151","W5":"A14","W4":"B14","W3":"C14","W2":"D14","W1":"E14"},{"CellLine":"PANC-1","Drug":"CUDC_305","minConc":1,"D1":-6.09,"D2":14.91,"D3":6.97,"D4":55.4,"D5":72.68,"IC50":437.74,"Slope":1.39,"max":72.68,"min":0,"Barcode":"H1FO4B-N7151","W5":"A15","W4":"B15","W3":"C15","W2":"D15","W1":"F15"},{"CellLine":"PANC-1","Drug":"Luminespib","minConc":0.1,"D1":-8.79,"D2":-12.06,"D3":24.86,"D4":62.97,"D5":54.58,"IC50":11.9,"Slope":2.5,"max":62.97,"min":0,"Barcode":"H1FO4B-N7151","W5":"G15","W4":"H15","W3":"I15","W2":"J15","W1":"K15"},{"CellLine":"PANC-1","Drug":"Ganetespib","minConc":0.1,"D1":-15.35,"D2":8.47,"D3":11.9,"D4":62.46,"D5":59.92,"IC50":17.76,"Slope":2.5,"max":62.46,"min":0,"Barcode":"H1FO4B-N7151","W5":"A16","W4":"C16","W3":"D16","W2":"E16","W1":"F16"},{"CellLine":"PANC-1","Drug":"Stattic","minConc":5,"D1":7.24,"D2":-11.85,"D3":-16.56,"D4":83.2,"D5":103.76,"IC50":2756.44,"Slope":2.5,"max":100,"min":0,"Barcode":"H1FO4B-N7151","W5":"F21","W4":"G21","W3":"H21","W2":"I21","W1":"J21"},{"CellLine":"PANC-1","Drug":"ONX_0914","minConc":1,"D1":6.17,"D2":-9.89,"D3":5.01,"D4":101.27,"D5":102.19,"IC50":272.39,"Slope":2.5,"max":100,"min":0,"Barcode":"H1FO4B-N7151","W5":"G23","W4":"H23","W3":"I23","W2":"J23","W1":"K23"},{"CellLine":"PANC-1","Drug":"SN_38","minConc":1,"D1":1.57,"D2":13.42,"D3":29.49,"D4":67.54,"D5":89.21,"IC50":231.22,"Slope":0.78,"max":89.21,"min":1.57,"Barcode":"H1FO4B-N8151","W5":"G2","W4":"H2","W3":"I2","W2":"J2","W1":"K2"},{"CellLine":"PANC-1","Drug":"Cabazitaxel","minConc":0.1,"D1":6.25,"D2":17.37,"D3":43.39,"D4":30.24,"D5":39.46,"IC50":1.4,"Slope":2.5,"max":43.39,"min":6.2,"Barcode":"H1FO4B-N8151","W5":"A3","W4":"B3","W3":"C3","W2":"D3","W1":"E3"},{"CellLine":"PANC-1","Drug":"OTSSP167","minConc":0.1,"D1":-0.74,"D2":10.57,"D3":-2.27,"D4":67.79,"D5":98.7,"IC50":73.18,"Slope":2.5,"max":98.7,"min":0,"Barcode":"H1FO4B-N8151","W5":"A5","W4":"B5","W3":"C5","W2":"E5","W1":"F5"},{"CellLine":"PANC-1","Drug":"Prexasertib","minConc":1,"D1":1.41,"D2":8.25,"D3":5.6,"D4":21.01,"D5":29.96,"IC50":434.78,"Slope":0.92,"max":29.96,"min":1.41,"Barcode":"H1FO4B-N8151","W5":"P6","W4":"O6","W3":"N6","W2":"M6","W1":"L6"},{"CellLine":"PANC-1","Drug":"Romidepsin","minConc":0.1,"D1":3.56,"D2":42.77,"D3":78.61,"D4":83.75,"D5":86.7,"IC50":1.07,"Slope":1.12,"max":86.7,"min":0,"Barcode":"H1FO4B-N8151","W5":"F18","W4":"G18","W3":"H18","W2":"I18","W1":"J18"},{"CellLine":"PANC-1","Drug":"Sabutoclax","minConc":2.5,"D1":3.16,"D2":1.79,"D3":49.01,"D4":83.43,"D5":99,"IC50":295.82,"Slope":0.98,"max":99,"min":0,"Barcode":"H1FO4B-N8151","W5":"F19","W4":"G19","W3":"I19","W2":"J19","W1":"K19"},{"CellLine":"PANC-1","Drug":"BCI","minConc":5,"D1":3.04,"D2":3.5,"D3":12.46,"D4":96.67,"D5":102.76,"IC50":1256.15,"Slope":2.42,"max":100,"min":3.04,"Barcode":"H1FO4B-N8151","W5":"A22","W4":"B22","W3":"C22","W2":"D22","W1":"E22"},{"CellLine":"MDA-PATC43","Drug":"PF_04691502","minConc":1,"D1":2.46,"D2":6.31,"D3":27.96,"D4":78.86,"D5":90.42,"IC50":208.66,"Slope":1.19,"max":90.42,"min":2.46,"Barcode":"H1FO4B-N1143","W5":"B2","W4":"C2","W3":"D2","W2":"E2","W1":"F2"},{"CellLine":"MDA-PATC43","Drug":"SNS_032","minConc":1,"D1":1.28,"D2":4.76,"D3":79.46,"D4":90.75,"D5":90.12,"IC50":42,"Slope":2.23,"max":90.75,"min":1.26,"Barcode":"H1FO4B-N1143","W5":"G9","W4":"H9","W3":"I9","W2":"J9","W1":"K9"},{"CellLine":"MDA-PATC43","Drug":"Alvocidib","minConc":1,"D1":0.27,"D2":1.08,"D3":34.35,"D4":92.53,"D5":93.82,"IC50":130.99,"Slope":2.06,"max":93.82,"min":0.27,"Barcode":"H1FO4B-N1143","W5":"P9","W4":"O9","W3":"N9","W2":"M9","W1":"L9"},{"CellLine":"MDA-PATC43","Drug":"KX2_391","minConc":1,"D1":0.67,"D2":4.27,"D3":70.85,"D4":73.79,"D5":71.6,"IC50":30.06,"Slope":2.5,"max":73.79,"min":0.28,"Barcode":"H1FO4B-N1143","W5":"F17","W4":"G17","W3":"H17","W2":"I17","W1":"J17"},{"CellLine":"MDA-PATC43","Drug":"AZD8055","minConc":1,"D1":2.4,"D2":6.37,"D3":51.05,"D4":79.54,"D5":81.91,"IC50":72.39,"Slope":1.42,"max":81.91,"min":2.03,"Barcode":"H1FO4B-N1143","W5":"G23","W4":"H23","W3":"I23","W2":"J23","W1":"K23"},{"CellLine":"MDA-PATC43","Drug":"PF_03758309","minConc":1,"D1":12.14,"D2":32.77,"D3":41.26,"D4":57.58,"D5":98.57,"IC50":376.57,"Slope":0.62,"max":98.57,"min":12.14,"Barcode":"H1FO4B-N2143","W5":"P5","W4":"O5","W3":"N5","W2":"M5","W1":"L5"},{"CellLine":"MDA-PATC43","Drug":"AZD_8330","minConc":1,"D1":21.82,"D2":46.43,"D3":67.9,"D4":74.78,"D5":73.28,"IC50":8.3,"Slope":0.85,"max":74.78,"min":13.1,"Barcode":"H1FO4B-N2143","W5":"F6","W4":"H6","W3":"I6","W2":"J6","W1":"K6"},{"CellLine":"MDA-PATC43","Drug":"Rigosertib","minConc":1,"D1":3.02,"D2":-1.71,"D3":73.61,"D4":72.51,"D5":75.25,"IC50":34.61,"Slope":2.5,"max":75.25,"min":0,"Barcode":"H1FO4B-N2143","W5":"F8","W4":"G8","W3":"H8","W2":"J8","W1":"K8"},{"CellLine":"MDA-PATC43","Drug":"Omipalisib","minConc":0.1,"D1":6.1,"D2":8.97,"D3":36.54,"D4":73.79,"D5":79.92,"IC50":13.35,"Slope":1.2,"max":79.92,"min":5.86,"Barcode":"H1FO4B-N2143","W5":"P17","W4":"O17","W3":"N17","W2":"M17","W1":"K17"},{"CellLine":"MDA-PATC43","Drug":"UCN_01","minConc":1,"D1":5.71,"D2":8.52,"D3":25.16,"D4":49.13,"D5":75.02,"IC50":397.01,"Slope":0.79,"max":75.02,"min":5.48,"Barcode":"H1FO4B-N2143","W5":"P21","W4":"O21","W3":"N21","W2":"M21","W1":"L21"},{"CellLine":"MDA-PATC43","Drug":"Dinaciclib","minConc":0.1,"D1":-4.9,"D2":-2.35,"D3":50.88,"D4":90.04,"D5":89.9,"IC50":9.01,"Slope":2.5,"max":90.04,"min":0,"Barcode":"H1FO4B-N3143","W5":"A3","W4":"B3","W3":"C3","W2":"D3","W1":"E3"},{"CellLine":"MDA-PATC43","Drug":"NVP_BGT226","minConc":0.1,"D1":8.39,"D2":-0.24,"D3":43.4,"D4":76.6,"D5":98.22,"IC50":16.47,"Slope":0.86,"max":98.22,"min":0,"Barcode":"H1FO4B-N3143","W5":"G12","W4":"H12","W3":"I12","W2":"J12","W1":"K12"},{"CellLine":"MDA-PATC43","Drug":"Gedatolisib","minConc":0.1,"D1":0.15,"D2":-1.74,"D3":13.27,"D4":75.38,"D5":85.09,"IC50":28.32,"Slope":1.63,"max":85.09,"min":0,"Barcode":"H1FO4B-N3143","W5":"P12","W4":"O12","W3":"N12","W2":"M12","W1":"L12"},{"CellLine":"MDA-PATC43","Drug":"PKI_402","minConc":0.25,"D1":4.64,"D2":7.81,"D3":21.1,"D4":75.62,"D5":82.11,"IC50":56.55,"Slope":1.58,"max":82.11,"min":4.64,"Barcode":"H1FO4B-N3143","W5":"A16","W4":"C16","W3":"D16","W2":"E16","W1":"F16"},{"CellLine":"MDA-PATC43","Drug":"AT7519","minConc":1,"D1":-0.89,"D2":2.88,"D3":12.33,"D4":82.48,"D5":90.09,"IC50":272.76,"Slope":1.83,"max":90.09,"min":0,"Barcode":"H1FO4B-N3143","W5":"F22","W4":"G22","W3":"H22","W2":"I22","W1":"J22"},{"CellLine":"MDA-PATC43","Drug":"GSK_461364","minConc":1,"D1":0.11,"D2":36.83,"D3":65.09,"D4":62.67,"D5":90.47,"IC50":34.23,"Slope":0.53,"max":90.47,"min":0,"Barcode":"H1FO4B-N4143","W5":"A6","W4":"B6","W3":"C6","W2":"D6","W1":"E6"},{"CellLine":"MDA-PATC43","Drug":"Daporinad","minConc":0.1,"D1":-1.05,"D2":-1.23,"D3":45.34,"D4":65.71,"D5":66.35,"IC50":7.36,"Slope":2.5,"max":66.35,"min":0,"Barcode":"H1FO4B-N4143","W5":"A14","W4":"B14","W3":"C14","W2":"D14","W1":"E14"},{"CellLine":"MDA-PATC43","Drug":"Mepacrine","minConc":5,"D1":3.34,"D2":9.51,"D3":15.1,"D4":65.84,"D5":102.4,"IC50":2850.05,"Slope":1.19,"max":100,"min":3.34,"Barcode":"H1FO4B-N4143","W5":"P20","W4":"O20","W3":"N20","W2":"M20","W1":"L20"},{"CellLine":"MDA-PATC43","Drug":"Camptothecin","minConc":0.5,"D1":6.57,"D2":12.68,"D3":48.92,"D4":81.17,"D5":97.09,"IC50":58.42,"Slope":0.82,"max":97.09,"min":3.58,"Barcode":"H1FO4B-N5143","W5":"O3","W4":"N3","W3":"M3","W2":"L3","W1":"K3"},{"CellLine":"MDA-PATC43","Drug":"Omacetaxine","minConc":1,"D1":7.25,"D2":6.66,"D3":98.87,"D4":96.72,"D5":99.17,"IC50":30.05,"Slope":2.5,"max":99.17,"min":4.29,"Barcode":"H1FO4B-N5143","W5":"A5","W4":"B5","W3":"C5","W2":"E5","W1":"F5"},{"CellLine":"MDA-PATC43","Drug":"Vinorelbine","minConc":1,"D1":12.55,"D2":67.11,"D3":77.1,"D4":75.66,"D5":83.6,"IC50":3.61,"Slope":1.31,"max":83.6,"min":0,"Barcode":"H1FO4B-N5143","W5":"P14","W4":"O14","W3":"N14","W2":"M14","W1":"L14"},{"CellLine":"MDA-PATC43","Drug":"Docetaxel","minConc":0.1,"D1":8.84,"D2":37.1,"D3":72.55,"D4":77.25,"D5":74.49,"IC50":1.22,"Slope":1.26,"max":77.25,"min":5.94,"Barcode":"H1FO4B-N5143","W5":"P17","W4":"O17","W3":"N17","W2":"M17","W1":"K17"},{"CellLine":"MDA-PATC43","Drug":"Carfilzomib","minConc":0.1,"D1":-5.23,"D2":4.95,"D3":84.79,"D4":96.76,"D5":101.5,"IC50":4.29,"Slope":2.03,"max":100,"min":0,"Barcode":"H1FO4B-N5143","W5":"A18","W4":"B18","W3":"C18","W2":"D18","W1":"E18"},{"CellLine":"MDA-PATC43","Drug":"Patupilone","minConc":0.1,"D1":5.02,"D2":6.3,"D3":68.88,"D4":70.7,"D5":63.36,"IC50":3.07,"Slope":2.5,"max":70.7,"min":3.83,"Barcode":"H1FO4B-N5143","W5":"F18","W4":"G18","W3":"H18","W2":"I18","W1":"J18"},{"CellLine":"MDA-PATC43","Drug":"Plicamycin","minConc":1,"D1":-3.55,"D2":9.64,"D3":46.78,"D4":98.68,"D5":93.36,"IC50":102.62,"Slope":1.3,"max":98.68,"min":0,"Barcode":"H1FO4B-N5143","W5":"P18","W4":"N18","W3":"M18","W2":"L18","W1":"K18"},{"CellLine":"MDA-PATC43","Drug":"Vincristine","minConc":0.1,"D1":3.86,"D2":-4.99,"D3":74.6,"D4":77.19,"D5":72.58,"IC50":3.76,"Slope":2.5,"max":77.19,"min":0,"Barcode":"H1FO4B-N5143","W5":"A19","W4":"B19","W3":"C19","W2":"D19","W1":"E19"},{"CellLine":"MDA-PATC43","Drug":"Dactinomycin","minConc":0.1,"D1":13.79,"D2":60.46,"D3":75.38,"D4":94.22,"D5":97.61,"IC50":0.78,"Slope":0.67,"max":97.61,"min":0,"Barcode":"H1FO4B-N5143","W5":"P19","W4":"O19","W3":"N19","W2":"M19","W1":"L19"},{"CellLine":"MDA-PATC43","Drug":"Vinblastine","minConc":0.1,"D1":1.54,"D2":-3.15,"D3":70.85,"D4":78.21,"D5":76.17,"IC50":4.34,"Slope":2.5,"max":78.21,"min":0,"Barcode":"H1FO4B-N5143","W5":"A21","W4":"B21","W3":"C21","W2":"D21","W1":"E21"},{"CellLine":"MDA-PATC43","Drug":"Paclitaxel","minConc":0.1,"D1":5.39,"D2":12.29,"D3":67.34,"D4":76.16,"D5":74.73,"IC50":3.4,"Slope":1.81,"max":76.16,"min":5.27,"Barcode":"H1FO4B-N5143","W5":"F21","W4":"G21","W3":"H21","W2":"I21","W1":"J21"},{"CellLine":"MDA-PATC43","Drug":"Oprozomib","minConc":0.25,"D1":0.73,"D2":-7.24,"D3":33.1,"D4":88.8,"D5":96.95,"IC50":40.15,"Slope":1.5,"max":96.95,"min":0,"Barcode":"H1FO4B-N5143","W5":"A22","W4":"B22","W3":"C22","W2":"D22","W1":"E22"},{"CellLine":"MDA-PATC43","Drug":"Bortezomib","minConc":0.1,"D1":-1.91,"D2":3.38,"D3":86.73,"D4":91.97,"D5":97.32,"IC50":4.09,"Slope":2.35,"max":97.32,"min":0,"Barcode":"H1FO4B-N5143","W5":"F22","W4":"G22","W3":"H22","W2":"I22","W1":"J22"},{"CellLine":"MDA-PATC43","Drug":"Filanesib","minConc":0.1,"D1":0.33,"D2":2.08,"D3":73.39,"D4":72.8,"D5":75.22,"IC50":3.13,"Slope":2.5,"max":75.22,"min":0,"Barcode":"H1FO4B-N5143","W5":"G23","W4":"H23","W3":"I23","W2":"J23","W1":"K23"},{"CellLine":"MDA-PATC43","Drug":"CUDC_907","minConc":1,"D1":1.53,"D2":27.26,"D3":99.7,"D4":100.78,"D5":97.27,"IC50":15.12,"Slope":2.5,"max":100,"min":1.4,"Barcode":"H1FO4B-N6143","W5":"F17","W4":"G17","W3":"H17","W2":"I17","W1":"J17"},{"CellLine":"MDA-PATC43","Drug":"YM155","minConc":1,"D1":3.51,"D2":26.05,"D3":95.83,"D4":102.14,"D5":104.21,"IC50":18.74,"Slope":1.86,"max":100,"min":3.1,"Barcode":"H1FO4B-N6143","W5":"F18","W4":"G18","W3":"H18","W2":"I18","W1":"J18"},{"CellLine":"MDA-PATC43","Drug":"AT101","minConc":10,"D1":1.73,"D2":-1.96,"D3":4,"D4":93.37,"D5":104.97,"IC50":3495.57,"Slope":2.5,"max":100,"min":0,"Barcode":"H1FO4B-N6143","W5":"A21","W4":"B21","W3":"C21","W2":"D21","W1":"E21"},{"CellLine":"MDA-PATC43","Drug":"Selinexor","minConc":1,"D1":4.56,"D2":6.06,"D3":66.78,"D4":85.87,"D5":87.73,"IC50":59.64,"Slope":2.11,"max":87.73,"min":4.36,"Barcode":"H1FO4B-N6143","W5":"P21","W4":"O21","W3":"N21","W2":"M21","W1":"L21"},{"CellLine":"MDA-PATC43","Drug":"BIIB021","minConc":1,"D1":7.61,"D2":14.98,"D3":48.26,"D4":80.36,"D5":90.27,"IC50":101.31,"Slope":0.91,"max":90.27,"min":6.2,"Barcode":"H1FO4B-N7143","W5":"P13","W4":"N13","W3":"M13","W2":"L13","W1":"K13"},{"CellLine":"MDA-PATC43","Drug":"Tanespimycin","minConc":1,"D1":2.53,"D2":-5.44,"D3":61.86,"D4":80.87,"D5":78.26,"IC50":62.96,"Slope":2.5,"max":80.87,"min":0,"Barcode":"H1FO4B-N7143","W5":"A14","W4":"B14","W3":"C14","W2":"D14","W1":"E14"},{"CellLine":"MDA-PATC43","Drug":"CUDC_305","minConc":1,"D1":-1.51,"D2":-9,"D3":12.02,"D4":75.99,"D5":82.26,"IC50":259.23,"Slope":1.89,"max":82.26,"min":0,"Barcode":"H1FO4B-N7143","W5":"A15","W4":"B15","W3":"C15","W2":"D15","W1":"F15"},{"CellLine":"MDA-PATC43","Drug":"Luminespib","minConc":0.1,"D1":5.5,"D2":8.89,"D3":55.02,"D4":78.28,"D5":77.69,"IC50":6.33,"Slope":1.66,"max":78.28,"min":5.5,"Barcode":"H1FO4B-N7143","W5":"G15","W4":"H15","W3":"I15","W2":"J15","W1":"K15"},{"CellLine":"MDA-PATC43","Drug":"Ganetespib","minConc":0.1,"D1":-1.86,"D2":-5.31,"D3":26.72,"D4":76.56,"D5":80.6,"IC50":14.96,"Slope":1.8,"max":80.6,"min":0,"Barcode":"H1FO4B-N7143","W5":"A16","W4":"C16","W3":"D16","W2":"E16","W1":"F16"},{"CellLine":"MDA-PATC43","Drug":"Stattic","minConc":5,"D1":12.58,"D2":0.27,"D3":-3.06,"D4":101.92,"D5":104.92,"IC50":1610.9,"Slope":2.5,"max":100,"min":0,"Barcode":"H1FO4B-N7143","W5":"F21","W4":"G21","W3":"H21","W2":"I21","W1":"J21"},{"CellLine":"MDA-PATC43","Drug":"ONX_0914","minConc":1,"D1":6.11,"D2":0.42,"D3":7.15,"D4":95.36,"D5":99.96,"IC50":291.24,"Slope":2.45,"max":99.96,"min":0.42,"Barcode":"H1FO4B-N7143","W5":"G23","W4":"H23","W3":"I23","W2":"J23","W1":"K23"},{"CellLine":"MDA-PATC43","Drug":"SN_38","minConc":1,"D1":5.36,"D2":35.06,"D3":60.75,"D4":94.79,"D5":92.18,"IC50":28.86,"Slope":0.7,"max":94.79,"min":0,"Barcode":"H1FO4B-N8143","W5":"G2","W4":"H2","W3":"I2","W2":"J2","W1":"K2"},{"CellLine":"MDA-PATC43","Drug":"Cabazitaxel","minConc":0.1,"D1":1.74,"D2":24.69,"D3":68.89,"D4":74.67,"D5":64.46,"IC50":1.68,"Slope":1.38,"max":74.67,"min":0.25,"Barcode":"H1FO4B-N8143","W5":"A3","W4":"B3","W3":"C3","W2":"D3","W1":"E3"},{"CellLine":"MDA-PATC43","Drug":"OTSSP167","minConc":0.1,"D1":5.49,"D2":4.91,"D3":18.72,"D4":91.1,"D5":96.8,"IC50":24.51,"Slope":1.93,"max":96.8,"min":4.91,"Barcode":"H1FO4B-N8143","W5":"A5","W4":"B5","W3":"C5","W2":"E5","W1":"F5"},{"CellLine":"MDA-PATC43","Drug":"Prexasertib","minConc":1,"D1":-5.89,"D2":-10.24,"D3":52.67,"D4":68.72,"D5":96.03,"IC50":145.3,"Slope":0.9,"max":96.03,"min":0,"Barcode":"H1FO4B-N8143","W5":"P6","W4":"O6","W3":"N6","W2":"M6","W1":"L6"},{"CellLine":"MDA-PATC43","Drug":"Romidepsin","minConc":0.1,"D1":1.46,"D2":24.49,"D3":96.91,"D4":99.39,"D5":101.06,"IC50":1.8,"Slope":2,"max":100,"min":1.16,"Barcode":"H1FO4B-N8143","W5":"F18","W4":"G18","W3":"H18","W2":"I18","W1":"J18"},{"CellLine":"MDA-PATC43","Drug":"Sabutoclax","minConc":2.5,"D1":-2.44,"D2":1.18,"D3":6.72,"D4":83.03,"D5":101.07,"IC50":1050.68,"Slope":1.84,"max":100,"min":0,"Barcode":"H1FO4B-N8143","W5":"F19","W4":"G19","W3":"I19","W2":"J19","W1":"K19"},{"CellLine":"MDA-PATC43","Drug":"BCI","minConc":5,"D1":-5.57,"D2":-5.99,"D3":7.02,"D4":101.53,"D5":101.44,"IC50":1273.9,"Slope":2.5,"max":100,"min":0,"Barcode":"H1FO4B-N8143","W5":"A22","W4":"B22","W3":"C22","W2":"D22","W1":"E22"},{"CellLine":"MDA-PATC50","Drug":"PF_04691502","minConc":1,"D1":-4.43,"D2":-7.03,"D3":16.57,"D4":67.8,"D5":99.8,"IC50":488.42,"Slope":1.14,"max":99.8,"min":0,"Barcode":"H1FO4B-N1193","W5":"B2","W4":"C2","W3":"D2","W2":"E2","W1":"F2"},{"CellLine":"MDA-PATC50","Drug":"SNS_032","minConc":1,"D1":-3.53,"D2":6.35,"D3":70.37,"D4":90.94,"D5":89.59,"IC50":47.92,"Slope":1.68,"max":90.94,"min":0,"Barcode":"H1FO4B-N1193","W5":"G9","W4":"H9","W3":"I9","W2":"J9","W1":"K9"},{"CellLine":"MDA-PATC50","Drug":"Alvocidib","minConc":1,"D1":-1.69,"D2":0.4,"D3":37.11,"D4":92.66,"D5":99.23,"IC50":144.68,"Slope":1.43,"max":99.23,"min":0,"Barcode":"H1FO4B-N1193","W5":"P9","W4":"O9","W3":"N9","W2":"M9","W1":"L9"},{"CellLine":"MDA-PATC50","Drug":"KX2_391","minConc":1,"D1":-2.93,"D2":-0.17,"D3":50.8,"D4":61.37,"D5":64.17,"IC50":58.79,"Slope":2.5,"max":64.17,"min":0,"Barcode":"H1FO4B-N1193","W5":"F17","W4":"G17","W3":"H17","W2":"I17","W1":"J17"},{"CellLine":"MDA-PATC50","Drug":"AZD8055","minConc":1,"D1":-2.34,"D2":1.55,"D3":31.95,"D4":55.28,"D5":60.94,"IC50":97.12,"Slope":1.24,"max":60.94,"min":0,"Barcode":"H1FO4B-N1193","W5":"G23","W4":"H23","W3":"I23","W2":"J23","W1":"K23"},{"CellLine":"MDA-PATC50","Drug":"PF_03758309","minConc":1,"D1":7.84,"D2":47.76,"D3":73.89,"D4":80.79,"D5":98.94,"IC50":17.46,"Slope":0.58,"max":98.94,"min":0,"Barcode":"H1FO4B-N2193","W5":"P5","W4":"O5","W3":"N5","W2":"M5","W1":"L5"},{"CellLine":"MDA-PATC50","Drug":"AZD_8330","minConc":1,"D1":-1.35,"D2":2.93,"D3":18.02,"D4":45.3,"D5":67.51,"IC50":367.62,"Slope":0.85,"max":67.51,"min":0,"Barcode":"H1FO4B-N2193","W5":"F6","W4":"H6","W3":"I6","W2":"J6","W1":"K6"},{"CellLine":"MDA-PATC50","Drug":"Rigosertib","minConc":1,"D1":5.37,"D2":4.09,"D3":55.73,"D4":66.34,"D5":63.29,"IC50":53.33,"Slope":2.5,"max":66.34,"min":4.09,"Barcode":"H1FO4B-N2193","W5":"F8","W4":"G8","W3":"H8","W2":"J8","W1":"K8"},{"CellLine":"MDA-PATC50","Drug":"Omipalisib","minConc":0.1,"D1":5.43,"D2":6.1,"D3":33.26,"D4":61.67,"D5":76,"IC50":16.38,"Slope":0.87,"max":76,"min":2.85,"Barcode":"H1FO4B-N2193","W5":"P17","W4":"O17","W3":"N17","W2":"M17","W1":"K17"},{"CellLine":"MDA-PATC50","Drug":"UCN_01","minConc":1,"D1":9.55,"D2":8.87,"D3":40.59,"D4":56.77,"D5":100.14,"IC50":418.82,"Slope":0.67,"max":100,"min":6.7,"Barcode":"H1FO4B-N2193","W5":"P21","W4":"O21","W3":"N21","W2":"M21","W1":"L21"},{"CellLine":"MDA-PATC50","Drug":"Dinaciclib","minConc":0.1,"D1":-3.43,"D2":-0.64,"D3":53.46,"D4":93.61,"D5":94.12,"IC50":8.97,"Slope":2.5,"max":94.12,"min":0,"Barcode":"H1FO4B-N3193","W5":"A3","W4":"B3","W3":"C3","W2":"D3","W1":"E3"},{"CellLine":"MDA-PATC50","Drug":"NVP_BGT226","minConc":0.1,"D1":1.16,"D2":0.39,"D3":29.79,"D4":56.54,"D5":98.77,"IC50":46.11,"Slope":0.76,"max":98.77,"min":0,"Barcode":"H1FO4B-N3193","W5":"G12","W4":"H12","W3":"I12","W2":"J12","W1":"K12"},{"CellLine":"MDA-PATC50","Drug":"Gedatolisib","minConc":0.1,"D1":1.37,"D2":0.09,"D3":13.1,"D4":52.75,"D5":77.19,"IC50":46.92,"Slope":1.09,"max":77.19,"min":0.09,"Barcode":"H1FO4B-N3193","W5":"P12","W4":"O12","W3":"N12","W2":"M12","W1":"L12"},{"CellLine":"MDA-PATC50","Drug":"PKI_402","minConc":0.25,"D1":5.64,"D2":6.78,"D3":7.73,"D4":52.32,"D5":71.6,"IC50":156.02,"Slope":1.88,"max":71.6,"min":5.64,"Barcode":"H1FO4B-N3193","W5":"A16","W4":"C16","W3":"D16","W2":"E16","W1":"F16"},{"CellLine":"MDA-PATC50","Drug":"AT7519","minConc":1,"D1":1.05,"D2":1.23,"D3":4.54,"D4":88.78,"D5":94.14,"IC50":337.63,"Slope":2.5,"max":94.14,"min":0.87,"Barcode":"H1FO4B-N3193","W5":"F22","W4":"G22","W3":"H22","W2":"I22","W1":"J22"},{"CellLine":"MDA-PATC50","Drug":"GSK_461364","minConc":1,"D1":0.91,"D2":13.75,"D3":29.44,"D4":34.04,"D5":98.95,"IC50":935.63,"Slope":0.68,"max":98.95,"min":0.91,"Barcode":"H1FO4B-N4193","W5":"A6","W4":"B6","W3":"C6","W2":"D6","W1":"E6"},{"CellLine":"MDA-PATC50","Drug":"Daporinad","minConc":0.1,"D1":-7.32,"D2":-6.88,"D3":43.15,"D4":90.18,"D5":93.03,"IC50":10.62,"Slope":2.5,"max":93.03,"min":0,"Barcode":"H1FO4B-N4193","W5":"A14","W4":"B14","W3":"C14","W2":"D14","W1":"E14"},{"CellLine":"MDA-PATC50","Drug":"Mepacrine","minConc":5,"D1":3.76,"D2":6.48,"D3":1.8,"D4":93.14,"D5":104.01,"IC50":1920.18,"Slope":2.5,"max":100,"min":1.8,"Barcode":"H1FO4B-N4193","W5":"P20","W4":"O20","W3":"N20","W2":"M20","W1":"L20"},{"CellLine":"MDA-PATC50","Drug":"Camptothecin","minConc":0.5,"D1":-4.65,"D2":2.4,"D3":43.56,"D4":87.26,"D5":97.2,"IC50":62.26,"Slope":1.17,"max":97.2,"min":0,"Barcode":"H1FO4B-N5193","W5":"O3","W4":"N3","W3":"M3","W2":"L3","W1":"K3"},{"CellLine":"MDA-PATC50","Drug":"Omacetaxine","minConc":1,"D1":-1.66,"D2":-5.02,"D3":35.25,"D4":92.19,"D5":102.07,"IC50":154.07,"Slope":1.5,"max":100,"min":0,"Barcode":"H1FO4B-N5193","W5":"A5","W4":"B5","W3":"C5","W2":"E5","W1":"F5"},{"CellLine":"MDA-PATC50","Drug":"Vinorelbine","minConc":1,"D1":0.3,"D2":48.84,"D3":62.34,"D4":67.81,"D5":85.36,"IC50":15.65,"Slope":0.6,"max":85.36,"min":0,"Barcode":"H1FO4B-N5193","W5":"P14","W4":"O14","W3":"N14","W2":"M14","W1":"L14"},{"CellLine":"MDA-PATC50","Drug":"Docetaxel","minConc":0.1,"D1":3.95,"D2":26.93,"D3":45.16,"D4":46.52,"D5":42.03,"IC50":0.83,"Slope":1.39,"max":46.52,"min":1.71,"Barcode":"H1FO4B-N5193","W5":"P17","W4":"O17","W3":"N17","W2":"M17","W1":"K17"},{"CellLine":"MDA-PATC50","Drug":"Carfilzomib","minConc":0.1,"D1":2.61,"D2":4.66,"D3":77.74,"D4":101.05,"D5":103.02,"IC50":5.78,"Slope":2.22,"max":100,"min":2.61,"Barcode":"H1FO4B-N5193","W5":"A18","W4":"B18","W3":"C18","W2":"D18","W1":"E18"},{"CellLine":"MDA-PATC50","Drug":"Patupilone","minConc":0.1,"D1":5.8,"D2":7.53,"D3":34.23,"D4":46.44,"D5":41,"IC50":6.12,"Slope":1.72,"max":46.44,"min":5.79,"Barcode":"H1FO4B-N5193","W5":"F18","W4":"G18","W3":"H18","W2":"I18","W1":"J18"},{"CellLine":"MDA-PATC50","Drug":"Plicamycin","minConc":1,"D1":-5.16,"D2":-1.7,"D3":98.98,"D4":98.15,"D5":99.24,"IC50":32.86,"Slope":2.5,"max":99.24,"min":0,"Barcode":"H1FO4B-N5193","W5":"P18","W4":"N18","W3":"M18","W2":"L18","W1":"K18"},{"CellLine":"MDA-PATC50","Drug":"Vincristine","minConc":0.1,"D1":2.72,"D2":6.18,"D3":59,"D4":68.85,"D5":63.88,"IC50":4.2,"Slope":2.01,"max":68.85,"min":2.69,"Barcode":"H1FO4B-N5193","W5":"A19","W4":"B19","W3":"C19","W2":"D19","W1":"E19"},{"CellLine":"MDA-PATC50","Drug":"Dactinomycin","minConc":0.1,"D1":6.27,"D2":47.63,"D3":64.41,"D4":96.05,"D5":101.23,"IC50":2.06,"Slope":0.62,"max":100,"min":0,"Barcode":"H1FO4B-N5193","W5":"P19","W4":"O19","W3":"N19","W2":"M19","W1":"L19"},{"CellLine":"MDA-PATC50","Drug":"Vinblastine","minConc":0.1,"D1":2.62,"D2":-1.64,"D3":55.15,"D4":64.8,"D5":62.28,"IC50":5.06,"Slope":2.5,"max":64.8,"min":0,"Barcode":"H1FO4B-N5193","W5":"A21","W4":"B21","W3":"C21","W2":"D21","W1":"E21"},{"CellLine":"MDA-PATC50","Drug":"Paclitaxel","minConc":0.1,"D1":11.23,"D2":11.9,"D3":40.67,"D4":42.14,"D5":47.62,"IC50":4.97,"Slope":2.04,"max":47.62,"min":10.91,"Barcode":"H1FO4B-N5193","W5":"F21","W4":"G21","W3":"H21","W2":"I21","W1":"J21"},{"CellLine":"MDA-PATC50","Drug":"Oprozomib","minConc":0.25,"D1":-2.63,"D2":-1.85,"D3":12.12,"D4":97.43,"D5":105.16,"IC50":56.09,"Slope":2.45,"max":100,"min":0,"Barcode":"H1FO4B-N5193","W5":"A22","W4":"B22","W3":"C22","W2":"D22","W1":"E22"},{"CellLine":"MDA-PATC50","Drug":"Bortezomib","minConc":0.1,"D1":1.47,"D2":2.15,"D3":96.55,"D4":101.32,"D5":102.06,"IC50":3.28,"Slope":2.5,"max":100,"min":0,"Barcode":"H1FO4B-N5193","W5":"F22","W4":"G22","W3":"H22","W2":"I22","W1":"J22"},{"CellLine":"MDA-PATC50","Drug":"Filanesib","minConc":0.1,"D1":1.5,"D2":2.51,"D3":30.97,"D4":29.49,"D5":32.66,"IC50":3.33,"Slope":2.5,"max":32.66,"min":1.28,"Barcode":"H1FO4B-N5193","W5":"G23","W4":"H23","W3":"I23","W2":"J23","W1":"K23"},{"CellLine":"MDA-PATC50","Drug":"CUDC_907","minConc":1,"D1":21.73,"D2":40.97,"D3":98.02,"D4":99.85,"D5":102.54,"IC50":17.1,"Slope":2.07,"max":100,"min":21.51,"Barcode":"H1FO4B-N6193","W5":"F17","W4":"G17","W3":"H17","W2":"I17","W1":"J17"},{"CellLine":"MDA-PATC50","Drug":"YM155","minConc":1,"D1":11.84,"D2":57.71,"D3":97.9,"D4":102.06,"D5":102.58,"IC50":7.2,"Slope":1.11,"max":100,"min":0,"Barcode":"H1FO4B-N6193","W5":"F18","W4":"G18","W3":"H18","W2":"I18","W1":"J18"},{"CellLine":"MDA-PATC50","Drug":"AT101","minConc":10,"D1":0.77,"D2":-0.95,"D3":9.7,"D4":98.22,"D5":104.33,"IC50":2408.25,"Slope":2.5,"max":100,"min":0,"Barcode":"H1FO4B-N6193","W5":"A21","W4":"B21","W3":"C21","W2":"D21","W1":"E21"},{"CellLine":"MDA-PATC50","Drug":"Selinexor","minConc":1,"D1":10.03,"D2":17.5,"D3":73.02,"D4":99.5,"D5":100,"IC50":54.68,"Slope":1.42,"max":100,"min":9.89,"Barcode":"H1FO4B-N6193","W5":"P21","W4":"O21","W3":"N21","W2":"M21","W1":"L21"},{"CellLine":"MDA-PATC50","Drug":"BIIB021","minConc":1,"D1":8.8,"D2":8.83,"D3":17.51,"D4":82.16,"D5":94.63,"IC50":355.22,"Slope":1.71,"max":94.63,"min":8.72,"Barcode":"H1FO4B-N7193","W5":"P13","W4":"N13","W3":"M13","W2":"L13","W1":"K13"},{"CellLine":"MDA-PATC50","Drug":"Tanespimycin","minConc":1,"D1":7.66,"D2":-1.89,"D3":28.09,"D4":89.31,"D5":90.93,"IC50":143.81,"Slope":2.22,"max":90.93,"min":0,"Barcode":"H1FO4B-N7193","W5":"A14","W4":"B14","W3":"C14","W2":"D14","W1":"E14"},{"CellLine":"MDA-PATC50","Drug":"CUDC_305","minConc":1,"D1":4.27,"D2":10.23,"D3":10.31,"D4":69.24,"D5":94.93,"IC50":545.21,"Slope":1.53,"max":94.93,"min":4.27,"Barcode":"H1FO4B-N7193","W5":"A15","W4":"B15","W3":"C15","W2":"D15","W1":"F15"},{"CellLine":"MDA-PATC50","Drug":"Luminespib","minConc":0.1,"D1":2.82,"D2":2.82,"D3":9.96,"D4":91.41,"D5":91.01,"IC50":24.71,"Slope":2.5,"max":91.41,"min":2.45,"Barcode":"H1FO4B-N7193","W5":"G15","W4":"H15","W3":"I15","W2":"J15","W1":"K15"},{"CellLine":"MDA-PATC50","Drug":"Ganetespib","minConc":0.1,"D1":6.05,"D2":7.99,"D3":2.34,"D4":85.01,"D5":92.88,"IC50":40.82,"Slope":2.5,"max":92.88,"min":2.34,"Barcode":"H1FO4B-N7193","W5":"A16","W4":"C16","W3":"D16","W2":"E16","W1":"F16"},{"CellLine":"MDA-PATC50","Drug":"Stattic","minConc":5,"D1":6.61,"D2":2.59,"D3":-3.83,"D4":36.99,"D5":101.05,"IC50":6187.21,"Slope":2.5,"max":100,"min":0,"Barcode":"H1FO4B-N7193","W5":"F21","W4":"G21","W3":"H21","W2":"I21","W1":"J21"},{"CellLine":"MDA-PATC50","Drug":"ONX_0914","minConc":1,"D1":0.28,"D2":1.77,"D3":24.02,"D4":102.23,"D5":102.92,"IC50":158.61,"Slope":2.5,"max":100,"min":0.28,"Barcode":"H1FO4B-N7193","W5":"G23","W4":"H23","W3":"I23","W2":"J23","W1":"K23"},{"CellLine":"MDA-PATC50","Drug":"SN_38","minConc":1,"D1":20.18,"D2":58.31,"D3":85.23,"D4":98.95,"D5":97.89,"IC50":6.36,"Slope":0.72,"max":98.95,"min":0,"Barcode":"H1FO4B-N8193","W5":"G2","W4":"H2","W3":"I2","W2":"J2","W1":"K2"},{"CellLine":"MDA-PATC50","Drug":"Cabazitaxel","minConc":0.1,"D1":5.59,"D2":25.58,"D3":37.66,"D4":39.92,"D5":42.31,"IC50":0.69,"Slope":0.88,"max":42.31,"min":0,"Barcode":"H1FO4B-N8193","W5":"A3","W4":"B3","W3":"C3","W2":"D3","W1":"E3"},{"CellLine":"MDA-PATC50","Drug":"OTSSP167","minConc":0.1,"D1":4.21,"D2":-4.32,"D3":-5.6,"D4":74.31,"D5":92.83,"IC50":58.13,"Slope":2.5,"max":92.83,"min":0,"Barcode":"H1FO4B-N8193","W5":"A5","W4":"B5","W3":"C5","W2":"E5","W1":"F5"},{"CellLine":"MDA-PATC50","Drug":"Prexasertib","minConc":1,"D1":8.31,"D2":7.03,"D3":69.96,"D4":69.87,"D5":36.73,"IC50":30.46,"Slope":2.5,"max":69.96,"min":5.96,"Barcode":"H1FO4B-N8193","W5":"P6","W4":"O6","W3":"N6","W2":"M6","W1":"L6"},{"CellLine":"MDA-PATC50","Drug":"Romidepsin","minConc":0.1,"D1":10.86,"D2":85.19,"D3":102.14,"D4":101.82,"D5":102.59,"IC50":0.52,"Slope":2.5,"max":100,"min":9.42,"Barcode":"H1FO4B-N8193","W5":"F18","W4":"G18","W3":"H18","W2":"I18","W1":"J18"},{"CellLine":"MDA-PATC50","Drug":"Sabutoclax","minConc":2.5,"D1":10.66,"D2":9.98,"D3":19.74,"D4":95.96,"D5":97.47,"IC50":569.71,"Slope":2.5,"max":97.47,"min":9.98,"Barcode":"H1FO4B-N8193","W5":"F19","W4":"G19","W3":"I19","W2":"J19","W1":"K19"},{"CellLine":"MDA-PATC50","Drug":"BCI","minConc":5,"D1":3.53,"D2":-0.89,"D3":4.39,"D4":97.78,"D5":104.63,"IC50":1484.4,"Slope":2.5,"max":100,"min":0,"Barcode":"H1FO4B-N8193","W5":"A22","W4":"B22","W3":"C22","W2":"D22","W1":"E22"},{"CellLine":"KPC-INK4","Drug":"PF_04691502","minConc":1,"D1":-7.43,"D2":15.51,"D3":56.88,"D4":92.92,"D5":95.27,"IC50":63.69,"Slope":1.04,"max":95.27,"min":0,"Barcode":"H1FO4B-N1169","W5":"B2","W4":"C2","W3":"D2","W2":"E2","W1":"F2"},{"CellLine":"KPC-INK4","Drug":"SNS_032","minConc":1,"D1":13.28,"D2":25.22,"D3":99.08,"D4":99.51,"D5":97.2,"IC50":20.52,"Slope":2.5,"max":99.51,"min":13.11,"Barcode":"H1FO4B-N1169","W5":"G9","W4":"H9","W3":"I9","W2":"J9","W1":"K9"},{"CellLine":"KPC-INK4","Drug":"Alvocidib","minConc":1,"D1":7.94,"D2":18.16,"D3":44.53,"D4":100.47,"D5":100.46,"IC50":126.15,"Slope":1.52,"max":100,"min":7.94,"Barcode":"H1FO4B-N1169","W5":"P9","W4":"O9","W3":"N9","W2":"M9","W1":"L9"},{"CellLine":"KPC-INK4","Drug":"KX2_391","minConc":1,"D1":6.33,"D2":-5.13,"D3":52.86,"D4":63.17,"D5":68.57,"IC50":62.27,"Slope":2.5,"max":68.57,"min":0,"Barcode":"H1FO4B-N1169","W5":"F17","W4":"G17","W3":"H17","W2":"I17","W1":"J17"},{"CellLine":"KPC-INK4","Drug":"AZD8055","minConc":1,"D1":18.7,"D2":43.54,"D3":73.31,"D4":99.12,"D5":98.37,"IC50":24.02,"Slope":0.74,"max":99.12,"min":11.83,"Barcode":"H1FO4B-N1169","W5":"G23","W4":"H23","W3":"I23","W2":"J23","W1":"K23"},{"CellLine":"KPC-INK4","Drug":"PF_03758309","minConc":1,"D1":1.13,"D2":49.66,"D3":63.65,"D4":71.98,"D5":101.8,"IC50":32,"Slope":0.48,"max":100,"min":0,"Barcode":"H1FO4B-N2169","W5":"P5","W4":"O5","W3":"N5","W2":"M5","W1":"L5"},{"CellLine":"KPC-INK4","Drug":"AZD_8330","minConc":1,"D1":-36.55,"D2":-22.89,"D3":1.67,"D4":17.77,"D5":25.04,"IC50":613.88,"Slope":1.84,"max":25.04,"min":0,"Barcode":"H1FO4B-N2169","W5":"F6","W4":"H6","W3":"I6","W2":"J6","W1":"K6"},{"CellLine":"KPC-INK4","Drug":"Rigosertib","minConc":1,"D1":-3.28,"D2":-8.45,"D3":55.99,"D4":61.03,"D5":64.66,"IC50":50.51,"Slope":2.5,"max":64.66,"min":0,"Barcode":"H1FO4B-N2169","W5":"F8","W4":"G8","W3":"H8","W2":"J8","W1":"K8"},{"CellLine":"KPC-INK4","Drug":"Omipalisib","minConc":0.1,"D1":-4.79,"D2":23.18,"D3":52.51,"D4":74.74,"D5":94.72,"IC50":7.93,"Slope":0.66,"max":94.72,"min":0,"Barcode":"H1FO4B-N2169","W5":"P17","W4":"O17","W3":"N17","W2":"M17","W1":"K17"},{"CellLine":"KPC-INK4","Drug":"UCN_01","minConc":1,"D1":18.56,"D2":23.41,"D3":76.53,"D4":85.55,"D5":102.8,"IC50":50.89,"Slope":1.13,"max":100,"min":15.26,"Barcode":"H1FO4B-N2169","W5":"P21","W4":"O21","W3":"N21","W2":"M21","W1":"L21"},{"CellLine":"KPC-INK4","Drug":"Dinaciclib","minConc":0.1,"D1":-19.69,"D2":-23.56,"D3":15.5,"D4":100.96,"D5":100.08,"IC50":19.57,"Slope":2.5,"max":100,"min":0,"Barcode":"H1FO4B-N3169","W5":"A3","W4":"B3","W3":"C3","W2":"D3","W1":"E3"},{"CellLine":"KPC-INK4","Drug":"NVP_BGT226","minConc":0.1,"D1":-7.25,"D2":5.83,"D3":68.15,"D4":88.7,"D5":101.65,"IC50":6.06,"Slope":1.38,"max":100,"min":0,"Barcode":"H1FO4B-N3169","W5":"G12","W4":"H12","W3":"I12","W2":"J12","W1":"K12"},{"CellLine":"KPC-INK4","Drug":"Gedatolisib","minConc":0.1,"D1":-2.29,"D2":12.76,"D3":49.32,"D4":92.85,"D5":100.55,"IC50":9.62,"Slope":0.98,"max":100,"min":0,"Barcode":"H1FO4B-N3169","W5":"P12","W4":"O12","W3":"N12","W2":"M12","W1":"L12"},{"CellLine":"KPC-INK4","Drug":"PKI_402","minConc":0.25,"D1":5.68,"D2":6.19,"D3":63.74,"D4":96.97,"D5":99.56,"IC50":19.29,"Slope":1.89,"max":99.56,"min":4.97,"Barcode":"H1FO4B-N3169","W5":"A16","W4":"C16","W3":"D16","W2":"E16","W1":"F16"},{"CellLine":"KPC-INK4","Drug":"AT7519","minConc":1,"D1":-7.82,"D2":10.29,"D3":78.95,"D4":97.23,"D5":99.14,"IC50":41.46,"Slope":1.55,"max":99.14,"min":0,"Barcode":"H1FO4B-N3169","W5":"F22","W4":"G22","W3":"H22","W2":"I22","W1":"J22"},{"CellLine":"KPC-INK4","Drug":"GSK_461364","minConc":1,"D1":-9.9,"D2":-1,"D3":43.49,"D4":68.08,"D5":101.22,"IC50":225.32,"Slope":0.79,"max":100,"min":0,"Barcode":"H1FO4B-N4169","W5":"A6","W4":"B6","W3":"C6","W2":"D6","W1":"E6"},{"CellLine":"KPC-INK4","Drug":"Daporinad","minConc":0.1,"D1":-12.88,"D2":-19.3,"D3":27.71,"D4":91.44,"D5":94.05,"IC50":14.23,"Slope":2.5,"max":94.05,"min":0,"Barcode":"H1FO4B-N4169","W5":"A14","W4":"B14","W3":"C14","W2":"D14","W1":"E14"},{"CellLine":"KPC-INK4","Drug":"Mepacrine","minConc":5,"D1":8.59,"D2":-0.97,"D3":13.52,"D4":103.5,"D5":105.03,"IC50":1022.78,"Slope":2.5,"max":100,"min":0,"Barcode":"H1FO4B-N4169","W5":"P20","W4":"O20","W3":"N20","W2":"M20","W1":"L20"},{"CellLine":"KPC-INK4","Drug":"Camptothecin","minConc":0.5,"D1":5.54,"D2":-1.8,"D3":8.3,"D4":27.86,"D5":70.56,"IC50":637.22,"Slope":1.45,"max":70.56,"min":0,"Barcode":"H1FO4B-N5169","W5":"O3","W4":"N3","W3":"M3","W2":"L3","W1":"K3"},{"CellLine":"KPC-INK4","Drug":"Omacetaxine","minConc":1,"D1":9.46,"D2":14.69,"D3":86.08,"D4":97.26,"D5":97.47,"IC50":38.97,"Slope":2.02,"max":97.47,"min":9.41,"Barcode":"H1FO4B-N5169","W5":"A5","W4":"B5","W3":"C5","W2":"E5","W1":"F5"},{"CellLine":"KPC-INK4","Drug":"Vinorelbine","minConc":1,"D1":2.24,"D2":59.52,"D3":68.51,"D4":71.71,"D5":87.17,"IC50":6.82,"Slope":1.01,"max":87.17,"min":0,"Barcode":"H1FO4B-N5169","W5":"P14","W4":"O14","W3":"N14","W2":"M14","W1":"L14"},{"CellLine":"KPC-INK4","Drug":"Docetaxel","minConc":0.1,"D1":8.65,"D2":12.69,"D3":52.89,"D4":69.16,"D5":65.59,"IC50":5.3,"Slope":1.58,"max":69.16,"min":8.59,"Barcode":"H1FO4B-N5169","W5":"P17","W4":"O17","W3":"N17","W2":"M17","W1":"K17"},{"CellLine":"KPC-INK4","Drug":"Carfilzomib","minConc":0.1,"D1":-6.26,"D2":5.46,"D3":90.27,"D4":95.9,"D5":99.07,"IC50":3.55,"Slope":2.25,"max":99.07,"min":0,"Barcode":"H1FO4B-N5169","W5":"A18","W4":"B18","W3":"C18","W2":"D18","W1":"E18"},{"CellLine":"KPC-INK4","Drug":"Patupilone","minConc":0.1,"D1":17.23,"D2":-5.72,"D3":62.02,"D4":64.08,"D5":58.25,"IC50":3.84,"Slope":2.5,"max":64.08,"min":0,"Barcode":"H1FO4B-N5169","W5":"F18","W4":"G18","W3":"H18","W2":"I18","W1":"J18"},{"CellLine":"KPC-INK4","Drug":"Plicamycin","minConc":1,"D1":16.11,"D2":15.17,"D3":8.04,"D4":85.7,"D5":98.77,"IC50":494.3,"Slope":2.5,"max":98.77,"min":8.04,"Barcode":"H1FO4B-N5169","W5":"P18","W4":"N18","W3":"M18","W2":"L18","W1":"K18"},{"CellLine":"KPC-INK4","Drug":"Vincristine","minConc":0.1,"D1":5.61,"D2":4.92,"D3":61.4,"D4":67.76,"D5":59.22,"IC50":4.25,"Slope":2.5,"max":67.76,"min":4.47,"Barcode":"H1FO4B-N5169","W5":"A19","W4":"B19","W3":"C19","W2":"D19","W1":"E19"},{"CellLine":"KPC-INK4","Drug":"Dactinomycin","minConc":0.1,"D1":15.67,"D2":54.45,"D3":82.73,"D4":100.22,"D5":99.24,"IC50":0.87,"Slope":0.73,"max":100,"min":0,"Barcode":"H1FO4B-N5169","W5":"P19","W4":"O19","W3":"N19","W2":"M19","W1":"L19"},{"CellLine":"KPC-INK4","Drug":"Vinblastine","minConc":0.1,"D1":-6.28,"D2":12.45,"D3":45.29,"D4":55.74,"D5":56.82,"IC50":3.12,"Slope":1.23,"max":56.82,"min":0,"Barcode":"H1FO4B-N5169","W5":"A21","W4":"B21","W3":"C21","W2":"D21","W1":"E21"},{"CellLine":"KPC-INK4","Drug":"Paclitaxel","minConc":0.1,"D1":2.23,"D2":2.33,"D3":46.16,"D4":70.27,"D5":63.17,"IC50":7.86,"Slope":2.5,"max":70.27,"min":2.08,"Barcode":"H1FO4B-N5169","W5":"F21","W4":"G21","W3":"H21","W2":"I21","W1":"J21"},{"CellLine":"KPC-INK4","Drug":"Oprozomib","minConc":0.25,"D1":6.42,"D2":6.36,"D3":17.35,"D4":93.49,"D5":99.29,"IC50":66.59,"Slope":2.05,"max":99.29,"min":6.33,"Barcode":"H1FO4B-N5169","W5":"A22","W4":"B22","W3":"C22","W2":"D22","W1":"E22"},{"CellLine":"KPC-INK4","Drug":"Bortezomib","minConc":0.1,"D1":4.71,"D2":-0.36,"D3":95.06,"D4":97.23,"D5":96.46,"IC50":3.35,"Slope":2.5,"max":97.23,"min":0,"Barcode":"H1FO4B-N5169","W5":"F22","W4":"G22","W3":"H22","W2":"I22","W1":"J22"},{"CellLine":"KPC-INK4","Drug":"Filanesib","minConc":0.1,"D1":5.12,"D2":2.95,"D3":51.07,"D4":43.67,"D5":41.83,"IC50":3.15,"Slope":2.5,"max":51.07,"min":2.87,"Barcode":"H1FO4B-N5169","W5":"G23","W4":"H23","W3":"I23","W2":"J23","W1":"K23"},{"CellLine":"KPC-INK4","Drug":"CUDC_907","minConc":1,"D1":-33.98,"D2":-54.88,"D3":91.68,"D4":98.89,"D5":100.88,"IC50":50.78,"Slope":2.5,"max":100,"min":0,"Barcode":"H1FO4B-N6169","W5":"F17","W4":"G17","W3":"H17","W2":"I17","W1":"J17"},{"CellLine":"KPC-INK4","Drug":"YM155","minConc":1,"D1":-1.34,"D2":-6.9,"D3":1.7,"D4":76.4,"D5":100.82,"IC50":602,"Slope":2.32,"max":100,"min":0,"Barcode":"H1FO4B-N6169","W5":"F18","W4":"G18","W3":"H18","W2":"I18","W1":"J18"},{"CellLine":"KPC-INK4","Drug":"AT101","minConc":10,"D1":-1.4,"D2":7.77,"D3":-7.96,"D4":80.5,"D5":100.23,"IC50":5770.64,"Slope":2.5,"max":100,"min":0,"Barcode":"H1FO4B-N6169","W5":"A21","W4":"B21","W3":"C21","W2":"D21","W1":"E21"},{"CellLine":"KPC-INK4","Drug":"Selinexor","minConc":1,"D1":1.29,"D2":8.51,"D3":63.02,"D4":89.33,"D5":94.1,"IC50":58.53,"Slope":1.27,"max":94.1,"min":0.2,"Barcode":"H1FO4B-N6169","W5":"P21","W4":"O21","W3":"N21","W2":"M21","W1":"L21"},{"CellLine":"KPC-INK4","Drug":"BIIB021","minConc":1,"D1":3.6,"D2":10.85,"D3":64.95,"D4":98.44,"D5":101.46,"IC50":65.76,"Slope":1.36,"max":100,"min":3.56,"Barcode":"H1FO4B-N7169","W5":"P13","W4":"N13","W3":"M13","W2":"L13","W1":"K13"},{"CellLine":"KPC-INK4","Drug":"Tanespimycin","minConc":1,"D1":-4.41,"D2":-16.1,"D3":74.4,"D4":97.79,"D5":99.24,"IC50":65.58,"Slope":2.5,"max":99.24,"min":0,"Barcode":"H1FO4B-N7169","W5":"A14","W4":"B14","W3":"C14","W2":"D14","W1":"E14"},{"CellLine":"KPC-INK4","Drug":"CUDC_305","minConc":1,"D1":-8.51,"D2":-6.95,"D3":31.81,"D4":96.42,"D5":100.86,"IC50":146.1,"Slope":2.05,"max":100,"min":0,"Barcode":"H1FO4B-N7169","W5":"A15","W4":"B15","W3":"C15","W2":"D15","W1":"F15"},{"CellLine":"KPC-INK4","Drug":"Luminespib","minConc":0.1,"D1":6.44,"D2":-2.01,"D3":82.71,"D4":101.08,"D5":99.35,"IC50":5.4,"Slope":2.5,"max":100,"min":0,"Barcode":"H1FO4B-N7169","W5":"G15","W4":"H15","W3":"I15","W2":"J15","W1":"K15"},{"CellLine":"KPC-INK4","Drug":"Ganetespib","minConc":0.1,"D1":-4.01,"D2":-10,"D3":62.03,"D4":102.37,"D5":103.75,"IC50":8.25,"Slope":2.5,"max":100,"min":0,"Barcode":"H1FO4B-N7169","W5":"A16","W4":"C16","W3":"D16","W2":"E16","W1":"F16"},{"CellLine":"KPC-INK4","Drug":"Stattic","minConc":5,"D1":-1.95,"D2":-5.83,"D3":-8.41,"D4":100.23,"D5":106.32,"IC50":1781.87,"Slope":2.5,"max":100,"min":0,"Barcode":"H1FO4B-N7169","W5":"F21","W4":"G21","W3":"H21","W2":"I21","W1":"J21"},{"CellLine":"KPC-INK4","Drug":"ONX_0914","minConc":1,"D1":11.67,"D2":0.54,"D3":4.25,"D4":98.88,"D5":101.11,"IC50":295.46,"Slope":2.5,"max":100,"min":0.54,"Barcode":"H1FO4B-N7169","W5":"G23","W4":"H23","W3":"I23","W2":"J23","W1":"K23"},{"CellLine":"KPC-INK4","Drug":"SN_38","minConc":1,"D1":5.1,"D2":10.88,"D3":21.6,"D4":86.33,"D5":100.45,"IC50":290.58,"Slope":1.43,"max":100,"min":5.1,"Barcode":"H1FO4B-N8169","W5":"G2","W4":"H2","W3":"I2","W2":"J2","W1":"K2"},{"CellLine":"KPC-INK4","Drug":"Cabazitaxel","minConc":0.1,"D1":6.1,"D2":15.58,"D3":62.63,"D4":62.04,"D5":56.59,"IC50":1.88,"Slope":2.5,"max":62.63,"min":6,"Barcode":"H1FO4B-N8169","W5":"A3","W4":"B3","W3":"C3","W2":"D3","W1":"E3"},{"CellLine":"KPC-INK4","Drug":"OTSSP167","minConc":0.1,"D1":-12.05,"D2":0.85,"D3":-4.18,"D4":58.14,"D5":100.58,"IC50":87.8,"Slope":2.5,"max":100,"min":0,"Barcode":"H1FO4B-N8169","W5":"A5","W4":"B5","W3":"C5","W2":"E5","W1":"F5"},{"CellLine":"KPC-INK4","Drug":"Prexasertib","minConc":1,"D1":-4.63,"D2":-6.46,"D3":67.09,"D4":69.62,"D5":72.12,"IC50":42.06,"Slope":2.5,"max":72.12,"min":0,"Barcode":"H1FO4B-N8169","W5":"P6","W4":"O6","W3":"N6","W2":"M6","W1":"L6"},{"CellLine":"KPC-INK4","Drug":"Romidepsin","minConc":0.1,"D1":-11.32,"D2":20.98,"D3":97.55,"D4":99.22,"D5":96.98,"IC50":1.7,"Slope":2.5,"max":99.22,"min":0,"Barcode":"H1FO4B-N8169","W5":"F18","W4":"G18","W3":"H18","W2":"I18","W1":"J18"},{"CellLine":"KPC-INK4","Drug":"Sabutoclax","minConc":2.5,"D1":-13.79,"D2":-7.33,"D3":51.63,"D4":81.32,"D5":92.59,"IC50":228.81,"Slope":2.5,"max":92.59,"min":0,"Barcode":"H1FO4B-N8169","W5":"F19","W4":"G19","W3":"I19","W2":"J19","W1":"K19"},{"CellLine":"KPC-INK4","Drug":"BCI","minConc":5,"D1":-12.18,"D2":-12.41,"D3":-18.07,"D4":101.79,"D5":105,"IC50":1886.07,"Slope":2.5,"max":100,"min":0,"Barcode":"H1FO4B-N8169","W5":"A22","W4":"B22","W3":"C22","W2":"D22","W1":"E22"},{"CellLine":"KPC-p53","Drug":"PF_04691502","minConc":1,"D1":1.19,"D2":11.05,"D3":17.81,"D4":93.5,"D5":98.01,"IC50":222.51,"Slope":1.94,"max":98.01,"min":1.19,"Barcode":"H1FO4B-N1185","W5":"B2","W4":"C2","W3":"D2","W2":"E2","W1":"F2"},{"CellLine":"KPC-p53","Drug":"SNS_032","minConc":1,"D1":4.6,"D2":-7.6,"D3":14.6,"D4":97.8,"D5":97.8,"IC50":199.36,"Slope":2.5,"max":97.8,"min":0,"Barcode":"H1FO4B-N1185","W5":"G9","W4":"H9","W3":"I9","W2":"J9","W1":"K9"},{"CellLine":"KPC-p53","Drug":"Alvocidib","minConc":1,"D1":8.54,"D2":17.27,"D3":36.22,"D4":97.98,"D5":101.69,"IC50":159.14,"Slope":1.73,"max":100,"min":8.54,"Barcode":"H1FO4B-N1185","W5":"P9","W4":"O9","W3":"N9","W2":"M9","W1":"L9"},{"CellLine":"KPC-p53","Drug":"KX2_391","minConc":1,"D1":-1.51,"D2":-6.95,"D3":57.93,"D4":67.92,"D5":69.77,"IC50":54.61,"Slope":2.5,"max":69.77,"min":0,"Barcode":"H1FO4B-N1185","W5":"F17","W4":"G17","W3":"H17","W2":"I17","W1":"J17"},{"CellLine":"KPC-p53","Drug":"AZD8055","minConc":1,"D1":9.35,"D2":23.61,"D3":62.49,"D4":91.59,"D5":97.19,"IC50":57.61,"Slope":0.89,"max":97.19,"min":7.42,"Barcode":"H1FO4B-N1185","W5":"G23","W4":"H23","W3":"I23","W2":"J23","W1":"K23"},{"CellLine":"KPC-p53","Drug":"PF_03758309","minConc":1,"D1":-0.92,"D2":40.6,"D3":65.16,"D4":67.12,"D5":95.15,"IC50":36.14,"Slope":0.52,"max":95.15,"min":0,"Barcode":"H1FO4B-N2185","W5":"P5","W4":"O5","W3":"N5","W2":"M5","W1":"L5"},{"CellLine":"KPC-p53","Drug":"AZD_8330","minConc":1,"D1":-11.99,"D2":-1.15,"D3":4.23,"D4":47.31,"D5":48.87,"IC50":256.66,"Slope":2.5,"max":48.87,"min":0,"Barcode":"H1FO4B-N2185","W5":"F6","W4":"H6","W3":"I6","W2":"J6","W1":"K6"},{"CellLine":"KPC-p53","Drug":"Rigosertib","minConc":1,"D1":-11.29,"D2":-16.24,"D3":61.84,"D4":62.65,"D5":59.4,"IC50":41.45,"Slope":2.5,"max":62.65,"min":0,"Barcode":"H1FO4B-N2185","W5":"F8","W4":"G8","W3":"H8","W2":"J8","W1":"K8"},{"CellLine":"KPC-p53","Drug":"Omipalisib","minConc":0.1,"D1":0.1,"D2":5.37,"D3":42.36,"D4":86.36,"D5":98.55,"IC50":13.51,"Slope":1.02,"max":98.55,"min":0,"Barcode":"H1FO4B-N2185","W5":"P17","W4":"O17","W3":"N17","W2":"M17","W1":"K17"},{"CellLine":"KPC-p53","Drug":"UCN_01","minConc":1,"D1":0.86,"D2":19.13,"D3":89.33,"D4":100.53,"D5":102.18,"IC50":25.47,"Slope":1.56,"max":100,"min":0.23,"Barcode":"H1FO4B-N2185","W5":"P21","W4":"O21","W3":"N21","W2":"M21","W1":"L21"},{"CellLine":"KPC-p53","Drug":"Dinaciclib","minConc":0.1,"D1":-2.24,"D2":5.44,"D3":32.84,"D4":101.17,"D5":101.2,"IC50":13.29,"Slope":2.5,"max":100,"min":0,"Barcode":"H1FO4B-N3185","W5":"A3","W4":"B3","W3":"C3","W2":"D3","W1":"E3"},{"CellLine":"KPC-p53","Drug":"NVP_BGT226","minConc":0.1,"D1":11.98,"D2":5.12,"D3":67.32,"D4":86.89,"D5":100.42,"IC50":6.94,"Slope":1.57,"max":100,"min":5.12,"Barcode":"H1FO4B-N3185","W5":"G12","W4":"H12","W3":"I12","W2":"J12","W1":"K12"},{"CellLine":"KPC-p53","Drug":"Gedatolisib","minConc":0.1,"D1":6.87,"D2":5.31,"D3":39.05,"D4":78.36,"D5":96.03,"IC50":18.16,"Slope":0.93,"max":96.03,"min":3.41,"Barcode":"H1FO4B-N3185","W5":"P12","W4":"O12","W3":"N12","W2":"M12","W1":"L12"},{"CellLine":"KPC-p53","Drug":"PKI_402","minConc":0.25,"D1":-8.36,"D2":-12.93,"D3":36.79,"D4":88.6,"D5":97.58,"IC50":35.76,"Slope":1.61,"max":97.58,"min":0,"Barcode":"H1FO4B-N3185","W5":"A16","W4":"C16","W3":"D16","W2":"E16","W1":"F16"},{"CellLine":"KPC-p53","Drug":"AT7519","minConc":1,"D1":-9.24,"D2":13.88,"D3":16.59,"D4":91.21,"D5":99.21,"IC50":250.13,"Slope":1.69,"max":99.21,"min":0,"Barcode":"H1FO4B-N3185","W5":"F22","W4":"G22","W3":"H22","W2":"I22","W1":"J22"},{"CellLine":"KPC-p53","Drug":"GSK_461364","minConc":1,"D1":-11.08,"D2":3.95,"D3":26.63,"D4":55.47,"D5":102.05,"IC50":525.31,"Slope":0.81,"max":100,"min":0,"Barcode":"H1FO4B-N4185","W5":"A6","W4":"B6","W3":"C6","W2":"D6","W1":"E6"},{"CellLine":"KPC-p53","Drug":"Daporinad","minConc":0.1,"D1":-19.64,"D2":-7.68,"D3":22.49,"D4":96.49,"D5":98.84,"IC50":17.04,"Slope":2.31,"max":98.84,"min":0,"Barcode":"H1FO4B-N4185","W5":"A14","W4":"B14","W3":"C14","W2":"D14","W1":"E14"},{"CellLine":"KPC-p53","Drug":"Mepacrine","minConc":5,"D1":4.44,"D2":-1.62,"D3":16.31,"D4":98.87,"D5":103.79,"IC50":960.33,"Slope":2.5,"max":100,"min":0,"Barcode":"H1FO4B-N4185","W5":"P20","W4":"O20","W3":"N20","W2":"M20","W1":"L20"},{"CellLine":"KPC-p53","Drug":"Camptothecin","minConc":0.5,"D1":-0.77,"D2":6.51,"D3":18.54,"D4":45.23,"D5":97.29,"IC50":432.79,"Slope":0.9,"max":97.29,"min":0,"Barcode":"H1FO4B-N5185","W5":"O3","W4":"N3","W3":"M3","W2":"L3","W1":"K3"},{"CellLine":"KPC-p53","Drug":"Omacetaxine","minConc":1,"D1":-5.35,"D2":-3.21,"D3":82.29,"D4":97.76,"D5":102.99,"IC50":54.74,"Slope":2.5,"max":100,"min":0,"Barcode":"H1FO4B-N5185","W5":"A5","W4":"B5","W3":"C5","W2":"E5","W1":"F5"},{"CellLine":"KPC-p53","Drug":"Vinorelbine","minConc":1,"D1":7.2,"D2":62.21,"D3":70.6,"D4":71.87,"D5":84.6,"IC50":5.02,"Slope":1.23,"max":84.6,"min":0,"Barcode":"H1FO4B-N5185","W5":"P14","W4":"O14","W3":"N14","W2":"M14","W1":"L14"},{"CellLine":"KPC-p53","Drug":"Docetaxel","minConc":0.1,"D1":-3.23,"D2":17.79,"D3":56.88,"D4":63.11,"D5":63.68,"IC50":2.03,"Slope":1.4,"max":63.68,"min":0,"Barcode":"H1FO4B-N5185","W5":"P17","W4":"O17","W3":"N17","W2":"M17","W1":"K17"},{"CellLine":"KPC-p53","Drug":"Carfilzomib","minConc":0.1,"D1":-7.49,"D2":-1.89,"D3":94.86,"D4":99.89,"D5":102.98,"IC50":3.72,"Slope":2.5,"max":100,"min":0,"Barcode":"H1FO4B-N5185","W5":"A18","W4":"B18","W3":"C18","W2":"D18","W1":"E18"},{"CellLine":"KPC-p53","Drug":"Patupilone","minConc":0.1,"D1":7.77,"D2":-2.17,"D3":17.44,"D4":54.79,"D5":51.15,"IC50":13.56,"Slope":2.5,"max":54.79,"min":0,"Barcode":"H1FO4B-N5185","W5":"F18","W4":"G18","W3":"H18","W2":"I18","W1":"J18"},{"CellLine":"KPC-p53","Drug":"Plicamycin","minConc":1,"D1":-15.99,"D2":6.73,"D3":9.92,"D4":90.98,"D5":99.45,"IC50":302.07,"Slope":1.97,"max":99.45,"min":0,"Barcode":"H1FO4B-N5185","W5":"P18","W4":"N18","W3":"M18","W2":"L18","W1":"K18"},{"CellLine":"KPC-p53","Drug":"Vincristine","minConc":0.1,"D1":-4.74,"D2":0.62,"D3":66.35,"D4":74.81,"D5":69.6,"IC50":4.45,"Slope":2.5,"max":74.81,"min":0,"Barcode":"H1FO4B-N5185","W5":"A19","W4":"B19","W3":"C19","W2":"D19","W1":"E19"},{"CellLine":"KPC-p53","Drug":"Dactinomycin","minConc":0.1,"D1":8.58,"D2":40.07,"D3":93.26,"D4":103.02,"D5":103.11,"IC50":1.52,"Slope":1.39,"max":100,"min":6.5,"Barcode":"H1FO4B-N5185","W5":"P19","W4":"O19","W3":"N19","W2":"M19","W1":"L19"},{"CellLine":"KPC-p53","Drug":"Vinblastine","minConc":0.1,"D1":-1.09,"D2":-1.87,"D3":58.23,"D4":65.65,"D5":73.51,"IC50":5.9,"Slope":2.5,"max":73.51,"min":0,"Barcode":"H1FO4B-N5185","W5":"A21","W4":"B21","W3":"C21","W2":"D21","W1":"E21"},{"CellLine":"KPC-p53","Drug":"Paclitaxel","minConc":0.1,"D1":8.42,"D2":15.5,"D3":42.34,"D4":71.57,"D5":56.23,"IC50":8.19,"Slope":1.06,"max":71.57,"min":8.42,"Barcode":"H1FO4B-N5185","W5":"F21","W4":"G21","W3":"H21","W2":"I21","W1":"J21"},{"CellLine":"KPC-p53","Drug":"Oprozomib","minConc":0.25,"D1":2.3,"D2":4.27,"D3":23.59,"D4":93.79,"D5":102.42,"IC50":52.43,"Slope":1.72,"max":100,"min":2.3,"Barcode":"H1FO4B-N5185","W5":"A22","W4":"B22","W3":"C22","W2":"D22","W1":"E22"},{"CellLine":"KPC-p53","Drug":"Bortezomib","minConc":0.1,"D1":15.33,"D2":7.69,"D3":98.57,"D4":99.79,"D5":101.78,"IC50":3.27,"Slope":2.5,"max":100,"min":7.69,"Barcode":"H1FO4B-N5185","W5":"F22","W4":"G22","W3":"H22","W2":"I22","W1":"J22"},{"CellLine":"KPC-p53","Drug":"Filanesib","minConc":0.1,"D1":8.1,"D2":5.09,"D3":55.86,"D4":54.23,"D5":49.53,"IC50":3.16,"Slope":2.5,"max":55.86,"min":5.09,"Barcode":"H1FO4B-N5185","W5":"G23","W4":"H23","W3":"I23","W2":"J23","W1":"K23"},{"CellLine":"KPC-p53","Drug":"CUDC_907","minConc":1,"D1":4.38,"D2":2.07,"D3":85.93,"D4":101.95,"D5":101.63,"IC50":49.37,"Slope":2.5,"max":100,"min":2.07,"Barcode":"H1FO4B-N6185","W5":"F17","W4":"G17","W3":"H17","W2":"I17","W1":"J17"},{"CellLine":"KPC-p53","Drug":"YM155","minConc":1,"D1":10.31,"D2":-3.73,"D3":6.11,"D4":28.26,"D5":103.39,"IC50":1445.8,"Slope":2.5,"max":100,"min":0,"Barcode":"H1FO4B-N6185","W5":"F18","W4":"G18","W3":"H18","W2":"I18","W1":"J18"},{"CellLine":"KPC-p53","Drug":"AT101","minConc":10,"D1":0.95,"D2":10.25,"D3":7.94,"D4":86.44,"D5":104.81,"IC50":3807.82,"Slope":1.91,"max":100,"min":0.95,"Barcode":"H1FO4B-N6185","W5":"A21","W4":"B21","W3":"C21","W2":"D21","W1":"E21"},{"CellLine":"KPC-p53","Drug":"Selinexor","minConc":1,"D1":11.86,"D2":24.51,"D3":63.53,"D4":94.08,"D5":95.19,"IC50":58.4,"Slope":1.04,"max":95.19,"min":11.61,"Barcode":"H1FO4B-N6185","W5":"P21","W4":"O21","W3":"N21","W2":"M21","W1":"L21"},{"CellLine":"KPC-p53","Drug":"BIIB021","minConc":1,"D1":-0.33,"D2":7.31,"D3":76.23,"D4":95.07,"D5":99.44,"IC50":47.77,"Slope":1.59,"max":99.44,"min":0,"Barcode":"H1FO4B-N7185","W5":"P13","W4":"N13","W3":"M13","W2":"L13","W1":"K13"},{"CellLine":"KPC-p53","Drug":"Tanespimycin","minConc":1,"D1":3.32,"D2":-3.09,"D3":47.35,"D4":88.28,"D5":97.5,"IC50":109.52,"Slope":1.39,"max":97.5,"min":0,"Barcode":"H1FO4B-N7185","W5":"A14","W4":"B14","W3":"C14","W2":"D14","W1":"E14"},{"CellLine":"KPC-p53","Drug":"CUDC_305","minConc":1,"D1":-2.9,"D2":8.76,"D3":40.43,"D4":96.29,"D5":88.86,"IC50":120.2,"Slope":1.46,"max":96.29,"min":0,"Barcode":"H1FO4B-N7185","W5":"A15","W4":"B15","W3":"C15","W2":"D15","W1":"F15"},{"CellLine":"KPC-p53","Drug":"Luminespib","minConc":0.1,"D1":6.9,"D2":3.51,"D3":81.3,"D4":96.8,"D5":92.6,"IC50":5.27,"Slope":2.5,"max":96.8,"min":3.51,"Barcode":"H1FO4B-N7185","W5":"G15","W4":"H15","W3":"I15","W2":"J15","W1":"K15"},{"CellLine":"KPC-p53","Drug":"Ganetespib","minConc":0.1,"D1":-5.95,"D2":2.76,"D3":55.09,"D4":95.8,"D5":98.12,"IC50":8.61,"Slope":1.64,"max":98.12,"min":0,"Barcode":"H1FO4B-N7185","W5":"A16","W4":"C16","W3":"D16","W2":"E16","W1":"F16"},{"CellLine":"KPC-p53","Drug":"Stattic","minConc":5,"D1":9.36,"D2":11.64,"D3":-3.82,"D4":101.47,"D5":105.82,"IC50":1641.74,"Slope":2.5,"max":100,"min":0,"Barcode":"H1FO4B-N7185","W5":"F21","W4":"G21","W3":"H21","W2":"I21","W1":"J21"},{"CellLine":"KPC-p53","Drug":"ONX_0914","minConc":1,"D1":15.11,"D2":13.37,"D3":16.1,"D4":101.67,"D5":104.75,"IC50":280.12,"Slope":2.5,"max":100,"min":12.94,"Barcode":"H1FO4B-N7185","W5":"G23","W4":"H23","W3":"I23","W2":"J23","W1":"K23"},{"CellLine":"KPC-p53","Drug":"SN_38","minConc":1,"D1":8.82,"D2":28.15,"D3":36.51,"D4":82.57,"D5":98.58,"IC50":184.55,"Slope":0.77,"max":98.58,"min":8.82,"Barcode":"H1FO4B-N8185","W5":"G2","W4":"H2","W3":"I2","W2":"J2","W1":"K2"},{"CellLine":"KPC-p53","Drug":"Cabazitaxel","minConc":0.1,"D1":9.45,"D2":6.58,"D3":62.81,"D4":70.12,"D5":54.36,"IC50":4.5,"Slope":2.5,"max":70.12,"min":6.58,"Barcode":"H1FO4B-N8185","W5":"A3","W4":"B3","W3":"C3","W2":"D3","W1":"E3"},{"CellLine":"KPC-p53","Drug":"OTSSP167","minConc":0.1,"D1":-0.91,"D2":1.71,"D3":16.9,"D4":67.07,"D5":102.36,"IC50":48.01,"Slope":1.09,"max":100,"min":0,"Barcode":"H1FO4B-N8185","W5":"A5","W4":"B5","W3":"C5","W2":"E5","W1":"F5"},{"CellLine":"KPC-p53","Drug":"Prexasertib","minConc":1,"D1":6.84,"D2":8.39,"D3":92.9,"D4":91.75,"D5":87.96,"IC50":28.57,"Slope":2.5,"max":92.9,"min":4.78,"Barcode":"H1FO4B-N8185","W5":"P6","W4":"O6","W3":"N6","W2":"M6","W1":"L6"},{"CellLine":"KPC-p53","Drug":"Romidepsin","minConc":0.1,"D1":5.41,"D2":13.82,"D3":99.18,"D4":102.81,"D5":102.18,"IC50":2.41,"Slope":2.5,"max":100,"min":4.92,"Barcode":"H1FO4B-N8185","W5":"F18","W4":"G18","W3":"H18","W2":"I18","W1":"J18"},{"CellLine":"KPC-p53","Drug":"Sabutoclax","minConc":2.5,"D1":-1.34,"D2":9.86,"D3":15.08,"D4":88.79,"D5":92.01,"IC50":541.56,"Slope":2.08,"max":92.01,"min":0,"Barcode":"H1FO4B-N8185","W5":"F19","W4":"G19","W3":"I19","W2":"J19","W1":"K19"},{"CellLine":"KPC-p53","Drug":"BCI","minConc":5,"D1":-5.41,"D2":3.29,"D3":8.27,"D4":102.56,"D5":101.74,"IC50":1208.77,"Slope":2.5,"max":100,"min":0,"Barcode":"H1FO4B-N8185","W5":"A22","W4":"B22","W3":"C22","W2":"D22","W1":"E22"},{"CellLine":"AsPC-1","Drug":"PF_04691502","minConc":1,"D1":-0.46,"D2":2.36,"D3":9.71,"D4":66.33,"D5":91.09,"IC50":480.26,"Slope":1.36,"max":91.09,"min":0,"Barcode":"H1FO4B-N1278","W5":"B2","W4":"C2","W3":"D2","W2":"E2","W1":"F2"},{"CellLine":"AsPC-1","Drug":"SNS_032","minConc":1,"D1":4.26,"D2":7.6,"D3":49.29,"D4":69.67,"D5":73.4,"IC50":64.11,"Slope":1.39,"max":73.4,"min":3.42,"Barcode":"H1FO4B-N1278","W5":"G9","W4":"H9","W3":"I9","W2":"J9","W1":"K9"},{"CellLine":"AsPC-1","Drug":"Alvocidib","minConc":1,"D1":7.03,"D2":5.67,"D3":32.93,"D4":70.81,"D5":74.63,"IC50":134.06,"Slope":1.5,"max":74.63,"min":5.67,"Barcode":"H1FO4B-N1278","W5":"P9","W4":"O9","W3":"N9","W2":"M9","W1":"L9"},{"CellLine":"AsPC-1","Drug":"KX2_391","minConc":1,"D1":5.83,"D2":13.93,"D3":53.41,"D4":64.53,"D5":76.09,"IC50":48.96,"Slope":0.84,"max":76.09,"min":1.49,"Barcode":"H1FO4B-N1278","W5":"F17","W4":"G17","W3":"H17","W2":"I17","W1":"J17"},{"CellLine":"AsPC-1","Drug":"AZD8055","minConc":1,"D1":5.35,"D2":11.09,"D3":49.21,"D4":58.53,"D5":69.25,"IC50":50.81,"Slope":0.92,"max":69.25,"min":1.86,"Barcode":"H1FO4B-N1278","W5":"G23","W4":"H23","W3":"I23","W2":"J23","W1":"K23"},{"CellLine":"AsPC-1","Drug":"PF_03758309","minConc":1,"D1":3.47,"D2":18.37,"D3":35.97,"D4":39.72,"D5":91.26,"IC50":498.8,"Slope":0.53,"max":91.26,"min":3.47,"Barcode":"H1FO4B-N2278","W5":"P5","W4":"O5","W3":"N5","W2":"M5","W1":"L5"},{"CellLine":"AsPC-1","Drug":"AZD_8330","minConc":1,"D1":19.41,"D2":37.67,"D3":43.23,"D4":50.25,"D5":40.34,"IC50":1.96,"Slope":0.53,"max":50.25,"min":0,"Barcode":"H1FO4B-N2278","W5":"F6","W4":"H6","W3":"I6","W2":"J6","W1":"K6"},{"CellLine":"AsPC-1","Drug":"Rigosertib","minConc":1,"D1":-5.24,"D2":-0.96,"D3":54.08,"D4":63.41,"D5":65.35,"IC50":53.87,"Slope":2.5,"max":65.35,"min":0,"Barcode":"H1FO4B-N2278","W5":"F8","W4":"G8","W3":"H8","W2":"J8","W1":"K8"},{"CellLine":"AsPC-1","Drug":"Omipalisib","minConc":0.1,"D1":6.8,"D2":5.93,"D3":9.09,"D4":58.01,"D5":71.49,"IC50":48.8,"Slope":1.88,"max":71.49,"min":5.93,"Barcode":"H1FO4B-N2278","W5":"P17","W4":"O17","W3":"N17","W2":"M17","W1":"K17"},{"CellLine":"AsPC-1","Drug":"UCN_01","minConc":1,"D1":2.17,"D2":-1.92,"D3":47.3,"D4":57.3,"D5":92.42,"IC50":213.06,"Slope":0.67,"max":92.42,"min":0,"Barcode":"H1FO4B-N2278","W5":"P21","W4":"O21","W3":"N21","W2":"M21","W1":"L21"},{"CellLine":"AsPC-1","Drug":"Dinaciclib","minConc":0.1,"D1":4.75,"D2":-0.06,"D3":14.72,"D4":74.78,"D5":77.01,"IC50":19.54,"Slope":2.15,"max":77.01,"min":0,"Barcode":"H1FO4B-N3278","W5":"A3","W4":"B3","W3":"C3","W2":"D3","W1":"E3"},{"CellLine":"AsPC-1","Drug":"NVP_BGT226","minConc":0.1,"D1":8.68,"D2":10.08,"D3":33.41,"D4":52.28,"D5":90.17,"IC50":48.94,"Slope":0.72,"max":90.17,"min":7.89,"Barcode":"H1FO4B-N3278","W5":"G12","W4":"H12","W3":"I12","W2":"J12","W1":"K12"},{"CellLine":"AsPC-1","Drug":"Gedatolisib","minConc":0.1,"D1":4.14,"D2":5.9,"D3":6.62,"D4":49.73,"D5":77.42,"IC50":74.67,"Slope":1.72,"max":77.42,"min":4.14,"Barcode":"H1FO4B-N3278","W5":"P12","W4":"O12","W3":"N12","W2":"M12","W1":"L12"},{"CellLine":"AsPC-1","Drug":"PKI_402","minConc":0.25,"D1":8.69,"D2":1.75,"D3":-3.03,"D4":41.72,"D5":73.39,"IC50":224.16,"Slope":2.5,"max":73.39,"min":0,"Barcode":"H1FO4B-N3278","W5":"A16","W4":"C16","W3":"D16","W2":"E16","W1":"F16"},{"CellLine":"AsPC-1","Drug":"AT7519","minConc":1,"D1":7.65,"D2":6.68,"D3":10.93,"D4":72.41,"D5":70.26,"IC50":265,"Slope":2.5,"max":72.41,"min":6.68,"Barcode":"H1FO4B-N3278","W5":"F22","W4":"G22","W3":"H22","W2":"I22","W1":"J22"},{"CellLine":"AsPC-1","Drug":"GSK_461364","minConc":1,"D1":4.34,"D2":20.43,"D3":43.26,"D4":42.87,"D5":88.55,"IC50":114.36,"Slope":0.54,"max":88.55,"min":4.34,"Barcode":"H1FO4B-N4278","W5":"A6","W4":"B6","W3":"C6","W2":"D6","W1":"E6"},{"CellLine":"AsPC-1","Drug":"Daporinad","minConc":0.1,"D1":10.52,"D2":-4.2,"D3":74.8,"D4":91.01,"D5":87.45,"IC50":5.5,"Slope":2.5,"max":91.01,"min":0,"Barcode":"H1FO4B-N4278","W5":"A14","W4":"B14","W3":"C14","W2":"D14","W1":"E14"},{"CellLine":"AsPC-1","Drug":"Mepacrine","minConc":5,"D1":-1.09,"D2":-6.82,"D3":-4.58,"D4":-8.64,"D5":-11.45,"IC50":50000,"Slope":1,"max":0,"min":0,"Barcode":"H1FO4B-N4278","W5":"P20","W4":"O20","W3":"N20","W2":"M20","W1":"L20"},{"CellLine":"AsPC-1","Drug":"Camptothecin","minConc":0.5,"D1":2.83,"D2":5.85,"D3":19.5,"D4":51.39,"D5":78.6,"IC50":228.68,"Slope":0.9,"max":78.6,"min":2.83,"Barcode":"H1FO4B-N5278","W5":"O3","W4":"N3","W3":"M3","W2":"L3","W1":"K3"},{"CellLine":"AsPC-1","Drug":"Omacetaxine","minConc":1,"D1":4.65,"D2":-2.27,"D3":-4.72,"D4":70.74,"D5":79.84,"IC50":465.22,"Slope":2.5,"max":79.84,"min":0,"Barcode":"H1FO4B-N5278","W5":"A5","W4":"B5","W3":"C5","W2":"E5","W1":"F5"},{"CellLine":"AsPC-1","Drug":"Vinorelbine","minConc":1,"D1":-6.34,"D2":36.57,"D3":61.24,"D4":76.11,"D5":99.8,"IC50":51.56,"Slope":0.58,"max":99.8,"min":0,"Barcode":"H1FO4B-N5278","W5":"P14","W4":"O14","W3":"N14","W2":"M14","W1":"L14"},{"CellLine":"AsPC-1","Drug":"Docetaxel","minConc":0.1,"D1":4.93,"D2":3.19,"D3":39.93,"D4":51.37,"D5":54.4,"IC50":6.82,"Slope":2.42,"max":54.4,"min":3.19,"Barcode":"H1FO4B-N5278","W5":"P17","W4":"O17","W3":"N17","W2":"M17","W1":"K17"},{"CellLine":"AsPC-1","Drug":"Carfilzomib","minConc":0.1,"D1":4.28,"D2":8.59,"D3":77.49,"D4":93.22,"D5":99.07,"IC50":5,"Slope":1.75,"max":99.07,"min":3.71,"Barcode":"H1FO4B-N5278","W5":"A18","W4":"B18","W3":"C18","W2":"D18","W1":"E18"},{"CellLine":"AsPC-1","Drug":"Patupilone","minConc":0.1,"D1":5.3,"D2":14.87,"D3":48.39,"D4":51.27,"D5":50.53,"IC50":2.12,"Slope":1.75,"max":51.27,"min":5.08,"Barcode":"H1FO4B-N5278","W5":"F18","W4":"G18","W3":"H18","W2":"I18","W1":"J18"},{"CellLine":"AsPC-1","Drug":"Plicamycin","minConc":1,"D1":0.8,"D2":-2.15,"D3":-1.63,"D4":80.01,"D5":83.51,"IC50":362.57,"Slope":2.5,"max":83.51,"min":0,"Barcode":"H1FO4B-N5278","W5":"P18","W4":"N18","W3":"M18","W2":"L18","W1":"K18"},{"CellLine":"AsPC-1","Drug":"Vincristine","minConc":0.1,"D1":4.26,"D2":2.52,"D3":47.76,"D4":73.26,"D5":76,"IC50":7.99,"Slope":2.07,"max":76,"min":2.52,"Barcode":"H1FO4B-N5278","W5":"A19","W4":"B19","W3":"C19","W2":"D19","W1":"E19"},{"CellLine":"AsPC-1","Drug":"Dactinomycin","minConc":0.1,"D1":-1.82,"D2":9.82,"D3":14.47,"D4":89.09,"D5":79.34,"IC50":19.15,"Slope":2.5,"max":89.09,"min":0,"Barcode":"H1FO4B-N5278","W5":"P19","W4":"O19","W3":"N19","W2":"M19","W1":"L19"},{"CellLine":"AsPC-1","Drug":"Vinblastine","minConc":0.1,"D1":4.08,"D2":-2.45,"D3":28.61,"D4":61.91,"D5":72.81,"IC50":16.42,"Slope":1.1,"max":72.81,"min":0,"Barcode":"H1FO4B-N5278","W5":"A21","W4":"B21","W3":"C21","W2":"D21","W1":"E21"},{"CellLine":"AsPC-1","Drug":"Paclitaxel","minConc":0.1,"D1":5.71,"D2":6.96,"D3":31.71,"D4":47.06,"D5":52.13,"IC50":8.2,"Slope":1.08,"max":52.13,"min":4.13,"Barcode":"H1FO4B-N5278","W5":"F21","W4":"G21","W3":"H21","W2":"I21","W1":"J21"},{"CellLine":"AsPC-1","Drug":"Oprozomib","minConc":0.25,"D1":1.72,"D2":-3.31,"D3":50.67,"D4":76.33,"D5":93.22,"IC50":26.52,"Slope":1.07,"max":93.22,"min":0,"Barcode":"H1FO4B-N5278","W5":"A22","W4":"B22","W3":"C22","W2":"D22","W1":"E22"},{"CellLine":"AsPC-1","Drug":"Bortezomib","minConc":0.1,"D1":-2.22,"D2":-0.56,"D3":72.24,"D4":89.01,"D5":95.32,"IC50":6.35,"Slope":2.5,"max":95.32,"min":0,"Barcode":"H1FO4B-N5278","W5":"F22","W4":"G22","W3":"H22","W2":"I22","W1":"J22"},{"CellLine":"AsPC-1","Drug":"Filanesib","minConc":0.1,"D1":-1.77,"D2":1.62,"D3":32.54,"D4":36.87,"D5":34.85,"IC50":4.03,"Slope":2.22,"max":36.87,"min":0,"Barcode":"H1FO4B-N5278","W5":"G23","W4":"H23","W3":"I23","W2":"J23","W1":"K23"},{"CellLine":"AsPC-1","Drug":"CUDC_907","minConc":1,"D1":-0.15,"D2":14.05,"D3":79.63,"D4":101.8,"D5":101.12,"IC50":37.15,"Slope":1.4,"max":100,"min":0,"Barcode":"H1FO4B-N6278","W5":"F17","W4":"G17","W3":"H17","W2":"I17","W1":"J17"},{"CellLine":"AsPC-1","Drug":"YM155","minConc":1,"D1":9.4,"D2":11.61,"D3":100.68,"D4":101.55,"D5":103.38,"IC50":27.75,"Slope":2.5,"max":100,"min":7.31,"Barcode":"H1FO4B-N6278","W5":"F18","W4":"G18","W3":"H18","W2":"I18","W1":"J18"},{"CellLine":"AsPC-1","Drug":"AT101","minConc":10,"D1":1.89,"D2":-2.24,"D3":1.55,"D4":86.74,"D5":104.63,"IC50":4727.4,"Slope":2.5,"max":100,"min":0,"Barcode":"H1FO4B-N6278","W5":"A21","W4":"B21","W3":"C21","W2":"D21","W1":"E21"},{"CellLine":"AsPC-1","Drug":"Selinexor","minConc":1,"D1":3.97,"D2":-1.01,"D3":17.72,"D4":67.54,"D5":80.72,"IC50":274.17,"Slope":1.28,"max":80.72,"min":0,"Barcode":"H1FO4B-N6278","W5":"P21","W4":"O21","W3":"N21","W2":"M21","W1":"L21"},{"CellLine":"AsPC-1","Drug":"BIIB021","minConc":1,"D1":9.09,"D2":7.73,"D3":38.01,"D4":79.48,"D5":87.31,"IC50":149.53,"Slope":1.24,"max":87.31,"min":7.14,"Barcode":"H1FO4B-N7278","W5":"P13","W4":"N13","W3":"M13","W2":"L13","W1":"K13"},{"CellLine":"AsPC-1","Drug":"Tanespimycin","minConc":1,"D1":5.57,"D2":13.01,"D3":74.04,"D4":86.45,"D5":89.93,"IC50":40.65,"Slope":1.61,"max":89.93,"min":5.19,"Barcode":"H1FO4B-N7278","W5":"A14","W4":"B14","W3":"C14","W2":"D14","W1":"E14"},{"CellLine":"AsPC-1","Drug":"CUDC_305","minConc":1,"D1":14.29,"D2":6.13,"D3":11.05,"D4":76.44,"D5":74.96,"IC50":260.19,"Slope":2.5,"max":76.44,"min":6.13,"Barcode":"H1FO4B-N7278","W5":"A15","W4":"B15","W3":"C15","W2":"D15","W1":"F15"},{"CellLine":"AsPC-1","Drug":"Luminespib","minConc":0.1,"D1":1.17,"D2":8.41,"D3":20.36,"D4":66,"D5":78.1,"IC50":24.37,"Slope":1.14,"max":78.1,"min":1.17,"Barcode":"H1FO4B-N7278","W5":"G15","W4":"H15","W3":"I15","W2":"J15","W1":"K15"},{"CellLine":"AsPC-1","Drug":"Ganetespib","minConc":0.1,"D1":7.91,"D2":6.12,"D3":6.69,"D4":70.83,"D5":78.84,"IC50":44.03,"Slope":2.5,"max":78.84,"min":6.12,"Barcode":"H1FO4B-N7278","W5":"A16","W4":"C16","W3":"D16","W2":"E16","W1":"F16"},{"CellLine":"AsPC-1","Drug":"Stattic","minConc":5,"D1":7.48,"D2":8.11,"D3":5.41,"D4":100.97,"D5":103.93,"IC50":1549.06,"Slope":2.5,"max":100,"min":5.41,"Barcode":"H1FO4B-N7278","W5":"F21","W4":"G21","W3":"H21","W2":"I21","W1":"J21"},{"CellLine":"AsPC-1","Drug":"ONX_0914","minConc":1,"D1":-1.55,"D2":0.22,"D3":11.74,"D4":90.69,"D5":99.3,"IC50":288.65,"Slope":1.9,"max":99.3,"min":0,"Barcode":"H1FO4B-N7278","W5":"G23","W4":"H23","W3":"I23","W2":"J23","W1":"K23"},{"CellLine":"AsPC-1","Drug":"SN_38","minConc":1,"D1":13.91,"D2":22.7,"D3":36,"D4":84.43,"D5":81.04,"IC50":150.05,"Slope":1.85,"max":84.43,"min":13.91,"Barcode":"H1FO4B-N8278","W5":"G2","W4":"H2","W3":"I2","W2":"J2","W1":"K2"},{"CellLine":"AsPC-1","Drug":"Cabazitaxel","minConc":0.1,"D1":1.41,"D2":8.8,"D3":47.47,"D4":53.61,"D5":50.03,"IC50":2.94,"Slope":1.65,"max":53.61,"min":1.21,"Barcode":"H1FO4B-N8278","W5":"A3","W4":"B3","W3":"C3","W2":"D3","W1":"E3"},{"CellLine":"AsPC-1","Drug":"OTSSP167","minConc":0.1,"D1":4.83,"D2":7.3,"D3":-5.1,"D4":0.55,"D5":74.91,"IC50":311.52,"Slope":2.5,"max":74.91,"min":0,"Barcode":"H1FO4B-N8278","W5":"A5","W4":"B5","W3":"C5","W2":"E5","W1":"F5"},{"CellLine":"AsPC-1","Drug":"Prexasertib","minConc":1,"D1":6.06,"D2":1.93,"D3":69.02,"D4":64.75,"D5":30.38,"IC50":31.62,"Slope":2.5,"max":69.02,"min":1.93,"Barcode":"H1FO4B-N8278","W5":"P6","W4":"O6","W3":"N6","W2":"M6","W1":"L6"},{"CellLine":"AsPC-1","Drug":"Romidepsin","minConc":0.1,"D1":9,"D2":24.38,"D3":98.02,"D4":100.4,"D5":99.85,"IC50":1.97,"Slope":2.34,"max":100,"min":8.92,"Barcode":"H1FO4B-N8278","W5":"F18","W4":"G18","W3":"H18","W2":"I18","W1":"J18"},{"CellLine":"AsPC-1","Drug":"Sabutoclax","minConc":2.5,"D1":8.25,"D2":12.22,"D3":16.24,"D4":46.78,"D5":97.26,"IC50":2897.41,"Slope":1.44,"max":97.26,"min":8.25,"Barcode":"H1FO4B-N8278","W5":"F19","W4":"G19","W3":"I19","W2":"J19","W1":"K19"},{"CellLine":"AsPC-1","Drug":"BCI","minConc":5,"D1":4.25,"D2":-5.67,"D3":-2.73,"D4":101.2,"D5":100.82,"IC50":1623.96,"Slope":2.5,"max":100,"min":0,"Barcode":"H1FO4B-N8278","W5":"A22","W4":"B22","W3":"C22","W2":"D22","W1":"E22"},{"CellLine":"HPAC","Drug":"PF_04691502","minConc":1,"D1":15.63,"D2":1.29,"D3":15.54,"D4":83.36,"D5":96.65,"IC50":307.83,"Slope":1.55,"max":96.65,"min":1.29,"Barcode":"H1FO4B-N1257","W5":"B2","W4":"C2","W3":"D2","W2":"E2","W1":"F2"},{"CellLine":"HPAC","Drug":"SNS_032","minConc":1,"D1":-10.15,"D2":11.61,"D3":25.12,"D4":83.26,"D5":83.18,"IC50":152.81,"Slope":1.9,"max":83.26,"min":0,"Barcode":"H1FO4B-N1257","W5":"G9","W4":"H9","W3":"I9","W2":"J9","W1":"K9"},{"CellLine":"HPAC","Drug":"Alvocidib","minConc":1,"D1":-4,"D2":-10.58,"D3":8.9,"D4":86.81,"D5":87.61,"IC50":234.96,"Slope":2.5,"max":87.61,"min":0,"Barcode":"H1FO4B-N1257","W5":"P9","W4":"O9","W3":"N9","W2":"M9","W1":"L9"},{"CellLine":"HPAC","Drug":"KX2_391","minConc":1,"D1":5.99,"D2":13.64,"D3":15.71,"D4":19.65,"D5":29.29,"IC50":106.43,"Slope":0.64,"max":29.29,"min":5.99,"Barcode":"H1FO4B-N1257","W5":"F17","W4":"G17","W3":"H17","W2":"I17","W1":"J17"},{"CellLine":"HPAC","Drug":"AZD8055","minConc":1,"D1":-13.62,"D2":-3.51,"D3":13.34,"D4":38.87,"D5":51.63,"IC50":316.91,"Slope":1.08,"max":51.63,"min":0,"Barcode":"H1FO4B-N1257","W5":"G23","W4":"H23","W3":"I23","W2":"J23","W1":"K23"},{"CellLine":"HPAC","Drug":"PF_03758309","minConc":1,"D1":-2.68,"D2":29.08,"D3":43.99,"D4":70.78,"D5":88.49,"IC50":76.83,"Slope":0.58,"max":88.49,"min":0,"Barcode":"H1FO4B-N2257","W5":"P5","W4":"O5","W3":"N5","W2":"M5","W1":"L5"},{"CellLine":"HPAC","Drug":"AZD_8330","minConc":1,"D1":9.45,"D2":30.98,"D3":40.35,"D4":55.04,"D5":48.08,"IC50":9.71,"Slope":0.57,"max":55.04,"min":0,"Barcode":"H1FO4B-N2257","W5":"F6","W4":"H6","W3":"I6","W2":"J6","W1":"K6"},{"CellLine":"HPAC","Drug":"Rigosertib","minConc":1,"D1":3.47,"D2":7.3,"D3":24.68,"D4":24.6,"D5":32.16,"IC50":36.92,"Slope":0.66,"max":32.16,"min":0,"Barcode":"H1FO4B-N2257","W5":"F8","W4":"G8","W3":"H8","W2":"J8","W1":"K8"},{"CellLine":"HPAC","Drug":"Omipalisib","minConc":0.1,"D1":-1.55,"D2":12.54,"D3":17.98,"D4":71.82,"D5":76.69,"IC50":20.58,"Slope":1.52,"max":76.69,"min":0,"Barcode":"H1FO4B-N2257","W5":"P17","W4":"O17","W3":"N17","W2":"M17","W1":"K17"},{"CellLine":"HPAC","Drug":"UCN_01","minConc":1,"D1":-0.73,"D2":7.5,"D3":31.96,"D4":63.17,"D5":87.43,"IC50":226.88,"Slope":0.76,"max":87.43,"min":0,"Barcode":"H1FO4B-N2257","W5":"P21","W4":"O21","W3":"N21","W2":"M21","W1":"L21"},{"CellLine":"HPAC","Drug":"Dinaciclib","minConc":0.1,"D1":10.32,"D2":2.27,"D3":4.83,"D4":86.62,"D5":84.18,"IC50":29.44,"Slope":2.5,"max":86.62,"min":2.27,"Barcode":"H1FO4B-N3257","W5":"A3","W4":"B3","W3":"C3","W2":"D3","W1":"E3"},{"CellLine":"HPAC","Drug":"NVP_BGT226","minConc":0.1,"D1":5.06,"D2":4.7,"D3":6.67,"D4":45.69,"D5":68.19,"IC50":71.49,"Slope":1.8,"max":68.19,"min":4.7,"Barcode":"H1FO4B-N3257","W5":"G12","W4":"H12","W3":"I12","W2":"J12","W1":"K12"},{"CellLine":"HPAC","Drug":"Gedatolisib","minConc":0.1,"D1":-7.11,"D2":1.88,"D3":-7.05,"D4":45.68,"D5":79.11,"IC50":88.5,"Slope":2.5,"max":79.11,"min":0,"Barcode":"H1FO4B-N3257","W5":"P12","W4":"O12","W3":"N12","W2":"M12","W1":"L12"},{"CellLine":"HPAC","Drug":"PKI_402","minConc":0.25,"D1":5.78,"D2":2.88,"D3":-5.17,"D4":46.03,"D5":72.54,"IC50":201.1,"Slope":2.5,"max":72.54,"min":0,"Barcode":"H1FO4B-N3257","W5":"A16","W4":"C16","W3":"D16","W2":"E16","W1":"F16"},{"CellLine":"HPAC","Drug":"AT7519","minConc":1,"D1":1.95,"D2":-9.87,"D3":-0.87,"D4":78.34,"D5":82.9,"IC50":371.92,"Slope":2.5,"max":82.9,"min":0,"Barcode":"H1FO4B-N3257","W5":"F22","W4":"G22","W3":"H22","W2":"I22","W1":"J22"},{"CellLine":"HPAC","Drug":"GSK_461364","minConc":1,"D1":10.26,"D2":1.71,"D3":10.37,"D4":7.71,"D5":70.88,"IC50":2447.78,"Slope":2.5,"max":70.88,"min":1.71,"Barcode":"H1FO4B-N4257","W5":"A6","W4":"B6","W3":"C6","W2":"D6","W1":"E6"},{"CellLine":"HPAC","Drug":"Daporinad","minConc":0.1,"D1":3.63,"D2":-9.79,"D3":-11.7,"D4":-19.72,"D5":-12.16,"IC50":1000,"Slope":2.5,"max":3.63,"min":0,"Barcode":"H1FO4B-N4257","W5":"A14","W4":"B14","W3":"C14","W2":"D14","W1":"E14"},{"CellLine":"HPAC","Drug":"Mepacrine","minConc":5,"D1":0.95,"D2":-8.23,"D3":-10.01,"D4":53.06,"D5":101.77,"IC50":4770.46,"Slope":2.5,"max":100,"min":0,"Barcode":"H1FO4B-N4257","W5":"P20","W4":"O20","W3":"N20","W2":"M20","W1":"L20"},{"CellLine":"HPAC","Drug":"Camptothecin","minConc":0.5,"D1":-7.17,"D2":-2.49,"D3":9.02,"D4":12.3,"D5":70.91,"IC50":928.52,"Slope":2.5,"max":70.91,"min":0,"Barcode":"H1FO4B-N5257","W5":"O3","W4":"N3","W3":"M3","W2":"L3","W1":"K3"},{"CellLine":"HPAC","Drug":"Omacetaxine","minConc":1,"D1":-3.55,"D2":4.75,"D3":43.31,"D4":85.69,"D5":85.53,"IC50":97.21,"Slope":1.55,"max":85.69,"min":0,"Barcode":"H1FO4B-N5257","W5":"A5","W4":"B5","W3":"C5","W2":"E5","W1":"F5"},{"CellLine":"HPAC","Drug":"Vinorelbine","minConc":1,"D1":-16.94,"D2":1.78,"D3":3.76,"D4":16.71,"D5":75.07,"IC50":1645.48,"Slope":2.5,"max":75.07,"min":0,"Barcode":"H1FO4B-N5257","W5":"P14","W4":"O14","W3":"N14","W2":"M14","W1":"L14"},{"CellLine":"HPAC","Drug":"Docetaxel","minConc":0.1,"D1":10.8,"D2":-2.08,"D3":68.34,"D4":11.29,"D5":1.04,"IC50":1000,"Slope":0.14,"max":68.34,"min":0,"Barcode":"H1FO4B-N5257","W5":"P17","W4":"O17","W3":"N17","W2":"M17","W1":"K17"},{"CellLine":"HPAC","Drug":"Carfilzomib","minConc":0.1,"D1":5.49,"D2":27.95,"D3":55.93,"D4":94.2,"D5":97.57,"IC50":5.64,"Slope":0.77,"max":97.57,"min":3.92,"Barcode":"H1FO4B-N5257","W5":"A18","W4":"B18","W3":"C18","W2":"D18","W1":"E18"},{"CellLine":"HPAC","Drug":"Patupilone","minConc":0.1,"D1":2.37,"D2":6.86,"D3":6.09,"D4":17.69,"D5":19.43,"IC50":22.68,"Slope":1.2,"max":19.43,"min":2.37,"Barcode":"H1FO4B-N5257","W5":"F18","W4":"G18","W3":"H18","W2":"I18","W1":"J18"},{"CellLine":"HPAC","Drug":"Plicamycin","minConc":1,"D1":8.32,"D2":-7.04,"D3":16.66,"D4":92.92,"D5":89.78,"IC50":183.26,"Slope":2.5,"max":92.92,"min":0,"Barcode":"H1FO4B-N5257","W5":"P18","W4":"N18","W3":"M18","W2":"L18","W1":"K18"},{"CellLine":"HPAC","Drug":"Vincristine","minConc":0.1,"D1":-1.02,"D2":76.62,"D3":7.33,"D4":15.23,"D5":10.88,"IC50":1000,"Slope":0.1,"max":76.62,"min":0,"Barcode":"H1FO4B-N5257","W5":"A19","W4":"B19","W3":"C19","W2":"D19","W1":"E19"},{"CellLine":"HPAC","Drug":"Dactinomycin","minConc":0.1,"D1":3.36,"D2":20.21,"D3":72.86,"D4":94.61,"D5":89.71,"IC50":3.46,"Slope":1.14,"max":94.61,"min":1.85,"Barcode":"H1FO4B-N5257","W5":"P19","W4":"O19","W3":"N19","W2":"M19","W1":"L19"},{"CellLine":"HPAC","Drug":"Vinblastine","minConc":0.1,"D1":-5.49,"D2":10.42,"D3":0.33,"D4":13.84,"D5":14.64,"IC50":42.39,"Slope":0.54,"max":18.23,"min":0,"Barcode":"H1FO4B-N5257","W5":"A21","W4":"B21","W3":"C21","W2":"D21","W1":"E21"},{"CellLine":"HPAC","Drug":"Paclitaxel","minConc":0.1,"D1":5.94,"D2":10.27,"D3":7.55,"D4":20.58,"D5":20.21,"IC50":22.45,"Slope":2.5,"max":20.58,"min":5.94,"Barcode":"H1FO4B-N5257","W5":"F21","W4":"G21","W3":"H21","W2":"I21","W1":"J21"},{"CellLine":"HPAC","Drug":"Oprozomib","minConc":0.25,"D1":-7.92,"D2":2.75,"D3":-12.94,"D4":88.14,"D5":95.14,"IC50":109.25,"Slope":2.5,"max":95.14,"min":0,"Barcode":"H1FO4B-N5257","W5":"A22","W4":"B22","W3":"C22","W2":"D22","W1":"E22"},{"CellLine":"HPAC","Drug":"Bortezomib","minConc":0.1,"D1":-3.54,"D2":7.85,"D3":88.8,"D4":94.38,"D5":98.64,"IC50":3.37,"Slope":2.02,"max":98.64,"min":0,"Barcode":"H1FO4B-N5257","W5":"F22","W4":"G22","W3":"H22","W2":"I22","W1":"J22"},{"CellLine":"HPAC","Drug":"Filanesib","minConc":0.1,"D1":-1.7,"D2":6.66,"D3":2.59,"D4":7.65,"D5":5.03,"IC50":1000,"Slope":0.41,"max":7.65,"min":0,"Barcode":"H1FO4B-N5257","W5":"G23","W4":"H23","W3":"I23","W2":"J23","W1":"K23"},{"CellLine":"HPAC","Drug":"CUDC_907","minConc":1,"D1":16.93,"D2":-1.46,"D3":38.1,"D4":84.97,"D5":94.24,"IC50":142.28,"Slope":1.23,"max":94.24,"min":0,"Barcode":"H1FO4B-N6257","W5":"F17","W4":"G17","W3":"H17","W2":"I17","W1":"J17"},{"CellLine":"HPAC","Drug":"YM155","minConc":1,"D1":2.62,"D2":26.39,"D3":87.85,"D4":100.39,"D5":103.3,"IC50":22.42,"Slope":1.33,"max":100,"min":1.07,"Barcode":"H1FO4B-N6257","W5":"F18","W4":"G18","W3":"H18","W2":"I18","W1":"J18"},{"CellLine":"HPAC","Drug":"AT101","minConc":10,"D1":22.15,"D2":-12.66,"D3":1.31,"D4":38.84,"D5":103.47,"IC50":11973.42,"Slope":2.5,"max":100,"min":0,"Barcode":"H1FO4B-N6257","W5":"A21","W4":"B21","W3":"C21","W2":"D21","W1":"E21"},{"CellLine":"HPAC","Drug":"Selinexor","minConc":1,"D1":-6.52,"D2":2.17,"D3":13.78,"D4":68.07,"D5":70.66,"IC50":201.71,"Slope":2.01,"max":70.66,"min":0,"Barcode":"H1FO4B-N6257","W5":"P21","W4":"O21","W3":"N21","W2":"M21","W1":"L21"},{"CellLine":"HPAC","Drug":"BIIB021","minConc":1,"D1":-6.65,"D2":0.86,"D3":32.45,"D4":77.06,"D5":80.94,"IC50":130.45,"Slope":1.54,"max":80.94,"min":0,"Barcode":"H1FO4B-N7257","W5":"P13","W4":"N13","W3":"M13","W2":"L13","W1":"K13"},{"CellLine":"HPAC","Drug":"Tanespimycin","minConc":1,"D1":8.6,"D2":3.75,"D3":30.37,"D4":75.42,"D5":77.32,"IC50":135.86,"Slope":1.86,"max":77.32,"min":3.75,"Barcode":"H1FO4B-N7257","W5":"A14","W4":"B14","W3":"C14","W2":"D14","W1":"E14"},{"CellLine":"HPAC","Drug":"CUDC_305","minConc":1,"D1":14.36,"D2":9.84,"D3":5.36,"D4":76.19,"D5":74.98,"IC50":316.12,"Slope":2.5,"max":76.19,"min":5.36,"Barcode":"H1FO4B-N7257","W5":"A15","W4":"B15","W3":"C15","W2":"D15","W1":"F15"},{"CellLine":"HPAC","Drug":"Luminespib","minConc":0.1,"D1":7.11,"D2":8.85,"D3":21.23,"D4":-0.25,"D5":51.03,"IC50":12.81,"Slope":0.34,"max":51.03,"min":0,"Barcode":"H1FO4B-N7257","W5":"G15","W4":"H15","W3":"I15","W2":"J15","W1":"K15"},{"CellLine":"HPAC","Drug":"Ganetespib","minConc":0.1,"D1":14.73,"D2":11.73,"D3":6.75,"D4":-5.41,"D5":61.48,"IC50":359.29,"Slope":2.5,"max":61.48,"min":0,"Barcode":"H1FO4B-N7257","W5":"A16","W4":"C16","W3":"D16","W2":"E16","W1":"F16"},{"CellLine":"HPAC","Drug":"Stattic","minConc":5,"D1":11.56,"D2":0.82,"D3":17.12,"D4":20.12,"D5":103.49,"IC50":8724.97,"Slope":2.5,"max":100,"min":0.82,"Barcode":"H1FO4B-N7257","W5":"F21","W4":"G21","W3":"H21","W2":"I21","W1":"J21"},{"CellLine":"HPAC","Drug":"ONX_0914","minConc":1,"D1":53.91,"D2":72.3,"D3":7.87,"D4":95.58,"D5":99.34,"IC50":355.92,"Slope":2.5,"max":100,"min":7.87,"Barcode":"H1FO4B-N7257","W5":"G23","W4":"H23","W3":"I23","W2":"J23","W1":"K23"},{"CellLine":"HPAC","Drug":"SN_38","minConc":1,"D1":4.36,"D2":26.38,"D3":31.42,"D4":71.87,"D5":86.82,"IC50":154.06,"Slope":0.67,"max":86.82,"min":4.36,"Barcode":"H1FO4B-N8257","W5":"G2","W4":"H2","W3":"I2","W2":"J2","W1":"K2"},{"CellLine":"HPAC","Drug":"Cabazitaxel","minConc":0.1,"D1":8.56,"D2":1.53,"D3":12.03,"D4":11.27,"D5":0.76,"IC50":1000,"Slope":-0.09,"max":12.03,"min":0,"Barcode":"H1FO4B-N8257","W5":"A3","W4":"B3","W3":"C3","W2":"D3","W1":"E3"},{"CellLine":"HPAC","Drug":"OTSSP167","minConc":0.1,"D1":14.16,"D2":-1.23,"D3":-1.77,"D4":7.43,"D5":89.58,"IC50":248.57,"Slope":2.5,"max":89.58,"min":0,"Barcode":"H1FO4B-N8257","W5":"A5","W4":"B5","W3":"C5","W2":"E5","W1":"F5"},{"CellLine":"HPAC","Drug":"Prexasertib","minConc":1,"D1":14.76,"D2":11.2,"D3":69.24,"D4":72.9,"D5":41.39,"IC50":37.06,"Slope":2.5,"max":72.9,"min":11.2,"Barcode":"H1FO4B-N8257","W5":"P6","W4":"O6","W3":"N6","W2":"M6","W1":"L6"},{"CellLine":"HPAC","Drug":"Romidepsin","minConc":0.1,"D1":4.32,"D2":-0.46,"D3":75.77,"D4":89.17,"D5":90.15,"IC50":5.18,"Slope":2.5,"max":90.15,"min":0,"Barcode":"H1FO4B-N8257","W5":"F18","W4":"G18","W3":"H18","W2":"I18","W1":"J18"},{"CellLine":"HPAC","Drug":"Sabutoclax","minConc":2.5,"D1":11.89,"D2":5.61,"D3":19.27,"D4":43.22,"D5":90.51,"IC50":2587.04,"Slope":1.07,"max":90.51,"min":5.61,"Barcode":"H1FO4B-N8257","W5":"F19","W4":"G19","W3":"I19","W2":"J19","W1":"K19"},{"CellLine":"HPAC","Drug":"BCI","minConc":5,"D1":6.41,"D2":2.15,"D3":-13.13,"D4":68.27,"D5":100.33,"IC50":3708.16,"Slope":2.5,"max":100,"min":0,"Barcode":"H1FO4B-N8257","W5":"A22","W4":"B22","W3":"C22","W2":"D22","W1":"E22"},{"CellLine":"MIA-PaCa-2","Drug":"PF_04691502","minConc":1,"D1":2.42,"D2":-1.42,"D3":5.35,"D4":55.85,"D5":78.04,"IC50":550.76,"Slope":1.56,"max":78.04,"min":0,"Barcode":"H1FO4B-N1211","W5":"B2","W4":"C2","W3":"D2","W2":"E2","W1":"F2"},{"CellLine":"MIA-PaCa-2","Drug":"SNS_032","minConc":1,"D1":6.63,"D2":15.72,"D3":83.84,"D4":88.29,"D5":86.59,"IC50":26.32,"Slope":2.14,"max":88.29,"min":6.56,"Barcode":"H1FO4B-N1211","W5":"G9","W4":"H9","W3":"I9","W2":"J9","W1":"K9"},{"CellLine":"MIA-PaCa-2","Drug":"Alvocidib","minConc":1,"D1":1.76,"D2":2.43,"D3":12.36,"D4":90.46,"D5":94.53,"IC50":250.84,"Slope":2.23,"max":94.53,"min":1.76,"Barcode":"H1FO4B-N1211","W5":"P9","W4":"O9","W3":"N9","W2":"M9","W1":"L9"},{"CellLine":"MIA-PaCa-2","Drug":"KX2_391","minConc":1,"D1":9.09,"D2":11.4,"D3":47.6,"D4":64.42,"D5":62.04,"IC50":61.95,"Slope":1.73,"max":64.42,"min":9.09,"Barcode":"H1FO4B-N1211","W5":"F17","W4":"G17","W3":"H17","W2":"I17","W1":"J17"},{"CellLine":"MIA-PaCa-2","Drug":"AZD8055","minConc":1,"D1":9.31,"D2":6,"D3":26.42,"D4":68.48,"D5":66.96,"IC50":133.45,"Slope":2.5,"max":68.48,"min":6,"Barcode":"H1FO4B-N1211","W5":"G23","W4":"H23","W3":"I23","W2":"J23","W1":"K23"},{"CellLine":"MIA-PaCa-2","Drug":"PF_03758309","minConc":1,"D1":4.99,"D2":8.4,"D3":37.18,"D4":63.91,"D5":96.88,"IC50":279.21,"Slope":0.7,"max":96.88,"min":2.48,"Barcode":"H1FO4B-N2211","W5":"P5","W4":"O5","W3":"N5","W2":"M5","W1":"L5"},{"CellLine":"MIA-PaCa-2","Drug":"AZD_8330","minConc":1,"D1":5.05,"D2":21.69,"D3":38.75,"D4":63.23,"D5":65.96,"IC50":46.4,"Slope":0.7,"max":65.96,"min":2.47,"Barcode":"H1FO4B-N2211","W5":"F6","W4":"H6","W3":"I6","W2":"J6","W1":"K6"},{"CellLine":"MIA-PaCa-2","Drug":"Rigosertib","minConc":1,"D1":10.03,"D2":-2.56,"D3":35.84,"D4":65.79,"D5":66.14,"IC50":93.61,"Slope":2.5,"max":66.14,"min":0,"Barcode":"H1FO4B-N2211","W5":"F8","W4":"G8","W3":"H8","W2":"J8","W1":"K8"},{"CellLine":"MIA-PaCa-2","Drug":"Omipalisib","minConc":0.1,"D1":1.16,"D2":1.48,"D3":16.32,"D4":34.92,"D5":58.86,"IC50":47.2,"Slope":0.8,"max":58.86,"min":0.52,"Barcode":"H1FO4B-N2211","W5":"P17","W4":"O17","W3":"N17","W2":"M17","W1":"K17"},{"CellLine":"MIA-PaCa-2","Drug":"UCN_01","minConc":1,"D1":3.71,"D2":3.62,"D3":6.32,"D4":33.87,"D5":85.17,"IC50":1268.67,"Slope":2.2,"max":85.17,"min":3.62,"Barcode":"H1FO4B-N2211","W5":"P21","W4":"O21","W3":"N21","W2":"M21","W1":"L21"},{"CellLine":"MIA-PaCa-2","Drug":"Dinaciclib","minConc":0.1,"D1":1.58,"D2":4.78,"D3":41.69,"D4":85.75,"D5":86.94,"IC50":10.71,"Slope":1.61,"max":86.94,"min":1.58,"Barcode":"H1FO4B-N3211","W5":"A3","W4":"B3","W3":"C3","W2":"D3","W1":"E3"},{"CellLine":"MIA-PaCa-2","Drug":"NVP_BGT226","minConc":0.1,"D1":2.87,"D2":6.59,"D3":24.16,"D4":58.63,"D5":97.7,"IC50":53.09,"Slope":0.86,"max":97.7,"min":2.87,"Barcode":"H1FO4B-N3211","W5":"G12","W4":"H12","W3":"I12","W2":"J12","W1":"K12"},{"CellLine":"MIA-PaCa-2","Drug":"Gedatolisib","minConc":0.1,"D1":3.78,"D2":-3.39,"D3":10.19,"D4":46.82,"D5":71.94,"IC50":55.55,"Slope":1.15,"max":71.94,"min":0,"Barcode":"H1FO4B-N3211","W5":"P12","W4":"O12","W3":"N12","W2":"M12","W1":"L12"},{"CellLine":"MIA-PaCa-2","Drug":"PKI_402","minConc":0.25,"D1":9.45,"D2":7.28,"D3":4.08,"D4":49.28,"D5":66.32,"IC50":169.33,"Slope":2.5,"max":66.32,"min":4.08,"Barcode":"H1FO4B-N3211","W5":"A16","W4":"C16","W3":"D16","W2":"E16","W1":"F16"},{"CellLine":"MIA-PaCa-2","Drug":"AT7519","minConc":1,"D1":4.65,"D2":1.02,"D3":5.93,"D4":85.93,"D5":90.87,"IC50":316.65,"Slope":2.47,"max":90.87,"min":1.02,"Barcode":"H1FO4B-N3211","W5":"F22","W4":"G22","W3":"H22","W2":"I22","W1":"J22"},{"CellLine":"MIA-PaCa-2","Drug":"GSK_461364","minConc":1,"D1":8.27,"D2":22.97,"D3":65.38,"D4":70.22,"D5":50.08,"IC50":20.61,"Slope":1.56,"max":70.22,"min":7.72,"Barcode":"H1FO4B-N4211","W5":"A6","W4":"B6","W3":"C6","W2":"D6","W1":"E6"},{"CellLine":"MIA-PaCa-2","Drug":"Daporinad","minConc":0.1,"D1":10.32,"D2":-4.73,"D3":9.29,"D4":59.86,"D5":68.78,"IC50":31.32,"Slope":1.66,"max":68.78,"min":0,"Barcode":"H1FO4B-N4211","W5":"A14","W4":"B14","W3":"C14","W2":"D14","W1":"E14"},{"CellLine":"MIA-PaCa-2","Drug":"Mepacrine","minConc":5,"D1":-4.91,"D2":-0.78,"D3":3.72,"D4":99.6,"D5":101.76,"IC50":1455.65,"Slope":2.5,"max":100,"min":0,"Barcode":"H1FO4B-N4211","W5":"P20","W4":"O20","W3":"N20","W2":"M20","W1":"L20"},{"CellLine":"MIA-PaCa-2","Drug":"Camptothecin","minConc":0.5,"D1":3.93,"D2":6.67,"D3":26.73,"D4":76.78,"D5":94.14,"IC50":133.8,"Slope":1.1,"max":94.14,"min":3.93,"Barcode":"H1FO4B-N5211","W5":"O3","W4":"N3","W3":"M3","W2":"L3","W1":"K3"},{"CellLine":"MIA-PaCa-2","Drug":"Omacetaxine","minConc":1,"D1":5.85,"D2":11.75,"D3":93.74,"D4":95.54,"D5":99.72,"IC50":31.7,"Slope":2.33,"max":99.72,"min":5.8,"Barcode":"H1FO4B-N5211","W5":"A5","W4":"B5","W3":"C5","W2":"E5","W1":"F5"},{"CellLine":"MIA-PaCa-2","Drug":"Vinorelbine","minConc":1,"D1":6.01,"D2":55.64,"D3":71.66,"D4":71.41,"D5":86.97,"IC50":7.73,"Slope":0.81,"max":86.97,"min":0,"Barcode":"H1FO4B-N5211","W5":"P14","W4":"O14","W3":"N14","W2":"M14","W1":"L14"},{"CellLine":"MIA-PaCa-2","Drug":"Docetaxel","minConc":0.1,"D1":10.56,"D2":23.82,"D3":59.37,"D4":64.89,"D5":53.72,"IC50":2.14,"Slope":1.42,"max":64.89,"min":9.85,"Barcode":"H1FO4B-N5211","W5":"P17","W4":"O17","W3":"N17","W2":"M17","W1":"K17"},{"CellLine":"MIA-PaCa-2","Drug":"Carfilzomib","minConc":0.1,"D1":1.43,"D2":2.67,"D3":11.12,"D4":99.67,"D5":102.72,"IC50":23.6,"Slope":2.5,"max":100,"min":1.43,"Barcode":"H1FO4B-N5211","W5":"A18","W4":"B18","W3":"C18","W2":"D18","W1":"E18"},{"CellLine":"MIA-PaCa-2","Drug":"Patupilone","minConc":0.1,"D1":10.16,"D2":7.8,"D3":43.83,"D4":64.36,"D5":47.55,"IC50":7.99,"Slope":2.5,"max":64.36,"min":7.8,"Barcode":"H1FO4B-N5211","W5":"F18","W4":"G18","W3":"H18","W2":"I18","W1":"J18"},{"CellLine":"MIA-PaCa-2","Drug":"Plicamycin","minConc":1,"D1":4.76,"D2":5.73,"D3":15.88,"D4":85.78,"D5":99.99,"IC50":344.37,"Slope":1.63,"max":99.99,"min":4.76,"Barcode":"H1FO4B-N5211","W5":"P18","W4":"N18","W3":"M18","W2":"L18","W1":"K18"},{"CellLine":"MIA-PaCa-2","Drug":"Vincristine","minConc":0.1,"D1":2.74,"D2":18.1,"D3":63.26,"D4":74.01,"D5":72.93,"IC50":2.58,"Slope":1.3,"max":74.01,"min":1.7,"Barcode":"H1FO4B-N5211","W5":"A19","W4":"B19","W3":"C19","W2":"D19","W1":"E19"},{"CellLine":"MIA-PaCa-2","Drug":"Dactinomycin","minConc":0.1,"D1":4.64,"D2":47.73,"D3":57.64,"D4":99.12,"D5":99.05,"IC50":2.4,"Slope":0.61,"max":99.12,"min":0,"Barcode":"H1FO4B-N5211","W5":"P19","W4":"O19","W3":"N19","W2":"M19","W1":"L19"},{"CellLine":"MIA-PaCa-2","Drug":"Vinblastine","minConc":0.1,"D1":2.91,"D2":7.09,"D3":51.33,"D4":74.05,"D5":73.67,"IC50":6.13,"Slope":1.56,"max":74.05,"min":2.91,"Barcode":"H1FO4B-N5211","W5":"A21","W4":"B21","W3":"C21","W2":"D21","W1":"E21"},{"CellLine":"MIA-PaCa-2","Drug":"Paclitaxel","minConc":0.1,"D1":10.85,"D2":7.87,"D3":40.01,"D4":70.03,"D5":65.44,"IC50":9.73,"Slope":2.5,"max":70.03,"min":7.87,"Barcode":"H1FO4B-N5211","W5":"F21","W4":"G21","W3":"H21","W2":"I21","W1":"J21"},{"CellLine":"MIA-PaCa-2","Drug":"Oprozomib","minConc":0.25,"D1":7.2,"D2":-1.38,"D3":-6.84,"D4":83.87,"D5":102.08,"IC50":132.6,"Slope":2.5,"max":100,"min":0,"Barcode":"H1FO4B-N5211","W5":"A22","W4":"B22","W3":"C22","W2":"D22","W1":"E22"},{"CellLine":"MIA-PaCa-2","Drug":"Bortezomib","minConc":0.1,"D1":-0.8,"D2":-2.14,"D3":76.85,"D4":98.42,"D5":100.55,"IC50":6.21,"Slope":2.5,"max":100,"min":0,"Barcode":"H1FO4B-N5211","W5":"F22","W4":"G22","W3":"H22","W2":"I22","W1":"J22"},{"CellLine":"MIA-PaCa-2","Drug":"Filanesib","minConc":0.1,"D1":3.44,"D2":1.67,"D3":66.58,"D4":66.62,"D5":70.11,"IC50":3.58,"Slope":2.5,"max":70.11,"min":1.28,"Barcode":"H1FO4B-N5211","W5":"G23","W4":"H23","W3":"I23","W2":"J23","W1":"K23"},{"CellLine":"MIA-PaCa-2","Drug":"CUDC_907","minConc":1,"D1":4.62,"D2":8.54,"D3":78.51,"D4":101.7,"D5":101.46,"IC50":52.57,"Slope":1.93,"max":100,"min":4.62,"Barcode":"H1FO4B-N6211","W5":"F17","W4":"G17","W3":"H17","W2":"I17","W1":"J17"},{"CellLine":"MIA-PaCa-2","Drug":"YM155","minConc":1,"D1":7.79,"D2":44.61,"D3":100.25,"D4":100.75,"D5":102.57,"IC50":11.75,"Slope":2.5,"max":100,"min":7.59,"Barcode":"H1FO4B-N6211","W5":"F18","W4":"G18","W3":"H18","W2":"I18","W1":"J18"},{"CellLine":"MIA-PaCa-2","Drug":"AT101","minConc":10,"D1":7.52,"D2":-0.66,"D3":4.05,"D4":78.27,"D5":103,"IC50":5195.37,"Slope":1.97,"max":100,"min":0,"Barcode":"H1FO4B-N6211","W5":"A21","W4":"B21","W3":"C21","W2":"D21","W1":"E21"},{"CellLine":"MIA-PaCa-2","Drug":"Selinexor","minConc":1,"D1":6.26,"D2":5.68,"D3":21.61,"D4":84.51,"D5":96.03,"IC50":278.57,"Slope":1.51,"max":96.03,"min":5.68,"Barcode":"H1FO4B-N6211","W5":"P21","W4":"O21","W3":"N21","W2":"M21","W1":"L21"},{"CellLine":"MIA-PaCa-2","Drug":"BIIB021","minConc":1,"D1":9.36,"D2":8.85,"D3":13.46,"D4":74.58,"D5":83.72,"IC50":380.28,"Slope":2.04,"max":83.72,"min":8.85,"Barcode":"H1FO4B-N7211","W5":"P13","W4":"N13","W3":"M13","W2":"L13","W1":"K13"},{"CellLine":"MIA-PaCa-2","Drug":"Tanespimycin","minConc":1,"D1":8.2,"D2":-4.02,"D3":46.86,"D4":79.35,"D5":71.75,"IC50":86.51,"Slope":2.5,"max":79.35,"min":0,"Barcode":"H1FO4B-N7211","W5":"A14","W4":"B14","W3":"C14","W2":"D14","W1":"E14"},{"CellLine":"MIA-PaCa-2","Drug":"CUDC_305","minConc":1,"D1":11.3,"D2":10.11,"D3":5.49,"D4":74.08,"D5":74.62,"IC50":321.35,"Slope":2.5,"max":74.62,"min":5.49,"Barcode":"H1FO4B-N7211","W5":"A15","W4":"B15","W3":"C15","W2":"D15","W1":"F15"},{"CellLine":"MIA-PaCa-2","Drug":"Luminespib","minConc":0.1,"D1":4.85,"D2":8.68,"D3":8.48,"D4":78.98,"D5":76.32,"IC50":27.88,"Slope":2.5,"max":78.98,"min":4.85,"Barcode":"H1FO4B-N7211","W5":"G15","W4":"H15","W3":"I15","W2":"J15","W1":"K15"},{"CellLine":"MIA-PaCa-2","Drug":"Ganetespib","minConc":0.1,"D1":6.45,"D2":12.24,"D3":6.08,"D4":72.18,"D5":76.64,"IC50":37.55,"Slope":2.5,"max":76.64,"min":6.08,"Barcode":"H1FO4B-N7211","W5":"A16","W4":"C16","W3":"D16","W2":"E16","W1":"F16"},{"CellLine":"MIA-PaCa-2","Drug":"Stattic","minConc":5,"D1":11.48,"D2":8.19,"D3":3.01,"D4":27.56,"D5":103.25,"IC50":7674.88,"Slope":2.5,"max":100,"min":3.01,"Barcode":"H1FO4B-N7211","W5":"F21","W4":"G21","W3":"H21","W2":"I21","W1":"J21"},{"CellLine":"MIA-PaCa-2","Drug":"ONX_0914","minConc":1,"D1":3.41,"D2":6.21,"D3":-2.58,"D4":100.59,"D5":103.01,"IC50":327.49,"Slope":2.5,"max":100,"min":0,"Barcode":"H1FO4B-N7211","W5":"G23","W4":"H23","W3":"I23","W2":"J23","W1":"K23"},{"CellLine":"MIA-PaCa-2","Drug":"SN_38","minConc":1,"D1":4.66,"D2":24.71,"D3":68.61,"D4":92.44,"D5":90.84,"IC50":32.15,"Slope":0.96,"max":92.44,"min":1.9,"Barcode":"H1FO4B-N8211","W5":"G2","W4":"H2","W3":"I2","W2":"J2","W1":"K2"},{"CellLine":"MIA-PaCa-2","Drug":"Cabazitaxel","minConc":0.1,"D1":4,"D2":20.99,"D3":53.44,"D4":65.09,"D5":39.41,"IC50":2.14,"Slope":0.91,"max":65.09,"min":0,"Barcode":"H1FO4B-N8211","W5":"A3","W4":"B3","W3":"C3","W2":"D3","W1":"E3"},{"CellLine":"MIA-PaCa-2","Drug":"OTSSP167","minConc":0.1,"D1":6.86,"D2":9.79,"D3":3.35,"D4":97.08,"D5":97.89,"IC50":32.19,"Slope":2.5,"max":97.89,"min":3.35,"Barcode":"H1FO4B-N8211","W5":"A5","W4":"B5","W3":"C5","W2":"E5","W1":"F5"},{"CellLine":"MIA-PaCa-2","Drug":"Prexasertib","minConc":1,"D1":7.88,"D2":12.56,"D3":58.62,"D4":66.68,"D5":71.83,"IC50":43.76,"Slope":1.6,"max":71.83,"min":7.39,"Barcode":"H1FO4B-N8211","W5":"P6","W4":"O6","W3":"N6","W2":"M6","W1":"L6"},{"CellLine":"MIA-PaCa-2","Drug":"Romidepsin","minConc":0.1,"D1":9.93,"D2":6.85,"D3":99.62,"D4":101.84,"D5":102.07,"IC50":3.13,"Slope":2.5,"max":100,"min":6.06,"Barcode":"H1FO4B-N8211","W5":"F18","W4":"G18","W3":"H18","W2":"I18","W1":"J18"},{"CellLine":"MIA-PaCa-2","Drug":"Sabutoclax","minConc":2.5,"D1":10.29,"D2":3.55,"D3":61.74,"D4":78.85,"D5":102.16,"IC50":220.58,"Slope":0.9,"max":100,"min":1.89,"Barcode":"H1FO4B-N8211","W5":"F19","W4":"G19","W3":"I19","W2":"J19","W1":"K19"},{"CellLine":"MIA-PaCa-2","Drug":"BCI","minConc":5,"D1":1.23,"D2":-2.35,"D3":-5.84,"D4":96.91,"D5":100.54,"IC50":1855.32,"Slope":2.5,"max":100,"min":0,"Barcode":"H1FO4B-N8211","W5":"A22","W4":"B22","W3":"C22","W2":"D22","W1":"E22"},{"CellLine":"SW1990","Drug":"PF_04691502","minConc":1,"D1":2.72,"D2":5.59,"D3":23.48,"D4":76.41,"D5":98.85,"IC50":327.76,"Slope":1.09,"max":98.85,"min":2.72,"Barcode":"H1FO4B-N1322","W5":"B2","W4":"C2","W3":"D2","W2":"E2","W1":"F2"},{"CellLine":"SW1990","Drug":"SNS_032","minConc":1,"D1":2.81,"D2":3.27,"D3":62.69,"D4":69.58,"D5":73.81,"IC50":51.01,"Slope":2.5,"max":73.81,"min":2.45,"Barcode":"H1FO4B-N1322","W5":"G9","W4":"H9","W3":"I9","W2":"J9","W1":"K9"},{"CellLine":"SW1990","Drug":"Alvocidib","minConc":1,"D1":7.4,"D2":0.3,"D3":17.85,"D4":71.63,"D5":74.8,"IC50":188,"Slope":1.87,"max":74.8,"min":0.3,"Barcode":"H1FO4B-N1322","W5":"P9","W4":"O9","W3":"N9","W2":"M9","W1":"L9"},{"CellLine":"SW1990","Drug":"KX2_391","minConc":1,"D1":1.2,"D2":8.2,"D3":28.68,"D4":48.56,"D5":61.87,"IC50":131.2,"Slope":0.72,"max":61.87,"min":0,"Barcode":"H1FO4B-N1322","W5":"F17","W4":"G17","W3":"H17","W2":"I17","W1":"J17"},{"CellLine":"SW1990","Drug":"AZD8055","minConc":1,"D1":9.67,"D2":18.2,"D3":45.33,"D4":59.35,"D5":63.37,"IC50":47.42,"Slope":0.93,"max":63.37,"min":7.98,"Barcode":"H1FO4B-N1322","W5":"G23","W4":"H23","W3":"I23","W2":"J23","W1":"K23"},{"CellLine":"SW1990","Drug":"PF_03758309","minConc":1,"D1":-0.72,"D2":8.47,"D3":26.62,"D4":50.86,"D5":99.09,"IC50":570.88,"Slope":0.72,"max":99.09,"min":0,"Barcode":"H1FO4B-N2322","W5":"P5","W4":"O5","W3":"N5","W2":"M5","W1":"L5"},{"CellLine":"SW1990","Drug":"AZD_8330","minConc":1,"D1":1.74,"D2":7.13,"D3":29.73,"D4":53.66,"D5":60.55,"IC50":107.96,"Slope":0.93,"max":60.55,"min":1.07,"Barcode":"H1FO4B-N2322","W5":"F6","W4":"H6","W3":"I6","W2":"J6","W1":"K6"},{"CellLine":"SW1990","Drug":"Rigosertib","minConc":1,"D1":-3.83,"D2":-3.35,"D3":25.15,"D4":55.47,"D5":55.36,"IC50":107.86,"Slope":2.5,"max":55.47,"min":0,"Barcode":"H1FO4B-N2322","W5":"F8","W4":"G8","W3":"H8","W2":"J8","W1":"K8"},{"CellLine":"SW1990","Drug":"Omipalisib","minConc":0.1,"D1":5.29,"D2":11.63,"D3":28.2,"D4":64.86,"D5":81.26,"IC50":23.9,"Slope":0.91,"max":81.26,"min":5.29,"Barcode":"H1FO4B-N2322","W5":"P17","W4":"O17","W3":"N17","W2":"M17","W1":"K17"},{"CellLine":"SW1990","Drug":"UCN_01","minConc":1,"D1":10.79,"D2":18.69,"D3":56.2,"D4":91.38,"D5":93.07,"IC50":80.94,"Slope":1.19,"max":93.07,"min":10.79,"Barcode":"H1FO4B-N2322","W5":"P21","W4":"O21","W3":"N21","W2":"M21","W1":"L21"},{"CellLine":"SW1990","Drug":"Dinaciclib","minConc":0.1,"D1":5.73,"D2":-1.75,"D3":25.31,"D4":75.68,"D5":80.11,"IC50":16.18,"Slope":1.63,"max":80.11,"min":0,"Barcode":"H1FO4B-N3322","W5":"A3","W4":"B3","W3":"C3","W2":"D3","W1":"E3"},{"CellLine":"SW1990","Drug":"NVP_BGT226","minConc":0.1,"D1":4.25,"D2":6.54,"D3":30.39,"D4":61.25,"D5":98.55,"IC50":42.59,"Slope":0.78,"max":98.55,"min":3.5,"Barcode":"H1FO4B-N3322","W5":"G12","W4":"H12","W3":"I12","W2":"J12","W1":"K12"},{"CellLine":"SW1990","Drug":"Gedatolisib","minConc":0.1,"D1":-0.58,"D2":1.53,"D3":18.8,"D4":62.25,"D5":85.09,"IC50":35.67,"Slope":1.03,"max":85.09,"min":0,"Barcode":"H1FO4B-N3322","W5":"P12","W4":"O12","W3":"N12","W2":"M12","W1":"L12"},{"CellLine":"SW1990","Drug":"PKI_402","minConc":0.25,"D1":5.56,"D2":-1.33,"D3":5.95,"D4":57.89,"D5":75.24,"IC50":117.53,"Slope":1.6,"max":75.24,"min":0,"Barcode":"H1FO4B-N3322","W5":"A16","W4":"C16","W3":"D16","W2":"E16","W1":"F16"},{"CellLine":"SW1990","Drug":"AT7519","minConc":1,"D1":0.12,"D2":2.45,"D3":0.97,"D4":79.35,"D5":72.87,"IC50":308.08,"Slope":2.5,"max":79.35,"min":0,"Barcode":"H1FO4B-N3322","W5":"F22","W4":"G22","W3":"H22","W2":"I22","W1":"J22"},{"CellLine":"SW1990","Drug":"GSK_461364","minConc":1,"D1":-3.72,"D2":1.28,"D3":31.59,"D4":36.94,"D5":47.87,"IC50":71.49,"Slope":1.1,"max":47.87,"min":0,"Barcode":"H1FO4B-N4322","W5":"A6","W4":"B6","W3":"C6","W2":"D6","W1":"E6"},{"CellLine":"SW1990","Drug":"Daporinad","minConc":0.1,"D1":-3.49,"D2":0.82,"D3":11.93,"D4":82.57,"D5":84.48,"IC50":21.12,"Slope":2.41,"max":84.48,"min":0,"Barcode":"H1FO4B-N4322","W5":"A14","W4":"B14","W3":"C14","W2":"D14","W1":"E14"},{"CellLine":"SW1990","Drug":"Mepacrine","minConc":5,"D1":1.03,"D2":1.69,"D3":4.49,"D4":71.43,"D5":101.93,"IC50":3092.94,"Slope":1.89,"max":100,"min":1.03,"Barcode":"H1FO4B-N4322","W5":"P20","W4":"O20","W3":"N20","W2":"M20","W1":"L20"},{"CellLine":"SW1990","Drug":"Camptothecin","minConc":0.5,"D1":7.53,"D2":8.58,"D3":45.83,"D4":75.39,"D5":88.29,"IC50":57.15,"Slope":0.92,"max":88.29,"min":4.06,"Barcode":"H1FO4B-N5322","W5":"O3","W4":"N3","W3":"M3","W2":"L3","W1":"K3"},{"CellLine":"SW1990","Drug":"Omacetaxine","minConc":1,"D1":8.89,"D2":-5.01,"D3":33.03,"D4":76.17,"D5":89.27,"IC50":175.12,"Slope":1.15,"max":89.27,"min":0,"Barcode":"H1FO4B-N5322","W5":"A5","W4":"B5","W3":"C5","W2":"E5","W1":"F5"},{"CellLine":"SW1990","Drug":"Vinorelbine","minConc":1,"D1":0.97,"D2":28.14,"D3":55.87,"D4":64.43,"D5":83.54,"IC50":41.75,"Slope":0.59,"max":83.54,"min":0,"Barcode":"H1FO4B-N5322","W5":"P14","W4":"O14","W3":"N14","W2":"M14","W1":"L14"},{"CellLine":"SW1990","Drug":"Docetaxel","minConc":0.1,"D1":8.97,"D2":21.27,"D3":35.16,"D4":45.88,"D5":39.05,"IC50":1.4,"Slope":0.61,"max":45.88,"min":1.42,"Barcode":"H1FO4B-N5322","W5":"P17","W4":"O17","W3":"N17","W2":"M17","W1":"K17"},{"CellLine":"SW1990","Drug":"Carfilzomib","minConc":0.1,"D1":4.32,"D2":3.72,"D3":101.39,"D4":100.41,"D5":103.28,"IC50":2.89,"Slope":2.5,"max":100,"min":1,"Barcode":"H1FO4B-N5322","W5":"A18","W4":"B18","W3":"C18","W2":"D18","W1":"E18"},{"CellLine":"SW1990","Drug":"Patupilone","minConc":0.1,"D1":2.32,"D2":2.97,"D3":17.91,"D4":39.78,"D5":39.98,"IC50":11.8,"Slope":2.09,"max":39.98,"min":2.32,"Barcode":"H1FO4B-N5322","W5":"F18","W4":"G18","W3":"H18","W2":"I18","W1":"J18"},{"CellLine":"SW1990","Drug":"Plicamycin","minConc":1,"D1":-0.27,"D2":4.45,"D3":70.82,"D4":81.31,"D5":85.5,"IC50":44.33,"Slope":1.92,"max":85.5,"min":0,"Barcode":"H1FO4B-N5322","W5":"P18","W4":"N18","W3":"M18","W2":"L18","W1":"K18"},{"CellLine":"SW1990","Drug":"Vincristine","minConc":0.1,"D1":3.04,"D2":-0.64,"D3":39.3,"D4":57.17,"D5":61.84,"IC50":7.68,"Slope":2.06,"max":61.84,"min":0,"Barcode":"H1FO4B-N5322","W5":"A19","W4":"B19","W3":"C19","W2":"D19","W1":"E19"},{"CellLine":"SW1990","Drug":"Dactinomycin","minConc":0.1,"D1":8.35,"D2":34.1,"D3":41.09,"D4":88.93,"D5":82.62,"IC50":7.76,"Slope":0.69,"max":88.93,"min":8.35,"Barcode":"H1FO4B-N5322","W5":"P19","W4":"O19","W3":"N19","W2":"M19","W1":"L19"},{"CellLine":"SW1990","Drug":"Vinblastine","minConc":0.1,"D1":6.72,"D2":9.58,"D3":32.2,"D4":54.85,"D5":58.62,"IC50":10.24,"Slope":1.14,"max":58.62,"min":6.33,"Barcode":"H1FO4B-N5322","W5":"A21","W4":"B21","W3":"C21","W2":"D21","W1":"E21"},{"CellLine":"SW1990","Drug":"Paclitaxel","minConc":0.1,"D1":9.08,"D2":7.63,"D3":21.88,"D4":43.11,"D5":47.76,"IC50":16.92,"Slope":1.2,"max":47.76,"min":7.63,"Barcode":"H1FO4B-N5322","W5":"F21","W4":"G21","W3":"H21","W2":"I21","W1":"J21"},{"CellLine":"SW1990","Drug":"Oprozomib","minConc":0.25,"D1":4.65,"D2":5.07,"D3":22.73,"D4":100.04,"D5":103.58,"IC50":44.56,"Slope":2.5,"max":100,"min":4.65,"Barcode":"H1FO4B-N5322","W5":"A22","W4":"B22","W3":"C22","W2":"D22","W1":"E22"},{"CellLine":"SW1990","Drug":"Bortezomib","minConc":0.1,"D1":-0.65,"D2":9.84,"D3":102.22,"D4":100.18,"D5":100.74,"IC50":2.31,"Slope":2.5,"max":100,"min":0,"Barcode":"H1FO4B-N5322","W5":"F22","W4":"G22","W3":"H22","W2":"I22","W1":"J22"},{"CellLine":"SW1990","Drug":"Filanesib","minConc":0.1,"D1":3.73,"D2":0.09,"D3":31.24,"D4":35.3,"D5":34.27,"IC50":4.51,"Slope":2.5,"max":35.3,"min":0.09,"Barcode":"H1FO4B-N5322","W5":"G23","W4":"H23","W3":"I23","W2":"J23","W1":"K23"},{"CellLine":"SW1990","Drug":"CUDC_907","minConc":1,"D1":2.85,"D2":11.67,"D3":59.82,"D4":95.03,"D5":97.75,"IC50":70.15,"Slope":1.2,"max":97.75,"min":2.7,"Barcode":"H1FO4B-N6322","W5":"F17","W4":"G17","W3":"H17","W2":"I17","W1":"J17"},{"CellLine":"SW1990","Drug":"YM155","minConc":1,"D1":7.79,"D2":18.1,"D3":88.26,"D4":100.49,"D5":102.31,"IC50":32.78,"Slope":1.73,"max":100,"min":7.59,"Barcode":"H1FO4B-N6322","W5":"F18","W4":"G18","W3":"H18","W2":"I18","W1":"J18"},{"CellLine":"SW1990","Drug":"AT101","minConc":10,"D1":8.08,"D2":4.44,"D3":0.63,"D4":53.04,"D5":103.79,"IC50":9563.62,"Slope":2.5,"max":100,"min":0.63,"Barcode":"H1FO4B-N6322","W5":"A21","W4":"B21","W3":"C21","W2":"D21","W1":"E21"},{"CellLine":"SW1990","Drug":"Selinexor","minConc":1,"D1":11.01,"D2":11.4,"D3":49.67,"D4":80.81,"D5":92.44,"IC50":112.21,"Slope":0.98,"max":92.44,"min":7.69,"Barcode":"H1FO4B-N6322","W5":"P21","W4":"O21","W3":"N21","W2":"M21","W1":"L21"},{"CellLine":"SW1990","Drug":"BIIB021","minConc":1,"D1":6.86,"D2":10.53,"D3":19.23,"D4":82.69,"D5":95.94,"IC50":323.09,"Slope":1.54,"max":95.94,"min":6.86,"Barcode":"H1FO4B-N7322","W5":"P13","W4":"N13","W3":"M13","W2":"L13","W1":"K13"},{"CellLine":"SW1990","Drug":"Tanespimycin","minConc":1,"D1":1.06,"D2":3.21,"D3":70.58,"D4":91.91,"D5":95.9,"IC50":58.82,"Slope":1.9,"max":95.9,"min":0.55,"Barcode":"H1FO4B-N7322","W5":"A14","W4":"B14","W3":"C14","W2":"D14","W1":"E14"},{"CellLine":"SW1990","Drug":"CUDC_305","minConc":1,"D1":-0.5,"D2":-5.03,"D3":0.15,"D4":75.41,"D5":88.9,"IC50":505.57,"Slope":2.5,"max":88.9,"min":0,"Barcode":"H1FO4B-N7322","W5":"A15","W4":"B15","W3":"C15","W2":"D15","W1":"F15"},{"CellLine":"SW1990","Drug":"Luminespib","minConc":0.1,"D1":5.61,"D2":3.05,"D3":28.11,"D4":89.95,"D5":87.38,"IC50":14.35,"Slope":2.5,"max":89.95,"min":3.05,"Barcode":"H1FO4B-N7322","W5":"G15","W4":"H15","W3":"I15","W2":"J15","W1":"K15"},{"CellLine":"SW1990","Drug":"Ganetespib","minConc":0.1,"D1":2.02,"D2":-5.44,"D3":2.46,"D4":87.21,"D5":90.6,"IC50":32.59,"Slope":2.5,"max":90.6,"min":0,"Barcode":"H1FO4B-N7322","W5":"A16","W4":"C16","W3":"D16","W2":"E16","W1":"F16"},{"CellLine":"SW1990","Drug":"Stattic","minConc":5,"D1":9.29,"D2":3.46,"D3":3.15,"D4":13.32,"D5":103.72,"IC50":11168.5,"Slope":2.5,"max":100,"min":3.15,"Barcode":"H1FO4B-N7322","W5":"F21","W4":"G21","W3":"H21","W2":"I21","W1":"J21"},{"CellLine":"SW1990","Drug":"ONX_0914","minConc":1,"D1":1.5,"D2":12.79,"D3":62.35,"D4":102.54,"D5":103.47,"IC50":64.19,"Slope":1.24,"max":100,"min":1.5,"Barcode":"H1FO4B-N7322","W5":"G23","W4":"H23","W3":"I23","W2":"J23","W1":"K23"},{"CellLine":"SW1990","Drug":"SN_38","minConc":1,"D1":6.16,"D2":43.42,"D3":60.49,"D4":81.12,"D5":79.59,"IC50":12.17,"Slope":0.72,"max":81.12,"min":0,"Barcode":"H1FO4B-N8322","W5":"G2","W4":"H2","W3":"I2","W2":"J2","W1":"K2"},{"CellLine":"SW1990","Drug":"Cabazitaxel","minConc":0.1,"D1":-4.25,"D2":9.56,"D3":31.02,"D4":42.94,"D5":32.33,"IC50":3.83,"Slope":1.06,"max":42.94,"min":0,"Barcode":"H1FO4B-N8322","W5":"A3","W4":"B3","W3":"C3","W2":"D3","W1":"E3"},{"CellLine":"SW1990","Drug":"OTSSP167","minConc":0.1,"D1":1.47,"D2":3.43,"D3":-2.15,"D4":77.35,"D5":75.09,"IC50":33.25,"Slope":2.5,"max":77.35,"min":0,"Barcode":"H1FO4B-N8322","W5":"A5","W4":"B5","W3":"C5","W2":"E5","W1":"F5"},{"CellLine":"SW1990","Drug":"Prexasertib","minConc":1,"D1":-2.98,"D2":16.34,"D3":43.45,"D4":38.04,"D5":55.01,"IC50":35.31,"Slope":0.7,"max":55.01,"min":0,"Barcode":"H1FO4B-N8322","W5":"P6","W4":"O6","W3":"N6","W2":"M6","W1":"L6"},{"CellLine":"SW1990","Drug":"Romidepsin","minConc":0.1,"D1":3.67,"D2":25.99,"D3":90.6,"D4":95.9,"D5":97.03,"IC50":2,"Slope":1.62,"max":97.03,"min":2.93,"Barcode":"H1FO4B-N8322","W5":"F18","W4":"G18","W3":"H18","W2":"I18","W1":"J18"},{"CellLine":"SW1990","Drug":"Sabutoclax","minConc":2.5,"D1":4.3,"D2":6.36,"D3":20.56,"D4":52.99,"D5":91.11,"IC50":1609.07,"Slope":0.93,"max":91.11,"min":4.3,"Barcode":"H1FO4B-N8322","W5":"F19","W4":"G19","W3":"I19","W2":"J19","W1":"K19"},{"CellLine":"SW1990","Drug":"BCI","minConc":5,"D1":5.64,"D2":4.23,"D3":-9.24,"D4":96.26,"D5":101.11,"IC50":1948.32,"Slope":2.5,"max":100,"min":0,"Barcode":"H1FO4B-N8322","W5":"A22","W4":"B22","W3":"C22","W2":"D22","W1":"E22"},{"CellLine":"HPAF-II","Drug":"PF_04691502","minConc":1,"D1":-4.72,"D2":-3.75,"D3":18.56,"D4":56.44,"D5":89.04,"IC50":502.75,"Slope":0.98,"max":89.04,"min":0,"Barcode":"H1FO4B-N1337","W5":"B2","W4":"C2","W3":"D2","W2":"E2","W1":"F2"},{"CellLine":"HPAF-II","Drug":"SNS_032","minConc":1,"D1":1.48,"D2":-4.64,"D3":36.77,"D4":63.47,"D5":65.23,"IC50":90.48,"Slope":2.5,"max":65.23,"min":0,"Barcode":"H1FO4B-N1337","W5":"G9","W4":"H9","W3":"I9","W2":"J9","W1":"K9"},{"CellLine":"HPAF-II","Drug":"Alvocidib","minConc":1,"D1":-3.51,"D2":7.61,"D3":16.08,"D4":64.35,"D5":68.81,"IC50":205.79,"Slope":1.58,"max":68.81,"min":0,"Barcode":"H1FO4B-N1337","W5":"P9","W4":"O9","W3":"N9","W2":"M9","W1":"L9"},{"CellLine":"HPAF-II","Drug":"KX2_391","minConc":1,"D1":4.95,"D2":4.19,"D3":41,"D4":46.14,"D5":53.57,"IC50":60.76,"Slope":2.12,"max":53.57,"min":4.06,"Barcode":"H1FO4B-N1337","W5":"F17","W4":"G17","W3":"H17","W2":"I17","W1":"J17"},{"CellLine":"HPAF-II","Drug":"AZD8055","minConc":1,"D1":8.76,"D2":2.47,"D3":28.36,"D4":50.6,"D5":57.68,"IC50":122.95,"Slope":1.07,"max":57.68,"min":2.47,"Barcode":"H1FO4B-N1337","W5":"G23","W4":"H23","W3":"I23","W2":"J23","W1":"K23"},{"CellLine":"HPAF-II","Drug":"PF_03758309","minConc":1,"D1":7.79,"D2":17.7,"D3":40.11,"D4":65.15,"D5":85.77,"IC50":160.16,"Slope":0.67,"max":85.77,"min":6.04,"Barcode":"H1FO4B-N2337","W5":"P5","W4":"O5","W3":"N5","W2":"M5","W1":"L5"},{"CellLine":"HPAF-II","Drug":"AZD_8330","minConc":1,"D1":8.73,"D2":23.33,"D3":16.86,"D4":39.65,"D5":43.53,"IC50":126.23,"Slope":0.59,"max":43.53,"min":8.73,"Barcode":"H1FO4B-N2337","W5":"F6","W4":"H6","W3":"I6","W2":"J6","W1":"K6"},{"CellLine":"HPAF-II","Drug":"Rigosertib","minConc":1,"D1":8.62,"D2":4.91,"D3":49.85,"D4":45.07,"D5":43.59,"IC50":31.63,"Slope":2.5,"max":49.85,"min":4.91,"Barcode":"H1FO4B-N2337","W5":"F8","W4":"G8","W3":"H8","W2":"J8","W1":"K8"},{"CellLine":"HPAF-II","Drug":"Omipalisib","minConc":0.1,"D1":10.36,"D2":16.03,"D3":38.2,"D4":53.96,"D5":58.68,"IC50":7,"Slope":0.9,"max":58.68,"min":9.11,"Barcode":"H1FO4B-N2337","W5":"P17","W4":"O17","W3":"N17","W2":"M17","W1":"K17"},{"CellLine":"HPAF-II","Drug":"UCN_01","minConc":1,"D1":7.25,"D2":11.06,"D3":32.34,"D4":54,"D5":79.15,"IC50":275.73,"Slope":0.7,"max":79.15,"min":5.73,"Barcode":"H1FO4B-N2337","W5":"P21","W4":"O21","W3":"N21","W2":"M21","W1":"L21"},{"CellLine":"HPAF-II","Drug":"Dinaciclib","minConc":0.1,"D1":2.73,"D2":4.68,"D3":19.21,"D4":62.27,"D5":65.36,"IC50":18.24,"Slope":1.7,"max":65.36,"min":2.73,"Barcode":"H1FO4B-N3337","W5":"A3","W4":"B3","W3":"C3","W2":"D3","W1":"E3"},{"CellLine":"HPAF-II","Drug":"NVP_BGT226","minConc":0.1,"D1":0.36,"D2":1,"D3":14.61,"D4":42.38,"D5":81.5,"IC50":76.89,"Slope":0.97,"max":81.5,"min":0.36,"Barcode":"H1FO4B-N3337","W5":"G12","W4":"H12","W3":"I12","W2":"J12","W1":"K12"},{"CellLine":"HPAF-II","Drug":"Gedatolisib","minConc":0.1,"D1":-0.2,"D2":8.45,"D3":17.81,"D4":55.47,"D5":67.46,"IC50":24.02,"Slope":1.01,"max":67.46,"min":0,"Barcode":"H1FO4B-N3337","W5":"P12","W4":"O12","W3":"N12","W2":"M12","W1":"L12"},{"CellLine":"HPAF-II","Drug":"PKI_402","minConc":0.25,"D1":-2.49,"D2":-4.54,"D3":-1.58,"D4":43.8,"D5":64.75,"IC50":186.47,"Slope":2.5,"max":64.75,"min":0,"Barcode":"H1FO4B-N3337","W5":"A16","W4":"C16","W3":"D16","W2":"E16","W1":"F16"},{"CellLine":"HPAF-II","Drug":"AT7519","minConc":1,"D1":-4.11,"D2":-0.31,"D3":8.56,"D4":66.46,"D5":63.31,"IC50":212.19,"Slope":2.5,"max":66.46,"min":0,"Barcode":"H1FO4B-N3337","W5":"F22","W4":"G22","W3":"H22","W2":"I22","W1":"J22"},{"CellLine":"HPAF-II","Drug":"GSK_461364","minConc":1,"D1":1.96,"D2":26.08,"D3":34.91,"D4":21.54,"D5":46.02,"IC50":29.88,"Slope":0.35,"max":46.02,"min":0,"Barcode":"H1FO4B-N4337","W5":"A6","W4":"B6","W3":"C6","W2":"D6","W1":"E6"},{"CellLine":"HPAF-II","Drug":"Daporinad","minConc":0.1,"D1":5.5,"D2":11.29,"D3":55.28,"D4":77.93,"D5":72.18,"IC50":5.7,"Slope":1.42,"max":77.93,"min":5.42,"Barcode":"H1FO4B-N4337","W5":"A14","W4":"B14","W3":"C14","W2":"D14","W1":"E14"},{"CellLine":"HPAF-II","Drug":"Mepacrine","minConc":5,"D1":-9.04,"D2":-0.14,"D3":8.76,"D4":69.57,"D5":92.41,"IC50":2341.06,"Slope":1.48,"max":92.41,"min":0,"Barcode":"H1FO4B-N4337","W5":"P20","W4":"O20","W3":"N20","W2":"M20","W1":"L20"},{"CellLine":"HPAF-II","Drug":"Camptothecin","minConc":0.5,"D1":4.21,"D2":3.1,"D3":7.13,"D4":52.31,"D5":71.81,"IC50":281.86,"Slope":1.62,"max":71.81,"min":3.1,"Barcode":"H1FO4B-N5337","W5":"O3","W4":"N3","W3":"M3","W2":"L3","W1":"K3"},{"CellLine":"HPAF-II","Drug":"Omacetaxine","minConc":1,"D1":3.15,"D2":-8.78,"D3":24.04,"D4":67.21,"D5":75.66,"IC50":184.05,"Slope":1.41,"max":75.66,"min":0,"Barcode":"H1FO4B-N5337","W5":"A5","W4":"B5","W3":"C5","W2":"E5","W1":"F5"},{"CellLine":"HPAF-II","Drug":"Vinorelbine","minConc":1,"D1":-7.29,"D2":30.84,"D3":51.32,"D4":51.16,"D5":81.04,"IC50":66.18,"Slope":0.51,"max":81.04,"min":0,"Barcode":"H1FO4B-N5337","W5":"P14","W4":"O14","W3":"N14","W2":"M14","W1":"L14"},{"CellLine":"HPAF-II","Drug":"Docetaxel","minConc":0.1,"D1":1,"D2":12.55,"D3":47.29,"D4":45.21,"D5":46.54,"IC50":1.55,"Slope":2.5,"max":47.29,"min":0.94,"Barcode":"H1FO4B-N5337","W5":"P17","W4":"O17","W3":"N17","W2":"M17","W1":"K17"},{"CellLine":"HPAF-II","Drug":"Carfilzomib","minConc":0.1,"D1":-3.63,"D2":-2.97,"D3":68.09,"D4":88.54,"D5":95.12,"IC50":6.94,"Slope":2.5,"max":95.12,"min":0,"Barcode":"H1FO4B-N5337","W5":"A18","W4":"B18","W3":"C18","W2":"D18","W1":"E18"},{"CellLine":"HPAF-II","Drug":"Patupilone","minConc":0.1,"D1":-4.08,"D2":-0.32,"D3":35.02,"D4":43.67,"D5":39.06,"IC50":5.74,"Slope":2.5,"max":43.67,"min":0,"Barcode":"H1FO4B-N5337","W5":"F18","W4":"G18","W3":"H18","W2":"I18","W1":"J18"},{"CellLine":"HPAF-II","Drug":"Plicamycin","minConc":1,"D1":-4.12,"D2":-6.25,"D3":57.06,"D4":73.93,"D5":69.28,"IC50":62.18,"Slope":2.5,"max":73.93,"min":0,"Barcode":"H1FO4B-N5337","W5":"P18","W4":"N18","W3":"M18","W2":"L18","W1":"K18"},{"CellLine":"HPAF-II","Drug":"Vincristine","minConc":0.1,"D1":1.63,"D2":2.46,"D3":43.55,"D4":49.06,"D5":46.43,"IC50":4.45,"Slope":2.5,"max":49.06,"min":1.49,"Barcode":"H1FO4B-N5337","W5":"A19","W4":"B19","W3":"C19","W2":"D19","W1":"E19"},{"CellLine":"HPAF-II","Drug":"Dactinomycin","minConc":0.1,"D1":-1.28,"D2":19.7,"D3":40.31,"D4":76.76,"D5":67.55,"IC50":6.27,"Slope":0.78,"max":76.76,"min":0,"Barcode":"H1FO4B-N5337","W5":"P19","W4":"O19","W3":"N19","W2":"M19","W1":"L19"},{"CellLine":"HPAF-II","Drug":"Vinblastine","minConc":0.1,"D1":2.37,"D2":-3.19,"D3":28.67,"D4":46.16,"D5":43.13,"IC50":8.23,"Slope":2.5,"max":46.16,"min":0,"Barcode":"H1FO4B-N5337","W5":"A21","W4":"B21","W3":"C21","W2":"D21","W1":"E21"},{"CellLine":"HPAF-II","Drug":"Paclitaxel","minConc":0.1,"D1":5.99,"D2":10.24,"D3":39.81,"D4":43.23,"D5":44.15,"IC50":3.17,"Slope":1.78,"max":44.15,"min":5.89,"Barcode":"H1FO4B-N5337","W5":"F21","W4":"G21","W3":"H21","W2":"I21","W1":"J21"},{"CellLine":"HPAF-II","Drug":"Oprozomib","minConc":0.25,"D1":-3.6,"D2":-3.02,"D3":-7.47,"D4":63.7,"D5":86.08,"IC50":165.98,"Slope":2.5,"max":86.08,"min":0,"Barcode":"H1FO4B-N5337","W5":"A22","W4":"B22","W3":"C22","W2":"D22","W1":"E22"},{"CellLine":"HPAF-II","Drug":"Bortezomib","minConc":0.1,"D1":1.53,"D2":0.13,"D3":64.16,"D4":78.8,"D5":91.82,"IC50":6.48,"Slope":1.85,"max":91.82,"min":0,"Barcode":"H1FO4B-N5337","W5":"F22","W4":"G22","W3":"H22","W2":"I22","W1":"J22"},{"CellLine":"HPAF-II","Drug":"Filanesib","minConc":0.1,"D1":5.66,"D2":0.14,"D3":34.15,"D4":35.9,"D5":37.47,"IC50":4.11,"Slope":2.5,"max":37.47,"min":0.14,"Barcode":"H1FO4B-N5337","W5":"G23","W4":"H23","W3":"I23","W2":"J23","W1":"K23"},{"CellLine":"HPAF-II","Drug":"CUDC_907","minConc":1,"D1":1.92,"D2":-10.01,"D3":33.13,"D4":78.67,"D5":89.57,"IC50":162.47,"Slope":1.34,"max":89.57,"min":0,"Barcode":"H1FO4B-N6337","W5":"F17","W4":"G17","W3":"H17","W2":"I17","W1":"J17"},{"CellLine":"HPAF-II","Drug":"YM155","minConc":1,"D1":-3.55,"D2":7.52,"D3":89.37,"D4":101.1,"D5":101.91,"IC50":34.81,"Slope":2.02,"max":100,"min":0,"Barcode":"H1FO4B-N6337","W5":"F18","W4":"G18","W3":"H18","W2":"I18","W1":"J18"},{"CellLine":"HPAF-II","Drug":"AT101","minConc":10,"D1":2.31,"D2":-2.37,"D3":9.15,"D4":45.33,"D5":104.22,"IC50":10884.66,"Slope":1.86,"max":100,"min":0,"Barcode":"H1FO4B-N6337","W5":"A21","W4":"B21","W3":"C21","W2":"D21","W1":"E21"},{"CellLine":"HPAF-II","Drug":"Selinexor","minConc":1,"D1":-3.51,"D2":5.4,"D3":15.28,"D4":73.92,"D5":79.69,"IC50":230.27,"Slope":1.7,"max":79.69,"min":0,"Barcode":"H1FO4B-N6337","W5":"P21","W4":"O21","W3":"N21","W2":"M21","W1":"L21"},{"CellLine":"HPAF-II","Drug":"BIIB021","minConc":1,"D1":8.34,"D2":8.99,"D3":27.17,"D4":66.68,"D5":71.59,"IC50":179.9,"Slope":1.45,"max":71.59,"min":8.18,"Barcode":"H1FO4B-N7337","W5":"P13","W4":"N13","W3":"M13","W2":"L13","W1":"K13"},{"CellLine":"HPAF-II","Drug":"Tanespimycin","minConc":1,"D1":1.69,"D2":-5.82,"D3":32.97,"D4":66.49,"D5":64.67,"IC50":100.85,"Slope":2.5,"max":66.49,"min":0,"Barcode":"H1FO4B-N7337","W5":"A14","W4":"B14","W3":"C14","W2":"D14","W1":"E14"},{"CellLine":"HPAF-II","Drug":"CUDC_305","minConc":1,"D1":3.68,"D2":-6.42,"D3":-2.53,"D4":51.53,"D5":55.59,"IC50":409.23,"Slope":2.5,"max":55.59,"min":0,"Barcode":"H1FO4B-N7337","W5":"A15","W4":"B15","W3":"C15","W2":"D15","W1":"F15"},{"CellLine":"HPAF-II","Drug":"Luminespib","minConc":0.1,"D1":5.53,"D2":3.86,"D3":5.83,"D4":58.52,"D5":60.31,"IC50":31.33,"Slope":2.5,"max":60.31,"min":3.86,"Barcode":"H1FO4B-N7337","W5":"G15","W4":"H15","W3":"I15","W2":"J15","W1":"K15"},{"CellLine":"HPAF-II","Drug":"Ganetespib","minConc":0.1,"D1":8.93,"D2":-3.28,"D3":-0.3,"D4":60.14,"D5":57.93,"IC50":31.96,"Slope":2.5,"max":60.14,"min":0,"Barcode":"H1FO4B-N7337","W5":"A16","W4":"C16","W3":"D16","W2":"E16","W1":"F16"},{"CellLine":"HPAF-II","Drug":"Stattic","minConc":5,"D1":3.28,"D2":5.89,"D3":-1.4,"D4":31.15,"D5":104.37,"IC50":6847.88,"Slope":2.5,"max":100,"min":0,"Barcode":"H1FO4B-N7337","W5":"F21","W4":"G21","W3":"H21","W2":"I21","W1":"J21"},{"CellLine":"HPAF-II","Drug":"ONX_0914","minConc":1,"D1":6.71,"D2":2.48,"D3":10.83,"D4":86.8,"D5":97.1,"IC50":335.99,"Slope":1.93,"max":97.1,"min":2.48,"Barcode":"H1FO4B-N7337","W5":"G23","W4":"H23","W3":"I23","W2":"J23","W1":"K23"},{"CellLine":"HPAF-II","Drug":"SN_38","minConc":1,"D1":12.09,"D2":18.96,"D3":46.77,"D4":69.12,"D5":72.9,"IC50":73.4,"Slope":1.01,"max":72.9,"min":11.49,"Barcode":"H1FO4B-N8337","W5":"G2","W4":"H2","W3":"I2","W2":"J2","W1":"K2"},{"CellLine":"HPAF-II","Drug":"Cabazitaxel","minConc":0.1,"D1":3.73,"D2":12.52,"D3":48.61,"D4":46.5,"D5":39.82,"IC50":1.75,"Slope":2.5,"max":48.61,"min":3.66,"Barcode":"H1FO4B-N8337","W5":"A3","W4":"B3","W3":"C3","W2":"D3","W1":"E3"},{"CellLine":"HPAF-II","Drug":"OTSSP167","minConc":0.1,"D1":3.21,"D2":0.77,"D3":2.02,"D4":39.52,"D5":80.21,"IC50":102.23,"Slope":2.12,"max":80.21,"min":0.77,"Barcode":"H1FO4B-N8337","W5":"A5","W4":"B5","W3":"C5","W2":"E5","W1":"F5"},{"CellLine":"HPAF-II","Drug":"Prexasertib","minConc":1,"D1":6.53,"D2":9.9,"D3":47.3,"D4":39.87,"D5":55.12,"IC50":38.64,"Slope":1.58,"max":55.12,"min":5.65,"Barcode":"H1FO4B-N8337","W5":"P6","W4":"O6","W3":"N6","W2":"M6","W1":"L6"},{"CellLine":"HPAF-II","Drug":"Romidepsin","minConc":0.1,"D1":5.74,"D2":1.08,"D3":53.31,"D4":76.84,"D5":83.28,"IC50":7.44,"Slope":1.79,"max":83.28,"min":1.08,"Barcode":"H1FO4B-N8337","W5":"F18","W4":"G18","W3":"H18","W2":"I18","W1":"J18"},{"CellLine":"HPAF-II","Drug":"Sabutoclax","minConc":2.5,"D1":8.99,"D2":-3.99,"D3":28.55,"D4":36.25,"D5":82.7,"IC50":1628.96,"Slope":0.69,"max":82.7,"min":0,"Barcode":"H1FO4B-N8337","W5":"F19","W4":"G19","W3":"I19","W2":"J19","W1":"K19"},{"CellLine":"HPAF-II","Drug":"BCI","minConc":5,"D1":3.53,"D2":-4.55,"D3":-6.76,"D4":83.53,"D5":102.12,"IC50":2674.45,"Slope":2.5,"max":100,"min":0,"Barcode":"H1FO4B-N8337","W5":"A22","W4":"B22","W3":"C22","W2":"D22","W1":"E22"},{"CellLine":"CFPAC-1","Drug":"PF_04691502","minConc":1,"D1":-0.37,"D2":-2.39,"D3":12.55,"D4":53.56,"D5":77.01,"IC50":463.92,"Slope":1.14,"max":77.01,"min":0,"Barcode":"H1FO4B-N1298","W5":"B2","W4":"C2","W3":"D2","W2":"E2","W1":"F2"},{"CellLine":"CFPAC-1","Drug":"SNS_032","minConc":1,"D1":-0.21,"D2":4.5,"D3":54.8,"D4":79.28,"D5":77.16,"IC50":60.08,"Slope":1.59,"max":79.28,"min":0,"Barcode":"H1FO4B-N1298","W5":"G9","W4":"H9","W3":"I9","W2":"J9","W1":"K9"},{"CellLine":"CFPAC-1","Drug":"Alvocidib","minConc":1,"D1":3.12,"D2":6.37,"D3":28.83,"D4":85.41,"D5":85.72,"IC50":141.55,"Slope":2.27,"max":85.72,"min":3.12,"Barcode":"H1FO4B-N1298","W5":"P9","W4":"O9","W3":"N9","W2":"M9","W1":"L9"},{"CellLine":"CFPAC-1","Drug":"KX2_391","minConc":1,"D1":-2.91,"D2":3.38,"D3":56.81,"D4":57.53,"D5":61.16,"IC50":33.56,"Slope":2.35,"max":61.16,"min":0,"Barcode":"H1FO4B-N1298","W5":"F17","W4":"G17","W3":"H17","W2":"I17","W1":"J17"},{"CellLine":"CFPAC-1","Drug":"AZD8055","minConc":1,"D1":5.44,"D2":1.47,"D3":13.28,"D4":40.02,"D5":46.42,"IC50":231.9,"Slope":1.24,"max":46.42,"min":1.47,"Barcode":"H1FO4B-N1298","W5":"G23","W4":"H23","W3":"I23","W2":"J23","W1":"K23"},{"CellLine":"CFPAC-1","Drug":"PF_03758309","minConc":1,"D1":7.45,"D2":25.1,"D3":35.43,"D4":54.98,"D5":89.07,"IC50":292.95,"Slope":0.56,"max":89.07,"min":7.45,"Barcode":"H1FO4B-N2298","W5":"P5","W4":"O5","W3":"N5","W2":"M5","W1":"L5"},{"CellLine":"CFPAC-1","Drug":"AZD_8330","minConc":1,"D1":7.59,"D2":16.01,"D3":11.8,"D4":22.27,"D5":25.1,"IC50":110.59,"Slope":0.51,"max":25.1,"min":7.59,"Barcode":"H1FO4B-N2298","W5":"F6","W4":"H6","W3":"I6","W2":"J6","W1":"K6"},{"CellLine":"CFPAC-1","Drug":"Rigosertib","minConc":1,"D1":9.3,"D2":5.43,"D3":55.32,"D4":51.62,"D5":56.16,"IC50":32.8,"Slope":2.5,"max":56.16,"min":5.43,"Barcode":"H1FO4B-N2298","W5":"F8","W4":"G8","W3":"H8","W2":"J8","W1":"K8"},{"CellLine":"CFPAC-1","Drug":"Omipalisib","minConc":0.1,"D1":8.87,"D2":10.58,"D3":35.3,"D4":47.64,"D5":53.81,"IC50":7.43,"Slope":0.98,"max":53.81,"min":6.87,"Barcode":"H1FO4B-N2298","W5":"P17","W4":"O17","W3":"N17","W2":"M17","W1":"K17"},{"CellLine":"CFPAC-1","Drug":"UCN_01","minConc":1,"D1":13.01,"D2":6.69,"D3":13.03,"D4":57.85,"D5":93.04,"IC50":746.14,"Slope":1.37,"max":93.04,"min":6.69,"Barcode":"H1FO4B-N2298","W5":"P21","W4":"O21","W3":"N21","W2":"M21","W1":"L21"},{"CellLine":"CFPAC-1","Drug":"Dinaciclib","minConc":0.1,"D1":-1.26,"D2":0.59,"D3":46.84,"D4":85.62,"D5":86.2,"IC50":9.24,"Slope":2.19,"max":86.2,"min":0,"Barcode":"H1FO4B-N3298","W5":"A3","W4":"B3","W3":"C3","W2":"D3","W1":"E3"},{"CellLine":"CFPAC-1","Drug":"NVP_BGT226","minConc":0.1,"D1":3.66,"D2":7.02,"D3":6.17,"D4":39.2,"D5":93.18,"IC50":121.23,"Slope":2.15,"max":93.18,"min":3.66,"Barcode":"H1FO4B-N3298","W5":"G12","W4":"H12","W3":"I12","W2":"J12","W1":"K12"},{"CellLine":"CFPAC-1","Drug":"Gedatolisib","minConc":0.1,"D1":4,"D2":-3.16,"D3":5.84,"D4":42.56,"D5":64.23,"IC50":59.98,"Slope":1.36,"max":64.23,"min":0,"Barcode":"H1FO4B-N3298","W5":"P12","W4":"O12","W3":"N12","W2":"M12","W1":"L12"},{"CellLine":"CFPAC-1","Drug":"PKI_402","minConc":0.25,"D1":6.43,"D2":1.62,"D3":2.22,"D4":38.34,"D5":54.29,"IC50":174.45,"Slope":2.32,"max":54.29,"min":1.62,"Barcode":"H1FO4B-N3298","W5":"A16","W4":"C16","W3":"D16","W2":"E16","W1":"F16"},{"CellLine":"CFPAC-1","Drug":"AT7519","minConc":1,"D1":-7.55,"D2":-3.85,"D3":-1.13,"D4":70.69,"D5":72.95,"IC50":349.98,"Slope":2.5,"max":72.95,"min":0,"Barcode":"H1FO4B-N3298","W5":"F22","W4":"G22","W3":"H22","W2":"I22","W1":"J22"},{"CellLine":"CFPAC-1","Drug":"GSK_461364","minConc":1,"D1":1.43,"D2":12.29,"D3":31.85,"D4":32.22,"D5":72.03,"IC50":119.46,"Slope":0.55,"max":72.03,"min":0.01,"Barcode":"H1FO4B-N4298","W5":"A6","W4":"B6","W3":"C6","W2":"D6","W1":"E6"},{"CellLine":"CFPAC-1","Drug":"Daporinad","minConc":0.1,"D1":5.4,"D2":8.66,"D3":68.31,"D4":88.01,"D5":86.8,"IC50":5.41,"Slope":1.89,"max":88.01,"min":5.38,"Barcode":"H1FO4B-N4298","W5":"A14","W4":"B14","W3":"C14","W2":"D14","W1":"E14"},{"CellLine":"CFPAC-1","Drug":"Mepacrine","minConc":5,"D1":2.25,"D2":-3.67,"D3":-1.11,"D4":91.22,"D5":99.42,"IC50":2035.37,"Slope":2.5,"max":99.42,"min":0,"Barcode":"H1FO4B-N4298","W5":"P20","W4":"O20","W3":"N20","W2":"M20","W1":"L20"},{"CellLine":"CFPAC-1","Drug":"Camptothecin","minConc":0.5,"D1":6.35,"D2":13.55,"D3":57.52,"D4":68.66,"D5":88.74,"IC50":33.22,"Slope":0.68,"max":88.74,"min":0,"Barcode":"H1FO4B-N5298","W5":"O3","W4":"N3","W3":"M3","W2":"L3","W1":"K3"},{"CellLine":"CFPAC-1","Drug":"Omacetaxine","minConc":1,"D1":3.3,"D2":3.29,"D3":18.06,"D4":74.49,"D5":84.84,"IC50":272.38,"Slope":1.49,"max":84.84,"min":3,"Barcode":"H1FO4B-N5298","W5":"A5","W4":"B5","W3":"C5","W2":"E5","W1":"F5"},{"CellLine":"CFPAC-1","Drug":"Vinorelbine","minConc":1,"D1":6.27,"D2":38.13,"D3":58.92,"D4":60.76,"D5":86.14,"IC50":30.48,"Slope":0.47,"max":86.14,"min":0,"Barcode":"H1FO4B-N5298","W5":"P14","W4":"O14","W3":"N14","W2":"M14","W1":"L14"},{"CellLine":"CFPAC-1","Drug":"Docetaxel","minConc":0.1,"D1":13.35,"D2":19.28,"D3":43,"D4":53.69,"D5":54.88,"IC50":4.43,"Slope":1.14,"max":54.88,"min":12.8,"Barcode":"H1FO4B-N5298","W5":"P17","W4":"O17","W3":"N17","W2":"M17","W1":"K17"},{"CellLine":"CFPAC-1","Drug":"Carfilzomib","minConc":0.1,"D1":4.99,"D2":4.14,"D3":91.01,"D4":96.67,"D5":98.56,"IC50":3.91,"Slope":2.5,"max":98.56,"min":3.1,"Barcode":"H1FO4B-N5298","W5":"A18","W4":"B18","W3":"C18","W2":"D18","W1":"E18"},{"CellLine":"CFPAC-1","Drug":"Patupilone","minConc":0.1,"D1":8.81,"D2":3.04,"D3":33.5,"D4":53.21,"D5":53.52,"IC50":8.46,"Slope":2.5,"max":53.52,"min":3.04,"Barcode":"H1FO4B-N5298","W5":"F18","W4":"G18","W3":"H18","W2":"I18","W1":"J18"},{"CellLine":"CFPAC-1","Drug":"Plicamycin","minConc":1,"D1":7.55,"D2":8.54,"D3":61.1,"D4":89.91,"D5":90.39,"IC50":75.39,"Slope":2.14,"max":90.39,"min":7.5,"Barcode":"H1FO4B-N5298","W5":"P18","W4":"N18","W3":"M18","W2":"L18","W1":"K18"},{"CellLine":"CFPAC-1","Drug":"Vincristine","minConc":0.1,"D1":7.25,"D2":6.5,"D3":37.95,"D4":56.32,"D5":58.4,"IC50":8.09,"Slope":2,"max":58.4,"min":6.5,"Barcode":"H1FO4B-N5298","W5":"A19","W4":"B19","W3":"C19","W2":"D19","W1":"E19"},{"CellLine":"CFPAC-1","Drug":"Dactinomycin","minConc":0.1,"D1":6.03,"D2":22.19,"D3":52.57,"D4":85.99,"D5":95.14,"IC50":7.6,"Slope":0.78,"max":95.14,"min":4.37,"Barcode":"H1FO4B-N5298","W5":"P19","W4":"O19","W3":"N19","W2":"M19","W1":"L19"},{"CellLine":"CFPAC-1","Drug":"Vinblastine","minConc":0.1,"D1":3.08,"D2":3.95,"D3":26.56,"D4":56.89,"D5":56.12,"IC50":11.2,"Slope":2.25,"max":56.89,"min":3.08,"Barcode":"H1FO4B-N5298","W5":"A21","W4":"B21","W3":"C21","W2":"D21","W1":"E21"},{"CellLine":"CFPAC-1","Drug":"Paclitaxel","minConc":0.1,"D1":9.1,"D2":13.88,"D3":37.99,"D4":50.71,"D5":58.01,"IC50":6.78,"Slope":0.82,"max":58.01,"min":6.67,"Barcode":"H1FO4B-N5298","W5":"F21","W4":"G21","W3":"H21","W2":"I21","W1":"J21"},{"CellLine":"CFPAC-1","Drug":"Oprozomib","minConc":0.25,"D1":1.78,"D2":-1.26,"D3":26.5,"D4":87.86,"D5":97.05,"IC50":50.06,"Slope":1.44,"max":97.05,"min":0,"Barcode":"H1FO4B-N5298","W5":"A22","W4":"B22","W3":"C22","W2":"D22","W1":"E22"},{"CellLine":"CFPAC-1","Drug":"Bortezomib","minConc":0.1,"D1":-0.93,"D2":2.34,"D3":87.92,"D4":95.47,"D5":96.43,"IC50":3.98,"Slope":2.5,"max":96.43,"min":0,"Barcode":"H1FO4B-N5298","W5":"F22","W4":"G22","W3":"H22","W2":"I22","W1":"J22"},{"CellLine":"CFPAC-1","Drug":"Filanesib","minConc":0.1,"D1":8.29,"D2":8.09,"D3":37.62,"D4":49.35,"D5":43.47,"IC50":6.91,"Slope":2.5,"max":49.35,"min":8.03,"Barcode":"H1FO4B-N5298","W5":"G23","W4":"H23","W3":"I23","W2":"J23","W1":"K23"},{"CellLine":"CFPAC-1","Drug":"CUDC_907","minConc":1,"D1":2.41,"D2":3.27,"D3":40.76,"D4":81.11,"D5":93.04,"IC50":133.83,"Slope":1.04,"max":93.04,"min":0,"Barcode":"H1FO4B-N6298","W5":"F17","W4":"G17","W3":"H17","W2":"I17","W1":"J17"},{"CellLine":"CFPAC-1","Drug":"YM155","minConc":1,"D1":3.32,"D2":13.29,"D3":68.29,"D4":99.62,"D5":102.62,"IC50":56.37,"Slope":1.29,"max":100,"min":3.28,"Barcode":"H1FO4B-N6298","W5":"F18","W4":"G18","W3":"H18","W2":"I18","W1":"J18"},{"CellLine":"CFPAC-1","Drug":"AT101","minConc":10,"D1":5.37,"D2":1.57,"D3":1.68,"D4":49.98,"D5":104.19,"IC50":10121.99,"Slope":2.5,"max":100,"min":1.57,"Barcode":"H1FO4B-N6298","W5":"A21","W4":"B21","W3":"C21","W2":"D21","W1":"E21"},{"CellLine":"CFPAC-1","Drug":"Selinexor","minConc":1,"D1":0.68,"D2":10.02,"D3":42.94,"D4":74.7,"D5":79.47,"IC50":83.16,"Slope":1.01,"max":79.47,"min":0.56,"Barcode":"H1FO4B-N6298","W5":"P21","W4":"O21","W3":"N21","W2":"M21","W1":"L21"},{"CellLine":"CFPAC-1","Drug":"BIIB021","minConc":1,"D1":10.23,"D2":9.11,"D3":29.01,"D4":57.37,"D5":64.79,"IC50":171.31,"Slope":1.13,"max":64.79,"min":8.65,"Barcode":"H1FO4B-N7298","W5":"P13","W4":"N13","W3":"M13","W2":"L13","W1":"K13"},{"CellLine":"CFPAC-1","Drug":"Tanespimycin","minConc":1,"D1":8.82,"D2":-9.41,"D3":41.48,"D4":57.93,"D5":51.93,"IC50":69.92,"Slope":2.5,"max":57.93,"min":0,"Barcode":"H1FO4B-N7298","W5":"A14","W4":"B14","W3":"C14","W2":"D14","W1":"E14"},{"CellLine":"CFPAC-1","Drug":"CUDC_305","minConc":1,"D1":8.6,"D2":2.2,"D3":7.51,"D4":48.8,"D5":49.13,"IC50":224.67,"Slope":2.5,"max":49.13,"min":2.2,"Barcode":"H1FO4B-N7298","W5":"A15","W4":"B15","W3":"C15","W2":"D15","W1":"F15"},{"CellLine":"CFPAC-1","Drug":"Luminespib","minConc":0.1,"D1":2.11,"D2":4.5,"D3":30.71,"D4":52.19,"D5":50.22,"IC50":8.24,"Slope":1.54,"max":52.19,"min":2.11,"Barcode":"H1FO4B-N7298","W5":"G15","W4":"H15","W3":"I15","W2":"J15","W1":"K15"},{"CellLine":"CFPAC-1","Drug":"Ganetespib","minConc":0.1,"D1":3.68,"D2":-1.82,"D3":10.73,"D4":54.58,"D5":47.44,"IC50":17.53,"Slope":2.5,"max":54.58,"min":0,"Barcode":"H1FO4B-N7298","W5":"A16","W4":"C16","W3":"D16","W2":"E16","W1":"F16"},{"CellLine":"CFPAC-1","Drug":"Stattic","minConc":5,"D1":4.64,"D2":4.52,"D3":-5.86,"D4":52.52,"D5":104.04,"IC50":4804.81,"Slope":2.5,"max":100,"min":0,"Barcode":"H1FO4B-N7298","W5":"F21","W4":"G21","W3":"H21","W2":"I21","W1":"J21"},{"CellLine":"CFPAC-1","Drug":"ONX_0914","minConc":1,"D1":0.44,"D2":-6.88,"D3":31.73,"D4":96.53,"D5":99.04,"IC50":138.1,"Slope":2.35,"max":99.04,"min":0,"Barcode":"H1FO4B-N7298","W5":"G23","W4":"H23","W3":"I23","W2":"J23","W1":"K23"},{"CellLine":"CFPAC-1","Drug":"SN_38","minConc":1,"D1":14.93,"D2":62.69,"D3":68.53,"D4":87.2,"D5":88.76,"IC50":5.33,"Slope":0.72,"max":88.76,"min":0,"Barcode":"H1FO4B-N8298","W5":"G2","W4":"H2","W3":"I2","W2":"J2","W1":"K2"},{"CellLine":"CFPAC-1","Drug":"Cabazitaxel","minConc":0.1,"D1":12.02,"D2":18.58,"D3":49.69,"D4":60.37,"D5":52.43,"IC50":3.84,"Slope":1.33,"max":60.37,"min":11.63,"Barcode":"H1FO4B-N8298","W5":"A3","W4":"B3","W3":"C3","W2":"D3","W1":"E3"},{"CellLine":"CFPAC-1","Drug":"OTSSP167","minConc":0.1,"D1":4.36,"D2":1.65,"D3":0.42,"D4":39.17,"D5":83.67,"IC50":105.69,"Slope":2.5,"max":83.67,"min":0.42,"Barcode":"H1FO4B-N8298","W5":"A5","W4":"B5","W3":"C5","W2":"E5","W1":"F5"},{"CellLine":"CFPAC-1","Drug":"Prexasertib","minConc":1,"D1":10.53,"D2":40.85,"D3":61.23,"D4":49.03,"D5":33.54,"IC50":4.86,"Slope":0.98,"max":61.23,"min":0,"Barcode":"H1FO4B-N8298","W5":"P6","W4":"O6","W3":"N6","W2":"M6","W1":"L6"},{"CellLine":"CFPAC-1","Drug":"Romidepsin","minConc":0.1,"D1":4.57,"D2":5.02,"D3":71.87,"D4":86.36,"D5":88.52,"IC50":5.72,"Slope":2.5,"max":88.52,"min":4.26,"Barcode":"H1FO4B-N8298","W5":"F18","W4":"G18","W3":"H18","W2":"I18","W1":"J18"},{"CellLine":"CFPAC-1","Drug":"Sabutoclax","minConc":2.5,"D1":14.61,"D2":11.51,"D3":16.05,"D4":45.03,"D5":86.2,"IC50":2795.3,"Slope":1.61,"max":86.2,"min":11.51,"Barcode":"H1FO4B-N8298","W5":"F19","W4":"G19","W3":"I19","W2":"J19","W1":"K19"},{"CellLine":"CFPAC-1","Drug":"BCI","minConc":5,"D1":3.65,"D2":3.48,"D3":-7.12,"D4":90.67,"D5":101.54,"IC50":2209.7,"Slope":2.5,"max":100,"min":0,"Barcode":"H1FO4B-N8298","W5":"A22","W4":"B22","W3":"C22","W2":"D22","W1":"E22"},{"CellLine":"HPNE","Drug":"PF_04691502","minConc":1,"D1":-4.39,"D2":6.31,"D3":18.65,"D4":77.42,"D5":89.05,"IC50":256.49,"Slope":1.36,"max":89.05,"min":0,"Barcode":"H1FO4B-N1310","W5":"B2","W4":"C2","W3":"D2","W2":"E2","W1":"F2"},{"CellLine":"HPNE","Drug":"SNS_032","minConc":1,"D1":0.33,"D2":4.74,"D3":64.05,"D4":86.58,"D5":87.7,"IC50":55.46,"Slope":1.68,"max":87.7,"min":0.17,"Barcode":"H1FO4B-N1310","W5":"G9","W4":"H9","W3":"I9","W2":"J9","W1":"K9"},{"CellLine":"HPNE","Drug":"Alvocidib","minConc":1,"D1":-10.02,"D2":5.81,"D3":31.54,"D4":88.34,"D5":92.43,"IC50":152.39,"Slope":1.5,"max":92.43,"min":0,"Barcode":"H1FO4B-N1310","W5":"P9","W4":"O9","W3":"N9","W2":"M9","W1":"L9"},{"CellLine":"HPNE","Drug":"KX2_391","minConc":1,"D1":3.49,"D2":-4.38,"D3":69.96,"D4":84.72,"D5":87.73,"IC50":58.46,"Slope":2.5,"max":87.73,"min":0,"Barcode":"H1FO4B-N1310","W5":"F17","W4":"G17","W3":"H17","W2":"I17","W1":"J17"},{"CellLine":"HPNE","Drug":"AZD8055","minConc":1,"D1":6.47,"D2":19.05,"D3":51.54,"D4":80.81,"D5":83.06,"IC50":62.82,"Slope":0.96,"max":83.06,"min":6.14,"Barcode":"H1FO4B-N1310","W5":"G23","W4":"H23","W3":"I23","W2":"J23","W1":"K23"},{"CellLine":"HPNE","Drug":"PF_03758309","minConc":1,"D1":12.59,"D2":31.77,"D3":41.22,"D4":47.47,"D5":96.94,"IC50":494.92,"Slope":0.51,"max":96.94,"min":12.59,"Barcode":"H1FO4B-N2310","W5":"P5","W4":"O5","W3":"N5","W2":"M5","W1":"L5"},{"CellLine":"HPNE","Drug":"AZD_8330","minConc":1,"D1":14.97,"D2":16.18,"D3":23.21,"D4":35.77,"D5":44.16,"IC50":307.88,"Slope":0.87,"max":44.16,"min":14.84,"Barcode":"H1FO4B-N2310","W5":"F6","W4":"H6","W3":"I6","W2":"J6","W1":"K6"},{"CellLine":"HPNE","Drug":"Rigosertib","minConc":1,"D1":7.37,"D2":9.96,"D3":76.33,"D4":88.17,"D5":89.97,"IC50":47.38,"Slope":2.17,"max":89.97,"min":7.29,"Barcode":"H1FO4B-N2310","W5":"F8","W4":"G8","W3":"H8","W2":"J8","W1":"K8"},{"CellLine":"HPNE","Drug":"Omipalisib","minConc":0.1,"D1":0.44,"D2":-1.01,"D3":30.19,"D4":68.09,"D5":78.45,"IC50":16.08,"Slope":1.14,"max":78.45,"min":0,"Barcode":"H1FO4B-N2310","W5":"P17","W4":"O17","W3":"N17","W2":"M17","W1":"K17"},{"CellLine":"HPNE","Drug":"UCN_01","minConc":1,"D1":0.88,"D2":13.78,"D3":81.78,"D4":93.56,"D5":95.94,"IC50":32.35,"Slope":1.54,"max":95.94,"min":0.37,"Barcode":"H1FO4B-N2310","W5":"P21","W4":"O21","W3":"N21","W2":"M21","W1":"L21"},{"CellLine":"HPNE","Drug":"Dinaciclib","minConc":0.1,"D1":7.95,"D2":3.36,"D3":39.88,"D4":90.29,"D5":90.57,"IC50":11.4,"Slope":2.5,"max":90.57,"min":3.36,"Barcode":"H1FO4B-N3310","W5":"A3","W4":"B3","W3":"C3","W2":"D3","W1":"E3"},{"CellLine":"HPNE","Drug":"NVP_BGT226","minConc":0.1,"D1":6.88,"D2":12.4,"D3":61.86,"D4":82.08,"D5":98.2,"IC50":6.94,"Slope":0.83,"max":98.2,"min":1.27,"Barcode":"H1FO4B-N3310","W5":"G12","W4":"H12","W3":"I12","W2":"J12","W1":"K12"},{"CellLine":"HPNE","Drug":"Gedatolisib","minConc":0.1,"D1":9.65,"D2":12.38,"D3":45.96,"D4":82.98,"D5":86.91,"IC50":10.9,"Slope":1.33,"max":86.91,"min":9.41,"Barcode":"H1FO4B-N3310","W5":"P12","W4":"O12","W3":"N12","W2":"M12","W1":"L12"},{"CellLine":"HPNE","Drug":"PKI_402","minConc":0.25,"D1":-3.5,"D2":4.13,"D3":42.19,"D4":77.68,"D5":86.39,"IC50":27.19,"Slope":1.11,"max":86.39,"min":0,"Barcode":"H1FO4B-N3310","W5":"A16","W4":"C16","W3":"D16","W2":"E16","W1":"F16"},{"CellLine":"HPNE","Drug":"AT7519","minConc":1,"D1":6.88,"D2":3.06,"D3":4.28,"D4":82.61,"D5":86.82,"IC50":349.88,"Slope":2.5,"max":86.82,"min":3.06,"Barcode":"H1FO4B-N3310","W5":"F22","W4":"G22","W3":"H22","W2":"I22","W1":"J22"},{"CellLine":"HPNE","Drug":"GSK_461364","minConc":1,"D1":0.92,"D2":29.77,"D3":53.11,"D4":67.01,"D5":83.78,"IC50":42.41,"Slope":0.59,"max":83.78,"min":0,"Barcode":"H1FO4B-N4310","W5":"A6","W4":"B6","W3":"C6","W2":"D6","W1":"E6"},{"CellLine":"HPNE","Drug":"Daporinad","minConc":0.1,"D1":-2.27,"D2":5.73,"D3":7.77,"D4":4.38,"D5":-9.1,"IC50":1000,"Slope":0.07,"max":7.77,"min":0,"Barcode":"H1FO4B-N4310","W5":"A14","W4":"B14","W3":"C14","W2":"D14","W1":"E14"},{"CellLine":"HPNE","Drug":"Mepacrine","minConc":5,"D1":-0.27,"D2":0.87,"D3":5.86,"D4":99.94,"D5":101.93,"IC50":1352.96,"Slope":2.5,"max":100,"min":0,"Barcode":"H1FO4B-N4310","W5":"P20","W4":"O20","W3":"N20","W2":"M20","W1":"L20"},{"CellLine":"HPNE","Drug":"Camptothecin","minConc":0.5,"D1":6.06,"D2":7.4,"D3":48.41,"D4":96.39,"D5":97.71,"IC50":54.25,"Slope":1.85,"max":97.71,"min":6.06,"Barcode":"H1FO4B-N5310","W5":"O3","W4":"N3","W3":"M3","W2":"L3","W1":"K3"},{"CellLine":"HPNE","Drug":"Omacetaxine","minConc":1,"D1":3.27,"D2":0.82,"D3":85.71,"D4":95.15,"D5":98.98,"IC50":48.12,"Slope":2.5,"max":98.98,"min":0.82,"Barcode":"H1FO4B-N5310","W5":"A5","W4":"B5","W3":"C5","W2":"E5","W1":"F5"},{"CellLine":"HPNE","Drug":"Vinorelbine","minConc":1,"D1":21.66,"D2":73.05,"D3":89.11,"D4":90.33,"D5":97.54,"IC50":3.53,"Slope":0.93,"max":97.54,"min":0,"Barcode":"H1FO4B-N5310","W5":"P14","W4":"O14","W3":"N14","W2":"M14","W1":"L14"},{"CellLine":"HPNE","Drug":"Docetaxel","minConc":0.1,"D1":-0.23,"D2":41,"D3":72.5,"D4":70.18,"D5":68.09,"IC50":0.9,"Slope":2.5,"max":72.5,"min":0,"Barcode":"H1FO4B-N5310","W5":"P17","W4":"O17","W3":"N17","W2":"M17","W1":"K17"},{"CellLine":"HPNE","Drug":"Carfilzomib","minConc":0.1,"D1":0.42,"D2":4.63,"D3":99.09,"D4":98.28,"D5":100.47,"IC50":2.86,"Slope":2.5,"max":100,"min":0,"Barcode":"H1FO4B-N5310","W5":"A18","W4":"B18","W3":"C18","W2":"D18","W1":"E18"},{"CellLine":"HPNE","Drug":"Patupilone","minConc":0.1,"D1":0.27,"D2":3.82,"D3":53.07,"D4":68.1,"D5":65.15,"IC50":4.97,"Slope":1.8,"max":68.1,"min":0.23,"Barcode":"H1FO4B-N5310","W5":"F18","W4":"G18","W3":"H18","W2":"I18","W1":"J18"},{"CellLine":"HPNE","Drug":"Plicamycin","minConc":1,"D1":-1.88,"D2":0.56,"D3":90.85,"D4":102.25,"D5":99.88,"IC50":41.23,"Slope":2.5,"max":100,"min":0,"Barcode":"H1FO4B-N5310","W5":"P18","W4":"N18","W3":"M18","W2":"L18","W1":"K18"},{"CellLine":"HPNE","Drug":"Vincristine","minConc":0.1,"D1":-0.53,"D2":-2.22,"D3":81.84,"D4":94,"D5":91.51,"IC50":4.78,"Slope":2.5,"max":94,"min":0,"Barcode":"H1FO4B-N5310","W5":"A19","W4":"B19","W3":"C19","W2":"D19","W1":"E19"},{"CellLine":"HPNE","Drug":"Dactinomycin","minConc":0.1,"D1":4.52,"D2":26.94,"D3":62.28,"D4":102.25,"D5":100.78,"IC50":4.79,"Slope":0.89,"max":100,"min":3.76,"Barcode":"H1FO4B-N5310","W5":"P19","W4":"O19","W3":"N19","W2":"M19","W1":"L19"},{"CellLine":"HPNE","Drug":"Vinblastine","minConc":0.1,"D1":-4.21,"D2":0.54,"D3":73.03,"D4":88.67,"D5":90.61,"IC50":5.67,"Slope":2.5,"max":90.61,"min":0,"Barcode":"H1FO4B-N5310","W5":"A21","W4":"B21","W3":"C21","W2":"D21","W1":"E21"},{"CellLine":"HPNE","Drug":"Paclitaxel","minConc":0.1,"D1":0.3,"D2":11.37,"D3":59.41,"D4":74.56,"D5":72.17,"IC50":3.6,"Slope":1.35,"max":74.56,"min":0,"Barcode":"H1FO4B-N5310","W5":"F21","W4":"G21","W3":"H21","W2":"I21","W1":"J21"},{"CellLine":"HPNE","Drug":"Oprozomib","minConc":0.25,"D1":-1.31,"D2":-6.45,"D3":14.24,"D4":98.05,"D5":101.46,"IC50":51.34,"Slope":2.5,"max":100,"min":0,"Barcode":"H1FO4B-N5310","W5":"A22","W4":"B22","W3":"C22","W2":"D22","W1":"E22"},{"CellLine":"HPNE","Drug":"Bortezomib","minConc":0.1,"D1":-3.92,"D2":-0.04,"D3":100.8,"D4":98.76,"D5":99.2,"IC50":3.12,"Slope":2.5,"max":100,"min":0,"Barcode":"H1FO4B-N5310","W5":"F22","W4":"G22","W3":"H22","W2":"I22","W1":"J22"},{"CellLine":"HPNE","Drug":"Filanesib","minConc":0.1,"D1":0.31,"D2":-9.15,"D3":64.78,"D4":62.76,"D5":59.92,"IC50":3.77,"Slope":2.5,"max":64.78,"min":0,"Barcode":"H1FO4B-N5310","W5":"G23","W4":"H23","W3":"I23","W2":"J23","W1":"K23"},{"CellLine":"HPNE","Drug":"CUDC_907","minConc":1,"D1":7.73,"D2":41.14,"D3":92.1,"D4":96.97,"D5":98.81,"IC50":14.26,"Slope":1.3,"max":98.81,"min":4.86,"Barcode":"H1FO4B-N6310","W5":"F17","W4":"G17","W3":"H17","W2":"I17","W1":"J17"},{"CellLine":"HPNE","Drug":"YM155","minConc":1,"D1":4.07,"D2":7.85,"D3":29.55,"D4":95.01,"D5":102.81,"IC50":182.58,"Slope":1.67,"max":100,"min":4.07,"Barcode":"H1FO4B-N6310","W5":"F18","W4":"G18","W3":"H18","W2":"I18","W1":"J18"},{"CellLine":"HPNE","Drug":"AT101","minConc":10,"D1":1.25,"D2":-5.4,"D3":4.03,"D4":79.3,"D5":103.41,"IC50":5104.24,"Slope":2.01,"max":100,"min":0,"Barcode":"H1FO4B-N6310","W5":"A21","W4":"B21","W3":"C21","W2":"D21","W1":"E21"},{"CellLine":"HPNE","Drug":"Selinexor","minConc":1,"D1":5.4,"D2":7.49,"D3":68.03,"D4":92.69,"D5":97.02,"IC50":63.78,"Slope":1.71,"max":97.02,"min":4.58,"Barcode":"H1FO4B-N6310","W5":"P21","W4":"O21","W3":"N21","W2":"M21","W1":"L21"},{"CellLine":"HPNE","Drug":"BIIB021","minConc":1,"D1":8.97,"D2":-1.88,"D3":30.66,"D4":88.92,"D5":91.87,"IC50":144.92,"Slope":1.88,"max":91.87,"min":0,"Barcode":"H1FO4B-N7310","W5":"P13","W4":"N13","W3":"M13","W2":"L13","W1":"K13"},{"CellLine":"HPNE","Drug":"Tanespimycin","minConc":1,"D1":3.5,"D2":-2.38,"D3":29.02,"D4":85.83,"D5":78.68,"IC50":130.84,"Slope":2.5,"max":85.83,"min":0,"Barcode":"H1FO4B-N7310","W5":"A14","W4":"B14","W3":"C14","W2":"D14","W1":"E14"},{"CellLine":"HPNE","Drug":"CUDC_305","minConc":1,"D1":2.93,"D2":-5.54,"D3":8.61,"D4":86.57,"D5":86,"IC50":234.55,"Slope":2.5,"max":86.57,"min":0,"Barcode":"H1FO4B-N7310","W5":"A15","W4":"B15","W3":"C15","W2":"D15","W1":"F15"},{"CellLine":"HPNE","Drug":"Luminespib","minConc":0.1,"D1":8.01,"D2":2.26,"D3":27.19,"D4":87.9,"D5":86.45,"IC50":14.27,"Slope":2.5,"max":87.9,"min":2.26,"Barcode":"H1FO4B-N7310","W5":"G15","W4":"H15","W3":"I15","W2":"J15","W1":"K15"},{"CellLine":"HPNE","Drug":"Ganetespib","minConc":0.1,"D1":1.97,"D2":-0.01,"D3":13.12,"D4":89.38,"D5":87.73,"IC50":20.07,"Slope":2.5,"max":89.38,"min":0,"Barcode":"H1FO4B-N7310","W5":"A16","W4":"C16","W3":"D16","W2":"E16","W1":"F16"},{"CellLine":"HPNE","Drug":"Stattic","minConc":5,"D1":10.53,"D2":6.44,"D3":2.75,"D4":52.29,"D5":103.07,"IC50":4922.08,"Slope":2.5,"max":100,"min":2.75,"Barcode":"H1FO4B-N7310","W5":"F21","W4":"G21","W3":"H21","W2":"I21","W1":"J21"},{"CellLine":"HPNE","Drug":"ONX_0914","minConc":1,"D1":7.08,"D2":7.94,"D3":21.76,"D4":100.25,"D5":101.97,"IC50":193.96,"Slope":2.5,"max":100,"min":7.08,"Barcode":"H1FO4B-N7310","W5":"G23","W4":"H23","W3":"I23","W2":"J23","W1":"K23"},{"CellLine":"HPNE","Drug":"SN_38","minConc":1,"D1":4.01,"D2":69.18,"D3":92.01,"D4":96.62,"D5":95.37,"IC50":5.83,"Slope":1.69,"max":96.62,"min":0,"Barcode":"H1FO4B-N8310","W5":"G2","W4":"H2","W3":"I2","W2":"J2","W1":"K2"},{"CellLine":"HPNE","Drug":"Cabazitaxel","minConc":0.1,"D1":2.17,"D2":38.09,"D3":69.31,"D4":66.51,"D5":65.31,"IC50":0.94,"Slope":2.5,"max":69.31,"min":1.92,"Barcode":"H1FO4B-N8310","W5":"A3","W4":"B3","W3":"C3","W2":"D3","W1":"E3"},{"CellLine":"HPNE","Drug":"OTSSP167","minConc":0.1,"D1":4.03,"D2":11.98,"D3":18.07,"D4":89.21,"D5":95.29,"IC50":24.75,"Slope":1.85,"max":95.29,"min":4.03,"Barcode":"H1FO4B-N8310","W5":"A5","W4":"B5","W3":"C5","W2":"E5","W1":"F5"},{"CellLine":"HPNE","Drug":"Prexasertib","minConc":1,"D1":2.05,"D2":66.32,"D3":88.94,"D4":85.01,"D5":80.75,"IC50":6.55,"Slope":2.5,"max":88.94,"min":1.26,"Barcode":"H1FO4B-N8310","W5":"P6","W4":"O6","W3":"N6","W2":"M6","W1":"L6"},{"CellLine":"HPNE","Drug":"Romidepsin","minConc":0.1,"D1":0.88,"D2":25.38,"D3":92.22,"D4":95.7,"D5":96.38,"IC50":1.79,"Slope":1.79,"max":96.38,"min":0.34,"Barcode":"H1FO4B-N8310","W5":"F18","W4":"G18","W3":"H18","W2":"I18","W1":"J18"},{"CellLine":"HPNE","Drug":"Sabutoclax","minConc":2.5,"D1":3.68,"D2":2.56,"D3":40.76,"D4":81.06,"D5":99.93,"IC50":419.01,"Slope":0.91,"max":99.93,"min":0,"Barcode":"H1FO4B-N8310","W5":"F19","W4":"G19","W3":"I19","W2":"J19","W1":"K19"},{"CellLine":"HPNE","Drug":"BCI","minConc":5,"D1":-2.83,"D2":-4.04,"D3":4.93,"D4":99.85,"D5":100.35,"IC50":1397.37,"Slope":2.5,"max":100,"min":0,"Barcode":"H1FO4B-N8310","W5":"A22","W4":"B22","W3":"C22","W2":"D22","W1":"E22"},{"CellLine":"Capan-2","Drug":"PF_04691502","minConc":1,"D1":-6.95,"D2":-5.12,"D3":-25.85,"D4":17.4,"D5":49.15,"IC50":1280.41,"Slope":2.5,"max":49.15,"min":0,"Barcode":"H1FO4B-N1345","W5":"B2","W4":"C2","W3":"D2","W2":"E2","W1":"F2"},{"CellLine":"Capan-2","Drug":"SNS_032","minConc":1,"D1":-13.8,"D2":1.41,"D3":32.78,"D4":43.96,"D5":55.01,"IC50":87.11,"Slope":1.11,"max":55.01,"min":0,"Barcode":"H1FO4B-N1345","W5":"G9","W4":"H9","W3":"I9","W2":"J9","W1":"K9"},{"CellLine":"Capan-2","Drug":"Alvocidib","minConc":1,"D1":-7.51,"D2":-0.61,"D3":-0.18,"D4":51.51,"D5":55.57,"IC50":390.51,"Slope":2.5,"max":55.57,"min":0,"Barcode":"H1FO4B-N1345","W5":"P9","W4":"O9","W3":"N9","W2":"M9","W1":"L9"},{"CellLine":"Capan-2","Drug":"KX2_391","minConc":1,"D1":-4.96,"D2":-1.65,"D3":37.53,"D4":63.67,"D5":58.16,"IC50":86.6,"Slope":2.5,"max":63.67,"min":0,"Barcode":"H1FO4B-N1345","W5":"F17","W4":"G17","W3":"H17","W2":"I17","W1":"J17"},{"CellLine":"Capan-2","Drug":"AZD8055","minConc":1,"D1":-1.14,"D2":-20.9,"D3":-35.35,"D4":-20.04,"D5":-2.78,"IC50":10000,"Slope":1,"max":0,"min":0,"Barcode":"H1FO4B-N1345","W5":"G23","W4":"H23","W3":"I23","W2":"J23","W1":"K23"},{"CellLine":"Capan-2","Drug":"PF_03758309","minConc":1,"D1":7.08,"D2":32.99,"D3":66.75,"D4":63.71,"D5":74.6,"IC50":12.7,"Slope":0.86,"max":74.6,"min":0,"Barcode":"H1FO4B-N2345","W5":"P5","W4":"O5","W3":"N5","W2":"M5","W1":"L5"},{"CellLine":"Capan-2","Drug":"AZD_8330","minConc":1,"D1":-10.15,"D2":-31.03,"D3":-51.92,"D4":-48.44,"D5":-46.74,"IC50":10000,"Slope":1,"max":0,"min":0,"Barcode":"H1FO4B-N2345","W5":"F6","W4":"H6","W3":"I6","W2":"J6","W1":"K6"},{"CellLine":"Capan-2","Drug":"Rigosertib","minConc":1,"D1":0.38,"D2":-3.11,"D3":58.18,"D4":68.8,"D5":68.58,"IC50":51.74,"Slope":2.5,"max":68.8,"min":0,"Barcode":"H1FO4B-N2345","W5":"F8","W4":"G8","W3":"H8","W2":"J8","W1":"K8"},{"CellLine":"Capan-2","Drug":"Omipalisib","minConc":0.1,"D1":8.07,"D2":13.83,"D3":-1.47,"D4":5.55,"D5":7.32,"IC50":1000,"Slope":-0.14,"max":13.83,"min":0,"Barcode":"H1FO4B-N2345","W5":"P17","W4":"O17","W3":"N17","W2":"M17","W1":"K17"},{"CellLine":"Capan-2","Drug":"UCN_01","minConc":1,"D1":4.46,"D2":-0.66,"D3":-35.78,"D4":8.43,"D5":71.56,"IC50":2214.89,"Slope":2.5,"max":71.56,"min":0,"Barcode":"H1FO4B-N2345","W5":"P21","W4":"O21","W3":"N21","W2":"M21","W1":"L21"},{"CellLine":"Capan-2","Drug":"Dinaciclib","minConc":0.1,"D1":-2.63,"D2":-0.97,"D3":22.62,"D4":52.35,"D5":55.97,"IC50":13.2,"Slope":1.5,"max":55.97,"min":0,"Barcode":"H1FO4B-N3345","W5":"A3","W4":"B3","W3":"C3","W2":"D3","W1":"E3"},{"CellLine":"Capan-2","Drug":"NVP_BGT226","minConc":0.1,"D1":-3.17,"D2":-20.04,"D3":-34.01,"D4":-14.53,"D5":-3.22,"IC50":1000,"Slope":1,"max":0,"min":0,"Barcode":"H1FO4B-N3345","W5":"G12","W4":"H12","W3":"I12","W2":"J12","W1":"K12"},{"CellLine":"Capan-2","Drug":"Gedatolisib","minConc":0.1,"D1":-6.86,"D2":-1.53,"D3":-7.01,"D4":-5.51,"D5":44.57,"IC50":371.92,"Slope":2.5,"max":44.57,"min":0,"Barcode":"H1FO4B-N3345","W5":"P12","W4":"O12","W3":"N12","W2":"M12","W1":"L12"},{"CellLine":"Capan-2","Drug":"PKI_402","minConc":0.25,"D1":-9.68,"D2":-4.44,"D3":-15.56,"D4":-25.26,"D5":18.48,"IC50":25,"Slope":1,"max":18.48,"min":0,"Barcode":"H1FO4B-N3345","W5":"A16","W4":"C16","W3":"D16","W2":"E16","W1":"F16"},{"CellLine":"Capan-2","Drug":"AT7519","minConc":1,"D1":1.53,"D2":-4.29,"D3":-9.34,"D4":49.85,"D5":43.79,"IC50":390.19,"Slope":2.5,"max":49.85,"min":0,"Barcode":"H1FO4B-N3345","W5":"F22","W4":"G22","W3":"H22","W2":"I22","W1":"J22"},{"CellLine":"Capan-2","Drug":"GSK_461364","minConc":1,"D1":-0.8,"D2":-2.58,"D3":7.56,"D4":-1.61,"D5":47.16,"IC50":3355.39,"Slope":2.5,"max":47.16,"min":0,"Barcode":"H1FO4B-N4345","W5":"A6","W4":"B6","W3":"C6","W2":"D6","W1":"E6"},{"CellLine":"Capan-2","Drug":"Daporinad","minConc":0.1,"D1":-7.1,"D2":-12.71,"D3":19.15,"D4":78.01,"D5":78.38,"IC50":15.73,"Slope":2.5,"max":78.38,"min":0,"Barcode":"H1FO4B-N4345","W5":"A14","W4":"B14","W3":"C14","W2":"D14","W1":"E14"},{"CellLine":"Capan-2","Drug":"Mepacrine","minConc":5,"D1":4.43,"D2":1.72,"D3":4.78,"D4":35.78,"D5":98.18,"IC50":6488.31,"Slope":2.31,"max":98.18,"min":1.72,"Barcode":"H1FO4B-N4345","W5":"P20","W4":"O20","W3":"N20","W2":"M20","W1":"L20"},{"CellLine":"Capan-2","Drug":"Camptothecin","minConc":0.5,"D1":1.75,"D2":1.8,"D3":-1.49,"D4":-18.72,"D5":67.51,"IC50":2048.52,"Slope":2.5,"max":67.51,"min":0,"Barcode":"H1FO4B-N5345","W5":"O3","W4":"N3","W3":"M3","W2":"L3","W1":"K3"},{"CellLine":"Capan-2","Drug":"Omacetaxine","minConc":1,"D1":0.86,"D2":17.44,"D3":50.55,"D4":31.53,"D5":40.01,"IC50":13.16,"Slope":2.5,"max":50.55,"min":0.78,"Barcode":"H1FO4B-N5345","W5":"A5","W4":"B5","W3":"C5","W2":"E5","W1":"F5"},{"CellLine":"Capan-2","Drug":"Vinorelbine","minConc":1,"D1":-3.92,"D2":22.29,"D3":65.78,"D4":72.45,"D5":92.42,"IC50":45.1,"Slope":0.76,"max":92.42,"min":0,"Barcode":"H1FO4B-N5345","W5":"P14","W4":"O14","W3":"N14","W2":"M14","W1":"L14"},{"CellLine":"Capan-2","Drug":"Docetaxel","minConc":0.1,"D1":-4.32,"D2":23.82,"D3":42.42,"D4":46.25,"D5":54.22,"IC50":1.92,"Slope":0.85,"max":54.22,"min":0,"Barcode":"H1FO4B-N5345","W5":"P17","W4":"O17","W3":"N17","W2":"M17","W1":"K17"},{"CellLine":"Capan-2","Drug":"Carfilzomib","minConc":0.1,"D1":0.5,"D2":-6.19,"D3":38.64,"D4":74.25,"D5":90.51,"IC50":15.99,"Slope":1.06,"max":90.51,"min":0,"Barcode":"H1FO4B-N5345","W5":"A18","W4":"B18","W3":"C18","W2":"D18","W1":"E18"},{"CellLine":"Capan-2","Drug":"Patupilone","minConc":0.1,"D1":-8.35,"D2":-3.68,"D3":30.76,"D4":41.41,"D5":43.31,"IC50":7.04,"Slope":2.5,"max":43.31,"min":0,"Barcode":"H1FO4B-N5345","W5":"F18","W4":"G18","W3":"H18","W2":"I18","W1":"J18"},{"CellLine":"Capan-2","Drug":"Plicamycin","minConc":1,"D1":-5.54,"D2":-4.77,"D3":48.76,"D4":56.51,"D5":60.75,"IC50":58.09,"Slope":2.5,"max":60.75,"min":0,"Barcode":"H1FO4B-N5345","W5":"P18","W4":"N18","W3":"M18","W2":"L18","W1":"K18"},{"CellLine":"Capan-2","Drug":"Vincristine","minConc":0.1,"D1":-5.36,"D2":-1.79,"D3":40.35,"D4":62.75,"D5":69.68,"IC50":8.52,"Slope":1.77,"max":69.68,"min":0,"Barcode":"H1FO4B-N5345","W5":"A19","W4":"B19","W3":"C19","W2":"D19","W1":"E19"},{"CellLine":"Capan-2","Drug":"Dactinomycin","minConc":0.1,"D1":-4.21,"D2":23.74,"D3":57.37,"D4":70.44,"D5":61.33,"IC50":2.15,"Slope":1.07,"max":70.44,"min":0,"Barcode":"H1FO4B-N5345","W5":"P19","W4":"O19","W3":"N19","W2":"M19","W1":"L19"},{"CellLine":"Capan-2","Drug":"Vinblastine","minConc":0.1,"D1":2.34,"D2":2.92,"D3":22.23,"D4":62.87,"D5":69.96,"IC50":19.24,"Slope":1.32,"max":69.96,"min":1.94,"Barcode":"H1FO4B-N5345","W5":"A21","W4":"B21","W3":"C21","W2":"D21","W1":"E21"},{"CellLine":"Capan-2","Drug":"Paclitaxel","minConc":0.1,"D1":0.64,"D2":14.61,"D3":33.91,"D4":48.82,"D5":51.89,"IC50":3.97,"Slope":0.78,"max":51.89,"min":0,"Barcode":"H1FO4B-N5345","W5":"F21","W4":"G21","W3":"H21","W2":"I21","W1":"J21"},{"CellLine":"Capan-2","Drug":"Oprozomib","minConc":0.25,"D1":-7.02,"D2":-3.83,"D3":-20.02,"D4":50.53,"D5":63.02,"IC50":150.46,"Slope":2.5,"max":63.02,"min":0,"Barcode":"H1FO4B-N5345","W5":"A22","W4":"B22","W3":"C22","W2":"D22","W1":"E22"},{"CellLine":"Capan-2","Drug":"Bortezomib","minConc":0.1,"D1":-6.18,"D2":-8.99,"D3":35.48,"D4":67.22,"D5":82.29,"IC50":15.88,"Slope":1.12,"max":82.29,"min":0,"Barcode":"H1FO4B-N5345","W5":"F22","W4":"G22","W3":"H22","W2":"I22","W1":"J22"},{"CellLine":"Capan-2","Drug":"Filanesib","minConc":0.1,"D1":-2.07,"D2":4.24,"D3":10.12,"D4":7.38,"D5":15.98,"IC50":15.48,"Slope":0.47,"max":15.98,"min":0,"Barcode":"H1FO4B-N5345","W5":"G23","W4":"H23","W3":"I23","W2":"J23","W1":"K23"},{"CellLine":"Capan-2","Drug":"CUDC_907","minConc":1,"D1":14.72,"D2":47.32,"D3":22.29,"D4":86.66,"D5":92.4,"IC50":291.31,"Slope":1.97,"max":92.4,"min":14.72,"Barcode":"H1FO4B-N6345","W5":"F17","W4":"G17","W3":"H17","W2":"I17","W1":"J17"},{"CellLine":"Capan-2","Drug":"YM155","minConc":1,"D1":8.54,"D2":22.72,"D3":66.24,"D4":101.25,"D5":103.46,"IC50":56.26,"Slope":1.09,"max":100,"min":8.54,"Barcode":"H1FO4B-N6345","W5":"F18","W4":"G18","W3":"H18","W2":"I18","W1":"J18"},{"CellLine":"Capan-2","Drug":"AT101","minConc":10,"D1":0.3,"D2":1.85,"D3":3.59,"D4":2.12,"D5":99.89,"IC50":30354.47,"Slope":2.5,"max":99.89,"min":0.3,"Barcode":"H1FO4B-N6345","W5":"A21","W4":"B21","W3":"C21","W2":"D21","W1":"E21"},{"CellLine":"Capan-2","Drug":"Selinexor","minConc":1,"D1":4.34,"D2":7.32,"D3":-39.01,"D4":-7.17,"D5":57.29,"IC50":100,"Slope":1,"max":57.29,"min":0,"Barcode":"H1FO4B-N6345","W5":"P21","W4":"O21","W3":"N21","W2":"M21","W1":"L21"},{"CellLine":"Capan-2","Drug":"BIIB021","minConc":1,"D1":4.4,"D2":7.78,"D3":-7.42,"D4":17.38,"D5":33.32,"IC50":970.67,"Slope":2.5,"max":33.32,"min":0,"Barcode":"H1FO4B-N7345","W5":"P13","W4":"N13","W3":"M13","W2":"L13","W1":"K13"},{"CellLine":"Capan-2","Drug":"Tanespimycin","minConc":1,"D1":3.09,"D2":-0.05,"D3":5.47,"D4":30.11,"D5":55.99,"IC50":841.26,"Slope":1.24,"max":55.99,"min":0,"Barcode":"H1FO4B-N7345","W5":"A14","W4":"B14","W3":"C14","W2":"D14","W1":"E14"},{"CellLine":"Capan-2","Drug":"CUDC_305","minConc":1,"D1":7.8,"D2":10.54,"D3":-1.92,"D4":14.25,"D5":25.79,"IC50":920.66,"Slope":2.5,"max":25.79,"min":0,"Barcode":"H1FO4B-N7345","W5":"A15","W4":"B15","W3":"C15","W2":"D15","W1":"F15"},{"CellLine":"Capan-2","Drug":"Luminespib","minConc":0.1,"D1":0.45,"D2":5.09,"D3":-19.6,"D4":15.89,"D5":24.02,"IC50":79.21,"Slope":2.5,"max":24.02,"min":0,"Barcode":"H1FO4B-N7345","W5":"G15","W4":"H15","W3":"I15","W2":"J15","W1":"K15"},{"CellLine":"Capan-2","Drug":"Ganetespib","minConc":0.1,"D1":1.1,"D2":6.53,"D3":-13.23,"D4":17.6,"D5":15.89,"IC50":47.23,"Slope":2.5,"max":17.6,"min":0,"Barcode":"H1FO4B-N7345","W5":"A16","W4":"C16","W3":"D16","W2":"E16","W1":"F16"},{"CellLine":"Capan-2","Drug":"Stattic","minConc":5,"D1":4.95,"D2":2.79,"D3":4.53,"D4":15.73,"D5":104.31,"IC50":10246.52,"Slope":2.5,"max":100,"min":2.79,"Barcode":"H1FO4B-N7345","W5":"F21","W4":"G21","W3":"H21","W2":"I21","W1":"J21"},{"CellLine":"Capan-2","Drug":"ONX_0914","minConc":1,"D1":7.28,"D2":1.63,"D3":-10.25,"D4":65.12,"D5":90.95,"IC50":696.91,"Slope":2.5,"max":90.95,"min":0,"Barcode":"H1FO4B-N7345","W5":"G23","W4":"H23","W3":"I23","W2":"J23","W1":"K23"},{"CellLine":"Capan-2","Drug":"SN_38","minConc":1,"D1":10.23,"D2":-0.74,"D3":4.13,"D4":54.98,"D5":56.52,"IC50":269.3,"Slope":2.5,"max":56.52,"min":0,"Barcode":"H1FO4B-N8345","W5":"G2","W4":"H2","W3":"I2","W2":"J2","W1":"K2"},{"CellLine":"Capan-2","Drug":"Cabazitaxel","minConc":0.1,"D1":2.26,"D2":16.45,"D3":45.77,"D4":48.19,"D5":49.96,"IC50":1.79,"Slope":1.35,"max":49.96,"min":1.27,"Barcode":"H1FO4B-N8345","W5":"A3","W4":"B3","W3":"C3","W2":"D3","W1":"E3"},{"CellLine":"Capan-2","Drug":"OTSSP167","minConc":0.1,"D1":-5.53,"D2":-1.8,"D3":-20.32,"D4":-32.29,"D5":39.46,"IC50":10,"Slope":1,"max":39.46,"min":0,"Barcode":"H1FO4B-N8345","W5":"A5","W4":"B5","W3":"C5","W2":"E5","W1":"F5"},{"CellLine":"Capan-2","Drug":"Prexasertib","minConc":1,"D1":0.1,"D2":1.75,"D3":29.03,"D4":17.97,"D5":-36.92,"IC50":100,"Slope":1,"max":29.03,"min":0,"Barcode":"H1FO4B-N8345","W5":"P6","W4":"O6","W3":"N6","W2":"M6","W1":"L6"},{"CellLine":"Capan-2","Drug":"Romidepsin","minConc":0.1,"D1":6.98,"D2":33.72,"D3":73.6,"D4":79.7,"D5":83.17,"IC50":1.54,"Slope":1.04,"max":83.17,"min":2.47,"Barcode":"H1FO4B-N8345","W5":"F18","W4":"G18","W3":"H18","W2":"I18","W1":"J18"},{"CellLine":"Capan-2","Drug":"Sabutoclax","minConc":2.5,"D1":4.95,"D2":1.19,"D3":7.93,"D4":19.25,"D5":32.66,"IC50":1444.6,"Slope":0.88,"max":32.66,"min":1.19,"Barcode":"H1FO4B-N8345","W5":"F19","W4":"G19","W3":"I19","W2":"J19","W1":"K19"},{"CellLine":"Capan-2","Drug":"BCI","minConc":5,"D1":-2.35,"D2":-6.21,"D3":8.49,"D4":76.9,"D5":101.17,"IC50":2332.02,"Slope":1.59,"max":100,"min":0,"Barcode":"H1FO4B-N8345","W5":"A22","W4":"B22","W3":"C22","W2":"D22","W1":"E22"}];
```
