## Supplemental Examples for "R/LinkedCharts: A novel approach for simple but powerful interactive data analysis": scoreData.js

```
drugs = ["Romidepsin","YM155","Bortezomib","Carfilzomib","CUDC_907","Dactinomycin","Vinorelbine","SN_38","Omacetaxine","SNS_032","Dinaciclib","PF_03758309","ONX_0914","Plicamycin","Prexasertib","Selinexor","Oprozomib","Tanespimycin","Alvocidib","Rigosertib","Vincristine","BCI","Daporinad","UCN_01","KX2_391","Cabazitaxel","Docetaxel","AT7519","Vinblastine","PF_04691502","BIIB021","AZD8055","Mepacrine","Luminespib","Patupilone","Paclitaxel","Sabutoclax","Omipalisib","AT101","NVP_BGT226","AZD_8330","CUDC_305","Camptothecin","Ganetespib","Stattic","GSK_461364","PKI_402","Filanesib","Gedatolisib","OTSSP167"];
cellLines = ["Pa16C","Pa14C","Pa02C","Pa18C","Pa03C","Pa01C","MDA-PATC53","Pa04C","PANC-1","MDA-PATC43","MDA-PATC50","KPC-INK4","KPC-p53","AsPC-1","HPAC","MIA-PaCa-2","SW1990","HPAF-II","CFPAC-1","HPNE","Capan-2"];
corMat = [[1,0.0796,0.319,0.2318,0.8328,0.4583,0.3559,0.3495,0.3147,0.5031,0.4803,0.3445,0.2211,0.3582,0.1577,0.4429,0.4339,0.4054,0.4657,0.2258,0.4257,0.3858,0.0113,0.3604,0.2899,0.2954,0.2649,0.3682,0.4278,0.3006,0.2722,0.4235,0.0764,0.3962,0.2099,0.2095,0.4071,0.2643,0.4642,0.3675,0.2599,0.4068,0.358,0.406,0.1892,0.2926,0.36,0.2729,0.3541,0.3859],[0.0796,1,-0.1353,-0.2729,0.0616,-0.3322,-0.4306,-0.0548,-0.2775,-0.12,-0.097,-0.1901,0.054,-0.0396,-0.231,-0.2785,-0.1042,-0.1733,-0.2024,-0.2515,-0.2937,-0.1732,0.1212,-0.4614,-0.3148,-0.282,-0.2465,-0.2744,-0.2012,-0.0966,-0.4149,-0.1928,-0.2908,-0.2866,-0.1405,-0.2977,0.0775,-0.1078,0.2722,-0.3824,0.0142,-0.3348,0.0589,-0.3296,-0.114,-0.1842,-0.3403,-0.2191,-0.2763,0.0775],[0.319,-0.1353,1,0.861,0.3991,0.4331,0.0875,0.5381,0.4576,0.5902,0.6023,-0.0556,0.7804,0.3823,0.4104,0.355,0.8433,0.481,0.7575,-0.2056,0.14,0.5821,-0.071,0.5503,0.0297,-0.1208,-0.0792,0.6075,0.1886,0.3038,0.4856,0.243,0.5423,0.5113,-0.162,-0.0583,0.3507,0.2575,0.3747,0.3178,0.2494,0.5487,0.5183,0.4166,0.3064,0.3514,0.2973,0.5478,0.3813,0.5226],[0.2318,-0.2729,0.861,1,0.3678,0.3819,0.2504,0.4783,0.2947,0.4303,0.4639,-0.0931,0.689,0.4115,0.5223,0.4347,0.8404,0.4858,0.6749,0.053,0.3132,0.6935,0.1562,0.6554,0.2955,0.0805,0.0904,0.5837,0.3385,0.3187,0.5428,0.2475,0.4698,0.5311,0.0343,0.1308,0.2417,0.3092,0.4045,0.2616,0.1567,0.559,0.4897,0.4704,0.3779,0.4404,0.3422,0.5619,0.4089,0.4598],[0.8328,0.0616,0.3991,0.3678,1,0.5345,0.5556,0.5337,0.4691,0.5547,0.5272,0.0557,0.2167,0.461,0.4979,0.629,0.4925,0.5781,0.5845,0.4571,0.6011,0.5224,0.0285,0.5118,0.5358,0.4953,0.4453,0.4202,0.6201,0.4886,0.4966,0.6215,0.1844,0.5505,0.432,0.4309,0.5848,0.5148,0.6678,0.5672,0.3677,0.6284,0.571,0.5647,0.295,0.5733,0.5821,0.5271,0.5929,0.5781],[0.4583,-0.3322,0.4331,0.3819,0.5345,1,0.5193,0.3136,0.7303,0.6791,0.5384,0.5482,0.0526,0.4121,0.5334,0.6736,0.2292,0.4835,0.7527,0.1672,0.4017,0.5029,0.0087,0.5121,0.3211,0.3885,0.4002,0.6891,0.4228,0.4215,0.636,0.4594,0.5945,0.5233,0.2267,0.4223,0.5712,0.443,0.3903,0.5083,0.2024,0.6812,0.2661,0.4683,0.2106,0.5993,0.5635,0.5458,0.56,0.5017],[0.3559,-0.4306,0.0875,0.2504,0.5556,0.5193,1,0.4393,0.4824,0.4463,0.2711,0.0713,0.047,0.3487,0.4508,0.7036,0.1919,0.4197,0.3661,0.8353,0.9294,0.5747,0.1321,0.4024,0.9166,0.888,0.8933,0.3295,0.8921,0.2115,0.5172,0.5379,0.3489,0.4914,0.7475,0.8937,0.4832,0.282,0.3223,0.5795,0.0538,0.5284,0.4588,0.6039,0.2697,0.7044,0.5246,0.692,0.5028,0.5238],[0.3495,-0.0548,0.5381,0.4783,0.5337,0.3136,0.4393,1,0.2435,0.4043,0.4707,-0.1901,0.6271,0.8111,0.4973,0.5909,0.5294,0.3525,0.4965,0.339,0.4708,0.3707,-0.1219,0.2434,0.4404,0.2885,0.3184,0.0959,0.4629,-0.0358,0.2336,0.0843,0.3193,0.3879,0.1346,0.277,0.3132,0.1254,0.3431,0.2436,0.1049,0.2514,0.9467,0.3607,-0.1213,0.4962,0.0844,0.4794,0.2002,0.523],[0.3147,-0.2775,0.4576,0.2947,0.4691,0.7303,0.4824,0.2435,1,0.6717,0.6477,0.2276,0.0177,0.0931,0.3078,0.432,0.2875,0.2365,0.6412,0.1563,0.4294,0.5804,-0.1404,0.3504,0.2555,0.3575,0.3471,0.6219,0.5396,0.3224,0.3716,0.3643,0.6971,0.4359,0.2729,0.4257,0.6116,0.2046,0.3293,0.4631,0.2148,0.5055,0.2167,0.4285,0.2856,0.4904,0.4936,0.6632,0.4398,0.6503],[0.5031,-0.12,0.5902,0.4303,0.5547,0.6791,0.4463,0.4043,0.6717,1,0.6708,0.2459,0.3464,0.1895,0.3253,0.4405,0.4407,0.4645,0.7933,0.1212,0.4169,0.4776,0.0113,0.3743,0.3089,0.3659,0.3521,0.8194,0.4436,0.4548,0.453,0.5418,0.5739,0.4223,0.4061,0.3764,0.6167,0.3412,0.3737,0.5467,0.2766,0.5859,0.3684,0.4675,0.33,0.6315,0.5345,0.6801,0.5576,0.4932],[0.4803,-0.097,0.6023,0.4639,0.5272,0.5384,0.2711,0.4707,0.6477,0.6708,1,0.2571,0.334,0.1885,0.3006,0.3743,0.5576,0.1175,0.8512,0.0105,0.3,0.5623,0.0499,0.3263,0.1928,0.2492,0.1937,0.5706,0.4922,0.2924,0.2354,0.2439,0.6296,0.3605,0.2025,0.2336,0.5491,0.2396,0.5536,0.3186,0.1912,0.382,0.4689,0.4162,0.257,0.4756,0.3156,0.5974,0.3013,0.4876],[0.3445,-0.1901,-0.0556,-0.0931,0.0557,0.5482,0.0713,-0.1901,0.2276,0.2459,0.2571,1,-0.2955,0.0818,0.017,0.2792,-0.269,-0.0417,0.2969,-0.1107,-0.0245,0.1049,0.1264,0.2072,-0.1041,0.0807,0.1027,0.3905,0.0209,0.2217,0.1567,0.1319,0.3626,0.0096,-0.0262,0.0677,0.2352,0.2001,0.0725,0.136,-0.1272,0.1945,-0.3127,0.0337,-0.0405,-0.0402,0.2321,-0.108,0.182,-0.0935],[0.2211,0.054,0.7804,0.689,0.2167,0.0526,0.047,0.6271,0.0177,0.3464,0.334,-0.2955,1,0.4511,0.1854,0.2942,0.7567,0.3858,0.4324,-0.1044,0.1156,0.2957,-0.008,0.2584,0.0509,-0.1506,-0.0645,0.2285,0.1229,0.0211,0.25,0.0944,0.2234,0.4532,-0.2207,-0.1232,0.1711,0.0667,0.2501,0.1824,-0.05,0.2375,0.6627,0.3934,0.1417,0.1701,0.0177,0.3365,0.1726,0.4494],[0.3582,-0.0396,0.3823,0.4115,0.461,0.4121,0.3487,0.8111,0.0931,0.1895,0.1885,0.0818,0.4511,1,0.4567,0.6445,0.2674,0.3742,0.3317,0.2594,0.324,0.271,-0.1649,0.3346,0.3085,0.2591,0.3416,0.0044,0.2901,0.0023,0.2973,0.0778,0.2345,0.3051,-0.0134,0.2326,0.1758,0.2275,0.1915,0.1987,0.0642,0.2384,0.7627,0.2214,-0.2612,0.3697,0.111,0.257,0.2633,0.4045],[0.1577,-0.231,0.4104,0.5223,0.4979,0.5334,0.4508,0.4973,0.3078,0.3253,0.3006,0.017,0.1854,0.4567,1,0.6077,0.29,0.5395,0.5071,0.3629,0.4745,0.4438,0.2151,0.6898,0.4628,0.2606,0.2248,0.4197,0.3862,0.4261,0.5444,0.3916,0.4057,0.4407,0.204,0.2271,0.4423,0.5152,0.548,0.3854,0.1124,0.6555,0.4226,0.4001,0.0372,0.5513,0.5334,0.3674,0.534,0.4156],[0.4429,-0.2785,0.355,0.4347,0.629,0.6736,0.7036,0.5909,0.432,0.4405,0.3743,0.2792,0.2942,0.6445,0.6077,1,0.2929,0.6535,0.5814,0.504,0.5842,0.5038,0.2128,0.6066,0.5744,0.533,0.5511,0.4581,0.6031,0.4876,0.71,0.6123,0.4483,0.7686,0.3381,0.5351,0.6504,0.6342,0.5561,0.6695,-0.1153,0.7203,0.5688,0.7622,0.1882,0.5346,0.6309,0.4626,0.6924,0.5422],[0.4339,-0.1042,0.8433,0.8404,0.4925,0.2292,0.1919,0.5294,0.2875,0.4407,0.5576,-0.269,0.7567,0.2674,0.29,0.2929,1,0.4347,0.6485,0.0193,0.2977,0.6173,0.0519,0.4031,0.259,0.0026,-0.0143,0.4413,0.3472,0.1577,0.3991,0.1786,0.2354,0.5538,0.0065,0.0529,0.2524,0.116,0.4685,0.2029,0.2317,0.4376,0.575,0.4937,0.4613,0.3711,0.157,0.5571,0.2054,0.4718],[0.4054,-0.1733,0.481,0.4858,0.5781,0.4835,0.4197,0.3525,0.2365,0.4645,0.1175,-0.0417,0.3858,0.3742,0.5395,0.6535,0.4347,1,0.488,0.1482,0.3663,0.4239,0.2467,0.6406,0.3377,0.2428,0.229,0.5086,0.2483,0.5313,0.7907,0.7266,0.0741,0.7412,0.1612,0.1883,0.4644,0.5867,0.4155,0.669,0.1928,0.8039,0.4041,0.6687,0.3723,0.5421,0.5812,0.4071,0.6909,0.2795],[0.4657,-0.2024,0.7575,0.6749,0.5845,0.7527,0.3661,0.4965,0.6412,0.7933,0.8512,0.2969,0.4324,0.3317,0.5071,0.5814,0.6485,0.488,1,0.0147,0.3188,0.6397,0.0802,0.5452,0.2729,0.2636,0.2333,0.7879,0.4416,0.4628,0.626,0.4646,0.614,0.5976,0.1979,0.2854,0.6519,0.4507,0.6103,0.5128,0.1908,0.6973,0.4765,0.5864,0.437,0.6484,0.498,0.6878,0.5482,0.4901],[0.2258,-0.2515,-0.2056,0.053,0.4571,0.1672,0.8353,0.339,0.1563,0.1212,0.0105,-0.1107,-0.1044,0.2594,0.3629,0.504,0.0193,0.1482,0.0147,1,0.8151,0.3033,0.1598,0.1708,0.9177,0.7732,0.7635,-0.0092,0.7439,0.1057,0.2262,0.3556,0.0507,0.2278,0.7428,0.7768,0.1804,0.1989,0.2167,0.3829,-0.0125,0.2331,0.3436,0.3635,0.1874,0.3869,0.3843,0.3636,0.3107,0.3524],[0.4257,-0.2937,0.14,0.3132,0.6011,0.4017,0.9294,0.4708,0.4294,0.4169,0.3,-0.0245,0.1156,0.324,0.4745,0.5842,0.2977,0.3663,0.3188,0.8151,1,0.6528,0.2585,0.3658,0.9126,0.8171,0.7999,0.3044,0.9312,0.1059,0.3461,0.4123,0.3307,0.4276,0.699,0.7872,0.4409,0.1509,0.4101,0.3999,0.1096,0.4184,0.5086,0.5474,0.2156,0.6701,0.3981,0.697,0.3702,0.6002],[0.3858,-0.1732,0.5821,0.6935,0.5224,0.5029,0.5747,0.3707,0.5804,0.4776,0.5623,0.1049,0.2957,0.271,0.4438,0.5038,0.6173,0.4239,0.6397,0.3033,0.6528,1,0.3628,0.5125,0.5622,0.4564,0.4609,0.5459,0.7217,0.2521,0.4595,0.3287,0.5499,0.437,0.388,0.4969,0.4433,0.2075,0.5177,0.2971,0.2662,0.551,0.4361,0.4493,0.4931,0.6111,0.3619,0.7633,0.3498,0.5519],[0.0113,0.1212,-0.071,0.1562,0.0285,0.0087,0.1321,-0.1219,-0.1404,0.0113,0.0499,0.1264,-0.008,-0.1649,0.2151,0.2128,0.0519,0.2467,0.0802,0.1598,0.2585,0.3628,1,0.0935,0.2783,0.1491,0.0555,0.2252,0.2073,0.1793,0.0618,0.1826,0.058,0.211,0.1989,0.0632,0.1347,0.2071,0.3897,-0.002,-0.1827,0.1421,-0.0131,0.277,0.3205,0.0824,0.1164,0.0674,0.0876,0.0061],[0.3604,-0.4614,0.5503,0.6554,0.5118,0.5121,0.4024,0.2434,0.3504,0.3743,0.3263,0.2072,0.2584,0.3346,0.6898,0.6066,0.4031,0.6406,0.5452,0.1708,0.3658,0.5125,0.0935,1,0.3008,0.2786,0.24,0.6332,0.3374,0.7023,0.7435,0.6653,0.4788,0.5793,0.1632,0.2243,0.4417,0.7021,0.3907,0.6299,0.1183,0.8365,0.2013,0.5723,0.2253,0.4148,0.79,0.3734,0.7988,0.3239],[0.2899,-0.3148,0.0297,0.2955,0.5358,0.3211,0.9166,0.4404,0.2555,0.3089,0.1928,-0.1041,0.0509,0.3085,0.4628,0.5744,0.259,0.3377,0.2729,0.9177,0.9126,0.5622,0.2783,0.3008,1,0.8516,0.8227,0.1987,0.8509,0.1415,0.406,0.4293,0.1784,0.3653,0.7917,0.8506,0.2905,0.2384,0.3375,0.4262,0.1058,0.406,0.4824,0.4908,0.3505,0.64,0.4125,0.6313,0.3731,0.4198],[0.2954,-0.282,-0.1208,0.0805,0.4953,0.3885,0.888,0.2885,0.3575,0.3659,0.2492,0.0807,-0.1506,0.2591,0.2606,0.533,0.0026,0.2428,0.2636,0.7732,0.8171,0.4564,0.1491,0.2786,0.8516,1,0.9523,0.2363,0.8383,0.2209,0.3681,0.5189,0.2804,0.2884,0.8537,0.9462,0.4115,0.3142,0.2448,0.4849,0.0911,0.3647,0.3665,0.4603,0.2324,0.6947,0.4785,0.6618,0.4698,0.382],[0.2649,-0.2465,-0.0792,0.0904,0.4453,0.4002,0.8933,0.3184,0.3471,0.3521,0.1937,0.1027,-0.0645,0.3416,0.2248,0.5511,-0.0143,0.229,0.2333,0.7635,0.7999,0.4609,0.0555,0.24,0.8227,0.9523,1,0.184,0.8289,0.18,0.3912,0.4803,0.3029,0.2339,0.8234,0.9576,0.3776,0.2751,0.1966,0.4757,0.14,0.3504,0.3971,0.3881,0.1701,0.6757,0.4286,0.6579,0.434,0.4139],[0.3682,-0.2744,0.6075,0.5837,0.4202,0.6891,0.3295,0.0959,0.6219,0.8194,0.5706,0.3905,0.2285,0.0044,0.4197,0.4581,0.4413,0.5086,0.7879,-0.0092,0.3044,0.5459,0.2252,0.6332,0.1987,0.2363,0.184,1,0.3475,0.667,0.6318,0.6026,0.6555,0.5569,0.2836,0.2648,0.6519,0.5307,0.4851,0.5166,0.1298,0.7569,0.0562,0.5787,0.4387,0.5079,0.6745,0.5714,0.6648,0.4057],[0.4278,-0.2012,0.1886,0.3385,0.6201,0.4228,0.8921,0.4629,0.5396,0.4436,0.4922,0.0209,0.1229,0.2901,0.3862,0.6031,0.3472,0.2483,0.4416,0.7439,0.9312,0.7217,0.2073,0.3374,0.8509,0.8383,0.8289,0.3475,1,0.1725,0.3482,0.4071,0.449,0.4443,0.7319,0.8445,0.5745,0.2007,0.54,0.393,0.0841,0.4204,0.5308,0.5772,0.2704,0.7014,0.4039,0.777,0.3769,0.6717],[0.3006,-0.0966,0.3038,0.3187,0.4886,0.4215,0.2115,-0.0358,0.3224,0.4548,0.2924,0.2217,0.0211,0.0023,0.4261,0.4876,0.1577,0.5313,0.4628,0.1057,0.1059,0.2521,0.1793,0.7023,0.1415,0.2209,0.18,0.667,0.1725,1,0.6737,0.8185,0.371,0.4378,0.3691,0.227,0.5171,0.9278,0.4609,0.6679,0.2392,0.794,-0.0253,0.4694,0.3207,0.2872,0.8859,0.2376,0.8335,0.1811],[0.2722,-0.4149,0.4856,0.5428,0.4966,0.636,0.5172,0.2336,0.3716,0.453,0.2354,0.1567,0.25,0.2973,0.5444,0.71,0.3991,0.7907,0.626,0.2262,0.3461,0.4595,0.0618,0.7435,0.406,0.3681,0.3912,0.6318,0.3482,0.6737,1,0.7745,0.3221,0.735,0.3005,0.4299,0.5388,0.7444,0.4379,0.754,0.1756,0.9379,0.2487,0.702,0.4824,0.5728,0.7388,0.529,0.79,0.3001],[0.4235,-0.1928,0.243,0.2475,0.6215,0.4594,0.5379,0.0843,0.3643,0.5418,0.2439,0.1319,0.0944,0.0778,0.3916,0.6123,0.1786,0.7266,0.4646,0.3556,0.4123,0.3287,0.1826,0.6653,0.4293,0.5189,0.4803,0.6026,0.4071,0.8185,0.7745,1,0.2272,0.6235,0.5035,0.4618,0.6043,0.8174,0.4253,0.9151,0.1336,0.8396,0.146,0.7044,0.5169,0.4704,0.911,0.4405,0.9305,0.301],[0.0764,-0.2908,0.5423,0.4698,0.1844,0.5945,0.3489,0.3193,0.6971,0.5739,0.6296,0.3626,0.2234,0.2345,0.4057,0.4483,0.2354,0.0741,0.614,0.0507,0.3307,0.5499,0.058,0.4788,0.1784,0.2804,0.3029,0.6555,0.449,0.371,0.3221,0.2272,1,0.2846,0.1997,0.354,0.5321,0.3318,0.2937,0.2804,0.0343,0.4439,0.2458,0.3231,0.0243,0.3707,0.4472,0.5832,0.4163,0.58],[0.3962,-0.2866,0.5113,0.5311,0.5505,0.5233,0.4914,0.3879,0.4359,0.4223,0.3605,0.0096,0.4532,0.3051,0.4407,0.7686,0.5538,0.7412,0.5976,0.2278,0.4276,0.437,0.211,0.5793,0.3653,0.2884,0.2339,0.5569,0.4443,0.4378,0.735,0.6235,0.2846,1,0.0897,0.2738,0.6715,0.5173,0.5979,0.6484,-0.216,0.7364,0.4343,0.9572,0.4472,0.434,0.5583,0.4839,0.6512,0.5768],[0.2099,-0.1405,-0.162,0.0343,0.432,0.2267,0.7475,0.1346,0.2729,0.4061,0.2025,-0.0262,-0.2207,-0.0134,0.204,0.3381,0.0065,0.1612,0.1979,0.7428,0.699,0.388,0.1989,0.1632,0.7917,0.8537,0.8234,0.2836,0.7319,0.3691,0.3005,0.5035,0.1997,0.0897,1,0.8775,0.3315,0.3423,0.2696,0.3821,0.3095,0.3436,0.2016,0.2786,0.2642,0.6327,0.4699,0.5764,0.3778,0.2096],[0.2095,-0.2977,-0.0583,0.1308,0.4309,0.4223,0.8937,0.277,0.4257,0.3764,0.2336,0.0677,-0.1232,0.2326,0.2271,0.5351,0.0529,0.1883,0.2854,0.7768,0.7872,0.4969,0.0632,0.2243,0.8506,0.9462,0.9576,0.2648,0.8445,0.227,0.4299,0.4618,0.354,0.2738,0.8775,1,0.409,0.2943,0.2405,0.4436,0.1494,0.3924,0.3392,0.4194,0.2709,0.6832,0.4512,0.7106,0.415,0.4219],[0.4071,0.0775,0.3507,0.2417,0.5848,0.5712,0.4832,0.3132,0.6116,0.6167,0.5491,0.2352,0.1711,0.1758,0.4423,0.6504,0.2524,0.4644,0.6519,0.1804,0.4409,0.4433,0.1347,0.4417,0.2905,0.4115,0.3776,0.6519,0.5745,0.5171,0.5388,0.6043,0.5321,0.6715,0.3315,0.409,1,0.5125,0.7833,0.5265,-0.1393,0.6664,0.3229,0.7061,0.227,0.5854,0.5559,0.5629,0.6129,0.6392],[0.2643,-0.1078,0.2575,0.3092,0.5148,0.443,0.282,0.1254,0.2046,0.3412,0.2396,0.2001,0.0667,0.2275,0.5152,0.6342,0.116,0.5867,0.4507,0.1989,0.1509,0.2075,0.2071,0.7021,0.2384,0.3142,0.2751,0.5307,0.2007,0.9278,0.7444,0.8174,0.3318,0.5173,0.3423,0.2943,0.5125,1,0.4889,0.6999,0.1516,0.7933,0.1495,0.5432,0.2629,0.3296,0.8541,0.2469,0.8577,0.209],[0.4642,0.2722,0.3747,0.4045,0.6678,0.3903,0.3223,0.3431,0.3293,0.3737,0.5536,0.0725,0.2501,0.1915,0.548,0.5561,0.4685,0.4155,0.6103,0.2167,0.4101,0.5177,0.3897,0.3907,0.3375,0.2448,0.1966,0.4851,0.54,0.4609,0.4379,0.4253,0.2937,0.5979,0.2696,0.2405,0.7833,0.4889,1,0.2723,0.0059,0.5659,0.4107,0.606,0.282,0.5,0.38,0.4395,0.4031,0.5649],[0.3675,-0.3824,0.3178,0.2616,0.5672,0.5083,0.5795,0.2436,0.4631,0.5467,0.3186,0.136,0.1824,0.1987,0.3854,0.6695,0.2029,0.669,0.5128,0.3829,0.3999,0.2971,-0.002,0.6299,0.4262,0.4849,0.4757,0.5166,0.393,0.6679,0.754,0.9151,0.2804,0.6484,0.3821,0.4436,0.5265,0.6999,0.2723,1,0.0527,0.7797,0.2504,0.7161,0.5009,0.4127,0.8634,0.4376,0.8999,0.3209],[0.2599,0.0142,0.2494,0.1567,0.3677,0.2024,0.0538,0.1049,0.2148,0.2766,0.1912,-0.1272,-0.05,0.0642,0.1124,-0.1153,0.2317,0.1928,0.1908,-0.0125,0.1096,0.2662,-0.1827,0.1183,0.1058,0.0911,0.14,0.1298,0.0841,0.2392,0.1756,0.1336,0.0343,-0.216,0.3095,0.1494,-0.1393,0.1516,0.0059,0.0527,1,0.2161,0.1605,-0.237,0.028,0.3474,0.1268,0.3275,0.0453,-0.0217],[0.4068,-0.3348,0.5487,0.559,0.6284,0.6812,0.5284,0.2514,0.5055,0.5859,0.382,0.1945,0.2375,0.2384,0.6555,0.7203,0.4376,0.8039,0.6973,0.2331,0.4184,0.551,0.1421,0.8365,0.406,0.3647,0.3504,0.7569,0.4204,0.794,0.9379,0.8396,0.4439,0.7364,0.3436,0.3924,0.6664,0.7933,0.5659,0.7797,0.2161,1,0.248,0.7172,0.4659,0.5887,0.8501,0.5527,0.8559,0.4081],[0.358,0.0589,0.5183,0.4897,0.571,0.2661,0.4588,0.9467,0.2167,0.3684,0.4689,-0.3127,0.6627,0.7627,0.4226,0.5688,0.575,0.4041,0.4765,0.3436,0.5086,0.4361,-0.0131,0.2013,0.4824,0.3665,0.3971,0.0562,0.5308,-0.0253,0.2487,0.146,0.2458,0.4343,0.2016,0.3392,0.3229,0.1495,0.4107,0.2504,0.1605,0.248,1,0.4188,-0.0166,0.5672,0.074,0.5736,0.2094,0.5845],[0.406,-0.3296,0.4166,0.4704,0.5647,0.4683,0.6039,0.3607,0.4285,0.4675,0.4162,0.0337,0.3934,0.2214,0.4001,0.7622,0.4937,0.6687,0.5864,0.3635,0.5474,0.4493,0.277,0.5723,0.4908,0.4603,0.3881,0.5787,0.5772,0.4694,0.702,0.7044,0.3231,0.9572,0.2786,0.4194,0.7061,0.5432,0.606,0.7161,-0.237,0.7172,0.4188,1,0.4818,0.4802,0.6235,0.5504,0.6989,0.564],[0.1892,-0.114,0.3064,0.3779,0.295,0.2106,0.2697,-0.1213,0.2856,0.33,0.257,-0.0405,0.1417,-0.2612,0.0372,0.1882,0.4613,0.3723,0.437,0.1874,0.2156,0.4931,0.3205,0.2253,0.3505,0.2324,0.1701,0.4387,0.2704,0.3207,0.4824,0.5169,0.0243,0.4472,0.2642,0.2709,0.227,0.2629,0.282,0.5009,0.028,0.4659,-0.0166,0.4818,1,0.209,0.4184,0.4344,0.4066,0.1702],[0.2926,-0.1842,0.3514,0.4404,0.5733,0.5993,0.7044,0.4962,0.4904,0.6315,0.4756,-0.0402,0.1701,0.3697,0.5513,0.5346,0.3711,0.5421,0.6484,0.3869,0.6701,0.6111,0.0824,0.4148,0.64,0.6947,0.6757,0.5079,0.7014,0.2872,0.5728,0.4704,0.3707,0.434,0.6327,0.6832,0.5854,0.3296,0.5,0.4127,0.3474,0.5887,0.5672,0.4802,0.209,1,0.3989,0.8367,0.4644,0.454],[0.36,-0.3403,0.2973,0.3422,0.5821,0.5635,0.5246,0.0844,0.4936,0.5345,0.3156,0.2321,0.0177,0.111,0.5334,0.6309,0.157,0.5812,0.498,0.3843,0.3981,0.3619,0.1164,0.79,0.4125,0.4785,0.4286,0.6745,0.4039,0.8859,0.7388,0.911,0.4472,0.5583,0.4699,0.4512,0.5559,0.8541,0.38,0.8634,0.1268,0.8501,0.074,0.6235,0.4184,0.3989,1,0.4039,0.953,0.3619],[0.2729,-0.2191,0.5478,0.5619,0.5271,0.5458,0.692,0.4794,0.6632,0.6801,0.5974,-0.108,0.3365,0.257,0.3674,0.4626,0.5571,0.4071,0.6878,0.3636,0.697,0.7633,0.0674,0.3734,0.6313,0.6618,0.6579,0.5714,0.777,0.2376,0.529,0.4405,0.5832,0.4839,0.5764,0.7106,0.5629,0.2469,0.4395,0.4376,0.3275,0.5527,0.5736,0.5504,0.4344,0.8367,0.4039,1,0.4444,0.6716],[0.3541,-0.2763,0.3813,0.4089,0.5929,0.56,0.5028,0.2002,0.4398,0.5576,0.3013,0.182,0.1726,0.2633,0.534,0.6924,0.2054,0.6909,0.5482,0.3107,0.3702,0.3498,0.0876,0.7988,0.3731,0.4698,0.434,0.6648,0.3769,0.8335,0.79,0.9305,0.4163,0.6512,0.3778,0.415,0.6129,0.8577,0.4031,0.8999,0.0453,0.8559,0.2094,0.6989,0.4066,0.4644,0.953,0.4444,1,0.4029],[0.3859,0.0775,0.5226,0.4598,0.5781,0.5017,0.5238,0.523,0.6503,0.4932,0.4876,-0.0935,0.4494,0.4045,0.4156,0.5422,0.4718,0.2795,0.4901,0.3524,0.6002,0.5519,0.0061,0.3239,0.4198,0.382,0.4139,0.4057,0.6717,0.1811,0.3001,0.301,0.58,0.5768,0.2096,0.4219,0.6392,0.209,0.5649,0.3209,-0.0217,0.4081,0.5845,0.564,0.1702,0.454,0.3619,0.6716,0.4029,1]];
scoreMat = [[31.76,32.88,30.91,29.35,25.27,19.51,24.53,22.86,18.24,21.16,15.94,13.06,23.08,18.72,1.92,14.47,23.58,15.2,15.61,18.33,17.23,16.93,6.27,11.24,18.18,15.52,17.66,14.27,15.65,12.55,17.34,12.79,12.78,13.89,14.69,16.88,9.5,10.32,10.81,10.41,17.18,12.07,14.13,11.74,16.71,7.43,8.69,16.55,9.14,15.08],[32.69,30.41,27.81,23.54,24.53,19.86,13.84,19.9,20.33,23.3,23.3,18.19,20.12,12.97,6.54,8.7,18.8,7.62,17.45,13.43,11.49,13.03,5.71,8.46,10.63,9.5,7.4,14.32,10.96,11.06,5.33,8.42,11.44,7.85,8.85,6.56,9.05,6.15,11.5,9.91,12.59,5.56,10.47,8.99,10.26,4.93,7.06,6.54,7.44,11],[33.16,27.98,30.23,29.43,28.84,30.83,27.71,19.68,25.44,28.8,22.67,26.36,16.94,19.16,25.22,16.31,16.65,17.34,20.43,17.37,19.76,19.78,17.58,26.56,18.49,22.27,24.19,21.28,17.9,22.78,17.83,22.01,23.52,2.14,22.63,18.93,11.89,20.23,11.69,17.02,37.62,17.08,10.64,4.18,12.63,20.09,20.73,18.88,17.46,10.39],[29.08,34.32,15.63,13.05,24.75,14.35,22.39,13.07,15.12,14.71,15.09,17.76,12.16,12.08,2.24,16.12,8.68,12.1,10.39,20.27,12.47,12.24,14.33,8.85,15.52,19.67,18.52,8.6,12.95,19.99,11.65,20.22,7.12,7.59,20.47,16.33,11.78,20.11,11.4,16.12,13.1,9.78,6.31,9.62,9.4,3.33,14.84,0.85,12.2,4.53],[35.05,36.8,30.81,27.24,31.63,24.11,17.57,21.99,25.66,22.37,19.98,16.57,19.1,18.84,17.73,17.36,20.16,22.58,17.56,14.33,14.07,17.9,21.68,20.54,13.29,12.08,7.17,16.17,11.55,17.67,12.98,19.91,13.62,18.52,7.53,7.53,12.89,16.06,12.97,18.37,12.87,13.76,13.3,14.51,17.65,4.04,16.72,9.53,16.36,14.48],[24.99,42.97,27.53,27.16,22.66,19.52,22.7,15.99,20.86,21.11,17.96,15.52,20.25,11.2,26.72,17.7,17.09,15.9,16.28,19.63,17.01,17.41,30.8,14.96,17.07,11.62,12.73,17.19,16.16,19.24,14.14,18.59,16.79,14.46,15.03,11.91,18.18,16.34,18.94,14.75,0,14.02,8.55,13.65,16.86,6.41,15.95,9.03,14.19,15.8],[33.69,30.15,29.69,31.85,32.79,27.17,26.93,27.77,23.54,20.9,21.42,15.92,18.22,21.01,39.06,20.43,22.42,17.99,18.55,20.16,23.79,18.66,23.53,25.27,20.02,17.2,13.86,18.63,20.94,19.01,15.79,11.89,19.38,18.6,16.87,14.81,17.93,18.14,21.77,4.89,20.6,15.98,16.77,15.55,7.9,17.77,14.11,14.99,11.43,17.76],[24.83,14.93,30.43,28.78,19.73,19.93,23.75,22.92,27.83,21.65,21.52,15.07,18.58,12.34,9.87,14.24,21.38,12.75,17.45,16.48,17.31,19.1,16.67,11.2,16.29,12.16,11.65,15.83,16.34,11.64,11.44,6.05,18.89,10.59,14.49,13.32,7.84,5.24,9.48,9.49,20.9,9.78,11.47,9.85,14.2,8.92,6.82,14.87,5.19,8.67],[30.94,41.9,29.48,28.25,24.3,23.62,16.92,16.34,25.25,22.04,24.93,22.82,18.78,14.54,3.56,15.55,20.43,12,21.17,9.72,11.39,19.19,18.42,14.01,10.89,15.47,13.97,18.56,17.27,15.69,13.15,11.32,19.07,15,12.54,13.6,21.95,12.07,19.02,6.47,5.1,11.39,10.86,13.47,14.95,15.27,7.81,16.41,9.64,13.25],[33.33,33.04,28.3,28.62,34.47,34.89,35.13,27.6,30.51,26.36,22.49,15.6,18.39,23.07,20.12,23.8,20.52,21.95,21.28,23.01,23.22,19.14,17.58,11.56,23.17,25.25,27.31,16.75,22.89,17.61,20.7,21,14.06,20.87,22.15,23.06,16.15,17.57,17.42,19.85,27.37,15.61,21.32,17.71,17.86,24.99,16.02,23.42,15.56,18.55],[40.26,37.52,30.27,27.09,33.64,30.58,27.32,36.56,21.36,25.47,23.46,30.87,21.71,30.05,21.97,26.9,19.81,20.27,21.48,18.78,20.21,18.31,22.35,15.31,17.83,15.52,17.47,16.65,18.55,14.81,16.12,14.48,16.91,17.76,12.43,13.49,19.28,15.38,19.44,14.67,10.64,13.85,21.57,15.47,10.57,11.47,10.12,9.66,11.55,13.67],[33.6,14.41,29.39,29.49,27.89,35.48,33.06,17.86,28.47,32.67,20.57,27.71,18.33,15.36,21.31,24.89,18.6,26.32,22.46,18.52,20.03,17.01,21.07,26.94,18.79,21.44,19.14,28.29,17.1,24.43,25.84,29.94,20.33,27.56,19.35,18.33,23,22.41,14.7,26.3,2.84,21.98,7,25.26,17.32,18.48,25.27,15.99,23.28,12.43],[31.93,9.73,30.28,29.58,28.05,33.81,33.91,19.21,27.49,20.05,22.67,25.99,18.62,17.92,28.99,24.87,19.84,22.56,21.35,18.65,21.84,19.42,21.03,31.17,19.63,20.48,20.93,18.73,20.32,19.28,27.63,25.17,20.67,26.88,12.64,17.49,18.38,21.23,16.72,25.73,9.46,21.87,11.48,24.18,17.76,14.23,21.35,17.51,19.04,14.88],[32.99,31.18,25.53,27.4,28.98,18.58,25.66,18.54,12.82,19.34,15.98,13.29,18.09,14.54,21.54,14.64,21.4,25.97,16.69,18.47,19.53,17.82,25.18,17.19,20.35,16.65,14.65,13.94,15.05,13.79,18.66,18.52,0,14.52,16.91,12.58,10.23,11.09,15.78,13.07,20.37,14.77,11.69,12.91,18.08,19.05,9.13,10.66,10.1,4.22],[25.25,31.64,29.55,25.11,20.33,27.56,6.82,17.55,20.65,18.23,16.05,20.9,17.32,19.55,22.05,14.51,15.47,17.47,17.31,7.61,0,13.34,0,16.06,4.98,0,0,14.32,0.81,17.08,18.22,8.55,11.98,9.56,2.31,3.17,9.89,15.33,10.75,9.08,18.59,13.9,5.96,2.94,8.72,4.89,9.47,0,9.89,6.14],[30.52,35.84,26.8,19.55,27.47,29.5,31.88,27.05,30.32,27.93,20.52,16.75,17.78,17.1,20.65,17.47,15.16,20.16,18.15,16.63,23.16,17.1,12.24,8.85,17.38,20.51,21.15,16.41,19.84,11.39,14.18,15.84,18.41,14.94,16.77,17.43,23.59,8.3,15.07,13.86,17.43,13.55,16.95,13.24,9.41,23.13,9.3,21.38,10.14,17.52],[31.62,29.92,32.18,30.95,24.76,21.21,22.38,27.55,18.19,20.99,17.15,13.69,25.84,24.54,15.14,20.8,21.08,25.83,15.56,13.52,16.1,16.83,17.19,22.89,12.83,11.92,14.35,14.56,13.62,16.7,16.7,17.16,14.09,20.17,9.22,9.62,12.06,14.89,11.96,15.03,13.58,13.8,19.7,16.23,7.41,11.84,11.66,10,13.94,13.88],[21.49,29.76,24.29,25.02,18.83,19.35,19.52,18.19,15.51,16.54,13.64,17.16,16.93,20.21,16.22,15.55,12.1,16.44,13.81,15.59,14.39,15.11,21.05,13.62,14.67,16.82,16.68,13.65,11.88,12.94,14.81,12.69,14.21,11.07,12.09,13.44,10.78,14.51,11.04,10.02,8.26,9.35,10.42,10.97,10.03,11.36,8.73,10.9,12.38,9.26],[24.35,26.6,28.26,28.92,20.13,22.99,23.99,33.74,15.59,21.31,21.37,15.09,22.21,23.33,24.5,19.27,19.32,15.55,19.27,17.48,15.01,16.15,24.24,11.9,18.9,17.6,15.3,12.91,13.67,11.62,13.23,8.54,16.51,13.06,13.78,14.26,9.3,13.17,11.66,9.88,3.41,9.89,21.87,11.8,11.94,15.13,7.42,13.24,8.89,9.63],[32.14,20.56,30.54,31.01,33.69,26.85,39.74,38.04,27.92,23.91,21.45,14.28,20.62,29.03,35.04,25.57,20.31,19.74,19.88,25.53,26.66,18.64,0,28.73,24.04,26.33,27.67,15.26,24.93,16.53,20.27,21.33,18.81,19.77,19.24,22,20.19,16.37,15.18,24.31,6.9,17.12,22.99,18.41,11.81,22.36,19.82,19.61,20.18,18.18],[27.91,26.35,17.19,18.76,16.9,22.5,24.88,10.85,18.33,13.18,12.53,25.57,12.51,16.94,5.71,13.16,9.25,6.53,9.49,19.57,17.56,15.44,17.32,5.33,16.3,16.63,17.14,8.48,14.08,5.04,3.67,0,10.08,2.54,11.48,14.08,3.55,0,5.8,0,0,2.63,2.82,1.78,7.87,2.31,2.52,0.7,1.91,8.6]];
```
