## Supplemental Examples for "R/LinkedCharts: A novel approach for simple but powerful interactive data analysis": histDens.js

```
var hist = {"value":{"1":{"1":{"1":[1]}},"2":{"1":{"1":[8]}},"3":{"1":{"1":[43]}},"4":{"1":{"1":[121]}},"5":{"1":{"1":[232]}},"6":{"1":{"1":[266]}},"7":{"1":{"1":[186]}},"8":{"1":{"1":[110]}},"9":{"1":{"1":[26]}},"10":{"1":{"1":[7]}},"__inds__":[-3.386,-2.6944,-2.0027,-1.3111,-0.6195,0.0721,0.7638,1.4554,2.147,2.8386]},"contScaleX":[true],"addColourScaleToLegend":[false],"groupWidth":[1],"groupIds":[-3.386,-2.6944,-2.0027,-1.3111,-0.6195,0.0721,0.7638,1.4554,2.147,2.8386],"barIds":[1],"stackIds":[1]};
var dens = {"x":[[-4.4153],[-4.3991],[-4.3829],[-4.3667],[-4.3505],[-4.3343],[-4.318],[-4.3018],[-4.2856],[-4.2694],[-4.2532],[-4.237],[-4.2208],[-4.2046],[-4.1884],[-4.1722],[-4.1559],[-4.1397],[-4.1235],[-4.1073],[-4.0911],[-4.0749],[-4.0587],[-4.0425],[-4.0263],[-4.0101],[-3.9938],[-3.9776],[-3.9614],[-3.9452],[-3.929],[-3.9128],[-3.8966],[-3.8804],[-3.8642],[-3.848],[-3.8318],[-3.8155],[-3.7993],[-3.7831],[-3.7669],[-3.7507],[-3.7345],[-3.7183],[-3.7021],[-3.6859],[-3.6697],[-3.6534],[-3.6372],[-3.621],[-3.6048],[-3.5886],[-3.5724],[-3.5562],[-3.54],[-3.5238],[-3.5076],[-3.4913],[-3.4751],[-3.4589],[-3.4427],[-3.4265],[-3.4103],[-3.3941],[-3.3779],[-3.3617],[-3.3455],[-3.3292],[-3.313],[-3.2968],[-3.2806],[-3.2644],[-3.2482],[-3.232],[-3.2158],[-3.1996],[-3.1834],[-3.1671],[-3.1509],[-3.1347],[-3.1185],[-3.1023],[-3.0861],[-3.0699],[-3.0537],[-3.0375],[-3.0213],[-3.005],[-2.9888],[-2.9726],[-2.9564],[-2.9402],[-2.924],[-2.9078],[-2.8916],[-2.8754],[-2.8592],[-2.8429],[-2.8267],[-2.8105],[-2.7943],[-2.7781],[-2.7619],[-2.7457],[-2.7295],[-2.7133],[-2.6971],[-2.6808],[-2.6646],[-2.6484],[-2.6322],[-2.616],[-2.5998],[-2.5836],[-2.5674],[-2.5512],[-2.535],[-2.5187],[-2.5025],[-2.4863],[-2.4701],[-2.4539],[-2.4377],[-2.4215],[-2.4053],[-2.3891],[-2.3729],[-2.3567],[-2.3404],[-2.3242],[-2.308],[-2.2918],[-2.2756],[-2.2594],[-2.2432],[-2.227],[-2.2108],[-2.1946],[-2.1783],[-2.1621],[-2.1459],[-2.1297],[-2.1135],[-2.0973],[-2.0811],[-2.0649],[-2.0487],[-2.0325],[-2.0162],[-2],[-1.9838],[-1.9676],[-1.9514],[-1.9352],[-1.919],[-1.9028],[-1.8866],[-1.8704],[-1.8541],[-1.8379],[-1.8217],[-1.8055],[-1.7893],[-1.7731],[-1.7569],[-1.7407],[-1.7245],[-1.7083],[-1.692],[-1.6758],[-1.6596],[-1.6434],[-1.6272],[-1.611],[-1.5948],[-1.5786],[-1.5624],[-1.5462],[-1.5299],[-1.5137],[-1.4975],[-1.4813],[-1.4651],[-1.4489],[-1.4327],[-1.4165],[-1.4003],[-1.3841],[-1.3678],[-1.3516],[-1.3354],[-1.3192],[-1.303],[-1.2868],[-1.2706],[-1.2544],[-1.2382],[-1.222],[-1.2057],[-1.1895],[-1.1733],[-1.1571],[-1.1409],[-1.1247],[-1.1085],[-1.0923],[-1.0761],[-1.0599],[-1.0436],[-1.0274],[-1.0112],[-0.995],[-0.9788],[-0.9626],[-0.9464],[-0.9302],[-0.914],[-0.8978],[-0.8816],[-0.8653],[-0.8491],[-0.8329],[-0.8167],[-0.8005],[-0.7843],[-0.7681],[-0.7519],[-0.7357],[-0.7195],[-0.7032],[-0.687],[-0.6708],[-0.6546],[-0.6384],[-0.6222],[-0.606],[-0.5898],[-0.5736],[-0.5574],[-0.5411],[-0.5249],[-0.5087],[-0.4925],[-0.4763],[-0.4601],[-0.4439],[-0.4277],[-0.4115],[-0.3953],[-0.379],[-0.3628],[-0.3466],[-0.3304],[-0.3142],[-0.298],[-0.2818],[-0.2656],[-0.2494],[-0.2332],[-0.2169],[-0.2007],[-0.1845],[-0.1683],[-0.1521],[-0.1359],[-0.1197],[-0.1035],[-0.0873],[-0.0711],[-0.0548],[-0.0386],[-0.0224],[-0.0062],[0.01],[0.0262],[0.0424],[0.0586],[0.0748],[0.091],[0.1073],[0.1235],[0.1397],[0.1559],[0.1721],[0.1883],[0.2045],[0.2207],[0.2369],[0.2531],[0.2694],[0.2856],[0.3018],[0.318],[0.3342],[0.3504],[0.3666],[0.3828],[0.399],[0.4152],[0.4315],[0.4477],[0.4639],[0.4801],[0.4963],[0.5125],[0.5287],[0.5449],[0.5611],[0.5773],[0.5935],[0.6098],[0.626],[0.6422],[0.6584],[0.6746],[0.6908],[0.707],[0.7232],[0.7394],[0.7556],[0.7719],[0.7881],[0.8043],[0.8205],[0.8367],[0.8529],[0.8691],[0.8853],[0.9015],[0.9177],[0.934],[0.9502],[0.9664],[0.9826],[0.9988],[1.015],[1.0312],[1.0474],[1.0636],[1.0798],[1.0961],[1.1123],[1.1285],[1.1447],[1.1609],[1.1771],[1.1933],[1.2095],[1.2257],[1.2419],[1.2582],[1.2744],[1.2906],[1.3068],[1.323],[1.3392],[1.3554],[1.3716],[1.3878],[1.404],[1.4203],[1.4365],[1.4527],[1.4689],[1.4851],[1.5013],[1.5175],[1.5337],[1.5499],[1.5661],[1.5824],[1.5986],[1.6148],[1.631],[1.6472],[1.6634],[1.6796],[1.6958],[1.712],[1.7282],[1.7445],[1.7607],[1.7769],[1.7931],[1.8093],[1.8255],[1.8417],[1.8579],[1.8741],[1.8903],[1.9066],[1.9228],[1.939],[1.9552],[1.9714],[1.9876],[2.0038],[2.02],[2.0362],[2.0524],[2.0686],[2.0849],[2.1011],[2.1173],[2.1335],[2.1497],[2.1659],[2.1821],[2.1983],[2.2145],[2.2307],[2.247],[2.2632],[2.2794],[2.2956],[2.3118],[2.328],[2.3442],[2.3604],[2.3766],[2.3928],[2.4091],[2.4253],[2.4415],[2.4577],[2.4739],[2.4901],[2.5063],[2.5225],[2.5387],[2.5549],[2.5712],[2.5874],[2.6036],[2.6198],[2.636],[2.6522],[2.6684],[2.6846],[2.7008],[2.717],[2.7333],[2.7495],[2.7657],[2.7819],[2.7981],[2.8143],[2.8305],[2.8467],[2.8629],[2.8791],[2.8954],[2.9116],[2.9278],[2.944],[2.9602],[2.9764],[2.9926],[3.0088],[3.025],[3.0412],[3.0575],[3.0737],[3.0899],[3.1061],[3.1223],[3.1385],[3.1547],[3.1709],[3.1871],[3.2033],[3.2196],[3.2358],[3.252],[3.2682],[3.2844],[3.3006],[3.3168],[3.333],[3.3492],[3.3654],[3.3816],[3.3979],[3.4141],[3.4303],[3.4465],[3.4627],[3.4789],[3.4951],[3.5113],[3.5275],[3.5437],[3.56],[3.5762],[3.5924],[3.6086],[3.6248],[3.641],[3.6572],[3.6734],[3.6896],[3.7058],[3.7221],[3.7383],[3.7545],[3.7707],[3.7869],[3.8031],[3.8193],[3.8355],[3.8517],[3.8679]],"y":[[0],[0],[0],[0],[0],[0.0001],[0.0001],[0.0001],[0.0001],[0.0001],[0.0001],[0.0002],[0.0002],[0.0002],[0.0002],[0.0003],[0.0003],[0.0004],[0.0004],[0.0005],[0.0005],[0.0006],[0.0006],[0.0007],[0.0008],[0.0008],[0.0009],[0.001],[0.0011],[0.0011],[0.0012],[0.0013],[0.0013],[0.0014],[0.0015],[0.0015],[0.0016],[0.0016],[0.0017],[0.0017],[0.0017],[0.0017],[0.0017],[0.0017],[0.0017],[0.0017],[0.0017],[0.0016],[0.0016],[0.0016],[0.0015],[0.0014],[0.0014],[0.0013],[0.0012],[0.0012],[0.0011],[0.001],[0.001],[0.0009],[0.0008],[0.0008],[0.0007],[0.0007],[0.0006],[0.0006],[0.0006],[0.0005],[0.0005],[0.0005],[0.0006],[0.0006],[0.0006],[0.0007],[0.0007],[0.0008],[0.0009],[0.001],[0.0011],[0.0013],[0.0014],[0.0016],[0.0018],[0.002],[0.0022],[0.0025],[0.0027],[0.003],[0.0033],[0.0037],[0.004],[0.0044],[0.0047],[0.0051],[0.0055],[0.0059],[0.0064],[0.0068],[0.0072],[0.0077],[0.0081],[0.0086],[0.0091],[0.0095],[0.01],[0.0105],[0.011],[0.0115],[0.0119],[0.0124],[0.0129],[0.0135],[0.014],[0.0145],[0.0151],[0.0157],[0.0163],[0.0169],[0.0176],[0.0183],[0.0191],[0.0199],[0.0208],[0.0217],[0.0226],[0.0237],[0.0248],[0.026],[0.0272],[0.0285],[0.0299],[0.0314],[0.0329],[0.0345],[0.0362],[0.038],[0.0398],[0.0417],[0.0436],[0.0456],[0.0476],[0.0497],[0.0518],[0.054],[0.0561],[0.0583],[0.0605],[0.0627],[0.065],[0.0672],[0.0694],[0.0716],[0.0738],[0.0759],[0.0781],[0.0802],[0.0824],[0.0845],[0.0865],[0.0886],[0.0907],[0.0927],[0.0947],[0.0967],[0.0988],[0.1008],[0.1028],[0.1048],[0.1068],[0.1089],[0.1109],[0.113],[0.1151],[0.1173],[0.1195],[0.1217],[0.124],[0.1264],[0.1288],[0.1313],[0.1339],[0.1365],[0.1392],[0.142],[0.145],[0.148],[0.1511],[0.1544],[0.1577],[0.1612],[0.1648],[0.1685],[0.1723],[0.1762],[0.1803],[0.1845],[0.1888],[0.1932],[0.1977],[0.2022],[0.2069],[0.2117],[0.2165],[0.2214],[0.2263],[0.2312],[0.2362],[0.2411],[0.2461],[0.251],[0.2559],[0.2607],[0.2655],[0.2701],[0.2747],[0.2792],[0.2836],[0.2879],[0.292],[0.296],[0.2999],[0.3037],[0.3073],[0.3108],[0.3141],[0.3173],[0.3204],[0.3234],[0.3262],[0.329],[0.3316],[0.3341],[0.3366],[0.339],[0.3413],[0.3436],[0.3459],[0.3481],[0.3503],[0.3525],[0.3546],[0.3568],[0.359],[0.3612],[0.3635],[0.3657],[0.3679],[0.3702],[0.3724],[0.3746],[0.3768],[0.379],[0.3811],[0.3831],[0.385],[0.3869],[0.3886],[0.3901],[0.3914],[0.3926],[0.3936],[0.3945],[0.395],[0.3953],[0.3954],[0.3953],[0.3949],[0.3943],[0.3935],[0.3924],[0.3911],[0.3897],[0.388],[0.3862],[0.3842],[0.3821],[0.3799],[0.3776],[0.3752],[0.3728],[0.3703],[0.3677],[0.3652],[0.3627],[0.3601],[0.3576],[0.3551],[0.3526],[0.3501],[0.3477],[0.3452],[0.3428],[0.3404],[0.338],[0.3356],[0.3332],[0.3308],[0.3284],[0.3259],[0.3235],[0.3209],[0.3184],[0.3158],[0.3131],[0.3104],[0.3077],[0.3049],[0.3021],[0.2992],[0.2963],[0.2934],[0.2905],[0.2876],[0.2846],[0.2817],[0.2788],[0.2759],[0.2731],[0.2703],[0.2676],[0.2649],[0.2623],[0.2598],[0.2573],[0.255],[0.2527],[0.2504],[0.2483],[0.2462],[0.2441],[0.2421],[0.2401],[0.2382],[0.2362],[0.2342],[0.2322],[0.2301],[0.228],[0.2258],[0.2236],[0.2212],[0.2187],[0.2162],[0.2135],[0.2107],[0.2078],[0.2048],[0.2016],[0.1984],[0.1951],[0.1917],[0.1883],[0.1847],[0.1812],[0.1776],[0.174],[0.1705],[0.1669],[0.1634],[0.1599],[0.1565],[0.1531],[0.1499],[0.1467],[0.1436],[0.1406],[0.1377],[0.1348],[0.1321],[0.1294],[0.1268],[0.1242],[0.1217],[0.1192],[0.1168],[0.1144],[0.112],[0.1096],[0.1072],[0.1048],[0.1023],[0.0999],[0.0974],[0.0949],[0.0923],[0.0897],[0.0871],[0.0845],[0.0819],[0.0792],[0.0765],[0.0739],[0.0712],[0.0685],[0.0659],[0.0633],[0.0608],[0.0583],[0.0558],[0.0534],[0.051],[0.0488],[0.0466],[0.0444],[0.0424],[0.0404],[0.0386],[0.0368],[0.0351],[0.0335],[0.032],[0.0306],[0.0293],[0.0281],[0.027],[0.0259],[0.0249],[0.024],[0.0232],[0.0225],[0.0218],[0.0211],[0.0206],[0.02],[0.0196],[0.0191],[0.0187],[0.0183],[0.018],[0.0176],[0.0173],[0.0169],[0.0166],[0.0162],[0.0159],[0.0155],[0.0152],[0.0148],[0.0144],[0.0139],[0.0135],[0.013],[0.0126],[0.0121],[0.0116],[0.0111],[0.0106],[0.0101],[0.0096],[0.0091],[0.0086],[0.0081],[0.0076],[0.0072],[0.0067],[0.0063],[0.0059],[0.0055],[0.0052],[0.0049],[0.0046],[0.0043],[0.004],[0.0038],[0.0036],[0.0034],[0.0032],[0.003],[0.0028],[0.0027],[0.0025],[0.0024],[0.0023],[0.0022],[0.002],[0.0019],[0.0018],[0.0017],[0.0016],[0.0015],[0.0014],[0.0013],[0.0012],[0.0011],[0.001],[0.0009],[0.0009],[0.0008],[0.0007],[0.0006],[0.0006],[0.0005],[0.0005],[0.0004],[0.0004],[0.0003],[0.0003],[0.0002],[0.0002],[0.0002],[0.0002],[0.0001],[0.0001],[0.0001],[0.0001],[0.0001],[0.0001],[0],[0],[0],[0],[0]],"nelements":[1],"nsteps":[512]};

		turnToScalar = function(v) {
			if(!v || (v.length === undefined && typeof v !== "object") || typeof v === "string") return v;
			if(Array.isArray(v)) {
				if(v.length == 1) return v[0];
				for(var i = 0; i < v.length; i++)
					v[i] = turnToScalar(v[i]);
			} else {
				var keys = Object.keys(v);
				for(var i = 0; i < keys.length; i++)
					v[keys[i]] = turnToScalar(v[keys[i]]);
			}
			return v;
		}

hist = turnToScalar(hist);
dens = turnToScalar(dens);
```
