## Supplemental Examples for "R/LinkedCharts: A novel approach for simple but powerful interactive data analysis": iris.js

```
var iris = [
    {"sepal_length": "5.1", "sepal_width": "3.5", "petal_length": "1.4", "petal_width": "0.2", "species": "setosa"},
    {"sepal_length": "4.9", "sepal_width": "3.0", "petal_length": "1.4", "petal_width": "0.2", "species": "setosa"},
    {"sepal_length": "4.7", "sepal_width": "3.2", "petal_length": "1.3", "petal_width": "0.2", "species": "setosa"},
    {"sepal_length": "4.6", "sepal_width": "3.1", "petal_length": "1.5", "petal_width": "0.2", "species": "setosa"},
    {"sepal_length": "5.0", "sepal_width": "3.6", "petal_length": "1.4", "petal_width": "0.2", "species": "setosa"},
    {"sepal_length": "5.4", "sepal_width": "3.9", "petal_length": "1.7", "petal_width": "0.4", "species": "setosa"},
    {"sepal_length": "4.6", "sepal_width": "3.4", "petal_length": "1.4", "petal_width": "0.3", "species": "setosa"},
    {"sepal_length": "5.0", "sepal_width": "3.4", "petal_length": "1.5", "petal_width": "0.2", "species": "setosa"},
    {"sepal_length": "4.4", "sepal_width": "2.9", "petal_length": "1.4", "petal_width": "0.2", "species": "setosa"},
    {"sepal_length": "4.9", "sepal_width": "3.1", "petal_length": "1.5", "petal_width": "0.1", "species": "setosa"},
    {"sepal_length": "5.4", "sepal_width": "3.7", "petal_length": "1.5", "petal_width": "0.2", "species": "setosa"},
    {"sepal_length": "4.8", "sepal_width": "3.4", "petal_length": "1.6", "petal_width": "0.2", "species": "setosa"},
    {"sepal_length": "4.8", "sepal_width": "3.0", "petal_length": "1.4", "petal_width": "0.1", "species": "setosa"},
    {"sepal_length": "4.3", "sepal_width": "3.0", "petal_length": "1.1", "petal_width": "0.1", "species": "setosa"},
    {"sepal_length": "5.8", "sepal_width": "4.0", "petal_length": "1.2", "petal_width": "0.2", "species": "setosa"},
    {"sepal_length": "5.7", "sepal_width": "4.4", "petal_length": "1.5", "petal_width": "0.4", "species": "setosa"},
    {"sepal_length": "5.4", "sepal_width": "3.9", "petal_length": "1.3", "petal_width": "0.4", "species": "setosa"},
    {"sepal_length": "5.1", "sepal_width": "3.5", "petal_length": "1.4", "petal_width": "0.3", "species": "setosa"},
    {"sepal_length": "5.7", "sepal_width": "3.8", "petal_length": "1.7", "petal_width": "0.3", "species": "setosa"},
    {"sepal_length": "5.1", "sepal_width": "3.8", "petal_length": "1.5", "petal_width": "0.3", "species": "setosa"},
    {"sepal_length": "5.4", "sepal_width": "3.4", "petal_length": "1.7", "petal_width": "0.2", "species": "setosa"},
    {"sepal_length": "5.1", "sepal_width": "3.7", "petal_length": "1.5", "petal_width": "0.4", "species": "setosa"},
    {"sepal_length": "4.6", "sepal_width": "3.6", "petal_length": "1.0", "petal_width": "0.2", "species": "setosa"},
    {"sepal_length": "5.1", "sepal_width": "3.3", "petal_length": "1.7", "petal_width": "0.5", "species": "setosa"},
    {"sepal_length": "4.8", "sepal_width": "3.4", "petal_length": "1.9", "petal_width": "0.2", "species": "setosa"},
    {"sepal_length": "5.0", "sepal_width": "3.0", "petal_length": "1.6", "petal_width": "0.2", "species": "setosa"},
    {"sepal_length": "5.0", "sepal_width": "3.4", "petal_length": "1.6", "petal_width": "0.4", "species": "setosa"},
    {"sepal_length": "5.2", "sepal_width": "3.5", "petal_length": "1.5", "petal_width": "0.2", "species": "setosa"},
    {"sepal_length": "5.2", "sepal_width": "3.4", "petal_length": "1.4", "petal_width": "0.2", "species": "setosa"},
    {"sepal_length": "4.7", "sepal_width": "3.2", "petal_length": "1.6", "petal_width": "0.2", "species": "setosa"},
    {"sepal_length": "4.8", "sepal_width": "3.1", "petal_length": "1.6", "petal_width": "0.2", "species": "setosa"},
    {"sepal_length": "5.4", "sepal_width": "3.4", "petal_length": "1.5", "petal_width": "0.4", "species": "setosa"},
    {"sepal_length": "5.2", "sepal_width": "4.1", "petal_length": "1.5", "petal_width": "0.1", "species": "setosa"},
    {"sepal_length": "5.5", "sepal_width": "4.2", "petal_length": "1.4", "petal_width": "0.2", "species": "setosa"},
    {"sepal_length": "4.9", "sepal_width": "3.1", "petal_length": "1.5", "petal_width": "0.1", "species": "setosa"},
    {"sepal_length": "5.0", "sepal_width": "3.2", "petal_length": "1.2", "petal_width": "0.2", "species": "setosa"},
    {"sepal_length": "5.5", "sepal_width": "3.5", "petal_length": "1.3", "petal_width": "0.2", "species": "setosa"},
    {"sepal_length": "4.9", "sepal_width": "3.1", "petal_length": "1.5", "petal_width": "0.1", "species": "setosa"},
    {"sepal_length": "4.4", "sepal_width": "3.0", "petal_length": "1.3", "petal_width": "0.2", "species": "setosa"},
    {"sepal_length": "5.1", "sepal_width": "3.4", "petal_length": "1.5", "petal_width": "0.2", "species": "setosa"},
    {"sepal_length": "5.0", "sepal_width": "3.5", "petal_length": "1.3", "petal_width": "0.3", "species": "setosa"},
    {"sepal_length": "4.5", "sepal_width": "2.3", "petal_length": "1.3", "petal_width": "0.3", "species": "setosa"},
    {"sepal_length": "4.4", "sepal_width": "3.2", "petal_length": "1.3", "petal_width": "0.2", "species": "setosa"},
    {"sepal_length": "5.0", "sepal_width": "3.5", "petal_length": "1.6", "petal_width": "0.6", "species": "setosa"},
    {"sepal_length": "5.1", "sepal_width": "3.8", "petal_length": "1.9", "petal_width": "0.4", "species": "setosa"},
    {"sepal_length": "4.8", "sepal_width": "3.0", "petal_length": "1.4", "petal_width": "0.3", "species": "setosa"},
    {"sepal_length": "5.1", "sepal_width": "3.8", "petal_length": "1.6", "petal_width": "0.2", "species": "setosa"},
    {"sepal_length": "4.6", "sepal_width": "3.2", "petal_length": "1.4", "petal_width": "0.2", "species": "setosa"},
    {"sepal_length": "5.3", "sepal_width": "3.7", "petal_length": "1.5", "petal_width": "0.2", "species": "setosa"},
    {"sepal_length": "5.0", "sepal_width": "3.3", "petal_length": "1.4", "petal_width": "0.2", "species": "setosa"},
    {"sepal_length": "7.0", "sepal_width": "3.2", "petal_length": "4.7", "petal_width": "1.4", "species": "versicolor"},
    {"sepal_length": "6.4", "sepal_width": "3.2", "petal_length": "4.5", "petal_width": "1.5", "species": "versicolor"},
    {"sepal_length": "6.9", "sepal_width": "3.1", "petal_length": "4.9", "petal_width": "1.5", "species": "versicolor"},
    {"sepal_length": "5.5", "sepal_width": "2.3", "petal_length": "4.0", "petal_width": "1.3", "species": "versicolor"},
    {"sepal_length": "6.5", "sepal_width": "2.8", "petal_length": "4.6", "petal_width": "1.5", "species": "versicolor"},
    {"sepal_length": "5.7", "sepal_width": "2.8", "petal_length": "4.5", "petal_width": "1.3", "species": "versicolor"},
    {"sepal_length": "6.3", "sepal_width": "3.3", "petal_length": "4.7", "petal_width": "1.6", "species": "versicolor"},
    {"sepal_length": "4.9", "sepal_width": "2.4", "petal_length": "3.3", "petal_width": "1.0", "species": "versicolor"},
    {"sepal_length": "6.6", "sepal_width": "2.9", "petal_length": "4.6", "petal_width": "1.3", "species": "versicolor"},
    {"sepal_length": "5.2", "sepal_width": "2.7", "petal_length": "3.9", "petal_width": "1.4", "species": "versicolor"},
    {"sepal_length": "5.0", "sepal_width": "2.0", "petal_length": "3.5", "petal_width": "1.0", "species": "versicolor"},
    {"sepal_length": "5.9", "sepal_width": "3.0", "petal_length": "4.2", "petal_width": "1.5", "species": "versicolor"},
    {"sepal_length": "6.0", "sepal_width": "2.2", "petal_length": "4.0", "petal_width": "1.0", "species": "versicolor"},
    {"sepal_length": "6.1", "sepal_width": "2.9", "petal_length": "4.7", "petal_width": "1.4", "species": "versicolor"},
    {"sepal_length": "5.6", "sepal_width": "2.9", "petal_length": "3.6", "petal_width": "1.3", "species": "versicolor"},
    {"sepal_length": "6.7", "sepal_width": "3.1", "petal_length": "4.4", "petal_width": "1.4", "species": "versicolor"},
    {"sepal_length": "5.6", "sepal_width": "3.0", "petal_length": "4.5", "petal_width": "1.5", "species": "versicolor"},
    {"sepal_length": "5.8", "sepal_width": "2.7", "petal_length": "4.1", "petal_width": "1.0", "species": "versicolor"},
    {"sepal_length": "6.2", "sepal_width": "2.2", "petal_length": "4.5", "petal_width": "1.5", "species": "versicolor"},
    {"sepal_length": "5.6", "sepal_width": "2.5", "petal_length": "3.9", "petal_width": "1.1", "species": "versicolor"},
    {"sepal_length": "5.9", "sepal_width": "3.2", "petal_length": "4.8", "petal_width": "1.8", "species": "versicolor"},
    {"sepal_length": "6.1", "sepal_width": "2.8", "petal_length": "4.0", "petal_width": "1.3", "species": "versicolor"},
    {"sepal_length": "6.3", "sepal_width": "2.5", "petal_length": "4.9", "petal_width": "1.5", "species": "versicolor"},
    {"sepal_length": "6.1", "sepal_width": "2.8", "petal_length": "4.7", "petal_width": "1.2", "species": "versicolor"},
    {"sepal_length": "6.4", "sepal_width": "2.9", "petal_length": "4.3", "petal_width": "1.3", "species": "versicolor"},
    {"sepal_length": "6.6", "sepal_width": "3.0", "petal_length": "4.4", "petal_width": "1.4", "species": "versicolor"},
    {"sepal_length": "6.8", "sepal_width": "2.8", "petal_length": "4.8", "petal_width": "1.4", "species": "versicolor"},
    {"sepal_length": "6.7", "sepal_width": "3.0", "petal_length": "5.0", "petal_width": "1.7", "species": "versicolor"},
    {"sepal_length": "6.0", "sepal_width": "2.9", "petal_length": "4.5", "petal_width": "1.5", "species": "versicolor"},
    {"sepal_length": "5.7", "sepal_width": "2.6", "petal_length": "3.5", "petal_width": "1.0", "species": "versicolor"},
    {"sepal_length": "5.5", "sepal_width": "2.4", "petal_length": "3.8", "petal_width": "1.1", "species": "versicolor"},
    {"sepal_length": "5.5", "sepal_width": "2.4", "petal_length": "3.7", "petal_width": "1.0", "species": "versicolor"},
    {"sepal_length": "5.8", "sepal_width": "2.7", "petal_length": "3.9", "petal_width": "1.2", "species": "versicolor"},
    {"sepal_length": "6.0", "sepal_width": "2.7", "petal_length": "5.1", "petal_width": "1.6", "species": "versicolor"},
    {"sepal_length": "5.4", "sepal_width": "3.0", "petal_length": "4.5", "petal_width": "1.5", "species": "versicolor"},
    {"sepal_length": "6.0", "sepal_width": "3.4", "petal_length": "4.5", "petal_width": "1.6", "species": "versicolor"},
    {"sepal_length": "6.7", "sepal_width": "3.1", "petal_length": "4.7", "petal_width": "1.5", "species": "versicolor"},
    {"sepal_length": "6.3", "sepal_width": "2.3", "petal_length": "4.4", "petal_width": "1.3", "species": "versicolor"},
    {"sepal_length": "5.6", "sepal_width": "3.0", "petal_length": "4.1", "petal_width": "1.3", "species": "versicolor"},
    {"sepal_length": "5.5", "sepal_width": "2.5", "petal_length": "4.0", "petal_width": "1.3", "species": "versicolor"},
    {"sepal_length": "5.5", "sepal_width": "2.6", "petal_length": "4.4", "petal_width": "1.2", "species": "versicolor"},
    {"sepal_length": "6.1", "sepal_width": "3.0", "petal_length": "4.6", "petal_width": "1.4", "species": "versicolor"},
    {"sepal_length": "5.8", "sepal_width": "2.6", "petal_length": "4.0", "petal_width": "1.2", "species": "versicolor"},
    {"sepal_length": "5.0", "sepal_width": "2.3", "petal_length": "3.3", "petal_width": "1.0", "species": "versicolor"},
    {"sepal_length": "5.6", "sepal_width": "2.7", "petal_length": "4.2", "petal_width": "1.3", "species": "versicolor"},
    {"sepal_length": "5.7", "sepal_width": "3.0", "petal_length": "4.2", "petal_width": "1.2", "species": "versicolor"},
    {"sepal_length": "5.7", "sepal_width": "2.9", "petal_length": "4.2", "petal_width": "1.3", "species": "versicolor"},
    {"sepal_length": "6.2", "sepal_width": "2.9", "petal_length": "4.3", "petal_width": "1.3", "species": "versicolor"},
    {"sepal_length": "5.1", "sepal_width": "2.5", "petal_length": "3.0", "petal_width": "1.1", "species": "versicolor"},
    {"sepal_length": "5.7", "sepal_width": "2.8", "petal_length": "4.1", "petal_width": "1.3", "species": "versicolor"},
    {"sepal_length": "6.3", "sepal_width": "3.3", "petal_length": "6.0", "petal_width": "2.5", "species": "virginica"},
    {"sepal_length": "5.8", "sepal_width": "2.7", "petal_length": "5.1", "petal_width": "1.9", "species": "virginica"},
    {"sepal_length": "7.1", "sepal_width": "3.0", "petal_length": "5.9", "petal_width": "2.1", "species": "virginica"},
    {"sepal_length": "6.3", "sepal_width": "2.9", "petal_length": "5.6", "petal_width": "1.8", "species": "virginica"},
    {"sepal_length": "6.5", "sepal_width": "3.0", "petal_length": "5.8", "petal_width": "2.2", "species": "virginica"},
    {"sepal_length": "7.6", "sepal_width": "3.0", "petal_length": "6.6", "petal_width": "2.1", "species": "virginica"},
    {"sepal_length": "4.9", "sepal_width": "2.5", "petal_length": "4.5", "petal_width": "1.7", "species": "virginica"},
    {"sepal_length": "7.3", "sepal_width": "2.9", "petal_length": "6.3", "petal_width": "1.8", "species": "virginica"},
    {"sepal_length": "6.7", "sepal_width": "2.5", "petal_length": "5.8", "petal_width": "1.8", "species": "virginica"},
    {"sepal_length": "7.2", "sepal_width": "3.6", "petal_length": "6.1", "petal_width": "2.5", "species": "virginica"},
    {"sepal_length": "6.5", "sepal_width": "3.2", "petal_length": "5.1", "petal_width": "2.0", "species": "virginica"},
    {"sepal_length": "6.4", "sepal_width": "2.7", "petal_length": "5.3", "petal_width": "1.9", "species": "virginica"},
    {"sepal_length": "6.8", "sepal_width": "3.0", "petal_length": "5.5", "petal_width": "2.1", "species": "virginica"},
    {"sepal_length": "5.7", "sepal_width": "2.5", "petal_length": "5.0", "petal_width": "2.0", "species": "virginica"},
    {"sepal_length": "5.8", "sepal_width": "2.8", "petal_length": "5.1", "petal_width": "2.4", "species": "virginica"},
    {"sepal_length": "6.4", "sepal_width": "3.2", "petal_length": "5.3", "petal_width": "2.3", "species": "virginica"},
    {"sepal_length": "6.5", "sepal_width": "3.0", "petal_length": "5.5", "petal_width": "1.8", "species": "virginica"},
    {"sepal_length": "7.7", "sepal_width": "3.8", "petal_length": "6.7", "petal_width": "2.2", "species": "virginica"},
    {"sepal_length": "7.7", "sepal_width": "2.6", "petal_length": "6.9", "petal_width": "2.3", "species": "virginica"},
    {"sepal_length": "6.0", "sepal_width": "2.2", "petal_length": "5.0", "petal_width": "1.5", "species": "virginica"},
    {"sepal_length": "6.9", "sepal_width": "3.2", "petal_length": "5.7", "petal_width": "2.3", "species": "virginica"},
    {"sepal_length": "5.6", "sepal_width": "2.8", "petal_length": "4.9", "petal_width": "2.0", "species": "virginica"},
    {"sepal_length": "7.7", "sepal_width": "2.8", "petal_length": "6.7", "petal_width": "2.0", "species": "virginica"},
    {"sepal_length": "6.3", "sepal_width": "2.7", "petal_length": "4.9", "petal_width": "1.8", "species": "virginica"},
    {"sepal_length": "6.7", "sepal_width": "3.3", "petal_length": "5.7", "petal_width": "2.1", "species": "virginica"},
    {"sepal_length": "7.2", "sepal_width": "3.2", "petal_length": "6.0", "petal_width": "1.8", "species": "virginica"},
    {"sepal_length": "6.2", "sepal_width": "2.8", "petal_length": "4.8", "petal_width": "1.8", "species": "virginica"},
    {"sepal_length": "6.1", "sepal_width": "3.0", "petal_length": "4.9", "petal_width": "1.8", "species": "virginica"},
    {"sepal_length": "6.4", "sepal_width": "2.8", "petal_length": "5.6", "petal_width": "2.1", "species": "virginica"},
    {"sepal_length": "7.2", "sepal_width": "3.0", "petal_length": "5.8", "petal_width": "1.6", "species": "virginica"},
    {"sepal_length": "7.4", "sepal_width": "2.8", "petal_length": "6.1", "petal_width": "1.9", "species": "virginica"},
    {"sepal_length": "7.9", "sepal_width": "3.8", "petal_length": "6.4", "petal_width": "2.0", "species": "virginica"},
    {"sepal_length": "6.4", "sepal_width": "2.8", "petal_length": "5.6", "petal_width": "2.2", "species": "virginica"},
    {"sepal_length": "6.3", "sepal_width": "2.8", "petal_length": "5.1", "petal_width": "1.5", "species": "virginica"},
    {"sepal_length": "6.1", "sepal_width": "2.6", "petal_length": "5.6", "petal_width": "1.4", "species": "virginica"},
    {"sepal_length": "7.7", "sepal_width": "3.0", "petal_length": "6.1", "petal_width": "2.3", "species": "virginica"},
    {"sepal_length": "6.3", "sepal_width": "3.4", "petal_length": "5.6", "petal_width": "2.4", "species": "virginica"},
    {"sepal_length": "6.4", "sepal_width": "3.1", "petal_length": "5.5", "petal_width": "1.8", "species": "virginica"},
    {"sepal_length": "6.0", "sepal_width": "3.0", "petal_length": "4.8", "petal_width": "1.8", "species": "virginica"},
    {"sepal_length": "6.9", "sepal_width": "3.1", "petal_length": "5.4", "petal_width": "2.1", "species": "virginica"},
    {"sepal_length": "6.7", "sepal_width": "3.1", "petal_length": "5.6", "petal_width": "2.4", "species": "virginica"},
    {"sepal_length": "6.9", "sepal_width": "3.1", "petal_length": "5.1", "petal_width": "2.3", "species": "virginica"},
    {"sepal_length": "5.8", "sepal_width": "2.7", "petal_length": "5.1", "petal_width": "1.9", "species": "virginica"},
    {"sepal_length": "6.8", "sepal_width": "3.2", "petal_length": "5.9", "petal_width": "2.3", "species": "virginica"},
    {"sepal_length": "6.7", "sepal_width": "3.3", "petal_length": "5.7", "petal_width": "2.5", "species": "virginica"},
    {"sepal_length": "6.7", "sepal_width": "3.0", "petal_length": "5.2", "petal_width": "2.3", "species": "virginica"},
    {"sepal_length": "6.3", "sepal_width": "2.5", "petal_length": "5.0", "petal_width": "1.9", "species": "virginica"},
    {"sepal_length": "6.5", "sepal_width": "3.0", "petal_length": "5.2", "petal_width": "2.0", "species": "virginica"},
    {"sepal_length": "6.2", "sepal_width": "3.4", "petal_length": "5.4", "petal_width": "2.3", "species": "virginica"},
    {"sepal_length": "5.9", "sepal_width": "3.0", "petal_length": "5.1", "petal_width": "1.8", "species": "virginica"}
];
```
