## Supplemental Examples for "R/LinkedCharts: A novel approach for simple but powerful interactive data analysis": JS_code_full.html

```
//scatter
lc.scatter()
   .x(i => +iris[i].sepal_length)
   .y(i => +iris[i].petal_length)
   .showPanel(false)
   .axisTitleX("Sepal length")
   .axisTitleY("Petal length")
   .colourValue(i => iris[i].species)
   .showLegend(false)
   .place("#A1");

//beeswarm
lc.beeswarm()
   .x(i => iris[i].species)
   .y(i => +iris[i].petal_length)
   .showPanel(false)
   .axisTitleX("Species")
   .axisTitleY("Petal length")
   .colourValue(i => +iris[i].sepal_length)
   .colourLegendTitle("Sepal length")
   .place("#A2");

//lines
var lines = lc.xLine()
   .showPanel(false)
   .lineFun(function(x) {return Math.sin(x)});

lc.xLine("abline", lines)
   .elementIds([-1, 1])
   .lineFun((x, i) => 2 *i * x + 1);

lc.xLine("hline", lines)
   .nelements(3)
   .lineFun((x, i) => i + 1);

lc.yLine("vline", lines)
   .nelements(3)
   .lineFun((y, i) => i + 1);

lc.pointLine("path", lines)
   .x(d3.range(30).map(el => 2 * Math.sin(el/4) - 3))
   .y(d3.range(30).map(el => 2 * Math.cos(el/4) - 3))
   .domainX([-6, 6])
   .domainY([-6, 6])
   .place("#A3");

//histogram and density plot are specfic to the ``rlc'' package
//JavaScript version of the library offers only barcharts and
//paths

//heatmap
//note that for this example the data are randomly generated
lc.heatmap()
   .ncols(30)
   .nrows(30)
   .colLabel(d => "Sample " + (d + 1))
   .rowLabel(d => "Feature " + (d + 1))
   .showPanel(false)
   .value(function(row, col) {
      var val = Math.random();
      if(row > 20)
         val += Math.random() * 2 + 1;
      if(col > 25)
         val += Math.random() * 4 + 0.5;
      return val;
   })
   .place("#B1");

//barchart
//the data for the example are randomly generated
var barData = d3.range(5).map(el1 => d3.range(3)
      .map(el2 => d3.range(2)
         .map(el3 => Math.ceil(Math.random() * 5))));

lc.barchart()
   .ngroups(5)
   .nbars(3)
   .nstacks(2)
   .showPanel(false)
   .showLegend(false)
   .value((d1, d2, d3) => barData[d1][d2][d3])
   .place("#B2");

//user inputs
//Though input charts exist in the LinkedCharts library,
//they are intended mainly for R users. In JavaScript we 
//suggest to use the <input> tag directly.

//colour-slider
//the slider has to be connected to a layer of some chart
lc.colourSlider()
   .linkedChart(beeswarm.layers.layer0)
   .place("#B3");
```
