## Supplemental Examples for "R/LinkedCharts: A novel approach for simple but powerful interactive data analysis": R_code_full.html

```
openPage(useViewer = FALSE, layout = "table2x4")

#scatter
lc_scatter(
  x = iris$Sepal.Length,
  y = iris$Petal.Length,
  showPanel = FALSE,
  axisTitleX = "Sepal length",
  axisTitleY = "Petal length",
  colourValue = iris$Species,
  showLegend = FALSE,
  place = "A1"
)

#beeswarm
lc_beeswarm(
  x = iris$Sepal.Length,
  y = iris$Species,
  showPanel = FALSE,
  axisTitleX = "Species",
  axisTitleY = "Petal length",
  colourValue = iris$Sepal.Length,
  colourLegendTitle = "Sepal length",
  place = "A2"
)

#lines
x <- seq(-1, 1, length.out = 100)
lc_line(
  x = x * 6,
  y = sin(x * 6),
  domainX = c(-6, 6),
  domainY = c(-6, 6),
  showPanel = FALSE,
  place = "A3"
)

lc_abLine(
  a = c(-2, 2),
  b = c(1, 1),
  chartId = "A3",
  addLayer = TRUE
)

lc_hLine(
  h = 1:3,
  chartId = "A3",
  addLayer = TRUE
)

lc_vLine(
  v = 1:3,
  chartId = "A3",
  addLayer = TRUE
)

lc_path(
  x = 2 * sin(x * 7) - 3,
  y = 2 * cos(x * 7) - 3,
  chartId = "A3",
  addLayer = TRUE
)

#histogram and density plot
#note that density plot has much smaller range across the Y-axis
#in the JavaScript example the values for the plot were multiplied
#by 100 for demonstration purposes
lc_hist(
  value = rnorm(100),
  showPanel = FALSE,
  place = "A4"
)

lc_dens(
  value = rnorm(1000),
  chartId = "A4",
  addLayer = TRUE
)

#heatmap
#Data are generated randomly
hData <- matrix(runif(900) + c(runif(600) * 2 + 1, rep(0, 300)), nrow = 30) + 
  t(matrix(c(runif(750) * 4 + 0.5, rep(0, 150)), nrow = 30))
lc_heatmap(
  value = hData,
  colLabel = paste0("Sample ", 1:30),
  rowLabel = paste0("Feature ", 1:30),
  showPanel = FALSE,
  place = "B1"
)

#barchart
lc_bars(
  value = sample(5, 30, replace = TRUE),
  groupIds = rep(1:5, each = 6),
  stackIds = rep(1:2, times = 15),
  barIds = rep(1:3, each = 2, times = 5),
  showPanel = FALSE,
  showLegend = FALSE,
  place = "B2"
)

#user input

#(optional) first add a table to arrange all inputs in two columns
lc_html(
  content = "

|  |  |
| --- | --- |
|  |  |

",
  place = "B3"  
)

lc_input(
  type = "text",
  labels = "Text box",
  value = "Some text",
  place = "B3a"
)
lc_input(
  type = "button",
  labels = c("Click me!", "No, better me!"),
  place = "B3b"
)
lc_input(
  type = "radio",
  labels = paste0("Option ", 1:3),
  value = 2,
  place = "B3a", chartId = "radio"
)
lc_input(
  type = "checkbox",
  labels = paste0("Option ", 1:3),
  value = c(TRUE, TRUE, FALSE),
  place = "B3a", chartId = "check"
)
lc_input(
  type = "range",
  width = 240,
  labels = paste0("Par", 1:4), 
  value = sample(100, 4),
  place = "B3b", chartId = "range"
)
lc_colourSlider(
  chart = "A2",
  place = "B3", chartId = "colour"
)

#html code
lc_html(content = paste0(
    "<center><p>Example of some</p>",
    "<h3>HTML code</h3>",
    "<i>You can add tables</i>",
    "<table border='1'><tr><td><b>colname1</b></td><td><b>colname2</b></td></tr>",
    "<tr><td>Element1</td><td>Element2</td></tr>",
    "<tr><td colspan='2'>More text</td></tr></table>",
    "Or images</center>",
    "<img src='https://via.placeholder.com/500x150.png' style = 'margin-left: 5px;'>",
    "<div style='background: lightgreen; padding: 5px; margin-top: 20px;'>",
    "Or any ohter HTML element</div>"),
  place = "B4")
```
