## Supplemental Examples for "R/LinkedCharts: A novel approach for simple but powerful interactive data analysis": heatmapData.js

```
var hData = {"corrMat":[[1,0.8928,0.7745,0.8964,0.8948,0.9015,0.882,0.8874,0.8655,0.8986,0.8975,0.8101,0.8926,0.8865,0.8917,0.8679,0.8485,0.8631,0.8663,0.8815,0.8921,0.8735,0.8656,0.8549,0.8692,0.8869,0.8451,0.8282,0.887,0.8856,0.7686,0.8705,0.8676,0.8757,0.8896,0.8819,0.8176,0.8759,0.8315,0.8734,0.8676,0.7463,0.8599,0.8633,0.7909,0.8634,0.7987,0.8173,0.8704,0.8602,0.8584,0.8818,0.8877,0.7622,0.869,0.8813,0.8289],[0.8928,1,0.7776,0.9026,0.9015,0.9087,0.8858,0.8945,0.8812,0.9118,0.9084,0.8083,0.8974,0.8969,0.9011,0.8783,0.8562,0.8756,0.8752,0.8862,0.9061,0.8784,0.8665,0.8582,0.8696,0.8943,0.8398,0.8328,0.8846,0.8939,0.7652,0.8776,0.8794,0.8768,0.8883,0.8842,0.8251,0.885,0.8335,0.8751,0.8723,0.7498,0.8641,0.8684,0.7947,0.8659,0.7939,0.808,0.8733,0.8661,0.8704,0.8807,0.8889,0.7628,0.8699,0.8863,0.8479],[0.7745,0.7776,1,0.7569,0.7539,0.7889,0.7557,0.7731,0.7858,0.7672,0.7801,0.8888,0.7896,0.7761,0.7764,0.7266,0.7313,0.7282,0.7251,0.7362,0.7434,0.7372,0.7355,0.7729,0.7588,0.7873,0.7979,0.7102,0.7913,0.7892,0.7188,0.7037,0.7388,0.7452,0.7819,0.7756,0.7008,0.737,0.7772,0.7283,0.7254,0.7616,0.7985,0.8321,0.8675,0.7703,0.837,0.8411,0.7435,0.7272,0.7737,0.7975,0.8153,0.8611,0.783,0.7535,0.7089],[0.8964,0.9026,0.7569,1,0.9363,0.9277,0.9119,0.9084,0.8743,0.9335,0.9246,0.7969,0.9134,0.9103,0.908,0.919,0.883,0.895,0.9194,0.9185,0.9284,0.9138,0.8848,0.8624,0.8982,0.899,0.8422,0.8484,0.8948,0.8986,0.7672,0.8979,0.8829,0.8984,0.9034,0.8939,0.8456,0.8974,0.8305,0.9111,0.8922,0.7421,0.8653,0.8641,0.7775,0.8929,0.7928,0.8094,0.8941,0.8836,0.8646,0.891,0.8948,0.7487,0.8904,0.9042,0.8483],[0.8948,0.9015,0.7539,0.9363,1,0.9242,0.9054,0.909,0.8741,0.9264,0.923,0.7969,0.9118,0.9093,0.9102,0.9036,0.8697,0.8864,0.9069,0.9129,0.926,0.9042,0.888,0.8639,0.8887,0.899,0.8439,0.8436,0.8987,0.8972,0.7705,0.8993,0.8837,0.9004,0.9089,0.8985,0.8402,0.8956,0.8291,0.9024,0.892,0.7389,0.8645,0.8614,0.7764,0.8835,0.7915,0.807,0.8935,0.8842,0.8651,0.889,0.8976,0.7455,0.8783,0.9014,0.8414],[0.9015,0.9087,0.7889,0.9277,0.9242,1,0.9036,0.9081,0.8888,0.922,0.9217,0.8276,0.9174,0.9133,0.9158,0.8917,0.8648,0.8915,0.8896,0.9046,0.9235,0.8938,0.8849,0.8751,0.8849,0.9067,0.8603,0.8442,0.9083,0.9139,0.7738,0.8875,0.8882,0.8924,0.9113,0.9074,0.8292,0.9,0.8515,0.8918,0.887,0.7609,0.8714,0.8762,0.8051,0.8779,0.8121,0.8354,0.8853,0.8762,0.8775,0.898,0.9064,0.7799,0.8843,0.9004,0.84],[0.882,0.8858,0.7557,0.9119,0.9054,0.9036,1,0.9181,0.871,0.9183,0.9167,0.802,0.9087,0.8985,0.8977,0.8982,0.8715,0.8816,0.9056,0.9074,0.9106,0.9093,0.8801,0.8561,0.8968,0.8998,0.8541,0.839,0.8992,0.8956,0.7718,0.8947,0.8775,0.9,0.9033,0.8939,0.837,0.8909,0.8273,0.9017,0.8813,0.7353,0.8731,0.8642,0.7766,0.8862,0.798,0.8104,0.8913,0.8755,0.8558,0.8962,0.8996,0.7579,0.8758,0.8888,0.8226],[0.8874,0.8945,0.7731,0.9084,0.909,0.9081,0.9181,1,0.8766,0.9199,0.9224,0.8166,0.917,0.9058,0.9062,0.8894,0.8653,0.8782,0.8983,0.9048,0.9124,0.9039,0.8819,0.8628,0.8948,0.908,0.8629,0.8386,0.9083,0.903,0.7742,0.8949,0.8811,0.9017,0.9127,0.9024,0.8326,0.894,0.8341,0.8968,0.8805,0.7398,0.8806,0.8754,0.791,0.8846,0.808,0.8216,0.892,0.8771,0.8617,0.901,0.911,0.7717,0.8778,0.8905,0.8236],[0.8655,0.8812,0.7858,0.8743,0.8741,0.8888,0.871,0.8766,1,0.8879,0.8913,0.8332,0.8739,0.8781,0.8791,0.8391,0.8236,0.8415,0.8522,0.864,0.8756,0.8534,0.8529,0.8593,0.8578,0.8859,0.8427,0.8033,0.874,0.8826,0.7609,0.8518,0.865,0.8576,0.8738,0.871,0.7995,0.8621,0.8272,0.8483,0.8427,0.7476,0.8569,0.8661,0.813,0.855,0.805,0.8197,0.8561,0.8528,0.8642,0.8665,0.8766,0.7758,0.8494,0.8603,0.8159],[0.8986,0.9118,0.7672,0.9335,0.9264,0.922,0.9183,0.9199,0.8879,1,0.9365,0.8156,0.9149,0.9101,0.9109,0.9128,0.8823,0.8935,0.9181,0.9161,0.9302,0.9135,0.8846,0.8671,0.9035,0.9104,0.8511,0.8518,0.9001,0.9033,0.7746,0.9044,0.8898,0.9041,0.9096,0.901,0.8471,0.8995,0.8353,0.9094,0.8917,0.7473,0.8804,0.8774,0.7876,0.8956,0.8006,0.8143,0.8953,0.8821,0.8719,0.8991,0.9015,0.7553,0.8922,0.9029,0.8497],[0.8975,0.9084,0.7801,0.9246,0.923,0.9217,0.9167,0.9224,0.8913,0.9365,1,0.8279,0.9251,0.9221,0.9215,0.9022,0.8737,0.8914,0.9073,0.9181,0.9327,0.9098,0.8892,0.8692,0.9011,0.9221,0.8712,0.8418,0.9172,0.9152,0.7741,0.9034,0.8957,0.9055,0.9245,0.914,0.8359,0.9095,0.8379,0.9029,0.8917,0.7438,0.8861,0.8854,0.7987,0.8904,0.8178,0.8265,0.9027,0.8894,0.8762,0.9033,0.9192,0.7816,0.8808,0.901,0.837],[0.8101,0.8083,0.8888,0.7969,0.7969,0.8276,0.802,0.8166,0.8332,0.8156,0.8279,1,0.8227,0.8134,0.8117,0.7589,0.7579,0.7639,0.7667,0.7816,0.7895,0.775,0.7801,0.8151,0.7987,0.8302,0.8329,0.7443,0.8269,0.8234,0.7491,0.7525,0.7809,0.7881,0.8219,0.8173,0.7362,0.7796,0.8132,0.7662,0.7642,0.7744,0.828,0.8573,0.8771,0.8056,0.853,0.8632,0.7837,0.7677,0.8125,0.8286,0.846,0.8536,0.8127,0.7898,0.7425],[0.8926,0.8974,0.7896,0.9134,0.9118,0.9174,0.9087,0.917,0.8739,0.9149,0.9251,0.8227,1,0.9221,0.9225,0.8933,0.8698,0.8886,0.8934,0.9022,0.9165,0.9041,0.8824,0.8645,0.892,0.9087,0.8681,0.8403,0.9156,0.9122,0.7764,0.8943,0.8842,0.9011,0.9182,0.9078,0.8299,0.9012,0.8412,0.898,0.8845,0.7508,0.8825,0.8785,0.8027,0.8838,0.8208,0.8353,0.8933,0.876,0.8657,0.905,0.9155,0.7915,0.8818,0.8965,0.8271],[0.8865,0.8969,0.7761,0.9103,0.9093,0.9133,0.8985,0.9058,0.8781,0.9101,0.9221,0.8134,0.9221,1,0.927,0.8951,0.8665,0.889,0.8902,0.9012,0.9156,0.894,0.8784,0.8594,0.8832,0.9056,0.8601,0.8351,0.9046,0.9053,0.7681,0.89,0.8837,0.892,0.9107,0.9016,0.8263,0.9022,0.834,0.8896,0.8804,0.7439,0.8714,0.8706,0.7936,0.8765,0.8129,0.824,0.889,0.878,0.8665,0.891,0.9028,0.7806,0.8713,0.893,0.831],[0.8917,0.9011,0.7764,0.908,0.9102,0.9158,0.8977,0.9062,0.8791,0.9109,0.9215,0.8117,0.9225,0.927,1,0.8832,0.8583,0.8815,0.8842,0.8995,0.9172,0.8901,0.8798,0.8626,0.8793,0.908,0.8593,0.8315,0.9075,0.9073,0.7681,0.89,0.8869,0.8905,0.9104,0.9025,0.8232,0.9022,0.8355,0.8885,0.8822,0.7431,0.871,0.8701,0.7935,0.8718,0.807,0.8215,0.888,0.8767,0.8688,0.8933,0.9047,0.7766,0.8697,0.8926,0.833],[0.8679,0.8783,0.7266,0.919,0.9036,0.8917,0.8982,0.8894,0.8391,0.9128,0.9022,0.7589,0.8933,0.8951,0.8832,1,0.9118,0.9116,0.909,0.8996,0.9058,0.9064,0.8669,0.8322,0.8805,0.872,0.8171,0.8666,0.8713,0.8699,0.7592,0.8856,0.8625,0.884,0.8853,0.8745,0.8506,0.882,0.8032,0.9017,0.8804,0.7295,0.8465,0.8383,0.7424,0.8737,0.766,0.7766,0.8762,0.868,0.8384,0.8692,0.8708,0.731,0.8804,0.8938,0.8423],[0.8485,0.8562,0.7313,0.883,0.8697,0.8648,0.8715,0.8653,0.8236,0.8823,0.8737,0.7579,0.8698,0.8665,0.8583,0.9118,1,0.887,0.8707,0.8691,0.8732,0.8751,0.8441,0.817,0.8552,0.8555,0.8096,0.8507,0.8529,0.8507,0.7618,0.8586,0.8455,0.8595,0.8589,0.8495,0.8373,0.8564,0.7906,0.8704,0.8551,0.7323,0.8364,0.8287,0.7448,0.8507,0.7682,0.7721,0.8543,0.8461,0.8271,0.8544,0.8538,0.7378,0.8581,0.8686,0.8265],[0.8631,0.8756,0.7282,0.895,0.8864,0.8915,0.8816,0.8782,0.8415,0.8935,0.8914,0.7639,0.8886,0.889,0.8815,0.9116,0.887,1,0.8712,0.8818,0.8981,0.8834,0.8584,0.8316,0.856,0.8697,0.8195,0.8583,0.8695,0.8749,0.76,0.8811,0.8656,0.8706,0.8789,0.8745,0.8324,0.8861,0.8172,0.878,0.8757,0.7367,0.8401,0.8308,0.7464,0.848,0.7671,0.783,0.8638,0.8548,0.8414,0.8631,0.8615,0.7354,0.8616,0.8844,0.8376],[0.8663,0.8752,0.7251,0.9194,0.9069,0.8896,0.9056,0.8983,0.8522,0.9181,0.9073,0.7667,0.8934,0.8902,0.8842,0.909,0.8707,0.8712,1,0.9141,0.9108,0.9148,0.872,0.8359,0.9029,0.884,0.8276,0.8241,0.8768,0.8781,0.7507,0.891,0.8691,0.8903,0.885,0.8707,0.8391,0.8836,0.8005,0.9097,0.873,0.7104,0.8529,0.8447,0.7482,0.8907,0.7722,0.7814,0.8852,0.8744,0.8376,0.8772,0.8766,0.7305,0.871,0.8795,0.8207],[0.8815,0.8862,0.7362,0.9185,0.9129,0.9046,0.9074,0.9048,0.864,0.9161,0.9181,0.7816,0.9022,0.9012,0.8995,0.8996,0.8691,0.8818,0.9141,1,0.9218,0.9076,0.8837,0.8524,0.8921,0.8962,0.8435,0.8333,0.8919,0.8913,0.7629,0.9001,0.8834,0.8945,0.8996,0.8885,0.8361,0.8968,0.8201,0.9035,0.8829,0.7223,0.8617,0.8546,0.7615,0.8794,0.7826,0.7998,0.8939,0.8859,0.855,0.8858,0.8916,0.7427,0.8675,0.8886,0.8252],[0.8921,0.9061,0.7434,0.9284,0.926,0.9235,0.9106,0.9124,0.8756,0.9302,0.9327,0.7895,0.9165,0.9156,0.9172,0.9058,0.8732,0.8981,0.9108,0.9218,1,0.9041,0.8882,0.86,0.8902,0.9077,0.8489,0.8426,0.8986,0.9008,0.7646,0.9081,0.8907,0.8983,0.9104,0.9031,0.8385,0.9144,0.8333,0.903,0.8923,0.7314,0.8635,0.8563,0.7655,0.8769,0.7854,0.8013,0.8951,0.8863,0.869,0.8884,0.8955,0.7399,0.8707,0.8979,0.8399],[0.8735,0.8784,0.7372,0.9138,0.9042,0.8938,0.9093,0.9039,0.8534,0.9135,0.9098,0.775,0.9041,0.894,0.8901,0.9064,0.8751,0.8834,0.9148,0.9076,0.9041,1,0.8854,0.8505,0.8977,0.891,0.8362,0.8393,0.8927,0.8925,0.7611,0.8957,0.8736,0.9054,0.9004,0.8885,0.8376,0.8865,0.8117,0.9065,0.8807,0.7216,0.8681,0.8551,0.7589,0.8858,0.7822,0.7936,0.89,0.8753,0.8443,0.8916,0.8913,0.7438,0.8738,0.886,0.8216],[0.8656,0.8665,0.7355,0.8848,0.888,0.8849,0.8801,0.8819,0.8529,0.8846,0.8892,0.7801,0.8824,0.8784,0.8798,0.8669,0.8441,0.8584,0.872,0.8837,0.8882,0.8854,1,0.8674,0.861,0.8776,0.8309,0.828,0.879,0.8827,0.7626,0.8756,0.8633,0.8871,0.8902,0.8876,0.8191,0.8732,0.8184,0.8719,0.8649,0.7291,0.8404,0.8372,0.7605,0.8524,0.7737,0.7914,0.8688,0.8673,0.8472,0.868,0.8781,0.7346,0.8486,0.8751,0.8141],[0.8549,0.8582,0.7729,0.8624,0.8639,0.8751,0.8561,0.8628,0.8593,0.8671,0.8692,0.8151,0.8645,0.8594,0.8626,0.8322,0.817,0.8316,0.8359,0.8524,0.86,0.8505,0.8674,1,0.8374,0.8628,0.8245,0.8118,0.8627,0.8683,0.7604,0.8405,0.8433,0.8606,0.8708,0.8706,0.7968,0.844,0.8286,0.8384,0.8362,0.752,0.8341,0.8425,0.7957,0.8399,0.7893,0.8128,0.8393,0.8381,0.8457,0.8545,0.8649,0.7544,0.8451,0.8585,0.8058],[0.8692,0.8696,0.7588,0.8982,0.8887,0.8849,0.8968,0.8948,0.8578,0.9035,0.9011,0.7987,0.892,0.8832,0.8793,0.8805,0.8552,0.856,0.9029,0.8921,0.8902,0.8977,0.861,0.8374,1,0.8948,0.8535,0.8152,0.8862,0.8842,0.759,0.8766,0.8662,0.8832,0.8876,0.8713,0.8239,0.8745,0.8125,0.8954,0.8619,0.7248,0.8669,0.8626,0.7777,0.8899,0.799,0.8097,0.88,0.8669,0.8434,0.8882,0.8875,0.7642,0.8709,0.8701,0.8075],[0.8869,0.8943,0.7873,0.899,0.899,0.9067,0.8998,0.908,0.8859,0.9104,0.9221,0.8302,0.9087,0.9056,0.908,0.872,0.8555,0.8697,0.884,0.8962,0.9077,0.891,0.8776,0.8628,0.8948,1,0.8854,0.8279,0.9165,0.9123,0.7738,0.888,0.8909,0.891,0.9108,0.8982,0.8174,0.8937,0.8315,0.8813,0.8761,0.7443,0.8877,0.887,0.8072,0.8772,0.8225,0.8302,0.8911,0.8793,0.8698,0.9001,0.9124,0.7936,0.864,0.8823,0.8215],[0.8451,0.8398,0.7979,0.8422,0.8439,0.8603,0.8541,0.8629,0.8427,0.8511,0.8712,0.8329,0.8681,0.8601,0.8593,0.8171,0.8096,0.8195,0.8276,0.8435,0.8489,0.8362,0.8309,0.8245,0.8535,0.8854,1,0.7821,0.8785,0.8703,0.7579,0.8292,0.8423,0.8409,0.8663,0.8572,0.7811,0.8446,0.8107,0.8323,0.8277,0.7336,0.8547,0.8617,0.8102,0.839,0.8287,0.8333,0.8463,0.8335,0.834,0.8673,0.8823,0.8124,0.8256,0.8332,0.7741],[0.8282,0.8328,0.7102,0.8484,0.8436,0.8442,0.839,0.8386,0.8033,0.8518,0.8418,0.7443,0.8403,0.8351,0.8315,0.8666,0.8507,0.8583,0.8241,0.8333,0.8426,0.8393,0.828,0.8118,0.8152,0.8279,0.7821,1,0.832,0.8292,0.7543,0.8379,0.822,0.8397,0.8414,0.8371,0.8081,0.825,0.786,0.8294,0.8344,0.728,0.8106,0.8024,0.7263,0.8131,0.7382,0.7535,0.8188,0.8146,0.806,0.8246,0.8239,0.7034,0.8346,0.8505,0.8135],[0.887,0.8846,0.7913,0.8948,0.8987,0.9083,0.8992,0.9083,0.874,0.9001,0.9172,0.8269,0.9156,0.9046,0.9075,0.8713,0.8529,0.8695,0.8768,0.8919,0.8986,0.8927,0.879,0.8627,0.8862,0.9165,0.8785,0.832,1,0.9189,0.7736,0.8837,0.8832,0.8965,0.92,0.9066,0.8122,0.8876,0.8309,0.8818,0.8779,0.7445,0.888,0.8857,0.8066,0.8739,0.8265,0.8366,0.8911,0.8759,0.8608,0.9054,0.9223,0.8025,0.8636,0.8816,0.8086],[0.8856,0.8939,0.7892,0.8986,0.8972,0.9139,0.8956,0.903,0.8826,0.9033,0.9152,0.8234,0.9122,0.9053,0.9073,0.8699,0.8507,0.8749,0.8781,0.8913,0.9008,0.8925,0.8827,0.8683,0.8842,0.9123,0.8703,0.8292,0.9189,1,0.7679,0.8789,0.8888,0.8887,0.9088,0.9047,0.8127,0.8958,0.8393,0.8795,0.8745,0.7484,0.8766,0.8788,0.8068,0.8711,0.8155,0.8323,0.8832,0.8723,0.8667,0.8981,0.9129,0.7983,0.8661,0.8824,0.8187],[0.7686,0.7652,0.7188,0.7672,0.7705,0.7738,0.7718,0.7742,0.7609,0.7746,0.7741,0.7491,0.7764,0.7681,0.7681,0.7592,0.7618,0.76,0.7507,0.7629,0.7646,0.7611,0.7626,0.7604,0.759,0.7738,0.7579,0.7543,0.7736,0.7679,1,0.7707,0.7641,0.771,0.7742,0.7707,0.7538,0.7589,0.755,0.7576,0.7622,0.712,0.7736,0.772,0.735,0.7595,0.7422,0.7511,0.7646,0.7557,0.7614,0.7764,0.7766,0.7129,0.7657,0.7698,0.7382],[0.8705,0.8776,0.7037,0.8979,0.8993,0.8875,0.8947,0.8949,0.8518,0.9044,0.9034,0.7525,0.8943,0.89,0.89,0.8856,0.8586,0.8811,0.891,0.9001,0.9081,0.8957,0.8756,0.8405,0.8766,0.888,0.8292,0.8379,0.8837,0.8789,0.7707,1,0.8886,0.8932,0.8904,0.8806,0.8332,0.8942,0.8118,0.8926,0.8821,0.7109,0.8586,0.838,0.7354,0.8606,0.7598,0.7748,0.8854,0.8757,0.8423,0.8737,0.8736,0.7105,0.851,0.8808,0.8231],[0.8676,0.8794,0.7388,0.8829,0.8837,0.8882,0.8775,0.8811,0.865,0.8898,0.8957,0.7809,0.8842,0.8837,0.8869,0.8625,0.8455,0.8656,0.8691,0.8834,0.8907,0.8736,0.8633,0.8433,0.8662,0.8909,0.8423,0.822,0.8832,0.8888,0.7641,0.8886,1,0.8715,0.8814,0.8754,0.82,0.8831,0.8204,0.8712,0.8704,0.7305,0.8529,0.8497,0.7666,0.8532,0.7761,0.792,0.8723,0.8689,0.8522,0.8703,0.878,0.7507,0.8453,0.8713,0.824],[0.8757,0.8768,0.7452,0.8984,0.9004,0.8924,0.9,0.9017,0.8576,0.9041,0.9055,0.7881,0.9011,0.892,0.8905,0.884,0.8595,0.8706,0.8903,0.8945,0.8983,0.9054,0.8871,0.8606,0.8832,0.891,0.8409,0.8397,0.8965,0.8887,0.771,0.8932,0.8715,1,0.911,0.8985,0.8277,0.88,0.8197,0.8886,0.875,0.7275,0.8651,0.8541,0.769,0.8734,0.7874,0.8017,0.8852,0.871,0.8491,0.8881,0.8935,0.7433,0.8644,0.8816,0.8176],[0.8896,0.8883,0.7819,0.9034,0.9089,0.9113,0.9033,0.9127,0.8738,0.9096,0.9245,0.8219,0.9182,0.9107,0.9104,0.8853,0.8589,0.8789,0.885,0.8996,0.9104,0.9004,0.8902,0.8708,0.8876,0.9108,0.8663,0.8414,0.92,0.9088,0.7742,0.8904,0.8814,0.911,1,0.9224,0.823,0.8927,0.8342,0.8874,0.8817,0.7442,0.8787,0.8783,0.7992,0.8767,0.8178,0.8301,0.8927,0.879,0.863,0.9006,0.9177,0.7804,0.871,0.8903,0.8176],[0.8819,0.8842,0.7756,0.8939,0.8985,0.9074,0.8939,0.9024,0.871,0.901,0.914,0.8173,0.9078,0.9016,0.9025,0.8745,0.8495,0.8745,0.8707,0.8885,0.9031,0.8885,0.8876,0.8706,0.8713,0.8982,0.8572,0.8371,0.9066,0.9047,0.7707,0.8806,0.8754,0.8985,0.9224,1,0.8165,0.8871,0.838,0.8755,0.8741,0.7456,0.867,0.8668,0.7938,0.8633,0.807,0.8244,0.878,0.8676,0.8601,0.8906,0.9075,0.7717,0.8612,0.8818,0.8144],[0.8176,0.8251,0.7008,0.8456,0.8402,0.8292,0.837,0.8326,0.7995,0.8471,0.8359,0.7362,0.8299,0.8263,0.8232,0.8506,0.8373,0.8324,0.8391,0.8361,0.8385,0.8376,0.8191,0.7968,0.8239,0.8174,0.7811,0.8081,0.8122,0.8127,0.7538,0.8332,0.82,0.8277,0.823,0.8165,1,0.8256,0.7824,0.8386,0.8298,0.7165,0.8005,0.7966,0.7234,0.8224,0.7415,0.7496,0.8254,0.8201,0.8043,0.8177,0.8156,0.7007,0.8197,0.8324,0.8052],[0.8759,0.885,0.737,0.8974,0.8956,0.9,0.8909,0.894,0.8621,0.8995,0.9095,0.7796,0.9012,0.9022,0.9022,0.882,0.8564,0.8861,0.8836,0.8968,0.9144,0.8865,0.8732,0.844,0.8745,0.8937,0.8446,0.825,0.8876,0.8958,0.7589,0.8942,0.8831,0.88,0.8927,0.8871,0.8256,1,0.8308,0.8865,0.8786,0.7257,0.8535,0.8468,0.7619,0.861,0.7847,0.8003,0.8819,0.8729,0.8562,0.8769,0.883,0.748,0.859,0.883,0.8231],[0.8315,0.8335,0.7772,0.8305,0.8291,0.8515,0.8273,0.8341,0.8272,0.8353,0.8379,0.8132,0.8412,0.834,0.8355,0.8032,0.7906,0.8172,0.8005,0.8201,0.8333,0.8117,0.8184,0.8286,0.8125,0.8315,0.8107,0.786,0.8309,0.8393,0.755,0.8118,0.8204,0.8197,0.8342,0.838,0.7824,0.8308,1,0.8102,0.8114,0.7526,0.8142,0.8197,0.7908,0.812,0.7821,0.8132,0.8083,0.7991,0.8254,0.8349,0.8365,0.7661,0.8231,0.8284,0.7828],[0.8734,0.8751,0.7283,0.9111,0.9024,0.8918,0.9017,0.8968,0.8483,0.9094,0.9029,0.7662,0.898,0.8896,0.8885,0.9017,0.8704,0.878,0.9097,0.9035,0.903,0.9065,0.8719,0.8384,0.8954,0.8813,0.8323,0.8294,0.8818,0.8795,0.7576,0.8926,0.8712,0.8886,0.8874,0.8755,0.8386,0.8865,0.8102,1,0.8815,0.7196,0.8568,0.8466,0.7515,0.8831,0.7769,0.7909,0.8856,0.8738,0.8433,0.8846,0.881,0.7346,0.8731,0.8839,0.8242],[0.8676,0.8723,0.7254,0.8922,0.892,0.887,0.8813,0.8805,0.8427,0.8917,0.8917,0.7642,0.8845,0.8804,0.8822,0.8804,0.8551,0.8757,0.873,0.8829,0.8923,0.8807,0.8649,0.8362,0.8619,0.8761,0.8277,0.8344,0.8779,0.8745,0.7622,0.8821,0.8704,0.875,0.8817,0.8741,0.8298,0.8786,0.8114,0.8815,1,0.729,0.8466,0.8391,0.748,0.852,0.7692,0.7827,0.8718,0.8654,0.841,0.8677,0.873,0.7321,0.8507,0.8782,0.8262],[0.7463,0.7498,0.7616,0.7421,0.7389,0.7609,0.7353,0.7398,0.7476,0.7473,0.7438,0.7744,0.7508,0.7439,0.7431,0.7295,0.7323,0.7367,0.7104,0.7223,0.7314,0.7216,0.7291,0.752,0.7248,0.7443,0.7336,0.728,0.7445,0.7484,0.712,0.7109,0.7305,0.7275,0.7442,0.7456,0.7165,0.7257,0.7526,0.7196,0.729,1,0.7411,0.7592,0.7609,0.7351,0.741,0.7579,0.7219,0.7145,0.7487,0.7496,0.7531,0.737,0.7557,0.7462,0.7272],[0.8599,0.8641,0.7985,0.8653,0.8645,0.8714,0.8731,0.8806,0.8569,0.8804,0.8861,0.828,0.8825,0.8714,0.871,0.8465,0.8364,0.8401,0.8529,0.8617,0.8635,0.8681,0.8404,0.8341,0.8669,0.8877,0.8547,0.8106,0.888,0.8766,0.7736,0.8586,0.8529,0.8651,0.8787,0.867,0.8005,0.8535,0.8142,0.8568,0.8466,0.7411,1,0.8875,0.812,0.8624,0.8282,0.833,0.8669,0.8449,0.8396,0.8883,0.8942,0.8058,0.8505,0.8502,0.793],[0.8633,0.8684,0.8321,0.8641,0.8614,0.8762,0.8642,0.8754,0.8661,0.8774,0.8854,0.8573,0.8785,0.8706,0.8701,0.8383,0.8287,0.8308,0.8447,0.8546,0.8563,0.8551,0.8372,0.8425,0.8626,0.887,0.8617,0.8024,0.8857,0.8788,0.772,0.838,0.8497,0.8541,0.8783,0.8668,0.7966,0.8468,0.8197,0.8466,0.8391,0.7592,0.8875,1,0.8417,0.8633,0.847,0.8506,0.86,0.8467,0.8519,0.8853,0.8991,0.8281,0.8552,0.8516,0.7987],[0.7909,0.7947,0.8675,0.7775,0.7764,0.8051,0.7766,0.791,0.813,0.7876,0.7987,0.8771,0.8027,0.7936,0.7935,0.7424,0.7448,0.7464,0.7482,0.7615,0.7655,0.7589,0.7605,0.7957,0.7777,0.8072,0.8102,0.7263,0.8066,0.8068,0.735,0.7354,0.7666,0.769,0.7992,0.7938,0.7234,0.7619,0.7908,0.7515,0.748,0.7609,0.812,0.8417,1,0.7905,0.8358,0.8432,0.7692,0.7566,0.7986,0.8097,0.8242,0.8361,0.7951,0.7738,0.7338],[0.8634,0.8659,0.7703,0.8929,0.8835,0.8779,0.8862,0.8846,0.855,0.8956,0.8904,0.8056,0.8838,0.8765,0.8718,0.8737,0.8507,0.848,0.8907,0.8794,0.8769,0.8858,0.8524,0.8399,0.8899,0.8772,0.839,0.8131,0.8739,0.8711,0.7595,0.8606,0.8532,0.8734,0.8767,0.8633,0.8224,0.861,0.812,0.8831,0.852,0.7351,0.8624,0.8633,0.7905,1,0.8095,0.8175,0.8699,0.8553,0.8435,0.8776,0.879,0.7659,0.8741,0.8662,0.8108],[0.7987,0.7939,0.837,0.7928,0.7915,0.8121,0.798,0.808,0.805,0.8006,0.8178,0.853,0.8208,0.8129,0.807,0.766,0.7682,0.7671,0.7722,0.7826,0.7854,0.7822,0.7737,0.7893,0.799,0.8225,0.8287,0.7382,0.8265,0.8155,0.7422,0.7598,0.7761,0.7874,0.8178,0.807,0.7415,0.7847,0.7821,0.7769,0.7692,0.741,0.8282,0.847,0.8358,0.8095,1,0.848,0.7942,0.7764,0.7977,0.8249,0.8394,0.832,0.8044,0.7863,0.732],[0.8173,0.808,0.8411,0.8094,0.807,0.8354,0.8104,0.8216,0.8197,0.8143,0.8265,0.8632,0.8353,0.824,0.8215,0.7766,0.7721,0.783,0.7814,0.7998,0.8013,0.7936,0.7914,0.8128,0.8097,0.8302,0.8333,0.7535,0.8366,0.8323,0.7511,0.7748,0.792,0.8017,0.8301,0.8244,0.7496,0.8003,0.8132,0.7909,0.7827,0.7579,0.833,0.8506,0.8432,0.8175,0.848,1,0.7996,0.7854,0.8138,0.8393,0.8499,0.8295,0.819,0.8032,0.7463],[0.8704,0.8733,0.7435,0.8941,0.8935,0.8853,0.8913,0.892,0.8561,0.8953,0.9027,0.7837,0.8933,0.889,0.888,0.8762,0.8543,0.8638,0.8852,0.8939,0.8951,0.89,0.8688,0.8393,0.88,0.8911,0.8463,0.8188,0.8911,0.8832,0.7646,0.8854,0.8723,0.8852,0.8927,0.878,0.8254,0.8819,0.8083,0.8856,0.8718,0.7219,0.8669,0.86,0.7692,0.8699,0.7942,0.7996,1,0.883,0.8519,0.883,0.8899,0.754,0.8542,0.8748,0.8121],[0.8602,0.8661,0.7272,0.8836,0.8842,0.8762,0.8755,0.8771,0.8528,0.8821,0.8894,0.7677,0.876,0.878,0.8767,0.868,0.8461,0.8548,0.8744,0.8859,0.8863,0.8753,0.8673,0.8381,0.8669,0.8793,0.8335,0.8146,0.8759,0.8723,0.7557,0.8757,0.8689,0.871,0.879,0.8676,0.8201,0.8729,0.7991,0.8738,0.8654,0.7145,0.8449,0.8467,0.7566,0.8553,0.7764,0.7854,0.883,1,0.8558,0.865,0.8754,0.737,0.8428,0.8719,0.8168],[0.8584,0.8704,0.7737,0.8646,0.8651,0.8775,0.8558,0.8617,0.8642,0.8719,0.8762,0.8125,0.8657,0.8665,0.8688,0.8384,0.8271,0.8414,0.8376,0.855,0.869,0.8443,0.8472,0.8457,0.8434,0.8698,0.834,0.806,0.8608,0.8667,0.7614,0.8423,0.8522,0.8491,0.863,0.8601,0.8043,0.8562,0.8254,0.8433,0.841,0.7487,0.8396,0.8519,0.7986,0.8435,0.7977,0.8138,0.8519,0.8558,1,0.856,0.8668,0.7676,0.8441,0.8567,0.819],[0.8818,0.8807,0.7975,0.891,0.889,0.898,0.8962,0.901,0.8665,0.8991,0.9033,0.8286,0.905,0.891,0.8933,0.8692,0.8544,0.8631,0.8772,0.8858,0.8884,0.8916,0.868,0.8545,0.8882,0.9001,0.8673,0.8246,0.9054,0.8981,0.7764,0.8737,0.8703,0.8881,0.9006,0.8906,0.8177,0.8769,0.8349,0.8846,0.8677,0.7496,0.8883,0.8853,0.8097,0.8776,0.8249,0.8393,0.883,0.865,0.856,1,0.913,0.8023,0.8723,0.8762,0.8079],[0.8877,0.8889,0.8153,0.8948,0.8976,0.9064,0.8996,0.911,0.8766,0.9015,0.9192,0.846,0.9155,0.9028,0.9047,0.8708,0.8538,0.8615,0.8766,0.8916,0.8955,0.8913,0.8781,0.8649,0.8875,0.9124,0.8823,0.8239,0.9223,0.9129,0.7766,0.8736,0.878,0.8935,0.9177,0.9075,0.8156,0.883,0.8365,0.881,0.873,0.7531,0.8942,0.8991,0.8242,0.879,0.8394,0.8499,0.8899,0.8754,0.8668,0.913,1,0.8243,0.8717,0.8812,0.8086],[0.7622,0.7628,0.8611,0.7487,0.7455,0.7799,0.7579,0.7717,0.7758,0.7553,0.7816,0.8536,0.7915,0.7806,0.7766,0.731,0.7378,0.7354,0.7305,0.7427,0.7399,0.7438,0.7346,0.7544,0.7642,0.7936,0.8124,0.7034,0.8025,0.7983,0.7129,0.7105,0.7507,0.7433,0.7804,0.7717,0.7007,0.748,0.7661,0.7346,0.7321,0.737,0.8058,0.8281,0.8361,0.7659,0.832,0.8295,0.754,0.737,0.7676,0.8023,0.8243,1,0.764,0.7458,0.6981],[0.869,0.8699,0.783,0.8904,0.8783,0.8843,0.8758,0.8778,0.8494,0.8922,0.8808,0.8127,0.8818,0.8713,0.8697,0.8804,0.8581,0.8616,0.871,0.8675,0.8707,0.8738,0.8486,0.8451,0.8709,0.864,0.8256,0.8346,0.8636,0.8661,0.7657,0.851,0.8453,0.8644,0.871,0.8612,0.8197,0.859,0.8231,0.8731,0.8507,0.7557,0.8505,0.8552,0.7951,0.8741,0.8044,0.819,0.8542,0.8428,0.8441,0.8723,0.8717,0.764,1,0.8809,0.828],[0.8813,0.8863,0.7535,0.9042,0.9014,0.9004,0.8888,0.8905,0.8603,0.9029,0.901,0.7898,0.8965,0.893,0.8926,0.8938,0.8686,0.8844,0.8795,0.8886,0.8979,0.886,0.8751,0.8585,0.8701,0.8823,0.8332,0.8505,0.8816,0.8824,0.7698,0.8808,0.8713,0.8816,0.8903,0.8818,0.8324,0.883,0.8284,0.8839,0.8782,0.7462,0.8502,0.8516,0.7738,0.8662,0.7863,0.8032,0.8748,0.8719,0.8567,0.8762,0.8812,0.7458,0.8809,1,0.8481],[0.8289,0.8479,0.7089,0.8483,0.8414,0.84,0.8226,0.8236,0.8159,0.8497,0.837,0.7425,0.8271,0.831,0.833,0.8423,0.8265,0.8376,0.8207,0.8252,0.8399,0.8216,0.8141,0.8058,0.8075,0.8215,0.7741,0.8135,0.8086,0.8187,0.7382,0.8231,0.824,0.8176,0.8176,0.8144,0.8052,0.8231,0.7828,0.8242,0.8262,0.7272,0.793,0.7987,0.7338,0.8108,0.732,0.7463,0.8121,0.8168,0.819,0.8079,0.8086,0.6981,0.828,0.8481,1]],"countMatrix":[[93,537,210,1315,1597,969,612,489,1046,3340,730,202,237,562,101,929,76,0,1030,210,485,516,12,123,212,70,0,182,0,0,0,40,0,0,114,120,0,222,140,50,123,0,0,291,0,1708,756,497,25,213,0,30,353,187,65,414,0],[0,0,179,6,0,0,0,0,0,202,0,0,0,128,15,0,0,0,0,0,0,0,194,61,0,0,0,0,101,0,0,0,0,278,208,363,0,0,0,86,0,0,0,0,0,0,426,0,0,0,0,0,0,0,26,0,0],[19850,24286,7026,39288,26004,33734,33858,36129,35554,70324,68394,9232,46014,42032,46415,44510,17861,21186,36446,42479,54244,49836,26554,21991,38556,33536,27151,12731,58477,50576,15038,25167,28027,22431,55571,41538,9799,68914,43965,39024,28553,20996,27748,20059,19367,38197,23053,21224,28429,19878,156051,27313,39468,10381,24033,24814,28350],[0,0,190,140,0,0,83,31,248,0,0,362,0,93,167,128,48,24,0,21,25,0,30,0,0,96,168,108,23,0,0,0,0,32,0,39,25,23,0,64,0,21,53,70,862,51,0,32,0,0,220,51,0,0,204,0,0],[74,25,1052,17,0,0,0,0,0,226,98,730,72,0,495,0,0,0,0,0,171,0,0,1194,0,101,0,0,1282,58,0,0,396,202,0,0,0,31,0,0,0,0,292,482,849,286,1405,0,331,1246,0,0,153,0,0,0,0],[0,50,1107,0,0,0,82,184,0,168,184,1013,573,0,137,0,0,0,0,0,0,77,0,1308,0,248,244,0,102,648,0,0,0,23,157,76,0,0,0,0,0,0,147,0,302,0,554,68,88,0,0,174,133,140,0,0,0],[5280,12238,16977,11905,6828,41603,15825,11959,11985,28068,22855,8132,23922,10405,10200,23194,5503,13598,26119,6940,20262,27370,6750,10034,11120,31599,18733,10106,9624,38579,13090,20087,10962,20399,9784,17990,2240,58658,18714,14406,8222,34017,61769,18601,26477,12390,28387,14435,11699,3351,48792,26817,12033,22267,6503,5231,10511],[0,0,31,23,0,0,0,0,0,0,0,87,0,0,0,0,12,0,0,0,0,0,0,0,0,0,0,0,0,0,0,0,0,0,0,0,0,0,0,0,0,0,0,0,0,528,0,0,0,0,0,0,0,108,75,0,0],[0,445,200,226,0,373,52,277,0,750,202,1073,0,130,320,101,0,0,43,200,0,454,110,0,101,0,0,103,274,101,0,0,0,491,0,369,371,0,0,0,0,0,2129,1123,28,875,0,99,0,0,5576,143,6,87,0,0,0],[422,122,1523,153,16,102,483,348,274,209,799,549,0,458,402,196,0,24,259,198,290,0,2057,0,96,0,714,0,175,258,0,200,202,575,735,3015,0,668,3830,733,0,0,30,231,1016,496,443,1867,0,0,0,323,643,301,210,0,0],[55831,160965,48024,167816,134965,238789,77318,103403,174894,386482,157479,68734,118179,162046,127194,157475,55661,126148,100884,82288,163245,153211,219157,440619,102757,247916,111392,139335,155418,309939,59553,99033,118567,235167,279111,289361,78583,251874,287636,47923,69161,42766,131488,59098,134210,156712,84059,111571,58323,66864,1140341,68148,70163,28360,124251,134172,181832],[138,200,3560,112,0,353,24,337,363,409,665,1176,472,265,425,161,51,35,0,396,343,42,30,1046,360,210,280,0,1094,735,0,0,58,161,534,91,37,0,924,0,0,1246,1490,611,881,81,1841,805,0,20,2019,388,682,3030,278,0,0],[343,483,1349,472,301,473,253,651,74,1703,177,863,458,102,101,168,180,497,993,704,369,202,84,40,364,606,1212,0,499,688,0,197,101,0,570,342,271,474,770,1297,0,142,0,283,784,312,1544,0,116,330,1051,344,382,1795,69,116,1933],[13202,23520,4487,22608,19758,17416,17328,20097,17399,48715,41836,5845,25299,14505,17304,22664,13543,18143,22615,21708,42929,34730,14294,14437,18085,42539,17988,1997,22013,48285,6945,21913,26530,26346,32853,27192,12338,74637,24952,19149,23707,4798,9828,9876,4378,18520,9381,11658,24165,13975,69618,19795,20771,2451,9646,12090,9963],[82261,97552,10217,192172,112941,103052,101022,105712,132839,344643,314236,22714,112902,120793,112806,119527,36151,60654,138860,127080,251076,168630,101505,125934,144290,270801,108029,34985,132770,230638,47635,86973,131361,120200,143074,138402,99028,234809,46541,112246,170975,25701,129205,73889,33013,135413,62823,55920,123936,68765,529763,94241,108787,12375,60339,59266,137913],[28740,47817,5994,26472,16860,15850,13300,22747,44339,61381,44253,11127,13514,20183,21579,31139,8904,16886,22253,15009,37561,35684,54167,39824,15123,43597,9424,3063,26784,41098,3313,14164,21708,36043,39305,40146,10530,62393,43872,12033,17728,3078,33442,15083,20990,19090,6725,20376,16228,19870,135793,21021,23963,6072,11095,8245,14519],[0,0,80,0,0,0,0,0,0,0,0,0,121,22,0,0,0,0,0,0,0,0,0,0,0,0,0,0,0,0,0,0,0,0,0,0,0,0,0,0,0,0,0,0,0,0,0,0,0,0,0,0,0,0,0,0,0],[2863,5151,2701,2888,2380,2959,2236,2634,3676,5636,6593,1550,5490,4200,4897,3450,1761,2417,916,1256,2055,3123,2552,6595,710,2154,1454,641,1985,3046,1998,4246,4854,4034,4720,7539,5673,5287,8840,3115,3476,4403,5565,2252,5698,3885,2422,5630,3089,1860,28246,3286,5256,960,2816,4085,2954],[106867,169350,56433,183109,136667,178813,81508,103953,322867,384970,233845,85769,97899,108232,113487,136981,56248,79031,111013,111686,204701,135626,176239,271426,80604,189039,89083,115877,184837,266125,49557,94600,100115,164606,228352,274112,75432,253201,245900,69346,118103,76605,221815,133689,185481,131246,93900,122664,101151,79112,1332162,83682,104421,37490,70756,91694,114822],[3630,5437,4703,22255,15657,7652,22405,24569,4430,53657,24309,7541,23271,14330,9019,30801,11074,3089,25946,9712,31956,41486,16104,12750,22466,23735,9297,10398,12788,8106,15297,23152,5316,32437,31866,18912,4918,18322,13223,19406,15748,4765,66807,15046,8186,23505,19506,13581,16428,9487,32864,17106,17355,801,12011,4934,6841],[0,257,1277,0,147,404,163,0,0,0,0,1898,739,400,94,28,0,0,0,0,0,0,0,0,0,0,201,0,199,795,0,0,0,0,1009,0,0,0,0,0,0,0,0,416,2390,0,597,2290,0,0,4374,167,785,1408,0,0,0],[758,660,337,2296,2014,933,631,1093,872,3154,1249,1177,1219,33,702,2437,255,2480,1669,813,2008,2166,3244,1031,3389,1897,342,933,615,925,6328,1269,1005,3442,3549,2261,6676,4363,1856,744,2120,878,3242,1493,2362,3529,120,15,2452,1173,36,2495,660,0,3117,591,96],[73732,119041,31147,172829,149741,296926,60155,75407,211659,193270,229470,42952,223782,256950,203838,154368,58585,144360,56232,97741,256539,128719,194327,171717,87936,213453,108077,128735,195548,364435,50926,84921,114070,106067,314078,294988,77832,457097,268506,63981,123078,49226,95126,47490,63073,89150,109171,135458,78578,73745,954496,91782,95173,52056,70498,99012,135017],[0,0,0,0,0,91,0,0,0,62,0,273,0,0,0,0,0,0,0,101,0,0,0,0,0,0,47,0,0,116,0,0,0,0,20,0,0,0,0,52,0,0,0,208,0,101,0,0,0,0,0,0,0,0,0,0,0],[19726,29619,8158,37193,24142,49265,26933,33449,49030,76646,79744,9315,27566,29328,37113,42076,21914,20178,36727,28209,44803,61014,63827,80322,24792,57948,23702,39784,78151,122601,19865,24884,26316,57282,94273,155772,13643,59677,32277,19892,32845,2881,20824,12885,7784,33410,17325,12091,18142,25724,164946,25657,37636,7135,17580,21669,10630],[0,299,138,196,0,0,0,198,177,461,510,425,0,545,130,0,0,0,0,128,0,2,0,0,0,461,0,0,0,170,0,197,0,0,651,402,0,601,0,324,0,0,300,29,505,199,110,0,202,0,2134,456,11,0,25,0,0],[494,219,448,1375,577,266,57,0,0,1656,796,0,1225,638,484,708,48,418,800,698,439,501,0,1032,287,33,0,0,348,740,0,0,834,286,432,124,0,14,1095,961,1514,0,321,25,0,1449,1661,0,130,215,1056,522,228,810,304,36,92],[82,0,294,89,204,45,0,0,0,88,118,185,216,0,0,164,0,86,133,0,40,0,0,0,0,0,0,0,39,0,0,0,0,0,163,0,0,0,54,0,0,0,0,0,0,0,0,200,44,0,0,0,0,0,33,0,0],[97154,89407,11784,200051,139326,185045,138183,158087,107838,408116,390771,19260,170867,119579,127962,229483,81727,102635,251893,165649,218709,318325,241703,116498,213390,264831,143497,69154,386483,329898,47387,147344,140229,201776,327860,251731,118952,282115,63234,169957,193076,35038,142699,72783,27157,174710,75305,59448,127079,116072,668264,135467,167991,30910,84072,62818,86869],[52665,126060,9523,41457,30303,64558,39430,32890,126258,253613,125399,12074,47024,76119,78827,101938,47708,148581,21664,101159,107209,93149,69212,49135,47854,287964,86313,44852,181913,278410,24897,120181,107064,117147,157279,130217,34795,272168,60755,29699,140292,31853,83365,24864,23801,28401,20826,26669,58297,23736,191195,77240,56506,12027,14677,22851,91574],[303,919,5233,512,140,1902,1456,1082,202,1709,1311,3717,2359,2726,801,1865,913,2023,386,828,858,1396,322,3135,1261,3040,4620,405,1533,2350,35,201,275,1654,2382,3300,37,1195,9978,706,91,9296,2835,1689,5626,859,13558,4242,996,1147,21181,2793,8747,5319,1425,276,0],[0,0,42,0,0,258,0,0,28,189,223,0,23,0,0,292,269,0,25,0,57,0,0,33,0,23,0,242,26,0,0,0,0,44,0,32,0,0,500,0,24,0,301,0,437,48,0,200,0,0,40,202,542,980,79,30,0],[6186,10342,6287,9551,4778,16945,4218,6096,6455,18036,9237,4101,9604,2396,4684,7569,7517,8580,4969,4263,8684,8769,6273,7980,4982,12024,11592,13160,10992,26537,1494,5109,6178,5145,12615,11509,252,14940,27998,7140,7473,10501,4512,3048,10914,1483,5599,16277,6052,2630,47544,7966,6814,5255,8716,2525,3960],[0,252,817,28,0,1088,0,0,202,0,8,1185,0,242,202,0,0,0,0,0,0,0,0,2810,404,124,0,0,0,0,0,0,739,0,154,0,0,0,0,0,0,0,397,613,1162,1086,5613,0,0,0,25,574,312,202,720,0,0],[18962,35973,9901,40105,30425,20228,18758,21118,28433,84001,50233,13046,31023,33378,35096,35511,15458,15982,37086,23664,48852,27757,16758,22679,28829,33310,6489,14204,30738,18346,13083,26354,17344,31582,30839,30215,11138,32118,21711,23904,18113,7551,42552,12640,13768,26345,20909,9644,20414,6314,65296,19150,14929,2324,10205,11468,24207],[27241,40721,3842,74414,50014,51968,46466,37273,40275,159826,117752,8331,39126,38437,41299,82441,32495,32221,92320,93118,69741,85254,58444,56014,70185,94343,39598,13468,69469,95240,15812,35060,61323,58216,66454,63936,26519,67500,15934,51577,57243,6047,67949,36952,13045,69745,22277,21242,48435,34471,336285,42347,43984,5363,32433,33187,45999],[4180,7568,9652,4610,5657,3139,662,2262,4610,9530,3220,4899,4895,1922,2756,5000,1265,2208,1098,898,2521,1518,2405,2945,892,1407,376,1552,1503,2878,2996,1569,4941,1584,2447,2986,1872,2833,9430,1542,5514,5231,2753,860,9339,1094,1826,1985,510,372,7720,1673,3201,1105,4241,2098,7449],[0,180,70,184,0,0,0,184,10,0,44,0,93,262,308,187,32,0,0,0,28,0,36,0,93,666,0,0,24,86,0,0,0,0,102,86,0,0,0,0,0,0,121,12,0,0,0,417,0,0,12,175,0,110,0,3,0],[232,28,399,0,37,659,0,626,202,682,902,2373,56,820,675,387,0,34,200,229,195,0,0,0,201,92,2159,23,388,1004,6572,0,661,1034,647,26,0,394,0,0,0,0,896,211,993,0,2942,419,401,0,117,514,24,2961,124,0,0],[82824,91343,8019,154608,110693,112677,78414,81007,62461,199158,297596,9578,85715,67620,68047,127149,30393,52859,118596,123928,196913,150921,134912,55412,100003,150415,86743,45971,164583,198902,29125,74226,125260,125363,169667,189717,58556,235442,25002,95799,113259,22265,98590,57174,17609,88378,61545,35390,83753,102973,544378,85596,112545,20007,36989,39758,47427],[100894,110338,21879,193918,186716,226949,93626,135337,191199,292062,412097,54080,185914,199629,198996,157834,44324,66620,122364,223761,265460,131546,238588,249196,109669,213335,156223,80620,274807,119673,105746,146291,141514,194517,474626,488459,126843,405878,247159,108472,154046,16507,192463,108722,60976,144576,141817,140053,104836,108462,1445824,93253,180363,16276,57197,69860,54836],[0,515,15,875,0,811,2250,1317,619,0,0,0,915,482,1111,63,0,0,1611,2131,3635,0,1225,1988,206,0,0,26,1322,2146,0,330,379,539,607,2300,0,3008,0,279,0,0,70,91,0,0,79,0,522,651,37,0,0,0,25,694,0],[1229,1464,1748,3009,1887,1249,2681,2840,1043,3979,5438,1163,2985,1556,1756,3686,969,2697,889,2562,4041,1949,1750,1264,2567,6232,4956,1807,2909,2773,0,2818,3152,1254,5172,5836,0,2931,2410,2074,1565,2782,5059,3060,2460,3986,1548,0,1285,261,5415,2532,2348,1148,1100,1371,517],[260,0,1766,0,95,1121,433,123,209,1341,483,2104,1338,465,163,58,0,0,0,102,0,281,0,362,110,550,705,46,506,1935,0,0,600,306,1320,689,0,0,43,0,585,42,541,1361,1879,1443,3432,1865,0,3,0,0,427,1007,252,259,0],[3977,5316,3662,11785,6406,9981,5014,5214,1956,9989,6015,3785,5403,5234,3654,9923,2550,2586,9179,4687,8439,8049,6920,4582,4902,4698,3487,370,6078,7204,80,4001,2869,3346,5830,10569,3628,5814,849,5699,6310,1420,3499,3045,4685,12788,5926,2622,3932,6088,19773,1844,5280,3259,3549,1284,6220],[3421,3854,1272,8508,6502,11464,3055,2755,16070,16575,4732,3676,5257,4886,6793,15034,3086,2911,4849,2863,4490,3510,7410,10608,5715,9334,1637,6148,6412,10905,599,1203,3032,6138,7102,14667,4426,4305,2527,5995,3376,1742,5605,5352,7978,8271,704,1884,1795,744,29506,1850,1399,1734,7403,6844,13396],[690,452,1860,0,108,202,0,495,402,1196,1075,1460,397,724,826,36,120,0,117,201,404,0,801,587,1006,1304,194,0,391,2472,0,404,0,0,1237,533,0,0,4746,0,0,3082,2340,1063,3107,731,4822,2833,1241,575,4,400,800,4741,0,159,0],[0,0,174,0,438,98,0,0,174,309,966,134,138,165,57,81,0,0,187,570,231,98,423,0,0,0,53,0,428,1400,0,60,0,364,824,404,38,0,0,0,0,0,0,232,0,268,502,0,0,0,747,0,513,314,0,988,0],[0,0,200,0,0,0,0,302,604,0,130,356,0,199,180,0,38,128,0,174,0,0,0,0,0,404,0,0,0,0,0,0,0,0,147,0,0,212,936,0,0,0,518,0,988,0,0,582,0,0,0,201,202,1199,0,70,0],[207,633,597,769,307,404,470,498,88,2168,1152,434,484,455,573,3016,768,278,468,358,708,225,543,366,164,134,591,178,516,860,792,72,34,705,484,613,265,414,315,127,0,0,391,223,564,444,798,28,154,54,1457,540,188,2439,939,240,31],[0,0,234,44,90,156,13,0,0,161,522,481,54,80,0,315,173,18,197,25,8,5,0,134,0,767,0,37,39,31,11,0,0,0,41,72,0,0,5,0,270,0,288,65,489,0,1431,499,18,0,44,0,0,5,2,5,447],[0,0,84,88,211,44,0,0,0,20,89,6,64,0,0,0,0,0,0,109,0,0,1285,0,0,0,0,0,0,459,135,84,0,392,26,8,0,0,0,0,0,0,0,0,75,57,0,0,0,0,146,97,219,0,64,160,0],[73,0,318,0,0,0,0,0,0,0,0,0,0,0,51,0,0,30,0,0,0,0,0,0,0,0,0,0,137,0,0,0,0,0,0,0,0,0,0,0,0,0,107,76,0,0,154,138,1,16,0,21,0,99,0,0,0],[1869,1922,0,7557,5350,1598,3777,4888,618,11656,16366,0,3564,2596,3421,5062,4736,1297,10012,5454,6593,11283,2255,396,7055,7882,2379,971,3057,3814,0,8043,5637,4480,2397,2815,865,10591,0,6516,2677,0,4340,2341,439,7008,866,0,5013,5749,12333,3829,6692,302,1364,1767,2783],[164,98,1006,92,0,340,442,20,117,191,1094,1449,331,511,299,65,0,0,244,0,299,1346,0,129,58,267,201,0,201,100,0,0,320,252,2335,761,0,1518,3369,35,0,0,804,882,2478,20,1777,646,404,20,0,572,569,844,350,927,0],[8,1479,4683,481,374,1243,569,1302,2786,857,819,1362,1787,2164,2419,2210,507,1421,965,362,459,55,5640,3379,0,1058,1208,253,854,6064,3334,0,401,1200,856,2143,63,2891,0,304,657,56,1330,98,4033,202,7363,945,1224,155,6267,1072,845,4009,158,631,70],[3569,2113,1105,3805,2472,3321,2319,4512,2599,10114,10724,1073,5802,3132,5854,7266,6157,3516,5380,4632,5422,3392,2847,8074,1195,3957,3920,1566,3113,6182,0,1374,1189,5721,3364,5708,9525,6223,5380,3621,8025,4,15064,7189,3098,3568,3119,421,5602,4339,7816,1556,5094,744,2860,1850,2685],[1397505,1866381,70751,2081045,1873274,1625065,1269347,1957672,2817818,6467149,3981686,445776,1383450,1559901,1806550,2787030,726525,1104268,1202883,753711,971518,3037593,1667205,3070122,1814554,1919055,907323,365823,2663898,2522149,27345,1073169,2298939,2264821,2728287,3185506,1290089,4159909,2285481,4048725,3074253,1827823,4368667,2198514,1095738,2950624,1259386,1626522,2105455,389805,10051838,1310948,1638889,213453,263757,644993,770474],[2321,1398,5924,4775,1749,2720,1430,2489,4114,3130,3544,3650,901,2581,1438,4241,3519,1180,1137,1773,1371,2206,329,4095,2557,2731,3366,1549,2308,3685,1255,382,890,2582,3951,3151,182,397,2699,390,1080,3431,3723,3291,7968,778,8054,3213,1385,438,8566,604,2401,3981,1767,492,164],[2638,1489,1246,2426,2539,780,1109,1594,140,5268,5857,687,2593,3674,2731,7806,1245,7451,1411,1881,1684,2567,5191,1161,3463,3406,1194,1654,1816,1959,0,1094,2804,2592,4177,2794,0,2515,100,1969,1393,0,651,1258,627,339,2567,1081,3518,1960,3996,2083,1378,969,551,525,2423],[15,112,0,0,226,0,0,0,0,6,173,0,17,0,0,36,18,0,203,173,35,2,0,0,42,365,107,0,93,0,0,385,699,0,0,0,260,660,0,65,0,0,100,0,0,0,0,60,266,0,4977,0,87,62,10,5,0],[0,200,25,0,0,0,0,0,0,0,0,0,0,0,44,0,0,0,0,0,0,0,0,0,0,0,0,0,0,0,0,0,0,0,0,0,0,0,0,0,0,0,70,0,0,0,0,149,0,0,0,0,0,0,210,0,0],[71499,108491,13596,116876,90606,103726,64940,82562,82900,282506,231085,25614,68631,72138,70522,115923,43297,81263,122632,91090,183003,204456,174828,180571,120066,191475,50846,64250,240458,355610,49573,108826,103880,163330,186928,213410,60950,164816,125481,82015,87785,16779,153685,83200,30922,102760,31375,48169,68139,73128,553718,80824,106575,14550,52275,58341,63125],[0,24,354,0,0,0,0,0,0,129,83,267,9,53,0,32,0,0,0,0,45,0,0,0,39,0,694,0,0,0,0,0,0,200,197,0,48,0,46,0,0,38,0,0,0,197,106,0,0,0,53,0,84,36,47,20,0],[1946,2359,1277,1817,782,1002,838,1635,2537,10979,7497,1070,2950,2175,2385,11361,6572,9423,2968,3198,6771,5589,1176,770,2889,3964,755,2160,1032,3767,0,2729,2554,3229,1130,3964,34,9150,3967,1645,408,0,8271,1179,1611,256,902,2979,4115,2543,7583,2335,2526,1876,395,578,541],[3350,6229,2842,17206,19176,9999,6316,6417,1547,41499,14089,3646,10060,3443,2246,8230,500,1958,16477,4619,11378,33246,17427,17065,16064,17641,4589,6168,14825,4631,7482,7256,8712,17913,15501,16253,0,5001,603,9308,515,10064,10488,7432,8706,22712,10298,3271,3982,4023,74800,5432,10192,2283,9652,2836,7180],[98135,130105,18788,189658,143431,134644,97681,120900,117313,409867,241226,32556,92598,100281,98448,108689,33451,58571,191393,126559,232110,217042,196241,261576,127165,209310,62871,67281,228818,328482,66267,110107,88433,212184,228610,254770,82822,166582,138145,86021,82896,19245,123098,80075,47958,147274,43084,64850,100802,83845,601795,96421,111159,23316,75530,68036,78019],[702644,999041,85633,1544615,1103422,813772,877736,953524,721633,3447778,2557099,199124,879442,941161,941409,941101,391198,546160,1438000,1134072,2548416,1605708,1233405,1133666,1228331,1856831,785122,420417,1663600,1871283,374648,859288,1014549,1305858,1774964,1674596,768800,1864550,826286,895584,1077106,230510,1264786,647595,413004,1335573,551718,497504,1008540,815555,6446101,754973,794062,117041,563548,508171,736073],[1600,2056,4499,754,520,1690,917,780,2402,1489,1997,4807,4497,1387,2122,3355,1347,4035,395,306,732,1468,1165,2233,1940,3588,4414,575,3135,2923,5181,659,1756,956,2172,3052,118,1741,4086,1275,1875,15023,6017,3919,12675,121,11401,12397,1521,43,22075,1110,2195,4713,6338,5173,3486],[70792,78111,10951,115102,91436,99471,75376,74073,62945,252583,181350,18611,84200,80156,76345,105880,30161,59927,113868,105428,167372,124540,80982,87741,95542,155130,56958,32332,125515,144744,22588,70550,67260,83134,146941,155898,71916,160375,53259,86328,71041,21187,117914,55166,29150,94583,48852,39807,65853,75800,332300,63806,79529,11691,44981,50942,30623],[5346,7311,2589,10313,8678,6757,8391,10762,6586,5322,2532,1597,1626,1751,1933,3580,494,777,27763,17407,22683,5865,3806,343,2935,2860,1594,220,3164,4266,101,1472,2170,14323,13360,7862,6726,10208,3270,1767,1838,2134,12067,9485,2161,2738,6005,2703,5788,7294,19802,4725,4819,1027,1490,1186,2058],[0,0,200,103,0,0,202,0,0,90,183,479,0,10,27,63,0,0,589,0,0,0,0,0,0,0,0,0,0,0,0,0,0,0,0,0,0,0,0,0,0,0,0,82,101,0,0,0,0,0,0,0,0,151,0,0,0],[0,60,508,0,0,0,228,88,35,36,10,454,226,7,171,0,0,435,0,0,74,44,0,0,138,0,169,0,30,112,859,0,93,133,195,0,0,0,0,0,0,0,413,278,909,0,772,489,0,0,0,0,357,729,442,0,0],[1110,2089,399,2599,1845,4550,620,576,754,4931,2301,166,1235,1519,1377,2577,0,3816,1546,904,2002,6387,5104,15202,1512,249,489,1366,1912,2206,0,978,739,8010,8349,13495,131,941,5173,2014,2494,1212,510,398,420,995,0,1636,160,157,1041,1133,1585,0,817,1065,765],[51619,92714,13417,77564,65403,91601,43643,53585,48911,184674,132522,17160,56429,69201,76818,70907,24995,53651,63878,67825,111453,116584,151251,73591,59170,203377,73303,21222,150827,734978,17437,68305,150784,80517,133867,136484,32460,149045,52996,48435,112316,23211,82554,49910,25638,48842,33467,27693,40917,46445,403378,54726,60984,16649,20166,28732,80761],[158,0,219,0,0,0,0,0,122,0,0,0,0,0,0,0,0,0,0,0,142,0,0,0,0,84,85,0,0,94,0,0,0,0,0,0,0,0,0,0,0,0,0,83,0,0,0,0,0,0,0,0,0,0,0,0,0],[588,0,2189,539,435,503,760,1468,605,1639,1195,1698,1757,1249,237,856,796,360,98,202,160,1596,0,1653,602,814,87,0,733,3626,0,403,25,906,341,1501,0,1804,0,0,0,0,7346,1503,2678,785,3480,1492,112,710,3419,2110,1458,623,624,508,0],[0,0,264,54,0,0,0,202,280,0,46,0,0,0,0,0,0,0,0,0,35,0,0,1010,0,0,0,0,0,209,0,0,0,183,0,0,0,0,0,0,0,0,0,102,745,0,0,0,0,0,0,0,0,97,0,0,0],[914,359,822,234,347,703,0,159,45,206,709,876,181,22,244,0,0,0,0,179,169,1014,86,684,571,249,805,0,906,627,6309,133,0,1524,170,1078,0,954,0,0,0,0,2048,450,321,0,3883,512,0,177,5945,203,530,631,0,131,0],[3646,4224,4824,1410,4632,7208,1338,1877,2761,4943,5409,5708,3981,5426,4856,5399,2051,2298,740,1916,4628,2055,4629,8228,1549,4066,4368,2556,3068,8543,1949,3529,1979,3612,10616,8017,1237,14162,14141,1215,6930,4536,7434,2807,6836,3562,4478,9330,220,312,13311,3103,2094,2070,1926,1946,0],[0,356,2069,1734,303,993,1136,768,2271,1328,1607,1927,566,1398,596,284,25,343,959,1195,1235,252,1210,0,808,492,453,23,833,1586,0,1776,1551,508,5634,3857,0,47,0,184,494,0,1896,1465,2611,296,3812,1336,394,0,0,1143,1298,696,765,658,0],[0,0,0,0,0,0,0,0,0,0,0,202,0,0,0,0,0,52,0,0,0,0,0,0,0,0,92,0,0,0,0,0,0,0,0,0,0,0,0,0,0,0,0,37,0,0,0,0,0,0,0,0,0,0,0,0,0],[654,1717,872,870,515,1286,291,2058,1182,2699,2749,1296,1199,981,1337,1391,1523,119,193,234,708,799,2726,4142,273,1433,2763,182,614,2462,200,207,519,858,1328,1691,94,267,2292,491,182,131,1586,1412,2352,530,1131,148,822,271,7764,223,1369,805,812,232,18],[523,1381,1551,1803,1722,1140,244,1350,573,2434,5780,951,2887,3149,4086,2557,720,1029,1183,1345,2331,3608,4079,2234,1284,2244,804,945,1535,3353,1254,557,592,1099,2194,4215,0,672,5945,1408,0,0,491,858,2162,1201,2616,1263,772,137,4983,2722,2279,1743,868,1229,47],[1079,728,768,1892,1121,1317,1030,1522,267,1457,1612,760,2626,2238,1507,2758,2589,2433,1705,1212,2239,276,1650,1314,1878,1091,603,990,1205,2825,1450,1500,321,1302,1321,4576,0,7147,1919,1298,2343,8,616,1082,2008,1819,2639,1550,1508,519,0,1107,754,506,2309,2276,0],[3845,5996,3840,9624,10349,11021,5910,5098,11485,16128,11321,4526,6082,7924,4763,34516,17439,8531,12065,9174,10122,10774,13024,6770,5732,9448,3094,3599,7710,15359,1752,4689,5822,6944,13780,12605,5415,8583,13596,6473,5367,4520,4004,3043,5534,8926,5194,7192,4444,5518,35771,7373,7084,4842,3567,3547,4504],[64,194,1406,0,0,1264,0,96,202,1083,443,751,0,0,0,0,0,0,191,0,0,137,0,2151,602,0,0,390,801,536,0,0,0,181,647,1316,0,0,0,0,0,0,1009,537,1540,601,383,1155,0,240,0,318,73,263,0,0,0],[0,0,321,0,83,404,202,156,0,0,0,940,0,0,0,0,0,0,0,57,0,0,0,29,0,248,0,0,202,0,0,0,0,0,0,0,0,0,0,0,0,0,58,0,256,0,598,0,0,0,0,0,163,7,0,0,0],[429,4593,1318,1124,1404,2896,114,1412,167,4982,917,240,1131,370,655,842,80,347,666,307,2494,368,0,205,265,170,158,62,250,690,30,382,366,301,497,402,456,1156,2041,197,1451,480,593,930,446,257,378,307,138,130,2018,717,1616,859,616,405,1603],[10112,21899,5849,11459,5332,21535,7589,5156,19885,14582,17654,5991,16392,10910,8314,5996,3145,3015,5956,7123,1596,51112,36161,20567,5421,30130,17609,8574,19326,56496,2253,18270,105371,49834,21518,40434,284,16083,7219,3588,7113,4735,25269,25192,12559,1922,3432,5450,1217,1114,2902,5036,13993,10139,3620,6428,16137],[105,161,0,0,78,118,0,94,114,267,93,302,237,0,87,0,0,31,0,46,98,0,0,108,100,0,0,0,86,180,38,0,0,62,174,269,0,37,295,0,75,219,104,104,326,44,0,0,0,0,994,0,384,82,0,86,0],[474,692,1114,2473,3327,883,1882,3722,2132,7298,4014,661,2653,3294,2449,3259,468,2141,6029,4996,3713,6240,3704,552,3392,1560,1360,0,3201,3571,420,2594,555,4202,4158,4565,1005,4246,599,4298,8652,133,3246,2251,402,5876,622,1684,3432,4371,5824,2311,1262,399,610,671,0],[0,0,44,200,180,163,156,478,0,402,392,0,0,208,101,0,0,0,0,0,0,391,404,0,0,0,2323,0,202,96,0,0,0,202,144,1926,0,0,0,0,0,0,401,0,0,0,324,0,306,0,0,338,0,1154,32,0,0],[0,238,1087,0,1,578,0,201,566,92,53,79,0,196,337,291,98,0,0,0,0,0,27,0,0,289,0,0,7,88,0,0,0,176,150,202,0,0,0,0,0,5923,605,192,295,0,426,495,0,0,2069,625,322,143,72,0,0],[8253,12393,3682,23481,16856,15709,14720,18735,8628,32277,30333,3420,18606,19433,16908,15803,4894,5909,27321,26878,38833,27993,19992,10117,18032,22267,14318,5389,32999,37206,11373,18814,11531,26507,40045,30542,20786,45562,28858,14304,8141,4705,14039,12633,3580,17629,11241,10035,12573,19041,36264,15665,22369,4286,5349,2261,10652],[0,0,535,134,0,538,166,6,331,672,402,386,278,0,0,0,0,0,204,0,42,472,0,0,402,0,0,0,0,0,0,0,0,0,378,0,0,0,0,0,0,0,0,132,1433,0,721,0,0,0,0,0,390,2222,0,55,0],[0,0,529,0,0,226,154,76,144,0,101,659,33,100,0,32,0,0,0,0,362,0,0,0,63,392,1412,0,764,791,0,0,0,0,0,1609,0,0,1795,0,0,2807,0,705,51,2,692,291,0,0,0,195,399,1365,922,0,0],[0,0,66,0,0,0,0,0,0,0,0,244,0,0,0,0,0,0,0,0,0,0,0,0,0,58,0,0,0,48,0,0,0,0,300,0,0,0,0,0,0,0,0,285,30,0,0,228,0,0,0,183,47,0,0,0,0],[33,805,1112,677,0,335,86,114,620,704,469,1665,37,604,0,125,139,0,0,83,1254,255,0,556,913,107,314,0,1568,263,0,0,906,0,288,2610,0,0,3182,0,0,8295,1154,433,2178,480,7182,1209,58,261,0,500,462,1302,52,0,0],[28223,73836,7973,119921,70014,53096,76818,82550,124403,257938,310636,14793,81413,89133,102654,94293,34928,44432,104989,78825,210264,139584,96932,115201,85952,234278,101507,7617,107352,178215,6698,63731,158949,67971,133282,125421,72473,176312,49793,104701,124377,6577,54823,47966,14070,66012,36409,21000,92490,59432,400340,62941,91001,13739,14620,22062,34003],[0,0,107,0,0,0,0,111,0,4,44,69,194,44,66,0,0,0,66,0,0,0,0,0,45,34,0,0,154,0,0,0,0,256,0,0,0,0,23,0,0,17,0,0,0,0,0,78,0,0,0,0,0,0,40,0,131],[0,0,596,60,0,0,0,0,201,1132,731,199,0,195,158,0,0,0,0,196,0,181,0,0,0,733,0,0,0,585,0,0,0,0,0,0,0,0,597,0,217,0,38,585,904,674,2012,1419,0,0,0,320,873,2379,0,0,0],[0,276,707,455,174,68,485,256,198,194,676,538,797,369,127,917,0,0,178,226,584,414,2273,1288,681,166,696,0,370,280,0,0,61,534,470,1469,0,1895,201,101,0,0,838,1021,172,887,763,264,151,0,1991,231,1812,454,64,81,0],[57070,91928,29299,137543,98373,183525,105081,126289,106220,298324,273345,49087,88691,149612,204193,139707,89520,126002,119971,155286,182775,160722,321390,366180,82294,259825,115993,59548,126630,269860,30181,80170,86161,117139,106865,189493,147975,260053,147750,89484,142046,94065,249322,73477,76831,92531,47389,81598,68934,55446,945269,116121,92755,53066,73125,58896,116259],[748,683,63,641,313,593,375,376,403,1291,715,0,838,764,428,688,12,596,768,882,1262,1720,618,67,357,931,36,488,585,1076,0,849,747,2128,804,1536,25,1344,0,323,685,0,884,1434,178,598,145,0,1583,2136,4126,571,727,64,50,0,0],[0,0,138,0,0,0,0,0,0,33,247,0,0,74,0,0,0,0,0,0,69,0,0,0,28,0,0,0,27,0,0,0,0,0,104,0,0,0,0,0,0,684,0,267,0,0,0,0,0,480,0,0,0,202,0,0,0],[7105,6349,3368,6551,5988,5334,3574,3361,2859,8450,7788,4759,2631,2553,2967,3385,1278,614,1830,6630,7496,6574,1786,6980,5135,5269,450,810,1477,2939,3245,3713,18209,5239,10667,2034,11231,5105,9825,2300,5790,10235,10231,4284,13098,5306,62,1318,4984,4861,113717,5016,4558,2068,4062,2378,11401],[533,403,3419,362,256,1405,1100,591,1667,2445,1902,2894,1594,1737,1999,685,56,1321,339,978,3858,1441,269,1305,525,2694,1670,71,1489,4662,0,680,262,1477,2584,2587,0,8809,1007,520,656,667,3013,1246,3351,830,7854,2782,874,42,4368,971,2759,5118,306,1513,0],[503,317,529,1597,881,338,899,1405,562,1433,2080,583,1238,425,1059,2411,1588,520,1008,430,1038,2340,798,6595,549,916,3459,5028,1291,1165,0,369,345,1497,5473,7213,1511,814,4879,1024,1635,579,1314,903,1063,1484,0,851,368,270,9138,1037,647,272,271,590,2462],[35670,28348,6926,114198,95565,67817,49858,57555,13893,159402,195481,6762,114912,78191,60820,94956,35180,56179,98371,67993,89574,178019,93566,39561,91932,86665,63597,16054,107948,98571,23738,69415,41591,91765,199175,159870,69873,115684,35195,83994,83680,7386,63051,31189,9804,79027,31636,28025,83198,51872,195065,66969,72264,9116,16877,24185,28911],[396,0,1926,117,0,0,311,403,0,536,735,2415,730,775,597,265,568,0,332,193,0,0,0,606,606,317,2205,0,520,543,26,0,2855,990,727,403,0,0,663,937,2199,0,1781,651,3337,0,3394,1511,0,202,1405,403,431,963,0,0,0],[2003,2339,8675,93,1279,1396,922,1113,1962,3241,1950,8421,1399,558,1089,2371,464,868,208,0,275,547,3002,5295,698,1690,3974,0,3402,3894,3232,43,758,615,3886,3036,28,323,5934,116,0,1631,4974,3987,12662,1433,15138,7689,986,740,14854,1228,3478,5273,818,503,221],[38507,32964,13633,82103,66319,77604,58704,60274,39660,148576,168086,11444,83855,65597,62585,69675,19900,48441,91716,70626,95113,114822,77600,52383,78419,86311,50466,14793,140553,91540,9570,42771,45798,86065,124334,105432,68070,157577,65419,72570,55151,15530,67761,36805,25717,81561,45756,33573,55849,46368,268215,63972,80957,29209,38810,28002,7893],[0,0,609,72,0,103,91,231,225,132,630,21,169,201,0,671,40,125,23,36,36,50,0,0,0,309,534,49,169,278,0,0,0,41,260,71,36,0,109,0,220,0,979,0,372,0,0,506,0,304,1968,231,199,919,0,92,124],[2170,2292,3825,3428,2557,4613,3211,1660,1718,4143,1747,4040,1819,1587,1572,1498,276,502,1911,1809,896,3366,1353,1689,1979,6153,4068,177,6057,5021,981,603,1041,1924,7669,4604,4990,6011,3413,1585,2789,2681,2933,1921,3442,4552,9229,1289,2695,909,22536,2162,3616,2975,1761,1187,1321],[9143,20061,10008,21685,21106,24850,13702,16707,22453,45798,32879,11598,20695,20112,21842,23687,13760,13752,20219,16721,19955,24678,27064,57516,33561,26324,11886,16889,19875,30827,10631,18144,12372,24809,33904,28380,21177,26326,38597,12419,14032,13483,18426,9804,29341,17046,15555,11761,10877,9838,118893,13936,19355,6352,13258,10548,15966],[426,945,658,2396,1261,1016,705,334,1126,2558,2536,672,331,887,277,854,0,1105,329,455,1160,119,112,2704,527,1479,335,0,1118,326,0,221,587,0,1025,1333,28,574,608,237,1743,0,541,735,2016,802,1104,460,33,312,6481,300,1286,0,509,0,0],[168,303,1095,414,88,802,49,302,402,957,305,779,0,245,168,1297,252,67,447,41,420,274,731,25,44,218,188,288,71,196,0,0,0,604,414,266,0,0,0,77,0,0,345,0,970,136,43,490,0,0,101,202,376,20,400,166,749],[1094,270,1184,253,178,850,146,0,602,712,540,1280,556,192,231,101,66,58,4,399,198,763,2030,3984,1408,729,404,106,591,976,261,0,0,374,644,2388,0,29,1504,180,508,0,200,1394,1598,934,1962,3659,0,71,47,788,1833,156,268,87,1548],[457,1106,33,1537,598,787,320,461,899,2292,1328,248,689,704,373,2107,55,254,378,403,1215,305,438,484,119,575,44,1251,690,882,0,131,397,1041,375,1275,6,3644,2068,533,430,3704,96,135,100,1119,879,162,659,147,3737,145,289,0,1017,791,4916],[0,257,812,365,0,0,96,1211,822,267,1904,202,202,572,2,84,721,0,1018,1150,3147,1395,0,0,111,1728,784,0,1383,1074,0,402,424,99,388,3102,0,661,0,0,450,0,1010,1296,0,285,1102,100,318,0,37,1300,773,388,0,396,0],[0,0,232,0,0,0,0,175,0,0,0,0,0,0,0,0,0,0,6,0,0,57,0,0,99,0,0,0,0,0,0,0,0,0,0,0,0,0,0,0,0,0,0,0,0,263,1788,0,0,0,0,0,0,60,0,0,0],[66,610,596,541,1232,1948,812,724,312,950,1492,643,345,134,255,326,196,94,483,1465,777,702,2859,1738,218,902,665,1871,1388,3367,1341,888,0,308,2031,1566,0,0,684,613,805,0,634,946,1702,915,833,1001,0,1842,607,154,608,576,0,407,0],[2948,4610,639,15737,8553,2289,10109,6556,1320,26202,46471,513,10851,6795,4626,11701,3224,5235,14246,5402,11527,19691,5390,2083,7875,14605,8012,3113,12640,6623,1997,5536,2933,9606,12705,4797,9329,10254,1935,6563,6658,1445,8948,13151,3333,12572,4621,3008,28915,17744,20407,6485,5584,1465,3780,2895,6296],[2376,2500,2395,2159,1327,2508,1125,1504,4100,6848,4091,1799,1173,1637,1588,676,20,263,2443,3104,4160,3723,10021,13114,3499,3566,3213,23,4345,6387,0,3287,4104,6980,4843,6930,2765,2716,5324,2382,2286,3880,5241,3953,2858,3245,1476,2636,1790,2571,31538,2410,2702,1020,57,581,164],[50911,84097,20042,101381,78972,101478,37821,47877,104868,172223,130345,39922,47304,62101,57366,49381,17156,33840,56484,73493,113145,73433,97742,213495,53182,96800,39948,13185,59063,112573,16312,49225,48039,57008,105609,159709,49056,119129,108060,30712,38499,23917,52647,54351,116820,71453,49796,49813,31508,23979,417093,36631,51824,9178,29640,30834,69961],[338,1913,6061,2798,2647,3666,1365,2523,851,4008,5239,3878,6795,2910,2861,5041,2184,5004,2476,1339,2874,3992,3362,4734,2126,5655,2675,0,7580,4537,6719,2933,1916,9051,9499,7396,0,3398,4638,1489,4878,5242,7447,3184,6645,2604,2487,6946,301,1044,3315,3752,6256,8958,2198,2365,0],[39354,59547,7474,71830,49559,59988,40283,44669,79249,131712,143813,13956,42426,48037,45386,62770,16793,39478,56341,52275,88114,72619,46243,40518,55236,119830,57505,17276,77095,112779,17462,50001,70219,48395,75126,77546,32780,114216,73057,46551,76190,6352,76288,41345,23643,38501,30982,33645,40964,37792,298816,38077,64949,11009,29221,27232,38017],[0,647,692,119,0,229,191,94,338,202,858,900,0,203,186,0,0,0,787,350,424,105,238,120,184,402,1043,0,587,268,0,0,196,435,770,1043,2322,0,161,233,0,1315,673,13,1312,16,704,0,288,196,0,636,74,289,373,0,0],[12308,17384,71911,12034,13853,11780,1463,3817,11914,32496,8613,25796,7732,4028,9930,7736,1795,4215,1980,1828,21843,3470,2996,5954,1259,2497,683,1184,2312,5926,1281,2000,3722,3539,5535,7366,4256,4816,17951,1893,7001,24730,3190,4200,32916,3819,3737,4523,2863,1286,19260,2080,3632,5214,11042,5243,12368],[0,30,121,182,0,76,34,52,0,48,394,201,0,0,192,3,0,0,144,473,0,0,0,0,44,100,0,0,0,0,0,0,0,0,0,0,0,0,738,0,0,0,0,0,427,286,0,0,0,0,1509,0,30,0,34,0,0],[7406,12484,1652,16334,9893,6086,7337,6577,21622,25286,22456,2945,5538,5379,5705,13143,5020,2996,8708,9326,19328,12209,7508,10801,7566,14381,4599,3738,10564,15684,8197,3592,10750,14205,10906,13173,13036,15984,8578,7372,6840,14,7955,6840,4027,15903,2672,1690,6332,4803,83228,5925,6625,652,4018,4101,6903],[0,0,302,67,4,26,144,201,441,166,113,140,17,0,42,33,0,74,0,0,96,0,0,300,0,0,138,8,0,202,890,0,0,139,149,709,0,110,0,0,0,33,66,172,39,639,1032,348,332,0,953,0,165,0,99,165,0],[16089,10218,11079,28187,18152,18463,12566,14993,29056,76713,33740,23284,15982,14651,21115,23150,9627,2862,25450,17721,24646,25391,47326,56431,19022,30817,20621,7331,30833,68270,21257,14762,23789,38150,35035,31829,27339,10805,21440,15997,14999,13102,22155,15057,39864,22297,16150,22492,11708,13067,262646,13660,15090,1765,9269,16649,11337],[456,716,532,889,533,955,690,205,842,2806,658,670,536,200,152,1216,8,661,1124,202,303,3306,2471,786,852,3103,2335,27,4492,2755,4639,490,1849,4114,1951,1490,435,0,122,732,1308,2,2200,2946,1675,675,5671,678,2467,1121,4440,1299,2110,42,177,655,0],[90,379,999,394,413,306,215,442,130,1033,572,1419,1077,677,419,2181,2163,902,139,249,212,490,144,1160,245,239,808,286,1374,778,1536,184,162,228,2282,583,0,80,106,0,538,0,281,1747,2966,401,967,1537,137,6437,4938,511,731,1308,604,160,1334],[54695,70925,13606,159292,111238,35170,5239,9809,5429,73350,74124,13060,23095,16412,23164,81115,9794,8286,34112,8747,80162,12256,8707,13865,94348,102754,40899,6580,14369,11897,6055,5547,7914,17874,35849,24048,1188,14643,7199,17390,22019,15329,18913,12186,23233,115123,42582,27804,20971,17432,109784,4045,16487,14155,27395,25603,79424],[0,0,0,2,0,0,0,0,0,0,3,0,0,0,0,0,0,0,0,0,0,0,0,0,0,0,0,0,0,0,0,0,0,0,175,0,0,0,0,0,0,0,0,0,0,0,0,0,0,0,0,85,81,0,0,0,0],[8,204,826,404,118,5,0,0,0,119,117,114,0,0,0,0,0,0,202,0,0,89,0,0,0,0,0,0,562,741,0,0,0,0,0,198,0,199,0,0,0,0,45,2,298,0,284,0,0,40,0,0,0,179,1195,0,0],[1219,576,1751,2416,3187,2006,1355,1736,0,7467,6059,1450,3686,2342,3433,9185,8782,3710,2315,1654,4628,1744,651,444,2961,1646,1481,691,2073,1772,13116,824,149,924,3657,5009,318,2632,5907,3901,1745,2775,10975,1118,431,2481,1267,3970,2096,867,9301,2680,1779,1515,456,2382,163],[3237,3931,4104,9675,6364,2621,2906,831,1547,15191,6934,4792,4235,2276,2839,2394,4807,4261,14960,2946,770,3872,11681,3220,1085,3879,651,2461,7093,6227,0,1071,1043,6110,9634,15451,13692,1572,3954,6196,222,6536,22965,4479,8200,2405,9440,3025,2095,3040,30884,13745,9053,4040,2007,909,0],[6249,5685,3398,10452,5850,11435,4250,4865,2814,11044,17173,4394,11164,14660,8311,16085,8460,8914,4028,3696,12632,5670,8811,17507,2421,8407,5706,10982,12655,11441,143,5143,10770,8238,14478,18749,3711,17864,7861,1974,8550,6907,3388,6624,5887,3072,7171,6893,4060,1560,31733,5438,8619,2363,6967,8355,4739],[636,382,1361,199,403,421,100,265,758,1007,1192,1397,672,525,382,741,61,223,28,0,262,605,0,1936,0,698,426,108,101,441,152,895,29,762,968,1364,0,35,2056,209,32,189,260,1000,771,173,1086,4242,293,297,441,1124,637,2311,635,25,0],[0,0,32,43,0,0,0,0,0,0,0,280,276,148,0,0,0,0,0,0,35,0,64,0,0,34,0,0,37,0,0,0,1085,0,362,0,0,0,0,0,0,874,0,208,184,0,0,35,0,0,0,0,0,89,0,0,0],[21737,19857,3835,49964,32725,22740,18784,18356,7444,75290,58632,7485,19290,13027,15737,45357,15137,16304,27867,19829,30986,28438,42601,52624,25514,53253,27430,5233,19619,37354,9785,20448,26330,31152,31966,40583,27198,28353,38470,21824,7949,7608,16138,25547,18755,27643,14390,14120,17830,33145,155225,11987,26520,2732,22624,20790,46696],[10272,14371,7885,13033,10705,13900,5571,7521,20645,26907,23419,9163,9074,9543,11866,16454,7105,4252,8399,9418,10134,15720,12815,23072,10356,16378,4669,2788,12652,20071,616,5730,8217,9007,13893,23478,10180,16746,29556,8408,6511,6256,10968,8738,21363,9253,6363,12240,20651,12434,91067,7033,9361,1360,6896,8240,4722],[632,458,101,103,601,304,559,307,729,1696,1953,793,77,245,518,1427,345,278,502,659,696,1419,1630,2206,101,637,303,63,777,495,0,101,69,1871,741,2625,431,1188,1795,268,519,502,201,551,973,1318,876,610,1010,562,2462,1243,476,38,1099,1017,17],[1109,947,7302,382,481,2873,601,1240,1262,1178,2738,3577,2259,661,1378,1642,37,640,761,202,578,122,1,1323,649,3248,2135,193,1981,4311,357,379,1391,540,1978,2507,0,1119,2607,0,324,8234,3440,2294,8845,1007,9766,4805,159,0,6614,670,2895,3599,899,501,49],[1733,3292,1107,4028,3867,4581,2596,2827,4527,5715,6888,1018,2052,3441,3547,7181,3748,4592,2828,2224,4158,4512,3404,2755,3135,4355,2417,674,2501,2480,57,2851,2765,1156,4927,4065,5548,4697,4773,1052,2369,1011,4169,3893,4599,1651,1954,1192,1849,3137,21574,2060,2533,2161,2183,1628,528],[51911,96696,13253,89948,66887,59308,44040,45329,117060,273083,127227,27480,40849,37764,40917,77621,25639,32572,98304,61042,127495,92296,86281,177170,49557,87129,25770,31098,61773,129303,19004,56393,44505,81863,68705,72069,65748,84592,57619,44103,56534,29450,79732,58534,61299,124845,46162,42122,69386,45897,446519,49758,43258,11848,56814,46136,67575],[0,0,0,0,0,0,0,0,0,0,0,0,0,0,34,0,0,0,0,0,38,0,0,0,0,0,0,0,0,0,0,21,0,0,0,0,0,0,0,0,0,0,90,0,0,0,0,0,0,0,0,0,0,0,0,0,0],[14961,14449,3352,19234,15974,18970,15450,22157,12429,38250,40154,2721,27817,26154,20547,28310,9856,18959,20495,19683,31111,41392,32606,28149,15965,26294,17731,2193,43814,56771,1218,17207,15256,26601,46435,57815,24894,48888,12539,21352,35508,2543,15872,5838,7548,16745,16038,5195,13475,12406,43813,14436,27696,6981,4724,7871,9061],[43870,47833,18589,67588,62010,87323,46136,56793,54746,101931,176444,17286,98107,124957,102145,127982,47486,80878,61379,68057,88897,111039,91375,82205,62253,82103,49257,35392,103110,98274,20524,56790,48359,99481,246568,191970,35298,169842,90526,62875,67713,17214,33845,21939,37565,64206,57533,37739,43824,44449,287292,48666,87959,19985,38302,36956,15479],[3,0,391,47,0,0,0,0,608,347,274,528,10,0,57,0,0,0,184,0,101,0,0,270,116,145,634,0,344,69,0,0,0,192,101,1,0,0,0,0,0,0,652,429,282,332,2783,607,0,0,0,577,17,347,0,84,0],[0,436,495,159,225,296,0,0,136,1185,456,562,360,0,746,745,248,188,101,96,178,66,166,188,0,50,279,49,240,328,0,0,0,121,264,929,0,0,36,196,106,1299,151,220,267,44,1488,1633,0,0,1321,49,129,378,11,0,0],[398,312,842,1668,1239,717,1206,2294,668,2348,1768,1390,907,900,1684,2462,613,620,2062,648,1207,754,782,4189,828,499,272,0,590,109,0,999,142,701,936,1890,0,1185,206,278,8,1569,147,912,3539,386,1454,223,1549,1041,13213,403,599,335,1385,1092,0],[97,509,209,192,384,767,197,67,0,2158,506,166,67,16,0,3151,65,514,420,324,446,935,0,140,0,90,0,162,151,471,0,59,299,383,700,394,0,39,0,97,124,0,302,39,43,358,393,765,0,90,0,200,290,0,346,377,201],[16436,60340,6554,59568,62179,42319,50221,54780,115122,144986,193308,22959,27812,32814,34611,16160,1254,6683,76638,43260,200227,60283,62539,122799,29600,72072,44193,1040,46611,74185,5196,14466,32150,40703,98253,140187,38982,119084,27428,31471,37445,1876,23340,31178,31478,36339,11554,21087,38358,37285,370889,15420,51826,8568,5212,6259,14679],[1327,1548,175,905,895,554,1266,2017,1905,4578,4060,945,1312,1610,1220,1771,577,450,402,1140,3306,3765,1725,40,1438,2709,2156,554,2494,2319,909,3702,2276,3449,5586,5149,0,3808,193,460,2653,0,2377,1711,755,1000,996,0,402,1412,0,626,2852,404,701,1206,151],[727,337,2475,484,1226,1559,553,456,4999,2882,2867,2508,1548,658,155,84,0,1087,617,593,948,3243,14399,6715,979,2312,1546,347,2279,3492,168,828,541,327,1123,5050,2014,2420,2059,698,189,615,1905,1803,3956,1173,3732,3211,1555,2437,13710,1252,1604,1665,314,624,0],[816,948,4709,11447,6870,4614,5357,4210,888,21893,6078,4635,4745,6857,5474,32968,8843,8001,7977,2681,6558,10493,8871,3210,9633,6521,6801,53,18916,7965,2480,1284,2984,5760,11294,9501,1235,4241,603,1865,2238,0,4109,3798,6435,5522,3768,2960,8466,13204,24562,7383,23742,7660,2576,6832,3875],[31354,52181,7724,53953,41414,51027,37905,30783,45415,113737,100642,7726,49153,45670,55027,66911,25327,38759,50452,51361,66896,78720,35001,30676,55774,98053,54419,20479,126470,91612,28000,70742,37030,41917,86975,92226,38281,101311,16020,47390,35915,222,143606,43275,7307,34878,31542,21805,55717,45426,262498,46238,62343,10153,12502,15934,30335],[33425,43504,7392,98459,63383,47188,46336,68444,6766,133556,204928,10396,89541,85916,93905,136820,49762,57665,93079,98582,186938,70855,89729,25063,62760,92855,37996,21986,72261,34919,26461,97717,28112,94902,183930,133602,79119,211497,21584,82990,96127,3392,63172,46864,10799,49427,49789,25811,53150,46064,99152,35403,56354,7117,25213,27776,22747],[1823,179,2618,6420,3569,3207,3066,3031,1036,4507,4676,1294,2627,3121,2483,6557,957,3032,6869,2635,2744,6137,4022,6488,4572,3495,806,1347,5692,4518,56,1478,623,3182,4853,8236,4433,1332,990,2089,1421,123,1551,2035,2236,4330,3482,3601,1325,1001,4496,1779,3377,2135,836,2301,91],[40,101,0,56,150,156,202,0,0,67,224,0,215,0,223,435,0,82,0,0,160,0,500,0,331,251,123,0,275,434,0,0,0,0,0,374,0,944,210,0,0,0,6,0,789,398,0,565,226,126,0,93,160,0,0,0,0],[15659,18884,4386,28229,23400,28979,15885,17647,28600,59491,47576,5819,22511,35188,36213,20517,5416,11605,28088,29300,29116,30062,19567,15961,21209,47838,22084,2734,34626,53062,3734,14101,19467,26663,42275,52104,14791,26613,12420,14349,20542,1358,16098,13296,9787,21093,12581,11133,13620,12942,100131,18545,23115,4620,8182,7636,10076],[2993,2435,3080,3357,3050,7119,1798,2504,4270,7605,9861,4422,6357,3604,4474,6864,930,5073,2101,3841,7805,1479,496,1737,1981,3354,2980,956,5238,5254,4172,3737,4778,592,3304,10660,2,18080,2951,2422,5664,4745,2559,1320,3246,2701,4633,688,791,2744,26780,1648,4562,4128,2128,1711,3614],[0,25,722,0,0,0,62,0,303,400,166,998,79,145,0,0,0,0,0,101,0,0,50,1008,0,520,617,161,207,300,0,0,0,183,470,1023,0,0,0,0,78,0,0,98,706,78,1232,188,99,0,336,598,89,205,73,104,0],[15695,17932,9643,25808,19749,25290,12355,14986,24619,46248,53820,11793,16978,22093,23217,20339,7245,6852,17220,13031,19731,25983,31047,44125,14379,25870,14693,5192,24981,36787,12893,11787,10469,19620,29256,41854,10073,8987,45644,14290,11804,10752,12560,8543,16661,23459,7586,14712,10157,9048,139472,10421,16155,4023,15132,14279,20398],[147,0,414,4,0,0,0,0,0,0,404,1509,0,57,202,32,0,0,0,0,237,0,0,0,0,0,133,27,0,0,0,0,0,0,356,0,0,0,0,0,0,0,3,31,1375,512,1764,254,0,0,0,998,183,992,0,0,0],[58,26,1316,0,0,0,174,41,0,407,778,0,202,0,190,0,0,0,0,101,25,201,0,936,404,40,0,0,301,559,0,0,96,1001,0,1871,0,669,0,0,0,0,0,765,202,24,1860,1226,0,0,0,0,434,0,0,0,0],[987,447,916,1253,1150,1003,197,791,243,3278,1217,1209,699,586,1048,6740,934,3598,825,838,1229,1122,858,243,267,1179,36,4096,528,1415,7378,143,237,521,562,3149,128,512,723,375,737,882,255,348,2541,939,2025,93,383,0,756,288,498,287,1717,1185,808],[856,1064,29,4189,3541,3126,1450,1164,1496,6977,3776,510,518,933,1335,14682,4960,7981,1367,801,1823,1838,1231,1560,561,521,379,4256,880,1354,246,609,2626,1610,887,2655,2297,3934,1588,35531,3962,3351,4488,1376,1327,2791,1639,21529,1847,639,12898,768,604,305,802,8216,1345],[33300,33966,9729,50955,35675,31683,32597,35646,10408,84952,42696,9381,44838,41954,40397,82498,26703,36617,38615,51042,86504,41191,33092,27569,33758,46311,21934,16431,27074,31542,18195,26990,16245,29042,38222,46112,34324,77570,63742,49371,30627,5953,41530,30132,28294,45742,26707,26408,37438,23640,135826,24896,32265,7584,24137,17663,22900],[101,68,1149,0,119,116,40,56,251,57,48,1243,401,0,202,0,69,0,71,0,0,806,0,108,0,0,445,0,307,534,0,0,0,0,0,136,0,0,0,0,64,2982,62,163,72,0,363,685,304,0,0,220,1531,568,40,0,1203],[3066,1787,7059,771,1007,3021,1147,1747,516,1396,1776,3698,2171,1294,2016,4310,1006,1680,1098,914,1187,2165,358,1831,2831,2729,4511,12323,3797,5919,6471,859,2723,2946,5984,5114,5742,2017,5692,321,753,9761,4901,3639,5398,3530,12930,5536,889,0,16067,1912,4720,5087,911,2295,0],[167,67,993,0,199,392,0,156,300,201,258,581,48,0,216,438,582,0,62,0,155,0,0,128,139,460,0,0,848,298,0,0,14,0,609,366,0,0,0,0,0,0,190,441,1080,0,737,494,0,0,2631,219,640,930,0,0,0],[425,168,3526,319,905,910,1172,531,1680,420,1298,988,555,1562,565,2958,475,0,202,175,665,1307,0,4787,2393,2493,323,123,2227,1767,0,573,187,288,796,1230,0,0,404,1062,1204,0,2441,834,3473,1798,1744,1184,0,215,10543,293,798,2648,139,0,0],[307,315,2497,41,787,489,323,202,235,268,623,1058,336,387,219,137,0,0,0,0,202,0,0,0,398,968,1449,71,63,236,0,132,0,0,2327,0,0,83,24,0,0,1043,1546,1351,2098,462,1326,770,78,0,1322,1090,554,428,38,320,0],[544,907,696,153,503,860,196,618,822,1377,602,873,1374,1336,1094,974,1812,681,443,210,119,103,1163,1065,842,1764,725,30,3608,4410,45,193,381,1716,2269,218,0,4745,685,270,128,864,147,176,2232,413,1450,4881,324,177,6209,1189,926,1134,61,478,3467],[11928,15479,3388,19436,13196,16446,14358,12894,7980,43094,42670,6084,23412,22582,21559,26975,6730,12792,19989,19499,36037,20395,18805,9956,12610,27145,13770,2181,26041,20282,14911,10450,9286,17898,40080,28962,6325,39946,16009,12484,17474,5527,7354,6052,8043,14942,12842,6235,10417,6701,69252,11555,21195,8804,3160,4863,17145],[101,27,157,0,14,430,13,111,159,103,62,695,56,96,301,282,0,77,0,0,0,0,599,188,0,215,0,470,1,341,0,44,92,0,187,32,0,0,2213,123,0,0,91,100,1444,0,0,248,0,0,166,116,159,97,594,0,0],[894,139,1353,417,0,327,868,100,763,542,515,1306,227,57,0,187,527,223,1041,0,180,523,0,0,624,581,1571,0,1577,41,0,0,991,728,419,27,0,0,1475,896,0,228,1365,1080,2267,94,2031,2385,0,0,0,375,392,441,448,223,0],[796,263,1719,792,880,928,184,1436,1752,2222,943,2467,486,636,920,2921,539,0,964,150,0,404,522,666,1599,5554,1942,302,2662,4804,34,196,11673,1527,4197,1545,0,1492,3246,883,46,3499,4568,1211,5630,2177,4257,1596,396,686,4117,933,1716,6446,746,1077,24],[1130,2009,845,2259,3326,2214,2649,3081,1592,8472,3908,2201,2628,2247,1702,4820,950,1067,2764,2019,7117,1558,1141,2063,2241,2729,3375,0,2988,4247,0,644,73,999,2530,2830,3274,6223,1064,2963,849,0,5571,1820,1836,3519,1558,6315,1323,1837,35340,1840,3698,1178,1205,800,0],[82659,118273,7938,248049,163714,112998,245959,210840,69439,508597,661990,12165,282792,126067,121110,375655,190335,198866,397725,226894,357209,855577,188032,109834,277047,371902,129334,73947,246555,291703,63145,212299,271901,335816,275220,245743,237817,400707,55811,217055,192299,10949,171202,98099,18450,235300,82152,36979,192291,99463,458067,170060,207787,15073,64558,72658,112590],[0,149,882,806,606,513,98,321,2,287,720,0,1748,1017,956,361,0,333,402,55,2537,715,8,0,198,131,905,0,1152,1045,0,1310,0,2,2367,1608,0,5259,5985,0,3294,0,37,0,0,136,800,500,639,0,0,152,1842,77,0,0,0],[3684,9170,19722,3703,4895,4569,1343,2949,2009,12379,5432,4903,5494,3392,5285,1965,1011,690,1061,501,2357,1294,3189,7712,1191,1737,114,1401,1323,2170,543,1025,1897,1492,1807,7527,4863,2416,12454,1528,2175,4517,2750,1804,6269,709,963,961,464,37,5859,2227,2860,8023,3582,2257,4821],[35657,15163,8077,92625,73982,60489,31938,50509,8081,108636,164125,26079,60509,69827,51773,69520,10638,34557,26661,34847,231296,60943,91496,116541,22446,48278,25250,17362,110271,68756,16349,51459,35966,87489,281249,290232,51336,146585,125579,32622,63612,5509,32263,45892,20961,26939,42913,46816,31526,29515,95196,33228,81426,3651,19992,23566,9968],[138,199,1407,576,183,349,499,537,178,637,106,420,340,376,272,1095,0,0,0,0,0,201,606,0,0,606,677,1459,185,618,0,0,0,0,630,506,0,0,0,0,35,0,1011,395,4197,0,2486,1711,0,400,398,0,745,1926,377,34,342],[18880,43058,10838,24058,16464,31752,11113,12775,95513,52457,27633,9234,19675,77357,73126,20558,9431,42550,7511,37611,23037,20522,42256,19431,8625,47877,25882,16950,59546,191988,8004,21020,57022,14081,31650,48814,4829,60677,30497,11300,18608,12843,23426,17232,32045,10162,27623,20521,10539,10327,180116,16644,15684,10991,13093,16220,59099],[398,0,76,66,0,50,0,0,0,0,118,265,107,0,92,0,0,0,0,0,0,0,0,0,0,202,0,0,3,0,0,0,0,0,198,0,0,0,0,0,0,0,0,641,0,0,202,0,0,0,0,0,0,0,5,0,0],[66,167,960,0,217,0,36,0,80,92,18,890,162,202,0,0,0,0,46,0,10,106,0,0,0,318,0,4,196,92,0,0,76,0,190,0,0,172,47,0,0,0,342,528,0,0,166,0,20,0,0,32,50,378,66,214,0],[24242,60428,6157,30176,27692,25549,31355,21709,30620,100369,139792,4569,28187,19894,22525,30695,8131,15998,45799,48547,110001,47291,39956,23620,63417,56987,29689,9772,56477,51302,23175,31752,17801,34863,60327,46955,14596,54296,11510,38773,43009,617,55414,25630,7511,40124,9831,11348,32929,28696,106584,26083,35285,4323,11614,11024,12166],[300,2725,1254,3447,2990,3175,1269,1923,1838,3845,2473,861,2056,2925,3549,582,384,1888,1805,1085,1764,3494,750,2523,1839,2419,700,195,2365,3827,2552,1370,1175,377,5078,2649,215,1470,94,1133,232,801,2586,1644,3590,4100,1449,4146,997,1372,10759,978,2533,134,320,734,644],[0,93,255,0,0,338,0,463,0,0,0,897,0,160,0,0,0,0,0,0,0,0,0,1212,0,0,0,0,755,400,0,0,0,0,0,777,0,1004,0,0,0,0,200,602,0,146,308,0,121,402,2216,742,202,175,0,0,156],[10509,6870,5007,10331,9495,11474,4261,4979,8112,12372,22667,3606,11201,12327,14252,11097,3158,6309,9986,12016,35180,19392,47817,29927,7754,12190,7684,3693,18379,34097,5566,3740,9070,12930,27437,46682,5936,20970,8354,9436,19309,1430,6647,7711,6206,12855,16686,9042,6561,14766,118508,9076,16954,4804,3580,5025,95],[869,608,186,781,788,618,909,636,692,1975,4378,218,681,661,1164,2213,185,1603,2141,484,3038,1056,687,720,981,2044,505,177,2529,3606,0,835,306,3574,1949,2273,2561,1563,0,625,206,38,964,431,175,0,148,1482,1443,40,20,373,1558,0,12,17,51],[132,1249,657,347,177,1026,81,241,32,2075,205,23,132,202,225,842,30,296,26,28,624,157,0,771,1,48,0,248,176,349,0,51,383,0,164,287,146,251,257,118,304,54,644,246,111,222,105,264,25,0,396,152,288,72,134,434,256],[33787,65342,29637,44388,29544,39397,25521,27211,86737,122234,73925,32240,30307,40637,39792,58811,31573,32721,29988,25969,55485,48200,49769,94459,27203,59852,22460,33125,56322,83103,10366,33633,35697,44633,51114,65804,46313,59156,75659,18377,33275,34403,52128,25527,47255,46962,22857,32622,17556,16683,333207,24051,26282,10364,33774,21610,32392],[27697,50615,827,83756,60827,37399,32475,23087,32212,126347,90467,6287,27732,27725,27398,26248,17966,17185,46030,28529,88606,33802,24609,19130,49987,104751,41633,10144,45800,54914,15360,67963,44555,33032,66749,56636,51249,68386,19357,31067,44641,8357,126128,32628,14818,35569,42108,15672,38156,19526,241798,34787,29176,4752,10323,8562,22296],[2537,574,13439,42563,17778,5208,443,1039,2490,17173,24342,6614,5687,3826,4594,7099,1303,1644,4813,6598,10500,302,1373,1294,3381,6897,860,925,1800,1061,630,4140,3905,3091,10449,2212,1007,775,1612,1210,811,395,798,2831,23753,1227,2135,2807,7895,4218,18093,8891,501,520,1841,887,5397],[0,102,0,0,0,0,0,0,0,216,0,0,0,0,82,0,0,0,0,0,0,0,0,0,0,0,0,0,130,0,0,0,0,0,24,0,0,0,584,0,222,0,0,318,365,286,0,0,38,0,0,0,0,0,0,0,0],[0,227,1136,154,232,229,217,111,862,665,1020,2351,484,0,0,102,45,0,0,0,0,0,96,28,0,344,977,39,0,1036,8520,0,30,0,889,1706,0,93,455,0,118,182,201,1176,3073,0,1797,894,0,99,103,772,202,755,309,0,0],[117,377,1448,155,398,540,560,546,306,116,270,501,31,0,75,432,27,768,21,15,15,248,0,1166,192,15,0,89,384,239,0,0,0,464,748,379,0,0,182,719,0,0,403,155,1550,264,3152,1248,296,0,0,256,171,215,875,0,0],[200,0,673,0,0,242,320,123,202,132,402,1341,94,202,191,97,23,200,312,0,182,404,0,984,0,480,993,0,639,606,0,0,780,0,930,1560,0,0,0,0,0,0,864,531,2893,202,3465,357,363,0,2427,129,1222,2124,991,80,0],[0,137,81,0,0,119,0,0,0,0,0,140,0,0,0,0,0,0,0,0,0,0,4,0,0,0,0,0,30,714,0,0,0,149,0,0,0,0,0,0,0,0,0,125,100,0,0,1269,0,0,0,0,74,0,0,0,0],[1309,1360,2897,480,252,3259,703,179,502,1445,3057,2325,1438,758,566,2128,483,834,23,975,1248,1614,1622,6237,805,1629,1404,3495,2434,4731,240,640,1158,584,1000,2788,124,5026,4043,594,674,5626,1946,1864,7004,1500,5865,4410,252,590,13065,2207,1888,1905,1634,847,87],[0,0,1183,0,0,720,0,0,0,0,337,201,0,74,0,125,0,0,0,0,0,0,0,0,0,0,202,0,0,0,0,0,0,595,201,0,1539,0,601,191,0,0,0,0,775,299,2358,0,139,0,0,77,346,709,0,0,1495],[8260,6442,3778,12783,12360,16236,7347,6729,9981,20419,25922,5205,13876,14032,10838,12949,2062,4749,14299,9518,23092,17453,24761,26246,5147,14446,7783,6419,17990,19171,12164,11788,7418,16124,29947,26422,2620,29912,16067,8918,9310,10137,9532,4648,8225,7857,19233,10354,3296,2979,58748,4194,7664,6690,5964,8019,2996],[5114,1551,1844,4460,5536,4885,3333,3602,5045,7650,6688,2755,4643,5597,4203,7073,753,2489,4205,2124,4618,4374,16049,16035,2039,4546,2320,2489,6968,6910,1724,1040,274,4479,5229,11535,496,6420,7689,3316,5123,5503,2390,1364,3770,1469,3131,944,2676,888,23818,2067,2524,840,1684,3445,1655],[375,761,135,654,278,947,371,72,243,728,820,120,226,298,999,1058,418,846,525,181,726,1020,0,410,0,425,249,63,256,1576,0,277,0,306,253,213,72,688,1270,0,1101,36,0,127,0,0,0,135,0,0,253,1321,859,593,103,279,89],[143,202,492,186,0,166,137,199,0,606,0,508,0,0,183,0,726,670,0,0,203,312,0,89,201,346,0,0,399,0,0,0,0,17,204,0,0,0,0,233,0,0,159,1695,701,0,1158,600,0,0,805,84,140,300,243,0,0],[1835,7769,3242,1495,565,1176,1037,871,12200,5878,2049,2169,1251,3186,3654,972,445,6098,1520,138,1469,1352,5144,3412,1876,4600,2229,555,12211,11628,224,208,757,2481,3048,1277,91,8727,2851,359,2655,2891,1826,2168,3773,0,2556,1636,294,1186,28602,586,1113,4721,1387,4238,19527],[1218,1244,612,1814,1166,998,1242,936,202,2140,4300,677,2137,2002,679,5040,1472,1040,507,1747,1725,1778,1806,1146,1880,2104,393,1330,3357,1509,0,410,393,1957,2102,2145,0,1059,0,864,2687,0,703,1034,749,1197,48,683,752,608,0,911,1263,401,566,1117,0],[0,164,804,0,0,0,166,149,630,606,517,807,190,0,168,1795,0,0,0,397,200,0,0,0,200,107,1562,0,398,399,0,0,0,0,201,1818,0,391,0,0,0,4576,965,301,2074,199,1206,1989,199,120,0,194,382,202,238,0,590],[362,1581,537,1980,955,2080,635,1011,2855,2354,923,731,2974,2336,1499,2361,74,1715,1434,1248,2309,196,1885,0,1622,3749,1672,700,3595,6620,0,3393,1554,546,1822,441,75,5484,486,547,1841,0,4702,2008,800,0,2600,562,1164,642,7925,202,2234,860,56,0,0],[2864,5606,9604,3590,4277,5967,1741,4728,6838,4932,7370,3262,5322,3902,4539,5232,1315,2935,3354,3142,4768,5589,7235,7033,1370,3305,3684,188,6122,9905,2630,2694,4897,3880,10691,8151,6253,11404,26761,1973,2683,6001,3250,4628,11361,3879,4754,11331,3738,691,57939,4388,4451,3512,3497,878,625],[0,0,0,0,17,74,0,0,0,267,117,217,188,0,193,0,0,0,94,35,0,135,0,0,0,225,260,0,0,63,0,0,0,45,0,464,0,0,256,0,0,0,0,0,0,0,0,0,0,0,0,0,0,0,0,0,425],[2627,3190,2977,2966,3327,6179,1478,2674,5114,9158,4486,2136,3897,4303,3006,2154,1507,2070,2646,2917,6950,2606,1392,3222,4234,3143,3707,2104,3103,9129,556,1168,2277,5106,4930,7919,1576,4610,1828,1668,3047,0,1704,3118,3824,2854,5207,1279,3171,1462,6145,3152,4680,1719,987,1314,2714],[1248,5570,8360,47,301,2677,980,948,1392,1703,1631,5713,1924,2092,470,285,49,162,360,928,713,2001,4670,11714,1186,1839,3715,1109,1910,11373,2053,68,2353,643,3556,24438,1204,1748,4189,0,0,3382,4009,6118,7559,1406,13647,3014,1398,2270,5577,3750,7015,7646,1340,352,0],[126,164,549,0,446,163,197,324,326,118,0,184,227,23,405,591,50,108,0,0,176,0,0,643,190,709,0,89,835,169,0,0,0,0,1063,0,0,0,0,0,0,0,543,365,859,919,1458,10,0,32,4006,0,19,703,69,15,0],[0,57,677,0,0,58,225,188,0,154,841,217,188,0,0,30,0,0,0,0,0,0,0,0,0,49,38,23,118,158,0,0,0,0,229,123,0,1624,0,0,0,0,102,581,1696,0,0,1760,744,0,948,32,85,941,0,0,86],[94647,233600,26523,259773,197022,248939,300845,313441,187125,781934,661692,48865,536046,271130,222012,590576,346807,400991,389903,271136,701184,573719,261397,240358,364308,703563,259816,326474,548810,844120,64951,295852,272529,378331,362576,335833,437016,825134,138962,329376,570375,49450,484676,209547,86637,482779,142559,90094,445749,250275,1405221,267058,209044,39672,117127,220462,294658],[808,558,5459,0,262,1264,411,1005,1216,926,1781,3035,877,623,1490,376,25,43,202,182,1629,530,1523,1212,477,974,5340,2789,2659,2553,0,0,40,207,1456,2754,30,40,342,0,0,1153,2145,2113,4361,906,3123,3238,497,0,6336,997,892,2932,912,608,0],[326,0,898,0,297,8,0,96,8,0,281,202,215,46,200,0,0,0,0,0,32,202,0,594,189,0,0,1038,553,0,0,0,0,202,308,220,0,310,999,103,0,0,183,380,2540,724,1194,1913,0,67,0,745,676,295,44,0,30],[179,2223,5583,734,395,1054,839,552,781,1408,2498,6232,938,589,1076,1027,556,516,568,276,402,153,119,6376,617,836,3987,501,3058,1701,574,0,630,719,2133,4877,48,1347,5794,275,1977,2375,4863,2269,4902,1533,10952,6994,94,0,23365,2211,2482,8580,1432,188,194],[29187,33778,10939,57005,48844,67181,43659,44044,55000,94945,122837,11153,64563,46007,49807,63295,30571,43118,34673,39256,96865,63188,64072,35438,41312,79297,55013,12862,61579,115874,27621,50990,45730,57703,105843,88712,36261,142053,73086,41338,72521,6882,29304,24892,16270,32300,31713,35960,53790,39387,183669,29185,40439,10212,24985,28746,16355],[1330,1197,1387,791,262,1444,444,662,2711,1011,1415,691,997,351,481,1772,658,3353,0,976,982,202,196,3573,802,2622,412,4459,2078,4266,0,397,606,288,350,4110,0,2598,4696,222,727,31,376,944,3340,310,3211,336,596,998,224,2182,1094,1302,198,51,4975],[0,106,0,24,38,89,0,0,0,0,0,0,0,0,0,28,0,0,58,0,0,0,0,0,0,290,38,0,0,0,0,0,0,0,0,0,0,0,0,0,0,0,0,0,0,0,389,0,0,0,0,0,0,218,0,0,0],[1375,100,595,242,1199,842,263,283,331,114,2557,1041,620,123,0,1233,370,945,150,763,508,247,3529,867,858,0,642,804,1171,1977,0,374,288,628,1527,736,0,647,176,311,2112,11,126,148,561,1315,201,540,641,135,125,896,1079,0,66,428,98],[733,3569,1424,2059,1347,2235,230,1207,614,4427,5049,694,1262,2347,2033,2050,393,614,679,9390,3752,409,606,387,1960,5711,2186,0,1904,4,0,1428,1896,1150,3232,1804,0,6354,2893,919,553,2,956,1021,661,467,2726,1201,2569,834,13994,413,1644,1286,396,1618,1057],[1665,2110,386,331,2465,3174,110,1013,309,1331,2053,282,1766,613,2939,78,32,301,0,348,2986,34,8639,4012,62,3976,3091,2237,26079,2833,2250,990,240,5443,12238,27481,24,457,975,0,2242,80,903,172,1244,33,277,204,329,242,2933,1671,10583,343,0,1108,0],[1801,2678,173,1863,89,1288,3621,1127,4172,6082,3099,577,1758,1841,1788,7914,2546,9809,1206,2007,1010,3728,4125,3037,4264,12617,4247,6544,4348,10250,903,2781,4392,2313,3166,4618,39,13783,1629,1912,4205,46,310,932,536,903,1496,885,1262,964,6832,1845,1788,686,1015,3220,8274],[0,840,271,689,179,1334,209,250,0,290,753,47,337,341,1093,58,5,20,134,144,805,720,774,1124,0,177,158,0,598,2093,0,170,32,0,582,456,0,1642,0,474,90,0,366,0,306,344,1310,101,0,0,0,0,0,65,11,625,2010],[186,7,259,178,333,285,0,152,122,183,173,340,237,0,193,831,80,96,0,20,81,7,0,12,0,27,0,84,26,88,0,0,7,18,252,165,32,14,14,237,120,48,300,0,1206,0,432,87,0,0,202,254,23,340,66,193,34],[38575,56417,7052,71752,45383,48840,45637,57341,57659,160829,153339,7686,75882,73417,73947,89634,17546,45278,73071,63386,112442,79373,61451,64187,52426,116954,52613,21506,97725,107156,9937,57889,58511,65303,111843,99245,42253,158772,24895,69869,65428,10456,65304,47145,12122,44172,26989,19463,60302,33807,448640,47018,67137,8146,16965,31236,42913],[0,44,81,70,188,98,133,166,75,264,171,269,198,0,81,75,0,0,161,0,0,0,0,0,196,110,132,0,58,67,0,0,0,62,726,17,0,0,0,0,0,0,0,0,71,147,576,0,165,101,0,69,101,0,0,27,0],[307,122,1106,136,98,0,0,240,0,234,337,877,171,0,0,0,330,0,0,476,372,0,264,130,118,455,616,0,395,211,885,0,0,186,423,1,0,0,0,272,0,698,87,943,1931,55,1716,706,111,484,5729,0,498,246,463,80,0],[61393,93034,9933,217481,169090,69004,266495,249683,78383,811566,487890,19882,294464,121115,104934,410102,213546,188228,511282,247629,384037,666361,201217,59951,383721,242072,87965,134996,244248,198059,30913,268160,80478,421281,294275,279295,214180,289779,40219,251074,174532,18874,317817,117649,25095,478116,111121,36659,164436,132104,505542,218112,146530,11502,165918,107702,91838],[61329,82237,8935,108656,68930,67932,58246,66086,46084,270678,273346,9103,90069,73347,91547,102661,38310,57118,93384,80759,131216,144631,103729,44234,96194,194026,99339,15533,116148,132969,18453,68909,72915,85296,143985,113811,63108,207879,22648,97825,100310,17899,71347,44024,19001,96136,45697,21541,74341,51480,353804,53378,84821,15863,35240,30690,94971],[6206,3247,3036,6534,5203,4121,3308,4862,1864,6878,8305,3118,7972,4769,3268,19662,11518,10374,5311,4910,2116,10009,3352,2110,4955,8343,5869,6899,13048,10611,0,1670,3785,10604,13432,6267,11531,10995,11413,4782,7568,3802,9388,2642,5346,2791,10986,8269,5038,2901,8660,5844,7996,3241,3097,4885,285],[653,921,44,1049,858,811,1972,1276,775,3239,2737,100,974,598,713,2266,2388,1068,1326,3363,3875,2044,180,0,3232,5190,1322,1002,5249,3721,0,2590,2253,1389,3209,1794,0,4711,7113,4016,6272,0,6471,3574,78,1711,2798,2496,5241,3187,4958,2235,3408,146,420,1105,0],[0,137,210,83,1076,1079,191,239,646,91,454,825,77,0,0,222,0,0,64,256,2,105,4844,6379,0,910,386,131,13,1075,0,0,146,242,90,1883,0,227,298,148,0,68,775,218,55,62,945,0,0,154,0,97,318,108,375,82,638],[202,110,200,0,16,121,0,0,0,86,254,383,0,202,0,249,0,0,0,58,0,0,0,0,0,343,652,0,0,0,0,590,0,370,0,0,0,0,0,0,0,1811,624,0,877,486,0,0,0,0,0,0,770,400,0,0,0],[44244,78084,15572,53740,27752,40707,17927,34883,15917,134779,44582,12880,42593,31056,34589,35673,4549,14481,34435,26172,35202,99017,38055,50853,48404,25090,5944,26004,54571,68857,4357,46272,62996,68765,93072,169233,19190,23429,51548,45514,29312,8947,37221,19406,10711,27580,11357,21358,8626,17931,121892,47623,38591,7526,13231,27668,26843],[40053,64907,7940,61041,41309,46434,40030,40235,21922,103091,130938,7710,43054,51094,51245,77386,22129,28737,63174,48095,87715,72300,51480,46111,47064,87360,40005,7731,72585,103178,7714,24318,38811,43341,78726,71987,23256,81669,20195,41744,58453,10952,28764,27721,19575,43585,23208,11350,45561,32983,197998,32323,60080,13038,22872,31145,35246],[39567,27606,10475,84566,82442,62625,84020,78859,21478,169539,205935,13086,100811,46916,51833,114382,35917,55489,125177,59091,169949,405240,122730,55796,101677,99971,47937,12514,92057,91949,12057,71440,36493,134437,168190,122782,67911,113829,50284,75985,94241,28380,180407,43237,21364,122765,32513,38892,100502,101578,158188,63844,115933,11397,41022,65379,16961],[12305,30521,8688,28767,18114,42819,9185,8450,42291,78460,45589,21384,24519,24596,28861,22843,17627,41974,5279,10611,13741,7247,7351,28584,12022,29907,17701,10636,20746,60877,16103,16973,40303,12219,10697,16368,2054,27043,81738,15894,13285,28996,29951,13005,42849,16078,17471,35238,8349,3407,105944,12236,12182,7798,21712,12647,24612],[407,125,592,245,192,501,195,400,547,2326,1272,0,502,927,333,379,285,403,586,185,440,1104,555,108,27,851,167,0,531,995,0,0,656,596,708,4,0,1378,624,287,199,3446,884,96,98,352,854,701,136,0,0,862,1145,1290,162,563,0],[409916,786419,29914,819732,589778,853422,504607,470660,983730,1527519,1206536,85740,314210,384606,367416,333719,120225,250978,497058,509457,1013331,545396,445669,382967,432989,1237404,626282,75832,978891,4033524,236161,374677,886110,444044,758746,898169,310583,980787,321107,320800,383225,147675,711737,316309,186230,499994,288507,242080,559301,442611,3950059,353511,535061,78156,173682,228085,339322],[34648,60369,9732,61115,33765,128459,18544,25838,53757,64093,39678,11634,40089,13109,16183,32955,14595,35830,11972,16861,32874,75846,50133,127507,29273,55348,30059,20605,93492,322016,16976,17956,31080,23619,41245,56501,26528,79222,83355,13638,40466,20842,17200,9951,25953,27917,15270,49958,10501,30172,171225,25942,26113,15716,34285,24197,13907],[402,161,494,156,163,134,135,0,403,1165,338,671,157,569,440,2701,0,587,228,661,559,266,204,256,404,202,0,88,453,0,0,0,0,0,0,1115,0,622,0,0,164,0,11,101,697,0,2960,933,0,445,0,223,198,284,251,1624,0],[185,453,3393,218,284,206,72,0,245,470,781,1206,188,530,127,92,30,0,0,604,647,202,1010,0,464,1529,86,74,1321,1150,0,0,0,79,1975,1271,0,0,2626,696,0,985,424,1683,7233,603,1300,2183,204,327,5138,863,1256,3340,92,246,0],[20781,20102,6316,86008,53774,48578,79641,75855,11023,117564,68869,4302,60849,44278,39436,101989,25152,24032,83332,61158,46628,176468,73092,23816,167664,156725,88723,6290,83910,74667,13368,54459,72136,96997,122170,55855,102821,83100,19563,69191,71475,6967,26359,33624,7268,123547,14465,3262,66044,77676,341359,66157,81177,9111,30804,29584,29237],[15095,29718,9918,51095,40304,26230,12543,17839,25709,67821,40510,14040,13411,13715,18077,17716,6521,4590,24297,15685,37510,27612,42669,52984,25508,42206,17615,4650,34723,45865,4722,17449,17830,19520,41469,43339,7740,29194,18835,15391,19980,6531,15693,15678,12297,18839,10825,11511,15050,14153,158752,12664,19568,4226,11643,10884,13123],[6080,5586,2027,17617,9871,7912,10481,12072,5820,44963,17489,3560,28990,13247,12088,12613,6768,6001,12674,10947,13771,24646,9477,10394,26831,25825,12185,3915,15555,7241,1718,15824,9739,26395,45556,21429,18037,16081,3001,9912,15116,6085,24695,22875,5134,22318,8628,6586,23389,10220,27339,25502,12821,1776,10214,4623,4160],[252,543,2045,1044,830,1510,692,673,1888,2924,1173,2542,922,861,824,6542,3450,3103,533,751,520,1280,129,1661,920,959,2573,2806,906,2255,104,65,0,855,2783,2295,82,527,2066,1037,2131,4556,1186,2704,7455,732,10626,5820,1008,206,6619,1810,1197,4186,2417,576,236],[11,18,1322,153,173,136,0,191,48,878,164,759,0,0,0,31,0,0,0,183,391,0,88,30,0,0,201,0,89,52,0,0,0,0,740,202,0,210,0,0,0,0,785,233,3076,1947,20,73,808,0,128,0,512,558,0,472,0],[19283,30880,2026,22926,16415,22540,10007,14531,40186,57913,43631,5354,14669,10624,12465,11981,6551,11709,13714,19142,32259,33339,36516,33321,13299,51795,16550,9811,52579,74908,9826,21725,19887,25141,38694,50196,12143,24402,16707,12647,10418,5684,36812,22025,6395,17056,12402,12360,17073,14586,150411,13130,21777,3394,7386,5767,13409],[174,822,1276,73,458,72,405,148,1411,266,697,1244,290,938,121,572,29,0,548,0,0,597,0,436,202,1340,1010,29,202,395,0,0,209,202,2278,1534,0,0,0,46,748,7351,1637,817,356,1895,1708,3880,0,540,27,325,1977,3889,931,2572,582],[0,0,0,0,0,0,29,0,0,0,0,167,110,0,0,0,0,267,0,0,0,0,0,0,0,0,107,0,170,0,0,0,0,0,0,0,0,0,0,0,0,0,0,0,0,0,1277,0,0,0,0,0,332,197,75,0,0],[6966,4936,142,7055,6345,3319,3001,3239,1120,15878,6155,107,5456,4836,2401,15335,6291,2192,5429,3895,6663,6129,2420,77,3324,2842,272,3024,3736,3562,1065,2543,1210,2055,4446,2321,3378,5516,1641,4141,2764,1209,3850,2042,563,3316,1020,1692,2687,849,4099,3813,3314,687,7729,1847,2612],[137,162,1069,1307,0,2,1072,510,366,404,201,631,679,402,0,0,0,0,872,0,0,1301,400,498,469,806,1008,238,858,604,0,0,0,402,400,285,0,0,0,1142,0,0,0,1017,957,919,907,1774,1395,1401,13682,582,2409,1000,1217,230,0],[0,600,0,0,197,0,0,202,0,566,0,198,142,202,0,9,0,0,197,0,0,0,0,0,161,0,603,0,200,0,0,0,0,153,162,0,0,0,0,0,0,3628,994,414,0,0,0,0,0,0,0,0,202,255,0,0,0],[8580,17485,5693,21894,19220,18017,16003,17075,15252,51196,54422,5639,13553,17953,24810,18730,3863,10321,28424,17325,36702,38032,19789,13540,24332,48226,12941,1642,33278,42421,1611,34533,13704,16827,33514,36043,21989,43143,15133,16048,22721,5615,26159,9487,4866,16243,9593,11634,19565,24100,86068,15610,19912,4704,5069,9880,9175],[15587,21670,5163,26081,17058,11828,11804,13770,17626,41599,44935,6407,18820,22885,19695,28267,10146,23598,19277,16768,26412,21291,15897,25819,12538,18758,9080,3531,32869,40238,1119,10293,9727,16106,30176,41194,33425,40348,17065,11780,12553,3650,19882,13318,9079,13172,15478,12916,13260,8555,96194,10610,14520,10183,7529,9637,8330],[0,7,659,0,0,0,0,0,115,0,0,1045,0,0,4,0,27,0,0,51,0,0,0,432,401,20,808,0,99,0,0,0,0,0,0,324,0,0,0,0,0,0,398,854,1073,0,972,177,0,0,0,389,183,0,234,0,0],[344,49,1096,1130,743,863,120,177,169,2951,1001,1258,628,279,291,1569,1203,370,834,2343,372,119,4729,1405,1065,44,867,230,973,93,2071,167,193,702,1459,1015,991,240,192,713,1733,676,812,444,796,519,0,554,16,526,4311,528,233,28,378,459,0],[436,452,42,188,28,522,396,287,0,523,596,77,198,136,0,501,29,137,167,74,365,13,388,1519,32,570,377,53,739,282,0,0,245,167,229,345,0,0,0,195,519,40,1357,231,1052,70,632,60,28,154,5997,537,5,3,463,0,0],[0,201,3137,396,156,171,0,0,170,571,509,607,0,0,0,0,25,0,600,0,348,0,0,0,578,0,546,0,132,597,0,0,600,0,142,0,0,794,2608,0,0,0,521,846,1892,201,682,1309,0,0,378,88,395,982,119,0,0],[22974,39926,3440,62208,42391,34124,46515,30954,43032,91738,82254,6381,27734,31334,27814,46022,10784,10076,78994,66340,75538,74742,36216,22181,84437,61169,34293,3687,51653,47737,16869,35685,41493,56568,88196,58355,48714,50850,19694,36754,30238,8022,35251,17377,11810,57649,24241,11994,27185,25071,148995,29955,23826,6389,13266,10193,4845],[3685,2647,2307,8458,6603,11240,5986,4941,2047,8232,14729,3294,13772,10354,5398,32174,13699,6664,9232,6200,11929,12201,9089,10607,5099,6161,7128,9291,9144,12531,86,5427,3320,8973,23068,26524,10357,21142,11610,8778,9518,12933,3352,2349,3164,11824,5767,10292,5738,1888,25292,5631,8869,4085,6758,4560,1944],[1087,4888,816,4610,3091,1720,2225,1403,954,7102,4897,1229,3727,1537,1355,4803,278,4239,5342,5432,8807,6688,3274,3330,6728,5417,2275,3859,3541,3803,3076,1228,744,2611,3608,2765,6695,2495,1188,4324,3295,0,8952,3688,2103,1819,4326,3474,4257,1941,4754,3637,2541,1680,1401,3023,10378],[0,9,0,31,25,91,160,21,51,646,362,0,32,212,62,1331,183,252,205,535,324,55,0,66,23,224,1384,148,1697,189,0,334,18,39,369,156,0,353,0,126,530,0,1024,259,0,62,0,0,0,184,0,9,266,71,12,79,65],[5898,7360,10439,7844,6146,10956,4863,4021,13245,15712,12181,7831,6296,4992,5533,8978,3676,5654,2349,3811,8526,2988,5694,20624,6717,5337,5108,939,7699,9481,843,5596,7557,4811,12276,19038,2459,8306,7597,2528,2119,2636,8696,9923,18505,5046,9840,8843,1448,650,29585,4552,9272,5349,5001,2175,2257],[588,11,3230,89,200,110,126,242,431,345,1087,1661,207,610,450,6,36,480,128,95,66,436,0,258,317,1131,501,181,1261,449,3223,100,195,242,664,1130,0,293,948,10,744,34,483,1440,4529,484,2911,1958,1,260,1143,844,762,3323,459,359,0],[308,200,2465,1175,199,967,760,989,789,8579,1123,2587,1424,700,184,2556,553,411,3820,183,1156,1013,3165,25,4659,7105,6212,2763,9621,60308,2486,687,16632,254,1572,206,26,7552,4587,1533,728,596,2989,7755,5774,2699,6744,5212,0,141,60,3854,1309,1203,2510,1739,0],[3004,3467,9873,9591,4234,11401,4290,3541,1556,13789,10049,6829,7837,5383,4687,24937,9440,8199,5045,4204,4496,15771,15008,8927,6938,9156,4522,909,14053,26780,8093,2570,3965,7960,16325,22083,1028,11704,11454,5846,7248,11449,18985,8259,9650,7214,11271,16714,4965,1280,16905,10365,12105,7504,5484,2907,866],[0,0,419,117,269,0,0,0,0,445,0,0,0,0,0,0,0,0,0,0,0,0,0,141,0,0,0,0,586,0,0,0,0,0,243,0,0,0,0,0,0,0,0,294,1075,0,1751,68,0,0,0,181,255,343,0,0,132],[2483,9281,7114,19227,21981,15661,11646,17689,7234,31680,33280,4058,17659,23716,19479,19084,5190,12620,24035,19585,47658,29710,34648,9122,13760,43118,17006,1713,10581,33118,6814,11416,8169,20470,36764,35000,15276,57648,13988,14806,25299,2835,9530,5485,10225,23944,7460,9276,17821,17586,152220,12315,15554,7360,5709,9143,6212],[93312,119121,16811,315932,222753,133416,183672,214826,260191,789966,446411,54779,182809,125499,128948,206771,79558,68566,384312,282957,476193,412875,256342,287488,291818,327406,110308,65379,184522,193703,85259,201900,165430,273265,216714,191697,206774,237424,151544,206750,139486,13523,419984,146019,49132,371237,100146,64714,194698,122759,630076,170243,143703,13767,137604,69226,104814],[0,0,0,0,32,33,0,1,0,67,0,164,110,0,69,38,58,0,0,0,33,0,0,0,60,0,0,0,0,0,0,0,0,219,0,0,165,0,814,0,0,434,13,34,0,0,0,0,0,0,0,0,248,70,0,0,0],[589,322,4276,210,728,486,723,1038,711,668,311,1993,62,624,848,1213,877,351,91,346,472,768,64,0,622,1137,1691,365,412,597,4378,340,0,216,120,1045,20,198,14,0,572,804,1258,2438,632,3,2818,2473,1470,0,372,604,862,4209,294,173,0],[94367,113999,29042,167166,142592,150364,78035,73606,106401,345185,257822,27410,93765,138872,127992,110310,52678,33756,143115,61507,64112,172969,94194,60937,213129,145704,42145,67268,98561,259888,25770,91674,45662,97955,151023,99602,55868,135729,54864,59073,96856,17019,142016,82540,43676,188416,67050,46325,92173,73703,282567,102581,96777,21857,64114,57947,46378],[104584,152396,51231,171656,111948,436721,79659,90410,181335,291558,176760,51031,202421,417430,365766,141263,63321,116468,120948,118489,233352,185380,352208,428161,124535,251344,191940,58516,341816,859213,50466,109693,135461,114456,104407,214081,122405,809670,636106,64023,116031,107106,105955,72863,121725,97569,46945,126614,102674,79607,1229122,158433,94840,79062,82094,60838,95014],[19074,38242,8282,35484,36471,33588,28703,27409,71994,112543,71268,14216,28181,21713,26695,26570,18377,12981,47166,38732,73476,46745,28421,30646,27582,50033,14008,11199,41706,35556,10346,56107,20137,34529,33672,35621,20978,40868,23361,24308,37287,9138,65117,20755,24592,48181,29026,13843,16113,16209,220861,20799,25104,4456,11721,12446,33578],[30457,30414,8422,45953,41567,67378,18194,26063,61024,81695,69823,10577,40670,36988,35069,59081,16051,33653,31975,33053,49348,43494,65649,113296,25627,74872,49556,21056,51369,79005,9831,37253,33802,57862,101297,133456,33828,76260,61316,26582,29610,18520,49891,31283,23782,26472,16080,27507,29373,26303,395029,32022,45248,7431,20163,25581,20157],[10771,15906,2805,12123,6930,8184,5992,5227,6277,19172,17584,2478,10021,12973,13067,10753,3891,7627,5807,4805,9548,8981,8261,16600,8212,16503,6555,4891,14101,21103,7280,7694,10176,13472,21436,23872,6938,13138,7397,4873,7738,7212,10273,6365,2109,12355,4416,3514,6156,5702,29388,4730,11533,2931,6763,8111,4445],[38197,20287,24616,90309,58004,76311,50937,57029,26297,94396,100093,16909,63649,48236,42953,88060,35056,45889,67092,90085,86160,105940,102186,84452,54961,49047,41663,32276,66736,84101,12851,23735,15181,44233,89363,100374,45446,93547,94099,64960,74582,21857,41106,33440,38626,78867,46853,73103,32359,20243,145881,44825,75119,11221,46068,49362,23396],[331,111,2062,2140,620,670,196,104,302,622,1798,3174,1689,317,230,28,31,0,0,1311,2682,10,0,800,649,2155,201,184,1457,279,0,331,0,388,30,344,0,760,515,1244,510,3458,1924,1242,3362,259,2036,1777,603,0,128,2147,1230,1764,176,0,0],[3,139,1009,8,22,0,99,24,260,8,40,136,203,1,0,50,5,1,0,20,23,0,193,35,0,12,0,0,507,34,0,2,0,0,150,22,3,10,18,0,18,9,9,0,151,96,3171,403,0,23,36,0,689,342,0,15,13],[50,0,375,51,0,0,0,0,0,0,0,496,0,0,0,314,0,0,0,0,0,135,0,0,0,0,0,0,0,0,0,0,9,0,0,268,0,0,0,50,0,0,0,31,0,0,0,0,0,0,0,0,0,44,0,0,0],[0,0,0,0,0,0,0,0,0,89,0,75,0,0,0,0,0,0,0,0,0,0,17,0,0,0,0,0,0,0,0,0,0,0,0,0,0,134,132,0,0,0,0,0,76,0,55,0,0,0,0,0,0,0,263,0,0],[351780,503069,66010,749847,444194,627668,447068,367045,478752,1321789,680644,78835,357358,380505,407369,871654,408068,401547,633939,505680,686378,997481,294819,161378,587249,723447,287247,252058,768588,882974,283520,688354,480691,762068,589889,420862,500799,766573,498959,444294,401178,84187,982971,342406,183715,530139,162651,270020,481611,327140,1930797,494125,522646,66023,354775,208675,278103],[1299,44,1786,263,328,147,228,502,92,1362,505,365,97,73,126,3351,2099,1503,127,129,711,361,104,575,0,0,201,54,535,601,44,15,160,640,829,2215,99,520,165,0,0,98,902,1172,2338,1014,1111,1140,31,24,7853,167,492,2313,185,31,59],[3232,2339,2832,7884,7015,6997,2469,2837,3035,22326,11183,3500,3064,4077,4224,5429,1731,858,9824,5067,9242,10717,4641,4484,3766,4710,2483,1651,7274,5565,0,4628,1949,6353,7284,8257,4112,7205,11213,3794,1408,2626,4601,5096,2837,2812,7385,7432,5200,1879,25524,2081,3869,1583,4088,2175,823],[396,0,441,0,0,0,0,187,202,0,231,133,57,0,0,0,0,106,0,126,202,0,0,0,0,302,0,0,1057,299,4832,0,0,0,405,836,0,0,0,0,0,0,599,425,1002,0,4237,1168,84,96,1440,0,110,886,0,0,0],[1630,2330,3265,1595,1424,1157,1885,1973,7904,3355,1817,2866,1529,1781,3075,1800,1198,2003,478,946,4278,2992,474,5298,1001,1408,1253,253,1452,2281,62,356,493,795,1967,2235,71,4474,6195,199,5043,692,6568,2667,5670,444,8629,2835,832,32,999,854,1895,4989,943,558,305],[0,0,18,0,242,95,0,39,0,17,0,0,0,0,0,0,0,0,0,0,6,160,0,0,0,135,420,0,10,123,0,431,352,0,164,0,0,0,0,65,0,2286,25,0,0,0,210,0,165,0,19,0,0,84,0,0,0],[34,488,640,538,590,211,318,644,418,1306,790,1121,202,0,395,216,110,637,0,141,1192,156,0,1660,939,696,0,311,591,458,33,253,0,197,296,503,398,547,264,49,144,1404,197,1002,2404,0,962,3747,806,89,4593,0,0,197,174,76,719],[23958,38990,5578,50840,36107,51706,39973,48803,42779,140707,128984,7636,58438,58285,63884,85206,29437,64112,55351,42102,83048,80405,43251,30714,57664,91181,37149,53839,105265,140961,23652,47197,47966,72299,110455,108512,46582,107359,42370,39266,68952,7576,63687,36441,11766,49470,29608,19317,46711,46270,245786,40137,65125,9858,19417,26180,28675],[190,1024,606,2066,784,768,32,169,185,2060,951,740,159,208,74,2192,113,100,501,268,497,1456,413,2540,35,620,152,118,1480,950,1009,0,246,1302,1729,384,513,607,37,39,605,240,0,1001,1457,1297,2253,1640,307,0,283,567,430,70,975,30,0],[158876,234271,53676,150934,123242,253428,91010,111084,410810,291431,206887,56706,143260,188641,189031,147234,48250,138016,90224,127783,245365,172081,304683,402887,87489,298861,137211,105776,246171,481714,72610,160755,211709,262285,287264,403207,161602,448317,562473,77894,165186,89965,225740,114418,148639,133575,89260,163684,84736,69897,1165145,142445,172731,52266,115503,104422,218369],[476,1079,614,246,281,322,194,176,863,2116,792,322,781,76,421,163,49,0,99,436,961,189,227,444,346,530,107,0,616,876,0,788,601,129,141,315,329,0,587,354,0,2565,491,80,615,436,460,385,365,66,3488,160,306,92,137,199,389],[9072,42869,3287,16080,13593,13475,20286,34336,37063,59580,86847,7434,14893,20934,16895,5394,420,1423,36025,27693,96423,25407,38601,82179,12171,31063,10024,848,21628,39877,6635,8487,9388,23964,51706,85071,13156,62241,36481,16176,31061,2900,3325,14772,21474,18757,6511,7288,25564,24779,315676,7001,36680,3578,3119,4727,10341],[1380,1092,5442,1825,725,2318,2910,2130,3732,2103,2837,6253,2213,814,1718,220,1211,0,1858,873,956,2808,3579,1540,2046,2456,2935,27,2382,8441,11374,1343,1923,2875,8411,3786,0,1010,5483,1317,3442,1092,7633,1808,3403,3171,9688,3117,654,1052,8569,1605,2857,5202,1596,967,0],[200,124,111,325,89,52,0,46,0,363,375,0,93,59,129,2056,38,192,26,59,0,187,78,0,0,222,0,295,0,0,0,0,0,88,118,0,0,0,0,0,0,0,0,110,0,0,752,0,0,0,0,0,32,108,161,0,266],[21764,60460,40731,35324,20504,37903,12746,24203,79292,99183,68904,47699,22145,31798,30851,28106,23070,27086,23235,33675,47757,43783,59221,100096,18902,87610,28091,14451,35729,97434,21602,26510,32325,39423,59616,52117,12286,41524,57691,15961,21682,23268,56444,28272,80563,21252,23339,28375,17137,27336,296626,24512,32087,14581,13753,29194,34169],[2076,1146,2607,2160,2546,1685,1159,2648,386,3351,8119,6176,4046,2120,1616,3182,1057,7539,1333,5697,11927,3981,5597,4726,419,2933,820,9517,9176,2728,788,4096,1709,7237,22840,15192,4919,12018,8749,4633,2480,24,2854,2677,5116,1655,6031,6883,3792,2329,4638,2787,3612,2429,183,2000,0],[9837,21095,3496,24915,14596,17254,8391,8692,36018,34840,32008,6558,12395,16984,15544,14597,4191,8010,13951,17010,23719,12427,22763,19257,8402,23481,19256,4115,23344,34460,11317,7701,14462,15543,23114,22840,13829,25159,20976,9881,14413,1876,7570,9351,11498,11740,9975,4715,9886,14563,159195,6702,14105,3648,7150,6264,29314],[0,0,0,114,0,189,135,127,479,286,168,174,7,0,0,192,0,185,0,122,166,0,72,0,392,123,0,0,167,106,0,0,0,304,0,786,0,0,0,64,0,0,0,324,126,0,894,772,0,108,0,236,262,56,69,36,0],[946,278,440,462,415,1055,811,938,0,145,782,509,1329,398,1000,1715,1212,1559,86,50,727,2053,469,0,231,1324,676,0,1070,4247,0,249,401,881,675,1226,0,552,0,406,0,0,1309,108,414,170,0,282,148,489,18041,201,923,477,334,0,0],[1272,587,319,905,380,1335,612,0,218,1491,854,669,744,787,1055,1301,499,483,385,389,1513,164,3092,1096,374,2018,0,0,1728,2424,0,350,1251,0,706,0,0,198,0,287,0,0,78,266,0,868,3987,0,972,308,2502,0,371,900,0,46,0],[1704,2294,1189,3942,2637,2041,1485,1722,645,3503,4356,1195,1764,2089,1068,6051,2734,1047,962,576,2206,1304,32,311,1268,1450,796,2498,853,816,577,1383,546,1574,3987,3030,7084,264,34,2750,1503,72,2064,921,556,4837,2943,741,2419,1223,5142,2452,692,0,1020,2519,5665],[63048,107329,35086,50265,43770,54099,14754,18109,107239,114890,66402,41040,15290,36466,34910,17834,5814,12889,22969,34124,54675,23820,44649,48951,22184,66865,21644,12655,45495,73824,10773,22429,42268,27747,59166,88240,20197,64245,60171,8318,22443,19276,51986,33393,68801,26591,18639,35263,28194,26208,362961,21517,27430,15862,12939,12548,32364],[0,0,24,0,0,0,0,4,145,0,71,227,0,0,0,11,0,39,0,0,0,27,0,0,0,0,0,0,0,0,0,0,0,0,0,0,45,0,0,0,0,16,0,121,0,0,48,376,0,0,0,0,0,0,0,0,0],[815,1381,2391,1983,1116,2391,2093,2421,696,4207,2809,1137,1450,2049,3010,890,137,915,620,1715,1976,3773,1975,1748,2643,3496,2336,1202,2358,4079,0,1706,865,1273,3118,3052,1007,5115,4291,2495,1021,268,1445,3326,1570,519,2776,1561,988,0,543,1114,2318,3225,538,245,2588],[315,665,375,734,502,686,1184,762,102,1930,1794,395,676,550,449,1456,260,1491,401,1380,604,1337,1539,1296,1395,1081,390,298,512,1712,302,697,253,1872,1747,982,592,1872,380,1587,174,13,2478,576,60,308,0,0,270,380,5122,1161,899,0,72,818,14],[4887,6685,2251,13115,10536,6113,7421,5954,13571,30781,23321,3738,8662,3321,2852,11100,978,3723,7557,6524,11364,10593,4668,4440,8589,9313,4276,1951,6733,10989,4341,3541,5313,5304,5317,4733,3321,13488,5821,8526,2604,1341,10359,7985,12002,12480,10898,7963,6791,5239,96872,6027,7261,2526,3681,3694,3806],[134,300,413,0,286,0,404,201,196,199,155,404,195,0,0,0,0,0,0,0,0,151,0,0,0,404,14,0,0,11,0,5,0,420,0,0,0,0,0,0,28,0,1202,0,0,0,0,202,0,0,0,0,0,31,0,0,0],[1162,1223,2462,894,2682,2029,1177,1725,2008,4742,3147,2010,1704,1035,1561,4042,307,1467,56,527,1605,1167,2540,2512,357,1279,757,976,3105,2630,186,174,1108,1561,3904,4546,113,2375,3645,492,2291,1005,1550,1610,1281,1134,3154,3078,432,325,5551,332,2308,959,1363,1252,561],[200,689,560,1411,812,392,1206,924,296,3839,1927,94,1686,290,120,3973,560,811,921,827,2641,2394,354,140,1685,707,458,93,453,293,0,1191,697,122,991,130,0,6775,0,2179,276,0,2186,297,0,375,0,1282,545,603,3180,471,866,202,1165,499,1152],[4690,6439,2173,11102,8331,10955,2753,4751,10896,29704,17852,8310,6532,4312,4492,9365,2620,5104,10062,6360,9799,7518,17148,18552,6875,11069,2509,2323,9051,12428,7568,6848,6524,13901,12972,16187,302,12833,12767,3612,2962,0,4640,3122,6255,5545,2251,5141,2960,2108,23539,3509,6568,1970,8455,6507,8784],[0,0,0,0,0,0,0,0,0,197,0,0,0,0,0,0,0,0,0,0,0,0,0,0,0,0,0,0,0,0,0,0,0,0,0,0,0,0,0,0,0,0,0,0,0,0,0,0,0,0,0,0,0,0,0,0,0],[0,0,250,141,0,0,0,0,0,29,0,0,0,111,0,380,178,43,0,0,0,25,0,0,0,0,0,85,141,0,0,0,0,0,0,11,17,207,606,0,0,0,8,0,450,17,0,4,0,0,0,0,0,1060,0,0,0],[2504,665,8107,917,991,1981,1688,1292,1881,2639,3142,5058,2671,1228,1345,2717,4097,754,1540,2362,1402,7112,266,3989,1510,2718,5428,648,3740,3369,9996,890,2216,3118,2876,7742,2476,537,1875,50,115,552,7801,2183,10373,2797,5916,6991,2101,114,6928,4254,4911,7691,2766,757,2867],[0,0,118,0,117,86,166,0,0,168,0,0,0,85,0,0,0,0,0,0,94,0,0,0,41,904,0,0,28,36,0,282,0,0,149,0,0,0,0,0,1320,0,307,58,0,84,0,251,0,0,0,0,0,0,0,0,0],[157,562,1561,3636,763,473,2860,2091,420,6671,3596,285,1092,1166,1635,3599,2810,1866,2899,699,4573,6035,5126,1852,931,1975,2114,4112,1199,5119,6413,2880,2715,2145,1015,2345,1195,2109,0,1120,183,0,750,1500,181,2195,2645,1183,0,144,3537,1201,1855,728,182,1442,553],[0,0,0,0,0,0,0,0,0,0,0,0,0,0,0,0,0,0,0,0,0,0,0,0,0,0,0,0,0,0,0,0,0,35,0,0,0,0,0,0,0,0,20,0,0,0,0,0,0,0,0,0,4,0,0,0,0],[9944,2752,614,2479,9392,20444,10804,34763,14896,7317,13851,1561,148952,14279,109276,219,524,292,101,518,3577,1928,35838,37918,1882,39023,48088,8302,109645,21375,5466,5526,1773,92699,198812,429312,234,2685,10481,1141,7574,378,2724,1473,1648,0,1078,1592,14189,1607,3820,22326,75440,294,0,4951,60],[21822,46285,7084,78226,47056,61523,36031,37798,49497,133510,116925,16203,49306,33227,35750,62260,14547,19305,67710,47542,66808,55002,42871,60383,52282,63341,36404,8510,87955,145464,17005,33071,47367,55581,66972,77861,26649,74924,43826,34982,24572,2526,29404,21344,44777,54339,32682,31973,30418,25901,264586,24449,33531,6157,18177,23671,50640],[0,0,0,0,0,0,0,0,0,0,0,28,4,0,100,0,0,0,0,2,0,0,196,241,0,0,0,0,0,0,0,0,0,0,0,10,0,0,56,0,0,0,0,0,14,0,0,0,0,0,0,0,0,0,0,0,1120],[2540,7807,5389,6122,4357,3351,5651,4095,13375,11030,17954,4752,6532,7352,7390,10948,1664,7613,4763,4964,18689,6981,5770,6611,3002,12511,8610,171,6382,16199,947,2548,5335,7008,9522,17099,4660,12673,15595,4263,5925,455,2868,4730,10446,2955,7402,6682,9964,6709,36528,3225,8764,5737,3011,4011,14389],[0,0,274,96,51,0,190,0,450,0,216,119,0,263,195,4177,0,1277,0,210,0,0,0,0,0,120,0,1844,0,246,152,0,0,74,0,0,0,1585,0,4,0,0,0,0,0,0,1113,104,0,0,0,0,10,666,396,163,0],[0,105,2964,179,202,402,202,404,202,917,791,1594,434,581,0,0,143,123,0,0,26,0,0,202,404,397,1808,27,1625,349,0,429,0,194,1305,2794,40,0,0,25,0,965,0,1578,3265,917,5214,3244,438,0,121,634,1064,1056,214,645,31],[8853,3557,4322,7720,6526,5098,5733,6571,6252,25820,20853,3038,16241,19146,14390,12542,8005,25298,6512,7785,23392,13704,8900,3468,7740,17001,8562,6524,35806,11774,1946,27232,5719,14438,29406,17349,13886,20804,3904,5063,14885,4676,26169,5956,2802,4189,3440,3202,6378,5411,28945,4614,17400,6737,2522,2049,3565],[36,0,260,301,282,43,42,223,739,362,685,0,294,247,93,110,318,126,311,411,553,38,0,404,217,839,102,370,583,45,0,333,176,217,90,80,664,841,0,626,0,0,631,50,0,34,0,0,90,45,504,17,26,0,757,0,0],[0,261,202,389,594,307,56,239,0,637,52,560,665,392,433,0,34,31,326,243,986,404,1118,0,166,528,156,0,442,0,0,152,0,326,297,831,0,0,0,0,0,0,1142,108,387,516,1203,308,0,0,0,296,149,0,414,273,0],[390,50,1067,214,0,108,0,0,403,229,27,628,337,0,366,178,0,0,96,351,104,0,0,0,0,0,0,0,814,0,7174,0,0,0,172,321,0,266,592,94,0,0,0,1013,229,43,1150,325,204,0,0,0,56,853,74,0,23],[3671,8232,2933,2813,3057,3338,668,1837,8494,9571,4420,2437,1027,2779,4042,1731,106,574,1812,2425,4241,2158,330,2311,685,1905,1356,645,2241,7904,1493,2219,4363,1884,4383,4167,111,5775,6869,749,1239,3200,5141,1240,3104,2695,397,1723,1727,274,14834,1367,2806,411,128,685,738],[0,46,73,41,0,0,66,0,37,529,155,0,84,0,10,78,0,882,519,18,107,32,0,0,816,585,967,0,38,360,0,488,466,383,565,218,0,0,0,303,0,0,579,0,0,0,0,0,708,34,2074,109,334,0,0,32,28],[236,27,19,0,0,120,0,5,17,28,133,0,22,20,14,46,0,0,0,88,66,80,0,816,0,45,0,0,244,0,0,50,16,87,35,0,0,294,0,0,139,0,0,0,0,0,0,36,0,14,0,427,0,0,0,110,0],[0,0,337,0,0,145,0,200,147,0,228,1158,27,0,0,0,0,0,0,334,0,0,0,0,0,197,0,0,404,76,0,0,184,404,792,1504,0,0,0,0,0,0,0,129,1514,0,820,1145,0,0,0,248,643,40,279,0,0],[3747,1636,474,12729,5294,5160,6629,7749,4625,12052,22699,251,8935,7023,8277,12748,1174,1821,18352,12109,18197,11411,21828,5826,12808,15235,11692,5460,8148,8113,2516,7623,7226,7194,10329,11240,2451,13909,916,16201,12178,353,8333,5595,2172,7446,7625,2398,7818,11212,54876,4757,18678,984,3003,4105,146],[0,146,606,55,202,0,106,404,198,712,196,851,0,24,0,59,0,0,0,0,184,0,0,975,0,22,320,0,227,31,0,0,0,0,101,1419,0,0,303,0,0,0,1044,383,3946,0,208,106,0,0,0,238,277,2910,0,0,0],[239,0,1137,316,107,41,0,0,0,655,176,288,275,100,0,163,41,143,0,48,486,536,95,112,402,276,396,61,401,85,0,80,92,0,1092,628,0,683,139,109,35,338,1027,165,255,0,75,101,99,0,424,24,238,961,516,41,134],[9475,24121,7718,34257,30529,25412,19879,23969,53085,83203,57528,10042,25584,27698,25374,61469,21509,42540,24923,24006,28376,32549,25412,40766,33705,55048,14813,35892,47900,43528,16402,29028,18634,51067,72343,76223,27481,39165,21018,18157,31125,10438,52010,14169,13748,34788,23105,17997,13347,7604,139008,23040,21074,6286,20536,13487,20998],[4722,5191,2581,6468,4911,5116,4547,7183,3686,22921,16229,2799,3573,7948,5203,1621,1594,3753,6026,8795,6991,9142,14084,5938,5341,7920,7888,0,5535,10789,7574,4951,2935,9810,10872,11524,2370,15891,552,1973,3117,7833,11682,4893,2402,4995,3922,3765,6780,1054,21718,2581,5622,1231,1553,616,2093],[1383,1485,482,2586,1167,2991,491,634,49,3646,1507,288,1492,672,547,3664,1391,2924,784,616,1587,1545,343,668,277,418,209,1497,171,771,98,1061,699,497,1643,2407,1917,1199,4087,331,3435,1185,2316,712,1408,196,665,278,605,683,1346,832,601,621,2964,470,4891],[347,201,1041,3385,1926,3811,785,508,1382,3444,1466,565,1162,885,251,2614,632,584,1439,749,1170,606,457,0,185,1314,297,0,1021,109,0,0,273,317,216,580,360,304,2665,442,948,0,755,390,389,634,2059,798,618,54,2085,436,339,1056,344,532,0],[275,809,3055,1189,1370,2620,1986,2676,2537,2902,2104,2813,4139,2782,2473,297,481,3154,200,1519,4065,1757,942,3869,171,7291,3707,131,3641,8271,0,2190,4283,4888,6811,3270,390,9187,2241,502,1792,3171,2169,259,2305,578,4656,4378,1481,2,0,2251,2345,6009,446,636,0],[22,13,511,101,0,84,48,33,21,0,49,122,48,190,21,60,43,226,0,0,0,87,4,126,0,63,197,52,57,0,385,0,7,129,71,18,57,0,39,0,1,30,37,0,17,29,392,22,409,21,351,255,15,21,18,99,0],[993290,2610802,204627,1446087,1185372,858160,1075881,783365,7744843,7334072,3435904,833246,789604,859023,721326,2727245,1811804,940966,1709589,1274074,1122537,1740436,2017204,844845,2625512,6163104,1419639,2092823,1656138,3047079,1120479,3392438,8194540,2178622,2439240,1698027,2050086,2390250,426604,1129643,2199106,391013,4299671,1953037,3724706,1423513,1036130,483914,2176637,1159446,9577236,2667465,1227435,448744,591770,383294,1310381],[3770,4843,576,11309,8872,7760,5487,4276,4515,17868,14502,2240,7725,6151,4037,5208,4646,3727,8136,7193,11187,8686,5992,15062,9439,10658,4963,2628,6460,14063,2537,1986,3713,10099,9398,13484,16229,9182,6892,6320,3906,0,7145,2814,2833,10898,4858,7087,3557,4310,34856,4144,5962,1233,5965,3846,9905],[733,904,160,3052,1566,686,1065,1192,2732,16908,7498,1656,1505,891,915,1682,0,604,600,0,804,1367,578,0,846,5555,2378,537,3883,2182,2358,7811,7534,2574,913,1945,43,753,2649,1912,3833,0,11711,4180,2442,1275,658,560,858,689,16805,4781,3934,1270,1167,84,33],[5,14,698,228,0,407,3,109,2,128,348,333,159,84,0,424,268,27,22,14,68,30,855,487,0,0,0,105,571,17,0,0,3,0,210,380,1,0,3,0,10,61,9,196,0,6,326,120,17,3,61,25,0,315,108,0,0],[2072,1432,2004,2584,1601,2831,2356,2269,2261,5668,4720,1610,980,736,964,822,274,1859,1174,1494,2459,1143,6257,5488,1211,2730,1377,259,1313,2633,1403,1352,2776,7433,7016,13655,3972,1158,9654,1430,2797,9266,1085,3664,6655,2337,2489,1174,336,517,12224,1182,1310,687,2339,1113,516],[404,995,6170,281,369,574,872,868,832,923,1484,3969,1389,1216,1141,1601,2026,751,203,266,607,406,475,1743,799,2266,1289,1087,2627,2028,1664,303,1033,735,2430,1890,218,312,4164,231,776,5633,3237,1184,6864,177,7632,6164,86,1260,9648,1585,2126,7032,1643,458,730],[1045,89,80,179,361,398,44,34,417,112,329,18,252,158,397,16,0,238,148,17,34,185,340,0,285,537,151,0,140,660,0,199,0,539,744,192,0,171,0,89,84,0,88,45,0,851,177,219,293,212,1157,323,39,171,98,97,0],[1112,2783,998,2755,1181,2453,1461,1915,200,5455,2287,630,2899,5181,2941,3830,161,1064,2564,2230,1203,2196,3562,2624,441,1898,768,2097,1086,1333,0,2133,1467,3569,5946,5114,0,1024,3439,1354,0,0,5643,3054,1052,3751,105,2813,1912,0,829,1524,1135,781,1776,630,2038],[74,0,120,0,0,0,0,0,0,8,0,524,0,80,0,78,0,0,0,0,0,174,0,885,0,0,0,0,114,0,0,0,0,116,0,198,0,0,0,0,0,0,300,256,0,800,1596,192,0,0,0,0,0,0,126,346,0],[104144,67437,17176,125020,83800,128015,70751,66982,77305,161134,101214,24588,104904,55681,56715,69447,41625,65349,44088,69828,67783,124737,156372,132253,72553,124596,73585,71922,208924,278604,61764,168508,80581,119747,139406,205976,47213,164766,135567,51828,128105,71382,281682,76244,59576,79496,56499,108983,50891,32421,265493,102308,78134,40543,54458,38428,18698],[0,326,201,0,0,261,202,0,202,736,188,475,0,0,0,26,0,0,0,0,0,0,0,913,0,187,0,0,0,0,0,0,111,0,100,0,0,0,0,0,0,0,320,0,1592,0,1306,0,0,0,0,394,290,0,0,0,0],[1113,465,2813,317,0,359,606,696,582,656,1686,2170,416,861,0,225,191,184,0,0,388,0,23,411,259,1179,201,0,736,758,616,172,0,753,681,506,0,202,68,0,0,1178,814,1560,4605,0,4549,3038,0,802,63,0,901,410,196,0,0],[294,492,195,711,228,428,541,275,407,1530,435,427,487,592,550,1439,1221,621,455,716,1027,187,0,33,210,296,331,14,109,327,119,72,121,32,45,597,26,2168,2810,898,147,0,1041,611,1188,781,80,1203,465,360,4835,178,382,285,670,292,628],[293,347,0,409,98,171,0,231,373,266,515,518,262,0,465,541,377,145,73,0,111,0,74,647,201,915,949,258,198,348,25,119,0,32,745,587,0,193,91,70,449,24,635,1844,0,0,55,0,127,34,676,0,314,354,128,132,0],[392,108,1219,128,239,0,144,92,164,103,115,661,346,960,91,176,217,78,0,129,358,0,0,0,0,628,0,0,202,80,2393,0,0,0,529,126,643,137,0,0,0,0,0,219,324,865,1380,1181,0,0,4663,194,192,837,477,193,0],[7245,4328,8907,11651,16808,16443,3053,5027,8730,10890,11854,10113,12552,10442,5108,16239,2417,11724,6258,6111,10904,10317,11318,26709,6462,8366,7915,7349,13755,44641,4911,8673,3834,7681,24399,16803,597,15616,8442,2796,9211,7072,4441,7365,8204,5723,13421,8964,4204,8491,60781,6466,7254,7434,7108,11007,3830],[31032,39955,6493,7351,44275,69328,20516,42001,11649,28210,75773,6126,85863,21512,136584,1242,448,3119,60,22391,41403,1310,128677,68368,781,76121,77061,19988,330010,46415,38031,18726,18839,59612,180808,463621,37,24834,25929,2346,31966,277,19009,9568,7719,1076,542,2473,10740,5920,62271,32673,269126,1214,112,10888,286],[0,113,190,201,0,0,0,0,323,0,0,210,383,0,0,0,0,0,0,0,0,199,0,0,0,0,0,0,0,0,0,0,0,0,0,54,0,0,0,0,0,0,0,0,1447,0,0,500,0,0,0,0,102,634,0,0,0],[601,465,2062,563,0,786,0,203,427,459,743,1230,484,374,560,165,0,0,125,367,0,100,0,20,3,1130,719,402,1089,200,0,0,104,0,1261,18,0,0,815,0,788,4367,425,989,1904,0,0,710,0,0,8162,0,1543,443,238,820,1676],[1348,172,173,1372,729,1140,1037,142,473,2294,1847,1201,505,1745,1094,351,436,316,1926,813,1334,306,622,1854,1504,678,554,0,1351,2651,0,844,918,624,2235,924,0,851,4784,509,1575,0,1326,354,1263,177,2973,2541,635,1855,6501,1480,1002,141,259,108,0],[2478,3658,2888,5607,4699,5340,5219,6235,2346,16412,12006,2372,8433,9073,8344,17150,4168,6751,6895,6651,11931,8957,3658,3591,6734,11089,5537,1239,5950,9469,795,3705,2036,4177,8179,11136,2871,15157,2561,7924,7603,5448,8163,6725,4679,7920,4589,3449,5591,4504,14698,4486,5123,2510,2597,3709,3679],[15080,1642,5272,29654,19205,36863,3994,6310,101593,15007,23764,8199,17374,52105,45935,5563,6467,13331,1862,4763,2698,5104,41277,6230,6283,8294,12485,2243,86359,70089,8987,9883,30327,6665,12003,12668,9970,46927,19908,3937,36301,13115,9447,10274,19653,6415,15431,10216,17991,27782,266155,11545,9073,12816,5007,12423,51632],[289669,136990,26176,114846,54276,515861,62792,89753,115071,229369,60699,28901,192899,51742,103408,21317,51889,137058,7221,18914,157588,20009,51354,139452,7697,510178,266665,27502,952462,683892,34564,99535,210637,22850,32601,61878,14691,533732,80013,27409,264907,31160,223464,255776,50735,13379,106088,122511,76518,37594,210927,183400,60566,50989,24031,78920,86653],[146586,355208,14681,458885,324420,298404,172867,189801,452992,1134642,1037325,46395,229710,180662,205828,263301,102782,148816,304712,256404,693926,316369,170577,132832,414621,712912,281371,81391,506817,438875,95692,414991,266077,208584,378003,322715,324609,591544,164491,256481,353464,56711,682656,314071,102153,370130,260523,141593,255658,145049,1571309,223727,248383,22539,63006,91433,149427],[30,0,0,0,0,0,96,0,0,282,106,0,58,9,107,0,0,0,0,0,0,0,0,910,0,0,0,0,0,0,0,0,0,0,20,0,0,30,0,144,0,0,72,72,732,0,0,0,0,0,1120,0,156,0,0,200,0],[0,0,114,0,0,0,0,93,0,347,51,404,199,0,88,0,405,630,0,0,0,0,0,0,0,0,0,0,0,400,0,0,0,0,212,0,0,0,0,0,0,0,0,0,0,0,794,0,0,0,0,0,740,144,107,0,0],[445,196,308,2091,986,430,742,2035,113,7929,6348,757,324,396,676,1132,0,110,3211,2749,760,2653,175,0,1677,845,204,1137,843,1687,0,1389,130,1585,2531,1198,5837,95,86,1011,4964,0,4066,1540,134,2062,320,97,542,553,8599,119,1992,201,99,83,1127],[73,311,526,851,1005,628,405,568,183,3939,76,363,520,630,0,869,0,41,1050,390,795,0,0,0,1528,0,0,0,694,243,0,0,0,0,114,593,0,0,83,225,1171,0,371,0,715,1920,1212,0,0,791,13472,792,390,57,276,0,0],[99124,186960,23456,308605,228722,246284,218908,203950,192578,589880,672895,31814,272977,207917,241606,334841,147567,219569,281005,252888,549850,413893,178386,145031,336797,482926,261782,84354,466698,535009,67215,228552,256816,282267,435814,387692,277057,775601,221011,253018,349070,91334,368220,187761,80895,287485,197286,173403,292705,206556,1522875,236995,285071,39540,78317,92717,85405],[909,101,1237,753,840,960,51,284,135,1083,830,1306,197,949,387,547,0,487,1317,0,437,804,0,0,51,931,242,0,995,1206,0,0,0,0,212,131,0,1797,0,359,1447,0,967,349,865,345,2841,377,0,202,4547,0,890,10,150,0,0],[150,198,72,206,50,237,0,42,0,329,189,0,36,79,38,363,0,672,0,26,89,259,0,0,38,0,59,69,62,87,0,0,0,106,318,643,0,0,0,56,107,0,0,44,252,0,0,0,0,0,54,223,0,0,35,0,257],[307,94,837,201,100,573,0,0,199,567,208,688,125,140,107,809,0,0,559,0,0,0,174,0,0,274,0,60,404,571,0,0,0,100,74,1207,0,0,0,0,84,0,1002,758,1753,177,0,960,0,0,0,722,396,1338,324,84,0],[0,0,0,0,0,0,0,0,0,174,0,0,56,0,0,0,0,54,0,0,0,0,0,0,0,0,0,0,0,357,0,0,0,0,140,0,0,0,0,0,0,2244,0,54,0,0,1109,796,0,0,0,0,0,0,0,0,0],[74,202,1747,36,269,240,143,0,199,1575,922,497,368,0,154,180,68,0,1017,496,188,2359,2827,0,913,296,808,0,662,258,0,12,151,485,2417,306,0,6,0,948,0,0,1862,1198,1174,1157,5111,423,86,47,0,431,924,1178,295,0,0],[0,70,133,95,0,155,0,0,0,0,558,113,0,0,0,27,249,0,0,0,171,0,0,68,0,53,1120,0,202,34,2709,0,0,0,298,36,0,0,0,0,0,0,528,274,606,0,601,0,0,0,1025,95,365,547,0,504,0],[1650,564,991,1304,632,1237,839,1452,1426,4429,3163,1522,1529,1695,1814,2965,2809,6541,362,1736,1393,1652,2188,4205,401,116,996,178,1764,1116,1250,1001,0,2258,3260,3091,0,1875,1705,1547,1199,7852,1047,1552,1476,309,132,1087,114,0,9748,913,1585,528,542,1171,102],[0,0,200,14,147,168,202,0,156,343,0,404,0,0,0,0,0,0,0,0,0,0,0,0,0,0,316,0,0,0,0,0,0,0,0,0,0,0,0,0,0,0,55,0,0,0,0,401,0,0,0,0,0,0,0,0,0],[10385,23333,1936,33667,24199,27889,15019,17324,23103,75666,65457,5010,12120,14374,12793,11953,5552,9443,26821,29541,57636,35543,28556,25369,18998,37665,17479,11828,32072,50767,5912,13479,11939,28698,42636,49314,5299,28832,12278,5990,13613,1766,15085,11527,5437,27841,9595,9789,18273,18551,141578,14051,21751,5025,5091,5803,4128],[0,29,203,74,0,0,0,0,52,0,0,297,0,0,183,0,0,0,0,154,0,172,0,0,161,100,0,0,101,194,0,0,0,0,0,100,0,0,0,0,0,0,0,139,487,0,0,0,103,0,2509,33,27,0,0,0,0],[1322,1200,5290,439,477,478,1209,786,825,3787,1142,2086,1741,1489,736,3048,220,845,561,1233,427,1051,808,833,2384,1630,805,960,1290,3724,5275,1405,872,427,1756,967,4019,2889,1385,663,4642,3139,202,444,2715,842,1561,5056,312,336,8146,956,1277,1680,544,868,549],[0,18,116,0,21,21,0,0,0,39,112,364,0,0,0,122,84,20,0,0,12,0,0,0,2,40,0,10,81,0,0,0,0,0,476,0,21,0,0,0,0,116,0,17,12,0,0,558,0,0,0,0,0,244,0,0,21],[15863,24623,6538,26710,17752,13558,23655,24686,19850,73046,57378,6072,20793,16129,12079,30934,7352,27424,29372,22226,31309,47427,27196,17916,24536,52618,17938,7151,33744,31618,368,23033,20234,33224,44077,42537,23753,44366,15945,20839,17571,410,28651,18858,12846,18133,8571,8859,20682,13051,64836,28345,31185,2666,9217,5843,13249],[0,0,0,0,0,0,0,0,0,0,54,9,0,0,0,0,0,0,0,0,0,0,0,0,0,0,0,0,0,132,0,0,0,0,0,19,0,0,0,0,0,0,0,5,0,0,0,0,0,0,0,0,0,0,0,0,0],[0,0,0,0,0,0,0,45,0,0,0,18,0,0,0,0,0,0,0,42,12,0,0,0,16,0,67,0,0,0,0,0,0,0,0,0,0,44,0,0,0,0,0,26,942,0,70,54,0,0,0,0,0,0,0,0,0],[6114,11832,3555,11620,9140,11992,6578,9982,11600,39549,22299,4217,7887,7256,5930,20739,13194,14350,4997,8024,13420,9024,5401,15942,10358,14263,4879,3925,14395,9517,8764,29313,13116,11535,17727,13399,1206,9153,6137,10077,11294,768,47297,14204,7700,10075,5662,6836,8478,7661,63880,10042,11052,1757,5964,8014,5959],[0,153,122,217,360,431,320,1027,457,912,187,83,555,310,156,961,460,24,631,318,658,452,282,360,744,1077,347,0,1165,1526,0,438,626,2872,1884,94,0,580,0,364,302,0,2845,591,0,1006,0,449,249,456,3026,966,648,960,112,0,0],[12915,32316,8514,31364,20259,33599,12197,12403,5169,100282,44605,6531,18810,9864,13996,77165,12152,35742,9688,6339,19737,42173,7250,18927,8398,9401,2949,23397,14272,21050,571,3232,4788,16331,39084,78576,16865,10371,12756,18422,20229,10556,9523,16209,5917,3080,3060,9094,2313,589,18550,10790,14531,3004,15960,36144,27434],[1329211,1833281,63328,2012511,1788410,1581900,1275265,1933267,2801530,6302836,3896391,426519,1359857,1523072,1779127,2797828,710346,1093627,1140768,718972,903858,3040428,1738185,3165208,1638127,1715855,859647,363379,2400564,2276738,30696,1018986,2229093,2228985,2741166,3257819,1225556,4165953,2225491,3908922,3065302,1845974,4306594,2139120,1124133,2964185,1326416,1600773,2147964,389591,10518490,1270716,1621337,200286,258904,626594,767330],[404,379,0,363,323,316,623,137,387,582,784,391,301,371,302,649,364,144,734,546,612,376,0,0,581,444,0,0,507,817,0,86,195,144,526,325,0,1313,0,82,805,0,215,78,0,0,1250,0,0,0,0,457,193,255,112,276,0],[0,0,1580,225,196,749,681,0,303,0,486,0,0,487,400,0,0,810,0,196,0,0,0,900,163,0,0,0,201,0,0,0,199,921,98,0,0,0,1154,0,0,0,401,207,2611,299,0,0,0,0,0,434,101,0,398,154,0],[0,0,1160,167,0,368,182,988,200,902,772,71,796,792,983,285,200,0,149,394,346,454,0,0,799,122,108,0,505,0,0,0,1345,734,84,0,0,0,183,888,0,0,475,600,1408,0,398,826,0,202,14938,1010,580,599,224,494,346],[942670,2663755,993463,1052344,959151,3627650,738953,970660,3450534,1862291,888105,359750,6413045,777323,1211814,345415,435575,1635261,365098,735502,3348231,934544,5409545,5678710,471634,3177877,2529745,551665,2350117,12861830,401994,1754802,2656938,349429,541052,1872734,439210,9698777,7548792,433158,1457092,1376458,152140,292598,2385081,224873,253522,1027692,407049,386304,4661403,1132827,637177,1923726,258769,612478,464238],[7434,5126,2537,10093,6017,13869,5194,4936,5828,9039,9556,2705,13551,3939,8237,5736,4502,8334,7706,9617,7588,8472,15671,6967,10059,8818,9118,1436,17080,60385,0,4088,10108,9314,10815,8343,1596,14307,3084,5046,6189,0,7252,2287,7332,4641,5835,8155,4628,3980,8468,8343,7191,3953,3099,1393,0],[232,41,262,449,300,546,140,127,276,632,208,161,35,209,271,7635,413,560,149,506,310,0,45,157,56,189,24,266,0,57,0,260,0,11,0,1173,0,44,943,0,0,0,132,140,85,49,154,30,0,269,3382,57,327,106,0,601,55],[29253,52815,4050,59544,45182,60180,34487,33445,62467,134622,122155,5986,32624,38223,39615,45066,15191,21178,51941,56999,90667,46736,43949,35075,31830,102214,37871,30765,82045,138219,8708,43846,43717,33329,58612,70062,23238,93015,32667,34809,53355,5450,76511,37420,8442,41739,34618,23831,41101,31873,242919,30185,43557,5463,14138,18266,15700],[0,0,110,0,0,0,0,0,0,0,0,39,0,0,0,64,62,0,0,0,71,0,0,31,0,0,0,0,0,88,0,31,0,0,0,0,0,0,2426,0,0,0,31,145,0,0,0,0,0,0,0,0,0,0,0,0,0],[41628,59014,18703,65246,43849,50481,27348,36080,78990,136002,79629,26716,47173,39822,44210,53783,24930,47969,35869,37656,47845,56476,65475,145140,34939,67079,24131,50239,47380,93358,27945,40325,55170,67926,94061,114489,48846,101838,129445,31024,44065,41035,57116,36973,44470,43728,32685,52143,28148,30566,344192,26928,41990,13890,57576,53192,44664],[1096,344,353,218,0,110,80,254,1031,189,161,186,326,0,363,471,177,0,0,145,0,0,0,1164,0,1109,642,0,794,740,0,665,0,0,509,0,0,50,0,0,0,0,162,204,0,0,0,1112,0,0,0,460,338,623,23,0,0],[131286,143784,13600,248078,187501,167046,182800,181374,200779,665593,500153,23711,197565,161688,170307,208797,96103,130906,280077,214911,279008,535283,276501,152042,263913,488905,194175,79555,397890,487313,116596,352774,306404,295755,344833,254037,134079,387528,86204,196190,279564,32020,463301,131880,40465,218467,79807,67987,173959,114712,935866,241514,224508,22441,91183,81691,159286],[777,806,3214,1809,1621,1477,677,1230,248,3078,3360,1458,1404,1053,962,7996,1404,5240,849,866,1726,850,307,1383,783,580,3597,968,2068,2901,3208,267,151,1344,4088,1694,1461,608,0,296,2776,5974,3202,1879,2226,742,1828,2360,316,110,8555,1132,811,2807,1289,181,2038],[25506,27439,6800,43516,31255,33808,30923,27486,25132,82886,54084,6162,31739,34736,25610,41207,24019,19537,44885,40820,70235,44577,29517,31560,32879,36225,26486,9846,52928,62480,8125,27689,23097,41091,40467,51193,31416,63723,34819,36399,31607,4511,39391,24168,22187,43134,24162,24593,28582,21790,131590,25590,37239,5948,17904,15462,11111],[6522,31416,5550,14479,9622,13510,13294,14262,10820,40297,20689,6480,20373,13447,15747,16200,5912,24514,15634,13877,25305,19920,17970,7142,24124,61383,22913,896,42291,28308,7438,20086,26019,17341,30319,40097,4580,51005,10797,14125,18289,12244,13994,10480,12418,11757,11767,7351,7125,9498,101556,18034,18969,8482,1807,5016,1928],[2322,3522,1963,9884,4687,5504,3398,4898,3462,19124,6318,3999,3454,3838,3295,11353,5058,2772,5294,4723,7068,13483,24272,28606,2106,2442,675,5327,2900,3074,4322,2465,2742,9483,8263,7482,4652,1354,4850,3130,1453,0,3050,1928,2976,3597,747,929,3327,3627,16942,4417,1365,181,7472,6671,5104],[97,330,52,541,937,645,201,94,72,1041,411,881,588,217,338,2682,39,894,686,469,863,145,191,117,56,738,941,81,324,1292,156,454,229,509,1002,432,0,323,1798,418,1855,0,1572,1404,792,245,606,4189,352,190,462,424,888,931,737,50,0],[12045,5358,9493,20861,12135,16172,22532,18061,1251,24512,3593,7523,2104,885,1048,7170,4806,11994,20012,24753,10511,16715,17084,11365,35360,9654,10503,663,13720,13230,5775,4704,3634,1025,2915,3232,328,8833,17129,21433,25137,7048,9575,5117,17917,14796,19441,18311,16532,7010,13436,34044,3383,7129,13914,15731,3859],[0,55,0,367,0,82,0,0,191,0,0,0,220,108,88,360,0,0,288,230,0,272,16,200,225,496,12,0,64,108,0,126,0,116,0,408,156,0,0,80,219,0,0,0,0,220,0,0,0,53,0,236,0,2,18,0,0],[0,193,81,221,689,482,270,282,0,0,703,126,392,0,0,265,220,0,62,316,0,530,0,654,0,0,0,0,512,118,0,127,52,358,174,882,1044,137,865,267,0,178,0,0,103,152,1356,164,344,0,1227,157,95,0,1133,0,0],[954,3245,608,4750,3247,4703,2427,1585,5762,5167,4964,1174,609,1246,1837,5338,152,1788,2704,2482,6325,1183,143,435,1519,1862,502,94,1541,2537,148,402,719,639,2436,706,0,2971,2849,1982,3651,36,2004,1181,835,582,1068,58,1353,1164,8728,699,1691,1564,0,1144,1053],[0,509,330,196,0,286,507,188,544,180,418,274,650,1613,974,408,232,0,621,0,1385,0,702,1655,737,4017,1754,0,75,1407,0,0,349,610,244,236,0,7498,220,0,0,2898,354,0,741,0,206,32,0,544,0,0,637,1718,250,300,0],[564,1637,3136,2066,1155,901,963,631,761,3097,2001,446,379,543,1461,1085,701,1838,1501,1259,1117,411,0,359,801,857,1036,1119,975,1687,4184,377,693,0,1005,1053,179,1628,3427,616,451,455,678,1425,838,789,3320,350,455,0,5066,904,710,1543,720,216,0],[10239,13306,12001,12237,10364,12965,11318,18657,15675,26452,41117,10289,13819,23735,36221,26671,6623,8921,13039,12558,23654,15816,15526,15169,9214,21552,14416,1288,20828,18208,1420,7638,6560,11946,21827,37100,6758,26342,3141,8728,22353,6969,14119,16890,12296,8062,23993,20027,22710,9272,68131,9194,18440,17865,2659,9179,17951],[14402,40789,4438,39322,33951,32374,20457,22983,24035,80653,87074,4617,28501,35210,30878,28985,10342,12775,38981,31869,86088,42226,31578,22952,37269,104621,37917,5090,63635,107670,2801,34337,51717,31662,49912,56328,16136,96251,17403,23503,27444,5853,50266,27849,9494,33248,11945,8278,35780,24651,100429,27623,33507,6222,4631,5430,37807],[2814,3775,457,3569,2559,1978,2337,2251,286,8116,7797,673,6446,1357,744,5339,4883,2680,4651,3968,3709,15036,7235,830,3629,2876,193,1250,7739,8793,87,4249,1620,10737,4187,4726,3984,1245,0,3703,553,331,3958,2139,509,6056,1806,1927,3426,1072,16533,2796,1740,834,5131,3365,846],[585,1686,812,2137,1086,2243,1561,1517,785,11958,6371,491,1102,1213,1152,13348,1069,2423,761,814,1390,1987,175,1071,1155,1773,115,746,2018,2275,389,48,1128,1537,1919,4376,500,761,1420,341,1037,320,1188,1562,1006,1144,287,370,633,150,4544,560,518,212,1317,830,1284],[7745,18268,21602,13653,15661,14948,3194,4957,11563,26676,15051,14254,10331,4550,12148,6253,2473,5888,4591,4814,10400,7374,15409,26402,2148,4645,704,681,5629,7909,3584,2335,10794,11246,14955,21791,3462,5472,22575,4311,12541,6937,4456,3123,20429,6193,2814,4631,1691,1958,42255,5781,9002,4400,5977,3983,20643],[0,0,640,182,0,0,101,0,0,0,269,1412,133,114,0,5,52,0,0,0,84,0,0,0,142,358,564,0,159,12,0,0,0,188,1058,376,0,0,0,0,0,0,1243,101,2073,0,90,311,0,0,0,334,64,0,0,26,0],[12459,8340,4861,21572,17563,28090,4691,6748,27943,32080,27748,9198,4539,11782,7409,14230,7235,7053,15847,11259,30187,13590,11382,38143,11553,22724,9064,8848,30193,34652,71,4091,18103,7665,12763,18812,7207,29136,23672,10909,7586,6870,5255,2010,9549,7518,3935,8778,5926,8191,84286,9053,7734,4033,10289,5690,15936],[5940,3992,22566,4910,7882,9875,4056,7617,7214,6601,13299,17702,8162,6700,4563,19721,4890,7045,2337,5569,6570,6979,9996,12656,8370,8983,9999,14964,23066,18391,5618,2303,3067,4343,22033,18092,3877,8611,20587,2277,3808,39348,25545,14699,29857,9413,44169,43038,3560,4097,42232,7782,10134,19972,6900,6067,2007],[1631,3387,512,2411,4113,2504,1769,1334,3038,11592,5974,1477,3727,1743,4734,7324,886,3752,2117,1330,4794,3803,1611,210,1729,2078,4934,0,3665,2959,8393,2492,1062,2367,3032,5334,28,8058,7940,4300,6464,0,3874,987,3336,2472,2870,450,2549,481,17057,1568,2725,697,851,1892,2504],[472,0,1275,23,23,181,202,0,0,198,0,428,202,253,123,411,225,115,48,257,59,56,0,60,0,381,0,96,56,0,0,0,0,31,436,62,0,0,0,0,428,59,643,101,1004,30,242,681,0,0,84,365,0,1579,123,74,0],[39339,94064,3736,116493,60449,39440,68714,58489,36848,269953,261908,5552,113909,84108,65883,224743,112014,69887,157593,51792,133488,155682,19140,13662,132894,162994,52082,36789,85673,73049,9159,49498,62267,91615,99046,54349,77774,169983,17242,119300,108168,3355,60379,36460,9590,113980,23550,11236,90394,64259,353894,65535,50266,5079,38146,63447,96726],[194727,278996,24685,484297,356224,339987,325274,318404,330348,876345,808939,34621,411105,343687,345173,577084,164761,253271,462166,400173,563389,593073,350620,177465,535062,602505,326472,77132,495190,598166,95662,264919,331262,387405,642401,639189,335661,891308,201113,379388,431925,45609,361962,215832,55347,368216,199525,147368,382915,254959,1427556,323497,487479,47064,126277,158216,138815],[3296,5113,2911,3074,1954,4210,1488,2892,947,2699,10728,2443,9889,4996,5673,2110,397,1527,3410,2900,3475,3178,3658,7069,1655,4759,1972,139,2530,3354,1604,355,2482,2395,3888,3553,262,6074,3539,3374,2375,594,5917,3579,1139,1959,4145,2744,1320,561,7825,3647,4891,1290,2143,1176,1100],[87,256,817,394,89,971,0,12,0,790,840,461,399,561,306,936,65,0,0,30,778,699,576,0,68,14,256,2,752,736,0,4,0,315,565,402,0,2876,0,389,0,0,55,101,420,43,1626,1504,0,0,1627,332,79,356,713,0,0],[0,988,0,478,196,318,439,74,396,2219,1973,267,1158,752,1552,258,500,4,41,0,364,514,91,0,0,69,414,11,1149,1500,0,140,99,0,348,560,27,204,0,0,718,26,371,371,164,0,295,1373,251,449,11016,350,1608,1133,1286,255,0],[0,148,2232,0,337,382,0,0,199,499,187,1562,742,388,0,0,0,796,0,0,0,240,0,990,398,679,351,0,577,806,0,0,0,0,457,1026,0,0,703,318,401,0,169,1217,4026,342,1040,1848,202,0,0,496,0,2196,30,0,0],[0,0,485,0,0,0,0,0,293,202,204,617,261,0,0,0,50,0,0,0,124,0,212,2200,0,204,60,0,735,0,0,388,184,0,43,38,0,0,0,0,0,1062,0,631,1871,0,82,285,0,0,0,0,172,230,0,96,0],[12,12,297,352,0,94,53,0,0,0,0,145,0,0,0,0,0,0,202,0,188,187,1188,362,0,0,0,1584,0,85,0,0,390,0,0,0,0,0,0,0,0,0,0,570,184,1362,0,14,0,0,1646,0,527,544,0,80,0],[394,653,701,2660,2960,1366,4076,3223,665,20947,6622,591,4500,1482,967,4549,1855,813,9787,1329,2849,11456,3362,0,7982,2067,195,740,740,1003,0,1378,0,4058,1456,350,3713,1947,0,5924,0,0,2704,552,590,22822,340,1358,385,89,329,3344,1999,955,6215,708,129],[121235,234044,36825,236878,157743,119785,116789,131254,235874,661532,511680,93505,158290,180938,190361,282553,115431,130698,169022,185885,264323,190893,147592,181718,156895,229051,86366,95166,168256,152838,38808,105957,137255,164739,223825,232620,173237,289653,232340,143285,130384,39007,252557,118325,227483,167228,102478,97364,98358,104662,1390026,104486,106086,22405,151141,111395,129177],[15875,28374,6946,25637,27378,44324,20272,30070,44999,74320,35843,11952,28711,32087,26926,50644,7575,21732,35315,27793,43333,64699,65499,83500,27982,39808,17689,13082,52010,46835,16398,22918,29900,67211,73850,87147,42675,34700,39267,20580,21007,11607,27422,14580,18274,13004,8005,15592,15784,15785,106867,28720,23360,7944,20480,18794,15232],[29294,22358,10842,55160,75826,73284,27031,40762,43590,77608,133044,15789,96918,128216,91796,64602,15809,62679,27326,53384,101399,35845,68438,60419,23047,63998,36188,21963,60325,63991,40341,63593,37620,90272,226863,181777,25843,173718,117390,36556,65651,10820,43492,21754,22033,40863,68085,119388,33397,19450,223028,26329,47193,10455,24537,30480,3640],[796,1526,5435,5865,5971,9480,1289,2524,3665,7947,16953,9635,4852,4469,3514,3083,938,2824,5757,3920,8272,1896,276,5418,4550,8587,5439,5108,5279,6357,976,702,10181,727,9710,9716,936,9207,6400,5406,7285,2004,6651,4351,8773,4191,11691,7258,3982,1091,16228,5099,22448,4884,3080,2003,176],[220687,343059,24116,485840,360468,222176,374124,364661,307550,1194655,757149,39952,306900,315817,332924,613161,258189,149317,769304,518173,588488,821339,292827,212914,617372,753383,335541,105126,592571,473103,100464,321164,301299,520628,567612,416197,338564,479730,105594,571656,430140,52143,742488,343834,131909,617042,229377,114351,450478,285910,1682487,357841,430582,32606,213899,162563,214228],[2038,1640,4267,306,1788,931,201,1359,1028,4312,6240,3682,452,3166,4343,270,54,0,635,459,6155,1158,29,2505,713,6225,3005,119,1594,3345,27,0,1455,159,2904,2110,0,2619,61,0,0,615,2704,3503,9454,795,1678,4152,840,8194,15101,2008,3207,8234,404,0,92],[23457,40600,5019,60854,54917,53377,36670,56735,23837,131281,140644,8426,52029,64649,60146,90338,37383,46648,72845,52758,70422,108355,56120,37536,68615,103357,48142,17959,100640,75328,12416,49807,31527,63937,108019,85785,34512,122423,27477,57707,89246,3277,51561,38173,14969,51782,39438,22511,54789,46208,158469,42317,70563,11027,20974,23947,39503],[0,0,133,338,941,174,182,0,0,0,0,495,0,30,0,173,0,400,0,206,157,447,200,202,202,0,101,0,15,606,0,0,0,202,0,0,0,0,0,166,0,0,331,79,666,651,0,0,170,180,3145,0,88,0,0,0,0],[1884,1194,177,1363,913,1173,1092,615,1246,2568,2195,302,427,1541,1171,473,132,419,806,1735,1196,1451,1294,33,419,3165,2077,31,1751,2345,0,933,477,1214,1850,701,131,3084,68,1135,124,57,443,527,385,1039,1956,1182,1200,800,2473,1209,1045,639,131,511,116],[324,567,589,1228,2076,493,599,362,610,3796,2597,356,654,199,259,511,445,297,636,515,878,1202,366,137,397,1344,797,26,820,618,0,0,472,498,1284,1703,443,3823,0,379,156,0,1981,324,143,762,567,2317,764,594,2292,84,1321,401,37,54,238],[15714,5208,2767,13140,10525,19958,9069,12868,6860,29013,20945,6286,9234,7595,9160,8566,4123,14363,8314,14815,34257,29018,23509,19705,9606,21999,8955,3821,22858,22237,12573,23725,10220,22843,36491,39216,15008,48566,54778,10974,11693,5024,13971,7030,7039,9677,11240,29613,7873,7929,49667,15746,15503,2463,9933,6723,2473],[77420,130599,27272,170154,151471,173998,78160,105803,189024,391824,263087,54500,111355,108954,116234,160664,65470,99017,122505,117400,292238,190969,188161,246376,137262,230330,76451,167049,203330,181642,99290,184136,115969,225525,331120,320414,89140,230867,176042,82998,130573,44134,254709,119006,88695,157851,123185,102495,124545,80185,589632,104562,103976,31904,105409,81108,99594],[1489,378,926,1792,1266,381,1063,528,867,2984,3005,404,667,1187,202,379,402,26,1013,704,431,1574,1135,1402,946,992,462,288,1429,424,0,1893,1412,1380,654,1185,0,3008,0,2004,826,1452,239,1302,36,1182,1591,169,1582,54,4577,475,1394,202,113,584,0],[0,23,0,0,0,27,0,17,0,0,0,0,308,58,0,0,0,0,0,0,0,0,0,0,0,0,0,0,0,0,0,0,0,0,0,0,0,0,0,0,0,0,0,0,0,0,107,526,0,0,7,0,221,116,0,0,0],[2768,2985,320,5270,5727,2565,3836,3833,4233,12310,13451,671,2998,6382,6544,4299,2776,1653,3403,2264,6339,7557,3628,1524,6636,10340,4829,3695,5858,6276,954,3703,2640,3454,3263,3754,4910,2880,1194,3778,5934,0,5568,3442,0,4672,3582,2191,3093,1199,17257,2177,2996,1302,789,561,6679],[22938,53078,8046,60520,32010,51116,25159,44186,31929,127668,94930,15533,41409,35669,42315,38292,13088,19562,53956,26252,91239,89675,59752,68319,47279,78058,24698,13940,48953,82399,19915,33074,33610,69987,69360,63161,20073,67311,40827,40576,44390,6516,43810,26190,19533,56071,18846,11055,34923,25866,287354,34753,38076,5834,22451,25544,65186],[0,0,20,0,0,215,0,0,0,86,0,0,16,0,0,76,0,0,0,0,0,0,0,0,0,404,0,0,0,93,0,0,0,0,0,0,0,0,0,0,126,0,0,110,0,0,0,0,0,0,0,0,0,0,0,0,0],[62,0,0,0,0,27,244,0,0,0,0,0,4,0,0,0,0,0,0,0,0,0,0,0,327,56,0,0,0,0,0,0,0,0,0,0,0,0,0,0,0,0,0,220,0,0,0,0,0,0,0,0,0,32,0,0,0],[605,0,267,0,0,0,0,168,0,199,201,533,153,0,0,120,0,0,0,0,0,0,0,0,0,0,0,0,0,0,0,0,0,600,0,200,0,0,0,0,0,0,1218,0,0,27,0,0,0,0,0,0,0,0,34,0,98],[2064,3167,2036,4859,4126,5176,374,1260,5900,5699,5713,2070,1940,1848,4841,1060,472,1622,967,3503,2516,4471,8768,15072,1209,4934,1105,3293,7775,8989,1202,3584,1227,4515,12142,13725,1324,3669,4906,872,3267,1901,2918,2092,1899,202,3115,3417,1784,1582,14809,2518,3359,1935,3446,3403,4210],[1036,1529,5005,1013,794,1099,115,507,766,1849,932,1466,920,363,586,1310,344,1007,315,149,819,303,243,394,145,368,30,565,838,608,156,195,148,358,1186,940,211,216,1161,170,878,1646,310,380,2919,209,505,360,31,0,1278,523,910,344,897,122,1122],[58,618,722,363,578,677,929,856,342,2014,1216,393,451,700,657,1630,1161,1893,397,537,445,3762,938,336,478,3313,738,2375,3573,995,0,4368,1136,3327,2042,178,247,2302,125,372,1027,201,4706,2807,0,38,0,2032,1774,1039,56,1178,1165,898,934,259,31],[40747,52643,12463,75804,61040,76110,68551,62791,80924,251849,73854,10573,67056,93399,95633,97780,72044,92841,150505,43525,108095,123742,151520,69070,95383,319299,124030,20422,234899,434804,9243,58015,116236,48278,138684,218740,84138,238108,48475,102875,130216,18991,142697,134872,34822,56275,43823,22338,58712,54489,402317,147871,161114,20215,20238,25875,72223],[12418,7351,5787,18935,18355,20971,10501,12847,4135,24933,36614,8167,30323,35800,24520,20746,6469,18798,10488,14935,33604,20112,38259,29384,15770,21473,11164,4512,20611,30111,3596,18141,12040,35586,67421,72850,19499,75835,73070,7100,10119,9959,16228,9401,14820,10120,26805,27849,12442,7405,54124,15722,25209,1786,9077,8987,6572],[0,200,1632,100,0,0,0,641,202,726,455,2396,290,91,533,0,182,133,25,0,0,404,55,939,727,1773,1849,0,1079,2399,0,200,0,579,202,3426,0,0,0,504,504,2152,3116,2503,3176,1620,5107,2256,401,383,0,600,840,3265,1179,0,0],[114832,11122,622,3747,32667,47599,47346,130973,4481,24383,28408,4864,30668,6556,36442,1090,1406,2153,0,3053,61340,3223,187960,77616,5271,152210,94796,23969,660511,94776,42493,26230,20683,156248,595557,721912,1730,8227,6847,3337,47665,1213,12936,5125,12898,1204,3161,6687,15193,5510,73625,41924,205207,721,926,20910,18272],[14949,34207,8641,36834,28048,41782,15929,18316,8463,126495,58968,8828,23380,11115,16793,89774,12114,38774,10719,6880,23588,39805,9289,22342,6551,10030,3381,17964,13820,24124,232,3939,3526,19154,44972,85051,19156,12266,13757,13076,24063,13330,12023,15597,12237,3925,3830,7316,1984,866,25249,10432,16693,2912,17567,27986,20476],[222463,449539,398654,557933,375973,401713,170537,171676,260919,912739,547152,206877,156309,144832,176598,709393,102867,272116,200388,130600,261280,462313,203765,486684,203211,163625,22604,228020,193469,213423,20896,91714,212587,382107,509490,834053,198928,172864,394465,700724,538740,373215,167566,125807,180200,137410,32287,85348,37896,49249,644618,139556,130086,53290,137469,125995,208131],[794,133,1293,1151,1289,1442,1,959,580,1456,1976,1400,644,1423,1251,1955,804,363,430,59,1991,849,995,5816,403,1023,604,143,368,1112,89,512,816,437,697,2125,802,0,939,245,2141,293,349,2166,328,446,47,3788,148,297,7138,1861,1923,839,1057,487,57],[2746,20476,2732,9961,4746,3855,6739,10224,2520,13943,11842,3278,8196,6084,3826,7740,1716,5162,8463,3063,10347,12061,7333,7153,4117,7140,3854,3003,2298,4925,0,2754,232,10190,4447,10352,2845,4315,5858,3049,400,394,8420,4693,4325,17838,2875,860,3057,1821,7022,3957,5760,344,8113,6768,13365],[1424,3648,15484,1801,592,1223,760,814,2086,2518,3648,9714,4305,2059,1225,1875,114,1709,1162,804,1747,2299,420,1375,9361,2916,5274,856,7924,2575,424,92,662,2065,2785,5326,113,827,9280,776,0,2221,5663,6705,8903,3291,13829,9225,1097,546,25547,3202,7007,14703,2325,1244,0],[0,89,0,0,0,205,36,0,0,44,0,188,0,84,202,0,0,0,0,0,0,0,0,0,0,96,0,0,0,505,0,0,121,0,158,0,0,74,0,0,884,0,531,334,147,0,0,300,0,0,4021,0,40,141,0,0,0],[584,0,1349,58,0,134,68,168,269,0,142,385,0,42,0,0,260,0,0,0,0,0,0,0,0,0,0,0,118,241,0,0,0,0,189,0,0,0,0,0,0,0,354,493,405,0,1728,0,0,0,0,328,119,0,63,0,0],[373,321,788,195,567,321,157,372,126,672,2091,29,1037,432,1122,259,112,82,385,277,553,1320,92,0,91,593,347,97,1292,624,0,613,1113,218,849,405,1936,1348,0,816,1518,12,1211,174,411,243,0,375,62,216,4044,179,941,0,405,976,0],[32,108,417,117,88,22,38,108,143,247,99,199,22,95,36,138,0,85,75,87,104,250,62,570,0,0,0,0,52,0,0,395,77,56,1048,30,0,81,13,207,28,0,121,19,0,475,0,0,126,49,306,322,55,202,334,120,13],[831,320,643,2490,1166,574,95,386,267,3205,508,333,0,68,77,509,643,0,3977,1310,334,906,0,1583,831,403,706,0,172,365,0,1601,0,0,1715,542,0,0,0,45,0,0,920,466,817,0,1820,20,1002,1684,4182,526,815,403,215,1235,0],[1209,1715,8851,461,441,3235,367,1346,1746,3086,2615,6512,6303,1994,1398,358,499,116,510,549,675,1830,411,3914,4036,1485,2692,2140,2613,3710,0,373,1195,690,3514,2412,0,6183,4210,897,2206,398,4780,4496,11788,2217,17577,6955,470,1029,5700,4404,4293,7678,890,774,121],[40347,75916,20591,69447,49957,73800,35174,50307,69591,165815,64426,16851,37995,45839,47136,43602,14133,14628,94806,68217,85471,95845,74945,74342,61867,67567,22860,6930,78407,124333,40373,60994,52535,101531,95094,70392,33291,61703,56072,35428,30602,7023,76115,44169,34334,65280,25597,27968,31208,34574,330938,52439,43823,4317,33743,21529,29460],[19985,60749,26110,33402,24271,75394,15447,23112,65462,61024,75636,40029,23146,35001,55891,19108,6056,36602,12228,62521,59626,25181,116283,158648,5528,107846,31324,6407,65755,95832,6478,17579,29192,25537,30224,70207,14011,84618,53666,7050,72184,61238,54648,26066,110388,21214,25545,43620,19475,33079,786947,21465,29712,32104,8327,38487,26399],[38006,99452,11308,83712,74622,45498,55160,60509,135808,332918,165740,28659,40794,62288,63971,20045,3737,4671,210084,90068,164750,255727,293058,294308,101190,140979,72580,3499,44830,34448,13015,73783,34105,225607,103680,83687,38032,124955,40518,61457,38029,13217,123089,45626,51583,187272,81062,28942,77956,61528,928591,64832,59746,9171,20201,15397,23172],[0,0,36,0,0,0,0,0,40,123,157,97,0,14,0,445,0,0,0,0,0,15,0,0,189,58,0,0,0,32,0,0,0,0,0,972,0,0,0,263,0,0,185,57,379,118,0,0,0,137,0,0,78,34,0,0,0],[0,0,1565,348,0,0,514,362,0,400,404,603,950,187,606,0,0,149,47,38,152,404,0,0,0,202,606,0,271,199,0,0,0,0,0,0,0,199,0,0,0,0,109,479,201,802,1764,1075,0,0,0,234,353,1038,0,0,0],[29254,34068,13624,97355,72809,71192,38575,47457,50508,138039,109188,14702,61501,75490,65425,110514,27340,28897,89868,73432,81376,95671,108442,90245,64272,81429,45879,18371,72384,94700,11569,51340,53039,77849,143596,147140,50635,111766,24770,53865,72146,5164,41241,30335,23211,87558,41383,42062,42961,59091,246474,45252,63030,15614,39160,26881,30286],[2396,417,964,2920,1534,2861,3035,2416,1823,4812,5716,454,2969,2988,3321,10821,8769,6958,5178,2285,2291,7043,1996,827,2539,4383,2956,5740,5600,4535,0,1681,1405,4733,5098,8830,10630,6096,128,2546,3230,0,1831,3127,1174,4066,2529,2012,1895,3082,5035,2211,4915,1507,2925,2660,1044],[1119,651,752,1301,1188,920,292,726,1627,803,1763,0,1393,538,1745,2301,1075,199,2156,1405,1193,2593,1676,575,1313,675,138,443,1417,1403,0,49,509,709,2730,3063,5770,2827,3709,3197,719,319,794,796,1206,2613,556,0,408,658,7076,783,1100,308,1221,1271,44],[56450,37593,18909,96327,73456,89048,69052,74259,29831,166060,139325,21212,89346,58163,40705,123450,33374,61405,93457,90519,79728,138166,80350,69577,78653,66034,61365,55930,181579,156971,22688,51271,35166,102837,169461,169812,68091,144078,84921,87949,77401,15966,72904,31723,30365,84952,63146,76130,48148,24728,183615,74873,77015,9885,80927,49541,22436],[404,0,1278,0,0,0,649,184,199,192,204,1072,160,0,0,55,0,0,98,28,52,0,0,344,0,0,404,0,1474,605,6076,0,0,201,767,189,0,0,0,0,0,0,2549,773,0,806,1719,905,0,711,0,35,575,672,178,0,0],[796,1697,338,3126,2798,2356,3435,3702,1265,6081,8836,1074,4707,2718,1418,3006,1871,2537,5378,2962,2397,9122,5674,3105,6274,6171,3364,2054,4278,5044,0,4991,1848,8883,4888,7297,681,5933,809,5129,3439,112,4698,1401,319,1895,2015,0,2134,4181,3520,3719,3902,317,2275,1402,6288],[672,60,58,61,3164,4581,0,947,0,9086,0,55,0,0,0,34,0,1138,0,0,3315,234,0,0,0,0,0,0,184,3420,0,0,0,0,0,420,0,0,0,0,0,0,0,0,0,2736,298,146,0,0,0,0,0,0,0,1067,0],[0,863,1715,266,72,289,669,182,727,190,388,1002,61,416,436,450,0,208,386,128,402,189,44,810,291,641,0,29,1057,387,0,99,88,1091,441,0,0,192,56,46,52,25,695,586,87,24,1245,510,735,237,4310,775,754,666,499,3,0],[898,1251,2636,764,250,184,0,197,674,1715,664,2012,2010,964,291,249,457,0,283,129,2793,0,444,328,1036,1066,1368,0,966,2585,0,459,401,812,1902,3040,0,1060,150,273,0,6855,55,1672,1936,1190,4527,3698,1922,2170,30374,375,1320,1923,69,675,4663],[407,0,250,243,106,11,169,166,0,212,586,345,374,560,680,303,159,0,62,441,814,319,2255,0,162,103,580,0,331,258,0,0,674,184,1257,1485,0,934,0,255,0,0,334,490,939,149,731,425,71,1346,0,197,814,0,45,584,351],[3359,7807,15888,4777,7759,7568,2069,2681,10508,9400,5307,8279,4835,1756,4223,3052,1132,2126,987,1141,3793,640,1651,2790,867,842,207,493,1357,2192,466,550,1737,1689,2254,4465,1982,2407,6200,1044,3820,7667,1478,1020,9724,1722,1543,1403,337,541,5961,1023,1404,1182,2491,1758,10605],[193,383,2050,282,1278,513,923,582,520,266,867,2207,782,0,521,1206,56,558,300,424,619,0,300,512,687,259,1001,23,1219,962,25,558,1145,0,289,3635,55,951,1486,453,0,10304,3094,1811,4632,1456,4390,1940,236,166,3324,983,1276,1806,559,184,325],[8549,10246,3813,11601,9750,16823,10967,12126,6786,22523,32104,1952,24859,20815,17470,16590,5534,18226,20832,18601,23017,39171,16420,9285,10499,22274,10463,9986,34082,38838,3990,11681,12449,16989,39202,39605,3963,39626,8410,15850,19281,6298,13058,3494,2907,9864,3556,12564,10876,5105,52438,20025,23486,6342,6054,8468,13343],[2072,1236,2700,806,28,723,0,209,1232,898,1827,2813,775,315,650,410,59,1,527,70,154,0,29,1637,86,806,3175,0,2265,1944,8,0,0,12,190,1625,0,171,1689,0,0,1142,188,1646,2756,2278,8832,2711,135,0,10,713,673,2011,1117,0,6],[22553,35823,3237,52357,33268,35525,30013,39464,35522,95371,72281,6281,33306,47913,39413,43180,19977,10827,61423,34720,53030,62849,40586,44142,47745,39314,20177,19102,45436,40521,6851,18790,22221,52460,53471,52865,51671,67298,28307,33629,37840,2203,34476,23999,19301,72419,33710,16327,20738,22690,187929,30960,34002,6565,26997,18453,16702],[199,0,1916,0,0,534,392,16,433,202,442,202,127,192,0,36,33,0,27,0,0,346,0,802,0,288,404,0,295,404,0,0,0,199,400,0,0,0,0,0,0,0,296,389,1160,522,0,1851,1148,0,7195,192,416,2107,0,0,0],[0,0,0,0,0,0,0,126,180,0,207,848,70,70,0,0,0,0,0,0,0,0,0,314,95,154,202,0,0,0,0,0,0,0,0,0,0,0,0,0,0,0,0,132,681,0,101,0,99,226,0,101,194,365,0,0,0],[0,31,61,6,0,371,206,170,635,241,243,176,158,450,254,34,0,489,139,0,362,0,0,49,36,0,240,503,117,326,1008,0,863,249,0,194,0,284,0,0,65,0,414,687,234,0,990,41,204,0,4312,63,57,309,757,224,62],[10309,24772,4615,31914,26895,38772,15102,15701,19976,47887,45980,7261,29496,27777,18976,36400,15590,15370,20676,23385,35383,27518,22781,11050,24957,48356,19522,5524,31974,87980,8027,13702,24727,18350,48506,69056,20545,76241,13101,23722,23775,10215,12956,17851,13748,18802,7673,9312,32040,22289,145628,13697,21588,5499,6002,15802,14729],[72585,86898,10852,174051,113713,124627,97661,105695,64801,225192,236678,18954,121842,111547,98247,197712,77497,84708,128744,100874,196607,173146,96044,101058,103136,202442,119611,11262,107546,159250,23158,93780,86520,95056,197683,170831,89536,271619,62679,108163,97012,15302,53128,46694,35143,109338,63675,57149,146813,81652,652542,64652,126915,13882,52984,45835,55988],[0,0,519,0,0,0,154,76,0,0,77,31,0,0,0,0,534,0,0,89,0,0,0,0,0,286,0,0,0,334,1064,0,0,0,0,0,0,0,0,0,0,0,0,303,0,0,559,0,0,0,0,0,325,0,0,0,0],[165,76,571,40,149,51,0,116,0,553,0,813,0,0,107,55,54,239,47,203,53,0,31,32,0,199,0,0,222,0,25,0,0,49,174,472,0,104,0,148,0,0,0,125,202,436,372,750,0,0,38,202,265,134,0,223,160],[1885,3786,893,3292,4085,3740,2651,3263,8105,8720,10866,656,5969,6933,4821,4076,2990,4004,5457,3617,8367,7557,3725,2861,2862,9841,4721,8,7983,10362,2309,4479,5833,4347,10628,8063,4126,9349,4862,5664,11663,5612,6121,2800,1504,4249,5435,6872,6707,7964,37100,4892,8222,1333,1077,1880,1040],[0,282,1344,0,0,615,0,124,0,0,662,235,497,582,404,0,0,0,0,202,114,0,0,402,0,757,2567,0,166,259,0,150,0,115,0,808,0,601,3804,0,0,4085,0,501,3770,0,3273,1720,0,0,0,16,669,520,300,0,0],[185,284,2160,180,434,369,0,579,525,1282,204,972,936,88,372,317,31,105,50,145,314,197,0,655,497,30,630,122,1360,626,0,149,232,1,793,230,0,0,87,79,421,44,757,739,1394,1225,5447,1751,60,271,393,651,893,1798,358,61,1614],[141,297,1147,699,195,860,60,583,962,914,1492,1109,1237,0,56,191,27,324,10,108,802,133,35,432,100,622,553,0,1109,392,418,0,59,188,390,1378,24,45,0,303,686,1347,955,1060,2708,267,4011,1045,731,0,6817,65,523,901,171,0,0],[14318,20632,5847,28074,19249,37460,14020,15785,11997,47413,42102,7269,42433,36447,34874,41792,14576,16548,15298,16857,32100,23574,19395,23497,11811,29775,15736,9787,33279,51840,0,11107,4309,11884,28236,56613,4873,34050,26712,19395,16215,54,12656,8475,7037,17172,15956,9395,16874,15878,86043,12666,20565,1315,13150,25010,11958],[0,621,480,548,128,583,491,137,188,116,241,997,956,367,184,208,0,0,208,27,189,510,850,746,862,16,504,0,0,135,0,501,0,106,1125,1651,0,155,0,509,0,439,2,150,508,655,1689,81,185,136,540,138,984,0,127,0,1687],[44,123,465,273,103,44,152,124,96,184,199,0,107,94,0,124,0,119,40,96,44,0,0,0,61,44,0,0,141,44,0,104,0,0,276,44,0,160,0,477,126,0,106,0,0,142,0,44,0,134,203,32,341,0,165,0,0],[22173,29805,14485,47334,60462,83242,33981,48082,18008,84331,85335,23609,67559,53143,46858,64920,28833,71154,15896,31327,89837,31272,65685,63764,18645,47087,25772,19978,67842,83665,24589,39393,18155,51643,149346,178534,42511,135263,147636,20828,42196,28027,33802,24714,32401,22860,63372,83849,25051,10461,200762,35025,45837,9184,22016,20272,7144],[3004,202,600,3616,1897,1116,572,298,114,2573,1136,106,3132,547,704,2742,869,232,3086,1908,1927,3585,534,193,1568,936,494,407,2610,4320,0,171,1177,1186,1255,3439,4934,1821,945,2510,927,2302,1398,617,159,1574,1557,982,1896,301,175,1439,2116,242,570,753,0],[179,10,1120,28,0,3,0,202,212,591,29,442,496,0,212,163,100,54,23,2,4,0,0,1157,0,453,9,52,635,404,0,0,0,2,840,259,0,0,230,2,0,0,1615,264,1223,295,119,746,0,0,6,451,181,975,290,0,0],[0,286,400,201,292,0,0,0,362,600,391,195,202,344,147,28,0,0,102,199,520,606,171,0,603,266,0,0,370,308,0,0,0,28,597,610,0,0,720,0,0,0,199,423,0,0,1401,334,0,0,0,252,201,0,93,0,0],[1896,3143,2520,2605,2160,2562,2011,3573,1122,8108,4395,945,5940,5827,2412,3793,3743,2231,3951,3681,5394,5401,6965,1635,3689,6359,2443,913,9403,5578,211,5015,1232,1574,6281,2693,15,5672,972,4113,4948,42,9257,4072,904,1850,814,2800,3619,2051,29205,2918,3180,3539,2670,5205,392],[201,190,519,428,87,522,0,0,0,906,299,545,234,0,0,92,0,0,237,0,296,0,0,392,42,356,1576,70,0,201,0,0,0,0,600,376,0,0,0,0,718,641,0,0,0,0,0,903,197,0,0,185,291,0,705,415,0],[24382,37370,11557,54111,33592,54716,30547,35150,52432,97477,58606,25200,32910,29458,36472,34460,15067,31028,32318,31759,55027,72283,80940,121928,29851,77604,22733,28534,69030,125577,17559,31318,30914,65831,93676,118432,9416,62523,80079,14940,25896,31506,36021,16496,39496,50878,20896,35147,17193,15199,315103,29050,36623,12776,28894,20893,21683],[748,0,603,112,281,1442,682,202,354,815,632,409,42,277,203,203,186,202,388,44,282,202,0,0,0,462,1010,0,0,816,0,84,0,482,601,74,0,2009,0,6,0,0,602,1009,320,0,1729,2969,0,0,0,275,138,456,38,40,0],[32781,41725,10092,57187,44634,49528,32603,28626,49869,124709,148135,11341,42584,50694,40686,85990,28074,45091,53910,42728,81082,54624,38807,31709,39834,103123,59873,21256,88310,99707,12815,43519,68193,39495,72881,101368,30275,110787,38846,38680,63817,17343,51347,24921,18124,33126,28249,23369,39473,29265,241091,32359,54216,18385,13050,21765,25280],[381,407,1488,942,345,807,3876,864,204,6308,3154,467,4181,1459,1494,2563,2095,1404,4180,767,1486,9144,1570,787,2338,3490,2203,875,1691,1053,110,1453,564,2878,1587,660,2536,2602,1702,4580,650,1298,2710,813,1435,5364,1529,807,969,763,2930,1628,2106,432,1036,1070,291],[0,0,18,0,0,148,0,0,0,47,960,219,0,470,21,12,0,0,0,0,158,0,0,0,0,0,104,0,0,0,0,0,62,0,216,765,0,198,0,0,0,0,0,318,0,0,218,99,0,0,3423,0,0,0,0,0,0],[10,441,4877,1219,2040,433,636,557,1652,2611,3699,5519,1364,607,728,1942,979,226,2168,773,1127,1144,1233,455,2071,2306,1455,32,2026,2470,3116,76,400,223,2830,3439,6,1226,508,323,690,5826,2158,3287,12102,1032,6060,4312,0,27,2037,620,1945,2250,1943,386,10],[53054,87103,29089,191274,136887,130842,100207,127449,86564,292300,320396,34801,136181,105270,105211,158191,53666,76463,144792,123456,249246,161920,130306,100259,107372,168830,90775,37133,148307,136391,47687,68470,78578,110544,201610,208331,152803,208589,122916,98655,90616,20794,76518,75247,97641,131914,66223,64119,92874,82538,603703,70030,121202,14666,61329,60851,84736],[183,0,1592,0,0,632,178,0,504,132,402,1452,644,202,202,0,0,0,0,0,302,539,0,0,0,0,156,0,403,937,0,0,101,400,372,1023,0,0,0,0,0,0,180,0,388,1751,239,666,0,0,6831,336,658,1476,84,37,0],[80,232,1016,10,303,38,157,66,0,842,113,877,768,575,308,441,0,0,0,516,795,0,221,0,532,133,155,0,954,1171,0,0,0,85,969,606,0,0,0,0,1902,5715,823,826,96,1212,419,880,0,0,196,451,827,2086,0,914,0],[0,0,226,0,0,0,201,0,0,0,0,21,43,0,0,0,0,0,0,0,0,0,0,0,0,0,144,0,0,0,0,0,0,0,0,0,0,0,0,0,0,0,0,0,0,0,0,0,0,0,0,92,0,0,0,0,0],[3051,4752,2332,5096,5467,12165,1972,3918,2379,7331,4664,3107,5336,1259,4506,5111,1421,433,2097,2070,1652,4170,7956,5690,2152,3883,1007,2060,6749,7134,1411,2328,1649,3688,6472,10288,3004,3248,3732,1320,2804,5485,4308,1604,2043,4121,1956,3554,50,588,23584,5041,2734,824,1635,1995,566],[31528,55374,17656,36120,27265,41565,27195,32969,28148,96486,55100,23933,27264,25135,29571,44804,14381,13820,33105,19981,45068,50911,33676,53399,17912,25615,9776,13829,30678,24592,3600,22087,18684,41294,40430,37908,15761,38401,30027,24675,16867,9631,22308,15522,34512,36531,16399,12782,21041,15917,131952,27706,25431,6777,26369,28757,10308],[96450,134698,24886,264663,189728,218902,110971,125011,147412,453671,317139,44498,149716,109680,120956,173735,63748,76976,163422,138618,188101,197436,137736,203091,118976,181832,90241,75537,195385,266149,67211,131808,111873,146718,236263,251292,106476,279648,202969,105002,113315,29460,204138,105400,86316,176595,101504,106787,86985,73908,818735,111354,143120,24817,105859,82961,100359],[567,1827,618,3041,2755,1609,923,1036,1707,4787,1480,553,739,1578,1507,1520,66,565,1764,1581,2104,3720,1782,3289,418,1357,343,159,1277,2433,0,631,728,2117,2445,6117,5944,383,2939,621,2752,1298,1211,692,1713,898,62,974,405,534,11678,667,773,359,951,896,527],[6077,7576,1321,5243,3228,3152,3179,5487,6518,29141,15152,2091,1613,1175,1918,5739,3446,1484,4775,6694,9847,17628,7962,7027,7573,25119,2457,7490,3776,5509,604,6111,16274,12717,5833,8060,6315,4136,8719,2314,2277,2142,7642,5336,4270,10738,5389,3093,2670,408,26582,5587,4492,321,3473,2569,9514],[0,795,150,159,774,191,202,358,32,1589,1247,311,405,570,917,902,0,78,51,303,293,40,1989,960,202,60,606,0,97,1014,0,46,0,403,605,819,0,1170,3807,0,0,0,929,415,1188,105,306,1651,199,0,1885,279,202,0,563,0,0],[16328,26256,4921,27525,21927,30427,17933,18980,36539,63818,53563,5867,15926,26267,26124,27124,13313,18526,23240,22781,48151,26034,17428,13050,20322,37275,15456,8897,33788,49980,1473,19863,16279,24001,28332,40927,16194,57230,21942,21412,14874,4063,23040,15972,14312,16809,15557,14791,18054,15723,116139,16664,27717,2009,10657,10805,11529],[0,921,1480,200,39,414,202,251,248,273,813,2401,406,1093,400,983,241,34,200,0,32,124,0,0,525,166,2103,2,337,2411,0,0,606,1129,819,202,0,3896,0,0,0,12482,522,1662,372,202,2918,343,1080,198,418,666,759,931,274,159,0],[7202,11384,4783,19462,13771,11811,14942,12955,10235,28355,21185,5361,17737,16724,13167,31050,7205,10209,14817,9255,18059,23869,19504,13981,12912,20752,15297,6947,19424,27195,17407,4772,5932,16548,29166,36745,11384,24409,14301,7280,9592,5436,16414,5270,11034,19599,18861,8876,11475,6417,68186,11850,12581,3037,11641,3724,7900],[0,165,540,286,202,288,366,0,265,779,32,686,118,569,457,61,0,24,193,229,344,324,2526,5442,76,569,2708,28,364,1398,0,0,35,271,1921,548,0,360,78,0,0,0,2950,244,2037,498,3852,299,472,0,0,381,2165,222,0,0,0],[70193,137172,1602,192702,146583,154122,155310,153656,120930,418250,269631,10011,105433,91874,91509,124363,28531,35390,354213,204236,306787,229542,104281,62397,241356,347458,240432,10367,278946,353136,74787,202883,227121,151774,202813,132070,101832,308389,41578,178402,122080,3760,241799,94776,32384,205406,77088,37571,182029,112369,663850,144013,161554,12516,35009,32248,16291],[2968,1206,5583,1469,2168,2133,1824,1231,1728,3691,4296,6263,4204,1207,1444,4302,472,512,283,2035,1239,4395,1170,1940,4761,2865,3189,1652,1601,4819,171,419,1387,3081,2810,5401,263,490,5430,1857,1859,6971,3334,3381,5943,2917,11510,6554,656,1434,12246,1630,1650,5214,386,2112,6210],[180,202,1018,68,0,120,0,0,606,303,164,933,532,0,0,97,0,0,0,0,251,0,0,0,144,526,1506,0,515,166,0,0,0,0,571,612,0,802,0,0,0,0,1570,413,1412,0,1570,1450,0,0,0,583,303,1450,0,212,0],[40246,40957,9902,98908,67659,69895,58374,62162,27786,144188,161943,7265,72843,60759,54906,121204,29471,35002,116351,80548,99173,124854,86046,33799,108522,100098,72080,14658,90727,79970,19397,54481,36448,91348,134225,105081,72755,125026,28853,82913,79874,5470,52132,34308,17502,96662,35420,34361,54513,43975,231362,55375,87876,16831,30609,29450,22740],[0,0,135,0,0,30,0,63,331,4,0,135,0,0,200,0,0,0,0,0,0,0,0,0,0,66,0,0,294,0,0,0,0,0,363,115,0,0,448,0,0,0,0,0,202,0,349,576,0,0,0,0,9,175,51,0,0],[4840,6850,2151,6827,9871,13587,3685,8733,15171,17807,10781,4248,5885,12787,10246,7224,922,5139,8329,11637,10027,10686,12106,23730,6884,18688,7260,6418,12109,10742,1510,12721,5406,7704,17262,18313,4508,17442,13143,5933,9716,4925,10319,5526,8061,6620,1291,9875,5901,4601,85867,7675,6483,2006,3623,2814,1749],[12582,7856,4893,10504,9966,19342,6334,9348,5120,25560,17759,5066,14213,7826,9108,11908,2058,4594,9266,14277,13933,22330,17403,10530,10907,18109,6189,7017,30209,42034,4564,7095,9386,11911,18665,28884,6193,24756,23802,5786,15113,8692,10686,8960,12240,10894,8624,16034,4131,3782,42177,11974,15548,4456,8385,4734,5752],[0,200,87,0,0,0,0,0,0,0,202,101,0,711,0,6,0,0,0,0,0,0,0,808,20,0,0,0,0,0,0,0,0,0,404,65,0,0,0,0,0,0,1040,200,0,0,0,0,0,0,0,32,196,286,0,0,0],[3864,3377,2792,2983,2085,5137,2772,3019,1136,5443,10245,2574,5373,4381,4456,2560,1684,3135,2067,2036,3184,5104,2165,10528,2166,5033,5007,5992,8313,9758,0,1703,3178,4093,5717,7633,0,8690,3180,5091,11634,4817,2114,2871,3624,1664,2799,1520,3492,1691,14924,3500,3581,3024,739,1730,1259],[51945,78570,19051,82963,67944,74772,31906,45951,92903,162555,119942,31827,54439,63847,64693,53938,14799,33884,62865,63793,102107,73080,100177,164991,56571,105350,43038,13794,113863,179628,23866,37208,57965,81887,125037,140163,34910,126974,96348,33886,50114,29230,52687,44496,53517,68388,34942,63902,41256,27264,322560,45639,56101,20431,30158,30118,35172],[54634,70413,7254,128664,79114,83739,85577,76877,71767,272738,284361,10704,97639,65837,79548,207037,60398,107070,144388,94881,205327,223833,130259,67250,103365,223284,89398,65247,139988,255128,12408,87744,147303,133628,164607,164523,73535,242469,36775,89617,117657,19650,85856,53724,13292,96986,48955,25105,77783,59284,234846,68577,79568,12106,46754,45168,68380],[125,666,109,1054,250,404,695,533,101,1296,2810,220,1789,1382,1245,1098,1224,470,287,379,1238,984,1026,1102,221,452,649,1270,278,639,77,1127,814,979,2050,1111,1395,993,205,376,976,358,3217,454,97,489,1522,579,161,58,1775,563,746,265,582,164,1737],[511,614,532,2579,673,1422,850,674,410,3346,1272,896,1731,1216,831,2432,1216,849,582,960,752,2100,393,3205,630,1224,273,399,447,3298,1888,1740,1169,4966,2625,3900,62,2611,836,387,932,52,1413,1330,3142,600,211,940,0,555,5117,719,1586,1360,1210,1944,90],[0,0,577,46,0,44,0,252,74,353,102,936,559,43,0,36,1,0,34,293,122,310,0,312,0,0,262,0,82,318,734,0,0,675,770,420,0,0,0,0,0,1632,561,308,345,0,1170,160,0,107,0,178,771,1258,234,0,0],[0,104,833,228,0,0,101,34,541,0,0,1166,212,117,265,81,26,0,59,0,33,42,0,0,337,162,0,39,603,158,78,0,0,62,849,0,0,0,0,35,0,3132,606,153,1138,37,398,2513,0,0,292,282,844,2047,188,116,0],[7655,11717,14778,12195,13948,49554,4461,5300,15644,10770,6933,4801,10453,6063,6034,4929,2007,8404,1614,5627,7998,7767,14931,39008,3696,8371,7871,163,15229,29101,1461,5104,13128,3606,8972,13882,12239,30488,55253,2331,6494,24690,10452,6397,40459,7295,8588,28545,5070,3171,27716,10341,7810,12302,2453,4162,2717],[546,635,511,238,146,665,192,624,434,1095,0,181,0,0,328,412,1668,1452,0,101,198,0,0,0,0,807,1369,0,860,343,0,0,159,0,0,588,0,0,340,0,61,5179,1558,1296,720,0,1360,0,382,72,5960,145,308,0,202,340,320],[388,144,1123,489,436,0,288,25,308,1084,22,1208,176,0,190,519,50,0,0,599,0,139,542,1639,886,1042,36,0,832,726,0,0,190,603,0,32,0,1419,0,444,4183,0,254,433,542,0,1898,161,381,0,0,754,445,941,236,565,46],[42954,46869,10473,83479,62746,68162,61050,58027,55572,132627,116330,14010,77950,59078,56952,109043,36678,37825,59823,55091,54167,79843,84598,78768,75176,79970,48919,35702,63374,94926,9380,44279,32482,67951,111322,148339,86833,91695,81812,70645,52609,5562,50424,38365,36593,77153,42494,39757,36729,41396,497065,42124,63352,9993,55055,46918,45545],[10617,12901,536,22876,13694,11337,7002,9399,14425,28350,12814,2263,7904,7852,10748,6237,3914,5105,10419,10108,14474,10281,22355,31294,9070,13672,5269,4779,12769,15259,0,5258,8613,11280,10198,13023,12325,12586,12564,6249,7668,4247,7162,9554,6915,8781,3976,5281,6306,5733,69331,3595,9468,1864,4799,5670,8874],[8566,15528,10744,26179,22991,23953,14373,11973,14014,45052,38902,6322,18116,19550,24902,24852,10160,15631,25293,19567,28825,30147,15830,15658,27974,35518,23838,10480,34984,63323,8012,8591,16754,17612,28390,38961,3081,47154,11566,18860,19591,4494,15837,12777,12013,14423,14872,10098,21503,12381,166579,17052,28477,10552,8795,13931,7171],[0,0,198,0,438,0,164,0,0,464,0,1210,256,0,0,0,0,0,0,0,0,0,0,333,0,202,201,0,0,202,0,0,0,0,124,0,0,0,2021,0,0,0,75,282,801,0,0,2111,0,0,0,0,0,481,0,0,0],[2170,1585,2158,6051,2988,2285,3002,3913,826,11457,4675,2008,1865,1884,1603,8411,3679,1261,4789,4316,3895,6160,10462,7671,4473,3952,3510,2284,1948,3682,1602,2670,4968,4868,8701,11490,2104,2042,3018,2912,6314,916,4960,1934,4983,8964,9649,4233,1068,2118,29492,5588,4051,565,5528,4507,1173],[95459,85268,21881,87221,90611,102552,41886,59007,104582,169039,251521,37785,62391,59389,58855,68595,34893,52965,55968,110539,244402,40788,70613,181778,56118,136248,54844,45048,209341,166891,33379,66786,75535,49072,173759,103219,36635,159622,76793,48188,114282,27237,64146,54929,59089,26878,24207,52656,69967,153041,746279,43074,70877,13967,27815,57950,82106],[12835,59656,12709,35360,22363,18943,19199,19293,31622,81028,41501,10295,24466,17803,19788,33681,15743,14117,29086,10128,35906,22405,14414,33221,25445,32625,2719,6400,11031,21367,5029,15367,21890,21413,15098,18777,47045,18240,33115,20373,19215,3563,34027,18799,13890,33815,2274,6424,14090,4057,98199,21614,11702,2197,26281,10143,24771],[318,48,413,12,194,138,0,216,0,274,151,735,449,152,1,0,0,0,127,0,5,0,0,259,303,576,400,0,0,0,0,0,189,0,459,110,0,0,5947,0,601,0,846,826,954,0,2196,1381,0,0,446,0,399,2424,36,0,0],[1156,290,2650,570,639,981,404,239,744,2997,644,1321,604,399,199,25,0,195,402,24,266,161,23,745,20,1770,343,56,507,654,3676,72,4,815,556,2803,0,230,6304,848,231,2629,1031,934,1713,569,436,4166,360,200,8951,1658,501,68,2984,349,0],[2361,4990,4437,2562,3938,4106,1366,1623,3846,14954,3708,4040,840,1122,1832,4355,708,1764,663,738,2669,1579,2988,2240,1004,1102,155,4011,760,3505,1054,533,184,2065,3227,4400,2917,2158,2221,240,4220,4981,1901,1067,4048,2266,1702,678,774,47,4150,857,1161,911,1995,621,1412],[1331,1498,1109,4370,2200,1405,2766,1837,1741,3193,7360,744,2758,1299,1050,5299,726,2564,3364,2280,3146,3933,4296,1639,2800,3638,837,2738,2866,2557,9,246,382,2021,6612,3229,0,3312,656,1425,7484,0,2669,2664,92,4314,88,907,2455,3366,490,1739,1838,702,1877,566,5794],[0,0,162,65,68,16,0,0,4,34,88,89,17,54,0,23,0,0,17,0,116,0,0,69,0,232,0,0,27,51,0,0,0,144,173,0,0,0,0,0,0,0,0,0,30,0,17,108,5,0,1392,38,17,222,0,0,0],[239,994,2715,960,998,806,415,602,1068,2909,1513,2395,176,284,472,89,174,45,142,213,358,1260,752,2017,260,654,24,348,890,375,0,76,0,1708,1628,1730,536,538,861,174,0,518,241,71,586,313,1224,1288,118,71,5168,548,406,627,356,64,35],[23572,32786,9253,51577,41330,46406,53770,39719,65916,134652,44545,6906,35154,58084,46963,41916,22197,19424,130001,38959,32766,172472,65867,31159,42230,101725,43154,22727,65773,147981,14761,33859,27205,81433,43287,55097,51885,85558,29710,39629,61306,6360,67183,24797,11127,66573,20207,15864,22785,10958,114990,36524,36057,13986,20772,14910,18344],[3024,1576,1485,2248,1744,2112,601,1103,543,3721,3283,1249,983,1462,1598,716,112,26,1200,980,1072,512,119,1672,641,1293,2950,880,1791,1457,0,86,2262,122,1058,4652,0,1072,638,168,0,0,288,827,3669,1029,1024,577,636,991,3199,1131,1601,810,762,2049,0],[2521,4427,950,2199,1968,1959,2883,2349,601,9906,3850,998,2576,1836,1397,2968,339,560,1571,1620,2391,6439,155,88,678,1547,893,165,1854,2157,46,1806,1077,1185,570,1366,31,965,0,1356,464,0,3676,936,1901,1834,1566,2809,632,0,4332,1200,2882,895,765,490,33],[0,423,0,199,280,709,0,174,200,494,246,159,572,553,1061,353,0,0,1027,201,0,0,807,204,397,998,0,0,202,18,0,313,101,100,197,514,0,1405,228,602,0,0,443,279,391,319,2057,98,200,0,0,0,681,0,56,0,0],[4816,12518,5991,20829,13334,16707,10464,14205,22163,29079,25418,5985,13415,18310,18247,15917,7811,10270,18128,9959,19805,13696,30274,42622,16730,15891,14983,133,20368,41433,11934,7741,13969,13538,24923,20632,5536,35183,19245,12479,9914,4297,3728,10020,17956,15266,9634,7899,14534,13230,44755,13062,15983,6005,9751,4210,5991],[0,595,1175,180,181,0,166,587,101,404,199,197,314,0,0,272,29,32,0,0,2,0,404,705,859,221,141,0,247,398,0,0,0,0,808,490,26,0,0,0,0,0,0,885,2543,0,1607,2162,788,0,61,361,197,197,0,45,0],[0,0,599,408,0,101,170,2,53,206,102,178,91,0,16,220,0,70,515,0,199,108,1193,0,221,168,0,0,0,84,0,0,135,0,0,521,0,0,0,229,0,1553,600,0,406,801,1623,187,0,0,1464,853,180,12,178,216,0],[100,0,252,119,0,0,0,0,0,0,0,0,0,0,373,0,0,5,0,0,80,0,0,0,41,0,44,0,50,0,0,0,0,0,0,0,0,0,0,0,0,0,0,0,225,0,0,0,0,0,5,0,0,0,0,0,0],[48754,52102,9873,101716,65848,65641,56361,66479,25015,169595,137862,10892,104038,86531,83076,131688,62092,67848,105878,88274,92478,156886,80153,42991,59880,154508,79578,32006,171563,200080,16401,64286,53410,73316,133597,102818,35922,146500,33518,77114,141694,6691,49673,54274,18176,62535,37510,23925,79442,60817,215542,59813,129654,15531,32289,52788,70243],[73,819,294,993,965,1246,298,794,96,834,956,204,457,1032,60,1130,672,690,424,716,894,840,708,659,1516,1320,1885,819,469,1167,0,80,669,956,1756,1555,730,2380,122,221,2527,0,998,0,0,297,2458,0,183,431,5255,526,675,148,733,929,48],[611,32,309,28,29,0,47,28,372,517,334,202,96,0,30,16,0,0,207,0,0,278,24,42,88,0,0,83,525,0,0,0,0,400,366,0,0,0,0,0,0,1448,0,689,752,0,70,99,27,806,0,0,10,20,741,0,0],[606,198,405,825,1729,1282,1201,471,0,2218,1696,367,342,398,1158,2757,0,1132,1033,491,1550,1023,0,473,872,2040,556,606,691,1119,0,499,18,1860,332,1495,60,998,42,261,2208,160,627,1780,335,162,1400,200,621,758,450,1008,481,331,1974,1420,0],[49,394,46,181,59,188,26,158,161,205,342,24,23,116,172,2913,1647,674,218,52,450,218,112,28,228,351,166,523,306,77,52,0,0,29,347,270,144,0,204,299,0,107,58,25,1767,134,1509,906,84,597,422,129,74,314,57,0,49],[1711,400,953,355,127,864,266,0,1004,1327,1089,1645,574,739,553,79,29,664,0,0,199,324,1470,354,0,425,0,1807,1941,2242,0,0,0,342,1880,4309,0,2557,1568,296,744,764,805,451,1436,202,4778,2212,600,0,0,829,1118,1081,78,256,0],[374,47,1260,0,381,312,118,0,492,90,749,985,180,89,561,936,0,0,137,402,254,384,496,1107,147,600,0,0,141,751,0,0,597,0,162,1520,0,0,0,0,0,3286,55,106,1379,632,265,917,0,0,25,0,785,517,415,0,0],[1278,1100,1523,883,319,4458,1013,158,822,1805,1470,1999,940,662,1773,3009,1151,1467,260,175,830,447,594,5556,394,1853,677,4190,2337,5641,0,99,1101,641,4994,2950,136,1336,1281,1018,4095,2402,3488,2478,2117,1657,3425,2239,245,168,1512,718,2964,2682,1132,656,615],[1290,2169,1825,794,1053,2558,1630,1112,520,1495,4057,1990,2589,2649,2513,780,56,2016,14,2266,3200,530,1083,2491,310,1705,2158,88,3339,2156,147,1213,80,659,7974,5735,201,5173,8642,348,4499,54,996,1598,3737,591,2026,5952,524,109,7776,367,2188,1484,482,1605,91],[0,0,0,0,0,24,0,0,154,158,0,161,0,45,0,0,156,0,0,0,0,155,0,0,0,9,0,0,0,0,0,0,0,0,0,0,0,0,0,0,0,0,456,0,235,0,0,0,0,0,0,0,0,168,0,0,0],[732,275,0,287,242,1113,403,299,1748,602,404,171,447,188,590,565,58,1662,215,510,714,545,808,76,886,792,265,0,9,418,0,179,0,172,538,1194,0,0,510,0,0,0,465,130,0,0,0,904,203,0,0,161,1549,614,88,168,0],[15521,50247,2164,46946,34771,29567,28741,30098,37677,106622,51875,3366,23003,24821,20954,25714,11693,10007,33173,25223,49730,25857,8059,8349,33450,40840,10346,1761,25854,49180,5639,15133,18849,17083,16978,22421,18643,54171,20972,33895,22530,1096,49673,29016,20139,39124,19776,17173,32503,10019,55894,17977,25999,2723,10001,11233,9405],[0,0,518,0,1109,57,0,0,0,431,312,324,232,288,257,23,0,0,0,86,771,793,0,51,0,284,471,0,187,0,7343,0,0,0,100,173,0,0,0,786,0,0,243,63,670,0,1173,0,0,0,0,56,202,246,0,0,0],[0,0,175,0,97,0,0,474,0,29,190,487,0,0,0,0,10,0,133,0,0,0,0,0,0,0,0,0,318,101,0,0,0,118,375,168,0,0,0,238,0,0,553,0,0,0,1632,1934,0,0,0,0,199,822,25,0,0],[33716,47989,3727,68477,52086,54311,42433,49755,115545,152598,161611,10865,46142,67705,76400,62644,15436,54716,58806,49781,105544,79688,66875,44934,44203,113501,37003,36362,92271,212467,25665,39435,75068,55421,96878,105585,42697,164790,30623,41338,63141,1684,49274,33360,14808,46654,24159,19199,52254,26880,178620,32942,64952,10266,22695,42515,45021],[1040,2297,2482,7212,2257,1449,8067,9674,448,15220,12017,4202,6295,2803,2639,5690,6553,725,8221,718,8472,3637,5091,1228,3210,7970,2397,345,4503,23556,0,2985,1359,5284,6933,14071,4307,6040,3932,6281,1214,222,5142,2999,3471,1756,825,2157,4959,2853,6738,11027,10368,2969,2580,511,0],[22555,41693,9977,67149,42388,27832,51935,50941,30177,139502,203471,7950,50789,52802,51992,111718,40590,38371,87737,46275,118776,100405,43201,9212,81589,116782,51705,18251,71123,85005,4691,46605,47488,54447,64525,44311,69672,141026,31291,85363,72048,2543,72428,36203,15377,62505,27992,10627,72259,27469,193636,38152,61976,7296,14394,23107,45638],[2670,2502,2107,3506,3344,6005,2081,2483,1156,5080,3827,2846,2825,3049,1500,1595,1217,948,2414,2336,3695,3958,3216,7529,1583,5287,2135,797,2437,3181,1797,1410,1277,4301,4855,4634,86,3362,6477,1400,4889,3298,2118,1007,1279,2270,1517,4668,881,703,16151,2192,834,210,1216,1907,1199],[63014,82485,16261,271189,193356,207445,104366,118634,29305,303082,331299,24029,91233,76668,67717,176567,89968,90801,173026,102628,281594,233128,181313,178717,134477,277929,112274,72370,246606,433565,35422,88174,110185,188793,199343,230393,115828,296648,116703,133781,179764,23477,130368,108947,42308,151086,44757,40479,98008,88000,180984,119202,140529,17016,89750,70283,44521],[653,1009,990,954,232,414,1398,692,327,904,747,1716,668,435,913,160,47,104,437,604,1259,1385,1749,2732,702,190,383,227,431,786,0,0,323,1836,1963,2944,152,0,29,308,1339,0,970,428,2365,280,3588,439,503,415,0,855,972,474,1829,405,0],[0,0,207,236,0,246,0,17,44,99,447,17,368,3,0,3,26,0,3,0,34,0,0,0,6,363,905,3,0,382,0,0,0,0,545,564,0,1075,0,0,0,0,202,279,1103,0,815,1102,144,0,0,114,299,361,6,4,0],[21000,15574,18571,48911,60918,69682,21954,31316,12405,64368,129848,29542,104866,101899,72727,54802,13380,41600,13356,37629,101879,38350,52043,63947,10464,38316,20230,43843,69528,59482,15167,71796,25468,70754,231558,167789,46348,131756,170166,18029,48450,16507,25646,14559,29944,28540,68285,97190,25699,10988,176405,26178,42754,8115,17799,32101,10371],[155,292,1692,1157,1742,1483,129,781,505,567,1966,1097,2504,3541,2070,852,1377,304,26,767,798,319,3047,0,241,798,404,2080,998,1881,0,994,309,1260,865,2334,0,5517,3511,0,601,58,1346,599,2865,1036,1159,3546,1028,0,86,544,1388,2442,1423,90,26],[34,327,1435,158,790,619,110,415,13559,38,768,430,249,3132,1433,117,24,0,58,0,0,0,3472,1462,0,1236,501,71,433,7602,99,0,0,0,498,606,49,0,0,0,0,0,398,407,3766,172,1221,184,184,2991,18882,981,4085,752,213,1156,4708],[300,21,194,0,0,326,314,0,356,0,16,404,388,292,2,493,0,0,158,0,0,0,0,0,0,0,0,0,318,0,9,0,0,373,400,540,0,324,792,0,0,4281,0,54,14,350,0,300,110,142,0,0,630,622,686,531,0],[18492,39943,7838,11603,8921,20562,7475,11440,45440,41611,20903,5107,7949,16350,18881,10932,7948,9470,7957,4468,17157,13826,35971,20150,8742,24165,10761,5948,15699,28473,3315,6513,7123,19632,34878,44137,2581,33350,12007,5782,12704,1738,26541,16417,8689,16562,2697,6383,7664,8129,136470,12627,7611,2585,10028,6253,15108],[76880,108626,22504,147815,100801,111410,69751,92694,42475,200493,179693,18173,118238,140561,116084,187478,75202,98533,133309,71750,139422,178248,119644,107536,151548,162095,99205,75821,126712,188817,24428,81696,106879,120831,211551,152100,91425,200997,65638,87777,96190,18514,39576,49818,36992,152457,54669,40373,73224,60793,318796,73678,114000,21414,95666,95974,256477],[2612,3875,856,3119,2283,3951,2863,2352,908,9963,2695,1025,2472,2390,1553,5087,618,1100,3287,2201,3517,6795,3026,4944,1469,1481,690,1341,2139,2024,0,1619,947,3199,6629,5841,153,963,1464,2328,280,194,1513,425,272,1351,986,1109,679,909,4261,2055,2424,190,3674,1023,1429],[139,154,0,407,0,1156,235,191,157,116,1021,0,359,259,456,498,64,0,3,291,248,816,265,351,0,23,0,28,557,529,0,0,191,0,490,33,0,0,1525,108,0,0,14,127,0,0,0,84,0,864,144,546,0,0,0,28,0],[7381,6138,1098,7955,4596,12519,5419,6502,4323,12364,20377,3186,9498,10687,9557,2359,4050,12167,2515,6011,10153,6476,16004,12865,2093,10190,7620,3991,13690,16169,9648,9255,7456,12699,20574,30002,1204,17765,12012,3923,4555,5557,8633,6584,1833,2165,9516,9792,3338,9832,30609,4221,5430,2989,2890,5852,5615],[0,0,30,355,124,258,51,0,0,455,96,0,99,0,0,514,35,1243,31,0,302,213,192,362,0,88,55,1484,255,53,0,0,0,48,207,351,0,25,24,0,4,0,452,100,0,0,252,0,0,0,52,0,0,0,29,70,336],[0,0,100,0,0,0,144,0,371,194,358,685,119,15,0,0,0,0,0,0,0,0,0,0,0,522,404,0,181,0,0,244,0,0,0,116,0,0,1415,0,0,30,22,94,871,0,40,815,0,0,2050,0,270,578,10,0,0],[0,58,296,0,0,0,0,58,0,74,0,58,146,83,132,0,0,0,111,6,0,288,0,0,0,0,0,0,0,0,0,0,0,0,0,0,0,522,0,14,0,0,0,0,0,0,0,32,103,0,35,0,0,0,0,40,0],[136,287,2544,506,0,315,104,0,87,799,971,1175,564,401,0,593,478,28,130,231,229,820,24,726,400,444,2114,137,995,148,33,0,0,360,670,848,0,495,0,0,478,4945,195,1262,3990,885,4046,3510,222,0,4849,604,1071,1900,158,0,2441],[273,373,718,536,1009,855,595,520,1598,1092,801,629,772,959,245,819,99,44,334,645,302,399,1989,1780,281,209,649,27,558,660,31,86,293,888,1648,727,4,420,168,158,178,566,177,237,874,143,1061,58,90,500,3319,537,549,623,924,233,0],[254,489,0,620,453,795,589,221,283,1227,474,151,134,431,65,1437,44,448,189,212,673,43,607,1144,0,0,0,128,567,938,0,69,115,142,581,1229,0,53,0,249,1402,0,209,0,0,76,0,24,318,0,596,216,196,0,327,136,183],[4196,6642,2074,9433,5164,6626,5516,5685,2616,17019,13015,2778,5657,4000,4428,8767,2044,1487,9054,4941,8772,12709,5754,6495,7015,9306,4561,5396,7640,10517,253,4832,5557,10409,14890,14992,6077,16743,1577,6414,4283,2301,7605,6474,4978,9182,2697,4474,4302,5180,30504,5806,7342,2134,4311,3799,8243],[13927,33911,16538,41603,28382,21303,18997,24426,59878,98404,54353,22785,25430,23027,19700,39136,13196,20362,34438,23416,33288,50949,36361,95473,28252,37927,19319,21879,27838,61907,18162,20524,17595,46544,43091,48997,14158,30927,58012,19527,13472,12857,30807,15535,39520,43764,10485,23746,19799,11429,177521,21976,21822,8216,21789,15115,18527],[45258,46688,42486,80586,54490,99861,35586,38536,97399,130132,68244,50291,44413,36876,44640,55686,28666,52836,43964,40123,61778,54958,54811,152407,38444,57563,37560,55339,42222,79493,21006,40256,56317,80213,80995,125697,27770,69646,379253,30291,29093,75084,55113,39863,98890,49614,36051,81431,30165,17029,416889,37836,28945,19547,89062,44443,44669],[6072,985,9330,12244,7968,6539,1234,550,6049,8637,17309,5022,5693,2064,2020,5364,501,1587,9792,2414,6639,8015,15164,12872,15157,2811,5325,240,4680,4158,49,2296,555,3806,9145,2924,3775,2347,10077,9268,8454,4097,17427,4240,17226,4683,14622,12580,2132,1465,10080,5870,2948,7158,5592,1921,1573],[6134,9310,872,13342,9720,15946,8149,6780,9540,22271,25128,2590,12882,12209,12279,14135,3313,15887,9030,7977,17535,14141,11145,16770,7820,16848,8347,5537,18818,31174,0,3714,11227,7306,15674,29097,8304,36919,8073,8510,13541,2299,5094,3200,1843,10148,1523,5413,8214,9397,43921,7900,13726,3012,2152,4587,2514],[282,66,60,0,704,29,101,80,0,0,0,0,0,0,0,302,0,110,0,0,27,184,0,0,0,0,0,0,0,0,0,0,0,0,573,0,0,0,0,0,0,2480,365,0,0,0,268,0,0,0,99,0,44,58,0,26,0],[68,100,155,261,275,62,66,176,1,940,418,0,244,55,117,342,133,38,0,0,182,13,2,57,12,4,78,12,110,168,86,92,6,35,27,11,3,185,63,1,76,7,263,228,159,752,201,1,35,171,783,537,292,439,72,49,0],[1165,1243,2349,2099,1922,2278,1542,1173,704,5343,2729,3516,3039,1355,920,3160,51,1556,2117,1437,2553,1205,467,3696,946,2276,886,903,2746,1944,667,897,430,3728,3104,3748,1003,1311,1912,1505,1122,1597,2171,2401,1980,2319,4045,2216,496,702,2978,1405,2365,1755,616,240,0],[38,147,249,771,231,187,138,281,93,639,1031,82,642,47,228,1136,186,248,603,829,534,1052,452,130,908,1362,30,403,974,419,0,628,241,271,456,1103,75,0,92,204,263,51,183,125,605,554,751,0,401,0,56,323,99,0,669,333,633],[198,0,1365,267,282,1154,54,313,0,790,101,1781,0,34,43,58,37,40,0,504,1166,403,0,202,0,273,202,137,1183,202,0,0,0,342,387,210,0,364,0,0,0,0,377,809,30,197,2797,739,18,93,2065,0,741,127,389,71,0],[648,273,80,116,698,3705,457,1047,192,201,592,487,1008,352,1088,15,32,0,0,45,569,88,5941,4105,0,2694,2945,1396,7546,907,0,85,130,1934,3587,16333,26,85,1654,61,861,105,355,183,174,121,2185,113,98,114,1988,1212,2415,7,181,383,462],[0,781,159,631,0,0,140,201,504,595,1418,462,209,339,268,624,0,86,0,0,404,127,0,1354,272,0,0,0,302,1682,0,0,0,0,808,0,0,296,0,478,1607,0,0,0,1463,1659,552,0,105,0,0,32,291,976,246,554,2586],[0,0,0,0,0,0,0,0,0,0,0,0,5,0,12,0,0,0,0,0,0,0,0,13,0,0,0,0,0,0,0,0,0,0,20,20,0,0,0,0,0,0,0,0,0,3,0,0,0,0,0,0,112,0,0,0,0],[1637,2247,6875,4798,5256,5890,2145,1979,3016,5992,9635,7730,3794,7477,8194,4704,968,1682,5222,5063,4583,5859,2548,11608,2260,7915,6591,2234,10925,11382,2479,2733,1829,3794,9578,9845,0,5420,6562,2613,5420,4748,3568,7784,6394,1756,6923,11307,2931,3861,29121,2961,5745,7056,2048,1600,528],[0,0,59,0,0,0,0,0,0,0,0,0,0,0,0,0,0,0,0,0,0,0,0,0,0,0,0,0,0,0,0,0,0,0,0,0,0,0,0,0,0,0,0,0,0,0,0,0,0,0,0,0,0,0,0,0,0],[0,138,723,0,0,0,68,0,26,106,225,235,0,274,0,0,0,49,0,0,0,0,0,0,503,752,2945,0,274,102,0,0,0,0,1009,199,0,0,0,0,0,0,594,82,1185,185,406,0,0,0,18,0,543,388,0,0,0],[112,164,116,94,66,100,91,86,0,0,417,302,196,9,90,16,176,0,396,0,93,399,718,880,0,171,0,0,0,0,0,0,0,28,100,30,0,0,0,98,0,0,350,366,0,0,339,0,0,0,0,164,45,188,0,0,0],[70,0,716,920,707,11,196,394,40,242,4243,705,113,284,201,5,0,219,301,35,715,255,0,590,174,288,339,0,0,301,0,366,30,877,436,467,0,0,0,35,290,0,1472,248,337,1243,4157,314,0,107,0,105,469,405,274,0,0],[7028,6372,4893,15251,10599,18615,36334,41153,9946,30233,19958,4567,21769,20174,24184,17628,3295,8134,33735,12783,26952,38104,27145,13974,54283,37331,24860,9144,24833,35694,12960,22247,9485,28456,56146,29565,14671,40489,26455,13335,28359,2539,17862,7848,4530,40642,13931,17772,3948,3989,43651,16047,12898,2874,8084,3324,1560],[143,0,498,0,0,233,0,0,398,0,156,312,0,0,0,0,0,0,0,0,152,0,0,0,0,0,1809,0,0,0,0,0,0,0,0,0,0,0,0,0,0,12,0,0,197,0,0,0,0,0,0,0,0,155,73,0,0],[3046,5434,2063,4918,3354,5582,2001,2816,5069,15361,9169,2723,4584,5193,5737,5409,1429,3857,3877,5847,7148,5202,2668,15134,5392,6660,3883,273,6512,11647,604,5435,3974,6174,6507,11711,1281,4896,6919,3310,4650,6623,4151,3563,2223,4747,2975,3631,4093,1931,17530,2096,3048,2481,1551,999,6860],[30333,29018,7076,68848,58424,40821,56242,43935,15152,130720,123357,4006,63515,36107,28289,75457,33384,33706,97918,67843,82076,135727,54501,33122,82379,59461,51759,8138,59657,64043,13937,27259,33108,79939,78760,65298,66611,69059,17486,76710,54764,10811,54376,33095,6653,69693,28940,20474,50731,17704,56775,45209,58689,13615,19046,17821,18349],[4524,6718,2399,12042,7911,7509,6161,5028,5923,18275,16263,2654,10310,6317,8929,16192,1751,6223,5493,6126,9264,11761,10189,7189,8284,15379,5722,2768,13079,14022,2543,6016,5701,5329,15339,13600,416,15288,1836,7756,6727,107,7302,6066,1536,2854,2670,1656,6571,7282,44993,8169,7655,201,3575,3708,1857],[362,0,218,0,159,85,21,17,218,0,492,695,186,1013,0,0,0,0,245,0,236,0,0,0,132,180,1126,0,48,52,0,0,0,0,45,0,0,0,0,0,0,1685,428,198,67,239,208,298,0,0,3159,0,333,565,0,0,0],[476,134,351,177,0,344,0,0,310,669,281,560,267,0,236,0,32,0,26,151,0,207,912,0,693,0,123,0,156,148,0,0,380,0,210,394,31,0,273,29,0,0,357,28,1350,0,462,232,382,83,0,402,87,1015,0,0,0],[2,0,8,0,0,0,0,22,0,0,62,12,0,0,0,0,0,0,0,0,0,0,0,0,0,0,109,42,0,92,0,0,0,0,0,230,0,0,0,0,0,0,0,0,0,0,0,0,0,0,6,0,0,0,0,7,0],[11369,18839,13528,33260,25094,30213,12669,16920,22102,53353,33147,13153,11300,13987,18845,15991,3875,8779,26068,14367,28512,27297,45412,46851,17510,25705,8516,3743,19609,37122,6101,11202,18076,21759,24838,39275,10093,31457,23962,18323,16358,2894,17170,7612,22708,20106,6888,9500,10194,16104,201432,14140,17301,3784,15472,7362,10711],[1266,6042,12065,11212,12886,2877,22288,12357,4003,64828,83639,10719,43719,11227,10360,21671,14421,2692,100760,43786,79898,18417,9978,8048,69873,22436,10291,18173,5143,4740,21266,35124,954,40260,11352,5199,6334,11197,4824,13888,1384,15810,15399,28771,24147,82543,35460,15967,5115,1408,30100,5992,9306,13226,44042,21072,4129],[2506,2994,3768,3290,4304,4436,988,1780,7875,7569,4016,4288,1724,1632,1520,1069,26,125,2514,2117,1215,3007,4579,14621,1175,2582,754,146,2877,5001,1689,2938,2262,2729,5143,6496,727,1906,11600,464,377,3825,5824,1919,8071,6427,1012,6671,1294,2645,21563,2677,1559,498,2999,2044,1853],[1222,111,587,511,451,926,706,523,252,1155,1135,0,432,177,474,282,62,1400,290,58,628,604,565,711,231,1677,654,9,920,1740,0,1412,280,1270,2092,1672,1151,878,1440,422,442,0,2681,1095,184,853,1090,0,228,0,0,670,531,452,451,244,79],[148,158,778,356,0,644,0,0,506,81,577,383,218,199,0,28,0,25,0,755,227,0,2786,5870,356,404,0,78,202,404,0,0,0,973,590,901,0,0,0,89,1743,0,120,0,174,216,1627,2751,1194,0,0,113,242,585,28,0,0],[17,0,0,0,71,0,0,0,0,0,0,195,0,0,0,0,3,0,0,0,0,0,0,142,9,0,0,0,0,0,0,0,0,0,0,204,0,0,0,0,0,0,0,1,0,0,186,0,0,0,0,177,171,197,0,0,0],[124981,152868,11535,286429,190401,296191,114417,119234,269470,352264,413888,19547,152101,184578,189533,207674,99467,154043,201893,306189,462131,257530,278400,320433,124254,583126,294603,51136,479365,675819,28550,162218,420959,154451,172486,229322,138558,694379,96888,148683,261634,27712,141709,90049,43263,112908,51536,66963,205001,249001,1271873,146980,195585,32852,47609,91743,133505],[0,95,110,0,0,62,0,0,109,23,153,0,92,0,0,0,23,55,0,0,153,0,0,0,7,0,0,0,0,47,0,0,0,0,24,37,0,0,0,0,0,0,276,217,0,0,0,434,0,6,0,78,0,0,0,0,0],[0,0,0,0,0,0,0,0,0,0,0,404,0,0,0,0,0,0,0,0,0,0,0,0,0,0,0,0,0,0,0,0,0,0,0,0,0,0,0,0,0,98,15,299,0,0,0,0,0,0,0,0,0,0,0,0,0],[0,850,634,298,0,314,0,0,74,0,845,245,46,76,26,918,0,0,439,14,0,145,1818,0,633,922,338,0,634,2369,0,12,0,260,25,1346,0,134,200,488,266,0,895,689,384,958,97,1254,0,734,2936,484,1071,1463,475,0,0],[2605,1773,898,2668,608,2488,1196,1394,1788,5890,2746,800,1611,1919,1663,1935,1849,2900,2180,3096,3101,1068,392,1112,1255,9433,6720,178,3454,5195,0,1307,1777,1733,3171,3352,540,9019,212,1686,3446,0,2713,1498,1098,949,249,2560,794,1775,1191,1620,1538,976,144,777,5285],[0,89,310,0,77,129,54,0,262,388,665,924,153,0,214,0,0,0,246,41,169,766,558,276,573,386,0,0,409,202,0,175,0,0,703,1180,0,0,0,0,0,1308,187,501,445,267,3557,660,0,0,0,417,931,0,25,397,0],[257,509,112,1076,489,682,1280,1303,12,1136,2135,0,588,293,320,624,44,0,1377,1060,2021,1503,1040,0,1335,1105,391,0,452,149,0,468,386,608,654,432,0,925,0,542,0,0,2981,346,107,1616,326,160,365,636,0,482,1064,14,68,0,182],[0,0,0,0,0,33,0,0,0,0,0,138,0,0,0,0,0,0,0,0,0,0,0,0,0,70,202,0,0,0,0,0,0,0,0,0,0,0,0,0,0,4192,0,0,61,0,0,0,0,0,0,0,414,290,0,0,0],[2626,6134,1249,5507,5322,7444,6484,7373,8553,19437,19932,541,9786,7151,8598,10198,2601,3927,9781,8806,14841,22141,16546,9032,8602,13231,8337,3980,9483,28762,2030,4638,3321,13798,13907,16732,9421,29986,3215,8332,5584,4175,4186,4509,1721,7678,2093,3321,3046,4640,15702,5007,7375,1892,2000,1640,20337],[1659,1626,1588,8648,5328,2215,3870,6387,1423,10482,7916,1251,4943,1395,1418,8159,2509,983,6945,432,8929,8468,5154,4719,6791,1706,217,3478,1925,702,1010,3996,75,14430,7041,4447,1516,1607,1912,1649,4339,3249,2294,1075,3586,12170,5538,9526,3897,1966,15698,1172,2837,819,7160,1412,336],[7599,4990,1775,14363,11922,6177,4045,6932,7810,24669,23886,2319,10927,7843,7034,8500,2076,6471,6106,9039,8868,13244,7438,5109,6843,13412,12415,263,9722,20131,3878,5126,13657,7828,21532,24183,7287,20671,5163,5780,9312,2098,7240,8128,2029,7334,7107,3898,5745,5831,36257,6598,7853,3338,3350,2106,920],[0,302,1604,0,87,570,302,418,600,1062,1174,2818,1319,322,235,882,0,325,0,208,284,424,257,80,297,1011,253,28,1238,1149,0,72,394,110,1551,1827,0,0,0,456,0,6789,1321,940,2850,338,3990,4731,161,40,3604,657,1330,4083,0,396,0],[20079,22835,8130,40734,27952,44538,24069,28090,38515,99535,71038,10818,30624,28290,25092,45125,20724,28862,49332,35662,63302,57122,35510,26675,44758,67086,30420,19962,65208,88304,12913,43969,47368,35923,57782,55744,31194,63591,28480,18262,44691,7210,75043,31820,19147,41195,23353,17344,25808,26924,203625,28825,35994,8834,8456,17920,22231],[202,0,887,0,0,0,201,0,146,0,0,367,318,0,0,0,0,0,0,0,0,0,0,300,106,0,131,0,255,0,0,0,0,0,0,728,0,0,111,0,0,0,826,54,1398,198,824,200,0,0,0,42,498,87,0,0,0],[109,352,794,159,0,69,67,86,146,100,233,654,414,332,478,268,0,0,0,0,142,202,530,0,164,76,24,240,180,496,0,0,46,0,433,276,746,51,0,0,726,0,44,204,1114,56,631,1264,450,0,3363,262,239,957,558,167,0],[3190,7833,2410,11077,7237,6372,3741,5221,8014,18547,4880,643,10566,16229,12054,10869,3065,6790,11438,7017,13754,12796,1342,3994,10932,11029,4477,6344,8589,15470,6897,7884,3332,4037,2686,2625,634,10246,1650,4697,5239,367,12131,8937,3348,8589,2208,1268,6543,6322,27428,6806,7313,3845,5084,2592,5383],[3153,936,6624,353,4,2982,421,262,2663,3934,1502,4589,306,1,2,2113,164,2497,520,75,94,1042,2082,8514,1743,2351,1540,3116,1618,3829,628,678,1063,1484,3359,4212,0,1860,5541,451,544,9593,2294,926,6962,107,280,3213,54,0,8738,1124,624,4067,860,27,303],[0,1267,98,945,2156,1314,1107,2522,5182,2358,5273,324,446,346,1345,1492,0,270,896,983,7471,2277,975,2753,1280,966,273,0,404,204,0,1214,199,804,2181,2708,0,0,802,1660,0,0,0,1003,8609,1158,1280,1005,31,0,8358,1508,1316,356,1162,5,0],[738,1400,336,1824,2070,2215,2081,1626,536,5125,3681,0,1254,2632,908,2317,1234,584,2033,1237,3487,2668,1020,41,453,1396,1869,446,3363,1492,38,1852,676,1852,2805,7457,59,4657,23,2695,833,101,2647,792,1075,1133,1919,4012,1455,97,7387,621,1586,12,321,1130,485],[841,242,1694,55,235,165,487,436,402,265,1441,1877,374,152,256,603,90,192,364,0,174,121,164,1654,466,530,1041,128,296,727,0,0,32,204,1096,586,0,256,3104,70,0,519,990,678,5067,267,3016,4240,162,680,9167,752,1313,1924,471,112,0],[322541,481186,30652,208401,169181,394360,281994,260109,356836,596496,353444,35160,287705,139808,184005,192326,99579,196778,232425,256667,385820,380886,403644,315092,374924,616938,424426,40990,489444,1005409,112218,215390,409267,185886,265372,325174,184958,1089389,403596,214873,288123,90525,270209,98297,126507,217637,126768,207821,265505,191499,1997879,379520,233548,122253,197908,123417,109179],[18994,23369,9312,49971,25881,58805,31070,35984,26800,76870,94563,12194,40402,34670,32108,116723,54455,62273,36738,38271,61281,71378,43314,37848,51013,93192,52124,34356,95654,162563,13418,33407,57196,56039,114096,107707,38139,122798,51551,32901,49066,17039,18929,13284,35049,35939,42370,31478,34579,26903,213284,40144,39922,14787,32954,24762,50027],[0,398,1120,135,1076,0,201,0,316,23,202,1348,0,27,310,0,28,186,659,0,0,480,0,0,202,569,0,53,0,1037,0,0,0,640,406,1585,0,0,2216,505,0,0,0,537,1475,0,808,598,0,0,0,0,0,202,997,0,0],[0,258,519,68,0,90,330,259,62,232,156,1498,287,160,45,78,23,0,192,0,0,0,0,0,0,90,0,38,158,0,115,0,0,420,201,1077,0,0,0,0,0,3918,222,0,0,1,1369,110,126,0,0,0,160,186,0,29,0],[147,99,770,64,15,10,163,65,76,452,127,160,4,235,62,30,6,38,80,83,0,47,0,110,46,200,707,107,343,69,0,0,11,248,234,44,0,44,172,6,7,34,1641,11,957,381,108,444,75,0,96,80,294,621,14,14,0],[71,81,1914,0,0,482,262,230,0,615,120,607,161,100,421,0,0,0,0,0,0,0,0,0,0,0,1510,0,362,168,0,0,0,974,123,1551,0,0,3205,0,0,29,480,1179,1526,0,1849,1014,0,0,0,0,851,1036,29,0,0],[81959,160600,10141,391286,329104,141300,287189,266116,75227,873795,725476,9354,265006,124849,111325,399514,112335,80716,635552,293880,523491,666608,118939,46433,642075,326175,133494,36187,186772,201461,68005,281676,111411,440899,296201,172978,214920,262567,38056,342972,194402,8289,258080,130433,23639,868420,163753,44183,351232,110252,362351,192047,204192,9620,150057,102613,84392],[2783,4111,7556,6052,3390,12564,1954,4013,2345,6764,5766,5993,4678,4275,2948,4787,846,9571,3356,2559,6302,7715,18018,34506,2581,9091,7153,10648,11512,22713,0,1841,11324,6389,10387,19662,6354,27522,16892,1520,946,3661,2340,1807,3824,3895,1141,8053,3497,3338,20816,5326,3599,8423,2126,5563,3875],[295,471,3084,276,84,292,404,199,522,844,582,1183,542,248,345,1492,90,0,238,247,1081,184,24,1485,246,1609,404,24,858,352,0,202,595,298,1480,1015,98,274,2305,264,294,30,226,883,3923,1166,3105,3410,197,1439,5522,998,1844,2297,424,322,0],[58,280,182,63,200,228,209,0,0,54,276,604,68,134,0,107,53,202,0,0,0,52,0,0,111,447,21,62,0,18,0,4,0,0,0,0,0,0,1689,0,0,0,0,0,0,0,646,88,0,199,0,136,368,404,23,404,23],[15000,11775,24182,16163,12074,23857,16678,21341,16293,50161,23882,18636,72815,47349,51785,12374,8762,31415,14073,20291,33866,24620,27933,16067,9734,43911,49864,5440,31282,95828,21264,20332,17117,15701,22491,32479,8006,130581,87976,14376,21551,19314,23064,16798,37486,6612,21043,29285,12744,5492,129773,25043,24069,23358,9834,6037,12168],[1316,2447,2399,10239,10430,14015,3419,4404,28945,17833,16139,4392,7483,13982,12134,7909,1393,4389,21017,13403,9190,32982,39371,46784,7494,7660,2850,9492,9346,22744,2818,3811,4068,23058,18224,33396,2860,9732,8058,7416,8242,6020,8514,6290,9400,11535,4561,8992,6449,3561,47955,8746,11924,1661,4022,4474,235],[112409,251860,59771,450533,318035,240989,194749,183258,353195,862346,598547,100927,216648,165416,196277,198373,88715,78431,326935,170450,432355,339125,258220,352798,286082,466050,188019,65123,187951,430906,98679,198861,398852,296849,231107,244479,156560,318831,186803,196327,165988,99088,284097,150031,199982,422298,139116,112821,181613,115321,1030218,146350,173325,31844,119756,126001,173549],[5766,3976,907,8336,5467,6860,5220,5026,8829,10569,6927,875,6014,7197,5677,11386,4082,4297,4657,8133,5750,11375,36584,23035,3795,3407,3186,3220,8029,16379,3779,3560,7054,9125,18572,15938,711,9214,3220,4901,9702,151,695,2769,1147,4151,3767,2746,2741,3824,9661,3474,6019,785,5704,2652,2533],[80842,227727,25219,257110,192981,251662,86561,104760,130562,351293,285296,32232,144200,169477,179747,148413,51896,66004,100590,217299,166677,150751,185353,148949,193178,363163,117758,62894,329984,598611,66623,111052,200090,120696,348595,213412,83060,325305,119919,76376,234711,45761,120146,143503,125626,136399,89173,83252,132117,125407,1260085,121202,88038,48779,41253,56573,102945],[0,235,467,0,0,8,9,89,113,0,0,290,0,16,53,0,58,0,0,0,32,0,0,0,0,89,101,0,0,175,0,0,0,0,161,222,0,0,246,0,0,0,0,0,140,425,227,773,0,0,117,37,1067,453,0,32,0],[2061,2203,388,7561,5203,13445,7184,6935,15507,14353,12466,2511,4965,4183,4124,5907,3368,5353,8521,7219,13238,11668,8249,8886,8113,11670,12557,1568,7218,12098,0,1951,9874,3431,8351,8260,11013,16706,3429,8056,13253,1402,3405,1537,1234,7717,3274,773,3581,1798,8150,4162,4884,1893,6057,1270,601],[297,0,867,71,0,173,33,377,201,1127,593,1333,476,218,77,24,0,0,0,73,187,0,0,606,0,264,0,724,89,1004,0,0,95,0,948,1412,0,0,105,0,977,0,1282,790,1089,0,2169,171,404,0,39,17,147,1060,970,0,0],[744,1222,1036,1034,610,702,304,436,218,1929,731,319,1085,238,364,3015,398,357,718,181,622,1775,500,504,971,942,0,193,596,157,35,310,0,648,2118,1644,56,200,186,873,283,0,241,0,1333,82,1203,74,252,152,756,610,376,276,644,502,66],[400,53,402,694,50,69,64,0,383,326,770,448,196,0,0,0,622,0,247,108,779,103,729,0,112,236,203,77,503,178,24,0,0,0,65,105,0,590,0,0,0,0,1209,503,0,401,1195,447,0,0,60,602,77,900,0,0,0],[1472,2013,2798,3363,1422,4167,1218,1010,1169,2629,2742,2756,977,1557,2999,2067,820,1095,61,1895,95,1469,426,1978,7605,18504,6844,4763,4241,4292,14211,3475,5231,144,6513,4884,3618,984,11867,66,7360,5251,3775,2502,4488,1545,10957,4300,1208,1495,2519,1515,3713,5031,2581,440,336],[3520,3941,776,5224,4618,2493,4525,4138,4585,8522,6797,1502,3227,2933,3427,7941,542,1385,2763,4392,4260,7041,6413,3401,4614,5197,862,265,2465,5014,1035,1133,4341,2651,4527,5656,0,8945,6371,5107,4810,86,5434,2616,4165,5714,1496,1898,3448,987,9553,2180,2700,962,2996,1227,6],[28,99,51,27,0,0,0,96,0,0,0,75,44,0,95,26,0,16,0,0,27,200,27,190,89,0,0,69,129,288,0,0,0,0,0,52,0,0,0,0,0,0,122,0,2,0,0,49,6,0,0,234,143,68,0,18,0],[9617,15829,8083,17521,17666,8091,17976,28360,1697,89305,67517,8674,13324,13813,9395,59054,60914,12517,68164,25586,17440,21786,20802,24700,35075,25153,10371,11509,8153,7362,11876,8249,3571,16777,19550,11574,39880,7167,7564,33892,20081,7420,14075,32290,8628,61186,19907,8960,23104,2042,11698,11474,15836,152238,12343,3539,2051],[0,40,101,60,0,0,170,116,0,13,126,0,153,128,125,0,0,0,106,20,0,0,0,0,11,28,0,0,316,97,0,65,0,213,572,475,0,76,0,0,8,0,129,38,0,24,80,465,130,9,227,0,82,10,10,0,0],[6643,12022,815,8741,6669,9642,4126,6182,9721,31858,13489,2330,4416,6017,7117,9715,2190,3604,10927,6052,13536,7317,5299,5545,8483,9804,3143,5396,8835,15659,1673,7128,7250,13503,11234,12691,2645,8318,4338,7111,7027,157,8937,4382,3244,6246,479,1008,4100,6299,40654,4430,6603,1249,4636,4683,7172],[0,0,404,0,0,0,0,30,0,162,199,0,0,0,0,0,0,0,0,0,0,0,0,0,0,202,0,0,0,354,0,0,0,94,0,402,0,0,0,0,0,0,0,0,0,0,0,184,0,0,0,0,0,334,0,0,0],[1485,3705,827,3891,3172,5394,4847,1986,2309,11317,10746,1900,2782,5862,5152,5101,2253,4734,4279,4064,9122,10076,5934,2967,3707,7252,7585,1955,9640,12441,3516,3068,2403,3430,12811,4854,3237,14815,4164,3463,6357,219,1771,996,1581,2603,1954,2632,2459,1493,25735,2879,4069,2056,731,2507,193],[10511,7174,3616,31997,16187,11207,8175,7559,4174,29316,27342,11830,6507,9979,9402,9254,3506,1807,9871,21219,20528,12301,32536,32440,5934,11375,10183,1776,7956,12732,6338,3802,5627,13789,12303,16539,11186,9693,15050,5958,5270,888,919,6363,11414,7933,5182,5088,4605,11580,62432,4232,8406,1903,10421,6896,9763],[1766,148,752,270,173,324,0,197,0,1251,3186,915,1488,2684,2008,568,0,570,0,0,720,603,1576,150,0,686,348,0,1040,1201,1999,1290,1983,2740,6737,3070,0,4245,0,603,0,2614,704,88,1108,687,2546,1701,454,1507,0,1337,913,1766,336,681,0],[134,788,814,241,333,223,166,0,586,637,123,404,321,120,235,634,64,363,0,0,59,201,0,292,0,0,257,59,1212,347,0,0,0,350,810,153,0,0,1828,138,95,0,76,600,712,202,0,0,0,0,0,522,70,1704,0,33,0],[109493,167394,26593,230436,170285,175861,140183,145111,327686,560063,447422,62447,152847,146342,143900,188789,55593,108964,176687,198910,279098,158920,132046,128769,173720,282285,154565,36003,245361,337219,104745,182193,182589,144848,252717,275073,173689,318777,209291,133760,200845,69266,268246,151549,145246,171299,120257,124191,137684,122819,1144033,128713,186675,29385,69808,81026,64053],[0,0,0,0,0,0,0,0,0,0,127,0,0,0,0,0,0,0,0,0,0,0,380,0,0,95,0,0,0,175,0,0,0,0,0,6,103,0,0,0,0,0,0,166,33,0,0,0,0,0,272,199,0,0,0,0,0],[144,168,316,148,98,102,0,70,205,199,201,287,0,57,0,94,29,0,0,0,0,0,0,514,140,271,808,74,0,0,53,0,0,0,0,763,26,0,136,0,0,215,2,384,2463,201,2063,1670,100,0,651,190,0,384,180,0,0],[85729,122400,18098,171660,128651,124808,131045,150940,134259,443085,351368,30231,115934,78800,67826,240000,93167,73340,223389,165234,252940,253164,167070,135255,172348,226670,118283,86192,222038,196096,74242,134684,128645,188191,241278,229403,139275,259187,108348,159048,141588,38576,216624,103393,46545,160193,68494,86867,135918,99091,691433,121769,152135,22018,79736,60532,154160],[2420,2244,737,4691,2947,1391,1169,1857,107,9572,6355,1459,2043,1090,1676,10837,1571,3886,1686,888,3734,2939,874,108,2394,1524,247,497,513,1232,0,513,549,1089,3490,3487,0,2058,0,2065,3016,0,1901,588,1117,488,293,2699,2529,478,919,1140,1127,1187,2214,1188,974],[468,563,374,1343,574,1659,1494,947,2745,2280,2094,1238,1719,1030,664,213,604,2618,471,399,971,1491,910,1455,1304,1140,887,245,2700,2899,0,491,24,52,1459,4266,598,0,2423,230,0,1,660,1049,1271,236,15,1105,201,1732,8487,918,975,45,387,845,554],[34,1195,0,1356,1427,1157,1274,1438,1797,4556,4493,576,1492,1856,314,2806,114,550,523,1528,1195,426,272,778,2491,1044,751,0,874,1268,0,849,684,171,853,2454,0,0,3737,3888,923,0,202,23,662,1186,2094,145,519,79,11750,963,1623,195,842,758,0],[282,0,0,3703,3003,1224,358,778,197,2911,1696,243,1263,1218,537,1139,346,900,1451,792,943,1303,0,103,409,0,0,1252,106,1271,0,897,947,116,1833,96,0,0,399,1975,1355,0,1967,608,475,868,382,1201,1653,796,1898,1423,1301,290,410,304,0],[0,0,0,0,0,0,0,0,0,0,0,0,0,0,0,0,0,0,0,0,0,0,0,0,0,0,0,0,0,0,0,0,0,0,0,0,0,0,0,0,0,0,0,0,87,0,0,0,0,0,0,0,0,0,0,0,0],[25855,29015,6772,40923,34676,57835,12147,23665,12966,48328,51364,12574,31654,24294,33051,17399,5675,22021,14358,15139,36310,25138,22840,21619,18479,47975,24828,15079,49156,70650,15408,21064,15718,24786,52521,55867,1951,52972,45782,11983,21019,17957,35564,18940,25096,16717,18619,25962,10473,7769,173686,22712,25959,4616,15251,14774,9371],[0,0,572,68,0,1,0,0,172,0,116,198,20,6,197,0,60,0,0,0,0,0,0,3,0,0,68,0,146,167,0,0,0,0,34,0,0,16,0,0,0,2810,0,126,230,0,1537,416,0,0,0,190,5,148,438,16,0],[14428,10218,2464,10957,10368,19136,12585,11253,18818,62755,39243,2795,8535,12393,9151,13842,8808,3616,12615,30456,17374,21510,15523,13491,18357,29028,10117,20787,25171,24826,7336,14514,8321,13111,20552,19722,10945,30600,8551,15415,26395,0,41478,14503,4299,8310,5727,5801,9716,12501,97116,12723,15216,2125,4612,9566,5252],[775,962,302,760,529,298,163,832,386,2341,1694,242,1002,950,1329,3918,163,880,423,247,829,913,29,127,305,340,808,381,570,789,26,495,202,87,732,1466,181,356,172,364,542,4556,430,217,1022,486,805,741,67,526,1407,261,815,1154,654,107,2027],[17299,21327,3510,26105,12415,18510,16107,15928,18436,44771,54139,3987,17428,24182,27467,18320,8048,11811,19267,17767,34711,29042,36409,16107,14819,53609,16251,15295,28540,47457,11795,19078,19985,23785,40082,50411,3138,42947,14987,11610,18446,6024,28471,11903,7250,15716,14650,5810,18343,22188,58924,16026,20312,4118,7663,7211,10260],[836,1369,2011,10556,7823,6933,4198,3676,1577,6873,4670,2566,6786,4722,3956,4187,2192,3094,8441,5244,5945,8943,8254,1414,4418,10574,9624,55,5559,18361,0,4650,5491,6720,11956,12088,4969,16903,4684,3412,5881,69,1651,2571,604,2560,2543,4606,4012,5359,16271,3732,9155,972,267,2832,88],[5658,9731,5575,12236,6372,14836,5240,7112,4817,26604,34686,5865,19404,16855,14631,43442,15313,15067,13217,9448,16668,15955,9258,12974,7622,18536,8746,6854,12923,19075,648,6455,10142,10811,11642,18941,7334,39186,13783,12450,10655,6423,13865,4378,7821,9726,11463,6326,12816,9789,66533,6508,7044,12069,16938,12147,27719],[44,66,186,88,42,11,0,0,119,28,83,18,24,83,0,0,0,0,0,12,0,0,0,0,242,0,0,0,52,0,0,0,72,8,0,0,0,0,1080,0,35,0,0,44,534,40,156,100,36,185,2155,184,0,164,0,0,0],[29489,42061,16033,58246,35005,61163,28965,37078,48255,84597,73505,19849,37055,42961,43963,35818,14212,25312,34082,31406,54983,49908,87688,110261,27831,62910,21225,13329,48951,82002,18056,30723,35558,46026,77346,71566,23691,81912,66852,27096,44440,16305,31173,27683,43929,33544,28753,35955,18208,21693,227808,27557,45833,7509,23711,22853,36461],[198634,236368,22617,297062,173877,243202,186033,202685,219811,892332,911865,34253,204077,149330,124999,409551,171103,164135,343896,279892,633570,436623,247792,244212,211869,501405,166614,339815,450911,358507,88581,306654,245959,259344,352195,323718,185553,414045,138297,281380,295665,42949,503963,300702,47039,238038,203228,107333,247404,164125,1093808,229748,308274,31436,150450,135415,185587],[0,0,420,0,0,0,0,0,0,232,202,917,0,0,2,0,35,0,0,0,0,0,0,0,308,228,68,0,301,0,0,0,0,186,0,294,0,0,0,402,0,0,0,385,0,776,0,24,0,0,0,0,120,1168,0,78,0],[0,0,0,0,0,0,0,0,0,0,0,0,0,0,0,41,0,0,0,0,0,0,0,0,0,165,0,0,0,0,0,0,0,0,0,0,0,100,0,0,0,0,0,0,534,0,0,54,0,0,0,0,100,0,0,0,0],[17800,23370,1731,29168,16140,14426,20071,20433,20645,70355,65303,7562,17064,9273,7110,35877,13029,9641,37085,13078,28002,63834,36223,31876,31471,37353,11282,17722,24709,33892,15569,15184,12674,50739,41106,56385,7130,23802,4926,22379,19376,32,27225,11182,5929,24851,14821,3847,10098,4990,64793,13496,17250,4439,19815,13172,6574],[3974,829,1539,3547,3171,2620,2038,2866,1738,2995,8235,1613,4654,3052,1751,5757,909,4511,4642,2541,3528,7713,6472,1302,1345,3310,3583,2055,10814,4218,1910,1677,1152,2668,7501,3514,48,3871,2077,1836,4242,23,1046,1674,746,1578,1882,0,2813,2853,3378,3844,4815,4657,478,1279,436],[4580,2356,2507,16494,13099,9918,5715,5300,1203,13201,8723,3209,6302,4202,3723,8943,1213,5263,9148,4698,5461,19386,9874,7376,8951,7932,4742,1561,11142,13428,0,1375,1775,7903,14257,11210,18969,6749,12099,7097,8425,1272,8394,3728,7018,6201,9722,7725,4377,1174,2746,7561,10888,2257,5603,2116,216],[0,0,116,0,0,0,0,5,0,0,262,0,206,0,0,0,0,0,199,0,0,0,0,0,0,0,0,0,179,0,0,0,0,0,0,57,0,0,0,478,0,0,0,49,0,208,0,1303,0,0,0,0,300,212,0,0,0],[513,1772,6997,457,637,1123,289,826,1383,1573,1572,2634,752,687,644,477,111,356,910,1619,1535,2641,5180,7394,2548,4200,1117,300,3225,4577,2393,627,511,4064,3801,1559,1553,427,6746,40,476,1286,606,6510,6141,824,2327,3843,1166,889,7950,1246,2820,3482,819,239,376],[597,0,353,984,565,0,403,594,161,1343,331,400,1444,688,600,1580,0,64,526,966,442,329,2609,183,379,592,1107,0,136,1132,144,298,903,602,711,690,0,0,0,64,95,26,896,0,781,0,0,297,24,887,1600,860,202,38,195,1065,600],[201,274,2442,831,0,280,202,0,335,862,477,1073,621,0,202,0,24,66,467,589,339,149,75,0,303,78,52,0,913,771,0,0,232,114,386,1624,0,47,4190,86,0,0,187,522,2551,56,134,1671,454,0,1162,529,213,714,0,206,0],[327,122,355,0,238,160,0,0,0,232,164,734,199,533,106,0,28,0,0,0,0,111,0,0,2,333,138,0,16,50,0,0,305,72,197,1260,0,0,0,0,0,0,187,357,303,134,300,266,512,0,1637,193,359,18,0,0,0],[158,0,1546,202,283,203,387,0,1002,594,604,1346,226,0,588,0,404,212,0,0,202,0,0,404,542,394,1198,0,248,552,499,0,0,0,795,1011,0,390,1007,0,210,0,1456,1697,3674,0,1413,2787,134,0,2580,58,1185,1870,0,0,7],[4564,3680,1318,5386,3585,2832,2856,2991,3685,12036,11400,829,3348,4307,4051,6597,2395,1285,5574,2717,5022,5190,4274,1945,4599,5820,1447,1011,4900,5637,0,1450,3175,4274,7323,6394,901,8298,309,4674,2630,310,3819,2073,0,2738,789,320,2864,2327,7448,4612,5132,797,2087,476,7079],[0,0,584,0,0,32,0,163,175,0,198,0,0,76,0,34,0,0,117,198,168,0,166,1040,0,194,0,0,114,0,0,275,402,318,166,0,336,0,831,0,68,0,0,54,198,0,741,1344,0,0,0,341,0,308,0,0,0],[388,3443,4493,1185,886,994,494,140,216,2405,1791,1698,880,241,716,725,265,320,137,387,352,405,607,1608,404,356,0,1303,672,751,52,0,450,129,671,2095,2745,354,824,63,617,665,497,590,1443,351,1060,254,23,0,5680,325,391,559,1126,641,877],[404,380,1212,985,597,1705,0,669,1979,1838,1498,1752,102,0,11,555,0,382,8,198,892,0,2451,6442,0,603,0,0,449,897,0,0,527,0,4430,2587,0,351,0,403,0,0,202,672,2590,176,857,0,176,399,3772,31,532,434,1425,1105,100],[2746,1191,1935,3240,860,1720,1951,1930,2385,3113,3757,2778,3111,922,1650,3516,1449,4814,3639,4259,2753,3361,599,5787,1842,3975,4776,978,4568,8969,0,1127,1528,3162,7629,6922,0,1413,5827,726,2177,1190,2890,2752,3907,1433,5290,3630,888,372,5549,5276,2923,1129,2941,3531,2977],[1691,219,1934,1035,0,1244,1052,1111,520,852,495,1850,347,1079,949,404,30,0,980,558,1773,555,844,0,709,1050,107,44,510,3398,135,1114,0,358,1024,2272,0,1041,1221,427,0,0,717,469,912,96,4387,979,107,0,9118,1829,2429,2017,320,227,0],[1671,1324,6492,2701,4066,2918,1885,5190,1059,9221,11759,4867,7043,9175,3958,5513,537,1472,4422,4760,9103,4411,9022,4237,5117,5083,1694,2009,9712,5843,4722,8399,2298,9043,16073,17396,982,8835,2506,1462,6911,0,8630,3749,3718,5922,8955,4440,3215,276,9208,3959,8103,6014,2328,2615,141],[36,0,258,285,102,0,504,88,62,198,332,335,182,0,255,124,36,26,434,652,151,237,1891,738,151,0,101,175,66,337,3821,0,1115,0,877,414,78,349,0,551,1180,0,0,1084,1656,263,187,906,279,0,0,717,290,570,487,94,0],[402,0,918,439,0,146,310,328,385,199,1069,261,150,0,0,0,0,23,0,0,0,0,26,985,112,588,720,0,0,404,0,0,0,0,58,0,0,105,0,0,0,0,338,579,1885,0,609,530,0,0,7184,182,508,724,117,0,0],[0,36,929,0,0,0,0,193,252,199,0,0,76,100,162,25,0,29,0,0,0,378,0,1,444,365,51,31,136,204,23,0,0,0,0,1608,0,702,105,23,0,0,215,210,806,0,0,903,0,0,1291,372,113,398,0,0,23],[203,678,864,1175,909,292,728,1862,82,1374,2896,1013,1914,851,1080,2051,1150,2947,1148,2482,1663,3604,2327,3291,667,1293,1989,0,4014,3280,0,320,0,455,1780,1311,0,2627,0,1621,1168,0,1762,507,147,405,883,163,380,874,4135,1832,1866,456,850,3002,404],[0,255,367,147,0,0,114,0,0,0,0,6,0,0,29,0,0,0,0,0,302,0,2823,0,0,98,0,0,0,0,0,0,0,0,0,0,0,0,0,0,0,0,0,49,568,193,2015,272,0,0,1411,175,94,931,0,0,0],[0,0,672,0,262,199,0,446,0,0,0,0,148,0,0,0,0,0,0,98,0,0,0,0,0,0,564,0,0,467,0,0,199,0,0,0,0,0,0,0,0,0,519,101,1027,0,151,1270,0,0,31,0,0,780,0,0,0],[213,557,329,1906,2719,2744,1129,1118,2383,5230,1800,867,1373,1132,1321,2273,300,980,4208,1458,4678,5538,5644,3280,4209,2899,1416,4168,3955,5351,0,1933,458,1971,1240,2203,1925,2737,1532,1355,2313,1783,2432,484,815,2011,909,2059,1433,402,4148,1142,829,1944,923,446,0],[35106,43679,8721,73851,37967,50176,50448,51328,32492,164794,152262,7409,57409,38926,38868,89946,42585,30267,94000,74097,107803,117896,68278,34240,74813,115945,43171,22875,91201,89116,6558,59670,55114,81301,108789,92390,40809,139088,33550,63617,90126,7792,104666,40828,12195,49419,23470,17241,58693,41193,178904,55754,73559,14572,16248,28458,65339],[17,33,40,181,5,26,7,184,247,122,204,0,0,61,38,58,0,26,36,89,78,10,30,0,50,159,19,0,59,9,0,30,0,13,31,55,0,128,0,66,0,0,12,0,55,174,0,0,19,2,0,6,39,0,0,11,1156],[25481,21449,8330,50413,47738,33358,27135,39320,23830,116938,119246,8211,68607,62500,57093,75943,24594,44622,52467,51127,81322,84465,62566,44223,39793,67197,25544,18200,71011,69114,2725,47230,34165,52954,115129,111294,28724,65896,16513,37879,52517,6241,37252,26338,14310,36687,26871,12453,42345,35954,205245,29446,48393,12072,18111,32924,47837],[701,83,1640,277,457,989,206,418,340,972,1122,2432,513,529,362,479,182,84,237,745,335,245,2767,839,184,1110,170,33,454,442,330,591,0,68,444,1071,0,1499,472,199,24,4246,2710,1345,4912,1055,3606,1802,1226,43,4999,1864,1217,1481,578,562,134],[0,261,1147,176,0,182,128,0,0,0,863,1578,180,0,0,0,0,0,0,156,0,177,0,0,0,491,178,0,654,0,0,0,0,0,1105,497,0,0,0,60,301,0,0,0,2146,602,2469,202,0,0,0,0,317,529,375,0,0],[25721,52783,7696,30260,27356,7592,50483,37875,11846,140413,62396,10522,28291,12506,9069,41821,31099,8135,81250,28290,23906,96070,27714,15499,55812,41947,25872,1937,23599,85907,10525,28792,39693,51181,19949,42199,53564,58166,8813,40975,12140,8258,86934,56997,9687,47573,20218,12433,49656,16557,52355,44942,61321,8203,15007,7061,8295],[64596,78487,4493,145528,107409,135071,86067,83790,88595,271694,335406,22727,63463,97946,92338,58567,16762,38124,138780,131719,247085,131723,101461,71811,89561,271593,114690,26479,176853,264952,14969,85298,119097,82912,158506,149731,43416,217361,61976,69974,100883,10462,129905,86231,33893,119485,103471,45132,87094,105871,869928,78590,134866,15375,20229,29193,76390],[531,75,1025,1010,0,236,386,141,51,266,407,651,178,295,48,631,0,25,0,0,180,783,327,0,100,574,1011,31,1059,1569,0,0,122,1059,321,352,0,54,1930,68,0,0,276,230,1249,508,1740,0,22,76,546,456,575,1159,0,0,27],[396,1079,230,1898,1886,378,1035,1535,71,1225,709,276,937,1157,2202,4166,934,1513,541,660,1567,3123,662,74,2103,3072,117,100,2065,1717,96,1432,1170,3273,3582,2421,1056,3891,0,1463,2250,1543,3850,1105,0,1822,0,0,584,777,99,469,2117,302,754,2960,84],[248,0,565,0,0,59,77,144,196,309,592,364,431,0,0,0,0,0,0,0,136,0,0,88,0,202,0,0,388,0,0,0,0,0,0,0,0,0,597,0,1352,0,51,516,3347,101,1206,2008,0,0,0,529,676,240,0,439,0],[2696,3084,4154,8846,8196,4903,9835,8571,1455,8769,18224,3960,7180,7443,3372,9828,4452,6905,8141,4226,26696,8304,5868,2643,2708,11986,4772,6529,13581,4021,4352,9377,2368,15421,38307,17690,22319,28646,9095,7018,14602,3594,7118,4094,9079,2386,8059,7882,6187,4142,5867,5007,8867,2327,1631,4870,11144],[51217,61195,10520,111570,94301,58918,42943,43725,33134,155000,194579,9282,60657,56519,65425,86889,40909,28892,96065,70979,121056,87635,47998,40717,135753,134948,52746,22841,76103,99606,21932,67552,41192,74235,104601,75400,63999,92843,28081,75881,65473,13019,51976,40089,19105,136755,42264,26895,68279,49964,216060,40687,68263,7260,51623,41147,65760],[286833,279032,30887,724060,582144,510571,402402,422388,304032,1361856,947549,59309,499006,374184,388563,733497,282959,290716,684842,577827,1031484,837861,499669,337708,543351,831176,381912,301107,1034672,904731,239271,558077,462496,578713,920141,791224,492656,950078,248590,552042,631425,90676,649253,317179,97153,507287,306393,189577,442600,435915,2078909,443556,509254,58361,252160,261499,318604],[2617,2261,10418,4231,2697,2640,1017,2353,3601,6228,8879,5622,4612,3382,2212,7707,2709,2636,2797,1547,2406,8712,9241,3241,3478,6531,4840,2312,8163,7070,5208,1927,6121,5892,5456,8354,167,4715,10230,1441,2999,6486,11504,5337,11149,4889,11283,3969,6612,4280,8056,2247,11954,9448,760,2172,644],[5565,1432,8707,14251,3670,9432,3586,2678,1951,36085,4628,12065,4690,2487,2449,5128,6310,2558,18571,3046,4067,10495,12002,4011,10889,10521,10979,6233,11739,17926,1559,3318,1228,2972,3811,9505,506,8855,18755,6794,2367,14279,10236,6509,11914,13245,10455,13823,2144,2794,32431,10295,19486,7233,7058,2457,3287],[1649,2671,630,3016,1996,2451,2192,1631,1379,4610,5451,455,3419,2000,1909,3472,1964,989,1628,2391,3583,2865,1450,2709,1784,4551,1775,34,2596,2936,58,1432,728,3031,3443,2414,2966,8184,4592,2269,2029,9,894,924,2653,1556,1218,2175,1850,1430,7883,2905,4217,462,1503,1507,1319],[9442,7763,7486,46295,29218,23820,23911,20817,56980,54347,31444,4916,22670,30304,23344,32767,6589,12582,86027,16691,17588,60674,36661,12767,78627,58113,39993,1432,21148,49879,5694,18977,38729,28992,20157,22980,34876,28543,7359,24160,25590,12663,18566,5200,13660,48141,11134,9138,13202,11279,160852,20961,17586,18803,21640,6339,5042],[31,22,178,242,98,125,110,0,181,453,12,101,0,319,0,2709,74,206,83,101,111,0,0,0,13,129,49,25,0,17,0,0,0,0,0,600,0,0,145,0,31,45,0,36,223,53,17,40,0,0,0,0,218,0,62,0,211],[0,0,96,48,0,0,178,128,0,64,0,0,50,0,238,0,0,0,0,14,0,120,396,12,57,1,0,0,0,123,0,0,0,18,0,0,0,0,0,0,122,0,0,38,0,293,0,167,0,0,0,126,0,0,0,0,0],[0,0,0,0,0,0,0,46,202,346,0,0,0,201,0,418,0,0,0,0,47,68,0,0,287,0,602,0,199,0,0,0,0,0,291,73,0,0,0,0,0,0,0,410,0,0,586,291,0,0,0,0,43,64,0,0,0],[8924251,13145858,1146039,25367079,13569262,14076662,19365255,20306728,8556238,48310690,43771106,851616,14581504,14076290,14343181,50366804,24821406,15133162,24195435,21646137,18648941,46888062,28487512,13774071,30628689,25995482,14338147,17818640,27982008,30176110,13181442,24311483,17352548,32161841,24093683,16281340,16376748,26803564,5214298,20143227,15566756,1510248,24798275,8883725,2631948,31416351,11423007,8226215,13713959,18214659,82042281,16364440,14572455,1192736,15902567,16568005,16294133],[0,0,474,0,10,363,0,0,0,105,152,312,0,338,0,169,0,427,0,0,328,303,0,976,0,193,176,0,313,783,0,0,0,0,804,82,0,0,1141,0,0,0,160,283,1737,267,1299,724,0,0,220,171,346,0,0,0,0],[279305,337148,38113,284606,243729,448572,161864,197546,448821,609728,507379,59959,260949,278005,269540,191662,71540,209670,176659,253038,335879,313867,392641,266021,223352,971238,432088,128076,654061,1492326,93199,259838,560200,237078,454159,466730,123164,749322,304035,168080,508461,118657,501938,312777,125022,135147,120203,135677,184175,144339,1626439,248303,248390,108443,127837,130706,226098],[0,0,71,0,0,106,0,0,0,0,0,63,0,0,0,0,0,0,0,0,0,0,0,0,0,0,0,0,0,0,0,0,0,0,88,0,0,0,0,0,0,0,455,78,0,82,0,0,0,0,0,0,0,0,0,0,0],[0,0,482,151,0,0,0,0,96,136,445,112,0,0,0,0,0,0,0,103,0,105,0,0,0,502,0,0,170,0,0,0,0,0,124,0,0,0,0,0,0,0,388,0,691,0,4437,0,0,0,15,468,372,873,514,16,0],[56,221,478,66,687,239,76,0,599,480,716,575,39,282,0,797,153,0,142,198,373,152,634,2216,86,606,605,0,0,1796,0,199,276,201,153,0,0,424,0,34,0,0,117,203,199,787,98,0,0,0,3283,76,248,0,462,78,728],[46,606,515,202,0,102,879,146,0,202,0,826,250,200,418,0,0,0,0,0,0,162,0,0,0,167,0,0,583,175,0,0,0,0,404,360,0,0,32,0,0,0,0,0,1024,0,1157,96,0,0,9537,2,388,1426,194,0,0],[121418,46875,33005,68363,91629,81484,62788,137709,237626,322766,204225,25737,158753,68822,142575,26342,16459,75004,62691,78741,105866,70391,510740,828772,61720,270052,52345,34061,238433,277471,48330,42155,113421,322297,282640,373706,40420,234864,774593,71421,231572,92570,177339,14230,166477,208164,39957,37940,281305,236749,1783500,271735,109173,79439,136572,77831,154444],[636,2932,2436,6478,9324,3708,4980,7383,1178,17755,19799,4670,5797,13554,15531,22655,2723,856,16689,13164,45459,11750,30738,6610,11092,20646,8603,1981,4015,4799,4,7265,16593,7791,19902,22547,3199,23593,8763,8443,4042,680,4880,5669,4453,11974,5744,1070,3479,12246,58014,2386,5660,2364,1984,1952,369],[20597,30724,10594,21640,12949,22462,6701,9307,43930,37466,39272,15903,15079,20318,15661,20814,8003,15504,8912,13683,34496,20972,35066,86695,8734,38261,15996,38980,21799,38536,11216,9899,16910,15635,27725,47089,19590,44959,39210,3076,12894,7448,6870,8211,18064,18323,21944,20384,5977,11372,227404,9816,14692,5009,16694,20246,40338],[123,0,105,0,76,0,0,0,0,0,170,50,0,0,181,123,86,0,0,100,0,0,122,0,0,804,7,0,56,80,0,0,0,65,0,24,0,0,0,224,0,0,334,149,0,0,235,0,0,0,0,0,349,100,150,0,0],[776,905,2753,2318,1081,1668,209,1035,1747,2633,7926,951,1805,1584,1421,3278,281,786,327,3214,5027,491,2171,4190,1023,3640,3470,1057,4082,3729,0,395,968,84,3117,2588,1095,5499,6,1344,510,2700,1782,3642,1586,1409,5605,6615,3020,1436,836,1059,4292,2172,1037,471,569],[217,413,112,1385,224,785,179,166,36,1738,647,56,548,56,277,1242,57,561,200,288,420,155,300,868,28,589,213,1191,198,303,195,365,104,442,558,1047,0,168,1140,61,180,0,0,54,84,423,20,0,207,3,72,112,0,28,902,280,486],[31909,44813,3998,82691,58251,69604,84279,77977,51229,216207,224442,10310,64028,75708,65976,104100,28373,33587,143754,49541,129851,160861,56600,45063,149715,94862,54276,18692,84727,88477,6948,60835,38023,112466,158233,91851,46816,109224,20675,89546,70180,3678,52174,49684,8526,119729,34795,17452,83788,43513,168953,48597,74522,11947,31692,27439,28519],[1074,1106,461,389,220,426,200,470,574,592,643,258,1227,414,468,951,76,105,531,496,195,495,9,1126,256,139,44,155,208,508,38,397,0,698,761,598,31,1052,270,199,0,0,0,724,56,586,0,370,120,152,3958,741,68,79,101,358,0],[241,1599,204,1664,472,1203,582,962,587,4952,2887,1194,533,987,905,657,29,592,551,765,1412,1310,42,3134,366,1122,406,132,809,1101,0,121,424,441,1168,2107,240,405,323,0,643,1871,887,312,216,535,116,0,582,1633,4595,295,192,0,596,370,899],[92,122,926,1158,667,886,473,182,118,2017,769,821,465,1233,38,22179,14400,5125,357,485,1573,807,1292,1026,38,200,91,709,159,158,145,19,140,1008,2167,1591,265,947,423,227,29,880,95,336,2435,145,1729,1783,490,198,2621,101,69,259,597,373,188],[255,0,979,86,0,58,202,0,0,0,153,204,136,200,0,217,33,0,87,0,305,0,0,125,768,630,2352,49,404,403,0,0,404,0,655,606,0,0,0,0,0,3828,404,1457,2264,0,0,710,538,0,0,750,0,404,81,0,0],[120186,210865,20611,261389,187707,186820,207160,229531,166429,628554,388497,19582,238762,231247,245392,460019,190656,191763,319451,250488,524717,416759,180806,94656,380422,402699,157856,218106,355229,387839,62482,240795,157449,208484,276248,240560,250322,556216,102332,300316,268471,25122,474302,178479,31624,281804,100928,67649,244897,138821,670343,203772,226640,37427,94463,93486,97456],[398420,963419,35234,6184925,4757990,533802,4476014,3188068,106750,18491298,6105623,192457,3092058,412637,467272,2607963,997885,344129,14485141,1356822,2350355,12006715,826128,363355,14863875,3145335,1792876,157913,486677,59158,1036529,3232171,193994,7519629,2418794,549117,3774938,730747,125388,4987285,948443,67237,3292434,720189,189275,28699562,3598389,699962,2268868,290182,1359871,2611770,854574,12274,1963904,448611,221378],[408,0,262,0,0,25,0,200,0,0,139,272,296,0,0,0,0,0,0,0,47,0,0,0,112,618,69,0,353,237,0,0,0,0,630,50,0,0,0,0,0,83,1656,38,168,66,0,260,0,0,0,167,0,142,228,104,0],[253,5655,512,1224,867,1715,1983,2183,6008,3417,3579,1010,966,1978,1227,2233,408,3962,729,1186,2024,1738,4338,2637,301,1020,2040,2059,2587,2554,196,1065,609,1970,1413,2406,3882,5162,2238,1726,2281,11,4787,2024,6102,503,3004,5336,2209,3808,53375,1722,843,764,1087,3295,291],[48026,75544,6316,76897,63456,59520,54681,63166,49848,184097,178179,11851,77430,79703,76463,92636,36412,42601,92214,93161,129352,104333,92758,62204,70331,110676,49851,20319,109749,96247,17505,48111,50777,81242,122634,118647,35495,159834,38556,73167,97600,5985,74159,37097,12665,60145,42685,20803,71755,29910,242495,51577,60551,7536,30119,33476,60632],[0,0,43,0,244,0,0,103,199,23,0,280,0,0,0,118,137,0,0,0,0,27,0,648,0,0,101,0,0,34,0,0,0,0,546,0,0,0,0,0,0,682,61,730,266,0,0,137,0,0,201,389,176,404,0,239,0],[0,251,691,2131,30,604,383,575,201,1477,932,669,1230,366,1203,2522,201,0,100,1639,3196,149,543,324,201,670,1613,0,3310,1469,0,431,590,903,3612,1401,0,730,282,892,0,1744,640,365,102,0,5474,402,684,802,0,87,829,1205,228,0,0],[52599,61829,40584,130229,94708,134662,50464,40560,180155,203512,65373,29281,102298,36407,45947,57615,13498,27280,116463,118189,71480,144948,156528,201489,161385,83396,53311,28035,130540,273931,35690,47528,54723,123058,88679,140501,68109,121499,217461,47576,49735,90047,64581,33669,87261,131714,24214,54646,38915,26482,362347,57616,39031,37725,67598,26086,28301],[72,0,1289,0,404,87,0,520,0,172,0,1098,297,43,129,0,3,0,0,0,225,0,0,0,0,280,1742,0,0,0,0,0,0,0,189,101,0,0,0,0,0,0,422,2452,779,302,637,833,109,768,9187,133,527,1444,95,62,0],[0,120,1691,29,0,157,0,330,0,534,0,1305,398,393,202,357,0,0,136,0,0,0,179,322,0,23,1459,0,317,177,0,0,0,676,104,0,0,179,0,0,0,0,43,1052,387,0,2218,2727,0,0,209,588,353,2799,0,0,0],[0,0,855,0,0,0,0,171,43,0,0,355,230,155,62,0,0,0,0,0,0,0,0,0,0,234,186,0,0,0,5270,164,0,0,524,114,0,551,0,0,0,0,15,499,13,0,3141,0,0,0,0,0,5,438,224,0,0],[1620,0,805,5468,5048,579,36621,19667,404,74214,6984,1079,24332,6094,1712,106956,37519,6511,28594,10643,5091,86407,12665,1976,29887,11480,2103,17234,1973,734,4692,6872,31,75699,3214,2697,16908,5626,0,24250,3325,20,12644,1987,894,51385,3219,1926,13385,1101,88,9797,5997,400,23878,3577,0],[977,322,651,701,88,933,201,560,164,994,998,542,24,635,684,122,0,0,399,0,476,0,273,965,0,287,0,0,802,1883,0,0,467,164,224,703,0,800,0,0,2250,0,1159,229,278,813,96,438,0,0,0,0,416,0,0,0,3405],[900,1030,2575,794,1049,1360,1451,378,1711,16108,3853,2365,1828,1838,1885,2991,3620,1013,1409,467,2191,2834,5587,1912,2778,4826,2397,0,4963,1967,1178,4906,1013,6047,5528,4245,2307,2772,0,472,501,234,6162,2750,3296,2141,6395,1075,3696,4518,7602,2118,1673,1156,1193,617,6140],[0,0,16,0,0,10,0,0,0,0,0,22,0,0,0,0,0,0,0,0,0,0,0,0,0,0,0,0,0,0,0,0,0,0,0,0,0,0,0,0,0,278,0,0,0,0,0,0,0,0,0,0,19,0,0,0,0],[4870,654,2446,10530,10359,6294,105,0,404,12784,23332,12561,1224,6482,3455,332,326,0,248,1526,20509,60,87,309,162,8541,1939,171,908,3678,56,1387,3665,940,7742,11551,0,9621,8245,977,6333,6151,737,16741,8748,506,1652,2732,827,2981,21229,1089,563,1077,529,4361,2400],[1367,1463,1325,2938,3636,5118,2503,2196,3118,4078,6310,1511,2178,2000,1594,2667,1543,3466,2144,3409,5857,4335,348,677,2409,1943,1310,71,3922,5666,3304,1457,2061,3123,4700,3377,2602,5052,921,2439,2543,0,2221,656,608,1395,1903,3367,1008,1301,2747,1164,3334,1656,1776,756,1411],[738,0,8,0,0,255,0,1,167,890,489,0,303,215,0,583,0,0,444,0,0,0,0,0,0,69,384,0,0,161,0,0,0,0,406,177,0,0,0,0,0,0,391,74,69,871,827,0,201,0,0,203,401,303,0,0,0],[59177,93863,25661,96472,67454,83318,49638,68391,117857,213338,93552,26848,59071,78801,79579,69023,25994,37842,72713,90651,117239,100914,109921,120484,51893,83158,37071,22865,97244,137404,25387,55946,45917,82610,93300,103476,40808,131122,111364,47023,59989,30313,86914,73030,70785,58356,30546,45651,53220,43518,435903,59201,59797,11244,43048,39623,62383],[1312,579,670,2161,1450,1803,734,1050,830,1704,1526,611,1777,544,914,1874,50,59,994,991,3175,394,654,3092,1549,877,849,1925,360,1212,2294,314,737,58,1329,1066,10245,588,3672,540,1138,415,190,96,1068,1654,2057,2099,342,0,168,542,749,552,405,542,0],[5001,4180,4294,13441,8113,12576,9212,8143,2734,14950,17313,6660,12275,11000,9119,17960,3856,7931,11359,10911,13077,24204,11266,8584,9988,8344,13310,8495,14603,20237,310,4418,7225,13661,15601,14196,19238,16585,6516,8193,16917,11776,9374,4934,6998,9435,18947,8452,7478,4261,8739,10563,13057,7880,6154,3877,4331],[141,47,218,594,393,129,94,295,569,387,469,636,789,101,370,147,167,78,154,177,213,149,102,346,818,618,38,14,317,170,169,304,814,670,303,429,75,95,655,51,135,57,162,233,1443,679,296,126,50,35,3640,343,435,14,193,391,15],[981,24,94,15,236,0,53,204,0,22,73,82,263,8,154,0,0,19,0,0,316,54,144,0,0,353,251,160,2761,296,0,0,0,268,1227,760,0,0,0,0,15,990,22,491,267,95,0,1362,34,98,34,145,1779,215,0,0,0],[0,0,0,0,121,276,182,167,173,103,485,83,0,155,333,0,0,0,0,0,0,143,0,0,0,0,0,0,70,36,0,1106,0,0,376,0,0,0,67,68,597,0,322,0,0,200,2201,0,0,173,0,0,0,0,0,0,0],[0,211,1237,372,99,0,0,0,0,0,97,1251,132,169,39,0,429,344,0,0,0,0,460,0,0,0,792,0,704,104,0,0,26,0,760,0,0,0,0,0,0,0,511,318,868,0,784,1977,0,0,0,0,198,897,0,0,0],[27,242,810,34,105,103,0,258,192,438,180,1236,374,96,0,728,58,489,36,254,484,0,440,629,271,43,344,42,60,404,303,0,0,0,36,520,0,0,24,56,0,132,0,293,1805,151,0,2228,156,0,9339,1274,51,634,169,0,86],[4581,6576,2915,9656,6541,9836,3988,5466,5064,7996,15945,2117,11329,11364,7533,11200,2815,7033,5097,8915,11542,6590,6041,2679,4135,7191,4143,6657,9876,18618,3178,4714,4200,5682,21597,28886,5888,27828,5616,8282,6592,8561,8908,3607,6382,1731,11671,7504,5212,8137,28831,3881,7881,3675,3415,3449,6050],[0,0,0,96,121,351,152,273,0,879,304,0,0,152,7,0,0,0,575,247,145,14,612,0,152,1031,0,0,304,109,0,3,440,89,24,182,0,0,748,24,214,0,0,254,232,1071,455,372,275,0,1357,75,0,0,195,0,4],[0,56,2434,210,168,249,0,115,202,348,332,604,852,151,0,238,0,0,49,0,25,0,0,124,477,0,606,0,584,256,0,0,1405,135,808,425,0,0,0,0,0,0,1405,505,971,425,983,814,0,0,2652,392,255,455,203,0,0],[347,871,2253,403,0,416,609,350,123,582,588,1148,871,496,101,0,25,35,295,82,137,167,1751,988,0,365,0,0,963,1218,3092,96,0,89,976,633,0,1201,0,4,0,78,1452,963,2933,349,1914,1507,101,0,7193,524,102,1142,48,3,1067],[529,1140,484,739,1110,481,634,517,623,2013,1861,214,1408,1089,1912,1692,1244,386,396,285,685,171,2643,24,280,102,610,214,1205,1367,128,1352,0,389,1717,2851,81,1046,24,455,1136,117,162,157,1324,0,23,907,287,326,444,304,0,346,0,130,52],[279,96,0,203,96,134,100,35,66,162,211,0,50,252,70,813,50,35,0,112,0,0,46,0,0,35,0,86,333,0,0,0,0,116,51,0,0,422,0,0,50,0,46,0,0,70,0,0,0,0,184,162,261,0,204,175,46],[108,0,1363,799,278,476,0,312,202,0,179,802,398,0,0,28,131,75,0,0,31,0,1400,0,0,255,0,78,199,199,0,0,0,0,2342,252,0,60,0,0,0,0,2547,420,805,0,912,1395,0,0,5745,415,207,795,278,243,0],[0,0,0,124,0,0,0,0,0,0,931,202,246,0,164,0,0,0,0,0,0,259,0,0,0,0,0,0,0,0,18,0,0,404,404,0,0,0,0,0,0,0,0,0,0,0,204,0,0,0,0,0,0,0,0,0,0],[189,462,0,0,0,0,23,152,0,171,223,248,128,0,0,0,0,384,0,21,0,0,0,0,0,189,0,0,340,178,0,0,0,0,40,0,0,4,0,0,0,0,256,67,283,0,200,132,0,0,159,0,535,86,0,0,0],[362,1089,3357,2742,2546,2854,313,804,2432,3747,1502,1929,567,219,709,1102,148,265,586,346,1521,212,465,362,346,293,0,315,136,1104,166,53,184,214,1254,457,171,875,916,258,426,1919,934,231,1683,889,83,791,200,65,2762,127,477,243,711,346,629],[79,87,47,0,0,0,24,110,238,249,121,203,5,85,0,66,0,0,0,13,0,0,54,0,0,0,17,0,0,0,90,0,0,0,0,130,0,0,0,7,99,0,0,31,0,0,0,0,0,0,0,66,0,15,10,0,0],[698,663,323,1624,1461,2016,1095,1535,680,3333,3245,292,2139,1342,1087,2767,1651,761,3082,2355,3368,3200,1204,1157,2683,3475,1822,204,2683,2704,0,1744,1544,2180,1996,4026,2391,2852,1319,2679,2475,0,3424,1171,3,1964,592,933,1180,1894,2854,1752,2842,639,999,288,622],[5410,1842,2836,11859,6046,4015,3843,3271,1331,16728,9245,2692,6430,4744,4626,12159,6675,9232,5436,7228,5837,9919,3960,1165,6453,23116,8937,3570,13238,4139,20002,16251,5463,6080,3794,7732,10978,5413,1801,6235,11603,5422,45081,12998,2921,2283,850,2689,8900,6350,6890,9390,15319,2039,2114,2887,3373],[116,0,284,0,0,0,62,138,0,55,94,177,0,0,0,0,26,33,0,0,0,0,0,108,0,0,292,0,0,38,644,0,0,0,0,0,0,0,0,0,0,0,27,98,199,49,375,328,0,113,0,0,0,162,2,0,0],[0,125,256,0,0,0,0,0,63,227,283,0,0,0,0,266,0,0,158,0,0,332,0,0,166,0,0,0,0,734,0,0,0,0,0,471,0,0,0,0,0,0,0,200,0,0,0,347,0,0,0,92,295,48,0,0,0],[168,0,0,0,0,0,0,0,36,370,217,0,27,0,12,5,0,0,0,0,0,0,0,0,0,27,0,0,0,87,0,0,0,0,76,108,0,0,0,145,0,0,0,0,293,0,0,273,0,0,31,0,0,159,725,0,0],[130,321,634,226,69,345,0,201,105,1116,292,524,69,296,23,48,23,92,0,0,69,0,107,54,0,23,0,0,69,176,2860,29,23,0,667,25,92,72,23,0,0,71,824,431,384,268,0,23,0,0,13143,0,201,747,160,48,69],[22201,28170,3079,49307,31436,43928,12969,25380,75158,62929,37278,13885,14935,21772,16428,14664,6103,9148,23557,25546,58863,28579,68123,83916,14592,42565,13988,14884,39713,40056,8340,13347,18033,42102,95685,98972,7263,30894,54205,10650,33393,13079,16099,21287,25330,19282,5932,14152,12224,19765,184035,13452,40232,5434,10350,22239,22370],[146,0,961,366,89,591,245,0,377,516,1948,1202,725,117,0,39,0,0,0,138,269,1046,2424,0,114,384,2000,2,874,1322,0,0,198,538,202,433,0,858,142,301,0,0,195,268,717,1001,2311,909,1114,0,52,0,1926,1081,707,53,0],[1056,399,0,448,491,869,18,45,0,66,662,6,729,289,90,1062,409,162,51,195,172,73,18,286,47,163,8,18,270,264,34,0,0,338,1033,88,6,2213,0,6,24,30,1,197,0,101,1750,173,0,653,2733,486,37,531,299,13,12],[830,649,6493,5164,2641,4885,2439,1100,2215,2962,10436,4358,1219,6378,5827,9220,6334,5390,823,4694,2987,1004,4131,4397,785,1995,2457,1337,5837,5889,111,1568,2849,317,5192,3664,374,2242,5776,976,3729,2595,5987,3060,7945,246,5346,5810,796,1637,12060,961,5808,2412,800,1471,121],[1665,3044,0,1440,1355,3600,2378,1684,3873,6303,3903,292,2870,2041,3206,37,1232,3127,2470,3685,4444,2489,2748,984,2503,4755,1833,4606,2507,5374,3414,6843,7416,1340,5182,4510,840,4564,6406,1169,3069,1045,3590,2352,2247,3901,199,5215,874,2025,3135,4943,4557,494,0,625,4477],[68878,133391,28684,221657,141967,81416,129150,131749,39880,377908,252790,47488,113426,58999,73641,171767,78441,48973,154541,84657,216064,239429,70457,135224,156078,217666,58598,33754,86582,197868,44757,155160,151531,219211,168213,115820,105807,163313,71877,110616,53537,19866,83983,40342,54936,218402,59387,49607,80048,60716,541914,104230,81978,8277,98487,57890,73523],[0,0,806,250,0,0,0,143,0,0,588,535,400,400,0,0,107,362,0,0,0,306,1455,1514,0,101,0,0,392,0,0,0,0,202,371,202,46,0,0,0,0,5020,0,249,609,0,0,0,0,40,240,0,194,378,24,236,0],[0,162,697,0,210,150,73,260,55,0,17,875,125,4,0,34,0,0,0,0,0,25,0,0,0,208,504,0,176,55,0,0,0,70,0,75,0,366,0,187,0,0,306,0,768,502,4293,827,0,0,0,0,166,1028,0,0,0],[19035,22591,6942,41533,37650,22647,32595,35876,29191,75230,64187,11824,39665,31975,23733,53554,10525,18442,34163,36163,46204,47702,42567,53972,42483,31486,15663,17243,34735,45520,4430,15458,23602,40592,49969,59031,44136,43591,28742,32051,20917,8769,51608,27019,19208,36747,9342,26177,26624,11185,103634,20669,26939,3403,27486,18389,19272],[296,201,311,262,522,148,36,159,330,406,577,605,572,428,114,613,0,0,163,0,1634,164,1180,2456,401,284,667,801,932,805,0,374,124,1120,491,1382,0,407,278,315,0,0,1040,772,258,561,2795,0,0,407,9064,327,857,102,547,0,28],[0,354,6,0,96,34,309,194,0,0,64,0,59,362,202,344,0,374,174,164,836,12,0,17,293,48,1072,0,38,288,0,0,0,0,333,18,0,87,0,0,0,0,150,44,0,456,0,0,0,54,0,13,53,325,0,43,0],[82,908,582,2920,1334,1466,2155,1704,1488,3581,4918,0,1914,2389,1342,1825,1292,1073,2067,1346,3220,1135,289,1108,1869,2652,3037,13,3342,3459,167,234,1274,306,3462,2683,1310,4664,2500,1400,3418,0,2415,1860,614,1751,2100,449,3580,944,8449,2178,1643,314,382,1258,0],[439,738,2323,12,916,3889,820,870,171,2819,2341,764,1502,2890,1070,1813,1656,2000,904,523,3258,2935,0,3531,1014,1816,251,1482,3707,2563,0,202,198,1214,2824,4823,0,3049,611,1053,4565,1322,495,338,1789,591,3052,1386,2574,1098,3431,618,1177,1285,1082,847,0],[387,0,196,0,0,0,0,0,0,80,0,111,0,0,0,0,0,0,0,0,0,0,0,0,0,0,0,0,0,0,0,0,0,0,0,0,0,47,0,0,327,0,0,73,0,0,1421,0,0,0,0,0,109,0,26,410,0],[18472,20059,4583,28316,27468,19121,21030,22009,3162,72471,43501,7803,28902,12389,11663,34901,15361,21648,36117,15077,32982,54152,48234,24456,36936,46281,17456,9192,35633,70351,0,15957,30135,25227,43170,37097,16361,47159,13527,29600,16354,4229,21368,10435,9306,17503,17229,13826,24165,17969,70074,19914,17657,4414,13875,13393,21544],[1362,0,675,447,1566,1057,86,603,296,2772,2465,341,1169,1913,1510,1056,361,1434,577,1381,183,1474,2519,436,910,450,270,4954,1838,1872,0,1268,1174,922,2747,266,0,2818,0,1566,5206,0,0,178,0,348,684,1355,1665,387,4344,817,664,294,996,837,2552],[0,0,0,0,0,0,0,0,0,97,83,83,0,0,0,0,0,0,0,0,0,0,0,0,0,0,0,0,0,0,0,0,0,76,202,0,0,0,0,0,0,0,0,0,1052,0,0,0,0,0,0,0,0,0,0,0,0],[5279,10175,6199,25667,19850,12153,26216,32003,10767,105147,56460,10545,32517,15315,14248,60509,11595,7816,45094,46705,38074,63292,29937,30189,20769,22982,8461,8199,32798,20875,20395,27941,10115,29968,19848,19430,11671,17822,16495,39138,23467,19399,47847,17565,17396,47816,35360,19142,40584,19650,73145,21462,34616,2355,15869,17763,7412],[14,578,539,253,19,403,940,374,307,2285,876,323,2112,418,0,590,228,171,388,118,167,1223,0,869,1387,825,602,87,153,60,0,2016,168,1545,629,44,0,51,813,404,0,349,1422,628,2015,1157,1597,145,856,195,4082,1235,608,16,491,10,0],[3510,5397,850,16377,14849,9159,12807,13012,11321,36921,28692,1144,15067,8789,9010,44046,17977,8964,27722,20733,17792,22123,6095,11970,27088,19730,6927,3617,12233,10303,2598,14040,8044,17178,13340,10023,23324,29570,2901,20782,5052,27,33518,15801,3446,29810,11548,7210,21545,17604,92168,14562,14237,958,13416,13208,6628],[24759,24246,9329,33163,21313,27497,33659,33025,29044,65824,96537,7836,26842,27296,31948,64173,17295,14251,46609,34888,47670,62988,28532,27594,44668,35647,16670,16837,32319,29419,8472,19954,19332,33194,51293,48282,29276,59346,13281,37343,48103,4459,24790,17585,7662,40744,20991,9104,34077,25078,131331,22766,36817,13656,15556,11397,13935],[2171,2788,1074,7114,9364,6468,3502,4201,22303,47070,111613,15594,44230,50327,65895,5934,2535,27110,2716,5793,257998,8655,26047,6295,1175,28103,9418,107,7419,36172,8878,18151,16385,10962,38029,113837,1195,109773,28535,5055,10973,0,7274,3167,6455,789,14967,11292,9753,6192,126369,3465,3621,3000,1958,4390,0],[412,2019,1236,2590,1295,2361,1047,2779,2625,1114,3684,1672,3318,4583,2508,2769,33,690,1121,2458,5244,2035,8438,4213,454,2775,714,784,2290,2375,0,125,379,2930,3507,9029,1264,2270,1776,2380,2774,0,1201,530,1674,2221,4073,2567,437,1190,2794,1237,1386,876,3419,732,231],[27241,34182,11433,79634,51274,45384,61646,50262,32404,125907,140548,8881,61107,30373,29536,77192,25187,37355,75340,58405,72079,108994,53114,27839,98463,120774,73110,6457,81076,83546,16057,58386,37789,75740,82654,105612,53119,107847,51108,58242,43752,17073,95623,23113,14609,79082,32495,22948,57472,36143,214360,55138,60699,12800,20071,19858,41129],[516,872,2221,436,162,202,50,352,339,436,387,1256,508,224,21,39,0,477,82,279,108,404,0,2481,558,720,188,0,1144,870,0,0,166,0,334,0,0,16,1900,71,0,0,3501,1721,2539,338,1875,359,0,56,522,956,2924,2357,510,263,378],[42974,96070,28242,103985,79950,71378,90414,88396,86744,291803,250509,29426,90414,96850,90336,119843,48789,65023,151387,98813,182904,250134,127748,128156,108059,186261,54171,84188,137270,181061,26141,102509,106469,139716,173393,140563,92477,165192,117970,100149,112169,22175,123539,52642,57584,107455,51587,44414,73094,68211,406899,85237,93590,22502,33175,55129,113568],[3892,13531,9104,9358,11870,10305,1888,3058,4628,18635,6969,6032,7520,3740,6096,9016,2158,8922,2606,2806,7629,2077,2290,4125,2085,4411,1958,1661,3733,7557,1464,2095,4714,3310,5760,6600,1419,5534,11877,1820,8197,3386,3226,3036,9129,2268,1050,2888,1309,564,9649,1825,4251,1167,5493,2391,5930],[0,26,1221,187,503,146,589,207,0,861,146,1014,43,78,56,1665,1069,399,83,25,149,94,56,244,404,258,856,194,24,252,32,0,54,24,898,26,117,26,27,81,1042,49,831,61,643,0,4241,137,58,0,949,394,26,756,25,85,2501],[130,169,1526,319,0,762,201,2,535,300,1291,2431,11,202,564,0,28,0,0,0,377,0,0,144,1572,3006,1455,82,1335,580,4028,0,874,306,944,0,0,0,4681,0,0,978,69,403,2258,401,4305,1514,99,0,10260,808,805,2142,0,0,0],[4839,4636,1904,7974,4562,9320,4972,3637,4022,14070,9011,3404,10018,4712,4928,13100,13358,20554,9022,4776,9333,13146,11322,1878,5311,23236,11919,3619,15669,42504,244,9740,17395,6504,12496,6185,1266,39958,5275,5800,11612,1205,2181,2928,1862,4287,5898,3624,5509,3296,8533,5154,5044,5149,2889,3826,6960],[0,378,1368,70,0,198,0,0,202,0,154,606,0,722,153,606,84,0,0,202,0,0,2865,0,0,208,70,0,48,1345,0,0,202,392,35,653,0,0,1956,0,0,0,0,0,0,0,0,0,0,606,6778,0,169,1051,0,0,0],[420,0,770,148,74,355,255,16,107,146,1010,969,358,0,74,473,0,0,319,177,26,1045,124,0,531,100,105,184,1016,860,1393,68,74,89,186,11,0,97,67,709,2,0,370,281,823,333,1283,1096,0,67,0,129,109,920,731,329,0],[9870,8216,4883,15967,24856,35512,10958,21107,7608,23747,32829,7020,37359,42445,24353,15672,5872,9722,7164,14832,50251,14530,24075,19443,4901,19671,16255,3935,42657,31502,20218,24611,11725,35298,92229,107959,16808,69301,66364,11315,18738,1634,25054,7989,7684,8213,30894,36300,10439,4448,42347,14351,29910,4763,2328,6267,1604],[17342,14307,12443,31517,23450,32398,12255,13815,7790,31758,40727,10726,27615,17430,20425,27012,9846,13232,18434,19779,18682,30856,41268,39618,19090,32120,21218,14512,45327,66084,11153,12827,50006,20656,21523,33912,12433,46738,40019,15438,26013,15137,23097,19831,28814,18159,26995,31690,20754,12831,42942,23090,35426,13260,17301,10716,10906],[448,0,310,125,0,78,0,254,0,462,228,0,220,359,132,0,0,0,0,132,235,0,1980,0,0,0,66,0,14,0,0,128,0,66,166,972,0,0,0,0,0,0,0,0,0,126,0,0,0,0,0,0,132,15,0,0,0],[728,289,1525,2083,3675,2923,577,1738,2269,1437,4592,919,1509,1200,2188,1982,1954,676,998,2249,2030,2194,3986,5859,876,2286,2153,611,3584,4594,0,187,1084,161,1308,2974,3509,3657,1118,501,0,0,1476,1634,2645,1007,1787,1098,747,1002,3065,681,2229,1010,3375,1889,451],[8473,17770,6488,9050,6350,16870,8863,8240,12608,25540,27901,4982,14371,13520,9727,18407,4995,12507,8084,9261,7507,11682,5589,13893,8649,22044,11475,4065,26213,22496,12560,23878,12067,15789,25660,25193,7593,13306,6283,6298,9704,3043,34445,8107,9569,7005,10937,8557,13456,6588,46763,11385,20707,3855,5919,4076,3571],[34113,62357,9671,60649,42812,64612,42015,42052,80181,113488,126216,12131,55144,155349,120764,64546,27725,38865,49286,42214,79870,82680,48013,51887,45302,88904,33223,18769,88581,118288,8216,37285,58657,46503,66703,85415,47278,153424,81335,48821,64924,31315,54491,33771,32387,44409,17444,15561,46254,38494,347070,37724,57871,20124,15844,31192,60934],[119,81,222,0,138,148,0,404,452,299,396,808,0,205,0,0,0,0,0,0,0,152,0,0,148,1012,0,0,88,0,3039,0,0,0,126,785,0,0,0,0,0,0,1342,515,1528,402,200,20,10,0,0,55,372,0,0,0,0],[0,0,39,0,0,0,0,0,0,80,0,0,58,0,0,0,0,0,0,69,0,75,0,0,74,0,0,0,0,0,0,0,0,112,0,0,0,0,0,0,0,0,0,0,0,0,0,0,0,0,0,29,72,0,0,0,0],[4350,2726,3711,8993,4342,9135,1912,5109,4364,13909,17092,4773,9261,15659,14564,8192,2669,7644,6149,4309,8582,5414,4294,1907,6942,12744,7501,4933,16905,17892,980,4570,5181,7360,10784,8496,2424,12332,2712,3920,17167,3251,7519,8167,5412,5960,5938,4071,3284,3534,31226,7943,4987,2755,3071,5874,6489],[590,715,543,741,881,331,72,512,0,649,61,292,19,772,220,0,25,20,202,1395,1214,126,890,1855,0,1259,1898,25,723,1796,0,61,72,228,852,860,0,377,1101,0,0,0,707,476,969,243,688,3477,77,359,1698,37,377,543,297,0,0],[0,0,0,0,0,116,0,0,0,0,0,128,0,0,0,0,0,0,0,0,0,0,0,0,102,0,74,0,38,0,0,0,0,0,154,0,0,0,0,0,0,0,153,182,0,0,0,0,0,0,0,0,115,0,0,0,0],[1292,853,1506,1685,1606,835,4435,1758,202,10287,2467,1683,4076,2094,2506,3923,2799,1859,4306,506,2351,22087,2333,1420,1285,3697,5129,934,2094,1309,1003,2161,490,3637,1139,1823,1436,3339,2557,6773,715,536,2210,1377,2077,11629,2835,1266,1122,794,3079,2394,2238,794,2019,2246,1394],[256,771,531,435,1539,615,1951,1206,750,2960,3615,1516,2733,3108,2475,2686,270,1447,84,207,1606,1155,1960,3950,318,2103,1546,504,4170,5568,1154,4281,1177,827,5570,4589,11259,2373,513,1489,1418,23,1268,171,2165,0,3583,3870,344,159,172,938,1616,2365,43,5,0],[1804,8016,0,11400,8755,1756,11984,13379,2288,37819,31693,357,10640,5210,3061,22274,8938,8765,36998,16313,19745,40634,8307,2317,17011,25894,4783,7522,6807,6894,9078,12018,2234,30712,12533,4973,3293,5136,1525,11466,5318,706,16630,7554,493,25966,5029,857,19886,15002,41982,10185,4807,0,5902,3793,4161],[0,101,917,146,0,0,0,200,0,0,182,346,0,256,182,715,0,0,0,146,0,73,0,0,98,0,950,0,135,19,0,0,0,0,288,0,0,0,0,0,0,23,228,101,410,0,890,604,0,0,0,180,0,959,0,0,0],[7459,16297,3941,14947,14308,13551,18444,16008,2394,38107,42155,5759,23755,28157,24616,33241,19796,34411,35030,39509,25543,36355,21702,10187,21178,29461,18483,10557,8382,22400,9632,8620,5424,20082,25993,20447,9999,26616,10009,35492,21621,8745,49483,9572,6302,26199,9861,6906,16980,9393,76871,21283,19457,5282,4185,8617,3116],[80027,111399,16337,98432,74348,55254,57544,77542,48116,343537,242120,23575,64813,43737,40965,117158,56871,70443,93441,67815,149886,127383,67588,101425,115329,78279,31668,64200,104463,100485,33001,157507,46714,100995,146624,129225,76717,104931,81303,56109,71434,12159,96914,62341,47144,128128,39135,42725,53378,44116,167748,71855,73321,8453,62169,44870,60450],[86524,80414,29865,142050,109670,160207,49771,61209,139759,215013,122768,37966,72490,57305,61827,111343,38518,50973,84945,98649,108975,127035,198266,354478,111439,99658,36509,72353,139691,163921,56876,74623,77437,153993,191987,196396,45620,95701,201641,52949,59811,34215,90119,74960,101131,75522,37999,73223,37866,47881,470994,81173,90561,31667,92393,73997,100751],[17887,20322,2322,67084,39614,30175,37591,39313,4972,154031,107778,1915,24283,43513,19498,1050439,185024,211847,26133,18689,29206,72616,28021,10095,20464,17676,6812,139740,33183,26444,10931,15582,9858,28481,35163,34094,13801,22534,5559,34573,31879,5923,18784,13159,900,29248,4923,4020,12679,12152,45985,12643,11629,1057,53400,42844,53951],[851,1736,2391,1527,1567,1799,129,811,884,3189,1522,2614,1337,792,1464,971,543,322,715,538,991,779,739,1599,673,832,146,557,962,1315,301,969,562,681,1295,1321,311,919,2206,444,1522,1054,714,337,2147,1293,1174,843,759,269,8256,1120,824,54,1458,509,2140],[4393,1846,2951,6509,2832,4047,3013,4718,4940,9040,11910,2975,5140,3744,3939,13384,2442,6569,5251,4332,11231,8303,9749,1643,7167,13139,6382,2619,8215,11157,31,3961,3813,5554,13849,11076,8010,19294,9778,4885,2427,2282,4854,4609,5055,3005,4317,8974,4554,6250,22117,5756,6281,3138,2238,4546,3061],[15542,32341,8820,62781,45544,61016,34,70,170,558,230,228,57639,50295,44203,68067,32837,56466,118,121,19,76632,37394,30438,41,53,0,36540,75810,103641,58,235,0,54425,71749,63456,116,198,177,48156,45263,2512,40219,25702,31072,44378,26630,18847,32980,20660,37569,33847,29547,5291,20017,31952,51753],[0,162,58,145,78,24,49,24,24,192,227,107,64,198,37,127,0,189,0,0,48,0,0,0,77,43,0,258,0,71,0,0,0,26,24,0,0,0,0,0,411,24,0,26,0,0,93,0,0,0,0,104,0,87,119,33,104],[6958,4393,2290,4965,2638,6458,5649,5784,6039,17690,19654,1467,8746,3982,6081,24834,20232,21710,8750,4681,8315,14310,5632,2163,9124,25533,12531,11626,16259,12897,720,11187,9098,9958,21226,11867,4044,16463,2521,11979,17763,1456,16974,12847,3134,1305,3808,3934,9751,13813,28519,5792,11144,2998,3720,5257,12638],[288274,585162,105630,816804,633345,685758,399953,415150,668545,2053888,1124594,210939,459067,440429,474648,694113,403080,429702,634268,503405,762620,976469,895916,1640365,536749,703129,325592,1023078,683274,1206168,230558,495692,518470,948214,980211,1253081,489773,1005204,663176,289049,352289,179508,566546,320452,504958,941300,370846,360937,382114,391354,3040277,385811,463561,97880,582315,465612,809339],[0,0,463,0,0,142,101,0,416,0,141,903,47,0,2,0,0,0,0,0,0,0,0,10,0,822,0,0,260,839,0,0,0,0,89,0,0,0,0,0,0,0,164,172,296,0,1466,1010,0,194,333,0,151,1077,0,0,0],[356,457,488,417,264,122,990,510,603,2463,393,799,1790,1960,664,3351,1688,560,1210,214,607,5299,5412,6122,1364,925,404,977,680,567,2545,238,196,1772,1338,1547,544,355,1079,2313,363,670,1865,657,521,2179,2161,2186,944,25,2446,1766,1380,182,4620,1228,2334],[3918,8110,4774,12283,10140,12808,10014,10119,7137,26973,35516,3271,14717,10443,11529,43100,7620,16317,15907,18329,18763,25054,10696,11052,15234,18944,10163,8468,37825,21036,0,20696,10428,15985,27942,34981,30112,34428,2901,16129,27959,436,16309,9844,5248,12020,14333,10386,19459,9045,18352,13232,16025,3947,8210,9093,8016],[104,0,796,895,1901,894,2610,536,0,4231,0,591,813,300,0,3599,0,556,3395,121,263,996,946,0,2998,1180,124,0,1061,558,0,0,0,1562,152,684,0,1056,0,2615,6,0,1900,279,0,6020,150,709,384,0,0,646,276,256,518,665,740],[120,150,430,99,0,0,101,440,411,99,391,1904,0,303,21,0,0,7,198,253,205,0,1006,732,0,0,0,7,226,416,0,0,0,0,1126,52,0,0,0,0,0,0,0,263,945,0,81,336,194,0,1058,0,219,10,0,63,70],[1112,901,464,169,0,726,450,113,3503,378,1240,196,227,4118,2975,146,1170,4482,0,32,745,152,13,151,99,3036,216,0,1076,4375,0,0,1400,159,196,2156,0,4986,1268,15,0,0,1400,220,75,0,927,0,184,0,2475,150,941,649,4,0,0],[0,357,998,0,0,80,0,0,0,197,385,0,0,694,1607,1002,0,0,605,2505,0,0,0,808,0,404,0,0,385,0,0,0,0,0,0,0,0,0,0,290,0,0,759,684,784,123,0,384,595,202,0,598,765,1521,0,0,8048],[208,54,303,5,6,1,7,7,3,159,21,155,603,6,568,0,6,0,10,7,2,13,811,110,14,225,198,23,963,8,0,4,7,478,543,1174,0,9,0,10,30,0,17,59,2584,603,0,748,8,14,125,395,1011,1,0,207,0],[98916,137307,16765,170380,118431,150281,117961,131588,156267,437853,313090,35658,157154,128052,134563,187915,78430,114306,162073,103903,201193,215844,153091,136797,141748,213400,99162,49926,176744,242774,26115,102145,136640,186110,288129,285071,168888,304629,172052,141307,151013,41648,127963,84038,65421,152040,82122,60234,118669,78080,683784,102952,138046,31139,92973,117507,288297],[0,202,0,30,0,155,0,0,352,402,384,0,199,0,0,54,0,0,0,190,309,0,201,992,0,198,202,0,556,0,0,0,0,60,402,0,0,0,0,0,3738,32,360,152,1069,0,1155,0,0,0,0,71,287,915,6,0,0],[476,2090,2917,1383,1427,1292,209,633,804,4335,2318,3242,1113,1895,1953,2031,1681,693,814,834,1830,821,399,1705,1425,1621,2256,3640,2236,2058,40,480,270,1069,2942,4097,134,336,1493,722,1917,835,3233,1957,3685,1434,8937,1425,761,2130,4158,1523,2695,4664,550,294,133],[2087,1376,103,3274,1301,2745,1902,3512,2294,6397,5751,215,1812,2317,3048,6384,2446,1694,4077,1425,2582,3643,7692,6077,3132,5550,1355,1872,6194,4810,2551,2386,3609,8164,6529,5038,3002,4435,2369,2081,2727,0,2932,2090,1420,4045,563,1134,2277,1763,20759,2173,4646,350,3513,2426,366],[12562,15886,7583,38836,24331,26336,16738,18039,4916,58491,26794,9350,20566,18003,17638,45748,11917,27342,20089,20410,23867,33246,22929,25215,17845,50594,22977,11892,37178,50358,5172,22885,26706,20090,43803,48565,12866,38415,12676,25557,13944,14198,22147,13458,11024,20179,18554,15791,15743,14289,50539,15296,18381,10256,14799,10309,13687],[12333,6097,3628,18901,8998,13572,6919,4132,4864,16367,14275,3943,21068,11054,5462,8222,6675,9878,6581,13513,5326,28277,10501,11165,9140,4501,4687,4210,11270,39038,8980,6250,19380,12432,19113,17146,2078,13565,18442,11158,9490,8386,11544,2495,7136,12132,12945,22711,6989,2050,15621,14138,12966,2305,19947,11692,9683],[126595,178263,19234,229043,141475,200431,156965,136791,88741,390340,597463,19230,174620,139585,151160,291500,155417,154093,200023,151310,314374,313491,235472,122935,184427,407118,153454,202784,345061,495726,23034,203910,382528,176098,234739,305038,170370,405666,77401,167807,201493,21891,178482,70292,32957,114214,53685,31004,156054,116448,492781,143194,185609,36190,94136,102116,246812],[99246,220055,44231,115440,77146,55674,41210,55463,129379,258840,234098,31862,94816,84621,87029,172638,34218,29991,109932,37472,129184,120276,95084,114010,126148,85844,27332,28042,66751,36845,20965,58599,24085,175383,167306,114504,93963,77572,47455,55260,28469,4187,85714,55467,71061,174045,39937,33578,62573,40561,513334,42968,47649,4149,93426,82586,121209],[0,1701,3117,1451,1356,667,1472,324,1485,8219,1868,2195,1328,556,2660,683,538,58,1448,2004,802,1492,1247,659,597,788,260,762,2705,6827,25,0,1565,1319,2412,2073,0,1089,736,872,118,235,4719,2022,3563,26,3135,3222,0,198,7977,637,2794,3390,909,603,0],[332,142,377,122,423,79,454,368,704,1742,512,1199,0,0,0,0,0,0,643,0,1571,0,0,0,0,509,64,0,29,0,0,0,395,194,684,0,0,1054,1552,0,0,0,0,151,68,207,263,0,202,0,0,471,0,232,62,0,0],[67633,110463,87838,89095,71275,70218,65188,71250,245362,262321,129070,59779,34234,49425,55453,82891,29284,23359,95335,76618,105069,96150,59304,67063,68156,74553,28019,19778,80536,80704,52801,62946,55766,75629,107541,101697,58005,81320,91506,59688,62563,59053,124744,64143,143728,81887,43457,47194,57828,40595,475662,64856,81261,21412,51146,42293,16177],[256,95,219,0,38,0,96,414,178,122,424,861,724,201,173,415,0,0,579,0,116,0,0,0,54,754,0,0,245,77,0,18,0,450,0,0,0,0,0,0,0,0,378,269,301,0,51,662,93,0,0,0,585,97,0,0,0],[319,202,1061,1554,1213,1024,2950,2857,192,3640,1046,1597,1519,945,764,6020,5789,675,3378,2222,774,3563,839,0,2379,2320,3664,9624,2107,335,0,0,1027,2744,2031,2489,0,2841,197,1705,819,0,1361,358,44,3368,6347,742,798,3428,25,742,1386,2383,2294,1911,1032],[120,258,503,571,425,465,61,174,233,520,335,548,222,97,65,342,46,175,396,101,452,156,565,824,100,187,0,108,71,253,183,270,101,70,304,919,143,282,25,152,192,30,128,357,1157,0,65,282,89,0,456,0,199,278,196,362,186],[240,149,871,227,945,419,227,138,1117,542,1455,444,335,549,472,502,25,35,1491,411,409,674,3454,1814,1150,1126,259,0,1015,151,0,240,673,1529,1747,1036,0,460,3004,13,860,0,292,335,561,54,0,254,328,544,857,404,1019,0,370,133,1109],[629,202,801,1438,477,346,924,733,0,804,1332,1457,2111,464,1144,848,605,0,742,866,1690,739,2587,0,772,1747,1473,0,1845,2550,0,473,527,465,2561,1447,0,3006,3035,1063,0,0,1172,0,0,720,2412,0,755,808,12245,1308,440,606,0,0,0],[449,32,544,170,89,63,106,129,0,495,1895,193,0,32,28,1934,96,275,252,61,32,0,235,402,0,248,0,422,368,250,0,0,442,0,73,0,0,37,0,0,43,0,790,954,0,0,1721,1168,0,231,210,198,182,1137,458,286,489],[0,491,1348,700,30,99,292,186,493,137,1363,1142,972,626,1127,231,0,1,0,1176,0,406,3020,6298,196,839,1324,0,0,976,0,398,0,0,919,903,0,111,1197,0,234,0,0,2035,795,14,3503,604,0,269,2,173,218,605,319,230,0],[26595,26075,9812,33529,26442,36977,17684,32474,29739,48716,46978,10003,33713,31424,32072,35497,13647,14497,25593,21360,35922,53248,53547,98222,16168,38583,14608,14204,42802,47076,7197,15597,14350,36081,57525,59894,25319,43076,48963,20638,32173,12993,12102,11618,34107,27909,16201,25709,16027,14832,110883,24554,36293,6179,15203,16374,21751],[9652,20157,5075,16274,15269,23220,8327,9884,8419,38019,62163,4228,11736,9815,15590,17579,6832,5043,12433,11988,15732,17816,26913,50076,5706,22823,9383,13675,22215,35579,6834,9255,7384,31895,35517,54570,292,19714,9382,5396,6340,0,10084,10170,9863,8692,5908,3465,5643,9984,68159,12018,33955,8869,7251,9208,10602],[460,319,0,424,0,336,286,0,0,874,558,0,362,557,147,3684,114,330,263,48,172,1022,108,0,34,0,0,48,0,531,0,0,0,424,860,263,0,0,0,0,504,4570,46,0,0,0,653,0,0,0,0,189,246,0,283,0,0],[56958,65017,12780,52477,30622,31359,17894,21012,14661,78060,21172,9394,20832,55587,135509,68183,16104,8467,65316,32094,16708,49393,44879,28641,57198,23689,15569,6400,15054,38026,7071,13518,6955,44265,60435,51217,44129,15011,9577,29068,17143,11047,6541,12753,19533,62822,10790,13570,17444,25460,170401,19349,23179,5126,41346,19909,10673],[455,793,2265,968,290,2878,1273,982,675,3531,2176,767,1649,2554,732,2552,1469,869,972,488,1962,1180,2070,4843,0,1457,468,0,2009,802,0,72,428,1992,1920,2845,916,3668,165,750,0,0,1682,171,5263,0,1260,303,1605,0,9392,1313,1335,1155,1107,1013,0],[395,175,90,209,0,282,160,52,0,582,486,0,350,107,309,223,27,239,0,297,7,218,671,1068,0,350,0,0,53,79,0,0,116,497,831,631,8,73,126,242,243,80,461,24,289,57,0,137,0,90,1052,158,125,606,474,98,178],[761,311,3190,1228,603,1156,568,537,674,2719,2285,2623,1209,1334,538,26285,15891,5202,458,374,1084,2174,1309,1562,241,755,219,1972,865,1562,5704,305,548,782,3483,3914,737,903,1120,648,485,3473,3861,1456,2215,834,999,2216,1255,118,6264,1173,1211,2059,1225,1008,1114],[0,0,50,30,120,193,202,336,0,220,463,884,0,0,366,17,0,18,0,197,99,90,0,0,65,447,0,16,778,0,0,0,0,0,303,430,0,0,0,0,0,0,356,444,135,0,2243,380,0,0,112,202,202,124,0,0,0],[2891,3203,631,53372,14873,48679,9690,4491,2030,69559,36869,3202,2087,4948,1280,58396,14500,33169,26473,16457,41378,35097,12072,12954,5956,7301,3909,39082,20890,23512,480,1617,487,16749,67290,87930,4743,804,1922,3285,7644,3116,1342,2907,779,4567,1655,1261,713,1111,3362,1033,1160,287,2954,2993,5704],[2479,2120,4994,5635,3302,3397,477,1625,1092,8312,3542,4676,1706,1190,657,2971,1460,1109,1135,415,4690,998,750,1823,832,513,669,379,460,831,35,354,426,1886,2331,1933,653,1397,1213,366,2675,1373,673,859,3754,1911,7368,623,1240,0,1180,152,375,302,3357,335,2976],[2605,11770,5169,17175,12194,7553,4903,6574,11887,27087,17294,4277,4910,10468,9719,16255,7306,7317,12630,13974,20899,14578,8998,22413,14322,18921,10491,1249,6837,9202,9329,9932,4458,9801,21787,23522,1185,24261,5915,15847,17082,1223,25829,15130,12417,5670,8857,4891,16907,8822,21335,9091,12781,3823,3655,4619,4342],[0,1,0,0,0,109,0,128,120,10,74,554,0,176,6,0,0,0,0,0,0,0,0,0,0,345,202,0,0,157,0,0,0,68,187,246,0,0,194,0,0,6,24,7,30,0,232,0,0,0,0,130,154,720,0,0,0],[43,0,203,382,0,35,0,100,0,1129,163,248,7,525,0,70,7,0,0,0,0,0,94,0,92,295,258,0,24,143,0,0,70,597,782,129,0,0,0,0,0,1881,0,127,1716,0,178,927,49,0,0,118,227,516,600,1,0],[7447,8144,2500,11614,8361,9745,11566,15027,14353,69045,31902,3568,8861,9497,10338,24098,13246,16205,18886,12515,20679,29901,16551,4958,30544,91894,19673,25660,57440,40238,10319,87010,42086,41410,29208,14383,5821,36683,18377,15872,31724,405,233161,91782,4811,12987,12241,5359,40717,22267,50524,31232,21465,5687,5142,5234,10010],[27995,26780,7025,59375,47532,40156,38048,47668,44699,74950,79409,9847,37739,41442,39971,124398,44006,33768,57510,76058,58463,76601,57036,63991,46799,48718,14193,50258,41839,49063,6550,11003,32430,62298,67603,59787,33512,50009,19382,27853,13349,1885,12648,22101,11599,60657,14224,13901,38126,27861,116564,35711,32832,5725,31890,33765,53649],[0,124,978,0,0,0,28,36,136,265,372,567,86,86,128,0,0,0,0,8,194,8,0,0,0,164,780,0,186,244,0,0,196,415,589,645,0,0,0,0,0,379,244,0,1052,119,796,317,0,77,0,0,350,879,56,0,0],[3169,3429,13279,7562,5311,5251,5113,7432,5140,11767,10666,10974,9719,7494,4274,55657,37334,15638,20365,5215,9734,11953,4945,4993,18279,10361,10218,15874,8485,6961,5883,3877,8453,3914,11525,15524,3889,5588,17982,8476,3275,11612,12627,6546,14420,11595,25451,11701,5471,3216,31641,7661,11007,15628,6132,3002,6483],[1247,2074,1536,1046,2032,4305,2683,2288,933,2544,2660,1071,2807,1785,2007,2091,767,1444,1738,640,2011,4504,3517,4180,1872,2503,174,767,3147,2758,1178,561,402,2498,7413,8650,0,1492,8433,1186,749,0,683,1612,1348,1801,2189,2440,991,2415,11769,1909,2216,682,2095,1859,4],[64485,83944,17802,96380,84317,118107,47451,49470,71917,173479,130842,15921,44015,54655,61467,32469,11850,20300,61438,99608,152987,104827,71040,99249,61326,127384,59882,10666,112020,128706,12730,39803,50218,69795,95487,119062,32173,121123,71465,41380,78249,25787,129412,84699,58052,66655,47309,57241,55684,59750,631814,46101,63769,8636,26121,29914,29979],[0,76,35,412,0,184,299,236,41,0,395,247,255,114,75,1866,312,637,477,468,128,0,0,38,0,131,0,38,86,0,0,0,0,182,329,1259,0,1367,400,0,0,25,0,31,0,81,222,236,0,0,0,0,257,409,0,194,43],[4520,2096,2122,3848,3520,4538,3027,2883,4754,5641,11480,1160,4477,3984,5542,14927,5570,5027,2221,2963,4576,4843,19886,36321,3455,6572,1204,6930,6454,6570,707,3384,8525,9718,12028,13336,776,13877,22181,2704,6146,8963,4967,2622,3123,5951,3256,2980,10603,6917,64980,4015,7032,631,7331,6923,8865],[706,0,3969,1141,838,1066,191,499,428,914,859,1608,1111,545,837,489,255,264,25,203,564,1078,0,1807,792,1006,525,0,787,6133,31,40,112,1804,3918,3924,0,51,2139,0,0,0,1169,1259,2947,752,2616,4245,606,0,703,1094,1587,1323,380,636,0],[0,0,0,0,0,0,0,0,0,0,0,0,0,0,0,0,0,0,0,0,0,0,0,0,0,16,0,0,0,0,0,0,0,0,0,0,0,0,0,0,0,10,0,0,0,0,0,0,0,0,0,0,0,0,0,0,0],[12978,11422,8193,22924,18738,24193,8732,9503,28060,51804,42758,15898,13656,14266,14750,22182,4989,9091,16849,14789,23417,15703,21401,43298,19064,22212,11975,14713,13543,25457,3297,12069,13721,19815,33870,35114,9269,29789,53244,9404,13467,5344,14965,9036,18676,18278,12688,14867,7009,8729,138092,10065,20050,5395,16286,18779,19301],[0,434,2579,73,184,304,401,1275,781,480,806,1638,55,637,490,2410,1225,576,198,0,0,1796,669,0,956,604,1194,0,2077,0,0,0,808,170,3201,3331,8230,1910,3869,379,1545,0,358,817,1338,552,401,0,402,603,8377,1699,1036,1317,199,992,0],[31293,39471,4019,62103,48824,48518,44303,53153,10514,146210,138550,7247,63382,51818,49260,99749,53000,46676,76568,41048,108985,107031,48739,40878,91873,86299,28685,17073,73495,72235,4211,46469,63092,64193,105534,63194,48785,97467,13660,66765,57589,7420,53817,21012,12838,58733,22438,12036,62777,33765,78771,42371,47448,5569,23139,29053,59545],[112,198,874,0,0,63,0,212,0,96,243,199,550,0,0,39,0,129,620,48,0,703,1950,848,0,0,0,1493,117,1437,0,0,0,480,253,2048,0,0,0,31,101,0,686,0,1232,0,1001,1187,31,0,0,169,520,202,0,0,0],[1673,28,127,116,192,811,0,79,186,251,559,590,0,190,199,27,0,0,144,0,0,0,0,85,0,213,456,0,202,224,0,0,0,42,121,606,0,111,0,101,0,0,106,536,2501,0,639,920,0,35,0,202,0,335,0,81,0],[1414,6078,10235,14323,16848,18058,8902,9633,8130,39721,21457,4425,20158,9140,9077,18356,7277,8858,19607,9208,13601,33908,15635,4845,26210,13374,10396,3031,33826,37199,2531,10555,15190,26230,17429,24854,9072,27351,18674,15915,14014,1999,12887,11980,9861,18134,9614,5457,13337,5093,29033,25992,24410,9879,5668,4137,819],[40,192,290,987,978,296,22,0,150,159,688,185,111,0,157,0,44,0,2132,15,248,0,53,0,300,0,208,30,78,23,20,0,100,1081,310,102,0,229,0,40,0,60,806,193,176,0,1114,90,489,0,1212,20,377,0,0,44,65],[2,229,280,415,48,11,230,138,53,491,135,0,130,31,74,385,0,0,15,168,66,168,0,0,327,12,0,0,202,96,0,42,0,0,128,0,0,0,0,0,0,30,0,0,1034,6,1496,159,0,168,14,0,0,0,12,5,0],[402,0,1126,0,0,0,0,402,202,222,202,178,77,165,202,0,0,0,0,0,202,0,0,288,0,0,736,0,321,590,0,0,0,0,350,392,0,0,727,100,0,993,402,909,1454,399,234,1205,0,0,9086,1391,606,1932,258,123,0],[8,247,1241,409,10,502,202,270,450,333,205,839,21,6,1222,53,17,439,0,32,202,65,356,1294,444,432,204,25,430,227,90,212,222,296,18,16,0,605,201,339,590,0,487,342,112,102,1174,1715,217,0,192,581,213,430,843,9,0],[2004,1482,2384,2194,1203,1625,1562,877,723,1602,1578,3183,949,1069,1670,560,0,2957,286,267,845,85,0,1680,1406,2034,4531,2167,1979,2596,2661,80,986,1375,2860,5959,0,4554,3540,202,180,0,1701,1372,2529,2042,3558,518,1932,260,230,2388,2920,2992,1396,827,0],[819,635,6064,2040,1273,2480,2756,1797,1242,6328,3714,2621,1946,763,2569,9048,5801,2208,560,1026,2656,2832,4649,3747,1432,2217,2380,980,5513,2620,78,27,435,742,2838,800,1712,5906,4509,56,2344,619,1422,1752,4465,893,9585,6562,1157,34,5444,2217,2686,2947,643,941,516],[1804,1395,7353,2162,1910,962,899,2192,687,3802,3351,4485,2014,2643,1258,2338,1097,1239,1268,683,3096,1746,760,8981,665,2742,5168,23,1589,1687,0,996,1180,1706,6537,4020,0,1554,6230,118,213,3622,3037,3768,10994,2520,16179,9433,2407,201,1815,2576,2041,4011,1342,1608,6608],[29,217,963,684,359,942,523,650,1120,2349,759,2049,869,703,855,1659,235,1029,0,67,360,181,1922,364,186,1039,405,795,1211,847,0,35,66,1117,1126,1536,146,58,223,0,425,33,1260,1844,1507,436,2240,3280,0,225,1165,334,195,563,1305,172,26],[695,231,2124,1431,1084,1268,412,977,404,2028,667,2872,598,965,313,1328,45,0,2135,992,599,4384,879,2091,1176,856,2466,0,1259,1088,0,350,200,839,726,656,96,46,0,130,0,0,2050,622,4780,1246,3046,3527,822,532,2890,663,1650,2321,906,475,6423],[1568,2492,4185,1889,2242,2220,794,2303,1980,4846,3570,3494,3522,2100,4206,3537,2886,594,1205,2094,1688,6590,1051,1185,3141,1558,2881,1478,5255,4912,123,425,1532,478,1428,3633,537,2087,1616,1560,1878,892,2635,2085,4439,5731,6442,5291,2680,1305,8479,1490,2286,3644,1445,2858,300],[6950,8706,1410,18059,12327,11756,6551,6382,8360,26168,20907,4700,12762,12692,11478,20795,9620,6408,10583,10017,17295,12308,9748,27346,8110,15612,7246,5608,13736,16279,5278,3334,9307,10784,14643,21584,12631,25152,13986,8528,7853,3845,6464,5809,7456,16562,15216,4247,5729,4689,55118,6944,13134,2671,8837,5303,12341],[9799,8310,2871,22446,12206,12537,10551,10494,8898,28191,21766,2774,12355,12881,10681,11881,10851,8724,11878,9655,14365,20021,17550,19358,15158,26918,5943,2601,14049,18671,516,5281,8241,10964,27940,29275,2937,21653,22024,10952,23899,2434,8220,8011,6771,11208,7479,4695,17907,10670,58437,11011,17313,3734,6174,3531,3713],[55552,3250,2457,64423,2794,234978,27171,6213,301,39877,6110,11645,12273,1271,2742,16015,2146,15889,2188,12758,56540,2282,3701,5627,109216,1597,14771,117,8183,6778,0,204,199,4031,4796,5932,999,28607,328146,158485,11408,15172,4112,1609,3458,22503,4338,101236,919,2218,101884,154900,9267,4867,120519,6146,3202],[0,1159,598,0,0,0,76,476,0,0,0,404,0,0,0,33,0,0,0,0,14,197,0,0,176,472,1309,0,0,368,0,0,0,0,630,0,0,0,0,0,0,0,0,514,400,549,941,0,0,0,260,0,247,603,361,236,0],[0,0,1232,82,0,320,202,0,404,26,871,2343,171,202,503,0,0,0,150,594,0,0,0,807,0,530,1707,0,1262,71,0,0,107,0,525,0,0,1405,1590,0,0,1105,924,2131,0,1813,2177,1278,0,0,5866,335,981,1384,180,638,0],[0,2,58,0,0,0,0,0,0,0,272,3,204,0,70,177,33,0,198,78,105,0,0,0,0,12,0,59,0,0,0,0,0,0,0,284,0,56,0,0,0,0,373,0,202,30,13,0,0,0,4,61,0,383,0,0,0],[858,146,659,492,99,223,494,30,130,584,621,206,672,363,533,284,129,396,202,189,333,171,528,325,535,806,199,0,0,702,0,233,0,0,2677,825,0,833,0,115,0,0,86,317,611,0,0,398,0,0,581,59,0,393,427,0,0],[1743,4132,2750,6011,3504,7277,2935,3367,5348,11463,10075,2848,4585,6145,5397,4007,2447,4707,2977,4251,7850,7490,3071,5521,4156,7519,2781,4009,6905,13757,540,5737,4598,5333,5572,7200,8122,9980,4099,2266,6106,513,3783,2932,5086,1493,4706,2613,2920,2568,20926,4577,4850,4011,1879,824,5609],[3201,1110,2650,6741,3885,7992,3496,3746,4555,12143,7122,2582,3877,4505,4545,19285,7385,11362,1794,1470,8682,4840,8044,4579,1253,7973,2239,3692,6836,5183,4188,3009,63,1724,7596,12326,3029,1416,98,2344,2265,3463,17133,4528,4617,1210,4066,3504,8271,3274,14127,758,3377,2632,2761,4897,223],[30846,29389,8568,34604,31274,40764,37898,33356,20640,67392,96937,8418,50225,42928,49798,46186,28495,68917,44445,51188,90937,80472,47848,25038,29150,70508,36525,20807,86663,91191,4958,44814,26871,43564,88643,83885,14978,139058,31294,31847,67201,6149,44961,21802,13427,26852,28706,17674,26077,16732,139912,32872,39018,7183,10991,13371,16459],[202,0,735,495,144,24,478,808,612,531,711,931,125,0,190,894,54,168,1074,793,387,2230,2394,2162,627,1494,1157,4213,177,170,0,200,200,585,739,2358,0,0,0,975,490,0,4717,2071,184,3038,1192,200,892,401,1553,375,1540,200,1024,234,0],[3110,3003,2018,3967,1996,4067,2811,3195,5702,10647,28489,2040,6081,9776,11561,7753,2477,1328,5292,6594,4382,7676,6822,5926,7458,7520,3222,1594,5412,8938,2124,3928,2120,8288,6908,13049,0,8550,2249,3522,3934,0,5506,6406,3991,3445,3075,3340,10311,3280,6624,6996,5347,2205,2453,2805,3645],[0,0,497,124,0,151,370,136,278,13,190,415,94,0,58,0,0,43,0,172,0,0,0,0,50,293,0,0,106,542,1748,0,0,0,332,354,0,0,302,0,0,377,74,370,2691,292,1323,2506,317,0,5938,0,430,1445,383,0,0],[1607,838,987,2155,1035,1314,2662,2316,2315,2605,4780,1104,2242,1948,2536,4725,219,790,3059,1999,1886,2395,9066,7046,3733,5686,2650,329,2559,1659,26,105,247,1581,3106,4870,5076,2098,3723,504,910,2226,1474,1963,260,1118,779,2801,558,1550,672,2454,3914,2969,1105,394,776],[172523,282963,31677,279915,212655,218947,204603,206773,182323,628908,368580,28136,205155,222356,201482,298539,112576,139147,303795,300689,404296,409561,233291,165315,276920,314287,151102,138631,294305,445003,74899,208504,154048,310526,305207,361250,145902,546331,151701,221239,193525,44535,394196,222987,112030,277216,119306,91286,226855,132675,919055,196721,241671,28899,88504,96722,92887],[30695,33215,4327,45389,34082,33788,44641,46344,22858,107915,121192,6863,40999,38170,42081,90570,24814,30483,69249,42351,54835,86614,29044,16922,54624,71919,27228,5049,63885,124975,6078,26312,32089,62822,84303,80883,45305,84825,13426,32258,49069,11792,40364,25218,11765,37566,17076,11098,47952,34798,173893,37659,54745,8240,12460,10917,12728],[1807,3867,1730,2581,2198,1692,1315,1421,1045,12487,2313,1639,1660,1182,1041,3736,2632,193,1155,255,1217,2463,2068,2473,1159,301,0,1639,305,872,0,1193,608,1452,1815,3521,5737,1050,3401,2817,908,474,778,1265,906,1393,98,1322,149,1072,2365,1493,893,379,1330,2032,833],[1774,1433,341,3858,3774,2675,2589,3508,1780,4684,5293,1737,2733,2827,1566,4341,810,1505,3840,3890,2124,2843,8181,2353,2926,4076,1564,446,7314,7936,0,651,1755,5531,8082,4515,0,6149,603,4235,4609,0,3488,3438,1400,2649,1594,1968,2911,2011,5349,4327,2295,261,725,1422,0],[13337,20776,10674,19939,16619,18275,11834,16160,7947,32403,31361,8679,18560,12232,16150,50230,25775,14195,18676,11576,25261,27867,8800,19451,10807,31509,14722,5845,17410,20359,9202,8681,10519,16446,23032,22822,8154,31817,18804,4882,10225,10584,17303,11186,13008,21488,17185,15629,19712,6466,63681,10989,16098,11476,6586,8808,31173],[6175,18492,5037,14124,11413,10978,4580,13457,7884,34259,15770,4582,10780,6434,8114,11767,2920,7895,16943,8464,14413,24379,9291,15330,9255,13057,6275,1661,9325,8413,86,6697,7032,18086,17249,15474,0,12339,4836,9119,11499,2779,13861,6241,3322,7547,1514,4023,7852,3092,25942,11858,11035,1918,4665,4597,10346],[0,0,6,0,0,0,0,0,40,0,198,0,399,40,80,0,4,0,0,0,0,60,0,6,40,0,532,0,39,0,0,0,0,0,7,1200,0,0,0,0,0,0,280,0,0,0,0,0,0,0,0,25,0,0,0,0,0],[0,13,113,0,0,0,175,0,0,146,202,492,122,0,0,0,0,144,0,0,0,0,0,0,193,0,387,0,0,99,0,0,0,35,0,0,0,184,0,0,0,283,554,71,734,0,342,0,55,0,0,0,131,92,1708,32,0],[37938,60751,70966,52799,43592,70141,32132,32578,103813,148200,72446,60047,36222,61880,52815,46349,21504,27716,34732,53301,43701,45770,52298,71890,33362,84566,54649,19538,57613,94423,24674,44057,32001,44085,64072,74449,15529,61377,152836,29913,19846,44033,92018,37276,121464,70450,46234,76121,23353,21179,544760,34866,35756,28575,38809,30921,37948],[0,62,773,174,0,171,0,271,562,299,773,1110,150,58,115,27,17,29,0,60,40,0,44,0,0,288,241,207,32,423,6,24,0,0,276,1549,0,0,758,0,37,0,0,1649,1044,0,290,101,0,294,53,565,679,253,170,112,189],[112159,108837,19442,155983,137307,193357,165134,169742,160679,361152,363055,30879,158290,195673,152331,246802,91765,242492,144594,139819,340805,248073,221800,140294,153423,291047,143946,107614,270442,293822,102638,177724,164625,186183,386954,418382,155278,575019,320217,154785,244041,50750,157199,96839,51318,125477,122964,157878,123452,74693,766811,225211,210796,31464,137287,93379,56662],[14740,17620,7152,33471,17578,25168,16791,16592,17074,47560,64364,6871,19274,21786,21777,18643,9501,27962,26842,25194,49094,35153,26867,24262,21650,46630,16106,6585,49603,57754,11311,12340,31107,24089,54936,44933,6400,38320,14152,12875,38178,1169,26420,20901,20012,19313,6614,10401,28303,22454,200288,12994,34903,7624,7544,10994,20755],[1490,1037,2260,2516,2386,1894,900,724,1460,3930,4557,641,2385,2287,2065,2415,1560,1016,546,393,611,3151,3107,1958,1400,1363,1079,0,5204,7618,3855,1375,1367,1363,3013,4217,4521,4394,2532,1785,2553,0,5354,1152,4447,1743,5420,3003,1711,193,8409,2805,4381,4078,1381,1346,0],[0,0,99,64,0,0,0,0,0,0,24,0,0,0,0,28,9,31,0,0,72,66,0,0,0,0,0,0,0,0,0,0,143,0,6,0,0,0,0,0,0,0,0,0,0,0,0,115,59,0,0,0,0,28,0,0,0],[0,0,56,160,0,0,16,0,136,0,180,0,28,186,0,44,0,0,40,0,0,0,76,0,0,171,0,0,160,0,0,0,0,0,0,429,0,0,0,0,0,0,42,0,0,96,280,0,172,0,0,0,288,0,34,0,0],[556,951,126,929,1877,1300,1354,2474,1206,3996,2412,173,1518,1967,2755,1991,31,623,676,698,3355,1722,1501,2865,1294,876,198,188,1470,1669,0,984,346,2987,2831,1885,0,4607,795,1168,1786,0,1577,479,610,1859,332,1065,950,1299,0,649,1513,0,650,2170,683],[0,0,1689,102,0,0,0,0,565,0,8,1560,364,0,796,160,0,0,24,0,293,0,0,0,598,200,202,0,196,461,0,0,0,0,1087,596,0,0,0,0,2302,3585,0,807,3922,331,195,1281,0,0,25,0,400,796,458,0,0],[0,0,89,0,48,426,0,18,0,0,0,66,0,81,32,0,0,0,0,0,0,0,0,0,0,0,64,0,7,0,0,0,50,0,0,154,0,0,381,0,0,0,43,5,0,0,0,150,0,0,0,0,86,94,150,0,0],[0,0,123,0,100,1532,0,104,0,200,0,435,197,0,142,0,0,240,0,0,120,79,0,111,0,0,960,36,360,0,0,0,0,119,66,464,867,0,360,0,0,0,0,16,0,0,0,240,0,0,0,120,58,0,0,98,720],[1664,2766,1351,2134,1852,2328,937,1000,583,6394,2798,1121,1192,911,1846,1459,1211,1232,767,949,2004,1111,4502,3088,824,950,337,600,1485,1265,710,811,2134,2247,3338,4260,1485,2227,8641,406,1886,1235,1315,359,1140,785,1332,685,923,250,4516,377,1151,330,1519,882,3268],[0,647,356,0,458,373,0,110,199,232,1312,569,303,542,0,199,49,0,0,162,121,0,0,0,0,425,1590,33,1151,133,656,0,0,32,1364,477,0,160,306,0,0,137,16,540,5574,0,796,603,1066,0,97,1658,881,1723,0,671,0],[28631,80423,3589,78830,56343,36566,52827,57501,42965,173516,247980,4681,53418,38201,35677,74044,34045,35374,70600,50377,280073,91156,24800,22925,67192,99744,35837,8128,73264,80560,6180,36534,39049,51706,66857,101235,39849,101188,23287,79597,67213,3932,51781,38327,17128,56804,34400,24807,81813,47637,493560,45720,65522,7604,7980,19965,29010],[4511,5181,544,9202,6439,8675,5681,5313,5547,18486,12867,786,6234,5769,4794,14340,5639,13735,8100,7796,12628,12443,7578,4372,6073,9996,4992,1978,10512,8892,8428,10179,7140,9073,14661,9996,4680,16310,4344,3562,8845,206,9884,7374,1953,9165,1979,6286,4258,592,24686,6888,7531,1197,6845,5708,12332],[2924,2221,1284,1540,2275,544,401,908,0,2635,2456,1094,1293,479,696,957,90,0,1285,943,2818,785,1020,276,1664,1634,0,79,0,333,1662,981,0,973,22,505,0,1186,0,663,0,1222,1692,552,1857,1272,1166,2170,582,0,1326,0,243,99,1015,847,1181],[107031,202657,20257,221967,155267,174989,173025,158924,179140,505546,469361,32928,158581,162207,159023,163418,75280,81718,243348,203310,305980,255776,113516,98396,221355,345227,178130,45704,233138,299056,75169,140643,168023,151909,258279,294252,133151,365379,192238,179941,143095,54319,205814,128677,78697,192446,99216,116242,133512,91992,733133,131816,176601,28243,50984,44317,74450],[0,0,87,26,0,0,0,0,160,0,0,0,401,0,0,0,0,0,0,0,0,0,0,0,0,78,0,0,0,76,1308,0,0,0,0,0,0,0,861,0,0,1406,206,348,114,502,511,0,0,0,1919,71,0,568,0,0,0],[789,1772,1307,1372,635,1731,621,632,532,3466,2837,962,775,1130,938,1180,901,306,1100,285,2316,363,920,1305,762,418,1370,173,409,1491,0,470,154,407,1003,900,163,1123,238,391,1228,29,1905,723,2726,1344,4656,773,0,128,3995,1495,701,621,876,795,342],[11108,10685,8237,14303,22406,22258,13415,14837,6335,11852,30617,5259,36083,22433,34291,24970,4666,12846,20140,9308,14268,83289,82084,28333,6700,32078,17378,8714,59458,78111,4355,4953,37654,22137,77548,121283,8148,47869,7415,9137,25943,3144,22328,13308,10168,10693,19508,9312,15869,16879,110957,14191,62579,11839,4790,14265,18945],[0,40,80,123,80,120,0,144,80,1370,160,93,0,93,0,903,0,80,0,0,40,429,0,80,144,246,0,0,126,0,0,0,18,0,163,656,283,0,0,0,0,0,127,0,0,0,264,0,40,0,981,102,0,107,138,0,676],[0,0,491,0,0,152,0,0,201,0,154,605,0,0,0,0,0,0,0,145,0,0,0,0,0,254,0,29,510,0,0,0,0,678,0,0,0,0,0,0,0,0,196,99,600,0,856,0,0,0,2203,198,489,431,84,0,0],[2553,497,2176,864,127,4729,752,758,9,2310,732,1416,266,294,0,1449,577,1146,0,515,919,57,0,80,862,23,116,2039,0,151,47,0,0,45,55,796,231,372,31394,3278,299,7885,68,287,3475,451,46,10926,92,24,12470,1161,56,204,6189,1332,877],[4776,7373,22865,8951,8713,10509,911,1540,2560,15600,7003,8787,3312,1056,3217,5961,1322,2253,834,966,6256,863,2427,1707,611,1718,654,2643,2289,3762,802,564,560,895,1718,2481,1458,4008,4206,886,5121,10167,1751,3215,9281,1305,2266,2107,297,339,6725,590,1108,1231,5065,1608,7813],[79,204,348,1117,993,1834,1117,472,54,2596,1548,857,264,155,944,483,314,854,354,625,2058,492,570,462,802,674,290,367,1688,673,826,175,395,241,509,206,606,24,1168,432,132,0,569,909,2626,1270,404,3045,132,368,177,174,494,367,242,0,1852],[612,481,907,176,1567,835,402,942,858,1238,1835,1976,843,1208,481,506,0,94,157,302,1166,483,363,2757,118,1433,798,0,1616,394,920,288,217,0,1296,2134,0,926,170,320,480,0,1606,709,1726,893,3074,2066,1537,200,4263,1486,1445,1244,248,176,1402],[121156,224086,12452,319304,219270,250539,201066,202158,98288,650279,434961,13560,269039,179197,182050,520254,333204,241014,361692,209345,382828,510021,226998,96175,367295,504258,186736,345314,388914,495454,71678,274056,172407,327197,387519,359744,277683,432811,69359,268289,163651,11278,264590,111649,23919,287922,64191,38520,247024,197370,969025,181949,202004,18239,201147,240541,406387],[0,0,229,0,98,111,0,0,95,0,0,63,0,107,0,0,0,0,0,0,0,162,0,198,0,588,0,0,199,170,0,0,0,0,0,964,0,0,0,0,0,0,631,0,384,0,176,0,0,0,0,521,0,624,0,0,0],[196,0,928,124,359,20,0,357,1965,390,1101,1678,98,537,0,547,454,374,10,25,250,16,5,0,0,450,0,15,0,132,5,0,201,15,1147,704,1,1,0,0,0,2,771,426,555,202,46,667,0,7,99,523,0,202,0,916,203],[0,767,294,92,34,0,618,202,596,390,657,214,0,610,435,330,242,0,519,21,713,606,0,0,128,375,0,0,0,202,0,397,0,0,404,1722,0,0,63,0,0,0,0,306,314,10,0,303,0,20,0,0,0,200,202,0,0],[0,130,489,0,0,0,0,0,0,0,0,161,153,0,0,0,0,0,0,0,0,0,0,0,0,0,395,0,0,136,0,0,0,0,428,0,0,0,150,65,0,0,0,87,457,0,0,594,0,0,0,0,0,189,0,0,0],[0,0,199,0,0,74,0,0,0,1106,135,305,0,232,0,0,0,0,0,0,0,0,0,0,3,0,3,0,199,271,0,0,0,0,0,150,0,100,0,0,0,0,398,99,1604,0,0,0,0,0,0,0,0,0,0,0,0],[346,31,796,45,0,898,134,608,0,26,42,212,119,71,28,251,26,0,0,0,0,0,0,73,0,370,0,0,426,141,1123,0,0,90,0,0,0,0,82,0,0,825,0,87,1219,543,526,989,0,0,0,126,225,1086,35,0,0],[0,2979,1483,179,556,1308,1240,1981,857,3615,1793,981,592,1947,3756,114,118,1904,2112,1572,937,1534,0,652,1278,3984,723,0,1775,9772,0,180,3995,809,105,4849,0,1969,0,1030,1472,2048,6552,1711,1319,2457,1423,2682,378,425,40436,310,1976,444,503,872,1740],[7586,5258,2191,8721,8771,8121,6073,7671,5911,20438,15907,2256,11430,8802,9206,24492,8349,8889,9195,8984,13800,12335,7105,8198,7877,13430,5152,12888,12794,12079,4342,12675,8586,7109,16367,17312,5847,30247,7069,7550,8654,2591,14469,4709,3017,6760,5194,5813,7627,4764,25487,5250,9932,1455,8546,10224,16091],[173,97,1945,331,373,409,203,75,798,1191,656,2141,440,1261,599,786,430,88,18,133,319,165,595,2424,0,1063,2725,224,2141,714,208,96,95,280,1800,1337,314,535,1408,223,46,4504,1875,1070,2334,323,0,3216,146,0,4882,681,807,738,1366,0,488],[120,0,1331,495,196,21,450,135,0,166,506,778,217,548,0,1883,545,302,75,0,12,282,376,0,309,1071,623,0,237,365,0,66,0,551,553,846,0,492,28,0,0,0,1435,583,1818,319,2698,460,79,0,7642,402,1434,1182,96,137,0],[820,1177,390,4450,2052,2802,1762,4063,86,1996,4528,518,6664,2832,1861,2492,1725,7158,1392,824,2076,2834,6390,2878,578,4476,5347,1573,4351,7165,48,776,1156,4318,7540,7734,256,7525,3697,154,2174,40,1658,844,2734,1410,496,4076,1277,1035,2258,3371,3076,826,2286,904,211],[0,0,386,109,62,0,0,145,101,352,257,196,101,379,294,30,0,34,455,0,0,629,300,0,466,0,101,0,327,102,0,101,0,0,218,197,97,0,0,277,425,0,239,99,431,0,0,101,0,0,0,83,262,463,0,75,0],[0,803,2722,278,0,134,0,0,613,1713,892,646,765,402,91,272,54,187,322,202,306,97,0,87,296,340,647,5,955,1010,10,0,462,800,1501,298,16,5,7,1123,0,692,1741,550,2006,889,1741,4504,158,0,1910,414,1531,2449,166,0,267],[7685,6638,2447,2769,1851,12436,10830,9510,19013,33226,5815,7403,2598,2613,2719,7135,3873,21093,3625,2018,1483,14512,2858,14734,2012,20376,13480,12200,30974,39258,24090,36935,42682,13924,11543,13162,0,13298,2619,471,3680,3704,32213,13204,6640,1206,3934,23295,10109,1083,8174,16294,6079,8199,8837,1052,1996],[1939,2085,6933,7890,2613,10381,1997,2081,2032,21939,10421,6369,1969,8563,7943,31105,9232,6754,8922,1065,6483,5101,1833,2650,16882,12374,14359,2889,4550,18142,389,36,6232,2388,7320,4830,359,2648,34919,9452,4726,4697,2508,1790,9856,21478,8157,9937,2198,924,14125,3547,1444,4121,10142,2047,91],[601,83,325,0,0,0,164,176,325,0,97,1173,0,98,191,0,0,0,0,0,38,0,0,0,0,0,0,0,30,70,259,0,0,0,385,0,0,0,0,0,91,0,800,247,575,60,473,293,0,0,0,0,199,0,0,0,0],[74145,65801,19883,103014,56817,104263,57026,73156,19849,127287,151827,15751,88698,57220,71003,77596,25922,70658,59603,69498,109419,151748,86433,58147,71132,130263,75638,23020,169155,127266,7850,48461,45039,75363,133948,140846,24680,160157,77324,57675,96896,8097,80917,36625,28815,33280,39619,55124,60454,44283,165921,102171,101102,20366,38742,33936,36991],[0,1114,2637,501,0,201,0,393,144,1357,663,1137,318,0,0,23,1160,0,0,605,114,971,30,0,1010,908,1009,2319,461,222,0,168,0,292,808,288,0,77,97,0,1200,3967,1151,1515,2749,803,426,938,0,198,407,707,309,1275,433,54,0],[380,0,2301,0,150,106,176,520,202,519,447,1022,476,37,0,80,0,0,228,31,50,320,94,852,110,2974,1696,128,1171,398,0,111,805,20,1069,0,0,168,408,213,0,731,769,1097,4209,1207,1104,2301,0,0,95,596,1094,2282,115,326,138],[236,349,2085,986,3379,4140,990,2706,642,2652,3590,2220,1023,402,500,1580,219,800,131,922,3167,272,0,305,1413,1835,3870,43,1045,1450,385,1973,1338,4,2092,661,0,341,819,282,69,100,3000,2173,3507,489,778,3006,507,0,11626,2671,361,3544,1093,42,960],[137,160,0,631,366,544,98,0,312,1326,391,643,77,65,229,1077,20,186,78,110,721,660,148,282,485,407,545,61,368,257,0,498,66,387,498,434,0,109,576,450,294,45,205,315,258,593,869,677,0,369,2110,270,662,77,83,53,353],[61554,166233,21294,156081,95164,74061,65390,107240,57296,257103,178541,13911,81124,73475,86201,210729,86368,62032,199122,107675,210926,241672,235821,163373,120189,199950,44808,90050,100491,205404,35011,81397,105526,205185,141242,186308,86589,184029,69607,98185,66822,27033,78655,48905,37598,137373,26060,27862,76707,90657,507828,78799,71191,16692,95580,61569,197114],[147864,278251,88560,555758,420454,242937,17700,44885,144125,575754,357829,92122,121890,92627,117107,70938,31757,29136,68254,129521,405647,12390,100294,137116,168426,210969,34037,23850,100592,40426,19940,61785,108032,66460,204800,66792,65048,107241,80589,48472,92046,67098,50720,44425,118214,90864,47049,44621,44013,83519,1012025,36805,52951,14747,35263,70750,146926],[1613,920,3967,5888,4145,4346,11426,7815,1906,21853,3198,2225,2695,2120,1135,8280,2736,3125,14779,5672,2167,30918,3882,4971,15913,3691,4227,4203,5553,4492,76,10359,1711,14159,4716,5939,4971,2915,5286,10644,2734,10603,16354,5452,3253,7161,6960,4540,3397,2016,16016,13329,3819,1354,4669,2267,545],[0,384,0,0,600,346,48,288,173,1188,1298,0,0,1094,1409,2229,102,324,418,444,772,998,608,1480,957,906,0,0,1138,189,661,1405,176,1901,1144,0,0,3523,0,508,47,0,1517,1279,1368,22,437,96,177,1009,584,654,230,115,527,424,0],[55,146,306,348,82,209,230,169,0,2336,303,0,189,245,157,1972,0,141,76,55,71,344,0,0,179,0,792,46,317,203,0,56,124,0,426,0,0,0,0,118,1384,0,512,313,241,0,410,0,0,0,935,32,273,261,212,4157,7324],[1188,1238,0,285,467,356,1609,2596,215,3004,4245,476,507,383,885,3224,407,1042,2709,2326,2755,5435,1335,1179,2330,552,667,390,2060,4208,0,3607,232,1780,7129,5538,46,3542,2423,2037,396,0,948,679,0,2324,0,285,474,302,1966,3079,2534,0,1312,491,0],[28947,36974,7809,46259,40082,44709,50232,44190,31678,124753,123336,5577,43749,50300,50242,70821,21918,28734,69094,55566,80718,98195,37063,23869,57537,91468,31767,13260,74145,64335,16977,49466,39838,68881,98851,59383,49216,112228,19360,54103,39171,1876,80546,32893,8208,42105,14541,19640,36444,21287,112015,45640,52478,8539,13015,15929,15214],[128,0,309,0,50,27,0,0,0,0,0,0,116,123,44,4,0,0,0,0,0,0,0,260,16,0,0,0,100,58,0,0,0,0,47,150,0,0,0,56,303,0,0,310,2307,0,1240,732,0,0,4060,798,175,122,0,86,0],[2236,261,3302,27056,39449,5182,9134,10866,288,55398,2743,5027,6772,1551,1319,6086,11686,4366,15538,1783,938,12630,6564,3232,2408,5018,1957,4769,22904,100376,8363,8697,863,3034,1805,4425,136,5411,854,6557,10066,381,10469,2221,4294,5145,2631,4408,1544,144,3845,13345,3240,1911,3077,1625,51],[0,0,336,0,0,196,3,108,152,1527,158,344,102,412,206,24,0,1176,0,0,494,202,0,0,350,0,0,0,837,295,0,0,0,776,207,1271,500,152,0,0,0,272,730,0,1042,685,2011,11,352,644,28,58,1194,978,134,85,0],[58424,86787,6851,103974,76123,66722,80508,75690,58514,279411,210467,12521,66595,76057,72881,180993,56221,50774,149682,134046,188714,147814,71774,48304,112477,192312,61835,32784,93683,89454,38720,78674,48967,139559,219849,151174,53606,166250,40082,111979,123710,16616,223139,96445,18133,88192,71864,46881,128483,87503,369013,78655,73510,12334,43794,32708,33477],[19496,19178,7750,27682,34857,72079,13992,21764,17861,46844,74295,22727,44878,60189,50395,40280,16393,87601,8497,59365,107634,32879,40765,42056,10363,61622,30501,34514,35970,143896,13333,33384,45744,31969,99496,149384,9200,171779,138007,12290,60179,38784,11144,6677,24660,13141,17720,32925,14251,20696,368368,21187,30539,14226,23161,26997,49380],[54133,38498,6818,44215,46444,86243,47576,54986,38515,131952,154354,11463,69436,59515,57390,117729,36974,106143,37582,52369,72717,105242,88692,49784,67683,153466,71010,90810,255772,133845,116779,259423,111200,154738,263729,204445,29855,179213,61671,59317,122982,11319,209597,61375,11138,41879,46799,56186,85909,33652,208151,116983,133497,9432,40994,33094,33035],[92935,106804,28733,323742,242024,200917,232689,237800,128944,664417,352342,56668,224064,176582,170430,283279,102724,111058,356191,261104,258839,404180,246830,257386,232786,215521,93198,112081,255321,315966,98811,205842,123269,395303,278654,344572,196311,340165,270128,213329,184686,68865,235840,84951,76640,276975,93714,107071,110856,63565,681354,203147,173365,31926,144046,80195,95124],[619,681,713,1688,526,1487,645,816,507,932,2187,1898,1020,2023,415,2036,1015,803,746,311,2276,367,1253,130,792,1665,403,47,1226,2078,522,0,277,0,1596,1550,9837,1634,0,752,1894,62,980,586,1940,173,0,1599,232,114,71,1062,2332,752,422,493,734],[0,261,631,238,39,399,421,110,374,106,0,416,174,0,54,59,0,0,0,0,0,0,24,0,18,124,497,8,0,269,0,0,0,274,98,101,0,36,0,47,0,0,0,506,313,5,2235,0,244,0,108,105,52,565,420,64,0],[0,0,854,127,0,51,199,0,0,0,241,525,273,0,0,194,0,0,159,0,0,0,0,0,212,350,308,0,0,199,0,0,0,92,722,0,0,0,867,0,0,0,1405,0,354,127,0,1017,101,0,0,0,220,1167,0,0,0],[2439,2941,3136,823,1258,2435,603,1196,1135,2267,1368,2602,3136,7603,4952,5512,3395,3445,1229,779,1127,774,483,2494,1174,3751,904,824,2325,6667,1775,720,1045,742,2165,4927,88,5161,4340,1017,3179,4326,3014,2250,5695,117,3359,2462,436,961,5660,1192,1632,675,740,114,106],[142,0,996,627,119,309,0,0,350,794,326,1368,2,164,517,0,16,0,0,304,800,146,329,288,71,232,509,0,385,217,2391,0,0,544,593,2166,0,1926,0,335,44,0,624,524,311,362,256,57,0,0,0,460,198,518,115,0,0],[99,1011,4792,8462,5274,2323,1221,2065,259,8506,4391,2952,2683,1692,1240,10942,1722,294,9614,2716,1151,3334,29993,3661,3758,2862,1714,1850,14172,2487,0,2714,863,4090,3984,476,2537,1408,2083,8357,9410,168,4562,1443,2328,7284,8596,2485,2259,341,5603,4802,12400,5286,1379,139,2],[0,30,1021,18,194,250,497,294,19,332,379,1804,101,37,154,0,0,0,95,0,97,67,0,0,507,0,1530,25,922,0,0,31,100,546,539,770,0,0,1612,0,195,0,881,1445,289,166,376,299,0,0,0,1,1117,1315,111,0,0],[202,97,740,4,0,0,62,37,128,0,199,391,0,0,23,35,63,180,0,0,0,177,0,0,102,13,0,0,683,81,0,289,99,0,589,0,0,38,0,101,0,0,0,210,243,173,458,194,0,0,0,0,185,576,0,0,1410],[190,68,1014,83,268,303,299,0,202,813,1563,513,441,332,370,34,0,166,169,231,1024,297,0,122,202,803,351,883,831,101,0,297,375,797,1059,101,0,0,0,0,461,0,528,302,1203,694,1229,303,560,181,70,200,200,168,65,100,34],[202,1443,961,85,0,1735,282,575,1418,526,852,921,688,3621,3640,75,4,1396,0,3,664,594,68,1017,312,2809,714,192,308,1544,105,354,2568,5,671,866,23,1764,1080,25,54,75,1381,486,1198,152,1021,400,132,0,4225,228,653,155,643,528,43],[170,201,3113,12,11,450,0,7,0,8,49,788,706,0,202,69,50,48,210,7,36,0,51,0,0,1338,1123,0,272,4,0,0,0,0,623,291,10,21,1041,0,0,144,625,403,98,0,0,1839,2,0,27,992,109,1023,290,609,2],[27,224,401,13364,2449,5974,258,96,117,3883,1514,348,210,991,92,31969,3100,11917,1370,923,3661,190,78,66,471,139,494,11462,543,388,34,0,167,208,167,447,590,147,261,172,752,529,99,408,87,310,373,91,57,167,4113,145,282,102,1325,543,2003],[0,202,593,139,0,87,632,110,152,150,160,551,372,160,138,248,148,0,0,0,600,0,1200,0,199,0,0,0,0,0,0,0,0,0,103,0,336,0,0,0,0,0,0,0,0,0,0,0,0,0,2207,0,86,0,1059,0,0],[0,50,905,43,0,0,0,0,183,31,297,167,0,0,351,0,0,0,0,0,98,0,0,330,0,0,42,0,232,0,0,0,0,0,0,0,0,0,0,0,0,0,0,0,204,0,66,56,0,0,0,26,0,143,0,153,0],[832,3716,943,4470,4741,4827,5060,3728,5973,6276,9194,1993,5534,4279,3460,8567,3077,2862,1498,2983,4524,7152,1095,4855,3317,5656,3479,1548,5034,7423,2596,4167,7946,2668,8388,5473,2923,5341,1733,2719,3644,5103,5593,4570,2573,2715,1457,2323,1747,1942,22298,2076,4345,1041,3398,4193,2841],[239,132,169,120,0,255,172,84,399,604,616,0,605,674,263,1412,78,1474,265,432,532,22,0,0,484,22,298,0,837,659,0,0,301,0,564,1111,1022,186,219,253,0,0,66,210,291,155,1689,0,233,0,0,33,200,38,23,0,0],[0,0,274,145,308,210,0,0,67,373,0,1024,171,51,0,0,0,588,0,30,0,0,0,0,0,108,0,0,210,314,0,90,0,47,187,638,0,0,0,0,0,1178,0,0,939,114,853,0,0,0,6686,0,252,335,0,0,0],[1287,2559,649,2994,1227,1342,2355,2353,1124,6384,6286,304,2321,3386,3625,6003,1149,4285,1656,3874,5976,4067,2799,131,4764,3918,813,2197,5709,5176,0,4694,2586,1871,7055,4738,8023,6406,135,2855,11533,0,3843,2629,600,2612,894,621,2945,2753,3814,2480,2349,738,431,1083,105],[10329,15948,1711,18108,15104,16503,10681,15810,18122,36564,23515,2999,9919,16296,11190,10870,3235,6878,20221,14585,16411,22564,20417,21582,10254,14105,5800,5376,19370,23029,34,12366,10979,24707,25685,25325,5851,44004,21655,11929,19859,6508,2530,4994,4669,13501,2829,4832,10600,14658,115495,10697,16777,3699,5815,4600,5428],[89894,163540,6514,252313,244906,118385,257780,209341,111144,747677,491175,15779,199192,126426,122136,279234,119691,104933,462885,255863,391141,700343,203032,144408,333972,294681,93100,89076,316842,310996,69823,310766,145735,391557,229151,215657,161032,270381,27053,219704,236376,9091,392576,154872,26116,366587,103756,31515,185290,220393,1127247,215622,244150,17030,82077,83821,141909],[798,372,957,341,0,17,405,321,388,2853,1157,1075,139,397,471,252,25,96,0,206,519,141,0,798,175,0,196,0,301,1214,0,0,1600,0,0,698,373,0,0,80,202,6925,404,590,0,0,0,700,139,0,0,106,606,754,1697,0,0],[1769,2665,3658,4361,3622,4302,1398,3138,4011,5817,6264,3524,2605,1451,2426,3071,2620,2118,3318,2621,2470,2744,4031,7676,1992,3804,963,1566,4689,4340,4133,3932,2649,2090,5661,7207,7266,2232,11075,381,1042,3296,3926,2683,6500,2263,196,2423,1786,3507,25388,2360,1493,1421,1480,2907,1491],[0,200,1492,0,0,278,356,398,297,0,436,0,100,354,0,87,0,0,0,0,128,0,0,916,202,396,202,0,104,1297,0,0,0,0,0,146,0,0,1806,0,0,0,50,501,1126,0,3541,0,193,0,0,103,202,1151,0,0,0],[28000,25421,22830,31105,18353,28971,10985,19580,27498,61943,29440,18630,19843,17057,19221,20266,6930,9599,24104,20181,21675,43821,61113,118343,19630,29912,12898,19021,33183,45553,18258,30371,20537,42221,59157,76799,18381,19561,75176,10503,15035,20572,28433,19575,43532,28005,7040,25261,7798,16052,160955,19390,19746,6971,30089,16570,16674],[104,0,1102,95,52,410,4,170,373,1069,183,466,322,202,393,0,0,176,0,165,0,0,0,0,101,249,48,103,288,0,0,0,72,143,400,168,0,0,0,103,0,45,0,143,162,134,0,889,0,0,185,138,28,0,0,0,112],[196,1154,811,540,901,1097,7,580,189,145,1486,458,998,566,482,1042,402,371,571,735,607,467,1720,513,122,1089,1864,0,712,2741,55,1127,529,553,506,1085,0,5871,0,828,1627,5028,1053,0,720,0,3703,914,260,544,2543,1,620,957,252,826,0],[12450,20481,8257,25627,20163,21431,15471,16958,38557,54467,41135,10752,15876,19052,19870,20046,8875,9671,19295,28676,27639,29133,37650,29043,26696,50441,25713,12225,31682,38482,8805,26261,27789,28924,34170,37777,9321,30399,21668,15560,22180,12987,45374,15501,22518,32882,16789,16255,19262,22654,176161,12886,15579,7044,10160,11693,20638],[132,585,433,744,202,618,606,650,557,647,1251,364,426,511,129,1461,261,269,325,373,933,35,545,334,501,983,202,690,556,384,73,305,79,279,772,897,49,997,487,563,284,409,320,753,587,388,586,0,179,163,3910,371,518,165,483,354,1733],[1347,532,634,2387,1082,1555,1467,363,6324,3998,1667,1512,3110,1263,1050,1282,1266,1152,3753,2491,1893,5876,1475,1574,7390,2221,3622,2127,3115,3500,469,2312,4240,3578,5918,1212,167,2472,120,1991,103,71,4388,852,1418,2445,1547,676,0,592,313,3866,2516,2741,664,0,16],[0,186,428,60,180,0,0,0,400,400,0,277,32,0,11,0,60,0,270,60,0,0,0,0,0,0,6,0,0,202,0,0,197,0,1096,752,0,0,0,0,0,0,804,0,244,0,0,615,60,0,466,0,73,746,188,0,0],[315,867,1369,1288,1912,2127,1679,1207,330,3537,3744,1104,1772,2477,1432,6638,800,2326,1342,1647,1568,2055,861,6103,1514,1963,1738,58,1898,3067,0,202,461,1823,4150,4607,645,2562,3081,1575,2462,3521,2196,901,3843,714,4380,835,1021,369,809,1704,1722,1645,2029,1454,211],[830,161,1354,616,945,2575,121,1296,1439,774,4508,512,2016,2768,2225,1670,1176,230,184,3369,1702,243,950,1260,352,1742,4641,2246,2979,985,2451,832,37,832,5241,6380,561,5353,1199,401,1454,6623,1728,1687,1721,454,5560,2948,1449,0,3495,1857,1157,981,116,1371,0],[0,0,0,0,0,0,0,70,0,0,0,0,0,0,0,0,0,0,0,0,0,0,0,0,0,0,548,0,19,160,0,260,0,0,0,0,0,0,34,0,0,0,0,0,0,28,0,0,0,0,2361,178,0,0,0,0,0],[0,0,201,57,0,0,0,0,0,0,0,762,0,493,0,0,0,81,0,0,0,0,0,0,199,354,101,0,0,0,0,0,206,0,311,0,0,0,0,284,0,0,283,42,582,0,132,42,0,0,0,0,566,63,0,0,351],[25461,44249,10870,68454,40794,45327,26642,28986,39929,126118,92241,11935,41525,44051,44172,51473,12992,25832,36784,24969,49832,38940,52086,91936,52552,70215,22192,18271,44162,77290,15118,41345,32496,47219,72466,74038,31685,112523,63834,27778,21079,14495,35257,38893,32168,42543,10352,12531,24174,37433,462380,28001,40842,5574,36899,45382,68138],[26574,30904,7380,57422,46036,40575,39652,47024,51995,121487,159084,15987,37186,48377,47607,68709,25927,37320,73449,45308,86235,85993,87807,68154,43462,91602,34999,7991,81651,97873,13644,26039,51521,47596,80201,67174,52314,71378,35540,45900,65665,7462,28085,37318,22045,42655,16519,22971,53537,42578,303039,32658,70860,8817,17544,25587,54113],[632,1346,290,3053,2117,1996,1801,3607,1698,5453,5386,957,2335,3138,1538,3792,807,2208,3649,4758,4468,2697,498,824,2152,1817,427,81,4947,2974,0,1016,1565,1992,1734,4029,6823,2177,6126,1721,7223,0,10290,1535,2385,1377,1287,3610,1932,5355,7344,2026,3857,326,949,408,0],[21472,26375,6960,53651,34786,39348,34513,30293,33695,82790,100581,10051,50046,47491,40229,70348,36681,49699,55016,41877,61983,81788,57138,38376,40144,70939,32130,15295,77063,112449,8183,31134,46036,45885,73727,56814,24252,86265,24857,45492,45934,33621,35174,14106,13935,29182,21402,10768,30047,31410,240858,40401,38422,7652,16486,20570,30546],[13350,21603,3560,35552,25726,28218,24071,23527,12096,57606,49561,6240,32174,25037,23127,39158,9804,8776,34699,26655,43280,56776,16578,27174,29249,42803,23299,2175,33314,42868,13486,17768,24886,33494,45807,35672,16532,46458,10651,28328,13676,11877,13301,8816,11330,36736,14529,13530,23593,18820,66461,16512,30274,6515,10326,11857,23498],[7945,13436,3673,21146,19158,21385,14669,13588,11105,30919,31693,4065,15356,11667,14044,12518,2334,3253,19924,15915,20735,26554,19050,9994,16755,31870,17098,4742,27149,45580,2033,9553,14546,11774,23638,21453,5938,31170,12840,15083,22121,8683,18392,9579,7620,19029,10015,11769,15683,7538,62367,9757,20848,6306,7757,6751,3951],[1359,6882,572,2740,1375,683,3594,4044,1290,7523,7338,1089,3206,1335,1461,8582,2987,2699,3490,1472,3456,6696,2526,2257,4307,2002,2300,5203,2382,1231,826,1098,563,7587,6747,3484,1678,7596,700,5526,3689,804,4088,1464,844,1274,1701,519,3407,1360,135,2164,3085,1990,703,815,1760],[163,288,525,129,938,258,1371,1753,934,2639,1986,1090,364,117,502,449,0,949,719,447,1657,5051,2668,0,497,323,1484,0,803,1265,0,917,1107,1179,454,132,0,2153,129,244,0,0,3336,1217,0,404,1234,1444,817,803,2749,1750,1771,1472,186,1001,0],[8852,19342,2846,13393,12547,19203,3660,7070,12098,13046,17856,1744,8345,15277,13465,3271,887,1790,4020,11535,23987,6129,18573,8907,1367,24137,10726,3344,5561,8632,7345,6301,7327,4752,16847,18096,1877,26988,3082,5638,6783,6597,3120,2438,2850,1910,7109,7081,8515,16514,62550,3656,20844,6091,1991,21,1833],[5602,17239,11530,19697,10700,15309,9053,11726,28932,44024,28953,26372,10853,16464,16349,20292,8262,9179,16579,13108,28857,25702,47629,59232,10343,27831,10511,9083,8400,31622,7625,10619,21836,26131,24286,31588,4430,15356,27985,6833,8192,4229,14619,7513,23924,16368,9540,9658,11781,10760,171864,9430,10562,4939,15399,10223,12620],[7214,1351,5104,4652,6670,15174,1341,3080,2932,6836,5679,4598,4261,10464,10507,4088,2529,5624,1587,4653,7115,3247,7385,8584,294,10550,5290,3429,4977,5570,3706,2905,2111,5133,11545,4491,116,10128,1253,757,3227,540,5496,3088,5177,2302,4938,17246,3548,2216,13970,2307,3046,2434,1092,3517,3567],[56121,76083,8982,176837,129030,138236,122754,109526,120833,350684,448150,19112,163119,156172,133654,273032,125297,168627,173291,154245,295135,195083,117756,65101,162859,287644,156066,72530,249367,206520,27302,128225,136740,123508,275421,210370,173053,375043,51769,155393,133432,29518,144194,83452,29740,139595,89989,54784,132989,84636,558644,117939,132949,23105,38328,52150,72800],[109976,123688,20961,183914,154123,218518,118174,127891,89058,287055,280005,21690,229257,210206,214910,188943,56494,207424,204283,130730,234139,354601,298299,146486,190333,420329,203802,88049,490816,467634,60222,151038,166256,155045,520877,474749,88869,554345,116359,133651,230314,30143,96334,86717,55301,189835,124814,82189,140614,114509,643167,162412,290747,39243,75081,95188,96625],[0,49,0,109,0,92,0,0,0,43,45,131,0,0,0,218,13,1,0,40,112,0,0,0,0,49,0,0,0,0,0,0,0,49,42,3,0,0,0,0,0,0,0,0,0,0,3,0,0,0,0,0,0,0,0,0,0],[3025,5798,4047,5080,4007,7672,3363,2509,3390,3496,6334,2823,6706,7040,9876,5777,1219,4017,2042,2216,4970,3781,5014,6128,2857,6489,3371,184,4729,18826,5557,1907,1238,2153,5335,9260,3678,11891,2080,4293,1547,3691,3280,1949,1993,2986,6688,2842,1992,1467,25715,1549,5491,2910,3355,1622,2697],[96435,155291,10700,269865,195329,184379,164983,164434,119801,686990,613239,18182,187737,180177,172926,257414,157363,132755,258611,302356,412782,391416,150945,143128,238376,357773,153005,295643,363630,326794,35819,255890,129449,267333,316144,277978,140725,344352,70542,161387,182858,9797,358195,136203,23594,222769,94028,52495,213617,106259,499093,151005,171247,14338,67954,75153,134289],[0,0,86,166,0,0,121,37,0,300,0,0,0,104,0,0,0,0,0,0,153,0,0,0,0,0,0,0,0,50,0,0,0,0,0,315,0,0,0,0,0,0,0,0,836,0,0,0,0,0,0,0,0,218,0,0,0],[0,0,0,0,0,0,38,0,0,180,0,12,67,0,0,0,0,0,0,0,0,0,0,0,0,0,0,0,0,0,0,0,0,101,0,183,0,0,0,0,0,0,0,0,0,0,0,0,0,0,0,0,0,0,0,204,0],[0,0,1731,0,0,99,0,0,352,225,0,328,981,0,169,0,0,0,28,200,0,144,0,238,200,0,603,21,111,721,4115,0,0,193,804,0,0,0,1209,0,0,33,2004,216,1152,146,3596,977,0,0,0,1204,368,1764,549,0,0],[606,545,949,0,0,0,0,0,32,0,659,798,0,0,0,32,32,0,0,0,0,0,0,0,404,0,278,0,0,242,0,0,0,24,429,0,0,0,1256,0,0,0,622,0,537,0,426,177,344,357,0,17,0,174,0,0,0],[136,652,2657,89,603,187,213,580,571,320,603,1601,777,188,482,91,189,0,196,399,202,173,0,994,202,879,413,0,889,1024,7304,168,0,0,528,200,0,1366,401,0,0,435,1413,1772,398,0,2995,395,0,199,9620,615,1605,1502,0,0,0],[939,2378,604,3608,3869,8113,1274,2968,8760,10322,9484,1850,4048,6105,4160,2421,865,946,2210,2444,7830,5918,14567,2529,2718,5635,1305,178,15392,17854,4010,1055,2844,7730,14052,16561,0,3872,413,2879,9431,382,3325,4363,5354,2398,4187,1697,6218,3949,21566,1375,9085,486,345,1133,2464],[125,102,176,0,258,129,0,201,192,375,0,210,155,0,126,0,30,0,0,52,128,40,0,808,40,93,0,0,16,554,0,0,99,32,318,0,0,0,80,0,0,0,0,53,2561,0,1018,1569,0,0,1503,0,62,389,78,126,0],[5214,4837,1071,8303,5555,5123,5020,6619,5389,19437,19924,1157,6376,5743,3498,15020,5366,1826,9452,6923,9454,12074,7728,3612,8524,11070,6916,1388,15323,5279,3060,10328,5930,3544,14108,15787,4856,7862,2583,6094,11596,0,16272,9005,4213,7357,4842,4232,4437,4314,12847,4185,7091,395,3543,2931,3636],[54,89,642,0,0,40,0,0,155,0,0,718,55,0,0,37,0,45,123,22,0,213,0,252,0,0,475,17,122,0,0,0,0,178,90,559,0,0,88,0,0,0,0,888,408,308,1515,0,0,0,91,40,2,544,91,66,0],[1185,11681,4507,3013,2954,7601,923,4777,61,24270,5568,1545,5640,2005,2597,3407,223,981,1543,914,5505,1207,333,742,389,873,252,944,1792,4166,24,625,1779,1177,2336,6163,673,1280,5476,914,4217,1279,2430,5993,1866,595,391,1323,647,659,7710,3904,5789,542,1915,2600,3092],[0,0,438,202,0,403,0,187,202,201,406,918,362,564,0,310,0,0,0,196,26,402,0,0,201,0,403,0,598,0,987,0,0,0,282,195,0,0,0,91,0,0,946,591,3339,0,3131,532,0,224,0,946,1396,873,0,0,0],[0,0,0,0,0,0,0,0,0,132,0,304,0,0,0,0,0,0,0,0,0,0,0,0,0,0,0,0,0,94,0,0,0,0,0,0,0,0,0,0,0,0,0,0,0,47,428,0,0,0,0,0,0,0,0,0,0],[9,11,186,0,0,0,8,202,0,0,101,386,0,0,2,11,0,0,0,0,0,0,0,558,0,0,84,0,212,0,0,0,0,201,0,0,0,0,36,0,0,0,111,304,170,0,101,0,0,0,0,204,658,116,0,0,0],[78,183,191,168,27,90,0,87,0,81,120,152,0,0,26,33,0,0,152,33,50,0,84,0,218,0,0,71,191,0,0,0,27,0,152,65,0,0,82,201,0,0,385,25,302,0,0,476,0,25,0,0,51,100,27,50,0],[9605,8426,6793,16221,8498,15344,6442,6049,8672,10146,14201,8569,18927,9638,10027,14115,5206,13153,5982,15885,12385,15988,17849,16493,7013,12899,16590,4261,18865,21294,124,3090,3106,11065,8563,11575,4330,31705,5234,4613,11627,8617,9758,7663,13557,11005,13065,11494,12639,7202,35634,11998,11152,6092,5496,10659,6228],[1618,1110,616,7867,5811,2315,0,0,69,0,116,136,6681,4872,4804,7388,2863,2152,0,0,0,20556,6362,5171,0,0,0,828,1807,4827,0,0,0,8447,4304,7015,0,0,0,2651,4147,0,3647,1989,1326,6251,2540,1637,6280,1012,195,3669,2990,402,3162,2329,404],[2699,1130,352,1133,1086,1804,1671,634,1172,2640,5928,985,1417,2098,1749,2012,1047,4299,468,1525,3563,1925,2029,1617,665,865,1847,0,1254,4817,49,1883,700,2190,2705,4795,0,2578,1199,961,0,0,784,887,0,423,275,180,636,1387,5606,1447,623,571,635,1008,1495],[0,0,0,0,0,202,0,117,0,0,0,599,2,0,0,0,0,0,0,0,0,0,0,136,0,945,202,0,184,0,0,0,0,200,0,0,0,0,0,0,0,0,0,101,0,0,0,0,0,0,0,0,0,163,0,0,0],[324,1198,1624,122,410,1745,402,381,1309,1409,1315,1023,417,437,474,29,0,1233,176,400,1561,570,0,1408,0,1565,3103,0,653,1479,0,198,0,0,756,605,0,933,1005,0,1397,0,199,1024,1592,1020,2179,1642,356,1006,0,1049,2125,374,0,93,2500],[0,150,19,0,0,29,0,0,0,0,233,0,0,0,0,0,0,0,158,0,0,124,83,0,0,0,0,0,0,100,0,0,0,0,20,0,0,0,0,0,0,0,1275,0,999,0,1876,1323,0,0,1155,198,263,96,0,391,0],[10317,35388,18775,41905,26319,67131,6851,10864,19805,29692,27799,9468,49843,10807,11061,6657,3825,23141,11811,25764,49114,32298,56507,144523,16027,38756,26019,9135,28559,207173,6834,16598,41748,26728,19340,29294,19178,93114,189325,13633,25133,27075,11854,8769,38803,17042,11293,59906,6396,6762,169027,27950,9767,31854,8129,9969,8363],[0,193,0,0,0,400,0,0,0,0,202,1206,0,0,202,0,0,0,0,0,0,57,0,0,0,0,0,0,0,634,0,0,0,693,0,198,0,196,0,404,0,0,0,200,0,0,0,699,0,0,0,0,0,126,650,402,0],[34097,63161,3001,83854,67742,75017,49737,48596,23292,218761,100885,6848,43008,41134,37532,183555,31588,46036,62392,47315,76682,73154,19112,19767,27713,34202,11595,24712,37189,59197,11811,24215,20137,48146,66630,73492,27120,76628,14719,48853,42447,1500,43427,26082,14694,38585,17256,15121,28157,8201,105130,29232,46225,3229,41284,29357,18374],[97,0,543,745,1077,404,510,841,69,847,134,440,128,150,410,515,34,118,147,0,549,562,1105,95,435,559,1079,115,717,403,0,140,268,48,1030,2189,0,0,205,586,256,476,460,220,351,0,0,654,52,282,63,65,705,2,119,237,64],[1991,1592,3727,2140,997,2121,3057,2584,1770,4959,1058,2658,4592,2478,1643,3166,441,145,2118,1171,2008,2953,680,2555,1508,967,1292,1039,2948,3658,6297,275,1603,5639,5176,4602,1789,4308,3075,2587,2361,2823,1699,4056,5061,1387,4977,2430,960,695,6414,2406,3480,1712,836,1183,33],[76,188,876,234,0,596,16,478,177,2029,1122,770,788,1085,227,192,165,0,296,712,571,458,308,0,0,202,755,0,519,278,4916,69,322,810,0,831,0,283,1492,66,192,0,1100,176,468,0,937,464,76,152,6045,317,475,470,384,60,0],[0,0,248,0,16,132,187,0,72,34,0,153,0,0,0,0,0,0,0,0,0,0,0,0,160,102,0,0,101,404,0,0,0,404,0,0,0,0,60,0,0,140,0,0,2137,0,175,0,0,0,0,94,435,525,0,0,0],[0,150,1625,202,134,47,532,378,347,171,729,2232,442,513,197,0,13,8,197,4,345,96,6,1590,1074,129,2389,0,190,414,0,0,165,744,520,483,0,24,2345,0,0,165,193,780,5582,599,1783,2593,33,0,309,100,877,2054,258,0,0],[1966,1277,683,1751,913,298,1340,1428,865,1470,1822,0,1882,2087,1260,2974,147,155,730,200,1233,125,0,1010,1017,2543,736,1000,2170,1013,1490,202,396,210,2580,2454,0,1650,344,993,2108,0,726,279,183,921,1344,2128,1033,0,5585,158,474,622,0,2425,0],[84371,112606,21621,161314,108029,86527,101976,97622,196155,298376,317012,21964,75670,72418,63068,192169,65945,78118,157937,141572,172948,187647,200700,165800,137769,149207,63191,67308,113457,129838,37644,89109,62065,169855,194068,189482,86166,165393,95472,95654,99556,24337,77820,62656,45551,124863,37025,41015,67478,74171,618973,72339,135530,10574,70931,62528,32384],[18706,9772,2708,32167,28771,20615,21931,28638,41398,75505,70861,5518,31977,31931,24830,37032,11279,9990,51264,25586,39158,47393,31873,27767,31134,53095,16045,13874,71579,45627,5744,15520,40261,45259,36423,33085,19008,43280,18811,34767,22246,6356,22365,17239,7080,30509,10147,7518,18935,17747,120599,24066,15419,5005,14759,9743,32606],[0,0,163,0,0,0,173,375,0,0,242,204,278,0,101,0,392,0,9,0,0,173,0,970,1041,255,1257,0,489,529,0,0,0,0,0,136,0,0,0,0,0,144,429,120,1167,0,138,952,0,0,0,294,0,1290,0,0,0],[60421,104854,3979,119796,107682,118761,86375,111881,86240,353807,126277,9433,79156,82408,78398,57975,14827,17604,209390,136458,165250,207851,92785,33495,133018,199599,46600,3402,139932,182525,28552,107916,88781,158571,167746,108946,62242,169984,45351,93791,49589,12786,146775,62148,25087,147144,51273,28304,92180,68455,524257,91845,94627,4867,27631,31715,22405],[403,0,332,0,734,363,406,412,0,216,977,1351,978,1068,0,865,1185,232,0,560,573,202,2417,517,370,454,0,1077,1862,573,0,517,268,142,640,966,0,652,486,200,61,0,892,0,1280,1002,1875,0,0,913,0,805,360,0,238,0,0],[0,262,475,365,115,403,237,186,21,1117,443,394,282,0,242,495,170,336,63,666,539,165,63,21,83,150,21,156,181,21,63,0,0,77,191,463,21,84,682,142,272,0,308,41,46,63,109,0,42,105,583,0,570,16,298,176,126],[77,1,319,49,127,249,93,16,120,443,186,146,185,0,218,26,0,184,0,0,90,0,1221,556,202,0,0,0,364,663,0,107,449,0,808,36,0,171,0,33,0,0,482,399,12,344,1125,417,0,0,2390,0,360,264,0,0,0],[0,46,266,48,0,20,48,149,0,48,18,424,0,0,29,150,0,0,0,24,0,0,0,45,0,44,0,0,94,0,0,0,47,0,0,18,0,220,0,24,0,0,0,300,29,0,0,891,0,78,0,0,0,123,0,28,48],[0,0,505,0,0,170,64,0,0,0,0,759,0,0,0,0,0,0,0,398,334,0,0,0,0,32,1236,0,404,0,0,0,0,248,0,0,0,0,0,0,0,0,398,86,596,0,1101,1773,0,0,0,221,0,0,0,43,0],[81277,84794,11343,168622,121954,128630,112121,91685,77655,299953,446023,15900,138781,132935,123109,154098,37392,75493,140041,183180,231807,201069,122549,87623,123267,265711,114954,39858,260683,235753,14208,97694,151200,146261,314673,364035,48298,316284,41883,115617,189044,10127,167764,108233,26679,88062,85436,57373,173271,114373,466599,103327,242724,27320,37489,57796,76998],[1342,828,1227,4255,1895,3458,631,1008,558,3569,544,638,1693,878,797,1505,464,288,1164,1902,2595,830,3192,5922,1321,931,1465,719,3780,2427,115,961,1402,761,2487,4588,4814,1493,3934,2847,4830,1107,2828,1800,151,1710,866,1799,1667,1612,4073,1489,2752,1175,573,467,1630],[24554,13414,7290,44644,30139,37943,25488,25590,8349,51016,26443,8013,34122,18533,19078,66156,44573,48073,44130,44917,33248,59119,89449,82367,59012,42059,27020,11296,48105,39299,19446,32224,23284,54525,43470,44899,37287,49781,64966,49949,32084,18306,21098,13105,15184,28409,26220,51545,18663,8876,37458,40260,29730,7306,45484,29499,9154],[15192,23305,13869,23437,20397,22432,14631,14224,23866,59702,35666,11002,18934,16655,14678,24907,6960,10659,24604,16682,20790,35509,47757,54015,40332,37543,12902,39671,43459,54135,2447,24009,20190,31919,40957,36987,22816,32453,37613,18224,17951,35284,17450,13349,26873,34559,18791,16391,11304,7183,133104,15484,21712,11679,29225,11207,13649],[449779,560277,89644,850577,668233,778319,376795,550475,473968,2495852,1046686,427679,559726,471478,586559,1032999,428555,909009,496032,507619,570927,701648,374935,704841,367223,293536,102883,771784,542500,709781,101553,267318,364785,734828,1243832,1560462,275017,430072,466913,410427,455087,288546,433179,223175,118436,709232,202744,226815,250142,102066,1410558,466830,515966,88248,491176,362566,524809],[2314,3765,1984,7838,9017,4242,14760,13492,1897,26526,46551,3837,3269,2302,989,13272,1675,920,18361,4939,21396,35446,36709,17330,7532,10744,5151,6731,11036,20261,9205,7182,1774,22246,19919,18758,3215,5443,4997,6621,2950,1524,5871,6011,2336,39692,9888,6398,13465,8108,31354,2723,13378,202,9760,8468,7987],[280,99,2884,913,292,611,0,160,583,683,216,668,493,475,331,567,320,560,159,5,252,212,389,91,431,620,1062,39,1021,115,1102,11,622,0,240,248,0,240,0,430,832,501,509,322,468,1248,2181,897,75,0,75,367,820,712,183,127,593],[0,101,34,0,0,64,0,0,59,0,0,23,51,150,0,0,0,0,0,0,7,0,0,87,147,144,67,0,0,0,0,0,0,18,35,79,0,0,0,0,0,0,0,0,90,0,218,0,0,19,0,0,22,0,0,0,0],[608,351,71,60,353,548,103,333,152,876,799,0,138,424,343,922,2820,838,532,160,989,2110,418,875,154,310,179,54,456,41,0,712,156,1438,1173,65,0,604,0,504,550,0,595,520,0,593,77,0,40,1040,918,511,372,0,124,148,282],[2511,3362,1081,4248,3611,5023,2414,2535,4190,6085,8661,202,3562,3824,5205,6038,1916,3869,1158,5258,6681,4377,4718,3190,2082,8320,3182,462,6941,7844,224,5466,2551,4771,6582,10399,0,15053,0,1952,3551,59,3972,3934,0,4262,4484,705,2769,3433,29177,2977,5741,1933,2051,693,1238],[62021,176712,6208,707208,555417,125335,272030,390498,12249,1899194,1980236,35946,498455,93908,130281,851523,407503,89380,1406517,375249,1626230,720106,178921,146303,1163185,545140,158197,126766,321011,83494,81186,372546,128067,244905,372828,96071,453938,277695,7964,629375,715496,23156,375856,324826,36357,1259526,234513,43370,423966,258517,742348,262436,230897,3661,190023,117044,59151],[24454,34066,7504,64289,56671,74803,19064,26248,30403,94232,66174,16457,32989,31104,39193,22221,4410,9928,45260,35040,78343,48709,59450,88174,29744,68822,32697,9861,90594,116042,8559,33586,38511,42044,76343,78634,28854,53452,57519,28794,30487,3101,33717,23143,22715,34511,17105,24071,20818,17814,208191,25742,40797,5742,13772,13357,35331],[3353,2084,2212,3041,3013,2044,1652,2122,1560,5297,11002,511,4580,2824,2779,6118,693,3200,1664,1132,4128,6174,5084,1816,1281,3928,1936,334,4703,4561,1877,4345,1884,3400,11301,5345,0,5539,0,1085,1953,0,6276,1828,835,912,1896,92,4703,1251,11,2261,2949,614,2150,1407,4330],[209,968,1292,606,1476,557,366,1605,1231,413,4026,1242,193,257,729,1333,139,119,809,1207,875,1105,1916,510,1630,2783,2031,56,1132,2283,0,204,1241,1515,1663,128,9059,1785,69,1984,0,208,464,1021,425,1406,787,1322,1377,365,31,1024,1423,744,299,172,3011],[482,972,675,993,300,931,1015,1070,1343,1753,950,947,1011,1382,692,210,768,0,942,782,781,1336,0,3557,605,486,69,192,766,1896,0,504,1092,245,2444,972,1317,1163,0,1118,0,1680,859,561,1801,2502,904,1421,91,172,7802,536,1437,0,824,679,0],[2703,1961,760,1310,1525,2253,2784,1580,1671,6579,1961,323,2579,2237,2643,5641,2090,903,2620,397,1919,3066,4145,2362,2095,1325,841,994,2391,1513,0,728,1485,3357,4424,4431,245,5522,462,1450,4674,728,884,403,0,1278,2143,910,816,499,8370,1391,2533,449,999,739,509],[0,350,62,186,0,148,202,0,226,802,404,603,319,0,0,0,0,0,0,199,0,31,660,0,0,145,202,0,624,0,0,0,199,0,135,0,0,0,0,0,0,0,396,1040,451,393,0,301,0,0,0,200,271,0,0,0,0],[302,1666,606,202,192,219,397,813,1071,1256,1247,220,280,476,0,254,23,0,0,0,200,718,61,1705,1591,1782,1503,0,1384,3859,0,1760,1002,808,1083,1816,0,0,396,968,4413,0,581,1976,2992,621,1184,271,1080,1058,539,118,1879,267,68,120,426],[144209,289575,4512,234303,160283,149546,192440,217054,221953,528925,352750,15809,193290,222630,203193,121940,43419,49673,378586,255308,332187,395095,298315,118630,332528,268258,108316,11778,323160,369547,47057,242106,129212,304368,310883,286607,169876,353417,96279,261967,136775,12715,259862,105718,45291,276763,111622,63237,170718,143929,966266,217927,210173,11965,85960,82229,91223],[0,0,0,0,0,0,321,233,0,80,44,0,0,0,42,0,0,0,0,0,0,142,58,0,380,256,0,0,88,0,0,0,0,0,0,0,0,969,0,0,0,0,156,0,0,0,0,0,0,0,0,0,146,0,0,0,0],[796,990,200,1520,762,972,903,853,0,3984,1571,0,1531,592,813,4546,320,674,2182,752,946,2435,1143,156,757,945,986,749,1403,899,55,390,339,708,1913,5576,97,355,0,296,1242,0,873,501,79,665,576,1000,570,90,536,797,996,9,1615,925,0],[126733,161247,10118,264102,210274,169944,183147,196479,120990,445927,453905,20136,243809,148918,157276,290827,109622,116299,289240,188571,333057,336506,214087,115692,267833,352949,211946,64000,382086,324483,52260,144394,102012,188006,323686,290872,177134,343229,117269,240065,246555,39435,200288,142063,49020,257204,124182,102594,216653,120381,515833,178129,289822,30228,65791,84199,129880],[521,1166,1875,5363,2522,2781,3015,1915,1573,11726,1661,2101,5019,2083,2541,10621,2263,524,6560,1749,939,14450,577,529,10143,3347,2538,1099,5951,1829,303,1972,643,6698,5483,4558,3293,1316,1289,10283,260,9551,8531,3075,5495,10826,8157,2969,2519,800,266,8988,5458,4872,2710,138,410],[74761,96534,16802,75051,46614,88332,31850,38191,105671,125461,71839,21395,38518,33617,40084,58660,12089,32210,67297,47392,86964,64226,55902,61424,66481,58875,30154,7706,91301,141590,9668,19352,33899,50303,82682,75252,20591,88235,297485,61495,43448,19557,37835,19370,24195,50070,26820,86342,37644,41656,532442,56734,44406,10989,62772,24450,14820],[0,0,876,166,0,723,188,288,0,600,279,1544,202,383,0,111,0,0,340,251,72,415,382,1204,280,1208,157,0,523,165,0,0,0,0,117,2163,0,0,4269,195,61,7841,775,540,398,0,3037,2650,0,354,0,358,519,628,321,0,0],[14,0,0,0,16,160,35,0,95,0,0,235,84,15,0,133,0,0,0,0,0,0,46,0,0,96,0,0,100,6,0,0,0,0,18,295,0,0,0,0,0,0,95,44,0,0,146,0,0,0,0,0,0,4,0,0,0],[0,26,306,92,34,14,0,226,44,648,93,7,82,148,309,0,0,0,466,0,134,45,18,212,0,0,0,0,4,20,0,144,50,0,32,0,421,0,0,0,65,0,359,9,7,1,0,373,0,0,0,81,173,0,19,534,0],[19826,20589,11772,26669,25897,27859,14587,16929,22850,53772,33829,10201,25221,21369,23614,15398,6951,12539,17742,13990,23160,33344,30671,61622,13619,28962,22213,10004,29835,45947,202,14390,21956,23392,40543,58785,5139,34873,54950,17835,26826,11870,17298,13832,17029,15393,10592,18003,6793,11432,140653,15814,30112,8716,10222,9312,18528],[0,0,100,0,0,0,88,0,0,0,270,0,0,0,0,0,0,0,0,0,0,0,0,0,0,123,55,0,177,0,0,0,0,0,0,246,0,0,0,0,0,0,0,9,525,0,496,0,0,0,0,0,133,0,0,0,0],[1407,2102,887,3524,1329,4943,868,1144,2006,2502,2835,652,1874,5277,3505,2309,2138,1276,1241,1218,2983,3268,2721,3055,1189,3865,1233,28,3320,5022,0,818,551,2225,3842,5978,1979,3057,2758,360,282,2655,997,1447,1348,2416,1408,438,2998,2194,1435,2716,4634,375,3183,1254,6135],[48847,66375,12864,76649,59865,60968,69222,71418,57592,222914,228506,13667,80140,73927,68893,141055,49650,82730,108420,81423,140793,145633,85883,61468,98339,184361,83385,52033,140292,152623,12082,100539,78337,92633,159700,137271,74973,187831,33655,85592,101026,10111,102818,44300,25779,71601,39867,36029,97684,51707,296117,68515,85679,15308,24641,43817,63523],[0,189,43,0,10,0,0,151,184,317,131,83,58,103,320,18,0,0,0,0,86,454,0,0,0,42,0,46,0,202,0,0,0,384,0,1035,0,0,0,0,0,0,216,12,0,0,12,0,0,0,35,202,174,152,149,0,0],[1090,981,0,719,955,1458,1726,1386,1283,3406,3037,76,2060,1412,433,1751,1244,1667,1358,1563,2254,4052,2627,449,2995,2495,782,0,2629,1695,0,3104,1620,1807,2615,2979,1176,6634,618,878,1954,0,1639,703,0,1365,1103,247,660,754,1011,668,1265,293,827,757,4768],[0,3,2,0,0,0,0,0,10,10,114,23,0,0,149,1095,11,42,0,0,43,69,29,3,0,3,37,225,3,0,0,0,0,0,169,160,13,10,0,0,89,2,195,0,0,0,39,0,0,0,153,0,188,61,0,258,6],[179,983,2415,167,23,574,0,185,179,108,1264,398,201,1084,381,797,578,92,38,591,46,122,0,539,0,1522,1816,127,1346,1217,0,0,464,0,453,295,23,46,0,91,46,946,3177,872,4631,0,6251,2977,0,0,23,1387,1228,2727,94,0,70],[0,0,102,0,0,0,202,0,0,0,101,0,0,0,0,0,0,0,0,0,0,0,0,0,105,210,88,0,0,0,0,0,0,0,0,0,0,0,402,0,0,0,82,101,1221,0,0,0,0,0,0,0,31,0,144,0,0],[0,175,1628,246,310,756,202,651,862,1467,607,2303,192,927,546,140,29,0,0,0,204,0,0,0,302,400,194,0,1079,360,0,0,0,0,202,1067,0,0,1134,0,188,147,1402,859,2126,0,1338,1901,188,0,2834,398,958,904,226,0,1285],[343,44,131,656,280,457,388,230,46,1746,20,46,0,0,0,1169,129,38,1230,788,3555,605,1059,129,0,1215,212,42,531,0,0,594,1726,94,662,1890,232,0,376,522,1000,0,388,103,1024,0,648,54,1434,1311,993,745,221,52,0,0,0],[58,565,2048,2042,1156,1009,1112,1877,519,2577,2094,1126,1082,675,406,1309,694,3082,486,102,1672,368,516,554,3922,5306,1968,101,2341,1076,0,30,395,892,1928,2469,375,475,2103,69,285,47,3087,1285,1192,2105,1089,4434,626,691,2098,2073,1418,1212,709,401,64],[404,204,0,0,696,598,360,695,509,589,690,424,1063,404,154,194,98,597,237,712,418,0,2424,707,312,678,210,0,148,927,0,564,402,183,0,865,832,0,21,799,0,0,502,184,0,409,0,0,1412,75,7369,204,472,72,196,435,0],[1024,114,940,469,102,85,0,462,0,414,525,0,828,135,808,364,0,0,0,123,0,0,392,0,130,0,207,29,594,392,0,0,0,0,163,101,0,0,0,230,0,0,1058,475,4850,181,3068,768,123,0,0,238,504,839,1146,0,788],[29947,72161,13860,42834,32573,193477,20568,30904,60104,56796,33807,19455,128545,51427,54079,9354,26213,67659,7778,12469,80182,28204,129279,329546,7936,75586,48078,33616,153322,270651,22232,25479,50139,29148,49054,166041,14565,294530,235764,13580,55979,29363,16079,17925,82749,5452,28907,53906,20214,22123,171322,24744,27044,27822,11022,26553,27225],[48,0,194,107,0,97,36,229,143,25,0,342,57,0,223,0,0,0,0,0,0,0,0,180,0,0,155,0,863,8,0,0,0,0,115,0,0,1018,0,0,0,0,0,484,503,242,0,0,0,0,0,0,0,1282,6,0,0],[248,493,228,1088,656,434,232,968,459,1705,1321,674,423,522,192,1963,320,404,316,113,308,1143,38,1448,204,711,96,312,411,991,18,452,185,821,847,265,3078,166,682,452,222,198,933,442,1274,497,221,1889,128,0,1286,737,402,196,386,1543,3073],[67899,123153,16482,204540,156396,140121,131802,122640,94029,327149,290125,17597,171536,151781,149526,161222,54322,109173,193210,196225,357713,210281,125301,64597,177756,257716,138708,43295,201574,267624,44041,130412,132616,140321,235406,193392,108798,416137,98128,161602,168802,28251,169700,92449,47700,213267,124481,86963,142889,137229,890450,125487,141626,25735,56618,56918,97327],[352,637,2146,397,73,244,62,62,182,1007,572,2569,1837,928,188,277,154,0,0,55,535,219,0,4396,1053,1238,821,8,687,959,0,83,594,0,1478,984,52,201,300,802,2116,357,754,686,2051,1705,4926,3694,83,0,1970,404,1097,2222,570,220,58],[0,0,0,0,0,0,39,0,0,0,0,0,0,0,0,37,0,0,0,0,0,0,0,255,0,123,518,0,41,0,0,0,0,0,252,0,0,0,798,0,246,0,0,0,116,0,0,0,0,0,0,0,0,50,0,0,0],[181,407,1997,153,241,583,131,161,220,1306,624,773,337,966,159,3292,1505,1694,337,283,438,690,202,0,1349,1649,1289,884,2398,3216,0,281,563,145,1508,1480,401,1730,0,2614,1604,3309,496,616,1288,64,4685,649,169,447,4735,683,1555,1061,1547,100,57],[191035,356355,18583,665846,376793,223779,350967,320471,440779,1542715,1240888,79651,314862,204488,246020,428179,147004,218445,589518,405966,1285013,564892,191415,137595,615569,1003303,509637,55824,331177,490555,155231,438358,747227,326209,394141,371443,329977,833809,200897,406886,405627,105523,672121,365886,104560,688121,385505,214922,435293,266981,1659769,283957,271745,34489,146966,120791,249859],[24981,49857,4122,25194,14450,38416,25027,26352,40607,63167,36067,8607,33942,17580,25996,25185,6359,46153,17029,30449,85440,61563,70283,92744,22239,83893,20442,49336,105010,140615,49058,78325,35565,47711,95545,101504,39031,84367,63227,15528,28831,11709,51223,41389,22863,12573,22335,32426,13485,9359,126446,41010,39546,27428,25833,21871,20306],[175,0,452,0,0,0,134,0,0,0,287,202,0,200,24,0,0,0,0,0,60,0,0,202,0,0,0,0,174,0,0,0,0,336,0,0,0,0,0,0,0,0,0,0,0,312,395,238,0,0,0,300,0,281,0,0,0],[427,116,0,130,134,179,6,0,140,256,1326,0,177,316,569,1704,75,0,0,0,137,606,0,1206,148,463,0,0,197,576,0,402,0,763,0,0,0,1670,0,131,0,1817,201,0,1214,1181,0,192,0,0,0,342,108,324,61,30,0],[902,3719,1178,1994,1096,2710,475,932,100,6957,1488,594,1515,1026,827,4870,439,2272,606,322,1613,729,134,281,306,728,195,3848,793,1676,0,1203,1460,475,1688,2027,226,217,653,337,769,716,699,294,109,503,102,176,30,86,527,563,843,137,1077,520,958],[9861,21394,4152,35600,24447,24662,28550,34035,20980,80053,52214,6042,23599,40044,41218,56992,20310,20398,57888,19983,45901,82051,41739,29606,31693,32210,9074,21910,37330,27020,4570,14985,18884,52265,41803,41718,20702,32980,20469,30626,12409,2675,25937,8084,13559,45986,10674,9737,19339,25845,139045,28100,24040,3916,17288,19137,52611],[38,15,54,114,244,23,202,102,162,272,79,824,118,191,0,1370,640,554,0,83,112,26,0,74,30,844,882,117,0,167,0,30,0,0,126,23,28,32,29,39,470,0,71,363,576,34,31,400,88,0,256,130,513,209,0,64,51],[900,175,846,102,202,471,0,398,0,228,1376,1520,376,206,147,1094,0,0,70,1060,301,171,1502,492,801,1179,0,29,877,739,1529,0,606,0,637,0,0,3993,0,400,0,0,723,598,0,697,0,693,306,0,29,37,503,2207,603,100,0],[817,1065,2355,3751,1078,4381,2883,2178,665,5158,4387,3210,4183,2935,2728,3633,487,1833,3574,2899,2342,4475,2521,4816,3791,2500,395,23,1631,1514,0,916,1439,1835,4012,6225,6190,4148,564,3186,2566,202,1105,2643,5136,3015,4018,1599,2264,937,5811,1911,3272,3081,1178,615,0],[0,0,0,0,0,40,0,0,0,0,0,0,0,0,0,0,0,0,0,0,0,0,0,0,0,0,0,0,0,0,0,0,0,0,0,700,0,0,0,0,0,1610,0,0,0,0,0,116,0,0,0,0,0,0,0,0,0],[306666,589165,119105,1002868,664545,787050,376137,410048,618649,2014156,796684,273159,401159,416462,451844,555443,279796,242084,599326,553863,804610,873146,893225,1511632,467252,1147148,288917,544459,446836,794000,193154,402724,323526,796794,666092,425466,412725,621781,243025,326130,287635,70331,598426,313967,468014,734087,216261,140308,415658,416286,4138712,412704,279271,47651,413356,643641,1031685],[0,0,196,179,0,0,0,0,185,179,199,538,702,671,0,0,0,0,12,0,0,0,0,0,0,0,313,0,225,0,5336,0,0,0,0,1200,0,316,0,0,0,0,397,396,401,0,0,0,97,0,0,0,313,424,0,172,361],[0,23,339,0,0,0,0,0,0,0,0,0,0,0,81,0,0,0,0,0,0,0,0,0,0,0,0,0,0,50,0,0,0,128,0,0,0,0,0,0,0,0,0,197,104,0,695,0,0,71,0,0,41,4,28,0,0],[536,264,426,1034,180,293,564,597,1043,1091,1288,409,698,436,250,2151,29,0,1893,258,1447,937,2792,0,378,855,803,31,561,357,0,397,443,706,562,352,194,1190,275,281,248,61,1125,1982,1970,2325,879,82,654,571,2470,0,1908,24,411,0,0],[4927,4977,748,5299,6303,3049,4618,7289,8137,20617,13197,883,5552,3036,4790,9523,1043,3628,6859,9655,6187,8517,14328,4498,7175,10087,4201,270,7613,9215,0,9693,6634,6477,11985,13826,2408,11561,4232,7033,10999,0,11340,4668,849,4944,550,4837,2633,3198,45255,5369,8057,1192,1389,3441,10515],[4566,4277,1047,4645,2013,5754,3157,3920,4513,8162,6223,1347,3758,2183,1951,9708,2061,2583,3236,4071,4000,3877,0,202,5660,4789,3443,1971,6518,6432,2437,4235,2558,3053,4271,7007,7486,3777,3567,4371,3400,682,9311,1357,1181,4555,1391,3050,6753,1045,7065,6625,3141,1426,2591,1973,0],[37010,34951,5412,55539,42330,41979,25618,30097,21539,98837,58914,5044,29663,29381,35983,61330,21254,20355,50705,37620,38179,46300,57105,36931,41367,48036,21921,11053,34115,47795,12407,25469,19133,42543,48709,54345,35801,58691,21587,41450,24001,9834,20827,15277,16678,41908,20262,14161,33097,33012,339315,18540,27561,5252,36111,30566,59474],[67525,75983,15037,93990,66441,106307,61033,70866,114386,155253,166881,31356,71894,67987,63677,99396,60638,84441,88876,89170,113700,92706,126981,301903,59993,150210,66065,85978,182803,354296,13109,79770,55398,91434,166331,190530,44838,99613,128968,74490,82429,61481,64163,56009,53247,80269,50359,81452,38593,47978,452769,60899,94610,17164,81154,82946,182354],[0,0,0,0,0,0,0,133,0,0,219,0,2,0,180,0,0,31,0,0,0,0,0,0,0,88,0,0,333,115,0,0,0,0,0,0,924,956,0,0,0,0,0,0,454,0,0,1016,0,0,0,0,0,37,0,0,0],[109329,223292,11676,280627,183489,163726,188873,199701,137139,592621,776633,16509,137417,154324,156815,156017,66615,51022,287949,185878,365035,304106,155731,118298,184633,409220,134580,30058,265062,297430,51444,145443,192308,190537,266392,201403,110750,329580,106100,184092,215407,22450,215881,131929,37395,159500,101443,50727,156238,114178,763607,131334,216685,9868,37804,57500,155718],[83896,109890,7041,197117,137857,143704,165268,152047,107687,437424,590727,11573,151500,106601,104251,203257,96837,120219,191783,176682,396848,348749,202958,73726,159847,368287,155201,52245,273412,218141,40641,163973,156042,206472,298930,307639,86416,315783,36149,149206,202148,9116,148097,83865,26046,118771,91124,39136,169584,107075,754059,134202,206303,16290,30437,50651,88965],[2491,6572,1658,13307,8334,6318,3157,5506,9638,22132,20129,2335,6299,5960,8337,12335,1909,7063,6511,5886,16210,8848,10494,5927,6409,18684,3474,6585,14110,15938,2832,9914,5231,6480,8872,8014,9373,8704,1558,3911,6378,1236,10200,13370,5204,4327,3149,3210,3210,2759,37188,5002,9371,2230,3967,3467,7023],[31,410,762,0,24,101,91,182,172,255,284,252,0,144,9,384,0,0,11,191,0,525,918,729,606,492,294,0,1167,652,0,732,0,928,581,297,12,49,24,202,156,573,835,392,814,235,1455,996,0,101,0,155,1044,667,57,91,28],[4805,3387,2654,3181,2776,20958,6056,4179,18968,28888,4699,4468,2325,4668,5853,1754,4744,33885,199,909,977,3675,3208,10464,264,35515,5630,7880,15097,16843,16701,56633,30188,7989,11338,41689,28,9673,3359,978,36421,0,93480,31010,4028,1305,5461,19785,8597,6128,17115,15617,14357,997,7210,2155,241],[0,46,33,227,132,126,45,416,0,152,1567,0,169,299,418,1256,261,244,695,239,198,689,807,74,168,0,675,1304,254,284,0,345,27,318,1035,1742,0,156,762,264,403,0,0,70,0,0,32,0,19,44,0,636,199,0,33,76,68],[36374,91720,29432,20017,27068,65533,14123,16315,5084,33093,14485,7919,26390,23446,23056,44734,11211,10882,17802,15817,26361,26580,18546,31896,29602,13134,5581,1298,30420,32224,2211,7800,9123,18580,25780,31999,6786,29630,23913,17632,13804,13399,13869,33439,28027,29988,4388,6831,10911,3504,9372,11396,10635,4525,31371,14480,40215],[10974,14708,2974,20116,11984,27677,8473,10978,28687,27349,17991,6948,18624,18363,20239,14465,6397,11992,12961,12272,22419,21760,37613,64864,12280,33140,9043,11577,27249,43008,1098,12143,23322,22682,37460,36788,11021,42729,28002,9754,17102,8667,9019,8149,19078,8002,10273,7455,10213,9389,99694,12910,22774,4236,13026,9545,23726],[72,0,200,0,205,24,0,0,0,18,131,0,0,738,14,0,0,0,0,0,0,0,0,597,0,227,0,0,150,0,0,0,0,244,0,0,0,0,0,0,0,0,0,0,132,0,0,40,0,0,0,0,0,115,0,297,0],[6131,12401,6590,8347,8619,7944,4832,6551,7750,9529,11659,4779,4126,3327,5286,2268,934,421,3026,10307,6244,4783,20964,26261,6489,13718,7671,4094,13203,44275,883,11507,8065,5453,20008,10401,590,26033,29112,2118,11998,10245,9561,8295,36087,9278,7178,26715,3377,2384,111647,4327,9044,6841,5888,4478,12801],[252,1700,3592,1809,553,917,1911,897,427,1180,3087,1409,1982,1101,940,3900,183,1311,1624,1647,1539,463,245,2057,2889,4401,3691,0,5040,2329,0,1192,732,1725,2842,4519,0,3312,11107,402,522,2703,4537,2455,6117,1549,3680,2261,1112,1120,3274,504,2774,5186,2310,395,78],[33860,31405,13091,77615,53765,43653,64789,59987,25606,140353,182953,10318,79879,56876,53712,104428,28450,49321,95251,57449,102354,191288,77785,28609,92233,102910,48536,20354,65913,76952,2964,53149,44424,93341,128768,111366,43399,160697,25128,72863,77287,3389,48409,21099,20036,60532,22396,14025,53169,38170,155610,60041,61269,10145,17298,14416,19778],[0,578,216,0,0,0,0,0,0,42,0,895,124,0,0,0,0,188,0,0,0,0,0,0,0,247,303,0,200,150,0,0,412,0,462,606,0,0,0,0,0,0,316,0,216,101,2347,1357,0,0,9504,199,61,194,0,234,178],[189,0,1232,185,192,134,96,0,142,485,152,790,796,183,0,0,0,0,0,0,0,123,0,526,202,156,50,0,245,384,0,0,0,24,355,225,0,0,0,0,183,0,404,136,606,934,1406,297,0,0,4903,0,128,133,26,0,0],[432,3,0,557,45,64,3,0,23,22,0,14,28,26,0,68,26,0,0,0,0,4,4,0,0,120,168,146,44,0,0,0,0,108,9,206,0,0,0,0,29,3,0,38,391,0,14,0,0,21,29,0,156,44,0,0,0],[8297,9439,4333,12226,8713,9975,6627,6574,6913,25846,18188,3145,8233,6751,4293,8763,2414,7397,7990,5377,22629,13264,17821,10186,6583,13245,4308,11140,19002,14802,5260,9473,4586,11050,14042,13578,5685,9026,7069,5526,5225,2265,10509,6419,10725,5554,14686,6012,4733,5456,32281,4687,6494,1897,4755,6307,5573],[346,279,984,420,1149,1832,649,1165,1398,756,2067,841,1549,1924,2232,1485,772,2478,1572,2357,2641,747,768,0,914,1592,1479,328,2788,2012,0,202,199,1379,1779,744,0,1421,0,951,368,0,1373,650,0,383,0,885,0,596,5530,0,1492,0,481,1778,1012],[0,0,399,0,0,90,56,0,319,194,333,525,0,384,360,0,0,0,0,160,0,372,0,0,0,119,62,0,130,350,0,0,0,0,0,0,0,0,0,0,0,0,944,303,31,634,1010,645,172,114,0,183,109,340,0,130,0],[0,26,233,39,0,27,0,98,0,0,68,279,265,0,98,86,0,1147,0,32,136,0,0,0,0,403,12,0,553,402,0,0,0,0,0,0,37,0,0,0,0,0,0,0,35,0,855,12,0,0,52,0,0,491,0,0,0],[0,65,3443,402,528,202,156,202,547,276,0,9,153,102,0,32,0,0,0,0,4,0,0,0,404,43,0,762,305,0,0,0,1396,0,995,0,0,0,0,0,0,0,928,455,903,0,782,460,0,0,1351,0,606,1357,96,56,0],[279,44,1370,534,113,265,449,394,687,1057,446,1170,401,953,1347,115,95,398,208,800,51,202,71,1722,841,405,639,54,165,1154,221,39,0,760,1097,1228,0,293,263,49,0,47,1137,202,1851,303,1814,2137,116,57,847,501,127,2060,204,707,0],[5848,4576,8236,15122,9899,10112,6866,7995,2638,12761,17038,5806,12801,12031,10114,9709,3588,4823,12648,10812,11607,20604,16627,4845,9988,12920,11277,3095,17984,23111,8800,2468,7650,13403,22918,13086,20770,19186,0,13834,14175,1586,6608,6863,8645,10167,15204,11991,8162,9638,27564,10550,11432,9000,2340,5253,1350],[14723,17118,6072,43315,32054,34520,21446,23062,18894,59410,49698,6941,19257,18652,20817,31609,11786,7999,32636,25543,47193,40605,63386,41819,26274,41570,19105,4413,24263,32138,1868,20893,18223,26809,39949,45573,17568,46944,25538,25542,20971,17637,19750,27429,16676,29074,18148,18903,25419,23316,116795,15471,30523,3308,11846,14769,20552],[4554,13832,4752,340,1829,7554,20,97,643,565,1946,514,1908,1408,636,529,617,1240,1415,1149,16868,625,266,1711,1009,4023,4188,101,974,17225,297,97,8482,29,454,2391,303,10475,3986,270,236,98,1351,6592,12177,573,1787,3750,349,298,48087,4543,3612,2133,380,215,844],[28842,37582,8188,73373,51165,30521,29323,34560,67699,136578,90850,25132,41091,51074,41489,42660,11260,7373,47049,31619,81014,68359,52317,84936,43977,71920,33488,8798,32219,51632,12205,30598,38951,54815,55169,56415,23136,74321,47872,35446,23199,15881,61115,29189,32806,63961,25335,32131,35653,25066,286795,23242,24801,4069,21637,18172,18061],[342,12,755,203,0,0,0,0,0,0,0,0,0,0,63,0,199,338,0,0,0,0,0,0,0,0,0,0,6,274,0,0,0,186,111,50,0,0,0,0,0,0,435,216,1599,164,3301,1008,0,0,92,0,266,739,33,0,0],[980,315,317,535,0,376,750,1560,369,1662,1077,611,1144,598,1176,1410,342,2998,1723,1005,1010,1000,0,0,780,2091,1824,0,1871,1465,0,1368,398,399,3071,1428,0,1565,0,144,0,0,1725,0,20,606,219,428,200,402,0,1011,893,201,373,935,0],[12151,21299,6474,20078,15852,29030,13692,9715,21922,36574,61472,6443,33654,29068,28313,28533,3513,10305,17148,38694,21138,28061,40087,19218,31781,61128,35991,2066,70345,74212,1661,11577,9593,16919,66313,52986,25167,33102,14106,23188,15088,9062,18584,26451,15260,22534,25777,19562,37097,39652,92006,28339,44025,14371,7912,10990,8273],[4182,1615,1059,356,2911,4308,428,1178,74,924,1674,250,1505,534,1758,73,88,114,0,587,1438,267,2219,2961,335,4162,3461,920,16221,1053,2889,876,146,3202,8645,12147,555,461,1269,24,1307,3870,794,627,843,0,428,0,696,591,2522,1686,12615,0,0,1513,105],[0,128,2018,477,636,302,200,469,0,606,304,760,23,186,146,58,248,0,24,342,199,0,0,629,61,230,162,0,957,693,354,0,0,0,718,806,0,0,420,340,0,0,179,800,3205,86,1473,198,495,0,3771,1123,245,1154,376,0,0],[146,0,62,0,113,522,404,0,0,554,500,1501,342,779,0,648,36,1014,46,251,762,532,0,0,0,529,790,0,825,48,0,524,0,386,701,0,0,644,140,61,730,764,304,258,550,194,0,957,0,0,0,0,519,660,623,151,348],[369,488,131,696,535,268,138,36,1087,1233,313,239,217,403,495,20,0,0,52,232,1119,1129,410,3557,488,545,251,153,192,1176,0,250,403,902,2097,1555,0,256,3869,258,0,640,319,80,1035,311,0,2591,269,175,8462,237,355,150,957,538,685],[80104,655767,54045,75251,137153,737003,30157,42984,3456392,589449,207430,58308,57049,573510,699395,30387,31672,458723,39823,608678,315217,45256,403018,359253,65825,3796248,940540,156897,866079,6919073,123131,412380,2962964,54310,165759,397203,28737,713499,355973,35840,660819,326853,684625,254251,722562,36961,127999,92894,80482,67904,3256628,348059,97381,1124422,16986,109588,668225],[504,2137,2977,2727,373,1531,1702,2950,1357,4094,3587,3033,1861,1186,1145,3494,2059,622,1982,1185,1622,4173,4032,5550,2202,1312,494,612,1906,2625,0,953,1266,4638,2434,4415,3369,882,224,1031,2419,3492,3673,2176,7171,2590,4632,3117,1048,141,14137,1338,2772,3867,2089,205,434],[567,223,0,701,525,714,278,631,132,1148,1317,129,1358,773,745,603,345,662,656,1065,1979,2105,1876,1099,1380,1510,582,28,1914,1222,0,61,1990,2863,1022,44,0,1217,0,390,1934,0,2585,930,0,259,0,0,128,1206,4879,1043,500,0,29,572,2844],[669968,1552194,80165,865153,501088,1097895,1092261,873038,2954821,3553470,2453785,200177,886984,741026,692822,5876640,3141521,3183499,500114,576905,1847815,1108924,1491011,1062279,915680,2495670,1125615,1189812,1222595,3066429,308269,2295961,1859937,1214505,1481265,2816141,1467114,4673399,864682,874310,2104723,461146,1717525,902633,540011,509932,358603,394603,852723,836592,6123079,940651,1006628,219409,375384,642541,646976],[149,681,0,171,126,357,183,273,68,371,0,173,0,310,180,584,31,139,209,2,0,202,26,642,206,179,0,774,0,128,0,63,0,111,136,639,0,57,0,0,309,0,0,190,0,139,96,605,68,0,0,0,0,0,224,26,0],[0,595,290,656,1062,1214,72,1221,989,2388,673,52,974,262,201,522,57,40,0,0,1059,288,98,588,303,849,80,160,982,306,3735,4413,1521,2345,2094,3036,0,2686,3480,0,0,1832,2734,1085,231,0,675,42,1579,0,826,703,1333,87,584,6,0],[73584,139107,20267,140479,102613,142607,80302,84351,222282,397413,267242,25848,107416,105738,101451,249421,103187,131433,96985,109578,167686,198540,140227,106376,113451,198456,71957,257815,186465,192570,75255,263814,133537,174672,230710,219092,76923,238855,119047,78444,133127,9096,307796,86514,37378,119084,75943,59342,106505,81224,475460,137980,105585,17753,69197,68881,117228],[0,0,370,58,0,0,0,0,0,156,97,84,73,78,0,718,0,234,0,128,0,38,0,349,0,0,72,146,374,0,0,212,0,39,130,0,0,84,0,0,56,0,380,344,322,118,0,0,0,0,0,66,322,44,71,0,261],[99791,226636,29903,283277,234215,145801,175930,172547,161957,780072,362517,56306,96925,101173,104076,198420,82368,57308,303555,123301,255841,468292,98778,107316,338775,265280,63424,67731,129516,116527,83190,209226,54296,382195,193416,227604,155210,195006,73433,160756,146767,9262,470239,151310,75219,532006,103408,46299,112547,66035,690476,128096,141876,11232,100411,42663,71467],[1741,970,188,4906,3533,6489,2713,4321,1226,7403,4184,2077,5191,5420,5534,12630,5933,1167,6537,2305,3816,6032,3951,3663,5629,3500,3990,710,6441,8922,5626,2307,5290,2769,6195,7040,3619,4782,1613,6034,6288,36,1969,482,1402,6173,2289,624,3891,2276,8537,1410,4996,3224,1837,2123,3356],[452,0,328,0,144,0,0,202,236,96,17,415,124,0,0,0,0,0,164,0,917,0,0,0,0,0,278,0,573,0,54,0,0,0,208,0,0,0,230,132,0,0,460,884,0,187,667,0,0,0,0,289,500,806,129,178,0],[24939,26972,11126,35768,33071,42900,25060,36809,23867,78302,73769,14419,57801,54151,48868,44900,10519,32109,31305,29975,66516,45719,47736,27755,24177,58907,30692,15825,86156,76481,13297,30363,29113,49969,108592,130453,16826,104080,42198,25897,33888,2709,43957,17990,18730,26392,39307,29881,35138,24276,215599,26684,47569,9990,23677,19005,20606],[11931,1162,5926,12451,3610,7302,8749,4722,805,31627,15190,5579,9771,7332,3019,7754,4007,3338,23981,4387,1531,41246,11794,2516,25093,20060,18506,2489,30973,30903,7165,12001,7572,16676,26592,6363,7933,11897,6188,15928,26533,645,17478,10059,4782,7173,7262,10920,9015,2184,6194,69685,11437,11313,6501,4741,47],[44227,63530,4444,94740,75985,87615,34747,54937,18123,223370,70481,6879,53590,44983,39293,152851,35685,36979,87673,59340,96183,69888,25567,24605,31927,38802,10916,28824,38116,48016,2939,22394,19181,38867,67139,61765,34820,78845,16546,53291,48917,4617,55544,27405,14421,43738,15888,12744,33492,14262,118281,28041,54494,5580,55128,37429,23412],[167293,252482,11349,369963,275029,379336,285160,267575,159975,866498,525977,32906,317722,301885,312960,409876,165531,353396,362298,314574,666850,637822,374845,212419,340334,1126606,366200,152742,555453,757304,127691,369235,430378,352990,645105,605033,234962,820331,133333,257569,409006,42640,436474,170227,33640,384647,145737,96242,268385,174554,972018,231272,309829,29626,84221,131917,190504],[65535,82089,10458,111568,94864,164137,81028,72409,65155,266924,176540,19854,114140,169260,163235,177646,82892,233125,82751,76889,227943,165896,174810,103714,96287,335912,130651,138951,218241,327125,78828,155172,144836,124436,227669,278075,81455,384890,110003,73544,181273,16085,122273,89862,29515,130972,73326,78714,60471,102849,529856,88840,111813,19379,58087,67760,91687],[0,0,100,0,0,0,34,0,0,68,0,76,0,0,0,0,34,0,0,0,0,110,0,0,0,0,0,0,207,0,0,0,0,427,0,84,0,0,0,0,0,0,0,373,477,0,194,0,0,0,0,0,257,20,0,0,0],[541,335,615,774,670,1474,903,1009,1246,2777,1970,1710,1365,497,622,1640,227,1010,707,399,1609,731,113,3152,126,225,647,1690,967,1750,2125,346,569,1371,2085,4507,148,1281,2839,610,441,156,997,778,1136,848,1787,1048,372,243,13422,443,573,1088,1604,2031,49],[1744,939,317,2,1771,387,418,2283,598,2084,3074,726,753,681,2418,0,28,9,59,0,436,81,3741,1490,0,4853,4104,116,22773,1730,1307,523,190,4208,8796,40780,0,0,126,58,2028,45,654,404,802,0,67,607,106,0,4242,1477,10639,370,0,15,0],[35,162,636,0,0,191,240,0,212,0,368,197,0,98,0,0,0,0,0,0,74,570,0,3,0,150,380,10,91,72,0,0,0,0,0,170,0,576,209,0,0,0,0,285,419,763,1512,0,173,0,4387,0,186,386,0,0,0],[21836,10570,10333,73293,52257,44208,42141,45238,11947,61485,132833,7647,57853,33279,37128,98538,26570,55332,56018,38944,65098,70936,51136,14805,47798,53580,47301,24691,69148,71575,7561,32844,25801,45111,108441,92318,32845,79826,24113,46546,54575,5699,37052,23638,10255,25935,33055,19709,44466,30260,52365,40204,36371,15123,9484,24033,21532],[0,0,363,0,467,258,0,240,274,532,0,1179,0,168,117,0,0,182,0,0,0,0,0,0,997,0,0,0,0,1612,0,0,0,0,256,0,0,0,853,176,0,40,937,462,1431,152,829,0,10,0,1884,202,141,80,0,0,0],[6430,2471,1485,11046,10082,8417,5283,7357,1155,8878,24331,1907,16673,10991,6557,23097,8496,16453,7808,6180,16269,11655,9524,2349,4666,13650,6031,10189,22761,11962,801,8020,6652,11738,25533,16039,11369,31742,17333,11237,10813,1533,4558,3511,4007,5691,14586,11191,6388,5694,17248,9220,9682,2234,4719,7194,3667],[64482,77661,10780,93666,68404,90769,51238,61733,62214,158555,138291,16950,81154,68466,66773,102146,49075,75349,91755,51672,96398,110538,113995,87133,77633,127202,54396,47433,120163,199188,27991,49115,78696,93957,103826,109169,60285,161508,75574,55030,70706,27959,47886,28659,34755,112152,39474,47794,49803,39268,401002,53363,71649,15584,84721,59691,91547],[167,73,1896,552,86,155,153,503,658,152,1585,1952,362,500,401,0,0,0,219,162,0,676,0,2828,802,420,1973,360,1276,453,1762,0,0,1282,800,675,0,120,0,0,90,1411,2219,809,2837,0,2114,1480,198,603,3979,800,198,1036,170,446,0],[14785,22274,9430,8331,7130,12689,2775,2940,2055,9478,4421,2021,3067,1343,636,3185,1405,332,8142,8825,3300,4413,24994,7560,7801,19980,10566,1148,8343,10448,0,4719,6122,6050,14627,22479,10883,1925,2026,2895,4258,2226,5590,3497,6748,10120,2689,2523,2369,5334,14833,4873,28859,3427,2974,506,0],[106,0,790,31,86,0,0,0,0,0,232,295,0,0,397,0,28,0,0,120,0,0,166,94,0,0,0,0,163,159,0,0,94,0,0,0,0,0,0,0,0,0,0,0,400,400,86,0,0,0,73,0,271,471,0,49,0],[425,1416,353,5627,5460,3417,95,0,871,6169,6586,643,2312,2649,1269,603,600,1164,7165,597,2150,1202,861,1300,4669,2181,96,0,547,407,0,1125,0,676,1748,0,0,5736,0,3714,100,0,82,0,933,7442,3648,859,0,36,10,95,171,254,415,580,3138],[0,0,0,0,0,0,0,0,0,0,89,134,0,0,0,0,0,0,0,0,0,0,0,0,0,6,0,0,0,0,0,0,0,0,57,0,0,0,0,0,0,0,0,0,0,0,0,0,0,0,0,0,0,0,0,0,0],[146795,264608,31256,228914,176778,322649,120745,173777,271531,493830,419712,66632,178727,198756,228590,219191,82805,143019,177551,177293,269251,267822,376500,430415,147593,465289,160391,260389,380622,849023,86629,183655,330883,282701,370618,487632,147797,431234,294433,121942,280987,106548,189218,109068,123233,120037,69770,97261,115094,151070,1390380,155187,242300,74294,119375,180067,282578],[40,514,776,265,29,375,0,0,40,328,124,29,290,69,198,274,61,26,0,0,0,29,80,578,0,274,0,57,130,158,55,40,0,0,58,96,57,0,324,29,0,203,64,798,55,81,117,439,26,25,186,0,135,369,40,28,54],[0,37,144,31,1,319,0,0,183,142,0,0,0,207,132,257,71,587,2,0,25,0,0,740,0,0,0,872,0,0,510,158,0,0,361,140,0,185,59,0,39,1,285,75,56,368,0,563,0,66,387,338,124,0,0,53,0],[24,76,1309,412,0,202,96,0,240,138,804,711,437,141,200,7,0,0,97,1,0,0,0,0,53,64,0,0,639,0,0,57,0,1143,582,445,6,0,3370,0,0,848,1215,628,945,857,1806,844,920,0,28,656,251,547,435,178,0],[29,47,20,269,53,30,124,14,0,165,0,49,0,65,0,893,20,162,24,24,200,0,0,0,0,0,0,49,0,0,0,0,0,29,0,0,0,0,0,29,0,0,0,0,212,0,0,0,0,0,29,0,0,0,82,24,25],[2269,641,2135,170,954,1449,409,765,1186,3318,3131,1649,1294,784,605,965,0,528,1308,667,1290,720,1094,2062,583,1131,3098,504,376,1010,0,0,200,884,602,570,0,1150,1339,1724,0,0,1466,2592,2887,997,5139,690,1008,0,188,1175,1936,398,1411,402,0],[0,0,0,7,21,0,0,0,0,0,0,26,0,0,27,46,104,0,0,0,16,1,11,454,0,0,0,74,0,0,0,0,0,0,22,15,0,0,2,0,0,0,0,0,0,0,0,0,2,0,0,83,0,1,0,0,750],[36474,3908,946,1053,7284,35302,1092,5018,329,4667,7159,4689,6536,613,5934,30,166,358,0,1826,3518,185,29935,7515,278,21251,25003,12265,74088,9722,5213,1367,642,20527,75331,136027,14,756,2230,105,8430,1169,2274,2407,2633,1209,234,2743,578,162,3268,6821,49855,1478,53,1096,0],[435,155,2468,343,222,110,299,358,236,1010,575,2531,1132,225,117,0,58,54,350,0,103,200,100,162,730,1221,2333,0,2302,2603,682,0,385,380,1700,375,0,34,0,140,840,2233,2902,3315,4754,531,6643,1670,87,178,268,975,1033,4213,139,65,917],[0,194,486,0,68,65,190,164,0,0,116,1276,126,202,0,0,0,0,0,0,0,202,0,0,0,64,170,0,617,473,168,0,0,0,70,493,0,139,0,0,0,0,62,391,637,0,4017,1506,0,202,1913,0,380,1026,792,0,0],[707,1329,116,2543,903,1462,849,726,793,7298,3885,386,912,1819,771,7770,1374,2563,1216,1017,2383,4118,786,1276,1449,1937,338,6486,1602,792,0,1385,686,1727,4456,4075,810,726,24,1456,2622,0,658,775,0,1097,0,0,1224,742,7805,540,846,22,342,707,854],[7994,5718,936,13692,8849,6905,7284,5868,3849,26046,22833,1644,10314,7637,8149,15486,7695,11517,10624,8678,8585,12901,5287,6038,7868,8345,4932,7892,8018,15142,5404,4519,4059,4827,14713,17704,10008,18983,2985,10532,11614,0,5070,5944,1613,7432,3386,3962,2472,6424,42188,5098,9035,1787,6220,4976,5662],[0,334,1023,0,0,152,404,146,0,0,297,1827,0,27,0,0,0,0,0,0,0,0,0,0,0,0,0,0,0,336,0,0,202,98,604,0,0,0,0,70,0,0,182,0,549,87,1183,0,0,0,201,0,194,164,138,0,0],[427,490,652,851,457,638,276,126,804,783,1874,977,100,416,1055,167,472,33,402,488,1435,230,0,98,511,999,398,0,620,598,32,482,0,12,515,637,0,0,2641,0,66,260,594,1853,2262,98,464,180,603,692,2428,129,354,13,150,130,576],[0,114,0,0,0,0,169,0,0,0,0,35,0,0,0,0,0,0,0,0,0,0,0,24,0,35,0,29,29,0,0,0,0,0,0,0,0,0,563,0,0,0,403,0,130,0,0,0,0,0,111,0,200,55,0,0,0],[99,103,482,122,45,45,96,8,307,58,389,483,0,0,0,0,0,0,0,0,99,41,0,3,0,0,0,0,0,27,0,0,0,0,393,487,39,0,0,0,0,38,105,273,148,0,772,0,40,45,0,0,132,0,40,0,0],[494,5171,1940,926,1948,3861,1468,1573,2320,11163,3776,4619,2580,3288,3498,640,427,3380,211,3550,7994,167,2764,884,0,9208,3526,302,3026,6346,1453,3088,479,1995,8242,10769,0,8259,2303,3096,6315,1413,11084,6702,3902,1004,9388,8206,1564,504,13495,4623,4018,4217,639,1208,2937],[263,935,5911,567,675,1004,520,663,3544,2187,1038,1139,844,1009,1106,3472,2448,834,180,265,954,133,211,4039,1609,4378,3683,1820,1702,5483,542,0,500,1319,2198,3742,80,1176,417,0,1577,162,1732,1729,3866,2010,3290,5049,0,643,15025,1246,1052,3821,833,311,0],[138,0,288,1326,877,913,158,75,201,153,180,101,73,136,0,845,0,46,93,0,119,142,0,0,0,69,0,377,94,595,0,0,0,0,270,1039,0,69,0,23,255,1493,376,430,336,0,1300,0,0,0,0,23,287,480,351,23,163],[9861,15456,4230,29602,26262,31933,23727,25011,19569,56746,58911,3435,27175,29755,23521,42463,14533,18697,49264,34539,33551,51288,29629,17083,34097,44959,28733,7654,36905,47089,2161,30956,26136,25694,50031,47097,1340,35956,10690,32752,20822,1744,16517,10203,8374,23496,10133,5973,19105,14696,109554,18938,28386,4877,8156,5324,20845],[92478,155695,31694,215205,161215,160729,117602,129214,170030,610471,493914,33834,126229,134299,129097,281310,101108,122364,259949,322739,321494,323596,232511,185177,189572,349522,154579,73856,231520,192123,68841,243655,185682,228087,327330,277939,136801,309835,92869,184034,213652,35690,282451,150170,108917,226511,98942,65624,188377,147735,509778,97299,133111,21172,71374,88201,76657],[165,490,198,310,0,202,201,358,598,1340,310,2260,202,202,59,217,404,0,559,202,399,900,0,1386,0,1003,444,28,555,1005,0,0,0,173,1029,88,0,3969,354,0,0,0,393,1950,114,730,2880,604,1008,762,3397,303,508,2731,685,511,0],[0,0,0,0,0,93,0,0,0,0,12,0,14,0,21,0,0,0,0,0,0,0,0,0,0,80,0,0,0,0,0,148,0,0,0,80,0,0,0,0,0,0,0,0,0,0,405,56,0,0,0,220,0,0,0,0,0],[6951,19270,4121,14156,14659,28143,11980,19015,22574,49842,45994,5223,21536,19522,20830,17603,4969,14034,24134,14310,31950,32318,20239,21344,18459,44755,18951,15185,44781,50736,7128,17278,24186,33918,35665,53859,20509,35165,22825,13303,31491,4347,20162,11811,6994,16418,4709,6539,12462,20060,95801,17411,30543,3886,7547,8861,12561],[185,624,314,278,351,506,104,349,897,1111,1998,72,180,74,224,1048,347,402,165,558,747,158,0,1266,217,341,0,0,971,1662,0,922,579,402,1469,591,0,495,0,431,518,0,186,58,0,605,493,692,185,181,6163,190,19,0,0,370,127],[354,387,751,623,369,1001,216,326,33,1648,503,351,152,230,58,2298,187,976,148,58,399,316,29,76,48,59,444,2073,157,496,29,88,29,67,568,301,81,54,0,147,213,371,88,14,683,212,78,0,0,33,2588,59,31,406,813,419,672],[55486,67158,6726,159217,121154,133962,104243,89966,99262,433409,340597,18228,120010,135241,142087,212344,101829,156548,121951,136311,295870,201347,100930,91487,132104,435770,166378,139616,280234,447006,19768,185984,592932,119607,222664,178647,163731,348646,36238,136497,289218,34650,139236,65224,21942,67842,54599,38785,97445,65945,495940,107689,83061,16253,30513,50660,63406],[37647,38482,8962,76709,55196,40903,71437,56633,64482,155921,235715,8262,72798,78847,86154,97758,45116,60842,89116,77146,137020,131650,80630,49955,59921,112126,31219,21536,95187,159457,14310,68044,92428,70607,80698,87165,46250,134141,11741,64688,57693,4765,60655,31218,7704,48277,13375,18947,66808,49589,175649,42305,56766,7845,11932,27201,87986],[106920,194861,16977,184880,121868,134310,103407,121191,68841,326659,580871,20571,155019,105725,103125,229952,98014,164432,142784,104297,439280,210472,89786,75222,132448,174603,76598,60533,170254,210476,45288,177667,87337,154598,226232,169565,89563,326553,63541,113376,136603,21636,115569,71076,31155,105114,56766,53923,124016,105374,408548,106181,129306,19613,51142,50390,88274],[309,655,1452,281,130,113,0,209,196,384,578,395,95,886,148,460,0,142,0,0,139,0,0,1687,0,156,42,0,96,586,0,0,605,0,975,620,0,0,1156,208,351,0,620,356,395,472,1047,202,0,0,0,463,330,761,588,65,132],[42,60,1004,0,0,345,130,202,64,196,457,201,114,132,0,0,23,0,0,0,68,13,0,0,306,991,0,29,272,55,0,0,0,269,165,0,0,0,388,178,0,0,454,101,28,136,2031,467,0,0,27,202,302,796,23,14,0],[1639,1260,10387,1173,588,2363,1731,882,1233,2118,3194,6340,3199,1964,2210,1157,447,137,629,117,514,37,495,3267,1683,4630,4650,325,2963,1622,2807,776,100,1798,4839,3895,26,39,7692,463,1261,3214,5863,4192,8798,771,8095,8963,2482,1145,9781,2973,5760,8711,1739,1434,408],[279,1067,3844,219,495,1735,435,864,1541,2809,1554,1922,1203,327,638,649,268,0,848,665,376,553,906,2632,450,1729,1932,248,750,1832,2934,347,1001,1003,1947,3157,74,558,124,175,200,5142,962,2064,2841,725,3833,3788,700,122,141,1474,1114,1784,1917,353,113],[3026,10650,2328,7527,8302,10784,3663,6155,13356,27999,41808,4229,11203,12168,11188,6241,1524,10223,11442,5904,30506,7275,8801,14893,4748,12464,2868,1362,16365,17888,4862,7542,3136,6583,14412,24044,404,38894,22362,7488,16798,3839,5752,5366,8950,3706,9347,5131,5633,5266,41293,3636,9528,2275,3753,5581,2064],[13749,24403,7906,22917,19147,23199,19864,15803,29413,44501,29613,8356,21553,19775,19122,38123,17475,17219,22529,21788,20726,41805,26753,55378,16740,27896,15428,25314,21747,39638,7568,20362,22592,21032,31409,48545,19204,26989,17837,16687,23015,9992,14512,9361,14797,24550,8794,9709,15777,12602,104395,21549,18363,5631,22436,12470,12007],[2570,2562,1963,8424,7712,9469,3451,3809,2017,16058,9312,3266,5854,4712,4319,23588,10893,7951,6347,5191,10857,7391,3282,11002,4084,3467,5251,5510,5452,3920,4272,1707,3702,3467,12014,14768,4087,7234,14546,3900,8264,2697,7469,3284,2895,3680,4014,10307,3278,2297,10229,3796,7055,2355,4550,2858,536],[0,171,238,445,197,213,0,100,0,1457,517,0,335,177,194,349,195,212,330,178,286,28,70,0,0,357,220,44,121,290,0,51,0,113,385,197,0,106,21,0,985,84,131,45,152,116,197,10,66,30,124,20,30,86,380,0,59],[202,202,603,264,199,572,404,0,0,615,1029,1209,0,375,41,81,24,23,404,0,0,1395,0,154,243,920,2411,26,1473,475,0,794,1595,620,284,840,0,202,25,0,401,6719,776,1185,0,0,45,806,0,0,0,184,0,1107,0,600,2010],[3885,10801,2280,26811,12268,7485,11736,10363,5605,34076,29220,2185,9807,13077,18413,12170,3284,2682,14546,8587,68008,10281,10825,19056,11214,20153,5462,101,13231,3662,3415,4736,3285,17512,14736,2908,18336,11486,0,7249,6393,3829,8652,9351,2469,11207,8044,2969,19465,14148,59911,5617,7048,1499,2894,4194,9346],[75,0,370,0,0,0,0,0,0,0,0,75,84,33,166,0,0,0,0,0,0,0,0,0,0,0,0,0,33,0,0,0,0,0,53,32,0,0,0,0,0,54,0,64,121,166,0,160,0,0,0,0,0,64,0,0,0],[0,297,1413,36,349,526,0,487,202,426,578,1007,522,0,357,109,0,57,296,202,830,0,0,1798,112,0,201,0,714,1121,0,197,0,0,0,0,0,883,0,93,1264,6288,1519,0,2142,154,1476,1945,399,0,0,946,249,510,9,0,0],[0,126,29,198,114,10,166,27,0,147,43,0,248,0,190,23,32,31,0,73,181,46,365,0,0,337,45,0,36,197,4352,144,0,0,218,392,31,0,0,0,0,0,805,6,370,0,29,0,45,25,25,0,161,169,28,30,0],[0,0,14,0,0,0,0,0,0,0,0,84,0,0,0,0,0,0,0,0,0,0,0,0,0,0,0,0,0,0,0,0,0,0,0,0,0,0,0,0,0,0,0,0,0,0,1123,0,0,0,0,0,0,0,0,0,0],[300,788,1856,1461,1197,457,915,841,466,4608,3171,1194,1985,995,1302,1738,819,939,932,1234,2767,4653,112,0,1426,2181,255,225,994,2958,0,1357,137,526,2875,1555,473,2287,2620,463,656,94,2273,1341,879,1282,2877,1056,3651,597,1342,1807,1277,941,1683,446,124],[703,535,1929,1403,759,1224,1065,709,666,1521,2797,1604,1967,1876,1388,3985,356,6275,835,988,2077,893,585,1642,1582,2506,1725,54,1770,1941,0,0,78,872,2258,836,0,412,788,543,1422,26,426,1485,2830,1871,2958,587,428,2252,7087,1663,2397,966,480,426,48],[1802,1448,8478,5121,2896,5355,2398,2686,2049,7529,8050,5734,7332,6714,4700,3992,1488,3951,3167,3594,4411,3621,3257,3273,3056,5942,4694,1056,6061,6466,5963,2876,4435,3101,10113,9117,1370,4413,7937,2204,1737,2369,5185,3677,12185,3058,13828,7267,4232,755,14843,3070,4330,11527,1464,1610,585],[11533,11674,6841,21007,21217,34856,8171,11092,18136,34274,20361,9476,16962,12388,10995,22670,6299,13225,12399,9447,19566,14468,15964,39239,10934,19664,16877,8762,36594,40575,4761,12205,12641,17183,29242,43862,16323,24951,28471,8904,18822,8179,16256,5807,15178,12219,12511,15703,14929,5539,71819,12901,12204,4883,11653,17942,10349],[0,818,34,651,431,112,205,980,311,931,492,0,1632,167,794,17,0,0,627,488,475,2615,2704,0,861,1264,596,0,2465,878,0,1178,316,346,1895,813,0,699,0,0,0,0,1618,166,0,256,0,514,1704,0,31,278,341,0,0,0,0],[0,0,24,147,0,36,0,5,69,5,61,44,2,10,0,16,0,0,0,0,18,2,440,0,2,210,0,0,0,20,0,136,8,60,1,56,0,0,0,0,160,0,100,83,260,0,0,0,99,0,5,0,28,0,42,6,360],[2592,1865,654,5157,2988,4826,4114,3203,1900,8361,18698,896,2700,3336,3092,3037,1340,495,4946,2162,6191,9669,3213,3321,6499,8591,4901,603,7800,11325,0,3708,4088,5326,7613,8041,1402,2523,4825,6619,6547,1870,2801,1988,1983,6543,1593,824,3454,2949,18102,1942,5722,914,609,934,83],[492,1685,3195,1909,1186,1459,2306,1466,1038,867,2288,1560,2036,1603,2155,4292,739,235,1005,652,1898,1760,0,1088,2601,2945,2800,4205,2472,5364,2652,0,799,990,3785,1507,4778,144,1375,2240,1402,0,279,2127,9019,1145,4568,4197,2280,1245,14993,991,1147,1058,373,1669,674],[0,0,0,0,0,0,0,0,0,0,0,0,40,40,0,0,0,0,0,0,20,0,60,0,0,0,0,0,21,0,0,0,0,42,120,463,0,0,0,0,0,0,0,0,0,0,0,0,0,0,140,60,0,0,0,0,0],[473,0,232,2137,496,328,504,319,0,1446,1813,0,413,749,667,2249,1762,0,1514,624,1746,1257,3703,0,1437,2225,250,5812,2127,453,0,173,255,2838,201,2930,3823,0,0,564,0,28,3887,1877,311,405,98,756,654,639,0,651,1194,0,58,84,0],[240,239,148,1379,128,1073,708,330,188,1911,1465,276,719,188,404,746,885,1974,233,1989,494,202,2275,1064,1152,934,407,3476,932,1193,0,133,28,0,240,870,0,176,0,147,716,0,602,166,0,25,1646,207,207,0,0,471,78,483,196,636,123],[44,1,203,0,0,184,0,0,187,533,174,108,0,0,0,0,0,0,0,0,265,0,0,0,0,0,452,0,0,16,0,0,173,0,368,0,0,0,0,0,0,0,0,95,0,0,548,0,0,0,0,132,0,0,0,35,0],[530338,1360646,126405,1642728,1020160,611906,348628,410248,384446,2650960,4533344,481952,512408,499502,450979,663179,199280,182623,418432,732072,1290113,509069,385034,339901,646273,1008569,339488,465031,745011,571202,144350,415851,499142,497404,1055820,672279,458291,1559852,156611,273489,726633,87191,325561,356343,299445,707963,435893,235177,874213,701464,5895579,322146,281919,37521,207917,342527,590604],[309260,248682,31366,298962,213886,133675,427689,378904,185575,1038746,675828,60020,93614,67907,68738,699745,187466,270182,487081,313952,246867,1242054,694546,487089,896581,1169827,268826,610311,882354,395239,393226,1088118,616711,1698932,1556389,887306,366750,712794,330147,225147,515524,48258,1386636,488770,75809,508987,183179,141473,515179,349252,704167,705997,479797,42284,233825,115078,93687],[1578,936,4653,566,0,1963,977,1130,1353,340,1202,2761,945,1690,473,576,1286,384,140,202,46,1291,284,1130,82,2904,941,0,1706,1146,1837,202,0,207,2045,3723,0,52,905,0,434,1749,1012,908,4045,1256,6743,5917,404,971,6289,1779,2322,4288,183,534,28],[102115,104025,3738,153552,106935,67838,15181,34961,102387,342334,544790,29904,54355,87553,109327,96898,23510,37564,59657,544724,373415,10620,45134,14340,59526,435039,46368,30632,124344,921,8325,50099,17023,8319,67800,29745,50781,142869,5210,26157,278366,8733,286085,63077,23012,27856,88561,37101,34399,29994,306952,27458,14873,771,6024,22636,26469],[19273,34174,11808,24803,20879,32946,14599,16830,40074,109756,55455,11982,43678,12442,13456,26198,14844,14636,18484,17863,62634,42025,29959,29798,22771,43018,9141,7391,47447,80851,21230,47407,24641,72921,48836,43188,35272,59778,23152,15603,18514,10937,56757,16579,26266,23399,15946,16125,15187,9308,82614,16835,17959,5795,12673,12046,26852],[0,98,356,318,0,412,0,0,486,947,531,0,0,40,158,114,0,0,0,0,0,0,0,796,197,6,129,0,1142,730,0,0,0,0,280,272,0,567,0,0,1408,0,174,323,423,0,1240,59,0,0,1002,0,482,585,0,0,80],[368,149,458,127,202,531,250,70,2000,529,184,1174,197,811,997,0,1458,50,0,0,233,312,0,0,919,119,200,0,579,4864,0,175,206,0,1179,3247,0,12234,271,0,0,0,0,0,2159,377,1386,666,272,120,0,1424,379,394,0,0,0],[200011,295159,67519,457629,312617,408367,187539,232233,422285,885047,648091,114122,259905,286387,278154,302051,125602,196865,275115,291674,500335,405099,589076,1008305,195930,417167,187698,347027,361562,494304,135264,277980,194257,355300,578265,707142,218451,552000,472503,172106,246616,115852,316520,313280,228839,288662,220449,236938,194511,197408,1641270,198510,325285,50630,214816,256006,422261],[465,0,608,97,0,58,97,0,154,647,102,795,64,5,2,63,53,418,0,70,0,169,0,84,0,314,149,0,161,2,0,4,0,27,397,126,0,185,344,0,135,0,179,120,926,163,294,234,6,0,0,257,200,438,0,0,0],[0,0,0,0,71,36,8,0,9,0,0,76,126,189,78,0,0,0,0,0,0,0,0,0,0,101,45,0,125,21,0,0,0,0,0,324,0,0,0,0,0,0,0,0,0,33,0,0,0,0,0,0,30,135,0,0,0],[239,881,2451,725,313,295,331,679,451,498,826,1356,1434,258,465,0,431,655,518,0,635,300,0,2206,603,1183,0,0,966,651,4917,0,0,1698,859,353,41,2546,296,0,0,11239,1985,1925,4349,1730,11396,3899,969,0,10522,1308,1394,2069,1794,26,0],[1401,858,38,889,545,606,457,867,539,1107,1180,589,412,210,451,2716,27,1478,574,359,2015,394,2455,103,1355,967,1269,2110,999,2155,0,188,743,207,1025,1687,460,2045,0,280,0,0,473,489,18,474,195,0,541,0,5529,47,651,194,294,25,187],[55553,36501,19961,134228,100525,54967,59733,58663,90351,149813,152698,19292,82039,61837,61224,231440,64195,37077,178151,78822,115181,194868,145632,167667,89788,126584,64872,74424,110112,100172,11901,57259,38068,82916,159370,132828,65127,147317,33318,102210,92421,12573,57821,55991,28153,92893,58670,36848,69112,59799,281730,62993,82809,18500,44458,93135,51264],[28344,48136,4477,87501,59539,68965,55503,59106,45622,172814,146960,9518,55949,61196,54821,108757,41349,47457,91439,65589,107956,101528,51353,42916,63808,109555,52692,16132,110279,178126,18358,55404,62107,59227,82472,69021,33656,125292,34608,60606,71296,15601,70971,51997,8437,65100,18019,17086,45578,28480,196922,52439,65171,5259,30029,33798,62106],[342,939,0,887,595,495,206,392,244,614,1496,100,414,287,301,283,113,864,1150,368,993,1614,140,0,916,985,0,0,692,1492,1794,848,1983,892,412,996,0,1397,0,1359,0,0,1315,247,0,21,51,25,1004,1014,0,311,476,0,0,160,0],[156,10,659,0,0,0,14,342,0,12,296,82,337,359,84,334,0,0,0,0,107,0,0,0,0,179,430,0,0,0,0,7,0,6,279,62,0,0,0,0,0,0,1091,310,283,49,2507,682,35,0,56,0,20,188,6,24,0],[982,929,197,4191,1611,3422,1511,1298,3432,3169,4792,1033,1605,1790,1899,2226,195,1464,3151,2468,3178,1877,4413,7002,1622,1515,1615,1307,443,1283,4030,1209,3125,2466,1956,3746,1583,1085,175,728,192,1264,2118,1770,1747,4599,1235,1266,873,458,8244,455,605,0,588,388,579],[3536,2834,201,3588,2757,1776,6645,6073,2007,3694,18408,1282,5179,4707,3540,6355,4425,1459,7010,4519,5689,14407,5094,1829,6579,6259,4388,2685,3084,6111,0,1820,3466,5892,14463,9790,4334,6459,1920,4146,4731,0,5953,2514,625,4178,3903,739,4425,2773,1848,6690,6472,48,968,1799,632],[56976,46553,6536,178045,114278,72413,87428,102710,39641,270706,390841,9236,148648,159027,174307,163704,49104,57193,180105,113713,216812,224690,135409,93443,185570,170306,77153,26487,155958,102228,57282,132605,79710,143892,172839,189026,91359,207251,28858,114431,111773,8034,104870,58984,13905,123457,58839,42238,113527,116625,508758,86047,105006,10234,51127,41980,39616],[11870,15849,7930,23043,17464,27547,11004,11545,14534,43647,27429,6097,26145,27787,28020,29366,10663,19458,9616,12208,21084,10092,18111,27785,6955,9626,6052,6394,23357,28806,9717,9741,8035,11778,25310,35303,1203,32362,18560,17819,23101,10336,20056,11007,12031,10598,11441,10860,9392,7517,80615,7107,11373,2843,8308,15606,8288],[2112,3491,3398,3869,4472,4144,3814,2683,6622,15021,10964,4322,6653,6241,5350,3222,1207,2837,4386,3661,4822,916,5100,1620,2181,6990,3212,135,2453,3985,53,1366,6242,1919,2299,3984,0,8585,2284,4003,5008,249,4319,4102,4936,5288,9353,4810,2395,1909,32392,1942,4137,4096,1506,2109,3274],[115,124,0,0,0,0,0,127,98,0,0,201,0,0,0,0,0,0,0,0,0,0,0,0,0,519,0,0,344,0,27,0,159,0,0,0,0,0,0,51,0,0,0,0,0,150,0,0,0,0,0,0,138,121,115,0,0],[29098,35943,12250,58743,47699,74495,31825,31725,49316,88633,89498,16367,53652,45978,36699,55256,17418,19829,42054,37668,60365,47249,49948,37887,45126,54009,28341,26700,111119,90320,16370,22510,37752,14582,38147,40104,6012,60767,44044,19851,20254,16704,38030,20571,34227,41347,33157,27360,22035,13494,142784,36014,43672,17376,27620,34728,44548],[52,169,2748,229,260,356,0,437,202,0,936,2232,648,4,415,150,5,4,0,0,0,295,0,1440,0,546,1122,60,2031,201,5374,0,64,404,228,7,0,0,1180,100,148,0,1260,1033,1815,0,2886,501,0,0,8768,1515,1677,3414,20,687,316],[11149,26734,14447,28990,26855,38836,14504,20024,41864,58505,33997,14631,17735,15321,18109,15904,3065,8669,26504,22439,38586,31440,55838,69225,18652,32619,11446,10136,17879,43335,7291,12729,16194,30637,44936,48952,8248,19423,56727,17457,22380,17530,13428,11584,25874,27891,19068,22236,9553,13685,162370,17932,21677,9202,10882,13128,8120],[68012,118380,10145,160643,125709,88741,70456,92894,23145,401967,142169,18077,81364,14273,18089,133579,57729,19281,122773,47149,195703,124598,43530,55234,123676,227622,46541,21905,55809,34377,15217,59299,7580,91728,97305,35353,88223,56601,16775,111685,64082,16392,115821,79713,13459,185955,58472,18515,103506,47367,432664,63563,49425,1788,34630,42772,84385],[0,173,0,0,336,548,0,515,392,490,380,1409,403,200,0,0,0,0,193,0,0,0,0,606,99,317,1206,25,0,594,0,0,0,0,0,350,0,0,0,0,0,49,1003,81,2003,0,0,0,0,0,0,0,178,174,0,0,0],[1075,66,184,577,994,1226,806,1005,397,1705,2702,130,1524,748,690,1221,240,120,793,701,716,2393,567,233,1636,905,622,1476,1827,3198,0,710,390,1418,1055,1530,2215,1687,711,1341,1056,0,436,310,387,1012,1368,204,942,178,573,1334,1265,165,1682,918,0],[7880,6844,5621,16825,14331,14054,7465,7462,6933,25928,32158,4902,15843,19719,15304,18130,7370,3795,9670,8065,20908,13004,16216,11943,5077,9497,8334,3305,14874,25268,0,3577,6619,8303,19037,25903,10097,15139,8038,11379,16240,7707,7997,3449,10033,11931,6077,4340,6473,3794,32383,4784,10683,1811,7068,6497,7528],[5634,7650,4191,16870,14777,12436,12767,12823,5060,17859,22709,3513,15836,13154,12724,20691,2722,5897,15940,16204,19415,20568,21705,16497,13083,18168,5204,2994,16555,23903,3864,4600,8591,11550,25668,25451,5425,30058,5034,10868,8342,1803,5424,3995,6319,9303,7560,8302,17182,10559,34540,9315,15397,7974,7934,6590,7470],[210,362,0,0,388,306,46,493,0,152,97,0,176,145,198,638,716,0,377,318,189,439,0,0,292,691,72,0,101,15,0,785,0,162,705,796,0,61,0,18,0,0,121,303,153,0,0,646,0,35,0,139,161,0,108,0,0],[0,109,644,0,0,0,0,0,0,9,145,218,178,32,138,0,32,0,0,0,0,66,0,0,0,8,0,0,53,0,0,0,0,0,444,0,0,92,0,0,0,0,0,0,946,0,371,219,0,159,0,16,42,270,268,0,0],[842,576,603,540,812,634,608,1286,871,1947,1593,236,564,1568,352,1262,219,1699,414,1401,613,1291,3018,0,1509,1266,142,218,1383,1894,0,1696,776,786,2025,946,0,4108,0,801,0,2410,2448,639,38,2093,2186,431,857,995,0,1123,953,518,94,268,0],[62353,60271,9899,170357,105103,122747,79473,81671,36623,235450,249461,9505,112762,95563,90137,210457,84342,91080,129061,90561,129119,209431,119425,69872,152694,140134,75622,58427,145020,176740,16018,79158,62062,129572,204445,185424,80600,211634,68160,98119,86402,12706,74425,46020,18465,98930,50363,29834,90506,73830,347848,78307,130783,22719,60763,45225,57018],[103,0,481,0,0,13,59,0,60,133,556,0,465,207,89,381,0,51,0,404,37,518,0,0,0,0,0,118,542,264,0,6,54,400,1257,498,0,1561,0,0,0,0,198,0,332,603,3399,1821,0,0,0,277,302,1228,0,0,68],[9287,18716,2396,27155,17710,29682,15807,16363,11121,65293,25643,4685,9597,13424,14635,4923,724,3792,27328,16227,46864,38970,44183,60510,17647,38876,13826,5960,25662,38606,2051,15595,10395,33650,40079,65667,18383,33391,17698,11887,11282,2731,23260,9484,8858,18868,7104,5459,19099,13054,174106,15118,18939,2399,1053,5097,8841],[10450,8490,1259,26682,22871,26522,16646,11937,1889,19617,126682,3308,26049,16429,18418,26769,4197,25011,19680,26972,69072,33482,25779,8376,17423,26896,15617,2765,33446,29196,10524,28018,25820,20654,53553,30650,20809,64089,27075,6782,22420,5311,25577,4711,2469,13180,10069,14527,23829,10715,23334,11464,22062,5108,6012,8922,9102],[31281,44254,11095,89189,67929,55724,46515,61242,109017,257784,229533,27400,53974,43787,41893,121784,59176,63843,95502,78309,127702,154333,134360,191489,95316,133563,40600,192993,149619,145464,19096,134118,53642,116099,130849,129143,96928,131651,56511,57302,79135,23134,152397,66319,21438,101351,39480,36395,41029,94322,479833,60249,59295,10557,72095,62338,81350],[0,0,105,0,0,81,0,0,90,125,146,0,0,111,0,28,0,0,0,0,0,0,0,290,0,0,0,0,0,189,0,0,0,0,0,0,0,0,0,101,0,0,0,0,0,0,0,245,0,0,0,0,184,308,0,0,0],[0,0,10,0,0,77,112,0,124,600,68,201,0,0,0,0,0,0,0,0,0,0,0,0,119,237,0,0,12,0,2352,0,0,78,312,0,0,0,0,246,0,0,0,199,1038,1467,193,0,0,0,576,0,129,196,100,0,0],[0,0,744,0,0,0,0,0,0,0,135,416,538,0,0,0,0,0,0,30,202,0,0,0,398,0,178,0,0,0,0,0,0,0,0,0,0,78,0,0,0,0,0,450,401,0,728,0,0,0,0,563,0,0,0,0,0],[0,0,0,0,0,0,115,0,71,0,38,0,0,0,0,0,0,0,0,0,0,0,0,0,0,0,18,0,152,0,0,0,0,0,0,0,0,107,0,0,0,0,0,420,0,0,0,0,0,0,0,0,0,0,0,0,0],[24495,57030,11922,60436,64254,119981,11602,17012,27018,57882,117831,15470,48436,111560,119830,23402,2881,66166,11092,34915,337122,15702,74603,171639,7691,56555,24244,11761,44247,142729,5636,48952,36843,33421,91904,86575,20512,185662,420847,14649,47341,7306,19864,9273,31400,9679,129402,190049,17467,15089,199642,47777,37919,16014,20462,25802,13705],[0,185,1257,0,0,319,0,0,0,188,0,927,1334,0,404,0,0,532,0,0,0,0,804,0,136,713,105,0,289,0,0,1404,177,896,0,606,0,808,0,0,0,0,1401,1033,1581,260,173,2618,0,0,0,0,423,536,0,0,0],[77,156,37,0,0,0,9,108,149,151,32,280,0,0,7,132,0,0,0,13,0,0,0,35,0,0,0,0,28,0,180,0,0,0,33,36,0,0,0,7,66,0,0,0,15,0,17,0,0,0,0,0,16,28,62,38,0],[1061,2657,2002,13148,10614,4182,155,287,4736,3907,22121,1481,975,115,209,7732,2084,655,3799,8179,6368,2821,20012,11697,2723,6079,3153,259,19235,18244,410,349,4060,1960,15651,2751,2938,970,1598,6001,24198,4914,1015,9836,5080,4307,8029,1164,3117,8144,17953,1181,18832,3645,2188,2679,450],[5178,8116,4114,14769,10855,14084,7452,7616,6570,22225,19435,6034,14274,12786,12820,7857,2040,7162,6958,9553,10968,18275,14993,12594,10735,12998,5215,3659,15395,23583,2635,13077,13946,18721,19942,20160,13414,28743,18292,11140,5341,6025,16886,12216,7638,9668,8268,6531,12176,10088,63096,7556,10881,3220,6772,6884,2529],[47,202,133,0,129,4,338,173,402,84,2685,385,136,50,415,28,0,0,0,0,162,0,0,192,252,225,355,0,373,58,551,527,353,0,1062,1827,0,84,16,0,0,0,301,281,659,47,1480,979,0,0,0,6,379,686,93,0,0],[14593,28096,2320,33107,34826,30963,32326,30085,59600,80498,82337,5162,32327,29456,26320,80470,24104,34123,57849,57507,59915,89782,46858,39749,40227,56484,18929,32460,75856,106993,11613,35004,46919,40831,71009,52974,13963,53665,15272,51280,49612,4856,41098,16596,2746,27282,20526,8744,53611,39996,143099,36532,39133,5217,11884,16641,37014],[0,2196,794,141,1423,1294,753,398,0,1793,2919,495,749,845,2078,917,5120,340,1932,2132,1848,280,808,990,935,1373,1220,0,1353,75,0,0,709,607,879,33,0,2263,0,483,0,0,597,1506,0,404,2860,1406,194,625,3005,1169,1201,2562,36,163,6760],[0,0,0,0,0,0,3,0,0,75,0,0,0,69,7,0,0,0,0,0,0,0,0,0,0,0,0,0,31,0,0,0,0,0,24,0,0,0,0,0,0,0,0,0,0,0,0,0,0,0,0,0,0,0,0,0,0],[401,617,3090,1846,2131,1941,812,1682,1843,2297,3855,3052,2011,2881,2066,1547,116,1304,721,1029,1344,2265,1143,3196,289,2725,3152,1738,3160,1558,3641,978,1145,2308,5229,10228,0,3646,11128,382,833,6219,1443,2923,6470,325,11432,7871,344,1236,23497,610,2794,3695,1225,405,261],[2,1209,82,583,660,181,142,533,408,1374,685,520,247,1059,110,860,0,136,322,907,1845,273,386,0,241,309,2899,0,927,1179,1909,60,403,432,486,686,0,1287,1098,66,1234,0,647,456,0,94,847,2262,820,39,660,196,1159,368,200,605,0],[658,474,156,1417,674,337,842,406,59,4113,2228,25,1389,128,217,2405,784,314,1767,374,558,2480,1451,53,1951,899,92,29,738,165,0,961,563,872,372,29,468,443,96,2169,1731,0,1292,831,204,2377,984,32,1606,6,56,455,1114,33,799,205,305],[171351,254194,38400,269067,218362,195473,193550,227384,390881,812720,658268,60438,188716,211835,207657,357555,144379,209339,295465,321908,438475,438276,206962,197509,306955,660550,226430,178395,480229,560103,72440,323249,378591,296243,412189,363794,221686,529528,155458,243341,201334,33152,546135,260624,146050,265392,132907,100902,287313,149793,1092756,201102,247785,36801,81055,87546,174917],[100,609,2752,542,353,406,199,306,598,422,801,1171,562,329,850,228,50,280,0,579,295,0,693,1699,313,277,2753,0,371,972,0,0,799,55,1003,1632,0,1549,0,53,208,0,2634,2801,7566,598,6332,5254,157,3387,10787,959,2421,1400,658,136,468],[2085,3147,478,4002,2738,3093,3498,3838,1302,5840,7537,81,5312,5131,3436,4053,225,1061,6317,3125,4698,5346,2389,2542,4903,7232,1324,0,5305,2040,3747,1637,1201,6896,7540,4384,12329,10766,4434,5369,3043,0,4614,3779,524,2788,2346,1153,3900,4000,9071,4904,5216,724,2546,893,4650],[1006,2799,1534,695,727,1235,1776,2089,1518,1498,5187,1190,2183,674,1396,3598,793,386,2926,1283,3420,2583,114,5073,2267,892,0,360,5174,1491,0,1419,52,3506,1957,2110,196,951,3819,2533,1811,0,2995,2132,606,3139,0,7,1576,1214,6729,2692,3603,845,850,270,3396],[84,113,1419,122,0,375,171,174,0,191,590,1142,103,36,165,301,390,22,66,0,33,0,222,1298,46,451,467,127,61,48,0,0,148,29,667,215,7,0,2028,38,0,2,472,411,1002,0,2585,225,0,0,25,819,547,2800,310,2,5],[33710,69176,5069,95764,60883,121580,37642,47967,59172,259603,142844,31305,40692,42025,47554,342003,73272,168426,23592,13350,31752,51749,19146,31433,26176,33618,14530,67064,35340,40964,418,14840,37801,21594,49551,75851,30669,88486,41434,67268,109863,91865,76688,33447,25045,69458,34864,35669,30175,6264,183460,17596,28810,4748,20481,26825,45765],[1492,3768,572,2171,1428,850,3241,2485,1042,3610,4964,91,2764,2893,2675,6862,5293,3619,4672,3256,5952,6632,3384,562,3541,6339,5038,4860,3566,4103,3775,2054,4901,5637,3252,3726,2729,8307,41,2978,642,0,2509,1343,77,4414,1311,1090,2003,1166,8778,1794,2037,214,2122,1970,5021],[0,0,0,84,0,96,0,0,32,44,67,350,0,0,0,112,0,84,0,0,0,292,0,0,0,167,0,0,167,86,0,0,0,0,0,0,0,0,0,0,0,0,97,70,137,537,12,990,120,0,24,223,95,0,116,0,0],[138,0,257,154,264,212,0,479,352,1108,261,469,176,337,281,0,284,0,196,186,158,166,275,23,245,448,327,0,0,581,0,110,64,0,406,531,0,457,0,251,0,1485,0,333,508,149,1504,526,403,30,0,16,426,772,366,299,0],[6438,21973,7096,5317,4570,16294,3410,4392,9636,13387,10644,2779,5065,5912,7639,6040,725,1139,4517,4174,7298,10138,17444,16241,4463,11003,1523,670,16708,11279,1797,3362,3058,9095,25599,32694,1381,9789,5725,1863,632,4298,11420,9315,5026,1792,4959,1734,4930,1656,33755,5328,8827,2040,837,2514,899],[120415,228791,32264,323462,277722,183964,236045,282690,126153,1029307,385680,55973,204422,147369,177555,180595,121466,137442,354093,439459,348456,894251,504868,522228,253944,543248,170209,134219,256707,314912,147466,411606,164574,487886,329582,419724,145162,459550,296652,249888,273388,51721,1144856,349313,147747,454590,146435,149820,297909,167347,1473551,315998,323616,28346,133196,119984,129378],[404,0,53,220,0,419,0,16,1468,0,246,1231,0,88,24,104,53,26,202,26,114,492,26,196,26,122,0,151,622,621,26,202,536,0,0,1120,0,402,0,0,0,3837,0,282,254,25,0,0,0,645,48,156,310,990,0,272,1198],[0,0,231,0,0,24,0,0,0,0,0,24,0,0,0,0,0,0,0,0,0,0,0,0,0,171,0,0,0,0,0,0,0,185,135,0,0,0,0,0,0,0,0,20,366,0,39,0,0,0,0,0,0,0,0,0,1],[2056,652,4057,255,169,1075,110,592,1101,807,2555,2739,2169,1148,387,280,938,48,30,0,244,1671,0,1210,404,2572,1715,0,1797,1461,0,24,400,813,1328,3471,0,619,1280,899,1411,6028,1489,2872,4938,323,5188,2893,71,16,5872,1785,2624,7134,2054,126,56],[0,220,0,0,202,0,0,228,202,0,338,690,219,192,0,0,0,110,0,231,0,256,0,0,184,177,0,0,0,393,0,0,0,0,594,7,0,0,0,0,0,2106,0,700,0,272,0,160,0,0,0,0,0,617,92,0,0],[206,827,975,370,289,579,253,183,210,861,263,1112,714,342,256,1427,1537,1033,427,85,498,652,1498,2190,630,429,91,707,156,104,350,272,398,484,906,984,717,400,882,383,825,1048,1218,569,2981,337,2311,4105,668,642,8991,482,1553,1209,899,441,692],[25971,56773,3570,88817,67494,69839,55271,43418,74698,178447,160361,10564,52970,52292,51452,54563,21628,42169,76385,74258,109120,98404,46705,36464,85724,113402,55444,19822,120844,128186,9311,48046,54917,54359,106973,119092,47044,100725,23519,72923,91905,10941,97913,45656,13515,60898,37085,32309,63651,71470,417094,48594,58211,12555,15721,20512,24429],[7075,8748,3012,12860,10833,15023,8003,10964,4186,29974,28224,2030,12350,11302,16603,13115,5642,9558,12366,16487,21566,19584,13059,9225,13117,18702,9977,6289,19813,19182,3018,12118,14108,7345,21848,14410,10852,29312,8120,11294,16253,3136,10259,10507,3482,13733,8211,5095,7355,14797,82070,6568,11590,2798,4699,6008,12539],[0,0,194,0,0,3,0,0,0,36,0,0,63,181,0,100,0,0,0,0,3,0,50,0,0,0,0,0,0,0,0,0,0,85,41,0,0,0,0,0,94,0,0,0,0,3,0,0,0,0,0,0,54,222,0,0,31],[4059,5016,13253,20439,11926,5657,10066,9923,3130,48132,32229,12721,8129,4038,2857,26757,6693,6763,27804,7325,4494,12431,8427,4706,14341,11324,5462,1329,19075,18162,5828,4619,16178,6153,15783,20571,23913,13899,5388,19233,4196,669,10652,9097,18432,19969,24789,13021,4907,8517,22873,18651,17602,8610,8849,4949,8880],[2664,1935,542,1953,1275,728,1467,675,1272,2208,3519,350,1165,2075,2256,5873,1536,927,817,799,3039,1009,3381,2097,2068,3360,163,10138,1340,4827,82,2605,1050,3150,3972,9309,0,3194,1080,795,211,0,2282,555,784,1231,557,2157,1304,404,2747,1120,1253,425,4443,442,625],[5943,5602,6032,18927,12985,10519,14680,17112,5326,27973,36547,3017,15232,11225,14561,35004,4742,7798,28622,18705,23440,39220,21938,9443,17661,22053,11132,3503,31250,24538,3508,18117,20997,17384,27815,28389,25184,23511,4535,24565,17943,708,10496,9556,6112,16388,8079,3284,18168,10698,29405,11068,23618,8764,3400,5558,2152],[395,248,232,712,349,320,146,45,0,864,724,730,370,348,373,661,136,499,84,193,182,484,102,73,156,101,101,136,926,312,40,0,0,225,238,1170,0,200,100,0,53,0,332,202,1142,248,1460,1041,306,235,912,110,625,779,78,100,141],[79,238,1256,112,0,112,268,745,263,2055,1975,1612,415,164,72,1002,334,56,821,0,749,101,0,1007,0,746,834,0,606,493,52,0,0,1194,1521,2055,1396,827,3564,0,0,10730,865,454,3363,577,2165,665,0,0,804,448,408,1830,399,159,0],[0,1597,1249,583,172,585,153,830,201,1465,598,2027,729,267,0,321,98,0,0,345,227,1208,69,102,85,370,206,25,398,664,0,0,1157,202,808,545,31,0,0,92,2054,4609,1483,1224,4966,910,2772,1614,602,0,4633,988,560,2375,74,262,0],[111,269,101,632,1923,1413,1328,276,3,863,1606,475,861,1044,844,903,198,60,1355,1866,852,1799,1,106,1207,1835,1145,140,1220,842,0,1696,402,104,1415,2357,0,2286,1143,1380,1855,0,1434,892,0,23,2163,0,1132,0,389,789,1041,2573,2,737,3180],[1175,1336,422,709,584,782,350,202,1746,2248,1985,1187,2594,1033,2057,1023,580,1575,532,848,908,45,3366,1729,94,725,0,0,972,1442,0,883,0,580,435,1193,334,543,1227,691,528,2660,404,490,856,1952,1606,325,281,434,1550,1441,664,719,758,758,626],[0,0,1733,0,0,15,159,0,283,197,0,291,174,184,8,1786,1550,0,207,53,27,0,0,546,160,466,533,0,403,164,0,0,0,42,0,1883,0,0,3,0,0,0,131,78,0,584,712,252,0,0,375,488,182,456,31,31,7],[591,1703,761,927,809,765,781,1575,111,5562,1389,686,469,699,344,2282,673,348,768,540,813,1160,1540,66,895,939,0,88,659,905,0,142,218,153,1494,530,64,465,161,627,799,0,258,432,1190,140,699,0,128,235,140,428,763,217,150,362,67],[202,308,3539,179,528,704,582,230,709,751,924,2197,1120,124,1321,68,0,0,110,0,163,145,98,858,970,1063,1113,0,2555,1005,495,23,140,526,2760,1313,0,383,1514,402,308,7425,926,2043,3867,366,4384,4204,114,366,2160,1056,2923,3584,561,168,1355],[205,52,320,36,0,0,0,0,18,0,0,202,540,319,56,0,0,0,0,172,0,186,0,632,8,381,575,0,190,199,0,0,0,0,93,0,0,0,0,0,250,0,410,135,2726,995,2002,1407,83,26,0,256,502,559,0,0,0],[0,0,620,0,0,208,603,34,206,238,277,659,445,162,592,820,42,1368,0,260,659,0,348,2166,0,134,1979,0,976,521,0,379,314,0,2229,201,0,0,2462,0,0,0,114,416,144,1156,123,254,0,0,2083,395,382,395,236,792,0],[0,0,0,0,0,105,0,0,1,0,0,0,0,0,0,0,0,0,0,0,0,0,0,0,0,6,0,0,0,0,0,6,5,0,0,375,0,0,0,0,0,0,0,0,0,0,0,0,0,0,0,0,94,0,0,0,0],[209,23,0,0,152,36,0,199,140,83,0,0,75,0,24,0,0,0,0,0,0,0,0,196,0,89,52,0,55,0,115,0,141,0,54,353,0,0,3,0,0,3225,0,200,228,0,269,7,0,0,516,296,167,0,0,75,0],[0,0,821,45,342,293,0,0,35,130,0,232,0,0,0,90,0,0,0,0,134,0,0,129,98,209,0,0,351,0,0,0,0,0,315,264,0,0,0,0,70,0,48,34,191,0,58,146,0,0,3029,0,0,42,192,0,0],[450,226,0,240,48,266,348,355,167,1281,254,606,532,132,214,173,118,232,123,0,264,266,58,544,251,0,148,206,14,616,0,0,0,92,517,560,0,104,1230,282,0,0,30,0,0,0,613,96,270,0,485,170,199,736,0,0,243],[0,0,297,0,58,0,0,0,392,0,80,256,130,0,0,0,0,0,0,0,0,0,979,0,179,182,0,0,0,2206,0,0,0,0,103,874,0,0,0,0,0,0,0,322,476,306,0,1412,0,0,0,109,117,31,0,0,0],[0,0,1642,509,141,154,125,202,13,450,239,849,235,79,478,82,0,0,0,44,0,272,0,742,634,469,113,0,713,756,0,0,0,4,979,388,0,383,0,0,0,2808,1149,35,3183,658,2544,1993,0,0,1942,848,611,1141,48,307,0],[214,455,104,563,275,124,114,0,799,147,1117,609,63,0,102,0,0,390,0,146,969,0,86,55,148,830,28,28,439,438,0,0,374,264,115,141,0,80,0,88,0,0,420,114,503,0,2725,72,0,137,5237,231,452,12,116,88,0],[0,0,102,0,0,0,146,0,0,0,0,654,0,85,0,255,0,0,0,0,0,0,0,0,0,0,0,0,48,0,0,0,0,0,620,0,0,0,0,0,0,276,0,0,0,0,997,0,125,0,1470,0,0,0,0,0,0],[25961,37670,10922,40964,32221,38210,18495,29551,74563,132785,82424,26652,27262,23485,30161,60538,15785,27295,22430,26945,66613,44324,68405,112319,27600,61668,22567,25823,35793,47544,21058,43360,36934,47660,64337,92784,20177,47805,44360,21369,35234,21950,83270,51951,40516,32926,22721,24055,24898,22753,182591,25899,38987,7758,21373,22213,37560],[0,198,919,0,0,1685,0,399,351,257,571,1064,267,201,109,0,0,88,223,0,201,896,318,363,197,402,0,0,578,243,0,0,0,510,205,515,0,0,0,367,344,349,0,152,3252,157,2706,2298,0,0,373,569,591,997,352,0,0],[10581,16104,3738,16304,15917,18353,14107,17294,17192,52919,50755,5551,11520,14186,16044,25844,9581,17023,20752,22784,31996,27824,32532,27367,10274,32539,11991,13224,27271,32090,485,12444,10548,20460,39654,28635,576,27975,13141,16025,21755,6859,25541,14345,7434,11132,12051,14982,18965,9171,53777,9696,17520,3451,6809,7357,15356],[334,354,2069,1326,836,720,555,438,2525,742,435,2341,366,360,1204,1088,128,0,751,0,0,733,4692,2279,712,471,1586,25,2069,1300,1994,0,1305,1209,1571,446,0,2906,6,1246,788,0,812,1345,5306,1105,3550,1302,2070,0,1229,777,985,2765,213,58,49],[3416,2720,9389,4162,4423,5661,2345,3013,3303,12466,12208,6874,6240,4106,4410,11658,4550,2845,3431,2716,4196,5181,4442,10936,4799,5264,4384,1023,4809,2695,2659,2857,3274,3563,6777,8608,1859,10144,7275,2510,1335,4600,5773,6270,12153,3675,15129,8320,3920,3254,22676,2805,3777,7043,3000,2968,4107],[909,17,1031,0,26,333,0,0,13,13,401,373,430,43,386,43,0,0,0,54,13,0,0,134,280,761,0,0,298,13,0,0,43,384,360,721,0,0,0,461,0,0,1166,439,2307,78,1389,1132,0,0,0,600,278,500,119,2,13],[10,0,61,12,174,16,78,29,6,0,329,27,663,662,94,108,35,402,2,0,352,0,0,568,0,0,176,325,95,67,0,49,0,80,498,214,23,330,12,18,0,1872,102,2,122,185,42,6,0,0,0,74,226,6,0,0,25],[1755,8254,2907,3906,2098,5783,683,2633,269,15045,2829,781,2465,1394,1897,1526,606,704,938,1019,3702,596,0,324,302,757,203,412,426,2308,24,425,643,347,1427,1416,515,1612,2810,727,2191,925,749,1751,369,359,852,298,224,227,2991,1892,2870,1557,1420,2447,2124],[0,44,399,0,41,0,0,76,0,84,0,226,67,0,0,0,25,0,0,0,0,602,0,0,0,202,96,0,0,478,0,0,0,0,974,0,0,0,0,0,0,0,747,0,0,0,201,827,0,0,45,202,78,0,218,0,0],[424,1783,1348,2248,1551,1443,195,299,711,4133,745,1111,316,119,814,829,202,599,288,183,934,232,0,501,0,419,448,28,584,497,52,0,226,122,396,362,25,326,343,32,428,728,197,353,849,141,226,1021,344,90,3133,261,130,334,499,248,370],[43826,39783,4606,104734,84366,42877,55936,69222,54803,176612,209024,10444,45739,35519,26199,58409,16580,22040,115374,76426,77973,115025,84608,54751,94240,88333,44160,11694,81525,85921,6465,29959,59178,69538,123298,74743,61865,72884,23291,68852,56547,6528,56596,70513,13554,93109,35637,30267,66671,58466,206673,49394,111792,8504,25242,35447,22820],[20779,33130,4836,46900,30168,37241,36366,34881,30136,92253,71980,7364,41646,35766,34286,55191,22893,19182,45273,30600,43996,78630,26952,30492,34173,48271,15295,9619,53107,64772,23745,36903,23519,43456,47783,56049,22197,62981,27175,47153,43686,11212,39183,20151,14024,40918,6441,13604,26337,17034,104588,30061,32097,7184,20540,15834,24021],[0,0,51,102,51,0,0,79,180,256,10,149,0,0,0,0,0,0,0,0,0,0,0,0,0,0,833,0,746,116,0,0,0,280,352,308,0,745,0,0,0,0,0,627,374,93,66,43,0,0,0,96,102,166,0,0,0],[11433,4068,3115,17606,11273,22218,13375,10714,0,39179,20309,4302,29791,13238,5893,37242,16111,18202,4848,11092,9915,38370,24411,8231,3307,6860,3740,8527,68979,53754,360,24465,7616,13146,34712,50849,33817,39998,7562,8641,29392,727,14876,6998,3713,4909,11180,8975,13388,4396,10557,21293,37340,2493,4748,99514,4840],[337,100,1241,1546,796,493,42,470,429,1574,1513,1185,542,372,198,573,119,232,600,477,209,250,99,387,1072,1432,922,0,698,753,608,100,189,1159,1769,3701,54,67,424,820,0,2558,101,302,643,332,1001,1290,0,0,1909,782,1955,1777,1124,178,134],[1516,2792,1200,2093,2403,1741,1495,1294,2349,9002,5904,1406,1451,1114,1348,1033,961,333,3043,1902,3549,4571,3915,890,2311,4199,2430,332,2121,3589,0,1633,2012,2647,3803,7910,0,2228,0,1360,337,0,1649,1149,352,1635,1367,191,584,596,4225,1814,2499,1275,326,331,311],[31,0,901,630,210,196,2072,742,0,5454,1266,82,714,90,0,6786,571,0,212,877,384,760,1146,1231,30,0,606,0,3652,0,0,298,875,1403,480,2477,102,102,48,2761,0,92,1481,700,0,1676,0,96,1749,764,72,592,1064,0,0,454,0],[256,249,200,542,373,864,0,162,663,2118,983,1377,376,488,323,1087,117,409,270,493,327,1397,631,70,37,92,93,103,226,309,0,55,199,780,1138,784,0,55,100,244,66,11670,57,76,1610,55,1097,1188,55,0,877,201,320,0,608,44,200],[0,594,952,0,0,97,0,453,0,176,397,172,35,56,232,0,0,0,0,0,0,780,0,0,0,0,168,0,996,148,0,0,0,0,34,368,0,0,0,0,0,0,473,344,93,0,1405,0,0,0,9036,0,457,507,259,219,0],[0,0,0,0,25,13,0,0,0,0,0,0,33,1,108,0,18,0,0,0,60,0,3,26,0,13,0,12,45,13,0,142,0,0,1,0,1,4,0,0,0,24,0,0,0,0,2,0,0,3,18,0,5,18,70,0,0],[137,192,316,0,0,496,112,159,0,0,126,9,0,276,379,0,0,0,201,0,0,0,0,600,66,134,0,0,0,101,2464,0,0,50,0,0,0,0,0,0,0,0,101,524,437,0,301,1206,152,239,0,101,244,966,0,0,0],[0,0,16,0,0,0,0,0,0,0,0,0,0,0,0,4,4,0,19,0,53,0,0,0,0,78,0,4,0,27,0,0,0,0,0,0,0,0,0,0,0,0,0,0,0,0,496,0,0,0,4,1,43,0,0,0,0],[172,502,821,346,314,430,0,303,29,559,311,722,122,321,105,1005,771,306,172,277,326,447,403,162,114,236,19,217,172,270,43,0,265,582,329,693,182,108,590,0,190,402,162,59,315,50,46,276,81,29,1257,410,385,50,255,66,461],[139,209,469,328,102,770,145,536,448,526,603,1458,363,0,483,541,35,309,137,88,204,186,0,70,299,640,220,51,541,760,0,46,348,200,344,1757,0,1053,90,136,51,0,3207,643,3907,887,4495,3583,88,0,358,181,704,560,677,100,128],[1436,1141,474,4461,2913,3236,2805,1906,1528,5514,7950,269,3426,4502,6848,7179,3169,4100,1858,4646,4094,5685,5733,665,3065,3841,1543,434,7111,7863,0,3954,5201,1805,4896,3226,2018,6604,962,6814,8227,0,2028,1631,185,1106,1139,329,3168,1131,5685,2223,2232,1190,1359,1862,1416],[293,199,535,346,973,394,236,0,198,2445,588,436,0,610,404,0,492,0,982,504,470,167,603,1217,0,201,0,0,670,452,592,0,38,606,490,196,1508,0,0,0,600,0,0,1467,1676,838,1446,3721,0,0,463,193,619,428,563,367,0],[15422,10944,3914,32757,22283,29081,18120,15347,9667,36254,42163,4348,31551,22196,20601,51225,10301,20531,29885,24135,42837,42171,20728,7356,28460,20419,21478,5798,43635,46029,3566,17907,10704,17363,44787,38975,13799,74742,11846,33546,31080,3290,14485,9307,1305,16062,17532,11515,20539,8806,26159,17251,26992,7262,8184,10528,5598],[14044,22270,4454,31860,28984,32751,14799,17379,28676,45787,43754,12042,21165,22269,25682,26164,7485,8853,18720,27001,34154,32204,49708,53712,20394,28278,15433,5385,29201,35973,11711,11309,13161,37528,57269,53569,21111,22549,24679,15113,28968,18235,15247,10606,17138,32807,8950,23343,15746,14514,90900,13331,24039,4289,12926,14383,12613],[238,0,1239,0,85,468,0,276,482,750,1,503,579,199,396,105,0,0,268,0,245,309,439,363,34,784,200,45,505,433,0,0,365,0,88,71,0,0,0,0,0,32,2490,113,1105,0,1169,508,0,0,2595,757,768,2989,304,107,0],[6960,41596,1869,18609,18077,16567,19785,27804,37184,56250,84601,5225,17002,23254,22437,6745,143,3165,24567,23153,59320,27824,39398,38321,14437,26929,10040,426,21897,35836,1408,9129,11153,26827,44523,41075,16038,51932,24309,12250,12381,456,9488,12422,7733,18256,2977,8154,22848,14674,110561,7598,27390,4344,2895,2521,11872],[555,1302,1887,2114,1400,3008,1394,2839,541,2049,2575,895,3762,3450,1736,5894,1963,6,4366,3121,3625,2569,2938,1047,2888,2159,2090,1904,2666,4135,0,1007,3867,1640,4731,2541,853,4431,207,2222,1234,1787,1551,240,1672,2106,2371,3219,1341,1343,3430,1267,3185,603,834,405,0],[1107,3082,303,803,725,1254,1209,2199,446,1865,1818,361,1175,1808,2959,1566,120,392,578,444,2042,397,1740,2316,1335,2454,269,188,2255,3307,0,516,583,2078,4371,3107,0,3470,1893,1799,149,0,621,279,70,90,0,370,336,254,2283,472,392,0,163,165,0],[1156,2992,1860,829,1316,1428,154,472,371,2582,2144,811,2195,882,396,905,205,385,1345,795,518,991,481,995,1535,748,2137,604,2088,1975,0,443,879,561,1746,802,0,1206,1208,999,236,0,4441,2424,1812,3882,7596,3326,2969,0,13149,1534,1056,1629,1330,929,1172],[89551,149048,18904,169102,131855,162639,88205,84616,185705,294459,370768,30720,90150,116615,137491,156848,34043,97690,117949,97843,219322,149802,123614,97776,114023,399158,163149,53258,300688,345217,44390,103053,177956,109730,254222,249074,131097,229016,62096,84665,166386,28892,251777,226685,57113,109220,75390,67844,202811,110842,698542,91163,226057,25773,33671,61451,133650],[72,143,140,264,106,242,60,245,92,376,457,238,52,166,112,1621,1214,763,244,272,608,134,161,60,172,314,220,216,191,338,4620,58,468,111,365,390,43,60,1582,56,373,183,128,54,660,33,159,438,0,109,2160,101,294,203,474,64,8],[390,1093,1431,5145,4679,4394,245,229,1187,1835,585,1195,546,332,562,137,73,193,1605,1370,4677,297,538,2318,542,429,0,142,444,1032,92,0,244,825,741,2690,244,404,1769,326,281,895,409,232,1575,335,122,542,65,0,2940,28,714,735,319,0,1202],[554,845,832,4883,1785,1679,520,3862,170,20200,953,13,1195,435,908,1903,1135,2562,1822,710,8239,1166,930,2522,292,308,143,1386,274,519,348,422,1970,1773,4153,5917,2112,1441,2619,454,2817,1500,2651,392,1203,2862,895,274,590,754,8647,618,306,69,3599,569,3081],[1334,1180,806,2346,508,2262,187,366,1313,1797,4005,2376,851,1481,3020,0,32,728,0,7337,3624,0,3867,8732,0,3458,1673,101,2854,2974,440,0,101,701,4481,2545,6828,6969,78,174,2145,11057,415,1095,2798,4994,4095,1222,2642,2194,30107,299,965,2023,43,1538,8576],[6265,3084,3224,16443,11780,11961,5039,7421,5265,16560,12072,4062,7651,14131,7637,9662,2457,5770,10076,5981,11910,13245,12170,11606,15995,14162,2770,6167,8015,6569,2240,8672,4259,6284,11947,12499,3055,11519,10007,6910,4231,5137,10490,6411,4991,7262,4147,3984,4437,4697,38455,6505,4604,2492,5577,3970,6106],[26841,19336,1678,44273,35038,43223,21589,23520,30039,79951,78340,3362,27021,25294,25844,24986,9921,13356,34629,23832,57435,39346,36880,24944,29654,53792,15521,4061,67035,74388,11420,32218,28069,36067,61385,50387,28542,44120,10696,25252,30479,3278,20941,13028,4132,20858,12288,11423,13886,13714,134273,19995,23601,5934,11227,13173,15786],[72,117,620,0,70,270,229,54,0,259,359,750,75,63,269,27,0,0,0,0,0,0,0,30,0,386,0,0,0,0,169,0,0,400,0,671,0,0,0,0,0,0,780,629,777,0,1984,70,0,0,31,0,91,14,10,0,0],[0,181,651,33,934,775,0,200,220,0,661,153,100,201,363,575,75,0,0,423,0,0,742,780,196,122,0,0,202,168,4419,0,74,154,185,0,0,0,45,0,0,0,980,198,2956,0,0,1568,0,0,70,0,461,186,207,306,0],[2999,4080,1803,8661,8499,8234,6049,5880,5533,17886,12769,3745,6860,4448,3916,8329,4447,5873,9985,7170,9598,14532,13957,17007,7167,8493,4589,4984,8183,10517,1579,4180,4490,7195,19573,17569,1193,9449,8065,7450,6985,4068,10488,4407,4461,8949,9170,7284,4605,4816,18194,5362,8070,1735,6983,6462,4548],[0,0,829,82,0,706,125,282,202,685,922,1448,209,156,202,1174,0,404,403,0,201,0,0,1530,202,600,0,0,788,403,0,0,0,937,639,198,0,59,1002,164,0,0,1926,120,1784,0,2339,0,0,0,0,339,401,199,282,44,0],[12023,28604,12099,56184,36819,47228,10672,10136,15137,86513,24895,12926,12046,10253,9195,100183,20925,28655,12514,8537,19354,22441,15038,17933,12910,7358,2933,27011,6711,9835,182,5882,8936,11489,19782,24176,6013,4147,7593,18162,29941,12803,10516,6146,10376,7911,1620,2377,2623,3022,15477,7733,8696,2248,12942,38934,52428],[17558,25131,3956,34330,26170,20386,26542,23583,24420,75729,52932,6092,24821,20305,24185,28853,10397,17510,48079,27778,74889,87878,24027,25605,36200,45744,25197,5823,32691,42982,8763,22778,27015,47146,32983,41524,15398,53693,24227,18734,29642,13234,32984,21835,8695,50390,16447,13089,20715,10850,162901,18428,26182,3919,15396,10926,13462],[3621,14030,1756,6727,6132,3017,1749,2881,20264,24785,24276,2142,17486,3622,2848,10042,3359,3852,15438,2161,7951,7882,2937,656,9391,8317,702,570,3166,14520,870,3218,6073,5953,3832,1310,319,4397,2007,6247,4611,435,3814,1740,6714,7859,6771,774,2747,2988,44991,5209,5514,1184,4922,4512,13954],[11106,14512,10949,54173,44547,31135,29291,32917,15678,117696,117741,16439,25956,13799,17264,54300,25506,14418,40649,26058,28079,52541,19720,11484,24692,25517,15025,9602,37799,41572,11327,29225,24486,54379,28075,29403,32310,51333,17928,32812,35001,5736,56494,25048,27289,28963,23526,18461,27543,20113,91278,24983,30186,9276,10088,15704,21529],[1198,1120,9999,1667,2128,3539,2938,570,953,4841,3347,5210,3613,2010,2163,4160,2085,1305,2529,406,1754,1821,727,6093,3253,6014,4282,728,6646,1415,20216,48,336,1617,5940,6139,427,3887,3243,2732,3124,1256,5615,4853,14686,2119,9926,11701,1169,107,20939,2905,3764,7049,3978,185,3512],[0,0,1869,0,0,234,0,0,0,0,621,1507,202,202,702,0,58,0,0,0,109,0,0,404,283,931,1151,0,1416,1274,0,0,196,852,604,709,0,1405,3099,0,6,0,2220,287,7795,765,463,1671,396,0,1604,962,1112,1761,623,303,0],[0,0,507,0,0,0,0,0,0,172,0,68,4,31,30,0,0,0,0,216,0,0,0,791,0,201,797,0,73,102,0,0,187,0,115,300,0,0,0,0,0,0,102,125,0,0,0,0,0,1009,1034,0,557,9,6,95,0],[15855,15216,11940,25596,14022,14733,19426,16602,8676,33673,32333,7073,12062,20217,18985,29394,12922,10898,22677,19597,9750,63764,63232,20095,19055,22617,19661,12201,26915,17490,6161,17263,39219,36420,41825,22161,12769,22117,7664,20196,30112,12255,16284,8833,13600,10407,14377,16468,11306,29595,157874,16856,41387,14681,8776,10182,3179],[0,0,833,307,55,140,0,0,116,116,155,449,474,0,42,0,30,200,0,257,0,462,24,41,0,0,12,0,116,0,0,77,688,255,1426,915,0,0,0,202,0,0,408,293,62,199,744,224,0,339,570,0,416,96,73,2,0],[0,26,250,0,0,0,0,0,0,0,50,0,0,0,0,48,0,23,129,43,0,0,0,0,0,0,0,49,0,0,0,0,0,0,0,0,0,0,0,0,0,0,0,114,0,112,0,0,0,0,0,0,0,0,0,0,0],[178,0,84,6,0,193,48,45,25,154,126,0,100,142,241,31,0,0,64,215,171,0,0,411,54,470,194,0,591,29,0,0,325,58,47,28,0,29,345,0,0,0,0,351,149,84,522,1118,0,218,0,0,345,295,48,401,0],[1466,1926,315,2494,1398,2480,1032,1118,1178,3199,1990,220,784,1033,489,1563,675,411,666,2468,2266,1885,3030,1633,825,3325,394,1255,2205,4223,0,152,544,1544,2591,3330,27,2808,391,347,813,811,4320,1077,1073,487,2177,776,606,2416,5952,609,1123,337,1531,701,325],[3759,4773,6654,12526,8565,19827,7605,7682,3938,25518,10492,2727,24250,9301,9728,10272,8010,6088,8999,5313,10939,10318,19429,6007,6843,17834,10424,4632,23570,20189,0,8450,8136,16448,16306,16188,790,7392,6794,9061,26905,10085,8106,3903,8956,9312,7802,5666,3937,3270,13559,11270,5897,5399,4222,2678,4207],[0,634,747,35,0,156,0,0,56,0,167,604,170,0,0,0,0,0,201,202,0,0,0,0,0,0,0,240,0,0,0,0,0,0,0,0,0,835,0,0,0,805,0,0,1796,0,593,200,0,0,0,0,0,171,164,0,608],[251,4024,1122,1798,838,1346,962,1887,5615,3768,4255,1914,3519,1591,2444,2167,768,27,2775,1794,3127,1203,2175,2855,2011,913,602,786,176,1759,0,2654,3431,3346,8348,3493,0,2342,3707,235,244,3013,1734,990,3489,1221,3364,1506,1011,1198,23735,2325,2213,782,151,631,3208],[298,320,1337,150,339,800,0,299,96,505,110,1990,168,706,300,103,0,889,918,0,207,292,6249,1195,6,722,390,246,1058,933,0,0,0,0,606,290,0,0,0,0,930,316,18,1594,286,485,294,2189,404,0,600,0,200,969,951,189,0],[0,0,124,0,0,0,0,0,0,0,6,59,0,0,0,0,0,0,4,0,0,0,0,0,0,0,0,0,25,0,0,0,492,276,84,0,0,0,0,0,0,0,0,12,0,0,16,370,0,0,0,16,82,0,0,0,0],[815,260,2486,193,99,67,193,467,72,864,390,584,394,282,0,354,0,419,201,23,23,269,0,733,54,794,588,0,553,1623,0,202,157,104,353,279,0,438,1369,831,0,705,837,994,141,102,1866,2681,0,0,12860,483,931,1224,430,0,603],[0,116,682,519,111,323,202,499,335,607,282,667,577,241,201,560,0,0,200,505,142,384,28,1212,0,1083,404,0,846,1622,0,200,0,167,404,100,0,326,0,0,202,0,115,499,985,199,0,2321,202,2315,0,0,93,403,176,0,0],[328,0,716,346,385,33,81,333,0,968,201,55,634,404,326,423,240,52,368,184,402,434,89,160,0,660,88,51,460,708,74,246,0,766,561,218,0,122,0,402,0,208,1030,0,0,320,0,0,0,0,196,317,248,402,0,29,0],[65,166,37,0,0,0,0,0,87,678,236,781,0,3,76,209,35,105,231,28,0,238,12,0,0,117,107,211,38,218,0,0,0,10,106,5,0,254,0,53,1119,2017,0,63,186,0,0,297,19,5,125,128,2,115,221,21,0],[7419,4126,3444,6301,3316,2729,2068,980,1991,6433,4420,3860,1034,5560,3723,2131,2668,648,4187,1561,7881,1043,241,784,2146,5609,1726,580,1467,430,4602,3648,709,3137,5325,729,582,1515,0,783,638,0,3376,1039,6495,1604,1744,416,676,649,15289,2083,536,0,2153,1306,6006],[54476,66098,14116,75755,56253,52788,27744,27658,76018,112222,75578,14201,40593,42176,46641,60444,21542,22004,43113,48565,38227,46508,66762,67878,39400,62931,33763,18160,58778,65607,11711,31525,29305,56741,73358,99054,28751,58857,50977,33967,28751,20295,50325,28383,53027,52986,19300,28128,37317,33523,350189,41067,42030,10462,37719,37707,56420],[3135,4981,5302,852,496,2735,548,1839,1292,7270,15485,5379,1766,1601,1351,10004,783,182,8861,12169,3840,15520,5010,3337,34746,7993,12033,1458,2777,8150,9516,4329,1443,3355,21517,17433,1263,1527,2363,4086,5748,7178,2788,13974,6780,25582,4749,2464,2079,6631,23306,2608,13090,6643,13418,988,0],[0,174,129,718,202,351,241,549,590,769,742,416,587,670,986,619,0,118,740,23,598,249,2378,199,0,656,0,0,841,4565,0,0,0,447,2211,505,606,438,0,496,0,0,0,4,294,58,201,810,525,829,0,203,401,486,128,0,532],[0,86,192,103,0,0,0,0,0,48,0,232,0,37,0,0,0,0,0,0,0,0,0,90,0,236,237,0,266,0,0,0,0,0,0,52,0,0,0,83,0,0,0,205,0,0,380,0,0,0,0,117,43,92,0,0,0],[44492,171697,18992,138307,161627,180033,68385,70355,115599,258560,161044,18108,40942,63191,55962,96440,26031,30359,133924,106558,146656,143128,95176,87637,72873,127100,40130,17316,112243,106852,37178,86142,46428,141083,109453,139006,41134,108669,70166,60690,67747,7097,136200,50789,24118,190542,30356,28790,71683,43811,335057,67164,83583,9513,68294,40597,51740],[928,1357,3912,570,1517,1176,757,653,1411,489,1698,3784,1736,1055,1020,833,232,1073,0,201,615,93,1313,3686,281,1703,1902,449,2030,1158,2229,305,315,460,1404,4146,0,3007,2564,1297,44,4057,3911,2721,8677,77,12326,6102,1895,0,13949,607,1972,2902,3356,1016,498],[34101,69667,6582,157422,110612,110376,66475,59418,108261,238224,234194,12180,61862,64524,53610,52623,11423,32605,100239,84823,171635,85241,28267,44024,82519,126240,77528,2344,75291,111202,22871,33762,79275,41351,63658,72619,80047,134739,59367,68581,78142,25666,80645,57697,31413,101959,60649,61516,71298,62561,392486,55206,89549,10892,16759,43637,55821],[980,2019,8418,2297,882,2498,1256,2261,1929,1804,3955,4577,5645,1659,1865,2970,1772,1279,1756,1837,3130,2823,1610,5141,1103,2911,5867,255,4040,8197,264,490,101,1208,2481,4378,57,3756,8327,1136,1764,9768,3652,5028,7374,3994,14151,12214,966,1043,4515,1705,3923,9233,1074,885,445],[606,80,920,30,169,96,0,200,0,32,299,1330,487,202,51,63,60,0,0,199,0,0,0,0,80,532,0,29,927,203,0,0,362,0,324,98,0,0,0,0,0,0,2160,470,355,637,2103,363,0,0,0,0,461,658,0,0,0],[4382,15430,12187,11772,7802,7390,7139,7737,8065,64331,13874,7472,6766,5081,3940,39831,5972,12137,3510,3010,4860,6675,2041,2852,4103,4097,1145,1988,6354,7429,366,1294,1390,3350,6904,9816,8335,6671,7598,10628,9611,13353,10161,9822,9303,6044,3534,6616,4460,4930,31967,3421,6254,2319,10965,21963,8628],[0,0,0,448,182,544,282,178,100,0,219,431,128,145,0,8,0,160,0,0,0,113,0,125,0,0,0,0,803,48,0,0,0,344,34,156,0,1039,712,0,78,0,0,0,0,1422,88,168,0,0,0,0,305,0,0,0,0],[1036,1207,2582,2415,1334,2305,1618,3644,1000,3860,4084,3462,2489,2476,1374,3250,440,850,3673,707,3273,2102,3676,2574,3207,4121,829,158,7517,2990,3782,1540,1066,2979,5023,9548,1366,751,7938,1454,0,2615,2706,1698,6889,2069,8663,4381,1181,131,9985,2487,3610,2228,920,759,0],[0,153,666,344,0,217,36,607,834,170,168,932,389,0,0,0,0,0,54,175,13,272,93,84,103,196,0,0,337,0,0,0,309,184,0,1632,0,0,0,0,0,0,0,405,1806,518,1072,203,335,0,9,765,401,1513,58,0,0],[52,227,46,726,377,337,75,46,202,384,584,16,199,396,631,84,68,120,438,872,404,1775,1932,703,11,498,1205,0,603,482,0,0,304,192,1028,74,0,681,0,32,0,0,0,101,14,279,0,160,15,26,117,0,323,743,377,120,41],[257,150,654,297,16,176,0,233,265,245,96,318,100,295,276,284,146,82,23,160,260,16,0,115,0,41,58,166,16,65,0,93,0,0,223,258,0,173,100,16,109,0,0,142,1236,1222,2113,0,0,0,691,76,94,180,115,132,209],[40541,59209,14977,75362,62100,72041,35109,39695,91537,144806,121699,24877,40224,46806,46062,37679,13475,25010,56957,60096,97580,62133,60515,120771,47345,82143,36117,16718,92100,162167,26260,40879,57846,64524,94207,104006,44069,103028,79291,39082,56670,30695,42568,34956,34706,60811,34001,58370,41126,28118,389617,44933,62406,16249,18881,20930,14512],[688,522,953,1163,460,1335,78,1511,202,1409,1648,511,498,1704,730,1749,1993,808,1186,387,2453,1944,2580,246,751,1732,698,87,1744,1241,0,213,594,1629,1523,1681,22,0,0,1778,96,0,1908,603,0,401,1004,1371,1482,1292,48,318,361,398,1099,98,3743],[100,201,430,34,138,23,226,404,482,111,434,744,648,125,0,0,0,0,0,292,0,404,0,2198,329,253,402,0,202,187,0,100,604,538,212,606,0,0,0,0,27,0,370,0,1437,744,780,49,0,447,0,101,161,118,196,751,0],[4915,1362,2546,3812,2673,3730,1628,1229,1715,10479,7076,3309,3221,2969,3652,1276,1048,2504,1112,1557,4025,1272,1005,3845,1895,3348,4977,42,3634,2855,2006,2137,1071,5327,7310,10863,995,5809,1533,1586,5063,0,10081,2770,2650,2335,2167,2828,2168,516,5528,2880,2608,769,2178,171,4337],[1813,1652,6183,1439,329,1662,327,476,2571,1740,945,2418,794,327,520,979,351,1302,616,1830,244,0,0,0,321,755,40,0,484,855,0,0,38,808,1432,841,0,2006,1053,951,1258,1152,1109,1148,4352,1522,3724,4622,263,604,3,3061,1507,3052,860,669,347],[0,0,189,0,0,0,0,0,404,340,258,273,0,0,0,29,0,0,0,0,0,0,0,1782,99,0,1615,0,1011,0,0,0,0,0,0,197,0,0,0,0,0,620,986,365,1168,0,1110,90,0,0,2102,202,164,149,441,0,0],[1780,2062,971,1964,2049,1945,745,977,2909,4288,2084,1583,1595,5255,2134,10044,2005,2024,1032,1260,1548,857,6207,5906,726,1195,502,884,2217,3303,97,56,579,666,951,2758,74,2376,1098,889,1428,0,1535,984,2512,119,3116,147,827,814,1759,1798,2953,1365,4505,951,358],[728,3297,1451,1733,1327,2994,240,1212,182,6083,1332,233,1278,247,742,1025,169,298,485,568,1872,429,200,624,606,743,275,371,1042,998,0,203,372,345,696,652,342,1510,1651,773,825,418,1594,912,102,102,252,107,230,118,1074,988,1560,340,751,993,699],[8,0,0,375,0,0,28,19,0,0,3,4,0,0,0,8,0,0,0,0,65,0,0,0,0,150,0,0,674,3,0,0,0,0,0,0,0,0,0,0,0,0,0,0,0,0,7,12,0,0,0,76,84,0,0,64,0],[14846,23760,16727,34216,24936,23177,27930,27617,39022,93540,39841,16736,17314,26546,23629,37547,16110,13837,37115,35867,46440,56607,43998,32215,33133,34023,16014,45223,23907,30928,20705,29930,13949,56923,38404,35078,35754,36387,54153,24880,25506,13006,86121,37137,33820,38402,14596,22787,21940,16848,166001,29639,19087,5401,22723,12612,30231],[226,0,613,357,24,90,3,0,0,139,91,424,437,0,113,99,0,34,0,149,0,32,0,0,49,72,216,31,0,84,0,0,276,0,1119,707,0,0,0,42,126,0,10,242,365,0,0,572,0,10,0,343,288,575,246,0,0],[953,1276,2065,3048,690,1585,2322,2758,2189,5818,7646,2382,5137,2516,2669,6661,659,1818,2608,3090,4152,4505,1697,2717,3741,3437,1019,1917,5645,4839,1354,453,1225,2647,2970,6307,4943,7361,801,4465,5297,48,4200,2626,4653,4770,6718,3152,1685,2699,24333,2717,4547,4012,782,2926,1],[494,629,2806,1758,2173,2318,1005,353,1897,3344,2168,2297,1022,1708,1271,1963,367,440,993,682,696,1052,663,5830,1123,2622,1438,1635,1099,606,0,387,1930,3596,4268,2520,56,2909,85,687,720,357,1401,3296,9961,1902,5381,3201,281,444,4305,1479,2717,3363,3209,901,1706],[0,0,1488,56,58,293,0,497,134,0,572,0,201,378,192,0,0,0,249,307,300,0,0,0,402,282,0,76,1181,592,0,0,0,0,1178,3354,0,0,0,0,0,0,1405,473,758,0,2006,501,0,825,35,0,1048,3732,0,25,0],[936,4796,1973,1276,763,1845,618,1100,411,8489,2164,1024,877,795,1200,2346,358,850,348,679,2493,1099,0,1074,403,969,114,493,297,1221,1,278,112,247,545,2417,593,1183,1582,466,894,133,546,1781,474,112,1034,1264,490,153,2800,914,2490,824,1030,1112,766],[16443,14701,10867,50956,34068,40302,29071,28077,21853,69798,74739,9271,33874,33342,35403,73740,29142,48612,33566,38689,52205,54923,45705,42559,26479,77360,48613,37432,57917,72154,8633,14756,30022,34674,109407,70932,35986,56766,32532,30325,57099,16758,22912,27812,19183,32433,23591,23208,30257,32387,201393,21984,47127,15541,24576,30120,50765],[8020,6896,4864,88255,67120,49525,24422,28832,39562,185513,64412,42716,17246,24529,30177,20591,4687,8878,59200,37781,42446,32718,36980,51630,19651,31193,6682,9391,145810,89529,15256,29317,21274,52604,44596,41989,13430,15387,55073,17369,28666,29341,45500,10804,8891,41390,19527,24200,13843,9139,100264,29954,29129,6386,5636,5567,1312],[995,258,802,642,624,691,1580,449,737,1445,0,361,656,703,324,565,0,3547,209,862,866,1862,3593,134,66,324,577,384,323,3060,0,136,0,66,2261,607,0,2199,719,639,3561,0,603,847,1377,767,69,75,788,170,3410,440,736,251,255,0,0],[674,3181,952,2531,1259,2077,2537,2905,2517,9755,3794,873,3209,1123,1351,5263,362,5518,494,715,1316,1390,1728,320,6919,6361,3804,4560,7258,5885,7933,6258,6337,4822,2861,1824,0,3101,512,2377,3030,2776,14701,6647,398,941,1487,0,4411,1678,499,8066,6587,744,2001,996,0],[49,32,0,52,0,101,0,0,109,85,222,20,10,20,166,56,0,10,0,0,51,0,0,0,4,4,0,0,14,18,0,15,31,0,0,24,4,66,0,0,11,0,0,0,0,0,3,257,12,10,4,93,22,0,0,0,0],[41479,51651,8192,64781,44877,76734,29743,32863,54433,129962,67859,13633,43985,51522,59217,56232,18601,43506,56534,36832,68101,76868,61632,53094,33063,124078,53548,16564,68303,218471,28551,53109,67814,63637,49355,69892,18730,122043,54742,31510,36825,9904,61594,24027,21512,51112,23615,26464,26003,25408,261073,39100,33866,13457,24652,19391,40421],[43666,67837,10209,101520,86923,75692,44903,63683,75132,208363,194910,28301,53362,54619,53298,53649,15893,25901,92321,70001,126909,88482,78109,109869,65170,110606,38100,7723,81405,106767,20480,50095,54598,83918,112019,123481,22827,111681,42618,47489,79796,22983,85577,69429,32727,63999,22443,28600,63440,52911,458797,45074,94351,12456,37137,31030,63524],[303,0,846,41,360,486,20,173,170,308,14,455,69,74,0,126,72,0,97,14,265,294,1386,357,502,205,0,0,118,26,0,0,0,294,130,159,0,19,65,134,0,170,64,642,1446,117,388,243,101,0,589,0,512,208,317,69,0],[20309,38050,1269,60658,46124,35184,42767,45217,24494,130377,121337,2269,52723,38363,41582,40045,13096,17076,84255,65086,91471,96363,28159,9037,63467,81427,35650,5601,66514,64040,14216,46119,38787,40390,78536,84232,26237,90403,17002,54053,53873,3263,82273,32933,6343,55198,17910,10053,40835,19675,93723,41647,53322,5460,9886,12350,6042],[9100,11505,1707,24773,25223,20424,14605,23871,7121,50929,45348,2676,18715,16600,19134,24199,5894,10496,26471,25183,46640,41365,34623,15871,28234,41686,16483,25267,28823,29433,12789,29017,23317,32143,53014,42017,10819,62007,13819,31181,17994,1638,135745,26707,1934,26862,15361,12678,19066,10381,49054,18203,26471,2017,8245,11235,14563],[1,162,0,63,0,0,4,0,136,216,9,84,31,28,9,59,17,4,0,105,20,0,36,2,52,0,0,5,0,11,5,3,7,2,183,0,10,0,0,0,0,0,318,0,398,4,0,0,0,0,15,38,76,0,65,16,27],[101948,201851,6152,312815,217600,272925,227259,223756,315459,681085,771612,19308,248385,201058,191611,306100,190750,161103,398030,224211,617792,557319,262052,118419,315573,451791,142590,75941,454157,673862,51754,256260,518397,311947,330112,275327,202705,503482,60569,246422,358190,25067,168511,99496,25836,278164,57744,30392,319555,220221,777127,197280,261868,22268,81055,153968,184125],[533,234,2266,590,111,1150,158,795,651,678,1273,2385,379,1150,1535,1125,149,375,27,200,172,506,1071,1495,544,1865,2101,54,1480,478,40,0,194,202,5253,2249,0,0,812,0,223,3105,2708,4655,4603,202,7463,3121,906,659,1528,2686,2186,3557,950,146,95],[1160,1133,1702,1894,1154,1373,2387,2006,431,2467,4839,417,1348,1230,254,1255,1616,1274,2021,1381,2643,4948,2401,3416,1735,2050,317,4122,3746,2466,0,333,623,2938,3079,3489,0,1375,731,1003,1814,0,1384,593,398,1127,2466,602,2244,2106,573,451,2189,427,70,894,489],[402,0,197,200,0,0,202,199,0,0,613,0,158,0,362,0,0,0,198,0,0,67,0,50,0,0,2005,0,95,0,0,108,0,0,20,1628,0,0,0,0,0,0,411,0,0,0,1166,398,0,0,0,0,99,190,306,0,0],[8010,7950,4618,15532,10517,13001,5255,8439,13750,16515,41019,4440,15163,10613,12153,20430,3555,12251,7060,16592,17316,21088,18042,20900,10098,21452,6657,1368,21356,28221,26,8774,13191,6636,20612,24402,2525,31134,16694,6220,20793,6262,4081,4568,10378,4417,11783,8171,9785,10535,62465,5902,26161,5493,5893,12716,10159],[49511,113861,8145,126791,89527,118878,85645,73185,53434,271129,305522,11262,98076,132799,138125,122576,44908,94884,121971,111005,179469,172653,60782,54632,118847,313454,107244,32598,221398,542724,8795,88333,187368,111407,177534,137606,70164,326970,38276,93574,133171,5263,100237,52364,22230,118212,52601,20825,103259,66734,368349,74402,112895,15100,22551,45079,101697],[698,654,2815,1726,2047,1803,1535,892,231,2204,5114,2984,1354,2034,1665,1073,1986,2591,3054,1093,1108,816,1859,680,2246,1415,1857,742,1576,2640,2807,199,199,859,4088,2796,0,1533,0,6,67,130,1942,4675,4725,1775,6690,1898,1090,259,14557,2284,2590,1290,1471,908,131],[0,368,610,260,164,489,1418,689,754,634,1483,199,2224,2084,534,6125,388,975,2283,929,742,5655,7449,340,1556,1332,1759,206,2311,2483,0,1464,1501,947,1350,1639,118,1921,0,1926,3643,0,1381,64,0,849,1010,0,173,1087,4757,1872,1849,907,430,138,272],[2687,1928,1148,1026,57,1537,841,745,2150,2279,1760,774,2086,785,272,1088,56,1723,1053,562,2180,2676,2843,5789,2712,1818,2114,0,3066,5685,73,1609,775,2294,2415,4149,1318,4394,3335,1382,2123,218,5514,655,4772,362,70,3920,1057,893,7870,1867,1889,1648,1287,1462,956],[1409,3304,24360,33579,9921,25997,786,2809,1192,21912,4176,8898,3309,2052,3637,18183,4614,12352,4204,2654,16187,1830,553,3395,723,613,143,21068,1385,3118,109,124,2019,2187,6715,10763,1040,698,8579,816,1148,27170,1279,6300,2824,2562,778,1808,416,404,9185,1100,638,2047,5447,2892,3988],[281,1356,2033,1007,362,922,260,285,577,1232,566,857,142,362,367,420,0,213,105,192,726,295,428,466,200,267,670,186,692,2359,0,0,1198,405,65,760,0,491,1616,151,677,0,2440,1745,2308,693,2427,866,153,137,3181,699,950,459,55,221,843],[122,94,907,44,326,0,335,832,721,65,328,664,230,90,82,308,0,0,456,246,667,161,0,0,124,523,312,0,1287,752,0,37,0,101,997,124,0,74,702,10,178,0,1577,159,1465,551,805,516,192,30,1015,892,635,275,0,225,0],[2075,2392,544,3041,1853,1947,1290,1072,2270,4242,5837,647,1206,1202,1968,1240,582,70,1383,1055,1789,1376,706,1258,865,4544,838,469,1110,1604,0,1481,1398,747,3827,2664,122,998,1812,893,3291,0,2199,1911,550,3211,686,956,732,1091,1927,930,1590,107,2136,985,6591],[8490,12732,10969,35648,27535,34746,32111,32853,20876,65025,86648,8748,44386,42383,45825,72262,23792,255087,38290,15163,48729,69342,44769,12786,40132,27059,31330,12523,44024,66741,23501,29842,28532,43966,88125,53491,30697,62496,20973,32335,36695,41801,24366,19293,15806,24907,29148,14978,31811,35022,156632,36356,49245,14731,12451,22984,6051],[480,0,541,270,0,0,0,0,146,0,140,1031,0,0,0,0,0,0,0,0,159,0,1530,802,201,0,105,0,596,213,0,0,264,0,0,4186,0,0,0,0,0,0,0,446,0,402,1847,557,0,0,0,404,0,197,0,0,0],[218,839,563,0,194,589,337,225,0,888,481,137,364,205,0,127,0,2725,0,0,217,245,87,920,0,172,0,132,1135,307,0,804,130,292,593,344,0,0,1690,42,0,0,802,918,592,64,147,237,0,602,0,372,896,387,0,64,0],[22,90,480,122,42,0,273,0,0,0,182,0,202,478,0,0,26,0,0,0,0,422,0,1160,19,146,0,100,0,635,52,29,0,77,184,68,0,28,0,0,0,0,0,164,0,179,34,25,0,236,0,0,70,16,111,0,0],[1450,511,6322,3030,2145,2972,1833,2189,2416,4272,3582,3623,2080,802,1551,266,138,700,1062,1277,723,6593,2315,3033,647,1030,389,1375,7899,11179,838,0,202,691,3146,3609,1866,2902,7565,2039,197,0,3285,2671,5776,1062,9335,5367,1345,1207,6901,4493,5158,5156,1870,1234,23],[1143,1153,1182,2385,1342,2093,726,550,381,3710,1702,384,2482,2833,4665,11266,4750,2634,1126,2488,2817,1982,789,3061,481,972,1465,1820,2740,2688,193,346,778,1667,2691,7658,621,853,1533,966,1805,470,191,431,1880,1569,547,3188,867,307,4421,1566,1634,0,901,1346,2093],[218815,396952,24710,423974,283063,318373,350058,388489,175885,893841,1279294,34303,433627,388094,458336,527214,261484,283649,434622,319270,641751,546814,539077,529250,282607,755678,305025,199876,578954,745620,118155,307652,387436,365563,628388,557117,384218,789937,114668,319327,437340,26351,243112,197272,76909,224191,141892,74093,389979,358577,1553013,255490,494262,49335,104522,221457,369330],[19619,27820,25719,48614,37920,63443,22195,26090,21969,72445,60232,25803,32902,38847,42121,22984,9520,20222,23777,32230,36554,28518,45340,52391,26829,113624,59983,3287,49942,85283,9371,25912,34332,24617,59990,69769,5422,57846,66750,14773,24888,27520,25135,24541,69457,25049,27998,34777,14419,19708,304816,24430,37233,16185,12625,24190,16748],[0,380,833,0,404,211,0,179,183,202,454,82,0,0,51,31,29,27,0,0,0,0,0,1353,0,543,0,311,404,0,0,0,0,133,53,0,0,0,0,0,0,34,0,0,0,401,0,1108,0,360,30,201,46,1574,36,0,0],[78,0,184,0,0,0,0,9,0,2,0,107,0,0,32,0,0,2,0,0,0,0,0,191,0,137,0,0,116,0,0,0,0,0,0,0,0,0,0,0,0,0,0,232,627,0,32,430,184,0,2031,26,280,111,0,0,0],[18460,5498,1333,82,617,2125,6649,11144,1084,821,1606,494,2770,1313,5976,58,505,39,0,4246,12772,3321,64044,69525,46,8212,6833,844,27422,4197,7959,7962,14667,6170,20270,53997,0,1303,266,357,1504,0,899,1014,1232,123,1879,423,76,0,4021,26230,87751,0,624,11432,140],[39729,62572,11742,39732,30687,36946,21257,26878,70268,93164,68541,14083,26030,31637,35610,61382,29364,44517,28532,43236,40339,39579,45050,66536,25074,48473,20306,29228,53187,73143,14008,22426,19093,39973,44330,58061,30553,65580,51286,21891,23950,10614,45717,21692,24338,24379,18734,23122,30779,21980,307060,28697,33645,9664,33094,29861,47270],[6247,5503,2469,7219,5887,8198,2669,4811,1789,14862,15089,1321,4064,3968,2807,9015,1434,2282,4437,4569,6743,4475,3022,8990,4793,4180,2005,1039,594
```
