## Supplemental Examples for "R/LinkedCharts: A novel approach for simple but powerful interactive data analysis": JS_code_full.html

```
var selGene = 1914;

lc.scatter()
   .x(i => maData.pvals[i].AveExpr)
   .y(i => maData.pvals[i].tissuetumour)
   .colour(i => maData.pvals[i]["adj.P.Val"] < 0.1 ? "red" : "black")
   .label(i => maData.geneNames[i])
   .size(1.3)
   .axisTitleY("log2 fc")
   .axisTitleX("Average expression")
   .on_click(i => {selGene = i; exprPlot.update();})
   //here, a CSS selector for an existing DOM element is used
   .place("#ma");

//the chart object is stored in a variable to update it later
var exprPlot = lc.scatter()
   //original data contains also information on dysplasia tissue sample
   //remove them for plot's clarity
   .elementIds(d3.range(maData.patients.length)
      .filter((e, i) => maData.tissue[i] != "dysplasia")
   )
   .x(i => maData.patients[i])
   //raw counts are normalised on-the-fly
   .y(i => maData.countMatrix[selGene][i]/maData.countSums[i] * 1e6 + .1)
   .logScaleY(10)
   .colourValue(i => maData.tissue[i])
   .title(() => maData.geneNames[selGene])
   .axisTitleY("CPM")
   .axisTitleX("Patients")
   .ticksRotateX(45)
   .colourLegendTitle("Tissue")
   .legend.width(120)
   //here, a CSS selector for an existing DOM element is used
   .place("#expr");
```
