## Supplemental Examples for "R/LinkedCharts: A novel approach for simple but powerful interactive data analysis": JS_code_min.html

```
var selGene = 1914;

lc.scatter()
   .x(i => maData.pvals[i].AveExpr)
   .y(i => maData.pvals[i].tissuetumour)
   .colour(i => maData.pvals[i]["adj.P.Val"] < 0.1 ? "red" : "black")
   .on_click(i => {selGene = i; exprPlot.update();})
   //here, a CSS selector for an existing DOM element is used
   .place("#ma");

var exprPlot = lc.scatter()
   .x(i => maData.patients[i])
   //raw counts are normalised on-the-fly
   .y(i => maData.countMatrix[selGene][i]/maData.countSums[i] * 1e6 + .1)
   .logScaleY(10)
   .colourValue(i => maData.tissue[i])
   .title(() => maData.geneNames[selGene])
   //here, a CSS selector for an existing DOM element is used
   .place("#expr");
```
