## Supplemental Examples for "R/LinkedCharts: A novel approach for simple but powerful interactive data analysis": maData.js

```
var maData = {"pvals":[{"tissuedysplasia":-0.6794,"tissuetumour":-1.176,"AveExpr":3.9404,"F":17.37,"P.Value":2.751e-06,"adj.P.Val":0.0001},{"tissuedysplasia":-2.2216,"tissuetumour":0.2941,"AveExpr":-4.6907,"F":4.1642,"P.Value":0.0221,"adj.P.Val":0.0735},{"tissuedysplasia":0.3354,"tissuetumour":0.1422,"AveExpr":3.5513,"F":1.6599,"P.Value":0.2018,"adj.P.Val":0.3391},{"tissuedysplasia":0.2533,"tissuetumour":0.0661,"AveExpr":4.213,"F":2.1529,"P.Value":0.1282,"adj.P.Val":0.2493},{"tissuedysplasia":0.3423,"tissuetumour":0.327,"AveExpr":3.3508,"F":3.2623,"P.Value":0.0477,"adj.P.Val":0.1265},{"tissuedysplasia":0.6687,"tissuetumour":1.0619,"AveExpr":2.6732,"F":7.0421,"P.Value":0.0022,"adj.P.Val":0.014},{"tissuedysplasia":0.0972,"tissuetumour":-0.3968,"AveExpr":6.2535,"F":1.7272,"P.Value":0.1914,"adj.P.Val":0.3276},{"tissuedysplasia":0.4812,"tissuetumour":0.4067,"AveExpr":3.1861,"F":5.2355,"P.Value":0.0091,"adj.P.Val":0.0388},{"tissuedysplasia":0.2558,"tissuetumour":-0.0271,"AveExpr":5.9582,"F":2.4765,"P.Value":0.0957,"adj.P.Val":0.2046},{"tissuedysplasia":0.3018,"tissuetumour":-0.11,"AveExpr":4.7115,"F":5.4503,"P.Value":0.0077,"adj.P.Val":0.0343},{"tissuedysplasia":0.0097,"tissuetumour":0.1029,"AveExpr":3.1409,"F":0.3194,"P.Value":0.7282,"adj.P.Val":0.806},{"tissuedysplasia":-0.0122,"tissuetumour":0.0066,"AveExpr":5.1223,"F":0.0151,"P.Value":0.985,"adj.P.Val":0.9895},{"tissuedysplasia":0.0744,"tissuetumour":0.6569,"AveExpr":4.5473,"F":19.8064,"P.Value":7.3409e-07,"adj.P.Val":0},{"tissuedysplasia":0.6171,"tissuetumour":0.4453,"AveExpr":1.5843,"F":1.9153,"P.Value":0.1594,"adj.P.Val":0.2892},{"tissuedysplasia":-0.601,"tissuetumour":-0.5704,"AveExpr":4.8491,"F":11.6899,"P.Value":0.0001,"adj.P.Val":0.0012},{"tissuedysplasia":-0.9799,"tissuetumour":-0.2983,"AveExpr":0.9076,"F":3.1256,"P.Value":0.0538,"adj.P.Val":0.1375},{"tissuedysplasia":0.2018,"tissuetumour":0.003,"AveExpr":5.5454,"F":2.6674,"P.Value":0.0806,"adj.P.Val":0.1817},{"tissuedysplasia":0.0501,"tissuetumour":-0.5107,"AveExpr":5.5162,"F":9.164,"P.Value":0.0005,"adj.P.Val":0.0044},{"tissuedysplasia":0.0951,"tissuetumour":-0.0043,"AveExpr":5.3078,"F":0.4433,"P.Value":0.6447,"adj.P.Val":0.7435},{"tissuedysplasia":0.08,"tissuetumour":-0.1436,"AveExpr":3.5096,"F":1.4712,"P.Value":0.2407,"adj.P.Val":0.3829},{"tissuedysplasia":1.2006,"tissuetumour":2.4322,"AveExpr":-0.2895,"F":8.694,"P.Value":0.0007,"adj.P.Val":0.0057},{"tissuedysplasia":0.1158,"tissuetumour":0.5503,"AveExpr":1.2126,"F":2.25,"P.Value":0.1174,"adj.P.Val":0.2351},{"tissuedysplasia":0.7189,"tissuetumour":1.3344,"AveExpr":5.1654,"F":22.8772,"P.Value":1.5434e-07,"adj.P.Val":9.6735e-06},{"tissuedysplasia":-0.2004,"tissuetumour":0.1418,"AveExpr":5.7103,"F":4.3453,"P.Value":0.019,"adj.P.Val":0.066},{"tissuedysplasia":-0.006,"tissuetumour":0.4944,"AveExpr":-12.1139,"F":0.7767,"P.Value":0.4661,"adj.P.Val":0.5975},{"tissuedysplasia":-0.3633,"tissuetumour":-0.7876,"AveExpr":3.1332,"F":6.1913,"P.Value":0.0043,"adj.P.Val":0.0224},{"tissuedysplasia":0.1226,"tissuetumour":0.5619,"AveExpr":0.625,"F":1.022,"P.Value":0.3683,"adj.P.Val":0.511},{"tissuedysplasia":0.433,"tissuetumour":0.4198,"AveExpr":-1.4719,"F":0.3896,"P.Value":0.6797,"adj.P.Val":0.7701},{"tissuedysplasia":0.9218,"tissuetumour":2.1756,"AveExpr":1.4306,"F":20.1951,"P.Value":5.9888e-07,"adj.P.Val":0},{"tissuedysplasia":0.3477,"tissuetumour":0.9395,"AveExpr":4.2892,"F":15.3191,"P.Value":8.9262e-06,"adj.P.Val":0.0002},{"tissuedysplasia":0.3079,"tissuetumour":1.0156,"AveExpr":6.0015,"F":44.7715,"P.Value":2.4648e-11,"adj.P.Val":1.7881e-08},{"tissuedysplasia":0.1502,"tissuetumour":0.366,"AveExpr":2.9819,"F":2.2262,"P.Value":0.12,"adj.P.Val":0.2383},{"tissuedysplasia":0.4161,"tissuetumour":1.249,"AveExpr":2.1996,"F":17.446,"P.Value":2.6367e-06,"adj.P.Val":0.0001},{"tissuedysplasia":0.2934,"tissuetumour":0.4415,"AveExpr":5.5407,"F":9.7676,"P.Value":0.0003,"adj.P.Val":0.0033},{"tissuedysplasia":0.2508,"tissuetumour":0.2715,"AveExpr":1.9122,"F":0.5119,"P.Value":0.6029,"adj.P.Val":0.711},{"tissuedysplasia":0.2101,"tissuetumour":1.0972,"AveExpr":3.5826,"F":9.0179,"P.Value":0.0005,"adj.P.Val":0.0048},{"tissuedysplasia":-0.2395,"tissuetumour":-0.5366,"AveExpr":2.8217,"F":2.3408,"P.Value":0.1081,"adj.P.Val":0.2223},{"tissuedysplasia":-0.7939,"tissuetumour":-1.2698,"AveExpr":1.7468,"F":14.2847,"P.Value":0,"adj.P.Val":0.0003},{"tissuedysplasia":0.3455,"tissuetumour":0.4944,"AveExpr":4.6141,"F":9.2969,"P.Value":0.0004,"adj.P.Val":0.0041},{"tissuedysplasia":0.6015,"tissuetumour":0.477,"AveExpr":4.7712,"F":6.235,"P.Value":0.0041,"adj.P.Val":0.0219},{"tissuedysplasia":0.2558,"tissuetumour":0.0931,"AveExpr":8.2263,"F":2.7307,"P.Value":0.0771,"adj.P.Val":0.1761},{"tissuedysplasia":-0.142,"tissuetumour":-0.3259,"AveExpr":5.0329,"F":1.9533,"P.Value":0.1539,"adj.P.Val":0.2825},{"tissuedysplasia":0.1995,"tissuetumour":0.0048,"AveExpr":3.4167,"F":0.8534,"P.Value":0.4329,"adj.P.Val":0.57},{"tissuedysplasia":-0.129,"tissuetumour":-0.2082,"AveExpr":6.895,"F":3.9224,"P.Value":0.0271,"adj.P.Val":0.0849},{"tissuedysplasia":-0.1167,"tissuetumour":-0.5808,"AveExpr":2.5118,"F":3.173,"P.Value":0.0516,"adj.P.Val":0.1335},{"tissuedysplasia":-0.3589,"tissuetumour":-0.993,"AveExpr":4.284,"F":9.4135,"P.Value":0.0004,"adj.P.Val":0.0039},{"tissuedysplasia":-0.0398,"tissuetumour":0.4608,"AveExpr":4.4988,"F":5.0344,"P.Value":0.0107,"adj.P.Val":0.0437},{"tissuedysplasia":-0.0266,"tissuetumour":0.3212,"AveExpr":3.6155,"F":2.1335,"P.Value":0.1305,"adj.P.Val":0.2525},{"tissuedysplasia":0.0699,"tissuetumour":0.2416,"AveExpr":4.3354,"F":2.1306,"P.Value":0.1309,"adj.P.Val":0.253},{"tissuedysplasia":0.4824,"tissuetumour":1.1384,"AveExpr":5.2767,"F":28.8409,"P.Value":9.9149e-09,"adj.P.Val":1.3832e-06},{"tissuedysplasia":0.1506,"tissuetumour":0.1827,"AveExpr":5.5321,"F":2.273,"P.Value":0.115,"adj.P.Val":0.2318},{"tissuedysplasia":1.272,"tissuetumour":1.7273,"AveExpr":3.8124,"F":21.1234,"P.Value":3.7102e-07,"adj.P.Val":0},{"tissuedysplasia":0.0823,"tissuetumour":0.2342,"AveExpr":5.9377,"F":1.6472,"P.Value":0.2042,"adj.P.Val":0.3419},{"tissuedysplasia":0.3694,"tissuetumour":0.5167,"AveExpr":5.6526,"F":10.0152,"P.Value":0.0003,"adj.P.Val":0.0029},{"tissuedysplasia":-0.0649,"tissuetumour":-0.1711,"AveExpr":6.8417,"F":1.4019,"P.Value":0.2569,"adj.P.Val":0.3999},{"tissuedysplasia":0.0884,"tissuetumour":0.6585,"AveExpr":3.1669,"F":6.2331,"P.Value":0.0041,"adj.P.Val":0.0219},{"tissuedysplasia":0.3808,"tissuetumour":0.4068,"AveExpr":4.7084,"F":3.4756,"P.Value":0.0397,"adj.P.Val":0.111},{"tissuedysplasia":0.1304,"tissuetumour":-0.2709,"AveExpr":4.0111,"F":3.3265,"P.Value":0.0451,"adj.P.Val":0.1215},{"tissuedysplasia":-0.8795,"tissuetumour":0.6801,"AveExpr":3.5824,"F":4.1561,"P.Value":0.0234,"adj.P.Val":0.0766},{"tissuedysplasia":-0.2215,"tissuetumour":0.5377,"AveExpr":1.7155,"F":4.2912,"P.Value":0.0198,"adj.P.Val":0.0682},{"tissuedysplasia":0.2711,"tissuetumour":0.1246,"AveExpr":3.0141,"F":1.0669,"P.Value":0.3528,"adj.P.Val":0.4965},{"tissuedysplasia":-0.8633,"tissuetumour":-1.1458,"AveExpr":5.2088,"F":6.7966,"P.Value":0.0029,"adj.P.Val":0.0171},{"tissuedysplasia":1.078,"tissuetumour":2.5265,"AveExpr":-3.0704,"F":4.4467,"P.Value":0.0174,"adj.P.Val":0.0621},{"tissuedysplasia":0.3342,"tissuetumour":1.7029,"AveExpr":-1.0247,"F":7.261,"P.Value":0.0019,"adj.P.Val":0.0124},{"tissuedysplasia":0.2636,"tissuetumour":1.2372,"AveExpr":0.4922,"F":5.2781,"P.Value":0.0088,"adj.P.Val":0.0379},{"tissuedysplasia":0.2716,"tissuetumour":1.9375,"AveExpr":-3.3947,"F":2.157,"P.Value":0.1277,"adj.P.Val":0.2487},{"tissuedysplasia":0.2912,"tissuetumour":-0.0111,"AveExpr":3.7649,"F":2.2942,"P.Value":0.1128,"adj.P.Val":0.2289},{"tissuedysplasia":0.3483,"tissuetumour":0.4617,"AveExpr":3.8692,"F":8.115,"P.Value":0.001,"adj.P.Val":0.0077},{"tissuedysplasia":0.2178,"tissuetumour":0.0424,"AveExpr":5.5311,"F":2.0199,"P.Value":0.1448,"adj.P.Val":0.271},{"tissuedysplasia":0.2898,"tissuetumour":0.9558,"AveExpr":0.5041,"F":5.1357,"P.Value":0.0099,"adj.P.Val":0.0411},{"tissuedysplasia":0.0593,"tissuetumour":1.4277,"AveExpr":-2.3169,"F":2.9184,"P.Value":0.0645,"adj.P.Val":0.1558},{"tissuedysplasia":0.1981,"tissuetumour":0.0645,"AveExpr":3.138,"F":0.8752,"P.Value":0.4239,"adj.P.Val":0.5618},{"tissuedysplasia":-0.0757,"tissuetumour":0.3627,"AveExpr":3.6876,"F":3.3954,"P.Value":0.0425,"adj.P.Val":0.1166},{"tissuedysplasia":-0.4622,"tissuetumour":-0.3836,"AveExpr":3.815,"F":1.2584,"P.Value":0.2941,"adj.P.Val":0.4385},{"tissuedysplasia":0.1444,"tissuetumour":0.6365,"AveExpr":5.5158,"F":15.6105,"P.Value":7.5225e-06,"adj.P.Val":0.0002},{"tissuedysplasia":1.1025,"tissuetumour":0.7652,"AveExpr":4.0684,"F":20.4899,"P.Value":5.1382e-07,"adj.P.Val":0},{"tissuedysplasia":0.1112,"tissuetumour":0.1286,"AveExpr":5.4663,"F":0.4232,"P.Value":0.6576,"adj.P.Val":0.7535},{"tissuedysplasia":0.6194,"tissuetumour":0.5509,"AveExpr":3.2402,"F":6.1437,"P.Value":0.0044,"adj.P.Val":0.023},{"tissuedysplasia":2.557,"tissuetumour":3.0794,"AveExpr":-4.889,"F":4.0261,"P.Value":0.0248,"adj.P.Val":0.0797},{"tissuedysplasia":0.131,"tissuetumour":0.353,"AveExpr":3.6756,"F":3.5158,"P.Value":0.0383,"adj.P.Val":0.1083},{"tissuedysplasia":0.1928,"tissuetumour":0.3805,"AveExpr":4.5116,"F":4.9575,"P.Value":0.0114,"adj.P.Val":0.0455},{"tissuedysplasia":0.5192,"tissuetumour":0.7947,"AveExpr":0.9823,"F":2.1403,"P.Value":0.1297,"adj.P.Val":0.2513},{"tissuedysplasia":0.11,"tissuetumour":0.2556,"AveExpr":3.0724,"F":0.6221,"P.Value":0.5415,"adj.P.Val":0.6612},{"tissuedysplasia":-0.0327,"tissuetumour":-0.1638,"AveExpr":5.4942,"F":1.6344,"P.Value":0.2067,"adj.P.Val":0.3448},{"tissuedysplasia":0.2067,"tissuetumour":-0.0664,"AveExpr":3.8602,"F":1.3964,"P.Value":0.2582,"adj.P.Val":0.4014},{"tissuedysplasia":0.4386,"tissuetumour":0.5135,"AveExpr":1.6766,"F":1.0851,"P.Value":0.3467,"adj.P.Val":0.4907},{"tissuedysplasia":0.3692,"tissuetumour":0.2538,"AveExpr":2.0262,"F":1.2998,"P.Value":0.2829,"adj.P.Val":0.4273},{"tissuedysplasia":0.1481,"tissuetumour":0.5832,"AveExpr":3.7167,"F":9.2106,"P.Value":0.0005,"adj.P.Val":0.0043},{"tissuedysplasia":0.1159,"tissuetumour":0.9206,"AveExpr":3.8857,"F":15.775,"P.Value":6.8338e-06,"adj.P.Val":0.0002},{"tissuedysplasia":0.0909,"tissuetumour":0.878,"AveExpr":3.6958,"F":12.834,"P.Value":0,"adj.P.Val":0.0007},{"tissuedysplasia":0.7281,"tissuetumour":1.5666,"AveExpr":2.2178,"F":23.0268,"P.Value":1.4345e-07,"adj.P.Val":9.1993e-06},{"tissuedysplasia":0.4029,"tissuetumour":-0.0711,"AveExpr":1.7544,"F":1.7143,"P.Value":0.1919,"adj.P.Val":0.3281},{"tissuedysplasia":0.2728,"tissuetumour":0.2951,"AveExpr":4.537,"F":4.0871,"P.Value":0.0235,"adj.P.Val":0.0769},{"tissuedysplasia":0.1809,"tissuetumour":0.1298,"AveExpr":7.0931,"F":2.329,"P.Value":0.1093,"adj.P.Val":0.2239},{"tissuedysplasia":0.1657,"tissuetumour":0.3404,"AveExpr":3.036,"F":0.8992,"P.Value":0.4142,"adj.P.Val":0.5529},{"tissuedysplasia":1.3924,"tissuetumour":2.2189,"AveExpr":-4.3287,"F":2.9527,"P.Value":0.0626,"adj.P.Val":0.1527},{"tissuedysplasia":0.9826,"tissuetumour":1.9477,"AveExpr":-2.3044,"F":3.6816,"P.Value":0.0332,"adj.P.Val":0.0978},{"tissuedysplasia":0.0874,"tissuetumour":-0.2469,"AveExpr":3.8475,"F":2.817,"P.Value":0.0706,"adj.P.Val":0.1659},{"tissuedysplasia":0.2469,"tissuetumour":1.0052,"AveExpr":2.2787,"F":11.1478,"P.Value":0.0001,"adj.P.Val":0.0016},{"tissuedysplasia":-0.2216,"tissuetumour":-0.8461,"AveExpr":4.2728,"F":8.4809,"P.Value":0.0008,"adj.P.Val":0.0064},{"tissuedysplasia":0.2467,"tissuetumour":0.5875,"AveExpr":1.435,"F":1.5216,"P.Value":0.2296,"adj.P.Val":0.3708},{"tissuedysplasia":0.0744,"tissuetumour":-0.2097,"AveExpr":7.1144,"F":5.7612,"P.Value":0.006,"adj.P.Val":0.0287},{"tissuedysplasia":0.2957,"tissuetumour":0.6594,"AveExpr":2.6155,"F":8.0278,"P.Value":0.0011,"adj.P.Val":0.0081},{"tissuedysplasia":1.0802,"tissuetumour":2.5614,"AveExpr":-3.1011,"F":4.0876,"P.Value":0.0235,"adj.P.Val":0.0769},{"tissuedysplasia":-0.0116,"tissuetumour":-0.3901,"AveExpr":5.5082,"F":6.475,"P.Value":0.0034,"adj.P.Val":0.0191},{"tissuedysplasia":0.2428,"tissuetumour":0.1604,"AveExpr":3.8955,"F":1.62,"P.Value":0.2095,"adj.P.Val":0.348},{"tissuedysplasia":0.0692,"tissuetumour":-0.3522,"AveExpr":8.3947,"F":6.2749,"P.Value":0.0043,"adj.P.Val":0.0226},{"tissuedysplasia":0.0865,"tissuetumour":-0.261,"AveExpr":5.1988,"F":2.9827,"P.Value":0.061,"adj.P.Val":0.15},{"tissuedysplasia":1.2017,"tissuetumour":1.0767,"AveExpr":4.1178,"F":11.7271,"P.Value":0.0001,"adj.P.Val":0.0012},{"tissuedysplasia":0.319,"tissuetumour":0.7457,"AveExpr":2.681,"F":6.1054,"P.Value":0.0046,"adj.P.Val":0.0235},{"tissuedysplasia":0.5356,"tissuetumour":1.5344,"AveExpr":6.4075,"F":18.9189,"P.Value":1.7887e-06,"adj.P.Val":0.0001},{"tissuedysplasia":0.1123,"tissuetumour":-0.2392,"AveExpr":5.3538,"F":4.23,"P.Value":0.0209,"adj.P.Val":0.0707},{"tissuedysplasia":0.096,"tissuetumour":-0.2444,"AveExpr":6.3759,"F":2.0002,"P.Value":0.1474,"adj.P.Val":0.2745},{"tissuedysplasia":0.748,"tissuetumour":1.9799,"AveExpr":-0.6767,"F":8.8525,"P.Value":0.0006,"adj.P.Val":0.0052},{"tissuedysplasia":-0.326,"tissuetumour":2.2716,"AveExpr":-2.4425,"F":5.1809,"P.Value":0.0095,"adj.P.Val":0.0401},{"tissuedysplasia":0.0387,"tissuetumour":-0.1196,"AveExpr":4.5088,"F":0.5547,"P.Value":0.5782,"adj.P.Val":0.6908},{"tissuedysplasia":0.1702,"tissuetumour":0.959,"AveExpr":2.7555,"F":14.1049,"P.Value":0,"adj.P.Val":0.0004},{"tissuedysplasia":0.0399,"tissuetumour":0.3744,"AveExpr":2.1735,"F":2.0111,"P.Value":0.146,"adj.P.Val":0.2725},{"tissuedysplasia":0.5508,"tissuetumour":1.11,"AveExpr":1.875,"F":10.3542,"P.Value":0.0002,"adj.P.Val":0.0024},{"tissuedysplasia":0.0582,"tissuetumour":0.3112,"AveExpr":2.4317,"F":1.3456,"P.Value":0.2709,"adj.P.Val":0.4149},{"tissuedysplasia":-0.4855,"tissuetumour":-0.503,"AveExpr":-0.8506,"F":0.5453,"P.Value":0.5835,"adj.P.Val":0.6951},{"tissuedysplasia":0.3529,"tissuetumour":-1.7321,"AveExpr":-3.265,"F":4.3265,"P.Value":0.0193,"adj.P.Val":0.0668},{"tissuedysplasia":0.0876,"tissuetumour":0.3474,"AveExpr":3.8429,"F":3.2626,"P.Value":0.0477,"adj.P.Val":0.1265},{"tissuedysplasia":0.49,"tissuetumour":1.914,"AveExpr":-2.1641,"F":5.6327,"P.Value":0.0066,"adj.P.Val":0.0309},{"tissuedysplasia":1.2467,"tissuetumour":3.2482,"AveExpr":-3.3676,"F":8.5367,"P.Value":0.0007,"adj.P.Val":0.0062},{"tissuedysplasia":1.2829,"tissuetumour":2.3465,"AveExpr":3.9499,"F":44.2508,"P.Value":2.9281e-11,"adj.P.Val":2.0724e-08},{"tissuedysplasia":0.3046,"tissuetumour":0.4828,"AveExpr":5.7077,"F":6.4394,"P.Value":0.0035,"adj.P.Val":0.0195},{"tissuedysplasia":0.0657,"tissuetumour":1.7113,"AveExpr":-4.5749,"F":1.9107,"P.Value":0.1601,"adj.P.Val":0.29},{"tissuedysplasia":0.3785,"tissuetumour":0.716,"AveExpr":4.173,"F":16.9513,"P.Value":3.4807e-06,"adj.P.Val":0.0001},{"tissuedysplasia":0.6851,"tissuetumour":0.7114,"AveExpr":2.9318,"F":5.5417,"P.Value":0.0071,"adj.P.Val":0.0326},{"tissuedysplasia":0.3744,"tissuetumour":0.967,"AveExpr":3.1385,"F":15.3977,"P.Value":8.5223e-06,"adj.P.Val":0.0002},{"tissuedysplasia":-0.1139,"tissuetumour":0.4325,"AveExpr":0.9681,"F":1.3153,"P.Value":0.2787,"adj.P.Val":0.4233},{"tissuedysplasia":0.782,"tissuetumour":2.3592,"AveExpr":3.6298,"F":26.1418,"P.Value":3.2925e-08,"adj.P.Val":3.2713e-06},{"tissuedysplasia":1.2973,"tissuetumour":0.9987,"AveExpr":-0.4215,"F":4.9599,"P.Value":0.0114,"adj.P.Val":0.0454},{"tissuedysplasia":-0.2099,"tissuetumour":-0.3132,"AveExpr":5.9578,"F":2.3357,"P.Value":0.1086,"adj.P.Val":0.2231},{"tissuedysplasia":-0.3269,"tissuetumour":-0.4553,"AveExpr":5.4796,"F":4.6072,"P.Value":0.0153,"adj.P.Val":0.0563},{"tissuedysplasia":0.1768,"tissuetumour":0.4642,"AveExpr":5.0282,"F":5.5439,"P.Value":0.0071,"adj.P.Val":0.0326},{"tissuedysplasia":0.2539,"tissuetumour":-0.0608,"AveExpr":6.4803,"F":2.2502,"P.Value":0.1174,"adj.P.Val":0.2351},{"tissuedysplasia":0.0019,"tissuetumour":0.6174,"AveExpr":2.1004,"F":4.725,"P.Value":0.0138,"adj.P.Val":0.0525},{"tissuedysplasia":0.2062,"tissuetumour":-0.0265,"AveExpr":4.2287,"F":1.8763,"P.Value":0.1652,"adj.P.Val":0.2964},{"tissuedysplasia":0.2555,"tissuetumour":0.8594,"AveExpr":2.2365,"F":7.6978,"P.Value":0.0014,"adj.P.Val":0.0097},{"tissuedysplasia":-0.4279,"tissuetumour":-0.7769,"AveExpr":5.5751,"F":12.7247,"P.Value":0,"adj.P.Val":0.0007},{"tissuedysplasia":0.3427,"tissuetumour":0.126,"AveExpr":4.6376,"F":3.9208,"P.Value":0.0271,"adj.P.Val":0.0849},{"tissuedysplasia":1.703,"tissuetumour":1.8113,"AveExpr":-5.3285,"F":1.2644,"P.Value":0.2925,"adj.P.Val":0.4371},{"tissuedysplasia":0.0023,"tissuetumour":-0.0136,"AveExpr":5.7522,"F":0.0144,"P.Value":0.9857,"adj.P.Val":0.99},{"tissuedysplasia":0.0736,"tissuetumour":1.8515,"AveExpr":-3.5484,"F":4.8366,"P.Value":0.0126,"adj.P.Val":0.049},{"tissuedysplasia":0.1289,"tissuetumour":-0.0525,"AveExpr":4.7216,"F":0.6733,"P.Value":0.5152,"adj.P.Val":0.6395},{"tissuedysplasia":0.6002,"tissuetumour":0.4793,"AveExpr":3.0503,"F":5.133,"P.Value":0.0099,"adj.P.Val":0.0412},{"tissuedysplasia":0.2514,"tissuetumour":1.9343,"AveExpr":-1.6572,"F":6.1631,"P.Value":0.0044,"adj.P.Val":0.0228},{"tissuedysplasia":-0.0985,"tissuetumour":-0.5897,"AveExpr":5.4668,"F":11.9705,"P.Value":0.0001,"adj.P.Val":0.0011},{"tissuedysplasia":1.2392,"tissuetumour":3.7786,"AveExpr":-6.788,"F":9.4313,"P.Value":0.0004,"adj.P.Val":0.0039},{"tissuedysplasia":0.1332,"tissuetumour":0.0267,"AveExpr":3.4123,"F":0.6296,"P.Value":0.5375,"adj.P.Val":0.6581},{"tissuedysplasia":-0.0339,"tissuetumour":0.1015,"AveExpr":4.6947,"F":0.4712,"P.Value":0.6274,"adj.P.Val":0.7305},{"tissuedysplasia":-0.061,"tissuetumour":0.2457,"AveExpr":4.7088,"F":2.7208,"P.Value":0.0769,"adj.P.Val":0.1758},{"tissuedysplasia":0.0881,"tissuetumour":0.1804,"AveExpr":5.3926,"F":1.7587,"P.Value":0.1842,"adj.P.Val":0.3194},{"tissuedysplasia":0.1639,"tissuetumour":0.8181,"AveExpr":1.7365,"F":6.448,"P.Value":0.0035,"adj.P.Val":0.0194},{"tissuedysplasia":0.1799,"tissuetumour":0.3873,"AveExpr":4.8019,"F":7.5442,"P.Value":0.0015,"adj.P.Val":0.0106},{"tissuedysplasia":-1.7971,"tissuetumour":-1.5964,"AveExpr":2.9438,"F":24.9157,"P.Value":5.8081e-08,"adj.P.Val":4.8087e-06},{"tissuedysplasia":-0.1515,"tissuetumour":-0.0056,"AveExpr":3.9939,"F":0.7976,"P.Value":0.4568,"adj.P.Val":0.5905},{"tissuedysplasia":0.1921,"tissuetumour":0.3068,"AveExpr":2.0582,"F":1.3052,"P.Value":0.2814,"adj.P.Val":0.426},{"tissuedysplasia":-0.5212,"tissuetumour":1.6422,"AveExpr":-0.6096,"F":10.8188,"P.Value":0.0002,"adj.P.Val":0.0019},{"tissuedysplasia":-0.0155,"tissuetumour":-0.365,"AveExpr":6.0056,"F":3.9449,"P.Value":0.0266,"adj.P.Val":0.0837},{"tissuedysplasia":0.3732,"tissuetumour":0.5896,"AveExpr":0.3126,"F":1.5958,"P.Value":0.2143,"adj.P.Val":0.3535},{"tissuedysplasia":0.4335,"tissuetumour":1.6394,"AveExpr":-1.5419,"F":3.4948,"P.Value":0.039,"adj.P.Val":0.1096},{"tissuedysplasia":1.1042,"tissuetumour":0.9314,"AveExpr":-3.6645,"F":0.8836,"P.Value":0.4205,"adj.P.Val":0.5585},{"tissuedysplasia":0.1913,"tissuetumour":0.6728,"AveExpr":4.4509,"F":11.7646,"P.Value":0.0001,"adj.P.Val":0.0012},{"tissuedysplasia":2.076,"tissuetumour":2.8434,"AveExpr":0.129,"F":11.578,"P.Value":0.0001,"adj.P.Val":0.0013},{"tissuedysplasia":-0.0937,"tissuetumour":0.6728,"AveExpr":2.7116,"F":12.0577,"P.Value":0.0001,"adj.P.Val":0.001},{"tissuedysplasia":1.5751,"tissuetumour":2.3041,"AveExpr":-0.7814,"F":9.0919,"P.Value":0.0005,"adj.P.Val":0.0046},{"tissuedysplasia":0.0295,"tissuetumour":-0.3219,"AveExpr":6.7846,"F":7.6353,"P.Value":0.0014,"adj.P.Val":0.0101},{"tissuedysplasia":0.8783,"tissuetumour":2.3907,"AveExpr":-1.2967,"F":4.5969,"P.Value":0.0154,"adj.P.Val":0.0566},{"tissuedysplasia":0.8343,"tissuetumour":1.995,"AveExpr":1.2715,"F":12.3356,"P.Value":0.0001,"adj.P.Val":0.0009},{"tissuedysplasia":0.0676,"tissuetumour":0.1411,"AveExpr":6.9958,"F":1.288,"P.Value":0.286,"adj.P.Val":0.4305},{"tissuedysplasia":0.8206,"tissuetumour":1.7685,"AveExpr":-3.6553,"F":2.6986,"P.Value":0.0784,"adj.P.Val":0.1782},{"tissuedysplasia":-0.6413,"tissuetumour":-0.9456,"AveExpr":5.6592,"F":23.0864,"P.Value":1.3933e-07,"adj.P.Val":9.0104e-06},{"tissuedysplasia":-0.2358,"tissuetumour":0.2745,"AveExpr":-1.1176,"F":0.3926,"P.Value":0.6777,"adj.P.Val":0.7687},{"tissuedysplasia":0.8104,"tissuetumour":1.5145,"AveExpr":-0.3853,"F":3.4841,"P.Value":0.0394,"adj.P.Val":0.1104},{"tissuedysplasia":-1.1094,"tissuetumour":-3.1367,"AveExpr":3.1254,"F":15.6353,"P.Value":9.1782e-06,"adj.P.Val":0.0002},{"tissuedysplasia":1.0544,"tissuetumour":1.0355,"AveExpr":-0.6444,"F":1.9465,"P.Value":0.1549,"adj.P.Val":0.2837},{"tissuedysplasia":-0.4464,"tissuetumour":2.0605,"AveExpr":-0.5314,"F":9.5605,"P.Value":0.0004,"adj.P.Val":0.0036},{"tissuedysplasia":0.1457,"tissuetumour":0.0575,"AveExpr":4.4345,"F":0.5853,"P.Value":0.5612,"adj.P.Val":0.6771},{"tissuedysplasia":1.3492,"tissuetumour":3.0712,"AveExpr":-4.6011,"F":5.5073,"P.Value":0.0073,"adj.P.Val":0.0333},{"tissuedysplasia":0.2624,"tissuetumour":0.9476,"AveExpr":1.9627,"F":5.422,"P.Value":0.0079,"adj.P.Val":0.0349},{"tissuedysplasia":0.7127,"tissuetumour":1.1797,"AveExpr":1.1817,"F":9.1972,"P.Value":0.0005,"adj.P.Val":0.0044},{"tissuedysplasia":0.0703,"tissuetumour":0.5468,"AveExpr":3.7994,"F":4.9769,"P.Value":0.0113,"adj.P.Val":0.045},{"tissuedysplasia":0.043,"tissuetumour":0.089,"AveExpr":5.6191,"F":0.3089,"P.Value":0.7358,"adj.P.Val":0.8119},{"tissuedysplasia":0.0368,"tissuetumour":0.0275,"AveExpr":1.0686,"F":0.007,"P.Value":0.993,"adj.P.Val":0.9951},{"tissuedysplasia":0.5695,"tissuetumour":0.7984,"AveExpr":1.5865,"F":8.1655,"P.Value":0.001,"adj.P.Val":0.0075},{"tissuedysplasia":0.4334,"tissuetumour":1.1189,"AveExpr":3.2027,"F":17.2694,"P.Value":2.9103e-06,"adj.P.Val":0.0001},{"tissuedysplasia":0.2591,"tissuetumour":1.0288,"AveExpr":2.476,"F":9.9221,"P.Value":0.0003,"adj.P.Val":0.003},{"tissuedysplasia":0.2721,"tissuetumour":0.5823,"AveExpr":4.9301,"F":12.2617,"P.Value":0.0001,"adj.P.Val":0.0009},{"tissuedysplasia":-0.1439,"tissuetumour":0.3034,"AveExpr":4.745,"F":5.2715,"P.Value":0.0089,"adj.P.Val":0.0381},{"tissuedysplasia":0.4361,"tissuetumour":0.1808,"AveExpr":2.8729,"F":0.8284,"P.Value":0.4434,"adj.P.Val":0.5793},{"tissuedysplasia":0.1986,"tissuetumour":0.5223,"AveExpr":4.6722,"F":12.9899,"P.Value":0,"adj.P.Val":0.0006},{"tissuedysplasia":0.3407,"tissuetumour":-0.7395,"AveExpr":2.1198,"F":6.4692,"P.Value":0.0034,"adj.P.Val":0.0192},{"tissuedysplasia":0.1043,"tissuetumour":-0.2879,"AveExpr":4.0116,"F":2.3511,"P.Value":0.1071,"adj.P.Val":0.2209},{"tissuedysplasia":0.518,"tissuetumour":1.5838,"AveExpr":-3.6581,"F":1.8704,"P.Value":0.1661,"adj.P.Val":0.2975},{"tissuedysplasia":0.0806,"tissuetumour":0.2646,"AveExpr":3.2127,"F":1.327,"P.Value":0.2757,"adj.P.Val":0.4201},{"tissuedysplasia":0.5008,"tissuetumour":0.5204,"AveExpr":4.8595,"F":8.5385,"P.Value":0.0007,"adj.P.Val":0.0062},{"tissuedysplasia":0.3394,"tissuetumour":0.8886,"AveExpr":-1.1896,"F":1.075,"P.Value":0.3501,"adj.P.Val":0.494},{"tissuedysplasia":0.1424,"tissuetumour":0.1812,"AveExpr":6.3228,"F":0.9631,"P.Value":0.3896,"adj.P.Val":0.5306},{"tissuedysplasia":-0.3786,"tissuetumour":-0.5524,"AveExpr":4.7965,"F":9.1153,"P.Value":0.0005,"adj.P.Val":0.0046},{"tissuedysplasia":0.0224,"tissuetumour":-0.2067,"AveExpr":1.084,"F":0.19,"P.Value":0.8276,"adj.P.Val":0.8791},{"tissuedysplasia":0.2985,"tissuetumour":0.9189,"AveExpr":4.3022,"F":27.8893,"P.Value":1.5025e-08,"adj.P.Val":1.8913e-06},{"tissuedysplasia":0.2723,"tissuetumour":1.1513,"AveExpr":4.7802,"F":34.8111,"P.Value":8.6173e-10,"adj.P.Val":2.263e-07},{"tissuedysplasia":-0.3397,"tissuetumour":0.0141,"AveExpr":0.1944,"F":0.2842,"P.Value":0.754,"adj.P.Val":0.8253},{"tissuedysplasia":0.2406,"tissuetumour":2.3538,"AveExpr":-3.6684,"F":5.3498,"P.Value":0.0083,"adj.P.Val":0.0363},{"tissuedysplasia":-0.4108,"tissuetumour":-0.5104,"AveExpr":3.2565,"F":8.3146,"P.Value":0.0009,"adj.P.Val":0.007},{"tissuedysplasia":0.0376,"tissuetumour":0.5479,"AveExpr":4.2581,"F":9.9176,"P.Value":0.0003,"adj.P.Val":0.003},{"tissuedysplasia":0.0772,"tissuetumour":0.2688,"AveExpr":2.1366,"F":0.4439,"P.Value":0.6444,"adj.P.Val":0.7432},{"tissuedysplasia":0.0679,"tissuetumour":-0.5623,"AveExpr":5.3413,"F":7.4879,"P.Value":0.0016,"adj.P.Val":0.0109},{"tissuedysplasia":-0.0565,"tissuetumour":0.3157,"AveExpr":4.5372,"F":5.2661,"P.Value":0.0089,"adj.P.Val":0.0382},{"tissuedysplasia":0.1035,"tissuetumour":-0.0542,"AveExpr":6.9944,"F":1.0485,"P.Value":0.3591,"adj.P.Val":0.5025},{"tissuedysplasia":0.9892,"tissuetumour":1.5698,"AveExpr":2.3203,"F":11.1375,"P.Value":0.0001,"adj.P.Val":0.0016},{"tissuedysplasia":0.0658,"tissuetumour":-0.7507,"AveExpr":3.992,"F":16.0064,"P.Value":5.9745e-06,"adj.P.Val":0.0002},{"tissuedysplasia":-0.6561,"tissuetumour":-0.8811,"AveExpr":3.4673,"F":16.8229,"P.Value":3.7428e-06,"adj.P.Val":0.0001},{"tissuedysplasia":0.3623,"tissuetumour":0.0867,"AveExpr":1.1402,"F":0.7149,"P.Value":0.4949,"adj.P.Val":0.6224},{"tissuedysplasia":-0.0352,"tissuetumour":0.7096,"AveExpr":2.2628,"F":6.733,"P.Value":0.0028,"adj.P.Val":0.0166},{"tissuedysplasia":-0.4568,"tissuetumour":-0.3605,"AveExpr":2.6468,"F":0.9808,"P.Value":0.3831,"adj.P.Val":0.5248},{"tissuedysplasia":-0.0929,"tissuetumour":-0.4257,"AveExpr":4.1478,"F":2.5326,"P.Value":0.091,"adj.P.Val":0.1976},{"tissuedysplasia":-0.9348,"tissuetumour":0.33,"AveExpr":-1.8504,"F":2.5029,"P.Value":0.0934,"adj.P.Val":0.2013},{"tissuedysplasia":-0.6901,"tissuetumour":-0.5759,"AveExpr":6.1964,"F":8.7319,"P.Value":0.0006,"adj.P.Val":0.0056},{"tissuedysplasia":-0.2044,"tissuetumour":0.0157,"AveExpr":5.5891,"F":1.2248,"P.Value":0.3036,"adj.P.Val":0.448},{"tissuedysplasia":1.0042,"tissuetumour":1.7695,"AveExpr":1.8632,"F":18.3236,"P.Value":1.6248e-06,"adj.P.Val":0.0001},{"tissuedysplasia":1.4133,"tissuetumour":2.4116,"AveExpr":4.0351,"F":27.4491,"P.Value":1.8261e-08,"adj.P.Val":2.1541e-06},{"tissuedysplasia":0.0577,"tissuetumour":0.6584,"AveExpr":5.2851,"F":8.7156,"P.Value":0.0006,"adj.P.Val":0.0057},{"tissuedysplasia":0.5609,"tissuetumour":1.0133,"AveExpr":0.4955,"F":2.5798,"P.Value":0.0872,"adj.P.Val":0.1918},{"tissuedysplasia":0.2678,"tissuetumour":0.7207,"AveExpr":4.6514,"F":19.6594,"P.Value":7.9325e-07,"adj.P.Val":0},{"tissuedysplasia":0.5912,"tissuetumour":1.4598,"AveExpr":3.0058,"F":22.6445,"P.Value":1.7305e-07,"adj.P.Val":0},{"tissuedysplasia":0.0593,"tissuetumour":-0.3459,"AveExpr":5.5102,"F":8.6741,"P.Value":0.0007,"adj.P.Val":0.0058},{"tissuedysplasia":0.3735,"tissuetumour":0.3964,"AveExpr":5.5345,"F":6.7737,"P.Value":0.0027,"adj.P.Val":0.0162},{"tissuedysplasia":0.182,"tissuetumour":0.0445,"AveExpr":7.0223,"F":1.2092,"P.Value":0.3082,"adj.P.Val":0.4527},{"tissuedysplasia":0.0246,"tissuetumour":-0.3636,"AveExpr":7.8137,"F":6.1655,"P.Value":0.0044,"adj.P.Val":0.0227},{"tissuedysplasia":0.2868,"tissuetumour":0.2899,"AveExpr":6.6783,"F":5.9025,"P.Value":0.0054,"adj.P.Val":0.0264},{"tissuedysplasia":-0.049,"tissuetumour":-0.56,"AveExpr":6.3596,"F":10.3142,"P.Value":0.0002,"adj.P.Val":0.0024},{"tissuedysplasia":0.0194,"tissuetumour":-0.2554,"AveExpr":2.9978,"F":0.5478,"P.Value":0.5821,"adj.P.Val":0.6941},{"tissuedysplasia":0.3848,"tissuetumour":1.4496,"AveExpr":-1.9915,"F":2.3191,"P.Value":0.1103,"adj.P.Val":0.2254},{"tissuedysplasia":0.1541,"tissuetumour":-0.3459,"AveExpr":1.7111,"F":1.9966,"P.Value":0.1479,"adj.P.Val":0.275},{"tissuedysplasia":0.2494,"tissuetumour":1.4017,"AveExpr":-0.0453,"F":6.4998,"P.Value":0.0034,"adj.P.Val":0.0189},{"tissuedysplasia":0.0445,"tissuetumour":0.1127,"AveExpr":3.6044,"F":0.2746,"P.Value":0.7612,"adj.P.Val":0.8306},{"tissuedysplasia":0.0196,"tissuetumour":-0.9196,"AveExpr":5.4394,"F":5.9193,"P.Value":0.0056,"adj.P.Val":0.0274},{"tissuedysplasia":0.1696,"tissuetumour":0.7679,"AveExpr":3.5912,"F":14.269,"P.Value":0,"adj.P.Val":0.0003},{"tissuedysplasia":0.0798,"tissuetumour":-0.4646,"AveExpr":4.5853,"F":8.2434,"P.Value":0.0009,"adj.P.Val":0.0072},{"tissuedysplasia":0.0457,"tissuetumour":0.8416,"AveExpr":-0.3148,"F":2.1261,"P.Value":0.1314,"adj.P.Val":0.2539},{"tissuedysplasia":0.283,"tissuetumour":0.5395,"AveExpr":6.7852,"F":18.3854,"P.Value":1.571e-06,"adj.P.Val":0.0001},{"tissuedysplasia":-0.5845,"tissuetumour":-1.5581,"AveExpr":6.1062,"F":35.5991,"P.Value":6.3643e-10,"adj.P.Val":1.8749e-07},{"tissuedysplasia":0.9086,"tissuetumour":1.5464,"AveExpr":1.3723,"F":11.4058,"P.Value":0.0001,"adj.P.Val":0.0014},{"tissuedysplasia":0.3156,"tissuetumour":0.237,"AveExpr":3.8874,"F":4.0056,"P.Value":0.0252,"adj.P.Val":0.0808},{"tissuedysplasia":0.4138,"tissuetumour":0.8724,"AveExpr":3.397,"F":19.2357,"P.Value":9.9335e-07,"adj.P.Val":0},{"tissuedysplasia":0.4231,"tissuetumour":0.6016,"AveExpr":2.1178,"F":3.5298,"P.Value":0.0379,"adj.P.Val":0.1074},{"tissuedysplasia":0.003,"tissuetumour":-0.3143,"AveExpr":7.5204,"F":6.6401,"P.Value":0.003,"adj.P.Val":0.0175},{"tissuedysplasia":-0.026,"tissuetumour":0.3579,"AveExpr":4.6422,"F":4.2939,"P.Value":0.0198,"adj.P.Val":0.0681},{"tissuedysplasia":0.4698,"tissuetumour":0.6106,"AveExpr":2.8849,"F":4.904,"P.Value":0.0119,"adj.P.Val":0.0469},{"tissuedysplasia":-0.2152,"tissuetumour":-0.6939,"AveExpr":6.9016,"F":14.471,"P.Value":0,"adj.P.Val":0.0003},{"tissuedysplasia":-0.7288,"tissuetumour":2.6548,"AveExpr":-3.3684,"F":6.7523,"P.Value":0.0028,"adj.P.Val":0.0164},{"tissuedysplasia":0.4904,"tissuetumour":0.7976,"AveExpr":5.4855,"F":15.8535,"P.Value":6.5287e-06,"adj.P.Val":0.0002},{"tissuedysplasia":0.5953,"tissuetumour":1.5033,"AveExpr":3.2512,"F":17.521,"P.Value":2.5289e-06,"adj.P.Val":0.0001},{"tissuedysplasia":0.2811,"tissuetumour":0.8095,"AveExpr":1.8746,"F":4.5105,"P.Value":0.0165,"adj.P.Val":0.0597},{"tissuedysplasia":-0.1341,"tissuetumour":0.1562,"AveExpr":1.8546,"F":0.7615,"P.Value":0.473,"adj.P.Val":0.6034},{"tissuedysplasia":-0.0804,"tissuetumour":0.2802,"AveExpr":2.3745,"F":0.9619,"P.Value":0.3901,"adj.P.Val":0.531},{"tissuedysplasia":-0.0951,"tissuetumour":0.299,"AveExpr":5.7227,"F":4.4629,"P.Value":0.0172,"adj.P.Val":0.0615},{"tissuedysplasia":0.3495,"tissuetumour":1.0614,"AveExpr":5.5785,"F":12.9364,"P.Value":0,"adj.P.Val":0.0007},{"tissuedysplasia":-0.0809,"tissuetumour":0.235,"AveExpr":2.0946,"F":1.3028,"P.Value":0.282,"adj.P.Val":0.4267},{"tissuedysplasia":0.2982,"tissuetumour":1.4751,"AveExpr":-1.448,"F":3.5008,"P.Value":0.0388,"adj.P.Val":0.1093},{"tissuedysplasia":0.0821,"tissuetumour":0.0082,"AveExpr":6.2481,"F":0.3001,"P.Value":0.7422,"adj.P.Val":0.8165},{"tissuedysplasia":-0.858,"tissuetumour":-1.1778,"AveExpr":2.5759,"F":8.5792,"P.Value":0.0007,"adj.P.Val":0.0061},{"tissuedysplasia":0.722,"tissuetumour":0.9811,"AveExpr":3.153,"F":19.4791,"P.Value":8.727e-07,"adj.P.Val":0},{"tissuedysplasia":0.7361,"tissuetumour":1.0602,"AveExpr":3.929,"F":11.6515,"P.Value":0.0001,"adj.P.Val":0.0012},{"tissuedysplasia":0.5582,"tissuetumour":1.1118,"AveExpr":1.3572,"F":6.3186,"P.Value":0.0039,"adj.P.Val":0.0209},{"tissuedysplasia":0.8874,"tissuetumour":0.9305,"AveExpr":2.5037,"F":10.5829,"P.Value":0.0002,"adj.P.Val":0.0021},{"tissuedysplasia":0.3806,"tissuetumour":0.6234,"AveExpr":2.5555,"F":5.1417,"P.Value":0.0098,"adj.P.Val":0.041},{"tissuedysplasia":0.1963,"tissuetumour":0.4448,"AveExpr":3.81,"F":5.3877,"P.Value":0.0081,"adj.P.Val":0.0355},{"tissuedysplasia":0.6417,"tissuetumour":0.4811,"AveExpr":5.3597,"F":4.996,"P.Value":0.0111,"adj.P.Val":0.0446},{"tissuedysplasia":0.1787,"tissuetumour":-0.038,"AveExpr":6.2889,"F":2.9346,"P.Value":0.0636,"adj.P.Val":0.1545},{"tissuedysplasia":-0.1482,"tissuetumour":-0.4803,"AveExpr":1.6068,"F":1.0447,"P.Value":0.3604,"adj.P.Val":0.5036},{"tissuedysplasia":0.0151,"tissuetumour":0.1605,"AveExpr":5.9012,"F":2.6977,"P.Value":0.0785,"adj.P.Val":0.1783},{"tissuedysplasia":0.1993,"tissuetumour":0.845,"AveExpr":3.0076,"F":12.1902,"P.Value":0.0001,"adj.P.Val":0.001},{"tissuedysplasia":0.1826,"tissuetumour":0.4542,"AveExpr":4.0733,"F":7.0777,"P.Value":0.0022,"adj.P.Val":0.0137},{"tissuedysplasia":-0.4654,"tissuetumour":0.5096,"AveExpr":4.9726,"F":10.7202,"P.Value":0.0002,"adj.P.Val":0.002},{"tissuedysplasia":0.0883,"tissuetumour":0.4097,"AveExpr":5.6106,"F":7.6439,"P.Value":0.0014,"adj.P.Val":0.01},{"tissuedysplasia":1.4738,"tissuetumour":3.5099,"AveExpr":-1.8895,"F":18.499,"P.Value":1.4769e-06,"adj.P.Val":0.0001},{"tissuedysplasia":0.4507,"tissuetumour":1.3964,"AveExpr":4.1621,"F":45.1805,"P.Value":2.1549e-11,"adj.P.Val":1.6959e-08},{"tissuedysplasia":-0.0596,"tissuetumour":-0.1996,"AveExpr":5.8033,"F":2.4095,"P.Value":0.1016,"adj.P.Val":0.2133},{"tissuedysplasia":0.4791,"tissuetumour":1.1391,"AveExpr":2.2191,"F":12.1694,"P.Value":0.0001,"adj.P.Val":0.001},{"tissuedysplasia":0.1679,"tissuetumour":0.3192,"AveExpr":2.3068,"F":1.6843,"P.Value":0.1973,"adj.P.Val":0.3342},{"tissuedysplasia":-0.3301,"tissuetumour":-0.8932,"AveExpr":4.0295,"F":15.0692,"P.Value":0,"adj.P.Val":0.0002},{"tissuedysplasia":0.2083,"tissuetumour":-0.0023,"AveExpr":2.4571,"F":0.3623,"P.Value":0.6982,"adj.P.Val":0.7835},{"tissuedysplasia":0.076,"tissuetumour":0.632,"AveExpr":4.7795,"F":11.4915,"P.Value":0.0001,"adj.P.Val":0.0014},{"tissuedysplasia":0.0392,"tissuetumour":-0.4081,"AveExpr":4.7845,"F":8.9953,"P.Value":0.0005,"adj.P.Val":0.0049},{"tissuedysplasia":0.2307,"tissuetumour":0.1121,"AveExpr":3.7279,"F":2.3314,"P.Value":0.109,"adj.P.Val":0.2237},{"tissuedysplasia":0.3447,"tissuetumour":0.5303,"AveExpr":5.3515,"F":11.4381,"P.Value":0.0001,"adj.P.Val":0.0014},{"tissuedysplasia":-0.0799,"tissuetumour":-0.464,"AveExpr":5.5334,"F":9.1582,"P.Value":0.0005,"adj.P.Val":0.0045},{"tissuedysplasia":0.0984,"tissuetumour":0.2482,"AveExpr":6.2497,"F":2.7651,"P.Value":0.0739,"adj.P.Val":0.1712},{"tissuedysplasia":0.9946,"tissuetumour":1.9316,"AveExpr":0.6013,"F":6.3335,"P.Value":0.0038,"adj.P.Val":0.0207},{"tissuedysplasia":0.4979,"tissuetumour":1.754,"AveExpr":2.4131,"F":13.982,"P.Value":0,"adj.P.Val":0.0004},{"tissuedysplasia":0.1065,"tissuetumour":0.1001,"AveExpr":3.5961,"F":0.4127,"P.Value":0.6644,"adj.P.Val":0.7587},{"tissuedysplasia":0.0869,"tissuetumour":-0.3321,"AveExpr":6.3021,"F":4.9913,"P.Value":0.0111,"adj.P.Val":0.0447},{"tissuedysplasia":0.3779,"tissuetumour":0.4582,"AveExpr":4.2017,"F":0.9059,"P.Value":0.412,"adj.P.Val":0.551},{"tissuedysplasia":0.6045,"tissuetumour":0.655,"AveExpr":5.2505,"F":6.1999,"P.Value":0.0042,"adj.P.Val":0.0223},{"tissuedysplasia":0.224,"tissuetumour":0.6618,"AveExpr":4.6971,"F":10.1142,"P.Value":0.0002,"adj.P.Val":0.0027},{"tissuedysplasia":-0.0321,"tissuetumour":-0.0774,"AveExpr":6.1901,"F":0.2516,"P.Value":0.7786,"adj.P.Val":0.8435},{"tissuedysplasia":0.3703,"tissuetumour":0.1517,"AveExpr":6.8463,"F":2.686,"P.Value":0.0799,"adj.P.Val":0.1806},{"tissuedysplasia":0.5866,"tissuetumour":1.2374,"AveExpr":1.5367,"F":11.2404,"P.Value":0.0001,"adj.P.Val":0.0015},{"tissuedysplasia":0.1257,"tissuetumour":0.4679,"AveExpr":4.586,"F":9.7305,"P.Value":0.0003,"adj.P.Val":0.0033},{"tissuedysplasia":0.0907,"tissuetumour":-0.3199,"AveExpr":5.1667,"F":7.7275,"P.Value":0.0013,"adj.P.Val":0.0096},{"tissuedysplasia":0.0098,"tissuetumour":-0.5743,"AveExpr":4.5021,"F":9.2753,"P.Value":0.0004,"adj.P.Val":0.0042},{"tissuedysplasia":0.4411,"tissuetumour":1.0443,"AveExpr":0.8613,"F":4.6276,"P.Value":0.015,"adj.P.Val":0.0556},{"tissuedysplasia":0.7282,"tissuetumour":1.3928,"AveExpr":2.6404,"F":9.8355,"P.Value":0.0003,"adj.P.Val":0.0031},{"tissuedysplasia":0.1078,"tissuetumour":0.1513,"AveExpr":4.1264,"F":0.2955,"P.Value":0.7456,"adj.P.Val":0.8191},{"tissuedysplasia":0.0241,"tissuetumour":0.4597,"AveExpr":0.4303,"F":1.0068,"P.Value":0.3737,"adj.P.Val":0.5162},{"tissuedysplasia":0.4695,"tissuetumour":0.498,"AveExpr":4.3357,"F":5.6474,"P.Value":0.0066,"adj.P.Val":0.0306},{"tissuedysplasia":0.0445,"tissuetumour":0.5603,"AveExpr":4.8512,"F":7.1472,"P.Value":0.0021,"adj.P.Val":0.0132},{"tissuedysplasia":1.651,"tissuetumour":1.7914,"AveExpr":1.0879,"F":7.9892,"P.Value":0.0011,"adj.P.Val":0.0083},{"tissuedysplasia":0.3908,"tissuetumour":1.3561,"AveExpr":1.0187,"F":8.1472,"P.Value":0.001,"adj.P.Val":0.0076},{"tissuedysplasia":0.1193,"tissuetumour":-0.0395,"AveExpr":3.6283,"F":0.7894,"P.Value":0.4604,"adj.P.Val":0.5937},{"tissuedysplasia":0.1069,"tissuetumour":0.3627,"AveExpr":2.9396,"F":2.3203,"P.Value":0.1101,"adj.P.Val":0.2252},{"tissuedysplasia":1.9574,"tissuetumour":1.4656,"AveExpr":1.7962,"F":2.7889,"P.Value":0.0745,"adj.P.Val":0.172},{"tissuedysplasia":0.5391,"tissuetumour":0.8959,"AveExpr":4.8853,"F":18.4439,"P.Value":1.5217e-06,"adj.P.Val":0.0001},{"tissuedysplasia":-0.8699,"tissuetumour":-0.0837,"AveExpr":-2.3351,"F":0.6917,"P.Value":0.5061,"adj.P.Val":0.6319},{"tissuedysplasia":0.5692,"tissuetumour":0.4984,"AveExpr":4.9634,"F":5.4704,"P.Value":0.0076,"adj.P.Val":0.0339},{"tissuedysplasia":0.4092,"tissuetumour":0.6332,"AveExpr":2.9679,"F":3.8997,"P.Value":0.0276,"adj.P.Val":0.086},{"tissuedysplasia":0.0915,"tissuetumour":-0.1298,"AveExpr":1.8535,"F":0.7963,"P.Value":0.4574,"adj.P.Val":0.591},{"tissuedysplasia":-0.0854,"tissuetumour":0.1001,"AveExpr":6.0515,"F":1.0752,"P.Value":0.35,"adj.P.Val":0.4939},{"tissuedysplasia":0.3432,"tissuetumour":0.1964,"AveExpr":3.7982,"F":2.9858,"P.Value":0.0608,"adj.P.Val":0.1497},{"tissuedysplasia":-0.0832,"tissuetumour":-0.4952,"AveExpr":5.9801,"F":7.5563,"P.Value":0.0015,"adj.P.Val":0.0105},{"tissuedysplasia":-0.8394,"tissuetumour":-1.1071,"AveExpr":4.5776,"F":22.3706,"P.Value":1.9814e-07,"adj.P.Val":0},{"tissuedysplasia":0.3472,"tissuetumour":0.5414,"AveExpr":5.2707,"F":6.5244,"P.Value":0.0033,"adj.P.Val":0.0186},{"tissuedysplasia":0.1358,"tissuetumour":0.5462,"AveExpr":4.7069,"F":9.181,"P.Value":0.0005,"adj.P.Val":0.0044},{"tissuedysplasia":0.3908,"tissuetumour":2.0162,"AveExpr":-0.0178,"F":11.9117,"P.Value":0.0001,"adj.P.Val":0.0011},{"tissuedysplasia":0.6105,"tissuetumour":1.1338,"AveExpr":1.2232,"F":6.2831,"P.Value":0.004,"adj.P.Val":0.0213},{"tissuedysplasia":-0.0072,"tissuetumour":0.3529,"AveExpr":4.4225,"F":4.7162,"P.Value":0.0139,"adj.P.Val":0.0527},{"tissuedysplasia":0.474,"tissuetumour":1.4145,"AveExpr":5.4509,"F":35.966,"P.Value":5.5344e-10,"adj.P.Val":1.7085e-07},{"tissuedysplasia":0.4746,"tissuetumour":0.7436,"AveExpr":5.0226,"F":16.0404,"P.Value":5.8582e-06,"adj.P.Val":0.0002},{"tissuedysplasia":0.2074,"tissuetumour":0.0289,"AveExpr":4.7262,"F":1.4176,"P.Value":0.2532,"adj.P.Val":0.3961},{"tissuedysplasia":0.734,"tissuetumour":0.6993,"AveExpr":1.0533,"F":3.0258,"P.Value":0.0587,"adj.P.Val":0.1461},{"tissuedysplasia":-0.0441,"tissuetumour":-0.4298,"AveExpr":6.4127,"F":7.9695,"P.Value":0.0011,"adj.P.Val":0.0084},{"tissuedysplasia":-0.0269,"tissuetumour":-0.3748,"AveExpr":3.897,"F":2.9446,"P.Value":0.0631,"adj.P.Val":0.1535},{"tissuedysplasia":0.4748,"tissuetumour":0.8959,"AveExpr":3.558,"F":16.6665,"P.Value":4.0905e-06,"adj.P.Val":0.0001},{"tissuedysplasia":0.1473,"tissuetumour":0.059,"AveExpr":4.2355,"F":0.8452,"P.Value":0.4363,"adj.P.Val":0.5731},{"tissuedysplasia":0.1885,"tissuetumour":0.7983,"AveExpr":3.411,"F":9.8242,"P.Value":0.0003,"adj.P.Val":0.0032},{"tissuedysplasia":0.4515,"tissuetumour":1.1271,"AveExpr":4.552,"F":30.5141,"P.Value":4.8627e-09,"adj.P.Val":8.3005e-07},{"tissuedysplasia":0.1357,"tissuetumour":0.1307,"AveExpr":3.2569,"F":0.6495,"P.Value":0.5273,"adj.P.Val":0.6501},{"tissuedysplasia":0.0837,"tissuetumour":0.0951,"AveExpr":5.5625,"F":0.0527,"P.Value":0.9487,"adj.P.Val":0.9646},{"tissuedysplasia":0.4295,"tissuetumour":0.904,"AveExpr":1.625,"F":7.13,"P.Value":0.0021,"adj.P.Val":0.0133},{"tissuedysplasia":0.9453,"tissuetumour":1.0012,"AveExpr":2.5228,"F":5.9083,"P.Value":0.0053,"adj.P.Val":0.0264},{"tissuedysplasia":0.5455,"tissuetumour":1.4236,"AveExpr":4.1472,"F":35.6063,"P.Value":6.3468e-10,"adj.P.Val":1.8749e-07},{"tissuedysplasia":-0.0222,"tissuetumour":-0.6204,"AveExpr":4.7022,"F":6.8414,"P.Value":0.0026,"adj.P.Val":0.0156},{"tissuedysplasia":0.8409,"tissuetumour":1.1586,"AveExpr":3.3918,"F":25.0862,"P.Value":5.3626e-08,"adj.P.Val":4.5435e-06},{"tissuedysplasia":0.1781,"tissuetumour":0.8299,"AveExpr":3.1123,"F":14.9478,"P.Value":0,"adj.P.Val":0.0003},{"tissuedysplasia":0.1187,"tissuetumour":0.2335,"AveExpr":6.6591,"F":1.6124,"P.Value":0.211,"adj.P.Val":0.3498},{"tissuedysplasia":-0.9243,"tissuetumour":-2.2324,"AveExpr":4.134,"F":29.4436,"P.Value":7.6503e-09,"adj.P.Val":1.1562e-06},{"tissuedysplasia":0.3045,"tissuetumour":0.1541,"AveExpr":4.0657,"F":3.7159,"P.Value":0.0323,"adj.P.Val":0.0959},{"tissuedysplasia":-0.0802,"tissuetumour":-0.4914,"AveExpr":5.4608,"F":6.7433,"P.Value":0.0028,"adj.P.Val":0.0165},{"tissuedysplasia":0.1473,"tissuetumour":0.7667,"AveExpr":3.0279,"F":6.5332,"P.Value":0.0033,"adj.P.Val":0.0185},{"tissuedysplasia":0.1797,"tissuetumour":0.0667,"AveExpr":4.5078,"F":1.4161,"P.Value":0.2535,"adj.P.Val":0.3964},{"tissuedysplasia":0.5805,"tissuetumour":0.6333,"AveExpr":5.9547,"F":10.515,"P.Value":0.0002,"adj.P.Val":0.0022},{"tissuedysplasia":0.3852,"tissuetumour":0.2661,"AveExpr":4.4117,"F":4.2897,"P.Value":0.0199,"adj.P.Val":0.0682},{"tissuedysplasia":-0.092,"tissuetumour":-0.4646,"AveExpr":5.5052,"F":9.3338,"P.Value":0.0004,"adj.P.Val":0.0041},{"tissuedysplasia":0.8519,"tissuetumour":1.236,"AveExpr":2.5477,"F":9.2753,"P.Value":0.0004,"adj.P.Val":0.0042},{"tissuedysplasia":2.9786,"tissuetumour":3.2361,"AveExpr":-4.348,"F":5.3018,"P.Value":0.0087,"adj.P.Val":0.0374},{"tissuedysplasia":0.8053,"tissuetumour":0.8058,"AveExpr":2.701,"F":8.3956,"P.Value":0.0008,"adj.P.Val":0.0067},{"tissuedysplasia":0.0766,"tissuetumour":-0.1091,"AveExpr":7.1868,"F":1.7637,"P.Value":0.1833,"adj.P.Val":0.3184},{"tissuedysplasia":0.4934,"tissuetumour":1.2446,"AveExpr":-2.4534,"F":1.4686,"P.Value":0.2413,"adj.P.Val":0.3836},{"tissuedysplasia":0.4339,"tissuetumour":0.7937,"AveExpr":3.8479,"F":9.9818,"P.Value":0.0003,"adj.P.Val":0.0029},{"tissuedysplasia":-0.0097,"tissuetumour":-0.2959,"AveExpr":4.9828,"F":4.0047,"P.Value":0.0252,"adj.P.Val":0.0808},{"tissuedysplasia":0.4092,"tissuetumour":1.2336,"AveExpr":-3.0883,"F":1.3794,"P.Value":0.2624,"adj.P.Val":0.4057},{"tissuedysplasia":0.2271,"tissuetumour":0.4864,"AveExpr":5.0981,"F":9.3633,"P.Value":0.0004,"adj.P.Val":0.004},{"tissuedysplasia":-0.3614,"tissuetumour":0.5531,"AveExpr":-2.6663,"F":0.5868,"P.Value":0.5604,"adj.P.Val":0.6764},{"tissuedysplasia":-0.1285,"tissuetumour":0.5269,"AveExpr":0.9729,"F":1.6685,"P.Value":0.2002,"adj.P.Val":0.3373},{"tissuedysplasia":0.0712,"tissuetumour":-0.6277,"AveExpr":2.6685,"F":3.8776,"P.Value":0.0281,"adj.P.Val":0.0872},{"tissuedysplasia":0.1655,"tissuetumour":1.1328,"AveExpr":-2.0135,"F":1.4022,"P.Value":0.2568,"adj.P.Val":0.3998},{"tissuedysplasia":-0.8224,"tissuetumour":-3.4497,"AveExpr":5.0386,"F":20.3322,"P.Value":1.093e-06,"adj.P.Val":0},{"tissuedysplasia":0.1616,"tissuetumour":-0.0644,"AveExpr":3.1525,"F":0.9254,"P.Value":0.404,"adj.P.Val":0.5439},{"tissuedysplasia":0.5293,"tissuetumour":0.3797,"AveExpr":6.313,"F":6.2942,"P.Value":0.0039,"adj.P.Val":0.0211},{"tissuedysplasia":-0.0106,"tissuetumour":-0.0007,"AveExpr":4.4069,"F":0.0007,"P.Value":0.9993,"adj.P.Val":0.9994},{"tissuedysplasia":0.1605,"tissuetumour":0.9606,"AveExpr":2.6826,"F":8.3996,"P.Value":0.0008,"adj.P.Val":0.0067},{"tissuedysplasia":0.1589,"tissuetumour":0.1518,"AveExpr":6.8136,"F":0.8867,"P.Value":0.4192,"adj.P.Val":0.5575},{"tissuedysplasia":-0.3786,"tissuetumour":-0.7483,"AveExpr":6.0024,"F":22.1905,"P.Value":2.1669e-07,"adj.P.Val":0},{"tissuedysplasia":0.7782,"tissuetumour":0.7462,"AveExpr":3.8819,"F":4.4953,"P.Value":0.0167,"adj.P.Val":0.0603},{"tissuedysplasia":0.5978,"tissuetumour":1.8323,"AveExpr":3.5905,"F":22.8435,"P.Value":1.5691e-07,"adj.P.Val":9.7818e-06},{"tissuedysplasia":0.0315,"tissuetumour":-0.1485,"AveExpr":5.9074,"F":1.6037,"P.Value":0.2127,"adj.P.Val":0.3517},{"tissuedysplasia":0.1895,"tissuetumour":0.3384,"AveExpr":5.3042,"F":1.381,"P.Value":0.262,"adj.P.Val":0.4052},{"tissuedysplasia":1.1007,"tissuetumour":2.9042,"AveExpr":-5.4067,"F":5.8089,"P.Value":0.0058,"adj.P.Val":0.0279},{"tissuedysplasia":0.3232,"tissuetumour":0.6126,"AveExpr":3.5855,"F":9.608,"P.Value":0.0003,"adj.P.Val":0.0035},{"tissuedysplasia":-1.1734,"tissuetumour":0.9439,"AveExpr":2.425,"F":7.1031,"P.Value":0.0023,"adj.P.Val":0.0141},{"tissuedysplasia":0.0923,"tissuetumour":-0.1808,"AveExpr":3.3643,"F":0.8313,"P.Value":0.4422,"adj.P.Val":0.5781},{"tissuedysplasia":0.2102,"tissuetumour":0.8164,"AveExpr":-0.6303,"F":1.4389,"P.Value":0.2481,"adj.P.Val":0.3908},{"tissuedysplasia":-0.5153,"tissuetumour":-0.3509,"AveExpr":0.3065,"F":0.9638,"P.Value":0.3894,"adj.P.Val":0.5304},{"tissuedysplasia":0.036,"tissuetumour":1.0535,"AveExpr":5.9075,"F":26.4406,"P.Value":2.8734e-08,"adj.P.Val":2.9257e-06},{"tissuedysplasia":0.2713,"tissuetumour":1.3792,"AveExpr":-0.6821,"F":4.8091,"P.Value":0.0129,"adj.P.Val":0.0498},{"tissuedysplasia":0.9948,"tissuetumour":0.3799,"AveExpr":3.6963,"F":5.3634,"P.Value":0.0082,"adj.P.Val":0.0361},{"tissuedysplasia":0.5111,"tissuetumour":0.8103,"AveExpr":3.6124,"F":9.4696,"P.Value":0.0004,"adj.P.Val":0.0038},{"tissuedysplasia":0.7946,"tissuetumour":2.5772,"AveExpr":-1.8895,"F":7.1865,"P.Value":0.002,"adj.P.Val":0.0129},{"tissuedysplasia":0.6267,"tissuetumour":1.0986,"AveExpr":4.9895,"F":10.9776,"P.Value":0.0001,"adj.P.Val":0.0018},{"tissuedysplasia":1.1643,"tissuetumour":1.5048,"AveExpr":9.2035,"F":18.1998,"P.Value":3.4209e-06,"adj.P.Val":0.0001},{"tissuedysplasia":0.3608,"tissuetumour":0.3706,"AveExpr":2.2942,"F":1.105,"P.Value":0.3402,"adj.P.Val":0.4843},{"tissuedysplasia":-0.0072,"tissuetumour":-0.4048,"AveExpr":5.9161,"F":15.9571,"P.Value":6.1474e-06,"adj.P.Val":0.0002},{"tissuedysplasia":0.5015,"tissuetumour":1.2047,"AveExpr":3.8358,"F":20.0997,"P.Value":6.2946e-07,"adj.P.Val":0},{"tissuedysplasia":0.013,"tissuetumour":0.4329,"AveExpr":4.4081,"F":6.1558,"P.Value":0.0044,"adj.P.Val":0.0229},{"tissuedysplasia":1.2742,"tissuetumour":1.0314,"AveExpr":-5.9262,"F":0.664,"P.Value":0.5199,"adj.P.Val":0.6437},{"tissuedysplasia":0.4437,"tissuetumour":0.7434,"AveExpr":3.6644,"F":12.009,"P.Value":0.0001,"adj.P.Val":0.001},{"tissuedysplasia":0.4038,"tissuetumour":0.4385,"AveExpr":3.2556,"F":2.5718,"P.Value":0.0878,"adj.P.Val":0.1927},{"tissuedysplasia":0.3533,"tissuetumour":1.0743,"AveExpr":4.5041,"F":9.8333,"P.Value":0.0003,"adj.P.Val":0.0031},{"tissuedysplasia":0.8981,"tissuetumour":1.0004,"AveExpr":3.6173,"F":8.5934,"P.Value":0.0007,"adj.P.Val":0.006},{"tissuedysplasia":0.1767,"tissuetumour":0.0743,"AveExpr":4.9489,"F":1.2192,"P.Value":0.3052,"adj.P.Val":0.4498},{"tissuedysplasia":0.185,"tissuetumour":0.3038,"AveExpr":1.066,"F":0.4464,"P.Value":0.6428,"adj.P.Val":0.7421},{"tissuedysplasia":-0.7652,"tissuetumour":-1.3458,"AveExpr":7.3713,"F":14.0893,"P.Value":0,"adj.P.Val":0.0005},{"tissuedysplasia":0.9689,"tissuetumour":1.8261,"AveExpr":-2.1263,"F":4.1145,"P.Value":0.023,"adj.P.Val":0.0756},{"tissuedysplasia":0.7553,"tissuetumour":1.438,"AveExpr":-2.662,"F":1.9322,"P.Value":0.1569,"adj.P.Val":0.2861},{"tissuedysplasia":0.1961,"tissuetumour":-0.0319,"AveExpr":4.3439,"F":1.9974,"P.Value":0.1478,"adj.P.Val":0.2749},{"tissuedysplasia":-0.0354,"tissuetumour":0.7079,"AveExpr":2.4768,"F":4.1546,"P.Value":0.0222,"adj.P.Val":0.0738},{"tissuedysplasia":0.0089,"tissuetumour":0.7173,"AveExpr":3.6745,"F":13.2184,"P.Value":0,"adj.P.Val":0.0006},{"tissuedysplasia":0.0349,"tissuetumour":-0.195,"AveExpr":6.1152,"F":3.3666,"P.Value":0.0436,"adj.P.Val":0.1185},{"tissuedysplasia":0.2269,"tissuetumour":0.2362,"AveExpr":2.5246,"F":0.4543,"P.Value":0.6378,"adj.P.Val":0.7383},{"tissuedysplasia":0.7023,"tissuetumour":2.5719,"AveExpr":-4.2612,"F":3.6834,"P.Value":0.0332,"adj.P.Val":0.0977},{"tissuedysplasia":-0.8534,"tissuetumour":-0.2419,"AveExpr":4.4826,"F":3.6658,"P.Value":0.0341,"adj.P.Val":0.0996},{"tissuedysplasia":0.1955,"tissuetumour":0.6458,"AveExpr":4.9393,"F":16.2645,"P.Value":5.148e-06,"adj.P.Val":0.0001},{"tissuedysplasia":0.3263,"tissuetumour":2.008,"AveExpr":-0.273,"F":10.9023,"P.Value":0.0001,"adj.P.Val":0.0018},{"tissuedysplasia":-0.0446,"tissuetumour":0.2644,"AveExpr":2.1205,"F":0.3778,"P.Value":0.6876,"adj.P.Val":0.7757},{"tissuedysplasia":0.5714,"tissuetumour":1.1156,"AveExpr":2.8759,"F":17.997,"P.Value":1.9432e-06,"adj.P.Val":0.0001},{"tissuedysplasia":-0.1819,"tissuetumour":-0.1828,"AveExpr":6.241,"F":4.297,"P.Value":0.0197,"adj.P.Val":0.068},{"tissuedysplasia":0.1952,"tissuetumour":0.226,"AveExpr":3.3347,"F":1.522,"P.Value":0.2295,"adj.P.Val":0.3706},{"tissuedysplasia":0.2179,"tissuetumour":-0.0048,"AveExpr":4.121,"F":1.7794,"P.Value":0.1807,"adj.P.Val":0.3153},{"tissuedysplasia":0.7382,"tissuetumour":1.5796,"AveExpr":2.8312,"F":21.334,"P.Value":3.3332e-07,"adj.P.Val":0},{"tissuedysplasia":-0.07,"tissuetumour":0.3253,"AveExpr":5.3901,"F":5.015,"P.Value":0.0109,"adj.P.Val":0.0441},{"tissuedysplasia":-0.0664,"tissuetumour":-0.1943,"AveExpr":6.4628,"F":2.4581,"P.Value":0.0973,"adj.P.Val":0.207},{"tissuedysplasia":0.1517,"tissuetumour":-0.2558,"AveExpr":6.2028,"F":2.623,"P.Value":0.0839,"adj.P.Val":0.1867},{"tissuedysplasia":0.1065,"tissuetumour":0.0231,"AveExpr":5.2683,"F":0.3531,"P.Value":0.7045,"adj.P.Val":0.7882},{"tissuedysplasia":-0.0044,"tissuetumour":0.0617,"AveExpr":3.4604,"F":0.0552,"P.Value":0.9464,"adj.P.Val":0.963},{"tissuedysplasia":1.1357,"tissuetumour":0.7025,"AveExpr":5.6838,"F":12.017,"P.Value":0.0001,"adj.P.Val":0.0011},{"tissuedysplasia":0.0365,"tissuetumour":-0.2788,"AveExpr":6.1819,"F":3.1562,"P.Value":0.0524,"adj.P.Val":0.1349},{"tissuedysplasia":-0.0349,"tissuetumour":0.0128,"AveExpr":1.2043,"F":0.016,"P.Value":0.9842,"adj.P.Val":0.989},{"tissuedysplasia":0.3391,"tissuetumour":0.2474,"AveExpr":2.8151,"F":0.7757,"P.Value":0.4666,"adj.P.Val":0.5979},{"tissuedysplasia":0.2345,"tissuetumour":0.0517,"AveExpr":4.179,"F":1.3741,"P.Value":0.2637,"adj.P.Val":0.4071},{"tissuedysplasia":0.2152,"tissuetumour":0.1779,"AveExpr":5.4106,"F":1.7362,"P.Value":0.188,"adj.P.Val":0.3239},{"tissuedysplasia":1.4186,"tissuetumour":2.065,"AveExpr":1.5926,"F":18.8598,"P.Value":1.2151e-06,"adj.P.Val":0},{"tissuedysplasia":0.4183,"tissuetumour":0.7693,"AveExpr":4.1552,"F":8.3259,"P.Value":0.0009,"adj.P.Val":0.0069},{"tissuedysplasia":0.6653,"tissuetumour":1.27,"AveExpr":4.0083,"F":14.9144,"P.Value":0,"adj.P.Val":0.0003},{"tissuedysplasia":-0.1183,"tissuetumour":-0.4519,"AveExpr":3.8668,"F":1.6997,"P.Value":0.1945,"adj.P.Val":0.331},{"tissuedysplasia":0.2213,"tissuetumour":-0.0238,"AveExpr":6.1335,"F":3.2714,"P.Value":0.0474,"adj.P.Val":0.1258},{"tissuedysplasia":0.2319,"tissuetumour":1.3435,"AveExpr":5.9543,"F":17.4866,"P.Value":3.3088e-06,"adj.P.Val":0.0001},{"tissuedysplasia":0.7293,"tissuetumour":0.572,"AveExpr":3.1779,"F":3.332,"P.Value":0.0449,"adj.P.Val":0.1212},{"tissuedysplasia":0.0299,"tissuetumour":0.2193,"AveExpr":2.037,"F":0.8647,"P.Value":0.4282,"adj.P.Val":0.5657},{"tissuedysplasia":-1.2242,"tissuetumour":-1.6409,"AveExpr":3.9934,"F":28.3823,"P.Value":1.2102e-08,"adj.P.Val":1.6073e-06},{"tissuedysplasia":0.0871,"tissuetumour":-0.2868,"AveExpr":2.3549,"F":2.6064,"P.Value":0.0852,"adj.P.Val":0.1886},{"tissuedysplasia":0.2593,"tissuetumour":0.3004,"AveExpr":5.4487,"F":7.7441,"P.Value":0.0013,"adj.P.Val":0.0095},{"tissuedysplasia":1.7184,"tissuetumour":1.9858,"AveExpr":0.5506,"F":15.5532,"P.Value":7.7789e-06,"adj.P.Val":0.0002},{"tissuedysplasia":0.6783,"tissuetumour":1.486,"AveExpr":2.2743,"F":14.6692,"P.Value":0,"adj.P.Val":0.0003},{"tissuedysplasia":1.1309,"tissuetumour":1.7874,"AveExpr":5.9478,"F":29.9651,"P.Value":7.5481e-09,"adj.P.Val":1.1498e-06},{"tissuedysplasia":0.3095,"tissuetumour":0.8755,"AveExpr":3.4838,"F":13.4976,"P.Value":0,"adj.P.Val":0.0005},{"tissuedysplasia":0.325,"tissuetumour":0.581,"AveExpr":2.9708,"F":6.319,"P.Value":0.0039,"adj.P.Val":0.0209},{"tissuedysplasia":0.2109,"tissuetumour":0.002,"AveExpr":5.7978,"F":2.3361,"P.Value":0.1086,"adj.P.Val":0.223},{"tissuedysplasia":0.086,"tissuetumour":0.1407,"AveExpr":6.4044,"F":1.03,"P.Value":0.3654,"adj.P.Val":0.5084},{"tissuedysplasia":0.3635,"tissuetumour":0.3849,"AveExpr":7.6972,"F":3.2991,"P.Value":0.0475,"adj.P.Val":0.1261},{"tissuedysplasia":0.3701,"tissuetumour":0.8701,"AveExpr":4.5343,"F":17.5068,"P.Value":2.5488e-06,"adj.P.Val":0.0001},{"tissuedysplasia":0.2163,"tissuetumour":-0.3738,"AveExpr":4.4178,"F":4.4344,"P.Value":0.0176,"adj.P.Val":0.0625},{"tissuedysplasia":0.5994,"tissuetumour":0.4854,"AveExpr":4.2215,"F":6.1932,"P.Value":0.0043,"adj.P.Val":0.0224},{"tissuedysplasia":0.4045,"tissuetumour":0.2368,"AveExpr":9.431,"F":3.6516,"P.Value":0.0359,"adj.P.Val":0.1033},{"tissuedysplasia":-0.7167,"tissuetumour":0.2748,"AveExpr":0.2894,"F":3.9892,"P.Value":0.0256,"adj.P.Val":0.0815},{"tissuedysplasia":0.4834,"tissuetumour":0.3878,"AveExpr":4.5878,"F":6.0416,"P.Value":0.0048,"adj.P.Val":0.0244},{"tissuedysplasia":1.182,"tissuetumour":1.0303,"AveExpr":4.1224,"F":11.094,"P.Value":0.0001,"adj.P.Val":0.0017},{"tissuedysplasia":0.9186,"tissuetumour":1.1418,"AveExpr":5.1754,"F":28.0429,"P.Value":1.4043e-08,"adj.P.Val":1.7983e-06},{"tissuedysplasia":-0.5033,"tissuetumour":-0.7639,"AveExpr":2.5349,"F":8.9472,"P.Value":0.0005,"adj.P.Val":0.005},{"tissuedysplasia":0.0672,"tissuetumour":-0.151,"AveExpr":5.9449,"F":2.59,"P.Value":0.0864,"adj.P.Val":0.1905},{"tissuedysplasia":0.2566,"tissuetumour":1.8794,"AveExpr":-0.8285,"F":9.3791,"P.Value":0.0004,"adj.P.Val":0.004},{"tissuedysplasia":0.7074,"tissuetumour":0.3177,"AveExpr":5.3695,"F":7.8309,"P.Value":0.0012,"adj.P.Val":0.0091},{"tissuedysplasia":-0.0411,"tissuetumour":0.2507,"AveExpr":2.6845,"F":1.2939,"P.Value":0.2844,"adj.P.Val":0.4289},{"tissuedysplasia":0.8366,"tissuetumour":1.6206,"AveExpr":1.9467,"F":10.8232,"P.Value":0.0002,"adj.P.Val":0.0019},{"tissuedysplasia":0.1095,"tissuetumour":-0.3664,"AveExpr":4.2013,"F":4.7009,"P.Value":0.0141,"adj.P.Val":0.0532},{"tissuedysplasia":0.7945,"tissuetumour":1.0163,"AveExpr":3.8669,"F":8.529,"P.Value":0.0007,"adj.P.Val":0.0062},{"tissuedysplasia":0.1476,"tissuetumour":0.0202,"AveExpr":6.0097,"F":1.1475,"P.Value":0.3268,"adj.P.Val":0.4709},{"tissuedysplasia":0.903,"tissuetumour":1.3259,"AveExpr":3.3168,"F":10.8993,"P.Value":0.0001,"adj.P.Val":0.0018},{"tissuedysplasia":0.287,"tissuetumour":0.5725,"AveExpr":5.6211,"F":8.3986,"P.Value":0.0008,"adj.P.Val":0.0067},{"tissuedysplasia":-0.0055,"tissuetumour":-0.0005,"AveExpr":5.9192,"F":0.0032,"P.Value":0.9968,"adj.P.Val":0.9978},{"tissuedysplasia":0.0977,"tissuetumour":-0.2707,"AveExpr":3.141,"F":1.9518,"P.Value":0.1541,"adj.P.Val":0.2828},{"tissuedysplasia":0.0704,"tissuetumour":-0.1674,"AveExpr":6.6589,"F":0.9225,"P.Value":0.4053,"adj.P.Val":0.5449},{"tissuedysplasia":0.0115,"tissuetumour":-0.3518,"AveExpr":6.7462,"F":8.2949,"P.Value":0.0009,"adj.P.Val":0.007},{"tissuedysplasia":0.3086,"tissuetumour":0.0532,"AveExpr":5.6252,"F":4.5314,"P.Value":0.0162,"adj.P.Val":0.0589},{"tissuedysplasia":0.4963,"tissuetumour":1.749,"AveExpr":2.3888,"F":13.7146,"P.Value":0,"adj.P.Val":0.0005},{"tissuedysplasia":0.523,"tissuetumour":3.5374,"AveExpr":-4.2388,"F":8.1731,"P.Value":0.001,"adj.P.Val":0.0075},{"tissuedysplasia":-0.1115,"tissuetumour":-0.2878,"AveExpr":2.7109,"F":1.2812,"P.Value":0.2879,"adj.P.Val":0.4324},{"tissuedysplasia":0.2948,"tissuetumour":0.0943,"AveExpr":5.8388,"F":3.4369,"P.Value":0.041,"adj.P.Val":0.1137},{"tissuedysplasia":0.5567,"tissuetumour":1.3058,"AveExpr":4.325,"F":24.5347,"P.Value":6.9494e-08,"adj.P.Val":5.4503e-06},{"tissuedysplasia":0.2555,"tissuetumour":0.0029,"AveExpr":5.2404,"F":3.2717,"P.Value":0.0474,"adj.P.Val":0.1258},{"tissuedysplasia":0.6348,"tissuetumour":1.1798,"AveExpr":3.1365,"F":24.6961,"P.Value":6.4396e-08,"adj.P.Val":5.1836e-06},{"tissuedysplasia":-0.2182,"tissuetumour":0.6792,"AveExpr":0.4403,"F":2.0973,"P.Value":0.1349,"adj.P.Val":0.2586},{"tissuedysplasia":-0.6236,"tissuetumour":-2.0817,"AveExpr":6.0153,"F":57.2924,"P.Value":5.6175e-13,"adj.P.Val":2.1735e-09},{"tissuedysplasia":-0.188,"tissuetumour":0.4552,"AveExpr":5.7912,"F":7.8041,"P.Value":0.0013,"adj.P.Val":0.0092},{"tissuedysplasia":0.0852,"tissuetumour":-0.3528,"AveExpr":4.9401,"F":6.3245,"P.Value":0.0039,"adj.P.Val":0.0208},{"tissuedysplasia":0.1381,"tissuetumour":0.1241,"AveExpr":4.9065,"F":0.4943,"P.Value":0.6134,"adj.P.Val":0.7196},{"tissuedysplasia":0.4607,"tissuetumour":0.0239,"AveExpr":1.3123,"F":2.3443,"P.Value":0.1078,"adj.P.Val":0.2219},{"tissuedysplasia":-0.1257,"tissuetumour":0.1897,"AveExpr":5.7309,"F":5.84,"P.Value":0.0056,"adj.P.Val":0.0274},{"tissuedysplasia":0.1601,"tissuetumour":0.3682,"AveExpr":4.5423,"F":4.5929,"P.Value":0.0154,"adj.P.Val":0.0568},{"tissuedysplasia":-0.0064,"tissuetumour":-0.1553,"AveExpr":5.4353,"F":1.4155,"P.Value":0.2536,"adj.P.Val":0.3965},{"tissuedysplasia":0.2854,"tissuetumour":0.5137,"AveExpr":3.2448,"F":4.9063,"P.Value":0.0119,"adj.P.Val":0.0469},{"tissuedysplasia":0.227,"tissuetumour":1.0768,"AveExpr":4.1712,"F":19.6151,"P.Value":8.1205e-07,"adj.P.Val":0},{"tissuedysplasia":0.0719,"tissuetumour":0.1967,"AveExpr":2.8159,"F":0.5091,"P.Value":0.6045,"adj.P.Val":0.7125},{"tissuedysplasia":0.488,"tissuetumour":0.9171,"AveExpr":2.7831,"F":12.0978,"P.Value":0.0001,"adj.P.Val":0.001},{"tissuedysplasia":0.165,"tissuetumour":0.064,"AveExpr":5.0185,"F":1.2153,"P.Value":0.3064,"adj.P.Val":0.4509},{"tissuedysplasia":0.3703,"tissuetumour":0.6034,"AveExpr":5.0278,"F":9.2964,"P.Value":0.0004,"adj.P.Val":0.0042},{"tissuedysplasia":0.6198,"tissuetumour":1.529,"AveExpr":4.3817,"F":47.8004,"P.Value":9.2868e-12,"adj.P.Val":1.1108e-08},{"tissuedysplasia":-0.1945,"tissuetumour":0.4389,"AveExpr":1.9669,"F":1.9509,"P.Value":0.1542,"adj.P.Val":0.2829},{"tissuedysplasia":0.4835,"tissuetumour":0.3991,"AveExpr":4.3054,"F":4.529,"P.Value":0.0163,"adj.P.Val":0.059},{"tissuedysplasia":0.2381,"tissuetumour":-0.2474,"AveExpr":6.5519,"F":5.1231,"P.Value":0.01,"adj.P.Val":0.0414},{"tissuedysplasia":-0.3772,"tissuetumour":-0.0234,"AveExpr":5.1409,"F":4.9167,"P.Value":0.0118,"adj.P.Val":0.0466},{"tissuedysplasia":0.0391,"tissuetumour":0.1273,"AveExpr":5.7757,"F":0.8998,"P.Value":0.414,"adj.P.Val":0.5527},{"tissuedysplasia":-0.0687,"tissuetumour":-0.513,"AveExpr":5.7845,"F":12.8038,"P.Value":0,"adj.P.Val":0.0007},{"tissuedysplasia":0.4844,"tissuetumour":0.178,"AveExpr":5.6487,"F":4.0815,"P.Value":0.0237,"adj.P.Val":0.0771},{"tissuedysplasia":0.9362,"tissuetumour":2.0914,"AveExpr":1.9646,"F":32.8489,"P.Value":1.8674e-09,"adj.P.Val":3.9494e-07},{"tissuedysplasia":-0.1253,"tissuetumour":0.0159,"AveExpr":4.8242,"F":0.5645,"P.Value":0.5727,"adj.P.Val":0.6865},{"tissuedysplasia":0.627,"tissuetumour":1.6184,"AveExpr":0.2905,"F":8.2617,"P.Value":0.0009,"adj.P.Val":0.0072},{"tissuedysplasia":0.5069,"tissuetumour":1.5683,"AveExpr":8.2499,"F":21.5406,"P.Value":6.8142e-07,"adj.P.Val":0},{"tissuedysplasia":0.5344,"tissuetumour":0.2907,"AveExpr":2.5956,"F":2.4513,"P.Value":0.0979,"adj.P.Val":0.2078},{"tissuedysplasia":0.4382,"tissuetumour":-0.1858,"AveExpr":6.0861,"F":8.9577,"P.Value":0.0005,"adj.P.Val":0.0049},{"tissuedysplasia":-0.0799,"tissuetumour":-0.5969,"AveExpr":3.0509,"F":5.2178,"P.Value":0.0093,"adj.P.Val":0.0392},{"tissuedysplasia":-0.0066,"tissuetumour":-0.1697,"AveExpr":4.1788,"F":0.4475,"P.Value":0.6421,"adj.P.Val":0.7416},{"tissuedysplasia":-1.6357,"tissuetumour":-1.7722,"AveExpr":-2.1452,"F":2.871,"P.Value":0.0673,"adj.P.Val":0.1603},{"tissuedysplasia":-0.0889,"tissuetumour":-0.2819,"AveExpr":5.1614,"F":4.253,"P.Value":0.0205,"adj.P.Val":0.0697},{"tissuedysplasia":0.2071,"tissuetumour":0.0007,"AveExpr":4.0972,"F":1.1589,"P.Value":0.3232,"adj.P.Val":0.4674},{"tissuedysplasia":0.2849,"tissuetumour":0.1986,"AveExpr":6.9035,"F":3.4836,"P.Value":0.0394,"adj.P.Val":0.1104},{"tissuedysplasia":0.3601,"tissuetumour":0.5367,"AveExpr":1.8463,"F":2.918,"P.Value":0.0646,"adj.P.Val":0.1559},{"tissuedysplasia":0.2889,"tissuetumour":0.5131,"AveExpr":4.0384,"F":2.1531,"P.Value":0.1282,"adj.P.Val":0.2492},{"tissuedysplasia":0.2325,"tissuetumour":-0.0224,"AveExpr":5.1562,"F":2.5613,"P.Value":0.0887,"adj.P.Val":0.194},{"tissuedysplasia":0.2391,"tissuetumour":-0.144,"AveExpr":4.9527,"F":3.4496,"P.Value":0.0406,"adj.P.Val":0.1127},{"tissuedysplasia":0.9228,"tissuetumour":0.8078,"AveExpr":-0.4973,"F":1.8409,"P.Value":0.1707,"adj.P.Val":0.3032},{"tissuedysplasia":-0.2378,"tissuetumour":0.4804,"AveExpr":6.4399,"F":9.9747,"P.Value":0.0003,"adj.P.Val":0.0029},{"tissuedysplasia":0.1143,"tissuetumour":0.096,"AveExpr":5.1403,"F":0.3791,"P.Value":0.6867,"adj.P.Val":0.7753},{"tissuedysplasia":0.2157,"tissuetumour":-0.1663,"AveExpr":5.1648,"F":4.4442,"P.Value":0.0175,"adj.P.Val":0.0622},{"tissuedysplasia":-0.2533,"tissuetumour":-1.0658,"AveExpr":6.3588,"F":31.0274,"P.Value":3.926e-09,"adj.P.Val":7.0982e-07},{"tissuedysplasia":-0.4077,"tissuetumour":1.5694,"AveExpr":0.603,"F":9.8388,"P.Value":0.0003,"adj.P.Val":0.0031},{"tissuedysplasia":1.1929,"tissuetumour":2.1756,"AveExpr":-6.4673,"F":1.9129,"P.Value":0.1597,"adj.P.Val":0.2897},{"tissuedysplasia":0.7552,"tissuetumour":1.7935,"AveExpr":0.6656,"F":9.8875,"P.Value":0.0003,"adj.P.Val":0.0031},{"tissuedysplasia":0.1612,"tissuetumour":-0.2053,"AveExpr":5.5632,"F":4.9103,"P.Value":0.0119,"adj.P.Val":0.0468},{"tissuedysplasia":0.5835,"tissuetumour":1.2712,"AveExpr":-0.4903,"F":3.3271,"P.Value":0.0451,"adj.P.Val":0.1215},{"tissuedysplasia":-0.7136,"tissuetumour":-0.5855,"AveExpr":3.4112,"F":5.8298,"P.Value":0.0057,"adj.P.Val":0.0276},{"tissuedysplasia":0.1011,"tissuetumour":2.5924,"AveExpr":-2.3177,"F":12.1973,"P.Value":0.0001,"adj.P.Val":0.0009},{"tissuedysplasia":0.4597,"tissuetumour":0.702,"AveExpr":3.2636,"F":8.2721,"P.Value":0.0009,"adj.P.Val":0.0071},{"tissuedysplasia":0.1028,"tissuetumour":0.2354,"AveExpr":1.9488,"F":0.799,"P.Value":0.4562,"adj.P.Val":0.5901},{"tissuedysplasia":0.6372,"tissuetumour":1.6709,"AveExpr":2.4045,"F":14.2688,"P.Value":0,"adj.P.Val":0.0003},{"tissuedysplasia":0.5606,"tissuetumour":0.5383,"AveExpr":5.9667,"F":8.3003,"P.Value":0.0009,"adj.P.Val":0.007},{"tissuedysplasia":-0.0691,"tissuetumour":-0.2359,"AveExpr":4.8733,"F":2.0238,"P.Value":0.1443,"adj.P.Val":0.2702},{"tissuedysplasia":0.0141,"tissuetumour":-0.05,"AveExpr":3.7112,"F":0.0985,"P.Value":0.9064,"adj.P.Val":0.9347},{"tissuedysplasia":0.2476,"tissuetumour":0.357,"AveExpr":4.0104,"F":3.7631,"P.Value":0.031,"adj.P.Val":0.0933},{"tissuedysplasia":0.2701,"tissuetumour":0.5269,"AveExpr":2.2696,"F":3.4114,"P.Value":0.0419,"adj.P.Val":0.1154},{"tissuedysplasia":1.9061,"tissuetumour":3.1617,"AveExpr":-0.5822,"F":17.9091,"P.Value":2.0396e-06,"adj.P.Val":0.0001},{"tissuedysplasia":-0.0272,"tissuetumour":-0.2503,"AveExpr":3.2919,"F":2.1019,"P.Value":0.1343,"adj.P.Val":0.2579},{"tissuedysplasia":0.1902,"tissuetumour":-0.0565,"AveExpr":4.4711,"F":1.1455,"P.Value":0.3274,"adj.P.Val":0.4716},{"tissuedysplasia":0.1032,"tissuetumour":0.0167,"AveExpr":6.8075,"F":0.4216,"P.Value":0.6586,"adj.P.Val":0.7542},{"tissuedysplasia":0.3818,"tissuetumour":0.3298,"AveExpr":4.7569,"F":3.7163,"P.Value":0.0323,"adj.P.Val":0.0959},{"tissuedysplasia":0.5327,"tissuetumour":1.974,"AveExpr":2.7553,"F":25.9582,"P.Value":3.5812e-08,"adj.P.Val":3.4478e-06},{"tissuedysplasia":0.2492,"tissuetumour":0.4962,"AveExpr":4.6514,"F":6.2629,"P.Value":0.004,"adj.P.Val":0.0215},{"tissuedysplasia":0.5172,"tissuetumour":0.8777,"AveExpr":6.833,"F":18.3704,"P.Value":1.5839e-06,"adj.P.Val":0.0001},{"tissuedysplasia":0.4205,"tissuetumour":1.1235,"AveExpr":6.834,"F":8.2134,"P.Value":0.0011,"adj.P.Val":0.0084},{"tissuedysplasia":0.13,"tissuetumour":0.195,"AveExpr":5.5018,"F":1.4892,"P.Value":0.2367,"adj.P.Val":0.3786},{"tissuedysplasia":0.583,"tissuetumour":1.1739,"AveExpr":2.9798,"F":11.0917,"P.Value":0.0001,"adj.P.Val":0.0017},{"tissuedysplasia":0.0671,"tissuetumour":-0.1167,"AveExpr":7.8864,"F":1.0129,"P.Value":0.372,"adj.P.Val":0.5145},{"tissuedysplasia":0.0322,"tissuetumour":-0.3609,"AveExpr":5.5055,"F":7.5041,"P.Value":0.0016,"adj.P.Val":0.0108},{"tissuedysplasia":0.6808,"tissuetumour":2.1531,"AveExpr":1.5966,"F":15.9518,"P.Value":6.1666e-06,"adj.P.Val":0.0002},{"tissuedysplasia":0.1309,"tissuetumour":-0.3453,"AveExpr":5.3186,"F":5.6327,"P.Value":0.0066,"adj.P.Val":0.0309},{"tissuedysplasia":0.0585,"tissuetumour":0.7555,"AveExpr":4.9186,"F":9.6232,"P.Value":0.0003,"adj.P.Val":0.0035},{"tissuedysplasia":-0.359,"tissuetumour":0.9438,"AveExpr":-0.7259,"F":1.9203,"P.Value":0.1587,"adj.P.Val":0.2884},{"tissuedysplasia":0.6141,"tissuetumour":1.2065,"AveExpr":4.7343,"F":9.3708,"P.Value":0.0004,"adj.P.Val":0.004},{"tissuedysplasia":0.0862,"tissuetumour":1.5816,"AveExpr":-3.2948,"F":3.2022,"P.Value":0.0503,"adj.P.Val":0.1312},{"tissuedysplasia":0.178,"tissuetumour":0.8452,"AveExpr":4.1173,"F":14.6335,"P.Value":0,"adj.P.Val":0.0003},{"tissuedysplasia":0.5615,"tissuetumour":1.4929,"AveExpr":-2.285,"F":2.4552,"P.Value":0.0975,"adj.P.Val":0.2073},{"tissuedysplasia":0.581,"tissuetumour":1.0449,"AveExpr":3.8163,"F":19.5726,"P.Value":8.3053e-07,"adj.P.Val":0},{"tissuedysplasia":-0.3767,"tissuetumour":-0.505,"AveExpr":3.755,"F":3.747,"P.Value":0.0314,"adj.P.Val":0.0942},{"tissuedysplasia":0.0944,"tissuetumour":-0.4705,"AveExpr":5.0034,"F":9.8642,"P.Value":0.0003,"adj.P.Val":0.0031},{"tissuedysplasia":0.1877,"tissuetumour":0.7674,"AveExpr":2.0086,"F":8.4779,"P.Value":0.0008,"adj.P.Val":0.0064},{"tissuedysplasia":0.2724,"tissuetumour":0.3671,"AveExpr":3.3468,"F":2.7149,"P.Value":0.0773,"adj.P.Val":0.1765},{"tissuedysplasia":0.769,"tissuetumour":1.5707,"AveExpr":-0.9903,"F":5.711,"P.Value":0.0062,"adj.P.Val":0.0295},{"tissuedysplasia":-0.0423,"tissuetumour":0.4806,"AveExpr":3.8524,"F":5.187,"P.Value":0.0095,"adj.P.Val":0.0399},{"tissuedysplasia":1.0623,"tissuetumour":1.6131,"AveExpr":-1.6282,"F":3.1914,"P.Value":0.0508,"adj.P.Val":0.1321},{"tissuedysplasia":0.624,"tissuetumour":0.6934,"AveExpr":5.2341,"F":17.0626,"P.Value":3.2688e-06,"adj.P.Val":0.0001},{"tissuedysplasia":-0.0542,"tissuetumour":-1.079,"AveExpr":5.4022,"F":22.1755,"P.Value":2.1831e-07,"adj.P.Val":0},{"tissuedysplasia":0.3095,"tissuetumour":0.2217,"AveExpr":5.1009,"F":1.952,"P.Value":0.1541,"adj.P.Val":0.2827},{"tissuedysplasia":0.5993,"tissuetumour":1.5619,"AveExpr":1.1873,"F":13.8789,"P.Value":0,"adj.P.Val":0.0004},{"tissuedysplasia":0.5883,"tissuetumour":0.4553,"AveExpr":5.2967,"F":10.6832,"P.Value":0.0002,"adj.P.Val":0.002},{"tissuedysplasia":0.1825,"tissuetumour":0.3609,"AveExpr":2.6207,"F":2.237,"P.Value":0.1188,"adj.P.Val":0.2369},{"tissuedysplasia":1.1153,"tissuetumour":2.4777,"AveExpr":7.9278,"F":19.5658,"P.Value":1.7501e-06,"adj.P.Val":0.0001},{"tissuedysplasia":0.2904,"tissuetumour":0.727,"AveExpr":2.7135,"F":9.728,"P.Value":0.0003,"adj.P.Val":0.0033},{"tissuedysplasia":0.3479,"tissuetumour":0.7767,"AveExpr":3.6287,"F":6.3956,"P.Value":0.0036,"adj.P.Val":0.02},{"tissuedysplasia":0.1876,"tissuetumour":0.3503,"AveExpr":3.6964,"F":3.7868,"P.Value":0.0304,"adj.P.Val":0.0921},{"tissuedysplasia":-0.0716,"tissuetumour":0.222,"AveExpr":2.8292,"F":2.0755,"P.Value":0.1376,"adj.P.Val":0.2619},{"tissuedysplasia":1.0143,"tissuetumour":1.8399,"AveExpr":-3.3645,"F":2.7881,"P.Value":0.0724,"adj.P.Val":0.1689},{"tissuedysplasia":-0.1287,"tissuetumour":-0.045,"AveExpr":2.3826,"F":0.2684,"P.Value":0.7658,"adj.P.Val":0.8339},{"tissuedysplasia":0.1683,"tissuetumour":0.2984,"AveExpr":4.4381,"F":2.4286,"P.Value":0.0999,"adj.P.Val":0.2108},{"tissuedysplasia":-0.1233,"tissuetumour":-0.2963,"AveExpr":4.6635,"F":2.5437,"P.Value":0.0901,"adj.P.Val":0.1962},{"tissuedysplasia":0.1144,"tissuetumour":0.6156,"AveExpr":4.6538,"F":10.3413,"P.Value":0.0002,"adj.P.Val":0.0024},{"tissuedysplasia":0.1115,"tissuetumour":0.3915,"AveExpr":3.6297,"F":4.4656,"P.Value":0.0172,"adj.P.Val":0.0614},{"tissuedysplasia":0.1013,"tissuetumour":1.2908,"AveExpr":1.454,"F":7.913,"P.Value":0.0012,"adj.P.Val":0.0086},{"tissuedysplasia":-0.5102,"tissuetumour":2.1471,"AveExpr":-4.6922,"F":4.2383,"P.Value":0.0207,"adj.P.Val":0.0703},{"tissuedysplasia":0.1094,"tissuetumour":1.0844,"AveExpr":1.5352,"F":8.5912,"P.Value":0.0007,"adj.P.Val":0.006},{"tissuedysplasia":0.7806,"tissuetumour":0.6146,"AveExpr":3.7518,"F":4.6606,"P.Value":0.0146,"adj.P.Val":0.0546},{"tissuedysplasia":-1.4394,"tissuetumour":-1.8544,"AveExpr":4.7605,"F":25.6585,"P.Value":4.1108e-08,"adj.P.Val":3.7631e-06},{"tissuedysplasia":0.2794,"tissuetumour":-0.2724,"AveExpr":5.2672,"F":6.1204,"P.Value":0.0045,"adj.P.Val":0.0233},{"tissuedysplasia":1.0405,"tissuetumour":1.4156,"AveExpr":4.8304,"F":27.0731,"P.Value":2.16e-08,"adj.P.Val":2.3772e-06},{"tissuedysplasia":2.1738,"tissuetumour":1.9608,"AveExpr":0.2321,"F":7.9861,"P.Value":0.0011,"adj.P.Val":0.0083},{"tissuedysplasia":0.9756,"tissuetumour":1.0933,"AveExpr":4.9418,"F":16.0849,"P.Value":5.7094e-06,"adj.P.Val":0.0002},{"tissuedysplasia":-0.0054,"tissuetumour":-0.1084,"AveExpr":3.9654,"F":0.2169,"P.Value":0.8059,"adj.P.Val":0.8637},{"tissuedysplasia":-0.004,"tissuetumour":0.7717,"AveExpr":-0.6115,"F":2.8799,"P.Value":0.0668,"adj.P.Val":0.1594},{"tissuedysplasia":1.4378,"tissuetumour":1.7976,"AveExpr":-9.8672,"F":1.7886,"P.Value":0.1791,"adj.P.Val":0.3136},{"tissuedysplasia":0.3773,"tissuetumour":1.0144,"AveExpr":3.9527,"F":23.7455,"P.Value":1.0125e-07,"adj.P.Val":7.2054e-06},{"tissuedysplasia":-0.2442,"tissuetumour":-0.5353,"AveExpr":7.1268,"F":12.4782,"P.Value":0.0001,"adj.P.Val":0.0008},{"tissuedysplasia":0.0648,"tissuetumour":0.1324,"AveExpr":4.1999,"F":0.7446,"P.Value":0.4808,"adj.P.Val":0.6102},{"tissuedysplasia":0.262,"tissuetumour":0.3414,"AveExpr":4.2517,"F":3.2217,"P.Value":0.0495,"adj.P.Val":0.1297},{"tissuedysplasia":-0.1011,"tissuetumour":-0.1612,"AveExpr":2.5124,"F":0.1411,"P.Value":0.8688,"adj.P.Val":0.9075},{"tissuedysplasia":0.2848,"tissuetumour":0.6968,"AveExpr":7.3673,"F":11.6379,"P.Value":0.0001,"adj.P.Val":0.0013},{"tissuedysplasia":0.6499,"tissuetumour":1.0001,"AveExpr":4.933,"F":6.878,"P.Value":0.0025,"adj.P.Val":0.0153},{"tissuedysplasia":0.1619,"tissuetumour":0.1402,"AveExpr":1.6399,"F":0.1915,"P.Value":0.8264,"adj.P.Val":0.8783},{"tissuedysplasia":0.0675,"tissuetumour":-0.2451,"AveExpr":4.7788,"F":2.7235,"P.Value":0.0767,"adj.P.Val":0.1755},{"tissuedysplasia":-0.0991,"tissuetumour":0.0202,"AveExpr":3.5794,"F":0.4091,"P.Value":0.6668,"adj.P.Val":0.7604},{"tissuedysplasia":1.2375,"tissuetumour":1.5899,"AveExpr":-5.061,"F":1.2851,"P.Value":0.2868,"adj.P.Val":0.4314},{"tissuedysplasia":-0.1161,"tissuetumour":-0.4103,"AveExpr":4.1651,"F":2.49,"P.Value":0.0945,"adj.P.Val":0.2029},{"tissuedysplasia":0.1892,"tissuetumour":0.1148,"AveExpr":4.352,"F":1.3943,"P.Value":0.2588,"adj.P.Val":0.4019},{"tissuedysplasia":-0.0862,"tissuetumour":-0.5305,"AveExpr":5.4778,"F":10.0509,"P.Value":0.0003,"adj.P.Val":0.0028},{"tissuedysplasia":0.1663,"tissuetumour":0.0084,"AveExpr":3.1108,"F":0.6708,"P.Value":0.5165,"adj.P.Val":0.6407},{"tissuedysplasia":0.2509,"tissuetumour":0.1209,"AveExpr":4.0585,"F":1.4074,"P.Value":0.2556,"adj.P.Val":0.3985},{"tissuedysplasia":0.2637,"tissuetumour":0.1417,"AveExpr":6.2201,"F":2.751,"P.Value":0.0749,"adj.P.Val":0.1726},{"tissuedysplasia":-0.282,"tissuetumour":-0.7363,"AveExpr":5.8997,"F":14.5448,"P.Value":0,"adj.P.Val":0.0003},{"tissuedysplasia":0.0859,"tissuetumour":-0.4746,"AveExpr":6.6881,"F":8.2658,"P.Value":0.0009,"adj.P.Val":0.0071},{"tissuedysplasia":0.1264,"tissuetumour":-0.2085,"AveExpr":7.205,"F":3.5355,"P.Value":0.0377,"adj.P.Val":0.107},{"tissuedysplasia":0.2377,"tissuetumour":0.1492,"AveExpr":3.3065,"F":0.2219,"P.Value":0.8019,"adj.P.Val":0.8611},{"tissuedysplasia":0.1091,"tissuetumour":0.1824,"AveExpr":4.6706,"F":1.4359,"P.Value":0.2488,"adj.P.Val":0.3914},{"tissuedysplasia":0.53,"tissuetumour":0.3984,"AveExpr":2.6679,"F":4.4959,"P.Value":0.0167,"adj.P.Val":0.0602},{"tissuedysplasia":-0.0625,"tissuetumour":-0.6259,"AveExpr":1.8718,"F":2.6951,"P.Value":0.0787,"adj.P.Val":0.1786},{"tissuedysplasia":0.0034,"tissuetumour":0.3646,"AveExpr":2.254,"F":1.418,"P.Value":0.2531,"adj.P.Val":0.396},{"tissuedysplasia":0.2469,"tissuetumour":0.0661,"AveExpr":3.8738,"F":1.9505,"P.Value":0.1543,"adj.P.Val":0.283},{"tissuedysplasia":0.0321,"tissuetumour":-0.6006,"AveExpr":3.0278,"F":8.3667,"P.Value":0.0008,"adj.P.Val":0.0068},{"tissuedysplasia":-0.0582,"tissuetumour":-0.1376,"AveExpr":5.334,"F":1.2803,"P.Value":0.2881,"adj.P.Val":0.4326},{"tissuedysplasia":-0.3143,"tissuetumour":-0.4494,"AveExpr":2.1471,"F":1.4808,"P.Value":0.2386,"adj.P.Val":0.3805},{"tissuedysplasia":0.0946,"tissuetumour":0.1223,"AveExpr":3.9678,"F":0.5337,"P.Value":0.5902,"adj.P.Val":0.7007},{"tissuedysplasia":-0.0026,"tissuetumour":0.4435,"AveExpr":7.8643,"F":12.9646,"P.Value":0,"adj.P.Val":0.0007},{"tissuedysplasia":0.2107,"tissuetumour":0.1316,"AveExpr":4.924,"F":1.6083,"P.Value":0.2118,"adj.P.Val":0.3507},{"tissuedysplasia":0.6754,"tissuetumour":3.1797,"AveExpr":-0.6172,"F":13.3011,"P.Value":0,"adj.P.Val":0.0006},{"tissuedysplasia":0.1182,"tissuetumour":0.1573,"AveExpr":4.328,"F":0.9037,"P.Value":0.4125,"adj.P.Val":0.5514},{"tissuedysplasia":0.5202,"tissuetumour":0.7673,"AveExpr":4.0018,"F":8.5255,"P.Value":0.0007,"adj.P.Val":0.0062},{"tissuedysplasia":0.176,"tissuetumour":0.7228,"AveExpr":4.0154,"F":15.4845,"P.Value":8.0989e-06,"adj.P.Val":0.0002},{"tissuedysplasia":-0.1113,"tissuetumour":-0.046,"AveExpr":3.6494,"F":0.3326,"P.Value":0.7188,"adj.P.Val":0.7988},{"tissuedysplasia":0.003,"tissuetumour":0.1284,"AveExpr":4.0457,"F":0.5731,"P.Value":0.568,"adj.P.Val":0.6824},{"tissuedysplasia":0.1602,"tissuetumour":0.0488,"AveExpr":3.9486,"F":0.6078,"P.Value":0.549,"adj.P.Val":0.6671},{"tissuedysplasia":0.0363,"tissuetumour":-0.457,"AveExpr":4.9658,"F":7.9186,"P.Value":0.0012,"adj.P.Val":0.0086},{"tissuedysplasia":1.0078,"tissuetumour":1.1905,"AveExpr":-0.7222,"F":3.5871,"P.Value":0.036,"adj.P.Val":0.1036},{"tissuedysplasia":0.0204,"tissuetumour":0.3605,"AveExpr":5.679,"F":3.6583,"P.Value":0.0339,"adj.P.Val":0.0992},{"tissuedysplasia":0.4489,"tissuetumour":0.9496,"AveExpr":1.9091,"F":3.093,"P.Value":0.0554,"adj.P.Val":0.1402},{"tissuedysplasia":-0.2198,"tissuetumour":-0.6597,"AveExpr":4.258,"F":7.79,"P.Value":0.0013,"adj.P.Val":0.0093},{"tissuedysplasia":-0.4335,"tissuetumour":2.2113,"AveExpr":-4.3387,"F":6.5736,"P.Value":0.0032,"adj.P.Val":0.0181},{"tissuedysplasia":0.1452,"tissuetumour":0.1716,"AveExpr":6.1188,"F":1.7763,"P.Value":0.1812,"adj.P.Val":0.3159},{"tissuedysplasia":-0.1998,"tissuetumour":-0.4594,"AveExpr":8.0164,"F":11.4209,"P.Value":0.0001,"adj.P.Val":0.0014},{"tissuedysplasia":0.4888,"tissuetumour":-0.0449,"AveExpr":5.8924,"F":4.9959,"P.Value":0.0111,"adj.P.Val":0.0446},{"tissuedysplasia":0.2477,"tissuetumour":0.3763,"AveExpr":6.6489,"F":7.6601,"P.Value":0.0014,"adj.P.Val":0.0099},{"tissuedysplasia":0.3011,"tissuetumour":0.1167,"AveExpr":5.2057,"F":3.5649,"P.Value":0.0367,"adj.P.Val":0.1051},{"tissuedysplasia":1.2412,"tissuetumour":2.1191,"AveExpr":0.8183,"F":12.3047,"P.Value":0.0001,"adj.P.Val":0.0009},{"tissuedysplasia":0.0101,"tissuetumour":-0.238,"AveExpr":4.4728,"F":0.8585,"P.Value":0.4308,"adj.P.Val":0.5679},{"tissuedysplasia":0.1016,"tissuetumour":-0.0716,"AveExpr":2.6817,"F":0.4779,"P.Value":0.6233,"adj.P.Val":0.7274},{"tissuedysplasia":0.9903,"tissuetumour":1.7083,"AveExpr":2.6625,"F":23.2287,"P.Value":1.3e-07,"adj.P.Val":8.6187e-06},{"tissuedysplasia":-0.0286,"tissuetumour":0.4768,"AveExpr":5.8164,"F":8.1872,"P.Value":0.0009,"adj.P.Val":0.0074},{"tissuedysplasia":0.0249,"tissuetumour":-0.0836,"AveExpr":4.9443,"F":0.2709,"P.Value":0.7639,"adj.P.Val":0.8326},{"tissuedysplasia":-0.0737,"tissuetumour":-0.3795,"AveExpr":5.0765,"F":3.6735,"P.Value":0.0335,"adj.P.Val":0.0983},{"tissuedysplasia":-0.1991,"tissuetumour":-0.3878,"AveExpr":1.8223,"F":1.1907,"P.Value":0.3136,"adj.P.Val":0.458},{"tissuedysplasia":1.0871,"tissuetumour":1.5071,"AveExpr":1.5076,"F":14.0703,"P.Value":0,"adj.P.Val":0.0004},{"tissuedysplasia":-0.2223,"tissuetumour":-0.2324,"AveExpr":3.9865,"F":0.9907,"P.Value":0.3794,"adj.P.Val":0.5215},{"tissuedysplasia":0.6904,"tissuetumour":1.8184,"AveExpr":5.6923,"F":23.7343,"P.Value":1.4371e-07,"adj.P.Val":9.2061e-06},{"tissuedysplasia":0.3913,"tissuetumour":0.6602,"AveExpr":4.9408,"F":8.5685,"P.Value":0.0007,"adj.P.Val":0.0061},{"tissuedysplasia":0.0648,"tissuetumour":0.2732,"AveExpr":3.8064,"F":0.2499,"P.Value":0.7801,"adj.P.Val":0.8444},{"tissuedysplasia":0.5168,"tissuetumour":0.6872,"AveExpr":2.829,"F":10.6905,"P.Value":0.0002,"adj.P.Val":0.002},{"tissuedysplasia":0.0528,"tissuetumour":-0.241,"AveExpr":7.5511,"F":9.4455,"P.Value":0.0004,"adj.P.Val":0.0038},{"tissuedysplasia":0.5979,"tissuetumour":1.0709,"AveExpr":5.3477,"F":28.9149,"P.Value":9.6027e-09,"adj.P.Val":1.3569e-06},{"tissuedysplasia":0.2755,"tissuetumour":0.3017,"AveExpr":5.0506,"F":2.0788,"P.Value":0.1372,"adj.P.Val":0.2615},{"tissuedysplasia":1.139,"tissuetumour":1.3003,"AveExpr":2.0681,"F":12.2997,"P.Value":0.0001,"adj.P.Val":0.0009},{"tissuedysplasia":0.1199,"tissuetumour":0.4269,"AveExpr":1.0076,"F":0.9941,"P.Value":0.3782,"adj.P.Val":0.5204},{"tissuedysplasia":0.4087,"tissuetumour":0.4228,"AveExpr":4.5947,"F":4.177,"P.Value":0.0218,"adj.P.Val":0.0729},{"tissuedysplasia":0.0754,"tissuetumour":-0.0111,"AveExpr":3.6145,"F":0.1641,"P.Value":0.8491,"adj.P.Val":0.8941},{"tissuedysplasia":-0.5303,"tissuetumour":0.3181,"AveExpr":0.7626,"F":2.6528,"P.Value":0.0817,"adj.P.Val":0.1833},{"tissuedysplasia":0.3071,"tissuetumour":0.4211,"AveExpr":3.8258,"F":4.1299,"P.Value":0.0227,"adj.P.Val":0.0749},{"tissuedysplasia":0.2577,"tissuetumour":1.2745,"AveExpr":4.7318,"F":22.5635,"P.Value":1.801e-07,"adj.P.Val":0},{"tissuedysplasia":0.457,"tissuetumour":0.8967,"AveExpr":6.1117,"F":16.8833,"P.Value":3.617e-06,"adj.P.Val":0.0001},{"tissuedysplasia":0.746,"tissuetumour":0.5237,"AveExpr":4.0948,"F":7.2537,"P.Value":0.0019,"adj.P.Val":0.0124},{"tissuedysplasia":-0.0037,"tissuetumour":-0.3472,"AveExpr":4.6205,"F":6.478,"P.Value":0.0034,"adj.P.Val":0.0191},{"tissuedysplasia":0.7411,"tissuetumour":0.8854,"AveExpr":6.5077,"F":12.2465,"P.Value":0.0001,"adj.P.Val":0.001},{"tissuedysplasia":0.1722,"tissuetumour":0.3746,"AveExpr":4.7858,"F":5.9081,"P.Value":0.0053,"adj.P.Val":0.0264},{"tissuedysplasia":1.1077,"tissuetumour":1.9539,"AveExpr":2.4259,"F":20.119,"P.Value":6.2314e-07,"adj.P.Val":0},{"tissuedysplasia":-0.1212,"tissuetumour":0.2136,"AveExpr":2.3336,"F":0.7817,"P.Value":0.4639,"adj.P.Val":0.5967},{"tissuedysplasia":-0.6528,"tissuetumour":-0.7745,"AveExpr":3.0912,"F":6.9528,"P.Value":0.0024,"adj.P.Val":0.0147},{"tissuedysplasia":1.8512,"tissuetumour":2.4879,"AveExpr":1.1651,"F":21.0197,"P.Value":3.912e-07,"adj.P.Val":0},{"tissuedysplasia":0.2314,"tissuetumour":-0.0411,"AveExpr":4.2203,"F":2.7246,"P.Value":0.0766,"adj.P.Val":0.1754},{"tissuedysplasia":0.2451,"tissuetumour":0.7663,"AveExpr":0.2845,"F":2.6703,"P.Value":0.0804,"adj.P.Val":0.1814},{"tissuedysplasia":0.5286,"tissuetumour":1.4239,"AveExpr":4.3288,"F":28.5318,"P.Value":1.1339e-08,"adj.P.Val":1.5304e-06},{"tissuedysplasia":0.0563,"tissuetumour":0.2656,"AveExpr":3.8544,"F":2.0662,"P.Value":0.1388,"adj.P.Val":0.2633},{"tissuedysplasia":0.6051,"tissuetumour":0.1339,"AveExpr":5.737,"F":6.4064,"P.Value":0.0036,"adj.P.Val":0.0199},{"tissuedysplasia":0.8731,"tissuetumour":2.012,"AveExpr":2.8359,"F":27.4538,"P.Value":1.8223e-08,"adj.P.Val":2.1541e-06},{"tissuedysplasia":0.5051,"tissuetumour":0.9195,"AveExpr":1.9198,"F":9.0336,"P.Value":0.0005,"adj.P.Val":0.0048},{"tissuedysplasia":0.6127,"tissuetumour":0.5794,"AveExpr":4.3688,"F":8.6343,"P.Value":0.0007,"adj.P.Val":0.0059},{"tissuedysplasia":0.1938,"tissuetumour":-0.2544,"AveExpr":5.1181,"F":5.619,"P.Value":0.0067,"adj.P.Val":0.0312},{"tissuedysplasia":0.8971,"tissuetumour":1.8728,"AveExpr":3.5128,"F":33.0794,"P.Value":1.7028e-09,"adj.P.Val":3.6637e-07},{"tissuedysplasia":-0.2224,"tissuetumour":-0.5183,"AveExpr":4.1078,"F":4.3854,"P.Value":0.0183,"adj.P.Val":0.0644},{"tissuedysplasia":-0.5158,"tissuetumour":-1.0954,"AveExpr":5.6379,"F":45.9438,"P.Value":1.6806e-11,"adj.P.Val":1.5633e-08},{"tissuedysplasia":0.5308,"tissuetumour":0.2464,"AveExpr":4.5131,"F":5.6297,"P.Value":0.0067,"adj.P.Val":0.0309},{"tissuedysplasia":-0.47,"tissuetumour":-0.7565,"AveExpr":6.2135,"F":17.4249,"P.Value":2.668e-06,"adj.P.Val":0.0001},{"tissuedysplasia":-0.1729,"tissuetumour":0.7517,"AveExpr":4.4647,"F":6.8345,"P.Value":0.0026,"adj.P.Val":0.0157},{"tissuedysplasia":0.2085,"tissuetumour":-0.2702,"AveExpr":4.6962,"F":6.3114,"P.Value":0.0039,"adj.P.Val":0.0209},{"tissuedysplasia":-0.0274,"tissuetumour":0.4738,"AveExpr":4.6351,"F":4.7626,"P.Value":0.0134,"adj.P.Val":0.0513},{"tissuedysplasia":0.2153,"tissuetumour":0.1239,"AveExpr":5.1284,"F":1.5112,"P.Value":0.2319,"adj.P.Val":0.373},{"tissuedysplasia":-0.24,"tissuetumour":-0.6836,"AveExpr":4.9374,"F":4.6794,"P.Value":0.0144,"adj.P.Val":0.0539},{"tissuedysplasia":0.0932,"tissuetumour":-0.1819,"AveExpr":6.0823,"F":3.3321,"P.Value":0.0449,"adj.P.Val":0.1212},{"tissuedysplasia":0.4666,"tissuetumour":0.859,"AveExpr":-0.1781,"F":1.3896,"P.Value":0.2599,"adj.P.Val":0.403},{"tissuedysplasia":-0.0527,"tissuetumour":-0.0748,"AveExpr":1.4072,"F":0.0148,"P.Value":0.9853,"adj.P.Val":0.9897},{"tissuedysplasia":-0.0198,"tissuetumour":-0.2534,"AveExpr":6.8304,"F":5.0145,"P.Value":0.0109,"adj.P.Val":0.0441},{"tissuedysplasia":-0.1623,"tissuetumour":-1.0138,"AveExpr":-1.2059,"F":1.3575,"P.Value":0.2679,"adj.P.Val":0.4115},{"tissuedysplasia":0.112,"tissuetumour":0.4102,"AveExpr":2.7358,"F":2.7317,"P.Value":0.0762,"adj.P.Val":0.1746},{"tissuedysplasia":0.1359,"tissuetumour":0.7859,"AveExpr":4.1365,"F":15.9014,"P.Value":6.3496e-06,"adj.P.Val":0.0002},{"tissuedysplasia":0.0432,"tissuetumour":0.2552,"AveExpr":2.977,"F":1.1227,"P.Value":0.3346,"adj.P.Val":0.4789},{"tissuedysplasia":0.2737,"tissuetumour":0.6556,"AveExpr":2.1098,"F":5.0032,"P.Value":0.011,"adj.P.Val":0.0444},{"tissuedysplasia":0.477,"tissuetumour":0.05,"AveExpr":5.3718,"F":4.1558,"P.Value":0.0222,"adj.P.Val":0.0738},{"tissuedysplasia":0.0325,"tissuetumour":0.4337,"AveExpr":2.8236,"F":2.4438,"P.Value":0.0985,"adj.P.Val":0.2088},{"tissuedysplasia":0.576,"tissuetumour":1.343,"AveExpr":7.393,"F":13.2992,"P.Value":0,"adj.P.Val":0.0008},{"tissuedysplasia":0.1349,"tissuetumour":0.1162,"AveExpr":4.8918,"F":0.8321,"P.Value":0.4419,"adj.P.Val":0.5777},{"tissuedysplasia":0.0856,"tissuetumour":0.0231,"AveExpr":4.7522,"F":0.3626,"P.Value":0.698,"adj.P.Val":0.7833},{"tissuedysplasia":0.5469,"tissuetumour":0.9689,"AveExpr":0.6397,"F":6.5424,"P.Value":0.0033,"adj.P.Val":0.0184},{"tissuedysplasia":0.1909,"tissuetumour":0.7375,"AveExpr":4.8931,"F":25.7844,"P.Value":3.879e-08,"adj.P.Val":3.6487e-06},{"tissuedysplasia":0.0153,"tissuetumour":0.2706,"AveExpr":6.729,"F":10.1691,"P.Value":0.0002,"adj.P.Val":0.0026},{"tissuedysplasia":-0.294,"tissuetumour":0.1594,"AveExpr":2.1631,"F":2.5734,"P.Value":0.0877,"adj.P.Val":0.1925},{"tissuedysplasia":-0.2853,"tissuetumour":0.6847,"AveExpr":-0.4658,"F":2.0416,"P.Value":0.1419,"adj.P.Val":0.2672},{"tissuedysplasia":0.5241,"tissuetumour":0.0963,"AveExpr":4.8353,"F":6.3478,"P.Value":0.0038,"adj.P.Val":0.0206},{"tissuedysplasia":0.1925,"tissuetumour":0.0305,"AveExpr":6.4587,"F":2.095,"P.Value":0.1352,"adj.P.Val":0.2589},{"tissuedysplasia":0.1483,"tissuetumour":0.3861,"AveExpr":3.1479,"F":1.4999,"P.Value":0.2343,"adj.P.Val":0.3758},{"tissuedysplasia":0.1747,"tissuetumour":0.0595,"AveExpr":4.7712,"F":1.2664,"P.Value":0.2919,"adj.P.Val":0.4364},{"tissuedysplasia":0.9136,"tissuetumour":3.0939,"AveExpr":-1.2952,"F":14.6683,"P.Value":0,"adj.P.Val":0.0003},{"tissuedysplasia":0.371,"tissuetumour":-0.0508,"AveExpr":2.4802,"F":2.1203,"P.Value":0.1321,"adj.P.Val":0.2548},{"tissuedysplasia":-0.0539,"tissuetumour":0.0452,"AveExpr":7.0549,"F":0.7839,"P.Value":0.4629,"adj.P.Val":0.5958},{"tissuedysplasia":0.3716,"tissuetumour":0.5764,"AveExpr":3.1745,"F":1.8862,"P.Value":0.1637,"adj.P.Val":0.2945},{"tissuedysplasia":0.1224,"tissuetumour":0.4096,"AveExpr":4.7071,"F":7.0274,"P.Value":0.0022,"adj.P.Val":0.0141},{"tissuedysplasia":0.2579,"tissuetumour":-0.0231,"AveExpr":9.1214,"F":3.2709,"P.Value":0.0492,"adj.P.Val":0.1293},{"tissuedysplasia":0.4411,"tissuetumour":0.475,"AveExpr":3.8345,"F":4.8359,"P.Value":0.0126,"adj.P.Val":0.049},{"tissuedysplasia":0.0972,"tissuetumour":-0.106,"AveExpr":5.6106,"F":1.1602,"P.Value":0.3228,"adj.P.Val":0.467},{"tissuedysplasia":0.7985,"tissuetumour":2.4913,"AveExpr":-6.2908,"F":2.1625,"P.Value":0.1271,"adj.P.Val":0.2479},{"tissuedysplasia":-0.3463,"tissuetumour":2.2156,"AveExpr":-5.5994,"F":4.3016,"P.Value":0.0197,"adj.P.Val":0.0678},{"tissuedysplasia":0.9248,"tissuetumour":3.1201,"AveExpr":-1.0443,"F":13.6591,"P.Value":0,"adj.P.Val":0.0005},{"tissuedysplasia":0.2577,"tissuetumour":0.5852,"AveExpr":5.4577,"F":16.0889,"P.Value":5.6961e-06,"adj.P.Val":0.0002},{"tissuedysplasia":0.4337,"tissuetumour":0.9606,"AveExpr":3.6354,"F":8.415,"P.Value":0.0008,"adj.P.Val":0.0066},{"tissuedysplasia":0.3132,"tissuetumour":0.397,"AveExpr":3.1903,"F":2.6423,"P.Value":0.0825,"adj.P.Val":0.1844},{"tissuedysplasia":0.211,"tissuetumour":-0.0139,"AveExpr":5.5369,"F":1.3782,"P.Value":0.2627,"adj.P.Val":0.406},{"tissuedysplasia":0.2509,"tissuetumour":0.6999,"AveExpr":5.4386,"F":28.4981,"P.Value":1.1506e-08,"adj.P.Val":1.5422e-06},{"tissuedysplasia":0.9132,"tissuetumour":2.2344,"AveExpr":-2.1036,"F":5.641,"P.Value":0.0066,"adj.P.Val":0.0308},{"tissuedysplasia":0.3191,"tissuetumour":0.0532,"AveExpr":5.3224,"F":3.9971,"P.Value":0.0254,"adj.P.Val":0.0812},{"tissuedysplasia":0.0118,"tissuetumour":-0.1778,"AveExpr":3.6158,"F":0.9547,"P.Value":0.3928,"adj.P.Val":0.5336},{"tissuedysplasia":1.0118,"tissuetumour":0.8523,"AveExpr":3.8252,"F":9.5467,"P.Value":0.0004,"adj.P.Val":0.0037},{"tissuedysplasia":0.1115,"tissuetumour":-0.2366,"AveExpr":6.2521,"F":4.4931,"P.Value":0.0168,"adj.P.Val":0.0603},{"tissuedysplasia":0.0598,"tissuetumour":0.0138,"AveExpr":4.2118,"F":0.0992,"P.Value":0.9057,"adj.P.Val":0.9343},{"tissuedysplasia":0.8101,"tissuetumour":0.9655,"AveExpr":6.7276,"F":15.1638,"P.Value":0,"adj.P.Val":0.0003},{"tissuedysplasia":-0.0167,"tissuetumour":0.4422,"AveExpr":5.8896,"F":6.3126,"P.Value":0.0039,"adj.P.Val":0.0209},{"tissuedysplasia":-0.0065,"tissuetumour":-0.3396,"AveExpr":8.1433,"F":12.3339,"P.Value":0.0001,"adj.P.Val":0.0009},{"tissuedysplasia":0.3263,"tissuetumour":0.1145,"AveExpr":2.8095,"F":0.905,"P.Value":0.412,"adj.P.Val":0.5509},{"tissuedysplasia":0.4027,"tissuetumour":0.2375,"AveExpr":-0.0308,"F":0.2951,"P.Value":0.746,"adj.P.Val":0.8194},{"tissuedysplasia":0.1267,"tissuetumour":-0.4407,"AveExpr":6.7848,"F":13.2029,"P.Value":0,"adj.P.Val":0.0006},{"tissuedysplasia":0.2117,"tissuetumour":-0.0218,"AveExpr":3.05,"F":1.4391,"P.Value":0.2481,"adj.P.Val":0.3908},{"tissuedysplasia":0.1661,"tissuetumour":1.1361,"AveExpr":2.0991,"F":7.0877,"P.Value":0.0021,"adj.P.Val":0.0136},{"tissuedysplasia":0.0869,"tissuetumour":0.6747,"AveExpr":-2.9397,"F":0.506,"P.Value":0.6064,"adj.P.Val":0.7141},{"tissuedysplasia":0.2893,"tissuetumour":0.0245,"AveExpr":0.6546,"F":0.3481,"P.Value":0.708,"adj.P.Val":0.7909},{"tissuedysplasia":0.1499,"tissuetumour":-0.1559,"AveExpr":5.8562,"F":4.0016,"P.Value":0.0253,"adj.P.Val":0.081},{"tissuedysplasia":0.2269,"tissuetumour":0.0346,"AveExpr":3.2248,"F":0.9875,"P.Value":0.3806,"adj.P.Val":0.5225},{"tissuedysplasia":0.8873,"tissuetumour":3.0112,"AveExpr":-3.245,"F":9.8924,"P.Value":0.0003,"adj.P.Val":0.0031},{"tissuedysplasia":-0.2691,"tissuetumour":0.0982,"AveExpr":-1.125,"F":0.2379,"P.Value":0.7893,"adj.P.Val":0.8514},{"tissuedysplasia":0.7625,"tissuetumour":0.8073,"AveExpr":4.3496,"F":9.781,"P.Value":0.0003,"adj.P.Val":0.0032},{"tissuedysplasia":0.2141,"tissuetumour":0.0912,"AveExpr":6.3212,"F":1.3961,"P.Value":0.2583,"adj.P.Val":0.4014},{"tissuedysplasia":-0.0028,"tissuetumour":0.0211,"AveExpr":1.4164,"F":0.0039,"P.Value":0.9961,"adj.P.Val":0.9974},{"tissuedysplasia":0.0407,"tissuetumour":0.0191,"AveExpr":4.8156,"F":0.0998,"P.Value":0.9052,"adj.P.Val":0.934},{"tissuedysplasia":0.2671,"tissuetumour":1.3332,"AveExpr":-1.5445,"F":2.157,"P.Value":0.1277,"adj.P.Val":0.2487},{"tissuedysplasia":0.3163,"tissuetumour":0.9921,"AveExpr":5.0441,"F":11.5264,"P.Value":0.0001,"adj.P.Val":0.0013},{"tissuedysplasia":-0.324,"tissuetumour":-1.5767,"AveExpr":6.5252,"F":7.7119,"P.Value":0.0016,"adj.P.Val":0.011},{"tissuedysplasia":0.6342,"tissuetumour":1.0984,"AveExpr":4.4261,"F":23.4483,"P.Value":1.1686e-07,"adj.P.Val":7.9231e-06},{"tissuedysplasia":0.1808,"tissuetumour":0.123,"AveExpr":3.9084,"F":0.2505,"P.Value":0.7795,"adj.P.Val":0.8441},{"tissuedysplasia":-0.2148,"tissuetumour":1.2396,"AveExpr":-3.0259,"F":2.36,"P.Value":0.1063,"adj.P.Val":0.2197},{"tissuedysplasia":1.1444,"tissuetumour":1.5023,"AveExpr":-1.8115,"F":2.4941,"P.Value":0.0942,"adj.P.Val":0.2024},{"tissuedysplasia":-0.0453,"tissuetumour":-0.3711,"AveExpr":6.92,"F":5.6469,"P.Value":0.0066,"adj.P.Val":0.0307},{"tissuedysplasia":-0.1187,"tissuetumour":-0.8033,"AveExpr":7.2691,"F":16.3516,"P.Value":4.8967e-06,"adj.P.Val":0.0001},{"tissuedysplasia":0.483,"tissuetumour":0.0629,"AveExpr":3.8557,"F":5.9664,"P.Value":0.0051,"adj.P.Val":0.0255},{"tissuedysplasia":0.641,"tissuetumour":0.2872,"AveExpr":4.3749,"F":5.3229,"P.Value":0.0085,"adj.P.Val":0.0369},{"tissuedysplasia":-0.1865,"tissuetumour":-0.4446,"AveExpr":4.7987,"F":5.5778,"P.Value":0.0069,"adj.P.Val":0.0319},{"tissuedysplasia":-0.0982,"tissuetumour":-0.5328,"AveExpr":4.6372,"F":8.7884,"P.Value":0.0006,"adj.P.Val":0.0054},{"tissuedysplasia":-0.0782,"tissuetumour":0.0076,"AveExpr":3.9164,"F":0.0917,"P.Value":0.9125,"adj.P.Val":0.9389},{"tissuedysplasia":0.0762,"tissuetumour":-0.7632,"AveExpr":7.2665,"F":11.5863,"P.Value":0.0001,"adj.P.Val":0.0015},{"tissuedysplasia":1.4888,"tissuetumour":3.3697,"AveExpr":6.6636,"F":17.0093,"P.Value":6.2269e-06,"adj.P.Val":0.0002},{"tissuedysplasia":0.9227,"tissuetumour":1.4739,"AveExpr":3.2262,"F":14.9594,"P.Value":0,"adj.P.Val":0.0003},{"tissuedysplasia":0.3816,"tissuetumour":0.4388,"AveExpr":6.3723,"F":7.86,"P.Value":0.0012,"adj.P.Val":0.0089},{"tissuedysplasia":0.1831,"tissuetumour":0.1084,"AveExpr":6.6288,"F":2.2691,"P.Value":0.1154,"adj.P.Val":0.2324},{"tissuedysplasia":1.0429,"tissuetumour":2.2809,"AveExpr":0.009,"F":10.7381,"P.Value":0.0002,"adj.P.Val":0.002},{"tissuedysplasia":-0.5509,"tissuetumour":0.8545,"AveExpr":0.6894,"F":5.5137,"P.Value":0.0073,"adj.P.Val":0.0331},{"tissuedysplasia":0.5217,"tissuetumour":1.0614,"AveExpr":4.0541,"F":10.1875,"P.Value":0.0002,"adj.P.Val":0.0026},{"tissuedysplasia":0.1852,"tissuetumour":0.1964,"AveExpr":5.0331,"F":1.8896,"P.Value":0.1632,"adj.P.Val":0.2939},{"tissuedysplasia":0.3414,"tissuetumour":0.3904,"AveExpr":7.6409,"F":6.2462,"P.Value":0.0041,"adj.P.Val":0.0218},{"tissuedysplasia":0.1734,"tissuetumour":-0.1068,"AveExpr":7.7602,"F":3.5925,"P.Value":0.0359,"adj.P.Val":0.1033},{"tissuedysplasia":0.0816,"tissuetumour":0.1615,"AveExpr":3.1125,"F":0.3555,"P.Value":0.7028,"adj.P.Val":0.7872},{"tissuedysplasia":0.1694,"tissuetumour":0.0246,"AveExpr":3.1162,"F":0.6391,"P.Value":0.5326,"adj.P.Val":0.6544},{"tissuedysplasia":0.4435,"tissuetumour":0.0348,"AveExpr":4.3067,"F":4.9632,"P.Value":0.0114,"adj.P.Val":0.0454},{"tissuedysplasia":0.3832,"tissuetumour":1.2256,"AveExpr":2.5084,"F":10.8488,"P.Value":0.0001,"adj.P.Val":0.0019},{"tissuedysplasia":0.5656,"tissuetumour":1.2416,"AveExpr":3.8596,"F":16.5713,"P.Value":4.3184e-06,"adj.P.Val":0.0001},{"tissuedysplasia":0.2782,"tissuetumour":0.6809,"AveExpr":3.519,"F":9.3412,"P.Value":0.0004,"adj.P.Val":0.0041},{"tissuedysplasia":0.1083,"tissuetumour":0.5376,"AveExpr":3.3841,"F":6.9076,"P.Value":0.0025,"adj.P.Val":0.0151},{"tissuedysplasia":0.9308,"tissuetumour":1.5231,"AveExpr":5.0837,"F":30.0179,"P.Value":5.9924e-09,"adj.P.Val":9.7095e-07},{"tissuedysplasia":0.1248,"tissuetumour":0.2715,"AveExpr":4.443,"F":1.7223,"P.Value":0.1905,"adj.P.Val":0.3267},{"tissuedysplasia":0.2577,"tissuetumour":0.1214,"AveExpr":5.2962,"F":3.1293,"P.Value":0.0536,"adj.P.Val":0.1372},{"tissuedysplasia":0.0586,"tissuetumour":0.4193,"AveExpr":3.8369,"F":4.1309,"P.Value":0.0227,"adj.P.Val":0.0749},{"tissuedysplasia":-0.3813,"tissuetumour":-0.9834,"AveExpr":7.7748,"F":9.4708,"P.Value":0.0005,"adj.P.Val":0.0045},{"tissuedysplasia":0.2135,"tissuetumour":-0.1981,"AveExpr":5.2073,"F":4.5567,"P.Value":0.0159,"adj.P.Val":0.058},{"tissuedysplasia":0.0842,"tissuetumour":0.2235,"AveExpr":3.0022,"F":1.1202,"P.Value":0.3353,"adj.P.Val":0.4796},{"tissuedysplasia":0.2302,"tissuetumour":0.2803,"AveExpr":5.1206,"F":0.908,"P.Value":0.4108,"adj.P.Val":0.5498},{"tissuedysplasia":0.0786,"tissuetumour":0.8774,"AveExpr":0.6399,"F":4.9786,"P.Value":0.0112,"adj.P.Val":0.045},{"tissuedysplasia":0.1964,"tissuetumour":0.4938,"AveExpr":3.9171,"F":5.0489,"P.Value":0.0106,"adj.P.Val":0.0433},{"tissuedysplasia":-0.3252,"tissuetumour":-0.373,"AveExpr":7.5509,"F":10.004,"P.Value":0.0003,"adj.P.Val":0.0029},{"tissuedysplasia":0.5463,"tissuetumour":0.2376,"AveExpr":2.171,"F":2.8559,"P.Value":0.0682,"adj.P.Val":0.1619},{"tissuedysplasia":-0.0277,"tissuetumour":-0.394,"AveExpr":8.7392,"F":11.3158,"P.Value":0.0001,"adj.P.Val":0.0015},{"tissuedysplasia":-0.4024,"tissuetumour":-0.3738,"AveExpr":-0.3907,"F":1.2847,"P.Value":0.2869,"adj.P.Val":0.4314},{"tissuedysplasia":0.0732,"tissuetumour":0.0213,"AveExpr":6.1676,"F":0.2473,"P.Value":0.782,"adj.P.Val":0.8458},{"tissuedysplasia":0.3719,"tissuetumour":1.5647,"AveExpr":4.2494,"F":34.8022,"P.Value":8.6471e-10,"adj.P.Val":2.263e-07},{"tissuedysplasia":-0.0923,"tissuetumour":0.025,"AveExpr":4.5504,"F":0.3898,"P.Value":0.6795,"adj.P.Val":0.7701},{"tissuedysplasia":0.975,"tissuetumour":3.1438,"AveExpr":-2.7736,"F":10.4102,"P.Value":0.0002,"adj.P.Val":0.0023},{"tissuedysplasia":0.3356,"tissuetumour":0.3397,"AveExpr":1.851,"F":1.0057,"P.Value":0.374,"adj.P.Val":0.5165},{"tissuedysplasia":0.3582,"tissuetumour":0.9142,"AveExpr":2.5485,"F":6.7977,"P.Value":0.0027,"adj.P.Val":0.016},{"tissuedysplasia":0.3249,"tissuetumour":0.5054,"AveExpr":4.2262,"F":7.947,"P.Value":0.0011,"adj.P.Val":0.0085},{"tissuedysplasia":0.5463,"tissuetumour":0.7783,"AveExpr":2.1481,"F":2.7258,"P.Value":0.0766,"adj.P.Val":0.1753},{"tissuedysplasia":0.3781,"tissuetumour":4.4926,"AveExpr":0.8912,"F":37.4046,"P.Value":3.2272e-10,"adj.P.Val":1.1562e-07},{"tissuedysplasia":0.1029,"tissuetumour":-0.6511,"AveExpr":6.3529,"F":19.5726,"P.Value":8.3051e-07,"adj.P.Val":0},{"tissuedysplasia":-0.1969,"tissuetumour":-1.098,"AveExpr":2.9358,"F":13.7278,"P.Value":0,"adj.P.Val":0.0005},{"tissuedysplasia":-0.2669,"tissuetumour":-0.2613,"AveExpr":4.4676,"F":3.0916,"P.Value":0.0554,"adj.P.Val":0.1403},{"tissuedysplasia":0.9489,"tissuetumour":1.7385,"AveExpr":3.6483,"F":21.3399,"P.Value":3.3232e-07,"adj.P.Val":0},{"tissuedysplasia":0.1978,"tissuetumour":0.8455,"AveExpr":5.5349,"F":22.2437,"P.Value":2.1103e-07,"adj.P.Val":0},{"tissuedysplasia":-0.0755,"tissuetumour":0.2295,"AveExpr":1.8524,"F":0.7106,"P.Value":0.4969,"adj.P.Val":0.6241},{"tissuedysplasia":-0.1895,"tissuetumour":0.4974,"AveExpr":1.7388,"F":2.7459,"P.Value":0.0752,"adj.P.Val":0.1731},{"tissuedysplasia":0.8905,"tissuetumour":2.3234,"AveExpr":-5.3059,"F":3.7239,"P.Value":0.0321,"adj.P.Val":0.0955},{"tissuedysplasia":0.682,"tissuetumour":1.3555,"AveExpr":0.7735,"F":6.7244,"P.Value":0.0028,"adj.P.Val":0.0167},{"tissuedysplasia":-0.1638,"tissuetumour":-0.614,"AveExpr":7.3111,"F":12.3033,"P.Value":0.0001,"adj.P.Val":0.0009},{"tissuedysplasia":0.2501,"tissuetumour":0.0067,"AveExpr":4.1036,"F":1.7034,"P.Value":0.1939,"adj.P.Val":0.3303},{"tissuedysplasia":0.8232,"tissuetumour":1.2603,"AveExpr":1.8882,"F":11.1153,"P.Value":0.0001,"adj.P.Val":0.0016},{"tissuedysplasia":0.183,"tissuetumour":0.2005,"AveExpr":6.013,"F":4.4609,"P.Value":0.0172,"adj.P.Val":0.0615},{"tissuedysplasia":0.445,"tissuetumour":1.5359,"AveExpr":5.4557,"F":35.4845,"P.Value":6.6493e-10,"adj.P.Val":1.9199e-07},{"tissuedysplasia":0.0492,"tissuetumour":-0.4173,"AveExpr":6.3659,"F":11.3674,"P.Value":0.0001,"adj.P.Val":0.0014},{"tissuedysplasia":1.0966,"tissuetumour":2.4038,"AveExpr":6.4947,"F":29.4636,"P.Value":2.0593e-08,"adj.P.Val":2.3118e-06},{"tissuedysplasia":-0.1604,"tissuetumour":-0.8663,"AveExpr":-2.4413,"F":0.9068,"P.Value":0.4112,"adj.P.Val":0.5502},{"tissuedysplasia":0.017,"tissuetumour":-0.4746,"AveExpr":5.9272,"F":6.6323,"P.Value":0.003,"adj.P.Val":0.0175},{"tissuedysplasia":-0.4888,"tissuetumour":-0.9175,"AveExpr":3.3937,"F":7.106,"P.Value":0.0021,"adj.P.Val":0.0135},{"tissuedysplasia":-0.1082,"tissuetumour":0.0239,"AveExpr":1.7585,"F":0.1864,"P.Value":0.8306,"adj.P.Val":0.881},{"tissuedysplasia":-0.0789,"tissuetumour":-0.2522,"AveExpr":6.2748,"F":4.1592,"P.Value":0.0222,"adj.P.Val":0.0736},{"tissuedysplasia":0.5969,"tissuetumour":1.3769,"AveExpr":-1.2437,"F":2.9225,"P.Value":0.0643,"adj.P.Val":0.1555},{"tissuedysplasia":0.3288,"tissuetumour":0.6837,"AveExpr":2.2384,"F":6.2258,"P.Value":0.0042,"adj.P.Val":0.022},{"tissuedysplasia":-0.1346,"tissuetumour":-0.4697,"AveExpr":4.6061,"F":9.542,"P.Value":0.0004,"adj.P.Val":0.0037},{"tissuedysplasia":0.1145,"tissuetumour":-0.2449,"AveExpr":6.1623,"F":4.2019,"P.Value":0.0214,"adj.P.Val":0.0719},{"tissuedysplasia":0.2781,"tissuetumour":0.4499,"AveExpr":8.888,"F":3.1882,"P.Value":0.053,"adj.P.Val":0.136},{"tissuedysplasia":-0.0047,"tissuetumour":-0.3016,"AveExpr":4.9078,"F":5.7612,"P.Value":0.006,"adj.P.Val":0.0287},{"tissuedysplasia":0.246,"tissuetumour":1.4159,"AveExpr":2.6916,"F":24.2373,"P.Value":8.0022e-08,"adj.P.Val":6.07e-06},{"tissuedysplasia":0.9468,"tissuetumour":1.4117,"AveExpr":0.5062,"F":6.1413,"P.Value":0.0044,"adj.P.Val":0.0231},{"tissuedysplasia":-0.1021,"tissuetumour":-0.2366,"AveExpr":7.9294,"F":4.3794,"P.Value":0.0184,"adj.P.Val":0.0646},{"tissuedysplasia":0.5112,"tissuetumour":0.8391,"AveExpr":0.4465,"F":2.0279,"P.Value":0.1437,"adj.P.Val":0.2695},{"tissuedysplasia":0.0088,"tissuetumour":0.809,"AveExpr":3.1408,"F":9.7519,"P.Value":0.0003,"adj.P.Val":0.0033},{"tissuedysplasia":0.2671,"tissuetumour":0.7799,"AveExpr":2.5862,"F":4.1483,"P.Value":0.0224,"adj.P.Val":0.0741},{"tissuedysplasia":-0.2507,"tissuetumour":-0.1293,"AveExpr":5.8742,"F":2.5847,"P.Value":0.0868,"adj.P.Val":0.1912},{"tissuedysplasia":-0.0401,"tissuetumour":0.578,"AveExpr":1.5253,"F":2.8635,"P.Value":0.0678,"adj.P.Val":0.1612},{"tissuedysplasia":-0.0819,"tissuetumour":0.7685,"AveExpr":2.7777,"F":13.4302,"P.Value":0,"adj.P.Val":0.0005},{"tissuedysplasia":0.2093,"tissuetumour":0.9004,"AveExpr":5.8712,"F":24.2719,"P.Value":7.8716e-08,"adj.P.Val":5.9954e-06},{"tissuedysplasia":-0.079,"tissuetumour":0.6189,"AveExpr":5.6907,"F":14.4201,"P.Value":0,"adj.P.Val":0.0003},{"tissuedysplasia":0.1834,"tissuetumour":0.3166,"AveExpr":4.2797,"F":3.0454,"P.Value":0.0577,"adj.P.Val":0.1444},{"tissuedysplasia":0.4016,"tissuetumour":0.2981,"AveExpr":2.2751,"F":2.1571,"P.Value":0.1277,"adj.P.Val":0.2487},{"tissuedysplasia":-0.2563,"tissuetumour":0.9534,"AveExpr":-10.2043,"F":0.8439,"P.Value":0.4369,"adj.P.Val":0.5736},{"tissuedysplasia":0.4627,"tissuetumour":0.5709,"AveExpr":3.3209,"F":4.4866,"P.Value":0.0169,"adj.P.Val":0.0606},{"tissuedysplasia":0.0836,"tissuetumour":-0.3141,"AveExpr":4.987,"F":7.8503,"P.Value":0.0012,"adj.P.Val":0.0089},{"tissuedysplasia":0.0456,"tissuetumour":0.7409,"AveExpr":5.581,"F":8.9825,"P.Value":0.0005,"adj.P.Val":0.0049},{"tissuedysplasia":0.2612,"tissuetumour":0.516,"AveExpr":1.7044,"F":1.3901,"P.Value":0.2598,"adj.P.Val":0.4029},{"tissuedysplasia":0.2398,"tissuetumour":0.6074,"AveExpr":3.1667,"F":6.1391,"P.Value":0.0045,"adj.P.Val":0.0231},{"tissuedysplasia":0.2302,"tissuetumour":0.4547,"AveExpr":4.8822,"F":10.8525,"P.Value":0.0001,"adj.P.Val":0.0019},{"tissuedysplasia":0.6913,"tissuetumour":0.3535,"AveExpr":5.0019,"F":12.8255,"P.Value":0,"adj.P.Val":0.0007},{"tissuedysplasia":-0.4816,"tissuetumour":-0.0701,"AveExpr":4.8414,"F":2.6363,"P.Value":0.0829,"adj.P.Val":0.1852},{"tissuedysplasia":0.2467,"tissuetumour":0.2694,"AveExpr":4.1298,"F":1.5299,"P.Value":0.2279,"adj.P.Val":0.3688},{"tissuedysplasia":-0.1129,"tissuetumour":-0.6518,"AveExpr":5.0261,"F":10.9361,"P.Value":0.0001,"adj.P.Val":0.0018},{"tissuedysplasia":-0.0095,"tissuetumour":0.0303,"AveExpr":3.0744,"F":0.0092,"P.Value":0.9908,"adj.P.Val":0.9934},{"tissuedysplasia":0.0307,"tissuetumour":0.3995,"AveExpr":1.8493,"F":1.6755,"P.Value":0.1989,"adj.P.Val":0.3359},{"tissuedysplasia":0.902,"tissuetumour":1.578,"AveExpr":0.0967,"F":5.9341,"P.Value":0.0052,"adj.P.Val":0.026},{"tissuedysplasia":0.3234,"tissuetumour":1.0744,"AveExpr":0.5674,"F":3.1796,"P.Value":0.0513,"adj.P.Val":0.133},{"tissuedysplasia":0.827,"tissuetumour":1.8842,"AveExpr":-3.712,"F":1.6696,"P.Value":0.2,"adj.P.Val":0.337},{"tissuedysplasia":0.5148,"tissuetumour":0.6925,"AveExpr":3.0789,"F":6.6417,"P.Value":0.003,"adj.P.Val":0.0174},{"tissuedysplasia":0.7618,"tissuetumour":0.8851,"AveExpr":3.8208,"F":9.1178,"P.Value":0.0005,"adj.P.Val":0.0046},{"tissuedysplasia":0.3176,"tissuetumour":0.7533,"AveExpr":2.7425,"F":4.2547,"P.Value":0.0205,"adj.P.Val":0.0696},{"tissuedysplasia":0.0298,"tissuetumour":-0.3766,"AveExpr":3.082,"F":1.4413,"P.Value":0.2476,"adj.P.Val":0.3903},{"tissuedysplasia":0.3627,"tissuetumour":0.2831,"AveExpr":5.4019,"F":4.1614,"P.Value":0.0221,"adj.P.Val":0.0736},{"tissuedysplasia":-0.3735,"tissuetumour":-0.3242,"AveExpr":7.37,"F":13.0272,"P.Value":0,"adj.P.Val":0.0006},{"tissuedysplasia":0.0913,"tissuetumour":0.2535,"AveExpr":5.631,"F":4.4176,"P.Value":0.0178,"adj.P.Val":0.0632},{"tissuedysplasia":0.2504,"tissuetumour":0.8886,"AveExpr":1.0904,"F":5.2342,"P.Value":0.0091,"adj.P.Val":0.0388},{"tissuedysplasia":0.1002,"tissuetumour":0.7493,"AveExpr":2.5193,"F":9.5609,"P.Value":0.0004,"adj.P.Val":0.0036},{"tissuedysplasia":0.5088,"tissuetumour":1.3867,"AveExpr":1.7818,"F":16.723,"P.Value":3.9611e-06,"adj.P.Val":0.0001},{"tissuedysplasia":0.7518,"tissuetumour":1.563,"AveExpr":1.9913,"F":15.2524,"P.Value":9.2845e-06,"adj.P.Val":0.0002},{"tissuedysplasia":0.3893,"tissuetumour":0.5462,"AveExpr":5.4609,"F":11.0009,"P.Value":0.0001,"adj.P.Val":0.0017},{"tissuedysplasia":0.0712,"tissuetumour":0.3038,"AveExpr":5.995,"F":8.7405,"P.Value":0.0006,"adj.P.Val":0.0056},{"tissuedysplasia":0.2774,"tissuetumour":0.4153,"AveExpr":5.0416,"F":4.3075,"P.Value":0.0196,"adj.P.Val":0.0676},{"tissuedysplasia":-0.3929,"tissuetumour":-0.7184,"AveExpr":3.7221,"F":8.3677,"P.Value":0.0008,"adj.P.Val":0.0068},{"tissuedysplasia":-1.1848,"tissuetumour":-2.0851,"AveExpr":3.1268,"F":21.2268,"P.Value":3.5197e-07,"adj.P.Val":0},{"tissuedysplasia":0.215,"tissuetumour":0.6256,"AveExpr":6.5918,"F":7.19,"P.Value":0.002,"adj.P.Val":0.0129},{"tissuedysplasia":-0.3206,"tissuetumour":0.2169,"AveExpr":3.9311,"F":3.2124,"P.Value":0.0499,"adj.P.Val":0.1305},{"tissuedysplasia":0.2702,"tissuetumour":0.202,"AveExpr":2.869,"F":0.7302,"P.Value":0.4876,"adj.P.Val":0.6162},{"tissuedysplasia":0.4146,"tissuetumour":0.3012,"AveExpr":3.4736,"F":3.9701,"P.Value":0.026,"adj.P.Val":0.0824},{"tissuedysplasia":0.1757,"tissuetumour":0.0106,"AveExpr":4.8039,"F":1.1405,"P.Value":0.3289,"adj.P.Val":0.4732},{"tissuedysplasia":0.1925,"tissuetumour":-0.003,"AveExpr":3.5093,"F":0.9451,"P.Value":0.3964,"adj.P.Val":0.537},{"tissuedysplasia":0.0154,"tissuetumour":-0.3772,"AveExpr":2.6994,"F":1.0352,"P.Value":0.3636,"adj.P.Val":0.5067},{"tissuedysplasia":-0.3159,"tissuetumour":0.2314,"AveExpr":-5.5827,"F":0.1624,"P.Value":0.8506,"adj.P.Val":0.8949},{"tissuedysplasia":0.6062,"tissuetumour":0.4829,"AveExpr":2.0226,"F":1.7315,"P.Value":0.1889,"adj.P.Val":0.3248},{"tissuedysplasia":0.2002,"tissuetumour":0.8202,"AveExpr":5.1164,"F":13.1016,"P.Value":0,"adj.P.Val":0.0006},{"tissuedysplasia":0.1105,"tissuetumour":-0.137,"AveExpr":3.5985,"F":1.347,"P.Value":0.2705,"adj.P.Val":0.4144},{"tissuedysplasia":0.1684,"tissuetumour":0.4868,"AveExpr":4.8566,"F":4.153,"P.Value":0.0223,"adj.P.Val":0.0739},{"tissuedysplasia":0.0672,"tissuetumour":-0.1143,"AveExpr":3.8115,"F":0.4282,"P.Value":0.6544,"adj.P.Val":0.7509},{"tissuedysplasia":0.1103,"tissuetumour":0.2872,"AveExpr":3.9127,"F":2.8379,"P.Value":0.0693,"adj.P.Val":0.1638},{"tissuedysplasia":0.335,"tissuetumour":0.4478,"AveExpr":5.2742,"F":10.6616,"P.Value":0.0002,"adj.P.Val":0.0021},{"tissuedysplasia":0.1414,"tissuetumour":0.6175,"AveExpr":6.3197,"F":23.4347,"P.Value":1.1763e-07,"adj.P.Val":7.9475e-06},{"tissuedysplasia":0.2729,"tissuetumour":0.5196,"AveExpr":3.8813,"F":5.9757,"P.Value":0.0051,"adj.P.Val":0.0253},{"tissuedysplasia":0.2943,"tissuetumour":1.4463,"AveExpr":2.9923,"F":19.4055,"P.Value":9.0747e-07,"adj.P.Val":0},{"tissuedysplasia":0.3575,"tissuetumour":0.89,"AveExpr":-0.121,"F":2.9037,"P.Value":0.0654,"adj.P.Val":0.1572},{"tissuedysplasia":0.4087,"tissuetumour":0.3605,"AveExpr":4.9452,"F":8.3318,"P.Value":0.0009,"adj.P.Val":0.0069},{"tissuedysplasia":-0.2792,"tissuetumour":-0.7825,"AveExpr":6.715,"F":14.3278,"P.Value":0,"adj.P.Val":0.0003},{"tissuedysplasia":0.0244,"tissuetumour":-0.1385,"AveExpr":5.8811,"F":0.9279,"P.Value":0.403,"adj.P.Val":0.5432},{"tissuedysplasia":0.0112,"tissuetumour":-0.5175,"AveExpr":4.8498,"F":5.0848,"P.Value":0.0103,"adj.P.Val":0.0424},{"tissuedysplasia":0.1296,"tissuetumour":-0.0267,"AveExpr":4.8385,"F":0.8045,"P.Value":0.4538,"adj.P.Val":0.5879},{"tissuedysplasia":-0.1411,"tissuetumour":-0.6424,"AveExpr":5.923,"F":11.9875,"P.Value":0.0001,"adj.P.Val":0.0011},{"tissuedysplasia":0.2742,"tissuetumour":0.8472,"AveExpr":1.5603,"F":7.0251,"P.Value":0.0023,"adj.P.Val":0.0141},{"tissuedysplasia":0.7442,"tissuetumour":1.1771,"AveExpr":4.908,"F":23.1874,"P.Value":1.3264e-07,"adj.P.Val":8.6981e-06},{"tissuedysplasia":0.178,"tissuetumour":0.4052,"AveExpr":3.7462,"F":1.8834,"P.Value":0.1641,"adj.P.Val":0.295},{"tissuedysplasia":0.5281,"tissuetumour":2.4297,"AveExpr":-1.6474,"F":8.1546,"P.Value":0.001,"adj.P.Val":0.0076},{"tissuedysplasia":0.6542,"tissuetumour":1.2162,"AveExpr":2.6169,"F":15.4983,"P.Value":8.0332e-06,"adj.P.Val":0.0002},{"tissuedysplasia":0.0365,"tissuetumour":0.2917,"AveExpr":7.3049,"F":4.3484,"P.Value":0.0189,"adj.P.Val":0.0659},{"tissuedysplasia":-0.4137,"tissuetumour":-0.6745,"AveExpr":6.2261,"F":13.7038,"P.Value":0,"adj.P.Val":0.0005},{"tissuedysplasia":0.6625,"tissuetumour":2.8359,"AveExpr":-1.5676,"F":12.1471,"P.Value":0.0001,"adj.P.Val":0.001},{"tissuedysplasia":0.7143,"tissuetumour":0.2362,"AveExpr":4.6327,"F":6.7405,"P.Value":0.0028,"adj.P.Val":0.0165},{"tissuedysplasia":0.3527,"tissuetumour":0.5768,"AveExpr":5.5302,"F":9.3689,"P.Value":0.0004,"adj.P.Val":0.004},{"tissuedysplasia":0.423,"tissuetumour":0.7135,"AveExpr":3.8854,"F":12.1257,"P.Value":0.0001,"adj.P.Val":0.001},{"tissuedysplasia":0.7351,"tissuetumour":1.3565,"AveExpr":3.5487,"F":34.2699,"P.Value":1.0637e-09,"adj.P.Val":2.6159e-07},{"tissuedysplasia":-0.0227,"tissuetumour":-0.2111,"AveExpr":3.6986,"F":1.588,"P.Value":0.2158,"adj.P.Val":0.3553},{"tissuedysplasia":0.3795,"tissuetumour":0.1,"AveExpr":3.2484,"F":3.746,"P.Value":0.0315,"adj.P.Val":0.0942},{"tissuedysplasia":0.083,"tissuetumour":0.1373,"AveExpr":7.5639,"F":1.686,"P.Value":0.197,"adj.P.Val":0.3338},{"tissuedysplasia":0.5007,"tissuetumour":0.8531,"AveExpr":5.7289,"F":13.6423,"P.Value":0,"adj.P.Val":0.0005},{"tissuedysplasia":0.0722,"tissuetumour":0.2305,"AveExpr":5.7153,"F":2.1273,"P.Value":0.1312,"adj.P.Val":0.2536},{"tissuedysplasia":-0.1737,"tissuetumour":-0.761,"AveExpr":5.0443,"F":16.9145,"P.Value":3.5537e-06,"adj.P.Val":0.0001},{"tissuedysplasia":0.1798,"tissuetumour":0.4827,"AveExpr":2.6196,"F":2.8827,"P.Value":0.0666,"adj.P.Val":0.1592},{"tissuedysplasia":0.5512,"tissuetumour":1.8595,"AveExpr":-1.212,"F":6.0177,"P.Value":0.0049,"adj.P.Val":0.0247},{"tissuedysplasia":-0.4067,"tissuetumour":-1.0171,"AveExpr":7.8586,"F":20.8943,"P.Value":6.2732e-07,"adj.P.Val":0},{"tissuedysplasia":0.448,"tissuetumour":0.3849,"AveExpr":4.129,"F":6.7902,"P.Value":0.0027,"adj.P.Val":0.0161},{"tissuedysplasia":0.3121,"tissuetumour":1.2203,"AveExpr":8.6642,"F":9.0729,"P.Value":0.0006,"adj.P.Val":0.0056},{"tissuedysplasia":-0.4479,"tissuetumour":-1.1705,"AveExpr":3.9302,"F":15.3937,"P.Value":8.5426e-06,"adj.P.Val":0.0002},{"tissuedysplasia":0.2445,"tissuetumour":0.1108,"AveExpr":3.145,"F":1.4294,"P.Value":0.2503,"adj.P.Val":0.3932},{"tissuedysplasia":1.0049,"tissuetumour":1.1826,"AveExpr":2.4523,"F":11.4089,"P.Value":0.0001,"adj.P.Val":0.0014},{"tissuedysplasia":-0.7334,"tissuetumour":-0.2486,"AveExpr":1.3549,"F":3.1865,"P.Value":0.051,"adj.P.Val":0.1324},{"tissuedysplasia":0.2123,"tissuetumour":0.8014,"AveExpr":3.9481,"F":18.0298,"P.Value":1.9084e-06,"adj.P.Val":0.0001},{"tissuedysplasia":0.3975,"tissuetumour":-0.0697,"AveExpr":4.7893,"F":4.3571,"P.Value":0.0188,"adj.P.Val":0.0655},{"tissuedysplasia":-0.249,"tissuetumour":-0.8588,"AveExpr":5.0696,"F":6.9242,"P.Value":0.0024,"adj.P.Val":0.0149},{"tissuedysplasia":0.3023,"tissuetumour":0.7613,"AveExpr":4.4855,"F":9.981,"P.Value":0.0003,"adj.P.Val":0.0029},{"tissuedysplasia":0.0828,"tissuetumour":-0.1927,"AveExpr":6.2583,"F":5.5493,"P.Value":0.0071,"adj.P.Val":0.0325},{"tissuedysplasia":0.4945,"tissuetumour":0.4515,"AveExpr":4.6029,"F":5.9248,"P.Value":0.0053,"adj.P.Val":0.0261},{"tissuedysplasia":0.6261,"tissuetumour":0.1522,"AveExpr":4.5924,"F":7.2012,"P.Value":0.002,"adj.P.Val":0.0128},{"tissuedysplasia":0.2943,"tissuetumour":0.4028,"AveExpr":4.4351,"F":6.1252,"P.Value":0.0045,"adj.P.Val":0.0233},{"tissuedysplasia":0.2391,"tissuetumour":0.6763,"AveExpr":7.6099,"F":7.7174,"P.Value":0.0015,"adj.P.Val":0.0105},{"tissuedysplasia":0.412,"tissuetumour":1.236,"AveExpr":3.0989,"F":12.8474,"P.Value":0,"adj.P.Val":0.0007},{"tissuedysplasia":-0.2027,"tissuetumour":-0.9122,"AveExpr":5.1098,"F":16.793,"P.Value":3.8069e-06,"adj.P.Val":0.0001},{"tissuedysplasia":0.2305,"tissuetumour":1.2466,"AveExpr":-0.2695,"F":2.9331,"P.Value":0.0637,"adj.P.Val":0.1546},{"tissuedysplasia":0.1207,"tissuetumour":-0.0931,"AveExpr":4.6438,"F":1.5274,"P.Value":0.2284,"adj.P.Val":0.3694},{"tissuedysplasia":0.1627,"tissuetumour":1.1995,"AveExpr":2.9684,"F":16.9784,"P.Value":3.4277e-06,"adj.P.Val":0.0001},{"tissuedysplasia":0.0581,"tissuetumour":-0.1073,"AveExpr":7.0003,"F":1.461,"P.Value":0.243,"adj.P.Val":0.3854},{"tissuedysplasia":0.3228,"tissuetumour":0.5207,"AveExpr":4.617,"F":9.4077,"P.Value":0.0004,"adj.P.Val":0.0039},{"tissuedysplasia":0.2634,"tissuetumour":0.4784,"AveExpr":5.2971,"F":6.2566,"P.Value":0.0041,"adj.P.Val":0.0216},{"tissuedysplasia":0.1387,"tissuetumour":0.0771,"AveExpr":4.7782,"F":0.7866,"P.Value":0.4617,"adj.P.Val":0.5947},{"tissuedysplasia":0.471,"tissuetumour":1.4943,"AveExpr":-2.7435,"F":2.2127,"P.Value":0.1214,"adj.P.Val":0.2403},{"tissuedysplasia":-1.0339,"tissuetumour":-0.6435,"AveExpr":2.04,"F":3.9185,"P.Value":0.0272,"adj.P.Val":0.0851},{"tissuedysplasia":-0.8455,"tissuetumour":-0.6711,"AveExpr":1.7888,"F":5.1001,"P.Value":0.0102,"adj.P.Val":0.042},{"tissuedysplasia":0.4075,"tissuetumour":-0.0529,"AveExpr":5.2023,"F":2.8584,"P.Value":0.0681,"adj.P.Val":0.1617},{"tissuedysplasia":0.954,"tissuetumour":0.5107,"AveExpr":4.7037,"F":7.8444,"P.Value":0.0012,"adj.P.Val":0.009},{"tissuedysplasia":-0.1151,"tissuetumour":0.1159,"AveExpr":5.1039,"F":1.5876,"P.Value":0.2159,"adj.P.Val":0.3554},{"tissuedysplasia":0.7509,"tissuetumour":1.1713,"AveExpr":2.8273,"F":11.9301,"P.Value":0.0001,"adj.P.Val":0.0011},{"tissuedysplasia":0.2249,"tissuetumour":0.8474,"AveExpr":1.7424,"F":4.3166,"P.Value":0.0194,"adj.P.Val":0.0672},{"tissuedysplasia":0.6832,"tissuetumour":1.1683,"AveExpr":0.8237,"F":4.364,"P.Value":0.0187,"adj.P.Val":0.0652},{"tissuedysplasia":0.1269,"tissuetumour":0.9561,"AveExpr":1.8358,"F":7.9431,"P.Value":0.0011,"adj.P.Val":0.0085},{"tissuedysplasia":0.273,"tissuetumour":-0.1375,"AveExpr":5.1189,"F":5.488,"P.Value":0.0074,"adj.P.Val":0.0336},{"tissuedysplasia":0.0059,"tissuetumour":0.1021,"AveExpr":5.9846,"F":0.6362,"P.Value":0.5341,"adj.P.Val":0.6557},{"tissuedysplasia":0.0608,"tissuetumour":-0.1,"AveExpr":4.513,"F":0.5319,"P.Value":0.5912,"adj.P.Val":0.7015},{"tissuedysplasia":-0.3846,"tissuetumour":-0.6315,"AveExpr":4.98,"F":7.1732,"P.Value":0.002,"adj.P.Val":0.013},{"tissuedysplasia":0.2336,"tissuetumour":-0.0582,"AveExpr":3.3101,"F":2.6224,"P.Value":0.084,"adj.P.Val":0.1868},{"tissuedysplasia":-0.4056,"tissuetumour":-1.0854,"AveExpr":2.894,"F":16.8086,"P.Value":3.7734e-06,"adj.P.Val":0.0001},{"tissuedysplasia":-0.0225,"tissuetumour":0.4979,"AveExpr":3.8516,"F":6.1434,"P.Value":0.0044,"adj.P.Val":0.023},{"tissuedysplasia":0.3034,"tissuetumour":0.0594,"AveExpr":3.226,"F":2.5869,"P.Value":0.0867,"adj.P.Val":0.1909},{"tissuedysplasia":0.1627,"tissuetumour":-0.0936,"AveExpr":5.2371,"F":3.031,"P.Value":0.0584,"adj.P.Val":0.1457},{"tissuedysplasia":0.3405,"tissuetumour":1.0973,"AveExpr":3.468,"F":21.1751,"P.Value":3.6137e-07,"adj.P.Val":0},{"tissuedysplasia":0.2929,"tissuetumour":1.0554,"AveExpr":0.2232,"F":4.4731,"P.Value":0.017,"adj.P.Val":0.0611},{"tissuedysplasia":0.0654,"tissuetumour":-0.2192,"AveExpr":5.8744,"F":5.0501,"P.Value":0.0106,"adj.P.Val":0.0433},{"tissuedysplasia":0.2075,"tissuetumour":-0.1908,"AveExpr":6.7675,"F":6.5661,"P.Value":0.0032,"adj.P.Val":0.0182},{"tissuedysplasia":0.6742,"tissuetumour":2.3419,"AveExpr":-0.8937,"F":10.8501,"P.Value":0.0001,"adj.P.Val":0.0019},{"tissuedysplasia":0.415,"tissuetumour":0.4922,"AveExpr":3.8993,"F":6.1974,"P.Value":0.0043,"adj.P.Val":0.0224},{"tissuedysplasia":0.5565,"tissuetumour":0.6977,"AveExpr":4.5536,"F":10.9761,"P.Value":0.0001,"adj.P.Val":0.0018},{"tissuedysplasia":0.3571,"tissuetumour":0.6245,"AveExpr":3.88,"F":5.754,"P.Value":0.006,"adj.P.Val":0.0288},{"tissuedysplasia":0.1625,"tissuetumour":-0.1032,"AveExpr":3.0421,"F":1.2825,"P.Value":0.2875,"adj.P.Val":0.432},{"tissuedysplasia":-0.0413,"tissuetumour":-0.3039,"AveExpr":6.4075,"F":6.8224,"P.Value":0.0026,"adj.P.Val":0.0158},{"tissuedysplasia":-0.1254,"tissuetumour":-0.7623,"AveExpr":3.5475,"F":6.697,"P.Value":0.0029,"adj.P.Val":0.0169},{"tissuedysplasia":-0.2786,"tissuetumour":0.2617,"AveExpr":5.1188,"F":5.3704,"P.Value":0.0082,"adj.P.Val":0.0359},{"tissuedysplasia":-0.0054,"tissuetumour":-0.7407,"AveExpr":4.617,"F":9.7609,"P.Value":0.0003,"adj.P.Val":0.0033},{"tissuedysplasia":0.6513,"tissuetumour":0.7808,"AveExpr":6.4816,"F":15.4305,"P.Value":8.3597e-06,"adj.P.Val":0.0002},{"tissuedysplasia":-0.2442,"tissuetumour":-0.9675,"AveExpr":8.4459,"F":33.038,"P.Value":3.2757e-09,"adj.P.Val":6.0739e-07},{"tissuedysplasia":-0.0531,"tissuetumour":-0.7218,"AveExpr":3.5096,"F":11.8417,"P.Value":0.0001,"adj.P.Val":0.0011},{"tissuedysplasia":-0.1259,"tissuetumour":-0.5925,"AveExpr":4.7308,"F":9.1171,"P.Value":0.0005,"adj.P.Val":0.0046},{"tissuedysplasia":0.2028,"tissuetumour":-0.0866,"AveExpr":5.9395,"F":1.9587,"P.Value":0.1531,"adj.P.Val":0.2816},{"tissuedysplasia":-0.0787,"tissuetumour":-0.1805,"AveExpr":3.7495,"F":0.3972,"P.Value":0.6746,"adj.P.Val":0.7665},{"tissuedysplasia":0.1724,"tissuetumour":0.639,"AveExpr":5.5474,"F":16.0288,"P.Value":5.8976e-06,"adj.P.Val":0.0002},{"tissuedysplasia":0.1296,"tissuetumour":1.6078,"AveExpr":2.2356,"F":22.8559,"P.Value":1.5596e-07,"adj.P.Val":9.7329e-06},{"tissuedysplasia":0.0512,"tissuetumour":-0.0979,"AveExpr":5.1327,"F":0.9213,"P.Value":0.4055,"adj.P.Val":0.5452},{"tissuedysplasia":0.3988,"tissuetumour":1.2025,"AveExpr":2.4041,"F":13.7096,"P.Value":0,"adj.P.Val":0.0005},{"tissuedysplasia":0.3532,"tissuetumour":1.0986,"AveExpr":8.6392,"F":24.8982,"P.Value":1.3942e-07,"adj.P.Val":9.0104e-06},{"tissuedysplasia":-0.0669,"tissuetumour":-0.3711,"AveExpr":5.0103,"F":4.0656,"P.Value":0.024,"adj.P.Val":0.0778},{"tissuedysplasia":0.1302,"tissuetumour":-0.2018,"AveExpr":5.3039,"F":2.7213,"P.Value":0.0769,"adj.P.Val":0.1758},{"tissuedysplasia":0.2332,"tissuetumour":0.7959,"AveExpr":3.5038,"F":11.5829,"P.Value":0.0001,"adj.P.Val":0.0013},{"tissuedysplasia":0.1493,"tissuetumour":0.0707,"AveExpr":5.8996,"F":1.3304,"P.Value":0.2748,"adj.P.Val":0.4192},{"tissuedysplasia":-0.018,"tissuetumour":0.5741,"AveExpr":1.3786,"F":2.3815,"P.Value":0.1042,"adj.P.Val":0.2169},{"tissuedysplasia":0.0199,"tissuetumour":0.0396,"AveExpr":4.2597,"F":0.0409,"P.Value":0.96,"adj.P.Val":0.9726},{"tissuedysplasia":-0.2636,"tissuetumour":-0.4962,"AveExpr":7.2386,"F":11.5915,"P.Value":0.0001,"adj.P.Val":0.0013},{"tissuedysplasia":0.2761,"tissuetumour":0.3596,"AveExpr":5.4084,"F":8.9449,"P.Value":0.0005,"adj.P.Val":0.005},{"tissuedysplasia":0.1747,"tissuetumour":0.4379,"AveExpr":1.1356,"F":2.245,"P.Value":0.1179,"adj.P.Val":0.2358},{"tissuedysplasia":-0.0903,"tissuetumour":0.1995,"AveExpr":3.8717,"F":2.5702,"P.Value":0.088,"adj.P.Val":0.1929},{"tissuedysplasia":-0.0766,"tissuetumour":-0.7091,"AveExpr":1.1427,"F":4.9257,"P.Value":0.0117,"adj.P.Val":0.0464},{"tissuedysplasia":0.0427,"tissuetumour":-0.1816,"AveExpr":5.4003,"F":1.3578,"P.Value":0.2678,"adj.P.Val":0.4114},{"tissuedysplasia":0.509,"tissuetumour":1.5262,"AveExpr":1.0201,"F":7.2156,"P.Value":0.0019,"adj.P.Val":0.0127},{"tissuedysplasia":0.5082,"tissuetumour":0.8787,"AveExpr":3.3959,"F":5.7033,"P.Value":0.0063,"adj.P.Val":0.0297},{"tissuedysplasia":0.0559,"tissuetumour":0.3588,"AveExpr":2.9133,"F":2.9342,"P.Value":0.0636,"adj.P.Val":0.1545},{"tissuedysplasia":0.0888,"tissuetumour":0.4991,"AveExpr":2.8162,"F":2.2857,"P.Value":0.1137,"adj.P.Val":0.2301},{"tissuedysplasia":0.0577,"tissuetumour":0.8529,"AveExpr":0.3846,"F":2.8091,"P.Value":0.0711,"adj.P.Val":0.1668},{"tissuedysplasia":0.4691,"tissuetumour":0.7804,"AveExpr":-0.8692,"F":1.3098,"P.Value":0.2802,"adj.P.Val":0.4247},{"tissuedysplasia":0.2903,"tissuetumour":0.2179,"AveExpr":6.6386,"F":3.6103,"P.Value":0.0353,"adj.P.Val":0.1022},{"tissuedysplasia":0.4133,"tissuetumour":0.2658,"AveExpr":5.2722,"F":5.9358,"P.Value":0.0052,"adj.P.Val":0.0259},{"tissuedysplasia":0.4258,"tissuetumour":0.0664,"AveExpr":3.1877,"F":3.7838,"P.Value":0.0305,"adj.P.Val":0.0922},{"tissuedysplasia":0.0682,"tissuetumour":0.3951,"AveExpr":3.1357,"F":3.2232,"P.Value":0.0494,"adj.P.Val":0.1296},{"tissuedysplasia":0.086,"tissuetumour":0.2634,"AveExpr":5.1469,"F":2.1688,"P.Value":0.1264,"adj.P.Val":0.2469},{"tissuedysplasia":0.3277,"tissuetumour":0.7455,"AveExpr":3.4473,"F":8.1223,"P.Value":0.001,"adj.P.Val":0.0077},{"tissuedysplasia":-0.0228,"tissuetumour":0.0786,"AveExpr":6.2771,"F":0.1175,"P.Value":0.8895,"adj.P.Val":0.9225},{"tissuedysplasia":1.2248,"tissuetumour":2.066,"AveExpr":2.4193,"F":38.2781,"P.Value":2.3407e-10,"adj.P.Val":9.1474e-08},{"tissuedysplasia":0.3888,"tissuetumour":0.4919,"AveExpr":3.4876,"F":7.6016,"P.Value":0.0015,"adj.P.Val":0.0103},{"tissuedysplasia":0.2934,"tissuetumour":0.9356,"AveExpr":4.0896,"F":14.0961,"P.Value":0,"adj.P.Val":0.0004},{"tissuedysplasia":-0.1586,"tissuetumour":0.2786,"AveExpr":3.99,"F":2.7998,"P.Value":0.0717,"adj.P.Val":0.1678},{"tissuedysplasia":0.3747,"tissuetumour":0.1012,"AveExpr":5.8612,"F":4.925,"P.Value":0.0117,"adj.P.Val":0.0464},{"tissuedysplasia":0.0869,"tissuetumour":0.4919,"AveExpr":4.7885,"F":7.3589,"P.Value":0.0017,"adj.P.Val":0.0117},{"tissuedysplasia":0.2626,"tissuetumour":0.5591,"AveExpr":4.479,"F":9.4182,"P.Value":0.0004,"adj.P.Val":0.0039},{"tissuedysplasia":0.1149,"tissuetumour":0.3778,"AveExpr":3.2819,"F":2.1972,"P.Value":0.1232,"adj.P.Val":0.2427},{"tissuedysplasia":0.4205,"tissuetumour":0.4703,"AveExpr":5.584,"F":3.5276,"P.Value":0.0379,"adj.P.Val":0.1075},{"tissuedysplasia":0.1342,"tissuetumour":0.2234,"AveExpr":3.5552,"F":1.3344,"P.Value":0.2738,"adj.P.Val":0.4181},{"tissuedysplasia":0.2413,"tissuetumour":0.4983,"AveExpr":2.5007,"F":2.9498,"P.Value":0.0628,"adj.P.Val":0.1529},{"tissuedysplasia":0.2343,"tissuetumour":0.5173,"AveExpr":5.0297,"F":10.076,"P.Value":0.0002,"adj.P.Val":0.0028},{"tissuedysplasia":0.3846,"tissuetumour":0.4867,"AveExpr":3.7493,"F":6.8858,"P.Value":0.0025,"adj.P.Val":0.0152},{"tissuedysplasia":0.6599,"tissuetumour":1.3434,"AveExpr":3.814,"F":33.6178,"P.Value":1.3747e-09,"adj.P.Val":3.1044e-07},{"tissuedysplasia":0.8201,"tissuetumour":1.1859,"AveExpr":1.8174,"F":5.5793,"P.Value":0.0069,"adj.P.Val":0.0319},{"tissuedysplasia":0.1528,"tissuetumour":0.4722,"AveExpr":5.6837,"F":9.249,"P.Value":0.0004,"adj.P.Val":0.0042},{"tissuedysplasia":0.0283,"tissuetumour":0.3757,"AveExpr":5.9686,"F":6.1148,"P.Value":0.0045,"adj.P.Val":0.0234},{"tissuedysplasia":0.099,"tissuetumour":-0.0291,"AveExpr":5.9395,"F":0.5564,"P.Value":0.5772,"adj.P.Val":0.6901},{"tissuedysplasia":0.1036,"tissuetumour":0.684,"AveExpr":5.0486,"F":15.4021,"P.Value":8.5004e-06,"adj.P.Val":0.0002},{"tissuedysplasia":-0.2339,"tissuetumour":-0.1607,"AveExpr":5.224,"F":2.0825,"P.Value":0.1367,"adj.P.Val":0.261},{"tissuedysplasia":0.2792,"tissuetumour":0.4177,"AveExpr":2.8094,"F":1.7274,"P.Value":0.1896,"adj.P.Val":0.3256},{"tissuedysplasia":0.2728,"tissuetumour":0.0985,"AveExpr":3.6616,"F":3.0456,"P.Value":0.0577,"adj.P.Val":0.1444},{"tissuedysplasia":0.2985,"tissuetumour":-0.0435,"AveExpr":4.8597,"F":5.3095,"P.Value":0.0086,"adj.P.Val":0.0373},{"tissuedysplasia":0.7527,"tissuetumour":1.0602,"AveExpr":2.3777,"F":4.3971,"P.Value":0.0182,"adj.P.Val":0.064},{"tissuedysplasia":0.409,"tissuetumour":0.0203,"AveExpr":6.6703,"F":7.3195,"P.Value":0.0018,"adj.P.Val":0.012},{"tissuedysplasia":0.2635,"tissuetumour":-0.1786,"AveExpr":4.3801,"F":3.1606,"P.Value":0.0522,"adj.P.Val":0.1346},{"tissuedysplasia":0.0274,"tissuetumour":0.2499,"AveExpr":4.0833,"F":1.7006,"P.Value":0.1944,"adj.P.Val":0.3308},{"tissuedysplasia":0.137,"tissuetumour":-0.0097,"AveExpr":4.1676,"F":0.5914,"P.Value":0.5579,"adj.P.Val":0.6744},{"tissuedysplasia":0.5296,"tissuetumour":1.0497,"AveExpr":3.3931,"F":20.7422,"P.Value":4.5107e-07,"adj.P.Val":0},{"tissuedysplasia":-1.6634,"tissuetumour":0.4015,"AveExpr":1.0551,"F":6.569,"P.Value":0.0032,"adj.P.Val":0.0181},{"tissuedysplasia":0.4405,"tissuetumour":1.7243,"AveExpr":-4.663,"F":3.8385,"P.Value":0.0291,"adj.P.Val":0.0892},{"tissuedysplasia":-0.6243,"tissuetumour":-1.2408,"AveExpr":6.2191,"F":17.0358,"P.Value":3.7391e-06,"adj.P.Val":0.0001},{"tissuedysplasia":0.1674,"tissuetumour":1.6635,"AveExpr":-3.5274,"F":1.9514,"P.Value":0.1542,"adj.P.Val":0.2828},{"tissuedysplasia":-0.3242,"tissuetumour":-0.9616,"AveExpr":7.7028,"F":34.6599,"P.Value":9.1376e-10,"adj.P.Val":2.3375e-07},{"tissuedysplasia":0.0219,"tissuetumour":0.1289,"AveExpr":4.1327,"F":0.4448,"P.Value":0.6438,"adj.P.Val":0.7428},{"tissuedysplasia":0.6859,"tissuetumour":1.7235,"AveExpr":3.7636,"F":24.2944,"P.Value":7.788e-08,"adj.P.Val":5.9551e-06},{"tissuedysplasia":-0.4536,"tissuetumour":2.1439,"AveExpr":-2.8356,"F":7.4144,"P.Value":0.0017,"adj.P.Val":0.0114},{"tissuedysplasia":0.2107,"tissuetumour":-0.0884,"AveExpr":4.6239,"F":2.4407,"P.Value":0.0988,"adj.P.Val":0.2091},{"tissuedysplasia":0.1027,"tissuetumour":0.1247,"AveExpr":7.6369,"F":1.0525,"P.Value":0.3577,"adj.P.Val":0.5011},{"tissuedysplasia":0.0522,"tissuetumour":-0.1998,"AveExpr":6.0725,"F":2.493,"P.Value":0.0943,"adj.P.Val":0.2026},{"tissuedysplasia":0.0449,"tissuetumour":-0.168,"AveExpr":5.4442,"F":1.3464,"P.Value":0.2707,"adj.P.Val":0.4146},{"tissuedysplasia":-0.1641,"tissuetumour":0.3231,"AveExpr":4.3863,"F":2.5188,"P.Value":0.0921,"adj.P.Val":0.1992},{"tissuedysplasia":0.8957,"tissuetumour":1.1434,"AveExpr":3.6528,"F":13.4486,"P.Value":0,"adj.P.Val":0.0005},{"tissuedysplasia":-0.0953,"tissuetumour":0.6054,"AveExpr":3.8237,"F":4.1649,"P.Value":0.0221,"adj.P.Val":0.0734},{"tissuedysplasia":1.1484,"tissuetumour":3.7695,"AveExpr":-5.7439,"F":6.9951,"P.Value":0.0023,"adj.P.Val":0.0143},{"tissuedysplasia":0.5689,"tissuetumour":0.0004,"AveExpr":5.2551,"F":8.0913,"P.Value":0.001,"adj.P.Val":0.0078},{"tissuedysplasia":0.1157,"tissuetumour":-0.2211,"AveExpr":2.1277,"F":1.6404,"P.Value":0.2055,"adj.P.Val":0.3435},{"tissuedysplasia":-1.2878,"tissuetumour":-1.6636,"AveExpr":1.6382,"F":12.4771,"P.Value":0.0001,"adj.P.Val":0.0008},{"tissuedysplasia":-0.2569,"tissuetumour":-1.0279,"AveExpr":7.0354,"F":25.6131,"P.Value":4.198e-08,"adj.P.Val":3.8248e-06},{"tissuedysplasia":0.827,"tissuetumour":1.3735,"AveExpr":-1.9673,"F":2.8327,"P.Value":0.0696,"adj.P.Val":0.1643},{"tissuedysplasia":-0.4181,"tissuetumour":-1.0219,"AveExpr":4.926,"F":12.2269,"P.Value":0.0001,"adj.P.Val":0.0009},{"tissuedysplasia":0.0679,"tissuetumour":0.2,"AveExpr":1.1319,"F":0.1289,"P.Value":0.8794,"adj.P.Val":0.9152},{"tissuedysplasia":-0.7405,"tissuetumour":-0.4016,"AveExpr":-2.1297,"F":1.0953,"P.Value":0.3434,"adj.P.Val":0.4874},{"tissuedysplasia":0.4318,"tissuetumour":0.8404,"AveExpr":4.5353,"F":10.662,"P.Value":0.0002,"adj.P.Val":0.0021},{"tissuedysplasia":0.6129,"tissuetumour":0.2545,"AveExpr":6.5579,"F":5.8279,"P.Value":0.0059,"adj.P.Val":0.0283},{"tissuedysplasia":0.1814,"tissuetumour":0.4822,"AveExpr":4.8797,"F":4.9898,"P.Value":0.0111,"adj.P.Val":0.0447},{"tissuedysplasia":0.4371,"tissuetumour":0.3356,"AveExpr":3.0076,"F":4.1757,"P.Value":0.0219,"adj.P.Val":0.073},{"tissuedysplasia":0.3007,"tissuetumour":-0.061,"AveExpr":6.8114,"F":2.6849,"P.Value":0.08,"adj.P.Val":0.1807},{"tissuedysplasia":0.4709,"tissuetumour":0.8825,"AveExpr":4.1764,"F":14.5259,"P.Value":0,"adj.P.Val":0.0003},{"tissuedysplasia":0.5146,"tissuetumour":0.3748,"AveExpr":3.9035,"F":6.168,"P.Value":0.0044,"adj.P.Val":0.0227},{"tissuedysplasia":0.1944,"tissuetumour":0.6472,"AveExpr":2.6633,"F":6.1111,"P.Value":0.0045,"adj.P.Val":0.0235},{"tissuedysplasia":0.3378,"tissuetumour":0.4361,"AveExpr":5.4422,"F":11.5192,"P.Value":0.0001,"adj.P.Val":0.0013},{"tissuedysplasia":0.0192,"tissuetumour":-0.1363,"AveExpr":5.3277,"F":1.0533,"P.Value":0.3574,"adj.P.Val":0.5009},{"tissuedysplasia":-0.3064,"tissuetumour":-1.0753,"AveExpr":2.5815,"F":7.8862,"P.Value":0.0012,"adj.P.Val":0.0088},{"tissuedysplasia":-0.395,"tissuetumour":1.8845,"AveExpr":-6.7975,"F":2.4439,"P.Value":0.0985,"adj.P.Val":0.2088},{"tissuedysplasia":0.0744,"tissuetumour":0.6049,"AveExpr":5.3423,"F":14.3155,"P.Value":0,"adj.P.Val":0.0003},{"tissuedysplasia":0.3939,"tissuetumour":0.5186,"AveExpr":5.0703,"F":11.0876,"P.Value":0.0001,"adj.P.Val":0.0017},{"tissuedysplasia":-0.8579,"tissuetumour":-1.709,"AveExpr":5.3185,"F":17.6341,"P.Value":2.8188e-06,"adj.P.Val":0.0001},{"tissuedysplasia":0.9435,"tissuetumour":1.3423,"AveExpr":4.0376,"F":13.6044,"P.Value":0,"adj.P.Val":0.0005},{"tissuedysplasia":-0.389,"tissuetumour":-0.6485,"AveExpr":5.7034,"F":12.4453,"P.Value":0.0001,"adj.P.Val":0.0008},{"tissuedysplasia":-0.288,"tissuetumour":0.9718,"AveExpr":4.2987,"F":7.1769,"P.Value":0.0021,"adj.P.Val":0.0134},{"tissuedysplasia":-0.0707,"tissuetumour":-1.0002,"AveExpr":5.433,"F":23.0931,"P.Value":1.3888e-07,"adj.P.Val":9.0104e-06},{"tissuedysplasia":-0.4919,"tissuetumour":-0.0956,"AveExpr":0.5678,"F":1.1327,"P.Value":0.3314,"adj.P.Val":0.4758},{"tissuedysplasia":0.2831,"tissuetumour":0.1927,"AveExpr":6.5628,"F":1.4972,"P.Value":0.2349,"adj.P.Val":0.3765},{"tissuedysplasia":0.2891,"tissuetumour":0.262,"AveExpr":3.8478,"F":1.6775,"P.Value":0.1986,"adj.P.Val":0.3355},{"tissuedysplasia":0.0123,"tissuetumour":0.2501,"AveExpr":5.8588,"F":5.5586,"P.Value":0.007,"adj.P.Val":0.0323},{"tissuedysplasia":0.1395,"tissuetumour":-0.0322,"AveExpr":5.6681,"F":1.3714,"P.Value":0.2644,"adj.P.Val":0.4077},{"tissuedysplasia":-0.0059,"tissuetumour":0.8379,"AveExpr":3.5684,"F":12.4096,"P.Value":0.0001,"adj.P.Val":0.0009},{"tissuedysplasia":0.2752,"tissuetumour":0.4239,"AveExpr":-1.1455,"F":0.3256,"P.Value":0.7238,"adj.P.Val":0.8027},{"tissuedysplasia":0.7811,"tissuetumour":2.9706,"AveExpr":2.9257,"F":64.8122,"P.Value":7.6521e-14,"adj.P.Val":5.5513e-10},{"tissuedysplasia":0.077,"tissuetumour":0.5125,"AveExpr":5.4109,"F":10.7486,"P.Value":0.0002,"adj.P.Val":0.002},{"tissuedysplasia":0.2193,"tissuetumour":0.8142,"AveExpr":3.4696,"F":12.1376,"P.Value":0.0001,"adj.P.Val":0.001},{"tissuedysplasia":-0.045,"tissuetumour":0.174,"AveExpr":5.7276,"F":1.0248,"P.Value":0.3673,"adj.P.Val":0.51},{"tissuedysplasia":0.2685,"tissuetumour":0.4634,"AveExpr":5.0869,"F":7.8483,"P.Value":0.0012,"adj.P.Val":0.009},{"tissuedysplasia":0.0673,"tissuetumour":0.4065,"AveExpr":3.6985,"F":5.8524,"P.Value":0.0056,"adj.P.Val":0.0272},{"tissuedysplasia":0.059,"tissuetumour":0.093,"AveExpr":3.5602,"F":0.2216,"P.Value":0.8021,"adj.P.Val":0.8612},{"tissuedysplasia":-0.3322,"tissuetumour":-1.3303,"AveExpr":2.5084,"F":8.3243,"P.Value":0.0009,"adj.P.Val":0.0069},{"tissuedysplasia":0.0292,"tissuetumour":0.0332,"AveExpr":4.3314,"F":0.0373,"P.Value":0.9634,"adj.P.Val":0.975},{"tissuedysplasia":-0.6447,"tissuetumour":-0.4828,"AveExpr":0.2855,"F":1.5121,"P.Value":0.2317,"adj.P.Val":0.3728},{"tissuedysplasia":0.0579,"tissuetumour":0.3394,"AveExpr":4.7858,"F":5.1024,"P.Value":0.0102,"adj.P.Val":0.0419},{"tissuedysplasia":-0.4007,"tissuetumour":-0.2911,"AveExpr":6.4409,"F":8.2114,"P.Value":0.0009,"adj.P.Val":0.0073},{"tissuedysplasia":0.0747,"tissuetumour":0.2734,"AveExpr":3.5095,"F":1.6921,"P.Value":0.1959,"adj.P.Val":0.3325},{"tissuedysplasia":0.3548,"tissuetumour":0.7585,"AveExpr":3.5087,"F":8.7245,"P.Value":0.0006,"adj.P.Val":0.0056},{"tissuedysplasia":2.4995,"tissuetumour":3.413,"AveExpr":1.5762,"F":22.7159,"P.Value":1.6707e-07,"adj.P.Val":0},{"tissuedysplasia":0.3896,"tissuetumour":-0.0106,"AveExpr":3.6718,"F":4.2464,"P.Value":0.0206,"adj.P.Val":0.07},{"tissuedysplasia":1.1338,"tissuetumour":3.3846,"AveExpr":-5.787,"F":5.0502,"P.Value":0.0106,"adj.P.Val":0.0433},{"tissuedysplasia":-0.3098,"tissuetumour":-0.7074,"AveExpr":9.6249,"F":13.2635,"P.Value":0,"adj.P.Val":0.0008},{"tissuedysplasia":0.0249,"tissuetumour":-0.3822,"AveExpr":2.9668,"F":3.4969,"P.Value":0.039,"adj.P.Val":0.1095},{"tissuedysplasia":0.5237,"tissuetumour":0.3718,"AveExpr":2.8505,"F":4.8429,"P.Value":0.0126,"adj.P.Val":0.0488},{"tissuedysplasia":0.3022,"tissuetumour":0.2803,"AveExpr":5.6072,"F":3.2899,"P.Value":0.0466,"adj.P.Val":0.1243},{"tissuedysplasia":-0.2828,"tissuetumour":-0.0252,"AveExpr":4.4505,"F":3.4269,"P.Value":0.0414,"adj.P.Val":0.1143},{"tissuedysplasia":0.3039,"tissuetumour":0.1714,"AveExpr":3.2188,"F":1.602,"P.Value":0.213,"adj.P.Val":0.3521},{"tissuedysplasia":-0.6543,"tissuetumour":0.7326,"AveExpr":0.891,"F":6.3453,"P.Value":0.0038,"adj.P.Val":0.0206},{"tissuedysplasia":0.2991,"tissuetumour":0.8085,"AveExpr":5.4114,"F":19.4293,"P.Value":8.9607e-07,"adj.P.Val":0},{"tissuedysplasia":0.1806,"tissuetumour":0.0941,"AveExpr":6.7017,"F":1.6161,"P.Value":0.2102,"adj.P.Val":0.3488},{"tissuedysplasia":0.3341,"tissuetumour":0.1506,"AveExpr":5.5604,"F":5.916,"P.Value":0.0053,"adj.P.Val":0.0262},{"tissuedysplasia":2.435,"tissuetumour":4.0591,"AveExpr":-4.0164,"F":12.5372,"P.Value":0,"adj.P.Val":0.0008},{"tissuedysplasia":0.9994,"tissuetumour":1.6956,"AveExpr":-0.5579,"F":5.6447,"P.Value":0.0066,"adj.P.Val":0.0307},{"tissuedysplasia":0.2127,"tissuetumour":0.0833,"AveExpr":3.5246,"F":0.9759,"P.Value":0.3849,"adj.P.Val":0.5264},{"tissuedysplasia":0.5114,"tissuetumour":0.0625,"AveExpr":5.7787,"F":8.8544,"P.Value":0.0006,"adj.P.Val":0.0052},{"tissuedysplasia":-0.169,"tissuetumour":-0.1871,"AveExpr":-3.8324,"F":0.0251,"P.Value":0.9752,"adj.P.Val":0.9829},{"tissuedysplasia":-0.1665,"tissuetumour":-0.4773,"AveExpr":5.8549,"F":7.943,"P.Value":0.0011,"adj.P.Val":0.0085},{"tissuedysplasia":0.1574,"tissuetumour":0.4239,"AveExpr":7.3543,"F":6.6055,"P.Value":0.0031,"adj.P.Val":0.0178},{"tissuedysplasia":0.2581,"tissuetumour":-0.0139,"AveExpr":4.492,"F":1.5868,"P.Value":0.2161,"adj.P.Val":0.3555},{"tissuedysplasia":-0.223,"tissuetumour":-0.3811,"AveExpr":7.3594,"F":4.91,"P.Value":0.0119,"adj.P.Val":0.0468},{"tissuedysplasia":-0.087,"tissuetumour":-0.0047,"AveExpr":6.9145,"F":0.3984,"P.Value":0.6738,"adj.P.Val":0.7658},{"tissuedysplasia":0.2667,"tissuetumour":-0.0937,"AveExpr":5.7779,"F":4.515,"P.Value":0.0165,"adj.P.Val":0.0595},{"tissuedysplasia":1.4801,"tissuetumour":3.4386,"AveExpr":-4.6774,"F":3.938,"P.Value":0.0267,"adj.P.Val":0.084},{"tissuedysplasia":-0.5035,"tissuetumour":-1.0878,"AveExpr":5.3471,"F":39.3359,"P.Value":1.5963e-10,"adj.P.Val":7.1266e-08},{"tissuedysplasia":0.1932,"tissuetumour":0.3911,"AveExpr":4.5805,"F":3.2046,"P.Value":0.0502,"adj.P.Val":0.1311},{"tissuedysplasia":0.2938,"tissuetumour":-0.2473,"AveExpr":2.8714,"F":3.9839,"P.Value":0.0257,"adj.P.Val":0.0818},{"tissuedysplasia":0.5411,"tissuetumour":1.1712,"AveExpr":4.2084,"F":21.4487,"P.Value":3.1448e-07,"adj.P.Val":0},{"tissuedysplasia":-0.2774,"tissuetumour":0.2169,"AveExpr":4.6151,"F":4.5614,"P.Value":0.0158,"adj.P.Val":0.0579},{"tissuedysplasia":0.2157,"tissuetumour":0.6728,"AveExpr":4.3852,"F":8.6991,"P.Value":0.0007,"adj.P.Val":0.0057},{"tissuedysplasia":0.2804,"tissuetumour":0.0626,"AveExpr":6.2639,"F":2.6992,"P.Value":0.0784,"adj.P.Val":0.1781},{"tissuedysplasia":1.0328,"tissuetumour":1.3722,"AveExpr":2.5216,"F":20.3516,"P.Value":5.5203e-07,"adj.P.Val":0},{"tissuedysplasia":-0.062,"tissuetumour":0.4037,"AveExpr":4.8978,"F":10.6821,"P.Value":0.0002,"adj.P.Val":0.002},{"tissuedysplasia":0.4775,"tissuetumour":1.076,"AveExpr":4.5138,"F":25.133,"P.Value":5.2465e-08,"adj.P.Val":4.4789e-06},{"tissuedysplasia":0.6099,"tissuetumour":0.6726,"AveExpr":4.6271,"F":9.0653,"P.Value":0.0005,"adj.P.Val":0.0047},{"tissuedysplasia":0.0528,"tissuetumour":0.4377,"AveExpr":5.7878,"F":12.0932,"P.Value":0.0001,"adj.P.Val":0.001},{"tissuedysplasia":-0.051,"tissuetumour":1.1223,"AveExpr":4.0285,"F":15.8985,"P.Value":6.3601e-06,"adj.P.Val":0.0002},{"tissuedysplasia":1.0308,"tissuetumour":0.3518,"AveExpr":-10.7786,"F":0.572,"P.Value":0.5685,"adj.P.Val":0.6829},{"tissuedysplasia":0.8183,"tissuetumour":0.6195,"AveExpr":4.0317,"F":7.5853,"P.Value":0.0015,"adj.P.Val":0.0103},{"tissuedysplasia":0.1575,"tissuetumour":1.1389,"AveExpr":3.9087,"F":27.4539,"P.Value":1.8222e-08,"adj.P.Val":2.1541e-06},{"tissuedysplasia":0.1516,"tissuetumour":0.5767,"AveExpr":3.065,"F":6.6658,"P.Value":0.003,"adj.P.Val":0.0172},{"tissuedysplasia":0.2563,"tissuetumour":0.9209,"AveExpr":4.7274,"F":23.4405,"P.Value":1.173e-07,"adj.P.Val":7.9439e-06},{"tissuedysplasia":1.0601,"tissuetumour":1.8073,"AveExpr":2.5688,"F":14.826,"P.Value":0,"adj.P.Val":0.0003},{"tissuedysplasia":-0.2849,"tissuetumour":-0.68,"AveExpr":5.7475,"F":9.0576,"P.Value":0.0005,"adj.P.Val":0.0047},{"tissuedysplasia":-0.7756,"tissuetumour":-1.0774,"AveExpr":1.75,"F":4.1126,"P.Value":0.023,"adj.P.Val":0.0757},{"tissuedysplasia":0.2443,"tissuetumour":-0.0624,"AveExpr":4.6241,"F":4.4677,"P.Value":0.0171,"adj.P.Val":0.0613},{"tissuedysplasia":0.0298,"tissuetumour":0.5178,"AveExpr":3.5322,"F":2.5965,"P.Value":0.0859,"adj.P.Val":0.1897},{"tissuedysplasia":-0.0837,"tissuetumour":-0.5774,"AveExpr":6.6219,"F":3.779,"P.Value":0.0321,"adj.P.Val":0.0956},{"tissuedysplasia":0.0336,"tissuetumour":0.5232,"AveExpr":5.0089,"F":9.3213,"P.Value":0.0004,"adj.P.Val":0.0041},{"tissuedysplasia":0.0817,"tissuetumour":0.5338,"AveExpr":4.1799,"F":3.436,"P.Value":0.0411,"adj.P.Val":0.1137},{"tissuedysplasia":0.8722,"tissuetumour":3.3352,"AveExpr":-3.9031,"F":8.3909,"P.Value":0.0008,"adj.P.Val":0.0067},{"tissuedysplasia":0.1252,"tissuetumour":1.1691,"AveExpr":6.7461,"F":21.8246,"P.Value":3.5736e-07,"adj.P.Val":0},{"tissuedysplasia":0.3364,"tissuetumour":0.47,"AveExpr":5.573,"F":10.3339,"P.Value":0.0002,"adj.P.Val":0.0024},{"tissuedysplasia":-1.2611,"tissuetumour":-0.8093,"AveExpr":0.8845,"F":6.8204,"P.Value":0.0026,"adj.P.Val":0.0158},{"tissuedysplasia":-0.4153,"tissuetumour":-0.6899,"AveExpr":5.711,"F":26.7047,"P.Value":2.5495e-08,"adj.P.Val":2.6612e-06},{"tissuedysplasia":0.2852,"tissuetumour":0.4241,"AveExpr":4.9387,"F":4.1644,"P.Value":0.0221,"adj.P.Val":0.0734},{"tissuedysplasia":0.0578,"tissuetumour":0.7706,"AveExpr":2.3658,"F":6.311,"P.Value":0.0039,"adj.P.Val":0.021},{"tissuedysplasia":0.8896,"tissuetumour":1.9889,"AveExpr":-1.0644,"F":7.0903,"P.Value":0.0021,"adj.P.Val":0.0136},{"tissuedysplasia":0.4599,"tissuetumour":2.3513,"AveExpr":2.8654,"F":27.962,"P.Value":1.4552e-08,"adj.P.Val":1.84e-06},{"tissuedysplasia":-0.5565,"tissuetumour":-1.1641,"AveExpr":3.8542,"F":25.3969,"P.Value":4.6401e-08,"adj.P.Val":4.0847e-06},{"tissuedysplasia":-0.478,"tissuetumour":-1.2524,"AveExpr":5.94,"F":31.6806,"P.Value":2.9989e-09,"adj.P.Val":5.6509e-07},{"tissuedysplasia":0.701,"tissuetumour":0.4096,"AveExpr":6.593,"F":10.2871,"P.Value":0.0002,"adj.P.Val":0.0025},{"tissuedysplasia":0.2999,"tissuetumour":0.7286,"AveExpr":5.3299,"F":6.5838,"P.Value":0.0032,"adj.P.Val":0.018},{"tissuedysplasia":-0.3872,"tissuetumour":0.3695,"AveExpr":-0.1881,"F":1.5878,"P.Value":0.2159,"adj.P.Val":0.3554},{"tissuedysplasia":-0.1587,"tissuetumour":-0.6283,"AveExpr":7.511,"F":13.4699,"P.Value":0,"adj.P.Val":0.0005},{"tissuedysplasia":-0.1115,"tissuetumour":-0.5513,"AveExpr":3.2618,"F":4.5469,"P.Value":0.016,"adj.P.Val":0.0584},{"tissuedysplasia":-0.1099,"tissuetumour":-0.5157,"AveExpr":5.4319,"F":9.2173,"P.Value":0.0005,"adj.P.Val":0.0043},{"tissuedysplasia":0.3809,"tissuetumour":1.2085,"AveExpr":2.147,"F":11.9721,"P.Value":0.0001,"adj.P.Val":0.0011},{"tissuedysplasia":0.2523,"tissuetumour":0.2892,"AveExpr":6.3672,"F":6.0767,"P.Value":0.0047,"adj.P.Val":0.0239},{"tissuedysplasia":-0.0736,"tissuetumour":0.1207,"AveExpr":2.9714,"F":0.5518,"P.Value":0.5799,"adj.P.Val":0.6922},{"tissuedysplasia":0.121,"tissuetumour":0.1972,"AveExpr":5.8829,"F":2.2509,"P.Value":0.1173,"adj.P.Val":0.235},{"tissuedysplasia":0.7357,"tissuetumour":0.666,"AveExpr":2.3209,"F":4.6439,"P.Value":0.0148,"adj.P.Val":0.0551},{"tissuedysplasia":0.0222,"tissuetumour":0.2172,"AveExpr":5.3847,"F":1.329,"P.Value":0.2752,"adj.P.Val":0.4196},{"tissuedysplasia":-0.131,"tissuetumour":0.1876,"AveExpr":1.2785,"F":0.2723,"P.Value":0.7629,"adj.P.Val":0.832},{"tissuedysplasia":0.3653,"tissuetumour":1.3397,"AveExpr":4.463,"F":26.2533,"P.Value":3.1291e-08,"adj.P.Val":3.1365e-06},{"tissuedysplasia":0.4113,"tissuetumour":0.7209,"AveExpr":3.2932,"F":5.3651,"P.Value":0.0082,"adj.P.Val":0.036},{"tissuedysplasia":0.1215,"tissuetumour":-0.2667,"AveExpr":6.0292,"F":8.5081,"P.Value":0.0008,"adj.P.Val":0.0063},{"tissuedysplasia":-0.0444,"tissuetumour":-0.2982,"AveExpr":3.8465,"F":2.1191,"P.Value":0.1322,"adj.P.Val":0.255},{"tissuedysplasia":-0.4231,"tissuetumour":-0.2997,"AveExpr":6.6933,"F":6.3035,"P.Value":0.0039,"adj.P.Val":0.021},{"tissuedysplasia":0.9156,"tissuetumour":1.2094,"AveExpr":4.83,"F":23.8912,"P.Value":9.4404e-08,"adj.P.Val":6.8486e-06},{"tissuedysplasia":-0.1787,"tissuetumour":-0.7619,"AveExpr":4.8896,"F":15.0431,"P.Value":0,"adj.P.Val":0.0002},{"tissuedysplasia":0.6903,"tissuetumour":0.9718,"AveExpr":4.0259,"F":10.3762,"P.Value":0.0002,"adj.P.Val":0.0024},{"tissuedysplasia":-0.2074,"tissuetumour":0.3426,"AveExpr":3.7513,"F":5.8222,"P.Value":0.0057,"adj.P.Val":0.0277},{"tissuedysplasia":0.061,"tissuetumour":1.0036,"AveExpr":4.672,"F":23.5167,"P.Value":1.1306e-07,"adj.P.Val":7.7706e-06},{"tissuedysplasia":0.2895,"tissuetumour":0.1589,"AveExpr":4.1508,"F":2.4632,"P.Value":0.0968,"adj.P.Val":0.2064},{"tissuedysplasia":0.6722,"tissuetumour":0.6877,"AveExpr":2.5954,"F":4.524,"P.Value":0.0163,"adj.P.Val":0.0592},{"tissuedysplasia":0.6363,"tissuetumour":0.938,"AveExpr":5.3356,"F":21.3683,"P.Value":3.2755e-07,"adj.P.Val":0},{"tissuedysplasia":0.2016,"tissuetumour":0.5504,"AveExpr":3.5159,"F":5.4887,"P.Value":0.0074,"adj.P.Val":0.0336},{"tissuedysplasia":-0.7286,"tissuetumour":1.134,"AveExpr":0.1907,"F":9.5671,"P.Value":0.0004,"adj.P.Val":0.0036},{"tissuedysplasia":0.171,"tissuetumour":0.6216,"AveExpr":4.2219,"F":5.9476,"P.Value":0.0052,"adj.P.Val":0.0258},{"tissuedysplasia":0.1846,"tissuetumour":0.7104,"AveExpr":4.2688,"F":9.7669,"P.Value":0.0003,"adj.P.Val":0.0033},{"tissuedysplasia":0.1468,"tissuetumour":0.721,"AveExpr":3.4889,"F":10.9309,"P.Value":0.0001,"adj.P.Val":0.0018},{"tissuedysplasia":0.6015,"tissuetumour":0.8989,"AveExpr":3.2995,"F":12.1586,"P.Value":0.0001,"adj.P.Val":0.001},{"tissuedysplasia":0.551,"tissuetumour":0.9513,"AveExpr":-2.1145,"F":1.2584,"P.Value":0.2941,"adj.P.Val":0.4385},{"tissuedysplasia":0.9875,"tissuetumour":1.4551,"AveExpr":3.5364,"F":34.9691,"P.Value":8.1066e-10,"adj.P.Val":2.1582e-07},{"tissuedysplasia":0.8497,"tissuetumour":2.137,"AveExpr":0.4769,"F":13.5876,"P.Value":0,"adj.P.Val":0.0005},{"tissuedysplasia":0.8805,"tissuetumour":1.4579,"AveExpr":-0.3084,"F":5.6054,"P.Value":0.0068,"adj.P.Val":0.0314},{"tissuedysplasia":0.1024,"tissuetumour":0.511,"AveExpr":2.7423,"F":6.1836,"P.Value":0.0043,"adj.P.Val":0.0225},{"tissuedysplasia":0.6955,"tissuetumour":0.5619,"AveExpr":8.1344,"F":11.5956,"P.Value":0.0001,"adj.P.Val":0.0016},{"tissuedysplasia":0.2152,"tissuetumour":0.0706,"AveExpr":4.7361,"F":1.1992,"P.Value":0.3111,"adj.P.Val":0.4555},{"tissuedysplasia":-0.3692,"tissuetumour":-0.1944,"AveExpr":4.5042,"F":2.5658,"P.Value":0.0883,"adj.P.Val":0.1934},{"tissuedysplasia":0.3653,"tissuetumour":0.1882,"AveExpr":4.5398,"F":2.9943,"P.Value":0.0604,"adj.P.Val":0.1491},{"tissuedysplasia":0.1827,"tissuetumour":0.3984,"AveExpr":1.3284,"F":0.5693,"P.Value":0.57,"adj.P.Val":0.6842},{"tissuedysplasia":-0.1341,"tissuetumour":0.2593,"AveExpr":2.3634,"F":1.5703,"P.Value":0.2194,"adj.P.Val":0.3594},{"tissuedysplasia":0.4888,"tissuetumour":0.9323,"AveExpr":5.2191,"F":17.494,"P.Value":2.567e-06,"adj.P.Val":0.0001},{"tissuedysplasia":0.1845,"tissuetumour":0.2293,"AveExpr":5.6265,"F":2.784,"P.Value":0.0727,"adj.P.Val":0.1694},{"tissuedysplasia":0.0773,"tissuetumour":0.4678,"AveExpr":-2.5838,"F":0.2022,"P.Value":0.8177,"adj.P.Val":0.8722},{"tissuedysplasia":0.529,"tissuetumour":0.7581,"AveExpr":4.2636,"F":13.6772,"P.Value":0,"adj.P.Val":0.0005},{"tissuedysplasia":0.1927,"tissuetumour":0.1346,"AveExpr":6.7084,"F":2.5001,"P.Value":0.0937,"adj.P.Val":0.2017},{"tissuedysplasia":0.1761,"tissuetumour":0.6159,"AveExpr":-1.569,"F":0.7406,"P.Value":0.4827,"adj.P.Val":0.6117},{"tissuedysplasia":0.632,"tissuetumour":0.3136,"AveExpr":5.5277,"F":10.113,"P.Value":0.0002,"adj.P.Val":0.0027},{"tissuedysplasia":0.1443,"tissuetumour":0.7131,"AveExpr":3.6497,"F":6.0138,"P.Value":0.0049,"adj.P.Val":0.0248},{"tissuedysplasia":0.9945,"tissuetumour":1.9519,"AveExpr":-0.5374,"F":6.9231,"P.Value":0.0024,"adj.P.Val":0.0149},{"tissuedysplasia":0.0501,"tissuetumour":-0.7748,"AveExpr":3.3299,"F":3.6062,"P.Value":0.0355,"adj.P.Val":0.1024},{"tissuedysplasia":0.1228,"tissuetumour":1.9852,"AveExpr":-3.1128,"F":5.0875,"P.Value":0.0103,"adj.P.Val":0.0423},{"tissuedysplasia":0.4074,"tissuetumour":0.5613,"AveExpr":3.7408,"F":0.8481,"P.Value":0.4358,"adj.P.Val":0.5727},{"tissuedysplasia":1.5575,"tissuetumour":2.8698,"AveExpr":4.8208,"F":28.4801,"P.Value":2.1092e-08,"adj.P.Val":2.3541e-06},{"tissuedysplasia":0.4546,"tissuetumour":0.2407,"AveExpr":5.092,"F":8.6955,"P.Value":0.0007,"adj.P.Val":0.0057},{"tissuedysplasia":0.8552,"tissuetumour":0.5788,"AveExpr":5.0153,"F":6.4067,"P.Value":0.0036,"adj.P.Val":0.0199},{"tissuedysplasia":0.6616,"tissuetumour":0.5319,"AveExpr":-0.3028,"F":1.1791,"P.Value":0.3171,"adj.P.Val":0.4614},{"tissuedysplasia":0.1312,"tissuetumour":0.4568,"AveExpr":1.2678,"F":1.8204,"P.Value":0.1739,"adj.P.Val":0.3071},{"tissuedysplasia":-0.3181,"tissuetumour":-0.3432,"AveExpr":4.4095,"F":2.581,"P.Value":0.0871,"adj.P.Val":0.1916},{"tissuedysplasia":0.2133,"tissuetumour":-0.0267,"AveExpr":6.8291,"F":1.8964,"P.Value":0.1622,"adj.P.Val":0.2926},{"tissuedysplasia":0.0962,"tissuetumour":-0.2,"AveExpr":4.6696,"F":3.3382,"P.Value":0.0447,"adj.P.Val":0.1207},{"tissuedysplasia":0.5872,"tissuetumour":1.9229,"AveExpr":2.6019,"F":21.5554,"P.Value":2.9795e-07,"adj.P.Val":0},{"tissuedysplasia":-0.0152,"tissuetumour":0.1444,"AveExpr":6.5427,"F":1.2588,"P.Value":0.294,"adj.P.Val":0.4384},{"tissuedysplasia":0.445,"tissuetumour":1.2585,"AveExpr":4.0104,"F":19.0069,"P.Value":1.1227e-06,"adj.P.Val":0},{"tissuedysplasia":-0.0386,"tissuetumour":0.3332,"AveExpr":6.8757,"F":3.7221,"P.Value":0.0321,"adj.P.Val":0.0956},{"tissuedysplasia":0.1885,"tissuetumour":0.1054,"AveExpr":5.3908,"F":0.7901,"P.Value":0.4601,"adj.P.Val":0.5935},{"tissuedysplasia":-0.0009,"tissuetumour":0.4721,"AveExpr":6.0843,"F":2.9945,"P.Value":0.061,"adj.P.Val":0.15},{"tissuedysplasia":0.4558,"tissuetumour":1.3262,"AveExpr":2.8722,"F":17.5585,"P.Value":2.4767e-06,"adj.P.Val":0.0001},{"tissuedysplasia":0.2588,"tissuetumour":0.3438,"AveExpr":4.3128,"F":3.723,"P.Value":0.0321,"adj.P.Val":0.0956},{"tissuedysplasia":0.9006,"tissuetumour":1.3246,"AveExpr":0.265,"F":7.1872,"P.Value":0.002,"adj.P.Val":0.0129},{"tissuedysplasia":-0.0319,"tissuetumour":2.6188,"AveExpr":-6.2838,"F":3.5291,"P.Value":0.0379,"adj.P.Val":0.1074},{"tissuedysplasia":-0.0275,"tissuetumour":-0.1044,"AveExpr":4.1857,"F":0.4104,"P.Value":0.6659,"adj.P.Val":0.7598},{"tissuedysplasia":0.0124,"tissuetumour":0.0117,"AveExpr":6.0636,"F":0.0121,"P.Value":0.988,"adj.P.Val":0.9915},{"tissuedysplasia":1.072,"tissuetumour":1.4112,"AveExpr":3.8497,"F":12.931,"P.Value":0,"adj.P.Val":0.0007},{"tissuedysplasia":0.1572,"tissuetumour":0.2673,"AveExpr":6.8996,"F":0.9794,"P.Value":0.3843,"adj.P.Val":0.5259},{"tissuedysplasia":-1.1007,"tissuetumour":-1.8632,"AveExpr":5.9619,"F":43.2458,"P.Value":4.0987e-11,"adj.P.Val":2.6727e-08},{"tissuedysplasia":-0.2451,"tissuetumour":-0.419,"AveExpr":3.0627,"F":1.3105,"P.Value":0.28,"adj.P.Val":0.4245},{"tissuedysplasia":0.0264,"tissuetumour":0.3372,"AveExpr":3.5061,"F":3.8879,"P.Value":0.0279,"adj.P.Val":0.0867},{"tissuedysplasia":0.7296,"tissuetumour":1.3527,"AveExpr":3.792,"F":31.7414,"P.Value":2.9251e-09,"adj.P.Val":5.5954e-07},{"tissuedysplasia":0.1977,"tissuetumour":-0.0059,"AveExpr":5.5716,"F":2.2736,"P.Value":0.1149,"adj.P.Val":0.2318},{"tissuedysplasia":0.3954,"tissuetumour":0.4588,"AveExpr":4.1262,"F":6.1916,"P.Value":0.0043,"adj.P.Val":0.0224},{"tissuedysplasia":0.4442,"tissuetumour":0.5246,"AveExpr":5.7776,"F":11.9248,"P.Value":0.0001,"adj.P.Val":0.0011},{"tissuedysplasia":0.5898,"tissuetumour":1.2239,"AveExpr":1.5862,"F":6.9738,"P.Value":0.0023,"adj.P.Val":0.0145},{"tissuedysplasia":-0.008,"tissuetumour":-0.2534,"AveExpr":6.0683,"F":2.0611,"P.Value":0.1394,"adj.P.Val":0.2641},{"tissuedysplasia":0.5409,"tissuetumour":0.6339,"AveExpr":5.1048,"F":20.018,"P.Value":6.5692e-07,"adj.P.Val":0},{"tissuedysplasia":0.623,"tissuetumour":0.0499,"AveExpr":3.4967,"F":5.4329,"P.Value":0.0078,"adj.P.Val":0.0347},{"tissuedysplasia":0.1261,"tissuetumour":0.1997,"AveExpr":6.6494,"F":3.9719,"P.Value":0.026,"adj.P.Val":0.0823},{"tissuedysplasia":0.3919,"tissuetumour":2.6125,"AveExpr":-5.9945,"F":4.8604,"P.Value":0.0124,"adj.P.Val":0.0483},{"tissuedysplasia":0.0734,"tissuetumour":0.4619,"AveExpr":8.0518,"F":7.4455,"P.Value":0.0018,"adj.P.Val":0.0118},{"tissuedysplasia":0.7821,"tissuetumour":2.3498,"AveExpr":0.2844,"F":14.86,"P.Value":0,"adj.P.Val":0.0003},{"tissuedysplasia":0.4589,"tissuetumour":1.0876,"AveExpr":3.1471,"F":21.3607,"P.Value":3.2883e-07,"adj.P.Val":0},{"tissuedysplasia":-0.3638,"tissuetumour":-0.1136,"AveExpr":3.5596,"F":1.548,"P.Value":0.224,"adj.P.Val":0.3645},{"tissuedysplasia":-0.095,"tissuetumour":-0.3114,"AveExpr":3.3775,"F":2.0964,"P.Value":0.135,"adj.P.Val":0.2588},{"tissuedysplasia":-0.4285,"tissuetumour":-0.2816,"AveExpr":5.0086,"F":7.0706,"P.Value":0.0022,"adj.P.Val":0.0137},{"tissuedysplasia":0.4652,"tissuetumour":-0.0243,"AveExpr":2.3067,"F":2.5,"P.Value":0.0937,"adj.P.Val":0.2017},{"tissuedysplasia":1.0892,"tissuetumour":1.152,"AveExpr":0.2896,"F":3.2091,"P.Value":0.05,"adj.P.Val":0.1307},{"tissuedysplasia":0.8466,"tissuetumour":2.9365,"AveExpr":-2.6707,"F":8.4302,"P.Value":0.0008,"adj.P.Val":0.0066},{"tissuedysplasia":-0.2412,"tissuetumour":-0.938,"AveExpr":0.5832,"F":2.3649,"P.Value":0.1058,"adj.P.Val":0.219},{"tissuedysplasia":0.6267,"tissuetumour":0.3789,"AveExpr":2.9495,"F":6.2163,"P.Value":0.0042,"adj.P.Val":0.0221},{"tissuedysplasia":-0.0869,"tissuetumour":0.0212,"AveExpr":2.3539,"F":0.2109,"P.Value":0.8107,"adj.P.Val":0.8671},{"tissuedysplasia":0.0917,"tissuetumour":-0.0647,"AveExpr":3.697,"F":0.4854,"P.Value":0.6187,"adj.P.Val":0.7236},{"tissuedysplasia":0.2103,"tissuetumour":0.2445,"AveExpr":6.8433,"F":4.041,"P.Value":0.0245,"adj.P.Val":0.079},{"tissuedysplasia":0.3506,"tissuetumour":0.218,"AveExpr":2.8457,"F":2.0434,"P.Value":0.1417,"adj.P.Val":0.267},{"tissuedysplasia":0.0503,"tissuetumour":0.6627,"AveExpr":3.081,"F":3.9488,"P.Value":0.0265,"adj.P.Val":0.0835},{"tissuedysplasia":0.5229,"tissuetumour":1.1284,"AveExpr":2.5201,"F":14.2665,"P.Value":0,"adj.P.Val":0.0003},{"tissuedysplasia":1.3542,"tissuetumour":1.9767,"AveExpr":4.9417,"F":15.1991,"P.Value":0,"adj.P.Val":0.0003},{"tissuedysplasia":0.1038,"tissuetumour":0.5276,"AveExpr":2.698,"F":5.5284,"P.Value":0.0072,"adj.P.Val":0.0329},{"tissuedysplasia":-0.1776,"tissuetumour":-0.2355,"AveExpr":3.7626,"F":1.7752,"P.Value":0.1814,"adj.P.Val":0.3161},{"tissuedysplasia":0.0838,"tissuetumour":0.4917,"AveExpr":4.8935,"F":6.664,"P.Value":0.003,"adj.P.Val":0.0172},{"tissuedysplasia":0.3867,"tissuetumour":0.9498,"AveExpr":6.4808,"F":18.0858,"P.Value":1.8507e-06,"adj.P.Val":0.0001},{"tissuedysplasia":0.8486,"tissuetumour":1.1759,"AveExpr":-2.2404,"F":1.8431,"P.Value":0.1703,"adj.P.Val":0.3028},{"tissuedysplasia":0.0781,"tissuetumour":-0.1967,"AveExpr":6.9971,"F":3.7306,"P.Value":0.0319,"adj.P.Val":0.0951},{"tissuedysplasia":0.1356,"tissuetumour":-0.2383,"AveExpr":2.742,"F":1.6385,"P.Value":0.2059,"adj.P.Val":0.3439},{"tissuedysplasia":-0.0244,"tissuetumour":0.0187,"AveExpr":3.1098,"F":0.0435,"P.Value":0.9574,"adj.P.Val":0.9709},{"tissuedysplasia":0.6926,"tissuetumour":1.962,"AveExpr":0.8037,"F":11.1768,"P.Value":0.0001,"adj.P.Val":0.0016},{"tissuedysplasia":0.5293,"tissuetumour":3.0553,"AveExpr":-4.2436,"F":4.5731,"P.Value":0.0157,"adj.P.Val":0.0575},{"tissuedysplasia":-0.3599,"tissuetumour":-0.6842,"AveExpr":6.3858,"F":20.9215,"P.Value":4.1137e-07,"adj.P.Val":0},{"tissuedysplasia":0.1151,"tissuetumour":0.7297,"AveExpr":3.5286,"F":8.6239,"P.Value":0.0007,"adj.P.Val":0.0059},{"tissuedysplasia":0.1267,"tissuetumour":-0.0054,"AveExpr":5.3502,"F":0.8676,"P.Value":0.427,"adj.P.Val":0.5647},{"tissuedysplasia":0.185,"tissuetumour":0.1813,"AveExpr":6.8154,"F":1.4551,"P.Value":0.2444,"adj.P.Val":0.3869},{"tissuedysplasia":-0.0088,"tissuetumour":-0.0996,"AveExpr":6.5724,"F":1.0635,"P.Value":0.354,"adj.P.Val":0.4976},{"tissuedysplasia":0.7568,"tissuetumour":1.2575,"AveExpr":6.7312,"F":27.6248,"P.Value":1.6889e-08,"adj.P.Val":2.0506e-06},{"tissuedysplasia":-0.4212,"tissuetumour":-0.4756,"AveExpr":8.0724,"F":10.6959,"P.Value":0.0002,"adj.P.Val":0.0021},{"tissuedysplasia":0.7663,"tissuetumour":2.3712,"AveExpr":-8.1817,"F":2.5112,"P.Value":0.0927,"adj.P.Val":0.2002},{"tissuedysplasia":-0.0615,"tissuetumour":1.2208,"AveExpr":-1.1425,"F":2.5693,"P.Value":0.088,"adj.P.Val":0.193},{"tissuedysplasia":0.1445,"tissuetumour":0.0816,"AveExpr":6.6502,"F":0.8147,"P.Value":0.4493,"adj.P.Val":0.5843},{"tissuedysplasia":0.0266,"tissuetumour":-0.2608,"AveExpr":6.251,"F":4.9445,"P.Value":0.0116,"adj.P.Val":0.0459},{"tissuedysplasia":0.9161,"tissuetumour":1.8663,"AveExpr":5.4687,"F":32.7436,"P.Value":1.948e-09,"adj.P.Val":4.0962e-07},{"tissuedysplasia":-0.0032,"tissuetumour":0.3805,"AveExpr":10.5236,"F":4.5987,"P.Value":0.0166,"adj.P.Val":0.06},{"tissuedysplasia":0.4545,"tissuetumour":0.4778,"AveExpr":3.4373,"F":4.752,"P.Value":0.0135,"adj.P.Val":0.0516},{"tissuedysplasia":-0.2233,"tissuetumour":-1.044,"AveExpr":6.6648,"F":25.597,"P.Value":4.2293e-08,"adj.P.Val":3.8473e-06},{"tissuedysplasia":-0.1969,"tissuetumour":-0.2835,"AveExpr":1.937,"F":0.1278,"P.Value":0.8804,"adj.P.Val":0.916},{"tissuedysplasia":0.4031,"tissuetumour":1.1945,"AveExpr":2.8674,"F":17.5732,"P.Value":2.4564e-06,"adj.P.Val":0.0001},{"tissuedysplasia":-0.0593,"tissuetumour":-0.4861,"AveExpr":6.9974,"F":9.0103,"P.Value":0.0005,"adj.P.Val":0.0048},{"tissuedysplasia":0.3058,"tissuetumour":0.2335,"AveExpr":7.7784,"F":2.8007,"P.Value":0.0725,"adj.P.Val":0.1691},{"tissuedysplasia":0.0057,"tissuetumour":0.0052,"AveExpr":4.8395,"F":0.0013,"P.Value":0.9987,"adj.P.Val":0.999},{"tissuedysplasia":0.3745,"tissuetumour":0.4427,"AveExpr":3.8834,"F":3.2778,"P.Value":0.0471,"adj.P.Val":0.1253},{"tissuedysplasia":0.2475,"tissuetumour":0.4743,"AveExpr":5.3246,"F":12.7254,"P.Value":0,"adj.P.Val":0.0007},{"tissuedysplasia":0.8657,"tissuetumour":0.4157,"AveExpr":3.0547,"F":5.4375,"P.Value":0.0078,"adj.P.Val":0.0346},{"tissuedysplasia":0.26,"tissuetumour":0.7368,"AveExpr":-0.5902,"F":2.2784,"P.Value":0.1144,"adj.P.Val":0.2311},{"tissuedysplasia":-0.2859,"tissuetumour":1.0976,"AveExpr":-2.119,"F":2.4162,"P.Value":0.101,"adj.P.Val":0.2125},{"tissuedysplasia":-0.1337,"tissuetumour":0.0564,"AveExpr":3.9122,"F":1.1988,"P.Value":0.3112,"adj.P.Val":0.4557},{"tissuedysplasia":0.3334,"tissuetumour":0.2798,"AveExpr":4.0329,"F":2.068,"P.Value":0.1386,"adj.P.Val":0.263},{"tissuedysplasia":-0.1081,"tissuetumour":-0.2705,"AveExpr":4.0105,"F":2.0639,"P.Value":0.1391,"adj.P.Val":0.2637},{"tissuedysplasia":0.5973,"tissuetumour":0.5378,"AveExpr":4.0839,"F":9.6504,"P.Value":0.0003,"adj.P.Val":0.0034},{"tissuedysplasia":-0.0901,"tissuetumour":-0.0054,"AveExpr":4.6655,"F":0.2334,"P.Value":0.7928,"adj.P.Val":0.8543},{"tissuedysplasia":0.0708,"tissuetumour":-0.0963,"AveExpr":3.8083,"F":0.7446,"P.Value":0.4808,"adj.P.Val":0.6102},{"tissuedysplasia":0.2721,"tissuetumour":0.598,"AveExpr":5.6131,"F":11.0886,"P.Value":0.0001,"adj.P.Val":0.0017},{"tissuedysplasia":0.0843,"tissuetumour":0.3267,"AveExpr":7.0333,"F":9.6579,"P.Value":0.0003,"adj.P.Val":0.0034},{"tissuedysplasia":-0.0436,"tissuetumour":0.4067,"AveExpr":4.9936,"F":7.8533,"P.Value":0.0012,"adj.P.Val":0.0089},{"tissuedysplasia":-0.1162,"tissuetumour":0.0431,"AveExpr":4.06,"F":0.9256,"P.Value":0.4039,"adj.P.Val":0.5438},{"tissuedysplasia":0.1868,"tissuetumour":0.3042,"AveExpr":3.4241,"F":2.3678,"P.Value":0.1055,"adj.P.Val":0.2186},{"tissuedysplasia":0.6253,"tissuetumour":0.1211,"AveExpr":1.4026,"F":2.322,"P.Value":0.11,"adj.P.Val":0.2249},{"tissuedysplasia":0.1104,"tissuetumour":-0.1048,"AveExpr":4.0182,"F":1.7595,"P.Value":0.184,"adj.P.Val":0.3193},{"tissuedysplasia":-0.6819,"tissuetumour":-0.4938,"AveExpr":1.5418,"F":4.6292,"P.Value":0.015,"adj.P.Val":0.0556},{"tissuedysplasia":0.0723,"tissuetumour":0.241,"AveExpr":3.1772,"F":0.548,"P.Value":0.582,"adj.P.Val":0.694},{"tissuedysplasia":-0.3619,"tissuetumour":-0.5986,"AveExpr":6.1225,"F":13.6261,"P.Value":0,"adj.P.Val":0.0005},{"tissuedysplasia":0.6023,"tissuetumour":0.6623,"AveExpr":3.8622,"F":9.541,"P.Value":0.0004,"adj.P.Val":0.0037},{"tissuedysplasia":0.0166,"tissuetumour":-0.1365,"AveExpr":5.9244,"F":0.9682,"P.Value":0.3877,"adj.P.Val":0.5288},{"tissuedysplasia":0.191,"tissuetumour":-0.465,"AveExpr":5.8069,"F":12.1193,"P.Value":0.0001,"adj.P.Val":0.001},{"tissuedysplasia":-0.2925,"tissuetumour":0.1311,"AveExpr":4.7609,"F":1.4594,"P.Value":0.2434,"adj.P.Val":0.3859},{"tissuedysplasia":0.2032,"tissuetumour":0.7794,"AveExpr":3.6495,"F":10.593,"P.Value":0.0002,"adj.P.Val":0.0021},{"tissuedysplasia":0.7374,"tissuetumour":0.7226,"AveExpr":5.0955,"F":12.4102,"P.Value":0.0001,"adj.P.Val":0.0009},{"tissuedysplasia":0.0562,"tissuetumour":0.0536,"AveExpr":3.5192,"F":0.1549,"P.Value":0.857,"adj.P.Val":0.8995},{"tissuedysplasia":0.8231,"tissuetumour":0.9499,"AveExpr":1.9687,"F":7.6143,"P.Value":0.0014,"adj.P.Val":0.0102},{"tissuedysplasia":-0.2557,"tissuetumour":-0.6676,"AveExpr":6.4148,"F":11.8603,"P.Value":0.0001,"adj.P.Val":0.0011},{"tissuedysplasia":0.3564,"tissuetumour":1.4155,"AveExpr":4.4604,"F":28.8581,"P.Value":9.8416e-09,"adj.P.Val":1.383e-06},{"tissuedysplasia":-0.6399,"tissuetumour":-0.1372,"AveExpr":1.9602,"F":3.4527,"P.Value":0.0405,"adj.P.Val":0.1125},{"tissuedysplasia":-0.3295,"tissuetumour":-1.1008,"AveExpr":5.0745,"F":21.3557,"P.Value":3.2966e-07,"adj.P.Val":0},{"tissuedysplasia":-0.2077,"tissuetumour":-0.0843,"AveExpr":2.9953,"F":0.4116,"P.Value":0.6651,"adj.P.Val":0.7591},{"tissuedysplasia":0.2253,"tissuetumour":0.6242,"AveExpr":2.2579,"F":4.2158,"P.Value":0.0211,"adj.P.Val":0.0712},{"tissuedysplasia":0.2574,"tissuetumour":0.572,"AveExpr":3.312,"F":6.3441,"P.Value":0.0038,"adj.P.Val":0.0206},{"tissuedysplasia":0.4602,"tissuetumour":1.2524,"AveExpr":5.8829,"F":16.2095,"P.Value":5.6609e-06,"adj.P.Val":0.0002},{"tissuedysplasia":0.1171,"tissuetumour":0.7784,"AveExpr":4.0788,"F":13.4611,"P.Value":0,"adj.P.Val":0.0005},{"tissuedysplasia":0.1248,"tissuetumour":0.6496,"AveExpr":4.3568,"F":9.8393,"P.Value":0.0003,"adj.P.Val":0.0031},{"tissuedysplasia":-3.4818,"tissuetumour":1.3177,"AveExpr":-5.8951,"F":6.342,"P.Value":0.0038,"adj.P.Val":0.0206},{"tissuedysplasia":0.4882,"tissuetumour":1.0108,"AveExpr":4.0881,"F":11.7901,"P.Value":0.0001,"adj.P.Val":0.0012},{"tissuedysplasia":-1.4586,"tissuetumour":-2.5745,"AveExpr":3.6317,"F":35.9749,"P.Value":5.5158e-10,"adj.P.Val":1.7085e-07},{"tissuedysplasia":1.2431,"tissuetumour":1.9503,"AveExpr":-4.316,"F":1.9903,"P.Value":0.1488,"adj.P.Val":0.2761},{"tissuedysplasia":-0.059,"tissuetumour":0.1619,"AveExpr":1.7066,"F":0.2951,"P.Value":0.7459,"adj.P.Val":0.8194},{"tissuedysplasia":-0.1902,"tissuetumour":-0.7307,"AveExpr":6.3715,"F":14.8663,"P.Value":0,"adj.P.Val":0.0003},{"tissuedysplasia":0.1357,"tissuetumour":-0.2063,"AveExpr":6.9901,"F":3.5788,"P.Value":0.0363,"adj.P.Val":0.1042},{"tissuedysplasia":0.6403,"tissuetumour":1.0301,"AveExpr":4.5617,"F":15.6377,"P.Value":7.4039e-06,"adj.P.Val":0.0002},{"tissuedysplasia":0.6159,"tissuetumour":0.8905,"AveExpr":0.8408,"F":3.0002,"P.Value":0.0601,"adj.P.Val":0.1484},{"tissuedysplasia":-0.0163,"tissuetumour":-0.2745,"AveExpr":6.251,"F":2.4101,"P.Value":0.1016,"adj.P.Val":0.2132},{"tissuedysplasia":0.275,"tissuetumour":0.328,"AveExpr":5.7403,"F":5.3073,"P.Value":0.0086,"adj.P.Val":0.0373},{"tissuedysplasia":-0.2047,"tissuetumour":-0.1703,"AveExpr":6.8947,"F":2.742,"P.Value":0.0755,"adj.P.Val":0.1735},{"tissuedysplasia":0.2975,"tissuetumour":0.6387,"AveExpr":5.955,"F":10.9768,"P.Value":0.0001,"adj.P.Val":0.0018},{"tissuedysplasia":0.1801,"tissuetumour":-0.051,"AveExpr":5.1585,"F":1.4978,"P.Value":0.2348,"adj.P.Val":0.3764},{"tissuedysplasia":0.3122,"tissuetumour":0.8031,"AveExpr":1.462,"F":2.749,"P.Value":0.075,"adj.P.Val":0.1728},{"tissuedysplasia":-0.5229,"tissuetumour":-1.0169,"AveExpr":1.2042,"F":2.3466,"P.Value":0.1076,"adj.P.Val":0.2216},{"tissuedysplasia":0.8194,"tissuetumour":1.8183,"AveExpr":-0.5936,"F":4.4865,"P.Value":0.0169,"adj.P.Val":0.0606},{"tissuedysplasia":0.1508,"tissuetumour":0.2418,"AveExpr":3.8377,"F":1.5721,"P.Value":0.2191,"adj.P.Val":0.359},{"tissuedysplasia":1.2483,"tissuetumour":3.3911,"AveExpr":-3.7715,"F":8.6812,"P.Value":0.0007,"adj.P.Val":0.0058},{"tissuedysplasia":0.3591,"tissuetumour":0.6109,"AveExpr":1.0882,"F":2.8132,"P.Value":0.0708,"adj.P.Val":0.1664},{"tissuedysplasia":-0.0363,"tissuetumour":0.1168,"AveExpr":5.5623,"F":0.747,"P.Value":0.4797,"adj.P.Val":0.6092},{"tissuedysplasia":0.8632,"tissuetumour":0.8451,"AveExpr":3.2366,"F":7.2019,"P.Value":0.002,"adj.P.Val":0.0128},{"tissuedysplasia":0.2167,"tissuetumour":0.5988,"AveExpr":4.3126,"F":5.6884,"P.Value":0.0063,"adj.P.Val":0.0299},{"tissuedysplasia":0.1823,"tissuetumour":-0.0354,"AveExpr":4.0751,"F":1.0293,"P.Value":0.3657,"adj.P.Val":0.5086},{"tissuedysplasia":-0.012,"tissuetumour":0.8637,"AveExpr":1.3223,"F":7.0055,"P.Value":0.0023,"adj.P.Val":0.0142},{"tissuedysplasia":0.1097,"tissuetumour":-0.009,"AveExpr":-1.8995,"F":0.0146,"P.Value":0.9855,"adj.P.Val":0.9899},{"tissuedysplasia":0.3486,"tissuetumour":0.5028,"AveExpr":3.8983,"F":6.9921,"P.Value":0.0023,"adj.P.Val":0.0143},{"tissuedysplasia":-2.4139,"tissuetumour":1.8168,"AveExpr":1.7427,"F":15.361,"P.Value":0,"adj.P.Val":0.0003},{"tissuedysplasia":0.0243,"tissuetumour":0.3324,"AveExpr":6.3691,"F":8.3621,"P.Value":0.0008,"adj.P.Val":0.0068},{"tissuedysplasia":0.476,"tissuetumour":0.0375,"AveExpr":2.8639,"F":4.8318,"P.Value":0.0127,"adj.P.Val":0.0491},{"tissuedysplasia":-0.0953,"tissuetumour":0.1557,"AveExpr":1.4265,"F":0.5078,"P.Value":0.6053,"adj.P.Val":0.7132},{"tissuedysplasia":-0.2297,"tissuetumour":-0.5838,"AveExpr":4.7331,"F":5.6847,"P.Value":0.0064,"adj.P.Val":0.03},{"tissuedysplasia":-0.3276,"tissuetumour":-0.5253,"AveExpr":4.491,"F":6.9554,"P.Value":0.0024,"adj.P.Val":0.0147},{"tissuedysplasia":0.1208,"tissuetumour":-0.1405,"AveExpr":4.8358,"F":1.2597,"P.Value":0.2938,"adj.P.Val":0.4383},{"tissuedysplasia":0.0563,"tissuetumour":0.3353,"AveExpr":5.2072,"F":2.1459,"P.Value":0.129,"adj.P.Val":0.2505},{"tissuedysplasia":0.8241,"tissuetumour":2.0118,"AveExpr":-2.3382,"F":5.0306,"P.Value":0.0108,"adj.P.Val":0.0437},{"tissuedysplasia":0.6228,"tissuetumour":0.7256,"AveExpr":0.7975,"F":3.1845,"P.Value":0.0511,"adj.P.Val":0.1326},{"tissuedysplasia":-0.1213,"tissuetumour":0.2495,"AveExpr":5.2006,"F":2.2959,"P.Value":0.1126,"adj.P.Val":0.2287},{"tissuedysplasia":0.6138,"tissuetumour":0.1213,"AveExpr":3.0721,"F":4.2803,"P.Value":0.02,"adj.P.Val":0.0686},{"tissuedysplasia":1.1489,"tissuetumour":0.7371,"AveExpr":1.0161,"F":7.6913,"P.Value":0.0014,"adj.P.Val":0.0098},{"tissuedysplasia":-0.815,"tissuetumour":-0.7149,"AveExpr":4.6986,"F":7.9256,"P.Value":0.0011,"adj.P.Val":0.0086},{"tissuedysplasia":0.102,"tissuetumour":-0.1907,"AveExpr":5.028,"F":3.0212,"P.Value":0.059,"adj.P.Val":0.1465},{"tissuedysplasia":0.409,"tissuetumour":0.9637,"AveExpr":1.4995,"F":3.4937,"P.Value":0.0391,"adj.P.Val":0.1097},{"tissuedysplasia":0.1859,"tissuetumour":0.293,"AveExpr":4.0813,"F":2.1779,"P.Value":0.1253,"adj.P.Val":0.2455},{"tissuedysplasia":0.1429,"tissuetumour":-0.4207,"AveExpr":4.4066,"F":7.4864,"P.Value":0.0016,"adj.P.Val":0.0109},{"tissuedysplasia":1.3057,"tissuetumour":1.302,"AveExpr":4.2415,"F":18.761,"P.Value":1.2815e-06,"adj.P.Val":0},{"tissuedysplasia":1.3039,"tissuetumour":2.4615,"AveExpr":3.2438,"F":21.1154,"P.Value":3.7254e-07,"adj.P.Val":0},{"tissuedysplasia":-0.5378,"tissuetumour":-1.4428,"AveExpr":6.299,"F":32.4362,"P.Value":2.205e-09,"adj.P.Val":4.5047e-07},{"tissuedysplasia":-0.0342,"tissuetumour":-0.2493,"AveExpr":4.8413,"F":1.8288,"P.Value":0.1726,"adj.P.Val":0.3055},{"tissuedysplasia":-0.0296,"tissuetumour":-0.1356,"AveExpr":6.1627,"F":0.635,"P.Value":0.5347,"adj.P.Val":0.6561},{"tissuedysplasia":0.1243,"tissuetumour":-0.1948,"AveExpr":5.4314,"F":4.5734,"P.Value":0.0157,"adj.P.Val":0.0575},{"tissuedysplasia":0.2264,"tissuetumour":-0.2884,"AveExpr":5.1851,"F":4.2725,"P.Value":0.0201,"adj.P.Val":0.0689},{"tissuedysplasia":0.5271,"tissuetumour":0.6055,"AveExpr":3.1597,"F":6.1526,"P.Value":0.0044,"adj.P.Val":0.0229},{"tissuedysplasia":-0.2723,"tissuetumour":-0.2482,"AveExpr":4.3516,"F":3.6313,"P.Value":0.0347,"adj.P.Val":0.1009},{"tissuedysplasia":0.1676,"tissuetumour":0.0583,"AveExpr":5.0667,"F":0.8007,"P.Value":0.4555,"adj.P.Val":0.5895},{"tissuedysplasia":-0.358,"tissuetumour":0.9905,"AveExpr":1.9606,"F":7.7139,"P.Value":0.0013,"adj.P.Val":0.0097},{"tissuedysplasia":0.2829,"tissuetumour":-0.0355,"AveExpr":6.4051,"F":7.4345,"P.Value":0.0017,"adj.P.Val":0.0112},{"tissuedysplasia":0.2426,"tissuetumour":0.3245,"AveExpr":2.8417,"F":1.8908,"P.Value":0.163,"adj.P.Val":0.2937},{"tissuedysplasia":-0.4881,"tissuetumour":1.4935,"AveExpr":0.3357,"F":8.894,"P.Value":0.0006,"adj.P.Val":0.0051},{"tissuedysplasia":0.2055,"tissuetumour":0.6034,"AveExpr":7.0162,"F":24.6514,"P.Value":6.5768e-08,"adj.P.Val":5.24e-06},{"tissuedysplasia":2.073,"tissuetumour":2.0892,"AveExpr":-3.0881,"F":2.7825,"P.Value":0.0728,"adj.P.Val":0.1695},{"tissuedysplasia":0.3319,"tissuetumour":-0.0492,"AveExpr":3.2821,"F":2.1629,"P.Value":0.1271,"adj.P.Val":0.2478},{"tissuedysplasia":-0.0796,"tissuetumour":-0.4971,"AveExpr":5.5666,"F":12.3138,"P.Value":0.0001,"adj.P.Val":0.0009},{"tissuedysplasia":0.0133,"tissuetumour":0.5203,"AveExpr":2.42,"F":3.4776,"P.Value":0.0396,"adj.P.Val":0.1108},{"tissuedysplasia":-0.6936,"tissuetumour":-0.6702,"AveExpr":1.3653,"F":2.8774,"P.Value":0.0669,"adj.P.Val":0.1597},{"tissuedysplasia":-0.2758,"tissuetumour":1.1617,"AveExpr":-3.6786,"F":1.333,"P.Value":0.2741,"adj.P.Val":0.4185},{"tissuedysplasia":0.7513,"tissuetumour":0.3585,"AveExpr":-0.2092,"F":1.2789,"P.Value":0.2885,"adj.P.Val":0.433},{"tissuedysplasia":-0.1683,"tissuetumour":-1.1596,"AveExpr":1.8294,"F":8.1139,"P.Value":0.001,"adj.P.Val":0.0077},{"tissuedysplasia":0.2246,"tissuetumour":0.0734,"AveExpr":6.525,"F":3.4884,"P.Value":0.0392,"adj.P.Val":0.1101},{"tissuedysplasia":-0.0411,"tissuetumour":0.0959,"AveExpr":4.5433,"F":0.3416,"P.Value":0.7125,"adj.P.Val":0.7941},{"tissuedysplasia":-0.0223,"tissuetumour":-0.4831,"AveExpr":6.9866,"F":8.277,"P.Value":0.0009,"adj.P.Val":0.0071},{"tissuedysplasia":0.1852,"tissuetumour":0.2212,"AveExpr":4.8152,"F":1.4747,"P.Value":0.2399,"adj.P.Val":0.382},{"tissuedysplasia":0.2779,"tissuetumour":0.1418,"AveExpr":4.8528,"F":2.608,"P.Value":0.085,"adj.P.Val":0.1884},{"tissuedysplasia":-1.2862,"tissuetumour":-0.4874,"AveExpr":-0.7151,"F":1.9966,"P.Value":0.1479,"adj.P.Val":0.275},{"tissuedysplasia":0.0345,"tissuetumour":-0.4741,"AveExpr":6.215,"F":8.0359,"P.Value":0.0011,"adj.P.Val":0.0081},{"tissuedysplasia":0.2199,"tissuetumour":0.0326,"AveExpr":2.011,"F":0.8612,"P.Value":0.4296,"adj.P.Val":0.567},{"tissuedysplasia":-0.2749,"tissuetumour":-0.5295,"AveExpr":6.085,"F":2.3766,"P.Value":0.1058,"adj.P.Val":0.219},{"tissuedysplasia":0.3056,"tissuetumour":1.3931,"AveExpr":5.04,"F":53.5674,"P.Value":1.6193e-12,"adj.P.Val":3.6063e-09},{"tissuedysplasia":0.1346,"tissuetumour":1.3398,"AveExpr":2.3795,"F":16.4109,"P.Value":4.7331e-06,"adj.P.Val":0.0001},{"tissuedysplasia":0.6842,"tissuetumour":3.9576,"AveExpr":-4.3417,"F":4.0217,"P.Value":0.0257,"adj.P.Val":0.0818},{"tissuedysplasia":0.4455,"tissuetumour":0.4825,"AveExpr":2.8089,"F":1.9187,"P.Value":0.1589,"adj.P.Val":0.2887},{"tissuedysplasia":-0.0194,"tissuetumour":-0.0153,"AveExpr":5.0315,"F":0.02,"P.Value":0.9802,"adj.P.Val":0.9863},{"tissuedysplasia":0.1561,"tissuetumour":-0.1573,"AveExpr":4.4516,"F":3.0866,"P.Value":0.0557,"adj.P.Val":0.1407},{"tissuedysplasia":-0.1168,"tissuetumour":1.2924,"AveExpr":-1.2129,"F":5.192,"P.Value":0.0095,"adj.P.Val":0.0398},{"tissuedysplasia":0.1159,"tissuetumour":0.3038,"AveExpr":4.0482,"F":0.9228,"P.Value":0.405,"adj.P.Val":0.5448},{"tissuedysplasia":0.5829,"tissuetumour":1.7306,"AveExpr":-0.7528,"F":5.5586,"P.Value":0.007,"adj.P.Val":0.0323},{"tissuedysplasia":0.1378,"tissuetumour":-0.1571,"AveExpr":5.0877,"F":2.3343,"P.Value":0.1088,"adj.P.Val":0.2232},{"tissuedysplasia":0.2676,"tissuetumour":0.5353,"AveExpr":0.824,"F":0.705,"P.Value":0.4996,"adj.P.Val":0.6263},{"tissuedysplasia":0.0712,"tissuetumour":0.0097,"AveExpr":3.4249,"F":0.0981,"P.Value":0.9068,"adj.P.Val":0.9351},{"tissuedysplasia":-0.1862,"tissuetumour":-0.4997,"AveExpr":4.8109,"F":6.4884,"P.Value":0.0034,"adj.P.Val":0.019},{"tissuedysplasia":0.1334,"tissuetumour":-0.2552,"AveExpr":6.4939,"F":5.0453,"P.Value":0.0106,"adj.P.Val":0.0434},{"tissuedysplasia":0.2143,"tissuetumour":-0.1704,"AveExpr":1.3679,"F":0.9018,"P.Value":0.4132,"adj.P.Val":0.5521},{"tissuedysplasia":1.1188,"tissuetumour":1.3988,"AveExpr":3.3448,"F":16.0399,"P.Value":5.8598e-06,"adj.P.Val":0.0002},{"tissuedysplasia":0.4819,"tissuetumour":0.5406,"AveExpr":5.164,"F":18.3764,"P.Value":1.5788e-06,"adj.P.Val":0.0001},{"tissuedysplasia":-0.3122,"tissuetumour":0.157,"AveExpr":6.9246,"F":3.2395,"P.Value":0.0496,"adj.P.Val":0.13},{"tissuedysplasia":0.2759,"tissuetumour":0.8112,"AveExpr":3.7932,"F":9.2222,"P.Value":0.0005,"adj.P.Val":0.0043},{"tissuedysplasia":0.1602,"tissuetumour":0.0048,"AveExpr":5.8979,"F":1.7149,"P.Value":0.1918,"adj.P.Val":0.328},{"tissuedysplasia":0.5799,"tissuetumour":0.2139,"AveExpr":3.171,"F":4.597,"P.Value":0.0154,"adj.P.Val":0.0566},{"tissuedysplasia":-0.1987,"tissuetumour":0.504,"AveExpr":5.2102,"F":12.1601,"P.Value":0.0001,"adj.P.Val":0.001},{"tissuedysplasia":-0.6103,"tissuetumour":-1.6328,"AveExpr":5.3231,"F":20.4849,"P.Value":5.1515e-07,"adj.P.Val":0},{"tissuedysplasia":0.296,"tissuetumour":0.1352,"AveExpr":4.1711,"F":2.5378,"P.Value":0.0906,"adj.P.Val":0.1969},{"tissuedysplasia":-0.5861,"tissuetumour":0.5393,"AveExpr":-4.366,"F":0.6654,"P.Value":0.5192,"adj.P.Val":0.643},{"tissuedysplasia":0.0991,"tissuetumour":0.9484,"AveExpr":5.4914,"F":14.7003,"P.Value":0,"adj.P.Val":0.0003},{"tissuedysplasia":0.3425,"tissuetumour":0.9497,"AveExpr":2.6087,"F":9.8307,"P.Value":0.0003,"adj.P.Val":0.0031},{"tissuedysplasia":0.2766,"tissuetumour":-0.0767,"AveExpr":5.3182,"F":2.6858,"P.Value":0.0793,"adj.P.Val":0.1797},{"tissuedysplasia":0.5275,"tissuetumour":1.1296,"AveExpr":0.1024,"F":3.0611,"P.Value":0.0569,"adj.P.Val":0.1429},{"tissuedysplasia":-0.162,"tissuetumour":-0.2463,"AveExpr":3.8058,"F":1.1182,"P.Value":0.336,"adj.P.Val":0.4803},{"tissuedysplasia":0.1262,"tissuetumour":1.9948,"AveExpr":-4.2588,"F":2.4846,"P.Value":0.095,"adj.P.Val":0.2036},{"tissuedysplasia":0.4553,"tissuetumour":0.8257,"AveExpr":4.2472,"F":17.8346,"P.Value":2.1252e-06,"adj.P.Val":0.0001},{"tissuedysplasia":-0.5667,"tissuetumour":2.2588,"AveExpr":-6.3107,"F":4.1434,"P.Value":0.0225,"adj.P.Val":0.0743},{"tissuedysplasia":0.3008,"tissuetumour":0.3031,"AveExpr":5.2765,"F":4.7778,"P.Value":0.0133,"adj.P.Val":0.0508},{"tissuedysplasia":-0.0603,"tissuetumour":0.4004,"AveExpr":3.6912,"F":2.9138,"P.Value":0.0648,"adj.P.Val":0.1563},{"tissuedysplasia":0.1136,"tissuetumour":-0.0969,"AveExpr":5.4742,"F":0.9361,"P.Value":0.3998,"adj.P.Val":0.5402},{"tissuedysplasia":0.041,"tissuetumour":0.2883,"AveExpr":6.3041,"F":6.7889,"P.Value":0.0027,"adj.P.Val":0.0161},{"tissuedysplasia":0.184,"tissuetumour":0.3069,"AveExpr":5.1262,"F":1.6946,"P.Value":0.1954,"adj.P.Val":0.332},{"tissuedysplasia":0.3532,"tissuetumour":1.7535,"AveExpr":0.2315,"F":9.484,"P.Value":0.0004,"adj.P.Val":0.0038},{"tissuedysplasia":0.335,"tissuetumour":-0.358,"AveExpr":3.8102,"F":3.8712,"P.Value":0.0283,"adj.P.Val":0.0875},{"tissuedysplasia":0.0221,"tissuetumour":-0.5887,"AveExpr":5.6203,"F":9.0673,"P.Value":0.0005,"adj.P.Val":0.0047},{"tissuedysplasia":0.2645,"tissuetumour":0.3628,"AveExpr":3.9075,"F":3.0529,"P.Value":0.0573,"adj.P.Val":0.1437},{"tissuedysplasia":0.332,"tissuetumour":0.5574,"AveExpr":3.8974,"F":7.2264,"P.Value":0.0019,"adj.P.Val":0.0126},{"tissuedysplasia":0.8167,"tissuetumour":1.0893,"AveExpr":0.4008,"F":4.1655,"P.Value":0.022,"adj.P.Val":0.0734},{"tissuedysplasia":3.0967,"tissuetumour":3.7519,"AveExpr":-0.1141,"F":14.5246,"P.Value":0,"adj.P.Val":0.0003},{"tissuedysplasia":-1.6609,"tissuetumour":-2.3338,"AveExpr":0.5106,"F":5.7792,"P.Value":0.006,"adj.P.Val":0.0286},{"tissuedysplasia":0.2503,"tissuetumour":0.5559,"AveExpr":4.1486,"F":9.4889,"P.Value":0.0004,"adj.P.Val":0.0038},{"tissuedysplasia":-0.0474,"tissuetumour":-0.7835,"AveExpr":6.8425,"F":18.0216,"P.Value":1.9172e-06,"adj.P.Val":0.0001},{"tissuedysplasia":-0.7683,"tissuetumour":0.4938,"AveExpr":-0.9679,"F":2.6191,"P.Value":0.0842,"adj.P.Val":0.1871},{"tissuedysplasia":0.4671,"tissuetumour":1.6133,"AveExpr":-1.7279,"F":3.1721,"P.Value":0.0517,"adj.P.Val":0.1336},{"tissuedysplasia":0.2141,"tissuetumour":0.1039,"AveExpr":3.6603,"F":0.7311,"P.Value":0.4871,"adj.P.Val":0.6159},{"tissuedysplasia":0.2661,"tissuetumour":0.0201,"AveExpr":6.8842,"F":3.7235,"P.Value":0.0321,"adj.P.Val":0.0955},{"tissuedysplasia":0.1664,"tissuetumour":-0.1795,"AveExpr":6.2498,"F":3.64,"P.Value":0.0344,"adj.P.Val":0.1003},{"tissuedysplasia":-0.371,"tissuetumour":0.2388,"AveExpr":0.3113,"F":0.7878,"P.Value":0.4612,"adj.P.Val":0.5943},{"tissuedysplasia":0.0521,"tissuetumour":-0.0295,"AveExpr":7.1051,"F":0.3074,"P.Value":0.7369,"adj.P.Val":0.8128},{"tissuedysplasia":1.3769,"tissuetumour":3.7202,"AveExpr":-4.7533,"F":6.4156,"P.Value":0.0036,"adj.P.Val":0.0198},{"tissuedysplasia":-0.1193,"tissuetumour":-0.7137,"AveExpr":3.9211,"F":9.0392,"P.Value":0.0005,"adj.P.Val":0.0047},{"tissuedysplasia":-0.1409,"tissuetumour":-0.5138,"AveExpr":3.1781,"F":4.4893,"P.Value":0.0168,"adj.P.Val":0.0604},{"tissuedysplasia":2.3879,"tissuetumour":3.4531,"AveExpr":-5.7986,"F":5.823,"P.Value":0.0057,"adj.P.Val":0.0277},{"tissuedysplasia":0.2345,"tissuetumour":2.006,"AveExpr":4.3607,"F":38.469,"P.Value":2.1834e-10,"adj.P.Val":8.7997e-08},{"tissuedysplasia":0.3292,"tissuetumour":0.7189,"AveExpr":7.0824,"F":12.9964,"P.Value":0,"adj.P.Val":0.0006},{"tissuedysplasia":0.0087,"tissuetumour":-0.2794,"AveExpr":4.9608,"F":1.3587,"P.Value":0.2676,"adj.P.Val":0.4112},{"tissuedysplasia":0.1763,"tissuetumour":1.7174,"AveExpr":-2.986,"F":2.3969,"P.Value":0.1028,"adj.P.Val":0.2149},{"tissuedysplasia":0.226,"tissuetumour":0.4857,"AveExpr":0.1684,"F":0.6556,"P.Value":0.5241,"adj.P.Val":0.6474},{"tissuedysplasia":0.355,"tissuetumour":1.2399,"AveExpr":-1.4208,"F":3.1762,"P.Value":0.0515,"adj.P.Val":0.1333},{"tissuedysplasia":0.1338,"tissuetumour":0.174,"AveExpr":5.4965,"F":1.9605,"P.Value":0.1529,"adj.P.Val":0.2813},{"tissuedysplasia":0.1733,"tissuetumour":-0.0274,"AveExpr":5.9192,"F":2.5131,"P.Value":0.0926,"adj.P.Val":0.1999},{"tissuedysplasia":0.4108,"tissuetumour":0.2927,"AveExpr":4.4933,"F":5.4493,"P.Value":0.0077,"adj.P.Val":0.0343},{"tissuedysplasia":0.0961,"tissuetumour":-0.5024,"AveExpr":6.0476,"F":10.8153,"P.Value":0.0002,"adj.P.Val":0.0019},{"tissuedysplasia":0.2218,"tissuetumour":0.5343,"AveExpr":3.3282,"F":4.9189,"P.Value":0.0118,"adj.P.Val":0.0466},{"tissuedysplasia":0.1986,"tissuetumour":0.1607,"AveExpr":8.8362,"F":0.591,"P.Value":0.559,"adj.P.Val":0.6754},{"tissuedysplasia":0.6786,"tissuetumour":0.5015,"AveExpr":4.1973,"F":12.462,"P.Value":0.0001,"adj.P.Val":0.0008},{"tissuedysplasia":0.192,"tissuetumour":0.8791,"AveExpr":5.506,"F":14.3496,"P.Value":0,"adj.P.Val":0.0003},{"tissuedysplasia":0.3384,"tissuetumour":0.8302,"AveExpr":0.6142,"F":2.2656,"P.Value":0.1157,"adj.P.Val":0.2329},{"tissuedysplasia":0.9386,"tissuetumour":1.4065,"AveExpr":-5.0555,"F":0.923,"P.Value":0.4049,"adj.P.Val":0.5446},{"tissuedysplasia":0.1845,"tissuetumour":0.5584,"AveExpr":3.1754,"F":3.8996,"P.Value":0.0276,"adj.P.Val":0.086},{"tissuedysplasia":0.6458,"tissuetumour":0.8145,"AveExpr":2.5326,"F":7.5332,"P.Value":0.0015,"adj.P.Val":0.0106},{"tissuedysplasia":0.129,"tissuetumour":0.0223,"AveExpr":4.8911,"F":0.6016,"P.Value":0.5524,"adj.P.Val":0.6697},{"tissuedysplasia":0.0312,"tissuetumour":-0.2246,"AveExpr":4.5897,"F":3.2362,"P.Value":0.0488,"adj.P.Val":0.1286},{"tissuedysplasia":-0.6231,"tissuetumour":-1.2829,"AveExpr":5.1393,"F":25.8506,"P.Value":3.7627e-08,"adj.P.Val":3.5741e-06},{"tissuedysplasia":0.4213,"tissuetumour":0.6888,"AveExpr":2.083,"F":1.9831,"P.Value":0.1498,"adj.P.Val":0.2772},{"tissuedysplasia":-0.4046,"tissuetumour":2.9706,"AveExpr":-4.0622,"F":13.9305,"P.Value":0,"adj.P.Val":0.0004},{"tissuedysplasia":0.8368,"tissuetumour":1.0692,"AveExpr":2.5977,"F":8.7482,"P.Value":0.0006,"adj.P.Val":0.0055},{"tissuedysplasia":0.0724,"tissuetumour":-0.1421,"AveExpr":4.7146,"F":0.7856,"P.Value":0.4621,"adj.P.Val":0.5951},{"tissuedysplasia":0.356,"tissuetumour":0.4187,"AveExpr":4.0477,"F":4.5744,"P.Value":0.0157,"adj.P.Val":0.0574},{"tissuedysplasia":0.292,"tissuetumour":1.4291,"AveExpr":0.4276,"F":6.5173,"P.Value":0.0033,"adj.P.Val":0.0187},{"tissuedysplasia":0.3497,"tissuetumour":1.2274,"AveExpr":0.7932,"F":6.7484,"P.Value":0.0028,"adj.P.Val":0.0164},{"tissuedysplasia":0.1619,"tissuetumour":-0.3299,"AveExpr":5.1125,"F":5.2733,"P.Value":0.0089,"adj.P.Val":0.038},{"tissuedysplasia":0.0241,"tissuetumour":0.0322,"AveExpr":3.7836,"F":0.0255,"P.Value":0.9748,"adj.P.Val":0.9826},{"tissuedysplasia":-0.1304,"tissuetumour":-0.4347,"AveExpr":4.0861,"F":5.0976,"P.Value":0.0102,"adj.P.Val":0.042},{"tissuedysplasia":0.1415,"tissuetumour":0.634,"AveExpr":4.4692,"F":3.363,"P.Value":0.0437,"adj.P.Val":0.1188},{"tissuedysplasia":0.9754,"tissuetumour":0.2384,"AveExpr":6.1489,"F":9.2874,"P.Value":0.0005,"adj.P.Val":0.0047},{"tissuedysplasia":-0.8568,"tissuetumour":2.4856,"AveExpr":-0.7013,"F":12.3251,"P.Value":0.0001,"adj.P.Val":0.0009},{"tissuedysplasia":-0.0811,"tissuetumour":-0.1133,"AveExpr":9.5849,"F":0.3136,"P.Value":0.7328,"adj.P.Val":0.8095},{"tissuedysplasia":0.2687,"tissuetumour":0.2219,"AveExpr":4.744,"F":2.5576,"P.Value":0.089,"adj.P.Val":0.1944},{"tissuedysplasia":0.7231,"tissuetumour":0.3035,"AveExpr":5.8988,"F":9.6906,"P.Value":0.0003,"adj.P.Val":0.0034},{"tissuedysplasia":0.111,"tissuetumour":0.0686,"AveExpr":3.7613,"F":0.3006,"P.Value":0.7419,"adj.P.Val":0.8163},{"tissuedysplasia":0.1861,"tissuetumour":-0.1095,"AveExpr":3.3177,"F":1.5516,"P.Value":0.2233,"adj.P.Val":0.3636},{"tissuedysplasia":0.5073,"tissuetumour":2.1579,"AveExpr":0.8162,"F":11.2196,"P.Value":0.0001,"adj.P.Val":0.0016},{"tissuedysplasia":-0.1982,"tissuetumour":-0.1293,"AveExpr":4.9816,"F":1.008,"P.Value":0.3732,"adj.P.Val":0.5158},{"tissuedysplasia":0.5097,"tissuetumour":1.1386,"AveExpr":1.9706,"F":13.2532,"P.Value":0,"adj.P.Val":0.0006},{"tissuedysplasia":0.2003,"tissuetumour":0.1441,"AveExpr":1.9067,"F":0.5787,"P.Value":0.5648,"adj.P.Val":0.6797},{"tissuedysplasia":0.0104,"tissuetumour":0.1066,"AveExpr":5.6814,"F":0.5257,"P.Value":0.5948,"adj.P.Val":0.7044},{"tissuedysplasia":0.0354,"tissuetumour":0.0093,"AveExpr":4.4207,"F":0.0529,"P.Value":0.9486,"adj.P.Val":0.9645},{"tissuedysplasia":0.7178,"tissuetumour":2.5223,"AveExpr":-1.9991,"F":7.0118,"P.Value":0.0023,"adj.P.Val":0.0142},{"tissuedysplasia":0.5234,"tissuetumour":0.7747,"AveExpr":3.3221,"F":10.1634,"P.Value":0.0002,"adj.P.Val":0.0027},{"tissuedysplasia":-0.0156,"tissuetumour":0.8779,"AveExpr":-0.9799,"F":1.4297,"P.Value":0.2503,"adj.P.Val":0.3931},{"tissuedysplasia":0.423,"tissuetumour":1.0688,"AveExpr":-0.5725,"F":1.7992,"P.Value":0.1774,"adj.P.Val":0.3114},{"tissuedysplasia":0.1348,"tissuetumour":0.4796,"AveExpr":4.702,"F":6.247,"P.Value":0.0041,"adj.P.Val":0.0217},{"tissuedysplasia":0.235,"tissuetumour":-0.0133,"AveExpr":3.073,"F":1.3245,"P.Value":0.2763,"adj.P.Val":0.4208},{"tissuedysplasia":-0.127,"tissuetumour":-0.432,"AveExpr":4.3686,"F":4.9261,"P.Value":0.0117,"adj.P.Val":0.0464},{"tissuedysplasia":0.1429,"tissuetumour":-0.3955,"AveExpr":5.5452,"F":9.0603,"P.Value":0.0005,"adj.P.Val":0.0047},{"tissuedysplasia":1.1425,"tissuetumour":1.4535,"AveExpr":3.0016,"F":11.5328,"P.Value":0.0001,"adj.P.Val":0.0013},{"tissuedysplasia":-0.2075,"tissuetumour":0.5777,"AveExpr":2.5938,"F":6.0394,"P.Value":0.0048,"adj.P.Val":0.0244},{"tissuedysplasia":0.261,"tissuetumour":0.1517,"AveExpr":4.3652,"F":2.3793,"P.Value":0.1044,"adj.P.Val":0.2172},{"tissuedysplasia":1.0838,"tissuetumour":0.9339,"AveExpr":5.4548,"F":13.8797,"P.Value":0,"adj.P.Val":0.0004},{"tissuedysplasia":0.785,"tissuetumour":1.1517,"AveExpr":-0.831,"F":2.4269,"P.Value":0.1,"adj.P.Val":0.211},{"tissuedysplasia":-0.2658,"tissuetumour":-0.3536,"AveExpr":2.2536,"F":1.6907,"P.Value":0.1961,"adj.P.Val":0.3328},{"tissuedysplasia":0.1603,"tissuetumour":-0.3566,"AveExpr":5.9928,"F":5.7661,"P.Value":0.006,"adj.P.Val":0.0286},{"tissuedysplasia":-0.3783,"tissuetumour":0.4406,"AveExpr":1.2971,"F":2.0565,"P.Value":0.14,"adj.P.Val":0.2648},{"tissuedysplasia":0.1745,"tissuetumour":0.0307,"AveExpr":2.1948,"F":0.3218,"P.Value":0.7265,"adj.P.Val":0.8046},{"tissuedysplasia":0.5292,"tissuetumour":0.3413,"AveExpr":4.314,"F":5.2141,"P.Value":0.0093,"adj.P.Val":0.0393},{"tissuedysplasia":0.4261,"tissuetumour":0.2208,"AveExpr":4.1944,"F":3.037,"P.Value":0.0581,"adj.P.Val":0.1452},{"tissuedysplasia":0.2629,"tissuetumour":-0.2946,"AveExpr":5.4039,"F":5.4071,"P.Value":0.0079,"adj.P.Val":0.0351},{"tissuedysplasia":-0.4561,"tissuetumour":-1.4767,"AveExpr":6.528,"F":25.5122,"P.Value":5.4024e-08,"adj.P.Val":4.5638e-06},{"tissuedysplasia":0.5077,"tissuetumour":1.5854,"AveExpr":0.1781,"F":5.1652,"P.Value":0.0097,"adj.P.Val":0.0404},{"tissuedysplasia":-0.5204,"tissuetumour":-0.6115,"AveExpr":5.1475,"F":7.696,"P.Value":0.0014,"adj.P.Val":0.0097},{"tissuedysplasia":-0.1488,"tissuetumour":-0.9678,"AveExpr":6.7792,"F":24.1442,"P.Value":8.365e-08,"adj.P.Val":6.2723e-06},{"tissuedysplasia":-0.2124,"tissuetumour":-0.4101,"AveExpr":4.595,"F":4.9775,"P.Value":0.0113,"adj.P.Val":0.045},{"tissuedysplasia":0.1401,"tissuetumour":1.2702,"AveExpr":-0.0102,"F":4.7912,"P.Value":0.0131,"adj.P.Val":0.0504},{"tissuedysplasia":-0.0477,"tissuetumour":-0.7144,"AveExpr":3.4763,"F":5.6236,"P.Value":0.0067,"adj.P.Val":0.0311},{"tissuedysplasia":0.763,"tissuetumour":0.641,"AveExpr":4.2473,"F":8.6337,"P.Value":0.0007,"adj.P.Val":0.0059},{"tissuedysplasia":0.2393,"tissuetumour":0.2879,"AveExpr":2.3636,"F":1.2157,"P.Value":0.3063,"adj.P.Val":0.4508},{"tissuedysplasia":-0.0554,"tissuetumour":-0.0414,"AveExpr":3.8951,"F":0.1066,"P.Value":0.8992,"adj.P.Val":0.9297},{"tissuedysplasia":0.591,"tissuetumour":1.6186,"AveExpr":-1.228,"F":4.6466,"P.Value":0.0148,"adj.P.Val":0.055},{"tissuedysplasia":0.4104,"tissuetumour":1.4702,"AveExpr":7.1989,"F":45.9067,"P.Value":2.3924e-11,"adj.P.Val":1.76e-08},{"tissuedysplasia":0.7153,"tissuetumour":1.5885,"AveExpr":-0.1159,"F":7.3179,"P.Value":0.0018,"adj.P.Val":0.012},{"tissuedysplasia":0.4241,"tissuetumour":0.1108,"AveExpr":6.2055,"F":10.431,"P.Value":0.0002,"adj.P.Val":0.0023},{"tissuedysplasia":0.1284,"tissuetumour":0.4883,"AveExpr":5.7826,"F":9.1526,"P.Value":0.0005,"adj.P.Val":0.0045},{"tissuedysplasia":0.1785,"tissuetumour":0.3696,"AveExpr":7.1225,"F":9.2703,"P.Value":0.0004,"adj.P.Val":0.0042},{"tissuedysplasia":0.254,"tissuetumour":0.1181,"AveExpr":6.6916,"F":3.7732,"P.Value":0.0307,"adj.P.Val":0.0928},{"tissuedysplasia":0.2657,"tissuetumour":0.9264,"AveExpr":-0.6987,"F":2.2691,"P.Value":0.1154,"adj.P.Val":0.2324},{"tissuedysplasia":0.5742,"tissuetumour":-0.0228,"AveExpr":4.8699,"F":7.4433,"P.Value":0.0016,"adj.P.Val":0.0112},{"tissuedysplasia":0.0108,"tissuetumour":0.8157,"AveExpr":1.8601,"F":3.3974,"P.Value":0.0425,"adj.P.Val":0.1164},{"tissuedysplasia":-0.4304,"tissuetumour":-3.5818,"AveExpr":6.0353,"F":6.2497,"P.Value":0.0047,"adj.P.Val":0.0239},{"tissuedysplasia":0.038,"tissuetumour":-0.5279,"AveExpr":5.5031,"F":11.0312,"P.Value":0.0001,"adj.P.Val":0.0017},{"tissuedysplasia":0.1571,"tissuetumour":0.1511,"AveExpr":4.3565,"F":1.3988,"P.Value":0.2576,"adj.P.Val":0.4007},{"tissuedysplasia":-0.3123,"tissuetumour":-0.573,"AveExpr":3.0743,"F":5.1896,"P.Value":0.0095,"adj.P.Val":0.0398},{"tissuedysplasia":0.0523,"tissuetumour":0.1541,"AveExpr":7.0159,"F":0.7372,"P.Value":0.4843,"adj.P.Val":0.6131},{"tissuedysplasia":-0.0647,"tissuetumour":-0.078,"AveExpr":7.132,"F":0.2152,"P.Value":0.8072,"adj.P.Val":0.8647},{"tissuedysplasia":0.0236,"tissuetumour":0.2411,"AveExpr":4.0009,"F":1.5864,"P.Value":0.2162,"adj.P.Val":0.3556},{"tissuedysplasia":-0.0587,"tissuetumour":-0.4556,"AveExpr":5.8729,"F":8.4954,"P.Value":0.0008,"adj.P.Val":0.0063},{"tissuedysplasia":0.2041,"tissuetumour":-0.1747,"AveExpr":6.3532,"F":4.8186,"P.Value":0.0128,"adj.P.Val":0.0495},{"tissuedysplasia":0.4985,"tissuetumour":1.828,"AveExpr":4.6206,"F":52.646,"P.Value":2.121e-12,"adj.P.Val":4.2448e-09},{"tissuedysplasia":1.229,"tissuetumour":1.4374,"AveExpr":1.3743,"F":7.9612,"P.Value":0.0011,"adj.P.Val":0.0084},{"tissuedysplasia":0.9203,"tissuetumour":1.236,"AveExpr":3.1842,"F":22.5898,"P.Value":1.7778e-07,"adj.P.Val":0},{"tissuedysplasia":0.457,"tissuetumour":0.3194,"AveExpr":5.7431,"F":8.4095,"P.Value":0.0008,"adj.P.Val":0.0066},{"tissuedysplasia":0.2031,"tissuetumour":-0.2025,"AveExpr":3.9195,"F":2.0549,"P.Value":0.1402,"adj.P.Val":0.265},{"tissuedysplasia":-0.2868,"tissuetumour":-0.7133,"AveExpr":4.1079,"F":9.3738,"P.Value":0.0004,"adj.P.Val":0.004},{"tissuedysplasia":-0.6835,"tissuetumour":1.2994,"AveExpr":-7.9315,"F":1.6133,"P.Value":0.2108,"adj.P.Val":0.3496},{"tissuedysplasia":0.1631,"tissuetumour":0.3715,"AveExpr":2.559,"F":1.7619,"P.Value":0.1836,"adj.P.Val":0.3188},{"tissuedysplasia":0.0688,"tissuetumour":-0.0515,"AveExpr":5.4543,"F":0.434,"P.Value":0.6506,"adj.P.Val":0.748},{"tissuedysplasia":-0.0803,"tissuetumour":-0.7252,"AveExpr":4.7024,"F":16.5617,"P.Value":4.3422e-06,"adj.P.Val":0.0001},{"tissuedysplasia":0.2914,"tissuetumour":0.2946,"AveExpr":2.8394,"F":2.5853,"P.Value":0.0868,"adj.P.Val":0.1911},{"tissuedysplasia":0.9756,"tissuetumour":3.2084,"AveExpr":-4.4022,"F":7.0652,"P.Value":0.0022,"adj.P.Val":0.0138},{"tissuedysplasia":0.2923,"tissuetumour":0.0205,"AveExpr":6.3581,"F":3.9952,"P.Value":0.0254,"adj.P.Val":0.0813},{"tissuedysplasia":-0.0398,"tissuetumour":0.4747,"AveExpr":1.9498,"F":2.7369,"P.Value":0.0758,"adj.P.Val":0.174},{"tissuedysplasia":0.1442,"tissuetumour":0.2785,"AveExpr":3.6555,"F":2.3932,"P.Value":0.1031,"adj.P.Val":0.2155},{"tissuedysplasia":0.2582,"tissuetumour":0.6226,"AveExpr":2.2734,"F":4.211,"P.Value":0.0212,"adj.P.Val":0.0715},{"tissuedysplasia":-0.0118,"tissuetumour":-0.2837,"AveExpr":1.7274,"F":0.8132,"P.Value":0.45,"adj.P.Val":0.5848},{"tissuedysplasia":0.1371,"tissuetumour":0.1252,"AveExpr":4.0173,"F":0.731,"P.Value":0.4872,"adj.P.Val":0.6159},{"tissuedysplasia":0.2907,"tissuetumour":0.4024,"AveExpr":2.5831,"F":1.3961,"P.Value":0.2583,"adj.P.Val":0.4014},{"tissuedysplasia":-0.0574,"tissuetumour":-0.2747,"AveExpr":5.7595,"F":3.3264,"P.Value":0.0452,"adj.P.Val":0.1215},{"tissuedysplasia":-0.0241,"tissuetumour":-0.1287,"AveExpr":6.2183,"F":0.5601,"P.Value":0.5751,"adj.P.Val":0.6884},{"tissuedysplasia":1.0262,"tissuetumour":1.3012,"AveExpr":7.7559,"F":16.4817,"P.Value":7.7946e-06,"adj.P.Val":0.0002},{"tissuedysplasia":-0.1499,"tissuetumour":-0.2014,"AveExpr":6.4752,"F":3.172,"P.Value":0.0517,"adj.P.Val":0.1336},{"tissuedysplasia":-0.2295,"tissuetumour":-0.8171,"AveExpr":4.0956,"F":10.7365,"P.Value":0.0002,"adj.P.Val":0.002},{"tissuedysplasia":0.7267,"tissuetumour":0.505,"AveExpr":5.8082,"F":9.8756,"P.Value":0.0003,"adj.P.Val":0.0031},{"tissuedysplasia":0.2598,"tissuetumour":0.7699,"AveExpr":3.644,"F":10.0855,"P.Value":0.0002,"adj.P.Val":0.0028},{"tissuedysplasia":0.3646,"tissuetumour":0.1875,"AveExpr":5.1844,"F":1.936,"P.Value":0.1564,"adj.P.Val":0.2856},{"tissuedysplasia":0.3388,"tissuetumour":0.2845,"AveExpr":6.065,"F":7.3397,"P.Value":0.0018,"adj.P.Val":0.0119},{"tissuedysplasia":0.0744,"tissuetumour":-0.0158,"AveExpr":3.2747,"F":0.2953,"P.Value":0.7458,"adj.P.Val":0.8192},{"tissuedysplasia":0.0567,"tissuetumour":-0.2457,"AveExpr":5.9225,"F":4.1641,"P.Value":0.0221,"adj.P.Val":0.0735},{"tissuedysplasia":-0.1954,"tissuetumour":-0.4301,"AveExpr":2.4549,"F":0.69,"P.Value":0.5069,"adj.P.Val":0.6325},{"tissuedysplasia":0.3367,"tissuetumour":0.6483,"AveExpr":5.9213,"F":22.4868,"P.Value":1.8706e-07,"adj.P.Val":0},{"tissuedysplasia":0.5031,"tissuetumour":0.3786,"AveExpr":6.8264,"F":8.4956,"P.Value":0.0008,"adj.P.Val":0.0063},{"tissuedysplasia":0.04,"tissuetumour":0.5896,"AveExpr":7.3991,"F":12.815,"P.Value":0,"adj.P.Val":0.0007},{"tissuedysplasia":0.2769,"tissuetumour":0.5088,"AveExpr":3.0796,"F":3.7777,"P.Value":0.0306,"adj.P.Val":0.0925},{"tissuedysplasia":1.8882,"tissuetumour":2.344,"AveExpr":-0.0302,"F":9.6731,"P.Value":0.0003,"adj.P.Val":0.0034},{"tissuedysplasia":0.2159,"tissuetumour":0.3098,"AveExpr":5.3739,"F":7.6962,"P.Value":0.0014,"adj.P.Val":0.0097},{"tissuedysplasia":0.0748,"tissuetumour":0.4308,"AveExpr":5.0487,"F":5.2575,"P.Value":0.009,"adj.P.Val":0.0383},{"tissuedysplasia":-0.3353,"tissuetumour":1.0412,"AveExpr":-1.5783,"F":2.9488,"P.Value":0.0628,"adj.P.Val":0.1531},{"tissuedysplasia":0.5901,"tissuetumour":2.0679,"AveExpr":5.1621,"F":47.7693,"P.Value":9.3783e-12,"adj.P.Val":1.1108e-08},{"tissuedysplasia":0.1474,"tissuetumour":0.4071,"AveExpr":5.8897,"F":6.9298,"P.Value":0.0024,"adj.P.Val":0.0149},{"tissuedysplasia":0.7195,"tissuetumour":2.4997,"AveExpr":0.3393,"F":18.6902,"P.Value":1.3315e-06,"adj.P.Val":0},{"tissuedysplasia":-0.3108,"tissuetumour":-1.0832,"AveExpr":6.2306,"F":41.9655,"P.Value":6.3388e-11,"adj.P.Val":3.6326e-08},{"tissuedysplasia":0.0691,"tissuetumour":0.0543,"AveExpr":4.158,"F":0.2045,"P.Value":0.8159,"adj.P.Val":0.8708},{"tissuedysplasia":-0.8351,"tissuetumour":-0.1087,"AveExpr":2.7853,"F":0.8087,"P.Value":0.453,"adj.P.Val":0.5874},{"tissuedysplasia":0.6821,"tissuetumour":1.8058,"AveExpr":2.2739,"F":22.1652,"P.Value":2.1944e-07,"adj.P.Val":0},{"tissuedysplasia":-0.3246,"tissuetumour":-1.153,"AveExpr":7.6759,"F":25.7433,"P.Value":6.5103e-08,"adj.P.Val":5.2115e-06},{"tissuedysplasia":-0.224,"tissuetumour":0.1377,"AveExpr":-0.352,"F":0.2691,"P.Value":0.7653,"adj.P.Val":0.8336},{"tissuedysplasia":-0.0859,"tissuetumour":-0.0321,"AveExpr":6.151,"F":0.4958,"P.Value":0.6125,"adj.P.Val":0.7189},{"tissuedysplasia":0.3699,"tissuetumour":0.706,"AveExpr":2.6204,"F":4.9075,"P.Value":0.0119,"adj.P.Val":0.0469},{"tissuedysplasia":0.3426,"tissuetumour":0.8666,"AveExpr":5.0219,"F":21.0944,"P.Value":3.7655e-07,"adj.P.Val":0},{"tissuedysplasia":-0.2447,"tissuetumour":-0.2857,"AveExpr":6.7864,"F":1.8301,"P.Value":0.1724,"adj.P.Val":0.3053},{"tissuedysplasia":0.3499,"tissuetumour":1.3005,"AveExpr":6.0261,"F":24.7947,"P.Value":6.1477e-08,"adj.P.Val":5.0232e-06},{"tissuedysplasia":0.0623,"tissuetumour":0.7143,"AveExpr":3.5595,"F":11.4192,"P.Value":0.0001,"adj.P.Val":0.0014},{"tissuedysplasia":0.1654,"tissuetumour":-0.0876,"AveExpr":7.0909,"F":1.9338,"P.Value":0.1567,"adj.P.Val":0.2859},{"tissuedysplasia":0.2411,"tissuetumour":-0.0606,"AveExpr":2.6209,"F":1.1598,"P.Value":0.3229,"adj.P.Val":0.4671},{"tissuedysplasia":0.3931,"tissuetumour":1.1353,"AveExpr":1.2434,"F":4.8524,"P.Value":0.0125,"adj.P.Val":0.0485},{"tissuedysplasia":0.4186,"tissuetumour":-0.2096,"AveExpr":3.7919,"F":5.9762,"P.Value":0.0051,"adj.P.Val":0.0253},{"tissuedysplasia":0.3318,"tissuetumour":0.0343,"AveExpr":3.6629,"F":1.1597,"P.Value":0.323,"adj.P.Val":0.4672},{"tissuedysplasia":0.0472,"tissuetumour":-0.6083,"AveExpr":4.3781,"F":6.9058,"P.Value":0.0025,"adj.P.Val":0.0151},{"tissuedysplasia":0.3093,"tissuetumour":0.1435,"AveExpr":4.437,"F":3.1904,"P.Value":0.0508,"adj.P.Val":0.1321},{"tissuedysplasia":-0.333,"tissuetumour":-0.7226,"AveExpr":4.5188,"F":12.2131,"P.Value":0.0001,"adj.P.Val":0.0009},{"tissuedysplasia":0.0334,"tissuetumour":-0.0874,"AveExpr":5.4481,"F":0.4223,"P.Value":0.6582,"adj.P.Val":0.7538},{"tissuedysplasia":-0.145,"tissuetumour":-0.8366,"AveExpr":4.8536,"F":16.5863,"P.Value":4.2816e-06,"adj.P.Val":0.0001},{"tissuedysplasia":0.3051,"tissuetumour":0.0875,"AveExpr":5.354,"F":2.279,"P.Value":0.1143,"adj.P.Val":0.231},{"tissuedysplasia":-0.0533,"tissuetumour":0.3332,"AveExpr":0.6769,"F":1.0024,"P.Value":0.3752,"adj.P.Val":0.5176},{"tissuedysplasia":0.3723,"tissuetumour":0.5737,"AveExpr":4.2803,"F":6.3558,"P.Value":0.0038,"adj.P.Val":0.0205},{"tissuedysplasia":-0.1055,"tissuetumour":-0.3921,"AveExpr":5.7849,"F":6.8205,"P.Value":0.0026,"adj.P.Val":0.0158},{"tissuedysplasia":0.4694,"tissuetumour":0.8612,"AveExpr":2.5216,"F":8.6494,"P.Value":0.0007,"adj.P.Val":0.0059},{"tissuedysplasia":0.3039,"tissuetumour":0.3324,"AveExpr":3.4932,"F":1.2571,"P.Value":0.2945,"adj.P.Val":0.4389},{"tissuedysplasia":0.5107,"tissuetumour":0.2498,"AveExpr":1.8753,"F":1.6688,"P.Value":0.2002,"adj.P.Val":0.3372},{"tissuedysplasia":0.3814,"tissuetumour":0.9849,"AveExpr":2.335,"F":9.1501,"P.Value":0.0005,"adj.P.Val":0.0045},{"tissuedysplasia":0.7809,"tissuetumour":1.5381,"AveExpr":3.437,"F":29.9994,"P.Value":6.0395e-09,"adj.P.Val":9.7095e-07},{"tissuedysplasia":-0.2247,"tissuetumour":-0.9216,"AveExpr":6.3232,"F":16.072,"P.Value":5.7522e-06,"adj.P.Val":0.0002},{"tissuedysplasia":0.2546,"tissuetumour":0.1489,"AveExpr":4.5902,"F":2.2203,"P.Value":0.1206,"adj.P.Val":0.2392},{"tissuedysplasia":0.9056,"tissuetumour":1.2327,"AveExpr":7.848,"F":9.642,"P.Value":0.0004,"adj.P.Val":0.0042},{"tissuedysplasia":1.1264,"tissuetumour":1.5777,"AveExpr":-1.943,"F":1.8717,"P.Value":0.1659,"adj.P.Val":0.2972},{"tissuedysplasia":0.1846,"tissuetumour":0.2738,"AveExpr":4.8352,"F":1.5545,"P.Value":0.2227,"adj.P.Val":0.363},{"tissuedysplasia":0.117,"tissuetumour":0.0628,"AveExpr":5.0268,"F":0.4938,"P.Value":0.6136,"adj.P.Val":0.7197},{"tissuedysplasia":-0.0576,"tissuetumour":-0.2947,"AveExpr":4.049,"F":1.7565,"P.Value":0.1845,"adj.P.Val":0.3198},{"tissuedysplasia":0.1325,"tissuetumour":-0.1921,"AveExpr":6.4317,"F":4.7141,"P.Value":0.014,"adj.P.Val":0.0528},{"tissuedysplasia":0.8935,"tissuetumour":1.3682,"AveExpr":2.0263,"F":17.8514,"P.Value":2.1055e-06,"adj.P.Val":0.0001},{"tissuedysplasia":-0.3015,"tissuetumour":-0.8003,"AveExpr":3.1782,"F":10.7307,"P.Value":0.0002,"adj.P.Val":0.002},{"tissuedysplasia":0.2189,"tissuetumour":0.6032,"AveExpr":4.1657,"F":7.547,"P.Value":0.0015,"adj.P.Val":0.0105},{"tissuedysplasia":0.2049,"tissuetumour":0.3557,"AveExpr":4.8705,"F":6.2516,"P.Value":0.0041,"adj.P.Val":0.0217},{"tissuedysplasia":0.18,"tissuetumour":0.1888,"AveExpr":5.1811,"F":2.3358,"P.Value":0.1086,"adj.P.Val":0.2231},{"tissuedysplasia":0.3237,"tissuetumour":0.7656,"AveExpr":3.7415,"F":13.6626,"P.Value":0,"adj.P.Val":0.0005},{"tissuedysplasia":0.6119,"tissuetumour":1.086,"AveExpr":2.792,"F":20.4496,"P.Value":5.2465e-07,"adj.P.Val":0},{"tissuedysplasia":0.1313,"tissuetumour":0.6237,"AveExpr":4.5766,"F":10.9958,"P.Value":0.0001,"adj.P.Val":0.0017},{"tissuedysplasia":0.0103,"tissuetumour":-0.0384,"AveExpr":5.752,"F":0.0369,"P.Value":0.9638,"adj.P.Val":0.9753},{"tissuedysplasia":0.1196,"tissuetumour":0.2641,"AveExpr":6.3366,"F":2.2396,"P.Value":0.1185,"adj.P.Val":0.2366},{"tissuedysplasia":0.0202,"tissuetumour":-0.185,"AveExpr":4.441,"F":0.7667,"P.Value":0.4707,"adj.P.Val":0.6012},{"tissuedysplasia":0.0183,"tissuetumour":-0.0453,"AveExpr":5.7252,"F":0.2277,"P.Value":0.7973,"adj.P.Val":0.8577},{"tissuedysplasia":0.9301,"tissuetumour":2.1394,"AveExpr":-1.4307,"F":9.3648,"P.Value":0.0004,"adj.P.Val":0.004},{"tissuedysplasia":-0.0038,"tissuetumour":-0.3571,"AveExpr":3.8126,"F":2.2787,"P.Value":0.1144,"adj.P.Val":0.231},{"tissuedysplasia":0.0049,"tissuetumour":-0.4405,"AveExpr":4.021,"F":6.3972,"P.Value":0.0036,"adj.P.Val":0.02},{"tissuedysplasia":0.7804,"tissuetumour":1.2903,"AveExpr":-0.2309,"F":4.1911,"P.Value":0.0216,"adj.P.Val":0.0723},{"tissuedysplasia":0.4026,"tissuetumour":0.4207,"AveExpr":5.8626,"F":8.7586,"P.Value":0.0006,"adj.P.Val":0.0055},{"tissuedysplasia":1.6727,"tissuetumour":2.223,"AveExpr":-2.2663,"F":5.3626,"P.Value":0.0082,"adj.P.Val":0.0361},{"tissuedysplasia":0.4548,"tissuetumour":1.366,"AveExpr":0.5153,"F":4.51,"P.Value":0.0165,"adj.P.Val":0.0597},{"tissuedysplasia":1.4486,"tissuetumour":1.615,"AveExpr":2.3815,"F":16.4065,"P.Value":4.7449e-06,"adj.P.Val":0.0001},{"tissuedysplasia":0.1038,"tissuetumour":-0.1059,"AveExpr":3.8727,"F":1.3813,"P.Value":0.2619,"adj.P.Val":0.4051},{"tissuedysplasia":0.2222,"tissuetumour":0.883,"AveExpr":2.6805,"F":8.9164,"P.Value":0.0006,"adj.P.Val":0.0051},{"tissuedysplasia":-2.161,"tissuetumour":-1.367,"AveExpr":-9.6283,"F":1.6354,"P.Value":0.2065,"adj.P.Val":0.3446},{"tissuedysplasia":-0.2089,"tissuetumour":-0.1608,"AveExpr":5.7336,"F":0.9155,"P.Value":0.4078,"adj.P.Val":0.5472},{"tissuedysplasia":-0.2804,"tissuetumour":-1.2762,"AveExpr":5.7268,"F":7.5742,"P.Value":0.0017,"adj.P.Val":0.0114},{"tissuedysplasia":0.062,"tissuetumour":-0.2117,"AveExpr":8.2951,"F":2.4226,"P.Value":0.1018,"adj.P.Val":0.2135},{"tissuedysplasia":0.239,"tissuetumour":0.2897,"AveExpr":4.7014,"F":5.1303,"P.Value":0.0099,"adj.P.Val":0.0412},{"tissuedysplasia":0.1381,"tissuetumour":-0.1273,"AveExpr":6.8066,"F":2.0698,"P.Value":0.1383,"adj.P.Val":0.2628},{"tissuedysplasia":0.5711,"tissuetumour":-0.1418,"AveExpr":3.2426,"F":6.9019,"P.Value":0.0025,"adj.P.Val":0.0151},{"tissuedysplasia":0.0247,"tissuetumour":-0.3576,"AveExpr":5.0025,"F":2.9591,"P.Value":0.0623,"adj.P.Val":0.1521},{"tissuedysplasia":0.5899,"tissuetumour":0.7135,"AveExpr":2.1189,"F":3.2642,"P.Value":0.0477,"adj.P.Val":0.1263},{"tissuedysplasia":0.214,"tissuetumour":-0.3789,"AveExpr":4.0065,"F":5.8488,"P.Value":0.0056,"adj.P.Val":0.0273},{"tissuedysplasia":0.0423,"tissuetumour":2.3447,"AveExpr":-1.434,"F":16.1998,"P.Value":5.3432e-06,"adj.P.Val":0.0001},{"tissuedysplasia":0.551,"tissuetumour":0.9919,"AveExpr":2.5021,"F":10.1557,"P.Value":0.0002,"adj.P.Val":0.0027},{"tissuedysplasia":0.0469,"tissuetumour":0.0701,"AveExpr":8.6424,"F":0.6505,"P.Value":0.5267,"adj.P.Val":0.6497},{"tissuedysplasia":0.1802,"tissuetumour":0.1748,"AveExpr":3.705,"F":0.6132,"P.Value":0.5462,"adj.P.Val":0.6649},{"tissuedysplasia":-0.2742,"tissuetumour":-0.3479,"AveExpr":4.0759,"F":3.4375,"P.Value":0.041,"adj.P.Val":0.1136},{"tissuedysplasia":-1.9223,"tissuetumour":2.4068,"AveExpr":0.0671,"F":15.0192,"P.Value":0,"adj.P.Val":0.0003},{"tissuedysplasia":0.9964,"tissuetumour":2.1728,"AveExpr":1.4535,"F":22.9407,"P.Value":1.4962e-07,"adj.P.Val":9.4798e-06},{"tissuedysplasia":-0.505,"tissuetumour":-0.6707,"AveExpr":3.8489,"F":10.5979,"P.Value":0.0002,"adj.P.Val":0.0021},{"tissuedysplasia":0.0979,"tissuetumour":-0.0063,"AveExpr":4.4052,"F":0.3221,"P.Value":0.7263,"adj.P.Val":0.8045},{"tissuedysplasia":-0.0952,"tissuetumour":0.0186,"AveExpr":6.8252,"F":0.329,"P.Value":0.7214,"adj.P.Val":0.8008},{"tissuedysplasia":0.7418,"tissuetumour":1.4676,"AveExpr":-0.8154,"F":4.2902,"P.Value":0.0199,"adj.P.Val":0.0682},{"tissuedysplasia":0.1596,"tissuetumour":0.5575,"AveExpr":4.3611,"F":8.0805,"P.Value":0.001,"adj.P.Val":0.0079},{"tissuedysplasia":0.3488,"tissuetumour":2.0648,"AveExpr":-0.287,"F":12.2708,"P.Value":0.0001,"adj.P.Val":0.0009},{"tissuedysplasia":0.6788,"tissuetumour":1.3288,"AveExpr":6.3892,"F":40.9514,"P.Value":9.0095e-11,"adj.P.Val":4.7535e-08},{"tissuedysplasia":0.1204,"tissuetumour":0.7057,"AveExpr":5.7214,"F":17.6178,"P.Value":2.3963e-06,"adj.P.Val":0.0001},{"tissuedysplasia":0.2967,"tissuetumour":1.0693,"AveExpr":2.981,"F":17.096,"P.Value":3.2079e-06,"adj.P.Val":0.0001},{"tissuedysplasia":0.2607,"tissuetumour":0.3204,"AveExpr":4.3407,"F":3.2079,"P.Value":0.0501,"adj.P.Val":0.1308},{"tissuedysplasia":0.095,"tissuetumour":0.5344,"AveExpr":4.1587,"F":9.8414,"P.Value":0.0003,"adj.P.Val":0.0031},{"tissuedysplasia":1.1482,"tissuetumour":2.5809,"AveExpr":5.1129,"F":34.9364,"P.Value":1.3035e-09,"adj.P.Val":2.9666e-07},{"tissuedysplasia":-0.7802,"tissuetumour":-2.4918,"AveExpr":5.2876,"F":13.076,"P.Value":0.0001,"adj.P.Val":0.0008},{"tissuedysplasia":0.0122,"tissuetumour":0.2901,"AveExpr":4.8687,"F":4.1008,"P.Value":0.0233,"adj.P.Val":0.0762},{"tissuedysplasia":0.5825,"tissuetumour":1.215,"AveExpr":3.8482,"F":30.3802,"P.Value":5.1438e-09,"adj.P.Val":8.6006e-07},{"tissuedysplasia":0.1946,"tissuetumour":0.272,"AveExpr":5.4225,"F":3.8374,"P.Value":0.0291,"adj.P.Val":0.0892},{"tissuedysplasia":0.153,"tissuetumour":0.2375,"AveExpr":4.4468,"F":2.4757,"P.Value":0.0957,"adj.P.Val":0.2047},{"tissuedysplasia":1.4523,"tissuetumour":2.6781,"AveExpr":-1.8786,"F":8.5296,"P.Value":0.0007,"adj.P.Val":0.0062},{"tissuedysplasia":0.5065,"tissuetumour":1.6444,"AveExpr":6.5886,"F":20.8189,"P.Value":7.914e-07,"adj.P.Val":0},{"tissuedysplasia":0.9108,"tissuetumour":1.5903,"AveExpr":-6.7762,"F":0.817,"P.Value":0.4483,"adj.P.Val":0.5835},{"tissuedysplasia":0.3592,"tissuetumour":0.3576,"AveExpr":4.2084,"F":2.6161,"P.Value":0.0844,"adj.P.Val":0.1876},{"tissuedysplasia":0.4594,"tissuetumour":0.9808,"AveExpr":2.6,"F":7.7196,"P.Value":0.0013,"adj.P.Val":0.0096},{"tissuedysplasia":-0.2905,"tissuetumour":0.1224,"AveExpr":4.9077,"F":7.8022,"P.Value":0.0013,"adj.P.Val":0.0092},{"tissuedysplasia":0.2594,"tissuetumour":0.8093,"AveExpr":5.9202,"F":15.6117,"P.Value":7.5172e-06,"adj.P.Val":0.0002},{"tissuedysplasia":0.0664,"tissuetumour":0.495,"AveExpr":5.5176,"F":8.488,"P.Value":0.0008,"adj.P.Val":0.0064},{"tissuedysplasia":-0.0332,"tissuetumour":0.2282,"AveExpr":6.8607,"F":9.3739,"P.Value":0.0004,"adj.P.Val":0.004},{"tissuedysplasia":-0.0162,"tissuetumour":-0.2578,"AveExpr":3.5514,"F":1.8746,"P.Value":0.1655,"adj.P.Val":0.2967},{"tissuedysplasia":0.0415,"tissuetumour":-0.435,"AveExpr":2.4217,"F":2.4368,"P.Value":0.0992,"adj.P.Val":0.2097},{"tissuedysplasia":0.1741,"tissuetumour":-0.1341,"AveExpr":5.0355,"F":1.954,"P.Value":0.1538,"adj.P.Val":0.2824},{"tissuedysplasia":0.7451,"tissuetumour":1.8454,"AveExpr":3.3455,"F":24.03,"P.Value":8.8338e-08,"adj.P.Val":6.531e-06},{"tissuedysplasia":0.2623,"tissuetumour":-0.0565,"AveExpr":3.8885,"F":2.8129,"P.Value":0.0709,"adj.P.Val":0.1664},{"tissuedysplasia":0.4221,"tissuetumour":0.9913,"AveExpr":6.6694,"F":47.1289,"P.Value":1.1488e-11,"adj.P.Val":1.258e-08},{"tissuedysplasia":0.403,"tissuetumour":0.4152,"AveExpr":4.6748,"F":6.5484,"P.Value":0.0032,"adj.P.Val":0.0184},{"tissuedysplasia":0.0483,"tissuetumour":0.3196,"AveExpr":7.2533,"F":10.6986,"P.Value":0.0002,"adj.P.Val":0.002},{"tissuedysplasia":0.2893,"tissuetumour":0.037,"AveExpr":5.824,"F":2.5886,"P.Value":0.0865,"adj.P.Val":0.1907},{"tissuedysplasia":1.7804,"tissuetumour":3.2979,"AveExpr":4.6728,"F":27.1469,"P.Value":4.4749e-08,"adj.P.Val":4.0107e-06},{"tissuedysplasia":0.9995,"tissuetumour":2.6945,"AveExpr":0.3484,"F":19.0654,"P.Value":1.088e-06,"adj.P.Val":0},{"tissuedysplasia":0.1969,"tissuetumour":-0.2397,"AveExpr":4.999,"F":3.0007,"P.Value":0.06,"adj.P.Val":0.1484},{"tissuedysplasia":0.0071,"tissuetumour":-0.2505,"AveExpr":5.2667,"F":2.3674,"P.Value":0.1056,"adj.P.Val":0.2187},{"tissuedysplasia":0.7596,"tissuetumour":1.3892,"AveExpr":2.9153,"F":22.3387,"P.Value":2.013e-07,"adj.P.Val":0},{"tissuedysplasia":0.61,"tissuetumour":0.506,"AveExpr":1.9194,"F":3.6885,"P.Value":0.033,"adj.P.Val":0.0975},{"tissuedysplasia":0.4265,"tissuetumour":0.1281,"AveExpr":2.2728,"F":1.935,"P.Value":0.1565,"adj.P.Val":0.2857},{"tissuedysplasia":0.0408,"tissuetumour":0.166,"AveExpr":1.6507,"F":0.3447,"P.Value":0.7103,"adj.P.Val":0.7926},{"tissuedysplasia":-0.1105,"tissuetumour":-0.0919,"AveExpr":3.5982,"F":0.2772,"P.Value":0.7592,"adj.P.Val":0.8292},{"tissuedysplasia":0.1092,"tissuetumour":0.2423,"AveExpr":3.3222,"F":1.3635,"P.Value":0.2664,"adj.P.Val":0.4097},{"tissuedysplasia":-0.5863,"tissuetumour":-0.9075,"AveExpr":5.3576,"F":8.3968,"P.Value":0.0008,"adj.P.Val":0.0067},{"tissuedysplasia":0.1989,"tissuetumour":-0.3593,"AveExpr":3.1044,"F":5.8816,"P.Value":0.0054,"adj.P.Val":0.0267},{"tissuedysplasia":0.1699,"tissuetumour":0.7001,"AveExpr":5.0383,"F":15.265,"P.Value":9.216e-06,"adj.P.Val":0.0002},{"tissuedysplasia":-0.0562,"tissuetumour":0.1219,"AveExpr":1.3219,"F":0.2808,"P.Value":0.7566,"adj.P.Val":0.8272},{"tissuedysplasia":0.2913,"tissuetumour":-0.0597,"AveExpr":4.8,"F":3.8467,"P.Value":0.0289,"adj.P.Val":0.0888},{"tissuedysplasia":0.3028,"tissuetumour":0.1728,"AveExpr":5.6538,"F":3.9265,"P.Value":0.027,"adj.P.Val":0.0846},{"tissuedysplasia":0.1651,"tissuetumour":0.5019,"AveExpr":6.3797,"F":10.0859,"P.Value":0.0002,"adj.P.Val":0.0028},{"tissuedysplasia":-0.0328,"tissuetumour":0.3738,"AveExpr":4.7833,"F":7.0646,"P.Value":0.0022,"adj.P.Val":0.0138},{"tissuedysplasia":-0.1724,"tissuetumour":0.3386,"AveExpr":4.0835,"F":5.7519,"P.Value":0.006,"adj.P.Val":0.0288},{"tissuedysplasia":0.5462,"tissuetumour":1.0524,"AveExpr":2.9848,"F":18.7239,"P.Value":1.3075e-06,"adj.P.Val":0},{"tissuedysplasia":1.3116,"tissuetumour":1.6626,"AveExpr":2.0778,"F":21.941,"P.Value":2.4543e-07,"adj.P.Val":0},{"tissuedysplasia":0.628,"tissuetumour":1.2066,"AveExpr":-1.3403,"F":2.3735,"P.Value":0.105,"adj.P.Val":0.218},{"tissuedysplasia":0.7228,"tissuetumour":1.0985,"AveExpr":4.8453,"F":24.1845,"P.Value":8.2058e-08,"adj.P.Val":6.193e-06},{"tissuedysplasia":1.1695,"tissuetumour":1.7446,"AveExpr":3.2621,"F":9.257,"P.Value":0.0004,"adj.P.Val":0.0043},{"tissuedysplasia":0.45,"tissuetumour":0.6594,"AveExpr":1.7131,"F":4.191,"P.Value":0.0216,"adj.P.Val":0.0723},{"tissuedysplasia":0.3413,"tissuetumour":-0.0754,"AveExpr":5.2196,"F":1.8182,"P.Value":0.1745,"adj.P.Val":0.3078},{"tissuedysplasia":0.6463,"tissuetumour":1.1821,"AveExpr":2.6035,"F":6.025,"P.Value":0.0049,"adj.P.Val":0.0246},{"tissuedysplasia":-0.5342,"tissuetumour":-1.0096,"AveExpr":7.547,"F":45.9138,"P.Value":1.697e-11,"adj.P.Val":1.5633e-08},{"tissuedysplasia":0.3526,"tissuetumour":0.4292,"AveExpr":5.2003,"F":5.3276,"P.Value":0.0085,"adj.P.Val":0.0369},{"tissuedysplasia":0.1171,"tissuetumour":0.041,"AveExpr":1.6681,"F":0.0999,"P.Value":0.9052,"adj.P.Val":0.9339},{"tissuedysplasia":-0.7872,"tissuetumour":-0.6226,"AveExpr":2.3371,"F":5.0552,"P.Value":0.0106,"adj.P.Val":0.0431},{"tissuedysplasia":0.5859,"tissuetumour":1.0881,"AveExpr":0.2441,"F":2.8726,"P.Value":0.0672,"adj.P.Val":0.1602},{"tissuedysplasia":-0.0623,"tissuetumour":0.018,"AveExpr":4.0557,"F":0.2074,"P.Value":0.8135,"adj.P.Val":0.8692},{"tissuedysplasia":-0.5972,"tissuetumour":-1.0249,"AveExpr":4.6565,"F":15.2313,"P.Value":9.4011e-06,"adj.P.Val":0.0002},{"tissuedysplasia":0.1342,"tissuetumour":0.3566,"AveExpr":1.91,"F":0.9801,"P.Value":0.3833,"adj.P.Val":0.525},{"tissuedysplasia":-0.1262,"tissuetumour":0.1845,"AveExpr":2.3384,"F":0.4245,"P.Value":0.6567,"adj.P.Val":0.7528},{"tissuedysplasia":0.2619,"tissuetumour":0.1777,"AveExpr":3.5784,"F":1.4454,"P.Value":0.2466,"adj.P.Val":0.3893},{"tissuedysplasia":2.231,"tissuetumour":3.1063,"AveExpr":-8.2834,"F":3.6932,"P.Value":0.0329,"adj.P.Val":0.0973},{"tissuedysplasia":0.0194,"tissuetumour":-0.2816,"AveExpr":4.9613,"F":4.9451,"P.Value":0.0116,"adj.P.Val":0.0458},{"tissuedysplasia":0.2921,"tissuetumour":0.4303,"AveExpr":5.7802,"F":4.6992,"P.Value":0.0141,"adj.P.Val":0.0533},{"tissuedysplasia":0.6083,"tissuetumour":1.117,"AveExpr":2.6666,"F":6.8191,"P.Value":0.0026,"adj.P.Val":0.0158},{"tissuedysplasia":0.5553,"tissuetumour":1.5396,"AveExpr":2.9525,"F":18.2404,"P.Value":1.7004e-06,"adj.P.Val":0.0001},{"tissuedysplasia":0.2555,"tissuetumour":0.231,"AveExpr":6.1617,"F":1.7868,"P.Value":0.1794,"adj.P.Val":0.3139},{"tissuedysplasia":0.1842,"tissuetumour":0.2895,"AveExpr":5.1741,"F":4.3338,"P.Value":0.0191,"adj.P.Val":0.0665},{"tissuedysplasia":0.0281,"tissuetumour":0.3692,"AveExpr":3.0886,"F":1.475,"P.Value":0.2399,"adj.P.Val":0.3819},{"tissuedysplasia":0.1937,"tissuetumour":0.2918,"AveExpr":5.3384,"F":3.0062,"P.Value":0.0597,"adj.P.Val":0.1479},{"tissuedysplasia":0.2051,"tissuetumour":0.2428,"AveExpr":3.5951,"F":1.726,"P.Value":0.1898,"adj.P.Val":0.3258},{"tissuedysplasia":0.2315,"tissuetumour":0.7553,"AveExpr":1.2946,"F":3.573,"P.Value":0.0365,"adj.P.Val":0.1045},{"tissuedysplasia":1.6381,"tissuetumour":3.2787,"AveExpr":0.9435,"F":20.83,"P.Value":4.3114e-07,"adj.P.Val":0},{"tissuedysplasia":0.0642,"tissuetumour":0.32,"AveExpr":6.9329,"F":4.8255,"P.Value":0.0127,"adj.P.Val":0.0493},{"tissuedysplasia":0.5067,"tissuetumour":1.5222,"AveExpr":1.2835,"F":14.1098,"P.Value":0,"adj.P.Val":0.0004},{"tissuedysplasia":0.6031,"tissuetumour":2.2005,"AveExpr":-3.2299,"F":4.6279,"P.Value":0.015,"adj.P.Val":0.0556},{"tissuedysplasia":0.397,"tissuetumour":0.2233,"AveExpr":5.5266,"F":3.8098,"P.Value":0.0298,"adj.P.Val":0.0909},{"tissuedysplasia":0.3822,"tissuetumour":0.0932,"AveExpr":3.632,"F":3.3168,"P.Value":0.0455,"adj.P.Val":0.1223},{"tissuedysplasia":-0.2157,"tissuetumour":-0.5832,"AveExpr":6.6092,"F":10.5271,"P.Value":0.0002,"adj.P.Val":0.0022},{"tissuedysplasia":0.1335,"tissuetumour":1.6761,"AveExpr":-1.7902,"F":5.6754,"P.Value":0.0064,"adj.P.Val":0.0302},{"tissuedysplasia":0.4174,"tissuetumour":0.3536,"AveExpr":6.062,"F":6.1782,"P.Value":0.0043,"adj.P.Val":0.0226},{"tissuedysplasia":-0.0233,"tissuetumour":-0.2328,"AveExpr":6.7661,"F":2.3563,"P.Value":0.1066,"adj.P.Val":0.2203},{"tissuedysplasia":1.6209,"tissuetumour":2.0013,"AveExpr":0.5121,"F":17.7858,"P.Value":2.1833e-06,"adj.P.Val":0.0001},{"tissuedysplasia":0.1753,"tissuetumour":0.0885,"AveExpr":5.3153,"F":1.5803,"P.Value":0.2174,"adj.P.Val":0.3571},{"tissuedysplasia":0.2236,"tissuetumour":0.3623,"AveExpr":3.1308,"F":1.7964,"P.Value":0.1778,"adj.P.Val":0.312},{"tissuedysplasia":0.2065,"tissuetumour":0.1713,"AveExpr":4.8274,"F":1.0464,"P.Value":0.3598,"adj.P.Val":0.5031},{"tissuedysplasia":-0.0191,"tissuetumour":-0.3155,"AveExpr":3.5268,"F":2.8994,"P.Value":0.0656,"adj.P.Val":0.1576},{"tissuedysplasia":0.2499,"tissuetumour":0.2299,"AveExpr":6.4482,"F":7.3652,"P.Value":0.0017,"adj.P.Val":0.0117},{"tissuedysplasia":0.2682,"tissuetumour":0.2194,"AveExpr":4.7188,"F":3.563,"P.Value":0.0368,"adj.P.Val":0.1052},{"tissuedysplasia":0.3794,"tissuetumour":0.665,"AveExpr":4.3796,"F":11.7844,"P.Value":0.0001,"adj.P.Val":0.0012},{"tissuedysplasia":0.5437,"tissuetumour":0.519,"AveExpr":4.0965,"F":6.3325,"P.Value":0.0038,"adj.P.Val":0.0207},{"tissuedysplasia":0.2999,"tissuetumour":0.5701,"AveExpr":1.4318,"F":1.6794,"P.Value":0.1982,"adj.P.Val":0.3351},{"tissuedysplasia":0.2033,"tissuetumour":0.4364,"AveExpr":5.1523,"F":7.1705,"P.Value":0.002,"adj.P.Val":0.013},{"tissuedysplasia":0.419,"tissuetumour":1.8735,"AveExpr":-0.6296,"F":8.6779,"P.Value":0.0007,"adj.P.Val":0.0058},{"tissuedysplasia":1.6988,"tissuetumour":3.3454,"AveExpr":-2.8794,"F":7.4886,"P.Value":0.0016,"adj.P.Val":0.0109},{"tissuedysplasia":0.4872,"tissuetumour":0.4212,"AveExpr":3.2449,"F":3.8675,"P.Value":0.0284,"adj.P.Val":0.0877},{"tissuedysplasia":0.7351,"tissuetumour":1.1535,"AveExpr":4.9814,"F":13.9279,"P.Value":0,"adj.P.Val":0.0004},{"tissuedysplasia":0.1086,"tissuetumour":0.5497,"AveExpr":6.4202,"F":21.3585,"P.Value":3.2919e-07,"adj.P.Val":0},{"tissuedysplasia":-0.1048,"tissuetumour":-0.0122,"AveExpr":7.6672,"F":0.5761,"P.Value":0.5663,"adj.P.Val":0.681},{"tissuedysplasia":0.4672,"tissuetumour":1.4901,"AveExpr":5.5256,"F":31.2062,"P.Value":3.6456e-09,"adj.P.Val":6.6745e-07},{"tissuedysplasia":0.893,"tissuetumour":0.949,"AveExpr":-3.4635,"F":0.779,"P.Value":0.4651,"adj.P.Val":0.5975},{"tissuedysplasia":0.9895,"tissuetumour":2.6424,"AveExpr":-0.2682,"F":16.6271,"P.Value":4.1834e-06,"adj.P.Val":0.0001},{"tissuedysplasia":0.3312,"tissuetumour":0.3582,"AveExpr":5.4111,"F":5.2239,"P.Value":0.0092,"adj.P.Val":0.039},{"tissuedysplasia":0.3014,"tissuetumour":0.1675,"AveExpr":3.8761,"F":1.1838,"P.Value":0.3157,"adj.P.Val":0.4601},{"tissuedysplasia":-0.2749,"tissuetumour":2.8425,"AveExpr":-0.1624,"F":13.992,"P.Value":0,"adj.P.Val":0.0004},{"tissuedysplasia":-0.3813,"tissuetumour":-0.5104,"AveExpr":6.0229,"F":16.6579,"P.Value":4.1106e-06,"adj.P.Val":0.0001},{"tissuedysplasia":0.9989,"tissuetumour":2.5385,"AveExpr":-1.362,"F":10.069,"P.Value":0.0003,"adj.P.Val":0.0028},{"tissuedysplasia":0.2441,"tissuetumour":0.0629,"AveExpr":4.1745,"F":1.3875,"P.Value":0.2604,"adj.P.Val":0.4035},{"tissuedysplasia":0.1884,"tissuetumour":0.5952,"AveExpr":5.2969,"F":18.9301,"P.Value":1.1699e-06,"adj.P.Val":0},{"tissuedysplasia":-0.1063,"tissuetumour":0.4342,"AveExpr":3.4305,"F":1.306,"P.Value":0.2812,"adj.P.Val":0.4258},{"tissuedysplasia":0.2298,"tissuetumour":0.3592,"AveExpr":7.4242,"F":6.1449,"P.Value":0.0044,"adj.P.Val":0.023},{"tissuedysplasia":0.0073,"tissuetumour":-0.3044,"AveExpr":3.8131,"F":2.8275,"P.Value":0.0699,"adj.P.Val":0.1648},{"tissuedysplasia":1.0031,"tissuetumour":1.5259,"AveExpr":4.8094,"F":35.6185,"P.Value":6.3173e-10,"adj.P.Val":1.8749e-07},{"tissuedysplasia":0.1778,"tissuetumour":0.1733,"AveExpr":3.6405,"F":0.8713,"P.Value":0.4255,"adj.P.Val":0.5632},{"tissuedysplasia":0.5754,"tissuetumour":1.4243,"AveExpr":3.7948,"F":23.2439,"P.Value":1.2904e-07,"adj.P.Val":8.5788e-06},{"tissuedysplasia":0.5068,"tissuetumour":0.579,"AveExpr":5.2143,"F":7.8595,"P.Value":0.0012,"adj.P.Val":0.0089},{"tissuedysplasia":0.142,"tissuetumour":0.4798,"AveExpr":2.2869,"F":2.1618,"P.Value":0.1272,"adj.P.Val":0.2479},{"tissuedysplasia":-0.0705,"tissuetumour":0.0503,"AveExpr":3.3666,"F":0.3047,"P.Value":0.7389,"adj.P.Val":0.8142},{"tissuedysplasia":0.8893,"tissuetumour":1.4092,"AveExpr":0.3999,"F":8.4903,"P.Value":0.0008,"adj.P.Val":0.0064},{"tissuedysplasia":1.187,"tissuetumour":1.8467,"AveExpr":-2.6798,"F":3.3747,"P.Value":0.0433,"adj.P.Val":0.118},{"tissuedysplasia":0.3353,"tissuetumour":0.8193,"AveExpr":6.4951,"F":27.2861,"P.Value":1.9637e-08,"adj.P.Val":2.2478e-06},{"tissuedysplasia":0.7415,"tissuetumour":1.333,"AveExpr":3.5284,"F":20.2376,"P.Value":5.8574e-07,"adj.P.Val":0},{"tissuedysplasia":0.2373,"tissuetumour":0.1334,"AveExpr":4.5082,"F":1.2694,"P.Value":0.2911,"adj.P.Val":0.4356},{"tissuedysplasia":0.7772,"tissuetumour":1.3222,"AveExpr":3.4542,"F":25.747,"P.Value":3.9464e-08,"adj.P.Val":3.6823e-06},{"tissuedysplasia":0.2208,"tissuetumour":1.0638,"AveExpr":1.0249,"F":5.4363,"P.Value":0.0078,"adj.P.Val":0.0346},{"tissuedysplasia":0.0667,"tissuetumour":0.2604,"AveExpr":5.0783,"F":2.2898,"P.Value":0.1132,"adj.P.Val":0.2295},{"tissuedysplasia":0.0593,"tissuetumour":-0.2553,"AveExpr":2.2919,"F":0.8479,"P.Value":0.4352,"adj.P.Val":0.572},{"tissuedysplasia":0.1462,"tissuetumour":0.2415,"AveExpr":6.3789,"F":3.0951,"P.Value":0.0552,"adj.P.Val":0.14},{"tissuedysplasia":0.4593,"tissuetumour":0.4147,"AveExpr":4.7774,"F":8.4408,"P.Value":0.0008,"adj.P.Val":0.0065},{"tissuedysplasia":0.1108,"tissuetumour":0.0196,"AveExpr":4.3478,"F":0.6789,"P.Value":0.5124,"adj.P.Val":0.6373},{"tissuedysplasia":0.4149,"tissuetumour":1.0098,"AveExpr":3.9562,"F":14.0533,"P.Value":0,"adj.P.Val":0.0004},{"tissuedysplasia":1.117,"tissuetumour":1.7103,"AveExpr":-1.3331,"F":4.033,"P.Value":0.0246,"adj.P.Val":0.0794},{"tissuedysplasia":-0.0267,"tissuetumour":-0.2511,"AveExpr":4.8136,"F":1.3103,"P.Value":0.2801,"adj.P.Val":0.4245},{"tissuedysplasia":-0.0558,"tissuetumour":-0.6254,"AveExpr":6.2837,"F":9.061,"P.Value":0.0005,"adj.P.Val":0.0047},{"tissuedysplasia":0.2923,"tissuetumour":0.1998,"AveExpr":6.2991,"F":4.3139,"P.Value":0.0195,"adj.P.Val":0.0673},{"tissuedysplasia":-0.3336,"tissuetumour":0.5348,"AveExpr":4.9598,"F":6.7991,"P.Value":0.0027,"adj.P.Val":0.016},{"tissuedysplasia":-0.4539,"tissuetumour":-0.5404,"AveExpr":3.0253,"F":4.9865,"P.Value":0.0112,"adj.P.Val":0.0448},{"tissuedysplasia":0.4169,"tissuetumour":1.239,"AveExpr":4.95,"F":27.9763,"P.Value":1.4461e-08,"adj.P.Val":1.8324e-06},{"tissuedysplasia":-0.0432,"tissuetumour":-0.2893,"AveExpr":4.9687,"F":1.9832,"P.Value":0.1497,"adj.P.Val":0.2771},{"tissuedysplasia":0.1469,"tissuetumour":-0.289,"AveExpr":7.0379,"F":7.5039,"P.Value":0.0016,"adj.P.Val":0.0108},{"tissuedysplasia":0.1188,"tissuetumour":0.2108,"AveExpr":5.7763,"F":2.6724,"P.Value":0.0803,"adj.P.Val":0.1811},{"tissuedysplasia":-0.0048,"tissuetumour":1.0148,"AveExpr":2.3694,"F":16.5446,"P.Value":4.3847e-06,"adj.P.Val":0.0001},{"tissuedysplasia":1.0245,"tissuetumour":0.3152,"AveExpr":6.3187,"F":7.7787,"P.Value":0.0015,"adj.P.Val":0.0105},{"tissuedysplasia":-0.1919,"tissuetumour":0.043,"AveExpr":4.2109,"F":1.2005,"P.Value":0.3107,"adj.P.Val":0.4552},{"tissuedysplasia":0.0952,"tissuetumour":0.3752,"AveExpr":2.8676,"F":2.9336,"P.Value":0.0637,"adj.P.Val":0.1546},{"tissuedysplasia":0.0047,"tissuetumour":0.037,"AveExpr":4.1668,"F":0.0664,"P.Value":0.9358,"adj.P.Val":0.9555},{"tissuedysplasia":0.1414,"tissuetumour":0.7442,"AveExpr":4.5132,"F":21.4866,"P.Value":3.0851e-07,"adj.P.Val":0},{"tissuedysplasia":0.1137,"tissuetumour":0.6303,"AveExpr":5.0201,"F":13.6513,"P.Value":0,"adj.P.Val":0.0005},{"tissuedysplasia":0.8471,"tissuetumour":1.4279,"AveExpr":4.1321,"F":14.8935,"P.Value":0,"adj.P.Val":0.0003},{"tissuedysplasia":-0.1195,"tissuetumour":0.5159,"AveExpr":5.1556,"F":10.8595,"P.Value":0.0001,"adj.P.Val":0.0019},{"tissuedysplasia":0.3468,"tissuetumour":0.1293,"AveExpr":4.8097,"F":1.7511,"P.Value":0.1855,"adj.P.Val":0.3209},{"tissuedysplasia":1.7917,"tissuetumour":1.2557,"AveExpr":6.1089,"F":11.362,"P.Value":0.0001,"adj.P.Val":0.0019},{"tissuedysplasia":-0.1745,"tissuetumour":3.0974,"AveExpr":-2.6425,"F":14.7834,"P.Value":0,"adj.P.Val":0.0003},{"tissuedysplasia":0.1935,"tissuetumour":0.6326,"AveExpr":3.1402,"F":12.2391,"P.Value":0.0001,"adj.P.Val":0.0009},{"tissuedysplasia":0.241,"tissuetumour":0.7891,"AveExpr":3.2394,"F":8.5691,"P.Value":0.0007,"adj.P.Val":0.0061},{"tissuedysplasia":-0.0247,"tissuetumour":0.1332,"AveExpr":3.7529,"F":1.1071,"P.Value":0.3396,"adj.P.Val":0.4836},{"tissuedysplasia":0.3475,"tissuetumour":0.372,"AveExpr":4.3994,"F":7.8331,"P.Value":0.0012,"adj.P.Val":0.009},{"tissuedysplasia":-0.9989,"tissuetumour":-1.5754,"AveExpr":0.8956,"F":6.6578,"P.Value":0.003,"adj.P.Val":0.0173},{"tissuedysplasia":0.4538,"tissuetumour":0.4276,"AveExpr":5.731,"F":6.0951,"P.Value":0.0046,"adj.P.Val":0.0237},{"tissuedysplasia":0.8987,"tissuetumour":1.4892,"AveExpr":5.0776,"F":8.575,"P.Value":0.0008,"adj.P.Val":0.0068},{"tissuedysplasia":0.9362,"tissuetumour":2.2459,"AveExpr":-2.5713,"F":4.37,"P.Value":0.0186,"adj.P.Val":0.065},{"tissuedysplasia":0.354,"tissuetumour":1.2308,"AveExpr":1.2209,"F":4.8185,"P.Value":0.0128,"adj.P.Val":0.0495},{"tissuedysplasia":0.4946,"tissuetumour":0.9144,"AveExpr":1.3794,"F":6.6064,"P.Value":0.0031,"adj.P.Val":0.0178},{"tissuedysplasia":0.4072,"tissuetumour":1.1256,"AveExpr":3.9239,"F":21.0421,"P.Value":3.8674e-07,"adj.P.Val":0},{"tissuedysplasia":0.4323,"tissuetumour":0.892,"AveExpr":4.0597,"F":21.2299,"P.Value":3.5143e-07,"adj.P.Val":0},{"tissuedysplasia":0.2226,"tissuetumour":-0.0139,"AveExpr":3.8976,"F":1.2984,"P.Value":0.2832,"adj.P.Val":0.4277},{"tissuedysplasia":0.0789,"tissuetumour":0.5116,"AveExpr":7.0418,"F":34.005,"P.Value":1.1801e-09,"adj.P.Val":2.8419e-07},{"tissuedysplasia":0.1953,"tissuetumour":0.6087,"AveExpr":6.3832,"F":7.823,"P.Value":0.0012,"adj.P.Val":0.0091},{"tissuedysplasia":0.631,"tissuetumour":1.4908,"AveExpr":2.4955,"F":15.3895,"P.Value":8.5639e-06,"adj.P.Val":0.0002},{"tissuedysplasia":-0.0007,"tissuetumour":-0.045,"AveExpr":4.6375,"F":0.058,"P.Value":0.9437,"adj.P.Val":0.9609},{"tissuedysplasia":0.035,"tissuetumour":0.7957,"AveExpr":2.4498,"F":11.204,"P.Value":0.0001,"adj.P.Val":0.0016},{"tissuedysplasia":0.2668,"tissuetumour":0.0851,"AveExpr":5.9257,"F":2.9291,"P.Value":0.0639,"adj.P.Val":0.155},{"tissuedysplasia":0.7777,"tissuetumour":1.4919,"AveExpr":5.0126,"F":29.7887,"P.Value":6.6039e-09,"adj.P.Val":1.0275e-06},{"tissuedysplasia":0.4001,"tissuetumour":0.5461,"AveExpr":5.6934,"F":16.5187,"P.Value":4.4502e-06,"adj.P.Val":0.0001},{"tissuedysplasia":0.6381,"tissuetumour":1.1644,"AveExpr":5.6506,"F":38.9481,"P.Value":1.8354e-10,"adj.P.Val":7.8903e-08},{"tissuedysplasia":0.2575,"tissuetumour":0.5909,"AveExpr":6.9584,"F":26.2081,"P.Value":3.1943e-08,"adj.P.Val":3.1908e-06},{"tissuedysplasia":0.6212,"tissuetumour":1.0112,"AveExpr":3.3645,"F":11.3801,"P.Value":0.0001,"adj.P.Val":0.0014},{"tissuedysplasia":0.0151,"tissuetumour":-0.0456,"AveExpr":6.6589,"F":0.2579,"P.Value":0.7738,"adj.P.Val":0.8398},{"tissuedysplasia":0.0519,"tissuetumour":0.4486,"AveExpr":5.2255,"F":3.0837,"P.Value":0.0558,"adj.P.Val":0.141},{"tissuedysplasia":0.8323,"tissuetumour":2.1206,"AveExpr":-5.3331,"F":2.5957,"P.Value":0.086,"adj.P.Val":0.1898},{"tissuedysplasia":0.584,"tissuetumour":1.8311,"AveExpr":-0.593,"F":8.2224,"P.Value":0.0009,"adj.P.Val":0.0073},{"tissuedysplasia":0.2934,"tissuetumour":0.7522,"AveExpr":3.9275,"F":11.8272,"P.Value":0.0001,"adj.P.Val":0.0011},{"tissuedysplasia":-0.0054,"tissuetumour":-0.6363,"AveExpr":5.2381,"F":8.072,"P.Value":0.001,"adj.P.Val":0.0079},{"tissuedysplasia":0.0372,"tissuetumour":-0.1082,"AveExpr":5.3689,"F":0.8968,"P.Value":0.4152,"adj.P.Val":0.5538},{"tissuedysplasia":-1.0356,"tissuetumour":0.9887,"AveExpr":-9.0863,"F":1.3278,"P.Value":0.2755,"adj.P.Val":0.4199},{"tissuedysplasia":-0.0773,"tissuetumour":-0.7474,"AveExpr":6.4434,"F":9.6929,"P.Value":0.0003,"adj.P.Val":0.0034},{"tissuedysplasia":0.0289,"tissuetumour":0.2115,"AveExpr":4.5912,"F":1.0872,"P.Value":0.346,"adj.P.Val":0.4901},{"tissuedysplasia":0.5999,"tissuetumour":0.8576,"AveExpr":0.8529,"F":3.4146,"P.Value":0.0418,"adj.P.Val":0.1152},{"tissuedysplasia":0.0658,"tissuetumour":0.0972,"AveExpr":4.5618,"F":0.3152,"P.Value":0.7313,"adj.P.Val":0.8082},{"tissuedysplasia":-2.0879,"tissuetumour":-3.4146,"AveExpr":2.4993,"F":22.9883,"P.Value":1.5416e-07,"adj.P.Val":9.6735e-06},{"tissuedysplasia":0.5976,"tissuetumour":0.5043,"AveExpr":3.6527,"F":2.8716,"P.Value":0.0673,"adj.P.Val":0.1603},{"tissuedysplasia":0.1333,"tissuetumour":0.5381,"AveExpr":6.7493,"F":6.6836,"P.Value":0.0029,"adj.P.Val":0.017},{"tissuedysplasia":0.6368,"tissuetumour":1.8298,"AveExpr":-0.7036,"F":7.4203,"P.Value":0.0017,"adj.P.Val":0.0113},{"tissuedysplasia":0.6255,"tissuetumour":2.2686,"AveExpr":-2.9017,"F":3.6918,"P.Value":0.0329,"adj.P.Val":0.0973},{"tissuedysplasia":0.1891,"tissuetumour":-0.1197,"AveExpr":5.3203,"F":2.3793,"P.Value":0.1044,"adj.P.Val":0.2172},{"tissuedysplasia":0.2818,"tissuetumour":0.0284,"AveExpr":4.9733,"F":3.2997,"P.Value":0.0462,"adj.P.Val":0.1235},{"tissuedysplasia":0.0564,"tissuetumour":-0.338,"AveExpr":4.916,"F":6.827,"P.Value":0.0026,"adj.P.Val":0.0157},{"tissuedysplasia":0.6748,"tissuetumour":1.2238,"AveExpr":0.6067,"F":3.1548,"P.Value":0.0524,"adj.P.Val":0.135},{"tissuedysplasia":0.157,"tissuetumour":0.1554,"AveExpr":1.9621,"F":0.2114,"P.Value":0.8103,"adj.P.Val":0.8669},{"tissuedysplasia":0.0213,"tissuetumour":-0.24,"AveExpr":5.3826,"F":1.6011,"P.Value":0.2132,"adj.P.Val":0.3523},{"tissuedysplasia":0.9809,"tissuetumour":1.5811,"AveExpr":8.2345,"F":13.1634,"P.Value":0.0001,"adj.P.Val":0.0008},{"tissuedysplasia":0.1052,"tissuetumour":0.1446,"AveExpr":3.2916,"F":0.3299,"P.Value":0.7207,"adj.P.Val":0.8004},{"tissuedysplasia":0.2196,"tissuetumour":0.7084,"AveExpr":5.9022,"F":8.1601,"P.Value":0.001,"adj.P.Val":0.0075},{"tissuedysplasia":-1.4442,"tissuetumour":2.0073,"AveExpr":-3.6852,"F":4.1274,"P.Value":0.0228,"adj.P.Val":0.075},{"tissuedysplasia":0.1035,"tissuetumour":-0.2769,"AveExpr":3.8326,"F":2.0775,"P.Value":0.1373,"adj.P.Val":0.2617},{"tissuedysplasia":1.2223,"tissuetumour":1.2809,"AveExpr":5.0279,"F":24.0545,"P.Value":8.7309e-08,"adj.P.Val":6.4881e-06},{"tissuedysplasia":-0.4215,"tissuetumour":0.0863,"AveExpr":2.1116,"F":1.3019,"P.Value":0.2823,"adj.P.Val":0.4269},{"tissuedysplasia":0.2621,"tissuetumour":0.0404,"AveExpr":7.1506,"F":4.5976,"P.Value":0.0154,"adj.P.Val":0.0566},{"tissuedysplasia":0.3717,"tissuetumour":1.4779,"AveExpr":0.7528,"F":8.945,"P.Value":0.0005,"adj.P.Val":0.005},{"tissuedysplasia":0.2757,"tissuetumour":0.707,"AveExpr":4.9865,"F":13.0515,"P.Value":0,"adj.P.Val":0.0006},{"tissuedysplasia":0.1259,"tissuetumour":2.9789,"AveExpr":-4.8356,"F":6.2995,"P.Value":0.0039,"adj.P.Val":0.0211},{"tissuedysplasia":0.3526,"tissuetumour":0.5637,"AveExpr":6.2447,"F":6.724,"P.Value":0.0028,"adj.P.Val":0.0167},{"tissuedysplasia":-0.3046,"tissuetumour":0.0073,"AveExpr":5.0404,"F":1.4771,"P.Value":0.2394,"adj.P.Val":0.3814},{"tissuedysplasia":0.0541,"tissuetumour":0.4303,"AveExpr":3.4816,"F":5.5548,"P.Value":0.0071,"adj.P.Val":0.0323},{"tissuedysplasia":0.5605,"tissuetumour":0.7421,"AveExpr":3.3896,"F":10.144,"P.Value":0.0002,"adj.P.Val":0.0027},{"tissuedysplasia":0.0228,"tissuetumour":0.6263,"AveExpr":3.9688,"F":3.7043,"P.Value":0.0326,"adj.P.Val":0.0966},{"tissuedysplasia":-0.4912,"tissuetumour":2.5078,"AveExpr":-2.9759,"F":7.4522,"P.Value":0.0016,"adj.P.Val":0.0111},{"tissuedysplasia":0.9626,"tissuetumour":2.2103,"AveExpr":-5.1981,"F":2.3357,"P.Value":0.1086,"adj.P.Val":0.2231},{"tissuedysplasia":0.4212,"tissuetumour":1.0999,"AveExpr":1.9238,"F":14.2213,"P.Value":0,"adj.P.Val":0.0004},{"tissuedysplasia":1.0671,"tissuetumour":1.2742,"AveExpr":2.6408,"F":14.4861,"P.Value":0,"adj.P.Val":0.0003},{"tissuedysplasia":0.293,"tissuetumour":0.7278,"AveExpr":1.996,"F":4.7439,"P.Value":0.0136,"adj.P.Val":0.0518},{"tissuedysplasia":0.3728,"tissuetumour":0.3134,"AveExpr":4.152,"F":3.3783,"P.Value":0.0432,"adj.P.Val":0.1177},{"tissuedysplasia":0.1901,"tissuetumour":0.9186,"AveExpr":2.3292,"F":7.5967,"P.Value":0.0015,"adj.P.Val":0.0103},{"tissuedysplasia":0.8487,"tissuetumour":1.3995,"AveExpr":1.476,"F":10.0076,"P.Value":0.0003,"adj.P.Val":0.0029},{"tissuedysplasia":0.229,"tissuetumour":0.5997,"AveExpr":6.4324,"F":2.1845,"P.Value":0.1266,"adj.P.Val":0.2473},{"tissuedysplasia":0.0706,"tissuetumour":-0.0213,"AveExpr":-1.2104,"F":0.0158,"P.Value":0.9843,"adj.P.Val":0.9891},{"tissuedysplasia":0.4748,"tissuetumour":0.6842,"AveExpr":4.822,"F":9.8838,"P.Value":0.0003,"adj.P.Val":0.0031},{"tissuedysplasia":0.2638,"tissuetumour":0.3108,"AveExpr":3.1781,"F":1.7708,"P.Value":0.1821,"adj.P.Val":0.317},{"tissuedysplasia":0.2425,"tissuetumour":0.2884,"AveExpr":3.0944,"F":2.1736,"P.Value":0.1258,"adj.P.Val":0.2462},{"tissuedysplasia":0.2209,"tissuetumour":0.9021,"AveExpr":-0.1841,"F":2.9961,"P.Value":0.0603,"adj.P.Val":0.1489},{"tissuedysplasia":-3.0819,"tissuetumour":1.9143,"AveExpr":2.0183,"F":13.2227,"P.Value":0,"adj.P.Val":0.0008},{"tissuedysplasia":1.8597,"tissuetumour":4.4275,"AveExpr":-5.7019,"F":5.8638,"P.Value":0.0055,"adj.P.Val":0.027},{"tissuedysplasia":0.4921,"tissuetumour":1.5163,"AveExpr":5.6249,"F":24.1447,"P.Value":8.3629e-08,"adj.P.Val":6.2723e-06},{"tissuedysplasia":0.1494,"tissuetumour":0.3045,"AveExpr":3.7561,"F":2.6588,"P.Value":0.0813,"adj.P.Val":0.1827},{"tissuedysplasia":-0.5009,"tissuetumour":0.4033,"AveExpr":1.3051,"F":3.6887,"P.Value":0.033,"adj.P.Val":0.0975},{"tissuedysplasia":0.3104,"tissuetumour":0.9721,"AveExpr":4.5729,"F":18.982,"P.Value":1.1378e-06,"adj.P.Val":0},{"tissuedysplasia":0.0453,"tissuetumour":-0.3106,"AveExpr":5.189,"F":3.3758,"P.Value":0.0433,"adj.P.Val":0.1179},{"tissuedysplasia":0.2067,"tissuetumour":0.1319,"AveExpr":3.9387,"F":1.1123,"P.Value":0.3379,"adj.P.Val":0.482},{"tissuedysplasia":-0.332,"tissuetumour":-0.7712,"AveExpr":8.0854,"F":25.7448,"P.Value":3.9505e-08,"adj.P.Val":3.6823e-06},{"tissuedysplasia":0.162,"tissuetumour":0.194,"AveExpr":7.7248,"F":2.4032,"P.Value":0.1022,"adj.P.Val":0.2141},{"tissuedysplasia":0.3739,"tissuetumour":0.7778,"AveExpr":0.5856,"F":1.7064,"P.Value":0.1933,"adj.P.Val":0.3298},{"tissuedysplasia":0.3218,"tissuetumour":0.5725,"AveExpr":6.1017,"F":15.2688,"P.Value":9.1952e-06,"adj.P.Val":0.0002},{"tissuedysplasia":-0.0207,"tissuetumour":-0.0832,"AveExpr":5.5865,"F":0.6341,"P.Value":0.5352,"adj.P.Val":0.6565},{"tissuedysplasia":0.5008,"tissuetumour":0.6792,"AveExpr":1.5292,"F":3.7984,"P.Value":0.0301,"adj.P.Val":0.0914},{"tissuedysplasia":-0.8188,"tissuetumour":-1.3581,"AveExpr":7.4821,"F":20.1359,"P.Value":1.1781e-06,"adj.P.Val":0},{"tissuedysplasia":0.0007,"tissuetumour":-0.1826,"AveExpr":5.2532,"F":2.056,"P.Value":0.1401,"adj.P.Val":0.2649},{"tissuedysplasia":2.1717,"tissuetumour":3.1703,"AveExpr":-3.1416,"F":7.4137,"P.Value":0.0017,"adj.P.Val":0.0114},{"tissuedysplasia":-0.9338,"tissuetumour":-0.3915,"AveExpr":-3.53,"F":1.5873,"P.Value":0.216,"adj.P.Val":0.3555},{"tissuedysplasia":-0.082,"tissuetumour":-0.2139,"AveExpr":4.3985,"F":0.6159,"P.Value":0.5448,"adj.P.Val":0.6638},{"tissuedysplasia":0.2226,"tissuetumour":0.083,"AveExpr":6.483,"F":3.4704,"P.Value":0.0399,"adj.P.Val":0.1113},{"tissuedysplasia":-0.8876,"tissuetumour":-0.5253,"AveExpr":3.9245,"F":10.1504,"P.Value":0.0002,"adj.P.Val":0.0027},{"tissuedysplasia":0.6303,"tissuetumour":0.7614,"AveExpr":3.3396,"F":8.2793,"P.Value":0.0009,"adj.P.Val":0.0071},{"tissuedysplasia":0.0853,"tissuetumour":0.5282,"AveExpr":4.2557,"F":9.3522,"P.Value":0.0004,"adj.P.Val":0.004},{"tissuedysplasia":0.2331,"tissuetumour":0.1528,"AveExpr":4.9017,"F":1.7715,"P.Value":0.182,"adj.P.Val":0.3169},{"tissuedysplasia":-0.3216,"tissuetumour":1.3435,"AveExpr":1.7463,"F":13.7794,"P.Value":0,"adj.P.Val":0.0004},{"tissuedysplasia":-0.2998,"tissuetumour":-0.8544,"AveExpr":2.9227,"F":4.1438,"P.Value":0.0224,"adj.P.Val":0.0743},{"tissuedysplasia":0.4582,"tissuetumour":1.1435,"AveExpr":-0.7173,"F":3.2664,"P.Value":0.0476,"adj.P.Val":0.1262},{"tissuedysplasia":-0.2086,"tissuetumour":-0.3863,"AveExpr":8.2105,"F":3.4554,"P.Value":0.0417,"adj.P.Val":0.1148},{"tissuedysplasia":-0.0281,"tissuetumour":0.1011,"AveExpr":7.3583,"F":2.1457,"P.Value":0.1291,"adj.P.Val":0.2505},{"tissuedysplasia":0.1725,"tissuetumour":0.1512,"AveExpr":6.0343,"F":1.4444,"P.Value":0.2469,"adj.P.Val":0.3895},{"tissuedysplasia":0.3193,"tissuetumour":1.5543,"AveExpr":-3.5015,"F":1.778,"P.Value":0.1809,"adj.P.Val":0.3156},{"tissuedysplasia":0.9754,"tissuetumour":1.6009,"AveExpr":3.5137,"F":48.6958,"P.Value":7.0157e-12,"adj.P.Val":9.2539e-09},{"tissuedysplasia":-0.2948,"tissuetumour":-0.6716,"AveExpr":5.6454,"F":21.4477,"P.Value":3.1463e-07,"adj.P.Val":0},{"tissuedysplasia":0.2362,"tissuetumour":0.5763,"AveExpr":3.6066,"F":4.6584,"P.Value":0.0146,"adj.P.Val":0.0546},{"tissuedysplasia":0.0379,"tissuetumour":0.2261,"AveExpr":4.9865,"F":3.1431,"P.Value":0.053,"adj.P.Val":0.136},{"tissuedysplasia":0.2944,"tissuetumour":0.2965,"AveExpr":6.2471,"F":3.1725,"P.Value":0.0516,"adj.P.Val":0.1336},{"tissuedysplasia":0.5806,"tissuetumour":0.9985,"AveExpr":3.1703,"F":9.235,"P.Value":0.0004,"adj.P.Val":0.0043},{"tissuedysplasia":0.3439,"tissuetumour":0.022,"AveExpr":4.7875,"F":3.8527,"P.Value":0.0287,"adj.P.Val":0.0884},{"tissuedysplasia":0.2472,"tissuetumour":0.0908,"AveExpr":6.4388,"F":2.5844,"P.Value":0.0869,"adj.P.Val":0.1912},{"tissuedysplasia":0.842,"tissuetumour":1.3102,"AveExpr":2.475,"F":23.188,"P.Value":1.326e-07,"adj.P.Val":8.6981e-06},{"tissuedysplasia":0.1454,"tissuetumour":0.3708,"AveExpr":4.6266,"F":3.7815,"P.Value":0.0305,"adj.P.Val":0.0923},{"tissuedysplasia":0.6952,"tissuetumour":0.4585,"AveExpr":4.3311,"F":2.8522,"P.Value":0.0684,"adj.P.Val":0.1623},{"tissuedysplasia":0.4246,"tissuetumour":1.2445,"AveExpr":3.8111,"F":23.5079,"P.Value":1.1354e-07,"adj.P.Val":7.7706e-06},{"tissuedysplasia":1.3716,"tissuetumour":1.0771,"AveExpr":1.963,"F":6.9804,"P.Value":0.0023,"adj.P.Val":0.0144},{"tissuedysplasia":0.5227,"tissuetumour":0.2007,"AveExpr":4.4245,"F":4.9971,"P.Value":0.0111,"adj.P.Val":0.0446},{"tissuedysplasia":-0.666,"tissuetumour":-1.0936,"AveExpr":6.4823,"F":32.9382,"P.Value":1.8017e-09,"adj.P.Val":3.8443e-07},{"tissuedysplasia":0.2918,"tissuetumour":0.4102,"AveExpr":4.3763,"F":7.1822,"P.Value":0.002,"adj.P.Val":0.0129},{"tissuedysplasia":-0.4187,"tissuetumour":-0.4612,"AveExpr":2.8254,"F":2.9321,"P.Value":0.0638,"adj.P.Val":0.1547},{"tissuedysplasia":0.0933,"tissuetumour":0.3684,"AveExpr":4.6533,"F":5.2434,"P.Value":0.0091,"adj.P.Val":0.0386},{"tissuedysplasia":1.0978,"tissuetumour":2.1734,"AveExpr":-5.0242,"F":2.0174,"P.Value":0.1451,"adj.P.Val":0.2714},{"tissuedysplasia":-0.7669,"tissuetumour":-0.8928,"AveExpr":1.8639,"F":1.4722,"P.Value":0.2413,"adj.P.Val":0.3835},{"tissuedysplasia":0.2403,"tissuetumour":-2.2746,"AveExpr":0.643,"F":12.4532,"P.Value":0.0001,"adj.P.Val":0.0008},{"tissuedysplasia":0.789,"tissuetumour":1.3074,"AveExpr":3.3244,"F":35.3566,"P.Value":6.9832e-10,"adj.P.Val":1.9569e-07},{"tissuedysplasia":-0.0939,"tissuetumour":-0.152,"AveExpr":2.331,"F":0.4486,"P.Value":0.6414,"adj.P.Val":0.741},{"tissuedysplasia":0.2003,"tissuetumour":0.0988,"AveExpr":3.9099,"F":1.2838,"P.Value":0.2872,"adj.P.Val":0.4317},{"tissuedysplasia":0.1294,"tissuetumour":0.7615,"AveExpr":3.4605,"F":18.3415,"P.Value":1.6091e-06,"adj.P.Val":0.0001},{"tissuedysplasia":0.1644,"tissuetumour":-0.1668,"AveExpr":7.0041,"F":5.086,"P.Value":0.0103,"adj.P.Val":0.0423},{"tissuedysplasia":-1.8261,"tissuetumour":-3.1054,"AveExpr":4.3557,"F":26.9392,"P.Value":4.4781e-08,"adj.P.Val":4.0107e-06},{"tissuedysplasia":-0.3806,"tissuetumour":-0.946,"AveExpr":5.8542,"F":12.052,"P.Value":0.0001,"adj.P.Val":0.001},{"tissuedysplasia":0.3482,"tissuetumour":-0.0135,"AveExpr":3.5692,"F":3.7837,"P.Value":0.0305,"adj.P.Val":0.0922},{"tissuedysplasia":-0.3388,"tissuetumour":-1.1201,"AveExpr":5.8955,"F":36.2864,"P.Value":4.9024e-10,"adj.P.Val":1.5711e-07},{"tissuedysplasia":0.0311,"tissuetumour":0.4473,"AveExpr":3.8351,"F":6.2146,"P.Value":0.0042,"adj.P.Val":0.0222},{"tissuedysplasia":-0.0991,"tissuetumour":0.2018,"AveExpr":5.3274,"F":2.2879,"P.Value":0.1134,"adj.P.Val":0.2298},{"tissuedysplasia":-0.2914,"tissuetumour":0.9304,"AveExpr":-0.2729,"F":2.8154,"P.Value":0.0707,"adj.P.Val":0.1661},{"tissuedysplasia":0.0777,"tissuetumour":0.189,"AveExpr":6.2701,"F":1.0398,"P.Value":0.3621,"adj.P.Val":0.5053},{"tissuedysplasia":0.8592,"tissuetumour":1.197,"AveExpr":3.4733,"F":12.5073,"P.Value":0.0001,"adj.P.Val":0.0008},{"tissuedysplasia":0.4729,"tissuetumour":0.6793,"AveExpr":2.2103,"F":3.7633,"P.Value":0.031,"adj.P.Val":0.0933},{"tissuedysplasia":0.2004,"tissuetumour":0.1189,"AveExpr":4.6497,"F":1.1029,"P.Value":0.3409,"adj.P.Val":0.4848},{"tissuedysplasia":-0.2322,"tissuetumour":0.3874,"AveExpr":5.0371,"F":3.3216,"P.Value":0.0453,"adj.P.Val":0.122},{"tissuedysplasia":0.0146,"tissuetumour":0.2564,"AveExpr":5.5357,"F":3.9705,"P.Value":0.026,"adj.P.Val":0.0823},{"tissuedysplasia":0.2006,"tissuetumour":0.8519,"AveExpr":2.2623,"F":5.2675,"P.Value":0.0089,"adj.P.Val":0.0381},{"tissuedysplasia":0.7099,"tissuetumour":1.0616,"AveExpr":1.7131,"F":8.4966,"P.Value":0.0008,"adj.P.Val":0.0063},{"tissuedysplasia":0.2038,"tissuetumour":0.5492,"AveExpr":4.3042,"F":10.0553,"P.Value":0.0003,"adj.P.Val":0.0028},{"tissuedysplasia":-0.1701,"tissuetumour":0.429,"AveExpr":4.4729,"F":6.3994,"P.Value":0.0036,"adj.P.Val":0.0199},{"tissuedysplasia":0.3733,"tissuetumour":1.1187,"AveExpr":2.5756,"F":13.3703,"P.Value":0,"adj.P.Val":0.0005},{"tissuedysplasia":0.1619,"tissuetumour":0.4924,"AveExpr":0.2547,"F":1.105,"P.Value":0.3402,"adj.P.Val":0.4843},{"tissuedysplasia":0.3533,"tissuetumour":0.3853,"AveExpr":3.4249,"F":4.8142,"P.Value":0.0129,"adj.P.Val":0.0496},{"tissuedysplasia":-0.0462,"tissuetumour":0.2337,"AveExpr":1.7356,"F":0.7395,"P.Value":0.4832,"adj.P.Val":0.6122},{"tissuedysplasia":-0.0274,"tissuetumour":-0.5837,"AveExpr":6.2934,"F":11.1796,"P.Value":0.0001,"adj.P.Val":0.0016},{"tissuedysplasia":0.1423,"tissuetumour":-0.2589,"AveExpr":3.7261,"F":2.2201,"P.Value":0.1206,"adj.P.Val":0.2392},{"tissuedysplasia":-0.6091,"tissuetumour":-1.6633,"AveExpr":4.2379,"F":39.1953,"P.Value":1.679e-10,"adj.P.Val":7.4384e-08},{"tissuedysplasia":0.4524,"tissuetumour":2.0247,"AveExpr":-0.3683,"F":10.0821,"P.Value":0.0002,"adj.P.Val":0.0028},{"tissuedysplasia":0.0695,"tissuetumour":0.5817,"AveExpr":-4.1364,"F":0.1875,"P.Value":0.8297,"adj.P.Val":0.8804},{"tissuedysplasia":0.5051,"tissuetumour":1.5211,"AveExpr":-0.057,"F":5.5725,"P.Value":0.007,"adj.P.Val":0.032},{"tissuedysplasia":-0.5604,"tissuetumour":-0.4419,"AveExpr":4.8471,"F":4.8391,"P.Value":0.0126,"adj.P.Val":0.0489},{"tissuedysplasia":-1.5278,"tissuetumour":-0.5868,"AveExpr":0.6581,"F":7.7815,"P.Value":0.0013,"adj.P.Val":0.0093},{"tissuedysplasia":-0.0615,"tissuetumour":-0.5115,"AveExpr":0.7377,"F":0.4537,"P.Value":0.6382,"adj.P.Val":0.7386},{"tissuedysplasia":0.8044,"tissuetumour":3.1367,"AveExpr":-0.276,"F":15.0715,"P.Value":0,"adj.P.Val":0.0002},{"tissuedysplasia":-0.177,"tissuetumour":0.7614,"AveExpr":0.0963,"F":1.8288,"P.Value":0.1726,"adj.P.Val":0.3055},{"tissuedysplasia":0.0102,"tissuetumour":-0.4329,"AveExpr":6.9693,"F":12.9763,"P.Value":0,"adj.P.Val":0.0006},{"tissuedysplasia":0.4365,"tissuetumour":0.3889,"AveExpr":3.1459,"F":2.9589,"P.Value":0.0623,"adj.P.Val":0.1521},{"tissuedysplasia":0.261,"tissuetumour":1.0974,"AveExpr":2.5868,"F":12.4614,"P.Value":0.0001,"adj.P.Val":0.0008},{"tissuedysplasia":0.1832,"tissuetumour":0.7345,"AveExpr":0.1324,"F":1.9063,"P.Value":0.1607,"adj.P.Val":0.2908},{"tissuedysplasia":1.4347,"tissuetumour":1.5588,"AveExpr":-4.0046,"F":1.2802,"P.Value":0.2881,"adj.P.Val":0.4327},{"tissuedysplasia":0.1001,"tissuetumour":-0.2587,"AveExpr":6.149,"F":6.3708,"P.Value":0.0037,"adj.P.Val":0.0203},{"tissuedysplasia":3.3946,"tissuetumour":4.5612,"AveExpr":-2.1671,"F":12.5982,"P.Value":0,"adj.P.Val":0.0008},{"tissuedysplasia":-1.5257,"tissuetumour":1.1692,"AveExpr":-5.8953,"F":2.7233,"P.Value":0.0767,"adj.P.Val":0.1755},{"tissuedysplasia":-3.0375,"tissuetumour":-4.6307,"AveExpr":2.0333,"F":23.7407,"P.Value":1.8807e-07,"adj.P.Val":0},{"tissuedysplasia":0.1174,"tissuetumour":-0.0412,"AveExpr":7.7338,"F":0.9922,"P.Value":0.3791,"adj.P.Val":0.5211},{"tissuedysplasia":-0.0179,"tissuetumour":-0.2074,"AveExpr":6.2493,"F":1.9068,"P.Value":0.1606,"adj.P.Val":0.2908},{"tissuedysplasia":0.06,"tissuetumour":0.4695,"AveExpr":4.9393,"F":8.8663,"P.Value":0.0006,"adj.P.Val":0.0052},{"tissuedysplasia":-0.0418,"tissuetumour":0.2211,"AveExpr":2.397,"F":0.7933,"P.Value":0.4587,"adj.P.Val":0.5922},{"tissuedysplasia":0.5297,"tissuetumour":2.4914,"AveExpr":-5.1461,"F":3.8617,"P.Value":0.0285,"adj.P.Val":0.088},{"tissuedysplasia":0.3339,"tissuetumour":-0.2,"AveExpr":2.2508,"F":2.2598,"P.Value":0.1164,"adj.P.Val":0.2337},{"tissuedysplasia":0.1307,"tissuetumour":0.2773,"AveExpr":4.4993,"F":1.5814,"P.Value":0.2172,"adj.P.Val":0.3568},{"tissuedysplasia":2.5597,"tissuetumour":3.2356,"AveExpr":-5.9721,"F":4.6888,"P.Value":0.0143,"adj.P.Val":0.0536},{"tissuedysplasia":0.1646,"tissuetumour":0.5594,"AveExpr":8.3165,"F":18.0276,"P.Value":1.9108e-06,"adj.P.Val":0.0001},{"tissuedysplasia":1.7873,"tissuetumour":2.2423,"AveExpr":-8.3456,"F":2.1889,"P.Value":0.1241,"adj.P.Val":0.2438},{"tissuedysplasia":0.1089,"tissuetumour":0.0616,"AveExpr":5.1763,"F":0.4484,"P.Value":0.6415,"adj.P.Val":0.7411},{"tissuedysplasia":0.582,"tissuetumour":1.1711,"AveExpr":7.0831,"F":45.7728,"P.Value":1.7764e-11,"adj.P.Val":1.5861e-08},{"tissuedysplasia":0.2762,"tissuetumour":-0.0763,"AveExpr":2.8714,"F":1.8341,"P.Value":0.1718,"adj.P.Val":0.3045},{"tissuedysplasia":-0.2571,"tissuetumour":-0.6252,"AveExpr":12.0244,"F":40.5037,"P.Value":5.9777e-10,"adj.P.Val":1.8069e-07},{"tissuedysplasia":-0.042,"tissuetumour":-0.1948,"AveExpr":4.298,"F":0.9881,"P.Value":0.3804,"adj.P.Val":0.5223},{"tissuedysplasia":0.1954,"tissuetumour":0.2402,"AveExpr":5.9996,"F":3.2862,"P.Value":0.0468,"adj.P.Val":0.1246},{"tissuedysplasia":-0.1054,"tissuetumour":-0.5827,"AveExpr":2.763,"F":4.1311,"P.Value":0.0227,"adj.P.Val":0.0749},{"tissuedysplasia":0.1473,"tissuetumour":-0.3127,"AveExpr":5.1256,"F":2.6725,"P.Value":0.0803,"adj.P.Val":0.1811},{"tissuedysplasia":1.2441,"tissuetumour":1.7105,"AveExpr":1.0602,"F":15.3704,"P.Value":8.6606e-06,"adj.P.Val":0.0002},{"tissuedysplasia":-0.0159,"tissuetumour":0.4081,"AveExpr":5.3056,"F":4.9212,"P.Value":0.0118,"adj.P.Val":0.0465},{"tissuedysplasia":0.4503,"tissuetumour":1.2814,"AveExpr":2.9922,"F":20.4198,"P.Value":5.3281e-07,"adj.P.Val":0},{"tissuedysplasia":-0.0552,"tissuetumour":-0.5677,"AveExpr":6.0146,"F":9.452,"P.Value":0.0004,"adj.P.Val":0.0038},{"tissuedysplasia":0.3348,"tissuetumour":0.3104,"AveExpr":2.8036,"F":2.3414,"P.Value":0.1081,"adj.P.Val":0.2222},{"tissuedysplasia":-0.437,"tissuetumour":-0.9835,"AveExpr":2.0475,"F":5.7115,"P.Value":0.0062,"adj.P.Val":0.0295},{"tissuedysplasia":-0.2085,"tissuetumour":-0.2299,"AveExpr":5.0829,"F":0.8271,"P.Value":0.444,"adj.P.Val":0.5797},{"tissuedysplasia":0.2105,"tissuetumour":0.1749,"AveExpr":7.3801,"F":0.4008,"P.Value":0.6727,"adj.P.Val":0.765},{"tissuedysplasia":-0.0449,"tissuetumour":0.5101,"AveExpr":1.7505,"F":4.401,"P.Value":0.0181,"adj.P.Val":0.0638},{"tissuedysplasia":-0.3108,"tissuetumour":-1.2393,"AveExpr":8.4812,"F":37.4745,"P.Value":9.0849e-10,"adj.P.Val":2.3375e-07},{"tissuedysplasia":-0.0277,"tissuetumour":-0.7873,"AveExpr":5.1408,"F":11.1177,"P.Value":0.0001,"adj.P.Val":0.0016},{"tissuedysplasia":0.0729,"tissuetumour":-0.3452,"AveExpr":5.6603,"F":3.7767,"P.Value":0.0306,"adj.P.Val":0.0926},{"tissuedysplasia":0.1205,"tissuetumour":0.2064,"AveExpr":5.8087,"F":1.3729,"P.Value":0.264,"adj.P.Val":0.4074},{"tissuedysplasia":-0.0768,"tissuetumour":0.0122,"AveExpr":0.9739,"F":0.1003,"P.Value":0.9048,"adj.P.Val":0.9337},{"tissuedysplasia":0.5925,"tissuetumour":0.7077,"AveExpr":0.6752,"F":2.7064,"P.Value":0.0779,"adj.P.Val":0.1773},{"tissuedysplasia":-0.0808,"tissuetumour":-0.6088,"AveExpr":5.9484,"F":9.9493,"P.Value":0.0003,"adj.P.Val":0.003},{"tissuedysplasia":0.7163,"tissuetumour":1.3144,"AveExpr":2.0463,"F":15.8688,"P.Value":6.4709e-06,"adj.P.Val":0.0002},{"tissuedysplasia":0.1272,"tissuetumour":0.2187,"AveExpr":2.2656,"F":0.2566,"P.Value":0.7748,"adj.P.Val":0.8407},{"tissuedysplasia":0.2789,"tissuetumour":0.2133,"AveExpr":5.405,"F":3.878,"P.Value":0.0281,"adj.P.Val":0.0872},{"tissuedysplasia":0.6414,"tissuetumour":1.0284,"AveExpr":3.1244,"F":13.4928,"P.Value":0,"adj.P.Val":0.0005},{"tissuedysplasia":0.023,"tissuetumour":0.1811,"AveExpr":4.4982,"F":0.6167,"P.Value":0.5443,"adj.P.Val":0.6634},{"tissuedysplasia":0.1293,"tissuetumour":0.5545,"AveExpr":-0.0503,"F":1.2202,"P.Value":0.305,"adj.P.Val":0.4495},{"tissuedysplasia":0.4728,"tissuetumour":0.9733,"AveExpr":6.3282,"F":24.9277,"P.Value":5.7753e-08,"adj.P.Val":4.7906e-06},{"tissuedysplasia":-0.0714,"tissuetumour":0.7837,"AveExpr":6.061,"F":9.6047,"P.Value":0.0004,"adj.P.Val":0.0037},{"tissuedysplasia":-0.1871,"tissuetumour":0.5351,"AveExpr":1.9922,"F":3.8899,"P.Value":0.0278,"adj.P.Val":0.0866},{"tissuedysplasia":0.0384,"tissuetumour":0.0988,"AveExpr":4.5993,"F":0.2483,"P.Value":0.7812,"adj.P.Val":0.8452},{"tissuedysplasia":0.2424,"tissuetumour":1.337,"AveExpr":3.4339,"F":17.5875,"P.Value":2.4369e-06,"adj.P.Val":0.0001},{"tissuedysplasia":0.3669,"tissuetumour":1.067,"AveExpr":1.1875,"F":6.429,"P.Value":0.0036,"adj.P.Val":0.0196},{"tissuedysplasia":0.4698,"tissuetumour":1.3862,"AveExpr":1.7643,"F":12.6792,"P.Value":0,"adj.P.Val":0.0008},{"tissuedysplasia":0.3877,"tissuetumour":1.0316,"AveExpr":4.597,"F":14.829,"P.Value":0,"adj.P.Val":0.0003},{"tissuedysplasia":0.3174,"tissuetumour":1.0632,"AveExpr":2.4673,"F":17.8995,"P.Value":2.0504e-06,"adj.P.Val":0.0001},{"tissuedysplasia":2.1244,"tissuetumour":3.0573,"AveExpr":-6.3049,"F":2.1663,"P.Value":0.1267,"adj.P.Val":0.2473},{"tissuedysplasia":-0.1868,"tissuetumour":0.6527,"AveExpr":-6.2831,"F":0.697,"P.Value":0.5035,"adj.P.Val":0.6297},{"tissuedysplasia":-0.0581,"tissuetumour":0.3914,"AveExpr":6.2045,"F":5.8899,"P.Value":0.0054,"adj.P.Val":0.0266},{"tissuedysplasia":-0.0412,"tissuetumour":0.7842,"AveExpr":3.1867,"F":9.0768,"P.Value":0.0005,"adj.P.Val":0.0047},{"tissuedysplasia":-0.2388,"tissuetumour":0.4747,"AveExpr":0.6436,"F":3.1997,"P.Value":0.0504,"adj.P.Val":0.1314},{"tissuedysplasia":0.2339,"tissuetumour":1.1541,"AveExpr":-0.1713,"F":4.8251,"P.Value":0.0127,"adj.P.Val":0.0493},{"tissuedysplasia":-0.581,"tissuetumour":0.1879,"AveExpr":-8.6259,"F":0.2312,"P.Value":0.7945,"adj.P.Val":0.8556},{"tissuedysplasia":0.1228,"tissuetumour":-0.4618,"AveExpr":3.4522,"F":5.1598,"P.Value":0.0097,"adj.P.Val":0.0405},{"tissuedysplasia":0.663,"tissuetumour":1.9119,"AveExpr":-4.5063,"F":2.0384,"P.Value":0.1424,"adj.P.Val":0.2677},{"tissuedysplasia":0.0764,"tissuetumour":0.2253,"AveExpr":6.4644,"F":3.8637,"P.Value":0.0285,"adj.P.Val":0.0879},{"tissuedysplasia":-0.0115,"tissuetumour":0.3823,"AveExpr":2.3531,"F":2.5014,"P.Value":0.0936,"adj.P.Val":0.2015},{"tissuedysplasia":0.1531,"tissuetumour":0.7658,"AveExpr":2.6578,"F":11.0304,"P.Value":0.0001,"adj.P.Val":0.0017},{"tissuedysplasia":-0.229,"tissuetumour":-0.2793,"AveExpr":5.0815,"F":2.8982,"P.Value":0.0657,"adj.P.Val":0.1577},{"tissuedysplasia":0.0761,"tissuetumour":0.4741,"AveExpr":3.5917,"F":7.0363,"P.Value":0.0022,"adj.P.Val":0.014},{"tissuedysplasia":-0.1709,"tissuetumour":0.5141,"AveExpr":3.4106,"F":8.2872,"P.Value":0.0009,"adj.P.Val":0.0071},{"tissuedysplasia":0.0634,"tissuetumour":0.1801,"AveExpr":5.5364,"F":0.916,"P.Value":0.4076,"adj.P.Val":0.5471},{"tissuedysplasia":2.3723,"tissuetumour":4.1327,"AveExpr":-0.3322,"F":21.2701,"P.Value":3.4431e-07,"adj.P.Val":0},{"tissuedysplasia":-0.0881,"tissuetumour":0.6658,"AveExpr":5.686,"F":11.7167,"P.Value":0.0001,"adj.P.Val":0.0012},{"tissuedysplasia":0.2062,"tissuetumour":0.5569,"AveExpr":4.3812,"F":9.2109,"P.Value":0.0005,"adj.P.Val":0.0043},{"tissuedysplasia":0.0916,"tissuetumour":0.8016,"AveExpr":3.0304,"F":9.0672,"P.Value":0.0005,"adj.P.Val":0.0047},{"tissuedysplasia":-0.223,"tissuetumour":0.7064,"AveExpr":-1.886,"F":1.4203,"P.Value":0.2525,"adj.P.Val":0.3954},{"tissuedysplasia":-0.0626,"tissuetumour":0.8327,"AveExpr":1.3982,"F":6.0329,"P.Value":0.0048,"adj.P.Val":0.0245},{"tissuedysplasia":0.854,"tissuetumour":1.6121,"AveExpr":-0.9267,"F":4.185,"P.Value":0.0217,"adj.P.Val":0.0726},{"tissuedysplasia":0.3559,"tissuetumour":1.0513,"AveExpr":0.8754,"F":4.293,"P.Value":0.0198,"adj.P.Val":0.0681},{"tissuedysplasia":-0.311,"tissuetumour":-0.0076,"AveExpr":4.0012,"F":2.2892,"P.Value":0.1133,"adj.P.Val":0.2296},{"tissuedysplasia":0.2726,"tissuetumour":1.2447,"AveExpr":1.0999,"F":7.5307,"P.Value":0.0015,"adj.P.Val":0.0106},{"tissuedysplasia":0.3339,"tissuetumour":1.027,"AveExpr":-3.367,"F":0.7463,"P.Value":0.48,"adj.P.Val":0.6095},{"tissuedysplasia":1.0996,"tissuetumour":3.8488,"AveExpr":-5.5262,"F":8.1905,"P.Value":0.0009,"adj.P.Val":0.0074},{"tissuedysplasia":-0.481,"tissuetumour":-0.308,"AveExpr":6.3473,"F":3.5819,"P.Value":0.0363,"adj.P.Val":0.1042},{"tissuedysplasia":0.3896,"tissuetumour":1.2143,"AveExpr":1.548,"F":7.5309,"P.Value":0.0015,"adj.P.Val":0.0106},{"tissuedysplasia":0.2982,"tissuetumour":0.8268,"AveExpr":3.4796,"F":13.5643,"P.Value":0,"adj.P.Val":0.0005},{"tissuedysplasia":0.2245,"tissuetumour":0.6614,"AveExpr":4.1608,"F":12.9922,"P.Value":0,"adj.P.Val":0.0006},{"tissuedysplasia":0.1361,"tissuetumour":0.8811,"AveExpr":3.0505,"F":10.1609,"P.Value":0.0002,"adj.P.Val":0.0027},{"tissuedysplasia":0.691,"tissuetumour":1.3074,"AveExpr":3.0945,"F":24.9267,"P.Value":5.7781e-08,"adj.P.Val":4.7906e-06},{"tissuedysplasia":0.2893,"tissuetumour":0.8103,"AveExpr":5.6127,"F":18.2145,"P.Value":1.7247e-06,"adj.P.Val":0.0001},{"tissuedysplasia":0.4055,"tissuetumour":0.6167,"AveExpr":1.4341,"F":2.7043,"P.Value":0.078,"adj.P.Val":0.1775},{"tissuedysplasia":0.0145,"tissuetumour":-0.0634,"AveExpr":4.4612,"F":0.3118,"P.Value":0.7337,"adj.P.Val":0.8103},{"tissuedysplasia":0.2745,"tissuetumour":0.297,"AveExpr":6.0837,"F":4.933,"P.Value":0.0117,"adj.P.Val":0.0462},{"tissuedysplasia":0.2958,"tissuetumour":0.6749,"AveExpr":4.5684,"F":10.9502,"P.Value":0.0001,"adj.P.Val":0.0018},{"tissuedysplasia":1.651,"tissuetumour":4.4588,"AveExpr":0.8636,"F":33.9291,"P.Value":1.2159e-09,"adj.P.Val":2.8795e-07},{"tissuedysplasia":0.7387,"tissuetumour":1.4775,"AveExpr":1.5928,"F":5.5241,"P.Value":0.0072,"adj.P.Val":0.0329},{"tissuedysplasia":0.3093,"tissuetumour":0.6613,"AveExpr":4.3688,"F":10.4012,"P.Value":0.0002,"adj.P.Val":0.0023},{"tissuedysplasia":0.561,"tissuetumour":1.7008,"AveExpr":-0.6699,"F":5.7056,"P.Value":0.0063,"adj.P.Val":0.0296},{"tissuedysplasia":1.1828,"tissuetumour":1.569,"AveExpr":2.0812,"F":9.606,"P.Value":0.0003,"adj.P.Val":0.0035},{"tissuedysplasia":0.3001,"tissuetumour":0.2106,"AveExpr":-2.5365,"F":0.0717,"P.Value":0.9309,"adj.P.Val":0.9518},{"tissuedysplasia":0.0519,"tissuetumour":0.3072,"AveExpr":5.2946,"F":4.2166,"P.Value":0.0211,"adj.P.Val":0.0712},{"tissuedysplasia":0.0899,"tissuetumour":0.6937,"AveExpr":2.69,"F":10.1759,"P.Value":0.0002,"adj.P.Val":0.0026},{"tissuedysplasia":0.0752,"tissuetumour":0.6274,"AveExpr":2.4515,"F":7.9819,"P.Value":0.0011,"adj.P.Val":0.0083},{"tissuedysplasia":0.1877,"tissuetumour":0.2564,"AveExpr":-2.2659,"F":0.082,"P.Value":0.9214,"adj.P.Val":0.9452},{"tissuedysplasia":0.8879,"tissuetumour":1.619,"AveExpr":-1.781,"F":2.0511,"P.Value":0.1407,"adj.P.Val":0.2656},{"tissuedysplasia":0.2045,"tissuetumour":0.5202,"AveExpr":5.6025,"F":13.075,"P.Value":0,"adj.P.Val":0.0006},{"tissuedysplasia":-0.057,"tissuetumour":0.1983,"AveExpr":4.2639,"F":1.9251,"P.Value":0.1579,"adj.P.Val":0.2876},{"tissuedysplasia":0.037,"tissuetumour":0.949,"AveExpr":1.4028,"F":8.3976,"P.Value":0.0008,"adj.P.Val":0.0067},{"tissuedysplasia":0.1479,"tissuetumour":0.4674,"AveExpr":5.823,"F":14.0715,"P.Value":0,"adj.P.Val":0.0004},{"tissuedysplasia":0.1004,"tissuetumour":1.1638,"AveExpr":3.2774,"F":20.3697,"P.Value":5.4685e-07,"adj.P.Val":0},{"tissuedysplasia":0.1148,"tissuetumour":0.8115,"AveExpr":2.7429,"F":10.8975,"P.Value":0.0001,"adj.P.Val":0.0018},{"tissuedysplasia":0.061,"tissuetumour":0.6176,"AveExpr":4.1008,"F":7.62,"P.Value":0.0014,"adj.P.Val":0.0102},{"tissuedysplasia":-0.025,"tissuetumour":0.737,"AveExpr":-1.6656,"F":1.2398,"P.Value":0.2993,"adj.P.Val":0.4437},{"tissuedysplasia":0.0481,"tissuetumour":-0.033,"AveExpr":5.864,"F":0.355,"P.Value":0.7032,"adj.P.Val":0.7874},{"tissuedysplasia":0.1235,"tissuetumour":0.223,"AveExpr":5.1114,"F":2.9075,"P.Value":0.0652,"adj.P.Val":0.1568},{"tissuedysplasia":0.5798,"tissuetumour":1.1408,"AveExpr":0.1552,"F":3.5489,"P.Value":0.0372,"adj.P.Val":0.1062},{"tissuedysplasia":0.3826,"tissuetumour":0.1189,"AveExpr":2.8049,"F":2.667,"P.Value":0.0807,"adj.P.Val":0.1818},{"tissuedysplasia":0.0741,"tissuetumour":0.2895,"AveExpr":4.8671,"F":2.6148,"P.Value":0.0845,"adj.P.Val":0.1877},{"tissuedysplasia":0.2979,"tissuetumour":1.3492,"AveExpr":4.7437,"F":30.8159,"P.Value":4.2867e-09,"adj.P.Val":7.5849e-07},{"tissuedysplasia":0.1255,"tissuetumour":-0.2469,"AveExpr":6.4565,"F":7.1861,"P.Value":0.002,"adj.P.Val":0.0129},{"tissuedysplasia":0.5798,"tissuetumour":1.0603,"AveExpr":1.0142,"F":6.6183,"P.Value":0.0031,"adj.P.Val":0.0177},{"tissuedysplasia":0.653,"tissuetumour":1.1215,"AveExpr":2.405,"F":6.6402,"P.Value":0.003,"adj.P.Val":0.0175},{"tissuedysplasia":0.0225,"tissuetumour":0.5558,"AveExpr":4.6876,"F":9.5666,"P.Value":0.0004,"adj.P.Val":0.0036},{"tissuedysplasia":0.2372,"tissuetumour":0.5981,"AveExpr":3.197,"F":3.905,"P.Value":0.0275,"adj.P.Val":0.0857},{"tissuedysplasia":0.2784,"tissuetumour":0.7677,"AveExpr":3.8183,"F":16.4753,"P.Value":4.5618e-06,"adj.P.Val":0.0001},{"tissuedysplasia":0.4314,"tissuetumour":0.2134,"AveExpr":-2.0246,"F":0.4654,"P.Value":0.631,"adj.P.Val":0.7333},{"tissuedysplasia":0.9343,"tissuetumour":1.2648,"AveExpr":3.6255,"F":13.2262,"P.Value":0,"adj.P.Val":0.0006},{"tissuedysplasia":0.3323,"tissuetumour":0.9432,"AveExpr":2.8449,"F":13.8069,"P.Value":0,"adj.P.Val":0.0004},{"tissuedysplasia":0.2345,"tissuetumour":0.5266,"AveExpr":3.9745,"F":6.2827,"P.Value":0.004,"adj.P.Val":0.0213},{"tissuedysplasia":0.9947,"tissuetumour":1.2041,"AveExpr":1.3878,"F":9.431,"P.Value":0.0004,"adj.P.Val":0.0039},{"tissuedysplasia":0.6417,"tissuetumour":1.8746,"AveExpr":3.5092,"F":32.2074,"P.Value":2.4191e-09,"adj.P.Val":4.8412e-07},{"tissuedysplasia":0.5068,"tissuetumour":0.8955,"AveExpr":2.221,"F":4.4756,"P.Value":0.017,"adj.P.Val":0.061},{"tissuedysplasia":0.211,"tissuetumour":0.641,"AveExpr":2.9363,"F":7.1912,"P.Value":0.002,"adj.P.Val":0.0129},{"tissuedysplasia":0.5702,"tissuetumour":0.6424,"AveExpr":4.5514,"F":11.1809,"P.Value":0.0001,"adj.P.Val":0.0016},{"tissuedysplasia":-0.2673,"tissuetumour":-0.2109,"AveExpr":2.3314,"F":0.454,"P.Value":0.638,"adj.P.Val":0.7385},{"tissuedysplasia":0.3976,"tissuetumour":0.0977,"AveExpr":0.7281,"F":1.1536,"P.Value":0.3249,"adj.P.Val":0.469},{"tissuedysplasia":-0.0225,"tissuetumour":0.309,"AveExpr":5.1357,"F":4.0263,"P.Value":0.0248,"adj.P.Val":0.0797},{"tissuedysplasia":0.2948,"tissuetumour":0.6687,"AveExpr":4.5117,"F":13.7466,"P.Value":0,"adj.P.Val":0.0004},{"tissuedysplasia":0.1339,"tissuetumour":0.6966,"AveExpr":4.3895,"F":11.3538,"P.Value":0.0001,"adj.P.Val":0.0014},{"tissuedysplasia":0.121,"tissuetumour":1.9753,"AveExpr":-0.5116,"F":13.685,"P.Value":0,"adj.P.Val":0.0005},{"tissuedysplasia":0.6075,"tissuetumour":2.299,"AveExpr":3.3177,"F":36.9629,"P.Value":3.8031e-10,"adj.P.Val":1.306e-07},{"tissuedysplasia":0.1386,"tissuetumour":0.3843,"AveExpr":5.1924,"F":6.5151,"P.Value":0.0033,"adj.P.Val":0.0187},{"tissuedysplasia":0.786,"tissuetumour":0.4411,"AveExpr":4.6046,"F":10.557,"P.Value":0.0002,"adj.P.Val":0.0022},{"tissuedysplasia":-0.1625,"tissuetumour":0.0139,"AveExpr":3.7856,"F":0.6623,"P.Value":0.5207,"adj.P.Val":0.6444},{"tissuedysplasia":0.2486,"tissuetumour":0.3512,"AveExpr":3.8685,"F":3.6656,"P.Value":0.0337,"adj.P.Val":0.0988},{"tissuedysplasia":0.4198,"tissuetumour":0.6537,"AveExpr":2.8908,"F":5.3181,"P.Value":0.0085,"adj.P.Val":0.0371},{"tissuedysplasia":-0.3741,"tissuetumour":-0.2517,"AveExpr":6.2034,"F":2.8509,"P.Value":0.0685,"adj.P.Val":0.1624},{"tissuedysplasia":0.3849,"tissuetumour":0.5152,"AveExpr":4.4045,"F":9.4765,"P.Value":0.0004,"adj.P.Val":0.0038},{"tissuedysplasia":-0.2316,"tissuetumour":0.4826,"AveExpr":0.9042,"F":3.6271,"P.Value":0.0348,"adj.P.Val":0.1011},{"tissuedysplasia":0.2031,"tissuetumour":0.7299,"AveExpr":-3.0272,"F":0.7771,"P.Value":0.4659,"adj.P.Val":0.5975},{"tissuedysplasia":-0.4725,"tissuetumour":1.7242,"AveExpr":-4.7587,"F":2.9235,"P.Value":0.0643,"adj.P.Val":0.1554},{"tissuedysplasia":0.0125,"tissuetumour":0.4017,"AveExpr":2.9184,"F":3.5487,"P.Value":0.0373,"adj.P.Val":0.1062},{"tissuedysplasia":-0.1438,"tissuetumour":-0.6255,"AveExpr":5.9738,"F":13.4155,"P.Value":0,"adj.P.Val":0.0005},{"tissuedysplasia":0.1733,"tissuetumour":0.1353,"AveExpr":4.1831,"F":0.7772,"P.Value":0.4659,"adj.P.Val":0.5975},{"tissuedysplasia":0.2435,"tissuetumour":0.7904,"AveExpr":6.0134,"F":15.9777,"P.Value":6.0745e-06,"adj.P.Val":0.0002},{"tissuedysplasia":0.1961,"tissuetumour":0.8469,"AveExpr":3.3026,"F":15.7181,"P.Value":7.0643e-06,"adj.P.Val":0.0002},{"tissuedysplasia":0.1889,"tissuetumour":0.0551,"AveExpr":0.5124,"F":0.1518,"P.Value":0.8596,"adj.P.Val":0.9012},{"tissuedysplasia":0.3762,"tissuetumour":1.3361,"AveExpr":1.8793,"F":18.9884,"P.Value":1.1339e-06,"adj.P.Val":0},{"tissuedysplasia":0.2156,"tissuetumour":0.3634,"AveExpr":4.2968,"F":4.5954,"P.Value":0.0154,"adj.P.Val":0.0567},{"tissuedysplasia":0.0384,"tissuetumour":0.2767,"AveExpr":3.1679,"F":1.8313,"P.Value":0.1722,"adj.P.Val":0.3051},{"tissuedysplasia":0.1658,"tissuetumour":0.1735,"AveExpr":4.4612,"F":0.5924,"P.Value":0.5574,"adj.P.Val":0.674},{"tissuedysplasia":0.2554,"tissuetumour":-0.225,"AveExpr":-2.7116,"F":0.1961,"P.Value":0.8227,"adj.P.Val":0.8758},{"tissuedysplasia":0.3528,"tissuetumour":1.0975,"AveExpr":0.0874,"F":5.0214,"P.Value":0.0109,"adj.P.Val":0.044},{"tissuedysplasia":0.2172,"tissuetumour":0.5291,"AveExpr":0.9628,"F":1.1049,"P.Value":0.3402,"adj.P.Val":0.4843},{"tissuedysplasia":-0.1576,"tissuetumour":-0.588,"AveExpr":3.9808,"F":3.7944,"P.Value":0.0302,"adj.P.Val":0.0916},{"tissuedysplasia":-0.4438,"tissuetumour":0.2473,"AveExpr":2.3643,"F":3.4646,"P.Value":0.0401,"adj.P.Val":0.1117},{"tissuedysplasia":0.8186,"tissuetumour":0.8468,"AveExpr":-0.6746,"F":2.0962,"P.Value":0.135,"adj.P.Val":0.2588},{"tissuedysplasia":0.3224,"tissuetumour":0.6182,"AveExpr":2.5297,"F":3.2131,"P.Value":0.0498,"adj.P.Val":0.1304},{"tissuedysplasia":0.0034,"tissuetumour":0.3277,"AveExpr":1.2427,"F":0.9751,"P.Value":0.3851,"adj.P.Val":0.5267},{"tissuedysplasia":0.1155,"tissuetumour":-0.0772,"AveExpr":9.2438,"F":2.3496,"P.Value":0.1086,"adj.P.Val":0.2231},{"tissuedysplasia":0.0702,"tissuetumour":0.4349,"AveExpr":0.0684,"F":0.8593,"P.Value":0.4305,"adj.P.Val":0.5676},{"tissuedysplasia":0.2108,"tissuetumour":0.1131,"AveExpr":6.5664,"F":1.8528,"P.Value":0.1688,"adj.P.Val":0.301},{"tissuedysplasia":0.4382,"tissuetumour":0.8486,"AveExpr":1.352,"F":7.4374,"P.Value":0.0016,"adj.P.Val":0.0112},{"tissuedysplasia":0.1675,"tissuetumour":0.7153,"AveExpr":2.4947,"F":8.5255,"P.Value":0.0007,"adj.P.Val":0.0062},{"tissuedysplasia":0.1485,"tissuetumour":0.4505,"AveExpr":5.2639,"F":8.2079,"P.Value":0.0009,"adj.P.Val":0.0073},{"tissuedysplasia":0.6321,"tissuetumour":0.2854,"AveExpr":-1.8046,"F":0.5912,"P.Value":0.558,"adj.P.Val":0.6745},{"tissuedysplasia":0.6174,"tissuetumour":0.7746,"AveExpr":5.9519,"F":5.7254,"P.Value":0.0064,"adj.P.Val":0.0303},{"tissuedysplasia":0.0738,"tissuetumour":0.2896,"AveExpr":4.6206,"F":3.1215,"P.Value":0.054,"adj.P.Val":0.1379},{"tissuedysplasia":0.3228,"tissuetumour":0.3445,"AveExpr":5.1063,"F":3.8464,"P.Value":0.0289,"adj.P.Val":0.0888},{"tissuedysplasia":0.2387,"tissuetumour":0.2156,"AveExpr":5.4328,"F":4.4706,"P.Value":0.0171,"adj.P.Val":0.0612},{"tissuedysplasia":0.1056,"tissuetumour":0.2725,"AveExpr":5.8955,"F":2.2291,"P.Value":0.1196,"adj.P.Val":0.238},{"tissuedysplasia":0.0853,"tissuetumour":0.3537,"AveExpr":4.6929,"F":4.4499,"P.Value":0.0174,"adj.P.Val":0.062},{"tissuedysplasia":0.2552,"tissuetumour":0.0001,"AveExpr":1.6158,"F":0.7223,"P.Value":0.4913,"adj.P.Val":0.6194},{"tissuedysplasia":-0.2035,"tissuetumour":0.1675,"AveExpr":6.6842,"F":2.656,"P.Value":0.0815,"adj.P.Val":0.183},{"tissuedysplasia":0.1126,"tissuetumour":0.4676,"AveExpr":5.1615,"F":11.0775,"P.Value":0.0001,"adj.P.Val":0.0017},{"tissuedysplasia":-0.1028,"tissuetumour":0.5313,"AveExpr":5.8989,"F":10.795,"P.Value":0.0002,"adj.P.Val":0.0019},{"tissuedysplasia":0.0148,"tissuetumour":0.2398,"AveExpr":6.0016,"F":2.4987,"P.Value":0.0938,"adj.P.Val":0.2019},{"tissuedysplasia":0.1054,"tissuetumour":0.6875,"AveExpr":5.414,"F":16.8293,"P.Value":3.7293e-06,"adj.P.Val":0.0001},{"tissuedysplasia":0.1432,"tissuetumour":0.5821,"AveExpr":1.7808,"F":2.4971,"P.Value":0.0939,"adj.P.Val":0.2021},{"tissuedysplasia":-0.0144,"tissuetumour":-0.4232,"AveExpr":-3.954,"F":0.1176,"P.Value":0.8893,"adj.P.Val":0.9223},{"tissuedysplasia":1.5039,"tissuetumour":2.6858,"AveExpr":-3.9596,"F":3.8546,"P.Value":0.0287,"adj.P.Val":0.0884},{"tissuedysplasia":0.4431,"tissuetumour":1.4103,"AveExpr":3.6266,"F":35.8899,"P.Value":5.6967e-10,"adj.P.Val":1.7401e-07},{"tissuedysplasia":0.6501,"tissuetumour":1.2955,"AveExpr":1.4367,"F":11.3731,"P.Value":0.0001,"adj.P.Val":0.0014},{"tissuedysplasia":0.1166,"tissuetumour":0.415,"AveExpr":5.2546,"F":8.53,"P.Value":0.0007,"adj.P.Val":0.0062},{"tissuedysplasia":0.5013,"tissuetumour":0.358,"AveExpr":0.8198,"F":1.9142,"P.Value":0.1595,"adj.P.Val":0.2894},{"tissuedysplasia":0.2089,"tissuetumour":0.7498,"AveExpr":-1.9349,"F":0.6832,"P.Value":0.5103,"adj.P.Val":0.6355},{"tissuedysplasia":-0.0454,"tissuetumour":0.0331,"AveExpr":6.1641,"F":0.2419,"P.Value":0.7862,"adj.P.Val":0.849},{"tissuedysplasia":0.2563,"tissuetumour":0.3315,"AveExpr":4.7395,"F":4.8394,"P.Value":0.0126,"adj.P.Val":0.0489},{"tissuedysplasia":-0.1814,"tissuetumour":0.167,"AveExpr":5.0702,"F":3.359,"P.Value":0.0439,"adj.P.Val":0.1191},{"tissuedysplasia":0.3509,"tissuetumour":0.844,"AveExpr":-0.444,"F":1.6928,"P.Value":0.1958,"adj.P.Val":0.3324},{"tissuedysplasia":0.3682,"tissuetumour":0.6975,"AveExpr":4.4501,"F":17.9723,"P.Value":1.9698e-06,"adj.P.Val":0.0001},{"tissuedysplasia":0.2386,"tissuetumour":0.2997,"AveExpr":0.922,"F":0.5146,"P.Value":0.6013,"adj.P.Val":0.7098},{"tissuedysplasia":0.0329,"tissuetumour":-0.2623,"AveExpr":3.6951,"F":1.2533,"P.Value":0.2956,"adj.P.Val":0.4399},{"tissuedysplasia":0.2305,"tissuetumour":0.475,"AveExpr":1.7685,"F":2.7087,"P.Value":0.0777,"adj.P.Val":0.1771},{"tissuedysplasia":0.3863,"tissuetumour":0.6689,"AveExpr":4.7915,"F":12.5012,"P.Value":0.0001,"adj.P.Val":0.0008},{"tissuedysplasia":0.7235,"tissuetumour":1.1155,"AveExpr":4.3712,"F":29.2929,"P.Value":8.1605e-09,"adj.P.Val":1.2113e-06},{"tissuedysplasia":1.4577,"tissuetumour":1.4355,"AveExpr":-2.9573,"F":1.6852,"P.Value":0.1972,"adj.P.Val":0.334},{"tissuedysplasia":0.112,"tissuetumour":0.9022,"AveExpr":2.7016,"F":13.4101,"P.Value":0,"adj.P.Val":0.0005},{"tissuedysplasia":-0.4644,"tissuetumour":-0.2072,"AveExpr":2.7252,"F":2.9803,"P.Value":0.0611,"adj.P.Val":0.1502},{"tissuedysplasia":0.7455,"tissuetumour":1.5148,"AveExpr":-0.6301,"F":4.3239,"P.Value":0.0193,"adj.P.Val":0.0669},{"tissuedysplasia":-0.0535,"tissuetumour":0.4725,"AveExpr":5.3694,"F":4.0849,"P.Value":0.0236,"adj.P.Val":0.077},{"tissuedysplasia":0.3063,"tissuetumour":0.3189,"AveExpr":3.7207,"F":1.5234,"P.Value":0.2292,"adj.P.Val":0.3704},{"tissuedysplasia":0.3498,"tissuetumour":1.3015,"AveExpr":2.565,"F":19.169,"P.Value":1.0293e-06,"adj.P.Val":0},{"tissuedysplasia":-0.2041,"tissuetumour":0.7009,"AveExpr":-2.0519,"F":1.4818,"P.Value":0.2384,"adj.P.Val":0.3803},{"tissuedysplasia":0.3709,"tissuetumour":0.9377,"AveExpr":0.1227,"F":2.2725,"P.Value":0.115,"adj.P.Val":0.2319},{"tissuedysplasia":-0.3464,"tissuetumour":-0.2513,"AveExpr":7.7596,"F":5.5853,"P.Value":0.0069,"adj.P.Val":0.0318},{"tissuedysplasia":0.581,"tissuetumour":0.8931,"AveExpr":1.7799,"F":5.8546,"P.Value":0.0056,"adj.P.Val":0.0272},{"tissuedysplasia":-0.1883,"tissuetumour":-0.018,"AveExpr":6.5658,"F":2.6098,"P.Value":0.0849,"adj.P.Val":0.1882},{"tissuedysplasia":-0.6346,"tissuetumour":-1.4131,"AveExpr":2.4633,"F":16.5758,"P.Value":4.3074e-06,"adj.P.Val":0.0001},{"tissuedysplasia":0.1319,"tissuetumour":0.6864,"AveExpr":3.2278,"F":10.0102,"P.Value":0.0003,"adj.P.Val":0.0029},{"tissuedysplasia":-0.0379,"tissuetumour":0.0497,"AveExpr":5.5447,"F":0.5324,"P.Value":0.5909,"adj.P.Val":0.7012},{"tissuedysplasia":-0.057,"tissuetumour":-0.0255,"AveExpr":5.975,"F":0.2857,"P.Value":0.7528,"adj.P.Val":0.8245},{"tissuedysplasia":0.3904,"tissuetumour":0.5677,"AveExpr":4.6659,"F":11.0901,"P.Value":0.0001,"adj.P.Val":0.0017},{"tissuedysplasia":0.9686,"tissuetumour":1.6703,"AveExpr":2.5504,"F":10.1248,"P.Value":0.0002,"adj.P.Val":0.0027},{"tissuedysplasia":0.3759,"tissuetumour":1.9917,"AveExpr":-5.5244,"F":2.8064,"P.Value":0.0713,"adj.P.Val":0.167},{"tissuedysplasia":1.7438,"tissuetumour":3.0428,"AveExpr":5.6984,"F":42.6536,"P.Value":1.8056e-10,"adj.P.Val":7.8203e-08},{"tissuedysplasia":0.6348,"tissuetumour":1.2594,"AveExpr":1.3524,"F":8.428,"P.Value":0.0008,"adj.P.Val":0.0066},{"tissuedysplasia":0.3735,"tissuetumour":1.2982,"AveExpr":9.6156,"F":26.5832,"P.Value":7.9462e-08,"adj.P.Val":6.0363e-06},{"tissuedysplasia":1.1317,"tissuetumour":1.2788,"AveExpr":-0.0117,"F":4.2795,"P.Value":0.02,"adj.P.Val":0.0686},{"tissuedysplasia":0.2324,"tissuetumour":0.3951,"AveExpr":4.075,"F":6.3533,"P.Value":0.0038,"adj.P.Val":0.0205},{"tissuedysplasia":-0.0159,"tissuetumour":0.4873,"AveExpr":3.5979,"F":7.386,"P.Value":0.0017,"adj.P.Val":0.0116},{"tissuedysplasia":0.7723,"tissuetumour":0.9329,"AveExpr":3.1145,"F":12.1921,"P.Value":0.0001,"adj.P.Val":0.001},{"tissuedysplasia":0.7126,"tissuetumour":0.9425,"AveExpr":3.1354,"F":6.1245,"P.Value":0.0045,"adj.P.Val":0.0233},{"tissuedysplasia":0.0401,"tissuetumour":0.1442,"AveExpr":5.9404,"F":0.8433,"P.Value":0.4371,"adj.P.Val":0.5738},{"tissuedysplasia":0.1798,"tissuetumour":0.0899,"AveExpr":7.4177,"F":2.4795,"P.Value":0.0954,"adj.P.Val":0.2042},{"tissuedysplasia":0.0149,"tissuetumour":0.4248,"AveExpr":4.7181,"F":6.103,"P.Value":0.0046,"adj.P.Val":0.0236},{"tissuedysplasia":0.2729,"tissuetumour":0.2495,"AveExpr":2.7151,"F":0.5372,"P.Value":0.5881,"adj.P.Val":0.6991},{"tissuedysplasia":1.4132,"tissuetumour":3.5667,"AveExpr":-6.0981,"F":3.8437,"P.Value":0.0289,"adj.P.Val":0.0889},{"tissuedysplasia":0.324,"tissuetumour":0.9663,"AveExpr":5.7408,"F":30.0253,"P.Value":5.9737e-09,"adj.P.Val":9.7095e-07},{"tissuedysplasia":0.8146,"tissuetumour":1.1682,"AveExpr":1.8595,"F":8.8896,"P.Value":0.0006,"adj.P.Val":0.0051},{"tissuedysplasia":0.8898,"tissuetumour":0.9322,"AveExpr":3.8169,"F":13.8703,"P.Value":0,"adj.P.Val":0.0004},{"tissuedysplasia":0.413,"tissuetumour":0.5843,"AveExpr":2.9843,"F":6.4588,"P.Value":0.0035,"adj.P.Val":0.0193},{"tissuedysplasia":-0.0471,"tissuetumour":0.6066,"AveExpr":-8.1535,"F":0.1547,"P.Value":0.8572,"adj.P.Val":0.8996},{"tissuedysplasia":0.1787,"tissuetumour":0.4374,"AveExpr":4.6697,"F":5.8607,"P.Value":0.0055,"adj.P.Val":0.0271},{"tissuedysplasia":0.0697,"tissuetumour":0.6258,"AveExpr":1.7729,"F":3.1546,"P.Value":0.0524,"adj.P.Val":0.135},{"tissuedysplasia":0.0815,"tissuetumour":-0.1315,"AveExpr":5.9209,"F":2.0746,"P.Value":0.1377,"adj.P.Val":0.2621},{"tissuedysplasia":1.3912,"tissuetumour":1.9554,"AveExpr":2.2577,"F":26.897,"P.Value":2.3378e-08,"adj.P.Val":2.5125e-06},{"tissuedysplasia":0.1149,"tissuetumour":-0.0103,"AveExpr":3.2332,"F":0.6119,"P.Value":0.5469,"adj.P.Val":0.6654},{"tissuedysplasia":0.0965,"tissuetumour":0.012,"AveExpr":7.5835,"F":1.3361,"P.Value":0.2733,"adj.P.Val":0.4177},{"tissuedysplasia":0.2669,"tissuetumour":0.8268,"AveExpr":4.2548,"F":21.3419,"P.Value":3.3198e-07,"adj.P.Val":0},{"tissuedysplasia":0.0195,"tissuetumour":0.5168,"AveExpr":-0.6715,"F":0.7637,"P.Value":0.472,"adj.P.Val":0.6024},{"tissuedysplasia":0.1371,"tissuetumour":1.0308,"AveExpr":6.0081,"F":15.5347,"P.Value":7.8636e-06,"adj.P.Val":0.0002},{"tissuedysplasia":0.2137,"tissuetumour":0.8004,"AveExpr":5.6847,"F":20.3636,"P.Value":5.4859e-07,"adj.P.Val":0},{"tissuedysplasia":0.5597,"tissuetumour":0.7093,"AveExpr":3.2511,"F":9.2743,"P.Value":0.0004,"adj.P.Val":0.0042},{"tissuedysplasia":0.2047,"tissuetumour":0.3319,"AveExpr":5.2853,"F":4.5229,"P.Value":0.0164,"adj.P.Val":0.0592},{"tissuedysplasia":0.1783,"tissuetumour":0.4445,"AveExpr":5.2161,"F":8.1441,"P.Value":0.001,"adj.P.Val":0.0076},{"tissuedysplasia":0.3651,"tissuetumour":0.6641,"AveExpr":4.2197,"F":10.6458,"P.Value":0.0002,"adj.P.Val":0.0021},{"tissuedysplasia":-0.4512,"tissuetumour":0.1231,"AveExpr":3.265,"F":3.9477,"P.Value":0.0265,"adj.P.Val":0.0835},{"tissuedysplasia":0.1532,"tissuetumour":0.2911,"AveExpr":5.1494,"F":3.2176,"P.Value":0.0496,"adj.P.Val":0.1301},{"tissuedysplasia":0.1769,"tissuetumour":0.0555,"AveExpr":4.8126,"F":1.1196,"P.Value":0.3355,"adj.P.Val":0.4799},{"tissuedysplasia":0.0013,"tissuetumour":0.0516,"AveExpr":5.5748,"F":0.1584,"P.Value":0.854,"adj.P.Val":0.8975},{"tissuedysplasia":0.079,"tissuetumour":0.1747,"AveExpr":5.0761,"F":1.4086,"P.Value":0.2553,"adj.P.Val":0.3983},{"tissuedysplasia":-0.3419,"tissuetumour":-0.1108,"AveExpr":-2.8991,"F":0.0816,"P.Value":0.9218,"adj.P.Val":0.9455},{"tissuedysplasia":0.4096,"tissuetumour":1.2867,"AveExpr":2.6882,"F":16.75,"P.Value":3.9009e-06,"adj.P.Val":0.0001},{"tissuedysplasia":-0.0768,"tissuetumour":-0.1577,"AveExpr":2.5928,"F":0.1273,"P.Value":0.8808,"adj.P.Val":0.9163},{"tissuedysplasia":-0.0434,"tissuetumour":0.0518,"AveExpr":3.8339,"F":0.2013,"P.Value":0.8184,"adj.P.Val":0.8728},{"tissuedysplasia":0.1144,"tissuetumour":0.5051,"AveExpr":6.058,"F":10.194,"P.Value":0.0002,"adj.P.Val":0.0026},{"tissuedysplasia":-0.0617,"tissuetumour":-0.2962,"AveExpr":3.9055,"F":2.6177,"P.Value":0.0843,"adj.P.Val":0.1873},{"tissuedysplasia":0.0796,"tissuetumour":0.1035,"AveExpr":4.2355,"F":0.3949,"P.Value":0.6761,"adj.P.Val":0.7676},{"tissuedysplasia":0.2474,"tissuetumour":0.1691,"AveExpr":-0.6319,"F":0.1252,"P.Value":0.8827,"adj.P.Val":0.9177},{"tissuedysplasia":1.071,"tissuetumour":1.2983,"AveExpr":-0.0521,"F":6.3196,"P.Value":0.0039,"adj.P.Val":0.0209},{"tissuedysplasia":1.4603,"tissuetumour":1.4117,"AveExpr":2.0085,"F":13.9037,"P.Value":0,"adj.P.Val":0.0004},{"tissuedysplasia":0.0051,"tissuetumour":-0.0222,"AveExpr":5.4057,"F":0.0415,"P.Value":0.9593,"adj.P.Val":0.9722},{"tissuedysplasia":0.043,"tissuetumour":0.2631,"AveExpr":4.3233,"F":2.14,"P.Value":0.1297,"adj.P.Val":0.2514},{"tissuedysplasia":0.2727,"tissuetumour":0.5012,"AveExpr":2.3475,"F":3.0281,"P.Value":0.0586,"adj.P.Val":0.1459},{"tissuedysplasia":0.1542,"tissuetumour":0.2927,"AveExpr":3.8461,"F":1.8098,"P.Value":0.1757,"adj.P.Val":0.3091},{"tissuedysplasia":0.4899,"tissuetumour":0.2645,"AveExpr":1.8426,"F":1.5882,"P.Value":0.2158,"adj.P.Val":0.3553},{"tissuedysplasia":0.2157,"tissuetumour":-0.0739,"AveExpr":2.4277,"F":0.9111,"P.Value":0.4095,"adj.P.Val":0.5487},{"tissuedysplasia":0.1132,"tissuetumour":-0.1691,"AveExpr":5.5217,"F":3.2358,"P.Value":0.0489,"adj.P.Val":0.1286},{"tissuedysplasia":-0.0127,"tissuetumour":0.3339,"AveExpr":3.2089,"F":3.7047,"P.Value":0.0326,"adj.P.Val":0.0966},{"tissuedysplasia":0.3629,"tissuetumour":0.2666,"AveExpr":6.7037,"F":6.7046,"P.Value":0.0029,"adj.P.Val":0.0168},{"tissuedysplasia":0.3907,"tissuetumour":0.7969,"AveExpr":5.715,"F":7.6992,"P.Value":0.0014,"adj.P.Val":0.0097},{"tissuedysplasia":0.1155,"tissuetumour":1.4217,"AveExpr":1.4531,"F":7.5641,"P.Value":0.0015,"adj.P.Val":0.0105},{"tissuedysplasia":0.1924,"tissuetumour":-0.0177,"AveExpr":5.6054,"F":1.92,"P.Value":0.1587,"adj.P.Val":0.2885},{"tissuedysplasia":0.6018,"tissuetumour":0.4988,"AveExpr":4.2531,"F":13.0672,"P.Value":0,"adj.P.Val":0.0006},{"tissuedysplasia":0.6581,"tissuetumour":0.5409,"AveExpr":5.1079,"F":10.086,"P.Value":0.0002,"adj.P.Val":0.0028},{"tissuedysplasia":0.9038,"tissuetumour":0.7946,"AveExpr":1.6015,"F":3.8353,"P.Value":0.0291,"adj.P.Val":0.0894},{"tissuedysplasia":0.2061,"tissuetumour":-0.2658,"AveExpr":4.4691,"F":3.9783,"P.Value":0.0258,"adj.P.Val":0.082},{"tissuedysplasia":0.1318,"tissuetumour":0.0782,"AveExpr":1.3811,"F":0.1194,"P.Value":0.8878,"adj.P.Val":0.9214},{"tissuedysplasia":-0.0207,"tissuetumour":-0.3588,"AveExpr":4.4968,"F":4.183,"P.Value":0.0217,"adj.P.Val":0.0727},{"tissuedysplasia":0.4288,"tissuetumour":1.878,"AveExpr":-0.9753,"F":11.2581,"P.Value":0.0001,"adj.P.Val":0.0015},{"tissuedysplasia":0.9678,"tissuetumour":1.7336,"AveExpr":2.7218,"F":15.5883,"P.Value":7.6209e-06,"adj.P.Val":0.0002},{"tissuedysplasia":0.0965,"tissuetumour":-0.1551,"AveExpr":2.0373,"F":0.8672,"P.Value":0.4272,"adj.P.Val":0.5649},{"tissuedysplasia":1.9453,"tissuetumour":3.6256,"AveExpr":-3.2791,"F":8.27,"P.Value":0.0009,"adj.P.Val":0.0071},{"tissuedysplasia":0.4996,"tissuetumour":0.1695,"AveExpr":5.1502,"F":6.5033,"P.Value":0.0034,"adj.P.Val":0.0188},{"tissuedysplasia":-0.0438,"tissuetumour":-0.0483,"AveExpr":5.8439,"F":0.211,"P.Value":0.8106,"adj.P.Val":0.867},{"tissuedysplasia":0.3106,"tissuetumour":-0.1343,"AveExpr":2.1863,"F":3.669,"P.Value":0.0336,"adj.P.Val":0.0986},{"tissuedysplasia":0.0628,"tissuetumour":0.4365,"AveExpr":3.2725,"F":4.0163,"P.Value":0.025,"adj.P.Val":0.0802},{"tissuedysplasia":0.0351,"tissuetumour":-0.014,"AveExpr":4.4496,"F":0.1005,"P.Value":0.9045,"adj.P.Val":0.9335},{"tissuedysplasia":0.419,"tissuetumour":0.3337,"AveExpr":4.6395,"F":5.6598,"P.Value":0.0065,"adj.P.Val":0.0304},{"tissuedysplasia":0.6181,"tissuetumour":1.5502,"AveExpr":0.1118,"F":11.1286,"P.Value":0.0001,"adj.P.Val":0.0016},{"tissuedysplasia":0.2548,"tissuetumour":0.5779,"AveExpr":5.3178,"F":11.0599,"P.Value":0.0001,"adj.P.Val":0.0017},{"tissuedysplasia":-0.1087,"tissuetumour":-0.3083,"AveExpr":6.6009,"F":3.4776,"P.Value":0.0396,"adj.P.Val":0.1108},{"tissuedysplasia":0.361,"tissuetumour":0.4922,"AveExpr":-3.1595,"F":0.1807,"P.Value":0.8353,"adj.P.Val":0.8843},{"tissuedysplasia":-0.2957,"tissuetumour":-0.4888,"AveExpr":6.3092,"F":17.4618,"P.Value":2.6136e-06,"adj.P.Val":0.0001},{"tissuedysplasia":0.2892,"tissuetumour":0.7768,"AveExpr":3.4817,"F":9.5486,"P.Value":0.0004,"adj.P.Val":0.0036},{"tissuedysplasia":0.2656,"tissuetumour":0.5884,"AveExpr":4.9478,"F":8.5893,"P.Value":0.0007,"adj.P.Val":0.006},{"tissuedysplasia":0.2364,"tissuetumour":0.2726,"AveExpr":2.2515,"F":1.6085,"P.Value":0.2117,"adj.P.Val":0.3507},{"tissuedysplasia":0.2254,"tissuetumour":0.3299,"AveExpr":4.8811,"F":3.0558,"P.Value":0.0572,"adj.P.Val":0.1434},{"tissuedysplasia":0.3209,"tissuetumour":1.4543,"AveExpr":-1.6764,"F":3.1045,"P.Value":0.0548,"adj.P.Val":0.1393},{"tissuedysplasia":0.5319,"tissuetumour":1.1535,"AveExpr":3.9646,"F":19.9946,"P.Value":6.6504e-07,"adj.P.Val":0},{"tissuedysplasia":0.4863,"tissuetumour":0.8639,"AveExpr":4.4007,"F":13.1101,"P.Value":0,"adj.P.Val":0.0006},{"tissuedysplasia":0.0751,"tissuetumour":-0.1677,"AveExpr":5.6327,"F":3.2055,"P.Value":0.0502,"adj.P.Val":0.131},{"tissuedysplasia":1.3434,"tissuetumour":2.4717,"AveExpr":1.1374,"F":20.475,"P.Value":5.178e-07,"adj.P.Val":0},{"tissuedysplasia":-0.0905,"tissuetumour":0.1457,"AveExpr":2.0183,"F":0.2391,"P.Value":0.7883,"adj.P.Val":0.8506},{"tissuedysplasia":1.271,"tissuetumour":2.0984,"AveExpr":-0.1397,"F":7.9404,"P.Value":0.0011,"adj.P.Val":0.0085},{"tissuedysplasia":0.3068,"tissuetumour":0.3228,"AveExpr":4.5325,"F":2.301,"P.Value":0.1121,"adj.P.Val":0.228},{"tissuedysplasia":0.3826,"tissuetumour":0.2427,"AveExpr":4.6044,"F":3.6576,"P.Value":0.0339,"adj.P.Val":0.0993},{"tissuedysplasia":0.3007,"tissuetumour":-0.0209,"AveExpr":7.5192,"F":3.708,"P.Value":0.0329,"adj.P.Val":0.0972},{"tissuedysplasia":0.2231,"tissuetumour":0.4622,"AveExpr":6.0947,"F":8.1668,"P.Value":0.001,"adj.P.Val":0.0075},{"tissuedysplasia":-0.329,"tissuetumour":-0.3973,"AveExpr":7.6096,"F":17.0973,"P.Value":3.2056e-06,"adj.P.Val":0.0001},{"tissuedysplasia":-0.6892,"tissuetumour":-0.9337,"AveExpr":1.5965,"F":4.9485,"P.Value":0.0115,"adj.P.Val":0.0458},{"tissuedysplasia":0.1059,"tissuetumour":-0.2833,"AveExpr":6.797,"F":5.0852,"P.Value":0.0103,"adj.P.Val":0.0424},{"tissuedysplasia":0.3039,"tissuetumour":2.9279,"AveExpr":-5.8046,"F":8.043,"P.Value":0.0011,"adj.P.Val":0.008},{"tissuedysplasia":0.1424,"tissuetumour":1.2684,"AveExpr":0.8238,"F":5.1244,"P.Value":0.01,"adj.P.Val":0.0414},{"tissuedysplasia":0.0392,"tissuetumour":-0.32,"AveExpr":7.4036,"F":9.9718,"P.Value":0.0003,"adj.P.Val":0.0029},{"tissuedysplasia":-0.2553,"tissuetumour":-0.227,"AveExpr":3.9468,"F":2.5041,"P.Value":0.0933,"adj.P.Val":0.2012},{"tissuedysplasia":0.2432,"tissuetumour":0.4388,"AveExpr":5.395,"F":6.1761,"P.Value":0.0043,"adj.P.Val":0.0226},{"tissuedysplasia":1.1492,"tissuetumour":1.8257,"AveExpr":-3.6873,"F":1.8699,"P.Value":0.1662,"adj.P.Val":0.2976},{"tissuedysplasia":0.0991,"tissuetumour":0.082,"AveExpr":5.6401,"F":0.4795,"P.Value":0.6223,"adj.P.Val":0.7265},{"tissuedysplasia":0.3152,"tissuetumour":1.3039,"AveExpr":2.6518,"F":16.5325,"P.Value":4.415e-06,"adj.P.Val":0.0001},{"tissuedysplasia":0.4861,"tissuetumour":0.2457,"AveExpr":6.6688,"F":7.9629,"P.Value":0.0011,"adj.P.Val":0.0084},{"tissuedysplasia":0.4278,"tissuetumour":0.1012,"AveExpr":1.0412,"F":1.4953,"P.Value":0.2354,"adj.P.Val":0.377},{"tissuedysplasia":-0.2104,"tissuetumour":-0.643,"AveExpr":3.192,"F":7.7405,"P.Value":0.0013,"adj.P.Val":0.0095},{"tissuedysplasia":0.288,"tissuetumour":0.2388,"AveExpr":3.3578,"F":1.5932,"P.Value":0.2148,"adj.P.Val":0.3541},{"tissuedysplasia":0.1834,"tissuetumour":0.2643,"AveExpr":4.3073,"F":3.5636,"P.Value":0.0368,"adj.P.Val":0.1052},{"tissuedysplasia":0.5945,"tissuetumour":1.0032,"AveExpr":4.5662,"F":16.5452,"P.Value":4.3833e-06,"adj.P.Val":0.0001},{"tissuedysplasia":0.347,"tissuetumour":0.0072,"AveExpr":5.3105,"F":3.2313,"P.Value":0.049,"adj.P.Val":0.1289},{"tissuedysplasia":0.0352,"tissuetumour":-0.1442,"AveExpr":6.4557,"F":1.8334,"P.Value":0.1719,"adj.P.Val":0.3046},{"tissuedysplasia":0.3556,"tissuetumour":0.3939,"AveExpr":2.0862,"F":1.8514,"P.Value":0.169,"adj.P.Val":0.3012},{"tissuedysplasia":0.395,"tissuetumour":0.5458,"AveExpr":3.9759,"F":8.1548,"P.Value":0.001,"adj.P.Val":0.0076},{"tissuedysplasia":0.1749,"tissuetumour":0.1519,"AveExpr":5.9155,"F":2.3214,"P.Value":0.11,"adj.P.Val":0.225},{"tissuedysplasia":0.0583,"tissuetumour":0.515,"AveExpr":7.3341,"F":15.8545,"P.Value":6.5248e-06,"adj.P.Val":0.0002},{"tissuedysplasia":0.174,"tissuetumour":0.009,"AveExpr":3.3105,"F":0.9203,"P.Value":0.4059,"adj.P.Val":0.5455},{"tissuedysplasia":-0.0359,"tissuetumour":-0.4159,"AveExpr":6.0256,"F":4.816,"P.Value":0.0128,"adj.P.Val":0.0496},{"tissuedysplasia":0.1263,"tissuetumour":0.344,"AveExpr":4.9757,"F":4.788,"P.Value":0.0131,"adj.P.Val":0.0505},{"tissuedysplasia":0.0169,"tissuetumour":0.5453,"AveExpr":6.6127,"F":17.3756,"P.Value":2.7424e-06,"adj.P.Val":0.0001},{"tissuedysplasia":-0.1984,"tissuetumour":-0.3354,"AveExpr":4.1343,"F":2.6976,"P.Value":0.0785,"adj.P.Val":0.1783},{"tissuedysplasia":-0.2763,"tissuetumour":-0.9532,"AveExpr":7.297,"F":19.3067,"P.Value":1.0424e-06,"adj.P.Val":0},{"tissuedysplasia":0.0747,"tissuetumour":0.3645,"AveExpr":3.2136,"F":4.2636,"P.Value":0.0203,"adj.P.Val":0.0693},{"tissuedysplasia":1.2248,"tissuetumour":2.8939,"AveExpr":-0.527,"F":24.2689,"P.Value":7.8829e-08,"adj.P.Val":5.996e-06},{"tissuedysplasia":0.2696,"tissuetumour":0.1162,"AveExpr":4.7494,"F":1.3146,"P.Value":0.2789,"adj.P.Val":0.4235},{"tissuedysplasia":0.7753,"tissuetumour":1.2918,"AveExpr":-0.1199,"F":4.0785,"P.Value":0.0237,"adj.P.Val":0.0772},{"tissuedysplasia":0.0916,"tissuetumour":-0.0486,"AveExpr":6.5827,"F":1.3432,"P.Value":0.2715,"adj.P.Val":0.4156},{"tissuedysplasia":0.5328,"tissuetumour":1.2844,"AveExpr":2.8898,"F":13.9938,"P.Value":0,"adj.P.Val":0.0004},{"tissuedysplasia":0.1861,"tissuetumour":0.2469,"AveExpr":5.1774,"F":3.4565,"P.Value":0.0403,"adj.P.Val":0.1123},{"tissuedysplasia":-0.0138,"tissuetumour":0.2204,"AveExpr":4.969,"F":2.9263,"P.Value":0.0641,"adj.P.Val":0.1552},{"tissuedysplasia":0.4119,"tissuetumour":0.607,"AveExpr":5.3329,"F":7.1504,"P.Value":0.002,"adj.P.Val":0.0131},{"tissuedysplasia":0.2278,"tissuetumour":0.1519,"AveExpr":6.4078,"F":1.8806,"P.Value":0.1646,"adj.P.Val":0.2956},{"tissuedysplasia":0.0329,"tissuetumour":0.6738,"AveExpr":1.3946,"F":4.9074,"P.Value":0.0119,"adj.P.Val":0.0469},{"tissuedysplasia":0.4964,"tissuetumour":0.8048,"AveExpr":4.7915,"F":18.4498,"P.Value":1.5169e-06,"adj.P.Val":0.0001},{"tissuedysplasia":0.1642,"tissuetumour":0.2178,"AveExpr":2.946,"F":0.8711,"P.Value":0.4256,"adj.P.Val":0.5633},{"tissuedysplasia":0.4507,"tissuetumour":1.0852,"AveExpr":3.1897,"F":16.0864,"P.Value":5.7043e-06,"adj.P.Val":0.0002},{"tissuedysplasia":0.1511,"tissuetumour":0.5466,"AveExpr":4.0886,"F":11.85,"P.Value":0.0001,"adj.P.Val":0.0011},{"tissuedysplasia":0.1949,"tissuetumour":0.3815,"AveExpr":5.1955,"F":2.3122,"P.Value":0.111,"adj.P.Val":0.2264},{"tissuedysplasia":0.0294,"tissuetumour":-0.4979,"AveExpr":5.0754,"F":14.0265,"P.Value":0,"adj.P.Val":0.0004},{"tissuedysplasia":0.074,"tissuetumour":-0.0886,"AveExpr":6.2918,"F":1.2322,"P.Value":0.3015,"adj.P.Val":0.446},{"tissuedysplasia":-0.1363,"tissuetumour":0.0926,"AveExpr":3.9175,"F":0.9226,"P.Value":0.405,"adj.P.Val":0.5448},{"tissuedysplasia":-0.1265,"tissuetumour":0.9922,"AveExpr":4.017,"F":11.9062,"P.Value":0.0001,"adj.P.Val":0.0011},{"tissuedysplasia":0.4511,"tissuetumour":1.0433,"AveExpr":4.7266,"F":19.936,"P.Value":6.8577e-07,"adj.P.Val":0},{"tissuedysplasia":0.2463,"tissuetumour":0.5149,"AveExpr":3.2881,"F":4.8763,"P.Value":0.0122,"adj.P.Val":0.0478},{"tissuedysplasia":0.2493,"tissuetumour":0.6282,"AveExpr":3.9469,"F":10.7467,"P.Value":0.0002,"adj.P.Val":0.002},{"tissuedysplasia":2.2042,"tissuetumour":4.331,"AveExpr":1.9719,"F":27.2217,"P.Value":2.0209e-08,"adj.P.Val":2.2887e-06},{"tissuedysplasia":0.5966,"tissuetumour":1.2339,"AveExpr":-3.0639,"F":0.7691,"P.Value":0.4696,"adj.P.Val":0.6003},{"tissuedysplasia":0.2911,"tissuetumour":0.4164,"AveExpr":5.1616,"F":8.797,"P.Value":0.0006,"adj.P.Val":0.0054},{"tissuedysplasia":0.3476,"tissuetumour":1.0568,"AveExpr":5.5379,"F":28.9204,"P.Value":9.5799e-09,"adj.P.Val":1.3569e-06},{"tissuedysplasia":-0.1163,"tissuetumour":0.1028,"AveExpr":4.5765,"F":2.0994,"P.Value":0.1346,"adj.P.Val":0.2583},{"tissuedysplasia":0.8814,"tissuetumour":1.7058,"AveExpr":2.1435,"F":19.7596,"P.Value":7.5241e-07,"adj.P.Val":0},{"tissuedysplasia":0.693,"tissuetumour":1.0165,"AveExpr":3.4519,"F":17.3822,"P.Value":2.7323e-06,"adj.P.Val":0.0001},{"tissuedysplasia":0.4889,"tissuetumour":0.9161,"AveExpr":3.5962,"F":12.3031,"P.Value":0.0001,"adj.P.Val":0.0009},{"tissuedysplasia":0.5531,"tissuetumour":0.8407,"AveExpr":2.8157,"F":10.9165,"P.Value":0.0001,"adj.P.Val":0.0018},{"tissuedysplasia":0.2869,"tissuetumour":0.449,"AveExpr":4.4339,"F":7.598,"P.Value":0.0015,"adj.P.Val":0.0103},{"tissuedysplasia":0.1074,"tissuetumour":0.1846,"AveExpr":5.8973,"F":1.0895,"P.Value":0.3453,"adj.P.Val":0.4893},{"tissuedysplasia":-0.3921,"tissuetumour":-0.056,"AveExpr":0.1388,"F":0.4714,"P.Value":0.6273,"adj.P.Val":0.7304},{"tissuedysplasia":0.1176,"tissuetumour":0.3134,"AveExpr":2.269,"F":1.7699,"P.Value":0.1823,"adj.P.Val":0.3172},{"tissuedysplasia":1.0625,"tissuetumour":2.0635,"AveExpr":3.1968,"F":51.0933,"P.Value":3.3682e-12,"adj.P.Val":5.7494e-09},{"tissuedysplasia":0.3794,"tissuetumour":2.1297,"AveExpr":-7.8369,"F":1.7956,"P.Value":0.178,"adj.P.Val":0.3121},{"tissuedysplasia":0.555,"tissuetumour":2.6508,"AveExpr":-1.3653,"F":10.905,"P.Value":0.0001,"adj.P.Val":0.0018},{"tissuedysplasia":0.1461,"tissuetumour":0.0546,"AveExpr":4.6084,"F":0.8215,"P.Value":0.4464,"adj.P.Val":0.5819},{"tissuedysplasia":0.3666,"tissuetumour":0.2196,"AveExpr":2.1976,"F":1.2544,"P.Value":0.2952,"adj.P.Val":0.4397},{"tissuedysplasia":0.0144,"tissuetumour":-0.2142,"AveExpr":4.6524,"F":2.3974,"P.Value":0.1027,"adj.P.Val":0.2149},{"tissuedysplasia":0.2286,"tissuetumour":0.0279,"AveExpr":4.7493,"F":0.6756,"P.Value":0.514,"adj.P.Val":0.6386},{"tissuedysplasia":-0.0409,"tissuetumour":1.4336,"AveExpr":-2.0853,"F":2.3615,"P.Value":0.1061,"adj.P.Val":0.2195},{"tissuedysplasia":0.3112,"tissuetumour":0.8598,"AveExpr":4.0462,"F":17.5047,"P.Value":2.5519e-06,"adj.P.Val":0.0001},{"tissuedysplasia":0.5367,"tissuetumour":0.3399,"AveExpr":6.5231,"F":11.4216,"P.Value":0.0001,"adj.P.Val":0.0014},{"tissuedysplasia":0.7794,"tissuetumour":2.2145,"AveExpr":0.1989,"F":18.1675,"P.Value":1.7696e-06,"adj.P.Val":0.0001},{"tissuedysplasia":0.2171,"tissuetumour":-0.0941,"AveExpr":6.4878,"F":5.0497,"P.Value":0.0106,"adj.P.Val":0.0433},{"tissuedysplasia":0.241,"tissuetumour":0.3617,"AveExpr":1.393,"F":0.4736,"P.Value":0.6259,"adj.P.Val":0.7294},{"tissuedysplasia":-0.1458,"tissuetumour":1.7825,"AveExpr":-1.6801,"F":6.1514,"P.Value":0.0044,"adj.P.Val":0.0229},{"tissuedysplasia":0.2539,"tissuetumour":0.1674,"AveExpr":3.7005,"F":2.7121,"P.Value":0.0775,"adj.P.Val":0.1768},{"tissuedysplasia":-0.0416,"tissuetumour":-0.9043,"AveExpr":3.8126,"F":7.7355,"P.Value":0.0013,"adj.P.Val":0.0095},{"tissuedysplasia":0.2223,"tissuetumour":0.2943,"AveExpr":3.4274,"F":1.8892,"P.Value":0.1632,"adj.P.Val":0.2939},{"tissuedysplasia":0.1948,"tissuetumour":0.8633,"AveExpr":5.7227,"F":20.8732,"P.Value":4.2169e-07,"adj.P.Val":0},{"tissuedysplasia":0.0275,"tissuetumour":0.428,"AveExpr":6.3909,"F":7.7709,"P.Value":0.0013,"adj.P.Val":0.0094},{"tissuedysplasia":0.1861,"tissuetumour":0.3978,"AveExpr":5.1482,"F":7.5629,"P.Value":0.0015,"adj.P.Val":0.0105},{"tissuedysplasia":0.8524,"tissuetumour":1.377,"AveExpr":4.7732,"F":24.4222,"P.Value":7.3296e-08,"adj.P.Val":5.6794e-06},{"tissuedysplasia":0.0675,"tissuetumour":0.6719,"AveExpr":5.5569,"F":17.8032,"P.Value":2.1624e-06,"adj.P.Val":0.0001},{"tissuedysplasia":0.5012,"tissuetumour":1.0067,"AveExpr":0.9026,"F":4.5653,"P.Value":0.0158,"adj.P.Val":0.0577},{"tissuedysplasia":0.0239,"tissuetumour":-0.4044,"AveExpr":4.9902,"F":3.7963,"P.Value":0.0301,"adj.P.Val":0.0916},{"tissuedysplasia":-0.4318,"tissuetumour":-0.4461,"AveExpr":-6.6724,"F":0.0908,"P.Value":0.9134,"adj.P.Val":0.9394},{"tissuedysplasia":0.4685,"tissuetumour":1.0918,"AveExpr":3.3188,"F":25.6617,"P.Value":4.1047e-08,"adj.P.Val":3.7631e-06},{"tissuedysplasia":0.3648,"tissuetumour":0.7583,"AveExpr":5.1479,"F":12.9441,"P.Value":0,"adj.P.Val":0.0007},{"tissuedysplasia":-0.3088,"tissuetumour":-0.2175,"AveExpr":2.9754,"F":1.4262,"P.Value":0.2511,"adj.P.Val":0.394},{"tissuedysplasia":1.6574,"tissuetumour":1.9408,"AveExpr":-2.9333,"F":3.3024,"P.Value":0.0461,"adj.P.Val":0.1233},{"tissuedysplasia":0.6593,"tissuetumour":0.9962,"AveExpr":2.2773,"F":8.3427,"P.Value":0.0008,"adj.P.Val":0.0069},{"tissuedysplasia":0.785,"tissuetumour":0.9537,"AveExpr":2.3489,"F":8.9554,"P.Value":0.0005,"adj.P.Val":0.005},{"tissuedysplasia":0.1289,"tissuetumour":0.2843,"AveExpr":6.2335,"F":6.1098,"P.Value":0.0046,"adj.P.Val":0.0235},{"tissuedysplasia":-0.1176,"tissuetumour":-0.1045,"AveExpr":5.5643,"F":0.8064,"P.Value":0.4529,"adj.P.Val":0.5873},{"tissuedysplasia":0.9472,"tissuetumour":1.3934,"AveExpr":1.888,"F":7.7193,"P.Value":0.0013,"adj.P.Val":0.0096},{"tissuedysplasia":-0.3386,"tissuetumour":0.6199,"AveExpr":-7.2287,"F":0.3517,"P.Value":0.7055,"adj.P.Val":0.789},{"tissuedysplasia":-0.2823,"tissuetumour":2.0647,"AveExpr":-4.8119,"F":4.4297,"P.Value":0.0177,"adj.P.Val":0.0627},{"tissuedysplasia":0.3124,"tissuetumour":1.2459,"AveExpr":4.4609,"F":38.3774,"P.Value":2.2575e-10,"adj.P.Val":8.9127e-08},{"tissuedysplasia":0.3116,"tissuetumour":1.6314,"AveExpr":-10.5984,"F":2.2027,"P.Value":0.1225,"adj.P.Val":0.2418},{"tissuedysplasia":0.7,"tissuetumour":2.3824,"AveExpr":-2.8442,"F":3.9386,"P.Value":0.0267,"adj.P.Val":0.084},{"tissuedysplasia":1.3345,"tissuetumour":2.6121,"AveExpr":-2.5064,"F":5.2352,"P.Value":0.0091,"adj.P.Val":0.0388},{"tissuedysplasia":-1.6139,"tissuetumour":0.69,"AveExpr":0.2564,"F":7.668,"P.Value":0.0014,"adj.P.Val":0.0099},{"tissuedysplasia":-0.821,"tissuetumour":-0.7488,"AveExpr":4.3861,"F":10.6484,"P.Value":0.0002,"adj.P.Val":0.0021},{"tissuedysplasia":0.4162,"tissuetumour":0.8367,"AveExpr":3.1608,"F":13.4902,"P.Value":0,"adj.P.Val":0.0005},{"tissuedysplasia":0.2076,"tissuetumour":1.1437,"AveExpr":2.0893,"F":14.9891,"P.Value":0,"adj.P.Val":0.0003},{"tissuedysplasia":0.7136,"tissuetumour":1.3026,"AveExpr":-5.6434,"F":0.6682,"P.Value":0.5178,"adj.P.Val":0.6418},{"tissuedysplasia":0.6508,"tissuetumour":1.3891,"AveExpr":5.6304,"F":27.9791,"P.Value":1.4442e-08,"adj.P.Val":1.8324e-06},{"tissuedysplasia":-0.5056,"tissuetumour":-0.7868,"AveExpr":3.0656,"F":5.5979,"P.Value":0.0068,"adj.P.Val":0.0316},{"tissuedysplasia":-0.1147,"tissuetumour":-0.0099,"AveExpr":4.6696,"F":0.5338,"P.Value":0.5901,"adj.P.Val":0.7007},{"tissuedysplasia":0.2006,"tissuetumour":0.863,"AveExpr":4.5554,"F":15.3582,"P.Value":8.7229e-06,"adj.P.Val":0.0002},{"tissuedysplasia":0.1845,"tissuetumour":0.3376,"AveExpr":3.4139,"F":2.303,"P.Value":0.1119,"adj.P.Val":0.2277},{"tissuedysplasia":0.8214,"tissuetumour":2.3945,"AveExpr":-4.1083,"F":4.3862,"P.Value":0.0183,"adj.P.Val":0.0644},{"tissuedysplasia":0.212,"tissuetumour":1.5846,"AveExpr":0.8474,"F":14.1687,"P.Value":0,"adj.P.Val":0.0004},{"tissuedysplasia":0.7988,"tissuetumour":0.53,"AveExpr":4.5784,"F":6.7702,"P.Value":0.0027,"adj.P.Val":0.0163},{"tissuedysplasia":0.1074,"tissuetumour":-0.1177,"AveExpr":5.7138,"F":2.675,"P.Value":0.0801,"adj.P.Val":0.1808},{"tissuedysplasia":0.4769,"tissuetumour":0.8926,"AveExpr":1.3837,"F":4.5964,"P.Value":0.0154,"adj.P.Val":0.0566},{"tissuedysplasia":0.2073,"tissuetumour":0.6688,"AveExpr":-1.9848,"F":1.0339,"P.Value":0.3641,"adj.P.Val":0.5071},{"tissuedysplasia":0.9893,"tissuetumour":1.0033,"AveExpr":-1.0561,"F":2.2432,"P.Value":0.1181,"adj.P.Val":0.2361},{"tissuedysplasia":0.0433,"tissuetumour":0.1879,"AveExpr":-2.2731,"F":0.0326,"P.Value":0.9679,"adj.P.Val":0.9782},{"tissuedysplasia":0.312,"tissuetumour":0.2896,"AveExpr":5.5047,"F":5.5437,"P.Value":0.0071,"adj.P.Val":0.0326},{"tissuedysplasia":1.6466,"tissuetumour":2.9128,"AveExpr":-6.6657,"F":3.5258,"P.Value":0.038,"adj.P.Val":0.1076},{"tissuedysplasia":0.0592,"tissuetumour":0.1973,"AveExpr":6.0021,"F":1.9346,"P.Value":0.1566,"adj.P.Val":0.2857},{"tissuedysplasia":-0.0633,"tissuetumour":-0.6471,"AveExpr":2.3821,"F":2.3326,"P.Value":0.1089,"adj.P.Val":0.2235},{"tissuedysplasia":0.4273,"tissuetumour":2.7784,"AveExpr":-2.8154,"F":8.0962,"P.Value":0.001,"adj.P.Val":0.0078},{"tissuedysplasia":0.9935,"tissuetumour":1.7169,"AveExpr":1.3683,"F":13.5304,"P.Value":0,"adj.P.Val":0.0005},{"tissuedysplasia":0.1986,"tissuetumour":-0.0242,"AveExpr":4.9019,"F":2.4448,"P.Value":0.0984,"adj.P.Val":0.2086},{"tissuedysplasia":0.543,"tissuetumour":0.6645,"AveExpr":5.9689,"F":5.3782,"P.Value":0.0082,"adj.P.Val":0.036},{"tissuedysplasia":-0.1884,"tissuetumour":0.0083,"AveExpr":-0.2861,"F":0.1212,"P.Value":0.8861,"adj.P.Val":0.9201},{"tissuedysplasia":1.1772,"tissuetumour":1.2374,"AveExpr":-5.6785,"F":1.0303,"P.Value":0.3653,"adj.P.Val":0.5083},{"tissuedysplasia":0.228,"tissuetumour":1.1365,"AveExpr":-1.0849,"F":2.445,"P.Value":0.0984,"adj.P.Val":0.2086},{"tissuedysplasia":-0.2537,"tissuetumour":-0.0389,"AveExpr":2.5536,"F":0.4866,"P.Value":0.618,"adj.P.Val":0.7229},{"tissuedysplasia":0.1638,"tissuetumour":1.1503,"AveExpr":3.84,"F":21.1108,"P.Value":3.7341e-07,"adj.P.Val":0},{"tissuedysplasia":1.1055,"tissuetumour":1.5578,"AveExpr":2.5897,"F":19.8641,"P.Value":7.1218e-07,"adj.P.Val":0},{"tissuedysplasia":0.1979,"tissuetumour":0.2566,"AveExpr":5.5373,"F":3.782,"P.Value":0.0305,"adj.P.Val":0.0923},{"tissuedysplasia":0.3144,"tissuetumour":0.247,"AveExpr":3.2825,"F":1.0109,"P.Value":0.3722,"adj.P.Val":0.5148},{"tissuedysplasia":0.1446,"tissuetumour":0.0141,"AveExpr":4.4764,"F":0.9945,"P.Value":0.3781,"adj.P.Val":0.5202},{"tissuedysplasia":0.3547,"tissuetumour":0.8756,"AveExpr":4.7754,"F":28.8519,"P.Value":9.8679e-09,"adj.P.Val":1.3832e-06},{"tissuedysplasia":0.3937,"tissuetumour":0.6191,"AveExpr":6.7728,"F":3.9014,"P.Value":0.0286,"adj.P.Val":0.0882},{"tissuedysplasia":0.1934,"tissuetumour":1.7397,"AveExpr":-1.2574,"F":5.1277,"P.Value":0.01,"adj.P.Val":0.0413},{"tissuedysplasia":0.5476,"tissuetumour":0.9473,"AveExpr":5.182,"F":38.665,"P.Value":2.0333e-10,"adj.P.Val":8.4898e-08},{"tissuedysplasia":-0.0386,"tissuetumour":0.7862,"AveExpr":0.804,"F":5.6761,"P.Value":0.0064,"adj.P.Val":0.0302},{"tissuedysplasia":0.1722,"tissuetumour":0.5308,"AveExpr":6.524,"F":9.063,"P.Value":0.0005,"adj.P.Val":0.0047},{"tissuedysplasia":0.0354,"tissuetumour":0.4144,"AveExpr":3.3613,"F":4.5534,"P.Value":0.0159,"adj.P.Val":0.0581},{"tissuedysplasia":0.3215,"tissuetumour":0.4018,"AveExpr":3.658,"F":3.9449,"P.Value":0.0266,"adj.P.Val":0.0837},{"tissuedysplasia":0.2855,"tissuetumour":0.1758,"AveExpr":5.9599,"F":3.915,"P.Value":0.0272,"adj.P.Val":0.0852},{"tissuedysplasia":0.0986,"tissuetumour":-0.0009,"AveExpr":7.6249,"F":0.44,"P.Value":0.6469,"adj.P.Val":0.7451},{"tissuedysplasia":0.1144,"tissuetumour":0.4194,"AveExpr":4.7256,"F":3.9443,"P.Value":0.0266,"adj.P.Val":0.0837},{"tissuedysplasia":-0.2995,"tissuetumour":1.0244,"AveExpr":1.7074,"F":16.0533,"P.Value":5.8146e-06,"adj.P.Val":0.0002},{"tissuedysplasia":1.0526,"tissuetumour":2.2908,"AveExpr":-7.9832,"F":1.3557,"P.Value":0.2683,"adj.P.Val":0.4119},{"tissuedysplasia":0.447,"tissuetumour":0.447,"AveExpr":4.2219,"F":9.2504,"P.Value":0.0004,"adj.P.Val":0.0042},{"tissuedysplasia":0.7507,"tissuetumour":1.6745,"AveExpr":1.1633,"F":16.9171,"P.Value":3.5486e-06,"adj.P.Val":0.0001},{"tissuedysplasia":0.3514,"tissuetumour":0.4568,"AveExpr":4.8618,"F":9.0895,"P.Value":0.0005,"adj.P.Val":0.0046},{"tissuedysplasia":0.1101,"tissuetumour":0.5957,"AveExpr":3.1214,"F":3.8182,"P.Value":0.0296,"adj.P.Val":0.0904},{"tissuedysplasia":0.0238,"tissuetumour":0.0618,"AveExpr":5.0495,"F":0.1836,"P.Value":0.8329,"adj.P.Val":0.8827},{"tissuedysplasia":1.4579,"tissuetumour":3.2833,"AveExpr":-1.8887,"F":9.2161,"P.Value":0.0005,"adj.P.Val":0.0043},{"tissuedysplasia":0.8277,"tissuetumour":2.2297,"AveExpr":-7.5987,"F":1.659,"P.Value":0.202,"adj.P.Val":0.3393},{"tissuedysplasia":0.5725,"tissuetumour":3.2156,"AveExpr":-2.5784,"F":13.2741,"P.Value":0,"adj.P.Val":0.0006},{"tissuedysplasia":0.1863,"tissuetumour":0.3889,"AveExpr":5.5398,"F":4.0884,"P.Value":0.0235,"adj.P.Val":0.0768},{"tissuedysplasia":-0.0347,"tissuetumour":-0.465,"AveExpr":-1.995,"F":0.2648,"P.Value":0.7686,"adj.P.Val":0.836},{"tissuedysplasia":0.8835,"tissuetumour":3.6679,"AveExpr":-4.9862,"F":8.6478,"P.Value":0.0007,"adj.P.Val":0.0059},{"tissuedysplasia":0.2311,"tissuetumour":0.4379,"AveExpr":2.3959,"F":1.7805,"P.Value":0.1805,"adj.P.Val":0.3151},{"tissuedysplasia":1.5946,"tissuetumour":1.4155,"AveExpr":-2.7392,"F":2.1861,"P.Value":0.1244,"adj.P.Val":0.2443},{"tissuedysplasia":-0.0102,"tissuetumour":-0.0806,"AveExpr":4.944,"F":0.2015,"P.Value":0.8182,"adj.P.Val":0.8727},{"tissuedysplasia":1.0231,"tissuetumour":0.7285,"AveExpr":3.7714,"F":8.3296,"P.Value":0.0009,"adj.P.Val":0.0069},{"tissuedysplasia":1.7493,"tissuetumour":5.1399,"AveExpr":-7.0298,"F":8.7983,"P.Value":0.0006,"adj.P.Val":0.0054},{"tissuedysplasia":-0.4189,"tissuetumour":-0.918,"AveExpr":4.0649,"F":9.3668,"P.Value":0.0004,"adj.P.Val":0.004},{"tissuedysplasia":1.1205,"tissuetumour":2.8231,"AveExpr":-1.3042,"F":10.8769,"P.Value":0.0001,"adj.P.Val":0.0018},{"tissuedysplasia":0.1886,"tissuetumour":-0.0155,"AveExpr":3.2815,"F":0.7703,"P.Value":0.469,"adj.P.Val":0.5998},{"tissuedysplasia":0.5233,"tissuetumour":0.9647,"AveExpr":2.9895,"F":13.6954,"P.Value":0,"adj.P.Val":0.0005},{"tissuedysplasia":-0.3054,"tissuetumour":-0.2015,"AveExpr":1.0049,"F":0.4347,"P.Value":0.6502,"adj.P.Val":0.7477},{"tissuedysplasia":0.8733,"tissuetumour":2.1864,"AveExpr":-1.8433,"F":5.0052,"P.Value":0.011,"adj.P.Val":0.0444},{"tissuedysplasia":0.3905,"tissuetumour":0.3276,"AveExpr":2.6922,"F":3.2466,"P.Value":0.0484,"adj.P.Val":0.1278},{"tissuedysplasia":-2.4743,"tissuetumour":1.5904,"AveExpr":-1.2001,"F":9.742,"P.Value":0.0003,"adj.P.Val":0.0033},{"tissuedysplasia":0.0288,"tissuetumour":0.3111,"AveExpr":3.318,"F":2.5687,"P.Value":0.0881,"adj.P.Val":0.1931},{"tissuedysplasia":0.5605,"tissuetumour":1.0284,"AveExpr":4.7827,"F":25.3334,"P.Value":4.7789e-08,"adj.P.Val":4.1645e-06},{"tissuedysplasia":-0.0143,"tissuetumour":0.3509,"AveExpr":0.7341,"F":0.8515,"P.Value":0.4337,"adj.P.Val":0.5707},{"tissuedysplasia":0.0313,"tissuetumour":0.0673,"AveExpr":4.7662,"F":0.1918,"P.Value":0.8262,"adj.P.Val":0.8781},{"tissuedysplasia":-0.1696,"tissuetumour":1.0898,"AveExpr":-1.2284,"F":2.6193,"P.Value":0.0842,"adj.P.Val":0.1871},{"tissuedysplasia":0.076,"tissuetumour":-0.1279,"AveExpr":4.6426,"F":1.2996,"P.Value":0.2829,"adj.P.Val":0.4274},{"tissuedysplasia":0.0624,"tissuetumour":0.2043,"AveExpr":3.4773,"F":0.8922,"P.Value":0.417,"adj.P.Val":0.5553},{"tissuedysplasia":0.2717,"tissuetumour":-0.039,"AveExpr":5.4568,"F":3.6195,"P.Value":0.0351,"adj.P.Val":0.1016},{"tissuedysplasia":0.0374,"tissuetumour":-0.3728,"AveExpr":6.2424,"F":7.1906,"P.Value":0.002,"adj.P.Val":0.0129},{"tissuedysplasia":0.5502,"tissuetumour":0.2603,"AveExpr":3.1668,"F":5.4312,"P.Value":0.0078,"adj.P.Val":0.0347},{"tissuedysplasia":0.2118,"tissuetumour":0.1087,"AveExpr":4.6758,"F":0.9477,"P.Value":0.3954,"adj.P.Val":0.536},{"tissuedysplasia":0.483,"tissuetumour":-0.0626,"AveExpr":6.9091,"F":6.3576,"P.Value":0.0039,"adj.P.Val":0.0211},{"tissuedysplasia":-0.9871,"tissuetumour":0.7632,"AveExpr":2.3853,"F":9.7065,"P.Value":0.0003,"adj.P.Val":0.0034},{"tissuedysplasia":0.0357,"tissuetumour":-0.1833,"AveExpr":6.0365,"F":1.4119,"P.Value":0.2545,"adj.P.Val":0.3974},{"tissuedysplasia":-0.1986,"tissuetumour":-0.4206,"AveExpr":2.8375,"F":1.5721,"P.Value":0.219,"adj.P.Val":0.359},{"tissuedysplasia":0.6282,"tissuetumour":2.3471,"AveExpr":0.1822,"F":15.2268,"P.Value":9.4259e-06,"adj.P.Val":0.0002},{"tissuedysplasia":0.4091,"tissuetumour":0.1602,"AveExpr":5.5724,"F":5.0249,"P.Value":0.0108,"adj.P.Val":0.0439},{"tissuedysplasia":0.2463,"tissuetumour":0.2544,"AveExpr":5.1139,"F":3.7181,"P.Value":0.0322,"adj.P.Val":0.0958},{"tissuedysplasia":0.0539,"tissuetumour":0.6796,"AveExpr":3.3453,"F":9.565,"P.Value":0.0004,"adj.P.Val":0.0036},{"tissuedysplasia":1.0656,"tissuetumour":1.0839,"AveExpr":3.5926,"F":6.0437,"P.Value":0.0048,"adj.P.Val":0.0244},{"tissuedysplasia":0.8384,"tissuetumour":2.6734,"AveExpr":2.3559,"F":22.4437,"P.Value":1.9109e-07,"adj.P.Val":0},{"tissuedysplasia":0.2866,"tissuetumour":-0.4692,"AveExpr":3.3256,"F":4.2271,"P.Value":0.0209,"adj.P.Val":0.0708},{"tissuedysplasia":-0.0356,"tissuetumour":-0.6,"AveExpr":6.8877,"F":9.533,"P.Value":0.0004,"adj.P.Val":0.0037},{"tissuedysplasia":-0.4944,"tissuetumour":1.8603,"AveExpr":-1.8856,"F":10.4314,"P.Value":0.0002,"adj.P.Val":0.0023},{"tissuedysplasia":0.0963,"tissuetumour":-0.1751,"AveExpr":4.0852,"F":1.5199,"P.Value":0.23,"adj.P.Val":0.3711},{"tissuedysplasia":0.2871,"tissuetumour":0.0617,"AveExpr":6.1648,"F":6.0247,"P.Value":0.0049,"adj.P.Val":0.0246},{"tissuedysplasia":0.5721,"tissuetumour":0.133,"AveExpr":5.0543,"F":4.1053,"P.Value":0.0232,"adj.P.Val":0.076},{"tissuedysplasia":-0.2673,"tissuetumour":-0.8655,"AveExpr":6.1205,"F":15.1732,"P.Value":9.7296e-06,"adj.P.Val":0.0002},{"tissuedysplasia":0.0828,"tissuetumour":0.115,"AveExpr":2.5916,"F":0.1472,"P.Value":0.8635,"adj.P.Val":0.9038},{"tissuedysplasia":-1.0485,"tissuetumour":1.5123,"AveExpr":-5.9419,"F":1.9593,"P.Value":0.1531,"adj.P.Val":0.2815},{"tissuedysplasia":0.1456,"tissuetumour":0.5827,"AveExpr":6.7281,"F":4.089,"P.Value":0.0244,"adj.P.Val":0.0787},{"tissuedysplasia":0.8818,"tissuetumour":1.4341,"AveExpr":0.8904,"F":7.5241,"P.Value":0.0015,"adj.P.Val":0.0107},{"tissuedysplasia":0.1456,"tissuetumour":0.201,"AveExpr":3.2052,"F":0.4946,"P.Value":0.6132,"adj.P.Val":0.7194},{"tissuedysplasia":0.1513,"tissuetumour":-0.0531,"AveExpr":2.5698,"F":0.6268,"P.Value":0.539,"adj.P.Val":0.6593},{"tissuedysplasia":0.537,"tissuetumour":0.776,"AveExpr":2.9526,"F":9.2329,"P.Value":0.0004,"adj.P.Val":0.0043},{"tissuedysplasia":0.2838,"tissuetumour":0.0021,"AveExpr":4.8589,"F":3.5153,"P.Value":0.0383,"adj.P.Val":0.1083},{"tissuedysplasia":0.514,"tissuetumour":0.0431,"AveExpr":-0.5097,"F":1.3197,"P.Value":0.2776,"adj.P.Val":0.4221},{"tissuedysplasia":0.1762,"tissuetumour":0.2497,"AveExpr":4.401,"F":1.2078,"P.Value":0.3086,"adj.P.Val":0.453},{"tissuedysplasia":0.1951,"tissuetumour":0.4144,"AveExpr":4.6785,"F":6.3267,"P.Value":0.0038,"adj.P.Val":0.0208},{"tissuedysplasia":0.1799,"tissuetumour":-0.1884,"AveExpr":4.6299,"F":2.6548,"P.Value":0.0816,"adj.P.Val":0.183},{"tissuedysplasia":0.2918,"tissuetumour":0.7219,"AveExpr":3.2756,"F":13.7534,"P.Value":0,"adj.P.Val":0.0004},{"tissuedysplasia":0.957,"tissuetumour":1.2758,"AveExpr":-5.6576,"F":0.7775,"P.Value":0.4658,"adj.P.Val":0.5975},{"tissuedysplasia":0.473,"tissuetumour":0.0664,"AveExpr":1.9961,"F":1.3261,"P.Value":0.2759,"adj.P.Val":0.4204},{"tissuedysplasia":-0.7342,"tissuetumour":2.2967,"AveExpr":-4.7347,"F":5.3857,"P.Value":0.0081,"adj.P.Val":0.0356},{"tissuedysplasia":-0.6285,"tissuetumour":2.4053,"AveExpr":-0.5502,"F":23.4118,"P.Value":1.1895e-07,"adj.P.Val":8.0177e-06},{"tissuedysplasia":0.4155,"tissuetumour":-0.2292,"AveExpr":0.4434,"F":1.3044,"P.Value":0.2816,"adj.P.Val":0.4263},{"tissuedysplasia":0.023,"tissuetumour":-0.1713,"AveExpr":5.1392,"F":1.2378,"P.Value":0.2999,"adj.P.Val":0.4443},{"tissuedysplasia":0.3245,"tissuetumour":-0.1917,"AveExpr":3.7438,"F":5.7959,"P.Value":0.0058,"adj.P.Val":0.0281},{"tissuedysplasia":0.8621,"tissuetumour":1.2863,"AveExpr":0.7912,"F":4.3731,"P.Value":0.0185,"adj.P.Val":0.0649},{"tissuedysplasia":-0.2028,"tissuetumour":-0.6209,"AveExpr":2.6909,"F":3.1587,"P.Value":0.0523,"adj.P.Val":0.1347},{"tissuedysplasia":0.1014,"tissuetumour":-0.2428,"AveExpr":5.1058,"F":2.492,"P.Value":0.0944,"adj.P.Val":0.2027},{"tissuedysplasia":0.0393,"tissuetumour":0.2935,"AveExpr":2.7006,"F":0.2797,"P.Value":0.7573,"adj.P.Val":0.8278},{"tissuedysplasia":0.0769,"tissuetumour":0.3242,"AveExpr":3.8318,"F":2.0575,"P.Value":0.1399,"adj.P.Val":0.2646},{"tissuedysplasia":0.5746,"tissuetumour":0.7422,"AveExpr":2.1202,"F":8.049,"P.Value":0.001,"adj.P.Val":0.008},{"tissuedysplasia":-0.2424,"tissuetumour":1.1563,"AveExpr":-7.0463,"F":0.9468,"P.Value":0.3958,"adj.P.Val":0.5364},{"tissuedysplasia":-0.6178,"tissuetumour":-1.0974,"AveExpr":1.9191,"F":3.7076,"P.Value":0.0325,"adj.P.Val":0.0964},{"tissuedysplasia":0.2134,"tissuetumour":1.0543,"AveExpr":-1.3402,"F":2.443,"P.Value":0.0986,"adj.P.Val":0.2089},{"tissuedysplasia":0.1332,"tissuetumour":0.0568,"AveExpr":6.213,"F":0.6767,"P.Value":0.5135,"adj.P.Val":0.6381},{"tissuedysplasia":0.2048,"tissuetumour":0.1126,"AveExpr":6.5185,"F":1.1746,"P.Value":0.3185,"adj.P.Val":0.4628},{"tissuedysplasia":0.344,"tissuetumour":0.3909,"AveExpr":4.5734,"F":3.4132,"P.Value":0.0419,"adj.P.Val":0.1153},{"tissuedysplasia":0.6464,"tissuetumour":1.4695,"AveExpr":-1.2583,"F":4.555,"P.Value":0.0159,"adj.P.Val":0.0581},{"tissuedysplasia":1.7826,"tissuetumour":2.3709,"AveExpr":-4.4987,"F":2.4364,"P.Value":0.0992,"adj.P.Val":0.2097},{"tissuedysplasia":0.6132,"tissuetumour":2.0233,"AveExpr":1.8134,"F":29.1744,"P.Value":8.5866e-09,"adj.P.Val":1.2533e-06},{"tissuedysplasia":0.2759,"tissuetumour":0.6398,"AveExpr":1.877,"F":2.9684,"P.Value":0.0618,"adj.P.Val":0.1512},{"tissuedysplasia":0.9975,"tissuetumour":1.5553,"AveExpr":-0.6618,"F":4.7435,"P.Value":0.0136,"adj.P.Val":0.0518},{"tissuedysplasia":0.1642,"tissuetumour":0.9126,"AveExpr":0.7818,"F":4.6942,"P.Value":0.0142,"adj.P.Val":0.0535},{"tissuedysplasia":0.1185,"tissuetumour":-0.123,"AveExpr":5.805,"F":1.5665,"P.Value":0.2202,"adj.P.Val":0.3602},{"tissuedysplasia":0.3567,"tissuetumour":0.3142,"AveExpr":-1.1184,"F":0.1942,"P.Value":0.8242,"adj.P.Val":0.8768},{"tissuedysplasia":1.152,"tissuetumour":3.6476,"AveExpr":-5.3402,"F":6.3088,"P.Value":0.0039,"adj.P.Val":0.021},{"tissuedysplasia":0.0664,"tissuetumour":-0.0805,"AveExpr":6.753,"F":0.2527,"P.Value":0.7779,"adj.P.Val":0.843},{"tissuedysplasia":-0.0462,"tissuetumour":0.7201,"AveExpr":4.1664,"F":15.2775,"P.Value":9.1482e-06,"adj.P.Val":0.0002},{"tissuedysplasia":0.4085,"tissuetumour":1.5044,"AveExpr":0.2074,"F":13.2265,"P.Value":0,"adj.P.Val":0.0006},{"tissuedysplasia":0.7259,"tissuetumour":1.3578,"AveExpr":4.8789,"F":22.1338,"P.Value":2.2289e-07,"adj.P.Val":0},{"tissuedysplasia":-0.0633,"tissuetumour":0.2791,"AveExpr":3.5554,"F":1.2591,"P.Value":0.2939,"adj.P.Val":0.4384},{"tissuedysplasia":0.0618,"tissuetumour":-0.2408,"AveExpr":0.3379,"F":0.268,"P.Value":0.7662,"adj.P.Val":0.8341},{"tissuedysplasia":1.081,"tissuetumour":1.4803,"AveExpr":-1.7383,"F":2.2908,"P.Value":0.1131,"adj.P.Val":0.2294},{"tissuedysplasia":0.042,"tissuetumour":0.0973,"AveExpr":1.7078,"F":0.0629,"P.Value":0.9391,"adj.P.Val":0.9577},{"tissuedysplasia":0.099,"tissuetumour":-0.1047,"AveExpr":5.31,"F":2.4023,"P.Value":0.1023,"adj.P.Val":0.2142},{"tissuedysplasia":1.4873,"tissuetumour":2.1295,"AveExpr":-9.6153,"F":2.1986,"P.Value":0.123,"adj.P.Val":0.2424},{"tissuedysplasia":0.7894,"tissuetumour":1.0352,"AveExpr":0.2574,"F":2.6649,"P.Value":0.0808,"adj.P.Val":0.182},{"tissuedysplasia":0.1848,"tissuetumour":1.6765,"AveExpr":-8.5251,"F":1.7661,"P.Value":0.1829,"adj.P.Val":0.3179},{"tissuedysplasia":0.1116,"tissuetumour":0.0864,"AveExpr":1.9336,"F":0.1246,"P.Value":0.8832,"adj.P.Val":0.9181},{"tissuedysplasia":0.1313,"tissuetumour":0.1248,"AveExpr":5.9273,"F":0.6489,"P.Value":0.5276,"adj.P.Val":0.6504},{"tissuedysplasia":0.9432,"tissuetumour":1.1065,"AveExpr":3.9327,"F":4.307,"P.Value":0.0201,"adj.P.Val":0.0687},{"tissuedysplasia":-0.0031,"tissuetumour":-0.7303,"AveExpr":0.1565,"F":1.6847,"P.Value":0.1972,"adj.P.Val":0.3341},{"tissuedysplasia":0.2916,"tissuetumour":0.6555,"AveExpr":2.7877,"F":6.4356,"P.Value":0.0035,"adj.P.Val":0.0196},{"tissuedysplasia":0.0279,"tissuetumour":0.2184,"AveExpr":3.0657,"F":0.9475,"P.Value":0.3955,"adj.P.Val":0.536},{"tissuedysplasia":0.0019,"tissuetumour":1.3399,"AveExpr":-2.3141,"F":3.1485,"P.Value":0.0527,"adj.P.Val":0.1355},{"tissuedysplasia":-0.336,"tissuetumour":1.3296,"AveExpr":-2.775,"F":4.5348,"P.Value":0.0162,"adj.P.Val":0.0588},{"tissuedysplasia":-0.1794,"tissuetumour":0.0666,"AveExpr":3.625,"F":1.1026,"P.Value":0.341,"adj.P.Val":0.485},{"tissuedysplasia":0.047,"tissuetumour":0.7753,"AveExpr":3.1899,"F":9.8697,"P.Value":0.0003,"adj.P.Val":0.0031},{"tissuedysplasia":-0.4219,"tissuetumour":0.2916,"AveExpr":-1.7987,"F":0.7374,"P.Value":0.4842,"adj.P.Val":0.613},{"tissuedysplasia":0.0259,"tissuetumour":0.4335,"AveExpr":6.5866,"F":3.6335,"P.Value":0.0348,"adj.P.Val":0.1011},{"tissuedysplasia":0.0337,"tissuetumour":1.2393,"AveExpr":-5.7417,"F":0.6284,"P.Value":0.5382,"adj.P.Val":0.6586},{"tissuedysplasia":0.1604,"tissuetumour":0.3514,"AveExpr":5.4813,"F":3.3894,"P.Value":0.0427,"adj.P.Val":0.117},{"tissuedysplasia":0.1387,"tissuetumour":0.8453,"AveExpr":5.2575,"F":11.9797,"P.Value":0.0001,"adj.P.Val":0.0011},{"tissuedysplasia":0.8456,"tissuetumour":1.2538,"AveExpr":1.0049,"F":7.3747,"P.Value":0.0017,"adj.P.Val":0.0116},{"tissuedysplasia":0.2962,"tissuetumour":0.9548,"AveExpr":2.8072,"F":13.802,"P.Value":0,"adj.P.Val":0.0004},{"tissuedysplasia":0.2192,"tissuetumour":0.8303,"AveExpr":3.6193,"F":13.7569,"P.Value":0,"adj.P.Val":0.0004},{"tissuedysplasia":-0.2317,"tissuetumour":-0.9292,"AveExpr":6.5975,"F":18.4991,"P.Value":1.4768e-06,"adj.P.Val":0.0001},{"tissuedysplasia":1.5032,"tissuetumour":2.5589,"AveExpr":-5.5651,"F":2.8999,"P.Value":0.0656,"adj.P.Val":0.1576},{"tissuedysplasia":-0.0088,"tissuetumour":-0.5446,"AveExpr":3.044,"F":4.4348,"P.Value":0.0176,"adj.P.Val":0.0625},{"tissuedysplasia":0.1382,"tissuetumour":-0.0315,"AveExpr":2.8725,"F":0.3178,"P.Value":0.7294,"adj.P.Val":0.8069},{"tissuedysplasia":-0.1116,"tissuetumour":0.0171,"AveExpr":5.149,"F":0.6803,"P.Value":0.5117,"adj.P.Val":0.6367},{"tissuedysplasia":0.2328,"tissuetumour":0.5965,"AveExpr":4.0644,"F":7.9612,"P.Value":0.0011,"adj.P.Val":0.0084},{"tissuedysplasia":0.8617,"tissuetumour":1.6775,"AveExpr":-0.7367,"F":5.7843,"P.Value":0.0059,"adj.P.Val":0.0283},{"tissuedysplasia":0.6143,"tissuetumour":2.0843,"AveExpr":-4.7244,"F":2.6398,"P.Value":0.0827,"adj.P.Val":0.1847},{"tissuedysplasia":0.0368,"tissuetumour":0.728,"AveExpr":5.0786,"F":23.0111,"P.Value":1.4455e-07,"adj.P.Val":9.2295e-06},{"tissuedysplasia":-0.729,"tissuetumour":0.7005,"AveExpr":-3.373,"F":1.1177,"P.Value":0.3361,"adj.P.Val":0.4805},{"tissuedysplasia":0.4184,"tissuetumour":-0.6728,"AveExpr":1.898,"F":5.0993,"P.Value":0.0102,"adj.P.Val":0.042},{"tissuedysplasia":0.5612,"tissuetumour":1.5891,"AveExpr":3.4579,"F":23.7442,"P.Value":1.0131e-07,"adj.P.Val":7.2054e-06},{"tissuedysplasia":0.4905,"tissuetumour":1.4917,"AveExpr":-0.7511,"F":4.3544,"P.Value":0.0188,"adj.P.Val":0.0656},{"tissuedysplasia":0.2656,"tissuetumour":-0.0132,"AveExpr":6.5781,"F":1.8271,"P.Value":0.1731,"adj.P.Val":0.3061},{"tissuedysplasia":0.2293,"tissuetumour":1.6339,"AveExpr":-0.2137,"F":8.2035,"P.Value":0.0009,"adj.P.Val":0.0074},{"tissuedysplasia":-0.1494,"tissuetumour":0.3967,"AveExpr":3.1296,"F":5.0441,"P.Value":0.0107,"adj.P.Val":0.0434},{"tissuedysplasia":-0.0968,"tissuetumour":0.1221,"AveExpr":2.5865,"F":0.9159,"P.Value":0.4076,"adj.P.Val":0.5471},{"tissuedysplasia":-0.361,"tissuetumour":-0.0356,"AveExpr":3.2626,"F":1.2971,"P.Value":0.2836,"adj.P.Val":0.428},{"tissuedysplasia":1.0157,"tissuetumour":1.886,"AveExpr":-1.5268,"F":4.7947,"P.Value":0.0131,"adj.P.Val":0.0503},{"tissuedysplasia":0.232,"tissuetumour":0.5585,"AveExpr":3.7301,"F":7.0495,"P.Value":0.0022,"adj.P.Val":0.0139},{"tissuedysplasia":-0.0581,"tissuetumour":-0.1812,"AveExpr":6.5735,"F":1.7684,"P.Value":0.1825,"adj.P.Val":0.3175},{"tissuedysplasia":-0.4585,"tissuetumour":-1.2277,"AveExpr":5.4282,"F":31.3497,"P.Value":3.4359e-09,"adj.P.Val":6.3105e-07},{"tissuedysplasia":-0.1429,"tissuetumour":0.2133,"AveExpr":3.7306,"F":1.8153,"P.Value":0.1748,"adj.P.Val":0.3081},{"tissuedysplasia":-0.7146,"tissuetumour":-0.733,"AveExpr":3.8178,"F":10.9655,"P.Value":0.0001,"adj.P.Val":0.0018},{"tissuedysplasia":-0.4566,"tissuetumour":-0.2111,"AveExpr":1.2767,"F":1.2159,"P.Value":0.3062,"adj.P.Val":0.4508},{"tissuedysplasia":0.6321,"tissuetumour":0.8309,"AveExpr":1.989,"F":3.1239,"P.Value":0.0539,"adj.P.Val":0.1377},{"tissuedysplasia":-0.0133,"tissuetumour":1.2386,"AveExpr":-0.1249,"F":4.703,"P.Value":0.0141,"adj.P.Val":0.0532},{"tissuedysplasia":0.6752,"tissuetumour":0.9324,"AveExpr":-0.8584,"F":1.4433,"P.Value":0.2471,"adj.P.Val":0.3898},{"tissuedysplasia":0.0418,"tissuetumour":0.3135,"AveExpr":2.9264,"F":1.8423,"P.Value":0.1705,"adj.P.Val":0.3029},{"tissuedysplasia":0.3992,"tissuetumour":0.3617,"AveExpr":2.3352,"F":1.8553,"P.Value":0.1684,"adj.P.Val":0.3005},{"tissuedysplasia":0.2399,"tissuetumour":0.6398,"AveExpr":3.9878,"F":5.2175,"P.Value":0.0093,"adj.P.Val":0.0392},{"tissuedysplasia":-1.0373,"tissuetumour":-1.4771,"AveExpr":2.1089,"F":15.874,"P.Value":6.4511e-06,"adj.P.Val":0.0002},{"tissuedysplasia":1.0979,"tissuetumour":0.8681,"AveExpr":1.434,"F":4.5402,"P.Value":0.0161,"adj.P.Val":0.0586},{"tissuedysplasia":0.1865,"tissuetumour":1.6744,"AveExpr":0.0787,"F":10.1158,"P.Value":0.0002,"adj.P.Val":0.0027},{"tissuedysplasia":-0.5982,"tissuetumour":-0.2766,"AveExpr":3.386,"F":2.9098,"P.Value":0.065,"adj.P.Val":0.1566},{"tissuedysplasia":0.3864,"tissuetumour":1.0204,"AveExpr":-0.7773,"F":1.279,"P.Value":0.2885,"adj.P.Val":0.433},{"tissuedysplasia":-0.0056,"tissuetumour":-0.429,"AveExpr":5.1713,"F":3.9884,"P.Value":0.0256,"adj.P.Val":0.0816},{"tissuedysplasia":0.6213,"tissuetumour":0.9703,"AveExpr":1.8084,"F":6.2717,"P.Value":0.004,"adj.P.Val":0.0214},{"tissuedysplasia":0.299,"tissuetumour":0.1651,"AveExpr":5.7851,"F":2.2271,"P.Value":0.1199,"adj.P.Val":0.2382},{"tissuedysplasia":2.917,"tissuetumour":3.5253,"AveExpr":-7.7945,"F":5.6747,"P.Value":0.0064,"adj.P.Val":0.0302},{"tissuedysplasia":-0.0887,"tissuetumour":0.1174,"AveExpr":2.304,"F":0.612,"P.Value":0.5468,"adj.P.Val":0.6654},{"tissuedysplasia":-0.9331,"tissuetumour":-1.6036,"AveExpr":2.8569,"F":24.8272,"P.Value":6.0544e-08,"adj.P.Val":4.9771e-06},{"tissuedysplasia":-0.2276,"tissuetumour":-0.8866,"AveExpr":8.0026,"F":23.8937,"P.Value":1.2077e-07,"adj.P.Val":8.1123e-06},{"tissuedysplasia":-0.3515,"tissuetumour":-0.578,"AveExpr":7.1825,"F":13.9464,"P.Value":0,"adj.P.Val":0.0004},{"tissuedysplasia":0.5699,"tissuetumour":1.097,"AveExpr":2.4944,"F":9.4563,"P.Value":0.0004,"adj.P.Val":0.0038},{"tissuedysplasia":0.0275,"tissuetumour":-0.183,"AveExpr":5.7509,"F":1.2814,"P.Value":0.2878,"adj.P.Val":0.4323},{"tissuedysplasia":0.0297,"tissuetumour":0.2114,"AveExpr":4.6594,"F":1.3036,"P.Value":0.2818,"adj.P.Val":0.4265},{"tissuedysplasia":0.0219,"tissuetumour":0.0703,"AveExpr":5.6976,"F":0.3253,"P.Value":0.724,"adj.P.Val":0.8028},{"tissuedysplasia":0.121,"tissuetumour":0.7936,"AveExpr":-1.085,"F":0.9396,"P.Value":0.3985,"adj.P.Val":0.5388},{"tissuedysplasia":-2.146,"tissuetumour":0.0904,"AveExpr":-1.3791,"F":8.5807,"P.Value":0.0007,"adj.P.Val":0.0061},{"tissuedysplasia":0.2483,"tissuetumour":-0.1381,"AveExpr":5.8257,"F":3.5563,"P.Value":0.037,"adj.P.Val":0.1057},{"tissuedysplasia":0.1814,"tissuetumour":-0.0589,"AveExpr":5.017,"F":1.7673,"P.Value":0.1827,"adj.P.Val":0.3176},{"tissuedysplasia":0.144,"tissuetumour":-0.1051,"AveExpr":3.041,"F":0.6325,"P.Value":0.536,"adj.P.Val":0.6572},{"tissuedysplasia":0.2039,"tissuetumour":0.4086,"AveExpr":1.2691,"F":0.9427,"P.Value":0.3973,"adj.P.Val":0.5378},{"tissuedysplasia":-0.0361,"tissuetumour":-0.5203,"AveExpr":5.0737,"F":6.1918,"P.Value":0.0043,"adj.P.Val":0.0224},{"tissuedysplasia":0.3302,"tissuetumour":0.3796,"AveExpr":4.6906,"F":4.2967,"P.Value":0.0197,"adj.P.Val":0.068},{"tissuedysplasia":0.49,"tissuetumour":-0.2254,"AveExpr":2.1429,"F":3.6042,"P.Value":0.0355,"adj.P.Val":0.1025},{"tissuedysplasia":0.1322,"tissuetumour":-0.0766,"AveExpr":5.2387,"F":1.315,"P.Value":0.2788,"adj.P.Val":0.4234},{"tissuedysplasia":0.2505,"tissuetumour":0.1139,"AveExpr":6.7526,"F":1.526,"P.Value":0.2287,"adj.P.Val":0.3698},{"tissuedysplasia":0.2396,"tissuetumour":0.5219,"AveExpr":4.2297,"F":7.9176,"P.Value":0.0012,"adj.P.Val":0.0086},{"tissuedysplasia":-0.1416,"tissuetumour":-0.4281,"AveExpr":6.8363,"F":7.2393,"P.Value":0.0019,"adj.P.Val":0.0125},{"tissuedysplasia":3.2131,"tissuetumour":3.9872,"AveExpr":-4.3341,"F":8.6356,"P.Value":0.0007,"adj.P.Val":0.0059},{"tissuedysplasia":-0.3142,"tissuetumour":-0.268,"AveExpr":2.591,"F":1.5314,"P.Value":0.2275,"adj.P.Val":0.3685},{"tissuedysplasia":0.0807,"tissuetumour":0.4648,"AveExpr":2.3421,"F":0.7345,"P.Value":0.4855,"adj.P.Val":0.6144},{"tissuedysplasia":0.0663,"tissuetumour":-0.1869,"AveExpr":6.3373,"F":1.5833,"P.Value":0.2168,"adj.P.Val":0.3564},{"tissuedysplasia":0.2465,"tissuetumour":0.1522,"AveExpr":5.173,"F":1.9092,"P.Value":0.1603,"adj.P.Val":0.2903},{"tissuedysplasia":2.6007,"tissuetumour":4.0462,"AveExpr":-2.9688,"F":11.7407,"P.Value":0.0001,"adj.P.Val":0.0012},{"tissuedysplasia":4.0982,"tissuetumour":5.3107,"AveExpr":-2.1081,"F":30.357,"P.Value":5.1941e-09,"adj.P.Val":8.6375e-07},{"tissuedysplasia":0.196,"tissuetumour":0.7078,"AveExpr":5.178,"F":9.5966,"P.Value":0.0003,"adj.P.Val":0.0036},{"tissuedysplasia":0.0668,"tissuetumour":0.7606,"AveExpr":3.8826,"F":14.2094,"P.Value":0,"adj.P.Val":0.0004},{"tissuedysplasia":0.2161,"tissuetumour":0.7883,"AveExpr":2.6146,"F":7.8928,"P.Value":0.0012,"adj.P.Val":0.0088},{"tissuedysplasia":0.1822,"tissuetumour":0.7175,"AveExpr":2.8726,"F":4.5861,"P.Value":0.0155,"adj.P.Val":0.057},{"tissuedysplasia":1.1037,"tissuetumour":1.3945,"AveExpr":3.1977,"F":13.5308,"P.Value":0,"adj.P.Val":0.0005},{"tissuedysplasia":-0.4288,"tissuetumour":-0.3692,"AveExpr":4.0664,"F":4.3045,"P.Value":0.0196,"adj.P.Val":0.0677},{"tissuedysplasia":-0.3202,"tissuetumour":-0.5351,"AveExpr":2.3528,"F":1.6196,"P.Value":0.2096,"adj.P.Val":0.3481},{"tissuedysplasia":0.3278,"tissuetumour":1.1434,"AveExpr":2.7258,"F":14.0225,"P.Value":0,"adj.P.Val":0.0004},{"tissuedysplasia":0.3,"tissuetumour":0.9059,"AveExpr":-7.5053,"F":0.3009,"P.Value":0.7417,"adj.P.Val":0.8161},{"tissuedysplasia":0.1108,"tissuetumour":-0.2018,"AveExpr":5.9505,"F":3.2073,"P.Value":0.0501,"adj.P.Val":0.1309},{"tissuedysplasia":-0.0797,"tissuetumour":-0.1083,"AveExpr":2.9612,"F":0.1862,"P.Value":0.8308,"adj.P.Val":0.8811},{"tissuedysplasia":0.3534,"tissuetumour":0.7306,"AveExpr":5.047,"F":14.0291,"P.Value":0,"adj.P.Val":0.0004},{"tissuedysplasia":0.1795,"tissuetumour":0.535,"AveExpr":4.1994,"F":9.5062,"P.Value":0.0004,"adj.P.Val":0.0037},{"tissuedysplasia":0.3025,"tissuetumour":0.7474,"AveExpr":4.7617,"F":11.1074,"P.Value":0.0001,"adj.P.Val":0.0016},{"tissuedysplasia":0.1199,"tissuetumour":0.7645,"AveExpr":1.8007,"F":7.2491,"P.Value":0.0019,"adj.P.Val":0.0125},{"tissuedysplasia":0.0569,"tissuetumour":-0.3015,"AveExpr":7.7939,"F":7.3854,"P.Value":0.0017,"adj.P.Val":0.0116},{"tissuedysplasia":0.1156,"tissuetumour":-0.1412,"AveExpr":6.5444,"F":5.1341,"P.Value":0.0099,"adj.P.Val":0.0411},{"tissuedysplasia":0.1917,"tissuetumour":0.113,"AveExpr":6.3282,"F":1.6941,"P.Value":0.1955,"adj.P.Val":0.3321},{"tissuedysplasia":0.9821,"tissuetumour":3.0495,"AveExpr":-2.821,"F":5.9627,"P.Value":0.0051,"adj.P.Val":0.0255},{"tissuedysplasia":0.2226,"tissuetumour":-0.0367,"AveExpr":4.2669,"F":2.7078,"P.Value":0.0778,"adj.P.Val":0.1772},{"tissuedysplasia":0.1993,"tissuetumour":0.0504,"AveExpr":4.465,"F":2.0603,"P.Value":0.1395,"adj.P.Val":0.2643},{"tissuedysplasia":-0.1187,"tissuetumour":0.6051,"AveExpr":2.5099,"F":5.2453,"P.Value":0.0091,"adj.P.Val":0.0386},{"tissuedysplasia":0.6727,"tissuetumour":0.843,"AveExpr":3.4529,"F":4.2142,"P.Value":0.0212,"adj.P.Val":0.0713},{"tissuedysplasia":0.1956,"tissuetumour":0.5379,"AveExpr":3.1627,"F":5.1899,"P.Value":0.0095,"adj.P.Val":0.0398},{"tissuedysplasia":0.7188,"tissuetumour":1.9277,"AveExpr":-3.3436,"F":2.6196,"P.Value":0.0842,"adj.P.Val":0.1871},{"tissuedysplasia":0.1793,"tissuetumour":0.1071,"AveExpr":5.2536,"F":2.0672,"P.Value":0.1386,"adj.P.Val":0.2631},{"tissuedysplasia":0.3488,"tissuetumour":1.0511,"AveExpr":2.0455,"F":9.1317,"P.Value":0.0005,"adj.P.Val":0.0045},{"tissuedysplasia":0.2386,"tissuetumour":0.3056,"AveExpr":3.8242,"F":3.7062,"P.Value":0.0325,"adj.P.Val":0.0965},{"tissuedysplasia":0.2,"tissuetumour":-1.2376,"AveExpr":-2.7332,"F":1.772,"P.Value":0.1819,"adj.P.Val":0.3169},{"tissuedysplasia":-0.0597,"tissuetumour":0.4034,"AveExpr":1.5737,"F":2.3764,"P.Value":0.1047,"adj.P.Val":0.2176},{"tissuedysplasia":0.7349,"tissuetumour":1.5417,"AveExpr":1.4092,"F":15.3235,"P.Value":8.9033e-06,"adj.P.Val":0.0002},{"tissuedysplasia":0.8303,"tissuetumour":2.3093,"AveExpr":5.0367,"F":44.8579,"P.Value":2.3957e-11,"adj.P.Val":1.76e-08},{"tissuedysplasia":0.4296,"tissuetumour":0.4939,"AveExpr":4.8511,"F":11.904,"P.Value":0.0001,"adj.P.Val":0.0011},{"tissuedysplasia":0.3028,"tissuetumour":0.743,"AveExpr":1.654,"F":2.4939,"P.Value":0.0942,"adj.P.Val":0.2025},{"tissuedysplasia":0.2538,"tissuetumour":1.0767,"AveExpr":-1.7757,"F":1.596,"P.Value":0.2142,"adj.P.Val":0.3535},{"tissuedysplasia":0.5514,"tissuetumour":1.278,"AveExpr":0.4786,"F":6.6767,"P.Value":0.0029,"adj.P.Val":0.0171},{"tissuedysplasia":0.2876,"tissuetumour":-0.0204,"AveExpr":4.8623,"F":1.0379,"P.Value":0.3627,"adj.P.Val":0.5058},{"tissuedysplasia":0.3627,"tissuetumour":0.4506,"AveExpr":5.491,"F":6.6384,"P.Value":0.003,"adj.P.Val":0.0175},{"tissuedysplasia":-0.0352,"tissuetumour":-0.4981,"AveExpr":4.8091,"F":12.1127,"P.Value":0.0001,"adj.P.Val":0.001},{"tissuedysplasia":0.4782,"tissuetumour":1.1794,"AveExpr":3.8165,"F":34.4155,"P.Value":1.005e-09,"adj.P.Val":2.5032e-07},{"tissuedysplasia":0.5368,"tissuetumour":0.9434,"AveExpr":4.7305,"F":24.7353,"P.Value":6.3218e-08,"adj.P.Val":5.1035e-06},{"tissuedysplasia":1.2571,"tissuetumour":0.6111,"AveExpr":5.546,"F":6.8305,"P.Value":0.0029,"adj.P.Val":0.0171},{"tissuedysplasia":0.3729,"tissuetumour":0.2079,"AveExpr":4.2407,"F":3.1504,"P.Value":0.0526,"adj.P.Val":0.1354},{"tissuedysplasia":0.2346,"tissuetumour":0.1753,"AveExpr":4.8125,"F":2.1453,"P.Value":0.1291,"adj.P.Val":0.2506},{"tissuedysplasia":-0.3901,"tissuetumour":-0.1614,"AveExpr":2.9803,"F":1.4974,"P.Value":0.2349,"adj.P.Val":0.3764},{"tissuedysplasia":0.2757,"tissuetumour":0.3806,"AveExpr":4.9731,"F":4.9511,"P.Value":0.0115,"adj.P.Val":0.0457},{"tissuedysplasia":0.3098,"tissuetumour":0.4588,"AveExpr":5.488,"F":6.7037,"P.Value":0.0029,"adj.P.Val":0.0169},{"tissuedysplasia":0.3607,"tissuetumour":1.3344,"AveExpr":1.6664,"F":17.7224,"P.Value":2.2613e-06,"adj.P.Val":0.0001},{"tissuedysplasia":-0.2589,"tissuetumour":-0.4197,"AveExpr":5.8379,"F":4.1196,"P.Value":0.0229,"adj.P.Val":0.0754},{"tissuedysplasia":0.6425,"tissuetumour":1.3167,"AveExpr":-0.5426,"F":4.3095,"P.Value":0.0195,"adj.P.Val":0.0675},{"tissuedysplasia":0.1444,"tissuetumour":0.2031,"AveExpr":5.0845,"F":1.6762,"P.Value":0.1988,"adj.P.Val":0.3357},{"tissuedysplasia":0.1877,"tissuetumour":0.4226,"AveExpr":5.1846,"F":8.3381,"P.Value":0.0009,"adj.P.Val":0.0069},{"tissuedysplasia":0.3105,"tissuetumour":0.4952,"AveExpr":4.0946,"F":7.2264,"P.Value":0.0019,"adj.P.Val":0.0126},{"tissuedysplasia":0.3819,"tissuetumour":1.1653,"AveExpr":4.1917,"F":23.5154,"P.Value":1.1313e-07,"adj.P.Val":7.7706e-06},{"tissuedysplasia":0.2807,"tissuetumour":1.151,"AveExpr":2.8468,"F":13.8302,"P.Value":0,"adj.P.Val":0.0004},{"tissuedysplasia":0.2446,"tissuetumour":0.8354,"AveExpr":3.7041,"F":14.7709,"P.Value":0,"adj.P.Val":0.0003},{"tissuedysplasia":0.105,"tissuetumour":0.241,"AveExpr":4.4792,"F":1.044,"P.Value":0.3606,"adj.P.Val":0.5038},{"tissuedysplasia":0.064,"tissuetumour":1.1771,"AveExpr":0.8656,"F":8.4374,"P.Value":0.0008,"adj.P.Val":0.0065},{"tissuedysplasia":0.1667,"tissuetumour":1.433,"AveExpr":-4.7617,"F":1.3686,"P.Value":0.2651,"adj.P.Val":0.4084},{"tissuedysplasia":0.2018,"tissuetumour":0.3112,"AveExpr":3.5262,"F":1.9876,"P.Value":0.1491,"adj.P.Val":0.2765},{"tissuedysplasia":0.2273,"tissuetumour":0.7446,"AveExpr":4.6261,"F":11.6429,"P.Value":0.0001,"adj.P.Val":0.0013},{"tissuedysplasia":0.2851,"tissuetumour":0.9314,"AveExpr":2.1567,"F":12.0277,"P.Value":0.0001,"adj.P.Val":0.001},{"tissuedysplasia":0.8086,"tissuetumour":2.0699,"AveExpr":1.3979,"F":23.623,"P.Value":1.074e-07,"adj.P.Val":7.5099e-06},{"tissuedysplasia":0.0959,"tissuetumour":0.8303,"AveExpr":-0.5975,"F":1.928,"P.Value":0.1575,"adj.P.Val":0.287},{"tissuedysplasia":0.8452,"tissuetumour":1.3092,"AveExpr":4.9239,"F":21.7925,"P.Value":2.6441e-07,"adj.P.Val":0},{"tissuedysplasia":0.2369,"tissuetumour":0.168,"AveExpr":5.2629,"F":2.7204,"P.Value":0.0769,"adj.P.Val":0.1759},{"tissuedysplasia":0.0843,"tissuetumour":0.4822,"AveExpr":5.29,"F":2.5232,"P.Value":0.0917,"adj.P.Val":0.1988},{"tissuedysplasia":0.1965,"tissuetumour":0.4429,"AveExpr":4.8807,"F":5.8668,"P.Value":0.0055,"adj.P.Val":0.027},{"tissuedysplasia":0.2047,"tissuetumour":0.9605,"AveExpr":2.6561,"F":11.4084,"P.Value":0.0001,"adj.P.Val":0.0014},{"tissuedysplasia":1.282,"tissuetumour":1.9099,"AveExpr":-11.0331,"F":2.8333,"P.Value":0.0696,"adj.P.Val":0.1643},{"tissuedysplasia":0.2291,"tissuetumour":1.1394,"AveExpr":2.7749,"F":18.2915,"P.Value":1.6536e-06,"adj.P.Val":0.0001},{"tissuedysplasia":0.341,"tissuetumour":0.54,"AveExpr":6.6921,"F":5.9808,"P.Value":0.005,"adj.P.Val":0.0252},{"tissuedysplasia":0.3275,"tissuetumour":0.6164,"AveExpr":4.6858,"F":17.9745,"P.Value":1.9674e-06,"adj.P.Val":0.0001},{"tissuedysplasia":0.138,"tissuetumour":0.3837,"AveExpr":2.2446,"F":1.0339,"P.Value":0.3641,"adj.P.Val":0.5071},{"tissuedysplasia":0.3131,"tissuetumour":0.3691,"AveExpr":1.8891,"F":0.9909,"P.Value":0.3794,"adj.P.Val":0.5215},{"tissuedysplasia":0.0809,"tissuetumour":0.7354,"AveExpr":3.0054,"F":10.1122,"P.Value":0.0002,"adj.P.Val":0.0027},{"tissuedysplasia":0.4722,"tissuetumour":0.8187,"AveExpr":2.9738,"F":15.7776,"P.Value":6.8232e-06,"adj.P.Val":0.0002},{"tissuedysplasia":0.6299,"tissuetumour":1.4456,"AveExpr":4.3998,"F":30.7453,"P.Value":4.4148e-09,"adj.P.Val":7.7878e-07},{"tissuedysplasia":0.2072,"tissuetumour":0.9653,"AveExpr":1.3186,"F":8.8905,"P.Value":0.0006,"adj.P.Val":0.0051},{"tissuedysplasia":0.9259,"tissuetumour":1.7296,"AveExpr":-1.14,"F":5.2083,"P.Value":0.0093,"adj.P.Val":0.0394},{"tissuedysplasia":0.4326,"tissuetumour":0.9739,"AveExpr":6.3242,"F":8.4183,"P.Value":0.0009,"adj.P.Val":0.0071},{"tissuedysplasia":0.2089,"tissuetumour":0.9926,"AveExpr":1.5709,"F":7.4873,"P.Value":0.0016,"adj.P.Val":0.0109},{"tissuedysplasia":0.1566,"tissuetumour":0.5752,"AveExpr":4.368,"F":5.5769,"P.Value":0.0069,"adj.P.Val":0.0319},{"tissuedysplasia":0.1489,"tissuetumour":0.6772,"AveExpr":4.508,"F":10.2557,"P.Value":0.0002,"adj.P.Val":0.0025},{"tissuedysplasia":1.2968,"tissuetumour":2.467,"AveExpr":-3.2428,"F":4.3037,"P.Value":0.0196,"adj.P.Val":0.0677},{"tissuedysplasia":0.4471,"tissuetumour":0.595,"AveExpr":2.6908,"F":5.5524,"P.Value":0.0071,"adj.P.Val":0.0324},{"tissuedysplasia":0.2306,"tissuetumour":0.3227,"AveExpr":4.9709,"F":5.2635,"P.Value":0.0089,"adj.P.Val":0.0382},{"tissuedysplasia":-0.3347,"tissuetumour":1.2582,"AveExpr":-2.7215,"F":2.4769,"P.Value":0.0956,"adj.P.Val":0.2046},{"tissuedysplasia":0.6971,"tissuetumour":1.7382,"AveExpr":2.1334,"F":17.5579,"P.Value":2.4775e-06,"adj.P.Val":0.0001},{"tissuedysplasia":-0.3534,"tissuetumour":0.1295,"AveExpr":1.0289,"F":1.297,"P.Value":0.2836,"adj.P.Val":0.428},{"tissuedysplasia":0.1289,"tissuetumour":0.0961,"AveExpr":5.6927,"F":0.6062,"P.Value":0.5499,"adj.P.Val":0.6678},{"tissuedysplasia":0.2406,"tissuetumour":1.0506,"AveExpr":4.2023,"F":10.5633,"P.Value":0.0002,"adj.P.Val":0.0022},{"tissuedysplasia":0.2489,"tissuetumour":1.0292,"AveExpr":6.5375,"F":27.6369,"P.Value":1.6799e-08,"adj.P.Val":2.0483e-06},{"tissuedysplasia":0.0218,"tissuetumour":-0.1539,"AveExpr":6.7814,"F":1.3694,"P.Value":0.2649,"adj.P.Val":0.4083},{"tissuedysplasia":0.569,"tissuetumour":0.3593,"AveExpr":1.4081,"F":2.4135,"P.Value":0.1013,"adj.P.Val":0.2128},{"tissuedysplasia":0.3785,"tissuetumour":0.4688,"AveExpr":3.4939,"F":4.6662,"P.Value":0.0145,"adj.P.Val":0.0544},{"tissuedysplasia":0.3021,"tissuetumour":0.5863,"AveExpr":4.9138,"F":14.6028,"P.Value":0,"adj.P.Val":0.0003},{"tissuedysplasia":1.9004,"tissuetumour":3.7123,"AveExpr":-7.4187,"F":3.7593,"P.Value":0.0311,"adj.P.Val":0.0935},{"tissuedysplasia":0.0539,"tissuetumour":0.2586,"AveExpr":3.7788,"F":1.7849,"P.Value":0.1797,"adj.P.Val":0.3143},{"tissuedysplasia":0.2389,"tissuetumour":0.334,"AveExpr":2.9259,"F":2.0601,"P.Value":0.1396,"adj.P.Val":0.2643},{"tissuedysplasia":0.3703,"tissuetumour":0.3559,"AveExpr":5.4687,"F":8.3737,"P.Value":0.0008,"adj.P.Val":0.0067},{"tissuedysplasia":0.6433,"tissuetumour":2.2047,"AveExpr":-6.3211,"F":1.6614,"P.Value":0.2016,"adj.P.Val":0.3388},{"tissuedysplasia":-0.3679,"tissuetumour":-0.6554,"AveExpr":4.1558,"F":12.7054,"P.Value":0,"adj.P.Val":0.0007},{"tissuedysplasia":0.2607,"tissuetumour":0.1293,"AveExpr":5.8134,"F":2.6041,"P.Value":0.0853,"adj.P.Val":0.1888},{"tissuedysplasia":0.0488,"tissuetumour":0.2539,"AveExpr":4.8233,"F":1.9988,"P.Value":0.1476,"adj.P.Val":0.2747},{"tissuedysplasia":0.2918,"tissuetumour":0.5746,"AveExpr":4.5917,"F":10.5194,"P.Value":0.0002,"adj.P.Val":0.0022},{"tissuedysplasia":0.4162,"tissuetumour":0.8048,"AveExpr":2.5356,"F":6.6313,"P.Value":0.003,"adj.P.Val":0.0175},{"tissuedysplasia":0.4407,"tissuetumour":0.9385,"AveExpr":4.4397,"F":29.2071,"P.Value":8.4667e-09,"adj.P.Val":1.244e-06},{"tissuedysplasia":0.8518,"tissuetumour":2.2097,"AveExpr":-1.5287,"F":5.8733,"P.Value":0.0055,"adj.P.Val":0.0269},{"tissuedysplasia":-0.0351,"tissuetumour":0.7891,"AveExpr":-1.0161,"F":1.6653,"P.Value":0.2008,"adj.P.Val":0.338},{"tissuedysplasia":0.8629,"tissuetumour":0.8974,"AveExpr":2.5321,"F":8.5779,"P.Value":0.0007,"adj.P.Val":0.0061},{"tissuedysplasia":0.3208,"tissuetumour":-0.0368,"AveExpr":6.4169,"F":8.1679,"P.Value":0.001,"adj.P.Val":0.0075},{"tissuedysplasia":0.6198,"tissuetumour":1.7424,"AveExpr":1.2197,"F":18.1999,"P.Value":1.7385e-06,"adj.P.Val":0.0001},{"tissuedysplasia":-0.539,"tissuetumour":-0.6331,"AveExpr":5.127,"F":12.8812,"P.Value":0,"adj.P.Val":0.0007},{"tissuedysplasia":0.3964,"tissuetumour":0.9086,"AveExpr":2.9414,"F":8.9757,"P.Value":0.0005,"adj.P.Val":0.0049},{"tissuedysplasia":0.3214,"tissuetumour":0.4527,"AveExpr":3.0815,"F":2.4563,"P.Value":0.0974,"adj.P.Val":0.2072},{"tissuedysplasia":0.491,"tissuetumour":1.5832,"AveExpr":5.5751,"F":44.9233,"P.Value":2.3446e-11,"adj.P.Val":1.76e-08},{"tissuedysplasia":0.3727,"tissuetumour":0.7255,"AveExpr":4.1435,"F":11.1235,"P.Value":0.0001,"adj.P.Val":0.0016},{"tissuedysplasia":0.3248,"tissuetumour":0.997,"AveExpr":3.5679,"F":13.4712,"P.Value":0,"adj.P.Val":0.0005},{"tissuedysplasia":0.2553,"tissuetumour":0.7067,"AveExpr":3.3799,"F":5.7545,"P.Value":0.006,"adj.P.Val":0.0288},{"tissuedysplasia":0.1401,"tissuetumour":0.3899,"AveExpr":4.2987,"F":5.9311,"P.Value":0.0052,"adj.P.Val":0.026},{"tissuedysplasia":0.3077,"tissuetumour":0.5635,"AveExpr":5.2535,"F":11.959,"P.Value":0.0001,"adj.P.Val":0.0011},{"tissuedysplasia":1.1573,"tissuetumour":1.7116,"AveExpr":2.3324,"F":17.5452,"P.Value":2.4949e-06,"adj.P.Val":0.0001},{"tissuedysplasia":0.0046,"tissuetumour":0.0516,"AveExpr":2.1097,"F":0.0174,"P.Value":0.9828,"adj.P.Val":0.988},{"tissuedysplasia":0.4129,"tissuetumour":0.7501,"AveExpr":2.1744,"F":1.5538,"P.Value":0.2228,"adj.P.Val":0.3632},{"tissuedysplasia":0.1027,"tissuetumour":0.0574,"AveExpr":4.2724,"F":0.3597,"P.Value":0.6999,"adj.P.Val":0.7848},{"tissuedysplasia":0.1655,"tissuetumour":0.1988,"AveExpr":4.6839,"F":1.9544,"P.Value":0.1537,"adj.P.Val":0.2823},{"tissuedysplasia":1.4868,"tissuetumour":3.0703,"AveExpr":0.4893,"F":20.2879,"P.Value":5.706e-07,"adj.P.Val":0},{"tissuedysplasia":0.251,"tissuetumour":0.7314,"AveExpr":5.3227,"F":17.4126,"P.Value":2.6864e-06,"adj.P.Val":0.0001},{"tissuedysplasia":0.2119,"tissuetumour":0.7082,"AveExpr":4.4423,"F":19.6426,"P.Value":8.0034e-07,"adj.P.Val":0},{"tissuedysplasia":1.0563,"tissuetumour":1.5881,"AveExpr":0.988,"F":7.9547,"P.Value":0.0011,"adj.P.Val":0.0084},{"tissuedysplasia":0.3825,"tissuetumour":0.509,"AveExpr":4.9989,"F":6.4873,"P.Value":0.0034,"adj.P.Val":0.019},{"tissuedysplasia":0.5719,"tissuetumour":0.7943,"AveExpr":1.5444,"F":2.919,"P.Value":0.0645,"adj.P.Val":0.1558},{"tissuedysplasia":0.5009,"tissuetumour":0.7425,"AveExpr":3.1808,"F":7.8418,"P.Value":0.0012,"adj.P.Val":0.009},{"tissuedysplasia":-0.1535,"tissuetumour":0.6916,"AveExpr":0.0533,"F":2.2131,"P.Value":0.1214,"adj.P.Val":0.2402},{"tissuedysplasia":-0.2423,"tissuetumour":-0.0167,"AveExpr":4.9779,"F":3.1382,"P.Value":0.0532,"adj.P.Val":0.1364},{"tissuedysplasia":0.1207,"tissuetumour":-0.058,"AveExpr":7.0698,"F":1.9363,"P.Value":0.1563,"adj.P.Val":0.2855},{"tissuedysplasia":0.2921,"tissuetumour":0.3326,"AveExpr":3.8195,"F":2.4963,"P.Value":0.094,"adj.P.Val":0.2022},{"tissuedysplasia":-0.0478,"tissuetumour":0.1592,"AveExpr":2.7889,"F":1.119,"P.Value":0.3357,"adj.P.Val":0.4801},{"tissuedysplasia":0.0435,"tissuetumour":-0.0182,"AveExpr":0.0435,"F":0.0161,"P.Value":0.984,"adj.P.Val":0.989},{"tissuedysplasia":0.5316,"tissuetumour":1.1375,"AveExpr":0.6452,"F":4.4935,"P.Value":0.0168,"adj.P.Val":0.0603},{"tissuedysplasia":0.1528,"tissuetumour":-0.1577,"AveExpr":3.8598,"F":1.5535,"P.Value":0.2229,"adj.P.Val":0.3632},{"tissuedysplasia":1.1045,"tissuetumour":2.1967,"AveExpr":1.7155,"F":16.051,"P.Value":5.8222e-06,"adj.P.Val":0.0002},{"tissuedysplasia":-0.3557,"tissuetumour":-0.944,"AveExpr":6.5752,"F":24.9533,"P.Value":5.7066e-08,"adj.P.Val":4.7571e-06},{"tissuedysplasia":0.7834,"tissuetumour":1.2775,"AveExpr":4.5947,"F":25.786,"P.Value":3.8761e-08,"adj.P.Val":3.6487e-06},{"tissuedysplasia":0.7338,"tissuetumour":1.1138,"AveExpr":-0.8837,"F":3.6978,"P.Value":0.0328,"adj.P.Val":0.097},{"tissuedysplasia":0.0655,"tissuetumour":0.1042,"AveExpr":3.8366,"F":0.2927,"P.Value":0.7477,"adj.P.Val":0.8207},{"tissuedysplasia":0.6548,"tissuetumour":2.077,"AveExpr":4.376,"F":45.9756,"P.Value":1.6634e-11,"adj.P.Val":1.5633e-08},{"tissuedysplasia":0.1726,"tissuetumour":0.6373,"AveExpr":2.4611,"F":4.9977,"P.Value":0.0111,"adj.P.Val":0.0445},{"tissuedysplasia":0.2034,"tissuetumour":0.6307,"AveExpr":4.9215,"F":2.6027,"P.Value":0.0855,"adj.P.Val":0.1891},{"tissuedysplasia":0.1748,"tissuetumour":0.8196,"AveExpr":3.7544,"F":11.4378,"P.Value":0.0001,"adj.P.Val":0.0014},{"tissuedysplasia":-0.1252,"tissuetumour":-0.3195,"AveExpr":5.2614,"F":2.7688,"P.Value":0.0737,"adj.P.Val":0.1708},{"tissuedysplasia":0.2294,"tissuetumour":0.3889,"AveExpr":5.3832,"F":2.7966,"P.Value":0.0719,"adj.P.Val":0.1682},{"tissuedysplasia":0.1339,"tissuetumour":0.0994,"AveExpr":5.4656,"F":0.6766,"P.Value":0.5135,"adj.P.Val":0.6381},{"tissuedysplasia":-0.0989,"tissuetumour":0.3516,"AveExpr":6.8126,"F":7.226,"P.Value":0.0019,"adj.P.Val":0.0126},{"tissuedysplasia":0.3804,"tissuetumour":0.3277,"AveExpr":4.3115,"F":5.0333,"P.Value":0.0108,"adj.P.Val":0.0437},{"tissuedysplasia":0.5595,"tissuetumour":0.8135,"AveExpr":3.2152,"F":5.4444,"P.Value":0.0077,"adj.P.Val":0.0344},{"tissuedysplasia":-0.3248,"tissuetumour":-0.0837,"AveExpr":-0.4465,"F":0.1891,"P.Value":0.8284,"adj.P.Val":0.8796},{"tissuedysplasia":0.1577,"tissuetumour":0.4079,"AveExpr":3.9004,"F":2.8021,"P.Value":0.0715,"adj.P.Val":0.1675},{"tissuedysplasia":-0.9531,"tissuetumour":-1.6092,"AveExpr":2.8085,"F":14.6929,"P.Value":0,"adj.P.Val":0.0003},{"tissuedysplasia":0.3311,"tissuetumour":2.1096,"AveExpr":-2.594,"F":4.2744,"P.Value":0.0201,"adj.P.Val":0.0688},{"tissuedysplasia":0.3692,"tissuetumour":0.1749,"AveExpr":4.712,"F":4.0369,"P.Value":0.0246,"adj.P.Val":0.0792},{"tissuedysplasia":-0.6415,"tissuetumour":-1.186,"AveExpr":7.7416,"F":39.2104,"P.Value":2.024e-10,"adj.P.Val":8.4898e-08},{"tissuedysplasia":1.0607,"tissuetumour":1.1393,"AveExpr":3.0843,"F":9.2625,"P.Value":0.0004,"adj.P.Val":0.0042},{"tissuedysplasia":0.4561,"tissuetumour":0.361,"AveExpr":5.3869,"F":4.9411,"P.Value":0.0116,"adj.P.Val":0.0459},{"tissuedysplasia":0.9497,"tissuetumour":1.2446,"AveExpr":-1.2686,"F":2.0146,"P.Value":0.1455,"adj.P.Val":0.2719},{"tissuedysplasia":-0.1399,"tissuetumour":0.2496,"AveExpr":3.5645,"F":3.0801,"P.Value":0.056,"adj.P.Val":0.1413},{"tissuedysplasia":-0.0509,"tissuetumour":-0.0292,"AveExpr":5.2787,"F":0.1125,"P.Value":0.8938,"adj.P.Val":0.9258},{"tissuedysplasia":0.1404,"tissuetumour":-0.1413,"AveExpr":6.5003,"F":4.6285,"P.Value":0.015,"adj.P.Val":0.0556},{"tissuedysplasia":-0.3579,"tissuetumour":0.0943,"AveExpr":5.3229,"F":2.8567,"P.Value":0.0682,"adj.P.Val":0.1618},{"tissuedysplasia":0.3233,"tissuetumour":1.0955,"AveExpr":3.6889,"F":26.9281,"P.Value":2.3053e-08,"adj.P.Val":2.4961e-06},{"tissuedysplasia":0.4007,"tissuetumour":0.2177,"AveExpr":0.6427,"F":0.8942,"P.Value":0.4162,"adj.P.Val":0.5546},{"tissuedysplasia":0.0318,"tissuetumour":-0.2111,"AveExpr":4.585,"F":3.1316,"P.Value":0.0535,"adj.P.Val":0.137},{"tissuedysplasia":0.3273,"tissuetumour":0.1956,"AveExpr":5.2556,"F":3.0103,"P.Value":0.0595,"adj.P.Val":0.1475},{"tissuedysplasia":0.4948,"tissuetumour":0.7522,"AveExpr":2.5817,"F":4.5537,"P.Value":0.0159,"adj.P.Val":0.0581},{"tissuedysplasia":0.4433,"tissuetumour":0.0776,"AveExpr":4.1055,"F":6.8394,"P.Value":0.0026,"adj.P.Val":0.0156},{"tissuedysplasia":0.028,"tissuetumour":0.0335,"AveExpr":2.6198,"F":0.0146,"P.Value":0.9855,"adj.P.Val":0.9899},{"tissuedysplasia":0.1826,"tissuetumour":0.1822,"AveExpr":4.7875,"F":1.4485,"P.Value":0.2459,"adj.P.Val":0.3886},{"tissuedysplasia":0.5878,"tissuetumour":0.2121,"AveExpr":2.6157,"F":3.0645,"P.Value":0.0568,"adj.P.Val":0.1426},{"tissuedysplasia":0.1084,"tissuetumour":0.3236,"AveExpr":3.2491,"F":1.8113,"P.Value":0.1754,"adj.P.Val":0.3089},{"tissuedysplasia":0.0717,"tissuetumour":0.051,"AveExpr":4.5844,"F":0.1412,"P.Value":0.8687,"adj.P.Val":0.9075},{"tissuedysplasia":0.0739,"tissuetumour":-0.0514,"AveExpr":4.0991,"F":0.2671,"P.Value":0.7668,"adj.P.Val":0.8347},{"tissuedysplasia":0.621,"tissuetumour":2.055,"AveExpr":0.5186,"F":10.0156,"P.Value":0.0003,"adj.P.Val":0.0029},{"tissuedysplasia":0.9873,"tissuetumour":1.334,"AveExpr":3.2872,"F":6.9184,"P.Value":0.0024,"adj.P.Val":0.015},{"tissuedysplasia":0.029,"tissuetumour":-0.308,"AveExpr":3.3405,"F":1.1827,"P.Value":0.316,"adj.P.Val":0.4603},{"tissuedysplasia":0.2708,"tissuetumour":0.3596,"AveExpr":2.8198,"F":2.9935,"P.Value":0.0604,"adj.P.Val":0.1491},{"tissuedysplasia":0.5246,"tissuetumour":0.2157,"AveExpr":2.7031,"F":2.3963,"P.Value":0.1028,"adj.P.Val":0.215},{"tissuedysplasia":-0.2258,"tissuetumour":-0.3222,"AveExpr":1.0888,"F":0.2491,"P.Value":0.7806,"adj.P.Val":0.8447},{"tissuedysplasia":0.3928,"tissuetumour":0.0972,"AveExpr":5.4207,"F":3.8808,"P.Value":0.028,"adj.P.Val":0.087},{"tissuedysplasia":0.8811,"tissuetumour":1.9826,"AveExpr":1.6568,"F":22.0521,"P.Value":2.3216e-07,"adj.P.Val":0},{"tissuedysplasia":0.3476,"tissuetumour":0.4402,"AveExpr":2.827,"F":1.6777,"P.Value":0.1985,"adj.P.Val":0.3354},{"tissuedysplasia":0.059,"tissuetumour":-0.2709,"AveExpr":4.6246,"F":3.2116,"P.Value":0.0499,"adj.P.Val":0.1305},{"tissuedysplasia":0.6061,"tissuetumour":1.4618,"AveExpr":1.5412,"F":8.0089,"P.Value":0.0011,"adj.P.Val":0.0082},{"tissuedysplasia":0.1592,"tissuetumour":-0.0048,"AveExpr":6.0487,"F":0.8025,"P.Value":0.4547,"adj.P.Val":0.5887},{"tissuedysplasia":-0.5964,"tissuetumour":-0.9374,"AveExpr":4.9079,"F":7.3332,"P.Value":0.0018,"adj.P.Val":0.0119},{"tissuedysplasia":0.9103,"tissuetumour":0.9941,"AveExpr":4.1148,"F":18.0621,"P.Value":1.8749e-06,"adj.P.Val":0.0001},{"tissuedysplasia":0.2894,"tissuetumour":0.049,"AveExpr":4.9685,"F":3.117,"P.Value":0.0542,"adj.P.Val":0.1382},{"tissuedysplasia":0.3871,"tissuetumour":0.6369,"AveExpr":3.5819,"F":6.1121,"P.Value":0.0045,"adj.P.Val":0.0235},{"tissuedysplasia":-0.0429,"tissuetumour":-0.8949,"AveExpr":5.3989,"F":13.7275,"P.Value":0,"adj.P.Val":0.0005},{"tissuedysplasia":0.0618,"tissuetumour":0.149,"AveExpr":6.0821,"F":0.5798,"P.Value":0.5642,"adj.P.Val":0.6792},{"tissuedysplasia":0.5834,"tissuetumour":0.4961,"AveExpr":4.5514,"F":2.61,"P.Value":0.0849,"adj.P.Val":0.1882},{"tissuedysplasia":-0.2435,"tissuetumour":0.371,"AveExpr":2.3392,"F":1.8751,"P.Value":0.1654,"adj.P.Val":0.2966},{"tissuedysplasia":-1.0744,"tissuetumour":-1.2013,"AveExpr":-2.2424,"F":2.2979,"P.Value":0.1124,"adj.P.Val":0.2284},{"tissuedysplasia":0.605,"tissuetumour":0.61,"AveExpr":4.7415,"F":10.414,"P.Value":0.0002,"adj.P.Val":0.0023},{"tissuedysplasia":0.1599,"tissuetumour":-0.0323,"AveExpr":3.0378,"F":0.5981,"P.Value":0.5543,"adj.P.Val":0.6713},{"tissuedysplasia":0.1008,"tissuetumour":-0.3896,"AveExpr":5.6506,"F":3.4285,"P.Value":0.0413,"adj.P.Val":0.1142},{"tissuedysplasia":0.0381,"tissuetumour":-0.3805,"AveExpr":6.5044,"F":5.0672,"P.Value":0.0105,"adj.P.Val":0.0428},{"tissuedysplasia":0.1152,"tissuetumour":-0.0934,"AveExpr":4.1678,"F":0.9164,"P.Value":0.4074,"adj.P.Val":0.5469},{"tissuedysplasia":-0.1553,"tissuetumour":-0.8172,"AveExpr":6.7504,"F":18.8151,"P.Value":1.2447e-06,"adj.P.Val":0},{"tissuedysplasia":-0.3361,"tissuetumour":-1.3115,"AveExpr":3.8997,"F":10.2174,"P.Value":0.0002,"adj.P.Val":0.0026},{"tissuedysplasia":-0.1972,"tissuetumour":0.1847,"AveExpr":9.4448,"F":1.2729,"P.Value":0.2923,"adj.P.Val":0.4369},{"tissuedysplasia":0.0008,"tissuetumour":-1.0802,"AveExpr":2.9178,"F":7.4722,"P.Value":0.0016,"adj.P.Val":0.011},{"tissuedysplasia":0.3698,"tissuetumour":-0.0814,"AveExpr":2.0082,"F":1.4192,"P.Value":0.2528,"adj.P.Val":0.3957},{"tissuedysplasia":0.1596,"tissuetumour":0.4911,"AveExpr":-0.494,"F":0.6945,"P.Value":0.5047,"adj.P.Val":0.6307},{"tissuedysplasia":0.4572,"tissuetumour":0.2254,"AveExpr":4.021,"F":3.983,"P.Value":0.0257,"adj.P.Val":0.0818},{"tissuedysplasia":-0.4848,"tissuetumour":-1.2776,"AveExpr":7.3507,"F":52.2013,"P.Value":2.419e-12,"adj.P.Val":4.4921e-09},{"tissuedysplasia":0.1473,"tissuetumour":-0.2463,"AveExpr":4.214,"F":2.9213,"P.Value":0.0644,"adj.P.Val":0.1556},{"tissuedysplasia":0.3469,"tissuetumour":0.2077,"AveExpr":4.1776,"F":4.3683,"P.Value":0.0186,"adj.P.Val":0.0651},{"tissuedysplasia":0.7249,"tissuetumour":0.7937,"AveExpr":3.1065,"F":12.8218,"P.Value":0,"adj.P.Val":0.0007},{"tissuedysplasia":-0.1129,"tissuetumour":-0.3868,"AveExpr":3.6448,"F":2.2678,"P.Value":0.1155,"adj.P.Val":0.2326},{"tissuedysplasia":2.3936,"tissuetumour":3.6628,"AveExpr":-4.7431,"F":5.1549,"P.Value":0.0097,"adj.P.Val":0.0406},{"tissuedysplasia":0.2826,"tissuetumour":0.2491,"AveExpr":7.5427,"F":4.3675,"P.Value":0.0186,"adj.P.Val":0.0651},{"tissuedysplasia":0.6387,"tissuetumour":1.5033,"AveExpr":3.2976,"F":32.427,"P.Value":2.2132e-09,"adj.P.Val":4.5047e-07},{"tissuedysplasia":-0.0494,"tissuetumour":0.2092,"AveExpr":3.9642,"F":1.6936,"P.Value":0.1956,"adj.P.Val":0.3322},{"tissuedysplasia":0.5153,"tissuetumour":1.5795,"AveExpr":0.8558,"F":8.6937,"P.Value":0.0007,"adj.P.Val":0.0057},{"tissuedysplasia":0.18,"tissuetumour":0.7683,"AveExpr":7.2306,"F":28.7864,"P.Value":1.0152e-08,"adj.P.Val":1.4006e-06},{"tissuedysplasia":1.1341,"tissuetumour":2.7514,"AveExpr":-2.6287,"F":6.9256,"P.Value":0.0024,"adj.P.Val":0.0149},{"tissuedysplasia":0.6922,"tissuetumour":0.8477,"AveExpr":6.1135,"F":7.3202,"P.Value":0.0019,"adj.P.Val":0.0126},{"tissuedysplasia":-0.4181,"tissuetumour":-1.0732,"AveExpr":3.9231,"F":14.8395,"P.Value":0,"adj.P.Val":0.0003},{"tissuedysplasia":0.0217,"tissuetumour":-0.38,"AveExpr":5.0597,"F":5.9051,"P.Value":0.0053,"adj.P.Val":0.0264},{"tissuedysplasia":0.3932,"tissuetumour":0.1258,"AveExpr":3.4217,"F":4.1698,"P.Value":0.022,"adj.P.Val":0.0732},{"tissuedysplasia":0.6374,"tissuetumour":1.3185,"AveExpr":4.4933,"F":16.703,"P.Value":4.0064e-06,"adj.P.Val":0.0001},{"tissuedysplasia":0.179,"tissuetumour":-0.2114,"AveExpr":4.2832,"F":5.4224,"P.Value":0.0079,"adj.P.Val":0.0348},{"tissuedysplasia":0.4619,"tissuetumour":-0.0886,"AveExpr":4.4339,"F":9.6616,"P.Value":0.0003,"adj.P.Val":0.0034},{"tissuedysplasia":0.1035,"tissuetumour":-0.1362,"AveExpr":3.591,"F":1.3362,"P.Value":0.2733,"adj.P.Val":0.4177},{"tissuedysplasia":0.3165,"tissuetumour":1.2771,"AveExpr":2.6383,"F":18.2073,"P.Value":1.7315e-06,"adj.P.Val":0.0001},{"tissuedysplasia":-0.0435,"tissuetumour":-0.4228,"AveExpr":5.0707,"F":4.0324,"P.Value":0.0247,"adj.P.Val":0.0794},{"tissuedysplasia":0.3967,"tissuetumour":0.1246,"AveExpr":3.8838,"F":3.1597,"P.Value":0.0522,"adj.P.Val":0.1346},{"tissuedysplasia":0.0932,"tissuetumour":-0.2528,"AveExpr":2.0973,"F":1.0076,"P.Value":0.3734,"adj.P.Val":0.516},{"tissuedysplasia":-0.1818,"tissuetumour":-0.7227,"AveExpr":4.8433,"F":10.7224,"P.Value":0.0002,"adj.P.Val":0.002},{"tissuedysplasia":0.1608,"tissuetumour":0.4911,"AveExpr":5.0005,"F":6.241,"P.Value":0.0041,"adj.P.Val":0.0218},{"tissuedysplasia":0.2686,"tissuetumour":1.7854,"AveExpr":1.25,"F":14.9569,"P.Value":0,"adj.P.Val":0.0003},{"tissuedysplasia":0.3571,"tissuetumour":0.4011,"AveExpr":2.9354,"F":2.3425,"P.Value":0.108,"adj.P.Val":0.2221},{"tissuedysplasia":0.0587,"tissuetumour":0.6391,"AveExpr":4.3947,"F":11.315,"P.Value":0.0001,"adj.P.Val":0.0015},{"tissuedysplasia":0.3427,"tissuetumour":1.0289,"AveExpr":4.893,"F":20.1594,"P.Value":6.1012e-07,"adj.P.Val":0},{"tissuedysplasia":0.5177,"tissuetumour":0.7463,"AveExpr":6.1129,"F":20.3159,"P.Value":5.6236e-07,"adj.P.Val":0},{"tissuedysplasia":0.0593,"tissuetumour":2.0094,"AveExpr":-5.0796,"F":3.3971,"P.Value":0.0425,"adj.P.Val":0.1165},{"tissuedysplasia":-0.0975,"tissuetumour":-0.6593,"AveExpr":5.0745,"F":14.5938,"P.Value":0,"adj.P.Val":0.0003},{"tissuedysplasia":0.4383,"tissuetumour":1.7336,"AveExpr":-2.7512,"F":3.5509,"P.Value":0.0372,"adj.P.Val":0.106},{"tissuedysplasia":-0.1466,"tissuetumour":-0.6753,"AveExpr":5.8282,"F":10.5822,"P.Value":0.0002,"adj.P.Val":0.0021},{"tissuedysplasia":-0.3934,"tissuetumour":-0.8934,"AveExpr":5.4822,"F":9.9327,"P.Value":0.0003,"adj.P.Val":0.003},{"tissuedysplasia":0.4436,"tissuetumour":0.9282,"AveExpr":4.7726,"F":19.9178,"P.Value":6.9237e-07,"adj.P.Val":0},{"tissuedysplasia":0.6708,"tissuetumour":1.0712,"AveExpr":1.0232,"F":4.0985,"P.Value":0.0233,"adj.P.Val":0.0763},{"tissuedysplasia":0.0256,"tissuetumour":-0.9754,"AveExpr":-0.9879,"F":1.819,"P.Value":0.1742,"adj.P.Val":0.3074},{"tissuedysplasia":0.071,"tissuetumour":0.6682,"AveExpr":5.4542,"F":19.7884,"P.Value":7.4109e-07,"adj.P.Val":0},{"tissuedysplasia":0.025,"tissuetumour":1.1471,"AveExpr":-7.0007,"F":0.5653,"P.Value":0.5722,"adj.P.Val":0.6862},{"tissuedysplasia":-0.0208,"tissuetumour":0.7969,"AveExpr":5.0338,"F":17.1832,"P.Value":3.0545e-06,"adj.P.Val":0.0001},{"tissuedysplasia":1.0496,"tissuetumour":0.9688,"AveExpr":2.7236,"F":7.2965,"P.Value":0.0018,"adj.P.Val":0.0122},{"tissuedysplasia":-0.3584,"tissuetumour":1.6729,"AveExpr":-7.2834,"F":1.3924,"P.Value":0.2592,"adj.P.Val":0.4024},{"tissuedysplasia":0.0894,"tissuetumour":-0.0979,"AveExpr":3.7341,"F":0.8213,"P.Value":0.4465,"adj.P.Val":0.582},{"tissuedysplasia":0.1112,"tissuetumour":0.1033,"AveExpr":6.9313,"F":1.0722,"P.Value":0.351,"adj.P.Val":0.4949},{"tissuedysplasia":0.037,"tissuetumour":0.7294,"AveExpr":3.1407,"F":10.2497,"P.Value":0.0002,"adj.P.Val":0.0025},{"tissuedysplasia":0.9143,"tissuetumour":1.5397,"AveExpr":-3.2061,"F":1.5294,"P.Value":0.228,"adj.P.Val":0.3689},{"tissuedysplasia":0.1993,"tissuetumour":3.3233,"AveExpr":-6.8954,"F":5.1815,"P.Value":0.0095,"adj.P.Val":0.0401},{"tissuedysplasia":-0.2451,"tissuetumour":0.7691,"AveExpr":0.3384,"F":5.3335,"P.Value":0.0084,"adj.P.Val":0.0367},{"tissuedysplasia":0.1299,"tissuetumour":0.6016,"AveExpr":2.5147,"F":5.6359,"P.Value":0.0066,"adj.P.Val":0.0308},{"tissuedysplasia":0.4708,"tissuetumour":1.6539,"AveExpr":0.5974,"F":11.1401,"P.Value":0.0001,"adj.P.Val":0.0016},{"tissuedysplasia":-0.0901,"tissuetumour":0.5784,"AveExpr":6.6098,"F":9.5146,"P.Value":0.0004,"adj.P.Val":0.0037},{"tissuedysplasia":0.1292,"tissuetumour":0.218,"AveExpr":5.1103,"F":2.0005,"P.Value":0.1474,"adj.P.Val":0.2744},{"tissuedysplasia":0.9121,"tissuetumour":1.3941,"AveExpr":2.4169,"F":13.6184,"P.Value":0,"adj.P.Val":0.0005},{"tissuedysplasia":0.0218,"tissuetumour":0.5423,"AveExpr":4.7294,"F":12.4485,"P.Value":0.0001,"adj.P.Val":0.0008},{"tissuedysplasia":0.1504,"tissuetumour":0.6137,"AveExpr":0.8651,"F":3.1469,"P.Value":0.0528,"adj.P.Val":0.1356},{"tissuedysplasia":0.0212,"tissuetumour":0.6112,"AveExpr":3.6898,"F":7.4607,"P.Value":0.0016,"adj.P.Val":0.0111},{"tissuedysplasia":0.0682,"tissuetumour":0.9264,"AveExpr":3.472,"F":10.7008,"P.Value":0.0002,"adj.P.Val":0.002},{"tissuedysplasia":-0.0466,"tissuetumour":0.6915,"AveExpr":3.8232,"F":13.1176,"P.Value":0,"adj.P.Val":0.0006},{"tissuedysplasia":-3.3434,"tissuetumour":0.7146,"AveExpr":0.6778,"F":5.5949,"P.Value":0.0075,"adj.P.Val":0.0339},{"tissuedysplasia":0.0391,"tissuetumour":0.2496,"AveExpr":5.1574,"F":1.9796,"P.Value":0.1502,"adj.P.Val":0.2778},{"tissuedysplasia":-0.2423,"tissuetumour":0.3354,"AveExpr":5.9782,"F":5.6836,"P.Value":0.0064,"adj.P.Val":0.03},{"tissuedysplasia":0.0062,"tissuetumour":0.2256,"AveExpr":-1.8196,"F":0.0827,"P.Value":0.9208,"adj.P.Val":0.9449},{"tissuedysplasia":-0.0595,"tissuetumour":0.6276,"AveExpr":3.5327,"F":8.8604,"P.Value":0.0006,"adj.P.Val":0.0052},{"tissuedysplasia":-0.0667,"tissuetumour":0.3025,"AveExpr":5.3932,"F":4.1498,"P.Value":0.0223,"adj.P.Val":0.0741},{"tissuedysplasia":0.0357,"tissuetumour":0.7066,"AveExpr":3.0953,"F":9.7668,"P.Value":0.0003,"adj.P.Val":0.0033},{"tissuedysplasia":0.7963,"tissuetumour":1.7569,"AveExpr":-0.6185,"F":5.5434,"P.Value":0.0071,"adj.P.Val":0.0326},{"tissuedysplasia":0.5423,"tissuetumour":1.3598,"AveExpr":2.2655,"F":19.1581,"P.Value":1.0353e-06,"adj.P.Val":0},{"tissuedysplasia":-0.1427,"tissuetumour":0.5795,"AveExpr":3.002,"F":5.1889,"P.Value":0.0095,"adj.P.Val":0.0399},{"tissuedysplasia":0.6459,"tissuetumour":1.1809,"AveExpr":3.6149,"F":11.6822,"P.Value":0.0001,"adj.P.Val":0.0012},{"tissuedysplasia":0.2774,"tissuetumour":0.8359,"AveExpr":3.9595,"F":14.2948,"P.Value":0,"adj.P.Val":0.0003},{"tissuedysplasia":-0.3811,"tissuetumour":0.9971,"AveExpr":-3.4832,"F":1.8682,"P.Value":0.1664,"adj.P.Val":0.2979},{"tissuedysplasia":0.3841,"tissuetumour":2.9045,"AveExpr":-4.6676,"F":9.3956,"P.Value":0.0004,"adj.P.Val":0.004},{"tissuedysplasia":0.1655,"tissuetumour":0.7525,"AveExpr":0.0282,"F":2.5026,"P.Value":0.0935,"adj.P.Val":0.2014},{"tissuedysplasia":0.3143,"tissuetumour":0.8027,"AveExpr":5.9601,"F":26.222,"P.Value":3.1741e-08,"adj.P.Val":3.1761e-06},{"tissuedysplasia":0.0211,"tissuetumour":0.6921,"AveExpr":3.417,"F":11.2027,"P.Value":0.0001,"adj.P.Val":0.0016},{"tissuedysplasia":0.078,"tissuetumour":0.6158,"AveExpr":2.8932,"F":7.2318,"P.Value":0.0019,"adj.P.Val":0.0126},{"tissuedysplasia":-1.3442,"tissuetumour":1.9847,"AveExpr":-9.6607,"F":5.0784,"P.Value":0.0104,"adj.P.Val":0.0425},{"tissuedysplasia":0.4127,"tissuetumour":1.4826,"AveExpr":-2.9034,"F":1.66,"P.Value":0.2018,"adj.P.Val":0.3391},{"tissuedysplasia":-0.1779,"tissuetumour":1.1768,"AveExpr":4.606,"F":20.3296,"P.Value":5.5838e-07,"adj.P.Val":0},{"tissuedysplasia":1.1599,"tissuetumour":2.5056,"AveExpr":-3.0122,"F":6.1079,"P.Value":0.0046,"adj.P.Val":0.0235},{"tissuedysplasia":0.1812,"tissuetumour":1.8085,"AveExpr":-3.625,"F":2.9557,"P.Value":0.0625,"adj.P.Val":0.1524},{"tissuedysplasia":0.2301,"tissuetumour":0.7985,"AveExpr":3.707,"F":11.6691,"P.Value":0.0001,"adj.P.Val":0.0012},{"tissuedysplasia":0.9372,"tissuetumour":2.0424,"AveExpr":1.4325,"F":22.4023,"P.Value":1.9504e-07,"adj.P.Val":0},{"tissuedysplasia":-0.3387,"tissuetumour":0.4494,"AveExpr":-1.8384,"F":1.1491,"P.Value":0.3263,"adj.P.Val":0.4704},{"tissuedysplasia":0.2575,"tissuetumour":0.5998,"AveExpr":3.1908,"F":6.4879,"P.Value":0.0034,"adj.P.Val":0.019},{"tissuedysplasia":0.1281,"tissuetumour":0.7264,"AveExpr":4.7133,"F":15.2148,"P.Value":9.4932e-06,"adj.P.Val":0.0002},{"tissuedysplasia":-0.034,"tissuetumour":0.375,"AveExpr":6.1872,"F":5.28,"P.Value":0.0088,"adj.P.Val":0.0379},{"tissuedysplasia":0.2328,"tissuetumour":1.0892,"AveExpr":1.705,"F":7.8969,"P.Value":0.0012,"adj.P.Val":0.0087},{"tissuedysplasia":0.0483,"tissuetumour":0.8768,"AveExpr":4.1218,"F":17.5909,"P.Value":2.4324e-06,"adj.P.Val":0.0001},{"tissuedysplasia":1.1448,"tissuetumour":2.2467,"AveExpr":-0.6083,"F":12.4292,"P.Value":0.0001,"adj.P.Val":0.0008},{"tissuedysplasia":1.3254,"tissuetumour":2.051,"AveExpr":1.6325,"F":21.5232,"P.Value":3.0284e-07,"adj.P.Val":0},{"tissuedysplasia":-0.0755,"tissuetumour":0.5261,"AveExpr":3.8872,"F":7.305,"P.Value":0.0018,"adj.P.Val":0.0121},{"tissuedysplasia":1.4081,"tissuetumour":2.4651,"AveExpr":-1.2186,"F":5.3488,"P.Value":0.0083,"adj.P.Val":0.0364},{"tissuedysplasia":-0.1226,"tissuetumour":0.9273,"AveExpr":0.6385,"F":5.8495,"P.Value":0.0056,"adj.P.Val":0.0273},{"tissuedysplasia":0.2743,"tissuetumour":0.5994,"AveExpr":3.1358,"F":9.0806,"P.Value":0.0005,"adj.P.Val":0.0046},{"tissuedysplasia":0.0461,"tissuetumour":0.5116,"AveExpr":3.7907,"F":9.0574,"P.Value":0.0005,"adj.P.Val":0.0047},{"tissuedysplasia":0.3177,"tissuetumour":1.1889,"AveExpr":0.766,"F":7.0465,"P.Value":0.0022,"adj.P.Val":0.0139},{"tissuedysplasia":0.9354,"tissuetumour":0.9341,"AveExpr":2.0202,"F":6.0174,"P.Value":0.0049,"adj.P.Val":0.0247},{"tissuedysplasia":0.587,"tissuetumour":1.0639,"AveExpr":1.5449,"F":12.1489,"P.Value":0.0001,"adj.P.Val":0.001},{"tissuedysplasia":-1.7138,"tissuetumour":2.3028,"AveExpr":2.2113,"F":27.7268,"P.Value":2.2961e-08,"adj.P.Val":2.4924e-06},{"tissuedysplasia":0.1993,"tissuetumour":0.41,"AveExpr":3.813,"F":4.0547,"P.Value":0.0242,"adj.P.Val":0.0783},{"tissuedysplasia":0.0954,"tissuetumour":0.0911,"AveExpr":4.5022,"F":0.5733,"P.Value":0.5678,"adj.P.Val":0.6824},{"tissuedysplasia":0.0743,"tissuetumour":0.5024,"AveExpr":5.5963,"F":8.4115,"P.Value":0.0008,"adj.P.Val":0.0066},{"tissuedysplasia":0.2305,"tissuetumour":0.4593,"AveExpr":2.8335,"F":4.8925,"P.Value":0.0121,"adj.P.Val":0.0472},{"tissuedysplasia":0.3373,"tissuetumour":0.5756,"AveExpr":3.2295,"F":7.9455,"P.Value":0.0011,"adj.P.Val":0.0085},{"tissuedysplasia":0.2817,"tissuetumour":0.8303,"AveExpr":2.2585,"F":3.809,"P.Value":0.0298,"adj.P.Val":0.0909},{"tissuedysplasia":1.2176,"tissuetumour":1.6905,"AveExpr":2.6633,"F":17.8243,"P.Value":2.1374e-06,"adj.P.Val":0.0001},{"tissuedysplasia":0.1622,"tissuetumour":0.2509,"AveExpr":4.8129,"F":2.9303,"P.Value":0.0639,"adj.P.Val":0.1549},{"tissuedysplasia":-0.4111,"tissuetumour":-1.199,"AveExpr":2.2219,"F":5.9008,"P.Value":0.0054,"adj.P.Val":0.0265},{"tissuedysplasia":-0.2226,"tissuetumour":-0.2465,"AveExpr":3.4132,"F":1.0053,"P.Value":0.3742,"adj.P.Val":0.5166},{"tissuedysplasia":0.2857,"tissuetumour":0.8944,"AveExpr":2.7048,"F":11.3973,"P.Value":0.0001,"adj.P.Val":0.0014},{"tissuedysplasia":0.0386,"tissuetumour":1.0826,"AveExpr":1.5814,"F":13.7184,"P.Value":0,"adj.P.Val":0.0005},{"tissuedysplasia":1.9801,"tissuetumour":1.9149,"AveExpr":2.2512,"F":7.6056,"P.Value":0.0016,"adj.P.Val":0.0107},{"tissuedysplasia":0.7538,"tissuetumour":2.4034,"AveExpr":-1.4227,"F":13.2471,"P.Value":0,"adj.P.Val":0.0006},{"tissuedysplasia":0.5407,"tissuetumour":0.9201,"AveExpr":-0.3364,"F":2.4805,"P.Value":0.0953,"adj.P.Val":0.2041},{"tissuedysplasia":0.2882,"tissuetumour":0.3426,"AveExpr":3.0913,"F":3.4453,"P.Value":0.0407,"adj.P.Val":0.113},{"tissuedysplasia":0.3542,"tissuetumour":0.9974,"AveExpr":1.4441,"F":5.2996,"P.Value":0.0087,"adj.P.Val":0.0375},{"tissuedysplasia":0.1355,"tissuetumour":-0.1119,"AveExpr":5.2716,"F":2.7025,"P.Value":0.0782,"adj.P.Val":0.1777},{"tissuedysplasia":-0.0247,"tissuetumour":-0.2037,"AveExpr":4.0393,"F":0.8278,"P.Value":0.4437,"adj.P.Val":0.5795},{"tissuedysplasia":0.0717,"tissuetumour":0.0931,"AveExpr":4.8558,"F":0.3167,"P.Value":0.7302,"adj.P.Val":0.8074},{"tissuedysplasia":-0.0364,"tissuetumour":-0.1322,"AveExpr":5.6491,"F":0.6537,"P.Value":0.5251,"adj.P.Val":0.6483},{"tissuedysplasia":0.1243,"tissuetumour":0.5404,"AveExpr":4.0417,"F":8.7207,"P.Value":0.0006,"adj.P.Val":0.0056},{"tissuedysplasia":1.0764,"tissuetumour":2.5102,"AveExpr":-8.1844,"F":2.4848,"P.Value":0.095,"adj.P.Val":0.2035},{"tissuedysplasia":-0.121,"tissuetumour":0.4859,"AveExpr":4.9351,"F":5.8639,"P.Value":0.0055,"adj.P.Val":0.027},{"tissuedysplasia":-0.1613,"tissuetumour":-0.056,"AveExpr":1.9476,"F":0.3398,"P.Value":0.7137,"adj.P.Val":0.795},{"tissuedysplasia":1.7375,"tissuetumour":3.6327,"AveExpr":-6.505,"F":5.3903,"P.Value":0.0081,"adj.P.Val":0.0355},{"tissuedysplasia":1.8484,"tissuetumour":2.7704,"AveExpr":-4.3741,"F":3.6698,"P.Value":0.0336,"adj.P.Val":0.0986},{"tissuedysplasia":-0.0588,"tissuetumour":0.4957,"AveExpr":7.0105,"F":11.1726,"P.Value":0.0001,"adj.P.Val":0.0016},{"tissuedysplasia":-0.1266,"tissuetumour":0.2917,"AveExpr":5.7252,"F":6.4615,"P.Value":0.0035,"adj.P.Val":0.0193},{"tissuedysplasia":0.3016,"tissuetumour":0.7575,"AveExpr":4.2014,"F":16.2813,"P.Value":5.0986e-06,"adj.P.Val":0.0001},{"tissuedysplasia":0.1201,"tissuetumour":0.8294,"AveExpr":0.6493,"F":3.2386,"P.Value":0.0487,"adj.P.Val":0.1284},{"tissuedysplasia":-0.0714,"tissuetumour":0.3211,"AveExpr":3.6283,"F":3.8873,"P.Value":0.0279,"adj.P.Val":0.0867},{"tissuedysplasia":-0.134,"tissuetumour":0.4109,"AveExpr":4.1088,"F":3.5476,"P.Value":0.0373,"adj.P.Val":0.1062},{"tissuedysplasia":1.335,"tissuetumour":1.8237,"AveExpr":-1.7948,"F":4.2894,"P.Value":0.0199,"adj.P.Val":0.0682},{"tissuedysplasia":0.0421,"tissuetumour":0.3294,"AveExpr":2.4029,"F":2.2165,"P.Value":0.121,"adj.P.Val":0.2397},{"tissuedysplasia":0.2399,"tissuetumour":1.6156,"AveExpr":-4.0812,"F":2.0232,"P.Value":0.1443,"adj.P.Val":0.2703},{"tissuedysplasia":0.0253,"tissuetumour":0.4489,"AveExpr":4.4322,"F":7.295,"P.Value":0.0018,"adj.P.Val":0.0122},{"tissuedysplasia":1.2462,"tissuetumour":2.071,"AveExpr":-1.6483,"F":6.203,"P.Value":0.0042,"adj.P.Val":0.0223},{"tissuedysplasia":-0.1301,"tissuetumour":-0.1072,"AveExpr":2.1985,"F":0.2726,"P.Value":0.7627,"adj.P.Val":0.8318},{"tissuedysplasia":0.7405,"tissuetumour":-0.2258,"AveExpr":-3.4918,"F":1.1962,"P.Value":0.312,"adj.P.Val":0.4565},{"tissuedysplasia":0.011,"tissuetumour":1.0196,"AveExpr":2.4271,"F":13.1243,"P.Value":0,"adj.P.Val":0.0006},{"tissuedysplasia":0.4912,"tissuetumour":1.4204,"AveExpr":-2.9279,"F":2.466,"P.Value":0.0966,"adj.P.Val":0.206},{"tissuedysplasia":0.5752,"tissuetumour":1.2267,"AveExpr":5.3539,"F":43.7236,"P.Value":3.4908e-11,"adj.P.Val":2.3835e-08},{"tissuedysplasia":0.2461,"tissuetumour":0.6444,"AveExpr":4.9384,"F":10.4614,"P.Value":0.0002,"adj.P.Val":0.0023},{"tissuedysplasia":-0.8418,"tissuetumour":-1.0397,"AveExpr":1.2132,"F":3.9198,"P.Value":0.0271,"adj.P.Val":0.085},{"tissuedysplasia":0.5572,"tissuetumour":1.6535,"AveExpr":-0.9974,"F":6.8609,"P.Value":0.0025,"adj.P.Val":0.0155},{"tissuedysplasia":0.3517,"tissuetumour":1.0226,"AveExpr":3.2894,"F":17.5564,"P.Value":2.4795e-06,"adj.P.Val":0.0001},{"tissuedysplasia":-0.0314,"tissuetumour":1.1921,"AveExpr":3.9527,"F":24.1899,"P.Value":8.1849e-08,"adj.P.Val":6.1852e-06},{"tissuedysplasia":0.1081,"tissuetumour":0.2988,"AveExpr":7.203,"F":6.2964,"P.Value":0.0039,"adj.P.Val":0.0211},{"tissuedysplasia":0.0429,"tissuetumour":0.4704,"AveExpr":4.3353,"F":4.7953,"P.Value":0.0131,"adj.P.Val":0.0502},{"tissuedysplasia":0.5929,"tissuetumour":1.6726,"AveExpr":0.7229,"F":16.0764,"P.Value":5.7374e-06,"adj.P.Val":0.0002},{"tissuedysplasia":0.3429,"tissuetumour":1.2574,"AveExpr":5.318,"F":22.9367,"P.Value":1.4991e-07,"adj.P.Val":9.4877e-06},{"tissuedysplasia":0.5844,"tissuetumour":0.9424,"AveExpr":5.2655,"F":13.4842,"P.Value":0,"adj.P.Val":0.0005},{"tissuedysplasia":0.0211,"tissuetumour":0.4435,"AveExpr":4.1418,"F":3.1065,"P.Value":0.0547,"adj.P.Val":0.1391},{"tissuedysplasia":1.1711,"tissuetumour":2.3248,"AveExpr":-0.574,"F":12.071,"P.Value":0.0001,"adj.P.Val":0.001},{"tissuedysplasia":-0.478,"tissuetumour":-0.0016,"AveExpr":4.8374,"F":4.4057,"P.Value":0.018,"adj.P.Val":0.0637},{"tissuedysplasia":-0.7151,"tissuetumour":-0.8258,"AveExpr":5.7115,"F":5.5979,"P.Value":0.0073,"adj.P.Val":0.033},{"tissuedysplasia":0.1871,"tissuetumour":0.5762,"AveExpr":3.3517,"F":6.0034,"P.Value":0.005,"adj.P.Val":0.0249},{"tissuedysplasia":0.64,"tissuetumour":1.3938,"AveExpr":-0.2284,"F":5.0711,"P.Value":0.0104,"adj.P.Val":0.0427},{"tissuedysplasia":2.3542,"tissuetumour":1.983,"AveExpr":-0.1987,"F":5.698,"P.Value":0.0063,"adj.P.Val":0.0298},{"tissuedysplasia":1.9344,"tissuetumour":4.3894,"AveExpr":-5.8981,"F":6.855,"P.Value":0.0026,"adj.P.Val":0.0155},{"tissuedysplasia":0.9292,"tissuetumour":2.2297,"AveExpr":-2.2473,"F":5.2035,"P.Value":0.0094,"adj.P.Val":0.0395},{"tissuedysplasia":-0.1941,"tissuetumour":0.0616,"AveExpr":6.7874,"F":2.7654,"P.Value":0.0739,"adj.P.Val":0.1712},{"tissuedysplasia":-0.0555,"tissuetumour":0.2918,"AveExpr":5.1347,"F":2.3401,"P.Value":0.1082,"adj.P.Val":0.2224},{"tissuedysplasia":1.5206,"tissuetumour":1.3958,"AveExpr":-0.644,"F":7.2172,"P.Value":0.0019,"adj.P.Val":0.0127},{"tissuedysplasia":1.2887,"tissuetumour":2.9693,"AveExpr":-1.288,"F":14.0781,"P.Value":0,"adj.P.Val":0.0004},{"tissuedysplasia":-0.0848,"tissuetumour":0.3087,"AveExpr":4.1181,"F":3.4349,"P.Value":0.0411,"adj.P.Val":0.1138},{"tissuedysplasia":0.3987,"tissuetumour":1.0818,"AveExpr":1.5509,"F":3.9752,"P.Value":0.0259,"adj.P.Val":0.0821},{"tissuedysplasia":-0.6328,"tissuetumour":-2.1182,"AveExpr":5.8363,"F":13.142,"P.Value":0,"adj.P.Val":0.0008},{"tissuedysplasia":0.8256,"tissuetumour":2.0525,"AveExpr":-0.4916,"F":9.2024,"P.Value":0.0005,"adj.P.Val":0.0043},{"tissuedysplasia":0.3186,"tissuetumour":0.6875,"AveExpr":3.642,"F":14.5692,"P.Value":0,"adj.P.Val":0.0003},{"tissuedysplasia":0.4346,"tissuetumour":1.0239,"AveExpr":4.8047,"F":25.6688,"P.Value":4.0913e-08,"adj.P.Val":3.7571e-06},{"tissuedysplasia":1.1712,"tissuetumour":3.3865,"AveExpr":-3.3593,"F":7.7434,"P.Value":0.0013,"adj.P.Val":0.0095},{"tissuedysplasia":0.12,"tissuetumour":0.912,"AveExpr":5.2454,"F":27.8336,"P.Value":1.5399e-08,"adj.P.Val":1.9138e-06},{"tissuedysplasia":-0.0406,"tissuetumour":0.222,"AveExpr":5.4745,"F":2.5927,"P.Value":0.0862,"adj.P.Val":0.1902},{"tissuedysplasia":0.2871,"tissuetumour":1.1274,"AveExpr":2.2624,"F":13.9794,"P.Value":0,"adj.P.Val":0.0004},{"tissuedysplasia":1.4249,"tissuetumour":2.4489,"AveExpr":-0.5408,"F":9.3903,"P.Value":0.0004,"adj.P.Val":0.004},{"tissuedysplasia":0.3957,"tissuetumour":1.5352,"AveExpr":2.6272,"F":15.4414,"P.Value":8.3063e-06,"adj.P.Val":0.0002},{"tissuedysplasia":0.0671,"tissuetumour":-0.018,"AveExpr":5.1623,"F":0.1547,"P.Value":0.8571,"adj.P.Val":0.8995},{"tissuedysplasia":-0.4459,"tissuetumour":-0.867,"AveExpr":6.5425,"F":4.3909,"P.Value":0.0194,"adj.P.Val":0.0672},{"tissuedysplasia":1.0358,"tissuetumour":1.1559,"AveExpr":-9.4588,"F":0.5922,"P.Value":0.5574,"adj.P.Val":0.674},{"tissuedysplasia":-0.0398,"tissuetumour":0.3978,"AveExpr":4.2632,"F":4.2265,"P.Value":0.0209,"adj.P.Val":0.0708},{"tissuedysplasia":0.1339,"tissuetumour":0.7855,"AveExpr":5.4986,"F":21.0297,"P.Value":3.8921e-07,"adj.P.Val":0},{"tissuedysplasia":-0.0743,"tissuetumour":0.356,"AveExpr":5.3594,"F":4.8195,"P.Value":0.0128,"adj.P.Val":0.0495},{"tissuedysplasia":0.0579,"tissuetumour":0.6865,"AveExpr":4.1775,"F":6.4701,"P.Value":0.0034,"adj.P.Val":0.0192},{"tissuedysplasia":1.7485,"tissuetumour":3.807,"AveExpr":0.3489,"F":43.6481,"P.Value":3.5802e-11,"adj.P.Val":2.4161e-08},{"tissuedysplasia":1.5344,"tissuetumour":2.53,"AveExpr":-6.5444,"F":2.1396,"P.Value":0.1298,"adj.P.Val":0.2515},{"tissuedysplasia":0.741,"tissuetumour":4.3243,"AveExpr":0.1773,"F":25.9576,"P.Value":3.5822e-08,"adj.P.Val":3.4478e-06},{"tissuedysplasia":0.3758,"tissuetumour":2.4286,"AveExpr":-1.114,"F":11.3912,"P.Value":0.0001,"adj.P.Val":0.0014},{"tissuedysplasia":0.5153,"tissuetumour":0.3033,"AveExpr":5.3214,"F":5.5995,"P.Value":0.0068,"adj.P.Val":0.0315},{"tissuedysplasia":0.3021,"tissuetumour":0.6853,"AveExpr":4.2637,"F":10.9859,"P.Value":0.0001,"adj.P.Val":0.0018},{"tissuedysplasia":0.2947,"tissuetumour":0.5353,"AveExpr":-0.9175,"F":0.5108,"P.Value":0.6035,"adj.P.Val":0.7116},{"tissuedysplasia":0.2261,"tissuetumour":0.2277,"AveExpr":2.8937,"F":1.3124,"P.Value":0.2795,"adj.P.Val":0.4241},{"tissuedysplasia":0.4517,"tissuetumour":1.3688,"AveExpr":2.4231,"F":16.7145,"P.Value":3.9804e-06,"adj.P.Val":0.0001},{"tissuedysplasia":0.9875,"tissuetumour":1.9637,"AveExpr":-0.5584,"F":7.6415,"P.Value":0.0014,"adj.P.Val":0.01},{"tissuedysplasia":0.8755,"tissuetumour":3.679,"AveExpr":-5.3992,"F":6.4256,"P.Value":0.0036,"adj.P.Val":0.0197},{"tissuedysplasia":0.2701,"tissuetumour":1.8017,"AveExpr":-2.4782,"F":3.6431,"P.Value":0.0343,"adj.P.Val":0.1002},{"tissuedysplasia":0.0098,"tissuetumour":-0.1556,"AveExpr":6.3178,"F":1.2523,"P.Value":0.2958,"adj.P.Val":0.4402},{"tissuedysplasia":0.301,"tissuetumour":0.3567,"AveExpr":2.3051,"F":1.3172,"P.Value":0.2782,"adj.P.Val":0.4227},{"tissuedysplasia":0.124,"tissuetumour":0.4564,"AveExpr":1.5008,"F":1.3885,"P.Value":0.2602,"adj.P.Val":0.4033},{"tissuedysplasia":0.0059,"tissuetumour":0.4673,"AveExpr":2.7936,"F":3.2413,"P.Value":0.0486,"adj.P.Val":0.1282},{"tissuedysplasia":0.2774,"tissuetumour":1.2903,"AveExpr":2.7353,"F":17.688,"P.Value":2.3048e-06,"adj.P.Val":0.0001},{"tissuedysplasia":0.0402,"tissuetumour":0.0557,"AveExpr":0.499,"F":0.011,"P.Value":0.9891,"adj.P.Val":0.9922},{"tissuedysplasia":0.379,"tissuetumour":1.1958,"AveExpr":1.9445,"F":13.1418,"P.Value":0,"adj.P.Val":0.0006},{"tissuedysplasia":0.0316,"tissuetumour":1.9741,"AveExpr":-4.6595,"F":2.8735,"P.Value":0.0672,"adj.P.Val":0.1601},{"tissuedysplasia":0.0362,"tissuetumour":1.3941,"AveExpr":-7.7135,"F":0.7799,"P.Value":0.4647,"adj.P.Val":0.5975},{"tissuedysplasia":0.2786,"tissuetumour":0.8985,"AveExpr":2.105,"F":8.1134,"P.Value":0.001,"adj.P.Val":0.0077},{"tissuedysplasia":0.2726,"tissuetumour":0.9962,"AveExpr":3.7401,"F":12.1353,"P.Value":0.0001,"adj.P.Val":0.001},{"tissuedysplasia":0.0991,"tissuetumour":1.3895,"AveExpr":3.1111,"F":16.2393,"P.Value":5.2231e-06,"adj.P.Val":0.0001},{"tissuedysplasia":-0.2007,"tissuetumour":0.1371,"AveExpr":5.527,"F":3.521,"P.Value":0.0382,"adj.P.Val":0.108},{"tissuedysplasia":0.0144,"tissuetumour":0.5737,"AveExpr":5.4148,"F":16.821,"P.Value":3.7469e-06,"adj.P.Val":0.0001},{"tissuedysplasia":0.0621,"tissuetumour":0.4983,"AveExpr":3.0233,"F":7.1384,"P.Value":0.0021,"adj.P.Val":0.0132},{"tissuedysplasia":0.7402,"tissuetumour":3.1171,"AveExpr":-5.4395,"F":3.2087,"P.Value":0.05,"adj.P.Val":0.1308},{"tissuedysplasia":0.0236,"tissuetumour":0.4918,"AveExpr":2.887,"F":3.6884,"P.Value":0.033,"adj.P.Val":0.0975},{"tissuedysplasia":0.35,"tissuetumour":1.0964,"AveExpr":2.0749,"F":9.6557,"P.Value":0.0003,"adj.P.Val":0.0034},{"tissuedysplasia":3.1247,"tissuetumour":1.0442,"AveExpr":-8.5965,"F":4.0909,"P.Value":0.0235,"adj.P.Val":0.0767},{"tissuedysplasia":0.4654,"tissuetumour":1.1186,"AveExpr":2.9302,"F":19.0461,"P.Value":1.0993e-06,"adj.P.Val":0},{"tissuedysplasia":0.7388,"tissuetumour":2.1341,"AveExpr":-1.3624,"F":8.1814,"P.Value":0.001,"adj.P.Val":0.0075},{"tissuedysplasia":0.4006,"tissuetumour":1.1446,"AveExpr":1.6952,"F":13.161,"P.Value":0,"adj.P.Val":0.0006},{"tissuedysplasia":0.2867,"tissuetumour":0.9367,"AveExpr":3.9929,"F":22.5915,"P.Value":1.7763e-07,"adj.P.Val":0},{"tissuedysplasia":0.0962,"tissuetumour":0.8628,"AveExpr":1.5278,"F":7.0627,"P.Value":0.0022,"adj.P.Val":0.0138},{"tissuedysplasia":1.048,"tissuetumour":1.2533,"AveExpr":3.9971,"F":9.6591,"P.Value":0.0003,"adj.P.Val":0.0034},{"tissuedysplasia":-0.1279,"tissuetumour":-0.0556,"AveExpr":4.9911,"F":0.5254,"P.Value":0.595,"adj.P.Val":0.7045},{"tissuedysplasia":1.0736,"tissuetumour":2.1348,"AveExpr":-1.9313,"F":7.1095,"P.Value":0.0021,"adj.P.Val":0.0134},{"tissuedysplasia":0.8593,"tissuetumour":1.9921,"AveExpr":-2.2297,"F":4.3805,"P.Value":0.0184,"adj.P.Val":0.0646},{"tissuedysplasia":0.4129,"tissuetumour":0.7273,"AveExpr":3.7394,"F":6.5834,"P.Value":0.0032,"adj.P.Val":0.018},{"tissuedysplasia":0.1467,"tissuetumour":0.3901,"AveExpr":4.8498,"F":3.2878,"P.Value":0.0467,"adj.P.Val":0.1245},{"tissuedysplasia":0.4983,"tissuetumour":1.0148,"AveExpr":-0.8911,"F":2.8198,"P.Value":0.0704,"adj.P.Val":0.1657},{"tissuedysplasia":0.0082,"tissuetumour":1.3548,"AveExpr":0.2522,"F":8.1846,"P.Value":0.001,"adj.P.Val":0.0074},{"tissuedysplasia":0.0113,"tissuetumour":0.6242,"AveExpr":1.2238,"F":3.3896,"P.Value":0.0427,"adj.P.Val":0.1169},{"tissuedysplasia":0.3522,"tissuetumour":0.7253,"AveExpr":3.9696,"F":9.2552,"P.Value":0.0004,"adj.P.Val":0.0042},{"tissuedysplasia":-0.0122,"tissuetumour":0.4992,"AveExpr":2.4237,"F":3.1002,"P.Value":0.055,"adj.P.Val":0.1396},{"tissuedysplasia":-0.1044,"tissuetumour":-0.1394,"AveExpr":2.669,"F":0.0645,"P.Value":0.9376,"adj.P.Val":0.9568},{"tissuedysplasia":-1.2919,"tissuetumour":-1.3742,"AveExpr":-1.5014,"F":4.6092,"P.Value":0.0152,"adj.P.Val":0.0562},{"tissuedysplasia":0.2044,"tissuetumour":0.6921,"AveExpr":3.5845,"F":12.6996,"P.Value":0,"adj.P.Val":0.0007},{"tissuedysplasia":0.2021,"tissuetumour":0.6068,"AveExpr":3.2461,"F":7.6081,"P.Value":0.0015,"adj.P.Val":0.0102},{"tissuedysplasia":1.0899,"tissuetumour":0.0738,"AveExpr":-3.7195,"F":0.9206,"P.Value":0.4058,"adj.P.Val":0.5454},{"tissuedysplasia":-0.4576,"tissuetumour":0.8957,"AveExpr":-2.6896,"F":1.8153,"P.Value":0.1748,"adj.P.Val":0.3081},{"tissuedysplasia":0.1759,"tissuetumour":0.6676,"AveExpr":2.6166,"F":6.9294,"P.Value":0.0024,"adj.P.Val":0.0149},{"tissuedysplasia":-0.307,"tissuetumour":-0.5519,"AveExpr":2.8118,"F":3.9652,"P.Value":0.0261,"adj.P.Val":0.0826},{"tissuedysplasia":0.0616,"tissuetumour":0.7509,"AveExpr":-0.593,"F":1.8626,"P.Value":0.1673,"adj.P.Val":0.299},{"tissuedysplasia":1.7731,"tissuetumour":0.9692,"AveExpr":-6.8698,"F":1.3223,"P.Value":0.2769,"adj.P.Val":0.4214},{"tissuedysplasia":0.7015,"tissuetumour":1.3327,"AveExpr":-6.3817,"F":0.7117,"P.Value":0.4964,"adj.P.Val":0.6237},{"tissuedysplasia":0.0464,"tissuetumour":1.5826,"AveExpr":-3.5703,"F":1.9267,"P.Value":0.1577,"adj.P.Val":0.2873},{"tissuedysplasia":0.1267,"tissuetumour":1.6034,"AveExpr":-1.8358,"F":3.7019,"P.Value":0.0327,"adj.P.Val":0.0967},{"tissuedysplasia":0.1975,"tissuetumour":0.4448,"AveExpr":5.5744,"F":9.3731,"P.Value":0.0004,"adj.P.Val":0.004},{"tissuedysplasia":0.493,"tissuetumour":1.3846,"AveExpr":3.9417,"F":26.1104,"P.Value":3.3401e-08,"adj.P.Val":3.3007e-06},{"tissuedysplasia":0.2547,"tissuetumour":0.6534,"AveExpr":4.439,"F":14.5099,"P.Value":0,"adj.P.Val":0.0003},{"tissuedysplasia":-0.0232,"tissuetumour":1.1758,"AveExpr":5.1692,"F":17.8028,"P.Value":2.1628e-06,"adj.P.Val":0.0001},{"tissuedysplasia":0.0172,"tissuetumour":0.3306,"AveExpr":3.6647,"F":3.5644,"P.Value":0.0368,"adj.P.Val":0.1051},{"tissuedysplasia":0.7425,"tissuetumour":2.3737,"AveExpr":-3.176,"F":5.5345,"P.Value":0.0072,"adj.P.Val":0.0327},{"tissuedysplasia":-0.2561,"tissuetumour":-0.3845,"AveExpr":1.6839,"F":1.1203,"P.Value":0.3353,"adj.P.Val":0.4796},{"tissuedysplasia":0.3818,"tissuetumour":1.8182,"AveExpr":0.7489,"F":9.8894,"P.Value":0.0003,"adj.P.Val":0.0031},{"tissuedysplasia":1.0044,"tissuetumour":1.22,"AveExpr":1.2789,"F":12.1317,"P.Value":0.0001,"adj.P.Val":0.001},{"tissuedysplasia":0.2408,"tissuetumour":1.3482,"AveExpr":3.6919,"F":16.6769,"P.Value":4.0664e-06,"adj.P.Val":0.0001},{"tissuedysplasia":0.1881,"tissuetumour":0.5392,"AveExpr":4.1553,"F":6.2001,"P.Value":0.0042,"adj.P.Val":0.0223},{"tissuedysplasia":0.2981,"tissuetumour":0.8332,"AveExpr":4.3573,"F":19.2803,"P.Value":9.7001e-07,"adj.P.Val":0},{"tissuedysplasia":-0.0592,"tissuetumour":0.2991,"AveExpr":2.5624,"F":1.8818,"P.Value":0.1644,"adj.P.Val":0.2954},{"tissuedysplasia":0.4327,"tissuetumour":1.4129,"AveExpr":-0.3267,"F":6.0694,"P.Value":0.0047,"adj.P.Val":0.024},{"tissuedysplasia":0.4089,"tissuetumour":0.7717,"AveExpr":5.3402,"F":17.8774,"P.Value":2.0755e-06,"adj.P.Val":0.0001},{"tissuedysplasia":0.0087,"tissuetumour":0.0951,"AveExpr":2.134,"F":0.0715,"P.Value":0.9311,"adj.P.Val":0.9519},{"tissuedysplasia":-0.4241,"tissuetumour":-0.0286,"AveExpr":2.3784,"F":3.3903,"P.Value":0.0427,"adj.P.Val":0.1169},{"tissuedysplasia":-0.0973,"tissuetumour":0.4316,"AveExpr":2.9338,"F":2.1058,"P.Value":0.1339,"adj.P.Val":0.2572},{"tissuedysplasia":0.4294,"tissuetumour":0.8311,"AveExpr":3.7924,"F":13.8095,"P.Value":0,"adj.P.Val":0.0004},{"tissuedysplasia":0.3269,"tissuetumour":0.3745,"AveExpr":5.6916,"F":10.0395,"P.Value":0.0003,"adj.P.Val":0.0028},{"tissuedysplasia":-0.0294,"tissuetumour":0.0565,"AveExpr":1.5809,"F":0.047,"P.Value":0.9541,"adj.P.Val":0.9685},{"tissuedysplasia":-0.0501,"tissuetumour":0.1886,"AveExpr":2.6781,"F":0.5815,"P.Value":0.5633,"adj.P.Val":0.6787},{"tissuedysplasia":0.1514,"tissuetumour":0.0696,"AveExpr":4.2442,"F":0.7718,"P.Value":0.4683,"adj.P.Val":0.5993},{"tissuedysplasia":0.2146,"tissuetumour":0.9149,"AveExpr":3.398,"F":11.2891,"P.Value":0.0001,"adj.P.Val":0.0015},{"tissuedysplasia":0.5748,"tissuetumour":0.7948,"AveExpr":6.6553,"F":9.0811,"P.Value":0.0005,"adj.P.Val":0.0049},{"tissuedysplasia":0.1398,"tissuetumour":-0.1998,"AveExpr":4.6125,"F":2.9764,"P.Value":0.0613,"adj.P.Val":0.1505},{"tissuedysplasia":0.267,"tissuetumour":0.3272,"AveExpr":4.8661,"F":3.5254,"P.Value":0.038,"adj.P.Val":0.1076},{"tissuedysplasia":1.4831,"tissuetumour":2.1375,"AveExpr":-1.2019,"F":3.5173,"P.Value":0.0383,"adj.P.Val":0.1082},{"tissuedysplasia":0.3868,"tissuetumour":0.4864,"AveExpr":2.5049,"F":2.3698,"P.Value":0.1053,"adj.P.Val":0.2184},{"tissuedysplasia":-0.1201,"tissuetumour":-1.0097,"AveExpr":7.6452,"F":16.5704,"P.Value":6.3587e-06,"adj.P.Val":0.0002},{"tissuedysplasia":0.2986,"tissuetumour":0.2315,"AveExpr":3.84,"F":2.5548,"P.Value":0.0892,"adj.P.Val":0.1947},{"tissuedysplasia":1.1055,"tissuetumour":0.7572,"AveExpr":3.1897,"F":9.2261,"P.Value":0.0005,"adj.P.Val":0.0043},{"tissuedysplasia":0.1343,"tissuetumour":0.6928,"AveExpr":0.0393,"F":2.1857,"P.Value":0.1244,"adj.P.Val":0.2443},{"tissuedysplasia":0.2078,"tissuetumour":-0.1104,"AveExpr":4.2591,"F":2.2501,"P.Value":0.1174,"adj.P.Val":0.2351},{"tissuedysplasia":0.387,"tissuetumour":0.0034,"AveExpr":6.306,"F":4.2567,"P.Value":0.0204,"adj.P.Val":0.0695},{"tissuedysplasia":-0.0701,"tissuetumour":-0.4448,"AveExpr":5.7004,"F":12.1846,"P.Value":0.0001,"adj.P.Val":0.001},{"tissuedysplasia":0.3112,"tissuetumour":0.4633,"AveExpr":2.3346,"F":2.1458,"P.Value":0.1291,"adj.P.Val":0.2505},{"tissuedysplasia":0.4565,"tissuetumour":0.121,"AveExpr":2.8806,"F":3.7906,"P.Value":0.0303,"adj.P.Val":0.0919},{"tissuedysplasia":0.1512,"tissuetumour":0.5744,"AveExpr":4.7938,"F":8.96,"P.Value":0.0005,"adj.P.Val":0.0049},{"tissuedysplasia":0.1713,"tissuetumour":0.472,"AveExpr":3.2887,"F":2.6846,"P.Value":0.0794,"adj.P.Val":0.1798},{"tissuedysplasia":0.1962,"tissuetumour":0.1936,"AveExpr":4.7092,"F":1.5156,"P.Value":0.2309,"adj.P.Val":0.3721},{"tissuedysplasia":0.7015,"tissuetumour":0.6043,"AveExpr":2.8043,"F":5.1345,"P.Value":0.0099,"adj.P.Val":0.0411},{"tissuedysplasia":-0.0715,"tissuetumour":-0.4759,"AveExpr":6.8932,"F":6.3948,"P.Value":0.0036,"adj.P.Val":0.02},{"tissuedysplasia":1.341,"tissuetumour":2.4829,"AveExpr":-0.0704,"F":20.0888,"P.Value":6.3304e-07,"adj.P.Val":0},{"tissuedysplasia":-1.5371,"tissuetumour":-2.9245,"AveExpr":2.719,"F":38.6728,"P.Value":2.0276e-10,"adj.P.Val":8.4898e-08},{"tissuedysplasia":0.0776,"tissuetumour":-0.0268,"AveExpr":3.9956,"F":0.2071,"P.Value":0.8137,"adj.P.Val":0.8694},{"tissuedysplasia":0.8115,"tissuetumour":1.7283,"AveExpr":3.4004,"F":29.0253,"P.Value":9.1558e-09,"adj.P.Val":1.3153e-06},{"tissuedysplasia":-2.2686,"tissuetumour":-2.4501,"AveExpr":2.0947,"F":30.6449,"P.Value":4.6037e-09,"adj.P.Val":8.0349e-07},{"tissuedysplasia":0.3927,"tissuetumour":0.2552,"AveExpr":6.5281,"F":6.6077,"P.Value":0.0031,"adj.P.Val":0.0178},{"tissuedysplasia":0.4434,"tissuetumour":0.575,"AveExpr":1.7002,"F":2.0583,"P.Value":0.1398,"adj.P.Val":0.2646},{"tissuedysplasia":0.2385,"tissuetumour":0.5161,"AveExpr":6.1349,"F":9.2656,"P.Value":0.0004,"adj.P.Val":0.0042},{"tissuedysplasia":1.1863,"tissuetumour":3.119,"AveExpr":-5.6801,"F":4.7329,"P.Value":0.0137,"adj.P.Val":0.0522},{"tissuedysplasia":0.4059,"tissuetumour":1.1007,"AveExpr":1.8697,"F":7.6164,"P.Value":0.0014,"adj.P.Val":0.0102},{"tissuedysplasia":0.7601,"tissuetumour":1.0514,"AveExpr":2.5387,"F":6.6387,"P.Value":0.003,"adj.P.Val":0.0175},{"tissuedysplasia":0.3313,"tissuetumour":0.1478,"AveExpr":5.5237,"F":6.7681,"P.Value":0.0027,"adj.P.Val":0.0163},{"tissuedysplasia":0.3482,"tissuetumour":0.5295,"AveExpr":5.1877,"F":3.8052,"P.Value":0.0299,"adj.P.Val":0.0911},{"tissuedysplasia":0.5648,"tissuetumour":1.432,"AveExpr":6.0462,"F":13.0089,"P.Value":0,"adj.P.Val":0.0008},{"tissuedysplasia":1.4148,"tissuetumour":1.7572,"AveExpr":5.4049,"F":24.1676,"P.Value":8.7751e-08,"adj.P.Val":6.496e-06},{"tissuedysplasia":0.0772,"tissuetumour":0.0857,"AveExpr":2.4669,"F":0.0945,"P.Value":0.91,"adj.P.Val":0.9372},{"tissuedysplasia":0.3255,"tissuetumour":0.1677,"AveExpr":5.3849,"F":3.4976,"P.Value":0.0389,"adj.P.Val":0.1095},{"tissuedysplasia":0.2841,"tissuetumour":-0.1443,"AveExpr":-0.0524,"F":0.6876,"P.Value":0.5081,"adj.P.Val":0.6336},{"tissuedysplasia":3.4272,"tissuetumour":6.2675,"AveExpr":-2.5923,"F":19.9461,"P.Value":6.8214e-07,"adj.P.Val":0},{"tissuedysplasia":1.9403,"tissuetumour":2.2388,"AveExpr":-0.2316,"F":9.9261,"P.Value":0.0003,"adj.P.Val":0.003},{"tissuedysplasia":0.0414,"tissuetumour":-0.2776,"AveExpr":7.0379,"F":3.0486,"P.Value":0.0576,"adj.P.Val":0.144},{"tissuedysplasia":1.3284,"tissuetumour":2.308,"AveExpr":-1.3854,"F":5.2221,"P.Value":0.0092,"adj.P.Val":0.0391},{"tissuedysplasia":0.9,"tissuetumour":1.4307,"AveExpr":0.1639,"F":6.7883,"P.Value":0.0027,"adj.P.Val":0.0161},{"tissuedysplasia":0.5805,"tissuetumour":1.2842,"AveExpr":2.7768,"F":13.8072,"P.Value":0,"adj.P.Val":0.0004},{"tissuedysplasia":2.3232,"tissuetumour":2.4884,"AveExpr":-1.4442,"F":8.8147,"P.Value":0.0006,"adj.P.Val":0.0053},{"tissuedysplasia":3.2498,"tissuetumour":3.3979,"AveExpr":-4.2536,"F":7.9793,"P.Value":0.0011,"adj.P.Val":0.0083},{"tissuedysplasia":0.3725,"tissuetumour":0.8704,"AveExpr":3.1329,"F":8.3392,"P.Value":0.0008,"adj.P.Val":0.0069},{"tissuedysplasia":0.7437,"tissuetumour":1.4691,"AveExpr":3.7782,"F":27.1319,"P.Value":2.1038e-08,"adj.P.Val":2.3525e-06},{"tissuedysplasia":0.6736,"tissuetumour":2.4302,"AveExpr":-2.5768,"F":8.2003,"P.Value":0.0009,"adj.P.Val":0.0074},{"tissuedysplasia":0.3588,"tissuetumour":1.3664,"AveExpr":-0.9586,"F":4.7959,"P.Value":0.0131,"adj.P.Val":0.0502},{"tissuedysplasia":0.1838,"tissuetumour":0.4837,"AveExpr":0.3926,"F":0.7403,"P.Value":0.4828,"adj.P.Val":0.6118},{"tissuedysplasia":0.183,"tissuetumour":0.1435,"AveExpr":4.6183,"F":1.5186,"P.Value":0.2303,"adj.P.Val":0.3714},{"tissuedysplasia":1.3733,"tissuetumour":2.4202,"AveExpr":-2.5449,"F":5.483,"P.Value":0.0075,"adj.P.Val":0.0337},{"tissuedysplasia":-0.2012,"tissuetumour":-0.0248,"AveExpr":3.0154,"F":0.5277,"P.Value":0.5936,"adj.P.Val":0.7034},{"tissuedysplasia":0.4679,"tissuetumour":2.532,"AveExpr":-4.9178,"F":3.6084,"P.Value":0.0354,"adj.P.Val":0.1023},{"tissuedysplasia":0.466,"tissuetumour":0.3932,"AveExpr":2.9465,"F":5.4979,"P.Value":0.0074,"adj.P.Val":0.0334},{"tissuedysplasia":-0.1679,"tissuetumour":-0.5599,"AveExpr":6.3036,"F":4.6249,"P.Value":0.015,"adj.P.Val":0.0557},{"tissuedysplasia":0.7555,"tissuetumour":0.5097,"AveExpr":3.4152,"F":5.4194,"P.Value":0.0079,"adj.P.Val":0.0349},{"tissuedysplasia":0.6342,"tissuetumour":0.9096,"AveExpr":2.2292,"F":6.6297,"P.Value":0.003,"adj.P.Val":0.0175},{"tissuedysplasia":0.2537,"tissuetumour":0.2724,"AveExpr":4.6449,"F":2.0686,"P.Value":0.1385,"adj.P.Val":0.263},{"tissuedysplasia":-0.0524,"tissuetumour":-0.105,"AveExpr":2.476,"F":0.1643,"P.Value":0.849,"adj.P.Val":0.8941},{"tissuedysplasia":-1.2072,"tissuetumour":-1.41,"AveExpr":3.8781,"F":20.7996,"P.Value":4.3793e-07,"adj.P.Val":0},{"tissuedysplasia":0.3114,"tissuetumour":0.5924,"AveExpr":3.8991,"F":12.8163,"P.Value":0,"adj.P.Val":0.0007},{"tissuedysplasia":0.1856,"tissuetumour":0.0921,"AveExpr":4.3078,"F":1.0555,"P.Value":0.3567,"adj.P.Val":0.5001},{"tissuedysplasia":0.8838,"tissuetumour":1.8987,"AveExpr":1.4181,"F":14.9972,"P.Value":0,"adj.P.Val":0.0002},{"tissuedysplasia":0.3689,"tissuetumour":0.6195,"AveExpr":4.1159,"F":11.9145,"P.Value":0.0001,"adj.P.Val":0.0011},{"tissuedysplasia":0.136,"tissuetumour":0.2617,"AveExpr":5.9347,"F":1.3141,"P.Value":0.2791,"adj.P.Val":0.4236},{"tissuedysplasia":0.3668,"tissuetumour":1.0352,"AveExpr":-1.0067,"F":1.8305,"P.Value":0.1723,"adj.P.Val":0.3052},{"tissuedysplasia":-0.2532,"tissuetumour":-0.729,"AveExpr":3.3008,"F":3.7079,"P.Value":0.0325,"adj.P.Val":0.0964},{"tissuedysplasia":0.7146,"tissuetumour":0.9106,"AveExpr":2.567,"F":10.3814,"P.Value":0.0002,"adj.P.Val":0.0024},{"tissuedysplasia":2.1892,"tissuetumour":3.5099,"AveExpr":-4.6208,"F":6.799,"P.Value":0.0027,"adj.P.Val":0.016},{"tissuedysplasia":0.1092,"tissuetumour":0.4399,"AveExpr":3.6086,"F":4.6918,"P.Value":0.0142,"adj.P.Val":0.0535},{"tissuedysplasia":0.3704,"tissuetumour":0.4675,"AveExpr":5.0543,"F":10.4849,"P.Value":0.0002,"adj.P.Val":0.0023},{"tissuedysplasia":0.3611,"tissuetumour":0.8012,"AveExpr":4.6435,"F":12.328,"P.Value":0.0001,"adj.P.Val":0.0009},{"tissuedysplasia":1.2045,"tissuetumour":2.7978,"AveExpr":-4.2763,"F":4.0782,"P.Value":0.0237,"adj.P.Val":0.0772},{"tissuedysplasia":0.1877,"tissuetumour":0.3214,"AveExpr":7.9698,"F":1.2445,"P.Value":0.2999,"adj.P.Val":0.4443},{"tissuedysplasia":0.5544,"tissuetumour":0.4604,"AveExpr":0.8698,"F":1.8246,"P.Value":0.1733,"adj.P.Val":0.3063},{"tissuedysplasia":0.1425,"tissuetumour":0.7668,"AveExpr":5.5467,"F":22.0306,"P.Value":2.3467e-07,"adj.P.Val":0},{"tissuedysplasia":0.1377,"tissuetumour":0.1591,"AveExpr":3.8641,"F":0.6504,"P.Value":0.5268,"adj.P.Val":0.6497},{"tissuedysplasia":0.1048,"tissuetumour":0.5251,"AveExpr":3.6536,"F":8.4859,"P.Value":0.0008,"adj.P.Val":0.0064},{"tissuedysplasia":-1.1344,"tissuetumour":-2.4416,"AveExpr":4.7854,"F":63.5785,"P.Value":1.0485e-13,"adj.P.Val":6.7611e-10},{"tissuedysplasia":-0.0325,"tissuetumour":0.0194,"AveExpr":6.7394,"F":0.2264,"P.Value":0.7983,"adj.P.Val":0.8584},{"tissuedysplasia":0.3157,"tissuetumour":0.7511,"AveExpr":6.6981,"F":19.7565,"P.Value":7.5365e-07,"adj.P.Val":0},{"tissuedysplasia":0.5234,"tissuetumour":1.3614,"AveExpr":3.792,"F":21.7911,"P.Value":2.6459e-07,"adj.P.Val":0},{"tissuedysplasia":0.7744,"tissuetumour":1.4682,"AveExpr":1.4296,"F":13.6445,"P.Value":0,"adj.P.Val":0.0005},{"tissuedysplasia":0.8631,"tissuetumour":1.2219,"AveExpr":6.5635,"F":21.087,"P.Value":4.4549e-07,"adj.P.Val":0},{"tissuedysplasia":0.2788,"tissuetumour":1.0679,"AveExpr":4.1742,"F":27.3609,"P.Value":1.8992e-08,"adj.P.Val":2.2015e-06},{"tissuedysplasia":0.1279,"tissuetumour":-0.428,"AveExpr":5.9741,"F":8.1667,"P.Value":0.001,"adj.P.Val":0.0075},{"tissuedysplasia":0.7125,"tissuetumour":1.5542,"AveExpr":-0.8867,"F":8.0026,"P.Value":0.0011,"adj.P.Val":0.0082},{"tissuedysplasia":1.3213,"tissuetumour":2.6239,"AveExpr":-3.0082,"F":6.2646,"P.Value":0.004,"adj.P.Val":0.0215},{"tissuedysplasia":0.4331,"tissuetumour":0.7254,"AveExpr":2.9424,"F":11.0468,"P.Value":0.0001,"adj.P.Val":0.0017},{"tissuedysplasia":0.5017,"tissuetumour":1.1057,"AveExpr":-0.4844,"F":3.3795,"P.Value":0.0431,"adj.P.Val":0.1176},{"tissuedysplasia":0.0343,"tissuetumour":0.9055,"AveExpr":3.2731,"F":9.7699,"P.Value":0.0003,"adj.P.Val":0.0033},{"tissuedysplasia":0.0714,"tissuetumour":0.5177,"AveExpr":1.3534,"F":2.3036,"P.Value":0.1118,"adj.P.Val":0.2276},{"tissuedysplasia":0.6733,"tissuetumour":1.8821,"AveExpr":-5.4857,"F":1.7164,"P.Value":0.1915,"adj.P.Val":0.3278},{"tissuedysplasia":0.4202,"tissuetumour":1.5204,"AveExpr":2.9752,"F":14.7103,"P.Value":0,"adj.P.Val":0.0003},{"tissuedysplasia":0.4175,"tissuetumour":1.1761,"AveExpr":-0.0688,"F":6.2799,"P.Value":0.004,"adj.P.Val":0.0213},{"tissuedysplasia":0.5964,"tissuetumour":2.3949,"AveExpr":-4.4275,"F":2.8637,"P.Value":0.0677,"adj.P.Val":0.1612},{"tissuedysplasia":0.272,"tissuetumour":0.4806,"AveExpr":5.9665,"F":11.9057,"P.Value":0.0001,"adj.P.Val":0.0011},{"tissuedysplasia":0.2933,"tissuetumour":0.3038,"AveExpr":4.304,"F":3.4456,"P.Value":0.0407,"adj.P.Val":0.113},{"tissuedysplasia":0.37,"tissuetumour":1.4237,"AveExpr":3.9993,"F":30.9639,"P.Value":4.0308e-09,"adj.P.Val":7.2426e-07},{"tissuedysplasia":-0.4767,"tissuetumour":-0.6022,"AveExpr":3.7929,"F":5.2046,"P.Value":0.0094,"adj.P.Val":0.0395},{"tissuedysplasia":0.456,"tissuetumour":0.3591,"AveExpr":2.9266,"F":3.269,"P.Value":0.0475,"adj.P.Val":0.126},{"tissuedysplasia":0.69,"tissuetumour":2.3982,"AveExpr":5.9592,"F":15.9356,"P.Value":0,"adj.P.Val":0.0002},{"tissuedysplasia":0.4345,"tissuetumour":1.0083,"AveExpr":2.5867,"F":13.1653,"P.Value":0,"adj.P.Val":0.0006},{"tissuedysplasia":0.6012,"tissuetumour":1.8293,"AveExpr":-2.7973,"F":4.1143,"P.Value":0.023,"adj.P.Val":0.0756},{"tissuedysplasia":-0.1703,"tissuetumour":-0.6101,"AveExpr":4.6863,"F":5.5693,"P.Value":0.007,"adj.P.Val":0.0321},{"tissuedysplasia":0.6461,"tissuetumour":1.8702,"AveExpr":3.2444,"F":33.9194,"P.Value":1.2205e-09,"adj.P.Val":2.8795e-07},{"tissuedysplasia":0.7254,"tissuetumour":1.4723,"AveExpr":2.5746,"F":27.5931,"P.Value":1.7129e-08,"adj.P.Val":2.0712e-06},{"tissuedysplasia":0.2866,"tissuetumour":0.8191,"AveExpr":3.8938,"F":16.0307,"P.Value":5.8911e-06,"adj.P.Val":0.0002},{"tissuedysplasia":0.3911,"tissuetumour":0.3557,"AveExpr":1.4158,"F":1.1119,"P.Value":0.338,"adj.P.Val":0.4822},{"tissuedysplasia":1.0003,"tissuetumour":2.9219,"AveExpr":-4.7614,"F":5.0719,"P.Value":0.0104,"adj.P.Val":0.0427},{"tissuedysplasia":0.6906,"tissuetumour":0.4036,"AveExpr":3.8939,"F":5.7021,"P.Value":0.0063,"adj.P.Val":0.0297},{"tissuedysplasia":0.0689,"tissuetumour":2.4988,"AveExpr":-5.8268,"F":5.2447,"P.Value":0.0091,"adj.P.Val":0.0386},{"tissuedysplasia":0.2268,"tissuetumour":0.2727,"AveExpr":5.7176,"F":3.8199,"P.Value":0.0295,"adj.P.Val":0.0903},{"tissuedysplasia":0.3324,"tissuetumour":0.3429,"AveExpr":3.3271,"F":3.6654,"P.Value":0.0337,"adj.P.Val":0.0988},{"tissuedysplasia":0.1692,"tissuetumour":-0.3298,"AveExpr":4.892,"F":5.1004,"P.Value":0.0102,"adj.P.Val":0.042},{"tissuedysplasia":0.2015,"tissuetumour":-0.0684,"AveExpr":4.4499,"F":1.609,"P.Value":0.2116,"adj.P.Val":0.3506},{"tissuedysplasia":0.6143,"tissuetumour":1.2205,"AveExpr":2.584,"F":14.1685,"P.Value":0,"adj.P.Val":0.0004},{"tissuedysplasia":-0.0128,"tissuetumour":-0.1154,"AveExpr":3.4001,"F":0.2385,"P.Value":0.7888,"adj.P.Val":0.851},{"tissuedysplasia":0.7303,"tissuetumour":2.9475,"AveExpr":0.8875,"F":22.9525,"P.Value":1.4876e-07,"adj.P.Val":9.4561e-06},{"tissuedysplasia":0.7126,"tissuetumour":0.8396,"AveExpr":2.0654,"F":4.5252,"P.Value":0.0163,"adj.P.Val":0.0591},{"tissuedysplasia":-0.5549,"tissuetumour":0.1664,"AveExpr":1.2758,"F":3.4459,"P.Value":0.0407,"adj.P.Val":0.1129},{"tissuedysplasia":0.1143,"tissuetumour":0.0447,"AveExpr":2.4752,"F":0.1177,"P.Value":0.8892,"adj.P.Val":0.9223},{"tissuedysplasia":0.3016,"tissuetumour":-0.1217,"AveExpr":3.6359,"F":3.8248,"P.Value":0.0294,"adj.P.Val":0.09},{"tissuedysplasia":1.8968,"tissuetumour":2.9468,"AveExpr":-0.1118,"F":15.605,"P.Value":7.5467e-06,"adj.P.Val":0.0002},{"tissuedysplasia":0.5091,"tissuetumour":0.5786,"AveExpr":3.4041,"F":2.6907,"P.Value":0.079,"adj.P.Val":0.1791},{"tissuedysplasia":0.5637,"tissuetumour":2.273,"AveExpr":1.8674,"F":35.3494,"P.Value":7.0027e-10,"adj.P.Val":1.9569e-07},{"tissuedysplasia":0.3969,"tissuetumour":3.4145,"AveExpr":-7.8167,"F":3.3594,"P.Value":0.0439,"adj.P.Val":0.1191},{"tissuedysplasia":1.4012,"tissuetumour":2.0482,"AveExpr":0.8938,"F":6.4679,"P.Value":0.0034,"adj.P.Val":0.0192},{"tissuedysplasia":0.4067,"tissuetumour":0.1758,"AveExpr":6.4337,"F":5.7311,"P.Value":0.0061,"adj.P.Val":0.0292},{"tissuedysplasia":0.2774,"tissuetumour":0.4253,"AveExpr":4.3957,"F":7.3046,"P.Value":0.0018,"adj.P.Val":0.0121},{"tissuedysplasia":0.8092,"tissuetumour":0.6131,"AveExpr":1.8413,"F":5.3005,"P.Value":0.0087,"adj.P.Val":0.0374},{"tissuedysplasia":0.652,"tissuetumour":1.3903,"AveExpr":3.4831,"F":21.5825,"P.Value":2.939e-07,"adj.P.Val":0},{"tissuedysplasia":0.5853,"tissuetumour":0.7959,"AveExpr":4.7823,"F":15.169,"P.Value":9.7537e-06,"adj.P.Val":0.0002},{"tissuedysplasia":0.4843,"tissuetumour":0.5498,"AveExpr":4.5732,"F":7.8467,"P.Value":0.0012,"adj.P.Val":0.009},{"tissuedysplasia":0.3176,"tissuetumour":0.1307,"AveExpr":3.4243,"F":1.866,"P.Value":0.1668,"adj.P.Val":0.2983},{"tissuedysplasia":0.4114,"tissuetumour":0.0826,"AveExpr":5.4782,"F":6.9102,"P.Value":0.0025,"adj.P.Val":0.015},{"tissuedysplasia":-0.1461,"tissuetumour":0.5056,"AveExpr":3.7073,"F":10.8205,"P.Value":0.0002,"adj.P.Val":0.0019},{"tissuedysplasia":0.3252,"tissuetumour":0.7387,"AveExpr":4.5588,"F":14.9224,"P.Value":0,"adj.P.Val":0.0003},{"tissuedysplasia":0.1964,"tissuetumour":0.0665,"AveExpr":5.5248,"F":1.8938,"P.Value":0.1626,"adj.P.Val":0.2931},{"tissuedysplasia":0.4142,"tissuetumour":0.3031,"AveExpr":2.2205,"F":3.6921,"P.Value":0.0329,"adj.P.Val":0.0973},{"tissuedysplasia":0.3023,"tissuetumour":0.3668,"AveExpr":4.296,"F":3.8032,"P.Value":0.03,"adj.P.Val":0.0912},{"tissuedysplasia":0.1435,"tissuetumour":0.3363,"AveExpr":3.52,"F":1.8416,"P.Value":0.1706,"adj.P.Val":0.3031},{"tissuedysplasia":0.4707,"tissuetumour":2.4325,"AveExpr":6.2337,"F":59.3938,"P.Value":1.0443e-12,"adj.P.Val":2.8861e-09},{"tissuedysplasia":0.1446,"tissuetumour":0.5733,"AveExpr":4.3202,"F":8.0065,"P.Value":0.0011,"adj.P.Val":0.0082},{"tissuedysplasia":2.4348,"tissuetumour":3.3946,"AveExpr":-7.4095,"F":4.1161,"P.Value":0.023,"adj.P.Val":0.0755},{"tissuedysplasia":0.4702,"tissuetumour":3.0595,"AveExpr":-3.9684,"F":6.5844,"P.Value":0.0032,"adj.P.Val":0.018},{"tissuedysplasia":0.1815,"tissuetumour":0.658,"AveExpr":4.3461,"F":12.3612,"P.Value":0.0001,"adj.P.Val":0.0009},{"tissuedysplasia":0.3644,"tissuetumour":1.0584,"AveExpr":5.4618,"F":36.38,"P.Value":4.7322e-10,"adj.P.Val":1.5507e-07},{"tissuedysplasia":0.2984,"tissuetumour":0.8648,"AveExpr":3.7518,"F":20.5371,"P.Value":5.0141e-07,"adj.P.Val":0},{"tissuedysplasia":0.5255,"tissuetumour":2.3823,"AveExpr":-2.6462,"F":5.211,"P.Value":0.0093,"adj.P.Val":0.0393},{"tissuedysplasia":0.4823,"tissuetumour":1.0548,"AveExpr":3.7388,"F":14.9226,"P.Value":0,"adj.P.Val":0.0003},{"tissuedysplasia":0.1084,"tissuetumour":0.0225,"AveExpr":6.5893,"F":0.9882,"P.Value":0.3804,"adj.P.Val":0.5223},{"tissuedysplasia":0.5723,"tissuetumour":1.6484,"AveExpr":4.2163,"F":47.1945,"P.Value":1.1251e-11,"adj.P.Val":1.258e-08},{"tissuedysplasia":-0.3317,"tissuetumour":-0.6247,"AveExpr":2.2439,"F":3.762,"P.Value":0.031,"adj.P.Val":0.0933},{"tissuedysplasia":-0.3294,"tissuetumour":-0.4874,"AveExpr":1.7807,"F":1.3133,"P.Value":0.2793,"adj.P.Val":0.4238},{"tissuedysplasia":2.7141,"tissuetumour":3.734,"AveExpr":-2.7936,"F":9.2742,"P.Value":0.0004,"adj.P.Val":0.0042},{"tissuedysplasia":0.2018,"tissuetumour":0.1302,"AveExpr":4.1778,"F":1.1803,"P.Value":0.3167,"adj.P.Val":0.4609},{"tissuedysplasia":0.3975,"tissuetumour":0.9114,"AveExpr":2.6438,"F":10.4325,"P.Value":0.0002,"adj.P.Val":0.0023},{"tissuedysplasia":-0.5839,"tissuetumour":-1.0068,"AveExpr":3.6089,"F":6.7489,"P.Value":0.0028,"adj.P.Val":0.0164},{"tissuedysplasia":0.0415,"tissuetumour":-0.1469,"AveExpr":5.1462,"F":0.9352,"P.Value":0.4002,"adj.P.Val":0.5405},{"tissuedysplasia":-0.1552,"tissuetumour":-0.5105,"AveExpr":3.6615,"F":4.0155,"P.Value":0.025,"adj.P.Val":0.0803},{"tissuedysplasia":0.1984,"tissuetumour":0.2394,"AveExpr":5.6465,"F":2.7819,"P.Value":0.0728,"adj.P.Val":0.1696},{"tissuedysplasia":-0.7051,"tissuetumour":1.1454,"AveExpr":3.3078,"F":17.0222,"P.Value":3.344e-06,"adj.P.Val":0.0001},{"tissuedysplasia":-0.3984,"tissuetumour":-0.6389,"AveExpr":5.3579,"F":9.1248,"P.Value":0.0005,"adj.P.Val":0.0045},{"tissuedysplasia":0.3365,"tissuetumour":0.4708,"AveExpr":5.3228,"F":7.0591,"P.Value":0.0022,"adj.P.Val":0.0138},{"tissuedysplasia":0.2946,"tissuetumour":0.4305,"AveExpr":4.6236,"F":3.2371,"P.Value":0.0488,"adj.P.Val":0.1285},{"tissuedysplasia":0.2814,"tissuetumour":0.2659,"AveExpr":5.6047,"F":1.3358,"P.Value":0.2734,"adj.P.Val":0.4177},{"tissuedysplasia":0.3539,"tissuetumour":0.1622,"AveExpr":2.8685,"F":1.4552,"P.Value":0.2444,"adj.P.Val":0.3869},{"tissuedysplasia":0.2341,"tissuetumour":0.1469,"AveExpr":2.5908,"F":0.5285,"P.Value":0.5932,"adj.P.Val":0.7031},{"tissuedysplasia":0.1401,"tissuetumour":-0.2603,"AveExpr":2.8527,"F":2.2319,"P.Value":0.1193,"adj.P.Val":0.2376},{"tissuedysplasia":0.7332,"tissuetumour":1.223,"AveExpr":3.9344,"F":8.111,"P.Value":0.001,"adj.P.Val":0.0077},{"tissuedysplasia":0.1832,"tissuetumour":0.1949,"AveExpr":4.217,"F":1.3033,"P.Value":0.2819,"adj.P.Val":0.4266},{"tissuedysplasia":0.2277,"tissuetumour":0.0288,"AveExpr":5.2152,"F":2.734,"P.Value":0.076,"adj.P.Val":0.1743},{"tissuedysplasia":-0.6463,"tissuetumour":0.2346,"AveExpr":1.154,"F":3.0681,"P.Value":0.0566,"adj.P.Val":0.1423},{"tissuedysplasia":-0.7224,"tissuetumour":-1.0472,"AveExpr":5.1516,"F":11.9595,"P.Value":0.0001,"adj.P.Val":0.0011},{"tissuedysplasia":0.3536,"tissuetumour":0.5742,"AveExpr":3.2928,"F":6.2763,"P.Value":0.004,"adj.P.Val":0.0214},{"tissuedysplasia":1.2531,"tissuetumour":1.4279,"AveExpr":-3.8476,"F":1.6752,"P.Value":0.199,"adj.P.Val":0.3359},{"tissuedysplasia":0.899,"tissuetumour":0.8626,"AveExpr":5.6145,"F":9.3232,"P.Value":0.0004,"adj.P.Val":0.0042},{"tissuedysplasia":0.8364,"tissuetumour":0.7141,"AveExpr":2.2269,"F":5.0619,"P.Value":0.0105,"adj.P.Val":0.043},{"tissuedysplasia":0.4689,"tissuetumour":1.276,"AveExpr":4.2087,"F":26.1078,"P.Value":3.3441e-08,"adj.P.Val":3.3007e-06},{"tissuedysplasia":0.5935,"tissuetumour":1.0764,"AveExpr":4.2698,"F":17.3324,"P.Value":2.8095e-06,"adj.P.Val":0.0001},{"tissuedysplasia":0.0494,"tissuetumour":1.0818,"AveExpr":3.0431,"F":16.2525,"P.Value":5.1835e-06,"adj.P.Val":0.0001},{"tissuedysplasia":-0.0807,"tissuetumour":-0.122,"AveExpr":3.8738,"F":0.3897,"P.Value":0.6796,"adj.P.Val":0.7701},{"tissuedysplasia":1.2614,"tissuetumour":1.9696,"AveExpr":-5.6068,"F":1.6428,"P.Value":0.2051,"adj.P.Val":0.3429},{"tissuedysplasia":0.2619,"tissuetumour":1.8779,"AveExpr":-4.6655,"F":1.8634,"P.Value":0.1672,"adj.P.Val":0.2988},{"tissuedysplasia":0.2856,"tissuetumour":0.2428,"AveExpr":2.0448,"F":0.7454,"P.Value":0.4805,"adj.P.Val":0.6099},{"tissuedysplasia":0.2572,"tissuetumour":1.0631,"AveExpr":3.5565,"F":17.6899,"P.Value":2.3024e-06,"adj.P.Val":0.0001},{"tissuedysplasia":0.0935,"tissuetumour":-0.2595,"AveExpr":5.7051,"F":3.9319,"P.Value":0.0268,"adj.P.Val":0.0844},{"tissuedysplasia":0.0429,"tissuetumour":-0.1752,"AveExpr":5.5306,"F":1.9751,"P.Value":0.1509,"adj.P.Val":0.2785},{"tissuedysplasia":0.5865,"tissuetumour":0.3891,"AveExpr":5.4672,"F":5.0597,"P.Value":0.0105,"adj.P.Val":0.043},{"tissuedysplasia":-0.6104,"tissuetumour":-0.8759,"AveExpr":6.8957,"F":23.8114,"P.Value":9.8089e-08,"adj.P.Val":7.0542e-06},{"tissuedysplasia":1.0162,"tissuetumour":1.7691,"AveExpr":1.0262,"F":10.8823,"P.Value":0.0001,"adj.P.Val":0.0018},{"tissuedysplasia":0.2874,"tissuetumour":1.3475,"AveExpr":4.2896,"F":21.1985,"P.Value":3.5708e-07,"adj.P.Val":0},{"tissuedysplasia":0.2816,"tissuetumour":0.7689,"AveExpr":3.2601,"F":9.328,"P.Value":0.0004,"adj.P.Val":0.0041},{"tissuedysplasia":-0.1669,"tissuetumour":0.3292,"AveExpr":0.421,"F":0.6851,"P.Value":0.5093,"adj.P.Val":0.6347},{"tissuedysplasia":1.3669,"tissuetumour":3.857,"AveExpr":-3.2788,"F":8.4367,"P.Value":0.0008,"adj.P.Val":0.0065},{"tissuedysplasia":0.0802,"tissuetumour":0.2134,"AveExpr":5.2117,"F":1.6288,"P.Value":0.2078,"adj.P.Val":0.346},{"tissuedysplasia":-1.0643,"tissuetumour":-1.7631,"AveExpr":-1.455,"F":2.4404,"P.Value":0.0988,"adj.P.Val":0.2092},{"tissuedysplasia":0.3872,"tissuetumour":0.646,"AveExpr":2.6846,"F":4.1331,"P.Value":0.0227,"adj.P.Val":0.0748},{"tissuedysplasia":0.037,"tissuetumour":0.7688,"AveExpr":5.1656,"F":17.8161,"P.Value":2.1471e-06,"adj.P.Val":0.0001},{"tissuedysplasia":-0.0138,"tissuetumour":0.319,"AveExpr":4.1247,"F":1.8878,"P.Value":0.1635,"adj.P.Val":0.2942},{"tissuedysplasia":0.5601,"tissuetumour":2.5072,"AveExpr":-5.9956,"F":2.82,"P.Value":0.0704,"adj.P.Val":0.1656},{"tissuedysplasia":0.9154,"tissuetumour":1.178,"AveExpr":4.6982,"F":9.6883,"P.Value":0.0003,"adj.P.Val":0.0034},{"tissuedysplasia":-0.2095,"tissuetumour":0.1906,"AveExpr":4.9229,"F":3.9773,"P.Value":0.0258,"adj.P.Val":0.082},{"tissuedysplasia":0.5947,"tissuetumour":1.1884,"AveExpr":0.5435,"F":3.4927,"P.Value":0.0391,"adj.P.Val":0.1098},{"tissuedysplasia":0.649,"tissuetumour":1.5524,"AveExpr":3.0816,"F":22.1875,"P.Value":2.1701e-07,"adj.P.Val":0},{"tissuedysplasia":-0.3426,"tissuetumour":-0.5627,"AveExpr":5.6127,"F":9.0575,"P.Value":0.0005,"adj.P.Val":0.0047},{"tissuedysplasia":0.4788,"tissuetumour":0.9821,"AveExpr":5.9424,"F":29.3432,"P.Value":7.9865e-09,"adj.P.Val":1.1952e-06},{"tissuedysplasia":-0.0834,"tissuetumour":0.8249,"AveExpr":2.1485,"F":6.8622,"P.Value":0.0025,"adj.P.Val":0.0154},{"tissuedysplasia":0.6986,"tissuetumour":0.4308,"AveExpr":2.5507,"F":3.9759,"P.Value":0.0259,"adj.P.Val":0.0821},{"tissuedysplasia":0.1912,"tissuetumour":0.019,"AveExpr":7.1456,"F":1.2481,"P.Value":0.297,"adj.P.Val":0.4414},{"tissuedysplasia":1.1038,"tissuetumour":2.5407,"AveExpr":-4.8517,"F":3.0539,"P.Value":0.0573,"adj.P.Val":0.1436},{"tissuedysplasia":-0.0253,"tissuetumour":-0.4693,"AveExpr":2.4003,"F":0.8412,"P.Value":0.438,"adj.P.Val":0.5746},{"tissuedysplasia":0.1596,"tissuetumour":0.6348,"AveExpr":3.5165,"F":5.4405,"P.Value":0.0077,"adj.P.Val":0.0345},{"tissuedysplasia":-0.2577,"tissuetumour":-0.2084,"AveExpr":4.041,"F":1.9321,"P.Value":0.1569,"adj.P.Val":0.2861},{"tissuedysplasia":-0.2149,"tissuetumour":-0.328,"AveExpr":5.8765,"F":3.7945,"P.Value":0.0302,"adj.P.Val":0.0916},{"tissuedysplasia":0.1023,"tissuetumour":-0.388,"AveExpr":3.5869,"F":3.5762,"P.Value":0.0364,"adj.P.Val":0.1043},{"tissuedysplasia":0.3615,"tissuetumour":-0.0409,"AveExpr":3.2617,"F":2.5765,"P.Value":0.0875,"adj.P.Val":0.1922},{"tissuedysplasia":-0.2081,"tissuetumour":-0.7457,"AveExpr":6.8012,"F":13.5367,"P.Value":0,"adj.P.Val":0.0005},{"tissuedysplasia":-0.2029,"tissuetumour":0.4624,"AveExpr":3.9872,"F":9.7898,"P.Value":0.0003,"adj.P.Val":0.0032},{"tissuedysplasia":0.2904,"tissuetumour":0.2672,"AveExpr":5.5733,"F":2.707,"P.Value":0.0778,"adj.P.Val":0.1773},{"tissuedysplasia":0.3602,"tissuetumour":0.0342,"AveExpr":3.2472,"F":2.7465,"P.Value":0.0752,"adj.P.Val":0.173},{"tissuedysplasia":0.4507,"tissuetumour":1.5894,"AveExpr":-5.2066,"F":1.0777,"P.Value":0.3492,"adj.P.Val":0.4931},{"tissuedysplasia":1.0219,"tissuetumour":0.9378,"AveExpr":2.1097,"F":7.6521,"P.Value":0.0014,"adj.P.Val":0.01},{"tissuedysplasia":-0.0991,"tissuetumour":-0.7935,"AveExpr":1.3598,"F":1.7943,"P.Value":0.1782,"adj.P.Val":0.3124},{"tissuedysplasia":0.3366,"tissuetumour":0.8796,"AveExpr":4.0404,"F":15.4027,"P.Value":8.4975e-06,"adj.P.Val":0.0002},{"tissuedysplasia":0.0865,"tissuetumour":0.4419,"AveExpr":1.8443,"F":1.7823,"P.Value":0.1802,"adj.P.Val":0.3148},{"tissuedysplasia":0.5666,"tissuetumour":0.8112,"AveExpr":4.526,"F":7.6447,"P.Value":0.0014,"adj.P.Val":0.01},{"tissuedysplasia":0.188,"tissuetumour":-0.0299,"AveExpr":5.5118,"F":2.0859,"P.Value":0.1363,"adj.P.Val":0.2605},{"tissuedysplasia":0.1597,"tissuetumour":-0.1939,"AveExpr":4.1669,"F":2.8242,"P.Value":0.0702,"adj.P.Val":0.1652},{"tissuedysplasia":0.1535,"tissuetumour":0.4875,"AveExpr":4.7062,"F":1.089,"P.Value":0.3463,"adj.P.Val":0.4903},{"tissuedysplasia":0.0783,"tissuetumour":-0.2693,"AveExpr":4.7755,"F":7.2385,"P.Value":0.0019,"adj.P.Val":0.0125},{"tissuedysplasia":0.0104,"tissuetumour":0.335,"AveExpr":7.8675,"F":7.4274,"P.Value":0.0017,"adj.P.Val":0.0113},{"tissuedysplasia":1.8226,"tissuetumour":2.9899,"AveExpr":-3.8426,"F":5.3076,"P.Value":0.0086,"adj.P.Val":0.0373},{"tissuedysplasia":0.4703,"tissuetumour":0.3705,"AveExpr":2.8817,"F":1.8087,"P.Value":0.1758,"adj.P.Val":0.3094},{"tissuedysplasia":0.2262,"tissuetumour":0.0644,"AveExpr":3.0627,"F":0.6751,"P.Value":0.5143,"adj.P.Val":0.6389},{"tissuedysplasia":0.247,"tissuetumour":0.1052,"AveExpr":4.3752,"F":2.042,"P.Value":0.1419,"adj.P.Val":0.2672},{"tissuedysplasia":0.0181,"tissuetumour":-0.4414,"AveExpr":4.7788,"F":7.6897,"P.Value":0.0014,"adj.P.Val":0.0098},{"tissuedysplasia":0.2719,"tissuetumour":0.0965,"AveExpr":5.1575,"F":3.3026,"P.Value":0.0461,"adj.P.Val":0.1233},{"tissuedysplasia":0.0508,"tissuetumour":-0.3779,"AveExpr":4.3404,"F":5.0305,"P.Value":0.0108,"adj.P.Val":0.0437},{"tissuedysplasia":-0.092,"tissuetumour":-0.4061,"AveExpr":3.8557,"F":3.522,"P.Value":0.0381,"adj.P.Val":0.1079},{"tissuedysplasia":0.0796,"tissuetumour":-0.406,"AveExpr":6.3981,"F":10.3756,"P.Value":0.0002,"adj.P.Val":0.0024},{"tissuedysplasia":0.221,"tissuetumour":1.1171,"AveExpr":3.3551,"F":13.3976,"P.Value":0,"adj.P.Val":0.0005},{"tissuedysplasia":0.4224,"tissuetumour":0.9214,"AveExpr":5.4322,"F":18.6508,"P.Value":1.3602e-06,"adj.P.Val":0},{"tissuedysplasia":0.0874,"tissuetumour":0.5568,"AveExpr":2.6704,"F":2.2696,"P.Value":0.1153,"adj.P.Val":0.2324},{"tissuedysplasia":0.4246,"tissuetumour":0.7226,"AveExpr":4.4126,"F":8.1753,"P.Value":0.001,"adj.P.Val":0.0075},{"tissuedysplasia":1.0481,"tissuetumour":1.2809,"AveExpr":4.1641,"F":9.0532,"P.Value":0.0005,"adj.P.Val":0.0047},{"tissuedysplasia":0.4497,"tissuetumour":0.5562,"AveExpr":3.8497,"F":8.9014,"P.Value":0.0006,"adj.P.Val":0.0051},{"tissuedysplasia":0.0556,"tissuetumour":-0.1301,"AveExpr":4.8719,"F":1.0039,"P.Value":0.3747,"adj.P.Val":0.5172},{"tissuedysplasia":-0.023,"tissuetumour":-0.5605,"AveExpr":6.1974,"F":14.9082,"P.Value":0,"adj.P.Val":0.0003},{"tissuedysplasia":0.1579,"tissuetumour":-0.1506,"AveExpr":4.3234,"F":3.5774,"P.Value":0.0363,"adj.P.Val":0.1042},{"tissuedysplasia":0.5909,"tissuetumour":0.7438,"AveExpr":2.1782,"F":5.2176,"P.Value":0.0093,"adj.P.Val":0.0392},{"tissuedysplasia":0.1006,"tissuetumour":0.8177,"AveExpr":4.4606,"F":12.3371,"P.Value":0.0001,"adj.P.Val":0.0009},{"tissuedysplasia":0.3993,"tissuetumour":0.3008,"AveExpr":3.7913,"F":2.0045,"P.Value":0.1468,"adj.P.Val":0.2737},{"tissuedysplasia":0.4955,"tissuetumour":-0.8636,"AveExpr":-1.091,"F":1.905,"P.Value":0.1609,"adj.P.Val":0.2911},{"tissuedysplasia":0.4735,"tissuetumour":0.8235,"AveExpr":2.697,"F":8.1735,"P.Value":0.001,"adj.P.Val":0.0075},{"tissuedysplasia":0.1519,"tissuetumour":0.7448,"AveExpr":3.8237,"F":8.2511,"P.Value":0.0009,"adj.P.Val":0.0072},{"tissuedysplasia":0.4898,"tissuetumour":1.0799,"AveExpr":4.4775,"F":23.4846,"P.Value":1.1482e-07,"adj.P.Val":7.8217e-06},{"tissuedysplasia":0.3054,"tissuetumour":0.607,"AveExpr":0.3899,"F":0.8224,"P.Value":0.446,"adj.P.Val":0.5815},{"tissuedysplasia":-0.0084,"tissuetumour":0.59,"AveExpr":4.1339,"F":7.064,"P.Value":0.0022,"adj.P.Val":0.0138},{"tissuedysplasia":-0.0508,"tissuetumour":0.3137,"AveExpr":-1.3693,"F":0.5634,"P.Value":0.5733,"adj.P.Val":0.6869},{"tissuedysplasia":0.3086,"tissuetumour":0.9037,"AveExpr":2.4994,"F":9.1492,"P.Value":0.0005,"adj.P.Val":0.0045},{"tissuedysplasia":0.0403,"tissuetumour":-0.5442,"AveExpr":5.9834,"F":10.7332,"P.Value":0.0002,"adj.P.Val":0.002},{"tissuedysplasia":0.5304,"tissuetumour":3.073,"AveExpr":-7.9412,"F":3.7699,"P.Value":0.0308,"adj.P.Val":0.0929},{"tissuedysplasia":0.1883,"tissuetumour":0.314,"AveExpr":6.1302,"F":8.1989,"P.Value":0.0009,"adj.P.Val":0.0074},{"tissuedysplasia":0.1939,"tissuetumour":0.2721,"AveExpr":6.4535,"F":2.088,"P.Value":0.136,"adj.P.Val":0.2601},{"tissuedysplasia":-0.4899,"tissuetumour":-0.9099,"AveExpr":4.3683,"F":16.9703,"P.Value":3.4434e-06,"adj.P.Val":0.0001},{"tissuedysplasia":0.5331,"tissuetumour":1.0767,"AveExpr":0.0679,"F":2.6796,"P.Value":0.0798,"adj.P.Val":0.1803},{"tissuedysplasia":0.2565,"tissuetumour":0.8615,"AveExpr":2.4609,"F":6.1439,"P.Value":0.0044,"adj.P.Val":0.023},{"tissuedysplasia":0.0119,"tissuetumour":0.1392,"AveExpr":4.1929,"F":0.4852,"P.Value":0.6188,"adj.P.Val":0.7236},{"tissuedysplasia":-0.1078,"tissuetumour":-0.2504,"AveExpr":4.4129,"F":1.1824,"P.Value":0.3161,"adj.P.Val":0.4604},{"tissuedysplasia":0.2838,"tissuetumour":0.0823,"AveExpr":4.0557,"F":2.6786,"P.Value":0.0798,"adj.P.Val":0.1804},{"tissuedysplasia":0.1822,"tissuetumour":0.1412,"AveExpr":2.9673,"F":0.4893,"P.Value":0.6164,"adj.P.Val":0.7217},{"tissuedysplasia":-0.0173,"tissuetumour":-0.208,"AveExpr":6.4225,"F":3.2902,"P.Value":0.0466,"adj.P.Val":0.1242},{"tissuedysplasia":0.3018,"tissuetumour":0.4547,"AveExpr":5.1398,"F":4.9382,"P.Value":0.0116,"adj.P.Val":0.046},{"tissuedysplasia":0.2529,"tissuetumour":0.2653,"AveExpr":4.532,"F":2.8662,"P.Value":0.0676,"adj.P.Val":0.1609},{"tissuedysplasia":0.2482,"tissuetumour":0.5233,"AveExpr":4.8226,"F":9.9295,"P.Value":0.0003,"adj.P.Val":0.003},{"tissuedysplasia":0.0032,"tissuetumour":-0.2412,"AveExpr":4.4042,"F":3.223,"P.Value":0.0494,"adj.P.Val":0.1296},{"tissuedysplasia":0.3091,"tissuetumour":-0.025,"AveExpr":1.5873,"F":1.2132,"P.Value":0.307,"adj.P.Val":0.4516},{"tissuedysplasia":0.1125,"tissuetumour":0.2549,"AveExpr":7.9296,"F":2.0768,"P.Value":0.1378,"adj.P.Val":0.2621},{"tissuedysplasia":0.1101,"tissuetumour":0.0687,"AveExpr":5.3675,"F":0.4336,"P.Value":0.6509,"adj.P.Val":0.7482},{"tissuedysplasia":-0.2341,"tissuetumour":-0.8458,"AveExpr":3.237,"F":4.7162,"P.Value":0.0139,"adj.P.Val":0.0527},{"tissuedysplasia":-0.0023,"tissuetumour":-1.0173,"AveExpr":4.379,"F":18.6463,"P.Value":1.3635e-06,"adj.P.Val":0},{"tissuedysplasia":0.9131,"tissuetumour":1.7787,"AveExpr":-0.2282,"F":5.2658,"P.Value":0.0089,"adj.P.Val":0.0382},{"tissuedysplasia":0.3043,"tissuetumour":0.9591,"AveExpr":1.6658,"F":7.0923,"P.Value":0.0021,"adj.P.Val":0.0136},{"tissuedysplasia":0.2492,"tissuetumour":0.3645,"AveExpr":4.7806,"F":6.6322,"P.Value":0.003,"adj.P.Val":0.0175},{"tissuedysplasia":-0.652,"tissuetumour":-0.2901,"AveExpr":1.6462,"F":1.274,"P.Value":0.2898,"adj.P.Val":0.4343},{"tissuedysplasia":0.3911,"tissuetumour":-0.0254,"AveExpr":6.3668,"F":5.9354,"P.Value":0.0052,"adj.P.Val":0.026},{"tissuedysplasia":0.4238,"tissuetumour":0.5556,"AveExpr":4.8867,"F":12.249,"P.Value":0.0001,"adj.P.Val":0.0009},{"tissuedysplasia":0.1918,"tissuetumour":0.045,"AveExpr":5.6162,"F":0.8589,"P.Value":0.4306,"adj.P.Val":0.5678},{"tissuedysplasia":0.0683,"tissuetumour":-0.2217,"AveExpr":5.6919,"F":3.4058,"P.Value":0.0421,"adj.P.Val":0.1158},{"tissuedysplasia":0.1722,"tissuetumour":0.0423,"AveExpr":4.1747,"F":1.3244,"P.Value":0.2764,"adj.P.Val":0.4208},{"tissuedysplasia":-0.2758,"tissuetumour":-0.7846,"AveExpr":6.4181,"F":22.9539,"P.Value":1.4865e-07,"adj.P.Val":9.4561e-06},{"tissuedysplasia":0.2368,"tissuetumour":-0.3079,"AveExpr":4.3315,"F":6.6115,"P.Value":0.0031,"adj.P.Val":0.0177},{"tissuedysplasia":0.0332,"tissuetumour":0.0329,"AveExpr":5.0362,"F":0.0844,"P.Value":0.9192,"adj.P.Val":0.9436},{"tissuedysplasia":0.1384,"tissuetumour":1.0456,"AveExpr":2.2975,"F":18.6109,"P.Value":1.3899e-06,"adj.P.Val":0.0001},{"tissuedysplasia":-0.1198,"tissuetumour":0.0684,"AveExpr":6.3554,"F":1.6984,"P.Value":0.1947,"adj.P.Val":0.3313},{"tissuedysplasia":-0.2855,"tissuetumour":-0.1614,"AveExpr":6.6615,"F":5.5401,"P.Value":0.0071,"adj.P.Val":0.0326},{"tissuedysplasia":0.5507,"tissuetumour":1.4359,"AveExpr":-1.8013,"F":4.4114,"P.Value":0.0179,"adj.P.Val":0.0635},{"tissuedysplasia":0.1631,"tissuetumour":0.0635,"AveExpr":5.8285,"F":0.369,"P.Value":0.6935,"adj.P.Val":0.7801},{"tissuedysplasia":0.4125,"tissuetumour":1.1033,"AveExpr":2.6455,"F":18.8949,"P.Value":1.1923e-06,"adj.P.Val":0},{"tissuedysplasia":-0.0286,"tissuetumour":0.0479,"AveExpr":6.7892,"F":0.287,"P.Value":0.7519,"adj.P.Val":0.8239},{"tissuedysplasia":-0.3065,"tissuetumour":0.4374,"AveExpr":0.8001,"F":1.1614,"P.Value":0.3225,"adj.P.Val":0.4667},{"tissuedysplasia":-0.152,"tissuetumour":-0.1807,"AveExpr":3.541,"F":0.8168,"P.Value":0.4484,"adj.P.Val":0.5836},{"tissuedysplasia":-0.4409,"tissuetumour":-2.0024,"AveExpr":4.5314,"F":40.2186,"P.Value":1.1657e-10,"adj.P.Val":5.6599e-08},{"tissuedysplasia":0.0478,"tissuetumour":-0.6408,"AveExpr":4.3743,"F":3.3717,"P.Value":0.0434,"adj.P.Val":0.1182},{"tissuedysplasia":0.129,"tissuetumour":0.1648,"AveExpr":-2.0723,"F":0.0375,"P.Value":0.9632,"adj.P.Val":0.9748},{"tissuedysplasia":0.0521,"tissuetumour":-0.2134,"AveExpr":7.4619,"F":2.8638,"P.Value":0.0677,"adj.P.Val":0.1611},{"tissuedysplasia":0.0784,"tissuetumour":0.6665,"AveExpr":4.6132,"F":9.8024,"P.Value":0.0003,"adj.P.Val":0.0032},{"tissuedysplasia":-0.1371,"tissuetumour":0.0141,"AveExpr":7.8836,"F":1.4054,"P.Value":0.2561,"adj.P.Val":0.399},{"tissuedysplasia":0.056,"tissuetumour":0.0626,"AveExpr":5.36,"F":0.1696,"P.Value":0.8446,"adj.P.Val":0.8909},{"tissuedysplasia":0.7029,"tissuetumour":1.1395,"AveExpr":0.8629,"F":4.937,"P.Value":0.0116,"adj.P.Val":0.0461},{"tissuedysplasia":0.0933,"tissuetumour":0.6743,"AveExpr":6.107,"F":23.4886,"P.Value":1.146e-07,"adj.P.Val":7.8156e-06},{"tissuedysplasia":0.8739,"tissuetumour":2.7097,"AveExpr":-0.9361,"F":7.8839,"P.Value":0.0012,"adj.P.Val":0.0088},{"tissuedysplasia":0.3691,"tissuetumour":0.9269,"AveExpr":5.042,"F":21.0093,"P.Value":3.9329e-07,"adj.P.Val":0},{"tissuedysplasia":0.0697,"tissuetumour":0.4475,"AveExpr":6.0372,"F":8.8302,"P.Value":0.0006,"adj.P.Val":0.0053},{"tissuedysplasia":-0.1367,"tissuetumour":-0.2779,"AveExpr":6.1833,"F":0.9331,"P.Value":0.4011,"adj.P.Val":0.5415},{"tissuedysplasia":0.943,"tissuetumour":1.2642,"AveExpr":1.534,"F":8.1112,"P.Value":0.001,"adj.P.Val":0.0077},{"tissuedysplasia":-0.2132,"tissuetumour":-0.3827,"AveExpr":4.1273,"F":2.6246,"P.Value":0.0838,"adj.P.Val":0.1866},{"tissuedysplasia":0.4421,"tissuetumour":0.3858,"AveExpr":1.1265,"F":1.2833,"P.Value":0.2873,"adj.P.Val":0.4318},{"tissuedysplasia":-0.1778,"tissuetumour":-0.0949,"AveExpr":1.5904,"F":0.1467,"P.Value":0.864,"adj.P.Val":0.9042},{"tissuedysplasia":0.3119,"tissuetumour":0.2007,"AveExpr":2.5518,"F":1.4831,"P.Value":0.2381,"a
```
