## Supplemental Examples for "R/LinkedCharts: A novel approach for simple but powerful interactive data analysis": R_code_full.html

```
countsums <- colSums(countMatrix)
openPage(useViewer = FALSE, layout = "table1x2")
selGene <- 1915

lc_scatter(
  x = voomResult$AveExpr,
  y = voomResult$tissuetumour,
  colour = ifelse(voomResult$adj.P.Val < 0.1, "red", "black"),
  label = rownames(voomResult),
  size = 1.3,
  axisTitleY = "log2 fc",
  axisTitleX = "Average expression",
  on_click = function(i) {
    selGene <<- i
    updateCharts("A2")
  },
  place = "A1")

#original data contains also information on dysplasia tissue sample
#remove them for plot's clarity
dyspl <- sampleTable$tissue == "dysplasia"

lc_scatter(dat(
    x = sampleTable$patient[!dyspl],
    y = countMatrix[selGene, !dyspl] / countsums[!dyspl] * 1e6 + .1,
    logScaleY = 10,
    colourValue = sampleTable$tissue[!dyspl],
    title = rownames(countMatrix)[selGene],
    axisTitleY = "CPM",
    axisTitleX = "Patients",
    ticksRotateX = 45,
    colourLegendTitle = "Tissue"),
  place = "A2")
```
