## Supplemental Examples for "R/LinkedCharts: A novel approach for simple but powerful interactive data analysis": R_code_min.html

```
countsums <- colSums(countMatrix)
openPage(useViewer = FALSE, layout = "table1x2")
selGene <- 1915

lc_scatter(
  x = voomResult$AveExpr,
  y = voomResult$tissuetumour,
  colour = ifelse(voomResult$adj.P.Val < 0.1, "red", "black"),
  on_click = function(i) {
    selGene <<- i
    updateCharts("A2")
  },
  place = "A1"
  )

lc_scatter(dat(
    x = sampleTable$patient,
    y = countMatrix[selGene, ] / countsums * 1e6 + .1,
    logScaleY = 10,
    colourValue = sampleTable$tissue,
    title = rownames(countMatrix)[selGene]),
  place = "A2")
```
