## Supplemental Examples for "R/LinkedCharts: A novel approach for simple but powerful interactive data analysis": JS_code_full.html

```
var xSample = 30, ySample = 31;

lc.heatmap()
  .nrows(hData.sampleNames.length)
  .ncols(hData.sampleNames.length)
  .value((i, j) => hData.corrMat[i][j])
  .rowLabel(i => hData.sampleNames[i])
  .colLabel(i => hData.sampleNames[i])
  .paddings({left: 55, top: 55, right: 30, bottom: 30})  
  .clusterRows(true)
  .clusterCols(true)
  .on_click(function(row, col) {
    xSample = row;
    ySample = col;
    sch.update();
  })
  .place("#heatmap");

var sch = lc.scatter()
//expression values are normalized on-the-fly.
//raw counts are used as input data
  .x(i => Math.log10(maData.countMatrix[i][xSample]/
    maData.countSums[xSample] * 1000000 + 0.1))
  .y(i => Math.log10(maData.countMatrix[i][ySample]/
    maData.countSums[ySample] * 1000000 + 0.1))
  .axisTitleX(() => hData.sampleNames[xSample])
  .axisTitleY(() => hData.sampleNames[ySample])
  .label(i => maData.geneNames[i])    
  .size(1.5)
  .opacity(0.5);

lc.xLine("line", sch)
  .lineFun(x => x)
  .colour("red")
  .place("#corplot");
```
