## Supplemental Examples for "R/LinkedCharts: A novel approach for simple but powerful interactive data analysis": R_code_full.html

```
xSample <- 31
ySample <- 32

colsums <- colSums(countMatrix)
normCounts <- t(log10(t(countMatrix)/colsums * 10^6 + 0.1))

openPage(FALSE, layout = "table1x2")

lc_heatmap(
  value = cor(normCounts, method = "spearman"),
  paddings = list(left = 55, top = 55, right = 30, bottom = 30),
  clusterRows = TRUE,
  clusterCols = TRUE,
  on_click = function(d) {
    xSample <<- d[1]
    ySample <<- d[2]
    updateCharts()
  }, place = "A1")

lc_scatter(dat(
  x = normCounts[, xSample],
  y = normCounts[, ySample],
  axisTitleX = colnames(normCounts)[xSample],
  axisTitleY = colnames(normCounts)[ySample]),
  size = 1.5,
  opacity = 0.5,
  place = "A2")

lc_abLine(a = 1, b = 0,
          colour = "red",
          chartId = "A2",
          addLayer = TRUE)
```
