## Supplemental Examples for "R/LinkedCharts: A novel approach for simple but powerful interactive data analysis": index.html

LinkedCharts - Supplement to the paper


### LinkedCharts - Supplement to the paper

This page provides live versions of the figures from the main paper. Each example is given as a *full* version with all the decorations, labels and titles and as a *minimalistic* app with only essential features.

For each example, one can look at the full code requried to generate the app both as an R script and as a JavaScript code. To run the R code, the "rlc" package must be installed from CRAN, or GitHub for the most recent version. The JavaScript code relies on the *linked-charts.js* library that can be downloaded from here. Links to all the required data for each example are given in the description below.
