## Supplemental Examples for "R/LinkedCharts: A novel approach for simple but powerful interactive data analysis": README.md

This directory contains the supplement for the paper "R/LinkedCharts: A novel approach for simple but powerful interactive data analysis". 

To view it, open the file index.html in any browser. Make sure that the subdirectories are available alongside the index.html file.

The content is also shown via GitHub Pages at https://anders-biostat.github.io/lc-paper/
