## Supplemental Examples for "R/LinkedCharts: A novel approach for simple but powerful interactive data analysis": OFL.txt

Copyright 2011 The Comfortaa Project Authors (https://github.com/alexeiva/comfortaa), with Reserved Font Name "Comfortaa".

This Font Software is licensed under the SIL Open Font License, Version 1.1.
This license is copied below, and is also available with a FAQ at:
http://scripts.sil.org/OFL


-----------------------------------------------------------
SIL OPEN FONT LICENSE Version 1.1 - 26 February 2007
-----------------------------------------------------------

PREAMBLE
The goals of the Open Font License (OFL) are to stimulate worldwide
development of collaborative font projects, to support the font creation
efforts of academic and linguistic communities, and to provide a free and
open framework in which fonts may be shared and improved in partnership
with others.

The OFL allows the licensed fonts to be used, studied, modified and
redistributed freely as long as they are not sold by themselves. The
fonts, including any derivative works, can be bundled, embedded, 
redistributed and/or sold with any software provided that any reserved
names are not used by derivative works. The fonts and derivatives,
however, cannot be released under any other type of license. The
requirement for fonts to remain under this license does not apply
to any document created using the fonts or their derivatives.

DEFINITIONS
"Font Software" refers to the set of files released by the Copyright
Holder(s) under this license and clearly marked as such. This may
include source files, build scripts and documentation.

"Reserved Font Name" refers to any names specified as such after the
copyright statement(s).

"Original Version" refers to the collection of Font Software components as
distributed by the Copyright Holder(s).

"Modified Version" refers to any derivative made by adding to, deleting,
or substituting -- in part or in whole -- any of the components of the
Original Version, by changing formats or by porting the Font Software to a
new environment.

"Author" refers to any designer, engineer, programmer, technical
writer or other person who contributed to the Font Software.

PERMISSION & CONDITIONS
Permission is hereby granted, free of charge, to any person obtaining
a copy of the Font Software, to use, study, copy, merge, embed, modify,
redistribute, and sell modified and unmodified copies of the Font
Software, subject to the following conditions:

1) Neither the Font Software nor any of its individual components,
in Original or Modified Versions, may be sold by itself.

2) Original or Modified Versions of the Font Software may be bundled,
redistributed and/or sold with any software, provided that each copy
contains the above copyright notice and this license. These can be
included either as stand-alone text files, human-readable headers or
in the appropriate machine-readable metadata fields within text or
binary files as long as those fields can be easily viewed by the user.

3) No Modified Version of the Font Software may use the Reserved Font
Name(s) unless explicit written permission is granted by the corresponding
Copyright Holder. This restriction only applies to the primary font name as
presented to the users.

4) The name(s) of the Copyright Holder(s) or the Author(s) of the Font
Software shall not be used to promote, endorse or advertise any
Modified Version, except to acknowledge the contribution(s) of the
Copyright Holder(s) and the Author(s) or with their explicit written
permission.

5) The Font Software, modified or unmodified, in part or in whole,
must be distributed entirely under this license, and must not be
distributed under any other license. The requirement for fonts to
remain under this license does not apply to any document created
using the Font Software.

TERMINATION
This license becomes null and void if any of the above conditions are
not met.

DISCLAIMER
THE FONT SOFTWARE IS PROVIDED "AS IS", WITHOUT WARRANTY OF ANY KIND,
EXPRESS OR IMPLIED, INCLUDING BUT NOT LIMITED TO ANY WARRANTIES OF
MERCHANTABILITY, FITNESS FOR A PARTICULAR PURPOSE AND NONINFRINGEMENT
OF COPYRIGHT, PATENT, TRADEMARK, OR OTHER RIGHT. IN NO EVENT SHALL THE
COPYRIGHT HOLDER BE LIABLE FOR ANY CLAIM, DAMAGES OR OTHER LIABILITY,
INCLUDING ANY GENERAL, SPECIAL, INDIRECT, INCIDENTAL, OR CONSEQUENTIAL
DAMAGES, WHETHER IN AN ACTION OF CONTRACT, TORT OR OTHERWISE, ARISING
FROM, OUT OF THE USE OR INABILITY TO USE THE FONT SOFTWARE OR FROM
OTHER DEALINGS IN THE FONT SOFTWARE.
