## Supplemental Examples for "R/LinkedCharts: A novel approach for simple but powerful interactive data analysis": README.txt

Comfortaa Variable Font
=======================

This download contains Comfortaa as both a variable font and static fonts.

Comfortaa is a variable font with this axis:
  wght

This means all the styles are contained in a single file:
  Comfortaa-VariableFont_wght.ttf

If your app fully supports variable fonts, you can now pick intermediate styles
that aren’t available as static fonts. Not all apps support variable fonts, and
in those cases you can use the static font files for Comfortaa:
  static/Comfortaa-Light.ttf
  static/Comfortaa-Regular.ttf
  static/Comfortaa-Medium.ttf
  static/Comfortaa-SemiBold.ttf
  static/Comfortaa-Bold.ttf

Get started
-----------

1. Install the font files you want to use

2. Use your app's font picker to view the font family and all the
available styles

Learn more about variable fonts
-------------------------------

  https://developers.google.com/web/fundamentals/design-and-ux/typography/variable-fonts
  https://variablefonts.typenetwork.com
  https://medium.com/variable-fonts

In desktop apps

  https://theblog.adobe.com/can-variable-fonts-illustrator-cc
  https://helpx.adobe.com/nz/photoshop/using/fonts.html#variable_fonts

Online

  https://developers.google.com/fonts/docs/getting_started
  https://developer.mozilla.org/en-US/docs/Web/CSS/CSS_Fonts/Variable_Fonts_Guide
  https://developer.microsoft.com/en-us/microsoft-edge/testdrive/demos/variable-fonts

Installing fonts

  MacOS: https://support.apple.com/en-us/HT201749
  Linux: https://www.google.com/search?q=how+to+install+a+font+on+gnu%2Blinux
  Windows: https://support.microsoft.com/en-us/help/314960/how-to-install-or-remove-a-font-in-windows

Android Apps

  https://developers.google.com/fonts/docs/android
  https://developer.android.com/guide/topics/ui/look-and-feel/downloadable-fonts

License
-------
Please read the full license text (OFL.txt) to understand the permissions,
restrictions and requirements for usage, redistribution, and modification.

You can use them freely in your products & projects - print or digital,
commercial or otherwise.

This isn't legal advice, please consider consulting a lawyer and see the full
license for all details.
