## Supplemental Examples for "R/LinkedCharts: A novel approach for simple but powerful interactive data analysis": linked-charts.min.js

```
//https://bl.ocks.org/Kcnarf/5c989173d0e0c74ab4b62161b33bb0a8 (original library was slightly modified)
(function(global,factory){typeof exports==="object"&&typeof module!=="undefined"?factory(exports):typeof define==="function"&&define.amd?define(["exports"],factory):factory(global.d3=global.d3||{})})(this,function(exports){"use strict";function SortedDirectAccessDoublyLinkedList(){this._valueAccessor=function(obj){return obj.value};this._idAccessor=function(obj){return obj.id};this._min=null;this._max=null;this._closestTo0=null;this.size=0;this._idToNode={}}SortedDirectAccessDoublyLinkedList.prototype.valueAccessor=function(_){if(!arguments.length){return this._valueAccessor}this._valueAccessor=_;return this};SortedDirectAccessDoublyLinkedList.prototype.idAccessor=function(_){if(!arguments.length){return this._idAccessor}this._idAccessor=_;return this};SortedDirectAccessDoublyLinkedList.prototype.min=function(){return this._min};SortedDirectAccessDoublyLinkedList.prototype.max=function(){return this._max};SortedDirectAccessDoublyLinkedList.prototype.closestTo0=function(){return this._closestTo0};SortedDirectAccessDoublyLinkedList.prototype.dln=function(datum){return this._idToNode[this._idAccessor(datum)]};SortedDirectAccessDoublyLinkedList.prototype.empty=function(){this._min=null;this._max=null;this._closestTo0=null;this.size=0;this._idToNode={};return this};SortedDirectAccessDoublyLinkedList.prototype.add=function(datum){var dln={datum:datum,value:this._valueAccessor(datum),prev:null,next:null};if(this.size===0){this._min=this._max=this._closestTo0=dln}else{if(dln.value<=this._min.value){dln.next=this._min;dln.next.prev=dln;this._min=dln}else if(dln.value>=this._max.value){dln.prev=this._max;dln.prev.next=dln;this._max=dln}else{var current=this._max;while(current.value>dln.value){current=current.prev}dln.prev=current;dln.next=current.next;dln.next.prev=dln;dln.prev.next=dln}if(Math.abs(dln.value)<Math.abs(this._closestTo0.value)){this._closestTo0=dln}}this._idToNode[this._idAccessor(datum)]=dln;this.size++;return this};SortedDirectAccessDoublyLinkedList.prototype.addMany=function(data){data.forEach(function(datum,item){this.add(datum)},this);return this};SortedDirectAccessDoublyLinkedList.prototype.remove=function(datum){var dln=this.dln(datum);if(this.size===1){this._min=this._max=this._closestTo0=null}else{if(dln===this._closestTo0){if(this._closestTo0.next===null){this._closestTo0=dln.prev}else if(this._closestTo0.prev===null){this._closestTo0=dln.prev}else{var prevAbsValue=Math.abs(this._closestTo0.prev.value);var nextAbsValue=Math.abs(this._closestTo0.next.value);if(prevAbsValue<nextAbsValue){this._closestTo0=this._closestTo0.prev}else{this._closestTo0=this._closestTo0.next}}}if(dln===this._min){this._min=this._min.next;this._min.prev=null}else if(dln===this._max){this._max=this._max.prev;this._max.next=null}else{dln.next.prev=dln.prev;dln.prev.next=dln.next}}delete this._idToNode[this._idAccessor(datum)];dln=null;this.size--;return this};function beeswarm(){var data=[];var radius=4;var orientation="horizontal";var side="symetric";var distributeOn=function(datum){return datum.x};var axis=function(datum){return datum.y};var minDistanceBetweenCircles;var minSquareDistanceBetweenCircles;var xBasedDataManager;var xBasedColliderManager;var yBasedColliderManager;var arrangement;var totalPossibleColliders,maxPossibleColliders,totalTestedPlacements,visitedColliderCount,totalVisitedColliders,maxVisitedColliders;function _beeswarm(){}_beeswarm.data=function(_){if(!arguments.length){return data}data=_;return _beeswarm};_beeswarm.radius=function(_){if(!arguments.length){return radius}radius=_;return _beeswarm};_beeswarm.orientation=function(_){if(!arguments.length){return orientation}if(_==="horizontal"||_==="vertical"){orientation=_}return _beeswarm};_beeswarm.side=function(_){if(!arguments.length){return side}if(_==="symetric"||_==="positive"||_==="negative"){side=_}return _beeswarm};_beeswarm.distributeOn=function(_){if(!arguments.length){return distributeOn}distributeOn=_;return _beeswarm};_beeswarm.axis=function(_){if(!arguments.length){return distributeOn}axis=_;return _beeswarm};_beeswarm.arrange=function(){initArrangement();arrangement.forEach(function(d){var bestYPosition=-Infinity,relativeYPos,xBasedPossibleColliders=gatherXBasedPossibleColliders(d);if(xBasedPossibleColliders.length===0){bestYPosition=d.axis}else{yBasedColliderManager.empty();yBasedColliderManager.addMany(xBasedPossibleColliders);d.free=d.axis;if(!collidesWithOther(d,yBasedColliderManager.closestTo0())){bestYPosition=d.axis;totalVisitedColliders+=visitedColliderCount;if(visitedColliderCount>maxVisitedColliders){maxVisitedColliders=visitedColliderCount}visitedColliderCount=0;totalTestedPlacements+=1}else{xBasedPossibleColliders.forEach(function(xbpc){relativeYPos=yPosRelativeToXbpc(xbpc,d);placeBelow(d,xbpc,relativeYPos);if(isAuthorizedPlacement(d)&&isBetterPlacement(d,bestYPosition)&&!collidesWithOther(d,yBasedColliderManager.dln(xbpc))){bestYPosition=d.free}totalVisitedColliders+=visitedColliderCount;if(visitedColliderCount>maxVisitedColliders){maxVisitedColliders=visitedColliderCount}visitedColliderCount=0;totalTestedPlacements+=1;placeAbove(d,xbpc,relativeYPos);if(isAuthorizedPlacement(d)&&isBetterPlacement(d,bestYPosition)&&!collidesWithOther(d,yBasedColliderManager.dln(xbpc))){bestYPosition=d.free}totalVisitedColliders+=visitedColliderCount;if(visitedColliderCount>maxVisitedColliders){maxVisitedColliders=visitedColliderCount}visitedColliderCount=0;totalTestedPlacements+=1})}}d.free=bestYPosition;if(orientation==="horizontal"){d.x=d.fixed;d.y=bestYPosition}else{d.x=bestYPosition;d.y=d.fixed}xBasedColliderManager.add(d)});return arrangement};_beeswarm.metrics=function(){return{totalPossibleColliders:totalPossibleColliders,maxPossibleColliders:maxPossibleColliders,totalTestedPlacements:totalTestedPlacements,visitedColliderCount:visitedColliderCount,totalVisitedColliders:totalVisitedColliders,maxVisitedColliders:maxVisitedColliders}};function initArrangement(){arrangement=data.map(function(d,i){return{datum:d,id:i,fixed:distributeOn(d),axis:axis(d),free:-Infinity}});minDistanceBetweenCircles=2*radius;minSquareDistanceBetweenCircles=Math.pow(minDistanceBetweenCircles,2);xBasedDataManager=(new SortedDirectAccessDoublyLinkedList).valueAccessor(function(d){return d.fixed}).addMany(arrangement);xBasedColliderManager=(new SortedDirectAccessDoublyLinkedList).valueAccessor(function(d){return d.fixed});yBasedColliderManager=(new SortedDirectAccessDoublyLinkedList).valueAccessor(function(d){return d.free});totalPossibleColliders=maxPossibleColliders=0;totalTestedPlacements=0;visitedColliderCount=totalVisitedColliders=maxVisitedColliders=0}function findNearestPossibleCollider(dln,visitedDln,direction){if(visitedDln===null){return null}else if(direction==="prev"?dln.value-visitedDln.value>minDistanceBetweenCircles:visitedDln.value-dln.value>minDistanceBetweenCircles){return null}else{if(isFinite(visitedDln.datum.free)){return visitedDln.datum}return findNearestPossibleCollider(dln,visitedDln[direction],direction)}}function visitToGatherXBasedPossibleColliders(dln,visitedDln,direction,xBasedPossibleColliders){if(visitedDln===null){return}else if(direction==="prev"?dln.value-visitedDln.value>minDistanceBetweenCircles:visitedDln.value-dln.value>minDistanceBetweenCircles){return}else{xBasedPossibleColliders.push(visitedDln.datum);visitToGatherXBasedPossibleColliders(dln,visitedDln[direction],direction,xBasedPossibleColliders)}}function gatherXBasedPossibleColliders(datum){var xBasedPossibleColliders=[];var dln=xBasedDataManager.dln(datum);var nearestXPrevAlreadyArrangedData=findNearestPossibleCollider(dln,dln.prev,"prev");var nearestXNextAlreadyArrangedData=findNearestPossibleCollider(dln,dln.next,"next");if(nearestXPrevAlreadyArrangedData!=null){dln=xBasedColliderManager.dln(nearestXPrevAlreadyArrangedData);visitToGatherXBasedPossibleColliders(dln,dln,"prev",xBasedPossibleColliders)}if(nearestXNextAlreadyArrangedData!=null){dln=xBasedColliderManager.dln(nearestXNextAlreadyArrangedData);visitToGatherXBasedPossibleColliders(dln,dln,"next",xBasedPossibleColliders)}totalPossibleColliders+=xBasedPossibleColliders.length;if(xBasedPossibleColliders.length>maxPossibleColliders){maxPossibleColliders=xBasedPossibleColliders.length}return xBasedPossibleColliders}function isAuthorizedPlacement(datum){if(side==="symetric"){return true}else if(side==="positive"){return datum.free>=0}else{return datum.free<=0}}function isBetterPlacement(datum,bestYPosition){return Math.abs(datum.free-datum.axis)<Math.abs(bestYPosition-datum.axis)}function yPosRelativeToXbpc(xbpc,d){return Math.sqrt(minSquareDistanceBetweenCircles-Math.pow(d.fixed-xbpc.fixed,2))+1e-6}function placeBelow(d,aad,relativeYPos){d.free=aad.free-relativeYPos}function placeAbove(d,aad,relativeYPos){d.free=aad.free+relativeYPos}function areCirclesColliding(d0,d1){visitedColliderCount++;return Math.pow(d1.free-d0.free,2)+Math.pow(d1.fixed-d0.fixed,2)<minSquareDistanceBetweenCircles}function visitToDetectCollisionWithOther(datum,visitedDln,direction,visitCount){if(visitedDln===null){return false}else if(direction==="prev"?datum.free-visitedDln.datum.free>minDistanceBetweenCircles:visitedDln.datum.free-datum.free>minDistanceBetweenCircles){return false}else if(areCirclesColliding(datum,visitedDln.datum)){return true}else{return visitToDetectCollisionWithOther(datum,visitedDln[direction],direction,visitCount++)}}function collidesWithOther(datum,visitedDln){var visitCount=0;if(visitToDetectCollisionWithOther(datum,visitedDln,"prev",visitCount++)){return true}else{return visitToDetectCollisionWithOther(datum,visitedDln,"next",visitCount++)}}return _beeswarm}exports.beeswarm=beeswarm;Object.defineProperty(exports,"__esModule",{value:true})});
// https://d3js.org Version 4.10.0. Copyright 2017 Mike Bostock.
(function(t,n){"object"==typeof exports&&"undefined"!=typeof module?n(exports):"function"==typeof define&&define.amd?define(["exports"],n):n(t.d3=t.d3||{})})(this,function(t){"use strict";function n(t){return function(n,e){return ss(t(n),e)}}function e(t,n){return[t,n]}function r(t,n,e){var r=(n-t)/Math.max(0,e),i=Math.floor(Math.log(r)/Math.LN10),o=r/Math.pow(10,i);return i>=0?(o>=Ts?10:o>=ks?5:o>=Ns?2:1)*Math.pow(10,i):-Math.pow(10,-i)/(o>=Ts?10:o>=ks?5:o>=Ns?2:1)}function i(t,n,e){var r=Math.abs(n-t)/Math.max(0,e),i=Math.pow(10,Math.floor(Math.log(r)/Math.LN10)),o=r/i;return o>=Ts?i*=10:o>=ks?i*=5:o>=Ns&&(i*=2),n<t?-i:i}function o(t){return t.length}function u(t){return"translate("+(t+.5)+",0)"}function a(t){return"translate(0,"+(t+.5)+")"}function c(t){return function(n){return+t(n)}}function s(t){var n=Math.max(0,t.bandwidth()-1)/2;return t.round()&&(n=Math.round(n)),function(e){return+t(e)+n}}function f(){return!this.__axis}function l(t,n){function e(e){var u=null==i?n.ticks?n.ticks.apply(n,r):n.domain():i,a=null==o?n.tickFormat?n.tickFormat.apply(n,r):Ls:o,y=Math.max(l,0)+p,g=n.range(),m=+g[0]+.5,x=+g[g.length-1]+.5,b=(n.bandwidth?s:c)(n.copy()),w=e.selection?e.selection():e,M=w.selectAll(".domain").data([null]),T=w.selectAll(".tick").data(u,n).order(),k=T.exit(),N=T.enter().append("g").attr("class","tick"),S=T.select("line"),E=T.select("text");M=M.merge(M.enter().insert("path",".tick").attr("class","domain").attr("stroke","#000")),T=T.merge(N),S=S.merge(N.append("line").attr("stroke","#000").attr(v+"2",d*l)),E=E.merge(N.append("text").attr("fill","#000").attr(v,d*y).attr("dy",t===qs?"0em":t===Ds?"0.71em":"0.32em")),e!==w&&(M=M.transition(e),T=T.transition(e),S=S.transition(e),E=E.transition(e),k=k.transition(e).attr("opacity",Fs).attr("transform",function(t){return isFinite(t=b(t))?_(t):this.getAttribute("transform")}),N.attr("opacity",Fs).attr("transform",function(t){var n=this.parentNode.__axis;return _(n&&isFinite(n=n(t))?n:b(t))})),k.remove(),M.attr("d",t===Os||t==Us?"M"+d*h+","+m+"H0.5V"+x+"H"+d*h:"M"+m+","+d*h+"V0.5H"+x+"V"+d*h),T.attr("opacity",1).attr("transform",function(t){return _(b(t))}),S.attr(v+"2",d*l),E.attr(v,d*y).text(a),w.filter(f).attr("fill","none").attr("font-size",10).attr("font-family","sans-serif").attr("text-anchor",t===Us?"start":t===Os?"end":"middle"),w.each(function(){this.__axis=b})}var r=[],i=null,o=null,l=6,h=6,p=3,d=t===qs||t===Os?-1:1,v=t===Os||t===Us?"x":"y",_=t===qs||t===Ds?u:a;return e.scale=function(t){return arguments.length?(n=t,e):n},e.ticks=function(){return r=Rs.call(arguments),e},e.tickArguments=function(t){return arguments.length?(r=null==t?[]:Rs.call(t),e):r.slice()},e.tickValues=function(t){return arguments.length?(i=null==t?null:Rs.call(t),e):i&&i.slice()},e.tickFormat=function(t){return arguments.length?(o=t,e):o},e.tickSize=function(t){return arguments.length?(l=h=+t,e):l},e.tickSizeInner=function(t){return arguments.length?(l=+t,e):l},e.tickSizeOuter=function(t){return arguments.length?(h=+t,e):h},e.tickPadding=function(t){return arguments.length?(p=+t,e):p},e}function h(){for(var t,n=0,e=arguments.length,r={};n<e;++n){if(!(t=arguments[n]+"")||t in r)throw new Error("illegal type: "+t);r[t]=[]}return new p(r)}function p(t){this._=t}function d(t,n){return t.trim().split(/^|\s+/).map(function(t){var e="",r=t.indexOf(".");if(r>=0&&(e=t.slice(r+1),t=t.slice(0,r)),t&&!n.hasOwnProperty(t))throw new Error("unknown type: "+t);return{type:t,name:e}})}function v(t,n){for(var e,r=0,i=t.length;r<i;++r)if((e=t[r]).name===n)return e.value}function _(t,n,e){for(var r=0,i=t.length;r<i;++r)if(t[r].name===n){t[r]=Is,t=t.slice(0,r).concat(t.slice(r+1));break}return null!=e&&t.push({name:n,value:e}),t}function y(t){return function(){var n=this.ownerDocument,e=this.namespaceURI;return e===Ys&&n.documentElement.namespaceURI===Ys?n.createElement(t):n.createElementNS(e,t)}}function g(t){return function(){return this.ownerDocument.createElementNS(t.space,t.local)}}function m(){return new x}function x(){this._="@"+(++Xs).toString(36)}function b(t,n,e){return t=w(t,n,e),function(n){var e=n.relatedTarget;e&&(e===this||8&e.compareDocumentPosition(this))||t.call(this,n)}}function w(n,e,r){return function(i){var o=t.event;t.event=i;try{n.call(this,this.__data__,e,r)}finally{t.event=o}}}function M(t){return t.trim().split(/^|\s+/).map(function(t){var n="",e=t.indexOf(".");return e>=0&&(n=t.slice(e+1),t=t.slice(0,e)),{type:t,name:n}})}function T(t){return function(){var n=this.__on;if(n){for(var e,r=0,i=-1,o=n.length;r<o;++r)e=n[r],t.type&&e.type!==t.type||e.name!==t.name?n[++i]=e:this.removeEventListener(e.type,e.listener,e.capture);++i?n.length=i:delete this.__on}}}function k(t,n,e){var r=Gs.hasOwnProperty(t.type)?b:w;return function(i,o,u){var a,c=this.__on,s=r(n,o,u);if(c)for(var f=0,l=c.length;f<l;++f)if((a=c[f]).type===t.type&&a.name===t.name)return this.removeEventListener(a.type,a.listener,a.capture),this.addEventListener(a.type,a.listener=s,a.capture=e),void(a.value=n);this.addEventListener(t.type,s,e),a={type:t.type,name:t.name,value:n,listener:s,capture:e},c?c.push(a):this.__on=[a]}}function N(n,e,r,i){var o=t.event;n.sourceEvent=t.event,t.event=n;try{return e.apply(r,i)}finally{t.event=o}}function S(){}function E(){return[]}function A(t,n){this.ownerDocument=t.ownerDocument,this.namespaceURI=t.namespaceURI,this._next=null,this._parent=t,this.__data__=n}function C(t,n,e,r,i,o){for(var u,a=0,c=n.length,s=o.length;a<s;++a)(u=n[a])?(u.__data__=o[a],r[a]=u):e[a]=new A(t,o[a]);for(;a<c;++a)(u=n[a])&&(i[a]=u)}function z(t,n,e,r,i,o,u){var a,c,s,f={},l=n.length,h=o.length,p=new Array(l);for(a=0;a<l;++a)(c=n[a])&&(p[a]=s=of+u.call(c,c.__data__,a,n),s in f?i[a]=c:f[s]=c);for(a=0;a<h;++a)(c=f[s=of+u.call(t,o[a],a,o)])?(r[a]=c,c.__data__=o[a],f[s]=null):e[a]=new A(t,o[a]);for(a=0;a<l;++a)(c=n[a])&&f[p[a]]===c&&(i[a]=c)}function P(t,n){return t<n?-1:t>n?1:t>=n?0:NaN}function R(t){return function(){this.removeAttribute(t)}}function L(t){return function(){this.removeAttributeNS(t.space,t.local)}}function q(t,n){return function(){this.setAttribute(t,n)}}function U(t,n){return function(){this.setAttributeNS(t.space,t.local,n)}}function D(t,n){return function(){var e=n.apply(this,arguments);null==e?this.removeAttribute(t):this.setAttribute(t,e)}}function O(t,n){return function(){var e=n.apply(this,arguments);null==e?this.removeAttributeNS(t.space,t.local):this.setAttributeNS(t.space,t.local,e)}}function F(t){return function(){this.style.removeProperty(t)}}function I(t,n,e){return function(){this.style.setProperty(t,n,e)}}function Y(t,n,e){return function(){var r=n.apply(this,arguments);null==r?this.style.removeProperty(t):this.style.setProperty(t,r,e)}}function B(t,n){return t.style.getPropertyValue(n)||uf(t).getComputedStyle(t,null).getPropertyValue(n)}function j(t){return function(){delete this[t]}}function H(t,n){return function(){this[t]=n}}function X(t,n){return function(){var e=n.apply(this,arguments);null==e?delete this[t]:this[t]=e}}function $(t){return t.trim().split(/^|\s+/)}function V(t){return t.classList||new W(t)}function W(t){this._node=t,this._names=$(t.getAttribute("class")||"")}function Z(t,n){for(var e=V(t),r=-1,i=n.length;++r<i;)e.add(n[r])}function G(t,n){for(var e=V(t),r=-1,i=n.length;++r<i;)e.remove(n[r])}function J(t){return function(){Z(this,t)}}function Q(t){return function(){G(this,t)}}function K(t,n){return function(){(n.apply(this,arguments)?Z:G)(this,t)}}function tt(){this.textContent=""}function nt(t){return function(){this.textContent=t}}function et(t){return function(){var n=t.apply(this,arguments);this.textContent=null==n?"":n}}function rt(){this.innerHTML=""}function it(t){return function(){this.innerHTML=t}}function ot(t){return function(){var n=t.apply(this,arguments);this.innerHTML=null==n?"":n}}function ut(){this.nextSibling&&this.parentNode.appendChild(this)}function at(){this.previousSibling&&this.parentNode.insertBefore(this,this.parentNode.firstChild)}function ct(){return null}function st(){var t=this.parentNode;t&&t.removeChild(this)}function ft(t,n,e){var r=uf(t),i=r.CustomEvent;"function"==typeof i?i=new i(n,e):(i=r.document.createEvent("Event"),e?(i.initEvent(n,e.bubbles,e.cancelable),i.detail=e.detail):i.initEvent(n,!1,!1)),t.dispatchEvent(i)}function lt(t,n){return function(){return ft(this,t,n)}}function ht(t,n){return function(){return ft(this,t,n.apply(this,arguments))}}function pt(t,n){this._groups=t,this._parents=n}function dt(){return new pt([[document.documentElement]],af)}function vt(){t.event.stopImmediatePropagation()}function _t(t,n){var e=t.document.documentElement,r=cf(t).on("dragstart.drag",null);n&&(r.on("click.drag",ff,!0),setTimeout(function(){r.on("click.drag",null)},0)),"onselectstart"in e?r.on("selectstart.drag",null):(e.style.MozUserSelect=e.__noselect,delete e.__noselect)}function yt(t,n,e,r,i,o,u,a,c,s){this.target=t,this.type=n,this.subject=e,this.identifier=r,this.active=i,this.x=o,this.y=u,this.dx=a,this.dy=c,this._=s}function gt(){return!t.event.button}function mt(){return this.parentNode}function xt(n){return null==n?{x:t.event.x,y:t.event.y}:n}function bt(){return"ontouchstart"in this}function wt(t,n){var e=Object.create(t.prototype);for(var r in n)e[r]=n[r];return e}function Mt(){}function Tt(t){var n;return t=(t+"").trim().toLowerCase(),(n=yf.exec(t))?(n=parseInt(n[1],16),new At(n>>8&15|n>>4&240,n>>4&15|240&n,(15&n)<<4|15&n,1)):(n=gf.exec(t))?kt(parseInt(n[1],16)):(n=mf.exec(t))?new At(n[1],n[2],n[3],1):(n=xf.exec(t))?new At(255*n[1]/100,255*n[2]/100,255*n[3]/100,1):(n=bf.exec(t))?Nt(n[1],n[2],n[3],n[4]):(n=wf.exec(t))?Nt(255*n[1]/100,255*n[2]/100,255*n[3]/100,n[4]):(n=Mf.exec(t))?Ct(n[1],n[2]/100,n[3]/100,1):(n=Tf.exec(t))?Ct(n[1],n[2]/100,n[3]/100,n[4]):kf.hasOwnProperty(t)?kt(kf[t]):"transparent"===t?new At(NaN,NaN,NaN,0):null}function kt(t){return new At(t>>16&255,t>>8&255,255&t,1)}function Nt(t,n,e,r){return r<=0&&(t=n=e=NaN),new At(t,n,e,r)}function St(t){return t instanceof Mt||(t=Tt(t)),t?(t=t.rgb(),new At(t.r,t.g,t.b,t.opacity)):new At}function Et(t,n,e,r){return 1===arguments.length?St(t):new At(t,n,e,null==r?1:r)}function At(t,n,e,r){this.r=+t,this.g=+n,this.b=+e,this.opacity=+r}function Ct(t,n,e,r){return r<=0?t=n=e=NaN:e<=0||e>=1?t=n=NaN:n<=0&&(t=NaN),new Rt(t,n,e,r)}function zt(t){if(t instanceof Rt)return new Rt(t.h,t.s,t.l,t.opacity);if(t instanceof Mt||(t=Tt(t)),!t)return new Rt;if(t instanceof Rt)return t;var n=(t=t.rgb()).r/255,e=t.g/255,r=t.b/255,i=Math.min(n,e,r),o=Math.max(n,e,r),u=NaN,a=o-i,c=(o+i)/2;return a?(u=n===o?(e-r)/a+6*(e<r):e===o?(r-n)/a+2:(n-e)/a+4,a/=c<.5?o+i:2-o-i,u*=60):a=c>0&&c<1?0:u,new Rt(u,a,c,t.opacity)}function Pt(t,n,e,r){return 1===arguments.length?zt(t):new Rt(t,n,e,null==r?1:r)}function Rt(t,n,e,r){this.h=+t,this.s=+n,this.l=+e,this.opacity=+r}function Lt(t,n,e){return 255*(t<60?n+(e-n)*t/60:t<180?e:t<240?n+(e-n)*(240-t)/60:n)}function qt(t){if(t instanceof Dt)return new Dt(t.l,t.a,t.b,t.opacity);if(t instanceof Ht){var n=t.h*Nf;return new Dt(t.l,Math.cos(n)*t.c,Math.sin(n)*t.c,t.opacity)}t instanceof At||(t=St(t));var e=Yt(t.r),r=Yt(t.g),i=Yt(t.b),o=Ot((.4124564*e+.3575761*r+.1804375*i)/Ef),u=Ot((.2126729*e+.7151522*r+.072175*i)/Af);return new Dt(116*u-16,500*(o-u),200*(u-Ot((.0193339*e+.119192*r+.9503041*i)/Cf)),t.opacity)}function Ut(t,n,e,r){return 1===arguments.length?qt(t):new Dt(t,n,e,null==r?1:r)}function Dt(t,n,e,r){this.l=+t,this.a=+n,this.b=+e,this.opacity=+r}function Ot(t){return t>Lf?Math.pow(t,1/3):t/Rf+zf}function Ft(t){return t>Pf?t*t*t:Rf*(t-zf)}function It(t){return 255*(t<=.0031308?12.92*t:1.055*Math.pow(t,1/2.4)-.055)}function Yt(t){return(t/=255)<=.04045?t/12.92:Math.pow((t+.055)/1.055,2.4)}function Bt(t){if(t instanceof Ht)return new Ht(t.h,t.c,t.l,t.opacity);t instanceof Dt||(t=qt(t));var n=Math.atan2(t.b,t.a)*Sf;return new Ht(n<0?n+360:n,Math.sqrt(t.a*t.a+t.b*t.b),t.l,t.opacity)}function jt(t,n,e,r){return 1===arguments.length?Bt(t):new Ht(t,n,e,null==r?1:r)}function Ht(t,n,e,r){this.h=+t,this.c=+n,this.l=+e,this.opacity=+r}function Xt(t){if(t instanceof Vt)return new Vt(t.h,t.s,t.l,t.opacity);t instanceof At||(t=St(t));var n=t.r/255,e=t.g/255,r=t.b/255,i=(Bf*r+If*n-Yf*e)/(Bf+If-Yf),o=r-i,u=(Ff*(e-i)-Df*o)/Of,a=Math.sqrt(u*u+o*o)/(Ff*i*(1-i)),c=a?Math.atan2(u,o)*Sf-120:NaN;return new Vt(c<0?c+360:c,a,i,t.opacity)}function $t(t,n,e,r){return 1===arguments.length?Xt(t):new Vt(t,n,e,null==r?1:r)}function Vt(t,n,e,r){this.h=+t,this.s=+n,this.l=+e,this.opacity=+r}function Wt(t,n,e,r,i){var o=t*t,u=o*t;return((1-3*t+3*o-u)*n+(4-6*o+3*u)*e+(1+3*t+3*o-3*u)*r+u*i)/6}function Zt(t,n){return function(e){return t+e*n}}function Gt(t,n,e){return t=Math.pow(t,e),n=Math.pow(n,e)-t,e=1/e,function(r){return Math.pow(t+r*n,e)}}function Jt(t,n){var e=n-t;return e?Zt(t,e>180||e<-180?e-360*Math.round(e/360):e):Jf(isNaN(t)?n:t)}function Qt(t){return 1==(t=+t)?Kt:function(n,e){return e-n?Gt(n,e,t):Jf(isNaN(n)?e:n)}}function Kt(t,n){var e=n-t;return e?Zt(t,e):Jf(isNaN(t)?n:t)}function tn(t){return function(n){var e,r,i=n.length,o=new Array(i),u=new Array(i),a=new Array(i);for(e=0;e<i;++e)r=Et(n[e]),o[e]=r.r||0,u[e]=r.g||0,a[e]=r.b||0;return o=t(o),u=t(u),a=t(a),r.opacity=1,function(t){return r.r=o(t),r.g=u(t),r.b=a(t),r+""}}}function nn(t){return function(){return t}}function en(t){return function(n){return t(n)+""}}function rn(t,n,e,r){function i(t){return t.length?t.pop()+" ":""}function o(t,r,i,o,u,a){if(t!==i||r!==o){var c=u.push("translate(",null,n,null,e);a.push({i:c-4,x:rl(t,i)},{i:c-2,x:rl(r,o)})}else(i||o)&&u.push("translate("+i+n+o+e)}function u(t,n,e,o){t!==n?(t-n>180?n+=360:n-t>180&&(t+=360),o.push({i:e.push(i(e)+"rotate(",null,r)-2,x:rl(t,n)})):n&&e.push(i(e)+"rotate("+n+r)}function a(t,n,e,o){t!==n?o.push({i:e.push(i(e)+"skewX(",null,r)-2,x:rl(t,n)}):n&&e.push(i(e)+"skewX("+n+r)}function c(t,n,e,r,o,u){if(t!==e||n!==r){var a=o.push(i(o)+"scale(",null,",",null,")");u.push({i:a-4,x:rl(t,e)},{i:a-2,x:rl(n,r)})}else 1===e&&1===r||o.push(i(o)+"scale("+e+","+r+")")}return function(n,e){var r=[],i=[];return n=t(n),e=t(e),o(n.translateX,n.translateY,e.translateX,e.translateY,r,i),u(n.rotate,e.rotate,r,i),a(n.skewX,e.skewX,r,i),c(n.scaleX,n.scaleY,e.scaleX,e.scaleY,r,i),n=e=null,function(t){for(var n,e=-1,o=i.length;++e<o;)r[(n=i[e]).i]=n.x(t);return r.join("")}}}function on(t){return((t=Math.exp(t))+1/t)/2}function un(t){return((t=Math.exp(t))-1/t)/2}function an(t){return((t=Math.exp(2*t))-1)/(t+1)}function cn(t){return function(n,e){var r=t((n=Pt(n)).h,(e=Pt(e)).h),i=Kt(n.s,e.s),o=Kt(n.l,e.l),u=Kt(n.opacity,e.opacity);return function(t){return n.h=r(t),n.s=i(t),n.l=o(t),n.opacity=u(t),n+""}}}function sn(t){return function(n,e){var r=t((n=jt(n)).h,(e=jt(e)).h),i=Kt(n.c,e.c),o=Kt(n.l,e.l),u=Kt(n.opacity,e.opacity);return function(t){return n.h=r(t),n.c=i(t),n.l=o(t),n.opacity=u(t),n+""}}}function fn(t){return function n(e){function r(n,r){var i=t((n=$t(n)).h,(r=$t(r)).h),o=Kt(n.s,r.s),u=Kt(n.l,r.l),a=Kt(n.opacity,r.opacity);return function(t){return n.h=i(t),n.s=o(t),n.l=u(Math.pow(t,e)),n.opacity=a(t),n+""}}return e=+e,r.gamma=n,r}(1)}function ln(){return El||(zl(hn),El=Cl.now()+Al)}function hn(){El=0}function pn(){this._call=this._time=this._next=null}function dn(t,n,e){var r=new pn;return r.restart(t,n,e),r}function vn(){ln(),++Ml;for(var t,n=Vf;n;)(t=El-n._time)>=0&&n._call.call(null,t),n=n._next;--Ml}function _n(){El=(Sl=Cl.now())+Al,Ml=Tl=0;try{vn()}finally{Ml=0,gn(),El=0}}function yn(){var t=Cl.now(),n=t-Sl;n>Nl&&(Al-=n,Sl=t)}function gn(){for(var t,n,e=Vf,r=1/0;e;)e._call?(r>e._time&&(r=e._time),t=e,e=e._next):(n=e._next,e._next=null,e=t?t._next=n:Vf=n);Wf=t,mn(r)}function mn(t){if(!Ml){Tl&&(Tl=clearTimeout(Tl));var n=t-El;n>24?(t<1/0&&(Tl=setTimeout(_n,n)),kl&&(kl=clearInterval(kl))):(kl||(Sl=El,kl=setInterval(yn,Nl)),Ml=1,zl(_n))}}function xn(t,n){var e=t.__transition;if(!e||!(e=e[n])||e.state>ql)throw new Error("too late");return e}function bn(t,n){var e=t.__transition;if(!e||!(e=e[n])||e.state>Dl)throw new Error("too late");return e}function wn(t,n){var e=t.__transition;if(!e||!(e=e[n]))throw new Error("too late");return e}function Mn(t,n,e){function r(c){var s,f,l,h;if(e.state!==Ul)return o();for(s in a)if((h=a[s]).name===e.name){if(h.state===Ol)return Pl(r);h.state===Fl?(h.state=Yl,h.timer.stop(),h.on.call("interrupt",t,t.__data__,h.index,h.group),delete a[s]):+s<n&&(h.state=Yl,h.timer.stop(),delete a[s])}if(Pl(function(){e.state===Ol&&(e.state=Fl,e.timer.restart(i,e.delay,e.time),i(c))}),e.state=Dl,e.on.call("start",t,t.__data__,e.index,e.group),e.state===Dl){for(e.state=Ol,u=new Array(l=e.tween.length),s=0,f=-1;s<l;++s)(h=e.tween[s].value.call(t,t.__data__,e.index,e.group))&&(u[++f]=h);u.length=f+1}}function i(n){for(var r=n<e.duration?e.ease.call(null,n/e.duration):(e.timer.restart(o),e.state=Il,1),i=-1,a=u.length;++i<a;)u[i].call(null,r);e.state===Il&&(e.on.call("end",t,t.__data__,e.index,e.group),o())}function o(){e.state=Yl,e.timer.stop(),delete a[n];for(var r in a)return;delete t.__transition}var u,a=t.__transition;a[n]=e,e.timer=dn(function(t){e.state=Ul,e.timer.restart(r,e.delay,e.time),e.delay<=t&&r(t-e.delay)},0,e.time)}function Tn(t,n){var e,r;return function(){var i=bn(this,t),o=i.tween;if(o!==e)for(var u=0,a=(r=e=o).length;u<a;++u)if(r[u].name===n){(r=r.slice()).splice(u,1);break}i.tween=r}}function kn(t,n,e){var r,i;if("function"!=typeof e)throw new Error;return function(){var o=bn(this,t),u=o.tween;if(u!==r){i=(r=u).slice();for(var a={name:n,value:e},c=0,s=i.length;c<s;++c)if(i[c].name===n){i[c]=a;break}c===s&&i.push(a)}o.tween=i}}function Nn(t,n,e){var r=t._id;return t.each(function(){var t=bn(this,r);(t.value||(t.value={}))[n]=e.apply(this,arguments)}),function(t){return wn(t,r).value[n]}}function Sn(t){return function(){this.removeAttribute(t)}}function En(t){return function(){this.removeAttributeNS(t.space,t.local)}}function An(t,n,e){var r,i;return function(){var o=this.getAttribute(t);return o===e?null:o===r?i:i=n(r=o,e)}}function Cn(t,n,e){var r,i;return function(){var o=this.getAttributeNS(t.space,t.local);return o===e?null:o===r?i:i=n(r=o,e)}}function zn(t,n,e){var r,i,o;return function(){var u,a=e(this);{if(null!=a)return(u=this.getAttribute(t))===a?null:u===r&&a===i?o:o=n(r=u,i=a);this.removeAttribute(t)}}}function Pn(t,n,e){var r,i,o;return function(){var u,a=e(this);{if(null!=a)return(u=this.getAttributeNS(t.space,t.local))===a?null:u===r&&a===i?o:o=n(r=u,i=a);this.removeAttributeNS(t.space,t.local)}}}function Rn(t,n){function e(){var e=this,r=n.apply(e,arguments);return r&&function(n){e.setAttributeNS(t.space,t.local,r(n))}}return e._value=n,e}function Ln(t,n){function e(){var e=this,r=n.apply(e,arguments);return r&&function(n){e.setAttribute(t,r(n))}}return e._value=n,e}function qn(t,n){return function(){xn(this,t).delay=+n.apply(this,arguments)}}function Un(t,n){return n=+n,function(){xn(this,t).delay=n}}function Dn(t,n){return function(){bn(this,t).duration=+n.apply(this,arguments)}}function On(t,n){return n=+n,function(){bn(this,t).duration=n}}function Fn(t,n){if("function"!=typeof n)throw new Error;return function(){bn(this,t).ease=n}}function In(t){return(t+"").trim().split(/^|\s+/).every(function(t){var n=t.indexOf(".");return n>=0&&(t=t.slice(0,n)),!t||"start"===t})}function Yn(t,n,e){var r,i,o=In(n)?xn:bn;return function(){var u=o(this,t),a=u.on;a!==r&&(i=(r=a).copy()).on(n,e),u.on=i}}function Bn(t){return function(){var n=this.parentNode;for(var e in this.__transition)if(+e!==t)return;n&&n.removeChild(this)}}function jn(t,n){var e,r,i;return function(){var o=B(this,t),u=(this.style.removeProperty(t),B(this,t));return o===u?null:o===e&&u===r?i:i=n(e=o,r=u)}}function Hn(t){return function(){this.style.removeProperty(t)}}function Xn(t,n,e){var r,i;return function(){var o=B(this,t);return o===e?null:o===r?i:i=n(r=o,e)}}function $n(t,n,e){var r,i,o;return function(){var u=B(this,t),a=e(this);return null==a&&(this.style.removeProperty(t),a=B(this,t)),u===a?null:u===r&&a===i?o:o=n(r=u,i=a)}}function Vn(t,n,e){function r(){var r=this,i=n.apply(r,arguments);return i&&function(n){r.style.setProperty(t,i(n),e)}}return r._value=n,r}function Wn(t){return function(){this.textContent=t}}function Zn(t){return function(){var n=t(this);this.textContent=null==n?"":n}}function Gn(t,n,e,r){this._groups=t,this._parents=n,this._name=e,this._id=r}function Jn(t){return dt().transition(t)}function Qn(){return++$l}function Kn(t){return((t*=2)<=1?t*t:--t*(2-t)+1)/2}function te(t){return((t*=2)<=1?t*t*t:(t-=2)*t*t+2)/2}function ne(t){return(1-Math.cos(Jl*t))/2}function ee(t){return((t*=2)<=1?Math.pow(2,10*t-10):2-Math.pow(2,10-10*t))/2}function re(t){return((t*=2)<=1?1-Math.sqrt(1-t*t):Math.sqrt(1-(t-=2)*t)+1)/2}function ie(t){return(t=+t)<Kl?ch*t*t:t<nh?ch*(t-=th)*t+eh:t<ih?ch*(t-=rh)*t+oh:ch*(t-=uh)*t+ah}function oe(t,n){for(var e;!(e=t.__transition)||!(e=e[n]);)if(!(t=t.parentNode))return _h.time=ln(),_h;return e}function ue(){t.event.stopImmediatePropagation()}function ae(t){return{type:t}}function ce(){return!t.event.button}function se(){var t=this.ownerSVGElement||this;return[[0,0],[t.width.baseVal.value,t.height.baseVal.value]]}function fe(t){for(;!t.__brush;)if(!(t=t.parentNode))return;return t.__brush}function le(t){return t[0][0]===t[1][0]||t[0][1]===t[1][1]}function he(n){function e(t){var e=t.property("__brush",a).selectAll(".overlay").data([ae("overlay")]);e.enter().append("rect").attr("class","overlay").attr("pointer-events","all").attr("cursor",Eh.overlay).merge(e).each(function(){var t=fe(this).extent;cf(this).attr("x",t[0][0]).attr("y",t[0][1]).attr("width",t[1][0]-t[0][0]).attr("height",t[1][1]-t[0][1])}),t.selectAll(".selection").data([ae("selection")]).enter().append("rect").attr("class","selection").attr("cursor",Eh.selection).attr("fill","#777").attr("fill-opacity",.3).attr("stroke","#fff").attr("shape-rendering","crispEdges");var i=t.selectAll(".handle").data(n.handles,function(t){return t.type});i.exit().remove(),i.enter().append("rect").attr("class",function(t){return"handle handle--"+t.type}).attr("cursor",function(t){return Eh[t.type]}),t.each(r).attr("fill","none").attr("pointer-events","all").style("-webkit-tap-highlight-color","rgba(0,0,0,0)").on("mousedown.brush touchstart.brush",u)}function r(){var t=cf(this),n=fe(this).selection;n?(t.selectAll(".selection").style("display",null).attr("x",n[0][0]).attr("y",n[0][1]).attr("width",n[1][0]-n[0][0]).attr("height",n[1][1]-n[0][1]),t.selectAll(".handle").style("display",null).attr("x",function(t){return"e"===t.type[t.type.length-1]?n[1][0]-p/2:n[0][0]-p/2}).attr("y",function(t){return"s"===t.type[0]?n[1][1]-p/2:n[0][1]-p/2}).attr("width",function(t){return"n"===t.type||"s"===t.type?n[1][0]-n[0][0]+p:p}).attr("height",function(t){return"e"===t.type||"w"===t.type?n[1][1]-n[0][1]+p:p})):t.selectAll(".selection,.handle").style("display","none").attr("x",null).attr("y",null).attr("width",null).attr("height",null)}function i(t,n){return t.__brush.emitter||new o(t,n)}function o(t,n){this.that=t,this.args=n,this.state=t.__brush,this.active=0}function u(){function e(){var t=Ks(w);!L||x||b||(Math.abs(t[0]-U[0])>Math.abs(t[1]-U[1])?b=!0:x=!0),U=t,m=!0,xh(),o()}function o(){var t;switch(y=U[0]-q[0],g=U[1]-q[1],T){case wh:case bh:k&&(y=Math.max(C-a,Math.min(P-p,y)),s=a+y,d=p+y),N&&(g=Math.max(z-l,Math.min(R-v,g)),h=l+g,_=v+g);break;case Mh:k<0?(y=Math.max(C-a,Math.min(P-a,y)),s=a+y,d=p):k>0&&(y=Math.max(C-p,Math.min(P-p,y)),s=a,d=p+y),N<0?(g=Math.max(z-l,Math.min(R-l,g)),h=l+g,_=v):N>0&&(g=Math.max(z-v,Math.min(R-v,g)),h=l,_=v+g);break;case Th:k&&(s=Math.max(C,Math.min(P,a-y*k)),d=Math.max(C,Math.min(P,p+y*k))),N&&(h=Math.max(z,Math.min(R,l-g*N)),_=Math.max(z,Math.min(R,v+g*N)))}d<s&&(k*=-1,t=a,a=p,p=t,t=s,s=d,d=t,M in Ah&&F.attr("cursor",Eh[M=Ah[M]])),_<h&&(N*=-1,t=l,l=v,v=t,t=h,h=_,_=t,M in Ch&&F.attr("cursor",Eh[M=Ch[M]])),S.selection&&(A=S.selection),x&&(s=A[0][0],d=A[1][0]),b&&(h=A[0][1],_=A[1][1]),A[0][0]===s&&A[0][1]===h&&A[1][0]===d&&A[1][1]===_||(S.selection=[[s,h],[d,_]],r.call(w),D.brush())}function u(){if(ue(),t.event.touches){if(t.event.touches.length)return;c&&clearTimeout(c),c=setTimeout(function(){c=null},500),O.on("touchmove.brush touchend.brush touchcancel.brush",null)}else _t(t.event.view,m),I.on("keydown.brush keyup.brush mousemove.brush mouseup.brush",null);O.attr("pointer-events","all"),F.attr("cursor",Eh.overlay),S.selection&&(A=S.selection),le(A)&&(S.selection=null,r.call(w)),D.end()}if(t.event.touches){if(t.event.changedTouches.length<t.event.touches.length)return xh()}else if(c)return;if(f.apply(this,arguments)){var a,s,l,h,p,d,v,_,y,g,m,x,b,w=this,M=t.event.target.__data__.type,T="selection"===(t.event.metaKey?M="overlay":M)?bh:t.event.altKey?Th:Mh,k=n===Nh?null:zh[M],N=n===kh?null:Ph[M],S=fe(w),E=S.extent,A=S.selection,C=E[0][0],z=E[0][1],P=E[1][0],R=E[1][1],L=k&&N&&t.event.shiftKey,q=Ks(w),U=q,D=i(w,arguments).beforestart();"overlay"===M?S.selection=A=[[a=n===Nh?C:q[0],l=n===kh?z:q[1]],[p=n===Nh?P:a,v=n===kh?R:l]]:(a=A[0][0],l=A[0][1],p=A[1][0],v=A[1][1]),s=a,h=l,d=p,_=v;var O=cf(w).attr("pointer-events","none"),F=O.selectAll(".overlay").attr("cursor",Eh[M]);if(t.event.touches)O.on("touchmove.brush",e,!0).on("touchend.brush touchcancel.brush",u,!0);else{var I=cf(t.event.view).on("keydown.brush",function(){switch(t.event.keyCode){case 16:L=k&&N;break;case 18:T===Mh&&(k&&(p=d-y*k,a=s+y*k),N&&(v=_-g*N,l=h+g*N),T=Th,o());break;case 32:T!==Mh&&T!==Th||(k<0?p=d-y:k>0&&(a=s-y),N<0?v=_-g:N>0&&(l=h-g),T=wh,F.attr("cursor",Eh.selection),o());break;default:return}xh()},!0).on("keyup.brush",function(){switch(t.event.keyCode){case 16:L&&(x=b=L=!1,o());break;case 18:T===Th&&(k<0?p=d:k>0&&(a=s),N<0?v=_:N>0&&(l=h),T=Mh,o());break;case 32:T===wh&&(t.event.altKey?(k&&(p=d-y*k,a=s+y*k),N&&(v=_-g*N,l=h+g*N),T=Th):(k<0?p=d:k>0&&(a=s),N<0?v=_:N>0&&(l=h),T=Mh),F.attr("cursor",Eh[M]),o());break;default:return}xh()},!0).on("mousemove.brush",e,!0).on("mouseup.brush",u,!0);lf(t.event.view)}ue(),jl(w),r.call(w),D.start()}}function a(){var t=this.__brush||{selection:null};return t.extent=s.apply(this,arguments),t.dim=n,t}var c,s=se,f=ce,l=h(e,"start","brush","end"),p=6;return e.move=function(t,e){t.selection?t.on("start.brush",function(){i(this,arguments).beforestart().start()}).on("interrupt.brush end.brush",function(){i(this,arguments).end()}).tween("brush",function(){function t(t){u.selection=1===t&&le(s)?null:f(t),r.call(o),a.brush()}var o=this,u=o.__brush,a=i(o,arguments),c=u.selection,s=n.input("function"==typeof e?e.apply(this,arguments):e,u.extent),f=cl(c,s);return c&&s?t:t(1)}):t.each(function(){var t=this,o=arguments,u=t.__brush,a=n.input("function"==typeof e?e.apply(t,o):e,u.extent),c=i(t,o).beforestart();jl(t),u.selection=null==a||le(a)?null:a,r.call(t),c.start().brush().end()})},o.prototype={beforestart:function(){return 1==++this.active&&(this.state.emitter=this,this.starting=!0),this},start:function(){return this.starting&&(this.starting=!1,this.emit("start")),this},brush:function(){return this.emit("brush"),this},end:function(){return 0==--this.active&&(delete this.state.emitter,this.emit("end")),this},emit:function(t){N(new mh(e,t,n.output(this.state.selection)),l.apply,l,[t,this.that,this.args])}},e.extent=function(t){return arguments.length?(s="function"==typeof t?t:gh([[+t[0][0],+t[0][1]],[+t[1][0],+t[1][1]]]),e):s},e.filter=function(t){return arguments.length?(f="function"==typeof t?t:gh(!!t),e):f},e.handleSize=function(t){return arguments.length?(p=+t,e):p},e.on=function(){var t=l.on.apply(l,arguments);return t===l?e:t},e}function pe(t){return function(n,e){return t(n.source.value+n.target.value,e.source.value+e.target.value)}}function de(){this._x0=this._y0=this._x1=this._y1=null,this._=""}function ve(){return new de}function _e(t){return t.source}function ye(t){return t.target}function ge(t){return t.radius}function me(t){return t.startAngle}function xe(t){return t.endAngle}function be(){}function we(t,n){var e=new be;if(t instanceof be)t.each(function(t,n){e.set(n,t)});else if(Array.isArray(t)){var r,i=-1,o=t.length;if(null==n)for(;++i<o;)e.set(i,t[i]);else for(;++i<o;)e.set(n(r=t[i],i,t),r)}else if(t)for(var u in t)e.set(u,t[u]);return e}function Me(){return{}}function Te(t,n,e){t[n]=e}function ke(){return we()}function Ne(t,n,e){t.set(n,e)}function Se(){}function Ee(t,n){var e=new Se;if(t instanceof Se)t.each(function(t){e.add(t)});else if(t){var r=-1,i=t.length;if(null==n)for(;++r<i;)e.add(t[r]);else for(;++r<i;)e.add(n(t[r],r,t))}return e}function Ae(t){return new Function("d","return {"+t.map(function(t,n){return JSON.stringify(t)+": d["+n+"]"}).join(",")+"}")}function Ce(t,n){var e=Ae(t);return function(r,i){return n(e(r),i,t)}}function ze(t){var n=Object.create(null),e=[];return t.forEach(function(t){for(var r in t)r in n||e.push(n[r]=r)}),e}function Pe(t,n,e,r){if(isNaN(n)||isNaN(e))return t;var i,o,u,a,c,s,f,l,h,p=t._root,d={data:r},v=t._x0,_=t._y0,y=t._x1,g=t._y1;if(!p)return t._root=d,t;for(;p.length;)if((s=n>=(o=(v+y)/2))?v=o:y=o,(f=e>=(u=(_+g)/2))?_=u:g=u,i=p,!(p=p[l=f<<1|s]))return i[l]=d,t;if(a=+t._x.call(null,p.data),c=+t._y.call(null,p.data),n===a&&e===c)return d.next=p,i?i[l]=d:t._root=d,t;do{i=i?i[l]=new Array(4):t._root=new Array(4),(s=n>=(o=(v+y)/2))?v=o:y=o,(f=e>=(u=(_+g)/2))?_=u:g=u}while((l=f<<1|s)==(h=(c>=u)<<1|a>=o));return i[h]=p,i[l]=d,t}function Re(t){return t[0]}function Le(t){return t[1]}function qe(t,n,e){var r=new Ue(null==n?Re:n,null==e?Le:e,NaN,NaN,NaN,NaN);return null==t?r:r.addAll(t)}function Ue(t,n,e,r,i,o){this._x=t,this._y=n,this._x0=e,this._y0=r,this._x1=i,this._y1=o,this._root=void 0}function De(t){for(var n={data:t.data},e=n;t=t.next;)e=e.next={data:t.data};return n}function Oe(t){return t.x+t.vx}function Fe(t){return t.y+t.vy}function Ie(t){return t.index}function Ye(t,n){var e=t.get(n);if(!e)throw new Error("missing: "+n);return e}function Be(t){return t.x}function je(t){return t.y}function He(t){return new Xe(t)}function Xe(t){if(!(n=vp.exec(t)))throw new Error("invalid format: "+t);var n,e=n[1]||" ",r=n[2]||">",i=n[3]||"-",o=n[4]||"",u=!!n[5],a=n[6]&&+n[6],c=!!n[7],s=n[8]&&+n[8].slice(1),f=n[9]||"";"n"===f?(c=!0,f="g"):dp[f]||(f=""),(u||"0"===e&&"="===r)&&(u=!0,e="0",r="="),this.fill=e,this.align=r,this.sign=i,this.symbol=o,this.zero=u,this.width=a,this.comma=c,this.precision=s,this.type=f}function $e(n){return _p=mp(n),t.format=_p.format,t.formatPrefix=_p.formatPrefix,_p}function Ve(){this.reset()}function We(t,n,e){var r=t.s=n+e,i=r-n,o=r-i;t.t=n-o+(e-i)}function Ze(t){return t>1?0:t<-1?rd:Math.acos(t)}function Ge(t){return t>1?id:t<-1?-id:Math.asin(t)}function Je(t){return(t=yd(t/2))*t}function Qe(){}function Ke(t,n){t&&wd.hasOwnProperty(t.type)&&wd[t.type](t,n)}function tr(t,n,e){var r,i=-1,o=t.length-e;for(n.lineStart();++i<o;)r=t[i],n.point(r[0],r[1],r[2]);n.lineEnd()}function nr(t,n){var e=-1,r=t.length;for(n.polygonStart();++e<r;)tr(t[e],n,1);n.polygonEnd()}function er(){Nd.point=ir}function rr(){or(Tp,kp)}function ir(t,n){Nd.point=or,Tp=t,kp=n,Np=t*=cd,Sp=hd(n=(n*=cd)/2+od),Ep=yd(n)}function or(t,n){n=(n*=cd)/2+od;var e=(t*=cd)-Np,r=e>=0?1:-1,i=r*e,o=hd(n),u=yd(n),a=Ep*u,c=Sp*o+a*hd(i),s=a*r*yd(i);Td.add(ld(s,c)),Np=t,Sp=o,Ep=u}function ur(t){return[ld(t[1],t[0]),Ge(t[2])]}function ar(t){var n=t[0],e=t[1],r=hd(e);return[r*hd(n),r*yd(n),yd(e)]}function cr(t,n){return t[0]*n[0]+t[1]*n[1]+t[2]*n[2]}function sr(t,n){return[t[1]*n[2]-t[2]*n[1],t[2]*n[0]-t[0]*n[2],t[0]*n[1]-t[1]*n[0]]}function fr(t,n){t[0]+=n[0],t[1]+=n[1],t[2]+=n[2]}function lr(t,n){return[t[0]*n,t[1]*n,t[2]*n]}function hr(t){var n=md(t[0]*t[0]+t[1]*t[1]+t[2]*t[2]);t[0]/=n,t[1]/=n,t[2]/=n}function pr(t,n){Dp.push(Op=[Ap=t,zp=t]),n<Cp&&(Cp=n),n>Pp&&(Pp=n)}function dr(t,n){var e=ar([t*cd,n*cd]);if(Up){var r=sr(Up,e),i=sr([r[1],-r[0],0],r);hr(i),i=ur(i);var o,u=t-Rp,a=u>0?1:-1,c=i[0]*ad*a,s=sd(u)>180;s^(a*Rp<c&&c<a*t)?(o=i[1]*ad)>Pp&&(Pp=o):(c=(c+360)%360-180,s^(a*Rp<c&&c<a*t)?(o=-i[1]*ad)<Cp&&(Cp=o):(n<Cp&&(Cp=n),n>Pp&&(Pp=n))),s?t<Rp?xr(Ap,t)>xr(Ap,zp)&&(zp=t):xr(t,zp)>xr(Ap,zp)&&(Ap=t):zp>=Ap?(t<Ap&&(Ap=t),t>zp&&(zp=t)):t>Rp?xr(Ap,t)>xr(Ap,zp)&&(zp=t):xr(t,zp)>xr(Ap,zp)&&(Ap=t)}else Dp.push(Op=[Ap=t,zp=t]);n<Cp&&(Cp=n),n>Pp&&(Pp=n),Up=e,Rp=t}function vr(){Ed.point=dr}function _r(){Op[0]=Ap,Op[1]=zp,Ed.point=pr,Up=null}function yr(t,n){if(Up){var e=t-Rp;Sd.add(sd(e)>180?e+(e>0?360:-360):e)}else Lp=t,qp=n;Nd.point(t,n),dr(t,n)}function gr(){Nd.lineStart()}function mr(){yr(Lp,qp),Nd.lineEnd(),sd(Sd)>ed&&(Ap=-(zp=180)),Op[0]=Ap,Op[1]=zp,Up=null}function xr(t,n){return(n-=t)<0?n+360:n}function br(t,n){return t[0]-n[0]}function wr(t,n){return t[0]<=t[1]?t[0]<=n&&n<=t[1]:n<t[0]||t[1]<n}function Mr(t,n){t*=cd;var e=hd(n*=cd);Tr(e*hd(t),e*yd(t),yd(n))}function Tr(t,n,e){Yp+=(t-Yp)/++Fp,Bp+=(n-Bp)/Fp,jp+=(e-jp)/Fp}function kr(){Ad.point=Nr}function Nr(t,n){t*=cd;var e=hd(n*=cd);Qp=e*hd(t),Kp=e*yd(t),td=yd(n),Ad.point=Sr,Tr(Qp,Kp,td)}function Sr(t,n){t*=cd;var e=hd(n*=cd),r=e*hd(t),i=e*yd(t),o=yd(n),u=ld(md((u=Kp*o-td*i)*u+(u=td*r-Qp*o)*u+(u=Qp*i-Kp*r)*u),Qp*r+Kp*i+td*o);Ip+=u,Hp+=u*(Qp+(Qp=r)),Xp+=u*(Kp+(Kp=i)),$p+=u*(td+(td=o)),Tr(Qp,Kp,td)}function Er(){Ad.point=Mr}function Ar(){Ad.point=zr}function Cr(){Pr(Gp,Jp),Ad.point=Mr}function zr(t,n){Gp=t,Jp=n,t*=cd,n*=cd,Ad.point=Pr;var e=hd(n);Qp=e*hd(t),Kp=e*yd(t),td=yd(n),Tr(Qp,Kp,td)}function Pr(t,n){t*=cd;var e=hd(n*=cd),r=e*hd(t),i=e*yd(t),o=yd(n),u=Kp*o-td*i,a=td*r-Qp*o,c=Qp*i-Kp*r,s=md(u*u+a*a+c*c),f=Ge(s),l=s&&-f/s;Vp+=l*u,Wp+=l*a,Zp+=l*c,Ip+=f,Hp+=f*(Qp+(Qp=r)),Xp+=f*(Kp+(Kp=i)),$p+=f*(td+(td=o)),Tr(Qp,Kp,td)}function Rr(t,n){return[t>rd?t-ud:t<-rd?t+ud:t,n]}function Lr(t,n,e){return(t%=ud)?n||e?zd(Ur(t),Dr(n,e)):Ur(t):n||e?Dr(n,e):Rr}function qr(t){return function(n,e){return n+=t,[n>rd?n-ud:n<-rd?n+ud:n,e]}}function Ur(t){var n=qr(t);return n.invert=qr(-t),n}function Dr(t,n){function e(t,n){var e=hd(n),a=hd(t)*e,c=yd(t)*e,s=yd(n),f=s*r+a*i;return[ld(c*o-f*u,a*r-s*i),Ge(f*o+c*u)]}var r=hd(t),i=yd(t),o=hd(n),u=yd(n);return e.invert=function(t,n){var e=hd(n),a=hd(t)*e,c=yd(t)*e,s=yd(n),f=s*o-c*u;return[ld(c*o+s*u,a*r+f*i),Ge(f*r-a*i)]},e}function Or(t,n,e,r,i,o){if(e){var u=hd(n),a=yd(n),c=r*e;null==i?(i=n+r*ud,o=n-c/2):(i=Fr(u,i),o=Fr(u,o),(r>0?i<o:i>o)&&(i+=r*ud));for(var s,f=i;r>0?f>o:f<o;f-=c)s=ur([u,-a*hd(f),-a*yd(f)]),t.point(s[0],s[1])}}function Fr(t,n){(n=ar(n))[0]-=t,hr(n);var e=Ze(-n[1]);return((-n[2]<0?-e:e)+ud-ed)%ud}function Ir(t,n,e,r){this.x=t,this.z=n,this.o=e,this.e=r,this.v=!1,this.n=this.p=null}function Yr(t){if(n=t.length){for(var n,e,r=0,i=t[0];++r<n;)i.n=e=t[r],e.p=i,i=e;i.n=e=t[0],e.p=i}}function Br(t,n,e,r){function i(i,o){return t<=i&&i<=e&&n<=o&&o<=r}function o(i,o,a,s){var f=0,l=0;if(null==i||(f=u(i,a))!==(l=u(o,a))||c(i,o)<0^a>0)do{s.point(0===f||3===f?t:e,f>1?r:n)}while((f=(f+a+4)%4)!==l);else s.point(o[0],o[1])}function u(r,i){return sd(r[0]-t)<ed?i>0?0:3:sd(r[0]-e)<ed?i>0?2:1:sd(r[1]-n)<ed?i>0?1:0:i>0?3:2}function a(t,n){return c(t.x,n.x)}function c(t,n){var e=u(t,1),r=u(n,1);return e!==r?e-r:0===e?n[1]-t[1]:1===e?t[0]-n[0]:2===e?t[1]-n[1]:n[0]-t[0]}return function(u){function c(t,n){i(t,n)&&w.point(t,n)}function s(){for(var n=0,e=0,i=h.length;e<i;++e)for(var o,u,a=h[e],c=1,s=a.length,f=a[0],l=f[0],p=f[1];c<s;++c)o=l,u=p,l=(f=a[c])[0],p=f[1],u<=r?p>r&&(l-o)*(r-u)>(p-u)*(t-o)&&++n:p<=r&&(l-o)*(r-u)<(p-u)*(t-o)&&--n;return n}function f(o,u){var a=i(o,u);if(h&&p.push([o,u]),x)d=o,v=u,_=a,x=!1,a&&(w.lineStart(),w.point(o,u));else if(a&&m)w.point(o,u);else{var c=[y=Math.max(Zd,Math.min(Wd,y)),g=Math.max(Zd,Math.min(Wd,g))],s=[o=Math.max(Zd,Math.min(Wd,o)),u=Math.max(Zd,Math.min(Wd,u))];Xd(c,s,t,n,e,r)?(m||(w.lineStart(),w.point(c[0],c[1])),w.point(s[0],s[1]),a||w.lineEnd(),b=!1):a&&(w.lineStart(),w.point(o,u),b=!1)}y=o,g=u,m=a}var l,h,p,d,v,_,y,g,m,x,b,w=u,M=Hd(),T={point:c,lineStart:function(){T.point=f,h&&h.push(p=[]),x=!0,m=!1,y=g=NaN},lineEnd:function(){l&&(f(d,v),_&&m&&M.rejoin(),l.push(M.result())),T.point=c,m&&w.lineEnd()},polygonStart:function(){w=M,l=[],h=[],b=!0},polygonEnd:function(){var t=s(),n=b&&t,e=(l=Cs(l)).length;(n||e)&&(u.polygonStart(),n&&(u.lineStart(),o(null,null,1,u),u.lineEnd()),e&&Vd(l,a,t,o,u),u.polygonEnd()),w=u,l=h=p=null}};return T}}function jr(){Kd.point=Kd.lineEnd=Qe}function Hr(t,n){Pd=t*=cd,Rd=yd(n*=cd),Ld=hd(n),Kd.point=Xr}function Xr(t,n){t*=cd;var e=yd(n*=cd),r=hd(n),i=sd(t-Pd),o=hd(i),u=r*yd(i),a=Ld*e-Rd*r*o,c=Rd*e+Ld*r*o;Qd.add(ld(md(u*u+a*a),c)),Pd=t,Rd=e,Ld=r}function $r(t,n){return!(!t||!ov.hasOwnProperty(t.type))&&ov[t.type](t,n)}function Vr(t,n){return 0===rv(t,n)}function Wr(t,n){var e=rv(t[0],t[1]);return rv(t[0],n)+rv(n,t[1])<=e+ed}function Zr(t,n){return!!Jd(t.map(Gr),Jr(n))}function Gr(t){return(t=t.map(Jr)).pop(),t}function Jr(t){return[t[0]*cd,t[1]*cd]}function Qr(t,n,e){var r=Ms(t,n-ed,e).concat(n);return function(t){return r.map(function(n){return[t,n]})}}function Kr(t,n,e){var r=Ms(t,n-ed,e).concat(n);return function(t){return r.map(function(n){return[n,t]})}}function ti(){function t(){return{type:"MultiLineString",coordinates:n()}}function n(){return Ms(pd(o/_)*_,i,_).map(h).concat(Ms(pd(s/y)*y,c,y).map(p)).concat(Ms(pd(r/d)*d,e,d).filter(function(t){return sd(t%_)>ed}).map(f)).concat(Ms(pd(a/v)*v,u,v).filter(function(t){return sd(t%y)>ed}).map(l))}var e,r,i,o,u,a,c,s,f,l,h,p,d=10,v=d,_=90,y=360,g=2.5;return t.lines=function(){return n().map(function(t){return{type:"LineString",coordinates:t}})},t.outline=function(){return{type:"Polygon",coordinates:[h(o).concat(p(c).slice(1),h(i).reverse().slice(1),p(s).reverse().slice(1))]}},t.extent=function(n){return arguments.length?t.extentMajor(n).extentMinor(n):t.extentMinor()},t.extentMajor=function(n){return arguments.length?(o=+n[0][0],i=+n[1][0],s=+n[0][1],c=+n[1][1],o>i&&(n=o,o=i,i=n),s>c&&(n=s,s=c,c=n),t.precision(g)):[[o,s],[i,c]]},t.extentMinor=function(n){return arguments.length?(r=+n[0][0],e=+n[1][0],a=+n[0][1],u=+n[1][1],r>e&&(n=r,r=e,e=n),a>u&&(n=a,a=u,u=n),t.precision(g)):[[r,a],[e,u]]},t.step=function(n){return arguments.length?t.stepMajor(n).stepMinor(n):t.stepMinor()},t.stepMajor=function(n){return arguments.length?(_=+n[0],y=+n[1],t):[_,y]},t.stepMinor=function(n){return arguments.length?(d=+n[0],v=+n[1],t):[d,v]},t.precision=function(n){return arguments.length?(g=+n,f=Qr(a,u,90),l=Kr(r,e,g),h=Qr(s,c,90),p=Kr(o,i,g),t):g},t.extentMajor([[-180,-90+ed],[180,90-ed]]).extentMinor([[-180,-80-ed],[180,80+ed]])}function ni(){sv.point=ei}function ei(t,n){sv.point=ri,qd=Dd=t,Ud=Od=n}function ri(t,n){cv.add(Od*t-Dd*n),Dd=t,Od=n}function ii(){ri(qd,Ud)}function oi(t,n){vv+=t,_v+=n,++yv}function ui(){Tv.point=ai}function ai(t,n){Tv.point=ci,oi(Yd=t,Bd=n)}function ci(t,n){var e=t-Yd,r=n-Bd,i=md(e*e+r*r);gv+=i*(Yd+t)/2,mv+=i*(Bd+n)/2,xv+=i,oi(Yd=t,Bd=n)}function si(){Tv.point=oi}function fi(){Tv.point=hi}function li(){pi(Fd,Id)}function hi(t,n){Tv.point=pi,oi(Fd=Yd=t,Id=Bd=n)}function pi(t,n){var e=t-Yd,r=n-Bd,i=md(e*e+r*r);gv+=i*(Yd+t)/2,mv+=i*(Bd+n)/2,xv+=i,bv+=(i=Bd*t-Yd*n)*(Yd+t),wv+=i*(Bd+n),Mv+=3*i,oi(Yd=t,Bd=n)}function di(t){this._context=t}function vi(t,n){zv.point=_i,Nv=Ev=t,Sv=Av=n}function _i(t,n){Ev-=t,Av-=n,Cv.add(md(Ev*Ev+Av*Av)),Ev=t,Av=n}function yi(){this._string=[]}function gi(t){return"m0,"+t+"a"+t+","+t+" 0 1,1 0,"+-2*t+"a"+t+","+t+" 0 1,1 0,"+2*t+"z"}function mi(t){return t.length>1}function xi(t,n){return((t=t.x)[0]<0?t[1]-id-ed:id-t[1])-((n=n.x)[0]<0?n[1]-id-ed:id-n[1])}function bi(t,n,e,r){var i,o,u=yd(t-e);return sd(u)>ed?fd((yd(n)*(o=hd(r))*yd(e)-yd(r)*(i=hd(n))*yd(t))/(i*o*u)):(n+r)/2}function wi(t){return function(n){var e=new Mi;for(var r in t)e[r]=t[r];return e.stream=n,e}}function Mi(){}function Ti(t,n,e){var r=n[1][0]-n[0][0],i=n[1][1]-n[0][1],o=t.clipExtent&&t.clipExtent();t.scale(150).translate([0,0]),null!=o&&t.clipExtent(null),Md(e,t.stream(dv));var u=dv.result(),a=Math.min(r/(u[1][0]-u[0][0]),i/(u[1][1]-u[0][1])),c=+n[0][0]+(r-a*(u[1][0]+u[0][0]))/2,s=+n[0][1]+(i-a*(u[1][1]+u[0][1]))/2;return null!=o&&t.clipExtent(o),t.scale(150*a).translate([c,s])}function ki(t,n,e){return Ti(t,[[0,0],n],e)}function Ni(t){return wi({point:function(n,e){n=t(n,e),this.stream.point(n[0],n[1])}})}function Si(t,n){function e(r,i,o,u,a,c,s,f,l,h,p,d,v,_){var y=s-r,g=f-i,m=y*y+g*g;if(m>4*n&&v--){var x=u+h,b=a+p,w=c+d,M=md(x*x+b*b+w*w),T=Ge(w/=M),k=sd(sd(w)-1)<ed||sd(o-l)<ed?(o+l)/2:ld(b,x),N=t(k,T),S=N[0],E=N[1],A=S-r,C=E-i,z=g*A-y*C;(z*z/m>n||sd((y*A+g*C)/m-.5)>.3||u*h+a*p+c*d<Uv)&&(e(r,i,o,u,a,c,S,E,k,x/=M,b/=M,w,v,_),_.point(S,E),e(S,E,k,x,b,w,s,f,l,h,p,d,v,_))}}return function(n){function r(e,r){e=t(e,r),n.point(e[0],e[1])}function i(){y=NaN,w.point=o,n.lineStart()}function o(r,i){var o=ar([r,i]),u=t(r,i);e(y,g,_,m,x,b,y=u[0],g=u[1],_=r,m=o[0],x=o[1],b=o[2],qv,n),n.point(y,g)}function u(){w.point=r,n.lineEnd()}function a(){i(),w.point=c,w.lineEnd=s}function c(t,n){o(f=t,n),l=y,h=g,p=m,d=x,v=b,w.point=o}function s(){e(y,g,_,m,x,b,l,h,f,p,d,v,qv,n),w.lineEnd=u,u()}var f,l,h,p,d,v,_,y,g,m,x,b,w={point:r,lineStart:i,lineEnd:u,polygonStart:function(){n.polygonStart(),w.lineStart=a},polygonEnd:function(){n.polygonEnd(),w.lineStart=i}};return w}}function Ei(t){return Ai(function(){return t})()}function Ai(t){function n(t){return t=f(t[0]*cd,t[1]*cd),[t[0]*_+a,c-t[1]*_]}function e(t){return(t=f.invert((t[0]-a)/_,(c-t[1])/_))&&[t[0]*ad,t[1]*ad]}function r(t,n){return t=u(t,n),[t[0]*_+a,c-t[1]*_]}function i(){f=zd(s=Lr(b,w,M),u);var t=u(m,x);return a=y-t[0]*_,c=g+t[1]*_,o()}function o(){return d=v=null,n}var u,a,c,s,f,l,h,p,d,v,_=150,y=480,g=250,m=0,x=0,b=0,w=0,M=0,T=null,k=Rv,N=null,S=uv,E=.5,A=Dv(r,E);return n.stream=function(t){return d&&v===t?d:d=Ov(k(s,A(S(v=t))))},n.clipAngle=function(t){return arguments.length?(k=+t?Lv(T=t*cd,6*cd):(T=null,Rv),o()):T*ad},n.clipExtent=function(t){return arguments.length?(S=null==t?(N=l=h=p=null,uv):Br(N=+t[0][0],l=+t[0][1],h=+t[1][0],p=+t[1][1]),o()):null==N?null:[[N,l],[h,p]]},n.scale=function(t){return arguments.length?(_=+t,i()):_},n.translate=function(t){return arguments.length?(y=+t[0],g=+t[1],i()):[y,g]},n.center=function(t){return arguments.length?(m=t[0]%360*cd,x=t[1]%360*cd,i()):[m*ad,x*ad]},n.rotate=function(t){return arguments.length?(b=t[0]%360*cd,w=t[1]%360*cd,M=t.length>2?t[2]%360*cd:0,i()):[b*ad,w*ad,M*ad]},n.precision=function(t){return arguments.length?(A=Dv(r,E=t*t),o()):md(E)},n.fitExtent=function(t,e){return Ti(n,t,e)},n.fitSize=function(t,e){return ki(n,t,e)},function(){return u=t.apply(this,arguments),n.invert=u.invert&&e,i()}}function Ci(t){var n=0,e=rd/3,r=Ai(t),i=r(n,e);return i.parallels=function(t){return arguments.length?r(n=t[0]*cd,e=t[1]*cd):[n*ad,e*ad]},i}function zi(t){function n(t,n){return[t*e,yd(n)/e]}var e=hd(t);return n.invert=function(t,n){return[t/e,Ge(n*e)]},n}function Pi(t,n){function e(t,n){var e=md(o-2*i*yd(n))/i;return[e*yd(t*=i),u-e*hd(t)]}var r=yd(t),i=(r+yd(n))/2;if(sd(i)<ed)return zi(t);var o=1+r*(2*i-r),u=md(o)/i;return e.invert=function(t,n){var e=u-n;return[ld(t,sd(e))/i*gd(e),Ge((o-(t*t+e*e)*i*i)/(2*i))]},e}function Ri(t){var n=t.length;return{point:function(e,r){for(var i=-1;++i<n;)t[i].point(e,r)},sphere:function(){for(var e=-1;++e<n;)t[e].sphere()},lineStart:function(){for(var e=-1;++e<n;)t[e].lineStart()},lineEnd:function(){for(var e=-1;++e<n;)t[e].lineEnd()},polygonStart:function(){for(var e=-1;++e<n;)t[e].polygonStart()},polygonEnd:function(){for(var e=-1;++e<n;)t[e].polygonEnd()}}}function Li(t){return function(n,e){var r=hd(n),i=hd(e),o=t(r*i);return[o*i*yd(n),o*yd(e)]}}function qi(t){return function(n,e){var r=md(n*n+e*e),i=t(r),o=yd(i),u=hd(i);return[ld(n*o,r*u),Ge(r&&e*o/r)]}}function Ui(t,n){return[t,vd(xd((id+n)/2))]}function Di(t){function n(){var n=rd*a(),u=o(jd(o.rotate()).invert([0,0]));return s(null==f?[[u[0]-n,u[1]-n],[u[0]+n,u[1]+n]]:t===Ui?[[Math.max(u[0]-n,f),e],[Math.min(u[0]+n,r),i]]:[[f,Math.max(u[1]-n,e)],[r,Math.min(u[1]+n,i)]])}var e,r,i,o=Ei(t),u=o.center,a=o.scale,c=o.translate,s=o.clipExtent,f=null;return o.scale=function(t){return arguments.length?(a(t),n()):a()},o.translate=function(t){return arguments.length?(c(t),n()):c()},o.center=function(t){return arguments.length?(u(t),n()):u()},o.clipExtent=function(t){return arguments.length?(null==t?f=e=r=i=null:(f=+t[0][0],e=+t[0][1],r=+t[1][0],i=+t[1][1]),n()):null==f?null:[[f,e],[r,i]]},n()}function Oi(t){return xd((id+t)/2)}function Fi(t,n){function e(t,n){o>0?n<-id+ed&&(n=-id+ed):n>id-ed&&(n=id-ed);var e=o/_d(Oi(n),i);return[e*yd(i*t),o-e*hd(i*t)]}var r=hd(t),i=t===n?yd(t):vd(r/hd(n))/vd(Oi(n)/Oi(t)),o=r*_d(Oi(t),i)/i;return i?(e.invert=function(t,n){var e=o-n,r=gd(i)*md(t*t+e*e);return[ld(t,sd(e))/i*gd(e),2*fd(_d(o/r,1/i))-id]},e):Ui}function Ii(t,n){return[t,n]}function Yi(t,n){function e(t,n){var e=o-n,r=i*t;return[e*yd(r),o-e*hd(r)]}var r=hd(t),i=t===n?yd(t):(r-hd(n))/(n-t),o=r/i+t;return sd(i)<ed?Ii:(e.invert=function(t,n){var e=o-n;return[ld(t,sd(e))/i*gd(e),o-gd(i)*md(t*t+e*e)]},e)}function Bi(t,n){var e=hd(n),r=hd(t)*e;return[e*yd(t)/r,yd(n)/r]}function ji(t,n,e,r){return 1===t&&1===n&&0===e&&0===r?uv:wi({point:function(i,o){this.stream.point(i*t+e,o*n+r)}})}function Hi(t,n){return[hd(n)*yd(t),yd(n)]}function Xi(t,n){var e=hd(n),r=1+hd(t)*e;return[e*yd(t)/r,yd(n)/r]}function $i(t,n){return[vd(xd((id+n)/2)),-t]}function Vi(t,n){return t.parent===n.parent?1:2}function Wi(t){return t.reduce(Zi,0)/t.length}function Zi(t,n){return t+n.x}function Gi(t){return 1+t.reduce(Ji,0)}function Ji(t,n){return Math.max(t,n.y)}function Qi(t){for(var n;n=t.children;)t=n[0];return t}function Ki(t){for(var n;n=t.children;)t=n[n.length-1];return t}function to(t){var n=0,e=t.children,r=e&&e.length;if(r)for(;--r>=0;)n+=e[r].value;else n=1;t.value=n}function no(t,n){if(t===n)return t;var e=t.ancestors(),r=n.ancestors(),i=null;for(t=e.pop(),n=r.pop();t===n;)i=t,t=e.pop(),n=r.pop();return i}function eo(t,n){var e,r,i,o,u,a=new uo(t),c=+t.value&&(a.value=t.value),s=[a];for(null==n&&(n=ro);e=s.pop();)if(c&&(e.value=+e.data.value),(i=n(e.data))&&(u=i.length))for(e.children=new Array(u),o=u-1;o>=0;--o)s.push(r=e.children[o]=new uo(i[o])),r.parent=e,r.depth=e.depth+1;return a.eachBefore(oo)}function ro(t){return t.children}function io(t){t.data=t.data.data}function oo(t){var n=0;do{t.height=n}while((t=t.parent)&&t.height<++n)}function uo(t){this.data=t,this.depth=this.height=0,this.parent=null}function ao(t){for(var n,e,r=t.length;r;)e=Math.random()*r--|0,n=t[r],t[r]=t[e],t[e]=n;return t}function co(t,n){var e,r;if(lo(n,t))return[n];for(e=0;e<t.length;++e)if(so(n,t[e])&&lo(vo(t[e],n),t))return[t[e],n];for(e=0;e<t.length-1;++e)for(r=e+1;r<t.length;++r)if(so(vo(t[e],t[r]),n)&&so(vo(t[e],n),t[r])&&so(vo(t[r],n),t[e])&&lo(_o(t[e],t[r],n),t))return[t[e],t[r],n];throw new Error}function so(t,n){var e=t.r-n.r,r=n.x-t.x,i=n.y-t.y;return e<0||e*e<r*r+i*i}function fo(t,n){var e=t.r-n.r+1e-6,r=n.x-t.x,i=n.y-t.y;return e>0&&e*e>r*r+i*i}function lo(t,n){for(var e=0;e<n.length;++e)if(!fo(t,n[e]))return!1;return!0}function ho(t){switch(t.length){case 1:return po(t[0]);case 2:return vo(t[0],t[1]);case 3:return _o(t[0],t[1],t[2])}}function po(t){return{x:t.x,y:t.y,r:t.r}}function vo(t,n){var e=t.x,r=t.y,i=t.r,o=n.x,u=n.y,a=n.r,c=o-e,s=u-r,f=a-i,l=Math.sqrt(c*c+s*s);return{x:(e+o+c/l*f)/2,y:(r+u+s/l*f)/2,r:(l+i+a)/2}}function _o(t,n,e){var r=t.x,i=t.y,o=t.r,u=n.x,a=n.y,c=n.r,s=e.x,f=e.y,l=e.r,h=r-u,p=r-s,d=i-a,v=i-f,_=c-o,y=l-o,g=r*r+i*i-o*o,m=g-u*u-a*a+c*c,x=g-s*s-f*f+l*l,b=p*d-h*v,w=(d*x-v*m)/(2*b)-r,M=(v*_-d*y)/b,T=(p*m-h*x)/(2*b)-i,k=(h*y-p*_)/b,N=M*M+k*k-1,S=2*(o+w*M+T*k),E=w*w+T*T-o*o,A=-(N?(S+Math.sqrt(S*S-4*N*E))/(2*N):E/S);return{x:r+w+M*A,y:i+T+k*A,r:A}}function yo(t,n,e){var r=t.x,i=t.y,o=n.r+e.r,u=t.r+e.r,a=n.x-r,c=n.y-i,s=a*a+c*c;if(s){var f=.5+((u*=u)-(o*=o))/(2*s),l=Math.sqrt(Math.max(0,2*o*(u+s)-(u-=s)*u-o*o))/(2*s);e.x=r+f*a+l*c,e.y=i+f*c-l*a}else e.x=r+u,e.y=i}function go(t,n){var e=n.x-t.x,r=n.y-t.y,i=t.r+n.r;return i*i-1e-6>e*e+r*r}function mo(t){var n=t._,e=t.next._,r=n.r+e.r,i=(n.x*e.r+e.x*n.r)/r,o=(n.y*e.r+e.y*n.r)/r;return i*i+o*o}function xo(t){this._=t,this.next=null,this.previous=null}function bo(t){if(!(i=t.length))return 0;var n,e,r,i,o,u,a,c,s,f,l;if(n=t[0],n.x=0,n.y=0,!(i>1))return n.r;if(e=t[1],n.x=-e.r,e.x=n.r,e.y=0,!(i>2))return n.r+e.r;yo(e,n,r=t[2]),n=new xo(n),e=new xo(e),r=new xo(r),n.next=r.previous=e,e.next=n.previous=r,r.next=e.previous=n;t:for(a=3;a<i;++a){yo(n._,e._,r=t[a]),r=new xo(r),c=e.next,s=n.previous,f=e._.r,l=n._.r;do{if(f<=l){if(go(c._,r._)){e=c,n.next=e,e.previous=n,--a;continue t}f+=c._.r,c=c.next}else{if(go(s._,r._)){(n=s).next=e,e.previous=n,--a;continue t}l+=s._.r,s=s.previous}}while(c!==s.next);for(r.previous=n,r.next=e,n.next=e.previous=e=r,o=mo(n);(r=r.next)!==e;)(u=mo(r))<o&&(n=r,o=u);e=n.next}for(n=[e._],r=e;(r=r.next)!==e;)n.push(r._);for(r=Hv(n),a=0;a<i;++a)n=t[a],n.x-=r.x,n.y-=r.y;return r.r}function wo(t){return null==t?null:Mo(t)}function Mo(t){if("function"!=typeof t)throw new Error;return t}function To(){return 0}function ko(t){return Math.sqrt(t.value)}function No(t){return function(n){n.children||(n.r=Math.max(0,+t(n)||0))}}function So(t,n){return function(e){if(r=e.children){var r,i,o,u=r.length,a=t(e)*n||0;if(a)for(i=0;i<u;++i)r[i].r+=a;if(o=bo(r),a)for(i=0;i<u;++i)r[i].r-=a;e.r=o+a}}}function Eo(t){return function(n){var e=n.parent;n.r*=t,e&&(n.x=e.x+t*n.x,n.y=e.y+t*n.y)}}function Ao(t){return t.id}function Co(t){return t.parentId}function zo(t,n){return t.parent===n.parent?1:2}function Po(t){var n=t.children;return n?n[0]:t.t}function Ro(t){var n=t.children;return n?n[n.length-1]:t.t}function Lo(t,n,e){var r=e/(n.i-t.i);n.c-=r,n.s+=e,t.c+=r,n.z+=e,n.m+=e}function qo(t){for(var n,e=0,r=0,i=t.children,o=i.length;--o>=0;)(n=i[o]).z+=e,n.m+=e,e+=n.s+(r+=n.c)}function Uo(t,n,e){return t.a.parent===n.parent?t.a:e}function Do(t,n){this._=t,this.parent=null,this.children=null,this.A=null,this.a=this,this.z=0,this.m=0,this.c=0,this.s=0,this.t=null,this.i=n}function Oo(t){for(var n,e,r,i,o,u=new Do(t,0),a=[u];n=a.pop();)if(r=n._.children)for(n.children=new Array(o=r.length),i=o-1;i>=0;--i)a.push(e=n.children[i]=new Do(r[i],i)),e.parent=n;return(u.parent=new Do(null,0)).children=[u],u}function Fo(t,n,e,r,i,o){for(var u,a,c,s,f,l,h,p,d,v,_,y=[],g=n.children,m=0,x=0,b=g.length,w=n.value;m<b;){c=i-e,s=o-r;do{f=g[x++].value}while(!f&&x<b);for(l=h=f,_=f*f*(v=Math.max(s/c,c/s)/(w*t)),d=Math.max(h/_,_/l);x<b;++x){if(f+=a=g[x].value,a<l&&(l=a),a>h&&(h=a),_=f*f*v,(p=Math.max(h/_,_/l))>d){f-=a;break}d=p}y.push(u={value:f,dice:c<s,children:g.slice(m,x)}),u.dice?Vv(u,e,r,i,w?r+=s*f/w:o):Jv(u,e,r,w?e+=c*f/w:i,o),w-=f,m=x}return y}function Io(t,n){return t[0]-n[0]||t[1]-n[1]}function Yo(t){for(var n=t.length,e=[0,1],r=2,i=2;i<n;++i){for(;r>1&&n_(t[e[r-2]],t[e[r-1]],t[i])<=0;)--r;e[r++]=i}return e.slice(0,r)}function Bo(t){this._size=t,this._call=this._error=null,this._tasks=[],this._data=[],this._waiting=this._active=this._ended=this._start=0}function jo(t){if(!t._start)try{Ho(t)}catch(n){if(t._tasks[t._ended+t._active-1])$o(t,n);else if(!t._data)throw n}}function Ho(t){for(;t._start=t._waiting&&t._active<t._size;){var n=t._ended+t._active,e=t._tasks[n],r=e.length-1,i=e[r];e[r]=Xo(t,n),--t._waiting,++t._active,e=i.apply(null,e),t._tasks[n]&&(t._tasks[n]=e||r_)}}function Xo(t,n){return function(e,r){t._tasks[n]&&(--t._active,++t._ended,t._tasks[n]=null,null==t._error&&(null!=e?$o(t,e):(t._data[n]=r,t._waiting?jo(t):Vo(t))))}}function $o(t,n){var e,r=t._tasks.length;for(t._error=n,t._data=void 0,t._waiting=NaN;--r>=0;)if((e=t._tasks[r])&&(t._tasks[r]=null,e.abort))try{e.abort()}catch(n){}t._active=NaN,Vo(t)}function Vo(t){if(!t._active&&t._call){var n=t._data;t._data=void 0,t._call(t._error,n)}}function Wo(t){if(null==t)t=1/0;else if(!((t=+t)>=1))throw new Error("invalid concurrency");return new Bo(t)}function Zo(t){return function(n,e){t(null==n?e:null)}}function Go(t){var n=t.responseType;return n&&"text"!==n?t.response:t.responseText}function Jo(t,n){return function(e){return t(e.responseText,n)}}function Qo(t){function n(n){var o=n+"",u=e.get(o);if(!u){if(i!==M_)return i;e.set(o,u=r.push(n))}return t[(u-1)%t.length]}var e=we(),r=[],i=M_;return t=null==t?[]:w_.call(t),n.domain=function(t){if(!arguments.length)return r.slice();r=[],e=we();for(var i,o,u=-1,a=t.length;++u<a;)e.has(o=(i=t[u])+"")||e.set(o,r.push(i));return n},n.range=function(e){return arguments.length?(t=w_.call(e),n):t.slice()},n.unknown=function(t){return arguments.length?(i=t,n):i},n.copy=function(){return Qo().domain(r).range(t).unknown(i)},n}function Ko(){function t(){var t=i().length,r=u[1]<u[0],l=u[r-0],h=u[1-r];n=(h-l)/Math.max(1,t-c+2*s),a&&(n=Math.floor(n)),l+=(h-l-n*(t-c))*f,e=n*(1-c),a&&(l=Math.round(l),e=Math.round(e));var p=Ms(t).map(function(t){return l+n*t});return o(r?p.reverse():p)}var n,e,r=Qo().unknown(void 0),i=r.domain,o=r.range,u=[0,1],a=!1,c=0,s=0,f=.5;return delete r.unknown,r.domain=function(n){return arguments.length?(i(n),t()):i()},r.range=function(n){return arguments.length?(u=[+n[0],+n[1]],t()):u.slice()},r.rangeRound=function(n){return u=[+n[0],+n[1]],a=!0,t()},r.bandwidth=function(){return e},r.step=function(){return n},r.round=function(n){return arguments.length?(a=!!n,t()):a},r.padding=function(n){return arguments.length?(c=s=Math.max(0,Math.min(1,n)),t()):c},r.paddingInner=function(n){return arguments.length?(c=Math.max(0,Math.min(1,n)),t()):c},r.paddingOuter=function(n){return arguments.length?(s=Math.max(0,Math.min(1,n)),t()):s},r.align=function(n){return arguments.length?(f=Math.max(0,Math.min(1,n)),t()):f},r.copy=function(){return Ko().domain(i()).range(u).round(a).paddingInner(c).paddingOuter(s).align(f)},t()}function tu(t){var n=t.copy;return t.padding=t.paddingOuter,delete t.paddingInner,delete t.paddingOuter,t.copy=function(){return tu(n())},t}function nu(t,n){return(n-=t=+t)?function(e){return(e-t)/n}:T_(n)}function eu(t){return function(n,e){var r=t(n=+n,e=+e);return function(t){return t<=n?0:t>=e?1:r(t)}}}function ru(t){return function(n,e){var r=t(n=+n,e=+e);return function(t){return t<=0?n:t>=1?e:r(t)}}}function iu(t,n,e,r){var i=t[0],o=t[1],u=n[0],a=n[1];return o<i?(i=e(o,i),u=r(a,u)):(i=e(i,o),u=r(u,a)),function(t){return u(i(t))}}function ou(t,n,e,r){var i=Math.min(t.length,n.length)-1,o=new Array(i),u=new Array(i),a=-1;for(t[i]<t[0]&&(t=t.slice().reverse(),n=n.slice().reverse());++a<i;)o[a]=e(t[a],t[a+1]),u[a]=r(n[a],n[a+1]);return function(n){var e=hs(t,n,1,i)-1;return u[e](o[e](n))}}function uu(t,n){return n.domain(t.domain()).range(t.range()).interpolate(t.interpolate()).clamp(t.clamp())}function au(t,n){function e(){return i=Math.min(a.length,c.length)>2?ou:iu,o=u=null,r}function r(n){return(o||(o=i(a,c,f?eu(t):t,s)))(+n)}var i,o,u,a=N_,c=N_,s=cl,f=!1;return r.invert=function(t){return(u||(u=i(c,a,nu,f?ru(n):n)))(+t)},r.domain=function(t){return arguments.length?(a=b_.call(t,k_),e()):a.slice()},r.range=function(t){return arguments.length?(c=w_.call(t),e()):c.slice()},r.rangeRound=function(t){return c=w_.call(t),s=sl,e()},r.clamp=function(t){return arguments.length?(f=!!t,e()):f},r.interpolate=function(t){return arguments.length?(s=t,e()):s},e()}function cu(t){var n=t.domain;return t.ticks=function(t){var e=n();return Ss(e[0],e[e.length-1],null==t?10:t)},t.tickFormat=function(t,e){return S_(n(),t,e)},t.nice=function(e){null==e&&(e=10);var i,o=n(),u=0,a=o.length-1,c=o[u],s=o[a];return s<c&&(i=c,c=s,s=i,i=u,u=a,a=i),(i=r(c,s,e))>0?i=r(c=Math.floor(c/i)*i,s=Math.ceil(s/i)*i,e):i<0&&(i=r(c=Math.ceil(c*i)/i,s=Math.floor(s*i)/i,e)),i>0?(o[u]=Math.floor(c/i)*i,o[a]=Math.ceil(s/i)*i,n(o)):i<0&&(o[u]=Math.ceil(c*i)/i,o[a]=Math.floor(s*i)/i,n(o)),t},t}function su(){var t=au(nu,rl);return t.copy=function(){return uu(t,su())},cu(t)}function fu(){function t(t){return+t}var n=[0,1];return t.invert=t,t.domain=t.range=function(e){return arguments.length?(n=b_.call(e,k_),t):n.slice()},t.copy=function(){return fu().domain(n)},cu(t)}function lu(t,n){return(n=Math.log(n/t))?function(e){return Math.log(e/t)/n}:T_(n)}function hu(t,n){return t<0?function(e){return-Math.pow(-n,e)*Math.pow(-t,1-e)}:function(e){return Math.pow(n,e)*Math.pow(t,1-e)}}function pu(t){return isFinite(t)?+("1e"+t):t<0?0:t}function du(t){return 10===t?pu:t===Math.E?Math.exp:function(n){return Math.pow(t,n)}}function vu(t){return t===Math.E?Math.log:10===t&&Math.log10||2===t&&Math.log2||(t=Math.log(t),function(n){return Math.log(n)/t})}function _u(t){return function(n){return-t(-n)}}function yu(){function n(){return o=vu(i),u=du(i),r()[0]<0&&(o=_u(o),u=_u(u)),e}var e=au(lu,hu).domain([1,10]),r=e.domain,i=10,o=vu(10),u=du(10);return e.base=function(t){return arguments.length?(i=+t,n()):i},e.domain=function(t){return arguments.length?(r(t),n()):r()},e.ticks=function(t){var n,e=r(),a=e[0],c=e[e.length-1];(n=c<a)&&(h=a,a=c,c=h);var s,f,l,h=o(a),p=o(c),d=null==t?10:+t,v=[];if(!(i%1)&&p-h<d){if(h=Math.round(h)-1,p=Math.round(p)+1,a>0){for(;h<p;++h)for(f=1,s=u(h);f<i;++f)if(!((l=s*f)<a)){if(l>c)break;v.push(l)}}else for(;h<p;++h)for(f=i-1,s=u(h);f>=1;--f)if(!((l=s*f)<a)){if(l>c)break;v.push(l)}}else v=Ss(h,p,Math.min(p-h,d)).map(u);return n?v.reverse():v},e.tickFormat=function(n,r){if(null==r&&(r=10===i?".0e":","),"function"!=typeof r&&(r=t.format(r)),n===1/0)return r;null==n&&(n=10);var a=Math.max(1,i*n/e.ticks().length);return function(t){var n=t/u(Math.round(o(t)));return n*i<i-.5&&(n*=i),n<=a?r(t):""}},e.nice=function(){return r(E_(r(),{floor:function(t){return u(Math.floor(o(t)))},ceil:function(t){return u(Math.ceil(o(t)))}}))},e.copy=function(){return uu(e,yu().base(i))},e}function gu(t,n){return t<0?-Math.pow(-t,n):Math.pow(t,n)}function mu(){var t=1,n=au(function(n,e){return(e=gu(e,t)-(n=gu(n,t)))?function(r){return(gu(r,t)-n)/e}:T_(e)},function(n,e){return e=gu(e,t)-(n=gu(n,t)),function(r){return gu(n+e*r,1/t)}}),e=n.domain;return n.exponent=function(n){return arguments.length?(t=+n,e(e())):t},n.copy=function(){return uu(n,mu().exponent(t))},cu(n)}function xu(){function t(){var t=0,o=Math.max(1,r.length);for(i=new Array(o-1);++t<o;)i[t-1]=As(e,t/o);return n}function n(t){if(!isNaN(t=+t))return r[hs(i,t)]}var e=[],r=[],i=[];return n.invertExtent=function(t){var n=r.indexOf(t);return n<0?[NaN,NaN]:[n>0?i[n-1]:e[0],n<i.length?i[n]:e[e.length-1]]},n.domain=function(n){if(!arguments.length)return e.slice();e=[];for(var r,i=0,o=n.length;i<o;++i)null==(r=n[i])||isNaN(r=+r)||e.push(r);return e.sort(ss),t()},n.range=function(n){return arguments.length?(r=w_.call(n),t()):r.slice()},n.quantiles=function(){return i.slice()},n.copy=function(){return xu().domain(e).range(r)},n}function bu(){function t(t){if(t<=t)return u[hs(o,t,0,i)]}function n(){var n=-1;for(o=new Array(i);++n<i;)o[n]=((n+1)*r-(n-i)*e)/(i+1);return t}var e=0,r=1,i=1,o=[.5],u=[0,1];return t.domain=function(t){return arguments.length?(e=+t[0],r=+t[1],n()):[e,r]},t.range=function(t){return arguments.length?(i=(u=w_.call(t)).length-1,n()):u.slice()},t.invertExtent=function(t){var n=u.indexOf(t);return n<0?[NaN,NaN]:n<1?[e,o[0]]:n>=i?[o[i-1],r]:[o[n-1],o[n]]},t.copy=function(){return bu().domain([e,r]).range(u)},cu(t)}function wu(){function t(t){if(t<=t)return e[hs(n,t,0,r)]}var n=[.5],e=[0,1],r=1;return t.domain=function(i){return arguments.length?(n=w_.call(i),r=Math.min(n.length,e.length-1),t):n.slice()},t.range=function(i){return arguments.length?(e=w_.call(i),r=Math.min(n.length,e.length-1),t):e.slice()},t.invertExtent=function(t){var r=e.indexOf(t);return[n[r-1],n[r]]},t.copy=function(){return wu().domain(n).range(e)},t}function Mu(t,n,e,r){function i(n){return t(n=new Date(+n)),n}return i.floor=i,i.ceil=function(e){return t(e=new Date(e-1)),n(e,1),t(e),e},i.round=function(t){var n=i(t),e=i.ceil(t);return t-n<e-t?n:e},i.offset=function(t,e){return n(t=new Date(+t),null==e?1:Math.floor(e)),t},i.range=function(e,r,o){var u=[];if(e=i.ceil(e),o=null==o?1:Math.floor(o),!(e<r&&o>0))return u;do{u.push(new Date(+e))}while(n(e,o),t(e),e<r);return u},i.filter=function(e){return Mu(function(n){if(n>=n)for(;t(n),!e(n);)n.setTime(n-1)},function(t,r){if(t>=t)if(r<0)for(;++r<=0;)for(;n(t,-1),!e(t););else for(;--r>=0;)for(;n(t,1),!e(t););})},e&&(i.count=function(n,r){return A_.setTime(+n),C_.setTime(+r),t(A_),t(C_),Math.floor(e(A_,C_))},i.every=function(t){return t=Math.floor(t),isFinite(t)&&t>0?t>1?i.filter(r?function(n){return r(n)%t==0}:function(n){return i.count(0,n)%t==0}):i:null}),i}function Tu(t){return Mu(function(n){n.setDate(n.getDate()-(n.getDay()+7-t)%7),n.setHours(0,0,0,0)},function(t,n){t.setDate(t.getDate()+7*n)},function(t,n){return(n-t-(n.getTimezoneOffset()-t.getTimezoneOffset())*R_)/L_})}function ku(t){return Mu(function(n){n.setUTCDate(n.getUTCDate()-(n.getUTCDay()+7-t)%7),n.setUTCHours(0,0,0,0)},function(t,n){t.setUTCDate(t.getUTCDate()+7*n)},function(t,n){return(n-t)/L_})}function Nu(t){if(0<=t.y&&t.y<100){var n=new Date(-1,t.m,t.d,t.H,t.M,t.S,t.L);return n.setFullYear(t.y),n}return new Date(t.y,t.m,t.d,t.H,t.M,t.S,t.L)}function Su(t){if(0<=t.y&&t.y<100){var n=new Date(Date.UTC(-1,t.m,t.d,t.H,t.M,t.S,t.L));return n.setUTCFullYear(t.y),n}return new Date(Date.UTC(t.y,t.m,t.d,t.H,t.M,t.S,t.L))}function Eu(t){return{y:t,m:0,d:1,H:0,M:0,S:0,L:0}}function Au(t){function n(t,n){return function(e){var r,i,o,u=[],a=-1,c=0,s=t.length;for(e instanceof Date||(e=new Date(+e));++a<s;)37===t.charCodeAt(a)&&(u.push(t.slice(c,a)),null!=(i=Py[r=t.charAt(++a)])?r=t.charAt(++a):i="e"===r?" ":"0",(o=n[r])&&(r=o(e,i)),u.push(r),c=a+1);return u.push(t.slice(c,a)),u.join("")}}function e(t,n){return function(e){var i=Eu(1900);if(r(i,t,e+="",0)!=e.length)return null;if("p"in i&&(i.H=i.H%12+12*i.p),"W"in i||"U"in i){"w"in i||(i.w="W"in i?1:0);var o="Z"in i?Su(Eu(i.y)).getUTCDay():n(Eu(i.y)).getDay();i.m=0,i.d="W"in i?(i.w+6)%7+7*i.W-(o+5)%7:i.w+7*i.U-(o+6)%7}return"Z"in i?(i.H+=i.Z/100|0,i.M+=i.Z%100,Su(i)):n(i)}}function r(t,n,e,r){for(var i,o,u=0,a=n.length,c=e.length;u<a;){if(r>=c)return-1;if(37===(i=n.charCodeAt(u++))){if(i=n.charAt(u++),!(o=T[i in Py?n.charAt(u++):i])||(r=o(t,e,r))<0)return-1}else if(i!=e.charCodeAt(r++))return-1}return r}var i=t.dateTime,o=t.date,u=t.time,a=t.periods,c=t.days,s=t.shortDays,f=t.months,l=t.shortMonths,h=Pu(a),p=Ru(a),d=Pu(c),v=Ru(c),_=Pu(s),y=Ru(s),g=Pu(f),m=Ru(f),x=Pu(l),b=Ru(l),w={a:function(t){return s[t.getDay()]},A:function(t){return c[t.getDay()]},b:function(t){return l[t.getMonth()]},B:function(t){return f[t.getMonth()]},c:null,d:Wu,e:Wu,H:Zu,I:Gu,j:Ju,L:Qu,m:Ku,M:ta,p:function(t){return a[+(t.getHours()>=12)]},S:na,U:ea,w:ra,W:ia,x:null,X:null,y:oa,Y:ua,Z:aa,"%":wa},M={a:function(t){return s[t.getUTCDay()]},A:function(t){return c[t.getUTCDay()]},b:function(t){return l[t.getUTCMonth()]},B:function(t){return f[t.getUTCMonth()]},c:null,d:ca,e:ca,H:sa,I:fa,j:la,L:ha,m:pa,M:da,p:function(t){return a[+(t.getUTCHours()>=12)]},S:va,U:_a,w:ya,W:ga,x:null,X:null,y:ma,Y:xa,Z:ba,"%":wa},T={a:function(t,n,e){var r=_.exec(n.slice(e));return r?(t.w=y[r[0].toLowerCase()],e+r[0].length):-1},A:function(t,n,e){var r=d.exec(n.slice(e));return r?(t.w=v[r[0].toLowerCase()],e+r[0].length):-1},b:function(t,n,e){var r=x.exec(n.slice(e));return r?(t.m=b[r[0].toLowerCase()],e+r[0].length):-1},B:function(t,n,e){var r=g.exec(n.slice(e));return r?(t.m=m[r[0].toLowerCase()],e+r[0].length):-1},c:function(t,n,e){return r(t,i,n,e)},d:Yu,e:Yu,H:ju,I:ju,j:Bu,L:$u,m:Iu,M:Hu,p:function(t,n,e){var r=h.exec(n.slice(e));return r?(t.p=p[r[0].toLowerCase()],e+r[0].length):-1},S:Xu,U:qu,w:Lu,W:Uu,x:function(t,n,e){return r(t,o,n,e)},X:function(t,n,e){return r(t,u,n,e)},y:Ou,Y:Du,Z:Fu,"%":Vu};return w.x=n(o,w),w.X=n(u,w),w.c=n(i,w),M.x=n(o,M),M.X=n(u,M),M.c=n(i,M),{format:function(t){var e=n(t+="",w);return e.toString=function(){return t},e},parse:function(t){var n=e(t+="",Nu);return n.toString=function(){return t},n},utcFormat:function(t){var e=n(t+="",M);return e.toString=function(){return t},e},utcParse:function(t){var n=e(t,Su);return n.toString=function(){return t},n}}}function Cu(t,n,e){var r=t<0?"-":"",i=(r?-t:t)+"",o=i.length;return r+(o<e?new Array(e-o+1).join(n)+i:i)}function zu(t){return t.replace(qy,"\\$&")}function Pu(t){return new RegExp("^(?:"+t.map(zu).join("|")+")","i")}function Ru(t){for(var n={},e=-1,r=t.length;++e<r;)n[t[e].toLowerCase()]=e;return n}function Lu(t,n,e){var r=Ry.exec(n.slice(e,e+1));return r?(t.w=+r[0],e+r[0].length):-1}function qu(t,n,e){var r=Ry.exec(n.slice(e));return r?(t.U=+r[0],e+r[0].length):-1}function Uu(t,n,e){var r=Ry.exec(n.slice(e));return r?(t.W=+r[0],e+r[0].length):-1}function Du(t,n,e){var r=Ry.exec(n.slice(e,e+4));return r?(t.y=+r[0],e+r[0].length):-1}function Ou(t,n,e){var r=Ry.exec(n.slice(e,e+2));return r?(t.y=+r[0]+(+r[0]>68?1900:2e3),e+r[0].length):-1}function Fu(t,n,e){var r=/^(Z)|([+-]\d\d)(?:\:?(\d\d))?/.exec(n.slice(e,e+6));return r?(t.Z=r[1]?0:-(r[2]+(r[3]||"00")),e+r[0].length):-1}function Iu(t,n,e){var r=Ry.exec(n.slice(e,e+2));return r?(t.m=r[0]-1,e+r[0].length):-1}function Yu(t,n,e){var r=Ry.exec(n.slice(e,e+2));return r?(t.d=+r[0],e+r[0].length):-1}function Bu(t,n,e){var r=Ry.exec(n.slice(e,e+3));return r?(t.m=0,t.d=+r[0],e+r[0].length):-1}function ju(t,n,e){var r=Ry.exec(n.slice(e,e+2));return r?(t.H=+r[0],e+r[0].length):-1}function Hu(t,n,e){var r=Ry.exec(n.slice(e,e+2));return r?(t.M=+r[0],e+r[0].length):-1}function Xu(t,n,e){var r=Ry.exec(n.slice(e,e+2));return r?(t.S=+r[0],e+r[0].length):-1}function $u(t,n,e){var r=Ry.exec(n.slice(e,e+3));return r?(t.L=+r[0],e+r[0].length):-1}function Vu(t,n,e){var r=Ly.exec(n.slice(e,e+1));return r?e+r[0].length:-1}function Wu(t,n){return Cu(t.getDate(),n,2)}function Zu(t,n){return Cu(t.getHours(),n,2)}function Gu(t,n){return Cu(t.getHours()%12||12,n,2)}function Ju(t,n){return Cu(1+Y_.count(oy(t),t),n,3)}function Qu(t,n){return Cu(t.getMilliseconds(),n,3)}function Ku(t,n){return Cu(t.getMonth()+1,n,2)}function ta(t,n){return Cu(t.getMinutes(),n,2)}function na(t,n){return Cu(t.getSeconds(),n,2)}function ea(t,n){return Cu(j_.count(oy(t),t),n,2)}function ra(t){return t.getDay()}function ia(t,n){return Cu(H_.count(oy(t),t),n,2)}function oa(t,n){return Cu(t.getFullYear()%100,n,2)}function ua(t,n){return Cu(t.getFullYear()%1e4,n,4)}function aa(t){var n=t.getTimezoneOffset();return(n>0?"-":(n*=-1,"+"))+Cu(n/60|0,"0",2)+Cu(n%60,"0",2)}function ca(t,n){return Cu(t.getUTCDate(),n,2)}function sa(t,n){return Cu(t.getUTCHours(),n,2)}function fa(t,n){return Cu(t.getUTCHours()%12||12,n,2)}function la(t,n){return Cu(1+ly.count(Ay(t),t),n,3)}function ha(t,n){return Cu(t.getUTCMilliseconds(),n,3)}function pa(t,n){return Cu(t.getUTCMonth()+1,n,2)}function da(t,n){return Cu(t.getUTCMinutes(),n,2)}function va(t,n){return Cu(t.getUTCSeconds(),n,2)}function _a(t,n){return Cu(py.count(Ay(t),t),n,2)}function ya(t){return t.getUTCDay()}function ga(t,n){return Cu(dy.count(Ay(t),t),n,2)}function ma(t,n){return Cu(t.getUTCFullYear()%100,n,2)}function xa(t,n){return Cu(t.getUTCFullYear()%1e4,n,4)}function ba(){return"+0000"}function wa(){return"%"}function Ma(n){return Cy=Au(n),t.timeFormat=Cy.format,t.timeParse=Cy.parse,t.utcFormat=Cy.utcFormat,t.utcParse=Cy.utcParse,Cy}function Ta(t){return new Date(t)}function ka(t){return t instanceof Date?+t:+new Date(+t)}function Na(t,n,e,r,o,u,a,c,s){function f(i){return(a(i)<i?v:u(i)<i?_:o(i)<i?y:r(i)<i?g:n(i)<i?e(i)<i?m:x:t(i)<i?b:w)(i)}function l(n,e,r,o){if(null==n&&(n=10),"number"==typeof n){var u=Math.abs(r-e)/n,a=fs(function(t){return t[2]}).right(M,u);a===M.length?(o=i(e/Hy,r/Hy,n),n=t):a?(o=(a=M[u/M[a-1][2]<M[a][2]/u?a-1:a])[1],n=a[0]):(o=i(e,r,n),n=c)}return null==o?n:n.every(o)}var h=au(nu,rl),p=h.invert,d=h.domain,v=s(".%L"),_=s(":%S"),y=s("%I:%M"),g=s("%I %p"),m=s("%a %d"),x=s("%b %d"),b=s("%B"),w=s("%Y"),M=[[a,1,Oy],[a,5,5*Oy],[a,15,15*Oy],[a,30,30*Oy],[u,1,Fy],[u,5,5*Fy],[u,15,15*Fy],[u,30,30*Fy],[o,1,Iy],[o,3,3*Iy],[o,6,6*Iy],[o,12,12*Iy],[r,1,Yy],[r,2,2*Yy],[e,1,By],[n,1,jy],[n,3,3*jy],[t,1,Hy]];return h.invert=function(t){return new Date(p(t))},h.domain=function(t){return arguments.length?d(b_.call(t,ka)):d().map(Ta)},h.ticks=function(t,n){var e,r=d(),i=r[0],o=r[r.length-1],u=o<i;return u&&(e=i,i=o,o=e),e=l(t,i,o,n),e=e?e.range(i,o+1):[],u?e.reverse():e},h.tickFormat=function(t,n){return null==n?f:s(n)},h.nice=function(t,n){var e=d();return(t=l(t,e[0],e[e.length-1],n))?d(E_(e,t)):h},h.copy=function(){return uu(h,Na(t,n,e,r,o,u,a,c,s))},h}function Sa(t){var n=t.length;return function(e){return t[Math.max(0,Math.min(n-1,Math.floor(e*n)))]}}function Ea(t){function n(n){var o=(n-e)/(r-e);return t(i?Math.max(0,Math.min(1,o)):o)}var e=0,r=1,i=!1;return n.domain=function(t){return arguments.length?(e=+t[0],r=+t[1],n):[e,r]},n.clamp=function(t){return arguments.length?(i=!!t,n):i},n.interpolator=function(e){return arguments.length?(t=e,n):t},n.copy=function(){return Ea(t).domain([e,r]).clamp(i)},cu(n)}function Aa(t){return t>1?0:t<-1?pg:Math.acos(t)}function Ca(t){return t>=1?dg:t<=-1?-dg:Math.asin(t)}function za(t){return t.innerRadius}function Pa(t){return t.outerRadius}function Ra(t){return t.startAngle}function La(t){return t.endAngle}function qa(t){return t&&t.padAngle}function Ua(t,n,e,r,i,o,u,a){var c=e-t,s=r-n,f=u-i,l=a-o,h=(f*(n-o)-l*(t-i))/(l*c-f*s);return[t+h*c,n+h*s]}function Da(t,n,e,r,i,o,u){var a=t-e,c=n-r,s=(u?o:-o)/lg(a*a+c*c),f=s*c,l=-s*a,h=t+f,p=n+l,d=e+f,v=r+l,_=(h+d)/2,y=(p+v)/2,g=d-h,m=v-p,x=g*g+m*m,b=i-o,w=h*v-d*p,M=(m<0?-1:1)*lg(cg(0,b*b*x-w*w)),T=(w*m-g*M)/x,k=(-w*g-m*M)/x,N=(w*m+g*M)/x,S=(-w*g+m*M)/x,E=T-_,A=k-y,C=N-_,z=S-y;return E*E+A*A>C*C+z*z&&(T=N,k=S),{cx:T,cy:k,x01:-f,y01:-l,x11:T*(i/b-1),y11:k*(i/b-1)}}function Oa(t){this._context=t}function Fa(t){return t[0]}function Ia(t){return t[1]}function Ya(t){this._curve=t}function Ba(t){function n(n){return new Ya(t(n))}return n._curve=t,n}function ja(t){var n=t.curve;return t.angle=t.x,delete t.x,t.radius=t.y,delete t.y,t.curve=function(t){return arguments.length?n(Ba(t)):n()._curve},t}function Ha(t){return t.source}function Xa(t){return t.target}function $a(t){function n(){var n,a=kg.call(arguments),c=e.apply(this,a),s=r.apply(this,a);if(u||(u=n=ve()),t(u,+i.apply(this,(a[0]=c,a)),+o.apply(this,a),+i.apply(this,(a[0]=s,a)),+o.apply(this,a)),n)return u=null,n+""||null}var e=Ha,r=Xa,i=Fa,o=Ia,u=null;return n.source=function(t){return arguments.length?(e=t,n):e},n.target=function(t){return arguments.length?(r=t,n):r},n.x=function(t){return arguments.length?(i="function"==typeof t?t:ig(+t),n):i},n.y=function(t){return arguments.length?(o="function"==typeof t?t:ig(+t),n):o},n.context=function(t){return arguments.length?(u=null==t?null:t,n):u},n}function Va(t,n,e,r,i){t.moveTo(n,e),t.bezierCurveTo(n=(n+r)/2,e,n,i,r,i)}function Wa(t,n,e,r,i){t.moveTo(n,e),t.bezierCurveTo(n,e=(e+i)/2,r,e,r,i)}function Za(t,n,e,r,i){var o=Tg(n,e),u=Tg(n,e=(e+i)/2),a=Tg(r,e),c=Tg(r,i);t.moveTo(o[0],o[1]),t.bezierCurveTo(u[0],u[1],a[0],a[1],c[0],c[1])}function Ga(t,n,e){t._context.bezierCurveTo((2*t._x0+t._x1)/3,(2*t._y0+t._y1)/3,(t._x0+2*t._x1)/3,(t._y0+2*t._y1)/3,(t._x0+4*t._x1+n)/6,(t._y0+4*t._y1+e)/6)}function Ja(t){this._context=t}function Qa(t){this._context=t}function Ka(t){this._context=t}function tc(t,n){this._basis=new Ja(t),this._beta=n}function nc(t,n,e){t._context.bezierCurveTo(t._x1+t._k*(t._x2-t._x0),t._y1+t._k*(t._y2-t._y0),t._x2+t._k*(t._x1-n),t._y2+t._k*(t._y1-e),t._x2,t._y2)}function ec(t,n){this._context=t,this._k=(1-n)/6}function rc(t,n){this._context=t,this._k=(1-n)/6}function ic(t,n){this._context=t,this._k=(1-n)/6}function oc(t,n,e){var r=t._x1,i=t._y1,o=t._x2,u=t._y2;if(t._l01_a>hg){var a=2*t._l01_2a+3*t._l01_a*t._l12_a+t._l12_2a,c=3*t._l01_a*(t._l01_a+t._l12_a);r=(r*a-t._x0*t._l12_2a+t._x2*t._l01_2a)/c,i=(i*a-t._y0*t._l12_2a+t._y2*t._l01_2a)/c}if(t._l23_a>hg){var s=2*t._l23_2a+3*t._l23_a*t._l12_a+t._l12_2a,f=3*t._l23_a*(t._l23_a+t._l12_a);o=(o*s+t._x1*t._l23_2a-n*t._l12_2a)/f,u=(u*s+t._y1*t._l23_2a-e*t._l12_2a)/f}t._context.bezierCurveTo(r,i,o,u,t._x2,t._y2)}function uc(t,n){this._context=t,this._alpha=n}function ac(t,n){this._context=t,this._alpha=n}function cc(t,n){this._context=t,this._alpha=n}function sc(t){this._context=t}function fc(t){return t<0?-1:1}function lc(t,n,e){var r=t._x1-t._x0,i=n-t._x1,o=(t._y1-t._y0)/(r||i<0&&-0),u=(e-t._y1)/(i||r<0&&-0),a=(o*i+u*r)/(r+i);return(fc(o)+fc(u))*Math.min(Math.abs(o),Math.abs(u),.5*Math.abs(a))||0}function hc(t,n){var e=t._x1-t._x0;return e?(3*(t._y1-t._y0)/e-n)/2:n}function pc(t,n,e){var r=t._x0,i=t._y0,o=t._x1,u=t._y1,a=(o-r)/3;t._context.bezierCurveTo(r+a,i+a*n,o-a,u-a*e,o,u)}function dc(t){this._context=t}function vc(t){this._context=new _c(t)}function _c(t){this._context=t}function yc(t){this._context=t}function gc(t){var n,e,r=t.length-1,i=new Array(r),o=new Array(r),u=new Array(r);for(i[0]=0,o[0]=2,u[0]=t[0]+2*t[1],n=1;n<r-1;++n)i[n]=1,o[n]=4,u[n]=4*t[n]+2*t[n+1];for(i[r-1]=2,o[r-1]=7,u[r-1]=8*t[r-1]+t[r],n=1;n<r;++n)e=i[n]/o[n-1],o[n]-=e,u[n]-=e*u[n-1];for(i[r-1]=u[r-1]/o[r-1],n=r-2;n>=0;--n)i[n]=(u[n]-i[n+1])/o[n];for(o[r-1]=(t[r]+i[r-1])/2,n=0;n<r-1;++n)o[n]=2*t[n+1]-i[n+1];return[i,o]}function mc(t,n){this._context=t,this._t=n}function xc(t,n){return t[n]}function bc(t){for(var n,e=0,r=-1,i=t.length;++r<i;)(n=+t[r][1])&&(e+=n);return e}function wc(t){return t[0]}function Mc(t){return t[1]}function Tc(){this._=null}function kc(t){t.U=t.C=t.L=t.R=t.P=t.N=null}function Nc(t,n){var e=n,r=n.R,i=e.U;i?i.L===e?i.L=r:i.R=r:t._=r,r.U=i,e.U=r,e.R=r.L,e.R&&(e.R.U=e),r.L=e}function Sc(t,n){var e=n,r=n.L,i=e.U;i?i.L===e?i.L=r:i.R=r:t._=r,r.U=i,e.U=r,e.L=r.R,e.L&&(e.L.U=e),r.R=e}function Ec(t){for(;t.L;)t=t.L;return t}function Ac(t,n,e,r){var i=[null,null],o=um.push(i)-1;return i.left=t,i.right=n,e&&zc(i,t,n,e),r&&zc(i,n,t,r),im[t.index].halfedges.push(o),im[n.index].halfedges.push(o),i}function Cc(t,n,e){var r=[n,e];return r.left=t,r}function zc(t,n,e,r){t[0]||t[1]?t.left===e?t[1]=r:t[0]=r:(t[0]=r,t.left=n,t.right=e)}function Pc(t,n,e,r,i){var o,u=t[0],a=t[1],c=u[0],s=u[1],f=0,l=1,h=a[0]-c,p=a[1]-s;if(o=n-c,h||!(o>0)){if(o/=h,h<0){if(o<f)return;o<l&&(l=o)}else if(h>0){if(o>l)return;o>f&&(f=o)}if(o=r-c,h||!(o<0)){if(o/=h,h<0){if(o>l)return;o>f&&(f=o)}else if(h>0){if(o<f)return;o<l&&(l=o)}if(o=e-s,p||!(o>0)){if(o/=p,p<0){if(o<f)return;o<l&&(l=o)}else if(p>0){if(o>l)return;o>f&&(f=o)}if(o=i-s,p||!(o<0)){if(o/=p,p<0){if(o>l)return;o>f&&(f=o)}else if(p>0){if(o<f)return;o<l&&(l=o)}return!(f>0||l<1)||(f>0&&(t[0]=[c+f*h,s+f*p]),l<1&&(t[1]=[c+l*h,s+l*p]),!0)}}}}}function Rc(t,n,e,r,i){var o=t[1];if(o)return!0;var u,a,c=t[0],s=t.left,f=t.right,l=s[0],h=s[1],p=f[0],d=f[1],v=(l+p)/2,_=(h+d)/2;if(d===h){if(v<n||v>=r)return;if(l>p){if(c){if(c[1]>=i)return}else c=[v,e];o=[v,i]}else{if(c){if(c[1]<e)return}else c=[v,i];o=[v,e]}}else if(u=(l-p)/(d-h),a=_-u*v,u<-1||u>1)if(l>p){if(c){if(c[1]>=i)return}else c=[(e-a)/u,e];o=[(i-a)/u,i]}else{if(c){if(c[1]<e)return}else c=[(i-a)/u,i];o=[(e-a)/u,e]}else if(h<d){if(c){if(c[0]>=r)return}else c=[n,u*n+a];o=[r,u*r+a]}else{if(c){if(c[0]<n)return}else c=[r,u*r+a];o=[n,u*n+a]}return t[0]=c,t[1]=o,!0}function Lc(t,n,e,r){for(var i,o=um.length;o--;)Rc(i=um[o],t,n,e,r)&&Pc(i,t,n,e,r)&&(Math.abs(i[0][0]-i[1][0])>sm||Math.abs(i[0][1]-i[1][1])>sm)||delete um[o]}function qc(t){return im[t.index]={site:t,halfedges:[]}}function Uc(t,n){var e=t.site,r=n.left,i=n.right;return e===i&&(i=r,r=e),i?Math.atan2(i[1]-r[1],i[0]-r[0]):(e===r?(r=n[1],i=n[0]):(r=n[0],i=n[1]),Math.atan2(r[0]-i[0],i[1]-r[1]))}function Dc(t,n){return n[+(n.left!==t.site)]}function Oc(t,n){return n[+(n.left===t.site)]}function Fc(){for(var t,n,e,r,i=0,o=im.length;i<o;++i)if((t=im[i])&&(r=(n=t.halfedges).length)){var u=new Array(r),a=new Array(r);for(e=0;e<r;++e)u[e]=e,a[e]=Uc(t,um[n[e]]);for(u.sort(function(t,n){return a[n]-a[t]}),e=0;e<r;++e)a[e]=n[u[e]];for(e=0;e<r;++e)n[e]=a[e]}}function Ic(t,n,e,r){var i,o,u,a,c,s,f,l,h,p,d,v,_=im.length,y=!0;for(i=0;i<_;++i)if(o=im[i]){for(u=o.site,a=(c=o.halfedges).length;a--;)um[c[a]]||c.splice(a,1);for(a=0,s=c.length;a<s;)d=(p=Oc(o,um[c[a]]))[0],v=p[1],l=(f=Dc(o,um[c[++a%s]]))[0],h=f[1],(Math.abs(d-l)>sm||Math.abs(v-h)>sm)&&(c.splice(a,0,um.push(Cc(u,p,Math.abs(d-t)<sm&&r-v>sm?[t,Math.abs(l-t)<sm?h:r]:Math.abs(v-r)<sm&&e-d>sm?[Math.abs(h-r)<sm?l:e,r]:Math.abs(d-e)<sm&&v-n>sm?[e,Math.abs(l-e)<sm?h:n]:Math.abs(v-n)<sm&&d-t>sm?[Math.abs(h-n)<sm?l:t,n]:null))-1),++s);s&&(y=!1)}if(y){var g,m,x,b=1/0;for(i=0,y=null;i<_;++i)(o=im[i])&&(x=(g=(u=o.site)[0]-t)*g+(m=u[1]-n)*m)<b&&(b=x,y=o);if(y){var w=[t,n],M=[t,r],T=[e,r],k=[e,n];y.halfedges.push(um.push(Cc(u=y.site,w,M))-1,um.push(Cc(u,M,T))-1,um.push(Cc(u,T,k))-1,um.push(Cc(u,k,w))-1)}}for(i=0;i<_;++i)(o=im[i])&&(o.halfedges.length||delete im[i])}function Yc(){kc(this),this.x=this.y=this.arc=this.site=this.cy=null}function Bc(t){var n=t.P,e=t.N;if(n&&e){var r=n.site,i=t.site,o=e.site;if(r!==o){var u=i[0],a=i[1],c=r[0]-u,s=r[1]-a,f=o[0]-u,l=o[1]-a,h=2*(c*l-s*f);if(!(h>=-fm)){var p=c*c+s*s,d=f*f+l*l,v=(l*p-s*d)/h,_=(c*d-f*p)/h,y=am.pop()||new Yc;y.arc=t,y.site=i,y.x=v+u,y.y=(y.cy=_+a)+Math.sqrt(v*v+_*_),t.circle=y;for(var g=null,m=om._;m;)if(y.y<m.y||y.y===m.y&&y.x<=m.x){if(!m.L){g=m.P;break}m=m.L}else{if(!m.R){g=m;break}m=m.R}om.insert(g,y),g||(em=y)}}}}function jc(t){var n=t.circle;n&&(n.P||(em=n.N),om.remove(n),am.push(n),kc(n),t.circle=null)}function Hc(){kc(this),this.edge=this.site=this.circle=null}function Xc(t){var n=cm.pop()||new Hc;return n.site=t,n}function $c(t){jc(t),rm.remove(t),cm.push(t),kc(t)}function Vc(t){var n=t.circle,e=n.x,r=n.cy,i=[e,r],o=t.P,u=t.N,a=[t];$c(t);for(var c=o;c.circle&&Math.abs(e-c.circle.x)<sm&&Math.abs(r-c.circle.cy)<sm;)o=c.P,a.unshift(c),$c(c),c=o;a.unshift(c),jc(c);for(var s=u;s.circle&&Math.abs(e-s.circle.x)<sm&&Math.abs(r-s.circle.cy)<sm;)u=s.N,a.push(s),$c(s),s=u;a.push(s),jc(s);var f,l=a.length;for(f=1;f<l;++f)s=a[f],c=a[f-1],zc(s.edge,c.site,s.site,i);c=a[0],(s=a[l-1]).edge=Ac(c.site,s.site,null,i),Bc(c),Bc(s)}function Wc(t){for(var n,e,r,i,o=t[0],u=t[1],a=rm._;a;)if((r=Zc(a,u)-o)>sm)a=a.L;else{if(!((i=o-Gc(a,u))>sm)){r>-sm?(n=a.P,e=a):i>-sm?(n=a,e=a.N):n=e=a;break}if(!a.R){n=a;break}a=a.R}qc(t);var c=Xc(t);if(rm.insert(n,c),n||e){if(n===e)return jc(n),e=Xc(n.site),rm.insert(c,e),c.edge=e.edge=Ac(n.site,c.site),Bc(n),void Bc(e);if(e){jc(n),jc(e);var s=n.site,f=s[0],l=s[1],h=t[0]-f,p=t[1]-l,d=e.site,v=d[0]-f,_=d[1]-l,y=2*(h*_-p*v),g=h*h+p*p,m=v*v+_*_,x=[(_*g-p*m)/y+f,(h*m-v*g)/y+l];zc(e.edge,s,d,x),c.edge=Ac(s,t,null,x),e.edge=Ac(t,d,null,x),Bc(n),Bc(e)}else c.edge=Ac(n.site,c.site)}}function Zc(t,n){var e=t.site,r=e[0],i=e[1],o=i-n;if(!o)return r;var u=t.P;if(!u)return-1/0;var a=(e=u.site)[0],c=e[1],s=c-n;if(!s)return a;var f=a-r,l=1/o-1/s,h=f/s;return l?(-h+Math.sqrt(h*h-2*l*(f*f/(-2*s)-c+s/2+i-o/2)))/l+r:(r+a)/2}function Gc(t,n){var e=t.N;if(e)return Zc(e,n);var r=t.site;return r[1]===n?r[0]:1/0}function Jc(t,n,e){return(t[0]-e[0])*(n[1]-t[1])-(t[0]-n[0])*(e[1]-t[1])}function Qc(t,n){return n[1]-t[1]||n[0]-t[0]}function Kc(t,n){var e,r,i,o=t.sort(Qc).pop();for(um=[],im=new Array(t.length),rm=new Tc,om=new Tc;;)if(i=em,o&&(!i||o[1]<i.y||o[1]===i.y&&o[0]<i.x))o[0]===e&&o[1]===r||(Wc(o),e=o[0],r=o[1]),o=t.pop();else{if(!i)break;Vc(i.arc)}if(Fc(),n){var u=+n[0][0],a=+n[0][1],c=+n[1][0],s=+n[1][1];Lc(u,a,c,s),Ic(u,a,c,s)}this.edges=um,this.cells=im,rm=om=um=im=null}function ts(t,n,e){this.target=t,this.type=n,this.transform=e}function ns(t,n,e){this.k=t,this.x=n,this.y=e}function es(t){return t.__zoom||hm}function rs(){t.event.stopImmediatePropagation()}function is(){return!t.event.button}function os(){var t,n,e=this;return e instanceof SVGElement?(t=(e=e.ownerSVGElement||e).width.baseVal.value,n=e.height.baseVal.value):(t=e.clientWidth,n=e.clientHeight),[[0,0],[t,n]]}function us(){return this.__zoom||hm}function as(){return-t.event.deltaY*(t.event.deltaMode?120:1)/500}function cs(){return"ontouchstart"in this}var ss=function(t,n){return t<n?-1:t>n?1:t>=n?0:NaN},fs=function(t){return 1===t.length&&(t=n(t)),{left:function(n,e,r,i){for(null==r&&(r=0),null==i&&(i=n.length);r<i;){var o=r+i>>>1;t(n[o],e)<0?r=o+1:i=o}return r},right:function(n,e,r,i){for(null==r&&(r=0),null==i&&(i=n.length);r<i;){var o=r+i>>>1;t(n[o],e)>0?i=o:r=o+1}return r}}},ls=fs(ss),hs=ls.right,ps=ls.left,ds=function(t){return null===t?NaN:+t},vs=function(t,n){var e,r,i=t.length,o=0,u=-1,a=0,c=0;if(null==n)for(;++u<i;)isNaN(e=ds(t[u]))||(c+=(r=e-a)*(e-(a+=r/++o)));else for(;++u<i;)isNaN(e=ds(n(t[u],u,t)))||(c+=(r=e-a)*(e-(a+=r/++o)));if(o>1)return c/(o-1)},_s=function(t,n){var e=vs(t,n);return e?Math.sqrt(e):e},ys=function(t,n){var e,r,i,o=t.length,u=-1;if(null==n){for(;++u<o;)if(null!=(e=t[u])&&e>=e)for(r=i=e;++u<o;)null!=(e=t[u])&&(r>e&&(r=e),i<e&&(i=e))}else for(;++u<o;)if(null!=(e=n(t[u],u,t))&&e>=e)for(r=i=e;++u<o;)null!=(e=n(t[u],u,t))&&(r>e&&(r=e),i<e&&(i=e));return[r,i]},gs=Array.prototype,ms=gs.slice,xs=gs.map,bs=function(t){return function(){return t}},ws=function(t){return t},Ms=function(t,n,e){t=+t,n=+n,e=(i=arguments.length)<2?(n=t,t=0,1):i<3?1:+e;for(var r=-1,i=0|Math.max(0,Math.ceil((n-t)/e)),o=new Array(i);++r<i;)o[r]=t+r*e;return o},Ts=Math.sqrt(50),ks=Math.sqrt(10),Ns=Math.sqrt(2),Ss=function(t,n,e){var i,o,u,a=n<t,c=-1;if(a&&(i=t,t=n,n=i),0===(u=r(t,n,e))||!isFinite(u))return[];if(u>0)for(t=Math.ceil(t/u),n=Math.floor(n/u),o=new Array(i=Math.ceil(n-t+1));++c<i;)o[c]=(t+c)*u;else for(t=Math.floor(t*u),n=Math.ceil(n*u),o=new Array(i=Math.ceil(t-n+1));++c<i;)o[c]=(t-c)/u;return a&&o.reverse(),o},Es=function(t){return Math.ceil(Math.log(t.length)/Math.LN2)+1},As=function(t,n,e){if(null==e&&(e=ds),r=t.length){if((n=+n)<=0||r<2)return+e(t[0],0,t);if(n>=1)return+e(t[r-1],r-1,t);var r,i=(r-1)*n,o=Math.floor(i),u=+e(t[o],o,t);return u+(+e(t[o+1],o+1,t)-u)*(i-o)}},Cs=function(t){for(var n,e,r,i=t.length,o=-1,u=0;++o<i;)u+=t[o].length;for(e=new Array(u);--i>=0;)for(n=(r=t[i]).length;--n>=0;)e[--u]=r[n];return e},zs=function(t,n){var e,r,i=t.length,o=-1;if(null==n){for(;++o<i;)if(null!=(e=t[o])&&e>=e)for(r=e;++o<i;)null!=(e=t[o])&&r>e&&(r=e)}else for(;++o<i;)if(null!=(e=n(t[o],o,t))&&e>=e)for(r=e;++o<i;)null!=(e=n(t[o],o,t))&&r>e&&(r=e);return r},Ps=function(t){if(!(i=t.length))return[];for(var n=-1,e=zs(t,o),r=new Array(e);++n<e;)for(var i,u=-1,a=r[n]=new Array(i);++u<i;)a[u]=t[u][n];return r},Rs=Array.prototype.slice,Ls=function(t){return t},qs=1,Us=2,Ds=3,Os=4,Fs=1e-6,Is={value:function(){}};p.prototype=h.prototype={constructor:p,on:function(t,n){var e,r=this._,i=d(t+"",r),o=-1,u=i.length;{if(!(arguments.length<2)){if(null!=n&&"function"!=typeof n)throw new Error("invalid callback: "+n);for(;++o<u;)if(e=(t=i[o]).type)r[e]=_(r[e],t.name,n);else if(null==n)for(e in r)r[e]=_(r[e],t.name,null);return this}for(;++o<u;)if((e=(t=i[o]).type)&&(e=v(r[e],t.name)))return e}},copy:function(){var t={},n=this._;for(var e in n)t[e]=n[e].slice();return new p(t)},call:function(t,n){if((e=arguments.length-2)>0)for(var e,r,i=new Array(e),o=0;o<e;++o)i[o]=arguments[o+2];if(!this._.hasOwnProperty(t))throw new Error("unknown type: "+t);for(o=0,e=(r=this._[t]).length;o<e;++o)r[o].value.apply(n,i)},apply:function(t,n,e){if(!this._.hasOwnProperty(t))throw new Error("unknown type: "+t);for(var r=this._[t],i=0,o=r.length;i<o;++i)r[i].value.apply(n,e)}};var Ys="http://www.w3.org/1999/xhtml",Bs={svg:"http://www.w3.org/2000/svg",xhtml:Ys,xlink:"http://www.w3.org/1999/xlink",xml:"http://www.w3.org/XML/1998/namespace",xmlns:"http://www.w3.org/2000/xmlns/"},js=function(t){var n=t+="",e=n.indexOf(":");return e>=0&&"xmlns"!==(n=t.slice(0,e))&&(t=t.slice(e+1)),Bs.hasOwnProperty(n)?{space:Bs[n],local:t}:t},Hs=function(t){var n=js(t);return(n.local?g:y)(n)},Xs=0;x.prototype=m.prototype={constructor:x,get:function(t){for(var n=this._;!(n in t);)if(!(t=t.parentNode))return;return t[n]},set:function(t,n){return t[this._]=n},remove:function(t){return this._ in t&&delete t[this._]},toString:function(){return this._}};var $s=function(t){return function(){return this.matches(t)}};if("undefined"!=typeof document){var Vs=document.documentElement;if(!Vs.matches){var Ws=Vs.webkitMatchesSelector||Vs.msMatchesSelector||Vs.mozMatchesSelector||Vs.oMatchesSelector;$s=function(t){return function(){return Ws.call(this,t)}}}}var Zs=$s,Gs={};t.event=null,"undefined"!=typeof document&&("onmouseenter"in document.documentElement||(Gs={mouseenter:"mouseover",mouseleave:"mouseout"}));var Js=function(){for(var n,e=t.event;n=e.sourceEvent;)e=n;return e},Qs=function(t,n){var e=t.ownerSVGElement||t;if(e.createSVGPoint){var r=e.createSVGPoint();return r.x=n.clientX,r.y=n.clientY,r=r.matrixTransform(t.getScreenCTM().inverse()),[r.x,r.y]}var i=t.getBoundingClientRect();return[n.clientX-i.left-t.clientLeft,n.clientY-i.top-t.clientTop]},Ks=function(t){var n=Js();return n.changedTouches&&(n=n.changedTouches[0]),Qs(t,n)},tf=function(t){return null==t?S:function(){return this.querySelector(t)}},nf=function(t){return null==t?E:function(){return this.querySelectorAll(t)}},ef=function(t){return new Array(t.length)};A.prototype={constructor:A,appendChild:function(t){return this._parent.insertBefore(t,this._next)},insertBefore:function(t,n){return this._parent.insertBefore(t,n)},querySelector:function(t){return this._parent.querySelector(t)},querySelectorAll:function(t){return this._parent.querySelectorAll(t)}};var rf=function(t){return function(){return t}},of="$",uf=function(t){return t.ownerDocument&&t.ownerDocument.defaultView||t.document&&t||t.defaultView};W.prototype={add:function(t){this._names.indexOf(t)<0&&(this._names.push(t),this._node.setAttribute("class",this._names.join(" ")))},remove:function(t){var n=this._names.indexOf(t);n>=0&&(this._names.splice(n,1),this._node.setAttribute("class",this._names.join(" ")))},contains:function(t){return this._names.indexOf(t)>=0}};var af=[null];pt.prototype=dt.prototype={constructor:pt,select:function(t){"function"!=typeof t&&(t=tf(t));for(var n=this._groups,e=n.length,r=new Array(e),i=0;i<e;++i)for(var o,u,a=n[i],c=a.length,s=r[i]=new Array(c),f=0;f<c;++f)(o=a[f])&&(u=t.call(o,o.__data__,f,a))&&("__data__"in o&&(u.__data__=o.__data__),s[f]=u);return new pt(r,this._parents)},selectAll:function(t){"function"!=typeof t&&(t=nf(t));for(var n=this._groups,e=n.length,r=[],i=[],o=0;o<e;++o)for(var u,a=n[o],c=a.length,s=0;s<c;++s)(u=a[s])&&(r.push(t.call(u,u.__data__,s,a)),i.push(u));return new pt(r,i)},filter:function(t){"function"!=typeof t&&(t=Zs(t));for(var n=this._groups,e=n.length,r=new Array(e),i=0;i<e;++i)for(var o,u=n[i],a=u.length,c=r[i]=[],s=0;s<a;++s)(o=u[s])&&t.call(o,o.__data__,s,u)&&c.push(o);return new pt(r,this._parents)},data:function(t,n){if(!t)return p=new Array(this.size()),s=-1,this.each(function(t){p[++s]=t}),p;var e=n?z:C,r=this._parents,i=this._groups;"function"!=typeof t&&(t=rf(t));for(var o=i.length,u=new Array(o),a=new Array(o),c=new Array(o),s=0;s<o;++s){var f=r[s],l=i[s],h=l.length,p=t.call(f,f&&f.__data__,s,r),d=p.length,v=a[s]=new Array(d),_=u[s]=new Array(d);e(f,l,v,_,c[s]=new Array(h),p,n);for(var y,g,m=0,x=0;m<d;++m)if(y=v[m]){for(m>=x&&(x=m+1);!(g=_[x])&&++x<d;);y._next=g||null}}return u=new pt(u,r),u._enter=a,u._exit=c,u},enter:function(){return new pt(this._enter||this._groups.map(ef),this._parents)},exit:function(){return new pt(this._exit||this._groups.map(ef),this._parents)},merge:function(t){for(var n=this._groups,e=t._groups,r=n.length,i=e.length,o=Math.min(r,i),u=new Array(r),a=0;a<o;++a)for(var c,s=n[a],f=e[a],l=s.length,h=u[a]=new Array(l),p=0;p<l;++p)(c=s[p]||f[p])&&(h[p]=c);for(;a<r;++a)u[a]=n[a];return new pt(u,this._parents)},order:function(){for(var t=this._groups,n=-1,e=t.length;++n<e;)for(var r,i=t[n],o=i.length-1,u=i[o];--o>=0;)(r=i[o])&&(u&&u!==r.nextSibling&&u.parentNode.insertBefore(r,u),u=r);return this},sort:function(t){t||(t=P);for(var n=this._groups,e=n.length,r=new Array(e),i=0;i<e;++i){for(var o,u=n[i],a=u.length,c=r[i]=new Array(a),s=0;s<a;++s)(o=u[s])&&(c[s]=o);c.sort(function(n,e){return n&&e?t(n.__data__,e.__data__):!n-!e})}return new pt(r,this._parents).order()},call:function(){var t=arguments[0];return arguments[0]=this,t.apply(null,arguments),this},nodes:function(){var t=new Array(this.size()),n=-1;return this.each(function(){t[++n]=this}),t},node:function(){for(var t=this._groups,n=0,e=t.length;n<e;++n)for(var r=t[n],i=0,o=r.length;i<o;++i){var u=r[i];if(u)return u}return null},size:function(){var t=0;return this.each(function(){++t}),t},empty:function(){return!this.node()},each:function(t){for(var n=this._groups,e=0,r=n.length;e<r;++e)for(var i,o=n[e],u=0,a=o.length;u<a;++u)(i=o[u])&&t.call(i,i.__data__,u,o);return this},attr:function(t,n){var e=js(t);if(arguments.length<2){var r=this.node();return e.local?r.getAttributeNS(e.space,e.local):r.getAttribute(e)}return this.each((null==n?e.local?L:R:"function"==typeof n?e.local?O:D:e.local?U:q)(e,n))},style:function(t,n,e){return arguments.length>1?this.each((null==n?F:"function"==typeof n?Y:I)(t,n,null==e?"":e)):B(this.node(),t)},property:function(t,n){return arguments.length>1?this.each((null==n?j:"function"==typeof n?X:H)(t,n)):this.node()[t]},classed:function(t,n){var e=$(t+"");if(arguments.length<2){for(var r=V(this.node()),i=-1,o=e.length;++i<o;)if(!r.contains(e[i]))return!1;return!0}return this.each(("function"==typeof n?K:n?J:Q)(e,n))},text:function(t){return arguments.length?this.each(null==t?tt:("function"==typeof t?et:nt)(t)):this.node().textContent},html:function(t){return arguments.length?this.each(null==t?rt:("function"==typeof t?ot:it)(t)):this.node().innerHTML},raise:function(){return this.each(ut)},lower:function(){return this.each(at)},append:function(t){var n="function"==typeof t?t:Hs(t);return this.select(function(){return this.appendChild(n.apply(this,arguments))})},insert:function(t,n){var e="function"==typeof t?t:Hs(t),r=null==n?ct:"function"==typeof n?n:tf(n);return this.select(function(){return this.insertBefore(e.apply(this,arguments),r.apply(this,arguments)||null)})},remove:function(){return this.each(st)},datum:function(t){return arguments.length?this.property("__data__",t):this.node().__data__},on:function(t,n,e){var r,i,o=M(t+""),u=o.length;{if(!(arguments.length<2)){for(a=n?k:T,null==e&&(e=!1),r=0;r<u;++r)this.each(a(o[r],n,e));return this}var a=this.node().__on;if(a)for(var c,s=0,f=a.length;s<f;++s)for(r=0,c=a[s];r<u;++r)if((i=o[r]).type===c.type&&i.name===c.name)return c.value}},dispatch:function(t,n){return this.each(("function"==typeof n?ht:lt)(t,n))}};var cf=function(t){return"string"==typeof t?new pt([[document.querySelector(t)]],[document.documentElement]):new pt([[t]],af)},sf=function(t,n,e){arguments.length<3&&(e=n,n=Js().changedTouches);for(var r,i=0,o=n?n.length:0;i<o;++i)if((r=n[i]).identifier===e)return Qs(t,r);return null},ff=function(){t.event.preventDefault(),t.event.stopImmediatePropagation()},lf=function(t){var n=t.document.documentElement,e=cf(t).on("dragstart.drag",ff,!0);"onselectstart"in n?e.on("selectstart.drag",ff,!0):(n.__noselect=n.style.MozUserSelect,n.style.MozUserSelect="none")},hf=function(t){return function(){return t}};yt.prototype.on=function(){var t=this._.on.apply(this._,arguments);return t===this._?this:t};var pf=function(t,n,e){t.prototype=n.prototype=e,e.constructor=t},df="\\s*([+-]?\\d+)\\s*",vf="\\s*([+-]?\\d*\\.?\\d+(?:[eE][+-]?\\d+)?)\\s*",_f="\\s*([+-]?\\d*\\.?\\d+(?:[eE][+-]?\\d+)?)%\\s*",yf=/^#([0-9a-f]{3})$/,gf=/^#([0-9a-f]{6})$/,mf=new RegExp("^rgb\\("+[df,df,df]+"\\)$"),xf=new RegExp("^rgb\\("+[_f,_f,_f]+"\\)$"),bf=new RegExp("^rgba\\("+[df,df,df,vf]+"\\)$"),wf=new RegExp("^rgba\\("+[_f,_f,_f,vf]+"\\)$"),Mf=new RegExp("^hsl\\("+[vf,_f,_f]+"\\)$"),Tf=new RegExp("^hsla\\("+[vf,_f,_f,vf]+"\\)$"),kf={aliceblue:15792383,antiquewhite:16444375,aqua:65535,aquamarine:8388564,azure:15794175,beige:16119260,bisque:16770244,black:0,blanchedalmond:16772045,blue:255,blueviolet:9055202,brown:10824234,burlywood:14596231,cadetblue:6266528,chartreuse:8388352,chocolate:13789470,coral:16744272,cornflowerblue:6591981,cornsilk:16775388,crimson:14423100,cyan:65535,darkblue:139,darkcyan:35723,darkgoldenrod:12092939,darkgray:11119017,darkgreen:25600,darkgrey:11119017,darkkhaki:12433259,darkmagenta:9109643,darkolivegreen:5597999,darkorange:16747520,darkorchid:10040012,darkred:9109504,darksalmon:15308410,darkseagreen:9419919,darkslateblue:4734347,darkslategray:3100495,darkslategrey:3100495,darkturquoise:52945,darkviolet:9699539,deeppink:16716947,deepskyblue:49151,dimgray:6908265,dimgrey:6908265,dodgerblue:2003199,firebrick:11674146,floralwhite:16775920,forestgreen:2263842,fuchsia:16711935,gainsboro:14474460,ghostwhite:16316671,gold:16766720,goldenrod:14329120,gray:8421504,green:32768,greenyellow:11403055,grey:8421504,honeydew:15794160,hotpink:16738740,indianred:13458524,indigo:4915330,ivory:16777200,khaki:15787660,lavender:15132410,lavenderblush:16773365,lawngreen:8190976,lemonchiffon:16775885,lightblue:11393254,lightcoral:15761536,lightcyan:14745599,lightgoldenrodyellow:16448210,lightgray:13882323,lightgreen:9498256,lightgrey:13882323,lightpink:16758465,lightsalmon:16752762,lightseagreen:2142890,lightskyblue:8900346,lightslategray:7833753,lightslategrey:7833753,lightsteelblue:11584734,lightyellow:16777184,lime:65280,limegreen:3329330,linen:16445670,magenta:16711935,maroon:8388608,mediumaquamarine:6737322,mediumblue:205,mediumorchid:12211667,mediumpurple:9662683,mediumseagreen:3978097,mediumslateblue:8087790,mediumspringgreen:64154,mediumturquoise:4772300,mediumvioletred:13047173,midnightblue:1644912,mintcream:16121850,mistyrose:16770273,moccasin:16770229,navajowhite:16768685,navy:128,oldlace:16643558,olive:8421376,olivedrab:7048739,orange:16753920,orangered:16729344,orchid:14315734,palegoldenrod:15657130,palegreen:10025880,paleturquoise:11529966,palevioletred:14381203,papayawhip:16773077,peachpuff:16767673,peru:13468991,pink:16761035,plum:14524637,powderblue:11591910,purple:8388736,rebeccapurple:6697881,red:16711680,rosybrown:12357519,royalblue:4286945,saddlebrown:9127187,salmon:16416882,sandybrown:16032864,seagreen:3050327,seashell:16774638,sienna:10506797,silver:12632256,skyblue:8900331,slateblue:6970061,slategray:7372944,slategrey:7372944,snow:16775930,springgreen:65407,steelblue:4620980,tan:13808780,teal:32896,thistle:14204888,tomato:16737095,turquoise:4251856,violet:15631086,wheat:16113331,white:16777215,whitesmoke:16119285,yellow:16776960,yellowgreen:10145074};pf(Mt,Tt,{displayable:function(){return this.rgb().displayable()},toString:function(){return this.rgb()+""}}),pf(At,Et,wt(Mt,{brighter:function(t){return t=null==t?1/.7:Math.pow(1/.7,t),new At(this.r*t,this.g*t,this.b*t,this.opacity)},darker:function(t){return t=null==t?.7:Math.pow(.7,t),new At(this.r*t,this.g*t,this.b*t,this.opacity)},rgb:function(){return this},displayable:function(){return 0<=this.r&&this.r<=255&&0<=this.g&&this.g<=255&&0<=this.b&&this.b<=255&&0<=this.opacity&&this.opacity<=1},toString:function(){var t=this.opacity;return(1===(t=isNaN(t)?1:Math.max(0,Math.min(1,t)))?"rgb(":"rgba(")+Math.max(0,Math.min(255,Math.round(this.r)||0))+", "+Math.max(0,Math.min(255,Math.round(this.g)||0))+", "+Math.max(0,Math.min(255,Math.round(this.b)||0))+(1===t?")":", "+t+")")}})),pf(Rt,Pt,wt(Mt,{brighter:function(t){return t=null==t?1/.7:Math.pow(1/.7,t),new Rt(this.h,this.s,this.l*t,this.opacity)},darker:function(t){return t=null==t?.7:Math.pow(.7,t),new Rt(this.h,this.s,this.l*t,this.opacity)},rgb:function(){var t=this.h%360+360*(this.h<0),n=isNaN(t)||isNaN(this.s)?0:this.s,e=this.l,r=e+(e<.5?e:1-e)*n,i=2*e-r;return new At(Lt(t>=240?t-240:t+120,i,r),Lt(t,i,r),Lt(t<120?t+240:t-120,i,r),this.opacity)},displayable:function(){return(0<=this.s&&this.s<=1||isNaN(this.s))&&0<=this.l&&this.l<=1&&0<=this.opacity&&this.opacity<=1}}));var Nf=Math.PI/180,Sf=180/Math.PI,Ef=.95047,Af=1,Cf=1.08883,zf=4/29,Pf=6/29,Rf=3*Pf*Pf,Lf=Pf*Pf*Pf;pf(Dt,Ut,wt(Mt,{brighter:function(t){return new Dt(this.l+18*(null==t?1:t),this.a,this.b,this.opacity)},darker:function(t){return new Dt(this.l-18*(null==t?1:t),this.a,this.b,this.opacity)},rgb:function(){var t=(this.l+16)/116,n=isNaN(this.a)?t:t+this.a/500,e=isNaN(this.b)?t:t-this.b/200;return t=Af*Ft(t),n=Ef*Ft(n),e=Cf*Ft(e),new At(It(3.2404542*n-1.5371385*t-.4985314*e),It(-.969266*n+1.8760108*t+.041556*e),It(.0556434*n-.2040259*t+1.0572252*e),this.opacity)}})),pf(Ht,jt,wt(Mt,{brighter:function(t){return new Ht(this.h,this.c,this.l+18*(null==t?1:t),this.opacity)},darker:function(t){return new Ht(this.h,this.c,this.l-18*(null==t?1:t),this.opacity)},rgb:function(){return qt(this).rgb()}}));var qf=-.14861,Uf=1.78277,Df=-.29227,Of=-.90649,Ff=1.97294,If=Ff*Of,Yf=Ff*Uf,Bf=Uf*Df-Of*qf;pf(Vt,$t,wt(Mt,{brighter:function(t){return t=null==t?1/.7:Math.pow(1/.7,t),new Vt(this.h,this.s,this.l*t,this.opacity)},darker:function(t){return t=null==t?.7:Math.pow(.7,t),new Vt(this.h,this.s,this.l*t,this.opacity)},rgb:function(){var t=isNaN(this.h)?0:(this.h+120)*Nf,n=+this.l,e=isNaN(this.s)?0:this.s*n*(1-n),r=Math.cos(t),i=Math.sin(t);return new At(255*(n+e*(qf*r+Uf*i)),255*(n+e*(Df*r+Of*i)),255*(n+e*(Ff*r)),this.opacity)}}));var jf,Hf,Xf,$f,Vf,Wf,Zf=function(t){var n=t.length-1;return function(e){var r=e<=0?e=0:e>=1?(e=1,n-1):Math.floor(e*n),i=t[r],o=t[r+1],u=r>0?t[r-1]:2*i-o,a=r<n-1?t[r+2]:2*o-i;return Wt((e-r/n)*n,u,i,o,a)}},Gf=function(t){var n=t.length;return function(e){var r=Math.floor(((e%=1)<0?++e:e)*n),i=t[(r+n-1)%n],o=t[r%n],u=t[(r+1)%n],a=t[(r+2)%n];return Wt((e-r/n)*n,i,o,u,a)}},Jf=function(t){return function(){return t}},Qf=function t(n){function e(t,n){var e=r((t=Et(t)).r,(n=Et(n)).r),i=r(t.g,n.g),o=r(t.b,n.b),u=Kt(t.opacity,n.opacity);return function(n){return t.r=e(n),t.g=i(n),t.b=o(n),t.opacity=u(n),t+""}}var r=Qt(n);return e.gamma=t,e}(1),Kf=tn(Zf),tl=tn(Gf),nl=function(t,n){var e,r=n?n.length:0,i=t?Math.min(r,t.length):0,o=new Array(r),u=new Array(r);for(e=0;e<i;++e)o[e]=cl(t[e],n[e]);for(;e<r;++e)u[e]=n[e];return function(t){for(e=0;e<i;++e)u[e]=o[e](t);return u}},el=function(t,n){var e=new Date;return t=+t,n-=t,function(r){return e.setTime(t+n*r),e}},rl=function(t,n){return t=+t,n-=t,function(e){return t+n*e}},il=function(t,n){var e,r={},i={};null!==t&&"object"==typeof t||(t={}),null!==n&&"object"==typeof n||(n={});for(e in n)e in t?r[e]=cl(t[e],n[e]):i[e]=n[e];return function(t){for(e in r)i[e]=r[e](t);return i}},ol=/[-+]?(?:\d+\.?\d*|\.?\d+)(?:[eE][-+]?\d+)?/g,ul=new RegExp(ol.source,"g"),al=function(t,n){var e,r,i,o=ol.lastIndex=ul.lastIndex=0,u=-1,a=[],c=[];for(t+="",n+="";(e=ol.exec(t))&&(r=ul.exec(n));)(i=r.index)>o&&(i=n.slice(o,i),a[u]?a[u]+=i:a[++u]=i),(e=e[0])===(r=r[0])?a[u]?a[u]+=r:a[++u]=r:(a[++u]=null,c.push({i:u,x:rl(e,r)})),o=ul.lastIndex;return o<n.length&&(i=n.slice(o),a[u]?a[u]+=i:a[++u]=i),a.length<2?c[0]?en(c[0].x):nn(n):(n=c.length,function(t){for(var e,r=0;r<n;++r)a[(e=c[r]).i]=e.x(t);return a.join("")})},cl=function(t,n){var e,r=typeof n;return null==n||"boolean"===r?Jf(n):("number"===r?rl:"string"===r?(e=Tt(n))?(n=e,Qf):al:n instanceof Tt?Qf:n instanceof Date?el:Array.isArray(n)?nl:"function"!=typeof n.valueOf&&"function"!=typeof n.toString||isNaN(n)?il:rl)(t,n)},sl=function(t,n){return t=+t,n-=t,function(e){return Math.round(t+n*e)}},fl=180/Math.PI,ll={translateX:0,translateY:0,rotate:0,skewX:0,scaleX:1,scaleY:1},hl=function(t,n,e,r,i,o){var u,a,c;return(u=Math.sqrt(t*t+n*n))&&(t/=u,n/=u),(c=t*e+n*r)&&(e-=t*c,r-=n*c),(a=Math.sqrt(e*e+r*r))&&(e/=a,r/=a,c/=a),t*r<n*e&&(t=-t,n=-n,c=-c,u=-u),{translateX:i,translateY:o,rotate:Math.atan2(n,t)*fl,skewX:Math.atan(c)*fl,scaleX:u,scaleY:a}},pl=rn(function(t){return"none"===t?ll:(jf||(jf=document.createElement("DIV"),Hf=document.documentElement,Xf=document.defaultView),jf.style.transform=t,t=Xf.getComputedStyle(Hf.appendChild(jf),null).getPropertyValue("transform"),Hf.removeChild(jf),t=t.slice(7,-1).split(","),hl(+t[0],+t[1],+t[2],+t[3],+t[4],+t[5]))},"px, ","px)","deg)"),dl=rn(function(t){return null==t?ll:($f||($f=document.createElementNS("http://www.w3.org/2000/svg","g")),$f.setAttribute("transform",t),(t=$f.transform.baseVal.consolidate())?(t=t.matrix,hl(t.a,t.b,t.c,t.d,t.e,t.f)):ll)},", ",")",")"),vl=Math.SQRT2,_l=function(t,n){var e,r,i=t[0],o=t[1],u=t[2],a=n[0],c=n[1],s=n[2],f=a-i,l=c-o,h=f*f+l*l;if(h<1e-12)r=Math.log(s/u)/vl,e=function(t){return[i+t*f,o+t*l,u*Math.exp(vl*t*r)]};else{var p=Math.sqrt(h),d=(s*s-u*u+4*h)/(2*u*2*p),v=(s*s-u*u-4*h)/(2*s*2*p),_=Math.log(Math.sqrt(d*d+1)-d),y=Math.log(Math.sqrt(v*v+1)-v);r=(y-_)/vl,e=function(t){var n=t*r,e=on(_),a=u/(2*p)*(e*an(vl*n+_)-un(_));return[i+a*f,o+a*l,u*e/on(vl*n+_)]}}return e.duration=1e3*r,e},yl=cn(Jt),gl=cn(Kt),ml=sn(Jt),xl=sn(Kt),bl=fn(Jt),wl=fn(Kt),Ml=0,Tl=0,kl=0,Nl=1e3,Sl=0,El=0,Al=0,Cl="object"==typeof performance&&performance.now?performance:Date,zl="object"==typeof window&&window.requestAnimationFrame?window.requestAnimationFrame.bind(window):function(t){setTimeout(t,17)};pn.prototype=dn.prototype={constructor:pn,restart:function(t,n,e){if("function"!=typeof t)throw new TypeError("callback is not a function");e=(null==e?ln():+e)+(null==n?0:+n),this._next||Wf===this||(Wf?Wf._next=this:Vf=this,Wf=this),this._call=t,this._time=e,mn()},stop:function(){this._call&&(this._call=null,this._time=1/0,mn())}};var Pl=function(t,n,e){var r=new pn;return n=null==n?0:+n,r.restart(function(e){r.stop(),t(e+n)},n,e),r},Rl=h("start","end","interrupt"),Ll=[],ql=0,Ul=1,Dl=2,Ol=3,Fl=4,Il=5,Yl=6,Bl=function(t,n,e,r,i,o){var u=t.__transition;if(u){if(e in u)return}else t.__transition={};Mn(t,e,{name:n,index:r,group:i,on:Rl,tween:Ll,time:o.time,delay:o.delay,duration:o.duration,ease:o.ease,timer:null,state:ql})},jl=function(t,n){var e,r,i,o=t.__transition,u=!0;if(o){n=null==n?null:n+"";for(i in o)(e=o[i]).name===n?(r=e.state>Dl&&e.state<Il,e.state=Yl,e.timer.stop(),r&&e.on.call("interrupt",t,t.__data__,e.index,e.group),delete o[i]):u=!1;u&&delete t.__transition}},Hl=function(t,n){var e;return("number"==typeof n?rl:n instanceof Tt?Qf:(e=Tt(n))?(n=e,Qf):al)(t,n)},Xl=dt.prototype.constructor,$l=0,Vl=dt.prototype;Gn.prototype=Jn.prototype={constructor:Gn,select:function(t){var n=this._name,e=this._id;"function"!=typeof t&&(t=tf(t));for(var r=this._groups,i=r.length,o=new Array(i),u=0;u<i;++u)for(var a,c,s=r[u],f=s.length,l=o[u]=new Array(f),h=0;h<f;++h)(a=s[h])&&(c=t.call(a,a.__data__,h,s))&&("__data__"in a&&(c.__data__=a.__data__),l[h]=c,Bl(l[h],n,e,h,l,wn(a,e)));return new Gn(o,this._parents,n,e)},selectAll:function(t){var n=this._name,e=this._id;"function"!=typeof t&&(t=nf(t));for(var r=this._groups,i=r.length,o=[],u=[],a=0;a<i;++a)for(var c,s=r[a],f=s.length,l=0;l<f;++l)if(c=s[l]){for(var h,p=t.call(c,c.__data__,l,s),d=wn(c,e),v=0,_=p.length;v<_;++v)(h=p[v])&&Bl(h,n,e,v,p,d);o.push(p),u.push(c)}return new Gn(o,u,n,e)},filter:function(t){"function"!=typeof t&&(t=Zs(t));for(var n=this._groups,e=n.length,r=new Array(e),i=0;i<e;++i)for(var o,u=n[i],a=u.length,c=r[i]=[],s=0;s<a;++s)(o=u[s])&&t.call(o,o.__data__,s,u)&&c.push(o);return new Gn(r,this._parents,this._name,this._id)},merge:function(t){if(t._id!==this._id)throw new Error;for(var n=this._groups,e=t._groups,r=n.length,i=e.length,o=Math.min(r,i),u=new Array(r),a=0;a<o;++a)for(var c,s=n[a],f=e[a],l=s.length,h=u[a]=new Array(l),p=0;p<l;++p)(c=s[p]||f[p])&&(h[p]=c);for(;a<r;++a)u[a]=n[a];return new Gn(u,this._parents,this._name,this._id)},selection:function(){return new Xl(this._groups,this._parents)},transition:function(){for(var t=this._name,n=this._id,e=Qn(),r=this._groups,i=r.length,o=0;o<i;++o)for(var u,a=r[o],c=a.length,s=0;s<c;++s)if(u=a[s]){var f=wn(u,n);Bl(u,t,e,s,a,{time:f.time+f.delay+f.duration,delay:0,duration:f.duration,ease:f.ease})}return new Gn(r,this._parents,t,e)},call:Vl.call,nodes:Vl.nodes,node:Vl.node,size:Vl.size,empty:Vl.empty,each:Vl.each,on:function(t,n){var e=this._id;return arguments.length<2?wn(this.node(),e).on.on(t):this.each(Yn(e,t,n))},attr:function(t,n){var e=js(t),r="transform"===e?dl:Hl;return this.attrTween(t,"function"==typeof n?(e.local?Pn:zn)(e,r,Nn(this,"attr."+t,n)):null==n?(e.local?En:Sn)(e):(e.local?Cn:An)(e,r,n+""))},attrTween:function(t,n){var e="attr."+t;if(arguments.length<2)return(e=this.tween(e))&&e._value;if(null==n)return this.tween(e,null);if("function"!=typeof n)throw new Error;var r=js(t);return this.tween(e,(r.local?Rn:Ln)(r,n))},style:function(t,n,e){var r="transform"==(t+="")?pl:Hl;return null==n?this.styleTween(t,jn(t,r)).on("end.style."+t,Hn(t)):this.styleTween(t,"function"==typeof n?$n(t,r,Nn(this,"style."+t,n)):Xn(t,r,n+""),e)},styleTween:function(t,n,e){var r="style."+(t+="");if(arguments.length<2)return(r=this.tween(r))&&r._value;if(null==n)return this.tween(r,null);if("function"!=typeof n)throw new Error;return this.tween(r,Vn(t,n,null==e?"":e))},text:function(t){return this.tween("text","function"==typeof t?Zn(Nn(this,"text",t)):Wn(null==t?"":t+""))},remove:function(){return this.on("end.remove",Bn(this._id))},tween:function(t,n){var e=this._id;if(t+="",arguments.length<2){for(var r,i=wn(this.node(),e).tween,o=0,u=i.length;o<u;++o)if((r=i[o]).name===t)return r.value;return null}return this.each((null==n?Tn:kn)(e,t,n))},delay:function(t){var n=this._id;return arguments.length?this.each(("function"==typeof t?qn:Un)(n,t)):wn(this.node(),n).delay},duration:function(t){var n=this._id;return arguments.length?this.each(("function"==typeof t?Dn:On)(n,t)):wn(this.node(),n).duration},ease:function(t){var n=this._id;return arguments.length?this.each(Fn(n,t)):wn(this.node(),n).ease}};var Wl=function t(n){function e(t){return Math.pow(t,n)}return n=+n,e.exponent=t,e}(3),Zl=function t(n){function e(t){return 1-Math.pow(1-t,n)}return n=+n,e.exponent=t,e}(3),Gl=function t(n){function e(t){return((t*=2)<=1?Math.pow(t,n):2-Math.pow(2-t,n))/2}return n=+n,e.exponent=t,e}(3),Jl=Math.PI,Ql=Jl/2,Kl=4/11,th=6/11,nh=8/11,eh=.75,rh=9/11,ih=10/11,oh=.9375,uh=21/22,ah=63/64,ch=1/Kl/Kl,sh=function t(n){function e(t){return t*t*((n+1)*t-n)}return n=+n,e.overshoot=t,e}(1.70158),fh=function t(n){function e(t){return--t*t*((n+1)*t+n)+1}return n=+n,e.overshoot=t,e}(1.70158),lh=function t(n){function e(t){return((t*=2)<1?t*t*((n+1)*t-n):(t-=2)*t*((n+1)*t+n)+2)/2}return n=+n,e.overshoot=t,e}(1.70158),hh=2*Math.PI,ph=function t(n,e){function r(t){return n*Math.pow(2,10*--t)*Math.sin((i-t)/e)}var i=Math.asin(1/(n=Math.max(1,n)))*(e/=hh);return r.amplitude=function(n){return t(n,e*hh)},r.period=function(e){return t(n,e)},r}(1,.3),dh=function t(n,e){function r(t){return 1-n*Math.pow(2,-10*(t=+t))*Math.sin((t+i)/e)}var i=Math.asin(1/(n=Math.max(1,n)))*(e/=hh);return r.amplitude=function(n){return t(n,e*hh)},r.period=function(e){return t(n,e)},r}(1,.3),vh=function t(n,e){function r(t){return((t=2*t-1)<0?n*Math.pow(2,10*t)*Math.sin((i-t)/e):2-n*Math.pow(2,-10*t)*Math.sin((i+t)/e))/2}var i=Math.asin(1/(n=Math.max(1,n)))*(e/=hh);return r.amplitude=function(n){return t(n,e*hh)},r.period=function(e){return t(n,e)},r}(1,.3),_h={time:null,delay:0,duration:250,ease:te};dt.prototype.interrupt=function(t){return this.each(function(){jl(this,t)})},dt.prototype.transition=function(t){var n,e;t instanceof Gn?(n=t._id,t=t._name):(n=Qn(),(e=_h).time=ln(),t=null==t?null:t+"");for(var r=this._groups,i=r.length,o=0;o<i;++o)for(var u,a=r[o],c=a.length,s=0;s<c;++s)(u=a[s])&&Bl(u,t,n,s,a,e||oe(u,n));return new Gn(r,this._parents,t,n)};var yh=[null],gh=function(t){return function(){return t}},mh=function(t,n,e){this.target=t,this.type=n,this.selection=e},xh=function(){t.event.preventDefault(),t.event.stopImmediatePropagation()},bh={name:"drag"},wh={name:"space"},Mh={name:"handle"},Th={name:"center"},kh={name:"x",handles:["e","w"].map(ae),input:function(t,n){return t&&[[t[0],n[0][1]],[t[1],n[1][1]]]},output:function(t){return t&&[t[0][0],t[1][0]]}},Nh={name:"y",handles:["n","s"].map(ae),input:function(t,n){return t&&[[n[0][0],t[0]],[n[1][0],t[1]]]},output:function(t){return t&&[t[0][1],t[1][1]]}},Sh={name:"xy",handles:["n","e","s","w","nw","ne","se","sw"].map(ae),input:function(t){return t},output:function(t){return t}},Eh={overlay:"crosshair",selection:"move",n:"ns-resize",e:"ew-resize",s:"ns-resize",w:"ew-resize",nw:"nwse-resize",ne:"nesw-resize",se:"nwse-resize",sw:"nesw-resize"},Ah={e:"w",w:"e",nw:"ne",ne:"nw",se:"sw",sw:"se"},Ch={n:"s",s:"n",nw:"sw",ne:"se",se:"ne",sw:"nw"},zh={overlay:1,selection:1,n:null,e:1,s:null,w:-1,nw:-1,ne:1,se:1,sw:-1},Ph={overlay:1,selection:1,n:-1,e:null,s:1,w:null,nw:-1,ne:-1,se:1,sw:1},Rh=Math.cos,Lh=Math.sin,qh=Math.PI,Uh=qh/2,Dh=2*qh,Oh=Math.max,Fh=Array.prototype.slice,Ih=function(t){return function(){return t}},Yh=Math.PI,Bh=2*Yh,jh=Bh-1e-6;de.prototype=ve.prototype={constructor:de,moveTo:function(t,n){this._+="M"+(this._x0=this._x1=+t)+","+(this._y0=this._y1=+n)},closePath:function(){null!==this._x1&&(this._x1=this._x0,this._y1=this._y0,this._+="Z")},lineTo:function(t,n){this._+="L"+(this._x1=+t)+","+(this._y1=+n)},quadraticCurveTo:function(t,n,e,r){this._+="Q"+ +t+","+ +n+","+(this._x1=+e)+","+(this._y1=+r)},bezierCurveTo:function(t,n,e,r,i,o){this._+="C"+ +t+","+ +n+","+ +e+","+ +r+","+(this._x1=+i)+","+(this._y1=+o)},arcTo:function(t,n,e,r,i){t=+t,n=+n,e=+e,r=+r,i=+i;var o=this._x1,u=this._y1,a=e-t,c=r-n,s=o-t,f=u-n,l=s*s+f*f;if(i<0)throw new Error("negative radius: "+i);if(null===this._x1)this._+="M"+(this._x1=t)+","+(this._y1=n);else if(l>1e-6)if(Math.abs(f*a-c*s)>1e-6&&i){var h=e-o,p=r-u,d=a*a+c*c,v=h*h+p*p,_=Math.sqrt(d),y=Math.sqrt(l),g=i*Math.tan((Yh-Math.acos((d+l-v)/(2*_*y)))/2),m=g/y,x=g/_;Math.abs(m-1)>1e-6&&(this._+="L"+(t+m*s)+","+(n+m*f)),this._+="A"+i+","+i+",0,0,"+ +(f*h>s*p)+","+(this._x1=t+x*a)+","+(this._y1=n+x*c)}else this._+="L"+(this._x1=t)+","+(this._y1=n);else;},arc:function(t,n,e,r,i,o){t=+t,n=+n;var u=(e=+e)*Math.cos(r),a=e*Math.sin(r),c=t+u,s=n+a,f=1^o,l=o?r-i:i-r;if(e<0)throw new Error("negative radius: "+e);null===this._x1?this._+="M"+c+","+s:(Math.abs(this._x1-c)>1e-6||Math.abs(this._y1-s)>1e-6)&&(this._+="L"+c+","+s),e&&(l<0&&(l=l%Bh+Bh),l>jh?this._+="A"+e+","+e+",0,1,"+f+","+(t-u)+","+(n-a)+"A"+e+","+e+",0,1,"+f+","+(this._x1=c)+","+(this._y1=s):l>1e-6&&(this._+="A"+e+","+e+",0,"+ +(l>=Yh)+","+f+","+(this._x1=t+e*Math.cos(i))+","+(this._y1=n+e*Math.sin(i))))},rect:function(t,n,e,r){this._+="M"+(this._x0=this._x1=+t)+","+(this._y0=this._y1=+n)+"h"+ +e+"v"+ +r+"h"+-e+"Z"},toString:function(){return this._}};be.prototype=we.prototype={constructor:be,has:function(t){return"$"+t in this},get:function(t){return this["$"+t]},set:function(t,n){return this["$"+t]=n,this},remove:function(t){var n="$"+t;return n in this&&delete this[n]},clear:function(){for(var t in this)"$"===t[0]&&delete this[t]},keys:function(){var t=[];for(var n in this)"$"===n[0]&&t.push(n.slice(1));return t},values:function(){var t=[];for(var n in this)"$"===n[0]&&t.push(this[n]);return t},entries:function(){var t=[];for(var n in this)"$"===n[0]&&t.push({key:n.slice(1),value:this[n]});return t},size:function(){var t=0;for(var n in this)"$"===n[0]&&++t;return t},empty:function(){for(var t in this)if("$"===t[0])return!1;return!0},each:function(t){for(var n in this)"$"===n[0]&&t(this[n],n.slice(1),this)}};var Hh=we.prototype;Se.prototype=Ee.prototype={constructor:Se,has:Hh.has,add:function(t){return t+="",this["$"+t]=t,this},remove:Hh.remove,clear:Hh.clear,values:Hh.keys,size:Hh.size,empty:Hh.empty,each:Hh.each};var Xh=function(t){function n(t,n){function e(){if(f>=s)return a;if(i)return i=!1,u;var n,e=f;if(34===t.charCodeAt(e)){for(var r=e;r++<s;)if(34===t.charCodeAt(r)){if(34!==t.charCodeAt(r+1))break;++r}return f=r+2,13===(n=t.charCodeAt(r+1))?(i=!0,10===t.charCodeAt(r+2)&&++f):10===n&&(i=!0),t.slice(e+1,r).replace(/""/g,'"')}for(;f<s;){var c=1;if(10===(n=t.charCodeAt(f++)))i=!0;else if(13===n)i=!0,10===t.charCodeAt(f)&&(++f,++c);else if(n!==o)continue;return t.slice(e,f-c)}return t.slice(e)}for(var r,i,u={},a={},c=[],s=t.length,f=0,l=0;(r=e())!==a;){for(var h=[];r!==u&&r!==a;)h.push(r),r=e();n&&null==(h=n(h,l++))||c.push(h)}return c}function e(n){return n.map(r).join(t)}function r(t){return null==t?"":i.test(t+="")?'"'+t.replace(/\"/g,'""')+'"':t}var i=new RegExp('["'+t+"\n\r]"),o=t.charCodeAt(0);return{parse:function(t,e){var r,i,o=n(t,function(t,n){if(r)return r(t,n-1);i=t,r=e?Ce(t,e):Ae(t)});return o.columns=i,o},parseRows:n,format:function(n,e){return null==e&&(e=ze(n)),[e.map(r).join(t)].concat(n.map(function(n){return e.map(function(t){return r(n[t])}).join(t)})).join("\n")},formatRows:function(t){return t.map(e).join("\n")}}},$h=Xh(","),Vh=$h.parse,Wh=$h.parseRows,Zh=$h.format,Gh=$h.formatRows,Jh=Xh("\t"),Qh=Jh.parse,Kh=Jh.parseRows,tp=Jh.format,np=Jh.formatRows,ep=function(t){return function(){return t}},rp=function(){return 1e-6*(Math.random()-.5)},ip=function(t,n,e,r,i){this.node=t,this.x0=n,this.y0=e,this.x1=r,this.y1=i},op=qe.prototype=Ue.prototype;op.copy=function(){var t,n,e=new Ue(this._x,this._y,this._x0,this._y0,this._x1,this._y1),r=this._root;if(!r)return e;if(!r.length)return e._root=De(r),e;for(t=[{source:r,target:e._root=new Array(4)}];r=t.pop();)for(var i=0;i<4;++i)(n=r.source[i])&&(n.length?t.push({source:n,target:r.target[i]=new Array(4)}):r.target[i]=De(n));return e},op.add=function(t){var n=+this._x.call(null,t),e=+this._y.call(null,t);return Pe(this.cover(n,e),n,e,t)},op.addAll=function(t){var n,e,r,i,o=t.length,u=new Array(o),a=new Array(o),c=1/0,s=1/0,f=-1/0,l=-1/0;for(e=0;e<o;++e)isNaN(r=+this._x.call(null,n=t[e]))||isNaN(i=+this._y.call(null,n))||(u[e]=r,a[e]=i,r<c&&(c=r),r>f&&(f=r),i<s&&(s=i),i>l&&(l=i));for(f<c&&(c=this._x0,f=this._x1),l<s&&(s=this._y0,l=this._y1),this.cover(c,s).cover(f,l),e=0;e<o;++e)Pe(this,u[e],a[e],t[e]);return this},op.cover=function(t,n){if(isNaN(t=+t)||isNaN(n=+n))return this;var e=this._x0,r=this._y0,i=this._x1,o=this._y1;if(isNaN(e))i=(e=Math.floor(t))+1,o=(r=Math.floor(n))+1;else{if(!(e>t||t>i||r>n||n>o))return this;var u,a,c=i-e,s=this._root;switch(a=(n<(r+o)/2)<<1|t<(e+i)/2){case 0:do{u=new Array(4),u[a]=s,s=u}while(c*=2,i=e+c,o=r+c,t>i||n>o);break;case 1:do{u=new Array(4),u[a]=s,s=u}while(c*=2,e=i-c,o=r+c,e>t||n>o);break;case 2:do{u=new Array(4),u[a]=s,s=u}while(c*=2,i=e+c,r=o-c,t>i||r>n);break;case 3:do{u=new Array(4),u[a]=s,s=u}while(c*=2,e=i-c,r=o-c,e>t||r>n)}this._root&&this._root.length&&(this._root=s)}return this._x0=e,this._y0=r,this._x1=i,this._y1=o,this},op.data=function(){var t=[];return this.visit(function(n){if(!n.length)do{t.push(n.data)}while(n=n.next)}),t},op.extent=function(t){return arguments.length?this.cover(+t[0][0],+t[0][1]).cover(+t[1][0],+t[1][1]):isNaN(this._x0)?void 0:[[this._x0,this._y0],[this._x1,this._y1]]},op.find=function(t,n,e){var r,i,o,u,a,c,s,f=this._x0,l=this._y0,h=this._x1,p=this._y1,d=[],v=this._root;for(v&&d.push(new ip(v,f,l,h,p)),null==e?e=1/0:(f=t-e,l=n-e,h=t+e,p=n+e,e*=e);c=d.pop();)if(!(!(v=c.node)||(i=c.x0)>h||(o=c.y0)>p||(u=c.x1)<f||(a=c.y1)<l))if(v.length){var _=(i+u)/2,y=(o+a)/2;d.push(new ip(v[3],_,y,u,a),new ip(v[2],i,y,_,a),new ip(v[1],_,o,u,y),new ip(v[0],i,o,_,y)),(s=(n>=y)<<1|t>=_)&&(c=d[d.length-1],d[d.length-1]=d[d.length-1-s],d[d.length-1-s]=c)}else{var g=t-+this._x.call(null,v.data),m=n-+this._y.call(null,v.data),x=g*g+m*m;if(x<e){var b=Math.sqrt(e=x);f=t-b,l=n-b,h=t+b,p=n+b,r=v.data}}return r},op.remove=function(t){if(isNaN(o=+this._x.call(null,t))||isNaN(u=+this._y.call(null,t)))return this;var n,e,r,i,o,u,a,c,s,f,l,h,p=this._root,d=this._x0,v=this._y0,_=this._x1,y=this._y1;if(!p)return this;if(p.length)for(;;){if((s=o>=(a=(d+_)/2))?d=a:_=a,(f=u>=(c=(v+y)/2))?v=c:y=c,n=p,!(p=p[l=f<<1|s]))return this;if(!p.length)break;(n[l+1&3]||n[l+2&3]||n[l+3&3])&&(e=n,h=l)}for(;p.data!==t;)if(r=p,!(p=p.next))return this;return(i=p.next)&&delete p.next,r?(i?r.next=i:delete r.next,this):n?(i?n[l]=i:delete n[l],(p=n[0]||n[1]||n[2]||n[3])&&p===(n[3]||n[2]||n[1]||n[0])&&!p.length&&(e?e[h]=p:this._root=p),this):(this._root=i,this)},op.removeAll=function(t){for(var n=0,e=t.length;n<e;++n)this.remove(t[n]);return this},op.root=function(){return this._root},op.size=function(){var t=0;return this.visit(function(n){if(!n.length)do{++t}while(n=n.next)}),t},op.visit=function(t){var n,e,r,i,o,u,a=[],c=this._root;for(c&&a.push(new ip(c,this._x0,this._y0,this._x1,this._y1));n=a.pop();)if(!t(c=n.node,r=n.x0,i=n.y0,o=n.x1,u=n.y1)&&c.length){var s=(r+o)/2,f=(i+u)/2;(e=c[3])&&a.push(new ip(e,s,f,o,u)),(e=c[2])&&a.push(new ip(e,r,f,s,u)),(e=c[1])&&a.push(new ip(e,s,i,o,f)),(e=c[0])&&a.push(new ip(e,r,i,s,f))}return this},op.visitAfter=function(t){var n,e=[],r=[];for(this._root&&e.push(new ip(this._root,this._x0,this._y0,this._x1,this._y1));n=e.pop();){var i=n.node;if(i.length){var o,u=n.x0,a=n.y0,c=n.x1,s=n.y1,f=(u+c)/2,l=(a+s)/2;(o=i[0])&&e.push(new ip(o,u,a,f,l)),(o=i[1])&&e.push(new ip(o,f,a,c,l)),(o=i[2])&&e.push(new ip(o,u,l,f,s)),(o=i[3])&&e.push(new ip(o,f,l,c,s))}r.push(n)}for(;n=r.pop();)t(n.node,n.x0,n.y0,n.x1,n.y1);return this},op.x=function(t){return arguments.length?(this._x=t,this):this._x},op.y=function(t){return arguments.length?(this._y=t,this):this._y};var up,ap=10,cp=Math.PI*(3-Math.sqrt(5)),sp=function(t,n){if((e=(t=n?t.toExponential(n-1):t.toExponential()).indexOf("e"))<0)return null;var e,r=t.slice(0,e);return[r.length>1?r[0]+r.slice(2):r,+t.slice(e+1)]},fp=function(t){return(t=sp(Math.abs(t)))?t[1]:NaN},lp=function(t,n){return function(e,r){for(var i=e.length,o=[],u=0,a=t[0],c=0;i>0&&a>0&&(c+a+1>r&&(a=Math.max(1,r-c)),o.push(e.substring(i-=a,i+a)),!((c+=a+1)>r));)a=t[u=(u+1)%t.length];return o.reverse().join(n)}},hp=function(t){return function(n){return n.replace(/[0-9]/g,function(n){return t[+n]})}},pp=function(t,n){var e=sp(t,n);if(!e)return t+"";var r=e[0],i=e[1];return i<0?"0."+new Array(-i).join("0")+r:r.length>i+1?r.slice(0,i+1)+"."+r.slice(i+1):r+new Array(i-r.length+2).join("0")},dp={"":function(t,n){t:for(var e,r=(t=t.toPrecision(n)).length,i=1,o=-1;i<r;++i)switch(t[i]){case".":o=e=i;break;case"0":0===o&&(o=i),e=i;break;case"e":break t;default:o>0&&(o=0)}return o>0?t.slice(0,o)+t.slice(e+1):t},"%":function(t,n){return(100*t).toFixed(n)},b:function(t){return Math.round(t).toString(2)},c:function(t){return t+""},d:function(t){return Math.round(t).toString(10)},e:function(t,n){return t.toExponential(n)},f:function(t,n){return t.toFixed(n)},g:function(t,n){return t.toPrecision(n)},o:function(t){return Math.round(t).toString(8)},p:function(t,n){return pp(100*t,n)},r:pp,s:function(t,n){var e=sp(t,n);if(!e)return t+"";var r=e[0],i=e[1],o=i-(up=3*Math.max(-8,Math.min(8,Math.floor(i/3))))+1,u=r.length;return o===u?r:o>u?r+new Array(o-u+1).join("0"):o>0?r.slice(0,o)+"."+r.slice(o):"0."+new Array(1-o).join("0")+sp(t,Math.max(0,n+o-1))[0]},X:function(t){return Math.round(t).toString(16).toUpperCase()},x:function(t){return Math.round(t).toString(16)}},vp=/^(?:(.)?([<>=^]))?([+\-\( ])?([$#])?(0)?(\d+)?(,)?(\.\d+)?([a-z%])?$/i;He.prototype=Xe.prototype,Xe.prototype.toString=function(){return this.fill+this.align+this.sign+this.symbol+(this.zero?"0":"")+(null==this.width?"":Math.max(1,0|this.width))+(this.comma?",":"")+(null==this.precision?"":"."+Math.max(0,0|this.precision))+this.type};var _p,yp=function(t){return t},gp=["y","z","a","f","p","n","µ","m","","k","M","G","T","P","E","Z","Y"],mp=function(t){function n(t){function n(t){var n,r,u,f=_,x=y;if("c"===v)x=g(t)+x,t="";else{var b=(t=+t)<0;if(t=g(Math.abs(t),d),b&&0==+t&&(b=!1),f=(b?"("===s?s:"-":"-"===s||"("===s?"":s)+f,x=x+("s"===v?gp[8+up/3]:"")+(b&&"("===s?")":""),m)for(n=-1,r=t.length;++n<r;)if(48>(u=t.charCodeAt(n))||u>57){x=(46===u?i+t.slice(n+1):t.slice(n))+x,t=t.slice(0,n);break}}p&&!l&&(t=e(t,1/0));var w=f.length+t.length+x.length,M=w<h?new Array(h-w+1).join(a):"";switch(p&&l&&(t=e(M+t,M.length?h-x.length:1/0),M=""),c){case"<":t=f+t+x+M;break;case"=":t=f+M+t+x;break;case"^":t=M.slice(0,w=M.length>>1)+f+t+x+M.slice(w);break;default:t=M+f+t+x}return o(t)}var a=(t=He(t)).fill,c=t.align,s=t.sign,f=t.symbol,l=t.zero,h=t.width,p=t.comma,d=t.precision,v=t.type,_="$"===f?r[0]:"#"===f&&/[boxX]/.test(v)?"0"+v.toLowerCase():"",y="$"===f?r[1]:/[%p]/.test(v)?u:"",g=dp[v],m=!v||/[defgprs%]/.test(v);return d=null==d?v?6:12:/[gprs]/.test(v)?Math.max(1,Math.min(21,d)):Math.max(0,Math.min(20,d)),n.toString=function(){return t+""},n}var e=t.grouping&&t.thousands?lp(t.grouping,t.thousands):yp,r=t.currency,i=t.decimal,o=t.numerals?hp(t.numerals):yp,u=t.percent||"%";return{format:n,formatPrefix:function(t,e){var r=n((t=He(t),t.type="f",t)),i=3*Math.max(-8,Math.min(8,Math.floor(fp(e)/3))),o=Math.pow(10,-i),u=gp[8+i/3];return function(t){return r(o*t)+u}}}};$e({decimal:".",thousands:",",grouping:[3],currency:["$",""]});var xp=function(t){return Math.max(0,-fp(Math.abs(t)))},bp=function(t,n){return Math.max(0,3*Math.max(-8,Math.min(8,Math.floor(fp(n)/3)))-fp(Math.abs(t)))},wp=function(t,n){return t=Math.abs(t),n=Math.abs(n)-t,Math.max(0,fp(n)-fp(t))+1},Mp=function(){return new Ve};Ve.prototype={constructor:Ve,reset:function(){this.s=this.t=0},add:function(t){We(nd,t,this.t),We(this,nd.s,this.s),this.s?this.t+=nd.t:this.s=nd.t},valueOf:function(){return this.s}};var Tp,kp,Np,Sp,Ep,Ap,Cp,zp,Pp,Rp,Lp,qp,Up,Dp,Op,Fp,Ip,Yp,Bp,jp,Hp,Xp,$p,Vp,Wp,Zp,Gp,Jp,Qp,Kp,td,nd=new Ve,ed=1e-6,rd=Math.PI,id=rd/2,od=rd/4,ud=2*rd,ad=180/rd,cd=rd/180,sd=Math.abs,fd=Math.atan,ld=Math.atan2,hd=Math.cos,pd=Math.ceil,dd=Math.exp,vd=Math.log,_d=Math.pow,yd=Math.sin,gd=Math.sign||function(t){return t>0?1:t<0?-1:0},md=Math.sqrt,xd=Math.tan,bd={Feature:function(t,n){Ke(t.geometry,n)},FeatureCollection:function(t,n){for(var e=t.features,r=-1,i=e.length;++r<i;)Ke(e[r].geometry,n)}},wd={Sphere:function(t,n){n.sphere()},Point:function(t,n){t=t.coordinates,n.point(t[0],t[1],t[2])},MultiPoint:function(t,n){for(var e=t.coordinates,r=-1,i=e.length;++r<i;)t=e[r],n.point(t[0],t[1],t[2])},LineString:function(t,n){tr(t.coordinates,n,0)},MultiLineString:function(t,n){for(var e=t.coordinates,r=-1,i=e.length;++r<i;)tr(e[r],n,0)},Polygon:function(t,n){nr(t.coordinates,n)},MultiPolygon:function(t,n){for(var e=t.coordinates,r=-1,i=e.length;++r<i;)nr(e[r],n)},GeometryCollection:function(t,n){for(var e=t.geometries,r=-1,i=e.length;++r<i;)Ke(e[r],n)}},Md=function(t,n){t&&bd.hasOwnProperty(t.type)?bd[t.type](t,n):Ke(t,n)},Td=Mp(),kd=Mp(),Nd={point:Qe,lineStart:Qe,lineEnd:Qe,polygonStart:function(){Td.reset(),Nd.lineStart=er,Nd.lineEnd=rr},polygonEnd:function(){var t=+Td;kd.add(t<0?ud+t:t),this.lineStart=this.lineEnd=this.point=Qe},sphere:function(){kd.add(ud)}},Sd=Mp(),Ed={point:pr,lineStart:vr,lineEnd:_r,polygonStart:function(){Ed.point=yr,Ed.lineStart=gr,Ed.lineEnd=mr,Sd.reset(),Nd.polygonStart()},polygonEnd:function(){Nd.polygonEnd(),Ed.point=pr,Ed.lineStart=vr,Ed.lineEnd=_r,Td<0?(Ap=-(zp=180),Cp=-(Pp=90)):Sd>ed?Pp=90:Sd<-ed&&(Cp=-90),Op[0]=Ap,Op[1]=zp}},Ad={sphere:Qe,point:Mr,lineStart:kr,lineEnd:Er,polygonStart:function(){Ad.lineStart=Ar,Ad.lineEnd=Cr},polygonEnd:function(){Ad.lineStart=kr,Ad.lineEnd=Er}},Cd=function(t){return function(){return t}},zd=function(t,n){function e(e,r){return e=t(e,r),n(e[0],e[1])}return t.invert&&n.invert&&(e.invert=function(e,r){return(e=n.invert(e,r))&&t.invert(e[0],e[1])}),e};Rr.invert=Rr;var Pd,Rd,Ld,qd,Ud,Dd,Od,Fd,Id,Yd,Bd,jd=function(t){function n(n){return n=t(n[0]*cd,n[1]*cd),n[0]*=ad,n[1]*=ad,n}return t=Lr(t[0]*cd,t[1]*cd,t.length>2?t[2]*cd:0),n.invert=function(n){return n=t.invert(n[0]*cd,n[1]*cd),n[0]*=ad,n[1]*=ad,n},n},Hd=function(){var t,n=[];return{point:function(n,e){t.push([n,e])},lineStart:function(){n.push(t=[])},lineEnd:Qe,rejoin:function(){n.length>1&&n.push(n.pop().concat(n.shift()))},result:function(){var e=n;return n=[],t=null,e}}},Xd=function(t,n,e,r,i,o){var u,a=t[0],c=t[1],s=0,f=1,l=n[0]-a,h=n[1]-c;if(u=e-a,l||!(u>0)){if(u/=l,l<0){if(u<s)return;u<f&&(f=u)}else if(l>0){if(u>f)return;u>s&&(s=u)}if(u=i-a,l||!(u<0)){if(u/=l,l<0){if(u>f)return;u>s&&(s=u)}else if(l>0){if(u<s)return;u<f&&(f=u)}if(u=r-c,h||!(u>0)){if(u/=h,h<0){if(u<s)return;u<f&&(f=u)}else if(h>0){if(u>f)return;u>s&&(s=u)}if(u=o-c,h||!(u<0)){if(u/=h,h<0){if(u>f)return;u>s&&(s=u)}else if(h>0){if(u<s)return;u<f&&(f=u)}return s>0&&(t[0]=a+s*l,t[1]=c+s*h),f<1&&(n[0]=a+f*l,n[1]=c+f*h),!0}}}}},$d=function(t,n){return sd(t[0]-n[0])<ed&&sd(t[1]-n[1])<ed},Vd=function(t,n,e,r,i){var o,u,a=[],c=[];if(t.forEach(function(t){if(!((n=t.length-1)<=0)){var n,e,r=t[0],u=t[n];if($d(r,u)){for(i.lineStart(),o=0;o<n;++o)i.point((r=t[o])[0],r[1]);i.lineEnd()}else a.push(e=new Ir(r,t,null,!0)),c.push(e.o=new Ir(r,null,e,!1)),a.push(e=new Ir(u,t,null,!1)),c.push(e.o=new Ir(u,null,e,!0))}}),a.length){for(c.sort(n),Yr(a),Yr(c),o=0,u=c.length;o<u;++o)c[o].e=e=!e;for(var s,f,l=a[0];;){for(var h=l,p=!0;h.v;)if((h=h.n)===l)return;s=h.z,i.lineStart();do{if(h.v=h.o.v=!0,h.e){if(p)for(o=0,u=s.length;o<u;++o)i.point((f=s[o])[0],f[1]);else r(h.x,h.n.x,1,i);h=h.n}else{if(p)for(s=h.p.z,o=s.length-1;o>=0;--o)i.point((f=s[o])[0],f[1]);else r(h.x,h.p.x,-1,i);h=h.p}s=(h=h.o).z,p=!p}while(!h.v);i.lineEnd()}}},Wd=1e9,Zd=-Wd,Gd=Mp(),Jd=function(t,n){var e=n[0],r=n[1],i=[yd(e),-hd(e),0],o=0,u=0;Gd.reset();for(var a=0,c=t.length;a<c;++a)if(f=(s=t[a]).length)for(var s,f,l=s[f-1],h=l[0],p=l[1]/2+od,d=yd(p),v=hd(p),_=0;_<f;++_,h=g,d=x,v=b,l=y){var y=s[_],g=y[0],m=y[1]/2+od,x=yd(m),b=hd(m),w=g-h,M=w>=0?1:-1,T=M*w,k=T>rd,N=d*x;if(Gd.add(ld(N*M*yd(T),v*b+N*hd(T))),o+=k?w+M*ud:w,k^h>=e^g>=e){var S=sr(ar(l),ar(y));hr(S);var E=sr(i,S);hr(E);var A=(k^w>=0?-1:1)*Ge(E[2]);(r>A||r===A&&(S[0]||S[1]))&&(u+=k^w>=0?1:-1)}}return(o<-ed||o<ed&&Gd<-ed)^1&u},Qd=Mp(),Kd={sphere:Qe,point:Qe,lineStart:function(){Kd.point=Hr,Kd.lineEnd=jr},lineEnd:Qe,polygonStart:Qe,polygonEnd:Qe},tv=function(t){return Qd.reset(),Md(t,Kd),+Qd},nv=[null,null],ev={type:"LineString",coordinates:nv},rv=function(t,n){return nv[0]=t,nv[1]=n,tv(ev)},iv={Feature:function(t,n){return $r(t.geometry,n)},FeatureCollection:function(t,n){for(var e=t.features,r=-1,i=e.length;++r<i;)if($r(e[r].geometry,n))return!0;return!1}},ov={Sphere:function(){return!0},Point:function(t,n){return Vr(t.coordinates,n)},MultiPoint:function(t,n){for(var e=t.coordinates,r=-1,i=e.length;++r<i;)if(Vr(e[r],n))return!0;return!1},LineString:function(t,n){return Wr(t.coordinates,n)},MultiLineString:function(t,n){for(var e=t.coordinates,r=-1,i=e.length;++r<i;)if(Wr(e[r],n))return!0;return!1},Polygon:function(t,n){return Zr(t.coordinates,n)},MultiPolygon:function(t,n){for(var e=t.coordinates,r=-1,i=e.length;++r<i;)if(Zr(e[r],n))return!0;return!1},GeometryCollection:function(t,n){for(var e=t.geometries,r=-1,i=e.length;++r<i;)if($r(e[r],n))return!0;return!1}},uv=function(t){return t},av=Mp(),cv=Mp(),sv={point:Qe,lineStart:Qe,lineEnd:Qe,polygonStart:function(){sv.lineStart=ni,sv.lineEnd=ii},polygonEnd:function(){sv.lineStart=sv.lineEnd=sv.point=Qe,av.add(sd(cv)),cv.reset()},result:function(){var t=av/2;return av.reset(),t}},fv=1/0,lv=fv,hv=-fv,pv=hv,dv={point:function(t,n){t<fv&&(fv=t),t>hv&&(hv=t),n<lv&&(lv=n),n>pv&&(pv=n)},lineStart:Qe,lineEnd:Qe,polygonStart:Qe,polygonEnd:Qe,result:function(){var t=[[fv,lv],[hv,pv]];return hv=pv=-(lv=fv=1/0),t}},vv=0,_v=0,yv=0,gv=0,mv=0,xv=0,bv=0,wv=0,Mv=0,Tv={point:oi,lineStart:ui,lineEnd:si,polygonStart:function(){Tv.lineStart=fi,Tv.lineEnd=li},polygonEnd:function(){Tv.point=oi,Tv.lineStart=ui,Tv.lineEnd=si},result:function(){var t=Mv?[bv/Mv,wv/Mv]:xv?[gv/xv,mv/xv]:yv?[vv/yv,_v/yv]:[NaN,NaN];return vv=_v=yv=gv=mv=xv=bv=wv=Mv=0,t}};di.prototype={_radius:4.5,pointRadius:function(t){return this._radius=t,this},polygonStart:function(){this._line=0},polygonEnd:function(){this._line=NaN},lineStart:function(){this._point=0},lineEnd:function(){0===this._line&&this._context.closePath(),this._point=NaN},point:function(t,n){switch(this._point){case 0:this._context.moveTo(t,n),this._point=1;break;case 1:this._context.lineTo(t,n);break;default:this._context.moveTo(t+this._radius,n),this._context.arc(t,n,this._radius,0,ud)}},result:Qe};var kv,Nv,Sv,Ev,Av,Cv=Mp(),zv={point:Qe,lineStart:function(){zv.point=vi},lineEnd:function(){kv&&_i(Nv,Sv),zv.point=Qe},polygonStart:function(){kv=!0},polygonEnd:function(){kv=null},result:function(){var t=+Cv;return Cv.reset(),t}};yi.prototype={_radius:4.5,_circle:gi(4.5),pointRadius:function(t){return(t=+t)!==this._radius&&(this._radius=t,this._circle=null),this},polygonStart:function(){this._line=0},polygonEnd:function(){this._line=NaN},lineStart:function(){this._point=0},lineEnd:function(){0===this._line&&this._string.push("Z"),this._point=NaN},point:function(t,n){switch(this._point){case 0:this._string.push("M",t,",",n),this._point=1;break;case 1:this._string.push("L",t,",",n);break;default:null==this._circle&&(this._circle=gi(this._radius)),this._string.push("M",t,",",n,this._circle)}},result:function(){if(this._string.length){var t=this._string.join("");return this._string=[],t}return null}};var Pv=function(t,n,e,r){return function(i,o){function u(n,e){var r=i(n,e);t(n=r[0],e=r[1])&&o.point(n,e)}function a(t,n){var e=i(t,n);_.point(e[0],e[1])}function c(){b.point=a,_.lineStart()}function s(){b.point=u,_.lineEnd()}function f(t,n){v.push([t,n]);var e=i(t,n);m.point(e[0],e[1])}function l(){m.lineStart(),v=[]}function h(){f(v[0][0],v[0][1]),m.lineEnd();var t,n,e,r,i=m.clean(),u=g.result(),a=u.length;if(v.pop(),p.push(v),v=null,a)if(1&i){if(e=u[0],(n=e.length-1)>0){for(x||(o.polygonStart(),x=!0),o.lineStart(),t=0;t<n;++t)o.point((r=e[t])[0],r[1]);o.lineEnd()}}else a>1&&2&i&&u.push(u.pop().concat(u.shift())),d.push(u.filter(mi))}var p,d,v,_=n(o),y=i.invert(r[0],r[1]),g=Hd(),m=n(g),x=!1,b={point:u,lineStart:c,lineEnd:s,polygonStart:function(){b.point=f,b.lineStart=l,b.lineEnd=h,d=[],p=[]},polygonEnd:function(){b.point=u,b.lineStart=c,b.lineEnd=s,d=Cs(d);var t=Jd(p,y);d.length?(x||(o.polygonStart(),x=!0),Vd(d,xi,t,e,o)):t&&(x||(o.polygonStart(),x=!0),o.lineStart(),e(null,null,1,o),o.lineEnd()),x&&(o.polygonEnd(),x=!1),d=p=null},sphere:function(){o.polygonStart(),o.lineStart(),e(null,null,1,o),o.lineEnd(),o.polygonEnd()}};return b}},Rv=Pv(function(){return!0},function(t){var n,e=NaN,r=NaN,i=NaN;return{lineStart:function(){t.lineStart(),n=1},point:function(o,u){var a=o>0?rd:-rd,c=sd(o-e);sd(c-rd)<ed?(t.point(e,r=(r+u)/2>0?id:-id),t.point(i,r),t.lineEnd(),t.lineStart(),t.point(a,r),t.point(o,r),n=0):i!==a&&c>=rd&&(sd(e-i)<ed&&(e-=i*ed),sd(o-a)<ed&&(o-=a*ed),r=bi(e,r,o,u),t.point(i,r),t.lineEnd(),t.lineStart(),t.point(a,r),n=0),t.point(e=o,r=u),i=a},lineEnd:function(){t.lineEnd(),e=r=NaN},clean:function(){return 2-n}}},function(t,n,e,r){var i;if(null==t)i=e*id,r.point(-rd,i),r.point(0,i),r.point(rd,i),r.point(rd,0),r.point(rd,-i),r.point(0,-i),r.point(-rd,-i),r.point(-rd,0),r.point(-rd,i);else if(sd(t[0]-n[0])>ed){var o=t[0]<n[0]?rd:-rd;i=e*o/2,r.point(-o,i),r.point(0,i),r.point(o,i)}else r.point(n[0],n[1])},[-rd,-id]),Lv=function(t,n){function e(t,n){return hd(t)*hd(n)>o}function r(t,n,e){var r=[1,0,0],i=sr(ar(t),ar(n)),u=cr(i,i),a=i[0],c=u-a*a;if(!c)return!e&&t;var s=o*u/c,f=-o*a/c,l=sr(r,i),h=lr(r,s);fr(h,lr(i,f));var p=l,d=cr(h,p),v=cr(p,p),_=d*d-v*(cr(h,h)-1);if(!(_<0)){var y=md(_),g=lr(p,(-d-y)/v);if(fr(g,h),g=ur(g),!e)return g;var m,x=t[0],b=n[0],w=t[1],M=n[1];b<x&&(m=x,x=b,b=m);var T=b-x,k=sd(T-rd)<ed,N=k||T<ed;if(!k&&M<w&&(m=w,w=M,M=m),N?k?w+M>0^g[1]<(sd(g[0]-x)<ed?w:M):w<=g[1]&&g[1]<=M:T>rd^(x<=g[0]&&g[0]<=b)){var S=lr(p,(-d+y)/v);return fr(S,h),[g,ur(S)]}}}function i(n,e){var r=u?t:rd-t,i=0;return n<-r?i|=1:n>r&&(i|=2),e<-r?i|=4:e>r&&(i|=8),i}var o=hd(t),u=o>0,a=sd(o)>ed;return Pv(e,function(t){var n,o,c,s,f;return{lineStart:function(){s=c=!1,f=1},point:function(l,h){var p,d=[l,h],v=e(l,h),_=u?v?0:i(l,h):v?i(l+(l<0?rd:-rd),h):0;if(!n&&(s=c=v)&&t.lineStart(),v!==c&&(!(p=r(n,d))||$d(n,p)||$d(d,p))&&(d[0]+=ed,d[1]+=ed,v=e(d[0],d[1])),v!==c)f=0,v?(t.lineStart(),p=r(d,n),t.point(p[0],p[1])):(p=r(n,d),t.point(p[0],p[1]),t.lineEnd()),n=p;else if(a&&n&&u^v){var y;_&o||!(y=r(d,n,!0))||(f=0,u?(t.lineStart(),t.point(y[0][0],y[0][1]),t.point(y[1][0],y[1][1]),t.lineEnd()):(t.point(y[1][0],y[1][1]),t.lineEnd(),t.lineStart(),t.point(y[0][0],y[0][1])))}!v||n&&$d(n,d)||t.point(d[0],d[1]),n=d,c=v,o=_},lineEnd:function(){c&&t.lineEnd(),n=null},clean:function(){return f|(s&&c)<<1}}},function(e,r,i,o){Or(o,t,n,i,e,r)},u?[0,-t]:[-rd,t-rd])};Mi.prototype={constructor:Mi,point:function(t,n){this.stream.point(t,n)},sphere:function(){this.stream.sphere()},lineStart:function(){this.stream.lineStart()},lineEnd:function(){this.stream.lineEnd()},polygonStart:function(){this.stream.polygonStart()},polygonEnd:function(){this.stream.polygonEnd()}};var qv=16,Uv=hd(30*cd),Dv=function(t,n){return+n?Si(t,n):Ni(t)},Ov=wi({point:function(t,n){this.stream.point(t*cd,n*cd)}}),Fv=function(){return Ci(Pi).scale(155.424).center([0,33.6442])},Iv=function(){return Fv().parallels([29.5,45.5]).scale(1070).translate([480,250]).rotate([96,0]).center([-.6,38.7])},Yv=Li(function(t){return md(2/(1+t))});Yv.invert=qi(function(t){return 2*Ge(t/2)});var Bv=Li(function(t){return(t=Ze(t))&&t/yd(t)});Bv.invert=qi(function(t){return t});Ui.invert=function(t,n){return[t,2*fd(dd(n))-id]};Ii.invert=Ii;Bi.invert=qi(fd);Hi.invert=qi(Ge);Xi.invert=qi(function(t){return 2*fd(t)});$i.invert=function(t,n){return[-n,2*fd(dd(t))-id]};uo.prototype=eo.prototype={constructor:uo,count:function(){return this.eachAfter(to)},each:function(t){var n,e,r,i,o=this,u=[o];do{for(n=u.reverse(),u=[];o=n.pop();)if(t(o),e=o.children)for(r=0,i=e.length;r<i;++r)u.push(e[r])}while(u.length);return this},eachAfter:function(t){for(var n,e,r,i=this,o=[i],u=[];i=o.pop();)if(u.push(i),n=i.children)for(e=0,r=n.length;e<r;++e)o.push(n[e]);for(;i=u.pop();)t(i);return this},eachBefore:function(t){for(var n,e,r=this,i=[r];r=i.pop();)if(t(r),n=r.children)for(e=n.length-1;e>=0;--e)i.push(n[e]);return this},sum:function(t){return this.eachAfter(function(n){for(var e=+t(n.data)||0,r=n.children,i=r&&r.length;--i>=0;)e+=r[i].value;n.value=e})},sort:function(t){return this.eachBefore(function(n){n.children&&n.children.sort(t)})},path:function(t){for(var n=this,e=no(n,t),r=[n];n!==e;)n=n.parent,r.push(n);for(var i=r.length;t!==e;)r.splice(i,0,t),t=t.parent;return r},ancestors:function(){for(var t=this,n=[t];t=t.parent;)n.push(t);return n},descendants:function(){var t=[];return this.each(function(n){t.push(n)}),t},leaves:function(){var t=[];return this.eachBefore(function(n){n.children||t.push(n)}),t},links:function(){var t=this,n=[];return t.each(function(e){e!==t&&n.push({source:e.parent,target:e})}),n},copy:function(){return eo(this).eachBefore(io)}};var jv=Array.prototype.slice,Hv=function(t){for(var n,e,r=0,i=(t=ao(jv.call(t))).length,o=[];r<i;)n=t[r],e&&fo(e,n)?++r:(e=ho(o=co(o,n)),r=0);return e},Xv=function(t){return function(){return t}},$v=function(t){t.x0=Math.round(t.x0),t.y0=Math.round(t.y0),t.x1=Math.round(t.x1),t.y1=Math.round(t.y1)},Vv=function(t,n,e,r,i){for(var o,u=t.children,a=-1,c=u.length,s=t.value&&(r-n)/t.value;++a<c;)(o=u[a]).y0=e,o.y1=i,o.x0=n,o.x1=n+=o.value*s},Wv="$",Zv={depth:-1},Gv={};Do.prototype=Object.create(uo.prototype);var Jv=function(t,n,e,r,i){for(var o,u=t.children,a=-1,c=u.length,s=t.value&&(i-e)/t.value;++a<c;)(o=u[a]).x0=n,o.x1=r,o.y0=e,o.y1=e+=o.value*s},Qv=(1+Math.sqrt(5))/2,Kv=function t(n){function e(t,e,r,i,o){Fo(n,t,e,r,i,o)}return e.ratio=function(n){return t((n=+n)>1?n:1)},e}(Qv),t_=function t(n){function e(t,e,r,i,o){if((u=t._squarify)&&u.ratio===n)for(var u,a,c,s,f,l=-1,h=u.length,p=t.value;++l<h;){for(c=(a=u[l]).children,s=a.value=0,f=c.length;s<f;++s)a.value+=c[s].value;a.dice?Vv(a,e,r,i,r+=(o-r)*a.value/p):Jv(a,e,r,e+=(i-e)*a.value/p,o),p-=a.value}else t._squarify=u=Fo(n,t,e,r,i,o),u.ratio=n}return e.ratio=function(n){return t((n=+n)>1?n:1)},e}(Qv),n_=function(t,n,e){return(n[0]-t[0])*(e[1]-t[1])-(n[1]-t[1])*(e[0]-t[0])},e_=[].slice,r_={};Bo.prototype=Wo.prototype={constructor:Bo,defer:function(t){if("function"!=typeof t)throw new Error("invalid callback");if(this._call)throw new Error("defer after await");if(null!=this._error)return this;var n=e_.call(arguments,1);return n.push(t),++this._waiting,this._tasks.push(n),jo(this),this},abort:function(){return null==this._error&&$o(this,new Error("abort")),this},await:function(t){if("function"!=typeof t)throw new Error("invalid callback");if(this._call)throw new Error("multiple await");return this._call=function(n,e){t.apply(null,[n].concat(e))},Vo(this),this},awaitAll:function(t){if("function"!=typeof t)throw new Error("invalid callback");if(this._call)throw new Error("multiple await");return this._call=t,Vo(this),this}};var i_=function(){return Math.random()},o_=function t(n){function e(t,e){return t=null==t?0:+t,e=null==e?1:+e,1===arguments.length?(e=t,t=0):e-=t,function(){return n()*e+t}}return e.source=t,e}(i_),u_=function t(n){function e(t,e){var r,i;return t=null==t?0:+t,e=null==e?1:+e,function(){var o;if(null!=r)o=r,r=null;else do{r=2*n()-1,o=2*n()-1,i=r*r+o*o}while(!i||i>1);return t+e*o*Math.sqrt(-2*Math.log(i)/i)}}return e.source=t,e}(i_),a_=function t(n){function e(){var t=u_.source(n).apply(this,arguments);return function(){return Math.exp(t())}}return e.source=t,e}(i_),c_=function t(n){function e(t){return function(){for(var e=0,r=0;r<t;++r)e+=n();return e}}return e.source=t,e}(i_),s_=function t(n){function e(t){var e=c_.source(n)(t);return function(){return e()/t}}return e.source=t,e}(i_),f_=function t(n){function e(t){return function(){return-Math.log(1-n())/t}}return e.source=t,e}(i_),l_=function(t,n){function e(t){var n,e=s.status;if(!e&&Go(s)||e>=200&&e<300||304===e){if(o)try{n=o.call(r,s)}catch(t){return void a.call("error",r,t)}else n=s;a.call("load",r,n)}else a.call("error",r,t)}var r,i,o,u,a=h("beforesend","progress","load","error"),c=we(),s=new XMLHttpRequest,f=null,l=null,p=0;if("undefined"==typeof XDomainRequest||"withCredentials"in s||!/^(http(s)?:)?\/\//.test(t)||(s=new XDomainRequest),"onload"in s?s.onload=s.onerror=s.ontimeout=e:s.onreadystatechange=function(t){s.readyState>3&&e(t)},s.onprogress=function(t){a.call("progress",r,t)},r={header:function(t,n){return t=(t+"").toLowerCase(),arguments.length<2?c.get(t):(null==n?c.remove(t):c.set(t,n+""),r)},mimeType:function(t){return arguments.length?(i=null==t?null:t+"",r):i},responseType:function(t){return arguments.length?(u=t,r):u},timeout:function(t){return arguments.length?(p=+t,r):p},user:function(t){return arguments.length<1?f:(f=null==t?null:t+"",r)},password:function(t){return arguments.length<1?l:(l=null==t?null:t+"",r)},response:function(t){return o=t,r},get:function(t,n){return r.send("GET",t,n)},post:function(t,n){return r.send("POST",t,n)},send:function(n,e,o){return s.open(n,t,!0,f,l),null==i||c.has("accept")||c.set("accept",i+",*/*"),s.setRequestHeader&&c.each(function(t,n){s.setRequestHeader(n,t)}),null!=i&&s.overrideMimeType&&s.overrideMimeType(i),null!=u&&(s.responseType=u),p>0&&(s.timeout=p),null==o&&"function"==typeof e&&(o=e,e=null),null!=o&&1===o.length&&(o=Zo(o)),null!=o&&r.on("error",o).on("load",function(t){o(null,t)}),a.call("beforesend",r,s),s.send(null==e?null:e),r},abort:function(){return s.abort(),r},on:function(){var t=a.on.apply(a,arguments);return t===a?r:t}},null!=n){if("function"!=typeof n)throw new Error("invalid callback: "+n);return r.get(n)}return r},h_=function(t,n){return function(e,r){var i=l_(e).mimeType(t).response(n);if(null!=r){if("function"!=typeof r)throw new Error("invalid callback: "+r);return i.get(r)}return i}},p_=h_("text/html",function(t){return document.createRange().createContextualFragment(t.responseText)}),d_=h_("application/json",function(t){return JSON.parse(t.responseText)}),v_=h_("text/plain",function(t){return t.responseText}),__=h_("application/xml",function(t){var n=t.responseXML;if(!n)throw new Error("parse error");return n}),y_=function(t,n){return function(e,r,i){arguments.length<3&&(i=r,r=null);var o=l_(e).mimeType(t);return o.row=function(t){return arguments.length?o.response(Jo(n,r=t)):r},o.row(r),i?o.get(i):o}},g_=y_("text/csv",Vh),m_=y_("text/tab-separated-values",Qh),x_=Array.prototype,b_=x_.map,w_=x_.slice,M_={name:"implicit"},T_=function(t){return function(){return t}},k_=function(t){return+t},N_=[0,1],S_=function(n,e,r){var o,u=n[0],a=n[n.length-1],c=i(u,a,null==e?10:e);switch((r=He(null==r?",f":r)).type){case"s":var s=Math.max(Math.abs(u),Math.abs(a));return null!=r.precision||isNaN(o=bp(c,s))||(r.precision=o),t.formatPrefix(r,s);case"":case"e":case"g":case"p":case"r":null!=r.precision||isNaN(o=wp(c,Math.max(Math.abs(u),Math.abs(a))))||(r.precision=o-("e"===r.type));break;case"f":case"%":null!=r.precision||isNaN(o=xp(c))||(r.precision=o-2*("%"===r.type))}return t.format(r)},E_=function(t,n){var e,r=0,i=(t=t.slice()).length-1,o=t[r],u=t[i];return u<o&&(e=r,r=i,i=e,e=o,o=u,u=e),t[r]=n.floor(o),t[i]=n.ceil(u),t},A_=new Date,C_=new Date,z_=Mu(function(){},function(t,n){t.setTime(+t+n)},function(t,n){return n-t});z_.every=function(t){return t=Math.floor(t),isFinite(t)&&t>0?t>1?Mu(function(n){n.setTime(Math.floor(n/t)*t)},function(n,e){n.setTime(+n+e*t)},function(n,e){return(e-n)/t}):z_:null};var P_=z_.range,R_=6e4,L_=6048e5,q_=Mu(function(t){t.setTime(1e3*Math.floor(t/1e3))},function(t,n){t.setTime(+t+1e3*n)},function(t,n){return(n-t)/1e3},function(t){return t.getUTCSeconds()}),U_=q_.range,D_=Mu(function(t){t.setTime(Math.floor(t/R_)*R_)},function(t,n){t.setTime(+t+n*R_)},function(t,n){return(n-t)/R_},function(t){return t.getMinutes()}),O_=D_.range,F_=Mu(function(t){var n=t.getTimezoneOffset()*R_%36e5;n<0&&(n+=36e5),t.setTime(36e5*Math.floor((+t-n)/36e5)+n)},function(t,n){t.setTime(+t+36e5*n)},function(t,n){return(n-t)/36e5},function(t){return t.getHours()}),I_=F_.range,Y_=Mu(function(t){t.setHours(0,0,0,0)},function(t,n){t.setDate(t.getDate()+n)},function(t,n){return(n-t-(n.getTimezoneOffset()-t.getTimezoneOffset())*R_)/864e5},function(t){return t.getDate()-1}),B_=Y_.range,j_=Tu(0),H_=Tu(1),X_=Tu(2),$_=Tu(3),V_=Tu(4),W_=Tu(5),Z_=Tu(6),G_=j_.range,J_=H_.range,Q_=X_.range,K_=$_.range,ty=V_.range,ny=W_.range,ey=Z_.range,ry=Mu(function(t){t.setDate(1),t.setHours(0,0,0,0)},function(t,n){t.setMonth(t.getMonth()+n)},function(t,n){return n.getMonth()-t.getMonth()+12*(n.getFullYear()-t.getFullYear())},function(t){return t.getMonth()}),iy=ry.range,oy=Mu(function(t){t.setMonth(0,1),t.setHours(0,0,0,0)},function(t,n){t.setFullYear(t.getFullYear()+n)},function(t,n){return n.getFullYear()-t.getFullYear()},function(t){return t.getFullYear()});oy.every=function(t){return isFinite(t=Math.floor(t))&&t>0?Mu(function(n){n.setFullYear(Math.floor(n.getFullYear()/t)*t),n.setMonth(0,1),n.setHours(0,0,0,0)},function(n,e){n.setFullYear(n.getFullYear()+e*t)}):null};var uy=oy.range,ay=Mu(function(t){t.setUTCSeconds(0,0)},function(t,n){t.setTime(+t+n*R_)},function(t,n){return(n-t)/R_},function(t){return t.getUTCMinutes()}),cy=ay.range,sy=Mu(function(t){t.setUTCMinutes(0,0,0)},function(t,n){t.setTime(+t+36e5*n)},function(t,n){return(n-t)/36e5},function(t){return t.getUTCHours()}),fy=sy.range,ly=Mu(function(t){t.setUTCHours(0,0,0,0)},function(t,n){t.setUTCDate(t.getUTCDate()+n)},function(t,n){return(n-t)/864e5},function(t){return t.getUTCDate()-1}),hy=ly.range,py=ku(0),dy=ku(1),vy=ku(2),_y=ku(3),yy=ku(4),gy=ku(5),my=ku(6),xy=py.range,by=dy.range,wy=vy.range,My=_y.range,Ty=yy.range,ky=gy.range,Ny=my.range,Sy=Mu(function(t){t.setUTCDate(1),t.setUTCHours(0,0,0,0)},function(t,n){t.setUTCMonth(t.getUTCMonth()+n)},function(t,n){return n.getUTCMonth()-t.getUTCMonth()+12*(n.getUTCFullYear()-t.getUTCFullYear())},function(t){return t.getUTCMonth()}),Ey=Sy.range,Ay=Mu(function(t){t.setUTCMonth(0,1),t.setUTCHours(0,0,0,0)},function(t,n){t.setUTCFullYear(t.getUTCFullYear()+n)},function(t,n){return n.getUTCFullYear()-t.getUTCFullYear()},function(t){return t.getUTCFullYear()});Ay.every=function(t){return isFinite(t=Math.floor(t))&&t>0?Mu(function(n){n.setUTCFullYear(Math.floor(n.getUTCFullYear()/t)*t),n.setUTCMonth(0,1),n.setUTCHours(0,0,0,0)},function(n,e){n.setUTCFullYear(n.getUTCFullYear()+e*t)}):null};var Cy,zy=Ay.range,Py={"-":"",_:" ",0:"0"},Ry=/^\s*\d+/,Ly=/^%/,qy=/[\\\^\$\*\+\?\|\[\]\(\)\.\{\}]/g;Ma({dateTime:"%x, %X",date:"%-m/%-d/%Y",time:"%-I:%M:%S %p",periods:["AM","PM"],days:["Sunday","Monday","Tuesday","Wednesday","Thursday","Friday","Saturday"],shortDays:["Sun","Mon","Tue","Wed","Thu","Fri","Sat"],months:["January","February","March","April","May","June","July","August","September","October","November","December"],shortMonths:["Jan","Feb","Mar","Apr","May","Jun","Jul","Aug","Sep","Oct","Nov","Dec"]});var Uy=Date.prototype.toISOString?function(t){return t.toISOString()}:t.utcFormat("%Y-%m-%dT%H:%M:%S.%LZ"),Dy=+new Date("2000-01-01T00:00:00.000Z")?function(t){var n=new Date(t);return isNaN(n)?null:n}:t.utcParse("%Y-%m-%dT%H:%M:%S.%LZ"),Oy=1e3,Fy=60*Oy,Iy=60*Fy,Yy=24*Iy,By=7*Yy,jy=30*Yy,Hy=365*Yy,Xy=function(t){return t.match(/.{6}/g).map(function(t){return"#"+t})},$y=Xy("1f77b4ff7f0e2ca02cd627289467bd8c564be377c27f7f7fbcbd2217becf"),Vy=Xy("393b795254a36b6ecf9c9ede6379398ca252b5cf6bcedb9c8c6d31bd9e39e7ba52e7cb94843c39ad494ad6616be7969c7b4173a55194ce6dbdde9ed6"),Wy=Xy("3182bd6baed69ecae1c6dbefe6550dfd8d3cfdae6bfdd0a231a35474c476a1d99bc7e9c0756bb19e9ac8bcbddcdadaeb636363969696bdbdbdd9d9d9"),Zy=Xy("1f77b4aec7e8ff7f0effbb782ca02c98df8ad62728ff98969467bdc5b0d58c564bc49c94e377c2f7b6d27f7f7fc7c7c7bcbd22dbdb8d17becf9edae5"),Gy=wl($t(300,.5,0),$t(-240,.5,1)),Jy=wl($t(-100,.75,.35),$t(80,1.5,.8)),Qy=wl($t(260,.75,.35),$t(80,1.5,.8)),Ky=$t(),tg=Sa(Xy("44015444025645045745055946075a46085c460a5d460b5e470d60470e6147106347116447136548146748166848176948186a481a6c481b6d481c6e481d6f481f70482071482173482374482475482576482677482878482979472a7a472c7a472d7b472e7c472f7d46307e46327e46337f463480453581453781453882443983443a83443b84433d84433e85423f854240864241864142874144874045884046883f47883f48893e49893e4a893e4c8a3d4d8a3d4e8a3c4f8a3c508b3b518b3b528b3a538b3a548c39558c39568c38588c38598c375a8c375b8d365c8d365d8d355e8d355f8d34608d34618d33628d33638d32648e32658e31668e31678e31688e30698e306a8e2f6b8e2f6c8e2e6d8e2e6e8e2e6f8e2d708e2d718e2c718e2c728e2c738e2b748e2b758e2a768e2a778e2a788e29798e297a8e297b8e287c8e287d8e277e8e277f8e27808e26818e26828e26828e25838e25848e25858e24868e24878e23888e23898e238a8d228b8d228c8d228d8d218e8d218f8d21908d21918c20928c20928c20938c1f948c1f958b1f968b1f978b1f988b1f998a1f9a8a1e9b8a1e9c891e9d891f9e891f9f881fa0881fa1881fa1871fa28720a38620a48621a58521a68522a78522a88423a98324aa8325ab8225ac8226ad8127ad8128ae8029af7f2ab07f2cb17e2db27d2eb37c2fb47c31b57b32b67a34b67935b77937b87838b9773aba763bbb753dbc743fbc7340bd7242be7144bf7046c06f48c16e4ac16d4cc26c4ec36b50c46a52c56954c56856c66758c7655ac8645cc8635ec96260ca6063cb5f65cb5e67cc5c69cd5b6ccd5a6ece5870cf5773d05675d05477d1537ad1517cd2507fd34e81d34d84d44b86d54989d5488bd6468ed64590d74393d74195d84098d83e9bd93c9dd93ba0da39a2da37a5db36a8db34aadc32addc30b0dd2fb2dd2db5de2bb8de29bade28bddf26c0df25c2df23c5e021c8e020cae11fcde11dd0e11cd2e21bd5e21ad8e219dae319dde318dfe318e2e418e5e419e7e419eae51aece51befe51cf1e51df4e61ef6e620f8e621fbe723fde725")),ng=Sa(Xy("00000401000501010601010802010902020b02020d03030f03031204041405041606051806051a07061c08071e0907200a08220b09240c09260d0a290e0b2b100b2d110c2f120d31130d34140e36150e38160f3b180f3d19103f1a10421c10441d11471e114920114b21114e22115024125325125527125829115a2a115c2c115f2d11612f116331116533106734106936106b38106c390f6e3b0f703d0f713f0f72400f74420f75440f764510774710784910784a10794c117a4e117b4f127b51127c52137c54137d56147d57157e59157e5a167e5c167f5d177f5f187f601880621980641a80651a80671b80681c816a1c816b1d816d1d816e1e81701f81721f817320817521817621817822817922827b23827c23827e24828025828125818326818426818627818827818928818b29818c29818e2a81902a81912b81932b80942c80962c80982d80992d809b2e7f9c2e7f9e2f7fa02f7fa1307ea3307ea5317ea6317da8327daa337dab337cad347cae347bb0357bb2357bb3367ab5367ab73779b83779ba3878bc3978bd3977bf3a77c03a76c23b75c43c75c53c74c73d73c83e73ca3e72cc3f71cd4071cf4070d0416fd2426fd3436ed5446dd6456cd8456cd9466bdb476adc4869de4968df4a68e04c67e24d66e34e65e44f64e55064e75263e85362e95462ea5661eb5760ec5860ed5a5fee5b5eef5d5ef05f5ef1605df2625df2645cf3655cf4675cf4695cf56b5cf66c5cf66e5cf7705cf7725cf8745cf8765cf9785df9795df97b5dfa7d5efa7f5efa815ffb835ffb8560fb8761fc8961fc8a62fc8c63fc8e64fc9065fd9266fd9467fd9668fd9869fd9a6afd9b6bfe9d6cfe9f6dfea16efea36ffea571fea772fea973feaa74feac76feae77feb078feb27afeb47bfeb67cfeb77efeb97ffebb81febd82febf84fec185fec287fec488fec68afec88cfeca8dfecc8ffecd90fecf92fed194fed395fed597fed799fed89afdda9cfddc9efddea0fde0a1fde2a3fde3a5fde5a7fde7a9fde9aafdebacfcecaefceeb0fcf0b2fcf2b4fcf4b6fcf6b8fcf7b9fcf9bbfcfbbdfcfdbf")),eg=Sa(Xy("00000401000501010601010802010a02020c02020e03021004031204031405041706041907051b08051d09061f0a07220b07240c08260d08290e092b10092d110a30120a32140b34150b37160b39180c3c190c3e1b0c411c0c431e0c451f0c48210c4a230c4c240c4f260c51280b53290b552b0b572d0b592f0a5b310a5c320a5e340a5f3609613809623909633b09643d09653e0966400a67420a68440a68450a69470b6a490b6a4a0c6b4c0c6b4d0d6c4f0d6c510e6c520e6d540f6d550f6d57106e59106e5a116e5c126e5d126e5f136e61136e62146e64156e65156e67166e69166e6a176e6c186e6d186e6f196e71196e721a6e741a6e751b6e771c6d781c6d7a1d6d7c1d6d7d1e6d7f1e6c801f6c82206c84206b85216b87216b88226a8a226a8c23698d23698f24699025689225689326679526679727669827669a28659b29649d29649f2a63a02a63a22b62a32c61a52c60a62d60a82e5fa92e5eab2f5ead305dae305cb0315bb1325ab3325ab43359b63458b73557b93556ba3655bc3754bd3853bf3952c03a51c13a50c33b4fc43c4ec63d4dc73e4cc83f4bca404acb4149cc4248ce4347cf4446d04545d24644d34743d44842d54a41d74b3fd84c3ed94d3dda4e3cdb503bdd513ade5238df5337e05536e15635e25734e35933e45a31e55c30e65d2fe75e2ee8602de9612bea632aeb6429eb6628ec6726ed6925ee6a24ef6c23ef6e21f06f20f1711ff1731df2741cf3761bf37819f47918f57b17f57d15f67e14f68013f78212f78410f8850ff8870ef8890cf98b0bf98c0af98e09fa9008fa9207fa9407fb9606fb9706fb9906fb9b06fb9d07fc9f07fca108fca309fca50afca60cfca80dfcaa0ffcac11fcae12fcb014fcb216fcb418fbb61afbb81dfbba1ffbbc21fbbe23fac026fac228fac42afac62df9c72ff9c932f9cb35f8cd37f8cf3af7d13df7d340f6d543f6d746f5d949f5db4cf4dd4ff4df53f4e156f3e35af3e55df2e661f2e865f2ea69f1ec6df1ed71f1ef75f1f179f2f27df2f482f3f586f3f68af4f88ef5f992f6fa96f8fb9af9fc9dfafda1fcffa4")),rg=Sa(Xy("0d088710078813078916078a19068c1b068d1d068e20068f2206902406912605912805922a05932c05942e05952f059631059733059735049837049938049a3a049a3c049b3e049c3f049c41049d43039e44039e46039f48039f4903a04b03a14c02a14e02a25002a25102a35302a35502a45601a45801a45901a55b01a55c01a65e01a66001a66100a76300a76400a76600a76700a86900a86a00a86c00a86e00a86f00a87100a87201a87401a87501a87701a87801a87a02a87b02a87d03a87e03a88004a88104a78305a78405a78606a68707a68808a68a09a58b0aa58d0ba58e0ca48f0da4910ea3920fa39410a29511a19613a19814a099159f9a169f9c179e9d189d9e199da01a9ca11b9ba21d9aa31e9aa51f99a62098a72197a82296aa2395ab2494ac2694ad2793ae2892b02991b12a90b22b8fb32c8eb42e8db52f8cb6308bb7318ab83289ba3388bb3488bc3587bd3786be3885bf3984c03a83c13b82c23c81c33d80c43e7fc5407ec6417dc7427cc8437bc9447aca457acb4679cc4778cc4977cd4a76ce4b75cf4c74d04d73d14e72d24f71d35171d45270d5536fd5546ed6556dd7566cd8576bd9586ada5a6ada5b69db5c68dc5d67dd5e66de5f65de6164df6263e06363e16462e26561e26660e3685fe4695ee56a5de56b5de66c5ce76e5be76f5ae87059e97158e97257ea7457eb7556eb7655ec7754ed7953ed7a52ee7b51ef7c51ef7e50f07f4ff0804ef1814df1834cf2844bf3854bf3874af48849f48948f58b47f58c46f68d45f68f44f79044f79143f79342f89441f89540f9973ff9983ef99a3efa9b3dfa9c3cfa9e3bfb9f3afba139fba238fca338fca537fca636fca835fca934fdab33fdac33fdae32fdaf31fdb130fdb22ffdb42ffdb52efeb72dfeb82cfeba2cfebb2bfebd2afebe2afec029fdc229fdc328fdc527fdc627fdc827fdca26fdcb26fccd25fcce25fcd025fcd225fbd324fbd524fbd724fad824fada24f9dc24f9dd25f8df25f8e125f7e225f7e425f6e626f6e826f5e926f5eb27f4ed27f3ee27f3f027f2f227f1f426f1f525f0f724f0f921")),ig=function(t){return function(){return t}},og=Math.abs,ug=Math.atan2,ag=Math.cos,cg=Math.max,sg=Math.min,fg=Math.sin,lg=Math.sqrt,hg=1e-12,pg=Math.PI,dg=pg/2,vg=2*pg;Oa.prototype={areaStart:function(){this._line=0},areaEnd:function(){this._line=NaN},lineStart:function(){this._point=0},lineEnd:function(){(this._line||0!==this._line&&1===this._point)&&this._context.closePath(),this._line=1-this._line},point:function(t,n){switch(t=+t,n=+n,this._point){case 0:this._point=1,this._line?this._context.lineTo(t,n):this._context.moveTo(t,n);break;case 1:this._point=2;default:this._context.lineTo(t,n)}}};var _g=function(t){return new Oa(t)},yg=function(){function t(t){var a,c,s,f=t.length,l=!1;for(null==i&&(u=o(s=ve())),a=0;a<=f;++a)!(a<f&&r(c=t[a],a,t))===l&&((l=!l)?u.lineStart():u.lineEnd()),l&&u.point(+n(c,a,t),+e(c,a,t));if(s)return u=null,s+""||null}var n=Fa,e=Ia,r=ig(!0),i=null,o=_g,u=null;return t.x=function(e){return arguments.length?(n="function"==typeof e?e:ig(+e),t):n},t.y=function(n){return arguments.length?(e="function"==typeof n?n:ig(+n),t):e},t.defined=function(n){return arguments.length?(r="function"==typeof n?n:ig(!!n),t):r},t.curve=function(n){return arguments.length?(o=n,null!=i&&(u=o(i)),t):o},t.context=function(n){return arguments.length?(null==n?i=u=null:u=o(i=n),t):i},t},gg=function(){function t(t){var n,f,l,h,p,d=t.length,v=!1,_=new Array(d),y=new Array(d);for(null==a&&(s=c(p=ve())),n=0;n<=d;++n){if(!(n<d&&u(h=t[n],n,t))===v)if(v=!v)f=n,s.areaStart(),s.lineStart();else{for(s.lineEnd(),s.lineStart(),l=n-1;l>=f;--l)s.point(_[l],y[l]);s.lineEnd(),s.areaEnd()}v&&(_[n]=+e(h,n,t),y[n]=+i(h,n,t),s.point(r?+r(h,n,t):_[n],o?+o(h,n,t):y[n]))}if(p)return s=null,p+""||null}function n(){return yg().defined(u).curve(c).context(a)}var e=Fa,r=null,i=ig(0),o=Ia,u=ig(!0),a=null,c=_g,s=null;return t.x=function(n){return arguments.length?(e="function"==typeof n?n:ig(+n),r=null,t):e},t.x0=function(n){return arguments.length?(e="function"==typeof n?n:ig(+n),t):e},t.x1=function(n){return arguments.length?(r=null==n?null:"function"==typeof n?n:ig(+n),t):r},t.y=function(n){return arguments.length?(i="function"==typeof n?n:ig(+n),o=null,t):i},t.y0=function(n){return arguments.length?(i="function"==typeof n?n:ig(+n),t):i},t.y1=function(n){return arguments.length?(o=null==n?null:"function"==typeof n?n:ig(+n),t):o},t.lineX0=t.lineY0=function(){return n().x(e).y(i)},t.lineY1=function(){return n().x(e).y(o)},t.lineX1=function(){return n().x(r).y(i)},t.defined=function(n){return arguments.length?(u="function"==typeof n?n:ig(!!n),t):u},t.curve=function(n){return arguments.length?(c=n,null!=a&&(s=c(a)),t):c},t.context=function(n){return arguments.length?(null==n?a=s=null:s=c(a=n),t):a},t},mg=function(t,n){return n<t?-1:n>t?1:n>=t?0:NaN},xg=function(t){return t},bg=Ba(_g);Ya.prototype={areaStart:function(){this._curve.areaStart()},areaEnd:function(){this._curve.areaEnd()},lineStart:function(){this._curve.lineStart()},lineEnd:function(){this._curve.lineEnd()},point:function(t,n){this._curve.point(n*Math.sin(t),n*-Math.cos(t))}};var wg=function(){return ja(yg().curve(bg))},Mg=function(){var t=gg().curve(bg),n=t.curve,e=t.lineX0,r=t.lineX1,i=t.lineY0,o=t.lineY1;return t.angle=t.x,delete t.x,t.startAngle=t.x0,delete t.x0,t.endAngle=t.x1,delete t.x1,t.radius=t.y,delete t.y,t.innerRadius=t.y0,delete t.y0,t.outerRadius=t.y1,delete t.y1,t.lineStartAngle=function(){return ja(e())},delete t.lineX0,t.lineEndAngle=function(){return ja(r())},delete t.lineX1,t.lineInnerRadius=function(){return ja(i())},delete t.lineY0,t.lineOuterRadius=function(){return ja(o())},delete t.lineY1,t.curve=function(t){return arguments.length?n(Ba(t)):n()._curve},t},Tg=function(t,n){return[(n=+n)*Math.cos(t-=Math.PI/2),n*Math.sin(t)]},kg=Array.prototype.slice,Ng={draw:function(t,n){var e=Math.sqrt(n/pg);t.moveTo(e,0),t.arc(0,0,e,0,vg)}},Sg={draw:function(t,n){var e=Math.sqrt(n/5)/2;t.moveTo(-3*e,-e),t.lineTo(-e,-e),t.lineTo(-e,-3*e),t.lineTo(e,-3*e),t.lineTo(e,-e),t.lineTo(3*e,-e),t.lineTo(3*e,e),t.lineTo(e,e),t.lineTo(e,3*e),t.lineTo(-e,3*e),t.lineTo(-e,e),t.lineTo(-3*e,e),t.closePath()}},Eg=Math.sqrt(1/3),Ag=2*Eg,Cg={draw:function(t,n){var e=Math.sqrt(n/Ag),r=e*Eg;t.moveTo(0,-e),t.lineTo(r,0),t.lineTo(0,e),t.lineTo(-r,0),t.closePath()}},zg=Math.sin(pg/10)/Math.sin(7*pg/10),Pg=Math.sin(vg/10)*zg,Rg=-Math.cos(vg/10)*zg,Lg={draw:function(t,n){var e=Math.sqrt(.8908130915292852*n),r=Pg*e,i=Rg*e;t.moveTo(0,-e),t.lineTo(r,i);for(var o=1;o<5;++o){var u=vg*o/5,a=Math.cos(u),c=Math.sin(u);t.lineTo(c*e,-a*e),t.lineTo(a*r-c*i,c*r+a*i)}t.closePath()}},qg={draw:function(t,n){var e=Math.sqrt(n),r=-e/2;t.rect(r,r,e,e)}},Ug=Math.sqrt(3),Dg={draw:function(t,n){var e=-Math.sqrt(n/(3*Ug));t.moveTo(0,2*e),t.lineTo(-Ug*e,-e),t.lineTo(Ug*e,-e),t.closePath()}},Og=-.5,Fg=Math.sqrt(3)/2,Ig=1/Math.sqrt(12),Yg=3*(Ig/2+1),Bg={draw:function(t,n){var e=Math.sqrt(n/Yg),r=e/2,i=e*Ig,o=r,u=e*Ig+e,a=-o,c=u;t.moveTo(r,i),t.lineTo(o,u),t.lineTo(a,c),t.lineTo(Og*r-Fg*i,Fg*r+Og*i),t.lineTo(Og*o-Fg*u,Fg*o+Og*u),t.lineTo(Og*a-Fg*c,Fg*a+Og*c),t.lineTo(Og*r+Fg*i,Og*i-Fg*r),t.lineTo(Og*o+Fg*u,Og*u-Fg*o),t.lineTo(Og*a+Fg*c,Og*c-Fg*a),t.closePath()}},jg=[Ng,Sg,Cg,qg,Lg,Dg,Bg],Hg=function(){};Ja.prototype={areaStart:function(){this._line=0},areaEnd:function(){this._line=NaN},lineStart:function(){this._x0=this._x1=this._y0=this._y1=NaN,this._point=0},lineEnd:function(){switch(this._point){case 3:Ga(this,this._x1,this._y1);case 2:this._context.lineTo(this._x1,this._y1)}(this._line||0!==this._line&&1===this._point)&&this._context.closePath(),this._line=1-this._line},point:function(t,n){switch(t=+t,n=+n,this._point){case 0:this._point=1,this._line?this._context.lineTo(t,n):this._context.moveTo(t,n);break;case 1:this._point=2;break;case 2:this._point=3,this._context.lineTo((5*this._x0+this._x1)/6,(5*this._y0+this._y1)/6);default:Ga(this,t,n)}this._x0=this._x1,this._x1=t,this._y0=this._y1,this._y1=n}};Qa.prototype={areaStart:Hg,areaEnd:Hg,lineStart:function(){this._x0=this._x1=this._x2=this._x3=this._x4=this._y0=this._y1=this._y2=this._y3=this._y4=NaN,this._point=0},lineEnd:function(){switch(this._point){case 1:this._context.moveTo(this._x2,this._y2),this._context.closePath();break;case 2:this._context.moveTo((this._x2+2*this._x3)/3,(this._y2+2*this._y3)/3),this._context.lineTo((this._x3+2*this._x2)/3,(this._y3+2*this._y2)/3),this._context.closePath();break;case 3:this.point(this._x2,this._y2),this.point(this._x3,this._y3),this.point(this._x4,this._y4)}},point:function(t,n){switch(t=+t,n=+n,this._point){case 0:this._point=1,this._x2=t,this._y2=n;break;case 1:this._point=2,this._x3=t,this._y3=n;break;case 2:this._point=3,this._x4=t,this._y4=n,this._context.moveTo((this._x0+4*this._x1+t)/6,(this._y0+4*this._y1+n)/6);break;default:Ga(this,t,n)}this._x0=this._x1,this._x1=t,this._y0=this._y1,this._y1=n}};Ka.prototype={areaStart:function(){this._line=0},areaEnd:function(){this._line=NaN},lineStart:function(){this._x0=this._x1=this._y0=this._y1=NaN,this._point=0},lineEnd:function(){(this._line||0!==this._line&&3===this._point)&&this._context.closePath(),this._line=1-this._line},point:function(t,n){switch(t=+t,n=+n,this._point){case 0:this._point=1;break;case 1:this._point=2;break;case 2:this._point=3;var e=(this._x0+4*this._x1+t)/6,r=(this._y0+4*this._y1+n)/6;this._line?this._context.lineTo(e,r):this._context.moveTo(e,r);break;case 3:this._point=4;default:Ga(this,t,n)}this._x0=this._x1,this._x1=t,this._y0=this._y1,this._y1=n}};tc.prototype={lineStart:function(){this._x=[],this._y=[],this._basis.lineStart()},lineEnd:function(){var t=this._x,n=this._y,e=t.length-1;if(e>0)for(var r,i=t[0],o=n[0],u=t[e]-i,a=n[e]-o,c=-1;++c<=e;)r=c/e,this._basis.point(this._beta*t[c]+(1-this._beta)*(i+r*u),this._beta*n[c]+(1-this._beta)*(o+r*a));this._x=this._y=null,this._basis.lineEnd()},point:function(t,n){this._x.push(+t),this._y.push(+n)}};var Xg=function t(n){function e(t){return 1===n?new Ja(t):new tc(t,n)}return e.beta=function(n){return t(+n)},e}(.85);ec.prototype={areaStart:function(){this._line=0},areaEnd:function(){this._line=NaN},lineStart:function(){this._x0=this._x1=this._x2=this._y0=this._y1=this._y2=NaN,this._point=0},lineEnd:function(){switch(this._point){case 2:this._context.lineTo(this._x2,this._y2);break;case 3:nc(this,this._x1,this._y1)}(this._line||0!==this._line&&1===this._point)&&this._context.closePath(),this._line=1-this._line},point:function(t,n){switch(t=+t,n=+n,this._point){case 0:this._point=1,this._line?this._context.lineTo(t,n):this._context.moveTo(t,n);break;case 1:this._point=2,this._x1=t,this._y1=n;break;case 2:this._point=3;default:nc(this,t,n)}this._x0=this._x1,this._x1=this._x2,this._x2=t,this._y0=this._y1,this._y1=this._y2,this._y2=n}};var $g=function t(n){function e(t){return new ec(t,n)}return e.tension=function(n){return t(+n)},e}(0);rc.prototype={areaStart:Hg,areaEnd:Hg,lineStart:function(){this._x0=this._x1=this._x2=this._x3=this._x4=this._x5=this._y0=this._y1=this._y2=this._y3=this._y4=this._y5=NaN,this._point=0},lineEnd:function(){switch(this._point){case 1:this._context.moveTo(this._x3,this._y3),this._context.closePath();break;case 2:this._context.lineTo(this._x3,this._y3),this._context.closePath();break;case 3:this.point(this._x3,this._y3),this.point(this._x4,this._y4),this.point(this._x5,this._y5)}},point:function(t,n){switch(t=+t,n=+n,this._point){case 0:this._point=1,this._x3=t,this._y3=n;break;case 1:this._point=2,this._context.moveTo(this._x4=t,this._y4=n);break;case 2:this._point=3,this._x5=t,this._y5=n;break;default:nc(this,t,n)}this._x0=this._x1,this._x1=this._x2,this._x2=t,this._y0=this._y1,this._y1=this._y2,this._y2=n}};var Vg=function t(n){function e(t){return new rc(t,n)}return e.tension=function(n){return t(+n)},e}(0);ic.prototype={areaStart:function(){this._line=0},areaEnd:function(){this._line=NaN},lineStart:function(){this._x0=this._x1=this._x2=this._y0=this._y1=this._y2=NaN,this._point=0},lineEnd:function(){(this._line||0!==this._line&&3===this._point)&&this._context.closePath(),this._line=1-this._line},point:function(t,n){switch(t=+t,n=+n,this._point){case 0:this._point=1;break;case 1:this._point=2;break;case 2:this._point=3,this._line?this._context.lineTo(this._x2,this._y2):this._context.moveTo(this._x2,this._y2);break;case 3:this._point=4;default:nc(this,t,n)}this._x0=this._x1,this._x1=this._x2,this._x2=t,this._y0=this._y1,this._y1=this._y2,this._y2=n}};var Wg=function t(n){function e(t){return new ic(t,n)}return e.tension=function(n){return t(+n)},e}(0);uc.prototype={areaStart:function(){this._line=0},areaEnd:function(){this._line=NaN},lineStart:function(){this._x0=this._x1=this._x2=this._y0=this._y1=this._y2=NaN,this._l01_a=this._l12_a=this._l23_a=this._l01_2a=this._l12_2a=this._l23_2a=this._point=0},lineEnd:function(){switch(this._point){case 2:this._context.lineTo(this._x2,this._y2);break;case 3:this.point(this._x2,this._y2)}(this._line||0!==this._line&&1===this._point)&&this._context.closePath(),this._line=1-this._line},point:function(t,n){if(t=+t,n=+n,this._point){var e=this._x2-t,r=this._y2-n;this._l23_a=Math.sqrt(this._l23_2a=Math.pow(e*e+r*r,this._alpha))}switch(this._point){case 0:this._point=1,this._line?this._context.lineTo(t,n):this._context.moveTo(t,n);break;case 1:this._point=2;break;case 2:this._point=3;default:oc(this,t,n)}this._l01_a=this._l12_a,this._l12_a=this._l23_a,this._l01_2a=this._l12_2a,this._l12_2a=this._l23_2a,this._x0=this._x1,this._x1=this._x2,this._x2=t,this._y0=this._y1,this._y1=this._y2,this._y2=n}};var Zg=function t(n){function e(t){return n?new uc(t,n):new ec(t,0)}return e.alpha=function(n){return t(+n)},e}(.5);ac.prototype={areaStart:Hg,areaEnd:Hg,lineStart:function(){this._x0=this._x1=this._x2=this._x3=this._x4=this._x5=this._y0=this._y1=this._y2=this._y3=this._y4=this._y5=NaN,this._l01_a=this._l12_a=this._l23_a=this._l01_2a=this._l12_2a=this._l23_2a=this._point=0},lineEnd:function(){switch(this._point){case 1:this._context.moveTo(this._x3,this._y3),this._context.closePath();break;case 2:this._context.lineTo(this._x3,this._y3),this._context.closePath();break;case 3:this.point(this._x3,this._y3),this.point(this._x4,this._y4),this.point(this._x5,this._y5)}},point:function(t,n){if(t=+t,n=+n,this._point){var e=this._x2-t,r=this._y2-n;this._l23_a=Math.sqrt(this._l23_2a=Math.pow(e*e+r*r,this._alpha))}switch(this._point){case 0:this._point=1,this._x3=t,this._y3=n;break;case 1:this._point=2,this._context.moveTo(this._x4=t,this._y4=n);break;case 2:this._point=3,this._x5=t,this._y5=n;break;default:oc(this,t,n)}this._l01_a=this._l12_a,this._l12_a=this._l23_a,this._l01_2a=this._l12_2a,this._l12_2a=this._l23_2a,this._x0=this._x1,this._x1=this._x2,this._x2=t,this._y0=this._y1,this._y1=this._y2,this._y2=n}};var Gg=function t(n){function e(t){return n?new ac(t,n):new rc(t,0)}return e.alpha=function(n){return t(+n)},e}(.5);cc.prototype={areaStart:function(){this._line=0},areaEnd:function(){this._line=NaN},lineStart:function(){this._x0=this._x1=this._x2=this._y0=this._y1=this._y2=NaN,this._l01_a=this._l12_a=this._l23_a=this._l01_2a=this._l12_2a=this._l23_2a=this._point=0},lineEnd:function(){(this._line||0!==this._line&&3===this._point)&&this._context.closePath(),this._line=1-this._line},point:function(t,n){if(t=+t,n=+n,this._point){var e=this._x2-t,r=this._y2-n;this._l23_a=Math.sqrt(this._l23_2a=Math.pow(e*e+r*r,this._alpha))}switch(this._point){case 0:this._point=1;break;case 1:this._point=2;break;case 2:this._point=3,this._line?this._context.lineTo(this._x2,this._y2):this._context.moveTo(this._x2,this._y2);break;case 3:this._point=4;default:oc(this,t,n)}this._l01_a=this._l12_a,this._l12_a=this._l23_a,this._l01_2a=this._l12_2a,this._l12_2a=this._l23_2a,this._x0=this._x1,this._x1=this._x2,this._x2=t,this._y0=this._y1,this._y1=this._y2,this._y2=n}};var Jg=function t(n){function e(t){return n?new cc(t,n):new ic(t,0)}return e.alpha=function(n){return t(+n)},e}(.5);sc.prototype={areaStart:Hg,areaEnd:Hg,lineStart:function(){this._point=0},lineEnd:function(){this._point&&this._context.closePath()},point:function(t,n){t=+t,n=+n,this._point?this._context.lineTo(t,n):(this._point=1,this._context.moveTo(t,n))}};dc.prototype={areaStart:function(){this._line=0},areaEnd:function(){this._line=NaN},lineStart:function(){this._x0=this._x1=this._y0=this._y1=this._t0=NaN,this._point=0},lineEnd:function(){switch(this._point){case 2:this._context.lineTo(this._x1,this._y1);break;case 3:pc(this,this._t0,hc(this,this._t0))}(this._line||0!==this._line&&1===this._point)&&this._context.closePath(),this._line=1-this._line},point:function(t,n){var e=NaN;if(t=+t,n=+n,t!==this._x1||n!==this._y1){switch(this._point){case 0:this._point=1,this._line?this._context.lineTo(t,n):this._context.moveTo(t,n);break;case 1:this._point=2;break;case 2:this._point=3,pc(this,hc(this,e=lc(this,t,n)),e);break;default:pc(this,this._t0,e=lc(this,t,n))}this._x0=this._x1,this._x1=t,this._y0=this._y1,this._y1=n,this._t0=e}}},(vc.prototype=Object.create(dc.prototype)).point=function(t,n){dc.prototype.point.call(this,n,t)},_c.prototype={moveTo:function(t,n){this._context.moveTo(n,t)},closePath:function(){this._context.closePath()},lineTo:function(t,n){this._context.lineTo(n,t)},bezierCurveTo:function(t,n,e,r,i,o){this._context.bezierCurveTo(n,t,r,e,o,i)}},yc.prototype={areaStart:function(){this._line=0},areaEnd:function(){this._line=NaN},lineStart:function(){this._x=[],this._y=[]},lineEnd:function(){var t=this._x,n=this._y,e=t.length;if(e)if(this._line?this._context.lineTo(t[0],n[0]):this._context.moveTo(t[0],n[0]),2===e)this._context.lineTo(t[1],n[1]);else for(var r=gc(t),i=gc(n),o=0,u=1;u<e;++o,++u)this._context.bezierCurveTo(r[0][o],i[0][o],r[1][o],i[1][o],t[u],n[u]);(this._line||0!==this._line&&1===e)&&this._context.closePath(),this._line=1-this._line,this._x=this._y=null},point:function(t,n){this._x.push(+t),this._y.push(+n)}};mc.prototype={areaStart:function(){this._line=0},areaEnd:function(){this._line=NaN},lineStart:function(){this._x=this._y=NaN,this._point=0},lineEnd:function(){0<this._t&&this._t<1&&2===this._point&&this._context.lineTo(this._x,this._y),(this._line||0!==this._line&&1===this._point)&&this._context.closePath(),this._line>=0&&(this._t=1-this._t,this._line=1-this._line)},point:function(t,n){switch(t=+t,n=+n,this._point){case 0:this._point=1,this._line?this._context.lineTo(t,n):this._context.moveTo(t,n);break;case 1:this._point=2;default:if(this._t<=0)this._context.lineTo(this._x,n),this._context.lineTo(t,n);else{var e=this._x*(1-this._t)+t*this._t;this._context.lineTo(e,this._y),this._context.lineTo(e,n)}}this._x=t,this._y=n}};var Qg=function(t,n){if((i=t.length)>1)for(var e,r,i,o=1,u=t[n[0]],a=u.length;o<i;++o)for(r=u,u=t[n[o]],e=0;e<a;++e)u[e][1]+=u[e][0]=isNaN(r[e][1])?r[e][0]:r[e][1]},Kg=function(t){for(var n=t.length,e=new Array(n);--n>=0;)e[n]=n;return e},tm=function(t){var n=t.map(bc);return Kg(t).sort(function(t,e){return n[t]-n[e]})},nm=function(t){return function(){return t}};Tc.prototype={constructor:Tc,insert:function(t,n){var e,r,i;if(t){if(n.P=t,n.N=t.N,t.N&&(t.N.P=n),t.N=n,t.R){for(t=t.R;t.L;)t=t.L;t.L=n}else t.R=n;e=t}else this._?(t=Ec(this._),n.P=null,n.N=t,t.P=t.L=n,e=t):(n.P=n.N=null,this._=n,e=null);for(n.L=n.R=null,n.U=e,n.C=!0,t=n;e&&e.C;)e===(r=e.U).L?(i=r.R)&&i.C?(e.C=i.C=!1,r.C=!0,t=r):(t===e.R&&(Nc(this,e),e=(t=e).U),e.C=!1,r.C=!0,Sc(this,r)):(i=r.L)&&i.C?(e.C=i.C=!1,r.C=!0,t=r):(t===e.L&&(Sc(this,e),e=(t=e).U),e.C=!1,r.C=!0,Nc(this,r)),e=t.U;this._.C=!1},remove:function(t){t.N&&(t.N.P=t.P),t.P&&(t.P.N=t.N),t.N=t.P=null;var n,e,r,i=t.U,o=t.L,u=t.R;if(e=o?u?Ec(u):o:u,i?i.L===t?i.L=e:i.R=e:this._=e,o&&u?(r=e.C,e.C=t.C,e.L=o,o.U=e,e!==u?(i=e.U,e.U=t.U,t=e.R,i.L=t,e.R=u,u.U=e):(e.U=i,i=e,t=e.R)):(r=t.C,t=e),t&&(t.U=i),!r)if(t&&t.C)t.C=!1;else{do{if(t===this._)break;if(t===i.L){if((n=i.R).C&&(n.C=!1,i.C=!0,Nc(this,i),n=i.R),n.L&&n.L.C||n.R&&n.R.C){n.R&&n.R.C||(n.L.C=!1,n.C=!0,Sc(this,n),n=i.R),n.C=i.C,i.C=n.R.C=!1,Nc(this,i),t=this._;break}}else if((n=i.L).C&&(n.C=!1,i.C=!0,Sc(this,i),n=i.L),n.L&&n.L.C||n.R&&n.R.C){n.L&&n.L.C||(n.R.C=!1,n.C=!0,Nc(this,n),n=i.L),n.C=i.C,i.C=n.L.C=!1,Sc(this,i),t=this._;break}n.C=!0,t=i,i=i.U}while(!t.C);t&&(t.C=!1)}}};var em,rm,im,om,um,am=[],cm=[],sm=1e-6,fm=1e-12;Kc.prototype={constructor:Kc,polygons:function(){var t=this.edges;return this.cells.map(function(n){var e=n.halfedges.map(function(e){return Dc(n,t[e])});return e.data=n.site.data,e})},triangles:function(){var t=[],n=this.edges;return this.cells.forEach(function(e,r){if(o=(i=e.halfedges).length)for(var i,o,u,a=e.site,c=-1,s=n[i[o-1]],f=s.left===a?s.right:s.left;++c<o;)u=f,f=(s=n[i[c]]).left===a?s.right:s.left,u&&f&&r<u.index&&r<f.index&&Jc(a,u,f)<0&&t.push([a.data,u.data,f.data])}),t},links:function(){return this.edges.filter(function(t){return t.right}).map(function(t){return{source:t.left.data,target:t.right.data}})},find:function(t,n,e){for(var r,i,o=this,u=o._found||0,a=o.cells.length;!(i=o.cells[u]);)if(++u>=a)return null;var c=t-i.site[0],s=n-i.site[1],f=c*c+s*s;do{i=o.cells[r=u],u=null,i.halfedges.forEach(function(e){var r=o.edges[e],a=r.left;if(a!==i.site&&a||(a=r.right)){var c=t-a[0],s=n-a[1],l=c*c+s*s;l<f&&(f=l,u=a.index)}})}while(null!==u);return o._found=r,null==e||f<=e*e?i.site:null}};var lm=function(t){return function(){return t}};ns.prototype={constructor:ns,scale:function(t){return 1===t?this:new ns(this.k*t,this.x,this.y)},translate:function(t,n){return 0===t&0===n?this:new ns(this.k,this.x+this.k*t,this.y+this.k*n)},apply:function(t){return[t[0]*this.k+this.x,t[1]*this.k+this.y]},applyX:function(t){return t*this.k+this.x},applyY:function(t){return t*this.k+this.y},invert:function(t){return[(t[0]-this.x)/this.k,(t[1]-this.y)/this.k]},invertX:function(t){return(t-this.x)/this.k},invertY:function(t){return(t-this.y)/this.k},rescaleX:function(t){return t.copy().domain(t.range().map(this.invertX,this).map(t.invert,t))},rescaleY:function(t){return t.copy().domain(t.range().map(this.invertY,this).map(t.invert,t))},toString:function(){return"translate("+this.x+","+this.y+") scale("+this.k+")"}};var hm=new ns(1,0,0);es.prototype=ns.prototype;var pm=function(){t.event.preventDefault(),t.event.stopImmediatePropagation()};t.version="4.10.0",t.bisect=hs,t.bisectRight=hs,t.bisectLeft=ps,t.ascending=ss,t.bisector=fs,t.cross=function(t,n,r){var i,o,u,a,c=t.length,s=n.length,f=new Array(c*s);for(null==r&&(r=e),i=u=0;i<c;++i)for(a=t[i],o=0;o<s;++o,++u)f[u]=r(a,n[o]);return f},t.descending=function(t,n){return n<t?-1:n>t?1:n>=t?0:NaN},t.deviation=_s,t.extent=ys,t.histogram=function(){function t(t){var o,u,a=t.length,c=new Array(a);for(o=0;o<a;++o)c[o]=n(t[o],o,t);var s=e(c),f=s[0],l=s[1],h=r(c,f,l);Array.isArray(h)||(h=i(f,l,h),h=Ms(Math.ceil(f/h)*h,Math.floor(l/h)*h,h));for(var p=h.length;h[0]<=f;)h.shift(),--p;for(;h[p-1]>l;)h.pop(),--p;var d,v=new Array(p+1);for(o=0;o<=p;++o)(d=v[o]=[]).x0=o>0?h[o-1]:f,d.x1=o<p?h[o]:l;for(o=0;o<a;++o)f<=(u=c[o])&&u<=l&&v[hs(h,u,0,p)].push(t[o]);return v}var n=ws,e=ys,r=Es;return t.value=function(e){return arguments.length?(n="function"==typeof e?e:bs(e),t):n},t.domain=function(n){return arguments.length?(e="function"==typeof n?n:bs([n[0],n[1]]),t):e},t.thresholds=function(n){return arguments.length?(r="function"==typeof n?n:bs(Array.isArray(n)?ms.call(n):n),t):r},t},t.thresholdFreedmanDiaconis=function(t,n,e){return t=xs.call(t,ds).sort(ss),Math.ceil((e-n)/(2*(As(t,.75)-As(t,.25))*Math.pow(t.length,-1/3)))},t.thresholdScott=function(t,n,e){return Math.ceil((e-n)/(3.5*_s(t)*Math.pow(t.length,-1/3)))},t.thresholdSturges=Es,t.max=function(t,n){var e,r,i=t.length,o=-1;if(null==n){for(;++o<i;)if(null!=(e=t[o])&&e>=e)for(r=e;++o<i;)null!=(e=t[o])&&e>r&&(r=e)}else for(;++o<i;)if(null!=(e=n(t[o],o,t))&&e>=e)for(r=e;++o<i;)null!=(e=n(t[o],o,t))&&e>r&&(r=e);return r},t.mean=function(t,n){var e,r=t.length,i=r,o=-1,u=0;if(null==n)for(;++o<r;)isNaN(e=ds(t[o]))?--i:u+=e;else for(;++o<r;)isNaN(e=ds(n(t[o],o,t)))?--i:u+=e;if(i)return u/i},t.median=function(t,n){var e,r=t.length,i=-1,o=[];if(null==n)for(;++i<r;)isNaN(e=ds(t[i]))||o.push(e);else for(;++i<r;)isNaN(e=ds(n(t[i],i,t)))||o.push(e);return As(o.sort(ss),.5)},t.merge=Cs,t.min=zs,t.pairs=function(t,n){null==n&&(n=e);for(var r=0,i=t.length-1,o=t[0],u=new Array(i<0?0:i);r<i;)u[r]=n(o,o=t[++r]);return u},t.permute=function(t,n){for(var e=n.length,r=new Array(e);e--;)r[e]=t[n[e]];return r},t.quantile=As,t.range=Ms,t.scan=function(t,n){if(e=t.length){var e,r,i=0,o=0,u=t[o];for(null==n&&(n=ss);++i<e;)(n(r=t[i],u)<0||0!==n(u,u))&&(u=r,o=i);return 0===n(u,u)?o:void 0}},t.shuffle=function(t,n,e){for(var r,i,o=(null==e?t.length:e)-(n=null==n?0:+n);o;)i=Math.random()*o--|0,r=t[o+n],t[o+n]=t[i+n],t[i+n]=r;return t},t.sum=function(t,n){var e,r=t.length,i=-1,o=0;if(null==n)for(;++i<r;)(e=+t[i])&&(o+=e);else for(;++i<r;)(e=+n(t[i],i,t))&&(o+=e);return o},t.ticks=Ss,t.tickIncrement=r,t.tickStep=i,t.transpose=Ps,t.variance=vs,t.zip=function(){return Ps(arguments)},t.axisTop=function(t){return l(qs,t)},t.axisRight=function(t){return l(Us,t)},t.axisBottom=function(t){return l(Ds,t)},t.axisLeft=function(t){return l(Os,t)},t.brush=function(){return he(Sh)},t.brushX=function(){return he(kh)},t.brushY=function(){return he(Nh)},t.brushSelection=function(t){var n=t.__brush;return n?n.dim.output(n.selection):null},t.chord=function(){function t(t){var o,u,a,c,s,f,l=t.length,h=[],p=Ms(l),d=[],v=[],_=v.groups=new Array(l),y=new Array(l*l);for(o=0,s=-1;++s<l;){for(u=0,f=-1;++f<l;)u+=t[s][f];h.push(u),d.push(Ms(l)),o+=u}for(e&&p.sort(function(t,n){return e(h[t],h[n])}),r&&d.forEach(function(n,e){n.sort(function(n,i){return r(t[e][n],t[e][i])})}),c=(o=Oh(0,Dh-n*l)/o)?n:Dh/l,u=0,s=-1;++s<l;){for(a=u,f=-1;++f<l;){var g=p[s],m=d[g][f],x=t[g][m],b=u,w=u+=x*o;y[m*l+g]={index:g,subindex:m,startAngle:b,endAngle:w,value:x}}_[g]={index:g,startAngle:a,endAngle:u,value:h[g]},u+=c}for(s=-1;++s<l;)for(f=s-1;++f<l;){var M=y[f*l+s],T=y[s*l+f];(M.value||T.value)&&v.push(M.value<T.value?{source:T,target:M}:{source:M,target:T})}return i?v.sort(i):v}var n=0,e=null,r=null,i=null;return t.padAngle=function(e){return arguments.length?(n=Oh(0,e),t):n},t.sortGroups=function(n){return arguments.length?(e=n,t):e},t.sortSubgroups=function(n){return arguments.length?(r=n,t):r},t.sortChords=function(n){return arguments.length?(null==n?i=null:(i=pe(n))._=n,t):i&&i._},t},t.ribbon=function(){function t(){var t,a=Fh.call(arguments),c=n.apply(this,a),s=e.apply(this,a),f=+r.apply(this,(a[0]=c,a)),l=i.apply(this,a)-Uh,h=o.apply(this,a)-Uh,p=f*Rh(l),d=f*Lh(l),v=+r.apply(this,(a[0]=s,a)),_=i.apply(this,a)-Uh,y=o.apply(this,a)-Uh;if(u||(u=t=ve()),u.moveTo(p,d),u.arc(0,0,f,l,h),l===_&&h===y||(u.quadraticCurveTo(0,0,v*Rh(_),v*Lh(_)),u.arc(0,0,v,_,y)),u.quadraticCurveTo(0,0,p,d),u.closePath(),t)return u=null,t+""||null}var n=_e,e=ye,r=ge,i=me,o=xe,u=null;return t.radius=function(n){return arguments.length?(r="function"==typeof n?n:Ih(+n),t):r},t.startAngle=function(n){return arguments.length?(i="function"==typeof n?n:Ih(+n),t):i},t.endAngle=function(n){return arguments.length?(o="function"==typeof n?n:Ih(+n),t):o},t.source=function(e){return arguments.length?(n=e,t):n},t.target=function(n){return arguments.length?(e=n,t):e},t.context=function(n){return arguments.length?(u=null==n?null:n,t):u},t},t.nest=function(){function t(n,i,u,a){if(i>=o.length)return null!=e&&n.sort(e),null!=r?r(n):n;for(var c,s,f,l=-1,h=n.length,p=o[i++],d=we(),v=u();++l<h;)(f=d.get(c=p(s=n[l])+""))?f.push(s):d.set(c,[s]);return d.each(function(n,e){a(v,e,t(n,i,u,a))}),v}function n(t,e){if(++e>o.length)return t;var i,a=u[e-1];return null!=r&&e>=o.length?i=t.entries():(i=[],t.each(function(t,r){i.push({key:r,values:n(t,e)})})),null!=a?i.sort(function(t,n){return a(t.key,n.key)}):i}var e,r,i,o=[],u=[];return i={object:function(n){return t(n,0,Me,Te)},map:function(n){return t(n,0,ke,Ne)},entries:function(e){return n(t(e,0,ke,Ne),0)},key:function(t){return o.push(t),i},sortKeys:function(t){return u[o.length-1]=t,i},sortValues:function(t){return e=t,i},rollup:function(t){return r=t,i}}},t.set=Ee,t.map=we,t.keys=function(t){var n=[];for(var e in t)n.push(e);return n},t.values=function(t){var n=[];for(var e in t)n.push(t[e]);return n},t.entries=function(t){var n=[];for(var e in t)n.push({key:e,value:t[e]});return n},t.color=Tt,t.rgb=Et,t.hsl=Pt,t.lab=Ut,t.hcl=jt,t.cubehelix=$t,t.dispatch=h,t.drag=function(){function n(t){t.on("mousedown.drag",e).filter(bt).on("touchstart.drag",o).on("touchmove.drag",u).on("touchend.drag touchcancel.drag",a).style("touch-action","none").style("-webkit-tap-highlight-color","rgba(0,0,0,0)")}function e(){if(!p&&d.apply(this,arguments)){var n=c("mouse",v.apply(this,arguments),Ks,this,arguments);n&&(cf(t.event.view).on("mousemove.drag",r,!0).on("mouseup.drag",i,!0),lf(t.event.view),vt(),l=!1,s=t.event.clientX,f=t.event.clientY,n("start"))}}function r(){if(ff(),!l){var n=t.event.clientX-s,e=t.event.clientY-f;l=n*n+e*e>x}y.mouse("drag")}function i(){cf(t.event.view).on("mousemove.drag mouseup.drag",null),_t(t.event.view,l),ff(),y.mouse("end")}function o(){if(d.apply(this,arguments)){var n,e,r=t.event.changedTouches,i=v.apply(this,arguments),o=r.length;for(n=0;n<o;++n)(e=c(r[n].identifier,i,sf,this,arguments))&&(vt(),e("start"))}}function u(){var n,e,r=t.event.changedTouches,i=r.length;for(n=0;n<i;++n)(e=y[r[n].identifier])&&(ff(),e("drag"))}function a(){var n,e,r=t.event.changedTouches,i=r.length;for(p&&clearTimeout(p),p=setTimeout(function(){p=null},500),n=0;n<i;++n)(e=y[r[n].identifier])&&(vt(),e("end"))}function c(e,r,i,o,u){var a,c,s,f=i(r,e),l=g.copy();if(N(new yt(n,"beforestart",a,e,m,f[0],f[1],0,0,l),function(){return null!=(t.event.subject=a=_.apply(o,u))&&(c=a.x-f[0]||0,s=a.y-f[1]||0,!0)}))return function t(h){var p,d=f;switch(h){case"start":y[e]=t,p=m++;break;case"end":delete y[e],--m;case"drag":f=i(r,e),p=m}N(new yt(n,h,a,e,p,f[0]+c,f[1]+s,f[0]-d[0],f[1]-d[1],l),l.apply,l,[h,o,u])}}var s,f,l,p,d=gt,v=mt,_=xt,y={},g=h("start","drag","end"),m=0,x=0;return n.filter=function(t){return arguments.length?(d="function"==typeof t?t:hf(!!t),n):d},n.container=function(t){return arguments.length?(v="function"==typeof t?t:hf(t),n):v},n.subject=function(t){return arguments.length?(_="function"==typeof t?t:hf(t),n):_},n.on=function(){var t=g.on.apply(g,arguments);return t===g?n:t},n.clickDistance=function(t){return arguments.length?(x=(t=+t)*t,n):Math.sqrt(x)},n},t.dragDisable=lf,t.dragEnable=_t,t.dsvFormat=Xh,t.csvParse=Vh,t.csvParseRows=Wh,t.csvFormat=Zh,t.csvFormatRows=Gh,t.tsvParse=Qh,t.tsvParseRows=Kh,t.tsvFormat=tp,t.tsvFormatRows=np,t.easeLinear=function(t){return+t},t.easeQuad=Kn,t.easeQuadIn=function(t){return t*t},t.easeQuadOut=function(t){return t*(2-t)},t.easeQuadInOut=Kn,t.easeCubic=te,t.easeCubicIn=function(t){return t*t*t},t.easeCubicOut=function(t){return--t*t*t+1},t.easeCubicInOut=te,t.easePoly=Gl,t.easePolyIn=Wl,t.easePolyOut=Zl,t.easePolyInOut=Gl,t.easeSin=ne,t.easeSinIn=function(t){return 1-Math.cos(t*Ql)},t.easeSinOut=function(t){return Math.sin(t*Ql)},t.easeSinInOut=ne,t.easeExp=ee,t.easeExpIn=function(t){return Math.pow(2,10*t-10)},t.easeExpOut=function(t){return 1-Math.pow(2,-10*t)},t.easeExpInOut=ee,t.easeCircle=re,t.easeCircleIn=function(t){return 1-Math.sqrt(1-t*t)},t.easeCircleOut=function(t){return Math.sqrt(1- --t*t)},t.easeCircleInOut=re,t.easeBounce=ie,t.easeBounceIn=function(t){return 1-ie(1-t)},t.easeBounceOut=ie,t.easeBounceInOut=function(t){return((t*=2)<=1?1-ie(1-t):ie(t-1)+1)/2},t.easeBack=lh,t.easeBackIn=sh,t.easeBackOut=fh,t.easeBackInOut=lh,t.easeElastic=dh,t.easeElasticIn=ph,t.easeElasticOut=dh,t.easeElasticInOut=vh,t.forceCenter=function(t,n){function e(){var e,i,o=r.length,u=0,a=0;for(e=0;e<o;++e)u+=(i=r[e]).x,a+=i.y;for(u=u/o-t,a=a/o-n,e=0;e<o;++e)(i=r[e]).x-=u,i.y-=a}var r;return null==t&&(t=0),null==n&&(n=0),e.initialize=function(t){r=t},e.x=function(n){return arguments.length?(t=+n,e):t},e.y=function(t){return arguments.length?(n=+t,e):n},e},t.forceCollide=function(t){function n(){for(var t,n,r,c,s,f,l,h=i.length,p=0;p<a;++p)for(n=qe(i,Oe,Fe).visitAfter(e),t=0;t<h;++t)r=i[t],f=o[r.index],l=f*f,c=r.x+r.vx,s=r.y+r.vy,n.visit(function(t,n,e,i,o){var a=t.data,h=t.r,p=f+h;if(!a)return n>c+p||i<c-p||e>s+p||o<s-p;if(a.index>r.index){var d=c-a.x-a.vx,v=s-a.y-a.vy,_=d*d+v*v;_<p*p&&(0===d&&(d=rp(),_+=d*d),0===v&&(v=rp(),_+=v*v),_=(p-(_=Math.sqrt(_)))/_*u,r.vx+=(d*=_)*(p=(h*=h)/(l+h)),r.vy+=(v*=_)*p,a.vx-=d*(p=1-p),a.vy-=v*p)}})}function e(t){if(t.data)return t.r=o[t.data.index];for(var n=t.r=0;n<4;++n)t[n]&&t[n].r>t.r&&(t.r=t[n].r)}function r(){if(i){var n,e,r=i.length;for(o=new Array(r),n=0;n<r;++n)e=i[n],o[e.index]=+t(e,n,i)}}var i,o,u=1,a=1;return"function"!=typeof t&&(t=ep(null==t?1:+t)),n.initialize=function(t){i=t,r()},n.iterations=function(t){return arguments.length?(a=+t,n):a},n.strength=function(t){return arguments.length?(u=+t,n):u},n.radius=function(e){return arguments.length?(t="function"==typeof e?e:ep(+e),r(),n):t},n},t.forceLink=function(t){function n(n){for(var e=0,r=t.length;e<p;++e)for(var i,a,c,f,l,h,d,v=0;v<r;++v)a=(i=t[v]).source,f=(c=i.target).x+c.vx-a.x-a.vx||rp(),l=c.y+c.vy-a.y-a.vy||rp(),f*=h=((h=Math.sqrt(f*f+l*l))-u[v])/h*n*o[v],l*=h,c.vx-=f*(d=s[v]),c.vy-=l*d,a.vx+=f*(d=1-d),a.vy+=l*d}function e(){if(a){var n,e,l=a.length,h=t.length,p=we(a,f);for(n=0,c=new Array(l);n<h;++n)(e=t[n]).index=n,"object"!=typeof e.source&&(e.source=Ye(p,e.source)),"object"!=typeof e.target&&(e.target=Ye(p,e.target)),c[e.source.index]=(c[e.source.index]||0)+1,c[e.target.index]=(c[e.target.index]||0)+1;for(n=0,s=new Array(h);n<h;++n)e=t[n],s[n]=c[e.source.index]/(c[e.source.index]+c[e.target.index]);o=new Array(h),r(),u=new Array(h),i()}}function r(){if(a)for(var n=0,e=t.length;n<e;++n)o[n]=+l(t[n],n,t)}function i(){if(a)for(var n=0,e=t.length;n<e;++n)u[n]=+h(t[n],n,t)}var o,u,a,c,s,f=Ie,l=function(t){return 1/Math.min(c[t.source.index],c[t.target.index])},h=ep(30),p=1;return null==t&&(t=[]),n.initialize=function(t){a=t,e()},n.links=function(r){return arguments.length?(t=r,e(),n):t},n.id=function(t){return arguments.length?(f=t,n):f},n.iterations=function(t){return arguments.length?(p=+t,n):p},n.strength=function(t){return arguments.length?(l="function"==typeof t?t:ep(+t),r(),n):l},n.distance=function(t){return arguments.length?(h="function"==typeof t?t:ep(+t),i(),n):h},n},t.forceManyBody=function(){function t(t){var n,a=i.length,c=qe(i,Be,je).visitAfter(e);for(u=t,n=0;n<a;++n)o=i[n],c.visit(r)}function n(){if(i){var t,n,e=i.length;for(a=new Array(e),t=0;t<e;++t)n=i[t],a[n.index]=+c(n,t,i)}}function e(t){var n,e,r,i,o,u=0;if(t.length){for(r=i=o=0;o<4;++o)(n=t[o])&&(e=n.value)&&(u+=e,r+=e*n.x,i+=e*n.y);t.x=r/u,t.y=i/u}else{(n=t).x=n.data.x,n.y=n.data.y;do{u+=a[n.data.index]}while(n=n.next)}t.value=u}function r(t,n,e,r){if(!t.value)return!0;var i=t.x-o.x,c=t.y-o.y,h=r-n,p=i*i+c*c;if(h*h/l<p)return p<f&&(0===i&&(i=rp(),p+=i*i),0===c&&(c=rp(),p+=c*c),p<s&&(p=Math.sqrt(s*p)),o.vx+=i*t.value*u/p,o.vy+=c*t.value*u/p),!0;if(!(t.length||p>=f)){(t.data!==o||t.next)&&(0===i&&(i=rp(),p+=i*i),0===c&&(c=rp(),p+=c*c),p<s&&(p=Math.sqrt(s*p)));do{t.data!==o&&(h=a[t.data.index]*u/p,o.vx+=i*h,o.vy+=c*h)}while(t=t.next)}}var i,o,u,a,c=ep(-30),s=1,f=1/0,l=.81;return t.initialize=function(t){i=t,n()},t.strength=function(e){return arguments.length?(c="function"==typeof e?e:ep(+e),n(),t):c},t.distanceMin=function(n){return arguments.length?(s=n*n,t):Math.sqrt(s)},t.distanceMax=function(n){return arguments.length?(f=n*n,t):Math.sqrt(f)},t.theta=function(n){return arguments.length?(l=n*n,t):Math.sqrt(l)},t},t.forceSimulation=function(t){function n(){e(),d.call("tick",o),u<a&&(p.stop(),d.call("end",o))}function e(){var n,e,r=t.length;for(u+=(s-u)*c,l.each(function(t){t(u)}),n=0;n<r;++n)null==(e=t[n]).fx?e.x+=e.vx*=f:(e.x=e.fx,e.vx=0),null==e.fy?e.y+=e.vy*=f:(e.y=e.fy,e.vy=0)}function r(){for(var n,e=0,r=t.length;e<r;++e){if(n=t[e],n.index=e,isNaN(n.x)||isNaN(n.y)){var i=ap*Math.sqrt(e),o=e*cp;n.x=i*Math.cos(o),n.y=i*Math.sin(o)}(isNaN(n.vx)||isNaN(n.vy))&&(n.vx=n.vy=0)}}function i(n){return n.initialize&&n.initialize(t),n}var o,u=1,a=.001,c=1-Math.pow(a,1/300),s=0,f=.6,l=we(),p=dn(n),d=h("tick","end");return null==t&&(t=[]),r(),o={tick:e,restart:function(){return p.restart(n),o},stop:function(){return p.stop(),o},nodes:function(n){return arguments.length?(t=n,r(),l.each(i),o):t},alpha:function(t){return arguments.length?(u=+t,o):u},alphaMin:function(t){return arguments.length?(a=+t,o):a},alphaDecay:function(t){return arguments.length?(c=+t,o):+c},alphaTarget:function(t){return arguments.length?(s=+t,o):s},velocityDecay:function(t){return arguments.length?(f=1-t,o):1-f},force:function(t,n){return arguments.length>1?(null==n?l.remove(t):l.set(t,i(n)),o):l.get(t)},find:function(n,e,r){var i,o,u,a,c,s=0,f=t.length;for(null==r?r=1/0:r*=r,s=0;s<f;++s)(u=(i=n-(a=t[s]).x)*i+(o=e-a.y)*o)<r&&(c=a,r=u);return c},on:function(t,n){return arguments.length>1?(d.on(t,n),o):d.on(t)}}},t.forceX=function(t){function n(t){for(var n,e=0,u=r.length;e<u;++e)(n=r[e]).vx+=(o[e]-n.x)*i[e]*t}function e(){if(r){var n,e=r.length;for(i=new Array(e),o=new Array(e),n=0;n<e;++n)i[n]=isNaN(o[n]=+t(r[n],n,r))?0:+u(r[n],n,r)}}var r,i,o,u=ep(.1);return"function"!=typeof t&&(t=ep(null==t?0:+t)),n.initialize=function(t){r=t,e()},n.strength=function(t){return arguments.length?(u="function"==typeof t?t:ep(+t),e(),n):u},n.x=function(r){return arguments.length?(t="function"==typeof r?r:ep(+r),e(),n):t},n},t.forceY=function(t){function n(t){for(var n,e=0,u=r.length;e<u;++e)(n=r[e]).vy+=(o[e]-n.y)*i[e]*t}function e(){if(r){var n,e=r.length;for(i=new Array(e),o=new Array(e),n=0;n<e;++n)i[n]=isNaN(o[n]=+t(r[n],n,r))?0:+u(r[n],n,r)}}var r,i,o,u=ep(.1);return"function"!=typeof t&&(t=ep(null==t?0:+t)),n.initialize=function(t){r=t,e()},n.strength=function(t){return arguments.length?(u="function"==typeof t?t:ep(+t),e(),n):u},n.y=function(r){return arguments.length?(t="function"==typeof r?r:ep(+r),e(),n):t},n},t.formatDefaultLocale=$e,t.formatLocale=mp,t.formatSpecifier=He,t.precisionFixed=xp,t.precisionPrefix=bp,t.precisionRound=wp,t.geoArea=function(t){return kd.reset(),Md(t,Nd),2*kd},t.geoBounds=function(t){var n,e,r,i,o,u,a;if(Pp=zp=-(Ap=Cp=1/0),Dp=[],Md(t,Ed),e=Dp.length){for(Dp.sort(br),n=1,o=[r=Dp[0]];n<e;++n)wr(r,(i=Dp[n])[0])||wr(r,i[1])?(xr(r[0],i[1])>xr(r[0],r[1])&&(r[1]=i[1]),xr(i[0],r[1])>xr(r[0],r[1])&&(r[0]=i[0])):o.push(r=i);for(u=-1/0,n=0,r=o[e=o.length-1];n<=e;r=i,++n)i=o[n],(a=xr(r[1],i[0]))>u&&(u=a,Ap=i[0],zp=r[1])}return Dp=Op=null,Ap===1/0||Cp===1/0?[[NaN,NaN],[NaN,NaN]]:[[Ap,Cp],[zp,Pp]]},t.geoCentroid=function(t){Fp=Ip=Yp=Bp=jp=Hp=Xp=$p=Vp=Wp=Zp=0,Md(t,Ad);var n=Vp,e=Wp,r=Zp,i=n*n+e*e+r*r;return i<1e-12&&(n=Hp,e=Xp,r=$p,Ip<ed&&(n=Yp,e=Bp,r=jp),(i=n*n+e*e+r*r)<1e-12)?[NaN,NaN]:[ld(e,n)*ad,Ge(r/md(i))*ad]},t.geoCircle=function(){function t(){var t=r.apply(this,arguments),a=i.apply(this,arguments)*cd,c=o.apply(this,arguments)*cd;return n=[],e=Lr(-t[0]*cd,-t[1]*cd,0).invert,Or(u,a,c,1),t={type:"Polygon",coordinates:[n]},n=e=null,t}var n,e,r=Cd([0,0]),i=Cd(90),o=Cd(6),u={point:function(t,r){n.push(t=e(t,r)),t[0]*=ad,t[1]*=ad}};return t.center=function(n){return arguments.length?(r="function"==typeof n?n:Cd([+n[0],+n[1]]),t):r},t.radius=function(n){return arguments.length?(i="function"==typeof n?n:Cd(+n),t):i},t.precision=function(n){return arguments.length?(o="function"==typeof n?n:Cd(+n),t):o},t},t.geoClipExtent=function(){var t,n,e,r=0,i=0,o=960,u=500;return e={stream:function(e){return t&&n===e?t:t=Br(r,i,o,u)(n=e)},extent:function(a){return arguments.length?(r=+a[0][0],i=+a[0][1],o=+a[1][0],u=+a[1][1],t=n=null,e):[[r,i],[o,u]]}}},t.geoContains=function(t,n){return(t&&iv.hasOwnProperty(t.type)?iv[t.type]:$r)(t,n)},t.geoDistance=rv,t.geoGraticule=ti,t.geoGraticule10=function(){return ti()()},t.geoInterpolate=function(t,n){var e=t[0]*cd,r=t[1]*cd,i=n[0]*cd,o=n[1]*cd,u=hd(r),a=yd(r),c=hd(o),s=yd(o),f=u*hd(e),l=u*yd(e),h=c*hd(i),p=c*yd(i),d=2*Ge(md(Je(o-r)+u*c*Je(i-e))),v=yd(d),_=d?function(t){var n=yd(t*=d)/v,e=yd(d-t)/v,r=e*f+n*h,i=e*l+n*p,o=e*a+n*s;return[ld(i,r)*ad,ld(o,md(r*r+i*i))*ad]}:function(){return[e*ad,r*ad]};return _.distance=d,_},t.geoLength=tv,t.geoPath=function(t,n){function e(t){return t&&("function"==typeof o&&i.pointRadius(+o.apply(this,arguments)),Md(t,r(i))),i.result()}var r,i,o=4.5;return e.area=function(t){return Md(t,r(sv)),sv.result()},e.measure=function(t){return Md(t,r(zv)),zv.result()},e.bounds=function(t){return Md(t,r(dv)),dv.result()},e.centroid=function(t){return Md(t,r(Tv)),Tv.result()},e.projection=function(n){return arguments.length?(r=null==n?(t=null,uv):(t=n).stream,e):t},e.context=function(t){return arguments.length?(i=null==t?(n=null,new yi):new di(n=t),"function"!=typeof o&&i.pointRadius(o),e):n},e.pointRadius=function(t){return arguments.length?(o="function"==typeof t?t:(i.pointRadius(+t),+t),e):o},e.projection(t).context(n)},t.geoAlbers=Iv,t.geoAlbersUsa=function(){function t(t){var n=t[0],e=t[1];return a=null,i.point(n,e),a||(o.point(n,e),a)||(u.point(n,e),a)}function n(){return e=r=null,t}var e,r,i,o,u,a,c=Iv(),s=Fv().rotate([154,0]).center([-2,58.5]).parallels([55,65]),f=Fv().rotate([157,0]).center([-3,19.9]).parallels([8,18]),l={point:function(t,n){a=[t,n]}};return t.invert=function(t){var n=c.scale(),e=c.translate(),r=(t[0]-e[0])/n,i=(t[1]-e[1])/n;return(i>=.12&&i<.234&&r>=-.425&&r<-.214?s:i>=.166&&i<.234&&r>=-.214&&r<-.115?f:c).invert(t)},t.stream=function(t){return e&&r===t?e:e=Ri([c.stream(r=t),s.stream(t),f.stream(t)])},t.precision=function(t){return arguments.length?(c.precision(t),s.precision(t),f.precision(t),n()):c.precision()},t.scale=function(n){return arguments.length?(c.scale(n),s.scale(.35*n),f.scale(n),t.translate(c.translate())):c.scale()},t.translate=function(t){if(!arguments.length)return c.translate();var e=c.scale(),r=+t[0],a=+t[1];return i=c.translate(t).clipExtent([[r-.455*e,a-.238*e],[r+.455*e,a+.238*e]]).stream(l),o=s.translate([r-.307*e,a+.201*e]).clipExtent([[r-.425*e+ed,a+.12*e+ed],[r-.214*e-ed,a+.234*e-ed]]).stream(l),u=f.translate([r-.205*e,a+.212*e]).clipExtent([[r-.214*e+ed,a+.166*e+ed],[r-.115*e-ed,a+.234*e-ed]]).stream(l),n()},t.fitExtent=function(n,e){return Ti(t,n,e)},t.fitSize=function(n,e){return ki(t,n,e)},t.scale(1070)},t.geoAzimuthalEqualArea=function(){return Ei(Yv).scale(124.75).clipAngle(179.999)},t.geoAzimuthalEqualAreaRaw=Yv,t.geoAzimuthalEquidistant=function(){return Ei(Bv).scale(79.4188).clipAngle(179.999)},t.geoAzimuthalEquidistantRaw=Bv,t.geoConicConformal=function(){return Ci(Fi).scale(109.5).parallels([30,30])},t.geoConicConformalRaw=Fi,t.geoConicEqualArea=Fv,t.geoConicEqualAreaRaw=Pi,t.geoConicEquidistant=function(){return Ci(Yi).scale(131.154).center([0,13.9389])},t.geoConicEquidistantRaw=Yi,t.geoEquirectangular=function(){return Ei(Ii).scale(152.63)},t.geoEquirectangularRaw=Ii,t.geoGnomonic=function(){return Ei(Bi).scale(144.049).clipAngle(60)},t.geoGnomonicRaw=Bi,t.geoIdentity=function(){function t(){return i=o=null,u}var n,e,r,i,o,u,a=1,c=0,s=0,f=1,l=1,h=uv,p=null,d=uv;return u={stream:function(t){return i&&o===t?i:i=h(d(o=t))},clipExtent:function(i){return arguments.length?(d=null==i?(p=n=e=r=null,uv):Br(p=+i[0][0],n=+i[0][1],e=+i[1][0],r=+i[1][1]),t()):null==p?null:[[p,n],[e,r]]},scale:function(n){return arguments.length?(h=ji((a=+n)*f,a*l,c,s),t()):a},translate:function(n){return arguments.length?(h=ji(a*f,a*l,c=+n[0],s=+n[1]),t()):[c,s]},reflectX:function(n){return arguments.length?(h=ji(a*(f=n?-1:1),a*l,c,s),t()):f<0},reflectY:function(n){return arguments.length?(h=ji(a*f,a*(l=n?-1:1),c,s),t()):l<0},fitExtent:function(t,n){return Ti(u,t,n)},fitSize:function(t,n){return ki(u,t,n)}}},t.geoProjection=Ei,t.geoProjectionMutator=Ai,t.geoMercator=function(){return Di(Ui).scale(961/ud)},t.geoMercatorRaw=Ui,t.geoOrthographic=function(){return Ei(Hi).scale(249.5).clipAngle(90+ed)},t.geoOrthographicRaw=Hi,t.geoStereographic=function(){return Ei(Xi).scale(250).clipAngle(142)},t.geoStereographicRaw=Xi,t.geoTransverseMercator=function(){var t=Di($i),n=t.center,e=t.rotate;return t.center=function(t){return arguments.length?n([-t[1],t[0]]):(t=n(),[t[1],-t[0]])},t.rotate=function(t){return arguments.length?e([t[0],t[1],t.length>2?t[2]+90:90]):(t=e(),[t[0],t[1],t[2]-90])},e([0,0,90]).scale(159.155)},t.geoTransverseMercatorRaw=$i,t.geoRotation=jd,t.geoStream=Md,t.geoTransform=function(t){return{stream:wi(t)}},t.cluster=function(){function t(t){var o,u=0;t.eachAfter(function(t){var e=t.children;e?(t.x=Wi(e),t.y=Gi(e)):(t.x=o?u+=n(t,o):0,t.y=0,o=t)});var a=Qi(t),c=Ki(t),s=a.x-n(a,c)/2,f=c.x+n(c,a)/2;return t.eachAfter(i?function(n){n.x=(n.x-t.x)*e,n.y=(t.y-n.y)*r}:function(n){n.x=(n.x-s)/(f-s)*e,n.y=(1-(t.y?n.y/t.y:1))*r})}var n=Vi,e=1,r=1,i=!1;return t.separation=function(e){return arguments.length?(n=e,t):n},t.size=function(n){return arguments.length?(i=!1,e=+n[0],r=+n[1],t):i?null:[e,r]},t.nodeSize=function(n){return arguments.length?(i=!0,e=+n[0],r=+n[1],t):i?[e,r]:null},t},t.hierarchy=eo,t.pack=function(){function t(t){return t.x=e/2,t.y=r/2,n?t.eachBefore(No(n)).eachAfter(So(i,.5)).eachBefore(Eo(1)):t.eachBefore(No(ko)).eachAfter(So(To,1)).eachAfter(So(i,t.r/Math.min(e,r))).eachBefore(Eo(Math.min(e,r)/(2*t.r))),t}var n=null,e=1,r=1,i=To;return t.radius=function(e){return arguments.length?(n=wo(e),t):n},t.size=function(n){return arguments.length?(e=+n[0],r=+n[1],t):[e,r]},t.padding=function(n){return arguments.length?(i="function"==typeof n?n:Xv(+n),t):i},t},t.packSiblings=function(t){return bo(t),t},t.packEnclose=Hv,t.partition=function(){function t(t){var u=t.height+1;return t.x0=t.y0=i,t.x1=e,t.y1=r/u,t.eachBefore(n(r,u)),o&&t.eachBefore($v),t}function n(t,n){return function(e){e.children&&Vv(e,e.x0,t*(e.depth+1)/n,e.x1,t*(e.depth+2)/n);var r=e.x0,o=e.y0,u=e.x1-i,a=e.y1-i;u<r&&(r=u=(r+u)/2),a<o&&(o=a=(o+a)/2),e.x0=r,e.y0=o,e.x1=u,e.y1=a}}var e=1,r=1,i=0,o=!1;return t.round=function(n){return arguments.length?(o=!!n,t):o},t.size=function(n){return arguments.length?(e=+n[0],r=+n[1],t):[e,r]},t.padding=function(n){return arguments.length?(i=+n,t):i},t},t.stratify=function(){function t(t){var r,i,o,u,a,c,s,f=t.length,l=new Array(f),h={};for(i=0;i<f;++i)r=t[i],a=l[i]=new uo(r),null!=(c=n(r,i,t))&&(c+="")&&(h[s=Wv+(a.id=c)]=s in h?Gv:a);for(i=0;i<f;++i)if(a=l[i],null!=(c=e(t[i],i,t))&&(c+="")){if(!(u=h[Wv+c]))throw new Error("missing: "+c);if(u===Gv)throw new Error("ambiguous: "+c);u.children?u.children.push(a):u.children=[a],a.parent=u}else{if(o)throw new Error("multiple roots");o=a}if(!o)throw new Error("no root");if(o.parent=Zv,o.eachBefore(function(t){t.depth=t.parent.depth+1,--f}).eachBefore(oo),o.parent=null,f>0)throw new Error("cycle");return o}var n=Ao,e=Co;return t.id=function(e){return arguments.length?(n=Mo(e),t):n},t.parentId=function(n){return arguments.length?(e=Mo(n),t):e},t},t.tree=function(){function t(t){var r=Oo(t);if(r.eachAfter(n),r.parent.m=-r.z,r.eachBefore(e),c)t.eachBefore(i);else{var s=t,f=t,l=t;t.eachBefore(function(t){t.x<s.x&&(s=t),t.x>f.x&&(f=t),t.depth>l.depth&&(l=t)});var h=s===f?1:o(s,f)/2,p=h-s.x,d=u/(f.x+h+p),v=a/(l.depth||1);t.eachBefore(function(t){t.x=(t.x+p)*d,t.y=t.depth*v})}return t}function n(t){var n=t.children,e=t.parent.children,i=t.i?e[t.i-1]:null;if(n){qo(t);var u=(n[0].z+n[n.length-1].z)/2;i?(t.z=i.z+o(t._,i._),t.m=t.z-u):t.z=u}else i&&(t.z=i.z+o(t._,i._));t.parent.A=r(t,i,t.parent.A||e[0])}function e(t){t._.x=t.z+t.parent.m,t.m+=t.parent.m}function r(t,n,e){if(n){for(var r,i=t,u=t,a=n,c=i.parent.children[0],s=i.m,f=u.m,l=a.m,h=c.m;a=Ro(a),i=Po(i),a&&i;)c=Po(c),(u=Ro(u)).a=t,(r=a.z+l-i.z-s+o(a._,i._))>0&&(Lo(Uo(a,t,e),t,r),s+=r,f+=r),l+=a.m,s+=i.m,h+=c.m,f+=u.m;a&&!Ro(u)&&(u.t=a,u.m+=l-f),i&&!Po(c)&&(c.t=i,c.m+=s-h,e=t)}return e}function i(t){t.x*=u,t.y=t.depth*a}var o=zo,u=1,a=1,c=null;return t.separation=function(n){return arguments.length?(o=n,t):o},t.size=function(n){return arguments.length?(c=!1,u=+n[0],a=+n[1],t):c?null:[u,a]},t.nodeSize=function(n){return arguments.length?(c=!0,u=+n[0],a=+n[1],t):c?[u,a]:null},t},t.treemap=function(){function t(t){return t.x0=t.y0=0,t.x1=i,t.y1=o,t.eachBefore(n),u=[0],r&&t.eachBefore($v),t}function n(t){var n=u[t.depth],r=t.x0+n,i=t.y0+n,o=t.x1-n,h=t.y1-n;o<r&&(r=o=(r+o)/2),h<i&&(i=h=(i+h)/2),t.x0=r,t.y0=i,t.x1=o,t.y1=h,t.children&&(n=u[t.depth+1]=a(t)/2,r+=l(t)-n,i+=c(t)-n,o-=s(t)-n,h-=f(t)-n,o<r&&(r=o=(r+o)/2),h<i&&(i=h=(i+h)/2),e(t,r,i,o,h))}var e=Kv,r=!1,i=1,o=1,u=[0],a=To,c=To,s=To,f=To,l=To;return t.round=function(n){return arguments.length?(r=!!n,t):r},t.size=function(n){return arguments.length?(i=+n[0],o=+n[1],t):[i,o]},t.tile=function(n){return arguments.length?(e=Mo(n),t):e},t.padding=function(n){return arguments.length?t.paddingInner(n).paddingOuter(n):t.paddingInner()},t.paddingInner=function(n){return arguments.length?(a="function"==typeof n?n:Xv(+n),t):a},t.paddingOuter=function(n){return arguments.length?t.paddingTop(n).paddingRight(n).paddingBottom(n).paddingLeft(n):t.paddingTop()},t.paddingTop=function(n){return arguments.length?(c="function"==typeof n?n:Xv(+n),t):c},t.paddingRight=function(n){return arguments.length?(s="function"==typeof n?n:Xv(+n),t):s},t.paddingBottom=function(n){return arguments.length?(f="function"==typeof n?n:Xv(+n),t):f},t.paddingLeft=function(n){return arguments.length?(l="function"==typeof n?n:Xv(+n),t):l},t},t.treemapBinary=function(t,n,e,r,i){function o(t,n,e,r,i,u,a){if(t>=n-1){var s=c[t];return s.x0=r,s.y0=i,s.x1=u,void(s.y1=a)}for(var l=f[t],h=e/2+l,p=t+1,d=n-1;p<d;){var v=p+d>>>1;f[v]<h?p=v+1:d=v}h-f[p-1]<f[p]-h&&t+1<p&&--p;var _=f[p]-l,y=e-_;if(u-r>a-i){var g=(r*y+u*_)/e;o(t,p,_,r,i,g,a),o(p,n,y,g,i,u,a)}else{var m=(i*y+a*_)/e;o(t,p,_,r,i,u,m),o(p,n,y,r,m,u,a)}}var u,a,c=t.children,s=c.length,f=new Array(s+1);for(f[0]=a=u=0;u<s;++u)f[u+1]=a+=c[u].value;o(0,s,t.value,n,e,r,i)},t.treemapDice=Vv,t.treemapSlice=Jv,t.treemapSliceDice=function(t,n,e,r,i){(1&t.depth?Jv:Vv)(t,n,e,r,i)},t.treemapSquarify=Kv,t.treemapResquarify=t_,t.interpolate=cl,t.interpolateArray=nl,t.interpolateBasis=Zf,t.interpolateBasisClosed=Gf,t.interpolateDate=el,t.interpolateNumber=rl,t.interpolateObject=il,t.interpolateRound=sl,t.interpolateString=al,t.interpolateTransformCss=pl,t.interpolateTransformSvg=dl,t.interpolateZoom=_l,t.interpolateRgb=Qf,t.interpolateRgbBasis=Kf,t.interpolateRgbBasisClosed=tl,t.interpolateHsl=yl,t.interpolateHslLong=gl,t.interpolateLab=function(t,n){var e=Kt((t=Ut(t)).l,(n=Ut(n)).l),r=Kt(t.a,n.a),i=Kt(t.b,n.b),o=Kt(t.opacity,n.opacity);return function(n){return t.l=e(n),t.a=r(n),t.b=i(n),t.opacity=o(n),t+""}},t.interpolateHcl=ml,t.interpolateHclLong=xl,t.interpolateCubehelix=bl,t.interpolateCubehelixLong=wl,t.quantize=function(t,n){for(var e=new Array(n),r=0;r<n;++r)e[r]=t(r/(n-1));return e},t.path=ve,t.polygonArea=function(t){for(var n,e=-1,r=t.length,i=t[r-1],o=0;++e<r;)n=i,i=t[e],o+=n[1]*i[0]-n[0]*i[1];return o/2},t.polygonCentroid=function(t){for(var n,e,r=-1,i=t.length,o=0,u=0,a=t[i-1],c=0;++r<i;)n=a,a=t[r],c+=e=n[0]*a[1]-a[0]*n[1],o+=(n[0]+a[0])*e,u+=(n[1]+a[1])*e;return c*=3,[o/c,u/c]},t.polygonHull=function(t){if((e=t.length)<3)return null;var n,e,r=new Array(e),i=new Array(e);for(n=0;n<e;++n)r[n]=[+t[n][0],+t[n][1],n];for(r.sort(Io),n=0;n<e;++n)i[n]=[r[n][0],-r[n][1]];var o=Yo(r),u=Yo(i),a=u[0]===o[0],c=u[u.length-1]===o[o.length-1],s=[];for(n=o.length-1;n>=0;--n)s.push(t[r[o[n]][2]]);for(n=+a;n<u.length-c;++n)s.push(t[r[u[n]][2]]);return s},t.polygonContains=function(t,n){for(var e,r,i=t.length,o=t[i-1],u=n[0],a=n[1],c=o[0],s=o[1],f=!1,l=0;l<i;++l)e=(o=t[l])[0],(r=o[1])>a!=s>a&&u<(c-e)*(a-r)/(s-r)+e&&(f=!f),c=e,s=r;return f},t.polygonLength=function(t){for(var n,e,r=-1,i=t.length,o=t[i-1],u=o[0],a=o[1],c=0;++r<i;)n=u,e=a,n-=u=(o=t[r])[0],e-=a=o[1],c+=Math.sqrt(n*n+e*e);return c},t.quadtree=qe,t.queue=Wo,t.randomUniform=o_,t.randomNormal=u_,t.randomLogNormal=a_,t.randomBates=s_,t.randomIrwinHall=c_,t.randomExponential=f_,t.request=l_,t.html=p_,t.json=d_,t.text=v_,t.xml=__,t.csv=g_,t.tsv=m_,t.scaleBand=Ko,t.scalePoint=function(){return tu(Ko().paddingInner(1))},t.scaleIdentity=fu,t.scaleLinear=su,t.scaleLog=yu,t.scaleOrdinal=Qo,t.scaleImplicit=M_,t.scalePow=mu,t.scaleSqrt=function(){return mu().exponent(.5)},t.scaleQuantile=xu,t.scaleQuantize=bu,t.scaleThreshold=wu,t.scaleTime=function(){return Na(oy,ry,j_,Y_,F_,D_,q_,z_,t.timeFormat).domain([new Date(2e3,0,1),new Date(2e3,0,2)])},t.scaleUtc=function(){return Na(Ay,Sy,py,ly,sy,ay,q_,z_,t.utcFormat).domain([Date.UTC(2e3,0,1),Date.UTC(2e3,0,2)])},t.schemeCategory10=$y,t.schemeCategory20b=Vy,t.schemeCategory20c=Wy,t.schemeCategory20=Zy,t.interpolateCubehelixDefault=Gy,t.interpolateRainbow=function(t){(t<0||t>1)&&(t-=Math.floor(t));var n=Math.abs(t-.5);return Ky.h=360*t-100,Ky.s=1.5-1.5*n,Ky.l=.8-.9*n,Ky+""},t.interpolateWarm=Jy,t.interpolateCool=Qy,t.interpolateViridis=tg,t.interpolateMagma=ng,t.interpolateInferno=eg,t.interpolatePlasma=rg,t.scaleSequential=Ea,t.creator=Hs,t.local=m,t.matcher=Zs,t.mouse=Ks,t.namespace=js,t.namespaces=Bs,t.select=cf,t.selectAll=function(t){return"string"==typeof t?new pt([document.querySelectorAll(t)],[document.documentElement]):new pt([null==t?[]:t],af)},t.selection=dt,t.selector=tf,t.selectorAll=nf,t.style=B,t.touch=sf,t.touches=function(t,n){null==n&&(n=Js().touches);for(var e=0,r=n?n.length:0,i=new Array(r);e<r;++e)i[e]=Qs(t,n[e]);return i},t.window=uf,t.customEvent=N,t.arc=function(){function t(){var t,s,f=+n.apply(this,arguments),l=+e.apply(this,arguments),h=o.apply(this,arguments)-dg,p=u.apply(this,arguments)-dg,d=og(p-h),v=p>h;if(c||(c=t=ve()),l<f&&(s=l,l=f,f=s),l>hg)if(d>vg-hg)c.moveTo(l*ag(h),l*fg(h)),c.arc(0,0,l,h,p,!v),f>hg&&(c.moveTo(f*ag(p),f*fg(p)),c.arc(0,0,f,p,h,v));else{var _,y,g=h,m=p,x=h,b=p,w=d,M=d,T=a.apply(this,arguments)/2,k=T>hg&&(i?+i.apply(this,arguments):lg(f*f+l*l)),N=sg(og(l-f)/2,+r.apply(this,arguments)),S=N,E=N;if(k>hg){var A=Ca(k/f*fg(T)),C=Ca(k/l*fg(T));(w-=2*A)>hg?(A*=v?1:-1,x+=A,b-=A):(w=0,x=b=(h+p)/2),(M-=2*C)>hg?(C*=v?1:-1,g+=C,m-=C):(M=0,g=m=(h+p)/2)}var z=l*ag(g),P=l*fg(g),R=f*ag(b),L=f*fg(b);if(N>hg){var q=l*ag(m),U=l*fg(m),D=f*ag(x),O=f*fg(x);if(d<pg){var F=w>hg?Ua(z,P,D,O,q,U,R,L):[R,L],I=z-F[0],Y=P-F[1],B=q-F[0],j=U-F[1],H=1/fg(Aa((I*B+Y*j)/(lg(I*I+Y*Y)*lg(B*B+j*j)))/2),X=lg(F[0]*F[0]+F[1]*F[1]);S=sg(N,(f-X)/(H-1)),E=sg(N,(l-X)/(H+1))}}M>hg?E>hg?(_=Da(D,O,z,P,l,E,v),y=Da(q,U,R,L,l,E,v),c.moveTo(_.cx+_.x01,_.cy+_.y01),E<N?c.arc(_.cx,_.cy,E,ug(_.y01,_.x01),ug(y.y01,y.x01),!v):(c.arc(_.cx,_.cy,E,ug(_.y01,_.x01),ug(_.y11,_.x11),!v),c.arc(0,0,l,ug(_.cy+_.y11,_.cx+_.x11),ug(y.cy+y.y11,y.cx+y.x11),!v),c.arc(y.cx,y.cy,E,ug(y.y11,y.x11),ug(y.y01,y.x01),!v))):(c.moveTo(z,P),c.arc(0,0,l,g,m,!v)):c.moveTo(z,P),f>hg&&w>hg?S>hg?(_=Da(R,L,q,U,f,-S,v),y=Da(z,P,D,O,f,-S,v),c.lineTo(_.cx+_.x01,_.cy+_.y01),S<N?c.arc(_.cx,_.cy,S,ug(_.y01,_.x01),ug(y.y01,y.x01),!v):(c.arc(_.cx,_.cy,S,ug(_.y01,_.x01),ug(_.y11,_.x11),!v),c.arc(0,0,f,ug(_.cy+_.y11,_.cx+_.x11),ug(y.cy+y.y11,y.cx+y.x11),v),c.arc(y.cx,y.cy,S,ug(y.y11,y.x11),ug(y.y01,y.x01),!v))):c.arc(0,0,f,b,x,v):c.lineTo(R,L)}else c.moveTo(0,0);if(c.closePath(),t)return c=null,t+""||null}var n=za,e=Pa,r=ig(0),i=null,o=Ra,u=La,a=qa,c=null;return t.centroid=function(){var t=(+n.apply(this,arguments)+ +e.apply(this,arguments))/2,r=(+o.apply(this,arguments)+ +u.apply(this,arguments))/2-pg/2;return[ag(r)*t,fg(r)*t]},t.innerRadius=function(e){return arguments.length?(n="function"==typeof e?e:ig(+e),t):n},t.outerRadius=function(n){return arguments.length?(e="function"==typeof n?n:ig(+n),t):e},t.cornerRadius=function(n){return arguments.length?(r="function"==typeof n?n:ig(+n),t):r},t.padRadius=function(n){return arguments.length?(i=null==n?null:"function"==typeof n?n:ig(+n),t):i},t.startAngle=function(n){return arguments.length?(o="function"==typeof n?n:ig(+n),t):o},t.endAngle=function(n){return arguments.length?(u="function"==typeof n?n:ig(+n),t):u},t.padAngle=function(n){return arguments.length?(a="function"==typeof n?n:ig(+n),t):a},t.context=function(n){return arguments.length?(c=null==n?null:n,t):c},t},t.area=gg,t.line=yg,t.pie=function(){function t(t){var a,c,s,f,l,h=t.length,p=0,d=new Array(h),v=new Array(h),_=+i.apply(this,arguments),y=Math.min(vg,Math.max(-vg,o.apply(this,arguments)-_)),g=Math.min(Math.abs(y)/h,u.apply(this,arguments)),m=g*(y<0?-1:1);for(a=0;a<h;++a)(l=v[d[a]=a]=+n(t[a],a,t))>0&&(p+=l);for(null!=e?d.sort(function(t,n){return e(v[t],v[n])}):null!=r&&d.sort(function(n,e){return r(t[n],t[e])}),a=0,s=p?(y-h*m)/p:0;a<h;++a,_=f)c=d[a],f=_+((l=v[c])>0?l*s:0)+m,v[c]={data:t[c],index:a,value:l,startAngle:_,endAngle:f,padAngle:g};return v}var n=xg,e=mg,r=null,i=ig(0),o=ig(vg),u=ig(0);return t.value=function(e){return arguments.length?(n="function"==typeof e?e:ig(+e),t):n},t.sortValues=function(n){return arguments.length?(e=n,r=null,t):e},t.sort=function(n){return arguments.length?(r=n,e=null,t):r},t.startAngle=function(n){return arguments.length?(i="function"==typeof n?n:ig(+n),t):i},t.endAngle=function(n){return arguments.length?(o="function"==typeof n?n:ig(+n),t):o},t.padAngle=function(n){return arguments.length?(u="function"==typeof n?n:ig(+n),t):u},t},t.areaRadial=Mg,t.radialArea=Mg,t.lineRadial=wg,t.radialLine=wg,t.pointRadial=Tg,t.linkHorizontal=function(){return $a(Va)},t.linkVertical=function(){return $a(Wa)},t.linkRadial=function(){var t=$a(Za);return t.angle=t.x,delete t.x,t.radius=t.y,delete t.y,t},t.symbol=function(){function t(){var t;if(r||(r=t=ve()),n.apply(this,arguments).draw(r,+e.apply(this,arguments)),t)return r=null,t+""||null}var n=ig(Ng),e=ig(64),r=null;return t.type=function(e){return arguments.length?(n="function"==typeof e?e:ig(e),t):n},t.size=function(n){return arguments.length?(e="function"==typeof n?n:ig(+n),t):e},t.context=function(n){return arguments.length?(r=null==n?null:n,t):r},t},t.symbols=jg,t.symbolCircle=Ng,t.symbolCross=Sg,t.symbolDiamond=Cg,t.symbolSquare=qg,t.symbolStar=Lg,t.symbolTriangle=Dg,t.symbolWye=Bg,t.curveBasisClosed=function(t){return new Qa(t)},t.curveBasisOpen=function(t){return new Ka(t)},t.curveBasis=function(t){return new Ja(t)},t.curveBundle=Xg,t.curveCardinalClosed=Vg,t.curveCardinalOpen=Wg,t.curveCardinal=$g,t.curveCatmullRomClosed=Gg,t.curveCatmullRomOpen=Jg,t.curveCatmullRom=Zg,t.curveLinearClosed=function(t){return new sc(t)},t.curveLinear=_g,t.curveMonotoneX=function(t){return new dc(t)},t.curveMonotoneY=function(t){return new vc(t)},t.curveNatural=function(t){return new yc(t)},t.curveStep=function(t){return new mc(t,.5)},t.curveStepAfter=function(t){return new mc(t,1)},t.curveStepBefore=function(t){return new mc(t,0)},t.stack=function(){function t(t){var o,u,a=n.apply(this,arguments),c=t.length,s=a.length,f=new Array(s);for(o=0;o<s;++o){for(var l,h=a[o],p=f[o]=new Array(c),d=0;d<c;++d)p[d]=l=[0,+i(t[d],h,d,t)],l.data=t[d];p.key=h}for(o=0,u=e(f);o<s;++o)f[u[o]].index=o;return r(f,u),f}var n=ig([]),e=Kg,r=Qg,i=xc;return t.keys=function(e){return arguments.length?(n="function"==typeof e?e:ig(kg.call(e)),t):n},t.value=function(n){return arguments.length?(i="function"==typeof n?n:ig(+n),t):i},t.order=function(n){return arguments.length?(e=null==n?Kg:"function"==typeof n?n:ig(kg.call(n)),t):e},t.offset=function(n){return arguments.length?(r=null==n?Qg:n,t):r},t},t.stackOffsetExpand=function(t,n){if((r=t.length)>0){for(var e,r,i,o=0,u=t[0].length;o<u;++o){for(i=e=0;e<r;++e)i+=t[e][o][1]||0;if(i)for(e=0;e<r;++e)t[e][o][1]/=i}Qg(t,n)}},t.stackOffsetDiverging=function(t,n){if((a=t.length)>1)for(var e,r,i,o,u,a,c=0,s=t[n[0]].length;c<s;++c)for(o=u=0,e=0;e<a;++e)(i=(r=t[n[e]][c])[1]-r[0])>=0?(r[0]=o,r[1]=o+=i):i<0?(r[1]=u,r[0]=u+=i):r[0]=o},t.stackOffsetNone=Qg,t.stackOffsetSilhouette=function(t,n){if((e=t.length)>0){for(var e,r=0,i=t[n[0]],o=i.length;r<o;++r){for(var u=0,a=0;u<e;++u)a+=t[u][r][1]||0;i[r][1]+=i[r][0]=-a/2}Qg(t,n)}},t.stackOffsetWiggle=function(t,n){if((i=t.length)>0&&(r=(e=t[n[0]]).length)>0){for(var e,r,i,o=0,u=1;u<r;++u){for(var a=0,c=0,s=0;a<i;++a){for(var f=t[n[a]],l=f[u][1]||0,h=(l-(f[u-1][1]||0))/2,p=0;p<a;++p){var d=t[n[p]];h+=(d[u][1]||0)-(d[u-1][1]||0)}c+=l,s+=h*l}e[u-1][1]+=e[u-1][0]=o,c&&(o-=s/c)}e[u-1][1]+=e[u-1][0]=o,Qg(t,n)}},t.stackOrderAscending=tm,t.stackOrderDescending=function(t){return tm(t).reverse()},t.stackOrderInsideOut=function(t){var n,e,r=t.length,i=t.map(bc),o=Kg(t).sort(function(t,n){return i[n]-i[t]}),u=0,a=0,c=[],s=[];for(n=0;n<r;++n)e=o[n],u<a?(u+=i[e],c.push(e)):(a+=i[e],s.push(e));return s.reverse().concat(c)},t.stackOrderNone=Kg,t.stackOrderReverse=function(t){return Kg(t).reverse()},t.timeInterval=Mu,t.timeMillisecond=z_,t.timeMilliseconds=P_,t.utcMillisecond=z_,t.utcMilliseconds=P_,t.timeSecond=q_,t.timeSeconds=U_,t.utcSecond=q_,t.utcSeconds=U_,t.timeMinute=D_,t.timeMinutes=O_,t.timeHour=F_,t.timeHours=I_,t.timeDay=Y_,t.timeDays=B_,t.timeWeek=j_,t.timeWeeks=G_,t.timeSunday=j_,t.timeSundays=G_,t.timeMonday=H_,t.timeMondays=J_,t.timeTuesday=X_,t.timeTuesdays=Q_,t.timeWednesday=$_,t.timeWednesdays=K_,t.timeThursday=V_,t.timeThursdays=ty,t.timeFriday=W_,t.timeFridays=ny,t.timeSaturday=Z_,t.timeSaturdays=ey,t.timeMonth=ry,t.timeMonths=iy,t.timeYear=oy,t.timeYears=uy,t.utcMinute=ay,t.utcMinutes=cy,t.utcHour=sy,t.utcHours=fy,t.utcDay=ly,t.utcDays=hy,t.utcWeek=py,t.utcWeeks=xy,t.utcSunday=py,t.utcSundays=xy,t.utcMonday=dy,t.utcMondays=by,t.utcTuesday=vy,t.utcTuesdays=wy,t.utcWednesday=_y,t.utcWednesdays=My,t.utcThursday=yy,t.utcThursdays=Ty,t.utcFriday=gy,t.utcFridays=ky,t.utcSaturday=my,t.utcSaturdays=Ny,t.utcMonth=Sy,t.utcMonths=Ey,t.utcYear=Ay,t.utcYears=zy,t.timeFormatDefaultLocale=Ma,t.timeFormatLocale=Au,t.isoFormat=Uy,t.isoParse=Dy,t.now=ln,t.timer=dn,t.timerFlush=vn,t.timeout=Pl,t.interval=function(t,n,e){var r=new pn,i=n;return null==n?(r.restart(t,n,e),r):(n=+n,e=null==e?ln():+e,r.restart(function o(u){u+=i,r.restart(o,i+=n,e),t(u)},n,e),r)},t.transition=Jn,t.active=function(t,n){var e,r,i=t.__transition;if(i){n=null==n?null:n+"";for(r in i)if((e=i[r]).state>Ul&&e.name===n)return new Gn([[t]],yh,n,+r)}return null},t.interrupt=jl,t.voronoi=function(){function t(t){return new Kc(t.map(function(r,i){var o=[Math.round(n(r,i,t)/sm)*sm,Math.round(e(r,i,t)/sm)*sm];return o.index=i,o.data=r,o}),r)}var n=wc,e=Mc,r=null;return t.polygons=function(n){return t(n).polygons()},t.links=function(n){return t(n).links()},t.triangles=function(n){return t(n).triangles()},t.x=function(e){return arguments.length?(n="function"==typeof e?e:nm(+e),t):n},t.y=function(n){return arguments.length?(e="function"==typeof n?n:nm(+n),t):e},t.extent=function(n){return arguments.length?(r=null==n?null:[[+n[0][0],+n[0][1]],[+n[1][0],+n[1][1]]],t):r&&[[r[0][0],r[0][1]],[r[1][0],r[1][1]]]},t.size=function(n){return arguments.length?(r=null==n?null:[[0,0],[+n[0],+n[1]]],t):r&&[r[1][0]-r[0][0],r[1][1]-r[0][1]]},t},t.zoom=function(){function n(t){t.property("__zoom",us).on("wheel.zoom",s).on("mousedown.zoom",f).on("dblclick.zoom",l).filter(cs).on("touchstart.zoom",p).on("touchmove.zoom",d).on("touchend.zoom touchcancel.zoom",v).style("touch-action","none").style("-webkit-tap-highlight-color","rgba(0,0,0,0)")}function e(t,n){return(n=Math.max(b,Math.min(w,n)))===t.k?t:new ns(n,t.x,t.y)}function r(t,n,e){var r=n[0]-e[0]*t.k,i=n[1]-e[1]*t.k;return r===t.x&&i===t.y?t:new ns(t.k,r,i)}function i(t,n){var e=t.invertX(n[0][0])-M,r=t.invertX(n[1][0])-T,i=t.invertY(n[0][1])-k,o=t.invertY(n[1][1])-S;return t.translate(r>e?(e+r)/2:Math.min(0,e)||Math.max(0,r),o>i?(i+o)/2:Math.min(0,i)||Math.max(0,o))}function o(t){return[(+t[0][0]+ +t[1][0])/2,(+t[0][1]+ +t[1][1])/2]}function u(t,n,e){t.on("start.zoom",function(){a(this,arguments).start()}).on("interrupt.zoom end.zoom",function(){a(this,arguments).end()}).tween("zoom",function(){var t=this,r=arguments,i=a(t,r),u=m.apply(t,r),c=e||o(u),s=Math.max(u[1][0]-u[0][0],u[1][1]-u[0][1]),f=t.__zoom,l="function"==typeof n?n.apply(t,r):n,h=A(f.invert(c).concat(s/f.k),l.invert(c).concat(s/l.k));return function(t){if(1===t)t=l;else{var n=h(t),e=s/n[2];t=new ns(e,c[0]-n[0]*e,c[1]-n[1]*e)}i.zoom(null,t)}})}function a(t,n){for(var e,r=0,i=C.length;r<i;++r)if((e=C[r]).that===t)return e;return new c(t,n)}function c(t,n){this.that=t,this.args=n,this.index=-1,this.active=0,this.extent=m.apply(t,n)}function s(){if(g.apply(this,arguments)){var t=a(this,arguments),n=this.__zoom,o=Math.max(b,Math.min(w,n.k*Math.pow(2,x.apply(this,arguments)))),u=Ks(this);if(t.wheel)t.mouse[0][0]===u[0]&&t.mouse[0][1]===u[1]||(t.mouse[1]=n.invert(t.mouse[0]=u)),clearTimeout(t.wheel);else{if(n.k===o)return;t.mouse=[u,n.invert(u)],jl(this),t.start()}pm(),t.wheel=setTimeout(function(){t.wheel=null,t.end()},R),t.zoom("mouse",i(r(e(n,o),t.mouse[0],t.mouse[1]),t.extent))}}function f(){if(!y&&g.apply(this,arguments)){var n=a(this,arguments),e=cf(t.event.view).on("mousemove.zoom",function(){if(pm(),!n.moved){var e=t.event.clientX-u,o=t.event.clientY-c;n.moved=e*e+o*o>L}n.zoom("mouse",i(r(n.that.__zoom,n.mouse[0]=Ks(n.that),n.mouse[1]),n.extent))},!0).on("mouseup.zoom",function(){e.on("mousemove.zoom mouseup.zoom",null),_t(t.event.view,n.moved),pm(),n.end()},!0),o=Ks(this),u=t.event.clientX,c=t.event.clientY;lf(t.event.view),rs(),n.mouse=[o,this.__zoom.invert(o)],jl(this),n.start()}}function l(){if(g.apply(this,arguments)){var o=this.__zoom,a=Ks(this),c=o.invert(a),s=i(r(e(o,o.k*(t.event.shiftKey?.5:2)),a,c),m.apply(this,arguments));pm(),E>0?cf(this).transition().duration(E).call(u,s,a):cf(this).call(n.transform,s)}}function p(){if(g.apply(this,arguments)){var n,e,r,i,o=a(this,arguments),u=t.event.changedTouches,c=u.length;for(rs(),e=0;e<c;++e)r=u[e],i=[i=sf(this,u,r.identifier),this.__zoom.invert(i),r.identifier],o.touch0?o.touch1||(o.touch1=i):(o.touch0=i,n=!0);if(_&&(_=clearTimeout(_),!o.touch1))return o.end(),void((i=cf(this).on("dblclick.zoom"))&&i.apply(this,arguments));n&&(_=setTimeout(function(){_=null},P),jl(this),o.start())}}function d(){var n,o,u,c,s=a(this,arguments),f=t.event.changedTouches,l=f.length;for(pm(),_&&(_=clearTimeout(_)),n=0;n<l;++n)o=f[n],u=sf(this,f,o.identifier),s.touch0&&s.touch0[2]===o.identifier?s.touch0[0]=u:s.touch1&&s.touch1[2]===o.identifier&&(s.touch1[0]=u);if(o=s.that.__zoom,s.touch1){var h=s.touch0[0],p=s.touch0[1],d=s.touch1[0],v=s.touch1[1],y=(y=d[0]-h[0])*y+(y=d[1]-h[1])*y,g=(g=v[0]-p[0])*g+(g=v[1]-p[1])*g;o=e(o,Math.sqrt(y/g)),u=[(h[0]+d[0])/2,(h[1]+d[1])/2],c=[(p[0]+v[0])/2,(p[1]+v[1])/2]}else{if(!s.touch0)return;u=s.touch0[0],c=s.touch0[1]}s.zoom("touch",i(r(o,u,c),s.extent))}function v(){var n,e,r=a(this,arguments),i=t.event.changedTouches,o=i.length;for(rs(),y&&clearTimeout(y),y=setTimeout(function(){y=null},P),n=0;n<o;++n)e=i[n],r.touch0&&r.touch0[2]===e.identifier?delete r.touch0:r.touch1&&r.touch1[2]===e.identifier&&delete r.touch1;r.touch1&&!r.touch0&&(r.touch0=r.touch1,delete r.touch1),r.touch0?r.touch0[1]=this.__zoom.invert(r.touch0[0]):r.end()}var _,y,g=is,m=os,x=as,b=0,w=1/0,M=-w,T=w,k=M,S=T,E=250,A=_l,C=[],z=h("start","zoom","end"),P=500,R=150,L=0;return n.transform=function(t,n){var e=t.selection?t.selection():t;e.property("__zoom",us),t!==e?u(t,n):e.interrupt().each(function(){a(this,arguments).start().zoom(null,"function"==typeof n?n.apply(this,arguments):n).end()})},n.scaleBy=function(t,e){n.scaleTo(t,function(){return this.__zoom.k*("function"==typeof e?e.apply(this,arguments):e)})},n.scaleTo=function(t,u){n.transform(t,function(){var t=m.apply(this,arguments),n=this.__zoom,a=o(t),c=n.invert(a);return i(r(e(n,"function"==typeof u?u.apply(this,arguments):u),a,c),t)})},n.translateBy=function(t,e,r){n.transform(t,function(){return i(this.__zoom.translate("function"==typeof e?e.apply(this,arguments):e,"function"==typeof r?r.apply(this,arguments):r),m.apply(this,arguments))})},n.translateTo=function(t,e,r){n.transform(t,function(){var t=m.apply(this,arguments),n=this.__zoom,u=o(t);return i(hm.translate(u[0],u[1]).scale(n.k).translate("function"==typeof e?-e.apply(this,arguments):-e,"function"==typeof r?-r.apply(this,arguments):-r),t)})},c.prototype={start:function(){return 1==++this.active&&(this.index=C.push(this)-1,this.emit("start")),this},zoom:function(t,n){return this.mouse&&"mouse"!==t&&(this.mouse[1]=n.invert(this.mouse[0])),this.touch0&&"touch"!==t&&(this.touch0[1]=n.invert(this.touch0[0])),this.touch1&&"touch"!==t&&(this.touch1[1]=n.invert(this.touch1[0])),this.that.__zoom=n,this.emit("zoom"),this},end:function(){return 0==--this.active&&(C.splice(this.index,1),this.index=-1,this.emit("end")),this},emit:function(t){N(new ts(n,t,this.that.__zoom),z.apply,z,[t,this.that,this.args])}},n.wheelDelta=function(t){return arguments.length?(x="function"==typeof t?t:lm(+t),n):x},n.filter=function(t){return arguments.length?(g="function"==typeof t?t:lm(!!t),n):g},n.extent=function(t){return arguments.length?(m="function"==typeof t?t:lm([[+t[0][0],+t[0][1]],[+t[1][0],+t[1][1]]]),n):m},n.scaleExtent=function(t){return arguments.length?(b=+t[0],w=+t[1],n):[b,w]},n.translateExtent=function(t){return arguments.length?(M=+t[0][0],T=+t[1][0],k=+t[0][1],S=+t[1][1],n):[[M,k],[T,S]]},n.duration=function(t){return arguments.length?(E=+t,n):E},n.interpolate=function(t){return arguments.length?(A=t,n):A},n.on=function(){var t=z.on.apply(z,arguments);return t===z?n:t},n.clickDistance=function(t){return arguments.length?(L=(t=+t)*t,n):Math.sqrt(L)},n},t.zoomTransform=es,t.zoomIdentity=hm,Object.defineProperty(t,"__esModule",{value:!0})});
// https://d3js.org/d3-scale-chromatic/ Version 1.1.1. Copyright 2017 Mike Bostock.
!function(f,e){"object"==typeof exports&&"undefined"!=typeof module?e(exports,require("d3-interpolate")):"function"==typeof define&&define.amd?define(["exports","d3-interpolate"],e):e(f.d3=f.d3||{},f.d3)}(this,function(f,e){"use strict";var d=function(f){for(var e=f.length/6|0,d=new Array(e),c=0;c<e;)d[c]="#"+f.slice(6*c,6*++c);return d},c=d("7fc97fbeaed4fdc086ffff99386cb0f0027fbf5b17666666"),a=d("1b9e77d95f027570b3e7298a66a61ee6ab02a6761d666666"),b=d("a6cee31f78b4b2df8a33a02cfb9a99e31a1cfdbf6fff7f00cab2d66a3d9affff99b15928"),r=d("fbb4aeb3cde3ccebc5decbe4fed9a6ffffcce5d8bdfddaecf2f2f2"),n=d("b3e2cdfdcdaccbd5e8f4cae4e6f5c9fff2aef1e2cccccccc"),t=d("e41a1c377eb84daf4a984ea3ff7f00ffff33a65628f781bf999999"),o=d("66c2a5fc8d628da0cbe78ac3a6d854ffd92fe5c494b3b3b3"),p=d("8dd3c7ffffb3bebadafb807280b1d3fdb462b3de69fccde5d9d9d9bc80bdccebc5ffed6f"),m=function(f){return e.interpolateRgbBasis(f[f.length-1])},s=new Array(3).concat("d8b365f5f5f55ab4ac","a6611adfc27d80cdc1018571","a6611adfc27df5f5f580cdc1018571","8c510ad8b365f6e8c3c7eae55ab4ac01665e","8c510ad8b365f6e8c3f5f5f5c7eae55ab4ac01665e","8c510abf812ddfc27df6e8c3c7eae580cdc135978f01665e","8c510abf812ddfc27df6e8c3f5f5f5c7eae580cdc135978f01665e","5430058c510abf812ddfc27df6e8c3c7eae580cdc135978f01665e003c30","5430058c510abf812ddfc27df6e8c3f5f5f5c7eae580cdc135978f01665e003c30").map(d),l=m(s),i=new Array(3).concat("af8dc3f7f7f77fbf7b","7b3294c2a5cfa6dba0008837","7b3294c2a5cff7f7f7a6dba0008837","762a83af8dc3e7d4e8d9f0d37fbf7b1b7837","762a83af8dc3e7d4e8f7f7f7d9f0d37fbf7b1b7837","762a839970abc2a5cfe7d4e8d9f0d3a6dba05aae611b7837","762a839970abc2a5cfe7d4e8f7f7f7d9f0d3a6dba05aae611b7837","40004b762a839970abc2a5cfe7d4e8d9f0d3a6dba05aae611b783700441b","40004b762a839970abc2a5cfe7d4e8f7f7f7d9f0d3a6dba05aae611b783700441b").map(d),u=m(i),h=new Array(3).concat("e9a3c9f7f7f7a1d76a","d01c8bf1b6dab8e1864dac26","d01c8bf1b6daf7f7f7b8e1864dac26","c51b7de9a3c9fde0efe6f5d0a1d76a4d9221","c51b7de9a3c9fde0eff7f7f7e6f5d0a1d76a4d9221","c51b7dde77aef1b6dafde0efe6f5d0b8e1867fbc414d9221","c51b7dde77aef1b6dafde0eff7f7f7e6f5d0b8e1867fbc414d9221","8e0152c51b7dde77aef1b6dafde0efe6f5d0b8e1867fbc414d9221276419","8e0152c51b7dde77aef1b6dafde0eff7f7f7e6f5d0b8e1867fbc414d9221276419").map(d),y=m(h),A=new Array(3).concat("998ec3f7f7f7f1a340","5e3c99b2abd2fdb863e66101","5e3c99b2abd2f7f7f7fdb863e66101","542788998ec3d8daebfee0b6f1a340b35806","542788998ec3d8daebf7f7f7fee0b6f1a340b35806","5427888073acb2abd2d8daebfee0b6fdb863e08214b35806","5427888073acb2abd2d8daebf7f7f7fee0b6fdb863e08214b35806","2d004b5427888073acb2abd2d8daebfee0b6fdb863e08214b358067f3b08","2d004b5427888073acb2abd2d8daebf7f7f7fee0b6fdb863e08214b358067f3b08").map(d),w=m(A),B=new Array(3).concat("ef8a62f7f7f767a9cf","ca0020f4a58292c5de0571b0","ca0020f4a582f7f7f792c5de0571b0","b2182bef8a62fddbc7d1e5f067a9cf2166ac","b2182bef8a62fddbc7f7f7f7d1e5f067a9cf2166ac","b2182bd6604df4a582fddbc7d1e5f092c5de4393c32166ac","b2182bd6604df4a582fddbc7f7f7f7d1e5f092c5de4393c32166ac","67001fb2182bd6604df4a582fddbc7d1e5f092c5de4393c32166ac053061","67001fb2182bd6604df4a582fddbc7f7f7f7d1e5f092c5de4393c32166ac053061").map(d),G=m(B),P=new Array(3).concat("ef8a62ffffff999999","ca0020f4a582bababa404040","ca0020f4a582ffffffbababa404040","b2182bef8a62fddbc7e0e0e09999994d4d4d","b2182bef8a62fddbc7ffffffe0e0e09999994d4d4d","b2182bd6604df4a582fddbc7e0e0e0bababa8787874d4d4d","b2182bd6604df4a582fddbc7ffffffe0e0e0bababa8787874d4d4d","67001fb2182bd6604df4a582fddbc7e0e0e0bababa8787874d4d4d1a1a1a","67001fb2182bd6604df4a582fddbc7ffffffe0e0e0bababa8787874d4d4d1a1a1a").map(d),R=m(P),Y=new Array(3).concat("fc8d59ffffbf91bfdb","d7191cfdae61abd9e92c7bb6","d7191cfdae61ffffbfabd9e92c7bb6","d73027fc8d59fee090e0f3f891bfdb4575b4","d73027fc8d59fee090ffffbfe0f3f891bfdb4575b4","d73027f46d43fdae61fee090e0f3f8abd9e974add14575b4","d73027f46d43fdae61fee090ffffbfe0f3f8abd9e974add14575b4","a50026d73027f46d43fdae61fee090e0f3f8abd9e974add14575b4313695","a50026d73027f46d43fdae61fee090ffffbfe0f3f8abd9e974add14575b4313695").map(d),O=m(Y),g=new Array(3).concat("fc8d59ffffbf91cf60","d7191cfdae61a6d96a1a9641","d7191cfdae61ffffbfa6d96a1a9641","d73027fc8d59fee08bd9ef8b91cf601a9850","d73027fc8d59fee08bffffbfd9ef8b91cf601a9850","d73027f46d43fdae61fee08bd9ef8ba6d96a66bd631a9850","d73027f46d43fdae61fee08bffffbfd9ef8ba6d96a66bd631a9850","a50026d73027f46d43fdae61fee08bd9ef8ba6d96a66bd631a9850006837","a50026d73027f46d43fdae61fee08bffffbfd9ef8ba6d96a66bd631a9850006837").map(d),S=m(g),v=new Array(3).concat("fc8d59ffffbf99d594","d7191cfdae61abdda42b83ba","d7191cfdae61ffffbfabdda42b83ba","d53e4ffc8d59fee08be6f59899d5943288bd","d53e4ffc8d59fee08bffffbfe6f59899d5943288bd","d53e4ff46d43fdae61fee08be6f598abdda466c2a53288bd","d53e4ff46d43fdae61fee08bffffbfe6f598abdda466c2a53288bd","9e0142d53e4ff46d43fdae61fee08be6f598abdda466c2a53288bd5e4fa2","9e0142d53e4ff46d43fdae61fee08bffffbfe6f598abdda466c2a53288bd5e4fa2").map(d),x=m(v),j=new Array(3).concat("e5f5f999d8c92ca25f","edf8fbb2e2e266c2a4238b45","edf8fbb2e2e266c2a42ca25f006d2c","edf8fbccece699d8c966c2a42ca25f006d2c","edf8fbccece699d8c966c2a441ae76238b45005824","f7fcfde5f5f9ccece699d8c966c2a441ae76238b45005824","f7fcfde5f5f9ccece699d8c966c2a441ae76238b45006d2c00441b").map(d),_=m(j),k=new Array(3).concat("e0ecf49ebcda8856a7","edf8fbb3cde38c96c688419d","edf8fbb3cde38c96c68856a7810f7c","edf8fbbfd3e69ebcda8c96c68856a7810f7c","edf8fbbfd3e69ebcda8c96c68c6bb188419d6e016b","f7fcfde0ecf4bfd3e69ebcda8c96c68c6bb188419d6e016b","f7fcfde0ecf4bfd3e69ebcda8c96c68c6bb188419d810f7c4d004b").map(d),q=m(k),D=new Array(3).concat("e0f3dba8ddb543a2ca","f0f9e8bae4bc7bccc42b8cbe","f0f9e8bae4bc7bccc443a2ca0868ac","f0f9e8ccebc5a8ddb57bccc443a2ca0868ac","f0f9e8ccebc5a8ddb57bccc44eb3d32b8cbe08589e","f7fcf0e0f3dbccebc5a8ddb57bccc44eb3d32b8cbe08589e","f7fcf0e0f3dbccebc5a8ddb57bccc44eb3d32b8cbe0868ac084081").map(d),M=m(D),z=new Array(3).concat("fee8c8fdbb84e34a33","fef0d9fdcc8afc8d59d7301f","fef0d9fdcc8afc8d59e34a33b30000","fef0d9fdd49efdbb84fc8d59e34a33b30000","fef0d9fdd49efdbb84fc8d59ef6548d7301f990000","fff7ecfee8c8fdd49efdbb84fc8d59ef6548d7301f990000","fff7ecfee8c8fdd49efdbb84fc8d59ef6548d7301fb300007f0000").map(d),C=m(z),E=new Array(3).concat("ece2f0a6bddb1c9099","f6eff7bdc9e167a9cf02818a","f6eff7bdc9e167a9cf1c9099016c59","f6eff7d0d1e6a6bddb67a9cf1c9099016c59","f6eff7d0d1e6a6bddb67a9cf3690c002818a016450","fff7fbece2f0d0d1e6a6bddb67a9cf3690c002818a016450","fff7fbece2f0d0d1e6a6bddb67a9cf3690c002818a016c59014636").map(d),F=m(E),H=new Array(3).concat("ece7f2a6bddb2b8cbe","f1eef6bdc9e174a9cf0570b0","f1eef6bdc9e174a9cf2b8cbe045a8d","f1eef6d0d1e6a6bddb74a9cf2b8cbe045a8d","f1eef6d0d1e6a6bddb74a9cf3690c00570b0034e7b","fff7fbece7f2d0d1e6a6bddb74a9cf3690c00570b0034e7b","fff7fbece7f2d0d1e6a6bddb74a9cf3690c00570b0045a8d023858").map(d),I=m(H),J=new Array(3).concat("e7e1efc994c7dd1c77","f1eef6d7b5d8df65b0ce1256","f1eef6d7b5d8df65b0dd1c77980043","f1eef6d4b9dac994c7df65b0dd1c77980043","f1eef6d4b9dac994c7df65b0e7298ace125691003f","f7f4f9e7e1efd4b9dac994c7df65b0e7298ace125691003f","f7f4f9e7e1efd4b9dac994c7df65b0e7298ace125698004367001f").map(d),K=m(J),L=new Array(3).concat("fde0ddfa9fb5c51b8a","feebe2fbb4b9f768a1ae017e","feebe2fbb4b9f768a1c51b8a7a0177","feebe2fcc5c0fa9fb5f768a1c51b8a7a0177","feebe2fcc5c0fa9fb5f768a1dd3497ae017e7a0177","fff7f3fde0ddfcc5c0fa9fb5f768a1dd3497ae017e7a0177","fff7f3fde0ddfcc5c0fa9fb5f768a1dd3497ae017e7a017749006a").map(d),N=m(L),Q=new Array(3).concat("edf8b17fcdbb2c7fb8","ffffcca1dab441b6c4225ea8","ffffcca1dab441b6c42c7fb8253494","ffffccc7e9b47fcdbb41b6c42c7fb8253494","ffffccc7e9b47fcdbb41b6c41d91c0225ea80c2c84","ffffd9edf8b1c7e9b47fcdbb41b6c41d91c0225ea80c2c84","ffffd9edf8b1c7e9b47fcdbb41b6c41d91c0225ea8253494081d58").map(d),T=m(Q),U=new Array(3).concat("f7fcb9addd8e31a354","ffffccc2e69978c679238443","ffffccc2e69978c67931a354006837","ffffccd9f0a3addd8e78c67931a354006837","ffffccd9f0a3addd8e78c67941ab5d238443005a32","ffffe5f7fcb9d9f0a3addd8e78c67941ab5d238443005a32","ffffe5f7fcb9d9f0a3addd8e78c67941ab5d238443006837004529").map(d),V=m(U),W=new Array(3).concat("fff7bcfec44fd95f0e","ffffd4fed98efe9929cc4c02","ffffd4fed98efe9929d95f0e993404","ffffd4fee391fec44ffe9929d95f0e993404","ffffd4fee391fec44ffe9929ec7014cc4c028c2d04","ffffe5fff7bcfee391fec44ffe9929ec7014cc4c028c2d04","ffffe5fff7bcfee391fec44ffe9929ec7014cc4c02993404662506").map(d),X=m(W),Z=new Array(3).concat("ffeda0feb24cf03b20","ffffb2fecc5cfd8d3ce31a1c","ffffb2fecc5cfd8d3cf03b20bd0026","ffffb2fed976feb24cfd8d3cf03b20bd0026","ffffb2fed976feb24cfd8d3cfc4e2ae31a1cb10026","ffffccffeda0fed976feb24cfd8d3cfc4e2ae31a1cb10026","ffffccffeda0fed976feb24cfd8d3cfc4e2ae31a1cbd0026800026").map(d),$=m(Z),ff=new Array(3).concat("deebf79ecae13182bd","eff3ffbdd7e76baed62171b5","eff3ffbdd7e76baed63182bd08519c","eff3ffc6dbef9ecae16baed63182bd08519c","eff3ffc6dbef9ecae16baed64292c62171b5084594","f7fbffdeebf7c6dbef9ecae16baed64292c62171b5084594","f7fbffdeebf7c6dbef9ecae16baed64292c62171b508519c08306b").map(d),ef=m(ff),df=new Array(3).concat("e5f5e0a1d99b31a354","edf8e9bae4b374c476238b45","edf8e9bae4b374c47631a354006d2c","edf8e9c7e9c0a1d99b74c47631a354006d2c","edf8e9c7e9c0a1d99b74c47641ab5d238b45005a32","f7fcf5e5f5e0c7e9c0a1d99b74c47641ab5d238b45005a32","f7fcf5e5f5e0c7e9c0a1d99b74c47641ab5d238b45006d2c00441b").map(d),cf=m(df),af=new Array(3).concat("f0f0f0bdbdbd636363","f7f7f7cccccc969696525252","f7f7f7cccccc969696636363252525","f7f7f7d9d9d9bdbdbd969696636363252525","f7f7f7d9d9d9bdbdbd969696737373525252252525","fffffff0f0f0d9d9d9bdbdbd969696737373525252252525","fffffff0f0f0d9d9d9bdbdbd969696737373525252252525000000").map(d),bf=m(af),rf=new Array(3).concat("efedf5bcbddc756bb1","f2f0f7cbc9e29e9ac86a51a3","f2f0f7cbc9e29e9ac8756bb154278f","f2f0f7dadaebbcbddc9e9ac8756bb154278f","f2f0f7dadaebbcbddc9e9ac8807dba6a51a34a1486","fcfbfdefedf5dadaebbcbddc9e9ac8807dba6a51a34a1486","fcfbfdefedf5dadaebbcbddc9e9ac8807dba6a51a354278f3f007d").map(d),nf=m(rf),tf=new Array(3).concat("fee0d2fc9272de2d26","fee5d9fcae91fb6a4acb181d","fee5d9fcae91fb6a4ade2d26a50f15","fee5d9fcbba1fc9272fb6a4ade2d26a50f15","fee5d9fcbba1fc9272fb6a4aef3b2ccb181d99000d","fff5f0fee0d2fcbba1fc9272fb6a4aef3b2ccb181d99000d","fff5f0fee0d2fcbba1fc9272fb6a4aef3b2ccb181da50f1567000d").map(d),of=m(tf),pf=new Array(3).concat("fee6cefdae6be6550d","feeddefdbe85fd8d3cd94701","feeddefdbe85fd8d3ce6550da63603","feeddefdd0a2fdae6bfd8d3ce6550da63603","feeddefdd0a2fdae6bfd8d3cf16913d948018c2d04","fff5ebfee6cefdd0a2fdae6bfd8d3cf16913d948018c2d04","fff5ebfee6cefdd0a2fdae6bfd8d3cf16913d94801a636037f2704").map(d),mf=m(pf);f.schemeAccent=c,f.schemeDark2=a,f.schemePaired=b,f.schemePastel1=r,f.schemePastel2=n,f.schemeSet1=t,f.schemeSet2=o,f.schemeSet3=p,f.interpolateBrBG=l,f.schemeBrBG=s,f.interpolatePRGn=u,f.schemePRGn=i,f.interpolatePiYG=y,f.schemePiYG=h,f.interpolatePuOr=w,f.schemePuOr=A,f.interpolateRdBu=G,f.schemeRdBu=B,f.interpolateRdGy=R,f.schemeRdGy=P,f.interpolateRdYlBu=O,f.schemeRdYlBu=Y,f.interpolateRdYlGn=S,f.schemeRdYlGn=g,f.interpolateSpectral=x,f.schemeSpectral=v,f.interpolateBuGn=_,f.schemeBuGn=j,f.interpolateBuPu=q,f.schemeBuPu=k,f.interpolateGnBu=M,f.schemeGnBu=D,f.interpolateOrRd=C,f.schemeOrRd=z,f.interpolatePuBuGn=F,f.schemePuBuGn=E,f.interpolatePuBu=I,f.schemePuBu=H,f.interpolatePuRd=K,f.schemePuRd=J,f.interpolateRdPu=N,f.schemeRdPu=L,f.interpolateYlGnBu=T,f.schemeYlGnBu=Q,f.interpolateYlGn=V,f.schemeYlGn=U,f.interpolateYlOrBr=X,f.schemeYlOrBr=W,f.interpolateYlOrRd=$,f.schemeYlOrRd=Z,f.interpolateBlues=ef,f.schemeBlues=ff,f.interpolateGreens=cf,f.schemeGreens=df,f.interpolateGreys=bf,f.schemeGreys=af,f.interpolatePurples=nf,f.schemePurples=rf,f.interpolateReds=of,f.schemeReds=tf,f.interpolateOranges=mf,f.schemeOranges=pf,Object.defineProperty(f,"__esModule",{value:!0})});
/*! @source http://purl.eligrey.com/github/FileSaver.js/blob/master/FileSaver.js */
var saveAs=saveAs||function(e){"use strict";if("undefined"==typeof navigator||!/MSIE [1-9]\./.test(navigator.userAgent)){var t=e.document,n=function(){return e.URL||e.webkitURL||e},o=t.createElementNS("http://www.w3.org/1999/xhtml","a"),r="download"in o,i=function(e){var t=new MouseEvent("click");e.dispatchEvent(t)},a=/Version\/[\d\.]+.*Safari/.test(navigator.userAgent),c=e.webkitRequestFileSystem,d=e.requestFileSystem||c||e.mozRequestFileSystem,u=function(t){(e.setImmediate||e.setTimeout)(function(){throw t},0)},s="application/octet-stream",f=0,l=4e4,v=function(e){var t=function(){"string"==typeof e?n().revokeObjectURL(e):e.remove()};setTimeout(t,l)},p=function(e,t,n){t=[].concat(t);for(var o=t.length;o--;){var r=e["on"+t[o]];if("function"==typeof r)try{r.call(e,n||e)}catch(i){u(i)}}},w=function(e){return/^\s*(?:text\/\S*|application\/xml|\S*\/\S*\+xml)\s*;.*charset\s*=\s*utf-8/i.test(e.type)?new Blob(["\uFEFF",e],{type:e.type}):e},y=function(t,u,l){l||(t=w(t));var y,m,S,h=this,R=t.type,O=!1,g=function(){p(h,"writestart progress write writeend".split(" "))},b=function(){if(m&&a&&"undefined"!=typeof FileReader){var o=new FileReader;return o.onloadend=function(){var e=o.result;m.location.href="data:attachment/file"+e.slice(e.search(/[,;]/)),h.readyState=h.DONE,g()},o.readAsDataURL(t),void(h.readyState=h.INIT)}if((O||!y)&&(y=n().createObjectURL(t)),m)m.location.href=y;else{var r=e.open(y,"_blank");void 0===r&&a&&(e.location.href=y)}h.readyState=h.DONE,g(),v(y)},E=function(e){return function(){return h.readyState!==h.DONE?e.apply(this,arguments):void 0}},N={create:!0,exclusive:!1};return h.readyState=h.INIT,u||(u="download"),r?(y=n().createObjectURL(t),void setTimeout(function(){o.href=y,o.download=u,i(o),g(),v(y),h.readyState=h.DONE})):(e.chrome&&R&&R!==s&&(S=t.slice||t.webkitSlice,t=S.call(t,0,t.size,s),O=!0),c&&"download"!==u&&(u+=".download"),(R===s||c)&&(m=e),d?(f+=t.size,void d(e.TEMPORARY,f,E(function(e){e.root.getDirectory("saved",N,E(function(e){var n=function(){e.getFile(u,N,E(function(e){e.createWriter(E(function(n){n.onwriteend=function(t){m.location.href=e.toURL(),h.readyState=h.DONE,p(h,"writeend",t),v(e)},n.onerror=function(){var e=n.error;e.code!==e.ABORT_ERR&&b()},"writestart progress write abort".split(" ").forEach(function(e){n["on"+e]=h["on"+e]}),n.write(t),h.abort=function(){n.abort(),h.readyState=h.DONE},h.readyState=h.WRITING}),b)}),b)};e.getFile(u,{create:!1},E(function(e){e.remove(),n()}),E(function(e){e.code===e.NOT_FOUND_ERR?n():b()}))}),b)}),b)):void b())},m=y.prototype,S=function(e,t,n){return new y(e,t,n)};return"undefined"!=typeof navigator&&navigator.msSaveOrOpenBlob?function(e,t,n){return n||(e=w(e)),navigator.msSaveOrOpenBlob(e,t||"download")}:(m.abort=function(){var e=this;e.readyState=e.DONE,p(e,"abort")},m.readyState=m.INIT=0,m.WRITING=1,m.DONE=2,m.error=m.onwritestart=m.onprogress=m.onwrite=m.onabort=m.onerror=m.onwriteend=null,S)}}("undefined"!=typeof self&&self||"undefined"!=typeof window&&window||this.content);"undefined"!=typeof module&&module.exports?module.exports.saveAs=saveAs:"undefined"!=typeof define&&null!==define&&null!==define.amd&&define([],function(){return saveAs});

!function(t,e){"object"==typeof exports&&"undefined"!=typeof module?e(exports):"function"==typeof define&&define.amd?define(["exports"],e):e(t.lc=t.lc||{})}(this,function(t){"use strict";function e(){var t={propList:[],add_property:function(e,r,a){t.propList.push(e);var n="get_"+e;return t[e]=function(e){return void 0===e?t[n]():("function"==typeof a&&(e=a(e)),t[n]="function"==typeof e?e:function(){return e},t.layers?t:t.chart?t.chart:t)},t[n]="function"==typeof r?r:function(){return r},t},wrapSetter:function(e,r){if("function"!=typeof r)throw"Error in 'wrapSetter': wrapper is not a function.";if(-1==t.propList.indexOf(e))throw"Error in 'wrapSetter': this object doesn't have "+e+" property";var a=t[e];return t[e]=function(){return 0==arguments.length?t["get_"+e]():r(a).apply(t,arguments)},t}};return t}function r(t){var e={};return function(){if("clear"!==arguments[0])return arguments in Object.keys(e)||0==arguments.length&&0!=Object.keys(e).length||(e[arguments]=t.apply(void 0,arguments)),e[arguments];e={}}}function a(t,e){if(t.length!=e.length)throw"Error in getEuclideanDistance: length of theinput vectors is not the same";for(var r=0,a=0;a<t.length;a++)r+=(t[a]-e[a])*(t[a]-e[a]);return Math.sqrt(r)}function n(t){var e,r,n,o=null,i=null,s=l(100),d=!1;t.svg.selectAll(".plotArea").append("rect").attr("class","clickPanel").attr("fill","transparent").lower();var c=t.updateSize;t.updateSize=function(){return c(),t.svg.selectAll(".clickPanel").attr("width",t.plotWidth()).attr("height",t.plotHeight()),t};var u=!1,p=function(e){var r=t.svg.select(".selection"),a=t.svg.select(".clickPanel").node().getBoundingClientRect(),o=[d3.max([d3.min([e.clientX-a.left,t.plotWidth()]),0]),d3.max([d3.min([e.clientY-a.top,t.plotHeight()]),0])];if(d)u||(u=!0,setTimeout(function(){u=!1},100),t.panMove(o));else{if(!r.empty()){r.attr("x",d3.min([o[0],n[0]])).attr("y",d3.min([n[1],o[1]])).attr("width",Math.abs(n[0]-o[0])).attr("height",Math.abs(n[1]-o[1]));var i=t.svg.select(".shadow");return i.empty()&&Math.abs((n[0]-o[0])*(n[1]-o[1]))>10&&(i=t.svg.selectAll(".plotArea").append("path").attr("class","shadow").attr("fill","#444").attr("opacity",.6)),void i.attr("d","M 0 0 h "+t.plotWidth()+" v "+t.plotHeight()+" h -"+t.plotWidth()+" v -"+t.plotHeight()+" M "+r.attr("x")+" "+r.attr("y")+" v "+r.attr("height")+" h "+r.attr("width")+" v -"+r.attr("height")+" h -"+r.attr("width"))}var l=[];t.container.selectAll("canvas").filter(function(){return d3.select(this).classed("active")}).each(function(){l.push(d3.select(this).attr("id"))}),l.length>0&&s.do(function(){var r=t.findElements(o,o);if(Array.isArray(r)){if(0==r.length)return void t.get_on_mouseout();t.container.selectAll(".inform").style("left",o[0]+10+t.paddings().left+"px").style("top",o[1]+10+t.paddings().top+"px").select(".value").html(function(){return t.get_informText(r[0][0],r[0][1])})}else{for(var a,n=Object.keys(r).length,i=!0;i&&n>0;)n--,a=Object.keys(r)[n],i=0==r[a].length;i||-1==l.indexOf(a)||d3.customEvent(e,t.get_layer(a).get_on_mouseover,this,[r[a][r[a].length-1]]),i&&t.get_on_mouseout&&t.get_layer(a).get_on_mouseout()}})}},f=function(n){var l=t.svg.select(".clickPanel").node().getBoundingClientRect(),s=[d3.max([d3.min([n.clientX-l.left,t.plotWidth()]),0]),d3.max([d3.min([n.clientY-l.top,t.plotHeight()]),0])];d3.event=n;var c=d3.event.shiftKey||t.selectMode();if(t.container.selectAll(".inform").classed("blocked",!1),!t.svg.select("rect.selection").empty())var u=+t.svg.selectAll("rect.selection").attr("x"),p=+t.svg.selectAll("rect.selection").attr("y"),f=[u,p],g=[u+ +t.svg.selectAll("rect.selection").attr("width"),p+ +t.svg.selectAll("rect.selection").attr("height")];var h=t.svg.selectAll(".plotArea");return t.svg.selectAll("rect.selection").remove(),t.svg.select(".shadow").remove(),i&&a(e,d3.mouse(document.body))<5?(window.clearTimeout(i),i=null,o&&a(r,d3.mouse(document.body))<5?(window.clearTimeout(o),o=null,h.on("dbl_click").apply(h,[c])):o=window.setTimeout((d3.event,function(){if(h.on("click").call(h,s,c),o=null,d)return d=!1,t.panMove(s),t.container.selectAll(".inform").classed("blocked",!1),void(t.transitionOff=!1)}),300),void(r=d3.mouse(document.body))):d?(d=!1,t.panMove(s),t.container.selectAll(".inform").classed("blocked",!1),void(t.transitionOff=!1)):(f&&(c?t.mark(t.findElements(f,g)):t.zoom(f,g)),document.onmousemove=null,void(document.onmouseup=null))};return t.svg.selectAll(".plotArea").on("mousedown",function(){t.container.selectAll(".hint").remove();var r=d3.mouse(this);navigator.userAgent.toLowerCase().indexOf("firefox")>-1&&(r[0]-=t.paddings().left,r[1]-=t.paddings().top),e=d3.mouse(document.body),n=r.slice(),i=window.setTimeout(function(){i=null},1e3),t.pan("mode")||t.svg.selectAll(".plotArea").append("rect").attr("class","selection").attr("x",r[0]).attr("y",r[1]).attr("width",1).attr("height",1),t.pan("mode")&&(d=!0,t.pan("down",n),t.transitionOff=!0),t.container.selectAll(".inform").classed("blocked",!0),document.addEventListener("mousemove",p,!1),document.addEventListener("mouseup",f,!1)},!0).on("dbl_click",function(e){e?t.mark("__clear__"):t.resetDomain()},!0).on("click",function(e,r){if(d3.event.stopPropagation(),void 0!==e){t.clickPosition&&t.clickPosition(e);var a=t.findElements(e,e);if(0!=a.length)if(r)t.mark(a);else{var n=[];if(t.container.selectAll("canvas").filter(function(){return d3.select(this).classed("active")}).each(function(){n.push(d3.select(this).attr("id"))}),n.length>0){if(Array.isArray(a))return void t.get_on_click(a[0][0],a[0][1]);for(var o,i=Object.keys(a).length,l=!0;l&&i>0;)if(i--,0!=a[o=Object.keys(a)[i]].length)if(t.clickSingle())l=!1,d3.customEvent(d3.event,t.get_layer(o).get_on_click,this,[a[o][0]]);else for(var s=0;s<a[o].length;s++)d3.customEvent(d3.event,t.get_layer(o).get_on_click,this,[a[o][s]])}else{var d=t.get_elements(a),c=d.filter(function(e){return e==t.container.selectAll(".inform").datum()});c.empty()||(d=c),t.clickSingle()&&(d=d3.select(d.node())),d.each(function(t){d3.select(this).on("click").call(this,t)})}}}},!0),t.container.node().addEventListener("mousemove",p,!1),t}function o(t,e){return 3.14*Math.pow(28.2*e/{Circle:30,Cross:35,Diamond:46,Square:36,Star:47,Triangle:44,Wye:37}[t],2)}function i(t){return t.replace(/[\[\]\/\{\}\(\)\*\+\?\.\\\^\$\|]/g,"\\$&")}function l(t){var e={interval:t,prev_time:-1/0,timer:null,do:function(){for(var r=arguments[0],a=[],n=1;n<arguments.length;n++)a.push(arguments[n]);e.timer&&e.timer.stop(),d3.now()-e.prev_time>=t?(r.call(this,a),e.prev_time=d3.now()):e.timer=d3.timeout(r,1.5*t)}};return e}function s(t,e){return"number_nonneg"==t?function(t){if("function"==typeof t)return t;if(d(+t)||(t=+t),"number"==typeof t){if(t>=0)return t;throw"Error in 'typeCheck' for property '"+e+"': negative values are not allowed."}throw"Error in 'typeCheck' for property '"+e+"': numeric value is required."}:"array"==t?function(t){if("function"==typeof t)return t;if(Array.isArray(t))return t;if("object"==typeof t){var r=[];for(el in t){if("string"!=typeof t[el]&&"number"!=typeof t[el])throw"Error in 'typeCheck' for property '"+e+"': the value is not an array.";r.push(t[el])}return r}if("number"==typeof t||"string"==typeof t)return[t];throw"Error in 'typeCheck' for property '"+e+"': the value is not an array."}:"array_fun"==t?function(t){return t&&"object"==typeof t?function(e){return t[e]}:t}:(t="matrix_fun")?function(t){if("function"==typeof t)return t;if(t&&"object"==typeof t){var r=Object.keys(t);return"object"==typeof t[r[0]]?function(e,r){return t[e][r]}:function(e){return t[e]}}throw"Error in 'typeCheck' for property '"+e+"': the value is not an array or an object."}:void 0}function d(t){return null==t||"NA"==t||"NaN"==t||"NAN"==t||"na"==t||"Infinity"==t||"Inf"==t}function c(t){var r=e().add_property("nelements",void 0,s("number_nonneg","nelements")).add_property("elementIds",void 0,s("array","elementIds")).add_property("label",function(t){return t},s("array_fun","label")).add_property("on_mouseover",function(){}).add_property("on_mouseout",function(){}).add_property("mode","svg").add_property("on_click",function(){}).add_property("layerDomainX",void 0,s("array","layerDomainX")).add_property("layerDomainY",void 0,s("array","layerDomainY")).add_property("contScaleX",function(){var t=r.layerDomainX();return void 0===t||2==t.length&&"number"==typeof t[0]&&"number"==typeof t[1]}).add_property("contScaleY",function(){var t=r.layerDomainY();return void 0===t||2==t.length&&"number"==typeof t[0]&&"number"==typeof t[1]}).add_property("colour",function(t){return r.colourScale(r.get_colourValue(t))},s("array_fun","colour")).add_property("addColourScaleToLegend",!0).add_property("palette",void 0,s("array","palette")).add_property("colourDomain",s("array","colourDomain")).add_property("colourValue",void 0,s("array_fun","colourValue")).add_property("colourLegendTitle","").add_property("opacity",1,s("array_fun","opacity")).add_property("dresser",function(){}).add_property("on_marked",function(){r.chart.on_marked()}).add_property("informText",function(t){return"<b>ID:</b> "+r.get_label(t)},s("array_fun","opacity"));r.propList=r.propList.concat(["updateElementStyle","updateElements","updateElementPosition"]),r.id=t,r.marked=[],r.wrapSetter("nelements",function(t){return function(){return r.get_elementIds=function(){return d3.range(t())},t.apply(r,arguments)}}),r.wrapSetter("elementIds",function(t){return function(){return r.get_nelements=function(){return t().length},t.apply(r,arguments)}}),r.wrapSetter("colour",function(t){return function(){return r.addColourScaleToLegend(!1),t.apply(r,arguments)}}),r.colourDomain(function(){var t=r.elementIds().map(t=>r.get_colourValue(t));if(0!=t.length){if(1==t.length)return void 0===t[0]?[]:t;var e,a=-1;if("number"==typeof t[1]&&"number"==typeof t[2])for(e=t[1]>t[0]?[t[0],t[1]]:[t[1],t[0]],a=2;a<t.length&&("number"==typeof t[a]||lc.isNaN(t[a]));)e[0]>t[a]&&(e[0]=t[a]),e[1]<t[a]&&(e[1]=t[a]),a++;if(a<t.length){e=[];for(a=0;a<t.length;a++)-1==e.indexOf(t[a])&&void 0!==t[a]&&e.push(t[a])}return e}}),r.colourScale=function(){return"black"},r.resetColourScale=function(){var t=r.colourDomain();if(void 0!==t&&0!=t.length){var e;if(r.chart.globalColourScale())for(var a in t=r.chart.globalColourDomain(r.id,t),r.chart.layers)e=r.chart.get_layer(a),a==r.id||e.globalScaleUpdate||(e.globalScaleUpdate=!0,e.updateElementStyle(),e.globalScaleUpdate=!1);var n=!0;t=t.filter(t=>!lc.isNaN(t));for(var o=0;o<t.length;o++)n=n&&"number"==typeof t[o];n&&t.sort(function(t,e){return t-e});var i=r.palette();if(n)if(t=t.filter(function(t){return"Infinity"!=t}),void 0==i&&(i=d3.interpolateSpectral?d3.interpolateSpectral:["red","yellow","green","blue"]),Array.isArray(i)){if((i=i.map(t=>/^#[0-9a-fA-F]{8,8}$/.test(t)?t.substring(0,7):t)).length!=t.length&&(t=[d3.min(t),d3.max(t)]),1==i.length&&i.push(i[0]),i.length>t.length){var l=[];for(o=0;o<i.length;o++)l.push(t[0]+o*(t[1]-t[0])/(i.length-1));t=l}r.colourValueScale=d3.scaleLinear().domain(t).range(i),r.colourScale=function(t){return r.colourValueScale(t)}}else{t=[d3.min(t),d3.max(t)];var s=[0,1];i.domain&&(s=i.domain()),r.colourValueScale=d3.scaleLinear().domain(t).range(s),r.colourScale=function(t){return null===t||void 0===t?"#999":i(r.colourValueScale(t))}}else if(void 0===i&&(i=d3.schemeCategory10),i.length){var d=i.length;for(o=0;o<t.length-d;o++)i.push(i[o%d]);r.colourValueScale=d3.scaleOrdinal().domain(t).range(i).unknown("black"),r.colourScale=function(t){return r.colourValueScale(t)}}else{s=[0,1];i.domain&&(s=i.domain()),r.colourValueScale=d3.scalePoint().domain(t).range(s),r.colourScale=function(t){return null===t||void 0===t?"#999":i(r.colourValueScale(t))}}if(r.colourScale.domain=r.colourValueScale.domain,r.chart.showLegend()&&r.addColourScaleToLegend()){var c={};c[r.id+"_colour"]=r.colourLegendTitle(),r.chart.legend.set_title(c),r.addLegendBlock(r.colourScale,"colour",r.id+"_colour")}}},r.legendBlocks=[],r.addLegendBlock=function(t,e,a){return r.chart.legend.add_block(t,e,a,r),-1==r.legendBlocks.indexOf(a)&&r.legendBlocks.push(a),r},r.update=function(){return r.updateStarted=!0,r.updateElements(),r.updateElementStyle(),r.updateElementPosition(),r.updateStarted=!1,r};var a=function(){return"default"==r.mode()?r.nelements()>2500?"canvas":"svg":r.mode()};return r.put_static_content=function(){r.g=r.chart.svg.selectAll(".plotArea").append("g").attr("class","chart_g").attr("id",r.id),r.canvas=r.chart.container.append("canvas").style("position","absolute").style("z-index",-5).attr("id",r.id),"svg"==a()?r.g.classed("active",!0):r.canvas.classed("active",!0)},r.updateElements=function(){},r.updateElementStyle=function(){return r.resetColourScale(),r.get_dresser(r.g.selectAll(".data_element")),0!=r.get_marked().length&&r.colourMarked(),r},r.updateElementPosition=function(){},r.findElements=function(){return[]},r.get_position=function(t){},r.on_mouseover(function(t){var e=r.chart.container.node().getBoundingClientRect(),a=[d3.max([d3.min([d3.event.clientX-e.left,r.chart.plotWidth()]),0]),d3.max([d3.min([d3.event.clientY-e.top,r.chart.plotHeight()]),0])];this&&this.prev_time||d3.select(this).attr("fill",function(t){return d3.rgb(r.get_colour(t)).darker(.5)}).classed("hover",!0),r.chart.container.selectAll(".inform").data([t]).style("left",a[0]+10+"px").style("top",a[1]+10+"px").select(".value").html(r.get_informText(t)),r.chart.container.selectAll(".inform").classed("hidden",!1)}),r.on_mouseout(function(t){if(!this.propList){var e=r.get_marked().length>0;d3.select(this).attr("fill",function(t){return e^d3.select(this).classed("marked")?"#aaa":r.get_colour(t)}).classed("hover",!1)}r.chart.container.selectAll(".inform").classed("hidden",!0)}),r.get_marked=function(){return"svg"==a()?r.g.selectAll(".data_element").filter(function(){return d3.select(this).classed("marked")}).data():r.marked},r.checkMode=function(){if("svg"==a()&&r.canvas.classed("active"))return r.canvas.classed("active",!1),r.g.classed("active",!0),r.canvas.node().getContext("2d").clearRect(0,0,r.chart.plotWidth(),r.chart.plotHeight()),!!r.updateStarted||(r.update(),!1);if("canvas"==a()&&r.g.classed("active"))for(r.canvas.classed("active",!0),r.g.classed("active",!1);r.g.node().firstChild;)r.g.node().removeChild(r.g.node().firstChild);return!0},r.mark=function(t,e){if("svg"==a()){if("__clear__"==t)return r.g.selectAll(".data_element.marked").classed("marked",!1),r.g.selectAll(".data_element").attr("opacity",1),e||r.on_marked(),r.colourMarked(),r.chart;if(!t.empty){var n={};n[r.id]=t,t=r.chart.get_elements(n)}if(t.empty())return r.chart;1==t.size()?t.classed("marked",!t.classed("marked")):t.classed("marked",!0),r.colourMarked()}else{if("__clear__"==t)r.marked=[];else for(var o,i=r.marked.map(function(t){return Array.isArray(t)?t.join("_"):t}),l=0;l<t.length;l++)-1==(o=t[l].join?i.indexOf(t[l].join("_")):i.indexOf(t[l]))?r.marked.push(t[l]):1==t.length&&r.marked.splice(o,1);r.updateCanvas()}e||r.on_marked()},r.colourMarked=function(){if("svg"==a()){var t={};t[r.id]=r.get_marked(),(t=r.chart.get_elements(t)).empty()?r.g.selectAll(".data_element").attr("fill",function(t){return r.get_colour(t)}):r.g.selectAll(".data_element").attr("fill",function(t){return d3.select(this).classed("marked")?r.get_colour(t):"#aaa"})}},r}function u(t){var r=e().add_property("x",function(){return t.width()-r.buttonSize()-5}).add_property("y",function(){return.05*t.paddings().top}).add_property("orientation","horizontal").add_property("height",function(){if("vertical"==r.orientation())return Math.floor(r.y()/r.buttonSize())*r.buttonSize()}).add_property("width",function(){if("horizontal"==r.orientation())return Math.floor(r.x()/r.buttonSize())*r.buttonSize()}).add_property("buttonSize",30);r.chart=t;var a,n={},o=[];function i(t,e,a){var i,l;if(a){for(l=d3.min([e,a]),i=1;Math.floor(e/l)*i<t;)i++,l=d3.min([a/i,l]);r.buttonSize(l)}else l=r.buttonSize(),i=Math.ceil(e/l);n={width:r.width(),height:r.height(),rowLength:d3.min([Math.floor(e/l),o.length])},"horizontal"==r.orientation()?n.width=l*n.rowLength:n.height=l*n.rowLength}function l(e){e&&t.container.append("div").attr("class","hint").style("left",r.chart.width()-105+"px").style("top",r.y()+r.g.node().getBBox().height+"px").text(e)}return r.wrapSetter("orientation",function(t){return function(){if(-1==["vertical","horizontal"].indexOf(t()))throw"Error in 'panel.orientation': value "+t()+" is not allowed for this property. Possible values are 'horizontal' or 'vertical'.";return t.apply(r,arguments)}}),r.put_static_content=function(){r.g=r.chart.svg.append("g").attr("class","panel_g"),function(){var t=r.g.append("def");a=r.buttonSize()-10;var e=t.append("g").attr("id","toggleOff");e.append("rect").attr("stroke-width",2).attr("width",a).attr("height",a).attr("fill","#aaa").attr("stroke","#444"),e.append("path").attr("d","M "+a/2+" "+Math.floor(a/3)+" L "+Math.ceil(2*a/3)+" "+Math.ceil(2*a/3)+" H "+Math.floor(a/3)+" L "+a/2+" "+Math.floor(a/3)).attr("fill","#444").attr("stroke-width",0),(e=t.append("g").attr("id","toggleOnHor")).append("rect").attr("stroke-width",2).attr("width",a).attr("height",a).attr("fill","#aaa").attr("stroke","#444"),e.append("path").attr("d","M "+Math.floor(a/3)+" "+a/2+" L "+Math.ceil(2*a/3)+" "+Math.floor(a/3)+" V "+Math.floor(2*a/3)+" L "+Math.floor(a/3)+" "+a/2).attr("fill","#444").attr("stroke-width",0),(e=t.append("g").attr("id","toggleOnVer")).append("rect").attr("stroke-width",2).attr("width",a).attr("height",a).attr("fill","#aaa").attr("stroke","#444"),e.append("path").attr("d","M "+a/2+" "+Math.ceil(2*a/3)+" L "+Math.floor(a/3)+" "+Math.floor(a/3)+" H "+Math.floor(2*a/3)+" L "+a/2+" "+Math.ceil(2*a/3)).attr("fill","#444").attr("stroke-width",0),(e=t.append("g").attr("id","save")).append("rect").attr("stroke-width",0).attr("width",a).attr("height",a).attr("fill","#444").attr("rx",a/10).attr("ry",a/10),e.append("path").attr("d","M "+Math.floor(4*a/5)+" 0 H "+a+" V "+Math.ceil(a/5)+" L "+Math.floor(4*a/5)+" 0").attr("fill","#fff").attr("stroke-width",0),e.append("rect").attr("x",Math.floor(a/3)).attr("height",Math.floor(a/3)).attr("width",Math.ceil(2*a/5)).attr("fill","#fff").attr("stroke-width",0),e.append("rect").attr("x",Math.floor(44*a/75)).attr("width",Math.ceil(2*a/25)).attr("height",Math.ceil(a/4)).attr("fill","#444").attr("stroke-width",0),e.append("rect").attr("x",Math.floor(a/4)).attr("width",Math.ceil(5*a/12)).attr("y",a/2).attr("height",a/2).attr("rx",a/10).attr("ry",a/10).attr("fill","#fff").attr("stroke-width",0),(e=t.append("g").attr("id","svg")).append("text").attr("font-size",1.5*a).attr("textLength",a).attr("lengthAdjust","spacingAndGlyphs").attr("fill","#444").attr("y",a).attr("font-weight",900).attr("font-family","Verdana").text("SVG"),(e=t.append("g").attr("id","selection")).append("rect").attr("fill-opacity",0).attr("width",Math.floor(2*a/3)).attr("height",Math.floor(2*a/3)).attr("x",Math.ceil(a/3)).attr("stroke","#444").attr("stroke-width",1).attr("stroke-dasharray",2),e.append("path").attr("fill","#444").attr("stroke-width",0).attr("stroke","#444").attr("d","M "+Math.ceil(a/3)+" "+Math.floor(2*a/3)+" L 0 "+Math.ceil(3*a/4)+" l "+a/12+" "+a/12+" L 0 "+9*a/10+" L "+a/10+" "+a+" L "+11*a/60+" "+11*a/12+" L "+Math.floor(a/4)+" "+a+" L "+Math.ceil(a/3)+" "+Math.floor(2*a/3)),e.append("circle").attr("cx",Math.floor(a/5)).attr("cy",Math.floor(a/5)).attr("r",2).attr("fill","#444"),e.append("circle").attr("cx",Math.floor(4*a/5)).attr("cy",Math.floor(4*a/5)).attr("r",2).attr("fill","#444"),e.append("circle").attr("cx",Math.floor(2*a/3)).attr("cy",Math.floor(a/3)).attr("r",3).attr("fill","#111"),(e=t.append("g").attr("id","zoomIn")).append("path").attr("fill","#444").attr("d","M "+2*a/5+" 0 h "+a/5+" v "+2*a/5+" h "+2*a/5+" v "+a/5+" h -"+2*a/5+" v "+2*a/5+" h -"+a/5+" v -"+2*a/5+" h -"+2*a/5+" v -"+a/5+" h "+2*a/5+" v -"+2*a/5),(e=t.append("g").attr("id","zoomOut")).append("rect").attr("y",3*a/8).attr("height",a/4).attr("width",a).attr("fill","#444"),(e=t.append("g").attr("id","home")).append("rect").attr("x",a/5).attr("y",2*a/5).attr("width",3*a/5).attr("height",3*a/5).attr("fill","#444"),e.append("rect").attr("x",2*a/5).attr("y",3*a/5).attr("width",a/5).attr("height",a/5).attr("fill","#fff"),e.append("path").attr("fill","#444").attr("d","M 0 "+2*a/5+" L "+a/2+" 0 L "+a+" "+2*a/5),e.append("path").attr("stroke","#fff").attr("stroke-width",1).attr("d","M 0 "+2*a/5+" L "+a+" "+2*a/5),(e=t.append("g").attr("id","pan")).append("path").attr("fill","#444").attr("d","M 0 "+a/2+" l "+a/5+" -"+a/5+" v "+a/10+" h "+a/5+" v -"+a/5+" h -"+a/10+" L "+a/2+" 0 l "+a/5+" "+a/5+" h -"+a/10+" v "+a/5+" h "+a/5+" v -"+a/10+" L "+a+" "+a/2+" l -"+a/5+" "+a/5+" v -"+a/10+" h -"+a/5+" v "+a/5+" h "+a/10+" L "+a/2+" "+a+" l -"+a/5+" -"+a/5+" h "+a/10+" v -"+a/5+" h -"+a/5+" v "+a/10+" L 0 "+a/2),(e=t.append("g").attr("id","fitSelected")).append("rect").attr("x",a/5).attr("y",a/5).attr("width",3*a/5).attr("height",3*a/5).attr("fill","#fff").attr("stroke","#444").attr("stroke-width",1).attr("stroke-dasharray",2),e.append("circle").attr("cx",a/5-3).attr("cy",a/2).attr("r",2).attr("fill","#444"),e.append("circle").attr("cx",3*a/5).attr("cy",a/5-3).attr("r",2).attr("fill","#444"),e.append("circle").attr("cx",4*a/5+3).attr("cy",2*a/5).attr("r",2).attr("fill","#444"),e.append("circle").attr("cx",2*a/5).attr("cy",4*a/5+3).attr("r",2).attr("fill","#444"),e.append("circle").attr("cx",4*a/5-3).attr("cy",a/5+3).attr("r",3).attr("fill","#111"),e.append("circle").attr("cx",a/5+3).attr("cy",4*a/5-3).attr("r",3).attr("fill","#111"),(e=t.append("g").attr("id","clusterRows")).append("rect").attr("x",2*a/5).attr("width",3*a/5).attr("height",a).attr("fill","#444"),e.append("rect").attr("stroke","#444").attr("stroke-width",1).attr("fill","#fff").attr("y",a/5).attr("height",a/2).attr("width",2*a/5),e.append("rect").attr("stroke","#444").attr("stroke-width",1).attr("fill","#fff").attr("y",a/10).attr("x",a/5).attr("height",a/5).attr("width",a/5),e.append("rect").attr("stroke","#444").attr("stroke-width",1).attr("fill","#fff").attr("x",a/10).attr("y",a/2).attr("height",3*a/10).attr("width",3*a/10),e.append("rect").attr("stroke","#444").attr("stroke-width",1).attr("fill","#fff").attr("x",a/5).attr("y",7*a/10).attr("height",a/5).attr("width",a/5),e.append("path").attr("stroke","#fff").attr("stroke-width",1).attr("d","M "+2*a/5+" "+a/5+" L "+a+" "+a/5),e.append("path").attr("stroke","#fff").attr("stroke-width",1).attr("d","M "+2*a/5+" "+2*a/5+" L "+a+" "+2*a/5),e.append("path").attr("stroke","#fff").attr("stroke-width",1).attr("d","M "+2*a/5+" "+3*a/5+" L "+a+" "+3*a/5),e.append("path").attr("stroke","#fff").attr("stroke-width",1).attr("d","M "+2*a/5+" "+4*a/5+" L "+a+" "+4*a/5),(e=t.append("g").attr("id","clusterCols")).append("rect").attr("y",2*a/5).attr("height",3*a/5).attr("width",a).attr("fill","#444"),e.append("rect").attr("stroke","#444").attr("stroke-width",1).attr("fill","#fff").attr("x",a/5).attr("width",a/2).attr("height",2*a/5),e.append("rect").attr("stroke","#444").attr("stroke-width",1).attr("fill","#fff").attr("x",a/10).attr("y",a/5).attr("width",a/5).attr("height",a/5),e.append("rect").attr("stroke","#444").attr("stroke-width",1).attr("fill","#fff").attr("y",a/10).attr("x",a/2).attr("width",3*a/10).attr("height",3*a/10),e.append("rect").attr("stroke","#444").attr("stroke-width",1).attr("fill","#fff").attr("y",a/5).attr("x",7*a/10).attr("width",a/5).attr("height",a/5),e.append("path").attr("stroke","#fff").attr("stroke-width",1).attr("d","M "+a/5+" "+2*a/5+" L "+a/5+" "+a),e.append("path").attr("stroke","#fff").attr("stroke-width",1).attr("d","M "+2*a/5+" "+2*a/5+" L "+2*a/5+" "+a),e.append("path").attr("stroke","#fff").attr("stroke-width",1).attr("d","M "+3*a/5+" "+2*a/5+" L "+3*a/5+" "+a),e.append("path").attr("stroke","#fff").attr("stroke-width",1).attr("d","M "+4*a/5+" "+2*a/5+" L "+4*a/5+" "+a),(e=t.append("g").attr("id","restoreOrder")).append("text").attr("font-size",1.5*a).attr("textLength",a).attr("lengthAdjust","spacingAndGlyphs").attr("fill","#444").attr("y",a).attr("font-weight",900).attr("font-family","Verdana").text("123"),t.selectAll("rect").attr("transform","translate(5, 5)"),t.selectAll("path").attr("transform","translate(5, 5)"),t.selectAll("text").attr("transform","translate(5, 5)"),t.selectAll("circle").attr("transform","translate(5, 5)"),t.selectAll("g").append("rect").attr("fill","transparent").attr("width",r.buttonSize()).attr("height",r.buttonSize()).lower()}(),r.g.append("use").attr("xlink:href","#toggleOff").attr("id","toggle").attr("opacity",.7).attr("title","Click to show instrument panel").on("mouseover",function(){d3.select(this).attr("opacity",1)}).on("mouseout",function(){d3.select(this).attr("opacity",.7)}).on("click",r.show).append("title").text("Click to show instrument panel"),r.g.append("g").attr("id","buttonPanel").attr("class","hidden")},r.updateSize=function(){"horizontal"==r.orientation()?(i(o.length,r.width(),r.height()),r.g.selectAll(".button").attr("transform",function(t,e){return"translate("+e%n.rowLength*r.buttonSize()+", "+Math.floor(e/n.rowLength)*r.buttonSize()+") scale("+(r.buttonSize()-10)/a+")"})):(i(o.length,r.height(),r.width()),r.g.selectAll(".button").attr("transform",function(t,e){return"translate("+Math.floor(e/n.rowLength)*r.buttonSize()+", "+(e%n.rowLength+1)*r.buttonSize()+") scale("+(r.buttonSize()-10)/a+")"})),"vertical"==r.orientation()?(r.g.attr("transform","translate("+r.x()+", "+r.y()+")"),r.g.select("#toggle").attr("transform","translate(0, 0)"),r.g.select("#buttonPanel").attr("transform","translate(0, "+r.buttonSize()+")")):(r.g.attr("transform","translate("+(r.x()-n.width)+", "+r.y()+")"),r.g.select("#toggle").attr("transform","translate("+n.width+", 0)"),r.g.select("#buttonPanel").attr("transform","translate(0, 0)"))},r.add_button=function(t,e,a,n){var i=!1;return o.push({name:t,icon:e,fun:function(t,e){i||(l(n),i=!0),a(t,e)}}),r.g.select("#buttonPanel").selectAll(".button").data(o,function(t){return t.name}).enter().append("use").attr("opacity",.6).attr("class","button").attr("id",function(t){return"b_"+t.icon.substr(1)}).attr("xlink:href",function(t){return t.icon}).on("click",function(t){t.fun(r.chart,d3.select(this))}).on("mouseover",function(){d3.select(this).attr("opacity",1)}).on("mouseout",function(){d3.select(this).attr("opacity",.6)}).append("title").text(function(t){return t.name}),r},r.show=function(){r.g.select("#toggle").attr("opacity",1).on("click",r.hide).on("mouseout",function(){}).select("title").text("Click to hide instrument panel"),"horizontal"==r.orientation()?r.g.select("#toggle").attr("xlink:href","#toggleOnHor"):r.g.select("#toggle").attr("xlink:href","#toggleOnVer"),r.g.select("#buttonPanel").classed("hidden",!1)},r.hide=function(){r.g.select("#toggle").attr("xlink:href","#toggleOff").attr("opacity",.7).on("click",r.show).on("mouseout",function(){d3.select(this).attr("opacity",.7)}).select("title").text("Click to show instrument panel"),r.g.select("#buttonPanel").classed("hidden",!0)},r}function p(t){var e=lc.base().add_property("width",100).add_property("height",function(){return t.height()}).add_property("sampleHeight",20).add_property("titles",{}).add_property("ncol").add_property("container",void 0,function(t){if("string"==typeof t&&(t=d3.select(t)),void 0===t.select)throw"Error in 'typeCheck' for property 'container': a D3 selection or a valid selector is required.";return t.empty()&&(t=d3.select("body")),t}),r={};e.chart=t,e.set_title=function(t){var r=e.titles();for(var a in t)r[a]=t[a];return e.titles(r),e},e.get_nblocks=function(){return Object.keys(r).length},e.get_blockIds=function(){return Object.keys(r)},e.add_block=function(t,n,o,i){if(e.chart.globalColourScale&&"colour"===n&&e.chart.globalColourScale()&&void 0===r[o]&&Object.keys(r).map(t=>r[t].type).indexOf("colour")>-1)return e.chart;var l={};if(l.scale="function"==typeof t?t:function(){return t},void 0!==i&&(l.layer=i),-1==["colour","size","symbol","dash"].indexOf(n))throw"Error in 'legend.add_block': "+n+" is not a suitable type of legend block. Please, use one of these: 'colour', 'size', 'symbol', 'dash'";return l.type=n,r[o]=l,e.container()&&a(),e.chart},e.updateScale=function(t,a){if(void 0===r[a])throw"Error in 'legend.updateScale': A block with ID "+a+" is not defined";return r[a].scale=t,e.updateBlock(a),e.chart};e.removeBlock=function(t){if(void 0===r[t])throw"Error in 'legend.remove': block with ID "+t+" doesn't exist";return void 0!==r[t].layer&&r[t].layer.legendBlocks.splice(r[t].layer.legendBlocks.indexOf(t),1),delete r[t],e.legendTable.select("#"+t).remove(),a(),e.chart},e.renameBlock=function(t,a){return r[a]=r[t],delete r[t],void 0!==r[a].layer&&r[a].layer.legendBlocks.splice(r[a].layer.legendBlocks.indexOf(t),1,a),e.legendTable&&(e.legendTable.select("#"+t).attr("id",a),e.updateBlock(a)),e.chart};var a=function(){var t,a,n=Object.keys(r).length;if(void 0===e.ncol()){var o,i=1+n;a=1;for(var l=2;l<=Math.ceil(Math.sqrt(n));l++)l+(o=Math.ceil(n/l))<=i&&(i=l+o,a=l);t=Math.ceil(n/a)}else t=e.ncol(),a=Math.ceil(n/t);for(var l in e.legendTable&&e.legendTable.remove(),e.legendTable=e.container().append("table").attr("class","legendTable linked-charts"),e.legendTable.selectAll("tr").data(d3.range(a)).enter().append("tr"),e.legendTable.selectAll("tr").selectAll("td").data(function(e){return d3.range(t).map(function(t){return[e,t]})}).enter().append("td").attr("id",function(e){try{return Object.keys(r)[e[0]*t+e[1]].replace(/[ .]/g,"_")}catch(t){return}}),r)e.updateBlock(l)};return e.updateBlock=function(t){if(void 0===r[t])throw"Error in 'legend.updateBlock': block with ID "+t+" is not defined";var a=function(t){var e,a;if("function"==typeof r[t].scale&&(e=r[t].scale()),(void 0===e||"function"!=typeof e&&void 0===e.splice)&&(e=r[t].scale),"function"!=typeof e){var n=!1,o=!1;if(1==e.length)throw"Error in 'legend.convertScale': range of the scale is not defined.";2==e[0].length&&"number"==typeof e[0][0]&&"number"==typeof e[0][1]&&(n=!0),"colour"==r[t].type&&e[0].length!=e[1].length&&(o=!0),2==e[1].length&&"number"==typeof e[0][0]&&"number"==typeof e[0][1]&&(o=!0),n&&o&&(a=d3.scaleLinear().domain(e[0]).range(e[1]),e.steps?a.steps=e.steps:a.steps=9),n&&!o&&((a=d3.scaleQuantize().domain(e[0]).range(e[1])).steps=e[1].length),!n&&o&&((a=d3.scalePoint().domain(e[0]).range(e[1])).steps=e[0].length),n||o||(e[0].length>e[1].length&&e[0].splice(e[1].length),e[1].length>e[0].length&&e[1].splice(e[0].length),(a=d3.scaleOrdinal().domain(e[0]).range(e[1])).steps=e[0].length),r[t].domain=e[0],void 0===a.domain&&(a.domain=r[t].domain)}else{if("function"!=typeof e)throw"Error in 'legend.convertScale': the type of scale argument is not suported. Scale should be an array or a function.";var i;if(void 0===(i="function"==typeof e.domain?e.domain():e.domain))throw"Error in 'legend.convertScale': the domain of the scale is not defined.";for(var l=0;l<i.length;)void 0===i[l]?i.splice(l,1):l++;if(r[t].domain=i,a=e,e.steps)a.steps=e.steps;else{var s=!0;for(l=0;s&&l<i.length;)s=s&&"number"==typeof i[l],l++;s&&(i=d3.extent(i)),2==i.length&&"number"==typeof i[0]?a.steps=9:a.steps=i.length}}return a}(t),n=e.legendTable.select("#"+t.replace(/[ .]/g,"_")),i=e.width()/e.legendTable.select("tr").selectAll("td").size(),l=a.steps,s=e.sampleHeight()*l,d=n.selectAll("svg");d.empty()&&(d=n.append("svg")),d.attr("width",i).attr("height",s);var c=d.select(".title");c.empty()&&(c=d.append("g").attr("class","title")).append("text").attr("class","plainText").attr("text-anchor","left");var u,p=d3.min([20,.2*i]),f=c.select("text").attr("font-size",p-3).attr("y",p-5).text(""==e.titles()[t]?"":e.titles()[t]||t).node().getBoundingClientRect().width;f>s&&c.select("text").attr("font-size",d3.max([10,(p-3)*s/f])),c.attr("transform","rotate(-90)translate(-"+s+", 0)"),u=r[t].domain.length==l?r[t].domain:d3.range(l).map(function(e){var a=r[t].domain.length-1;return r[t].domain[a]-e*(r[t].domain[a]-r[t].domain[0])/(l-1)});for(var g,h=[],m=0;m<u.length;m++)"number"==typeof u[m]&&(Math.abs(u[m])>=100?u[m]=u[m].toFixed(0):Math.abs(u[m])>=10?u[m]=u[m].toFixed(1):Math.abs(u[m]>=1||0==u[m])?u[m]=u[m].toFixed(2):(g=1-Math.floor(Math.log(Math.abs(u[m]))/Math.log(10)),u[m]=g>3?u[m].toExponential(2):u[m].toFixed(g))),h.push([u[m]]);var _=d.selectAll(".sample").data(h);if(_.exit().remove(),_.enter().append("g").attr("class","sample").merge(_).attr("transform",function(t,r){return"translate("+(p+1)+", "+r*e.sampleHeight()+")"}),"colour"==r[t].type){var y=d.selectAll(".sample").selectAll("rect").data(function(t){return t});y.enter().append("rect").merge(y).attr("width",p).attr("height",e.sampleHeight()).attr("fill",function(t){return a(t)})}if("symbol"==r[t].type){var v=d3.min([e.sampleHeight()/2,p/2]),x=d.selectAll(".sample").selectAll("path").data(function(t){return t});x.enter().append("path").merge(x).attr("d",function(t){return d3.symbol().type(d3["symbol"+a(t)]).size(o(a(t),v))()}).attr("transform","translate("+v+", "+v+")")}if("dash"==r[t].type){var b=d.selectAll(".sample").selectAll("line").data(function(t){return t});b.enter().append("line").style("stroke","black").merge(b).attr("x1",0).attr("x2",p).attr("y1",e.sampleHeight()/2).attr("y2",e.sampleHeight()/2).attr("stroke-dasharray",function(t){return a(t)})}d.selectAll(".sample").selectAll("g").data(function(t){return"number"==typeof t[0]?[t[0].toString()]:t}).enter().append("g").append("text").attr("class","plainText").attr("text-anchor","left").attr("dominant-baseline","middle"),d.selectAll(".sample").selectAll("g").attr("transform","translate("+(p+5)+", 0)");var w,k=d.selectAll(".sample").selectAll("g").selectAll("text").attr("font-size",e.sampleHeight()-5).attr("y",e.sampleHeight()/2).text(t=>t),S=[0];k.each(function(){S[0]<this.getBoundingClientRect().width&&(S[0]=this.getBoundingClientRect().width)}),S[0]>i-2*p-5&&(w=(i-2*p-5)/S[0],k.attr("font-size",d3.max([(e.sampleHeight()-5)*w,10])))},e.update=function(){a()},e}function f(){var t=e().add_property("width",500,s("number_nonneg","width")).add_property("height",500,s("number_nonneg","height")).add_property("paddings",{top:35,right:10,bottom:50,left:50},function(t){if("function"==typeof t)return t;if(void 0===t.top)throw"Error in 'typeCheck' for property 'paddings': top-padding is not defined";if(void 0===t.left)throw"Error in 'typeCheck' for property 'paddings': left-padding is not defined";if(void 0===t.bottom)throw"Error in 'typeCheck' for property 'paddings': bottom-padding is not defined";if(void 0===t.right)throw"Error in 'typeCheck' for property 'paddings': right-padding is not defined";return t}).add_property("title","").add_property("titleX",function(){return t.width()/2},s("number_nonneg","titleX")).add_property("titleY",function(){return d3.min([22,.9*t.paddings().top])},s("number_nonneg","titleY")).add_property("titleSize",function(){return d3.min([20,.8*t.paddings().top])},s("number_nonneg","titleSize")).add_property("transitionDuration",1e3,s("number_nonneg","transitionDuration")).add_property("on_marked",function(){}).add_property("showPanel",!0).add_property("clickSingle",!0).add_property("showLegend",!0).add_property("plotWidth",void 0,s("number_nonneg","plotWidth")).add_property("plotHeight",void 0,s("number_nonneg","plotHeight"));t.legend=p(t);var r=function(){return t.height()-(t.paddings().top+t.paddings().bottom)},a=function(){return t.width()-(t.paddings().right+t.paddings().left)};t.plotWidth(a).plotHeight(r),t.wrapSetter("width",function(e){return function(){return t.get_plotWidth=a,e.apply(t,arguments)}}),t.wrapSetter("height",function(e){return function(){return t.get_plotHeight=r,e.apply(t,arguments)}}),t.wrapSetter("plotWidth",function(e){return function(){return t.get_width=function(){return e()+t.paddings().left+t.paddings().right},e.apply(t,arguments)}}),t.wrapSetter("plotHeight",function(e){return function(){return t.get_height=function(){return e()+t.paddings().top+t.paddings().bottom},e.apply(t,arguments)}});var n=!1;t.selectMode=function(e){return void 0==e?n:(n=e=1==e,t)};var o={mode:!1,down:void 0};return t.pan=function(e,r){return void 0==r?o[e]:(o[e]=r,t)},t.set_paddings=function(e){return void 0===e.top&&(e.top=t.paddings().top),void 0===e.bottom&&(e.bottom=t.paddings().bottom),void 0===e.left&&(e.left=t.paddings().left),void 0===e.right&&(e.right=t.paddings().right),t.paddings(e),t},t.put_static_content=function(e){t.container=e.append("div").style("position","relative").attr("class","linked-charts"),t.container.node().ondragstart=function(){return!1},t.svg=t.container.append("table").attr("class","mainTable").append("tr").append("td").append("svg"),void 0===t.legend.container()&&t.legend.container(t.container.selectAll("tr").append("td").attr("id","legend")),t.container.append("div").attr("class","inform hidden").append("p").attr("class","value"),t.svg.append("text").attr("class","title plainText").attr("text-anchor","middle"),t.svg.append("svg").attr("class","plotArea"),t.showPanel()&&(void 0==t.panel&&(t.panel=u(t),t.panel.put_static_content()),t.panel.add_button("Save plot as png","#save",g),t.panel.add_button("Save plot as svg","#svg",h),t.panel.add_button("Select elements","#selection",m,"You can also select elements by pressing 'Shift'"),t.panel.add_button("Togle pan mode","#pan",_),t.panel.add_button("Reset scales","#home",function(t){t.resetDomain()},"You can also use double click to reset scales"),t.panel.add_button("Fit selected","#fitSelected",y))},t.mark=function(e,r){return"__clear__"==e?(t.svg.selectAll(".data_element.marked").classed("marked",!1),t.svg.selectAll(".data_element").attr("opacity",1).style("filter","none"),r||t.on_marked(),t):(e.empty||(e=t.get_elements(e)),t.svg.selectAll(".data_element.marked").empty()&&t.svg.selectAll(".data_element").attr("opacity",.3).style("filter","saturate(0.3) brightness(0.3)"),e.classed("switch",!0),e.size()<2&&e.filter(function(){return d3.select(this).classed("marked")}).classed("switch",!1).classed("marked",!1).attr("opacity",.3).style("filter","saturate(0.3) brightness(0.3)"),e.filter(function(){return d3.select(this).classed("switch")}).classed("marked",!0).classed("switch",!1).attr("opacity",1).style("filter","none"),t.svg.selectAll(".data_element.marked").empty()&&t.svg.selectAll(".data_element").attr("opacity",1).style("filter","none"),r||t.on_marked(),t)},t.place=function(e){if(void 0===e&&(e="body"),"string"==typeof e){var r=e;if(0==(e=d3.selectAll(r)).size())throw"Error in function 'place': DOM selection for string '"+r+"' did not find a node."}return t.transitionOff=!0,t.put_static_content(e),t.update(),t.transitionOff=!1,t},t.updateSize=function(){if(t.transitionDuration()>0&&!t.transitionOff){var e=d3.transition("size").duration(t.transitionDuration());t.svg.transition(e).attr("width",t.width()).attr("height",t.height()),t.svg.selectAll(".title").transition(e).attr("font-size",t.titleSize()).attr("x",t.titleX()).attr("y",t.titleY()),t.svg.selectAll(".plotArea").transition(e).attr("x",t.paddings().left).attr("y",t.paddings().top).attr("width",t.plotWidth()).attr("height",t.plotHeight())}else t.svg.attr("width",t.width()).attr("height",t.height()),t.svg.selectAll(".title").attr("font-size",t.titleSize()).attr("x",t.titleX()).attr("y",t.titleY()),t.svg.selectAll(".plotArea").attr("x",t.paddings().left).attr("y",t.paddings().top).attr("width",t.plotWidth()).attr("height",t.plotHeight());return t.showPanel()&&t.panel.updateSize(),t},t.updateTitle=function(){return t.svg.selectAll(".title").text(t.title()),t},t.get_elements=function(e){return e.splice||(e=[e]),e=e.map(function(t){return i(t).replace(/[ .]/g,"_")}),t.svg.selectAll("#p"+e.join(", #p"))},t.update=function(){return t.updateSize(),t.updateTitle(),t.showLegend()&&t.legend.get_nblocks()>0&&t.legend.update(),t},t}function g(t){var e=document.createElement("canvas");void 0===t.legend?(e.height=t.svg.attr("height"),e.width=t.svg.attr("width")):(e.height=d3.max([t.height(),t.legend.height()]),e.width=t.width()+t.legend.width());for(var r=e.getContext("2d"),a=t.container.selectAll("canvas").filter(function(){return d3.select(this).classed("active")}).nodes(),n=0;n<a.length;n++)r.drawImage(a[n],t.paddings().left,t.paddings().top);t.svg.selectAll("text").attr("fill","black");var o=t.svg.node().cloneNode(!0);for(n=0;n<o.childNodes.length;n++)"panel_g"==o.childNodes[n].classList[0]&&o.removeChild(o.childNodes[n]);!function(e,r,a,n){var o;if(void 0!==n){var i,l,s,d=0;o="<g>"+e.innerHTML+"</g>",n.container().selectAll("tr").each(function(t,e){i=0,s=[],d3.select(this).selectAll("td").each(function(t,e){l=d3.select(this).selectAll("svg"),o+="<g transform='translate("+(n.chart.width()+i)+", "+d+")'>"+l.node().innerHTML+"</g>",s.push(l.attr("height")),i+=+l.attr("width")}),d+=+d3.max(s)}),o="<svg xmlns='http://www.w3.org/2000/svg'>"+o+"</svg>"}else o=(new XMLSerializer).serializeToString(e);var c=new Image;c.onload=function(){r.drawImage(this,0,0),t.canvas&&t.canvas.classed("active")&&r.drawImage(t.canvas.node(),t.paddings().left,t.paddings().top),a()},c.src="data:image/svg+xml; charset=utf8, "+encodeURIComponent(o)}(o,e.getContext("2d"),function(){for(var t=e.toDataURL("image/png"),r=atob(t.substring("data:image/png;base64,".length)),a=new Uint8Array(r.length),n=0,o=r.length;n<o;++n)a[n]=r.charCodeAt(n);var i=new Blob([a.buffer],{type:"image/png"});saveAs(i,"export_"+Date.now()+".png")},t.legend)}function h(t){for(var e,r=t.svg.node().cloneNode(!0),a=0;a<r.childNodes.length;a++)"panel_g"==r.childNodes[a].classList[0]&&r.removeChild(r.childNodes[a]);if(!t.container.selectAll("canvas").filter(function(){return d3.select(this).classed("active")}).empty())return t.container.append("div").attr("class","hint").attr("id","errMessage").style("left",t.width()/3+"px").style("top",t.height()/3+"px").style("width",t.width()/3+"px").text("Chart in canvas mode cannot be saved as SVG."),void setTimeout(function(){t.container.select("#errMessage").remove()},2e3);if(void 0!==t.legend){var n,o,i,l=0;e="<g>"+r.innerHTML+"</g>",t.legend.container().selectAll("tr").each(function(){n=0,i=[],d3.select(this).selectAll("td").each(function(){o=d3.select(this).selectAll("svg"),e+="<g transform='translate("+(t.width()+n)+", "+l+")'>"+o.node().innerHTML+"</g>",i.push(o.attr("height")),n+=+o.attr("width")}),l+=+d3.max(i)}),e="<svg xmlns='http://www.w3.org/2000/svg'>"+e+"</svg>"}else e=t.svg.attr("xmlns","http://www.w3.org/2000/svg").node().parentNode.innerHTML;var s=new Blob([e],{type:"image/svg+xml"});saveAs(s,"export_"+Date.now()+".svg")}function m(t,e){if(e.classed("clicked"))e.classed("clicked",!1).attr("opacity",.6).on("mouseout",function(){d3.select(this).attr("opacity",.6)}),t.selectMode(!1);else{e.classed("clicked",!0).attr("opacity",1).on("mouseout",function(){}),t.selectMode(!0);var r=t.panel.g.selectAll("#b_pan");r.classed("clicked")&&r.on("click").call(r.node(),r.datum())}}function _(t,e){if(e.classed("clicked"))e.classed("clicked",!1).attr("opacity",.6).on("mouseout",function(){d3.select(this).attr("opacity",.6)}),t.pan("mode",!1);else{e.classed("clicked",!0).attr("opacity",1).on("mouseout",function(){}),t.pan("mode",!0);var r=t.panel.g.selectAll("#b_selection");r.classed("clicked")&&r.on("click").call(r.node(),r.datum())}}function y(t){var e=t.get_marked();t.layers&&(e=Object.keys(t.layers).map(t=>e[t].map(e=>[t,e])).flat()),0==e.length&&(e=t.layers?Object.keys(t.layers).map(e=>t.get_layer(e).elementIds().map(t=>[e,t])).flat():t.elementIds());var r={x:[],y:[]};e.map(function(e){var a=t.get_position(e);a&&(r.x.push(a[0]),r.y.push(a[1]))});var a=d3.extent(r.x),n=d3.extent(r.y);t.zoom([a[0],n[0]],[a[1],n[1]])}function v(){var t=M();t.add_property("singleScaleX",!0).add_property("singleScaleY",!0).add_property("domainX",void 0,s("array","domainX")).add_property("domainY",void 0,s("array","domainY")).add_property("aspectRatio",void 0,s("number_nonneg","aspectRatio")).add_property("axisTitleX","").add_property("axisTitleY","").add_property("axisTitlePos",{x:{across:"up",along:"end"},y:{across:"up",along:"end"}},function(t){var e={x:{across:"up",along:"end"},y:{across:"up",along:"end"}};if(t.x){if(!t.x.split)throw"Error in 'typeCheck' for property 'axisTitlePos': 'x' must be a string.";var r=t.x.split(" ").map(t=>t.toLowerCase());r.indexOf("down")>-1&&(e.x.across="down"),r.indexOf("middle")>-1&&(e.x.along="middle"),r.indexOf("start")>-1&&(e.x.along="start")}if(t.y){if(!t.y.split)throw"Error in 'typeCheck' for property 'axisTitlePos': 'y' must be a string.";var a=t.y.split(" ").map(t=>t.toLowerCase());a.indexOf("down")>-1&&(e.y.across="down"),a.indexOf("middle")>-1&&(e.y.along="middle"),a.indexOf("start")>-1&&(e.y.along="start")}return e}).add_property("ticksX",void 0).add_property("ticksY",void 0).add_property("ticksRotateX",0,function(t){if(isNaN(t))throw"Error in 'typeCheck' for property 'ticksRotateX': the value is not a number.";return t%91}).add_property("ticksRotateY",0,function(t){if(isNaN(t))throw"Error in 'typeCheck' for property 'ticksRotateY': the value is not a number.";return t%91}).add_property("logScaleX",!1).add_property("logScaleY",!1).add_property("on_clickPosition",function(t,e){}),t.axes={};var e=function(e){return function(){var r;if(t["get_singleScale"+e]()){var a=!0;for(var n in t.layers)a=a&&t.get_layer(n)["get_contScale"+e]();if(a)for(var n in t.layers)void 0!==t.get_layer(n)["get_layerDomain"+e]()&&(void 0===r?r=t.get_layer(n)["get_layerDomain"+e]():(r[0]=d3.min([r[0],t.get_layer(n)["get_layerDomain"+e]()[0]]),r[1]=d3.max([r[1],t.get_layer(n)["get_layerDomain"+e]()[1]])));else for(var n in t.layers)void 0!==t.get_layer(n)["get_layerDomain"+e]()&&(r=void 0===r?t.get_layer(n)["get_layerDomain"+e]():r.concat(t.get_layer(n)["get_layerDomain"+e]().filter(function(t){return r.indexOf(t)<0})))}return void 0===r&&(r=[0,1]),a&&(t["logScale"+e]()?(r[1]=2*r[1],r[0]=r[0]/2):(r[1]=r[1]+.03*(r[1]-r[0]),r[0]=r[0]-.03*(r[1]-r[0]))),r}};t.get_domainX=e("X"),t.get_domainY=e("Y"),t.domainX=function(e){return"reset"==e?(t.domainX(t.origDomainX),t):("function"==typeof e&&(t.get_domainX=e),e.splice&&(t.get_domainX=function(){return e}),t)},t.domainY=function(e){return"reset"==e?(t.domainY(t.origDomainY),t):("function"==typeof e&&(t.get_domainY=e),e.splice&&(t.get_domainY=function(){return e}),t)},t.get_position=function(e){return t.get_layer(e[0]).get_position(e[1])},t.zoom=function(e,r){if(e[0]!=r[0]&&e[1]!=r[1]){if(t.axes.scale_x.invert)t.domainX([t.axes.scale_x.invert(e[0]),t.axes.scale_x.invert(r[0])]);else{for(var a=[],n=t.get_domainX(),o=0;t.axes.scale_x(n[o])<=r[0];)t.axes.scale_x(n[o])>=e[0]&&a.push(n[o]),o++;a.length>0&&t.domainX(a)}if(t.axes.scale_y.invert)t.domainY([t.axes.scale_y.invert(r[1]),t.axes.scale_y.invert(e[1])]);else{var i=[],l=t.get_domainY();for(o=0;t.axes.scale_y(l[o])<=r[1];)t.axes.scale_y(l[o])>=e[1]&&i.push(l[o]),o++;i.length>0&&t.domainY(i)}t.updateAxes()}},t.resetDomain=function(){t.domainX("reset"),t.domainY("reset"),t.updateAxes()};var r=t.put_static_content;t.put_static_content=function(e){r(e);var a=t.svg.append("g").attr("class","axes_g");t.axes.x_g=a.append("g").attr("class","x axis"),t.axes.x_label=t.axes.x_g.append("text").attr("class","label"),t.axes.y_g=a.append("g").attr("class","y axis"),t.axes.y_label=t.axes.y_g.append("text").attr("class","label").attr("transform","rotate(-90)"),t.origDomainY=t.get_domainY,t.origDomainX=t.get_domainX,t.showPanel()&&(t.panel.add_button("Zoom in","#zoomIn",function(t){var e=t.axes.scale_x.domain(),r=t.axes.scale_y.domain();if(2==e.length&&"number"==typeof e[0]&&(e[1],1))t.domainX([(4*e[0]+e[1])/5,(e[0]+4*e[1])/5]);else{var a=Math.ceil(.1*e.length);e.splice(0,a),e.splice(e.length-a),e.length>0&&t.domainX(e)}if(2==r.length&&"number"==typeof r[0]&&(r[1],1))t.domainY([(4*r[0]+r[1])/5,(r[0]+4*r[1])/5]);else{a=Math.ceil(.1*r.length);r.splice(0,a),r.splice(r.length-a),r.length>0&&t.domainY(r)}t.updateAxes()},"Double click to return to original scales"),t.panel.add_button("Zoom out","#zoomOut",function(t){var e=t.axes.scale_x.domain(),r=t.axes.scale_y.domain();if(2==e.length&&"number"==typeof e[0]&&(e[1],1))t.domainX([(6*e[0]-e[1])/5,(-e[0]+6*e[1])/5]);else{for(var a=Math.ceil(.1*e.length),n=t.origDomainX(),o=n.indexOf(e[0]),i=n.indexOf(e[e.length-1]),l=o-1;l>=d3.max([0,o-a]);l--)e.unshift(n[l]);for(l=i+1;l<d3.min([n.length,i+a+1]);l++)e.push(n[l]);t.domainX(e)}if(2==r.length&&"number"==typeof r[0]&&(r[1],1))t.domainY([(6*r[0]-r[1])/5,(-r[0]+6*r[1])/5]);else{a=Math.ceil(.1*r.length);var s=t.origDomainY();for(o=s.indexOf(r[0]),i=s.indexOf(r[r.length-1]),l=s[o-1];l>=d3.max([0,o-a]);l--)r.unshift(s[l]);for(l=i+1;l<d3.min([s.length,i+a+1]);l++)r.push(s[l]);t.domainY(r)}t.updateAxes()},"Double click to return to original scales"))};var a=t.updateSize;t.updateSize=function(){if(a(),t.transitionDuration()>0&&!t.transitionOff){var e=d3.transition("size").duration(t.transitionDuration());t.svg.selectAll(".axes_g").transition(e).attr("transform","translate("+t.paddings().left+", "+t.paddings().top+")"),t.axes.x_g.transition(e).attr("transform","translate(0,"+t.get_plotHeight()+")")}else t.svg.selectAll(".axes_g").attr("transform","translate("+t.paddings().left+", "+t.paddings().top+")"),t.axes.x_g.attr("transform","translate(0,"+t.get_plotHeight()+")");return t.updateAxes(),t};var n=function(e){var r={tickValues:null,tickFormat:null},a=t["ticks"+e]();if(a){if(void 0===a.splice)throw"Error in 'get_ticks': new tick values and labels should be passed as an array";void 0===a[0].splice&&(a=[a]);for(var n=1;n<a.length;n++)if(a[0].length!=a[n].length)throw"Error in 'get_ticks': the amount of tick labels must be equal to the amount of tick values";if(r.tickValues=a[0],1==a.length)return r;for(var o=a.length>2,i=1;o&&i<a.length-1;){for(n=0;o&&n<a[i].length;)o=a[i][n]==a[i+1][n],n++;i++}o&&a.splice(2);var l,s=[];if(a.length>2)for(n=0;n<a[0].length;n++){l="";for(i=1;i<a.length;i++)l+="<tspan x = 0.5 dy = 1.1em",a.colour?l+=" fill = '"+a.colour[i-1]+"'>":l+=">",l+=a[i][n]+"</tspan>";s.push(l)}else s=a[1];r.tickFormat=function(t){return s[r.tickValues.indexOf(t)]}}return r};t.panMove=function(e){var r=t.axes.scale_x.domain(),a=t.axes.scale_y.domain();if(t.axes.scale_x.invert){var n=t.axes.scale_x.invert(e[0])-t.axes.scale_x.invert(t.pan("down")[0]);t.pan("down")[0]=e[0];var o=[r[0]-n,r[1]-n];t.logScaleX()&&(o[0]<=0&&(o[0]=r[0]),o[1]<=0&&(o[1]=r[1])),t.domainX(o)}else{n=e[0]-t.pan("down")[0];if((u=Math.floor(Math.abs(n)/t.axes.scale_x.step()*2))>0){t.pan("down")[0]+=Math.sign(n)*u*t.axes.scale_x.step()/2;var i=t.origDomainX(),l=i.indexOf(r[0]),s=i.indexOf(r[r.length-1]);n<0?(r.splice(0,u),r=r.concat(i.slice(s+1,d3.min([i.length,s+u+1])))):(r.splice(r.length-u),r=i.slice(d3.max([0,l-u]),l).concat(r)),r.length>0&&t.domainX(r)}}if(t.axes.scale_y.invert){var d=t.axes.scale_y.invert(e[1])-t.axes.scale_y.invert(t.pan("down")[1]);t.pan("down")[1]=e[1];var c=[a[0]-d,a[1]-d];t.logScaleY()&&(c[0]<=0&&(c[0]=a[0]),c[1]<=0&&(c[1]=a[1])),t.domainY(c)}else{var u;d=e[1]-t.pan("down")[1];if((u=Math.floor(Math.abs(d)/t.axes.scale_y.step()*2))>0){t.pan("down")[1]+=Math.sign(d)*u*t.axes.scale_y.step()/2;var p=t.origDomainY();l=p.indexOf(a[0]),s=p.indexOf(a[a.length-1]);d>0?(a.splice(0,u),a=a.concat(p.slice(s+1,d3.min([p.length,s+u+1])))):(a.splice(a.length-u),a=p.slice(d3.max([0,l-u]),l).concat(a)),a.length>0&&t.domainY(a)}}t.updateAxes()};var o=function(e){var r=t["get_domain"+e]();if(void 0===t.axes["scale_"+e.toLowerCase()]&&(t.axes["scale_"+e.toLowerCase()]={},t.axes["shiftScale"+e]=d3.scaleLinear().domain([0,1])),2==r.length&&"number"==typeof(r[0]+r[1])){var a=t["logScale"+e]();a&&a>0&&1!=a?(void 0===t.axes["scale_"+e.toLowerCase()].base&&(t.axes["scale_"+e.toLowerCase()]=d3.scaleLog()),t.axes["scale_"+e.toLowerCase()].base(a)):void 0!==t.axes["scale_"+e.toLowerCase()].clamp&&void 0==t.axes["scale_"+e.toLowerCase()].base||(t.axes["scale_"+e.toLowerCase()]=d3.scaleLinear()),t.axes["scale_"+e.toLowerCase()].nice()}else void 0===t.axes["scale_"+e.toLowerCase()].padding&&(t.axes["scale_"+e.toLowerCase()]=d3.scalePoint().padding(.3))},i=function(e){var r=t["get_domain"+e]();return 2==r.length&&"number"==typeof r[0]&&"number"==typeof r[1]&&r[0]==r[1]&&(r[0]=r[0]-.5,r[1]=r[1]+.5),r},l=function(e){var r=[0],a=t.axes["scale_"+e.toLowerCase()];a.padding&&r.push(a(a.domain()[1])-a(a.domain()[0])||a(a.domain()[0])),a.base&&r.push(a(t["logScale"+e]())-a(1)),a.clamp&&!a.base&&r.push(a(1)-a(0)),t.axes["shiftScale"+e].range(r)};return t.updateAxesTitles=function(){var e={x:{x:5,y:-5,anchor:"start",dy:0},y:{x:0,y:6,anchor:"end",dy:"1.8ex"}};if("middle"==t.axisTitlePos().x.along&&(e.x.x=t.plotWidth()/2,e.x.anchor="middle"),"end"==t.axisTitlePos().x.along&&(e.x.x=t.plotWidth(),e.x.anchor="end"),"down"==t.axisTitlePos().x.across&&(t.axes.x_label.text(""),e.x.y=t.axes.x_g.node().getBBox().height,e.x.dy="1.8ex"),"middle"==t.axisTitlePos().y.along&&(e.y.x=-t.plotHeight()/2,e.y.anchor="middle"),"start"==t.axisTitlePos().y.along&&(e.y.x=-t.plotHeight(),e.y.anchor="start"),"down"==t.axisTitlePos().y.across&&(t.axes.y_label.text(""),e.y.y=-t.axes.y_g.node().getBBox().width,e.y.dy=-6),t.axes.x_label.html(t.axisTitleX()),t.axes.y_label.html(t.axisTitleY()),t.transitionDuration()>0&&!t.transitionOff){var r=d3.transition("axes").duration(t.transitionDuration());t.axes.x_label.transition(r).attr("x",e.x.x).attr("y",e.x.y).attr("dy",e.x.dy).style("text-anchor",e.x.anchor),t.axes.y_label.transition(r).attr("x",e.y.x).attr("y",e.y.y).attr("dy",e.y.dy).style("text-anchor",e.y.anchor)}else t.axes.x_label.attr("x",e.x.x).attr("y",e.x.y).attr("dy",e.x.dy).style("text-anchor",e.x.anchor),t.axes.y_label.attr("x",e.y.x).attr("y",e.y.y).attr("dy",e.y.dy).style("text-anchor",e.y.anchor);return t},t.updateAxes=function(){o("X"),o("Y"),t.axes.scale_x.range([0,t.get_plotWidth()]),t.axes.scale_y.range([t.get_plotHeight(),0]),t.axes.scale_x.domain(i("X")),t.axes.scale_y.domain(i("Y")),t.aspectRatio()&&!t.axes.scale_x.base&&!t.axes.scale_y.base&&t.axes.scale_x.clamp&&t.axes.scale_y.clamp&&function(t,e,r){var a=(t.range()[1]-t.range()[0])/(t.domain()[1]-t.domain()[0]),n=(e.range()[1]-e.range()[0])/(e.domain()[1]-e.domain()[0]),o=Math.abs(a/n);if(o>r){var i=(t.domain()[1]-t.domain()[0])*(o/r-1)/2;t.domain([t.domain()[0]-i,t.domain()[1]+i])}else i=(e.domain()[1]-e.domain()[0])*(r/o-1)/2,e.domain([e.domain()[0]-i,e.domain()[1]+i])}(t.axes.scale_x,t.axes.scale_y,t.get_aspectRatio()),l("X"),l("Y");var e=n("X"),r=n("Y");if(t.transitionDuration()>0&&!t.transitionOff){var a=d3.transition("axes").duration(t.transitionDuration());d3.axisBottom().scale(t.axes.scale_x).tickValues(e.tickValues).tickFormat(e.tickFormat)(t.axes.x_g.transition(a)),d3.axisLeft().scale(t.axes.scale_y).tickValues(r.tickValues).tickFormat(r.tickFormat)(t.axes.y_g.transition(a))}else d3.axisBottom().scale(t.axes.scale_x).tickValues(e.tickValues).tickFormat(e.tickFormat)(t.axes.x_g),d3.axisLeft().scale(t.axes.scale_y).tickValues(r.tickValues).tickFormat(r.tickFormat)(t.axes.y_g);var s=function(){t.axes.x_g.selectAll(".tick").selectAll("text").html(e.tickFormat)};e.tickFormat&&(t.transitionDuration()>0&&!t.transitionOff?setTimeout(s,t.transitionDuration()):s());var d=function(){t.axes.y_g.selectAll(".tick").selectAll("text").html(r.tickFormat)};for(var c in r.tickFormat&&(t.transitionDuration()>0&&!t.transitionOff?setTimeout(d,t.transitionDuration()):d()),t.axes.y_g.selectAll(".tick").selectAll("text").attr("transform","rotate(-"+t.ticksRotateY()+", -9, 0)").style("text-anchor",t.ticksRotateY()>69?"middle":"end").attr("dy",t.ticksRotateY()>69?0:"0.32em"),t.axes.x_g.selectAll(".tick").selectAll("text").attr("transform","rotate(-"+t.ticksRotateX()+")").style("text-anchor",t.ticksRotateX()>19?"end":"middle").attr("dx",t.ticksRotateX()>19?"-.8em":0).attr("dy",t.ticksRotateX()>19?.55-.01*(t.ticksRotateX()-20)+"em":".71em"),t.updateAxesTitles(),t.layers)t.get_layer(c).updateElementPosition();return t},t.clickPosition=function(e){var r,a,n=function(t,e){return t.domain()[Math.round((e-t.padding()*t.step())/t.step())]};return r=t.axes.scale_x.invert?t.axes.scale_x.invert(e[0]):n(t.axes.scale_x,e[0]),a=t.axes.scale_y.invert?t.axes.scale_y.invert(e[1]):n(t.axes.scale_y,t.axes.scale_y.range()[0]-e[1]),t.get_on_clickPosition(r,a),t},t}function x(t,e){void 0===e&&(e=v()),void 0===t&&(t="layer"+e.get_nlayers());var r=e.create_layer(t).get_layer(t).add_property("lineFun",void 0,function(t){if("function"==typeof t)return t;if("string"==typeof t)return new Function("return "+t)();throw"Error in 'typeCheck' for property 'lineFun': value must be a function ora string that can be evaluated."}).add_property("nsteps",100).add_property("lineWidth",1.5,s("array_fun","lineWidth")).add_property("dasharray",void 0,s("array_fun","dasharray")).add_property("fill","none",s("array_fun","fill"));return e.syncProperties(r),r.nelements(1),r.updateElements=function(){var t=r.g.selectAll(".data_element").data(r.elementIds(),function(t){return t});t.exit().remove(),t.enter().append("path").attr("class","data_element").merge(t).attr("id",function(t){return"p"+(r.id+"_"+t).replace(/[ .]/g,"_")}).on("click",r.get_on_click).on("mouseover",r.get_on_mouseover).on("mouseout",r.get_on_mouseout)},r.dresser(function(t){t.attr("stroke",function(t){return r.get_colour(t)}).attr("stroke-width",function(t){return r.get_lineWidth(t)}).attr("stroke-dasharray",function(t){return r.get_dasharray(t)}).attr("opacity",function(t){return r.get_opacity(t)}).attr("fill",function(t){return r.get_fill(t)})}),r.findElements=function(t,e){var a=d3.max([10,2*r.lineWidth()]);return t[0]-e[0]<a&&t[1]-e[1]<a&&(t=[t[0]+a/2,t[1]+a/2],e=[e[0]-a/2,e[1]-a/2]),r.g.selectAll("path").filter(function(){for(var r,a,n,o,i,l=d3.select(this).attr("d").substr(1).split("L").map(function(t){return t.split(",")}),s=!1,d=0;!s&&d<l.length-1;)n=l[d+1][0]-l[d][0],o=l[d][1]-l[d+1][1],i=l[d+1][1]*l[d][0]-l[d][1]*l[d+1][0],(l[d][0]-t[0])*(l[d+1][0]-t[0])<0?(r=-(t[0]*o+i)/n,s=(t[1]-r)*(e[1]-r)<0):(l[d][0]-e[0])*(l[d+1][0]-e[0])<0?(r=-(e[0]*o+i)/n,s=(t[1]-r)*(e[1]-r)<0):(l[d][1]-t[1])*(l[d+1][1]-t[1])<0?(a=-(t[1]*n+i)/o,s=(t[0]-a)*(e[0]-a)<0):(l[d][1]-e[1])*(l[d+1][1]-e[1])<0&&(a=-(e[1]*n+i)/o,s=(t[0]-a)*(e[0]-a)<0),d++;return s}).data()},r.updateElementPosition=function(){var t=d3.line().x(function(t){return r.chart.axes.scale_x(t.x)}).y(function(t){return r.chart.axes.scale_y(t.y)});return r.chart.transitionDuration()>0&&!r.chart.transitionOff?r.g.selectAll(".data_element").transition("elementPosition").duration(r.chart.transitionDuration()).attr("d",function(e){return t(r.get_data(e))}):r.g.selectAll(".data_element").attr("d",function(e){return t(r.get_data(e))}),r.chart},r.on_mouseover(function(t){var e=d3.mouse(r.chart.container.node());d3.select(this).attr("stroke",function(t){return d3.rgb(r.get_colour(t)).darker(.5)}).classed("hover",!0),r.chart.container.selectAll(".inform").data([t]).style("left",e[0]+10+"px").style("top",e[1]+10+"px").select(".value").html(r.get_informText(t)),r.chart.container.selectAll(".inform").classed("hidden",!1)}),r.on_mouseout(function(t){var e=r.get_marked().length>0;d3.select(this).attr("stroke",function(t){return e^d3.select(this).classed("marked")?"#aaa":r.get_colour(t)}).classed("hover",!1),r.chart.container.selectAll(".inform").classed("hidden",!0)}),r.colourMarked=function(){var t={};t[r.id]=r.get_marked(),(t=r.chart.get_elements(t)).empty()?r.g.selectAll(".data_element").attr("stroke",function(t){return r.get_colour(t)}):r.g.selectAll(".data_element").attr("stroke",function(t){return d3.select(this).classed("marked")?r.get_colour(t):"#aaa"})},e}function b(t,e){var r=x(t,e).activeLayer();return r.type="xLine",r.get_data=function(t){var e=r.layerDomainX();void 0===e&&(e=r.chart.axes.scale_x.domain());for(var a=(e[1]-e[0])/r.get_nsteps(t),n=[],o=e[0];o<e[1];o+=a)n.push({x:o,y:r.get_lineFun(o,t)});d3.line().x(function(t){return r.chart.axes.scale_x(t.x)}).y(function(t){return r.chart.axes.scale_y(t.y)});return n},r.layerDomainY(function(){if(r.layerDomainX()){for(var t=r.elementIds(),e=[],a=0;a<t.length;a++)e=e.concat(d3.extent(r.get_data(t[a]).map(function(t){return t.y})));return d3.extent(e)}}),r.chart}function w(t,e){var r=x(t,e).activeLayer();return r.type="yLine",r.get_data=function(t){var e=r.layerDomainY();void 0===e&&(e=r.chart.axes.scale_y.domain());for(var a=(e[1]-e[0])/r.get_nsteps(t),n=[],o=e[0];o<e[1];o+=a)n.push({y:o,x:r.get_lineFun(o,t)});d3.line().x(function(t){return r.chart.axes.scale_x(t.x)}).y(function(t){return r.chart.axes.scale_y(t.y)});return n},r.layerDomainX(function(){if(r.layerDomainY()){for(var t=r.elementIds(),e=[],a=0;a<t.length;a++)e=e.concat(d3.extent(r.get_data(t[a]).map(function(t){return t.x})));return d3.extent(e)}}),r.chart}function k(t,e){var r=x(t,e).activeLayer();return r.type="paramCurve",r.add_property("xFunction").add_property("yFunction").add_property("paramRange",[0,1]),r.chart.syncProperties(r),r.get_data=function(t){var e=r.paramRange();e[1]<e[0]&&(e=[e[1],e[0]]);for(var a=(e[1]-e[0])/r.get_nsteps(),n=[],o=e[0];o<e[1];o+=a)n.push({x:r.get_xFunction(o,t),y:r.get_yFunction(o,t)});return n},r.layerDomainX(function(){for(var t=r.elementIds(),e=[],a=0;a<t.length;a++)e=e.concat(d3.extent(r.get_data(t[a]).map(function(t){return t.x})));return d3.extent(e)}),r.layerDomainY(function(){for(var t=r.elementIds(),e=[],a=0;a<t.length;a++)e=e.concat(d3.extent(r.get_data(t[a]).map(function(t){return t.y})));return d3.extent(e)}),r.chart}function S(t,e){var r=x(t,e).activeLayer();return r.type="pointLine",r.add_property("x",void 0,s("matrix_fun","x")).add_property("y",void 0,s("matrix_fun","y")),r.chart.syncProperties(r),r.get_data=function(t){for(var e=[],a=0;a<r.nsteps();a++)e.push({x:r.get_x(a,t),y:r.get_y(a,t)});return e},r.layerDomainX(function(){var t=[];return r.elementIds().map(function(t){return d3.extent(d3.range(r.nsteps()).map(function(e){return r.get_x(e,t)}).filter(function(t){return!lc.isNaN(t)}))}).forEach(function(e){t=t.concat(e)}),d3.extent(t)}),r.layerDomainY(function(){var t=[];return r.elementIds().map(function(t){return d3.extent(d3.range(r.nsteps()).map(function(e){return r.get_y(e,t)}).filter(function(t){return!lc.isNaN(t)}))}).forEach(function(e){t=t.concat(e)}),d3.extent(t)}),r.chart}function A(t,e){void 0===e&&(e=v()),void 0===t&&(t="layer"+e.get_nlayers());var r=e.create_layer(t).get_layer(t).add_property("x",void 0,s("array_fun","x")).add_property("y",void 0,s("array_fun","y")).add_property("size",6,s("array_fun","size")).add_property("stroke",function(t){return d3.rgb(r.get_colour(t)).darker(.8)},s("array_fun","stroke")).add_property("strokeWidth",function(t){return.1*r.get_size(t)},s("array_fun","strokeWidth")).add_property("symbol","Circle",s("array_fun","symbol")).add_property("symbolValue",void 0,s("array_fun","symbolValue")).add_property("symbolLegendTitle","").add_property("shiftX",0,s("array_fun","shiftX")).add_property("shiftY",0,s("array_fun","shiftY"));["X","Y"].forEach(function(t){r.wrapSetter("shift"+t,function(a){return function(){return r["get_scaledShift"+t]=function(e){return r.chart.axes["shiftScale"+t](r["get_shift"+t](e))},a.apply(e,arguments)}})}),e.syncProperties(r),r.type="scatter",r.mode("default"),r.chart.informText(function(t){var e=r.get_x(t),a=r.get_y(t);return e.toFixed&&(e=e.toFixed(2)),a.toFixed&&(a=a.toFixed(2)),"ID: <b>"+r.get_label(t)+"</b>;<br>x = "+e+";<br>y = "+a}),r.nelements(function(){for(var t,e=0;e<1e5;e++){try{t=r.get_x(e)}catch(t){return e>=0?e:0}if(void 0===t)return e>=0?e:0}throw"There seem to be very many data points. Please supply a number via 'nelements'."}),r.get_scaledShiftX=function(){return 0},r.get_scaledShiftY=function(){return 0},r.findElements=function(t,e){return r.elementIds().filter(function(a){var n=[r.chart.axes.scale_x(r.get_x(a))+r.get_scaledShiftX(a),r.chart.axes.scale_y(r.get_y(a))+r.get_scaledShiftY(a)];return n[0]-r.get_size(a)-1<=e[0]&&n[1]-r.get_size(a)-1<=e[1]&&n[0]+r.get_size(a)+1>=t[0]&&n[1]+r.get_size(a)+1>=t[1]})},r.get_position=function(t){return[r.chart.axes.scale_x(r.get_x(t))+r.get_scaledShiftX(t),r.chart.axes.scale_y(r.get_y(t))+r.get_scaledShiftY(t)]},r.layerDomainX(function(){var t=r.elementIds().map(function(t){return r.get_x(t)}).filter(function(t){return!lc.isNaN(t)}),e=!0,a=0;if(r.chart.logScaleX())return d3.extent(t.filter(function(t){return t>0}));for(;e&&a<t.length;)"number"!=typeof t[a]&&(e=!1),a++;return e?d3.extent(t):t}),r.layerDomainY(function(){var t=r.elementIds().map(function(t){return r.get_y(t)}).filter(function(t){return!lc.isNaN(t)}),e=!0,a=0;if(r.chart.logScaleY())return d3.extent(t.filter(function(t){return t>0}));for(;e&&a<t.length;)"number"!=typeof t[a]&&(e=!1),a++;return e?d3.extent(t):t}),r.resetSymbolScale=function(){for(var t=[],e=r.elementIds(),a=0;a<e.length;a++)-1==t.indexOf(r.get_symbolValue(e[a]))&&t.push(r.get_symbolValue(e[a]));if(r.symbolScale=d3.scaleOrdinal().domain(t).range(["Circle","Cross","Diamond","Square","Star","Triangle","Wye"]),t[0]&&(r.symbol(function(t){return r.symbolScale(r.get_symbolValue(t))}),r.chart.showLegend())){var n={};n[r.id+"_symbol"]=r.symbolLegendTitle(),r.chart.legend.set_title(n),r.addLegendBlock(r.symbolScale,"symbol",r.id+"_symbol")}};var a=function(){return"default"==r.mode()?r.nelements()>2500?"canvas":"svg":r.mode()};return r.updateElementPosition=function(){if(!r.checkMode())return e;if("svg"==a()){var t=function(t){var e=r.chart.axes.scale_x(r.get_x(t))+r.get_scaledShiftX(t),a=r.chart.axes.scale_y(r.get_y(t))+r.get_scaledShiftY(t);return void 0==e||void 0==a||Number.isNaN(e)||Number.isNaN(a)?"translate(-100, 0)":"translate("+e+", "+a+")"};r.chart.transitionDuration()>0&&!r.chart.transitionOff?r.g.selectAll(".data_element").transition("elementPosition").duration(r.chart.transitionDuration()).attr("transform",t):r.g.selectAll(".data_element").attr("transform",t);var n=r.chart.axes.scale_x.domain(),o=r.chart.axes.scale_y.domain(),i=r.g.selectAll(".data_element").filter(function(t){return void 0!=r.chart.axes.scale_x.invert&&(r.get_x(t)>n[1]||r.get_x(t)<n[0])||void 0!=r.chart.axes.scale_y.invert&&(r.get_y(t)>o[1]||r.get_y(t)<o[0])}).data(),l=r.g.selectAll(".out_tick").data(i,function(t){return t});l.exit().remove(),l.enter().append("rect").attr("class","out_tick").attr("fill",function(t){return r.get_colour(t)}).merge(l).attr("width",function(t){return void 0!=r.chart.axes.scale_y.invert&&(r.get_y(t)>o[1]||r.get_y(t)<o[0])?4:12}).attr("height",function(t){return void 0!=r.chart.axes.scale_x.invert&&(r.get_x(t)>n[1]||r.get_x(t)<n[0])?4:12}).attr("x",function(t){return void 0!=r.chart.axes.scale_x.invert?d3.max([r.chart.axes.scale_x(n[0]),r.chart.axes.scale_x(d3.min([r.get_x(t),n[1]]))-d3.select(this).attr("width")]):r.chart.axes.scale_x(r.get_x(t))}).attr("y",function(t){return void 0!=r.chart.axes.scale_y.invert?d3.min([r.chart.axes.scale_y(o[0])-d3.select(this).attr("height"),r.chart.axes.scale_y(d3.min([r.get_y(t),o[1]]))]):r.chart.axes.scale_y(r.get_y(t))}).on("mousedown",function(t){void 0!=r.chart.axes.scale_x.invert&&r.chart.domainX([d3.min([n[0],r.get_x(t)]),d3.max([n[1],r.get_x(t)])]),void 0!=r.chart.axes.scale_y.invert&&r.chart.domainY([d3.min([o[0],r.get_y(t)]),d3.max([o[1],r.get_y(t)])]),r.chart.updateAxes()})}else r.updateStarted||r.updateCanvas();return e},r.updateElementStyle=function(){if(!r.checkMode())return e;if(r.resetColourScale(),r.resetSymbolScale(),"svg"==a()){var t=r.g.selectAll(".data_element");r.chart.transitionDuration()>0&&!r.chart.transitionOff&&0==r.get_marked().length&&(t=t.transition("elementStyle").duration(r.chart.transitionDuration())),t.attr("d",function(t){return d3.symbol().type(d3["symbol"+r.get_symbol(t)]).size(o(r.get_symbol(t),r.get_size(t)))()}).attr("fill",function(t){return r.get_colour(t)}).attr("stroke",function(t){return r.get_stroke(t)}).attr("stroke-width",function(t){return r.get_strokeWidth(t)}).attr("opacity",function(t){return r.get_opacity(t)}),0!=r.get_marked().length&&r.colourMarked()}else r.updateStarted||r.updateCanvas();return e},r.dresser(function(t){t.attr("fill",function(t){return r.get_colour(t)}).attr("r",function(t){return r.get_size(t)})}),r.updateElements=function(){if(!r.checkMode())return e;if("svg"==a()){var t=r.g.selectAll(".data_element").data(r.elementIds(),function(t){return t});t.exit().remove(),t.enter().append("path").attr("class","data_element").merge(t).attr("id",function(t){return"p"+(r.id+"_"+t).replace(/[ .]/g,"_")}).on("click",r.get_on_click).on("mouseover",r.get_on_mouseover).on("mouseout",r.get_on_mouseout)}else r.updateStarted||r.updateCanvas();return e},r.updateCanvas=function(){var t,a,n,i=r.elementIds(),l=r.canvas.node().getContext("2d");l.clearRect(0,0,e.plotWidth(),e.plotHeight());for(var s=0;s<i.length;s++)0!=r.marked.length&&-1==r.marked.indexOf(i[s])?l.fillStyle="#aaa":l.fillStyle=r.get_colour(i[s]),l.globalAlpha=r.get_opacity(i[s]),a=r.chart.axes.scale_x(r.get_x(i[s]))+r.get_scaledShiftX(i[s]),n=r.chart.axes.scale_y(r.get_y(i[s]))+ +r.get_scaledShiftY(i[s]),a>=0&&a<=r.chart.plotWidth()&&n>=0&&n<=r.chart.plotHeight()?(l.strokeStyle=r.get_stroke(i[s]),l.lineWidth=r.get_strokeWidth(i[s]),t=d3.symbol().type(d3["symbol"+r.get_symbol(i[s])]).size(o(r.get_symbol(i[s]),r.get_size(i[s])))()):(l.strokeStyle=r.get_colour(i[s]),l.lineWidth=2*r.get_strokeWidth(i[s]),t="M 0 0 L ",a<0?(a=0,t+="15 "):a>r.chart.plotWidth()?(a=r.chart.plotWidth(),t+="-15 "):t+="0 ",n<0?(n=0,t+="15"):n>r.chart.plotHeight()?(n=r.chart.plotHeight(),t+="-15"):t+="0"),l.translate(a,n),t=new Path2D(t),l.stroke(t),l.fill(t),l.translate(-a,-n)},e}function C(t,e){void 0===e&&(e=v()),void 0===t&&(t="layer"+e.get_nlayers()),A(t,e);var r=e.get_layer(t);r.add_property("valueAxis","y",function(t){if("x"!=t&&"y"!=t&&"function"!=typeof t)throw"Error in 'typeCheck' for 'valueAxis': axis nameshould be 'x' or 'y'";return t}),e.syncProperties(r),r.type="beeswarm";var a=r.updateElementPosition;return r.updateElementPosition=function(){r.valueAxis();var t=d3.beeswarm().data(r.elementIds().filter(function(t){var e="x"==r.valueAxis()?"y":"x";return void 0!==r.chart.axes["scale_"+e](r["get_"+e](t))}).sort(function(t,e){return r["get_"+r.valueAxis()](t)-r["get_"+r.valueAxis()](e)})).distributeOn(function(t){return r.chart.axes["scale_"+r.valueAxis()](r["get_"+r.valueAxis()](t))}).axis(function(t){return"x"==r.valueAxis()?r.chart.axes.scale_y(r.get_y(t)):r.chart.axes.scale_x(r.get_x(t))}).radius(r.size()).arrange();t.res={};for(var e=0;e<t.length;e++)t.res[t[e].datum]=t[e];return"y"==r.valueAxis()?r.get_scaledShiftX=function(e){return t.res[e]?t.res[e].axis-t.res[e].y:-10}:r.get_scaledShiftY=function(e){return t.res[e]?t.res[e].axis-t.res[e].y:-10},a()},e}function L(t,e){void 0===e&&(e=v()),void 0===t&&(t="layer"+e.get_nlayers());var r=e.create_layer(t).get_layer(t).add_property("ngroups",void 0,s("number_nonneg","ngroups")).add_property("groupIds",void 0,s("array","groupIds")).add_property("nbars",void 0,s("number_nonneg","nbars")).add_property("barIds",void 0,s("array","barIds")).add_property("nstacks",void 0,s("number_nonneg","nstacks")).add_property("stackIds",void 0,s("array","stackIds")).add_property("value",void 0,function(t){if("function"==typeof t)return t;var e;if("object"==typeof t){t.__inds__&&(e=t.__inds__,t.__inds__=null);var r=Object.keys(t);if("object"==typeof t[r[0]]){var a=Object.keys(t[r[0]]);if("object"==typeof t[r[0]][a[0]]){var n=Object.keys(t[r[0]][a[0]]);return"object"==typeof t[r[0]][a[0]][n[0]]?function(r,a,n){return e&&(r=e.indexOf(r)+1),t[r][a][n][0]}:function(r,a,n){return e&&(r=e.indexOf(r)+1),t[r][a][n]}}}}throw"Error in 'typeCheck' for property 'value': the value is not an array or an object."}).add_property("groupWidth",.6).add_property("stroke","#444").add_property("strokeWidth",0);return r.chart.syncProperties(r),r.type="barchart",r.chart.informText(function(t,e,a){var n=t;return r.nbars()>1&&(n+=", "+e),r.nstacks()>1&&(n+=", "+a),"ID: <b>"+n+"</b>;<br>value = "+r.get_value(t,e,a).toFixed(2)}),["group","bar","stack"].forEach(function(t){r.wrapSetter("n"+t+"s",function(e){return function(){return r["get_"+t+"Ids"]=function(){return d3.range(e())},e.apply(r,arguments)}}),r.wrapSetter(t+"Ids",function(e){return function(){return r["get_n"+t+"s"]=function(){return e().length},e.apply(r,arguments)}})}),r.nbars(1),r.nstacks(1),r.contScaleX(!1),r.elementIds(function(){if(1==r.nbars()&&(0==r.barIds()[0]||1==r.barIds()[0]))return r.stackIds();for(var t=[],e=r.barIds(),a=r.stackIds(),n=0;n<r.nbars();n++)for(var o=0;o<r.nstacks();o++)t.push(e[n]+", "+a[o]);return t}),r.colourValue(function(t){return!t.split||1!=r.nstacks()||0!=r.stackIds()[0]&&1!=r.stackIds()[0]?t.toString():t.split(", ")[0].toString()}),r.colour(function(t,e,a){return 1!=r.nbars()||0!=e&&1!=e?r.colourScale(r.get_colourValue(e+", "+a)):r.colourScale(r.get_colourValue(a))}),r.addColourScaleToLegend(!0),r.layerDomainX(function(){var t=r.groupIds();return r.contScaleX()?[d3.min(t),d3.max(t)]:r.groupIds()}),r.layerDomainY(function(){for(var t,e=r.barIds(),a=r.groupIds(),n=r.stackIds(),o=0,i=0;i<r.ngroups();i++)for(var l=0;l<r.nbars();l++){t=0;for(var s=0;s<r.nstacks();s++)t+=r.get_value(a[i],e[l],n[s]);t>o&&(o=t)}return[0,o]}),r.on_mouseover(function(t){var a=d3.mouse(e.container.node());d3.select(this).attr("fill",function(t){return d3.rgb(r.get_colour(t[0],t[1],t[2])).darker(.5)}).classed("hover",!0),r.chart.container.select(".inform").style("left",a[0]+10+"px").style("top",a[1]+10+"px").select(".value").html(r.get_informText(t[0],t[1],t[2])),r.chart.container.select(".inform").classed("hidden",!1)}),r.on_mouseout(function(t){var e=r.get_marked().length>0;d3.select(this).attr("fill",function(t){return e^d3.select(this).classed("marked")?"#aaa":r.get_colour(t[0],t[1],t[2])}).classed("hover",!1),r.chart.container.select(".inform").classed("hidden",!0)}),r.findElements=function(t,e){return r.g.selectAll(".data_element").filter(function(){var r=+d3.select(this).attr("x"),a=+d3.select(this).attr("y"),n=+d3.select(this).attr("width"),o=+d3.select(this).attr("height");return t[0]<=r+n&&e[0]>r&&t[1]<=a+o&&e[1]>a}).data()},r.get_position=function(t){return[r.g.select("#p"+t.join("_")).attr("x"),r.g.select("#p"+t.join("_")).attr("y")]},r.updateElementPosition=function(){var t,e=r.groupIds(),a=(t=r.contScaleX()?Math.abs(r.chart.axes.scale_x(e[1])-r.chart.axes.scale_x(e[0]))*r.groupWidth():r.chart.axes.scale_x.step()*r.groupWidth())/r.nbars(),n=Math.abs(r.chart.axes.scale_y(1)-r.chart.axes.scale_y(0)),o=d3.scaleLinear().domain([0,r.nbars()-1]).range([-t/2,t/2-a]),i=r.barIds(),l=r.stackIds();return r.chart.transitionDuration()>0&&!r.chart.transitionOff?r.g.selectAll(".data_element").transition("elementPosition").duration(r.chart.transitionDuration()).attr("width",a).attr("height",function(t){return r.get_value(t[0],t[1],t[2])*n}).attr("x",function(t){return void 0==r.chart.axes.scale_x(t[0])?-500:o(i.indexOf(t[1]))+r.chart.axes.scale_x(t[0])}).attr("y",function(t){for(var e=0,a=0;a<=l.indexOf(t[2]);a++)e+=r.get_value(t[0],t[1],l[a]);return r.chart.axes.scale_y(e)}):r.g.selectAll(".data_element").attr("width",a).attr("height",function(t){return r.get_value(t[0],t[1],t[2])*n}).attr("x",function(t){return void 0==r.chart.axes.scale_x(t[0])?-500:o(i.indexOf(t[1]))+r.chart.axes.scale_x(t[0])}).attr("y",function(t){for(var e=0,a=0;a<=l.indexOf(t[2]);a++)e+=r.get_value(t[0],t[1],l[a]);return r.chart.axes.scale_y(e)}),r},r.updateElementStyle=function(){r.resetColourScale();var t=r.g.selectAll(".data_element");r.chart.transitionDuration()>0&&!r.chart.transitionOff&&0==r.get_marked().length&&(t=t.transition("elementStyle").duration(r.chart.transitionDuration())),t.attr("fill",function(t){return r.get_colour(t[0],t[1],t[2])}).attr("stroke",function(t){return r.get_stroke(t[0],t[1],t[2])}).attr("stroke-width",function(t){return r.get_strokeWidth(t[0],t[1],t[2])}).attr("opacity",function(t){return r.get_opacity(t[0],t[1],t[2])}),0!=r.get_marked().length&&r.colourMarked()},r.updateElements=function(){var t=r.g.selectAll(".group").data(r.groupIds(),function(t){return t});t.exit().remove(),t.enter().append("g").attr("class","group");var e=r.g.selectAll(".group").selectAll(".bar").data(function(t){return r.barIds().map(function(e){return[t,e]})},function(t){return t});e.exit().remove(),e.enter().append("g").attr("class","bar");var a=r.g.selectAll(".group").selectAll(".bar").selectAll(".data_element").data(function(t){return r.stackIds().map(function(e){return t.concat(e)})},function(t){return t});a.exit().remove(),a.enter().append("rect").attr("class","data_element").merge(a).attr("id",function(t){return"p"+r.id+"_"+t.join("_").replace(/[ .]/g,"_")}).on("click",function(t){r.get_on_click(t[0],t[1],t[2])}).on("mouseover",r.get_on_mouseover).on("mouseout",r.get_on_mouseout)},r.colourMarked=function(){var t={};t[r.id]=r.get_marked(),(t=r.chart.get_elements(t)).empty()?r.g.selectAll(".data_element").attr("fill",function(t){return r.get_colour(t[0],t[1],t[2])}):r.g.selectAll(".data_element").attr("fill",function(t){return d3.select(this).classed("marked")?r.get_colour(t[0],t[1],t[2]):"#aaa"})},r.chart}function M(){var t=f().add_property("activeLayer",void 0).add_property("layerIds",function(){return Object.keys(t.layers)},s("array","layerIds")).add_property("layerType",function(e){return t.get_layer(e).type},s("array_fun","layerType")).add_property("globalColourScale",!0);t.layers={};var e=function(e){return function(){if(t.get_activeLayer()[e])return t.get_activeLayer()[e].apply(t,arguments);for(var r in t.layers)if(t.layers[r][e])return t.layers[r][e].apply(t,arguments)}};t.syncProperties=function(r){for(var a=0;a<r.propList.length;a++)void 0===t[r.propList[a]]&&(t[r.propList[a]]=e(r.propList[a]),t["get_"+r.propList[a]]=e("get_"+r.propList[a]))},t.get_nlayers=function(){return Object.keys(t.layers).length},t.get_layer=function(e){if(-1==Object.keys(t.layers).indexOf(e))throw"Error in 'get_layer': layer with id "+e+" is not defined";return t.layers[e]},t.create_layer=function(e){void 0===e&&(e="layer"+t.get_nlayers());var r=c(e);return r.chart=t,t.layers[e]=r,t.activeLayer(t.get_layer(e)),t},t.add_layer=function(e){var r;void 0===e&&(e="layer"+t.get_nlayers());try{r=t.get_layerType(e)}catch(t){}return void 0===r?t.create_layer(e):("scatter"==r&&A(e,t),"xLine"==r&&b(e,t),"yLine"==r&&w(e,t),"paramCurve"==r&&k(e,t),"bar"==r&&L(e,t),"beeswarm"==r&&C(e,t)),t},t.remove_layer=function(e){if(-1==Object.keys(t.layers).indexOf(e))return-1;for(var r in t.layers[e].legendBlocks)t.legend.removeBlock(t.layers[e].legendBlocks[r]);try{t.layers[e].g.remove(),t.layers[e].canvas.remove()}catch(t){}return delete t.layers[e],0},t.select_layers=function(e){void 0===e&&(e=t.layerIds());for(var r,a={layers:{}},n=0;n<e.length;n++)-1!=t.layerIds().indexOf(e[n])?(void 0===t.layers[e[n]]&&(t.add_layer(e[n]),t.svg&&t.get_layer(e[n]).put_static_content()),a.layers[e[n]]=t.get_layer(e[n])):(e.splice(n,1),n--);if(0==Object.keys(a.layers).length){for(n in t)a[n]=function(){return a};return a}for(var o=0;o<e.length;o++)for(n=0;n<a.layers[e[o]].propList.length;n++)r=a.layers[e[o]].propList[n],void 0===a[r]&&(a[r]=function(t){return function(r){var n;n="function"!=typeof r?function(){return r}:r;for(var o=0;o<e.length;o++)void 0!==a.layers[e[o]][t]&&a.layers[e[o]][t](function(t){return function(){for(var e=[],r=0;r<arguments.length;r++)e.push(arguments[r]);return e.unshift(t),n.apply(void 0,e)}}(e[o]));return a}}(r));return 0==a.length?t:a};var r={};t.globalColourDomain=function(e,a){void 0!==e&&(r[e]=a);var n=[];for(var o in t.layers)r[o]||(r[o]=t.get_layer(o).colourDomain()),n=n.concat(r[o]);return n.filter(function(t,e,r){return void 0!==t&&r.indexOf(t)===e}),n},t.place_layer=function(e){t.get_layer(e).put_static_content(),t.get_layer(e).update()};var a=t.put_static_content;t.put_static_content=function(e){for(var r in a(e),n(t),t.layers)t.get_layer(r).put_static_content()},t.findElements=function(e,r){var a={};return Object.keys(t.layers).forEach(function(n){a[n]=t.get_layer(n).findElements(e,r)}),a},t.get_elements=function(e){if("object"!=typeof e&&(e=[e]),Array.isArray(e)){if(1!=t.get_nlayers())throw"Error in 'get_elements': layerId is not defined";var r=e;(e={})[Object.keys(t.layers)[0]]=r}var a=[],n={};for(var o in e)t.get_layer(o).canvas&&t.get_layer(o).canvas.classed("active")?n[o]=e[o]:a=a.concat(e[o].map(function(t){return"object"==typeof t&&(t=t.join("_")),("p"+o+"_"+t).replace(/[ .]/g,"_")}));var i=0==a.length?d3.select("_______"):t.svg.selectAll(".chart_g").selectAll("#"+a.join(",#"));return i.canvas=n,i},t.get_marked=function(){var e={};for(var r in t.layers)e[r]=t.get_layer(r).get_marked();return e};var o=t.mark;t.mark=function(e,r){if(Array.isArray(e))o(e,r);else if(!e.empty||e.canvas&&0!=Object.keys(e.canvas))if("__clear__"!=e)for(var a in e.canvas&&(e=e.canvas),e)t.get_layer(a).mark(e[a],r);else for(var a in t.layers)t.get_layer(a).mark(e,r);else o(e,r)};var i=t.update;t.update=function(){for(var e=t.layerIds(),r=0;r<e.length;r++)void 0===t.layers[e[r]]&&t.add_layer(e[r]);for(var a in t.layers)-1==e.indexOf(a)?t.remove_layer(a):(t.get_layer(a).updateStarted=!0,t.get_layer(a).updateElements(),t.get_layer(a).updateElementStyle(),t.get_layer(a).updateStarted=!1);return i(),t};var l=t.updateSize;return t.updateSize=function(){for(var e in l(),t.layers)t.get_layer(e).canvas.style("left",t.paddings().left+"px").style("top",t.paddings().top+"px").attr("width",t.plotWidth()).attr("height",t.plotHeight())},t}var I=function(t,e){for(var r=[],a=[t,e],n=[t.length,e.length],o=n.indexOf(Math.min.apply(null,n)),i=0==o?1:0,l=0;l<a[o].length;l++)-1!=a[i].indexOf(a[o][l])&&r.push(a[o][l]);return r};function O(t,e){this.id=t,this.val=e,this.left=null,this.right=null,this.parent=null,this.height=0,this.x=null,this.val_inds=[t]}function z(t){var r=e().add_property("orientation","horizontal").add_property("height",300).add_property("width",500).add_property("nelements").add_property("elementIds",function(){}).add_property("paddings",{left:20,top:20,bottom:20,right:20}).add_property("distance",function(t,e){return lc.getEuclideanDistance(t,e)}).add_property("data").add_property("lineColours",["black","red"]).add_property("on_click",function(){});r.wrapSetter("nelements",function(t){return function(){return r.get_elementIds=function(){return d3.range(t())},t.apply(r,arguments)}}),r.wrapSetter("elementIds",function(t){return function(){return r.get_nelements=function(){return t().length},t.apply(r,arguments)}}),r.heatmap=t,r.clusters=void 0,r.set_paddings=function(t){return void 0===t.top&&(t.top=r.paddings().top),void 0===t.bottom&&(t.bottom=r.paddings().bottom),void 0===t.left&&(t.left=r.paddings().left),void 0===t.right&&(t.right=r.paddings().right),r.paddings(t),r};var a=0,n=function(t){if(null==t.right&&null==t.left)return t.x=a,void a++;null==t.left.x&&n(t.left),null==t.right.x&&n(t.right),t.x=(t.right.x+t.left.x)/2},o=function(t,e,a){if(null!=t.right&&null!=t.left){e.append("line").attr("x1",a[0](t.left.x)).attr("x2",a[0](t.right.x)).attr("y1",a[1](t.height)).attr("y2",a[1](t.height)).attr("stroke-width",3).attr("id",t.id).attr("orient","h").attr("stroke",r.lineColours()[0]);for(var n=[t.left,t.right],i=0;i<n.length;i++)e.append("line").attr("x1",a[0](n[i].x)).attr("x2",a[0](n[i].x)).attr("y1",a[1](t.height)).attr("y2",a[1](n[i].height)).attr("stroke-width",3).attr("id",n[i].id).attr("orient","v").attr("stroke",r.lineColours()[0]);return o(t.left,e,a),void o(t.right,e,a)}if(null!=t.right||null!=t.left){var l=t.right||t.left;e.append("line").attr("x1",a[0](t.x)).attr("x2",a[0](t.x)).attr("y1",a[1](t.height)).attr("y2",a[1](l.height)).attr("stroke-width",3).attr("id",l.id).attr("orient","v").attr("stroke",r.lineColours()[0]),o(l,e,a)}},i=function(t,e){return null!=t?t.id==e?t:i(t.left,e)||i(t.right,e):null},l=function(t,e){e.push(t.id),null!=t.left&&l(t.left,e),null!=t.right&&l(t.right,e)},s=function(t,e,a){return e.selectAll("line").on("click",function(n){!function(t,e,a,n){var o=i(t,e[0]),s=[];if(l(o,s),"h"==e[1]&&e[0],void 0!=r.heatmap){if("horizontal"==r.orientation()){var d=I(s,r.heatmap.colIds());r.heatmap.cluster("Row",d),r.heatmap.drawDendogram("Row")}"vertical"==r.orientation()&&(d=I(s,r.heatmap.rowIds()),r.heatmap.cluster("Col",d),r.heatmap.drawDendogram("Col")),r.heatmap.updateLabelPosition()}else r.get_on_click(I(s,r.elementIds()));!function(t,e,r,a){t.selectAll("line").attr("stroke",function(t){return function(t,e){for(var r=0;r<e.length;r++)if(e[r]==t)return!0;return!1}(this.id,e)?r[1]:r[0]})}(a,s,n)}(t,[this.getAttribute("id"),this.getAttribute("orient")],e,a)}),r};r.cluster=function(){r.get_data("clear");var t=r.elementIds();r.bucket=[];for(var e=r.elementIds(),a=0;a<t.length;a++)r.bucket[a]=new O(t[a],r.get_data(t[a]));var n=function(t,r){for(var a,n=o[e.indexOf(t[0])][e.indexOf(r[0])],i=0;i<t.length;i++)for(var l=0;l<r.length;l++)(a=o[e.indexOf(t[i])][e.indexOf(r[l])])>n&&(n=a);return n},o=d();r.clusters=function(){r.bucket.length;for(var t=JSON.parse(JSON.stringify(r.bucket));t.length>1;){for(var e=[t[0],t[1]],a=n(t[0].val_inds,t[1].val_inds),o=[0,1],i=0;i<t.length;i++)for(var l=i+1;l<t.length;l++){var s=n(t[i].val_inds,t[l].val_inds);s<a&&(a=s,e=[t[i],t[l]],o=[i,l])}var d=new O(e[0].id+"_"+e[1].id,null);d.left=e[0],d.right=e[1],d.height=a,d.val_inds=d.left.val_inds.concat(d.right.val_inds),o.sort(function(t,e){return e-t}),t.splice(o[0],1),t.splice(o[1],1),t.push(d)}return t[0]}()};var d=function(){for(var t=r.elementIds(),e=new Array(t.length),a=0;a<t.length;a++)e[a]=new Array(t.length);for(a=0;a<t.length;a++)for(var n=a;n<t.length;n++){var o=r.get_distance(r.get_data(t[a]),r.get_data(t[n]));e[a][n]=o,e[n][a]=o}return e};r.draw=function(){n(r.clusters),a=0,d3.selectAll(r.g.node().childNodes).remove(),"vertical"==r.orientation()?(r.heatmap?(r.width(r.heatmap.height()).height(r.heatmap.paddings().left).paddings({left:r.heatmap.paddings().top,top:0,bottom:0,right:r.heatmap.paddings().bottom}),r.svg.select(".row").attr("transform","translate("+(r.heatmap.paddings().left+ +r.heatmap.plotWidth()+5)+", "+r.heatmap.paddings().top+")").selectAll("text").style("text-anchor","start")):r.svg.attr("width",r.width()).attr("height",r.height()),r.g.attr("transform","rotate(90) translate(0, -"+r.height()+")")):r.heatmap?(r.width(r.heatmap.width()).height(r.heatmap.paddings().top).paddings({top:0,left:r.heatmap.paddings().left,right:r.heatmap.paddings().right,bottom:0}),r.svg.select(".col").attr("transform","translate("+ +r.paddings().left+", "+(r.heatmap.paddings().top+ +r.heatmap.plotHeight()+5)+")").selectAll("text").style("text-anchor","end")):r.svg.attr("width",r.width()).attr("height",r.height());var t,e,i,l,d,u,p=c();if(void 0!==p)return r.scales=(t=-1,e=0,i=r.paddings(),l=r.elementIds().length,d=(r.width()-i.right-i.left)/l,u=d3.scaleLinear().domain([0,l-1]).range([i.left+d/2,r.width()-i.right-d/2]),"vertical"==r.get_orientation()&&(e=r.height(),t=1),[u,d3.scaleLinear().domain([0,r.clusters.height]).range([r.height()+i.top*t-e,e-i.bottom*t])]),o(p,r.g,r.scales),s(p,r.g,r.lineColours()),r};var c=function(){for(var t=r.elementIds(),e={id:r.clusters.id,left:null,right:null,val_inds:r.clusters.val_inds,x:r.clusters.x,val:r.clusters.val,original:r.clusters,height:r.clusters.height,parent:null},a=function(t,e){if(0!=t.height)if(t.left&&-1!=t.left.val_inds.indexOf(e))a(t.left,e);else if(t.right&&-1!=t.right.val_inds.indexOf(e))a(t.right,e);else{if(-1!=t.original.left.val_inds.indexOf(e))return t.left={id:t.original.left.id,left:null,right:null,val_inds:t.original.left.val_inds,x:t.original.left.x,val:t.original.left.val,original:t.original.left,height:t.original.left.height,parent:t},void a(t.left,e);t.right={id:t.original.right.id,left:null,right:null,val_inds:t.original.right.val_inds,x:t.original.right.x,val:t.original.right.val,original:t.original.right,height:t.original.right.height,parent:t},a(t.right,e)}},o=0;o<t.length;o++){if(-1==r.clusters.val_inds.indexOf(t[o]))return r.heatmap?void r.remove():(r.cluster(),n(r.clusters),r.clusters);a(e,t[o])}var i=0,l=function(t){if(null!==t){if(l(t.left),l(t.right),0==t.height)return t.x=i,void i++;t.left&&t.right&&(t.x=(t.left.x+t.right.x)/2),null!=t.parent&&(null==t.left&&(t.x=t.right.x),null==t.right&&(t.x=t.left.x))}};l(e);var s=function(t){return t.left&&t.right?t:t.left?s(t.left):t.right?s(t.right):void 0};return s(e)};return r.put_static_content=function(e){return r.heatmap?(r.g=r.heatmap.svg.append("g").attr("class","dendogram"),r.svg=t.svg,r.container=t.container):(r.container=e.append("div").style("position","relative"),r.svg=r.container.append("svg"),r.g=r.svg.append("g")),r},r.remove=function(){var t;if(r.g.remove(),r.heatmap){var e=r.heatmap;if(t="horizontal"==r.orientation()?"Col":"Row",e.showDendogram(t,!1),e["dendogram"+t]=void 0,e.transitionDuration()>0&&!e.transitionOff){var a=d3.transition("remove").duration(e.transitionDuration());e.svg.selectAll(".label_panel."+t.toLowerCase()).transition(a).attr("transform","translate("+e.paddings().left+", "+e.paddings().top+")"),"Row"==t?e.svg.select(".row").selectAll("text").transition(a).style("text-anchor","end"):e.svg.select(".col").selectAll("text").transition(a).style("text-anchor","start")}else e.svg.selectAll(".label_panel."+t.toLowerCase()).attr("transform","translate("+e.paddings().left+", "+e.paddings().top+")"),"Row"==t?e.svg.select(".row").selectAll("text").style("text-anchor","end"):e.svg.select(".col").selectAll("text").style("text-anchor","start")}},r.update=function(){return r.cluster(),r.draw(),r},r.place=function(t){if(void 0===t&&(t="body"),"string"==typeof t){var e=t;if(0==(t=d3.select(e)).size())throw"Error in function 'place': DOM selection for string '"+e+"' did not find a node."}return r.put_static_content(t),void 0===r.heatmap&&r.update(),r},r}function D(t,e,r){return 1/(1+Math.exp(-r*(t-e)))}function R(){var t=f().add_property("content","");t.width(0).height(0).paddings({top:5,left:5,bottom:5,right:5}).showPanel(!1);var e=t.put_static_content;return t.put_static_content=function(r){e(r),t.svg.remove(),t.container.style("overflow","auto"),t.container.select("table").remove(),t.container.select("div").remove()},t.updateSize=function(){return t.container.style("width",0!=t.width()?t.width():void 0).style("height",0!=t.height()?t.height():void 0).style("padding-top",t.paddings().top).style("padding-left",t.paddings().left).style("padding-right",t.paddings().right).style("padding-bottom",t.paddings().bottom),t},t.inherited_update=t.update,t.update=function(){return t.inherited_update(),t.container.node().innerHTML=t.content(),t},t}t.base=e,t.layerBase=c,t.layerChart=M,t.chartBase=f,t.axesChart=v,t.scatter=A,t.xLine=b,t.yLine=w,t.parametricCurve=k,t.pointLine=S,t.pointRibbon=function(t,e){var r=S(t,e).activeLayer();return r.add_property("ymax",void 0,s("matrix_fun","ymax")).add_property("ymin",void 0,s("matrix_fun","ymin")),r.type="pointRibbon",r.chart.syncProperties(r),r.opacity(.33).lineWidth(0),r.get_data=function(t){for(var e=[],a=0;a<r.nsteps();a++)e.push({x:r.get_x(a,t),y:r.get_ymin(a,t)});for(a=r.nsteps()-1;a>=0;a--)e.push({x:r.get_x(a,t),y:r.get_ymax(a,t)});return e},r.dresser(function(t){t.attr("fill",function(t){return r.get_colour(t)}).attr("stroke-width",function(t){return r.get_lineWidth(t)}).attr("stroke-dasharray",function(t){return r.get_dasharray(t)}).attr("opacity",function(t){return r.get_opacity(t)})}),r.layerDomainX(function(){var t=[];return r.elementIds().map(function(e){return d3.range(r.nsteps()).map(function(a){0==t.length?t=[r.get_x(a,e),r.get_x(a,e)]:(t[0]=d3.min([r.get_x(a,e),t[0]]),t[1]=d3.max([r.get_x(a,e),t[1]]))})}),t}),r.layerDomainY(function(){var t=[];return r.elementIds().map(function(e){return d3.range(r.nsteps()).map(function(a){0==t.length?t=[r.get_y(a,e),r.get_y(a,e)]:(t[0]=d3.min([t[0],r.get_ymax(a,e),r.get_ymin(a,e)]),t[1]=d3.max([t[1],r.get_ymax(a,e),r.get_ymin(a,e)]))})}),t}),r.chart},t.heatmap=function(t,e){var o;(e=f().add_property("nrows",void 0,s("number_nonneg","nrows")).add_property("ncols",void 0,s("number_nonneg","ncols")).add_property("colLabel",function(t){return t},s("array_fun","colLabel")).add_property("rowLabel",function(t){return t},s("array_fun","rowLabel")).add_property("colIds",void 0,s("array","colIds")).add_property("rowIds",void 0,s("array","rowIds")).add_property("dispColIds",function(){return e.colIds()},s("array","dispColIds")).add_property("dispRowIds",function(){return e.rowIds()},s("array","dispRowIds")).add_property("heatmapRow",function(t){return e.dispRowIds().indexOf(t)},s("array_fun","heatmapRow")).add_property("heatmapCol",function(t){return e.dispColIds().indexOf(t)},s("array_fun","heatmapCol")).add_property("showDendogramRow",!0).add_property("showDendogramCol",!0).add_property("value",void 0,s("matrix_fun","value")).add_property("mode","default").add_property("colour").add_property("palette",t=>d3.interpolateRdYlBu(1-t),function(t){if("string"==typeof t){if(!d3[t])throw"Error in 'typeCheck' for 'palette': invalid palette name, must be one of d3 interpolators (e.g. 'interpolateOrRd').";return d3[t]}if(Array.isArray(t))return t=t.map(t=>/^#[0-9a-fA-F]{8,8}$/.test(t)?t.substring(0,7):t),d3.scaleLinear().domain(d3.range(t.length).map(function(e){return e/t.length})).range(t);if("function"==typeof t)return t;throw"Error in 'typeCheck' for 'palette': invalid type. Palette must be a name of a d3 interpolator, an array of colours or a function."}).add_property("colourDomain",function(){return e.dataRange()},s("array","colourDomain")).add_property("clusterRowMetric",a).add_property("clusterColMetric",a).add_property("on_click",function(){}).add_property("rowTitle","").add_property("showValue",!1).add_property("valueTextColour",(t,r)=>{let a=d3.hsl(e.get_colour(e.get_value(t,r)));return a.h=(a.h+180)%360,a.s=Math.sqrt(a.s),a.l=1-a.l,a},function(t){if("function"==typeof t)return t;if("string"==typeof t)return t;throw"Error in 'typeCheck' for 'valueTextColour': invalid type. valueTextColour must be a colour name or a function."}).add_property("showInform",!0).add_property("colTitle","").add_property("on_mouseover").add_property("on_mouseout").add_property("on_labelClickRow").add_property("on_labelClickCol").add_property("clusterRows",!1).add_property("clusterCols",!1).add_property("informText",function(t,r){var a=e.get_value(t,r);return"number"==typeof a&&(a=a.toFixed(2)),"Row: <b>"+e.get_rowLabel(t)+"</b>;<br>Col: <b>"+e.get_colLabel(r)+"</b>;<br>value = "+a},s("matrix_fun","value"))).paddings({top:100,left:100,right:10,bottom:40}),e.legend.width(75),["col","row"].forEach(function(t){e.wrapSetter("n"+t+"s",function(r){return function(){return e["get_"+t+"Ids"]=function(){return d3.range(r())},r.apply(e,arguments)}}),e.wrapSetter(t+"Ids",function(r){return function(){e["get_n"+t+"s"]=function(){return r().length};var a=t[0].toUpperCase()+t.slice(1);return r.apply(e,arguments),e["get_disp"+a+"Ids"]=e["get_"+t+"Ids"],e}})}),e.wrapSetter("colour",function(t){return function(){var r=t.apply(e,arguments),a=e.get_colour;return e.get_colour=function(){return 1==arguments.length&&void 0===arguments[0]?"#eee":a.apply(e,arguments)},r}}),["col","row"].forEach(function(t){var r="col"==t?"bottom":"right";e.wrapSetter(t+"Title",function(t){return function(){let a={};return a[r]=d3.max([e.paddings()[r],20]),e.set_paddings(a),t.apply(e,arguments)}})}),e.colour(function(t){return e.colourScale(t)}),e.axes={},e.marked=[],o={Row:e.showDendogramRow(),Col:e.showDendogramCol()},e.showDendogram=function(t,r){return void 0===r?o[t]:(o[t]=r&&e["showDendogram"+t](),e)},e.showDendogram("Row",!1).showDendogram("Col",!1);var l=e.put_static_content;e.put_static_content=function(t){l(t),n(e),e.svg.append("g").attr("class","row label_panel"),e.svg.append("g").attr("class","col label_panel"),e.canvas=e.container.append("canvas").style("position","absolute").style("z-index",-5).attr("id","hCanvas"),e.g=e.svg.select(".plotArea").append("g").attr("class","chart_g"),e.text=e.g.append("g").attr("class","text_g"),e.axes.x_label=e.svg.append("text").attr("class","axisLabel").attr("text-anchor","end"),e.axes.y_label=e.svg.append("text").attr("class","axisLabel").attr("text-anchor","end").attr("transform","rotate(-90)"),"svg"==d()?e.g.classed("active",!0):e.canvas.classed("active",!0),e.svg.select(".clickPanel").on("mouseover",function(){e.container.select(".inform").classed("hidden",!1)}).on("mouseout",function(){e.container.select(".inform").classed("hidden",!0)}),e.showPanel()&&(e.panel.add_button("Zoom in","#zoomIn",function(t){var e=-Math.ceil(.1*t.dispRowIds().length),r=-Math.ceil(.1*t.dispColIds().length);t.dispRowIds(p(e,"top")),t.dispRowIds(p(e,"bottom")),t.dispColIds(p(r,"left")),t.dispColIds(p(r,"right")),t.updateStarted=!0,t.updateLabels(),t.updateLabelPosition(),t.updateStarted=!1},"Double click to return to original scales"),e.panel.add_button("Zoom out","#zoomOut",function(t){var e=Math.ceil(.1*t.dispRowIds().length),r=Math.ceil(.1*t.dispColIds().length);t.dispRowIds(p(e,"top")),t.dispRowIds(p(e,"bottom")),t.dispColIds(p(r,"left")),t.dispColIds(p(r,"right")),t.updateStarted=!0,t.updateLabels(),t.updateLabelPosition(),t.updateCellColour(),t.updateLabelText(),t.updateStarted=!1},"Double click to return to original scales"),e.panel.add_button("Cluster rows","#clusterRows",function(t){t.cluster("Row"),t.showDendogram("Row",!0),t.updateLabelPosition()}),e.panel.add_button("Cluster columns","#clusterCols",function(t){t.cluster("Col"),t.showDendogram("Col",!0),t.updateLabelPosition()}),e.panel.add_button("Restore original order","#restoreOrder",function(t){var e=t.rowIds().slice(),r=t.colIds().slice();t.reorder("Row",function(t,r){return e.indexOf(t)-e.indexOf(r)}),t.reorder("Col",function(t,e){return r.indexOf(t)-r.indexOf(e)}),t.dendogramRow&&t.dendogramRow.remove(),t.dendogramCol&&t.dendogramCol.remove(),t.updateStarted=!0,t.updateLabels().updateLabelPosition().updateLabelText().updateCellColour(),t.updateStarted=!1}))};var d=function(){return"default"==e.mode()?e.dispColIds().length*e.dispRowIds().length>2500?"canvas":"svg":e.mode()};function c(){d3.select(this).classed("hover",!0)}function u(){d3.select(this).classed("hover",!1)}function p(t,r){var a,n,o;if("top"!=r&&"bottom"!=r||(-1==(a=e.get_heatmapRow("__order__"))&&(a=e.rowIds()),n=e.dispRowIds()),"left"!=r&&"right"!=r||(-1==(a=e.get_heatmapCol("__order__"))&&(a=e.colIds()),n=e.dispColIds()),0==t)return n;if(t<0){var i,l;if(t=-t,"top"==r||"left"==r){for(i=0;t>0&&n.length>1;)-1!=(l=n.indexOf(a[i]))&&(t--,n.splice(l,1)),i++;return n}for(i=a.length-1;t>0&&n.length>1;)-1!=(l=n.indexOf(a[i]))&&(t--,n.splice(l,1)),i--;return n}if("top"==r||"left"==r){o=a.length;for(var s=0;s<n.length;s++)o>a.indexOf(n[s])&&(o=a.indexOf(n[s]));for(s=o-1;s>=d3.max([0,o-t]);s--)n.unshift(a[s])}else{for(o=-1,s=0;s<n.length;s++)o<a.indexOf(n[s])&&(o=a.indexOf(n[s]));for(s=o+1;s<d3.min([a.length,o+t+1]);s++)n.push(a[s])}return n}e.findElements=function(t,r){var a=[];if("svg"==d())a=e.g.selectAll(".data_element").filter(function(){var a=[this.x.baseVal.value,this.y.baseVal.value];return a[0]<=r[0]&&a[1]<=r[1]&&a[0]+e.cellSize.width>=t[0]&&a[1]+e.cellSize.height>=t[1]}).data();else for(var n=e.svg.select(".col").selectAll(".label").filter(function(){var a=this.y.baseVal[0].value;return a>=t[0]&&a<=r[0]+e.cellSize.width}).data(),o=e.svg.select(".row").selectAll(".label").filter(function(){var a=this.y.baseVal[0].value;return a>=t[1]&&a<=r[1]+e.cellSize.height}).data(),i=0;i<o.length;i++)for(var l=0;l<n.length;l++)a.push([o[i],n[l]]);return a},e.get_position=function(t){return[e.axes.scale_x(e.get_heatmapCol(t[1]))+e.cellSize.width/2,e.axes.scale_y(e.get_heatmapRow(t[0]))+e.cellSize.height/2]},e.dataRange=function(){var t,r,a=0,n=e.get_rowIds(),o=e.get_colIds();do{r=d3.extent(o,function(t){return e.get_value(n[a],t)}),void 0===t&&(t=r),r[0]<t[0]&&(t[0]=r[0]),r[1]>t[1]&&(t[1]=r[1]),a++}while(a<e.get_nrows());return t},e.reorder=function(t,r){if("flip"==r)return e["get_heatmap"+t]("__flip__"),e.updateLabelPosition(),e;r.domain=e["disp"+t+"Ids"]().slice();var a=e["get_heatmap"+t]("__order__");-1==a&&(a=e[t.toLowerCase()+"Ids"]().slice());var n=a.slice(),o=function(a,o){return-1!=r.domain.indexOf(a)&&-1!=r.domain.indexOf(o)?r(a,o):-1!=n.indexOf(a)&&-1!=n.indexOf(o)?n.indexOf(a)-n.indexOf(o):e[t.toLowerCase()+"Ids"]().indexOf(a)-e[t.toLowerCase()+"Ids"]().indexOf(o)},i=e["disp"+t+"Ids"]().slice().sort(r);a.sort(o);var l,s=1;return e["heatmap"+t](function(r){return"__flip__"==r?(s*=-1,void a.reverse()):"__order__"==r?a:"__sort__"!=r?i.indexOf(r):void(i=e["disp"+t+"Ids"]().slice().sort(function(t,e){return s*o(t,e)}))}),e.svg&&(l="Row"==t?"col":"row",e.svg.select("."+l).selectAll(".sorted").classed("selected",!1).classed("sorted",!1)),e};var g=e.updateSize;function h(t){var r=d3.mouse(e.container.node());d3.select(this).attr("fill",function(t){return d3.rgb(e.get_colour(e.get_value(t[0],t[1]))).darker(.5)}).classed("hover",!0),e.svg.select(".col").selectAll(".label").filter(function(e){return e==t[1]}).classed("hover",!0),e.svg.select(".row").selectAll(".label").filter(function(e){return e==t[0]}).classed("hover",!0),e.container.select(".inform").style("left",r[0]+10+"px").style("top",r[1]+10+"px").select(".value").html(function(){return e.get_informText(t[0],t[1])}),e.container.select(".inform").classed("hidden",!1),e.get_on_mouseover(t[0],t[1])}function m(t){d3.select(this).attr("fill",function(t){return e.get_colour(e.get_value(t[0],t[1]))}).classed("hover",!1),e.svg.selectAll(".label").classed("hover",!1),e.container.select(".inform").classed("hidden",!0),e.get_on_mouseout(t[0],t[1])}function _(t){var r;r=d3.select(this.parentNode).classed("row")?"row":"col",d3.select(this).classed("sorted")?"col"==r?e.reorder("Row","flip"):e.reorder("Col","flip"):("col"==r?e.get_on_labelClickCol(t):e.get_on_labelClickRow(t),e.updateLabelPosition()),d3.select(this.parentNode).selectAll(".label").classed("selected",!1),d3.select(this).classed("selected",!0)}function y(){if("svg"==d()&&e.canvas.classed("active"))return e.canvas.classed("active",!1),e.g.classed("active",!0),e.canvas.node().getContext("2d").clearRect(0,0,e.plotWidth(),e.plotHeight()),!!e.updateStarted||(e.updateStarted=!0,e.updateLabels().updateLabelText().updateCellColour(),e.updateStarted=!1,e.mark(e.marked.map(function(t){return"p"+t.join("_-sep-_")}),!0),e.marked=[],!1);if("canvas"==d()&&e.g.classed("active"))for(e.canvas.classed("active",!0),e.marked=e.g.selectAll(".marked").data(),e.g.classed("active",!1);e.g.node().firstChild;)e.g.node().removeChild(e.g.node().firstChild);return!0}e.updateSize=function(){g();var t=d3.max([d3.min([e.paddings().bottom-4,15]),3]),r=d3.max([d3.min([e.paddings().right-4,15]),3]),a=d3.min([e.height(),e.plotHeight()+t+e.paddings().top]),n=d3.min([e.width(),e.plotWidth()+r+e.paddings().left]);if(e.transitionDuration()>0&&!e.transitionOff){var o=d3.transition("size").duration(e.transitionDuration());e.showDendogram("Row")||e.svg.selectAll(".label_panel.row").transition(o).attr("transform","translate("+e.paddings().left+", "+e.paddings().top+")"),e.showDendogram("Col")||e.svg.selectAll(".label_panel.col").transition(o).attr("transform","translate("+e.paddings().left+", "+e.paddings().top+")"),e.axes.x_label.transition(o).attr("font-size",t).attr("x",e.plotWidth()+e.paddings().left).attr("y",a),e.axes.y_label.transition(o).attr("font-size",r).attr("x",-e.paddings().top).attr("y",n)}else e.showDendogram("Row")||e.svg.selectAll(".label_panel.row").attr("transform","translate("+e.paddings().left+", "+e.paddings().top+")"),e.showDendogram("Col")||e.svg.selectAll(".label_panel.col").attr("transform","translate("+e.paddings().left+", "+e.paddings().top+")"),e.axes.x_label.attr("font-size",t).attr("x",e.get_plotWidth()+e.paddings().left).attr("y",a),e.axes.y_label.attr("font-size",r).attr("x",-e.paddings().top).attr("y",n);return e.canvas.style("left",e.paddings().left+"px").style("top",e.paddings().top+"px").attr("width",e.plotWidth()).attr("height",e.plotHeight()),e.updateLabelPosition(),e},e.updateLabelPosition=function(){var t=e.dispColIds().length,r=e.dispRowIds().length;if(e.get_heatmapRow("__sort__"),e.get_heatmapCol("__sort__"),e.cellSize={width:e.plotWidth()/t,height:e.plotHeight()/r},e.axes.scale_x=d3.scaleLinear().domain([0,t-1]).range([0,e.plotWidth()-e.cellSize.width]),e.axes.scale_y=d3.scaleLinear().domain([0,r-1]).range([0,e.plotHeight()-e.cellSize.height]),e.transitionDuration()>0&&!e.transitionOff){var a=d3.transition("labelPosition").duration(e.transitionDuration());e.svg.select(".col").selectAll(".label").transition(a).attr("font-size",d3.min([e.cellSize.width-2,14])).attr("y",function(t){return e.axes.scale_x(e.get_heatmapCol(t)+1)}).attr("dy",-2),e.svg.select(".row").selectAll(".label").transition(a).attr("font-size",d3.min([e.cellSize.height-2,14])).attr("y",function(t){return e.axes.scale_y(e.get_heatmapRow(t)+1)}).attr("dy",-2)}else e.svg.select(".col").selectAll(".label").attr("font-size",d3.min([e.cellSize.width-2,14])).attr("y",function(t){return e.axes.scale_x(e.get_heatmapCol(t)+1)}).attr("dy",-2),e.svg.select(".row").selectAll(".label").attr("font-size",d3.min([e.cellSize.height-2,14])).attr("y",function(t){return e.axes.scale_y(e.get_heatmapRow(t)+1)}).attr("dy",-2);return e.updateCellPosition(),e.showDendogram("Col")&&e.drawDendogram("Col"),e.showDendogram("Row")&&e.drawDendogram("Row"),e},e.updateLabels=function(){var t=e.svg.select(".col").selectAll(".label").data(e.get_dispColIds(),function(t){return t});t.exit().remove();var r=e.svg.select(".row").selectAll(".label").data(e.get_dispRowIds(),function(t){return t});return r.exit().remove(),t.enter().append("text").attr("class","label plainText").attr("transform","rotate(-90)").style("text-anchor","start").attr("dx",2).merge(t).attr("id",function(t){return t.toString().replace(/[ .]/g,"_")}).on("mouseover",c).on("mouseout",u).on("click",_),r.enter().append("text").attr("class","label plainText").style("text-anchor","end").attr("dx",-2).merge(r).attr("id",function(t){return t.toString().replace(/[ .]/g,"_")}).on("mouseover",c).on("mouseout",u).on("click",_),e.updateCells(),e},e.updateLabelText=function(){if(e.transitionDuration()>0&&!e.transitionOff){var t=d3.transition("labelText").duration(e.transitionDuration());e.svg.select(".col").selectAll(".label").transition(t).text(function(t){return e.get_colLabel(t)}),e.svg.select(".row").selectAll(".label").transition(t).text(function(t){return e.get_rowLabel(t)})}else e.svg.select(".col").selectAll(".label").text(function(t){return e.get_colLabel(t)}),e.svg.select(".row").selectAll(".label").text(function(t){return e.get_rowLabel(t)});return e},e.zoom=function(t,r){var a=e.findElements(t,r);if(!(a.length<2)){var n=[],o=[];a.map(function(t){n.push(t[0]),o.push(t[1])});for(var i=[],l=[],s=0;s<n.length;s++)-1==i.indexOf(n[s])&&i.push(n[s]);for(s=0;s<o.length;s++)-1==l.indexOf(o[s])&&l.push(o[s]);return i.length>0&&e.dispRowIds(i),e.dispColIds(l),e.updateLabels(),e.updateLabelPosition(),e}},e.resetDomain=function(){return e.dispColIds(e.colIds()),e.dispRowIds(e.rowIds()),e.updateStarted=!0,e.updateLabels().updateLabelPosition().updateCellColour().updateLabelText(),e.updateStarted=!1,e},e.resetColourScale=function(){var t=e.colourDomain();2==t.length&&"number"==typeof(t[0]+t[1])?e.colourScale=d3.scaleSequential(e.get_palette).domain(t):(e.colourScale=d3.scaleOrdinal().domain(t),e.get_palette().splice?e.colourScale.range(e.get_palette()):"function"==typeof e.get_palette.range?e.colourScale.range(e.get_palette.range()):e.colourScale.range(d3.schemeSet1)),e.showLegend()&&e.legend&&e.legend.set_title({heatmap:""}).add_block(e.colourScale,"colour","heatmap")},e.on_labelClickCol(function(t){e.reorder("Row",function(r,a){return e.get_value(a,t)-e.get_value(r,t)}),e.dendogramRow&&e.dendogramRow.remove()}),e.on_labelClickRow(function(t){e.reorder("Col",function(r,a){return e.get_value(t,a)-e.get_value(t,r)}),e.dendogramCol&&e.dendogramCol.remove()}),e.updateCellColour=function(){return y()?("svg"==d()?(e.transitionDuration()>0&&!e.transitionOff?e.g.selectAll(".data_element").transition("cellColour").duration(e.transitionDuration()).attr("fill",function(t){return e.get_colour(e.get_value(t[0],t[1]))}):e.g.selectAll(".data_element").attr("fill",function(t){return e.get_colour(e.get_value(t[0],t[1]))}),e.svg.selectAll(".sorted").classed("selected",!1).classed("sorted",!1),e.get_showValue()&&e.updateTextValues()):e.updateStarted||e.updateCanvas(),e.svg.selectAll(".sorted").filter(function(t){return!function(t){var r,a=d3.select(t).datum(),n=!0,o=1;if(d3.select(t.parentNode).classed("row")){for(r=e.dispColIds();n&&o<r.length;)e.get_value(a,r[o])>e.get_value(a,r[o-1])&&(n=!1),o++;if(n)return n;for(o=1,n=!0;n&&o<r.length;)e.get_value(a,r[o])<e.get_value(a,r[o-1])&&(n=!1),o++;return n}for(r=e.dispRowIds();n&&o<r.length;)e.get_value(r[o],a)>e.get_value(r[o-1],a)&&(n=!1),o++;if(n)return n;for(o=1,n=!0;n&&o<r.length;)e.get_value(r[o],a)<e.get_value(r[o-1],a)&&(n=!1),o++;return n}(this)}).classed("sorted",!1).classed("selected",!1),e):e},e.updateCells=function(){if(!y())return e;var t=e.get_marked().length;if("svg"==d()){var r=e.g.selectAll(".data_row").data(e.get_dispRowIds(),function(t){return t});r.exit().remove(),r.enter().append("g").attr("class","data_row");var a=e.g.selectAll(".data_row").selectAll(".data_element").data(function(t){return e.get_dispColIds().map(function(e){return[t,e]})},function(t){return t});a.exit().remove(),a.enter().append("rect").attr("class","data_element").merge(a).attr("id",function(t){return"p"+(t[0]+"_-sep-_"+t[1]).replace(/[ .]/g,"_")}).attr("rowId",function(t){return t[0]}).attr("colId",function(t){return t[1]}).on("mouseover",h).on("mouseout",m).on("click",function(t){e.get_on_click.apply(this,[t[0],t[1]])}),e.get_showValue()&&e.updateTexts()}else{for(var n=e.dispRowIds(),o=e.dispColIds(),i=0;i<e.marked.length;)-1==n.indexOf(e.marked[i][0])||-1==o.indexOf(e.marked[i][1])?e.marked.splice(i,1):i++;e.updateStarted||e.updateCanvas()}var l=e.get_marked().length;return t>l&&e.on_marked(),0==l&&e.g.selectAll(".data_element").attr("opacity",1),e},e.updateCellPosition=function(){return y()?("svg"==d()?(e.transitionDuration()>0&&!e.transitionOff?e.g.selectAll(".data_element").transition("cellPosition").duration(e.transitionDuration()).attr("x",function(t){return e.axes.scale_x(e.get_heatmapCol(t[1]))}).attr("width",e.cellSize.width).attr("height",e.cellSize.height).attr("y",function(t){return e.axes.scale_y(e.get_heatmapRow(t[0]))}):e.g.selectAll(".data_element").attr("x",function(t){return e.axes.scale_x(e.get_heatmapCol(t[1]))}).attr("width",e.cellSize.width).attr("height",e.cellSize.height).attr("y",function(t){return e.axes.scale_y(e.get_heatmapRow(t[0]))}),e.get_showValue()&&e.updateTextPosition()):e.updateCanvas(),e):e},e.cluster=function(t,a){if("Row"!=t&&"Col"!=t)throw"Error in 'cluster': type "+t+" cannot be recognised. Please, use either 'Row' or 'Col'";e.showDendogram(t,!0),void 0===e["dendogram"+t]&&(e["dendogram"+t]=z(e),"Row"==t?e["dendogram"+t].orientation("vertical"):e["dendogram"+t].orientation("horizontal")),e["dendogram"+t].elementIds(function(){return e["disp"+t+"Ids"]()}),void 0===a&&(a="Row"==t?e.dispColIds():e.dispRowIds()),"Row"==t?e["dendogram"+t].data(r(function(t){return a.map(function(r){return e.get_value(t,r)})})):e["dendogram"+t].data(r(function(t){return a.map(function(r){return e.get_value(r,t)})})),e["dendogram"+t].distance(function(r,a){return e["get_cluster"+t+"Metric"](r,a)}).cluster();var n=e["dendogram"+t].clusters.val_inds;return e.reorder(t,function(t,e){return n.indexOf(t)-n.indexOf(e)}),e},e.drawDendogram=function(t){return void 0===e["dendogram"+t]?e:(void 0===e["dendogram"+t].g?e["dendogram"+t].place():e["dendogram"+t].g.selectAll("line").remove(),e["dendogram"+t].draw(),e)},e.updateTexts=function(){var t=e.g.selectAll(".text_row").data(e.get_dispRowIds(),function(t){return t});t.exit().remove(),t.enter().append("g").attr("class","text_row");var r=e.g.selectAll(".text_row").selectAll(".tval").data(function(t){return e.get_dispColIds().map(function(e){return[t,e]})},function(t){return t});return r.exit().remove(),r.enter().append("text").attr("class","tval").attr("text-anchor","middle").on("click",function(t){var r=e.svg.selectAll("#p"+(t[0]+"_-sep-_"+t[1]).replace(/[ .]/g,"_")).node();e.get_on_click.apply(r,[t[0],t[1]])}).on("mouseover",function(t){var r=e.svg.selectAll("#p"+(t[0]+"_-sep-_"+t[1]).replace(/[ .]/g,"_")).node();h.apply(r,[t])}).on("mouseout",function(t){var r=e.svg.selectAll("#p"+(t[0]+"_-sep-_"+t[1]).replace(/[ .]/g,"_")).node();m.apply(r,[t])}),e},e.updateTextPosition=function(){return e.transitionDuration()>0&&!e.transitionOff?e.g.selectAll(".tval").transition("textPosition").duration(e.transitionDuration()).attr("x",function(t){return e.axes.scale_x(e.get_heatmapCol(t[1]))+e.cellSize.width/2}).attr("font-size",.4*e.cellSize.height).attr("y",function(t){return e.axes.scale_y(e.get_heatmapRow(t[0]))+.75*e.cellSize.height}):e.g.selectAll(".tval").attr("x",function(t){return e.axes.scale_x(e.get_heatmapCol(t[1]))+e.cellSize.width/2}).attr("font-size",.4*e.cellSize.height).attr("y",function(t){return e.axes.scale_y(e.get_heatmapRow(t[0]))+.75*e.cellSize.height}),e},e.updateTextValues=function(){return e.transitionDuration()>0&&!e.transitionOff?e.g.selectAll(".tval").transition("textValues").duration(e.transitionDuration()).attr("fill",t=>e.get_valueTextColour(t[0],t[1])).text(function(t){var r=e.get_value(t[0],t[1]);return r.toFixed?r.toFixed(1):"NA"}):e.g.selectAll(".tval").attr("fill",t=>e.get_valueTextColour(t[0],t[1])).text(function(t){var r=e.get_value(t[0],t[1]);return r.toFixed?r.toFixed(1):"NA"}),e},e.updateCanvas=function(){var t=e.canvas.node().getContext("2d");t.clearRect(0,0,e.plotWidth(),e.plotHeight());var r,a,n=e.dispRowIds(),o=e.dispColIds(),i=o.length,l=n.length,s=document.createElement("canvas");s.width=i,s.height=l;var d=new ImageData(i,l);if(0==e.marked.length)for(var c=0;c<l;c++)for(var u=0;u<i;u++)r=d3.rgb(e.get_colour(e.get_value(n[c],o[u]))),a=e.get_heatmapRow(n[c])*i*4+4*e.get_heatmapCol(o[u]),d.data[a]=r.r,d.data[a+1]=r.g,d.data[a+2]=r.b,d.data[a+3]=255;else{for(c=0;c<l;c++)for(u=0;u<i;u++)r=d3.rgb(e.get_colour(e.get_value(n[c],o[u]))),a=e.get_heatmapRow(n[c])*i*4+4*e.get_heatmapCol(o[u]),d.data[a]=r.r/4,d.data[a+1]=r.g/4,d.data[a+2]=r.b/4,d.data[a+3]=55;for(c=0;c<e.marked.length;c++)a=e.get_heatmapRow(e.marked[c][0])*i*4+4*e.get_heatmapCol(e.marked[c][1]),d.data[a]*=4,d.data[a+1]*=4,d.data[a+2]*=4,d.data[a+3]=255}if(s.getContext("2d").putImageData(d,0,0),t.imageSmoothingEnabled=!1,t.drawImage(s,0,0,i,l,0,0,e.plotWidth(),e.plotHeight()),e.showValue())for(t.font=.4*e.cellSize.height+"px arial",t.textAlign="center",c=0;c<l;c++)for(u=0;u<i;u++){let r=e.axes.scale_x(e.get_heatmapCol(c))+e.cellSize.width/2,a=e.axes.scale_y(e.get_heatmapRow(u))+.75*e.cellSize.height,n=e.get_value(u,c),o=d3.hsl(e.get_colour(e.get_value(u,c)));o.h=(o.h+180)%360,o.s=Math.floor(100*Math.sqrt(o.s)),o.l=Math.floor(100*(1-o.l)),t.fillStyle="hsl("+o.h+","+o.s+"%,"+o.l+"%)",n=n.toFixed?n.toFixed(1):"NA",t.fillText(n,r,a,e.cellSize.width)}},e.get_elements=function(t){return 2==t.length&&t[0].substr&&(t=[t]),t=t.map(function(t){return"p"+t.join("_-sep-_")}),"svg"==d()?t.length>0?e.svg.selectAll("#"+i(t.join(",#").replace(/[ .]/g,"_"))):e.svg.selectAll("______"):t},e.get_marked=function(){if("svg"==d()){var t=[];return e.svg.selectAll(".marked").each(function(){t.push(d3.select(this).datum())}),t}return e.marked};var v=e.mark;return e.mark=function(t,r){if("svg"==d())v(t,r);else if("__clear__"==t)e.marked=[];else{t.length&&t[0].substr&&(t=t.map(function(t){return t.substr(1).split("_-sep-_")}));for(var a,n=e.marked.map(function(t){return t.join("_")}),o=0;o<t.length;o++)-1==(a=n.indexOf(t[o].join("_")))?e.marked.push(t[o]):(e.marked.splice(a,1),n.splice(a,1))}return"canvas"==d()&&e.updateCanvas(),r||e.on_marked(),e},e.panMove=function(t){var r=[t[0]-e.pan("down")[0],t[1]-e.pan("down")[1]],a=Math.floor(Math.abs(r[1]/e.cellSize.height)),n=Math.floor(Math.abs(r[0]/e.cellSize.width));e.pan("down")[0]+=Math.sign(r[0])*n*e.cellSize.width,e.pan("down")[1]+=Math.sign(r[1])*a*e.cellSize.height,e.dispColIds(p(-Math.sign(r[0])*n,"right")),e.dispColIds(p(Math.sign(r[0])*n,"left")),e.dispRowIds(p(Math.sign(r[1])*a,"top")),e.dispRowIds(p(-Math.sign(r[1])*a,"bottom")),Math.abs(a)+Math.abs(n)>0&&(e.updateStarted=!0,e.updateLabels(),e.updateLabelPosition(),e.updateCellColour(),e.updateLabelText(),e.updateStarted=!1)},e.update=function(){return e.updateTitle(),e.resetColourScale(),e.dispColIds(function(){return e.colIds()}),e.dispRowIds(function(){return e.rowIds()}),e.axes.x_label.text(e.get_colTitle()),e.axes.y_label.text(e.get_rowTitle()),e.updateStarted=!0,e.updateLabels(),e.clusterRows()&&e.cluster("Row"),e.clusterCols()&&e.cluster("Col"),e.updateSize().updateLabelText().updateCellColour(),e.updateStarted=!1,e},e},t.cache=r,t.separateBy=function t(e,r,a){if("object"!=typeof e)throw"Error in function 'separateBy': first argument is not an object";if(a=void 0===e.length?"obj":"arr","number"!=typeof r&&"string"!=typeof r||(r=[r]),void 0===r.length)throw"Error in function 'separateBy': "+r.toString()+" is not a property name";if(0==r.length)return e;for(var n,o,i={},l=Object.keys(e),s=!1,d=0;!s&&d<l.length;)"undefined"!==e[l[d]][r[0]]&&(s=!0),d++;if(!s)return r.shift(),t(e,r);for(d=0;d<l.length;d++)void 0!==(n=e[l[d]][r[0]])&&(o={},Object.assign(o,e[l[d]]),delete o[r[0]],void 0===i[n]&&(i[n]="obj"==a?{}:[]),"obj"==a&&(i[n][l[d]]={}),"obj"==a?i[n][l[d]]=o:i[n].push(o));if("arr"==a){for(var c=!0,u=(d=0,Object.keys(i));c&&d<u.length;)c=1==i[u[d]].length,d++;var p;if(c)for(d=0;d<u.length;d++)p={},Object.assign(p,i[u[d]][0]),i[u[d]]={},Object.assign(i[u[d]],p)}r.shift();for(d=0;d<u.length;d++)i[u[d]]=t(i[u[d]],r.slice());return i},t.getEuclideanDistance=a,t.add_click_listener=n,t.pearsonCorr=function(t,e){for(var r=0,a=0,n=0;n<t.length;n++)r+=t[n],a+=e[n];var o=r/t.length,i=a/e.length,l=0,s=0,d=0;for(n=0;n<t.length;n++)l+=(t[n]-o)*(e[n]-i),s+=(t[n]-o)*(t[n]-o),d+=(e[n]-i)*(e[n]-i);return l/Math.sqrt(s*d)},t.get_symbolSize=o,t.escapeRegExp=i,t.check=s,t.call_pacer=l,t.isNaN=d,t.random=function(t,e){return t=t||1,e=e||0,Math.seed=(9301*Math.seed+49297)%233280,e+Math.seed/233280*(t-e)},t.colourSlider=function(){var t=f().add_property("straightColourScale").add_property("midpoint",void 0).add_property("slopeWidth",void 0).add_property("linkedChart",void 0,function(t){if("string"==typeof t){var e=t.split(".");t=window[e[0]];for(var r=1;r<e.length;r++)t=t[e[r]]}return t}).add_property("on_drag",function(){}).add_property("on_change",function(){}).paddings({top:20,right:10,bottom:5,left:10}).height(80).transitionDuration(0);t.showPanel(!1),t.straightColourScale(d3.scaleLinear().range(["white","darkblue"])),t.setStraightColourScale=function(){t.linkedChart()&&t.linkedChart().colourScale&&(t.straightColourScale(t.linkedChart().colourScale),t.linkedChart().updateElementStyle&&t.on_drag(t.linkedChart().updateElementStyle),t.linkedChart().updateCellColour&&t.on_drag(t.linkedChart().updateCellColour))};var e=function(){var e=d3.min(t.get_straightColourScale.domain()),r=d3.max(t.get_straightColourScale.domain());t.get_midpoint()<e&&t.midpoint(e),t.get_midpoint()>r&&t.midpoint(r),t.midpoint()-e<t.slopeWidth()/2&&t.slopeWidth(2*(t.midpoint()-e)),r-t.midpoint()<t.slopeWidth()/2&&t.slopeWidth(2*(r-t.midpoint()))},r=t.put_static_content;t.put_static_content=function(a){r(a);var n=t.svg.append("g").attr("class","sigmoidColorSlider").attr("transform","translate("+t.paddings().left+", "+t.paddings().top+")");t.axis=n.append("g").attr("class","axis");var o=n.append("defs");t.gradient=o.append("linearGradient").attr("id","scaleGradient"+Math.random().toString(36).substring(2,6)).attr("x1","0%").attr("y1","0%").attr("x2","100%").attr("y2","0%"),t.gradient.selectAll("stop").data(d3.range(100)).enter().append("stop").attr("offset",function(t){return t+"%"});var i=t.gradient.attr("id");t.colorBar=n.append("rect").attr("x","0").attr("y","5").attr("height",20).attr("fill","url(#"+i+")").style("stroke","black").style("stroke-width","1"),o.append("path").attr("id","mainMarker").attr("d","M 0 0 L 8 5 L 8 25 L -8 25 L -8 5 Z").style("fill","gray").style("stroke","black"),o.append("path").attr("id","rightMarker").attr("d","M 0 0 L 5 5 L 5 15 L 0 15 Z").style("fill","lightgray").style("stroke","black"),o.append("path").attr("id","leftMarker").attr("d","M 0 0 L -5 5 L -5 15 L 0 15 Z").style("fill","lightgray").style("stroke","black"),t.mainMarker=n.append("use").attr("xlink:href","#mainMarker").attr("y",28).call(d3.drag().on("drag",function(){t.midpoint(t.pos_scale.invert(t.pos_scale(t.get_midpoint())+d3.event.dx)),e(),t.get_on_drag(),t.update()}).on("end",function(){t.get_on_change()})),t.rightMarker=n.append("use").attr("xlink:href","#rightMarker").attr("y",30).call(d3.drag().on("drag",function(){t.slopeWidth(t.pos_scale.invert(t.pos_scale(t.slopeWidth())+2*d3.event.dx)),e(),t.update(),t.get_on_drag()}).on("end",function(){t.get_on_change()})),t.leftMarker=n.append("use").attr("xlink:href","#leftMarker").attr("y",30).call(d3.drag().on("drag",function(){t.slopeWidth(t.pos_scale.invert(t.pos_scale(t.slopeWidth())-2*d3.event.dx)),e(),t.update(),t.get_on_drag()}).on("end",function(){t.get_on_change()}))};var a=t.update;return t.update=function(){a(),t.setStraightColourScale(),void 0==t.get_straightColourScale.domain&&(t.get_straightColourScale.domain=function(){return[0,1]});for(var e=t.get_straightColourScale.domain(),r=[],n=[],o=!0,i=0;i<e.length;i++)r.push(100*i/(e.length-1)),n.push(i*t.plotWidth()/(e.length-1)),o="number"==typeof e[i]&&o;if(!o)return t.colourScale=t.get_straightColourScale,t;var l,s,d,c,u,p,f,g=d3.scaleLinear().domain(r).range(e);void 0==t.get_midpoint()&&t.midpoint(g(50)),void 0==t.get_slopeWidth()&&t.slopeWidth(Math.abs(g(70)-g(0))),t.pos_scale=d3.scaleLinear().range(n).domain(e),d3.axisTop().scale(t.pos_scale)(t.axis),t.colorBar.attr("width",t.get_plotWidth()),t.the_sigmoid=(l=t.get_midpoint(),s=1.38/t.slopeWidth(),d=d3.min(t.get_straightColourScale.domain()),c=d3.max(t.get_straightColourScale.domain()),u=D(d,l,s),p=D(c,l,s),f=Math.min(u,p),function(t){return(D(t,l,s)-f)/Math.abs(p-u)}),t.gradient.selectAll("stop").data(d3.range(100)).style("stop-color",function(e){return t.get_straightColourScale(g(100*t.the_sigmoid(g(e))))}),t.colourScale=function(e){return t.get_straightColourScale(g(100*t.the_sigmoid(e)))},t.linkedChart()&&t.linkedChart().colour&&t.linkedChart().colour(function(e){return t.linkedChart().colourValue&&(e=t.linkedChart().get_colourValue(e)),t.colourScale(e)}),t.mainMarker.attr("x",t.pos_scale(t.get_midpoint())),t.rightMarker.attr("x",t.pos_scale(t.get_midpoint()+t.slopeWidth()/2)),t.leftMarker.attr("x",t.pos_scale(t.get_midpoint()-t.slopeWidth()/2))},t},t.table=function(){var t=f().add_property("record",{});t.showPanel(!1);var e=t.put_static_content;t.put_static_content=function(r){e(r),t.svg.remove(),t.table=t.container.append("table")},t.updateSize=function(){return t.table.style("width",t.width()),t.table.selectAll("td").style("height",t.height()/Object.keys(t.record()).length),t};var r=t.update;return t.update=function(){r();var e=t.table.selectAll("tr").data(Object.keys(t.record()));return e.exit().remove(),e.enter().append("tr").merge(e).html(function(e){return"<td>"+e+"</td><td>"+t.get_record()[e]+"</td>"}),t.table.selectAll("td").style("border-bottom","1px solid #ddd"),t},t},t.html=R,t.input=function(){var t=R().add_property("type","text",function(t){if(-1==["text","radio","checkbox","button","range"].indexOf(t))throw"Error in 'typeCheck' for property 'type': the supported types are 'text', 'radio', 'checkbox', 'button', 'range'.";return t}).add_property("label",t=>t,s("array_fun","label")).add_property("nelements",1,s("number_nonneg","nelements")).add_property("elementIds",[0],s("array","elementIds")).add_property("value",function(){},s("array_fun","value")).add_property("min",0,s("array_fun","min")).add_property("max",100,s("array_fun","max")).add_property("step",1,s("array_fun","step")).add_property("fontSize",17,s("number_nonneg","fontSize")).add_property("ncols",1,s("number_nonneg","ncols")).add_property("nrows",function(){return t.nelements()},s("number_nonneg","nrows")).add_property("on_change",function(){});t.wrapSetter("nelements",function(e){return function(){return t.get_elementIds=function(){return d3.range(e())},e.apply(t,arguments)}}),t.wrapSetter("elementIds",function(e){return function(){return t.get_nelements=function(){return e().length},e.apply(t,arguments)}}),t.wrapSetter("nrows",function(e){return function(){return t.get_ncols=function(){return Math.ceil(t.nelements()/e())},e.apply(t,arguments)}}),t.wrapSetter("ncols",function(e){return function(){return t.get_nrows=function(){return Math.ceil(t.nelements()/e())},e.apply(t,arguments)}}),t.value(e=>{switch(t.type()){case"text":return"";case"button":return e;case"range":case"radio":return;case"checkbox":return!1}}),t.name="n"+Math.random().toString(36).substring(5),t.width(200);var e=t.put_static_content;return t.put_static_content=function(r){return e(r),t.container.style("display","grid").style("grid-template-columns","repeat(auto-fit, fit-content(500px) 1fr fit-content(20px))").style("column-gap","9px").style("align-items","center").append("p").style("grid-row",1).style("grid-column","1/"+(3*t.ncols()+1)).style("text-align","center").style("font-weight","bold").attr("id","title"),t},t.updateTitle=function(){return t.container.select("#title").text(t.title()),t},t.updateElements=function(){var e=t.container.selectAll("input").data(t.elementIds()),r=t.container.selectAll("label").data(t.elementIds());if(e.enter().append("input").attr("type",t.type()).attr("name",t.name).style("grid-row",(e,r)=>r%t.nrows()+2).style("grid-column",(e,r)=>3*Math.floor(r/t.nrows())+("text"==t.type()||"range"==t.type()?2:1)).attr("id",e=>t.name+"_in_"+e).property("value",e=>"radio"==t.type()?e:void 0).style("width","text"==t.type()||"range"==t.type()?"100%":void 0).on("button"==t.type()?"click":"change",function(){t.get_on_change(function(e){if("button"==t.type()||"radio"==t.type())return e.value;if("checkbox"==t.type())return d3.select(e.parentNode).selectAll("input").nodes().map(t=>t.checked);var r={};return"range"!=t.type()&&"text"!=t.type()||d3.select(e.parentNode).selectAll("input").nodes().forEach(function(e){r[e.id.substring(t.name.length+1)]=+e.value?+e.value:e.value}),r}(this))}),e.exit().remove(),"button"!=t.type()?(r.exit().remove(),r.enter().append("label").attr("for",e=>t.name+"_in_"+e).style("grid-row",(e,r)=>r%t.nrows()+2).style("grid-column",(e,r)=>3*Math.floor(r/t.nrows())+("text"==t.type()||"range"==t.type()?1:2)).merge(r).text(e=>t.get_label(e)).style("font-size",t.fontSize())):t.container.selectAll("input").style("font-size",t.fontSize()).property("value",e=>t.get_label(e)),"range"==t.type()){var a=t.container.selectAll(".currentValue").data(t.elementIds());a.enter().append("p").attr("class","currentValue").attr("id",e=>t.name+"_in_"+e).style("grid-row",(e,r)=>r%t.nrows()+2).style("grid-column",(e,r)=>3*Math.floor(r/t.nrows())+3).style("margin-top",0).style("margin-bottom",0),a.exit().remove(),t.container.selectAll("input").on("input",function(e){t.container.select(".currentValue#"+this.id).text(this.value)})}else t.container.selectAll(".currentValue").remove();return t},t.updateState=function(){return"radio"==t.type()&&t.container.selectAll("input").filter(function(){return this.id==t.name+"_in_"+t.value()}).attr("checked",!0),"checkbox"==t.type()&&t.container.selectAll("input").property("checked",e=>!!t.get_value(e)),"text"==t.type()&&t.container.selectAll("input").property("value",e=>t.get_value(e)),"range"==t.type()&&(t.container.selectAll("input").attr("max",e=>t.get_max(e)).attr("min",e=>t.get_min(e)).attr("step",e=>t.get_step(e)).nodes().forEach((e,r)=>{e.value=t.get_value(t.elementIds()[r])}),t.container.selectAll(".currentValue").text(e=>t.get_value(e))),t},t.update=function(){return t.inherited_update(),t.updateElements(),t.updateState(),t},t},t.image=function(){var t=f().add_property("src",void 0,function(t){if(void 0!==t&&"string"!=typeof t)throw"Error in 'typeCheck' for property 'src': must be a string or undefined.";return t}).showPanel(!1).showLegend(!1).paddings({top:35,right:10,bottom:10,left:10}),e=t.put_static_content;t.put_static_content=function(r){e(r),t.svg.select(".plotArea").append("image")},t.updateElements=function(){return t.svg.select("image").attr("href",t.src()),t};var r=t.updateSize;t.updateSize=function(){return r(),t.svg.select("image").attr("width",t.plotWidth()).attr("height",t.plotHeight()),t};var a=t.update;return t.update=function(){return a(),t.updateElements(),t},t},t.barchart=L,t.dendogram=z,t.beeswarm=C,Object.defineProperty(t,"__esModule",{value:!0})});
```
