## Supplemental Examples for "R/LinkedCharts: A novel approach for simple but powerful interactive data analysis": prism.js

```
/* PrismJS 1.15.0
https://prismjs.com/download.html#themes=prism-coy&languages=markup+css+clike+javascript+r&plugins=line-numbers */
var _self="undefined"!=typeof window?window:"undefined"!=typeof WorkerGlobalScope&&self instanceof WorkerGlobalScope?self:{},Prism=function(){var e=/\blang(?:uage)?-([\w-]+)\b/i,t=0,n=_self.Prism={manual:_self.Prism&&_self.Prism.manual,disableWorkerMessageHandler:_self.Prism&&_self.Prism.disableWorkerMessageHandler,util:{encode:function(e){return e instanceof a?new a(e.type,n.util.encode(e.content),e.alias):"Array"===n.util.type(e)?e.map(n.util.encode):e.replace(/&/g,"&amp;").replace(/</g,"&lt;").replace(/\u00a0/g," ")},type:function(e){return Object.prototype.toString.call(e).slice(8,-1)},objId:function(e){return e.__id||Object.defineProperty(e,"__id",{value:++t}),e.__id},clone:function(e,t){var a=n.util.type(e);switch(t=t||{},a){case"Object":if(t[n.util.objId(e)])return t[n.util.objId(e)];var r={};t[n.util.objId(e)]=r;for(var l in e)e.hasOwnProperty(l)&&(r[l]=n.util.clone(e[l],t));return r;case"Array":if(t[n.util.objId(e)])return t[n.util.objId(e)];var r=[];return t[n.util.objId(e)]=r,e.forEach(function(e,a){r[a]=n.util.clone(e,t)}),r}return e}},languages:{extend:function(e,t){var a=n.util.clone(n.languages[e]);for(var r in t)a[r]=t[r];return a},insertBefore:function(e,t,a,r){r=r||n.languages;var l=r[e],i={};for(var o in l)if(l.hasOwnProperty(o)){if(o==t)for(var s in a)a.hasOwnProperty(s)&&(i[s]=a[s]);a.hasOwnProperty(o)||(i[o]=l[o])}var u=r[e];return r[e]=i,n.languages.DFS(n.languages,function(t,n){n===u&&t!=e&&(this[t]=i)}),i},DFS:function(e,t,a,r){r=r||{};for(var l in e)e.hasOwnProperty(l)&&(t.call(e,l,e[l],a||l),"Object"!==n.util.type(e[l])||r[n.util.objId(e[l])]?"Array"!==n.util.type(e[l])||r[n.util.objId(e[l])]||(r[n.util.objId(e[l])]=!0,n.languages.DFS(e[l],t,l,r)):(r[n.util.objId(e[l])]=!0,n.languages.DFS(e[l],t,null,r)))}},plugins:{},highlightAll:function(e,t){n.highlightAllUnder(document,e,t)},highlightAllUnder:function(e,t,a){var r={callback:a,selector:'code[class*="language-"], [class*="language-"] code, code[class*="lang-"], [class*="lang-"] code'};n.hooks.run("before-highlightall",r);for(var l,i=r.elements||e.querySelectorAll(r.selector),o=0;l=i[o++];)n.highlightElement(l,t===!0,r.callback)},highlightElement:function(t,a,r){for(var l,i,o=t;o&&!e.test(o.className);)o=o.parentNode;o&&(l=(o.className.match(e)||[,""])[1].toLowerCase(),i=n.languages[l]),t.className=t.className.replace(e,"").replace(/\s+/g," ")+" language-"+l,t.parentNode&&(o=t.parentNode,/pre/i.test(o.nodeName)&&(o.className=o.className.replace(e,"").replace(/\s+/g," ")+" language-"+l));var s=t.textContent,u={element:t,language:l,grammar:i,code:s};if(n.hooks.run("before-sanity-check",u),!u.code||!u.grammar)return u.code&&(n.hooks.run("before-highlight",u),u.element.textContent=u.code,n.hooks.run("after-highlight",u)),n.hooks.run("complete",u),void 0;if(n.hooks.run("before-highlight",u),a&&_self.Worker){var g=new Worker(n.filename);g.onmessage=function(e){u.highlightedCode=e.data,n.hooks.run("before-insert",u),u.element.innerHTML=u.highlightedCode,n.hooks.run("after-highlight",u),n.hooks.run("complete",u),r&&r.call(u.element)},g.postMessage(JSON.stringify({language:u.language,code:u.code,immediateClose:!0}))}else u.highlightedCode=n.highlight(u.code,u.grammar,u.language),n.hooks.run("before-insert",u),u.element.innerHTML=u.highlightedCode,n.hooks.run("after-highlight",u),n.hooks.run("complete",u),r&&r.call(t)},highlight:function(e,t,r){var l={code:e,grammar:t,language:r};return n.hooks.run("before-tokenize",l),l.tokens=n.tokenize(l.code,l.grammar),n.hooks.run("after-tokenize",l),a.stringify(n.util.encode(l.tokens),l.language)},matchGrammar:function(e,t,a,r,l,i,o){var s=n.Token;for(var u in a)if(a.hasOwnProperty(u)&&a[u]){if(u==o)return;var g=a[u];g="Array"===n.util.type(g)?g:[g];for(var c=0;c<g.length;++c){var h=g[c],f=h.inside,d=!!h.lookbehind,m=!!h.greedy,p=0,y=h.alias;if(m&&!h.pattern.global){var v=h.pattern.toString().match(/[imuy]*$/)[0];h.pattern=RegExp(h.pattern.source,v+"g")}h=h.pattern||h;for(var b=r,k=l;b<t.length;k+=t[b].length,++b){var w=t[b];if(t.length>e.length)return;if(!(w instanceof s)){if(m&&b!=t.length-1){h.lastIndex=k;var _=h.exec(e);if(!_)break;for(var j=_.index+(d?_[1].length:0),P=_.index+_[0].length,A=b,x=k,O=t.length;O>A&&(P>x||!t[A].type&&!t[A-1].greedy);++A)x+=t[A].length,j>=x&&(++b,k=x);if(t[b]instanceof s)continue;I=A-b,w=e.slice(k,x),_.index-=k}else{h.lastIndex=0;var _=h.exec(w),I=1}if(_){d&&(p=_[1]?_[1].length:0);var j=_.index+p,_=_[0].slice(p),P=j+_.length,N=w.slice(0,j),S=w.slice(P),C=[b,I];N&&(++b,k+=N.length,C.push(N));var E=new s(u,f?n.tokenize(_,f):_,y,_,m);if(C.push(E),S&&C.push(S),Array.prototype.splice.apply(t,C),1!=I&&n.matchGrammar(e,t,a,b,k,!0,u),i)break}else if(i)break}}}}},tokenize:function(e,t){var a=[e],r=t.rest;if(r){for(var l in r)t[l]=r[l];delete t.rest}return n.matchGrammar(e,a,t,0,0,!1),a},hooks:{all:{},add:function(e,t){var a=n.hooks.all;a[e]=a[e]||[],a[e].push(t)},run:function(e,t){var a=n.hooks.all[e];if(a&&a.length)for(var r,l=0;r=a[l++];)r(t)}}},a=n.Token=function(e,t,n,a,r){this.type=e,this.content=t,this.alias=n,this.length=0|(a||"").length,this.greedy=!!r};if(a.stringify=function(e,t,r){if("string"==typeof e)return e;if("Array"===n.util.type(e))return e.map(function(n){return a.stringify(n,t,e)}).join("");var l={type:e.type,content:a.stringify(e.content,t,r),tag:"span",classes:["token",e.type],attributes:{},language:t,parent:r};if(e.alias){var i="Array"===n.util.type(e.alias)?e.alias:[e.alias];Array.prototype.push.apply(l.classes,i)}n.hooks.run("wrap",l);var o=Object.keys(l.attributes).map(function(e){return e+'="'+(l.attributes[e]||"").replace(/"/g,"&quot;")+'"'}).join(" ");return"<"+l.tag+' class="'+l.classes.join(" ")+'"'+(o?" "+o:"")+">"+l.content+"</"+l.tag+">"},!_self.document)return _self.addEventListener?(n.disableWorkerMessageHandler||_self.addEventListener("message",function(e){var t=JSON.parse(e.data),a=t.language,r=t.code,l=t.immediateClose;_self.postMessage(n.highlight(r,n.languages[a],a)),l&&_self.close()},!1),_self.Prism):_self.Prism;var r=document.currentScript||[].slice.call(document.getElementsByTagName("script")).pop();return r&&(n.filename=r.src,n.manual||r.hasAttribute("data-manual")||("loading"!==document.readyState?window.requestAnimationFrame?window.requestAnimationFrame(n.highlightAll):window.setTimeout(n.highlightAll,16):document.addEventListener("DOMContentLoaded",n.highlightAll))),_self.Prism}();"undefined"!=typeof module&&module.exports&&(module.exports=Prism),"undefined"!=typeof global&&(global.Prism=Prism);
Prism.languages.markup={comment:/<!--[\s\S]*?-->/,prolog:/<\?[\s\S]+?\?>/,doctype:/<!DOCTYPE[\s\S]+?>/i,cdata:/<!\[CDATA\[[\s\S]*?]]>/i,tag:{pattern:/<\/?(?!\d)[^\s>\/=$<%]+(?:\s+[^\s>\/=]+(?:=(?:("|')(?:\\[\s\S]|(?!\1)[^\\])*\1|[^\s'">=]+))?)*\s*\/?>/i,greedy:!0,inside:{tag:{pattern:/^<\/?[^\s>\/]+/i,inside:{punctuation:/^<\/?/,namespace:/^[^\s>\/:]+:/}},"attr-value":{pattern:/=(?:("|')(?:\\[\s\S]|(?!\1)[^\\])*\1|[^\s'">=]+)/i,inside:{punctuation:[/^=/,{pattern:/(^|[^\\])["']/,lookbehind:!0}]}},punctuation:/\/?>/,"attr-name":{pattern:/[^\s>\/]+/,inside:{namespace:/^[^\s>\/:]+:/}}}},entity:/&#?[\da-z]{1,8};/i},Prism.languages.markup.tag.inside["attr-value"].inside.entity=Prism.languages.markup.entity,Prism.hooks.add("wrap",function(a){"entity"===a.type&&(a.attributes.title=a.content.replace(/&amp;/,"&"))}),Prism.languages.xml=Prism.languages.markup,Prism.languages.html=Prism.languages.markup,Prism.languages.mathml=Prism.languages.markup,Prism.languages.svg=Prism.languages.markup;
Prism.languages.css={comment:/\/\*[\s\S]*?\*\//,atrule:{pattern:/@[\w-]+?.*?(?:;|(?=\s*\{))/i,inside:{rule:/@[\w-]+/}},url:/url\((?:(["'])(?:\\(?:\r\n|[\s\S])|(?!\1)[^\\\r\n])*\1|.*?)\)/i,selector:/[^{}\s][^{};]*?(?=\s*\{)/,string:{pattern:/("|')(?:\\(?:\r\n|[\s\S])|(?!\1)[^\\\r\n])*\1/,greedy:!0},property:/[-_a-z\xA0-\uFFFF][-\w\xA0-\uFFFF]*(?=\s*:)/i,important:/!important\b/i,"function":/[-a-z0-9]+(?=\()/i,punctuation:/[(){};:,]/},Prism.languages.css.atrule.inside.rest=Prism.languages.css,Prism.languages.markup&&(Prism.languages.insertBefore("markup","tag",{style:{pattern:/(<style[\s\S]*?>)[\s\S]*?(?=<\/style>)/i,lookbehind:!0,inside:Prism.languages.css,alias:"language-css",greedy:!0}}),Prism.languages.insertBefore("inside","attr-value",{"style-attr":{pattern:/\s*style=("|')(?:\\[\s\S]|(?!\1)[^\\])*\1/i,inside:{"attr-name":{pattern:/^\s*style/i,inside:Prism.languages.markup.tag.inside},punctuation:/^\s*=\s*['"]|['"]\s*$/,"attr-value":{pattern:/.+/i,inside:Prism.languages.css}},alias:"language-css"}},Prism.languages.markup.tag));
Prism.languages.clike={comment:[{pattern:/(^|[^\\])\/\*[\s\S]*?(?:\*\/|$)/,lookbehind:!0},{pattern:/(^|[^\\:])\/\/.*/,lookbehind:!0,greedy:!0}],string:{pattern:/(["'])(?:\\(?:\r\n|[\s\S])|(?!\1)[^\\\r\n])*\1/,greedy:!0},"class-name":{pattern:/((?:\b(?:class|interface|extends|implements|trait|instanceof|new)\s+)|(?:catch\s+\())[\w.\\]+/i,lookbehind:!0,inside:{punctuation:/[.\\]/}},keyword:/\b(?:if|else|while|do|for|return|in|instanceof|function|new|try|throw|catch|finally|null|break|continue)\b/,"boolean":/\b(?:true|false)\b/,"function":/\w+(?=\()/,number:/\b0x[\da-f]+\b|(?:\b\d+\.?\d*|\B\.\d+)(?:e[+-]?\d+)?/i,operator:/--?|\+\+?|!=?=?|<=?|>=?|==?=?|&&?|\|\|?|\?|\*|\/|~|\^|%/,punctuation:/[{}[\];(),.:]/};
Prism.languages.javascript=Prism.languages.extend("clike",{"class-name":[Prism.languages.clike["class-name"],{pattern:/(^|[^$\w\xA0-\uFFFF])[_$A-Z\xA0-\uFFFF][$\w\xA0-\uFFFF]*(?=\.(?:prototype|constructor))/,lookbehind:!0}],keyword:[{pattern:/((?:^|})\s*)(?:catch|finally)\b/,lookbehind:!0},/\b(?:as|async|await|break|case|class|const|continue|debugger|default|delete|do|else|enum|export|extends|for|from|function|get|if|implements|import|in|instanceof|interface|let|new|null|of|package|private|protected|public|return|set|static|super|switch|this|throw|try|typeof|var|void|while|with|yield)\b/],number:/\b(?:(?:0[xX][\dA-Fa-f]+|0[bB][01]+|0[oO][0-7]+)n?|\d+n|NaN|Infinity)\b|(?:\b\d+\.?\d*|\B\.\d+)(?:[Ee][+-]?\d+)?/,"function":/[_$a-zA-Z\xA0-\uFFFF][$\w\xA0-\uFFFF]*(?=\s*\(|\.(?:apply|bind|call)\()/,operator:/-[-=]?|\+[+=]?|!=?=?|<<?=?|>>?>?=?|=(?:==?|>)?|&[&=]?|\|[|=]?|\*\*?=?|\/=?|~|\^=?|%=?|\?|\.{3}/}),Prism.languages.javascript["class-name"][0].pattern=/(\b(?:class|interface|extends|implements|instanceof|new)\s+)[\w.\\]+/,Prism.languages.insertBefore("javascript","keyword",{regex:{pattern:/((?:^|[^$\w\xA0-\uFFFF."'\])\s])\s*)\/(\[(?:[^\]\\\r\n]|\\.)*]|\\.|[^\/\\\[\r\n])+\/[gimyu]{0,5}(?=\s*($|[\r\n,.;})\]]))/,lookbehind:!0,greedy:!0},"function-variable":{pattern:/[_$a-z\xA0-\uFFFF][$\w\xA0-\uFFFF]*(?=\s*[=:]\s*(?:async\s*)?(?:\bfunction\b|(?:\([^()]*\)|[_$a-z\xA0-\uFFFF][$\w\xA0-\uFFFF]*)\s*=>))/i,alias:"function"},parameter:[{pattern:/(function(?:\s+[_$a-z\xA0-\uFFFF][$\w\xA0-\uFFFF]*)?\s*\(\s*)[^\s()][^()]*?(?=\s*\))/,lookbehind:!0,inside:Prism.languages.javascript},{pattern:/[_$a-z\xA0-\uFFFF][$\w\xA0-\uFFFF]*(?=\s*=>)/,inside:Prism.languages.javascript},{pattern:/(\(\s*)[^\s()][^()]*?(?=\s*\)\s*=>)/,lookbehind:!0,inside:Prism.languages.javascript},{pattern:/((?:\b|\s|^)(?!(?:as|async|await|break|case|catch|class|const|continue|debugger|default|delete|do|else|enum|export|extends|finally|for|from|function|get|if|implements|import|in|instanceof|interface|let|new|null|of|package|private|protected|public|return|set|static|super|switch|this|throw|try|typeof|var|void|while|with|yield)(?![$\w\xA0-\uFFFF]))(?:[_$a-z\xA0-\uFFFF][$\w\xA0-\uFFFF]*\s*)\(\s*)[^\s()][^()]*?(?=\s*\)\s*\{)/,lookbehind:!0,inside:Prism.languages.javascript}],constant:/\b[A-Z][A-Z\d_]*\b/}),Prism.languages.insertBefore("javascript","string",{"template-string":{pattern:/`(?:\\[\s\S]|\${[^}]+}|[^\\`])*`/,greedy:!0,inside:{interpolation:{pattern:/\${[^}]+}/,inside:{"interpolation-punctuation":{pattern:/^\${|}$/,alias:"punctuation"},rest:Prism.languages.javascript}},string:/[\s\S]+/}}}),Prism.languages.markup&&Prism.languages.insertBefore("markup","tag",{script:{pattern:/(<script[\s\S]*?>)[\s\S]*?(?=<\/script>)/i,lookbehind:!0,inside:Prism.languages.javascript,alias:"language-javascript",greedy:!0}}),Prism.languages.js=Prism.languages.javascript;
Prism.languages.r={comment:/#.*/,string:{pattern:/(['"])(?:\\.|(?!\1)[^\\\r\n])*\1/,greedy:!0},"percent-operator":{pattern:/%[^%\s]*%/,alias:"operator"},"boolean":/\b(?:TRUE|FALSE)\b/,ellipsis:/\.\.(?:\.|\d+)/,number:[/\b(?:NaN|Inf)\b/,/(?:\b0x[\dA-Fa-f]+(?:\.\d*)?|\b\d+\.?\d*|\B\.\d+)(?:[EePp][+-]?\d+)?[iL]?/],keyword:/\b(?:if|else|repeat|while|function|for|in|next|break|NULL|NA|NA_integer_|NA_real_|NA_complex_|NA_character_)\b/,operator:/->?>?|<(?:=|<?-)?|[>=!]=?|::?|&&?|\|\|?|[+*\/^$@~]/,punctuation:/[(){}\[\],;]/};
!function(){if("undefined"!=typeof self&&self.Prism&&self.document){var e="line-numbers",t=/\n(?!$)/g,n=function(e){var n=r(e),s=n["white-space"];if("pre-wrap"===s||"pre-line"===s){var l=e.querySelector("code"),i=e.querySelector(".line-numbers-rows"),a=e.querySelector(".line-numbers-sizer"),o=l.textContent.split(t);a||(a=document.createElement("span"),a.className="line-numbers-sizer",l.appendChild(a)),a.style.display="block",o.forEach(function(e,t){a.textContent=e||"\n";var n=a.getBoundingClientRect().height;i.children[t].style.height=n+"px"}),a.textContent="",a.style.display="none"}},r=function(e){return e?window.getComputedStyle?getComputedStyle(e):e.currentStyle||null:null};window.addEventListener("resize",function(){Array.prototype.forEach.call(document.querySelectorAll("pre."+e),n)}),Prism.hooks.add("complete",function(e){if(e.code){var r=e.element.parentNode,s=/\s*\bline-numbers\b\s*/;if(r&&/pre/i.test(r.nodeName)&&(s.test(r.className)||s.test(e.element.className))&&!e.element.querySelector(".line-numbers-rows")){s.test(e.element.className)&&(e.element.className=e.element.className.replace(s," ")),s.test(r.className)||(r.className+=" line-numbers");var l,i=e.code.match(t),a=i?i.length+1:1,o=new Array(a+1);o=o.join("<span></span>"),l=document.createElement("span"),l.setAttribute("aria-hidden","true"),l.className="line-numbers-rows",l.innerHTML=o,r.hasAttribute("data-start")&&(r.style.counterReset="linenumber "+(parseInt(r.getAttribute("data-start"),10)-1)),e.element.appendChild(l),n(r),Prism.hooks.run("line-numbers",e)}}}),Prism.hooks.add("line-numbers",function(e){e.plugins=e.plugins||{},e.plugins.lineNumbers=!0}),Prism.plugins.lineNumbers={getLine:function(t,n){if("PRE"===t.tagName&&t.classList.contains(e)){var r=t.querySelector(".line-numbers-rows"),s=parseInt(t.getAttribute("data-start"),10)||1,l=s+(r.children.length-1);s>n&&(n=s),n>l&&(n=l);var i=n-s;return r.children[i]}}}}}();
```
